# A Collaborative DNA Barcode Library for the Diptera of Churchill, Canada: Exploration of Biodiversity and Biogeography

**DOI:** 10.1101/2024.03.27.586819

**Authors:** S.E. Majoros, T.A. Elliott, J. Wang, V. Levesque-Beaudin, C.J. Borkent, F. Brodo, S.E. Brooks, S. Boucher, J.M. Cumming, D.C. Currie, T. Ekrem, P.D.N. Hebert, J. Savage, P. Schaefer, B.J. Sinclair, J.H. Skevington, A.M. Solecki, E. Stur, T.A. Wheeler, T.S. Woodcock, A. D. Young, C. Young, S.J. Adamowicz

**Affiliations:** Department of Integrative Biology, University of Guelph, Guelph, Ontario, Canada; China Agricultural University, Beijing, China; Centre for Biodiversity Genomics, University of Guelph, Guelph, Ontario, Canada; Biological Control Program and California State Collection of Arthropods, California Department of Food and Agriculture, Sacramento, California, USA; Canadian Museum of Nature, Ottawa, Ontario, Canada; Agriculture and Agri-Food Canada, Ottawa, Ontario, Canada; McGill University, Ste-Anne-de-Bellevue, Quebec, Canada; Royal Ontario Museum, Toronto, Ontario, Canada; University of Toronto, Ontario, Canada; Department of Natural History, NTNU University Museum, Norwegian University of Science and Technology, Trondheim, Norway; Bishop University, Sherbrooke, Quebec, Canada; Canadian Food Inspection Agency, Ottawa, Ontario, Canada; Carleton University, Ottawa, Ontario, Canada; Carnegie Museum of Natural History, Pittsburgh, Pennsylvania, USA

**Keywords:** Subarctic, Cytochrome c oxidase subunit 1 (CO1), Boreal habitats, Biodiversity

## Abstract

Churchill, Manitoba, Canada is a diverse subarctic region that has been the focus of DNA barcoding efforts and research for decades. Despite this effort, there are still taxa in this region that are underrepresented in public databases, particularly hyper diverse insect groups. We present a collaborative large-scale DNA barcode reference library for Diptera based on molecular data and expert taxonomic identification, and we explore habitat occupancy, biogeographic patterns, and molecular evolution. The reference library contains 16786 specimens sampled around Churchill from 2005 to 2011. Selected specimens were identified by morphological means, sequenced for the animal barcode marker, and sorted into 2225 Molecular Operational Taxonomic Units (using Barcode Index Numbers: BINs) representing 68 families and 1216 named species. Eighty-five species shared a BIN with another species, 143 species were split across several BINs, and 990 species were assigned to a unique BIN, including 129 BINs new to the Barcode of Life Data System. The intraspecific and nearest neighbour distances varied across families, but most species can be easily distinguished from their nearest neighbours. By combining molecular data with taxonomic expertise, the barcode library provides species-level information for 12608 specimens, supplying future researchers with detailed taxonomic information and the opportunity to perform barcode-based specimen identifications for large-scale studies through sequence matching. Comparison of the Diptera composition of Churchill to other subarctic regions reflected likely postglacial colonization northwards as well as a Beringian and circumpolar component. This study also provides an initial investigation of molecular evolution in the best-represented species and finds variability in the rate of molecular evolution, opening intriguing avenues for further investigation. This study provides a publicly available, detailed reference library for use in future research, as well as new insights into the current diversity and taxonomic composition of Diptera present among the diverse habitats of Churchill.

## Introduction

DNA barcoding, the process by which standardized gene regions are used to identify specimens and to discover and delineate species, is a useful tool that can allow us to document further the diversity of different regions and ecosystems (Floyd et al. 2002; Hebert et al., 2003). These barcodes also enable further analysis and research, allowing us to study biodiversity and temporal and spatial trends in community composition.

There has been a concentrated effort to barcode specimens in various areas globally, including Costa Rica (Janzen & Hallwachs, 2011, Janzen et al., 2009), Finland (Pohjoismäki et al., 2016, Roslin et al., 2021), Greenland (Wirta et al., 2016), and Nunavut (deWaard, 2023, Pentinsaari et al., 2020; Nowosad et al. 2024). Regional barcode studies and libraries provide a valuable resource and allow for the biodiversity of regions to be elucidated, as well as provide the opportunity for further research. Wirta et al. (2016), for example, provided a barcode library including over 403 terrestrial animal species and 160 vascular plant species for the Zackenberg Valley in Northeast Greenland. This library has been used for a variety of novel studies, including to determine the prey of spiders and investigate niche overlap (Wirta et al., 2015a), to elucidate Arctic food webs and interactions between predators and prey (Wirta et al., 2015b), and to investigate the role of flies in the pollination of Arctic plant species (Tiusanen et al., 2016). By continuing to barcode specimens and building these regional libraries, we can expand our knowledge of the communities and ecological processes within these regions.

Since the 1800s, the northern regions of Canada have been the focus of various biological surveys and sampling events. One notable sampling event that spanned several insect orders was the Canadian Northern Insect Survey, conducted from 1947 – 1962 (Freeman, 1959, Danks, 1981, Freeman, 1958, Huckett et al., 1965a, McAlpine, 1965). In 2010-2011, the Northern Biodiversity Program resampled the sites of the Northern Insect Survey with the goal of comparing their more recent survey to the historical one (Northern Biodiversity Program, n.d.).

A taxon that has been of special research interest, both in historical and recent studies, is Diptera (true flies), particularly biting flies and those that act as vectors of disease. In a global malaise trap barcoding study, Srivathsan et al. (2023) found that 20 insect families, 10 of which were from Diptera, made up 50% of the local diversity of flying insects at their sites worldwide. Diptera is a diverse and ecologically important group that contributes to ecosystem functioning, and it has a major impact on agriculture and public health (Adler & Courtney, 2019, Bedini et al., 2017, Bragard et al., 2020, Mesmin et al., 2019, Molinatto et al., 2020, Olafson et al., 2021, Shamshad, 2010, Wearing et al., 2013), and act as useful bioindicators (Czechowski et al., 2020, Hamerlík & Bitušík, 2009, Medeiros et al., 2020, Rossaro et al., 2022). Diptera have been found to act as an important pollinator group in sites across the Arctic, including the Canadian high Arctic (Kevan, 1972), Greenland (Cirtwill et al., 2022, Tiusanen et al., 2016) and northern Finland (Zoller et al., 2020), typically represented by members of the Muscidae, Syrphidae, Scathophagidae, and Empididae families. Flies are also vectors for several important, widespread diseases, including Malaria, Zika virus, Dengue, Chikungunya virus, and West Nile virus (Bonizzoni et al., 2013, World Health Organization, 2018, World Health Organization, 2020). There have been Zika virus outbreaks in over 30 countries, and Aedes aegypti (Linnaeus), the principal vector of this virus, as well as Dengue, Chikungunya, and yellow fever, is a highly adaptive and widespread species (World Health Organization, 2018). Similarly, Aedes albopictus (Skuse), a vector of Dengue and Chikungunya, is an extremely invasive species affecting an increasing proportion of the world (Bonizzoni et al., 2013). Developing a capacity for monitoring potential vectors of disease is important, as species may shift their geographic distributions in response to climate change. Non-biting fly species can also be a public health concern, transmitting microorganisms through their mouthparts, feces, and body surface, and acting as vectors of parasites, disease, and food-borne pathogens (Förster et al., 2007, Graczyk et al., 2001, Pava-Ripoll et al., 2012).

Observations on Diptera diversity have been recorded in northern North America since the 1800s. Huckett et al. (1965a) provided a study of Muscidae (Diptera) specimens from Arctic and subarctic regions of North America, Greenland, and Europe, and described the historical collection events and expeditions that have occurred in these regions. Various historical collection events, occurring from 1835 – 1954, recorded Muscidae flies in northern regions, such as the Hudson Bay region, Alaska, the Bering Sea region, Northwest Territories, Ellesmere Island, Yukon Territory, and the Labrador coast (Aldrich, 1918, Coquillett, 1900, Huckett et al., 1965a, James and Huckett, 1952, Johnson, 1929, Malloch 1919, 1920a, 1921, 1923, Stein, 1898, 1920, Walker, 1849, Weber, 1950, 1954).

Another area that has been the focus of past sampling is Churchill, Manitoba, Canada. Churchill is a diverse subarctic region containing a wide range of wildlife and plant species. Churchill is in the Coastal Hudson Bay Lowland ecoregion (Smith et al., 1998) (Fig 1a), an area characterized by short, cool summers; long, very cold winters; and widespread permafrost (Smith et al., 1998). This region contains a wide range of habitat types, including peat bogs, fens, tundra, wetlands, and boreal forests, and is bordered by Hudson Bay, where sandy beaches and rocky outcrops can be found (Smith et al., 1998). The Churchill River is also an important water feature, in addition to several small, unconnected lakes throughout the region (Smith et al., 1998). The diversity of habitats makes it a place of interest for investigating the diversity of subarctic species and for research into northern ecology and climate change. The establishment of the Churchill Northern Studies Centre in 1976 facilitated the usage of the Churchill area as a hub for northern research (Churchill Northern Studies Centre, 2022). There have been several historical studies and surveys that included Churchill fly species, such as the Biting Fly Survey and Experimental Control Program and the Canadian Northern Insect Survey (Hocking et al., 1950, Miller, 1951, Twinn et al., 1948, Danks, 1981, Freeman, 1958, Huckett et al., 1965a).

**Figure 1.**
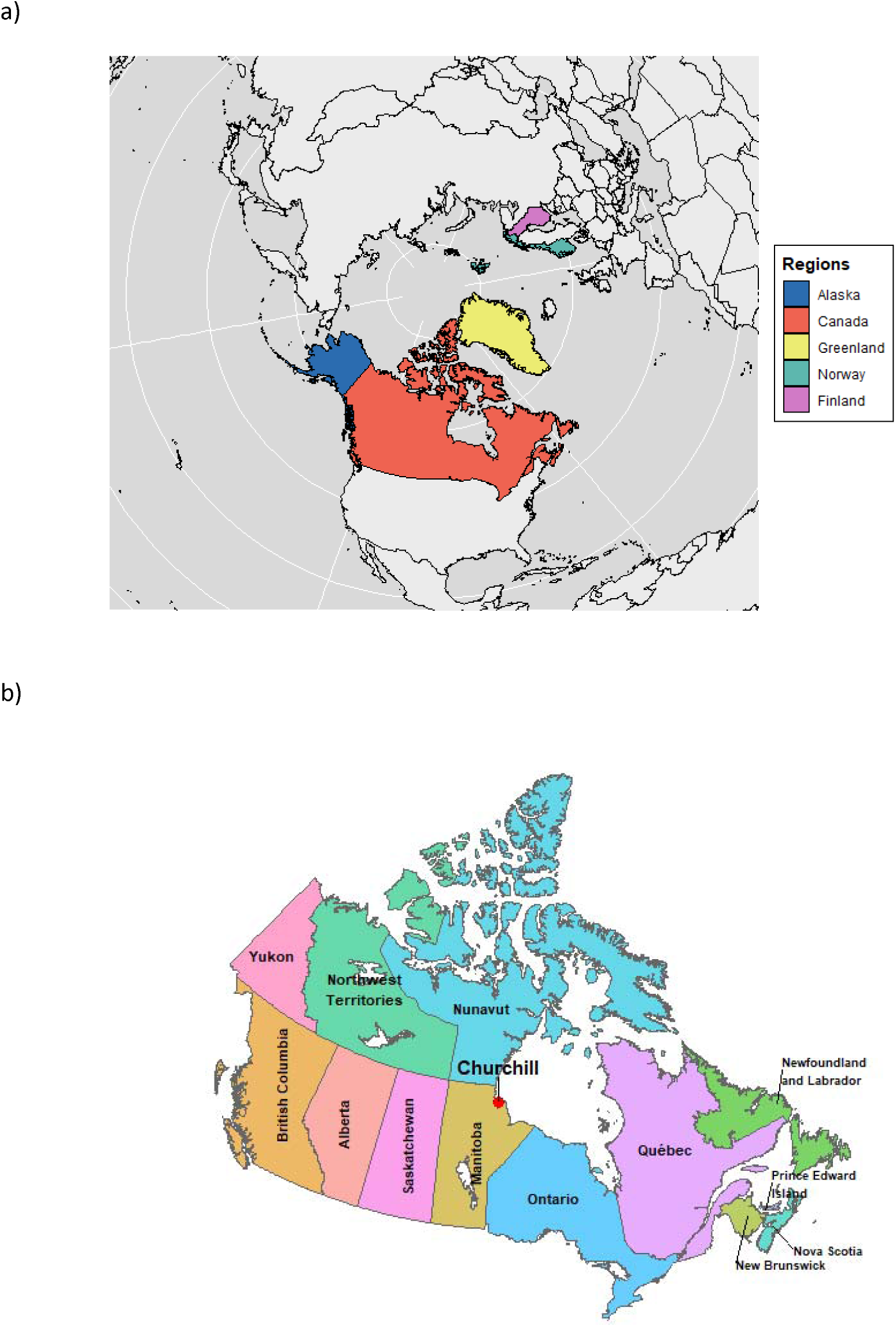
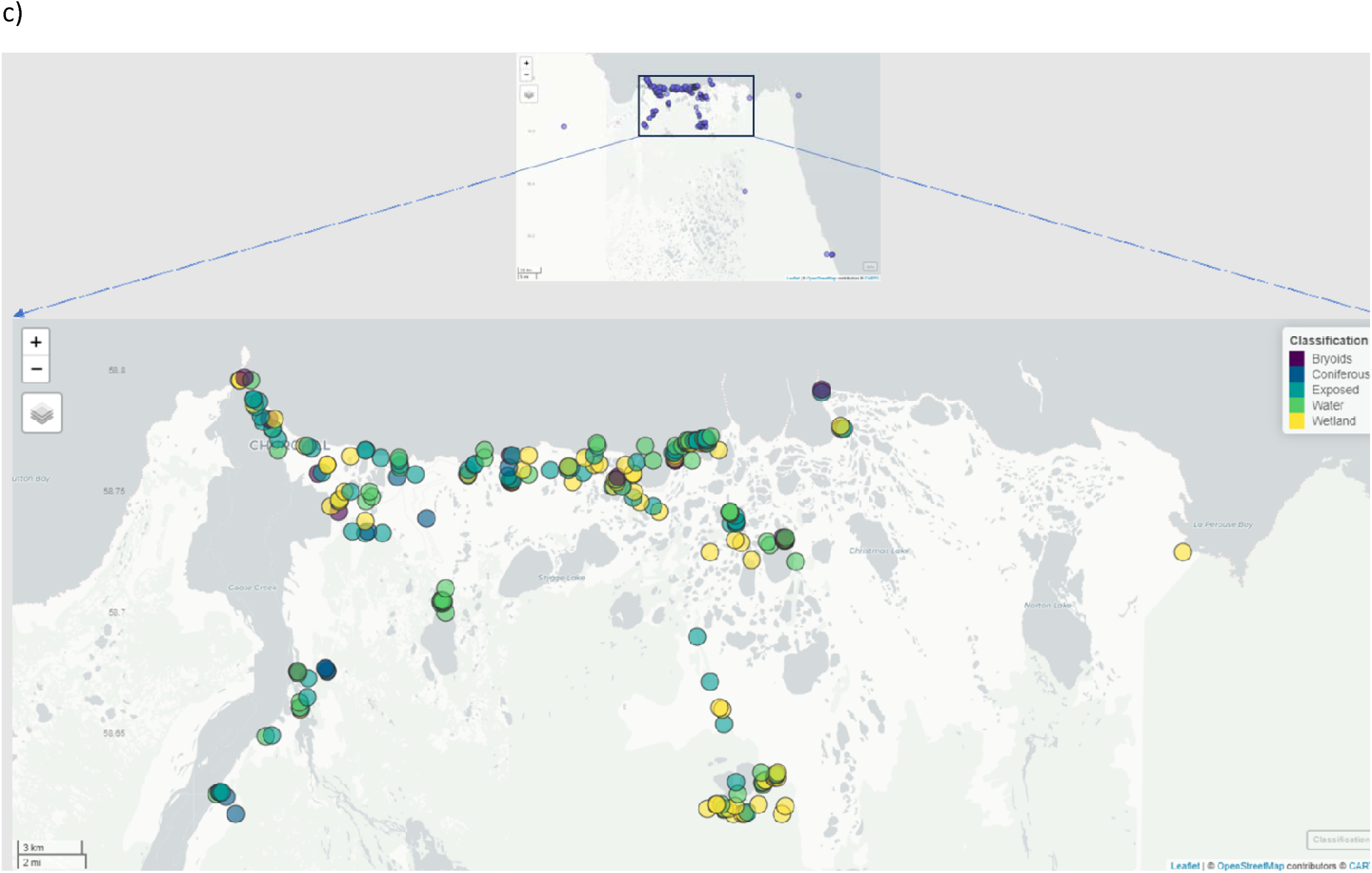
The location of Churchill, Manitoba, and the study sites within the region. **a**) shows the placement of Canada in relation to other northern countries and regions. b) shows the location of Churchill within Canada. Churchill is located on the coast of Hudson Bay in northern Manitoba. It is a Subarctic region located within the Coastal Hudson Bay Lowland ecoregion. c) shows the locations where samples were collected in the Churchill region and the land-use classification categories of sampling sites, with the most samples inferred using the EOSD classifications from satellite data. For details on the classification status for each site, please refer to Supplementary File 6.

Despite this past sampling, work is still needed to fully understand the diversity of species present in Churchill and other northern regions. By continuing sampling efforts, we can develop a solid baseline which can be used by future researchers. Historical sampling events and present-day efforts provide information on Diptera diversity across northern regions and the opportunity to see how diversity changes over time. Renaud et al. (2012a) compared the historical surveys conducted during the Canadian Northern Insect Survey (1947 – 1962) to a more recent Diptera survey conducted in 2007. The sampling methods differed between these time periods, as historical samples were primarily collected using a sweep net, but the 2007 survey was done using Malaise traps and pan traps, in addition to sweep nets (Renaud et al., 2012a). This survey compared 10294 specimens representing 155 species of Muscidae from the 2007 survey to 2881 historical specimens representing 105 species (Renaud et al., 2012a). Eighty-seven percent of the species found in the historical records were rediscovered in the 2007 survey, and Spilogona Schnabl remained the most diverse genus of muscids in both time periods, with 38 species as of 1965 and 51 species as of 2007 (Renaud et al., 2012a). With consistent sampling over time, we can document how diversity changes and continue to add to our knowledge and number of known species.

Across geographic regions, DNA barcoding has been shown to be a helpful and successful approach for Diptera, and it is a useful tool for identifying and delineating species within this group (Adeniran et al., 2019, Banerjee et al., 2015, Brodin et al., 2012, Changbunjong et al., 2018, Cywinska et al., 2010, de Souza Pinto, 2023, Machado et al., 2017, Posada-López et al., 2023, Rivera & Currie, 2009, Votýpka et al., 2019). Renaud et al. (2012b) added to our knowledge of Diptera diversity in Churchill by conducting a barcoding study of Muscidae that included 1114 individuals and 160 species. In addition, Churchill has been the focus of many barcoding studies for arthropod taxa such as Coleoptera (Woodcock et al. 2013), Ephemeroptera, Plecoptera, Trichoptera (Zhou et al. 2009, 2010), Hymenoptera (Fernandez-Triana et al., 2011, Stahlhut et al. 2013), Collembola (Porco et al. 2013), Araneae (Blagoev et al. 2013), Ostracoda (Laforest et al. 2013), Crustacea (Jeffery et al. 2011), and Acari (Young et al. 2012).

To add to this barcoding effort, we conducted a large-scale barcoding study of Diptera species in Churchill. Despite previous surveys, there are still gaps in our understanding of Diptera diversity, and there are many families which have not yet been the focus of dedicated barcoding efforts. Previous work was focused on handful of species from a select few families (e.g. Culicidae, Tabanidae, Simuliidae, Chaoboridae) or more extensive surveys of single families such as Muscidae (Hocking et al., 1950, Miller, 1951, Twinn et al., 1948, Huckett et al., 1965a. Renaud et al., 2012a, Renaud et al., 2012b). In this paper, we present a large survey of Diptera, including 16786 specimens and 2235 Barcode Index Numbers (BINs; Ratnasingham and Hebert, 2013) representing 68 families and 1211 species. Through a collaborative approach, we have conducted extensive field collections and created a taxonomically linked reference library that includes molecular data and expert taxonomic identifications. This approach is necessary due to the complex patterns of DNA sequence divergence within Diptera and the challenges with differentiating species using morphological characters alone in certain families, such as Tabanidae and Chironomidae (eg. Mackerras, 1954, Cranston, 2008, Proulx et al., 2013). For example, Lin et al. (2015) determined that a divergence threshold of 4–5% was suitable to delineate species within Chironomidae, reflecting the high genetic diversity within species of the family. However, Renaud et al. (2012b) found that a divergence threshold of 1.2% was more suitable for Muscidae, to optimize between cases of taxonomic lumping vs. over-splitting. Due to these complex and differing patterns of divergence and genetic diversity within Diptera, along with the difficulty in morphological differentiation of species in certain groups, it is valuable to construct an extensive DNA barcode library so that new sequences can be linked with named species and their associated information. In this study, we provide a taxonomically linked reference library, outline the diversity and composition of Diptera currently present in Churchill across diverse habitat types, and contribute barcodes for thousands of individuals to the Barcode of Life Data (BOLD) Systems (https://boldsystems.org/; Ratnasingham & Hebert, 2007).

## Materials and Methods

### Study Site

Samples were collected from sites around Churchill, Manitoba (Latitude: 58.73, Longitude: – 93.82 for the Churchill Northern Studies Centre) (Figure 1b) from 2005 to 2011. Sampling intensity varied across years, with the most sampling occurring in 2010 and the least occurring in 2011 (Supplementary File 1: Supplementary Figure 1.). Sampling was conducted across several areas within the Churchill region, and areas closer to the town of Churchill and the Churchill Northern Studies Centre were sampled more frequently. Two hundred and sixty-seven sites were sampled, based on unique pairs of latitude and longitude coordinates. Sites were chosen to represent a wide variety of habitat types in the Churchill area, and accessibility of the sites was also considered.

### Sampling Methods

There were two main approaches used for sampling. The first strategy is general collection, using a wide variety of methods and conducted by a variety of expert investigators, towards the reference library. This approach is expected to yield a more complete species list for the Churchill region. The field collector, identifier, and other metadata are available for each specimen through BOLD. Specimens were collected using various methods, including baited traps, Berlese traps, dip nets, fan traps, hand collection, intercept traps, light traps, Malaise traps, pan traps, pitfall traps, sweep nets, and UV light traps. As Diptera species vary in their microhabitats and behaviours, the sampling method used can result in a bias as to what types of species are collected. Therefore, by using different trap types and sampling methods, a wider variety of the species that are present in the community should be detected. Most specimens were caught using Malaise traps (Figure 2). The second strategy is sampling conducted using Malaise traps by a single investigator (J. Wang) to obtain detailed data from five sites, in addition to the sampling of a number of other sites using a variety of methods. These sites were repeatedly sampled (five Malaise: 26-29 times; other sites: 1-25 times, median of 5 times) throughout the summer of 2010. The minor variability in the number of sampling events per Malaise site related to weather and likely polar bear-inflicted damage to traps. This subset of the data could be queried for a temporal analysis over the summer season by future researchers.

**Figure 2.**
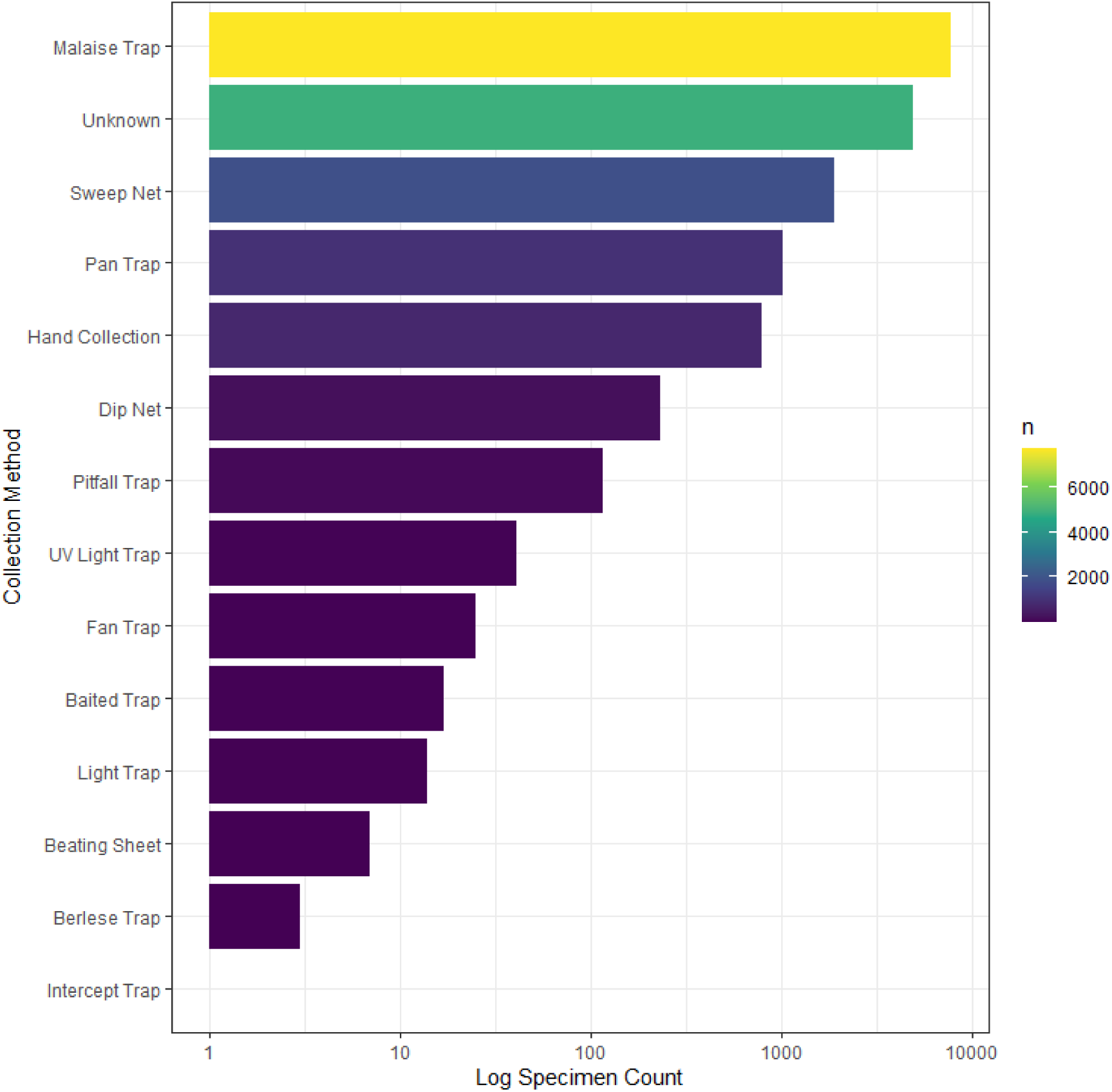
The number of Diptera specimens sampled with each collection method depicted with a log scale for specimen count. The majority of specimens were caught using Malaise traps.

### Barcoding Methods

DNA barcoding was conducted at the Canadian Centre for DNA Barcoding (https://ccdb.ca/) located at the University of Guelph. For sampling strategy 1, experts who conducted the field collecting focused upon their respective taxonomic groups and pulled specimens from any bulk samples (e.g. pitfall traps, etc.) for sequencing, with the goal of including all species present in the samples. All collected specimens were typically sequenced, except in cases where particular species were hyper-abundant in the samples. For sampling strategy 2, three individuals per morphospecies per collecting event (i.e. per Malaise trap, sweep net, pan trap, hand collection, or light trap sample) were generally selected by J. Wang for molecular analysis. Additional specimens (>= 10 per morphospecies) were sequenced from the Malaise trap samples for the following diverse families: Cecidomyiidae, Scathophagidae, Ulidiidae, Ceratopogonidae, Culicidae, Tabanidae, Phoridae, Muscidae, Anthomyiidae, Simuliidae, and Psychodidae. DNA was extracted from each adult specimen using a single leg, and from each larvae using a section of the specimen. For some small-bodied specimens, the protocol from Porco et al. (2010) was used to extract DNA from the whole specimen. The protocol from Ivanova et al. (2006) was used for DNA extraction, and the final elution volume was 50 µl. The primers used for the amplification of the DNA barcode region of COI are included in Supplementary File 2. The PCR reaction volume was 12.5 µl and included 1.25 µl of 10x reaction buffer, 1.25 pmol of primer, 0.3U of Platinum Taq DNA polymerase, 5% trehalose, 2.5 mM of MgCl_2_, 50 um of dNTP, and 2 µl of genomic DNA. The E-gel 96 System (Invitrogen, Carlsbad, Ca, USA) was used to visualize the PCR products on 2% agarose gels. The PCR products were bidirectionally sequenced using the protocols from Hajibabaei et al. (2005) and BigDye v3.1 on an ABI 3730xl DNA Analyzer. Using CodonCode Aligner v.3.0.2 (CodonCode Corporation), the forward and reverse sequences were used to assemble consensus sequences for each individual.

### Specimen Identification

Specimens were identified using morphological characteristics to the family level (McAlpine, 1981). After sequencing, 1-5 specimens from each BIN were identified by experts to species level for 19 of the families based on the specimen’s morphology. BINs are clusters of barcode sequences similar to species and are the operational taxonomic units (OTUs) used by BOLD (Ratnasingham and Hebert, 2013). The goal was to identify a minimum of three specimens per BIN and per species, but some BINs were singletons or doubletons. Of the 16786 records, the taxonomy of 9159 records were assigned using morphological identification. The remaining 7303 records were identified using barcodes matching to BINs on the BOLD database, 5590 of which belonged to BINs which were verified by a taxonomic expert. The reference materials used to identify the specimens are included in Table 1.

**Table 1.**
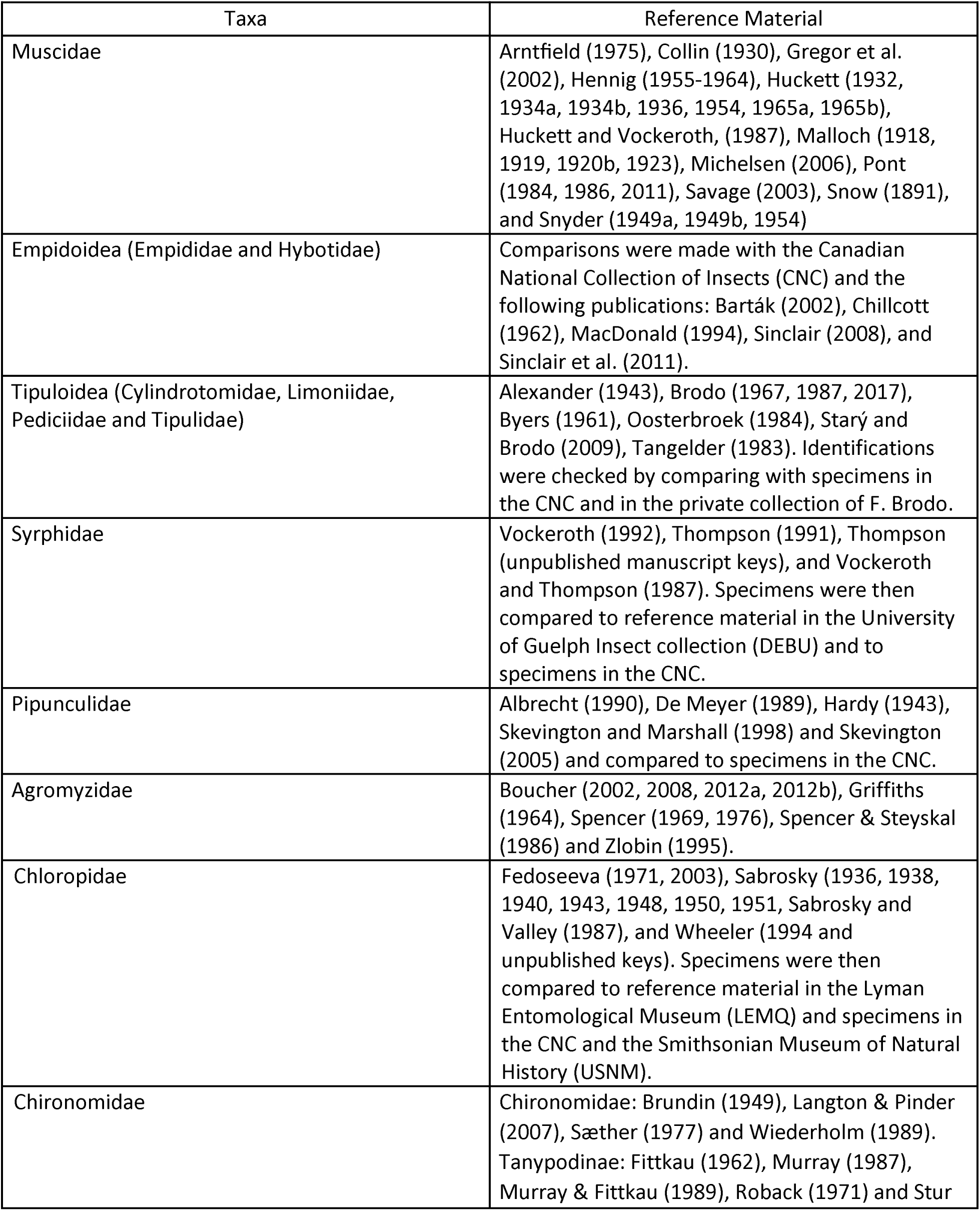
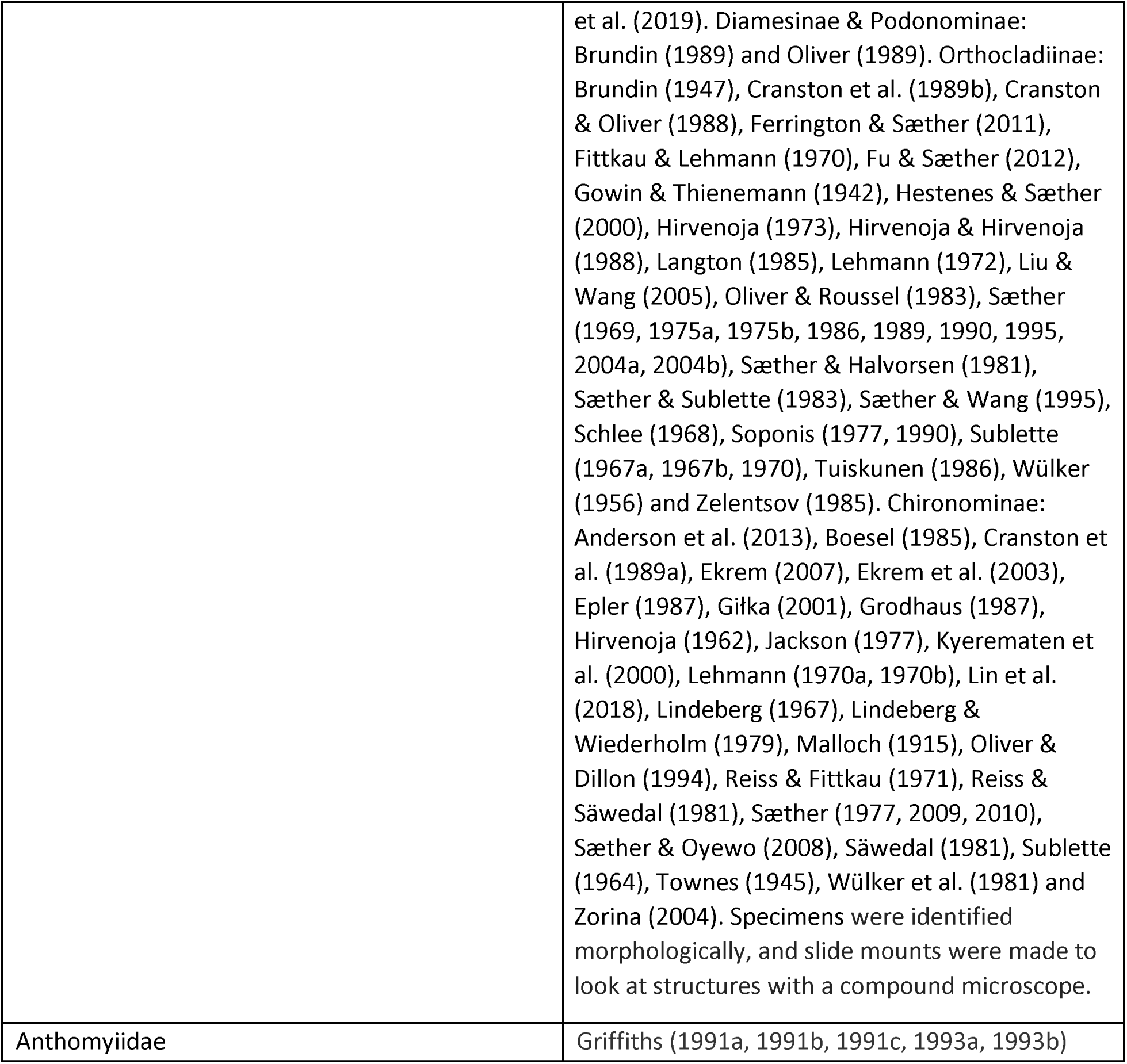
The reference materials used to identify specimens for each Diptera family.

### Specimen and Data Storage

The specimens are stored in 95% ethanol, pinned or slide mounted material in various institutions including the Centre for Biodiversity Genomics (13222 specimens), the Wallis Roughley Museum of Entomology at the University of Manitoba (92 specimens), the Carnegie Museum of Natural History (251 specimens), the Insect Collection at Bishop’s University (868 specimens), the Canadian National Collection of Insects, Arachnids and Nematodes (1312 specimens), and the Department of Natural History at NTNU University Museum (1042 specimens).

The specimens were each assigned a unique specimen ID and photographed before the data were uploaded to BOLD. The data are available in the dataset DS-CDIPTERA (dx.doi.org/10.5883/DS-CDIPTERA) (Majoros et al., 2023) and are also available on GenBank. A portion of the data has been previously published in association with Renaud et al. (2012b). The full dataset including BINs, taxonomic assignment, voucher storage information, and sequences is available in Supplementary File 3. All sequences found to be contaminated or flagged by BOLD for errors or stop codons associated with the COI markers, were removed from the dataset.

### Quality Checking, Analyses and Summary Statistics

A phenogram was created using the Neighbor-Joining method (Saitou & Nei, 1987) for species in the dataset using the Taxon ID Tree tool available on BOLD version 4.0 (Ratnasingham & Hebert, 2007). For the phenogram, Kimura 2 parameter (K2P) was used as the model for calculating distances between DNA sequences, and pairwise deletion was used for ambiguous base/gap handling. The K2P distance model is the one most commonly employed for neighbour-joining phylogram construction using BOLD, is commonly used in barcoding (allowing for comparisons between studies), and, despite criticisms of its use (Srivathsan and Meier, 2011) success rates of specimen identification to species even if it may not be the most appropriate mutation model (Collins et al., 2012). The sequences were aligned using the BOLD aligner (amino acid based hidden Markov model (HMM) (Eddy, 1998)). Known contaminants, records with stop codons, and sequences flagged by BOLD were excluded, and the phenogram was coloured by family. This phenogram was used to look for any misplaced sequences at the family level, and any long branches not matching the typical patterns of genetic diversity in the families. It is not equivalent or intended as a phylogeny. The mean and maximum intraspecific distances for each species and nearest neighbour distances (the shortest distance between a sequence in one species to a sequence in another species) were calculated using the Barcode Gap Analysis tool on BOLD. For these distance measures per species, pairwise distance (p-distance) was used as the sequence distance measure for comparative purposes, as this method is used by the BIN algorithm (Ratnasingham & Hebert, 2013). Discordant BINs (BINs where more than one taxonomic name was provided for the included records, at any taxonomic rank) were found using the BIN Discordance tool on BOLD, using records with a sequence length over 100 base pairs and excluding known contaminants, records with stop codons, and flagged records.

Other analyses and all summary statistics were completed in the R programming language version 4.3.0 (R Core Team, 2023). Various plots were created using the package ggplot2 version 3.4.2 (Wickham, 2016), including bar plots and scatterplots, to summarize and investigate different aspects of the dataset, as follows. Using the categories outlined in Ratnasingham and Hebert (2013), we determined the number of matches (species assigned to a unique BIN), splits (a single species split across several BINs), and merges (species that share a BIN with another species) in the dataset. A linear regression was performed to determine if there is a relationship between the number of specimens sequenced of each species and the number of BINs associated with that species. Only species with two or more specimens were included in the regression. Species and BIN accumulation curves were created using the iNEXT function, and Chao1 estimators (Chao, 1984) were calculated using the ChaoRichness function from the iNEXT package version 3.0.0 (Chao et al., 2014). As typically only 1-5 specimens per morphospecies per collecting event were selected during sampling, depending upon specimen availability, there is a bias in analyses that rely on abundance data. Nevertheless, we expect the specimen counts to be correlated with abundance. In order to account for lack of full abundance data, both abundance and incidence-based richness values were calculated. For the iNEXT function, datatype was set to “abundance” and nboot was set to 1000. For the ChaoRichness function, datatype was set to “abundance” to determine abundance-based richness, and to “incidence_freq” to determine incidence-based richness.

To compare the intraspecific distances across families and species, a Maximum Likelihood (ML) tree was created using the 50 species with the most records (specimens). The ML tree used in this study is not intended to be equivalent to a phylogeny, as it is based upon a single molecular marker and a subset of species, but to provide a visualization tool showing the intraspecific distances across families. Sequences with that were flagged with issues, such as stop codons or contamination, were excluded, as well as sequences with more than 1% ambiguous bases or gaps, and leading and trailing ambiguous bases and gaps were trimmed from each end. A centroid sequence (the sequence with the minimum average distance to others in the species) was chosen for each species. These sequences were aligned using DECIPHER version 2.28.0, following, its recommended steps (Wright, 2016). The AlignSeqs function uses the profile-to-profile method to align the sequences following a guide tree. The profile-to-profile method merges profiles along a guide tree until all the sequences in the set are aligned. As no guide tree was provided and the default parameters were used, the sequences were aligned first using a single-linkage guide tree based on a distance matrix of shared k-mers. The initial alignment was then re-aligned using an UPGMA guide tree. Finally, the alignment subsets were re-aligned to the rest of the alignment. Two alignments were created by this method, and the best was selected by the function using the sum-of-pairs score. The maximum likelihood tree was created using phangorn version 2.11.1 (Schliep, 2011). Models were tested using the ModelTest function from phangorn, and the best model of molecular evolution for the data was determined to be GTG+G(4)+I, as this model had the lowest score for both the Bayesian Information Criterion (BIC) and the Akaike Information Criterion (AIC). The intraspecific distances were mapped to the ML tree using the species names and the function full_join from the package dplyr version 1.1.2 (Wickham et al. 2023). Upon constructing the tree and observing long branch lengths, several steps were taken to further investigate the sequences. The quality of the alignment created using DECIPHER was checked by viewing it in MEGA-X: Molecular Evolutionary Genetics Analysis version 10.2.0 (Tamura et al., 2021). The identities of the sequences were confirmed using the Basic Local Alignment Search Tool (BLAST) (Altschul et al., 1990). We compared these sequences to others of the same species available on BOLD and constructed trees using other sequences to check for long branch lengths. We also constructed trees based on the first, second and third codon positions, as well as an amino acid tree, in MEGA-X. After these diagnostic checks, these long branches were determined to be real and not artifactual.

### Comparison of BIN composition among geographic regions

The dataset was compared to public data from other northern and southern regions to determine how many BINs were shared. Data from Canadian provinces and territories, as well as Alaska, Greenland, Finland, and Norway, were pulled directly into the R environment using BOLD’s application programming interface (API). Data was downloaded on March 21 and 22 2026. The number of shared BINs between each geographic region and Churchill was determined. The alpha diversity was calculated for each region using the Shannon Index and the Simpson Index, with the expectation that these indices are slightly over/underestimated due to typical sorting procedures especially for Malaise trap samples. Specimen selection may underestimate abundance values, specifically of hyper-abundant species, in favour of detecting as many species as possible in the bulk samples. Alpha diversity was obtained using the function diversity from the R package vegan version 2.6.4. (Oksanen et al., 2022), rarefied based on the number of samples in the lowest sampled region. Beta diversity between regions was obtained using the function vegdist from the same package to calculate the Bray-Curtis index using relative abundance. A list of BINs shared between Churchill and each region is available in Supplementary File 4.

### Land-use classification, visualization, and taxonomic breakdown by land-use class

We used an exploratory, graphical approach to investigate the taxonomic composition of Churchill dipterans among major habitat types. We determined the family-level taxonomic composition across the varied landscape of Churchill based on the most common sequences encountered in different land-use and land-type categories, such as wetlands and coniferous forest. Land-use classification for the area surrounding Churchill was determined by extracting the Earth Observation for Sustainable Development (EOSD) categories for each unique set of GPS coordinates from the LUC_EOSD_Northeast dataset from the provincial government of Manitoba Conservation and Water Stewardship website (Manitoba Land Initiative, accessed June 16 2023). Briefly, the “GCS_North_American_1983” Geographic Coordinate System was selected to ensure accurate spatial reference. Next, the unique GPS coordinates were inputted into ArcGIS, and the “Longitude” and “Latitude” attributes were defined (Environmental Systems Research Institute, 2015). The “Spatial Analyst” toolbar was then accessed, and the “Extract Values to Points” tool was selected from the “Extraction” toolset. The shapefile served as the input point layer, while the raster layer was chosen as the input raster. To store the extracted land-classification values, specific output fields were specified and extracted. Classification data were then joined to the rest of the metadata for the unique sites using tidyverse v. 1.3.2 (Wickham et al., 2019). Definitions for various land-use categories can be found in Supplementary File 1. The map itself (see Figure 1) was generated in mapview v. 2.11 in R (Appelhans et al., 2023). For the visualization, specimens lacking GPS coordinates were filtered out, along with seven sites that had <3 specimens and were geographically distant from the primary cluster of sampling sites closest to Churchill itself.

The specimen count by family and for each unique sampling site was determined by grouping data in R using the tidyverse v. 1.3.2 (Wickham et al., 2019) and janitor v. 2.2 (Firke, 2023) suite of data manipulation and data cleaning packages. A table containing the specimen count/site along with their respective EOSD classification is included in Supplementary File 5. Pie charts for visualization were also created in R using ggplot2 v. 3.4.1 (Wickham, 2016) and viridis v. 0.6.3 (Garnier et al., 2023). To illustrate the phylogenetic relationships between different families, taxa within the same infraorder or superfamily (as determined by the phylogeny in Wiegmann et al (2011)), were assigned different shades of a similar colour using hex codes in R.

## Results

### Dataset Characteristics

We created a reference library containing 16786 specimens and 2235 BINs representing 68 families and 1211 species. These were collected by 45 named individuals, including trainees and taxonomists, along with a number of unnamed undergraduate students from field courses. Specimens were identified by 43 individuals, and in 5507 cases specimens were assigned identifications based on matches to BINs that had been verified by a relevant taxonomic expert. Nearly half the dataset was collected by a single collector in 2010 (J. Wang, n=7919), and most of the specimens were collected via Malaise traps (n=8626), followed by sweep netting (n=3295). Not all BINs were identified to the species level, and some species were split across multiple BINs. A phenogram showing the relationship between the taxa collected in this study shows that COI sequences within families generally group together for the Churchill area (Supplementary File 6). However, the families within superfamily Tipuloidea were separated on the phenogram, as well as species within the family Limoniidae, despite being closely related. As this is not a phylogeny and Churchill is a small northern region with lower species richness, these results are not unusual. Muscidae was the most represented family in regard to specimen number, with 2478 specimens collected, while Chironomidae had the highest species richness, with 214 species recorded (Figure 3). A total of 12608 specimens have been identified to species level. Species such as Aedes nigripes (Zetterstedt), Aedes communis (De Geer), Aedes excrucians (Walker), Aedes campestris Dyar & Knab, Aedes flavescens (Muller), Aedes cinereus (Meigen), Chrysops nigripes Zetterstedt, Chrysops mitis Osten Sacken, Hybomitra affinis (Kirby), Hybomitra zonalis (Kirby), and Hybomitra hearlei (Philip) have been previously sampled in historical studies (Hocking et al. 1950, Miller 1951, Twinn et al. 1948) and have been sampled again in our 2005-2011 survey.

**Figure 3.**
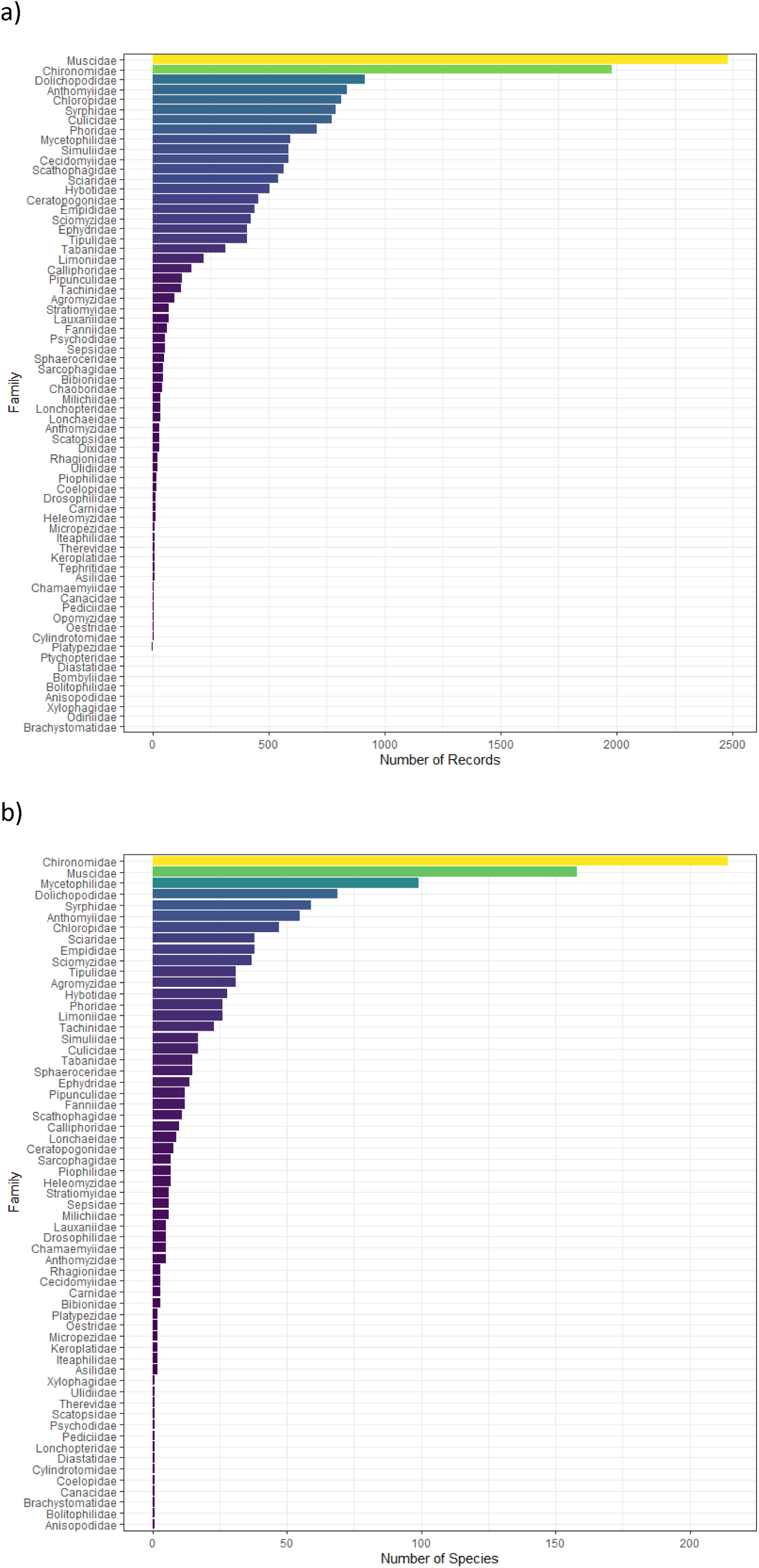
The number of a) records and b) named species collected for each family. In a), some families (Ptychopteridae to Brachystomatidae) have less than 3 records. Muscidae was the most represented family in our dataset in regard to specimens, with 2478 specimens collected, while Chironomidae had the highest species richness, with 214 named species recorded.

Using the categories outlined in Ratnasingham and Hebert (2013), 990 matches (species assigned to a single BIN), 143 splits (a single species split across several BINs), and 85 merges (species that share a BIN with another species) were recorded. Out of the matches, 693 matches were for species represented by more than one sequence. Most species were assigned to a unique BIN. There was a significant relationship between the number of specimens sequenced of each species and the number of BINs associated with that species (p-value = 2.2ex10^-16^ R^2^ = 0.07), and species with more specimens appear to be more likely to be split across BINs (Supplementary File 1: Supplementary Figure 2.). This relationship persists even when excluding species with 4 or greater BINs and over 200 samples, although in both cases the amount of variation explained is small. A total of 948 BINs did not have a species-level identification. Of these, 491 have a genus-level identification, and all have a family-level identification. The majority of BINs are not unique to this study, but contain sequences from BINs already on BOLD that were found in previous works and other geographic regions (Figure 4). Our study contains 129 BINs new to BOLD (as of March 17^th^ 2026), representing 41 species across 23 families. The families Cecidomyiidae and Chironomidae had the most new BINs, with 37 and 25 new BINs, respectively. Some of these may be species new to science; they may be species that have not been registered in BOLD with taxonomic metadata; or these may represent newly detected mitochondrial phylogroups within known species. Thirty-nine discordant BINs were found, most of which appear to contain two species within the same genus.

**Figure 4.**
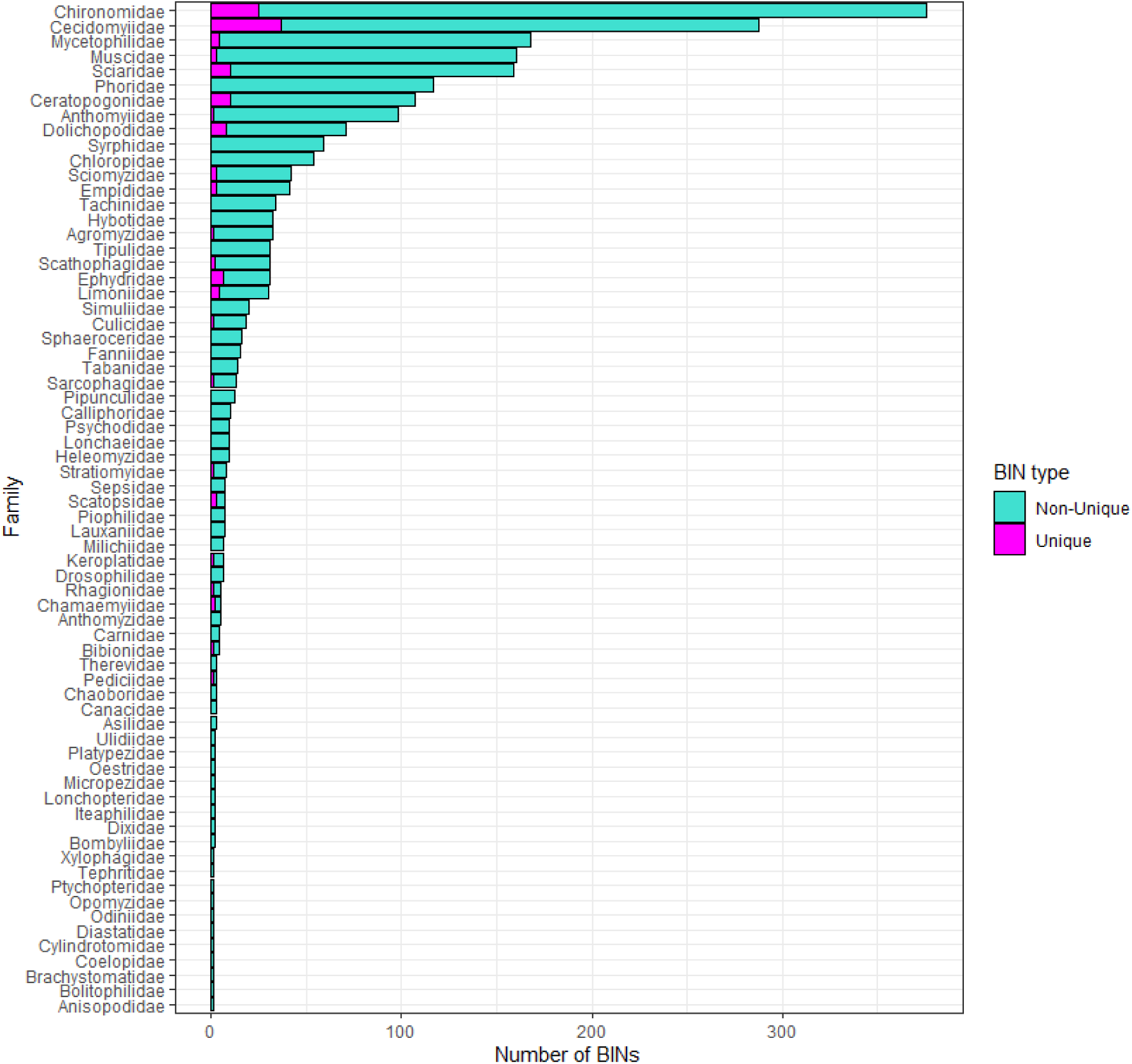
The number of BINs unique to the project on BOLD (i.e. new to BOLD) and the number of non-unique BINs (i.e. already represented in BOLD) for each family. The families Cecidomyiidae and Chironomidae had the most unique BINs, with 37 and 25, respectively.

### Completeness of Sampling

A species accumulation curve and a BIN accumulation curve were created, and Chao1 estimators were calculated (Figure 5). Both curves point to a slowing rate of accumulation. For species, 79.73% of the expected value was observed, and for BINs, 66.5% of the expected value was observed. These results were consistent for both abundance-based and incidence-based richness. As mentioned in the methods, only 1 –5 specimens per morphospecies were selected for molecular analysis, so the species accumulation curve is overly conservative due to the sampling bias. These results indicate that the region is beginning to be well sampled, although further work is still required to capture the full species richness.

**Figure 5.**
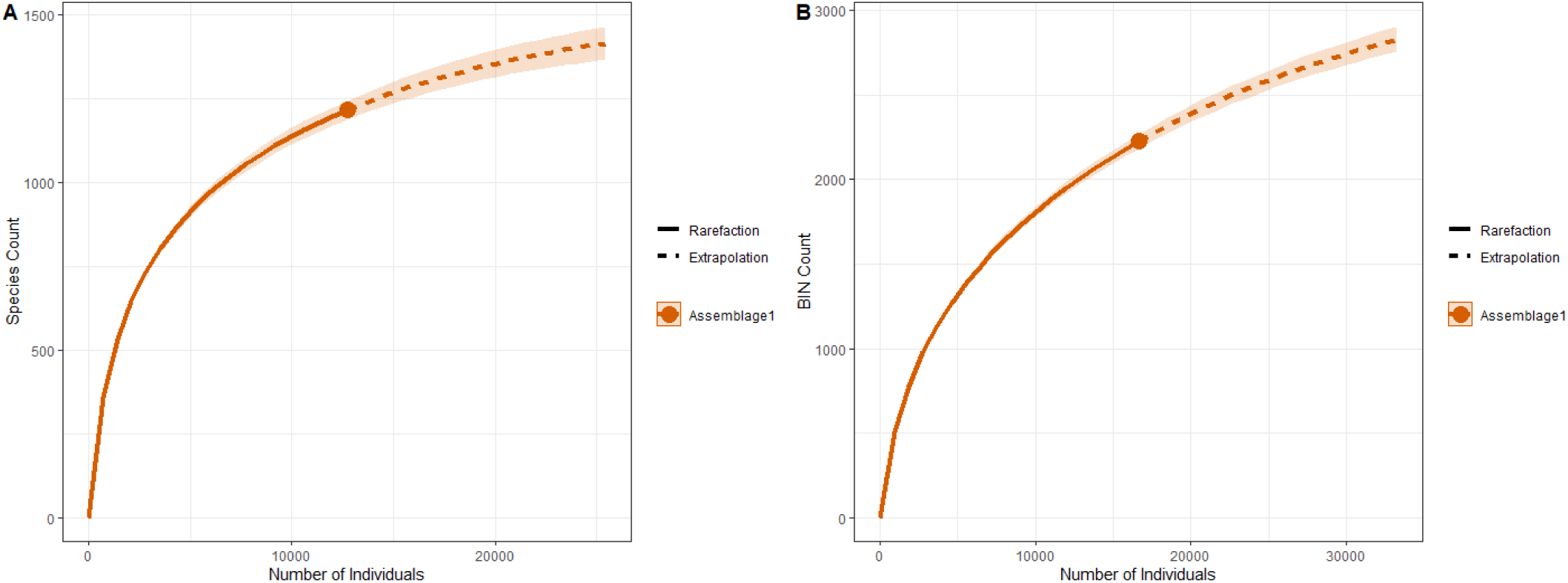
Accumulation curves for the sampled Churchill region. Both curves appear to be slowing, but have yet to plateau, in the rate at which species/BINs are accumulating. A) shows the named species accumulation. The observed value was 1216, and the estimated value was 1525. B) shows the BIN accumulation. The observed value was 2225, and the estimated value was 3344. These results indicate that the region is reasonably well sampled.

### Patterns of Sequence Divergence

The distance to nearest neighbour and maximum and mean intraspecific distances were investigated for each family included in DS-CDIPTERA. The species within Coelopidae had the largest nearest neighbour distances, while Calliphoridae had the lowest (Figure 6). For the mean intraspecific distances, the value for Canacidae was the highest, while Ulidiidae, Lonchopteridae, Chamaemyiidae, and Bolitophilidae all had mean intraspecific distances of zero. There is little overlap between the nearest neighbour distances and mean intraspecific distances, with most mean intraspecific distances being below 5%, and most nearest neighbour distances being above 5%. The max intraspecific distances ranged from 0–16.06% and the nearest neighbour distances ranged from 0–16.23%. Most species appear to be easily distinguished from their nearest neighbours, with 1150 out of 1216 species having a greater interspecific distance than intraspecific (Figure 7). In Figure 8, we present maximum likelihood trees showing the maximum and mean intraspecific distances for the 50 best-sampled species. Genetic distances vary across the tree. The species with the highest maximum intraspecific distance (highlighted green and yellow on the figure) are candidates for examination for potential cryptic species. The full results of the barcode gap analysis are available in Supplementary File 7.

**Figure 6.**
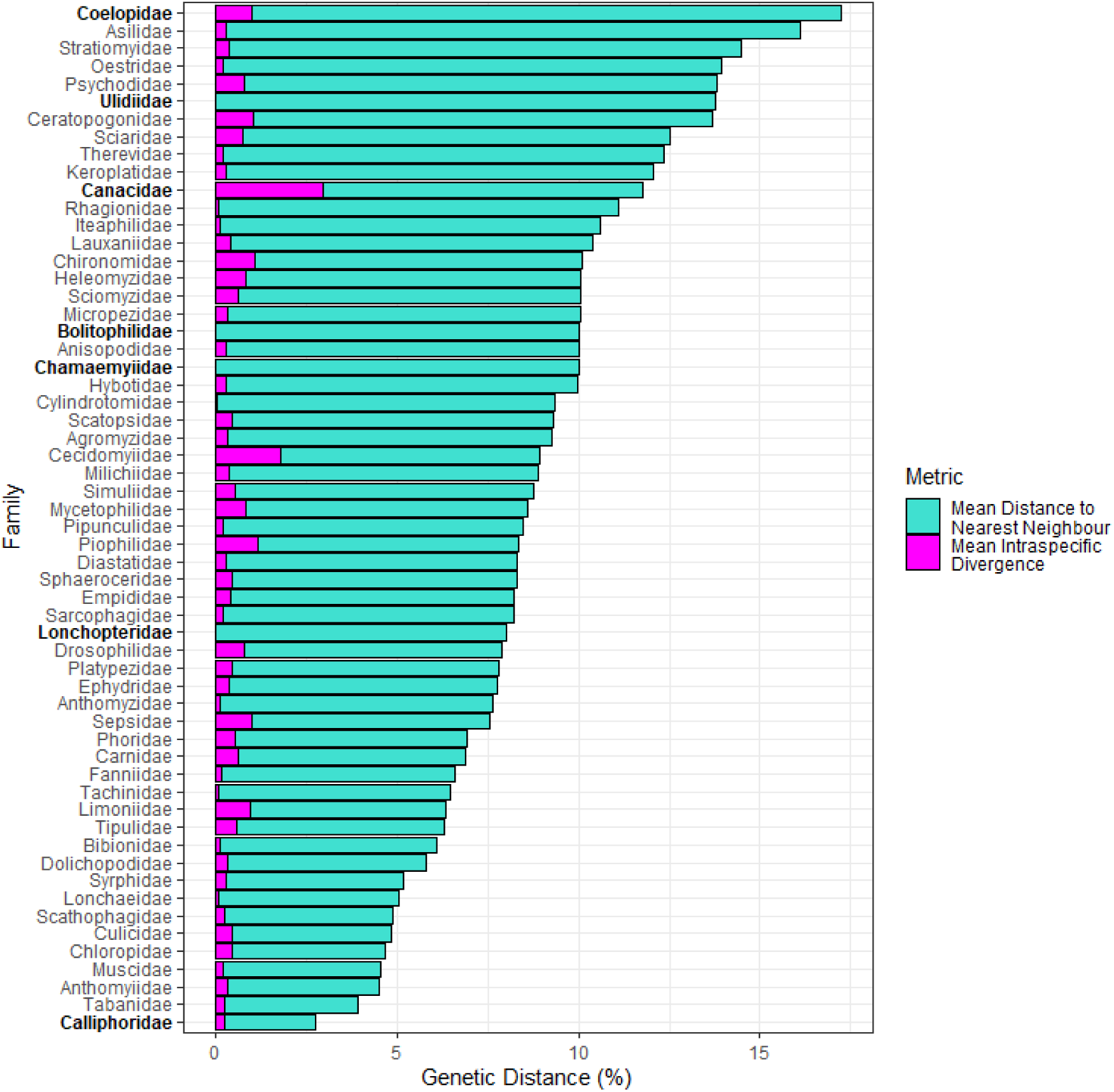
The mean distance to nearest neighbour and mean intraspecific distance for species within each family. The species within Coelopidae had the largest nearest neighbour distances, while Calliphoridae had the lowest. For the intraspecific divergences, the value for Canacidae was the highest, while Ulidiidae, Lonchopteridae, Chamaemyiidae, and Bolitophilidae all had mean intraspecific distances of zero. The families mentioned are indicated in bold.

**Figure 7.**
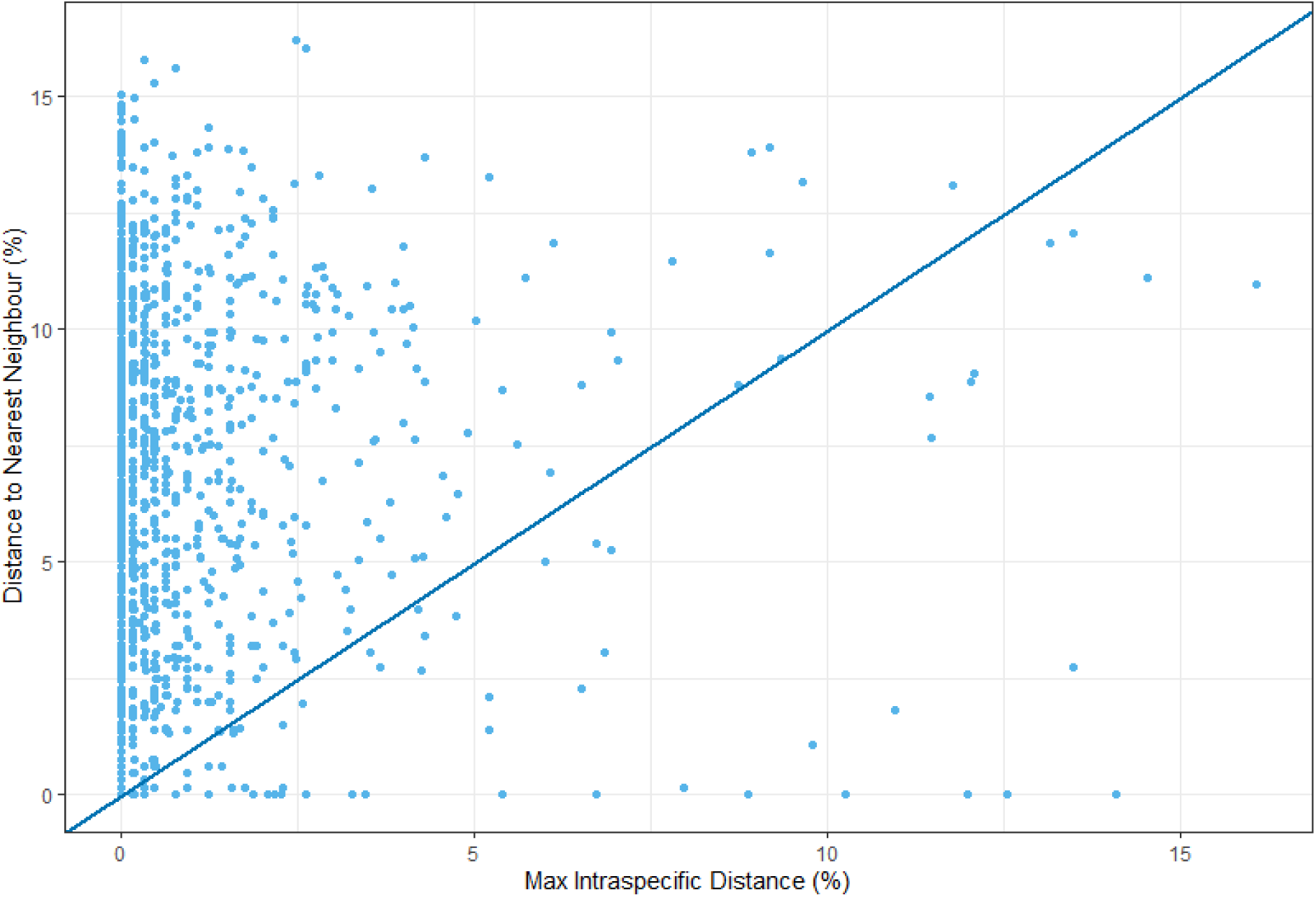
The overlap of the maximum intraspecific distances and the distances to nearest neighbours. Each dot represents a species. Out of the total sample size of 1216 species, 1150 lie above the line, and 66 lie below the line. This indicates that most species have a larger interspecific distance than intraspecific distance, and most species (94.6%) can be easily distinguished from their nearest neighbour.

**Figure 8.**
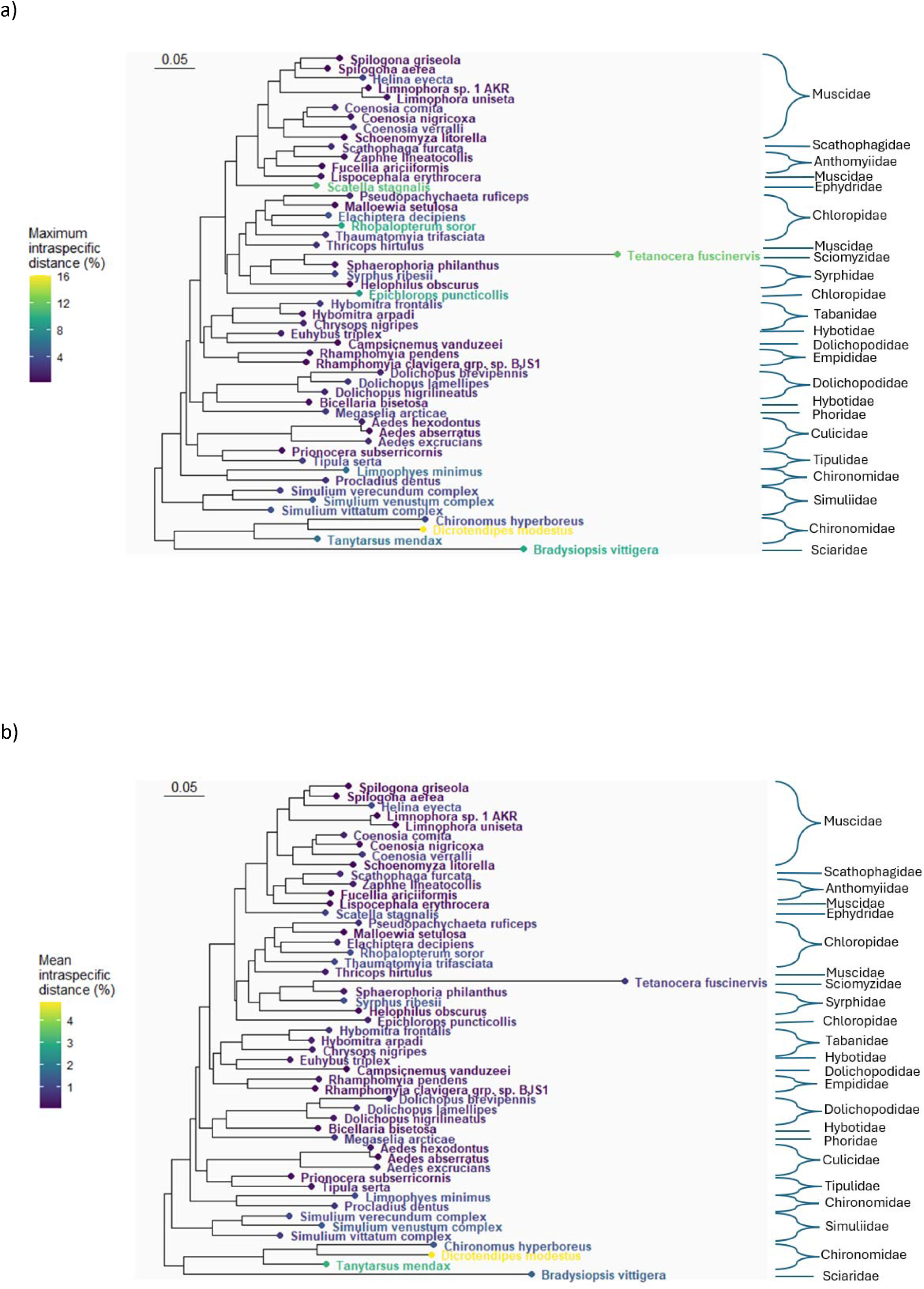
Maximum Likelihood tree of the 51 best-sampled species with an assignment to a Linnean species in the dataset, for comparing intraspecific divergence patterns across families. The a) maximum intraspecific distance (p-distance in %) and b) mean intraspecific distance (p-distance in %) for each species is shown on the tips using the colour scale. The sequences were aligned using DECIPHER version 2.28.0 (Wright, 2016), and the maximum likelihood tree was created using phangorn version 2.11.1 (Schliep, 2011). Higher divergence values were present in scattered locations on the tree, including Dicrotendipes modestus (Chironomidae).

### Comparing BIN Composition Across Geographic Regions

The BINs included in this study were also shared across several other northern and southern regions (Figure 9). When looking at total counts, most BINs were shared with Yukon and Alberta, with 1404 BINs (20.18% of Yukon BINs, 63.1% of Churchill BINs) and 1255 BINs (15.65% of Alberta BINs, 56.4% of Churchill BINs) being shared, respectively. Alaska (261), Northwest Territories (1067) and Greenland (95) all shared ∼28% of their BINs with Churchill, the highest for any of the regions. For Alpha diversity, Finland was the most diverse when using the Shannon Index (7.0) and Simpson Index (0.998), followed by Ontario (6.71, 0.997) and British Columbia (6.64, 0.997). For beta diversity, Churchill was most similar to Alberta (0.832) and Northwest Territories (0.836), followed by Alaska (0.859).

**Figure 9.**
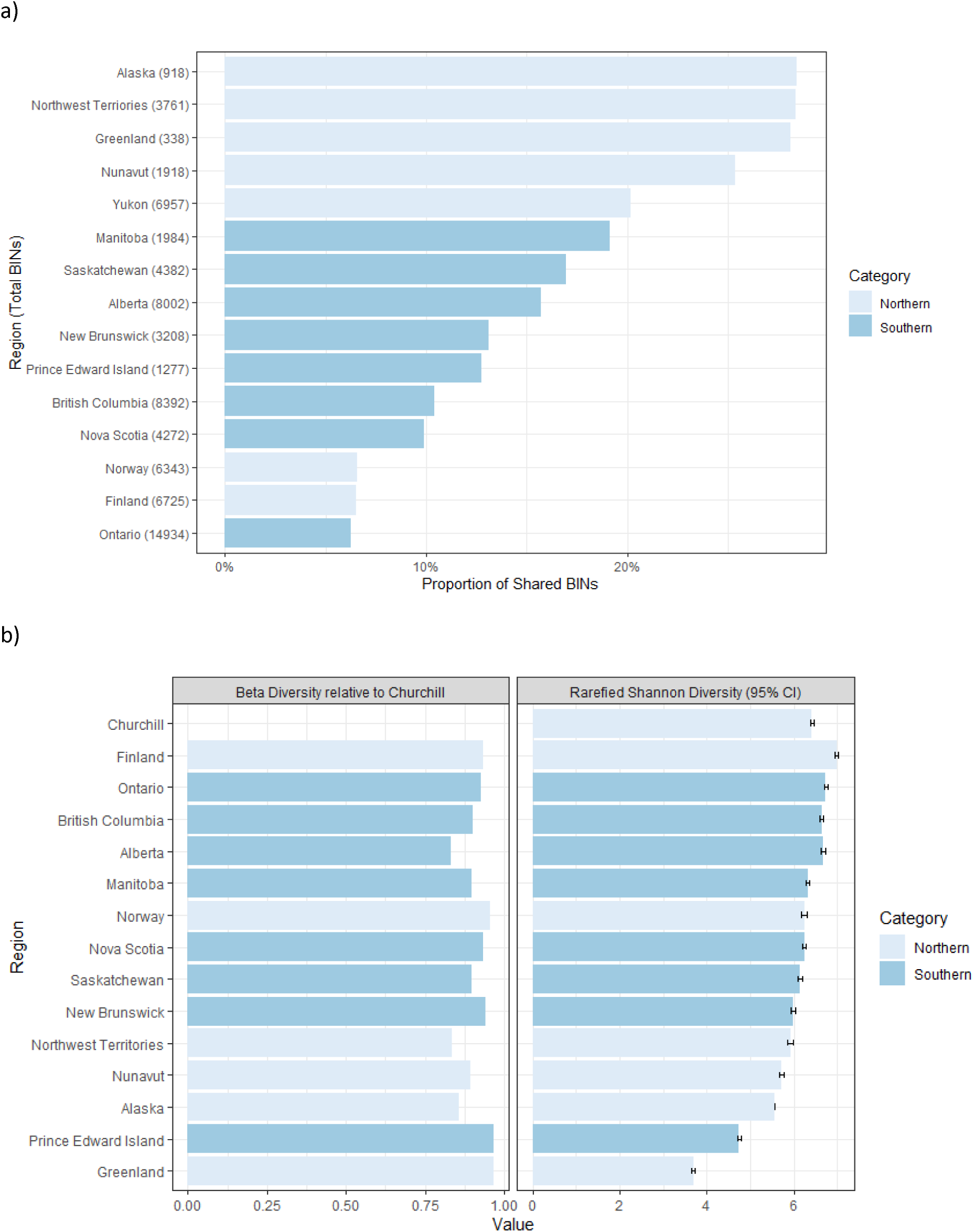
The number of BINs shared between our Churchill barcode library and other geographic regions. **a**) shows the total count of BINs in each geographic region and the proportion of shared BINs between the region and Churchill, while b) shows the alpha diversity (Shannon diversity rarefied 100 times by the sample number in the lowest sampled region) of the geographic regions and the beta diversity (Bray-Curtis index using relative abundance) between the region and Churchill. For a), most BINs were shared with Yukon, with 1404 BINs (20.18% of Yukon BINs, 63.1% of Churchill BINs) being shared. b) indicates that Churchill is most similar to Alberta and the Northwest Territories with a beta diversity of 0.836. Finland and Ontario had the highest alpha diversity. Churchill shared BINs with both northern and southern regions. For total counts, Churchill shared more with northern regions, and when calculating beta diversity, Churchill was more similar to other northern regions.

### Taxonomic Composition by Habitat

Wetlands yielded the most sequenced specimens, with a total of 6723 specimens and 68 families represented with at least one specimen. This was followed by water sites (n=5107); exposed/barren areas (n=3876), which were composed primarily of human adjacent/impacted areas and rock pool bluffs overlooking Hudson Bay; and coniferous (n=707) sites. Muscidae specimens were in the top 10 most-sampled families by specimen number of all land-use areas, and the most represented in our dataset in two of the areas (wetlands, water), with Chironomidae being the second most represented in water and wetland, and the most represented in exposed/barren and bryoid sites (Figure 10). This trend continued with Dolichopodidae, Syrphidae, Chloropidae, and Anthomyiidae found in four of the five areas. Bryoids were less densely sampled (n=198), with Chironomidae and Muscidae dominating in number of specimens sampled as well.

**Figure 10.**
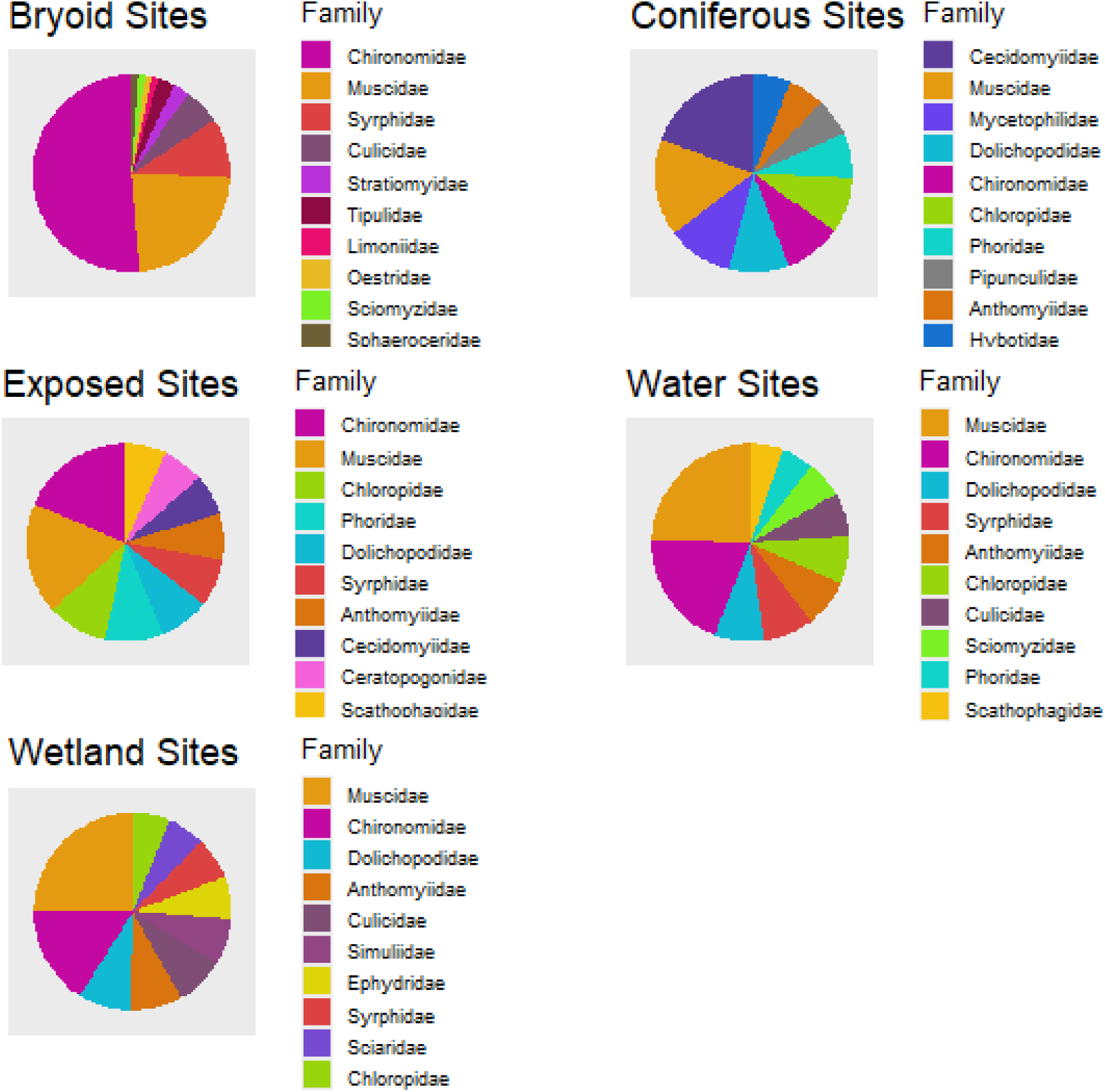
The top 10 families for each land type classification in our dataset DS-CDIPTERA. Each pie diagram depicts the number of specimens in the top 10 most represented families in that site classification by specimen number, and the order of the associated legend proceeds from the most to least represented. Portions appear in each pie diagram from most to least represented in a counter-clockwise manner.

## Discussion

### New Reference Library for Subarctic Diptera

In this study, we present a DNA barcode reference library containing 16786 records for 1211 named species and 2235 BINs of Diptera collected near Churchill, Manitoba, Canada. The library includes representatives of 68 families, with Muscidae being most represented in our dataset. All sequences and metadata are easily accessed as they are assembled into a single dataset on BOLD (DS-CDIPTERA, dx.doi.org/10.5883/DS-CDIPTERA). Through this study, we show the current diversity and taxonomic composition of Diptera within Churchill and provide opportunities for further research into this community using the barcode information provided.

One hundred and forty-two BINs were unique to this project, i.e. new to BOLD. Most unique BINs showed low representation in this dataset, containing fewer than 10 records, except for one BIN identified only to the family level as Ephydridae, which contained 32 records, and the BIN identified as Stictochironomus unguiculatus, which contained 25 records. The unique BINs represented 25 families, and within these, species of the families Cecidomyiidae and Chironomidae were most represented, with 38 and 29 unique BINs recorded. Almost all of the Cecidomyiidae BINs were only identified to family and reflected the taxonomic challenge that this family represents and the limited expertise available to identify the specimens. Cecidomyiidae and Chironomidae differ in habitat, with Cecidomyiidae having mostly terrestrial larvae, and Chironomidae mostly aquatic (Gagné, 1981, Oliver, 1981). Regarding diet, Cecidomyiidae larvae are often phytophagous or mycophagous, and frequently form galls on plants, while Chironomidae larvae are mostly microphagous, feeding on small particles of detritus, plants, and microorganisms (Gagné, 1981, Oliver, 1981). Chironomidae was the best-sampled family in our study, and it is possible that with increased sampling and research, more unique BINs could be discovered for other families.

### Geographic Affiliation of Churchill Diptera

Most of the BINs (2096 of 2225) were not unique to this study but contained members from other studies of Churchill and other geographic regions. Yukon (1404, 20.2% of Yukon, 63.1% of Churchill) and Alberta (1255, 15.7% of Alberta, 56.4% of Churchill) shared the most BINs with Churchill when looking at total counts; and when calculating beta diversity, Churchill BIN composition was most similar to Alberta and the Northwest Territories. Some BINs were also shared with the northern Palearctic region, such as Finland (440, 6.5% of Finland, 19.8% of Churchill) and Norway (417, 6.7% of Norway, 18.7% of Churchill), reflecting broader polar distributions. Churchill shared more BINs with southern regions when looking at total BIN counts (5192 vs 4170 BINs), but a high northern affinity was observed when calculating beta diversity. The beta diversity results, which consider total richness, show that Churchill is most similar to other northern regions in Canada, as well as nearby southern regions, and is more similar to distant northern regions, such as Finland and Norway, than some Canadian southern regions, such as Ontario. The similarity to other northern regions, such as the Northwest Territories, may reflect post-glacial dispersal from the Beringia glacial refuge, while similarity to nearby southern regions may reflect northward colonization following deglaciation of the Churchill region and may include ongoing north-south dispersal. The similarity in BIN composition across northern regions has been documented in other studies, such as Ekrem et al. (2018), who compared the species and BIN composition of Greenland Chironomidae to other Nearctic and Palearctic regions. They found that most taxa in Greenland were shared with at least one other geographic region, and that, when looking at taxon counts, Greenland was more similar to northern North America than to Europe (Ekrem et al., 2018). While Greenland, a harsh environment with lowest richness (338 species) among the regions examined, was one of the least-similar regions compared to Churchill in our study (Bray-Curtis = 0.965), 95 BINs out of 338 total (28% for Greenland, 4.2% for Churchill) in Greenland were still shared between them, and the overall sharing of dipteran BINs across Arctic regions here augments the findings for chironomids from Ekrem et al. (2018).

### Genetic Divergence Patterns across Studies and Taxa

Based on the intraspecific distances, most species can be easily distinguished from their nearest neighbours using COI. For the dataset (DS-CDIPTERA), across all Diptera families, the maximum intraspecific distances ranged from 0-16.06% and the nearest neighbour distances ranged from 0–16%. The high distance values, particularly those over 5% on the basis of other studies (e.g. Lin et al., 2015), are candidates for examination for cryptic species, particularly in understudied taxa. These distance values could also be high due to the biology of the species. The distances within individual families varied. For example, the mean intraspecific divergence was 0.06% in Tachinidae, with a nearest neighbour distance of 6.97%. This value is comparable to previous research by Pohjioismaki et al. (2016), who determined that the mean nearest neighbour distance within the family Tachinidae was 5.52%. The maximum intraspecific distance for Drosophilidae ranged from 0–2.47%, similar to Machado et al. (2017) who found that the intraspecific distances ranged from 0–5.3%. The maximum intraspecific distances for Muscidae ranged from 0–4.25% here, similar to the results of Renaud et al. (2012b), who found distances of 0–3.01% for Churchill Muscidae, though there is some dataset overlap between our studies. Variation in intraspecific divergence may be associated with effective population size differences between taxa, where shifts between aquatic-terrestrial, and flighted-flightless lifestyles have resulted in significant increases in the non-synonymous-to-synonymous mitochondrial substitution rate in a variety of insect orders (Mitterboeck et al., 2013, 2016).

Some of the differences between our study and literature could be due to the geographic scope of the studies. The intraspecific divergence range can differ based on the area included in the study. Lin et al. (2015) found an average intraspecific divergence of 2.14% in a global scale study of Chironomidae, with maximum divergences reaching 21.1% due to cryptic species complexes. This is similar to our study, which found a maximum intraspecific divergence of 16.06%, but a much lower average of 1.56%. When focusing on the Baltic Sea region, Brodin et al. (2012) found that Chironomidae had intraspecific distances ranging from 0– 3.15%. For Psychodidae, the distances ranged from 0– 4.9% within the family in Brazil and increased to 0– 8.32% when including multiple countries in the Neotropical region (de Souza Pinto et al., 2022, Posada-Lopez et al., 2023). When focusing only on Mexico, Adeniran et al. (2019) found intraspecific distances of 0.12–1.22%. This range increased to 0–9.48% when including other sequences from North and South America (Adeniran et al., 2019). The maximum intraspecific distance was 1.07% for Psychodidae in our study, which is similar to other studies that focused on small geographic regions. Tabanidae has also been studied across several different countries with varying intraspecific distances, such as Thailand (0–4.4%), Africa (0–6.5%), India (0–6.34%), and Canada (0–3.3%) (Banerjee et al., 2015, Changbunjong et al., 2018, Cywinska et al., 2010, Votýpka et al., 2019). Our results were similar to previous Tabanidae results from Canada (Cywinska et al., 2010), with maximum intraspecific distances that ranged from 0–2.5%.

A family where our results differed from the literature was Mycetophilidae, which had intraspecific distances of 0–13.46% in our dataset. Jürgenstein et al. (2015) conducted a study of Mycetophila ruficollis Meigen using 116 specimens collected throughout Europe from 1984 to 2014. Their study revealed intraspecific distances with a range of 0.08–0.8% (Jürgenstein et al., 2015), which is much lower than that of our study. The average intraspecific distance from our study, 0.7%, aligned more with Jürgenstein et al. (2015). Similarly, Kjærandsen (2022) found a mean intraspecific distance of 0.81% for Sciaroidea. This suggests that while the average genetic diversity in this family is low, there are some potential cryptic species to consider.

Nearest neighbour distances and intraspecific distances vary across insect orders. Lepidoptera has been shown to have lower levels of intraspecific variation than Diptera, with average intraspecific distances of 1.08% being found in species from Finland and Austria (Huemer et al., 2014), divergences ranging from 0-5.92% found in species from Nigeria (Nneji et al., 2020), and 0.43% in species from eastern North America, including Churchill (Hebert et al., 2009). Alos, a threshold of <2% was found to be appropriate to differentiate most Canadian Noctuoidea species, though 7.7% of species had deeper divergences (Zahiri et al., 2014). Ephemeroptera, Plecoptera and Trichoptera of Churchill were found to have average intraspecific distances of 0.31%, and average nearest neighbour distances of 13.09% (Zhou et al., 2009). The mean divergence between Nordic and Baltic species of Lepidoptera was reported as 5.73% (Pentinsaari, 2016) and between Coleoptera was 11.99% for the same regions (Pentinsaari et al., 2014). The mean intraspecific divergence and mean nearest neighbour distance reported in this study was 0.91% and 7.22%, respectively. Though Lepidoptera has been reported to be lower than Diptera in the literature, the mean intraspecific distance in our study is comparable to that seen in Lepidoptera. The mean nearest neighbour distance, however, appears to be lower in Lepidoptera. Both nearest neighbour and intraspecific distances appear to be higher in Coleoptera than Diptera, which aligns with what has been previously seen in the literature (Pentinsaari et al., 2014).

### Taxonomic Composition by Habitat

We found a strong presence of families associated with aquatic habitats in the top 10 families for each land type classification, from aquatic larvae (e.g. Chironomidae, Culicidae, Simuliidae, Tipulidae) to predatory adults feeding near water (e.g. Dolichopodidae, Empididae) (Cook, 1981, Oliver, 1981, Stone, 1981, Peterson, 1981, Alexander & Byers, 1981, Robinson & Vockeroth, 1981, Steyskal & Knutson, 1981). This is not surprising considering that Churchill is located on the Hudson’s Bay coast and is surrounded by diverse water body types like the Churchill River and a plethora of ponds and lakes. Despite the high occurrence of water in the area, shore flies (Ephydridae), which can be found around coastal areas, maritime marshes and even salt lakes, only made the top 10 in the wetlands habitat type. Furthermore, fungus gnats (Mycetophilidae, Sciaridae) were only abundant in the coniferous and wetlands sites. As their larvae are typically fungivores, we can suspect a moist forest with abundant bracket fungi among others. Overall, the top 10 families are similar across the different habitats (with the exception of bryoids), which can suggest that habitat patchiness and edge effects may act to homogenize the family composition detected using our trapping methods in each habitat type.

### Findings for Further Investigation

Several of the findings outlined above could be elaborated upon in future studies, such as the comparison between Churchill and other geographic regions or the differences in Diptera community composition across different habitat types in the Churchill region. Statistically significant differences in composition across land-use categories were not evaluated here, as samples were collected over multiple seasons by a variety of people using different sampling methods and approaches as to what was sampled and later sequenced. We therefore used a graphical approach to explore the biodiversity across Churchill habitats. Future research could focus on comparing these different Churchill habitats with more consistent methods and timespan (such as specifically our repeated Malaise trap samples from consistent sites across the summer season in 2010) to determine if there are any significant differences in community composition.

A wide variety of collection methods were used to target a wider diversity of taxa and to obtain the specimens for this reference library. This resulted in a stronger reference library, and the benefits of using different trap types have been discussed in various studies (Ernst et al., 2015, Pentinsaari et al., 2020). However, we acknowledge there could still be biases and taxonomic groups potentially missed or underrepresented. Different families are caught most often using different methods. As Dipterans are strong flyers, most specimens were caught using Malaise traps (51.38% of sequenced specimens). Hence, they are commonly used to collect these taxa, but there can be some biases depending on the type of Malaise trap used, the material used to make the trap, and the type of attractant used (Betts, 2010, Martinez et al., 2020, Skvarla et al., 2020, Uhler et al., 2022). Despite the potential limitations of each, we advance the strategy of combining a variety of trap types and sampling methods, and field involvement of expert collectors, allowing us to construct a robust reference library for the region that is as thorough as feasible. Further research could be done to determine the effectiveness of trap type and their effect on species composition in Churchill.

While many specimens were collected, there are still groups missing from the barcode data. About 20% of the known Churchill species of Empidoidea (Freeman, 1959, Danks, 1981, Northern Biodiversity Program, n.d.) were not accounted for in the barcode data. It is possible that the habitats associated with these groups, such as running water habitats, were not well sampled, likely resulting in some species being missed. Future research could focus on the continued sampling of Churchill and other northern regions to further expand the DNA barcode data available to other researchers.

Further research could also be done regarding the long branch lengths observed in the ML tree. After following the steps outlined in the methods, we did not find any issues with the quality or identification of the sequences. These sequences translated correctly without stop codons, passed other quality control steps, and when BLASTed, returned the expected results and were similar to other sequences within the species. For Tetanocera fuscinervis (Zetterstedt), all codon positions showed the long branch length, but for Bradysiopsis vittigera (Zetterstedt), the long branch length only appeared when using the third codon position and could suggest a more rapid rate of evolution. Overall, the branch lengths do not appear to be caused by an issue with the sequences or alignment, but instead are something to do with the species itself. Tetanocera fuscinervis is a member of the family Sciomyzidae and is a predator of snails on shorelines (Foote, 1996). There is little information readily available regarding Bradysiopsis vittigera, but this species is a member of the family Sciaridae, where many larvae are saprophagous (Scudder & Cannings, 2006, Steffan, 1981). These species are the only members of their families on the tree. There is possibly something about these species and families that results in the long branch lengths and genetic differences. Further research could look into the reasons for these branch lengths and the relationships between these species and other Diptera, and to see if other molecular markers recapitulate these distances. To look more closely at the phylogenetic relationships between the taxa recorded in Churchill, a range of nuclear and mitochondrial markers would need to be combined with morphological characters in order to get a more accurate picture, such as the phylogeny presented in Wiegmann et al. (2011), which uses nuclear loci, mitochondrial genomes, and morphological characters.

### Opportunities for Using New Library

As outlined earlier in the discussion, there are differences across and within insect orders regarding the intraspecific divergence. The families within Diptera differed in their intraspecific distances, and these differences may be considered when clustering specimens into species. A single clustering threshold is not optimal for all Diptera when performing de novo clustering of sequences into taxonomic units as close to species as possible, but rather this threshold should differ based on the genetic diversity present in the family. If a threshold is used that is too large, this could result in closely related species being grouped together, or if one is too small, this could result in species being split across operational taxonomic units. These differences can impact estimates of species richness. However, through this collaboration between molecular analysts and taxonomic experts, researchers are not solely reliant on specific divergence thresholds. By using well-populated reference libraries, future researchers can perform identifications despite variation in genetic diversity within and among species. When sorting sequenced specimens into species, we recommend accessing reference libraries when possible.

References libraries are valuable resources that can help researchers more accurately identify and delineate species and provide opportunities for future research. The barcode library presented in this study reflects the current diversity of Diptera in Churchill, and the barcode sequences are publicly available on BOLD and accompanied by their voucher, taxonomic, and collection information. This library is an important resource for any researcher in need of species-level information. Taxonomic experts identified the specimens, and 12719 specimens were assigned species-level identification. While molecular information and BINs are an important resource, BINs do not always correspond to species. By combining molecular data with taxonomic identification, we can reduce the number of specimens that are misidentified and provide more accurate and detailed information to future researchers. Trait information is usually associated with species names, so having this taxonomic information available allows future researchers to make use of the molecular data to a greater extent. Future research could include comparing our reference library to past Churchill studies and future studies to see how these communities are changing. This region can also be compared to other northern regions, such as those mentioned above. Overall, this barcode library can be applied to a wide range of studies investigating this northern region and the ecology and evolution of Diptera. However, further work is needed, and future studies could focus on a larger geographic scale, and include other northern regions, as well as temperate and tropical regions, so we can further our knowledge of Diptera diversity and barcode variability at a global scale.

## Conclusion

In conclusion, we present a large-scale barcoding study of Diptera from Churchill and contribute to the growing amount of barcode data available for research. We provide a detailed reference library, making use of molecular data and taxonomic knowledge, and make this library publicly available for future research. Our work demonstrates the utility of broad, multi-year collecting programs as well as collaborations to build well-populated DNA barcode reference libraries that will enable identification at the species level even for hyper-diverse arthropod groups. We outline the diversity of Diptera and contribute to the knowledge of this ecologically important group, including taxa that interact closely with humans and those that can be used for freshwater ecosystem health monitoring. By studying northern Diptera communities and understanding current diversity, we can study the change over time, especially as northern regions are impacted by the changing climate. By collecting and releasing barcode libraries we open the door for future research and allow other researchers to build off our work and further investigate taxa and regions of interest.

## Supporting information

The primers used for the amplification of the DNA barcode region of COI are included in Supplementary File 2.

The full dataset including BINs, taxonomic assignment, voucher storage information, and sequences is available in Supplementary File 3.

A list of BINs shared between Churchill and each region is available in Supplementary File 4.

A table containing the specimen count/site along with their respective EOSD classification is included in Supplementary File 5.

A phenogram showing the relationship between the taxa collected in this study shows that COI sequences within families generally group together for th

The full results of the barcode gap analysis are available in Supplementary File 7.

Sampling intensity varied across years, with the most sampling occurring in 2010 and the least occurring in 2011 (Supplementary File 1

## Acknowledgements

We dedicate this paper to Terry Wheeler, who was an Associate Professor and Director of the Lyman Entomological Museum, McGill University, and to Rob Roughley, who was a beetle specialist from the University of Manitoba. Terry was an excellent scientist, educator, mentor, and friend to many who contributed significantly to this research before he passed away. Rob was a major partner in the overall insect biodiversity survey of Churchill, and he contributed field work and important specimens to the present paper. The authors are also grateful to the staff of the Churchill Northern Studies Centre, with special thanks to LeeAnn Fishback for supporting this project. Colleagues at the Centre for Biodiversity Genomics, University of Guelph, provided critical assistance in field collecting and in conducting the DNA sequencing work. We are grateful to Anais Renaud, Jostein Kjaerandsen, Terry L. Whitworth, Kai Heller, Norm E. Woodley, Kevin Barber, Brian V. Brown, Art Borkent, Ludovic Jolicoeur, Mikko Pentinsaari, Ayman K. Elsayed, Jeremy deWaard, Jukka E. Salmela, Michelle Locke, Ole A. Saether, Alyssa M. Anderson, Michel Rapinski, Xiaolong Lin, Susan E. Gresens, Lucas E. Roscoe, and Jin-hua Ran for conducting additional specimen identifications. Thank you to Angela Telfer, Kate Perez, Allison Brown, Renee Miskie, and staff at the Centre for Biodiversity and Genomics for adding barcode-based IDs to portions of the data. Thank you to Peter Kevan for contributions to the field collections. Thank you to Jiaojiao Diao and Jessica Castellanos-Labarcena for assisting with assigning land-use classification to sites and generating figures. This work was conducted under permits issued by the Manitoba Conservation Wildlife and Ecosystem Protection Branch from Winnipeg, MB to the Churchill Northern Studies Centre (CNSC) for conducting research in the Churchill Wildlife Management Area. Funding for this work was provided by a Natural Sciences and Engineering Research Council of Canada (NSERC) Research Network grant and a NSERC International Polar Year (IPY) grant to PDNH, NSERC Discovery Grants to SJA, grants in Genomics and Bioinformatics and Computational Biology from Genome Canada through the Ontario Genomics Institute and from the Government of Ontario to SJA and PDNH, by the Government of Canada through the Canada First Research Excellence Fund to SJA and PDNH, and through a NSERC Strategic Project Grant (Ecological Structure of Northern Arthropods: Adaptation to a Changing Environment) awarded to C.M. Buddle, TAW and DCC, plus their supporting partners and collaborators.

## Data and Accessibility and Benefit-Sharing Section

### Data Accessibility Statement

The DNA barcode reference library is publicly available on the Barcode of Life Data Systems in the dataset DS-CDIPTERA (dx.doi.org/10.5883/DS-CDIPTERA) (Majoros et al., 2023). The sequences are also available on GenBank. A list of process IDs and GenBank accessions included in the dataset is available in supplementary file 3.

### Benefit-Sharing Statement

Benefits from this research accrue from the public sharing of the DNA barcode reference library on the Barcode of Life Data Systems.

### Author Contributions

All authors reviewed and edited the manuscript. SEM, TAE and SJA wrote the manuscript. SEM,TAE and SJA designed the analyses, while SEM and TAE performed the analyses and created the figures. FB and CY identified crane flies from the families Cylindrotomidae, Limoniidae, Pediciidae and Tipulidae. JW conducted field work, identified specimens, processed the specimens for barcoding, did family-level determinations, and species-level determinations for Chloropidae. VLB did family-level determinations, co-ordination of sending representative vouchers to collaborators for determination, and contributed to manuscript writing. CJB identified the Mycetophilidae specimens. SEB, JMC and BJS identified the Empidoidea (Empididae and Dolichopodidae) specimens. SB identified the Agromyzidae specimens. DCC and PS did collecting and determinations for Simuliidae, Culicidae, and Tabanidae. TE and ES did collecting and determinations for Chironomidae. PDNH and RER contributed to the collecting and conception of the project. JS did Muscidae determinations for the 2010 clusters and the Anthomyiidae determinations. JHS did Syrphidae and Pipunculidae collecting and determinations. AMS conducted field work and specimen identifications. TAW did the Chloropidae, Piophilidae and Heleomyzidae identifications. TSW contributed to collecting and sorting the malaise traps to family. ADY identified Syrphidae specimens.

## References

1. Adeniran, A.A., Fernández-Santos, N.A., Rodríguez-Rojas, J.J., Treviño-Garza, N., Huerta-Jiménez, H., Mis-Ávila, P.C., Pérez-Pech, W.A., Hernández-Triana, L.M. & Rodríguez-Pérez, M.A. (2019). Identification of phlebotomine sand flies (Diptera: Psychodidae) from leishmaniasis endemic areas in southeastern Mexico using DNA barcoding. Ecology and Evolution, 9(23), 13543–13554. doi: 10.1002/ece3.5811

2. Adler, P.H. & Courtney, G.W. (2019). Ecological and societal services of aquatic Diptera. Insects, 10(3), 70. doi: 10.3390/insects10030070

3. Albrecht, A. (1990). Revision, phylogeny and classification of the genus Dorylomorpha (Diptera, Pipunculidae). Acta Zoologica Fennica, 188, 1–240.

4. Aldrich, J.M. (1918). The kelp flies of North America. Proceedings of the California Academy of Sciences, 8, 157–179.

5. Alexander, C.P. (1943). Family Tipulidae. Pp. 196–486 In Crampton, G.C. et al. (Eds.) Guide to the insects of Connecticut. Part VI. The Diptera or true flies of Connecticut. First Fascicle. Bulletin Connecticut State Geological and Natural History Survey, 64.

6. Alexander, C.P. & Byers, G.W. (1981). Tipulidae. In McAlpine, J.F., Peterson, B.V., Shewell, G.E., Teskey, H.J., Vockeroth, J.R. & Wood, D.M. (Eds), Manual of Nearctic Diptera. Volume 1. (pp. 247–256). Research Branch Agriculture Canada.

7. Altschul, S.F., Gish, W., Miller, W., Myers, E.W. & Lipman, D.J. (1990). Basic local alignment search tool. Journal of Molecular Biology, 215(3), 403–410. doi: 10.1016/S0022-2836(05)80360-2

8. Anderson, A. M., Stur, E., & Ekrem, T. (2013). Molecular and morphological methods reveal cryptic diversity and three new species of Nearctic Micropsectra (Diptera: Chironomidae). Freshwater Science, 32(3), 892–921. doi: 10.1899/12-026.1

9. Appelhans, T., Detsch, F., Reudenbach, C. & Woellauer, S. (2023). mapview: Interactive Viewing of Spatial Data in R. R package version 2.11. https://github.com/r-spatial/mapview.

10. Arntfield, P.W. (1975). A revision of Graphomya Robineau-Desvoidy (Diptera: Muscidae) from North America. The Canadian Entomologist, 107(3), 257–302. doi: 10.4039/Ent107257-3

11. Banerjee, D., Kumar, V., Maity, A., Ghosh, B., Tyagi, K., Singha, D., Kundu, S., Laskar, B.A., Naskar, A. & Rath, S. (2015). Identification through DNA barcoding of Tabanidae (Diptera) vectors of surra disease in India. Acta Tropica, 150, 52–58. doi: 10.1016/j.actatropica.2015.06.023

12. Barták, M. (2002). Nearctic species of Rhamphomyia subgenus Megacyttarus (Diptera: Empididae). Acta Universitatis Carolinae, Biologica, 46, 3–215.

13. Bedini, S., Flamini, G., Cosci, F., Ascrizzi, R., Echeverria, M.C., Guidi, L., Landi, M., Lucchi, A. & Conti, B. (2017). Artemisia spp. essential oils against the disease-carrying blowfly Calliphora vomitoria. Parasites & Vectors, 10(1), 80. doi: 10.1186/s13071-017-2006-y

14. Betts, D.J. (2010). Investigation of potential trapping bias in Malaise traps due to mesh gauge, in two Habitats [Master’s Thesis, Brigham Young University]. https://scholarsarchive.byu.edu/etd/2561

15. Blagoev, G.A., Nikolova, N.I., Sobel, C.N., Hebert, P.D.N. & Adamowicz, S.J. (2013). Spiders (Araneae) of Churchill, Manitoba: DNA barcodes and morphology reveal high species diversity and new Canadian records. BMC Ecology, 13(1), 44. doi: 10.1186/1472-6785-13-44

16. Boesel, M. W. (1985). A brief review if the genus Polypedilum in Ohio, with keys to known stages of species occurring in northeastern United States (Diptera, Chironomidae). Ohio Journal of Science, 85(5), 245–262.

17. Bonizzoni, M., Gasperi, G., Chen, X. & James, A.A. (2013). The invasive mosquito species Aedes albopictus: current knowledge and future perspectives. Trends in Parasitology, 29(9), 460–468. doi: 10.1016/j.pt.2013.07.003

18. Boucher, S. (2002). Revision of Nearctic species of Cerodontha (Cerodontha) (Diptera: Agromyzidae). The Canadian Entomologist 134, 577–603

19. Boucher, S. (2008). A new species and taxonomic notes on northern Nearctic Cerodontha (Icteromyza) (Diptera: Agromyzidae). The Canadian Entomologist 140, 557–562

20. Boucher, S. (2012a). Revision of the Nearctic species of Cerodontha (Icteromyza) (Diptera: Agromyzidae). The Canadian Entomologist 144, 122–157

21. Boucher, S. (2012b). Revision of the Canadian species of Amauromyza Hendel (Diptera: Agromyzidae). The Canadian Entomologist 144, 733–757

22. Bragard, C., Dehnen-Schmutz, K., Di Serio, F., Gonthier, P., Jacques, M., Jaques Miret, J..A., Justesen, A.F., Magnusson, C.S., Milonas, P., Navas-Cortes, J.A., Parnell, S., Potting, R., Reignault, P.L., Thulke, H., Van der Werf, W., Vicent Civera, A., Yuen, J., Zappalà, L., Czwienczek, E., Streissl, F. & MacLeod, A. (2020). Pest categorisation of Liriomyza bryoniae. EFSA Journal, 18 (3), p.e06038-n/a. doi: 10.2903/j.efsa.2020.6038

23. Brodin, Y., Ejdung, G., Strandberg, J., Lyrholm, T. (2012). Improving environmental and biodiversity monitoring in the Baltic Sea using DNA barcoding of Chironomidae (Diptera). Molecular Ecology Resources, 13(6), 996–1004. doi: 10.1111/1755-0998.12053

24. Brodo, F. (1967). A review of the subfamily Cylindrotominae in North America (Diptera, Tipulidae). The University of Kansas Science Bulletin, 47, 71–115.

25. Brodo, F. (1987). A revision of the genus Prionocera (Diptera: Tipulidae). Evolutionary Monograph, 8, 1–93.

26. Brodo, F. (2017). Taxonomic review of Angarotipula Savchenko, 1961 (Diptera: Tipulidae) in North America. Canadian Entomologist, 150, 12–34.

27. Brundin, L. (1947) Zur Kenntnis der schwedischen Chironomiden. Arkiv för zoologi, 39 A(3), 1–95.

28. Brundin, L. (1949) Chironomiden und andere Bodentiere der südschwedischen Urgebirgsseen. Ein Beitrag zur Kenntnis der bodenfaunistischen Charakterzüge schwedischer oligotropher Seen. Report/Institute of Freshwater Research, Drottningholm, 30, 915.

29. Brundin, L. (1989) The adult males of Podonominae (Diptera: Chironomidae) of the Holarctic region – Keys and diagnoses. In: Wiederholm, T. (ed.): Chironomidae of the Holarctic region. Keys and diagnoses. Part 3 – Adult males. Entomologica Scandinavica, Suppl., 34, 23–36.

30. Byers, G.W. (1961). The crane fly genus Dolichopeza in North America. The University of Kansas Science Bulletin, 42, 665–924.

31. Changbunjong, T., Bhusri, B., Sedwisai, P., Weluwanarak, T., Nitiyamatawat, E., Chareonviriyaphap, T. & Ruangsittichai, J. (2018). Species identification of horse flies (Diptera: Tabanidae) in Thailand using DNA barcoding. Veterinary Parasitology, 259, 35–43. doi: 10.1016/j.vetpar.2018.07.002

32. Chao, A. (1984). Nonparametric estimation of the number of classes in a population. Scandinavian Journal of Statistics, 11(4), 265–270.

33. Chao, A., Gotelli, N.J., Hsieh, T.C., Sander, E.L., Ma, K.H., Colwell, R.K. & Ellison, A.M. (2014). Rarefaction and extrapolation with Hill numbers: a framework for sampling and estimation in species diversity studies. Ecological Monographs, 84(1), 45–67. doi: 10.1890/13-0133.1

34. Chillcott, J.G. (1962). A revision of the Platypalpus juvenis complex in North America (Diptera: Empididae). The Canadian Entomologist, 94(2), 113–143. doi: 0.4039/Ent94113-2

35. Churchill Northern Studies Centre. (2022). The Centre. Churchill Northern Studies Centre. https://churchillscience.ca/about/the-centre/

36. Cirtwill, A.R., Kaartinen, R., Rasmussen, C., Redr, D., Wirta, H., Olesen, J.M., Tiusanen, M., Ballantyne, G., Cunnold, H., Stone, G.N., Schmidt, N.M. & Roslin, T. (2022). Stable pollination service in a generalist high Arctic community despite the warming climate. Ecological Monographs, 93(1). doi: 10.1002/ecm.1551

37. Collin, J.E. (1930). A revision of the Greenland species of the anthomyid genus Limnophora sens. lat. (Diptera), with figures of the male genitalia of these and many other Palearctic species. Transactions of the Entomological Society of London 78(2), 255–281. doi: 10.1111/j.1365-2311.1930.tb00387.x

38. Collins, R.A., Boykin, L.M., Cruickshank, R.H. & Armstrong, K.F. (2012) Barccoding’s next top model: an evaluation of nucleotide substitution models for species identification. Methods in Ecology and Evolution, 3(2), 457–465

39. Cook, E.F. (1981). Chaoboridae. In McAlpine, J.F., Peterson, B.V., Shewell, G.E., Teskey, H.J., Vockeroth, J.R. & Wood, D.M. (Eds), Manual of Nearctic Diptera. Volume 1. (pp. 257–292). Research Branch Agriculture Canada.

40. Coquillett, D.W. (1900). Papers from the Harriman Alaska Expedition. IX. Entomological Results (3): Diptera. Proceedings of the Washington Academy of Sciences, 2, 389–464.

41. Cranston, P. S., Oliver, D. R. (1988) Additions and corrections to the Nearctic Orthocladiinae (Diptera: Chironomidae). Canadian entomologist, 120(5), 425–462. doi: 10.4039/Ent120425-5

42. Cranston, P. S., Dillon, M. E., Pinder, L. C. V., Reiss, F. (1989a). The adult males of Chironominae (Diptera: Chironomidae) of the Holarctic region – Keys and diagnoses. In: Wiederholm, T. (ed.): Chironomidae of the Holarctic region. Keys and diagnoses. Part 3 – Adult males. Entomologica Scandinavica, Suppl., 34, 353–502.

43. Cranston, P. S., Oliver, D. R., Sæther, O. A. (1989b) The adult males of Orthocladiinae (Diptera: Chironomidae) of the Holarctic region – Keys and diagnoses. – In: Wiederholm, T. (ed.): Chironomidae of the Holarctic region. Keys and diagnoses. Part 3 – Adult males. Entomologica Scandinavica, Suppl., 34, 165–352.

44. Cranston, P. (2008) Keys to the adult male Chironomidae of Britain and Ireland—edited by P.H. Langton and L.C.V. Pinder and a guide to the identification of genera of Chironomid Pupal Exuviae occurring in Britain and Ireland (including common genera from Northern Europe) and their use in monitoring Lotic and Lentic fresh waters—edited by R.S. Wilson and L.P. Ruse. Systematic Entomology, 33, 215–217

45. Cywinska, A., Hannan, M.A., Kevan, P.G., Roughley, R.E., Iranpour, M. & Hunter, F.F. (2010). Evaluation of DNA barcoding and identification of new haplomorphs in Canadian deerflies and horseflies. Medical and Veterinary Entomology, 24(4), 382–410. doi: 10.1111/j.1365-2915.2010.00896.x

46. Czechowski, P., Stevens, M.I., Madden, C. & Weinstein, P. (2020). Steps towards a more efficient use of chironomids as bioindicators for freshwater bioassessment: Exploiting eDNA and other genetic tools. Ecological Indicators, 110. doi: 10.1016/j.ecolind.2019.105868

47. Danks, H.V. (1981). Arctic arthropods, a review of systematics and ecology with particular reference to the North American fauna. Entomological Society of Canada.

48. De Meyer, M. (1989). Systematics of the Nearctic species of the genus Cephalops Fallén (Diptera, Pipunculidae). Bulletin de l’Institut Royal des Sciences Naturelles de Belgique, Entomologie, 59, 99–130.

49. de Souza Pinto, I., Rodrigues, B.L., de Araujo-Pereira, T., Shimabukuro, P.H.F., de Pita-Pereira, D., Britto, C. & Brazil, R.P. (2023). DNA barcoding of sand flies (Diptera, Psychodidae, Phlebotominae) from the western Brazilian Amazon. PLOS ONE, 18(2). doi: 10.1371/journal.pone.0281289

50. deWaard, J. (2023). Welcome to Arctic BIOSCAN (ARCBIO). Arctic BIOSCAN. https://arcticbioscan.ca/

51. Eddy, S.R. (1998). Profile hidden Markov models. Bioinformatics, 14(9), 755–763. doi: 10.1093/bioinformatics/14.9.755

52. Ekrem, T. (2007). A taxonomic revision of the genus Stempellinella (Diptera: Chironomidae). Journal of Natural History, 41(21-24), 1367–1465. doi: 10.1080/00222930701437360

53. Ekrem, T., Sublette, M. F., & Sublette, J. E. (2003). North American Tanytarsus I. Descriptions and keys to species in the eminulus, gregarius, lugens and mendax species groups (Diptera: Chironomidae). Annals of the Entomological Society of America, 96(3), 265–328. doi: 10.1603/0013-8746(2003)096[0265:NATIDA]2.0.CO;2

54. Ekrem, T., Stur, E., Orton, M.G. & Adamowicz, S.J. (2018). DNA barcode data reveal biogeographic trends in Arctic non-biting midges. Genome, 61(11), 787–796. doi: 10.1139/gen-2018-0100

55. Environmental Systems Research Institute. (2015). ArcGIS Desktop: Release 10.3 [Software]. Redlands, CA: Environmental Systems Research Institute.

56. Epler, J. H. (1987). Revision of the Nearctic Dicrotendipes Kieffer, 1913 (Diptera: Chironomidae). Evolutionary Monographs, 9, 1–102

57. Ernst, C.M., Loboda, S. & Buddle, C.M. (2015). Capturing northern biodiversity: diversity of arctic, Subarctic and north boreal beetles and spiders are affected by trap type and habitat. Insect Conservation and Diversity, 9(1), 63–73. doi: 10.1111/icad.12143

58. Fedoseeva, L.I. 1971. A revision of the Nearctic species of grass flies of the genus Meromyza Mg. (Diptera: Chloropidae). Entomological Review, 50(4): 520—529.

59. Fedoseeva, L.I. 2003 Chloropid flies of the genus Meromyza Meigen, 1830 (Diptera: Chloropidae) of the fauna of North America. Russian Entomological Journal, 12(4): 425—430.

60. Fernandez-Triana, J., Smith, M.A., Boudreault, C., Goulet, H., Hebert, P.D.N., Smith, A.C. & Roughley, R. (2011). A poorly known high-latitude parasitoid wasp community: unexpected diversity and dramatic changes through time. PLOS ONE, 6(8). doi: 10.1371/journal.pone.0023719

61. Ferrington, L. C. Jr & Sæther, O. A. (2011) A revision of the genera Pseudosmittia Edwards, 1932, Allocladius Kieffer, 1913, and Hydrosmittia gen. n. (Diptera: Chironomidae, Orthocladiinae). Zootaxa, 2849(1), 1–314. doi: 10.11646/zootaxa.2849.1.1

62. Firke, S. (2023). janitor: Simple Tools for Examining and Cleaning Dirty Data. R package version 2.2. https://github.com/sfirke/janitor

63. Fittkau, E. J. (1962) Die Tanypodinae (Diptera, Chironomidae). Die Tribus Anatopyniini, Macropeloplini und Pentaneurini. Abh. Larvalsyst. Insekten, 6, 1–453.

64. Fittkau, E. J. & Lehmann, J. (1970) Revision der Gattung Microcricotopus Thien. u. Harn. (Dipt., Chironomidae). Internationale Revue der gesamten Hydrobiologie, 55(3), 391–402.

65. Floyd R, Abebe E, Papert A, Blaxter M (2002) Molecular barcodes for soil nematode identification. Molecular Ecology 11:839–850. 10.1046/j.1365-294X.2002.01485.x

66. Folmer, O., Black, M., Hoeh, W., Lutz, R. & Vrijenhoek, R. (1994). DNA primers for amplification of mitochondrial cytochrome c oxidase subunit I from diverse metazoan invertebrates. Mitochondrial DNA. Part B. Resources, 5(2), 1622–1624. doi: 10.1080/23802359.2020.1745100

67. Foote, B.A. (1996). Biology and immature stages of snail-killing flies belonging to the genus Tetanocera (Insecta: Diptera: Sciomyzidae). 1. Introduction and life histories of predators of shoreline snails. Annals of the Carnegie Museum, 65, 1–12. doi: 10.5962/p.226629

68. Förster, M., Klimpel, S., Mehlhorn, H., Sievert, K., Messler, S. & Pfeffer, K. (2007). Pilot study on synanthropic flies (e.g. Musca, Sarcophaga, Calliphora, Fannia, Lucilia, Stomoxys) as vectors of pathogenic microorganisms. Parasitology Research, 101(1), 243–246. doi: 10.1007/s00436-007-0522-y

69. Freeman, T.N. (1958). A historical account of insect collecting in northern Canada. Proceedings of the 10th International Congress of Entomology.

70. Freeman, T.N. (1959). The Canadian Northern Insect Survey, 1947-57. Polar Record, 9(61), 299–307. doi: 10.1017/S0032247400066006

71. Fu, Y. & Sæther, O. A. (2012) Corynoneura Winnertz and Thienemanniella Kieffer from the Nearctic Region (Diptera: Chironomidae: Orthocladiinae). Zootaxa, 3536(1), 1–61. doi: 10.11646/zootaxa.3536.1.1

72. Gagné, R.J. (1981). Cecidomyiidae. In McAlpine, J.F., Peterson, B.V., Shewell, G.E., Teskey, H.J., Vockeroth, J.R. & Wood, D.M. (Eds), Manual of Nearctic Diptera. Volume 1. (pp. 257–292). Research Branch Agriculture Canada.

73. Garnier, S., Ross, N., Rudis, R., Camargo, P.A., Sciaini, M. & Scherer, C. (2023). Viridis: Colorblind-Friendly Color Maps for R. R package version 0.6.3. https://sjmgarnier.github.io/viridis/

74. Giłka, W. (2001). A review of Polish Cladotanytarsus Kieffer (Diptera: Chironomidae) with description of three new species. Polish Journal of Entomology, 70, 307–328.

75. Gowin, F. & Thienemann, A. (1942) Zwei neue Orthocladiinen-Arten aus Lunz (Niederdonau). Chironomiden aus dem Lunzer Seengebiet VII. Zoologischer Anzeiger, 140(7/8), 101–109.

76. Graczyk, T.K., Knight, R., Gilman, R.H. & Cranfield, M.R. (2001). The role of non-biting flies in the epidemiology of human infectious diseases. Microbes and Infection, 3(3), 231–235. doi: 10.1016/S1286-4579(01)01371-5

77. Gregor, F., Rozkošný, R., Barták, M. & Vaňhara, J. (2002). The Muscidae (Diptera) of Central Europe. Folia Facultatis Scientiarium Naturalium Universitatis Masarykianae Brunensis. Biol. 107, 1–280.

78. Griffiths, G.C.D. (1991a). Anthomyiidae [part]. In Griffiths, G.C.D. (Eds), Flies of the Nearctic Region, (pp. 1049–1240). E. Schweizerbart, Stuttgart, Germany.

79. Griffiths, G.C.D. (1991b). Anthomyiidae [part]. In Griffiths, G.C.D. (Eds), Flies of the Nearctic Region, (pp. 1633–1872). E. Schweizerbart, Stuttgart, Germany.

80. Griffiths, G.C.D. (1991c). Anthomyiidae [part]. In Griffiths, G.C.D. (Eds), Flies of the Nearctic Region, (pp. 2289–2484). E. Schweizerbart, Stuttgart, Germany.

81. Griffiths, G.C.D. (1993a). Anthomyiidae [part]. In Griffiths, G.C.D. (Eds), Flies of the Nearctic Region, (pp. 1417–1632). E. Schweizerbart, Stuttgart, Germany.

82. Griffiths, G.C.D. (1993b). Anthomyiidae [part]. In Griffiths, G.C.D. (Eds), Flies of the Nearctic Region, (pp. 1873–2120). E. Schweizerbart, Stuttgart, Germany.

83. Griffiths, G.C.D. (1964). The Agromyzid fauna of Iceland and the Faroes, with Appendices on the Phytomyza millii and robustella Groups. Entomologiske Meddelelser 32, 393–450

84. Grodhaus, G. (1987). Endochironomus Kieffer, Tribelos Townes, Synendotendipes, n. gen., and Endotribelos, n. gen. (Diptera: Chironomidae) of the Nearctic Region. Journal of the Kansas Entomological Society, 60(2), 167–247.

85. Hajibabaei, M., DeWaard, J.R., Ivanova, N.V., Ratnasingham, S., Dooh, R.T., Kirk, S.L., Mackie, P.M. & Hebert, P.D.N. (2005). Critical factors for assembling a high volume of DNA Barcodes. Philosophical Transactions of the Royal Society B: Biological Sciences, 360(1462), 1959–1967. doi: 10.1098/rstb.2005.1727

86. Hajibabaei, M., Janzen, D.H., Burns, J.M., Hallwachs, W. & Hebert, P.D.N. (2006). DNA barcodes distinguish species of tropical Lepidoptera. Proceedings of the National Academy of Sciences, 103(4), 968–971. doi: 10.1073/pnas.0510466103

87. Hamerlík, L. & Bitušík, P. (2009). The distribution of littoral chironomids along an altitudinal gradient in High Tatra Mountain lakes: Could they be used as indicators of climate change? Annales de Limnologie, 45(3), 145–156. doi: 10.1051/limn/2009021

88. Hardy, D.E. (1943). A revision of Nearctic Dorilaidae (Pipunculidae). University of Kansas Science Bulletin, 29, 1–231.

89. Hebert, P.D.N., Cywinska, A., Ball, S.L. & DeWaard, J.R. (2003). Biological identifications through DNA barcodes. Proceedings of the Royal Society B: Biological Sciences, 270(1512), 313– 21. doi: 10.1098/rspb.2002.2218

90. Hebert, P.D.N., Stoeckle, M.Y., Zemlak, T.S. & Francis, C.M. (2004). Identification of birds through DNA barcodes. PLOS Biology, 2(10). doi: 10.1371/journal.pbio.0020312

91. Hebert, P.D.N., deWaard, J.R. & Landry, J.-F. (2009) DNA barcodes of 1/1000 of the animal kingdom. Biology Letters, 6(3). doi: 10.1098/rsbl.2009.0848

92. Hennig, W. (1955–1964). Muscidae. In E. Lindner (Eds.), Die Fliegen der palaearktischen Region 63b 7(2). (pp. 1–1110). Schweizerbart’sche Verlagsbuchhandlung, Stuttgart.

93. Hestenes, T. C. & Sæther, O. A. (2000) Three new Nearctic Thienemanniella Kieffer species with a review of the Nearctic species. In: Hoffrichter, O. (ed.): Late 20th Century Research on Chironomidae: an Anthology from the 13th International Symposium on Chironomidae, pp. 103–127. Shaker Verl., Aachen.

94. Hirvenoja, M. (1962) Cladotanytarsus-Arten (Dipt., Chironomidae) aus Finnisch-Lappland. Annales Entomologici Fennici, 28(4), 173–181.

95. Hirvenoja, M. (1973) Revision der Gattung Cricotopus van der Wulp und ihrer Verwandten (Diptera, Chironomidae). Annales Zoologici Fennici, 10(1), 1–363.

96. Hirvenoja, M. & Hirvenoja, E. (1988) Corynoneura brundini spec. nov. Ein Beitrag zur Systematik der Gattung Corynoneura (Diptera, Chironomidae). In: Fittkau, E. J. (ed.): Festschrift zu Ehren von Lars Brundin. Spixiana Suppl., 14, 213–238.

97. Hocking, B., Richards, W.R. & Twinn, C.R. (1950). Observations on the bionomics of some northern mosquito species (Culicidae: Diptera). Canadian Journal of Research, 28(2), 58–80. doi: 10.1139/cjr50d-006

98. Huckett, H.C. (1932). The North American species of the genus Limnophora Robineau-Desvoidy with descriptions of new species (Muscidae, Diptera). Journal of the New York Entomological Society, 40(3), 25–76, 107-158, 279-339.

99. Huckett, H.C (1934a). Revision of the North American species belonging to the genus Coenosia Meigen and related genera (Diptera: Muscidae). Part I. The subgenera Neodexiopsis, Coenosia, Hoplogaster and related genera Allognota, Bithoracochaeta and Schoenomyza. Transactions of the American Entomological Society, 60(1), 57–119.

100. Huckett, H.C. (1934b). Revision of the North American species belonging to the genus Coenosia Meigen and related genera. (Diptera: Muscidae). Part II. The subgenus Limosia (Coenosia of authors). Transactions of the American Entomological Society, 60(2), 133–198.

101. Huckett, H.C. (1936). A revision of connectant forms between coenosian and limnophorine genera occurring in North America (Diptera: Muscidae). Journal of the New York Entomological Society, 44(3), 187–223.

102. Huckett, H.C. (1954). A review of the North American species belonging to the genus Hydrotaea Robineau-Desvoidy (Diptera: Muscidae). Annals of the Entomological Society of America, 47(2), 316–342. doi: 10.1093/aesa/47.2.316

103. Huckett, H.C. (1965a). The Muscidae of northern Canada, Alaska, and Greenland (Diptera). Memoirs of the Entomological Society of Canada, 97, 5–369. doi: 10.4039/entm9742fv

104. Huckett, H.C. (1965b). Family Muscidae. In A. Stone, C.W. Sabrosky, W.W. Wirth, R.H. Foote, and J.T.P. Coulson (Eds.), A catalog of the Diptera of America north of Mexico. Handbook no. 276. (pp. 869–915). U.S. Department of Agriculture, Washington, DC.

105. Huckett, H.C. & Vockeroth, J.R. (1987). Muscidae. In J.F. McAlpine, B.V. Peterson, G.E. Shewell, H.J. Teskey, J.R. Vockeroth, and D.M. Wood (Eds), Manual of Nearctic Diptera. Vol. 2. (pp. 1115–1131). Agriculture Canada Monograph No. 28.

106. Huemer, P., Mutanen, M., Sefc, K.M. & Hebert, P.D.N. (2014). Testing DNA barcode performance in 1000 species of European Lepidoptera: large geographic distances have small genetic impacts. PLOS ONE, 9(12). doi: 10.1371/journal.pone.0115774

107. Ivanova, N.V., DeWaard, J.R. & Hebert, P.D.N. (2006). An inexpensive, automation-friendly protocol for recovering high-quality DNA. Molecular Ecology Notes, 6(4), 998–1002. doi: 10.1111/j.1471-8286.2006.01428.x

108. Jackson, G. A. (1977) Nearctic and Palaearctic Paracladopelma Harnisch and Saetheria n. gen. (Diptera: Chironomidae). Journal of the Fisheries Research Board of Canada, 34(9), 1321–1359.

109. James, M.T. & Huckett, H.C. (1952). The Diptera collected by I.O. Buss in southwestern Yukon Territory during the summer of 1950. The Canadian Entomologist, 84(9), 265–269.

110. Janzen, D.H. & Hallwachs, W. (2011). Joining inventory by parataxonomists with DNA barcoding of a large complex tropical conserved wildland in Northwestern Costa Rica. PLOS ONE, 6(8). doi: 10.1371/journal.pone.0018123

111. Janzen, D.H., Hallwachs, W., Blandin, P., Burns, J.M., Cadiou, J-M., Chacon, I., Dapkey, T., Deans, A.R., Epstein, M.E., Espinoza, B., Franclemont, J.G., Haber, W.A., Hajibabaei, M., Hall, J.P.W., Hebert, P.D.N., Gauld, I.D., Harvey, D.J., Hausmann, A., Kitching, I.J., Lafontaine, D., Landry, J-F., Lemaire, C., Miller, J.Y., Miller, J.S., Miller, L., Miller, S.E., Montero, J., Munroe, E., Green, S.R., Ratnasingham, S., Rawlins, J.E., Robbins, R.K., Rodriguez, J.J., Rougerie, R., Sharkey, M.J., Smith, M.A., Solis, M.A., Sullivan, J.B., Thiaucourt, P., Wahl, D.B., Weller, S.J., Whitfield, J.B., Willmott, K.R., Wood, D.M., Woodley, N.E. & Wilson, J.J. (2009) Integration of DNA barcoding into an ongoing inventory of complex tropical biodiversity. Molecular Ecology Resources, 9(1), 1–16. doi: 10.1111/j.1755-0998.2009.02628.x

112. Jeffery, N.W., Elías-Gutiérrez, M. & Adamowicz, S.J. (2011). Species diversity and phylogeographical affinities of the Branchiopoda (Crustacea) of Churchill, Manitoba, Canada. PLOS ONE, 6(5). doi: 10.1371/journal.pone.0018364

113. Johnson, C.W. (1929). Diptera of Labrador. Psyche, 36(2), 129–146. doi: 10.1155/1929/84643

114. Jürgenstein, S., Kurina, O. & Põldmaa, K. (2015). The Mycetophila ruficollis Meigen (Diptera, Mycetophilidae) group in Europe: elucidating species delimitation with COI and ITS2 sequence data. ZooKeys, 508(508), 15–51. doi: 10.3897/zookeys.508.9814

115. Kevan, P.G. (1972). Insect pollination in high Arctic flowers. Journal of Ecology, 60(3), 831–847.

116. Kjærandsen, J. (2022). Current state of DNA barcoding of Sciaroidea (Diptera)—Highlighting the need to build the reference library. Insects, 13(2), 147. doi: 10.3390/insects13020147

117. Kyerematen, R. A. K., Sæther, O. A., & Andersen, T. (2000). A review of the Rheotanytarsus pellucidus group (Diptera: Chironomidae). In O. Hoffrichter (Ed.), Late 20th century research on Chironomidae: an anthology from the 13th international symposium on Chironomidae (pp. 147–170). Shaker Verlag.

118. Laforest, B.J., Winegardner, A.K., Zaheer, O.A., Jeffery, N.W., Boyle, E.E. & Adamowicz, S.J. (2013). Insights into biodiversity sampling strategies for freshwater microinvertebrate faunas through bioblitz campaigns and DNA barcoding. BMC Ecology, 13(1), 13. doi: 10.1186/1472-6785-13-13

119. Langton, P. H. (1985) Review of type specimens of the limbatellus group, with a provisional key to known females of Psectrocladius Kieffer (Diptera: Chironomidae). Entomologica Scandinavica, 15, 477–485.

120. Langton, P. H. & Pinder, L. C. V. (2007) Keys to the adult male Chironomidae of Britain and Ireland; 2 vols. Freshwater Biological Association, 64, 239+168.

121. Lehmann, J. (1970). Revision der Europäischen Arten (Imagines) der Gattung Parachironomus Lenz (Diptera, Chironomidae). Hydrobiologia, 36,(1), 129–158. doi: 10.1007/bf00751287

122. Lehmann, J. (1970). Revision der europäischen Arten (Imaginees und Puppen) der Gattung Rheotanytarsus Bause (Diptera, Chironomidae). Zoologischer Anzeiger, 185(5/6), 343–378.

123. Lehmann, J. (1972). Revision der europäischen Arten (Puppen ♂♂ und Imagines ♂♂) der Gattung Eukiefferiella Thienemann (Diptera: Chironomidae). Beiträge zur Entomologie, 22(7/8), 347–405.

124. Lin, X., Stur, E. & Ekrem, T. (2015). Exploring genetic divergence in a species-rich insect genus using 2790 DNA barcodes. PLoS One, 10(9), p.e0138993. doi: 10.1371/journal.pone.0138993

125. Lin, X.-L., Stur, E., & Ekrem, T. (2018). DNA barcodes and morphology reveal unrecognized species in Chironomidae (Diptera). Insect Systematics & Evolution, 49(4), 329–398. doi: 10.1163/1876312X-00002172

126. Lindeberg, B. (1967). Sibling species delimitation in the Tanytarsus lestagei aggregate (Diptera, Chironomidae). Annales Zoologici Fennici, 4(1), 45–86.

127. Lindeberg, B., & Wiederholm, T. (1979). Notes on the taxonomy of European species of Chironomus (Diptera: Chironomidae). Entomologica scandinavica Suppl., 10, 99–116.

128. Liu, Y. & Wang, X. (2005) Synorthocladius Thienemann from China, with a review on the genus (Diptera: Chironomidae: Orthocladiinae). Zootaxa, 1057(1), 51–60. doi: 10.11646/zootaxa.1057.1.3

129. MacDonald, J.F. (1994). Review of the Nearctic species of the genus Chelifera Macquart (Diptera: Empididae; Hemerodromiinae) of America north of Mexico. Proceedings of the Entomological Society of Washington, 96(2), 236–275.

130. Machado, S., dos Santos, J.P.J., Fonseca, P.M., Bolzan, A.R., David, J., da Silva Loreto, E.L., Gottschalk, M.S. & Robe, L.J. (2017). Neotropical mycophagous drosophilids (Diptera: Drosophilidae): DNA barcoding as a way of overcoming the taxonomic impediment. Insect Conservation and Diversity, 10(3), 271–281. doi: 10.1111/icad.12223

131. [dataset] Majoros, S.E., Wang, J., Levesque-Beaudin, V., Borkent, C.J., Brodo, F., Brooks, S.E., Boucher, S., Cumming, J.M., Currie, D.C., Ekrem, T., Elliott, T.A., Hebert, P.D.N., Savage, J., Schaefer, P., Sinclair, B.J., Skevington, J.H., Solecki, A.M., Stur, E., Wheeler, T.A., Woodcock, T.S., Young, A.D., Young, C. & Adamowicz, S.J.; 2023; DS-CDIPTERA Collaborative DNA barcode library for Diptera of Churchill, Canada: A resource for northern research; Barcode of Life Data Systems; dx.doi.org/10.5883/DS-CDIPTERA

132. Mackerras, I.M. (1954) The classification and distribution of Tabanidae (Diptera). I. General review. Australian Journal of Zoology, 2, 431–454

133. Malloch, J. R. (1915) The Chironomidae, or midges, of Illinois, with particular reference to the species occuring in the Illinois River. Bulletin of the Illinois State Laboratory of Natural History, 10, 273–591.

134. Malloch, J.R. (1918). Diptera from southwestern United States. Part IV. Anthomyiidae. Transactions of the American Entomological Society, 44, 263–319.

135. Malloch, J.R. (1919). Report of the Canadian Arctic Expedition 1913-18, 3. Pt. C, Dipt., pp. 60c–90c. Ottawa.

136. Malloch, J.R. (1920a). Scientific results of the Katmai Expedition of the National Geographic Society. XII. Descriptions of Diptera of the families Anthomyidae and Scatophagidae. The Ohio Journal of Science, 20, 267–288.

137. Malloch, J.R. (1920b). Descriptions of new North American Anthomyiidae (Diptera). Transactions of the American Entomological Society, 46(2), 133–196.

138. Malloch, J.R. (1921). Dipterous insects of the family Anthomyiidae from the Pribilof Islands, Alaska. Proceedings of the California Academy of Sciences, 11, 178–182.

139. Malloch, J.R. (1923). Flies of the Anthomyiid genus Phaonia Robineau-Desvoidy and related genera, known to occur in North America. Transactions of the American Entomological Society, 48(3), 227–282.

140. Manitoba Land Initiative (n.d.) LUC_EOSD_Northeast [Dataset]. Manitoba Land Initiative. Accessed from https://mli.gov.mb.ca/landuse/index.html

141. Martinez, A., Reeves, N., Perry, R. & Horton, M. (2020). Do attractants bias the results of Malaise trap research? Journal of Emerging Investigators, 3.

142. McAlpine, J.F. (1965). Insects and related terrestrial invertebrates of Ellef Ringnes Island. Arctic, 18(2), 73–103. doi: 10.14430/arctic3455

143. McAlpine, J.F. (1981). Key to families-adults. In McAlpine, J.F., Peterson, B.V., Shewell, G.E., Teskey, H.J., Vockeroth, J.R. & Wood, D.M (Eds), Manual of Nearctic Diptera. Volume 1. (pp 89–124). Research Branch Agriculture Canada.

144. Medeiros, A.S., Milošević, Đ., Francis, D.R., Maddison, E., Woodroffe, S., Long, A., Walker, I.R., Hamerlík, L., Quinlan, R., Langdon, P., Brodersen, K.P. & Axford, Y. (2020). Arctic chironomids of the northwest North Atlantic reflect environmental and biogeographic gradients. Journal of Biogeography, 48(3), 511–525. doi: 10.1111/jbi.14015

145. Mesmin, X., Vincent, M., Tricault, Y., Estorgues, V., Daniel, L., Cortesero, A., Faloya, V. & Le Ralec, A. (2019). Assessing the relationship between pest density and plant damage: a case study with the belowground herbivore Delia radicum (Diptera: Anthomyiidae) on broccoli. Applied Entomology and Zoology, 54(2), 155–165. doi: 10.1007/s13355-019-00607-3

146. Messing, J. (1983). New M13 vectors for cloning. Methods in Enzymology, 101, 20–78. 10.1016/0076-6879(83)01005-8

147. Michelsen, V. (2006). Annotated catalogue of the Anthomyiidae, Fanniidae, Muscidae and Scathophagidae (Diptera: Muscoidea) of Greenland. Steenstrupia, 29, 105–126.

148. Miller, L.A. (1951). Observations on the bionomics of some northern species of Tabanidae (Diptera). Canadian Journal of Zoology, 29(3), 240–265. doi: 10.1139/z51-023

149. Mitterboeck, F.T. & Adamowicz, S.J. (2013) Flight loss linked to faster molecular evolution in insects. Proceedings of the Royal Society B, 280(1767), 20131128. doi: 10.1098/rspb.2013.1128

150. Mitterboeck, F.T., Fu, J. & Adamowicz, S.J. (2016) Rates and patterns of molecular evolution in freshwater versus terrestrial insects. Genome, 59(11), 968–980. doi: 10.1139/gen-2016-0030

151. Molinatto, G., Demichelis, S., Bodino, N., Giorgini, M., Mori, N. & Bosco, D. (2020). Biology and prevalence in northern Italy of Verrallia aucta (Diptera, Pipunculidae), a parasitoid of Philaenus spumarius (Hemiptera, Aphrophoridae), the main vector of Xylella fastidiosa in Europe. Insects, 11(9), 607. doi: 10.3390/insects11090607

152. Murray, D. A. (1987) Conchapelopia aagaardi n. sp., a new species of Tanypodinae (Diptera: Chironomidae) from Norway. In: Sæther, O. A. (ed.): A conspectus of contemporary studies in Chironomidae (Diptera). Contributions from the IXth International Symposium on Chironomidae, Bergen, Norway, 1985. Entomologica Scandinavica, Suppl., 29, 161–166.

153. Murray, D. A., Fittkau, E. J. (1989) The adult males of Tanypodinae (Diptera: Chironomidae) of the Holarctic region – Keys and diagnoses. In: Wiederholm, T. (ed.): Chironomidae of the Holarctic region. Keys and diagnoses. Part 3. Adult males. Entomologica Scandinavica, Suppl., 34, 37–123.

154. Nneji, L.M., Adeola, A.C., Ayoola, A.O., Oladipo, S.O., Wang, Y-Y., Malann, Y.D., Anyaele, O., Nneji, I.C., Rahman, M.M. & Olory, C.S. (2020). DNA barcoding and species delimitation of butterflies (Lepidoptera) from Nigeria. Molecular Biology Reports, 47(12), 9441–9457. doi: 10.1007/s11033-020-05984-5

155. Northern Biodiversity Program. (n.d.) About the NMP. Northern Biodiversity Program. Retrieved from https://northernbiodiversity.wordpress.com/about-the-nbp/

156. Nowosad DSJ, Hogg ID, Cottenie K, et al (2024) High diversity of freshwater invertebrates on Inuinnait Nuna, the Canadian arctic, revealed using mitochondrial DNA barcodes. Polar Biol 47:1475–1490. 10.1007/s00300-024-03311-3

157. Nugent, C.M., Elliott, T.A., Ratnasingham, S., Adamowicz, S.J. (2020). Coil: an R package for cytochrome c oxidase I (CO1) DNA barcode data cleaning, translation, and error evaluation. Genome, 63(6), 291–305. doi: 10.1139/gen-2019-0206

158. Oksanen, J., Simpson, G., Blanchet, F., Kindt, R., Legendre, P., Minchin, P., O’Hara, R., Solymos, P., Stevens, M., Szoecs, E., Wagner, H., Barbour, M., Bedward, M., Bolker, B., Borcard, D., Carvalho, G., Chirico, M., De Caceres, M., Durand, S., Evangelista, H., FitzJohn, R., Friendly, M., Furneaux, B., Hannigan, G., Hill, M., Lahti, L., McGlinn, D., Ouellette, M., Ribeiro Cunha, E., Smith, T., Stier, A., Ter Braak, C. & Weedon, J. (2022). vegan: Community Ecology Package. R package version 2.6–4. https://CRAN.R-project.org/package=vegan.

159. Olafson, PU., Aksoy, S., Attardo, G.M., Buckmeier, G., Chen, X., Coates, C.J., Davis, M., Dykema, J., Emrich, S.J., Friedrich, M., Holmes, C.J., Ioannidis, P., Jansen, E.N., Jennings, E.C., Lawson, D., Martinson, E.O., Maslen, G.L., Meisel, R.P., Murphy, T.D., Nayduch, D., Nelson, D.R., Oyen, K.J., Raszick, T.J., Ribeiro, J.M.C., Robertson, H.M., Rosendale, A.J., Sackton, T.B., Saelao, P., Swiger, S.L., Sze, S., Tarone, A.M., Taylor, D.B., Warren, W.C., Waterhouse, R.M., Weirauch, M.T., Werren, J.H., Wilson, R.K., Zdobnov, E.M. & Benoit, J.B. (2021). The genome of the stable fly, Stomoxys calcitrans, reveals potential mechanisms underlying reproduction, host interactions, and novel targets for pest control. BMC Biology, 19(1), 41. doi: 10.1186/s12915-021-00975-9

160. Oliver, D.R. (1981). Chironomidae. In McAlpine, J.F., Peterson, B.V., Shewell, G.E., Teskey, H.J., Vockeroth, J.R. & Wood, D.M. (Eds), Manual of Nearctic Diptera. Volume 1. (pp. 423–458). Research Branch Agriculture Canada

161. Oliver, D. R. (1989). The adult males of Diamesinae (Diptera: Chironomidae) of the Holarctic region – Keys and diagnoses. In: Wiederholm, T. (ed.): Chironomidae of the Holarctic region. Keys and diagnoses. Part 3 – Adult males. Entomologica Scandinavica, Suppl., 34, 129–154.

162. Oliver, D. R., & Dillon, M. E. (1994). Systematics of some species of Micropsectra (Diptera: Chironomidae) living in low-order streams in southern Ontario, Canada. The Canadian Entomologist, 126(2), 199–217. doi: 10.4039/Ent126199-2

163. Oliver, D. R. & Roussel, M. E. (1983) Redescription of Brillia Kieffer (Diptera: Chironomidae) with descriptions of Nearctic species. Canadian entomologist, 115, 257–279. doi: 10.4039/Ent115257-3

164. Oosterbroek, P. (1984). A revision of the crane-fly genus Nephrotoma Meigen, 1803, in North America (Diptera, Tipulidae). Part II. The non-dorsalis species-groups. Beaufortia, 34, 117–180.

165. Pava-Ripoll, M., Pearson, R.E.G., Miller, A.K. & Ziobro, G.C. (2012). Prevalence and relative risk of Cronobacter spp., Salmonella spp., and Listeria monocytogenes associated with the body surfaces and guts of individual filth flies. Applied and Environmental Microbiology, 78(22), 7891–7902. doi: 10.1128/AEM.02195-12

166. Pentinsaari, M. (2016). Utility of DNA barcodes in identification and delimitation of beetle species, with insights into COI protein structure across the animal kingdom. [Doctoral Dissertation, University of Oulu]. http://jultika.oulu.fi/files/isbn9789526212104.pdf

167. Pentinsaari, M., Blagoev, G.A., Hogg, I.D., Levesque-Beaudin, V., Perez, K., Sobel, C.N., Vandenbrink, B. & Borisenko, A. (2020). A DNA barcoding survey of an Arctic arthropod community: Implications for future monitoring. Insects, 11(1), 46. doi: 10.3390/insects11010046

168. Pentinsaari, M., Hebert, P.D.N. & Mutanen, M. (2014). Barcoding beetles: a regional survey of 1872 species reveals high identification success and unusually deep interspecific divergences. PLOS ONE, 9(9). doi: 10.1371/journal.pone.0108651

169. Peterson, B.V. (1981). Simuliidae. In McAlpine, J.F., Peterson, B.V., Shewell, G.E., Teskey, H.J., Vockeroth, J.R. & Wood, D.M. (Eds), Manual of Nearctic Diptera. Volume 1. (pp. 247–256). Research Branch Agriculture Canada.

170. Pohjoismäki, J.L.O., Kahanpää, J. & Mutanen, M. (2016). DNA barcodes for the northern European Tachinid flies (Diptera: Tachinidae). PLOS ONE, 11(11). doi: 10.1371/journal.pone.0164933

171. Pont, A.C. (1984). A revision of the Fanniidae and Muscidae (Diptera) described by Fallén. Entomologica Scandinavica, 15, 277–297.

172. Pont, A.C. (1986). Family Muscidae. In Soós Á, Papp L. (Eds.), Catalogue of Palaearctic Diptera, volume 11:Scathophagidae-Hypodermatidae. (pp. 55–215). Budapest: Akadémiai Kiadó.

173. Pont, A.C. (2011). The Muscidae described by J.W. Zetterstedt (Insecta: Diptera). Zootaxa, 2852(1), 1–83. doi: 10.11646/zootaxa.2852.1.1

174. Porco, D., Rougerie, R., Deharveng, L. & Hebert, P.D.N. (2010). Coupling non-destructive DNA extraction and voucher retrieval for small soft-bodied Arthropods in a high-throughput context: the example of Collembola. Molecular Ecology Resources, 10(6), 942–945. doi: 10.1111/j.1755-0998.2010.2839.x

175. Porco, D., Skarżyński, D., Decaëns, T., Hebert, P.D.N. & Deharveng, L. (2013). Barcoding the Collembola of Churchill: a molecular taxonomic reassessment of species diversity in a Sub-Arctic area. Molecular Ecology Resources, 14(2), 249–261. doi: 10.1111/1755-0998.12172

176. Posada-López, L., Rodrigues, B.L., Velez, I.D. & Uribe, S. (2023). Improving the COI DNA barcoding library for Neotropical phlebotomine sand flies (Diptera: Psychodidae). Parasites & Vectors, 16(1), 198. doi: 10.1186/s13071-023-05807-z

177. Proulx, I., Martin, J., Carew, M. & Hare, L. (2013) Using various lines of evidence to identify Chironomus species (Diptera: Chironomidae) in eastern Canadian lakes. Zootaxa, 3741(4), 401–458

178. R Core Team. (2023). R: A language and environment for statistical computing. R Foundation for Statistical Computing, Vienna, Austria. URL http://www.R-project.org/

179. Ratnasingham, S. & Hebert, P.D.N. (2007). BOLD: the barcode of life data system. Molecular Ecology Notes, 7(3), 355–364. doi:10.1111/j.1471-8286.2007.01678.

180. Ratnasingham, S. & Hebert, P.D.N. (2013). A DNA-based registry for all animal species: The Barcode Index Number (BIN) system. PLoS ONE, 8(7), e66213.

181. Reiss, F., & Fittkau, E. J. (1971). Taxonomie und Ökologie europäisch verbreiteter Tanytarsus-Arten (Chironomidae, Diptera). Archiv für Hydrobiologie Suppl., 40, 75–200.

182. Reiss, F., & Säwedal, L. (1981). Keys to males and pupae of the Palaearctic (excl. Japan) Paratanytarsus Thienemann & Bause, 1913, n. comb., with descriptions of three new species (Diptera: Chironomidae). Entomologica Scandinavica, Suppl., 15, 73–104.

183. Renaud, A.K., Savage, J. & Roughley, R.E. (2012a). Muscidae (Diptera) diversity in Churchill, Canada, between two time periods: evidence for limited changes since the Canadian Northern Insect Survey. The Canadian Entomologist, 144(1), 29–51. doi: 10.4039/tce.2012.6

184. Renaud, A.K., Savage, J. & Adamowicz, S.J. (2012b). DNA barcoding of northern Nearctic Muscidae (Diptera) reveals high correspondence between morphological and molecular species limits. BMC Ecology, 12(1), 24. doi: 10.1186/1472-6785-12-24

185. Rivera, J. & Currie, D.C. (2009). Identification of Nearctic black flies using DNA barcodes (Diptera: Simuliidae). Molecular Ecology Resources, 9(s1), 224–236. doi: 10.1111/j.1755-0998.2009.02648.x

186. Roback, S. S. (1971) The adults of the subfamily Tanypodinae (= Pelopiinae) in North America (Diptera: Chironomidae). Monographs, Academy of Natural Sciences of Philadelphia, 17, 1–410.

187. Robinson, H. & Vockeroth, J.R. (1981). Dolichopodidae. In McAlpine, J.F., Peterson, B.V., Shewell, G.E., Teskey, H.J., Vockeroth, J.R. & Wood, D.M. (Eds), Manual of Nearctic Diptera. Volume 1. (pp. 247–256). Research Branch Agriculture Canada.

188. Roslin, T., Somervuo, P., Pentinsaari, M., Hebert, P.D.N., Agda, J., Ahlroth, P., Anttonen, P., Aspi, J., Blagoev, G., Blanco, S., Chan, D., Clayhills, T., deWaard, J., deWaard, S., Elliot, T., Elo, R., Haapala, S., Helve, E., Ilmonen, J., Hirvonen, P., Ho, C., Itämies, J., Ivanov, V., Jakovlev, J., Juslén, A., Jussila, R., Kahanpää, J., Kaila, L., Jari-PekkaKaitila, Kakko, A., Kakko, I., Karhu, A., Karjalainen, S., Kjaerandsen, J., Koskinen, J., Laasonen, E.M., Laasonen, L., Laine, E., Lampila, P., Levesque-Beaudin, V., Lu, L., Lähteenaro, M., Majuri, P., Malmberg, S., Manjunath, R., Martikainen, P., Mattila, J., McKeown, J., Metsälä, P., Miklasevskaja, M., Miller, M., Miskie, R., Muinonen, A., Veli-MattiMukkala, Naik, S., Nikolova, N., Nupponen, K., Ovaskainen, O., Österblad, I., Paasivirta, L., Pajunen, T., Parkko, P., Paukkunen, J., Penttinen, R., Perez, K., Pohjoismäki, J., Prosser, S., Raekunnas, M., Rahulan, M., Rannisto, M., Ratnasingham, S., Raukko, P., Rinne, A., Rintala, T., Romo, S.M., Salmela, J., Salokannel, J., Savolainen, R., Schulman, L., Sihvonen, P., Soliman, D., Sones, J., Steinke, C., Ståhls, G., Tabell, J., Tiusanen, M., Várkonyi, G., Vesterinen, E.J., Viitanen, E., Vikberg, V., Viitasaari, M., Vilen, J., Warne, C., Wei, C., Winqvist, K., Zakharov, E. & Mutanen, M. (2021). A molecular-based identification resource for the arthropods of Finland. Molecular Ecology Resources, 22(2), 803–822. doi: 10.1111/1755-0998.13510

189. Rossaro, B., Marziali, L. & Boggero, A. (2022). Response of Chironomids to key environmental factors: perspective for biomonitoring. Insects, 13(10), 911. doi: 10.3390/insects13100911

190. Sabrosky, C.W. 1936. A synopsis of the Nearctic species of Oscinella and Madiza, based on a study of the types (Diptera, Chloropidae). Annals Entomological Society of America, 29: 707—728.

191. Sabrosky, C.W. 1938. Taxonomic notes on the dipterous family Chloropidae. Journal of the New York Entomological Society. 46(4): 417—434.

192. Sabrosky, C.W. 1940. Twelve New North American species of Oscinella (Diptera: Chloropidae). The Canadian Entomologist. 72(11): 214—230.

193. Sabrosky, C.W. 1943 A revised synopsis of Nearctic Thaumatomyia (=Chloropisca) (Diptera: Chloropidae). The Canadian Entomologist, 75(6): 109–117.

194. Sabrosky, C.W. 1948. A synopsis of the Nearctic species of Elachiptera and related genera (Diptera: Chloropidae). Journal of the Washington Academy of Sciences, 38(11): 365—382.

195. Sabrosky, C.W. 1950 A new species of Eribolus from California (Diptera, Chloropidae). The Pan-Pacific Entomologist, 26: 91--92.

196. Sabrosky, C.W. 1951 A revision of nearctic species of Gaurax (Diptera: Chloropidae) The American Midland Naturalist, 45(2): 407—431.

197. Sabrosky, C.W. and Valley, K.R. 1987. A new Elachiptera from salt marshes, with a redescription of E. penita and partially revised key to nearctic Elachiptera (Diptera: Chloropidae). Proceedings of the Entomological Society of Washington. 89: 581—586.

198. Sæther, O. A. (1969). Some nearctic Podonominae, Diamesinae, and Orthocladiinae (Diptera: Chironomidae). Bulletin/Fisheries Research Board of Canada, 170, 1–154.

199. Sæther, O. A. (1975a). Nearctic and Palaearctic Heterotrissocladius (Diptera: Chironomidae). Bulletin/Fisheries Research Board of Canada, 193, 67.

200. Sæther, O. A. (1975b). Twelve new species of Limnophyes Eaton, with keys to Nearctic males of the genus (Diptera: Chironomidae). Canadian entomologist, 107(10), 1029–1056. doi: 10.4039/Ent1071029-10

201. Sæther, O. A. (1977). Taxonomic studies on Chironomidae: Nanocladius, Pseudochironomus, and the Harnischia complex. Bulletin/Fisheries Research Board of Canada, 196, 1–143.

202. Sæther, O. A. (1986). A review of the genus Rheocricotopus Thienemann & Harnisch, 1932, with the description of three new species (Diptera, Chironomidae) In: Fittkau, E. J. (ed.): Beiträge zur Systematik der Chironomidae, Diptera. Spixiana Suppl, 11, 59–108.

203. Sæther, O. A. (1989). Metriocnemus van der Wulp: a new species and a revision of species described by Meigen, Zetterstedt, Stæger, Holmgren, Lundström and Strenzke (Diptera: Chironomidae). Entomologica Scandinavica, 19(4), 393–430.

204. Sæther, O. A. (1990). A review of the genus Limnophyes Eaton from the Holarctic and Afrotropical regions (Diptera: Chironomidae, Orthocladiinae). Entomologica Scandinavica, Suppl., 35, 135.

205. Sæther, O. A. (1995). Metriocnemus van der Wulp: Seven new species, revision of species, and new records (Diptera: Chironomidae). Annales de limnologie, 31(1), 35–64. doi: 10.1051/limn/1995002

206. Sæther, O. A. (2004a). A review of Orthocladius subgenus Symposiocladius Cranston (Diptera: Chironomidae). Aquatic Insects, 25(4), 281–317. doi: 10.1076/aqin.25.4.281.26221

207. Sæther, O. A. (2004b). Three new species of Orthocladius subgenus Eudactylocladius (Diptera: Chironomidae) from Norway. Zootaxa, 508(1), 1–12. doi: 10.11646/zootaxa.508.1.1

208. Sæther, O. A. (2009). Cryptochironomus Kieffer from Lake Winnipeg, Canada, with a review if Nearctic species (Diptera: Chironomidae). Zootaxa, 2208(1), 1–24. doi: 10.11646/zootaxa.2208.1.1

209. Sæther, O. A. (2010).Cryptotendipes Lenz from Manitoba, Canada, with keys to known immatures of the genus (Diptera: Chironomidae). Zootaxa, 2412, 1–2.

210. Sæther, O. A. & Halvorsen, G. A. (1981). Diagnoses of Tvetenia Kieff. emend., Dratnalia n. gen., and Eukiefferiella Thien. emend., with a phylogeny of the Cardiocladius group (Diptera: Chironomidae). Entomologica Scandinavica, Suppl., 15, 269–285.

211. Sæther, O. A. & Oyewo, E. A. (2008). Keys, phylogenies and biogeography of Polypedilum subgenus Uresipedilum Oyewo and Sæther (Diptera, Chironomidae). Zootaxa, 1806, 1–34.

212. Sæther, O. A. & Sublette, J. E. (1983). A review of the genera Doithrix n. gen., Georthocladius Strenzke, Parachaetocladius Wülker and Pseudorthocladius Goetghebuer (Diptera: Chironomidae, Orthocladiinae). Entomologica Scandinavica, Suppl., 20, 100.

213. Sæther, O. A. & Wang, X. (1995). Revision of the genus Paraphaenocladius Thienemann, 1924 of the world (Diptera: Chironomidae, Orthocladiinae). Entomologica Scandinavica, Suppl., 48, 1–69.

214. Saitou, N. & Nei, M. (1987). The neighbor-joining method: a new method for reconstructing phylogenetic trees. Molecular Biology and Evolution, 4(4), 406–425. doi: 10.1093/oxfordjournals.molbev.a040454

215. Savage, J. (2003). Revision of the genus Thricops Rondani (Diptera: Muscidae). Insect Systematics and Evolution supplement, 61, 3–143.

216. Säwedal, L. (1981). Description of Micropsectra tori n. sp. from Greenland, with notes on the recurvata-group (Diptera: Chironomidae). Entomologica Scandinavica, 12, 27–30.

217. Schlee, D. (1968) Vergleichende Merkmalsanalyse zur Morphologie und Phylogenie der Corynoneura-Gruppe (Diptera, Chironomidae). Zugleich eine Allgemeine Morphologie der Chironomiden-Imago (?) Stuttgarter Beiträge zur Naturkunde, 180, 150.

218. Schliep K.P. (2011). phangorn: phylogenetic analysis in R. Bioinformatics, 27(4), 592–593. doi: 10.1093/bioinformatics/btq706

219. Scudder, G.G.E. & Cannings, R.A. (2006). The Diptera Families of British Columbia. University of British Columbia.

220. Shamshad, A. (2010). The development of integrated pest management for the control of mushroom sciarid flies, Lycoriella ingenua (Dufour) and Bradysia ocellaris (Comstock), in cultivated mushrooms. Pest Management Science, 66(10), 1063–1074. doi: 10.1002/ps.1987

221. Sinclair, B.J. (2008). The systematics of New World Clinocera Meigen (Diptera: Empididae: Clinocerinae). NRC Research Press, Ottawa. viii + 245 pp.

222. Sinclair, B.J., Brooks, S.E., Cumming, J.M., & Coovert, G.A. (2011). Revision of the Nearctic species of Heleodromia (Diptera: Empidoidea: Brachystomatidae). The Canadian Entomologist, 143(6), 629–649. doi: 10.4039/n11-036

223. Skevington, J. & Marshall, S.A. (1998). Systematics of New World Pipunculus (Diptera: Pipunculidae). Thomas Say Publications in Entomology: Monographs. Lanham, Maryland: Entomological Society of America. 201 pp.

224. Skevington, J.H. (2005). Revision of Nearctic Nephrocerus Zetterstedt (Diptera: Pipunculidae). Zootaxa, 977, 1–36.

225. Skvarla, M.J., Larson, J.L., Fisher, J.R. & Dowling, A.P.G. (2020). A review of terrestrial and canopy Malaise traps. Annals of the Entomological Society of America, 114(1), 27–47. doi: 10.1093/aesa/saaa044

226. Smith, R.E., Veldhuis, H., Mills, G.F., Eilers, R.G., Fraser, W.R. & Lelyk, G.W. (1998). Terrestrial Ecozones, Ecoregions, and Ecodistricts of Manitoba: An Ecological Stratification of Manitoba’s Natural Landscapes. Agriculture and Agri-Food Canada. Available at: https://sis.agr.gc.ca/cansis/publications/ecostrat/provDescriptions/mbteee/mbteee_report.pdf

227. Snow, W.A. (1891). The moose fly – a new Haematobia. The Canadian Entomologist, 23(4), 87–89. doi: 10.4039/Ent2387-4

228. Snyder, F.M. (1949a). Nearctic Helina Robineau-Desvoidy (Diptera: Muscidae). American Museum of Natural History, 94, 112–159.

229. Snyder, F.M. (1949b). Review of Nearctic Mydaea, sensu stricto, and Xenomydaea (Diptera: Muscidae). American Museum Novitates, 1401, 1–38.

230. Snyder, F.M. (1954). A review of Nearctic Lispe Latreille (Diptera: Muscidae). American Museum Novitates, 1675, 1–40.

231. Soponis, A. R. (1977) A revision of the Nearctic species of Orthocladius (Orthocladius) van der Wulp (Diptera: Chironomidae). Memoirs of the Entomological Society of Canada, 102, 1–187.

232. Soponis, A. R. (1990) A revision of the Holarctic species of Orthocladius (Euorthocladius) (Diptera: Chironomidae). Spixiana Suppl., 13, 1–56.

233. Spencer, K.A. (1969). The Agromyzidae of Canada and Alaska. Memoirs of the Entomological Society of Canada 64, 1–311

234. Spencer, K.A. (1976). The Agromyzidae of Fennoscandia and Denmark. Part II. Fauna Entomologica Scandinavica, 5, 305–606

235. Spencer, K.A. & Steyskal, G.C. (1986). Manual of the Agromyzidae (Diptera) of the United States. U. S. Department of Agriculture, Agriculture Handbook 638. 478 pp

236. Srivathsan, A., Ang, Y., Heraty, J.M., Hwang, W.S., Jusoh, W.F.A., Kutty, S.N., Puniamoorthy, J., Yeo, D., Roslin, T. & Meier, R. (2023). Convergence of dominance and neglect in flying insect diversity. Nature Ecology & Evolution.

237. Srivathsan, A. & Meier, R. (2011) On the inappropriate use of Kimura-2-parameter (K2P) divergences in the DNA-barcoding literature. Cladistics, 28(2), 190–194

238. Stahlhut, J.K., Fernández-Triana, J., Adamowicz, S.J., Buck, M., Goulet, H., Hebert, P.D.N., Huber, J.T., Merilo, M.T., Sheffield, C.S., Woodcock, T. & Smith, M.A. (2013). DNA barcoding reveals diversity of Hymenoptera and the dominance of parasitoids in a Sub-Arctic environment. BMC Ecology, 13(1), 2. doi: 10.1186/1472-6785-13-2

239. Starý, J. and Brodo, F. (2009). Arctic species the subgenus Symplecta sensu stricto (Diptera: Limoniidae). Canadian Entomologist, 141, 1–30.

240. Steffan, W.A. (1981). Sciaridae. In McAlpine, J.F., Peterson, B.V., Shewell, G.E., Teskey, H.J., Vockeroth, J.R. & Wood, D.M. (Eds), Manual of Nearctic Diptera. Volume 1. (pp. 247–256). Research Branch Agriculture Canada.

241. Stein, P. (1898). Nordamerikanische Anrhomyiden. Bcitrag zur Dipterenfauna der Vereinigten Staaten. Berliner Entomolog. Zeitschr. (1897). 42, 161–288

242. Stein, P. (1920). Nordamerikanische Anthomyiden, 2. Beitrag. Arch. Naturgesch. (1918). 84A(9), 1–106

243. Steyskal, G.C. & Knutson, L.V. (1981). Empididae. In McAlpine, J.F., Peterson, B.V., Shewell, G.E., Teskey, H.J., Vockeroth, J.R. & Wood, D.M. (Eds), Manual of Nearctic Diptera. Volume 1. (pp. 247–256). Research Branch Agriculture Canada.

244. Stone, A. (1981). Culicidae. In McAlpine, J.F., Peterson, B.V., Shewell, G.E., Teskey, H.J., Vockeroth, J.R. & Wood, D.M. (Eds), Manual of Nearctic Diptera. Volume 1. (pp. 247–256). Research Branch Agriculture Canada.

245. Stur, E., Silva, F. L. da, Ekrem, T. (2019) Back from the past: DNA barcodes and morphology support Ablabesmyia americana Fittkau as a valid species (Diptera: Chironomidae). Diversity [Basle], 11(9, paper 173), 15.

246. Sublette, J. E. (1964). Chironomidae (Diptera) of Lousiana I. Systematics and immature stages of some lentic chironomids of West-Central Lousiana. Tulane Studies in Zoology, 11(4), 109–150. doi: 10.5962/bhl.part.7051

247. Sublette, J. E. (1967a). Type specimens of Chironomidae (Diptera) in the Canadian National Collections, Ottawa. Journal of the Kansas Entomological Society, 40(3), 290–331.

248. Sublette, J. E. (1967b). Type specimens of Chironomidae (Diptera) in the Cornell University collection. Journal of the Kansas Entomological Society, 40(4), 477–564.

249. Sublette, J. E. (1970). Type specimens of Chironomidae (Dipt.) in the Illinois Natural History Survey collection, Urbana. Journal of the Kansas Entomological Society, 43(1), 44–95.

250. Tamura, K., Stecher, G. & Kumar, S. (2021). MEGA11: Molecular Evolutionary Genetics Analysis version 11. Molecular Biology and Evolution, 38(7), 3022–3027. doi: 10.1093/molbev/msab120

251. Tangelder, I. (1983). A revision of the Crane Fly genus Nephrotoma Meigen, 1803, in North America (Diptera, Tipulidae), Part 1, the dorsalis species-group. Beaufortia, 33, 111–205.

252. Thompson, F.C. A Conspectus of Nearctic Syrphidae. Unpublished.

253. Thompson, F. C. (1991). A conspectus of the flower flies (Diptera: Syrphidae) of the Nearctic Region. Unpublished manuscript.

254. Tiusanen, M., Hebert, P.D.N., Schmidt, N.M. & Roslin, T. (2016). One fly to rule them all – muscid flies are the key pollinators in the Arctic. Proceedings of the Royal Society: Biological Sciences, 283(1839), 20161271. doi: 10.1098/rspb.2016.1271

255. Townes, H. K. Jr. (1945). The Nearctic species of Tendipedini [Diptera; Tendipedidae (= Chironomidae)]. The American Midland Naturalist Journal, 34(1), 1–206.

256. Tuiskunen, J. (1986). The Fennoscandian species of Parakiefferiella Thienemann (Diptera, Chironomidae, Orthocladiinae). Annales Zoologici Fennici, 23(2), 175–196.

257. Twinn, C.R., Hocking, B., McDuffie, Wm.C. & Cross, H.F. (1948). A preliminary account of the biting flies at Churchill, Manitoba. Canadian Journal of Research, 26(6), 334–357. doi: 10.1139/cjr48d-025

258. Uhler, J., Haase, P., Hoffmann, L., Hothorn, T., Schmidl, J., Stoll, S., Welti, E.A.R., Buse, J. & Müller, J. (2022). A comparison of different Malaise trap types. Insect Conservation and Diversity, 15(6), 666–672. doi: 10.1111/icad.12604

259. Vockeroth, J.R. (1992) The flower flies of the subfamily Syrphinae of Canada, Alaska and Greenland (Vol. 18). Ottawa: Canada Communications Group – Publishing. 456 pp.

260. Vockeroth, J.R. & Thompson, F.C. (1987). Manual of Nearctic Diptera.

261. Votýpka, J., Brzoňová, J., Ježek, J. & Modrý, D. (2019). Horse flies (Diptera: Tabanidae) of three West African countries: A faunistic update, barcoding analysis and trypanosome occurrence. Acta Tropica, 197. doi: 10.1016/j.actatropica.2019.105069

262. Walker, F. (1849). List of the specimens of dipterous insects in the collection of the British Museum. Pt. 4. pp. 923–972. London.

263. Wearing, C. H., Marshall, R. R., Attfield, B. & Colhoun, C. (2013). Phenology and distribution of the apple leafcurling midge (Dasineura mali (Kieffer)) (Diptera: Cecidomyiidae) and its natural enemies on apples under biological and integrated pest management in Central Otago, New Zealand. New Zealand Entomologist, 36(2), 87–106. doi: 10.1080/00779962.2012.712887

264. Weber, N.A. (1950). A survey of the insects and related arthropods of arctic Alaska. Part 1. Transactions of the American Entomological Society, 76(3), 147–206.

265. Weber, N.A. (1954). Arctic Alaskan Diptera. Proceedings of the Entomological Society of Washington, 56, 86–91.

266. Wheeler, T.A. (1994). A revision of the genus Epichlorops Becker (Diptera: Chloropidae). Insect Systematics and Evolution, 25(4): 393—414.

267. Wickham, H. (2016). ggplot2: Elegant graphics for data analysis. Springer-Verlag New York. https://ggplot2.tidyverse.org

268. Wickham, H., François, R., Henry, L., Müller, K. & Vaughan, D. (2023). dplyr: A Grammar of Data Manipulation. https://dplyr.tidyverse.org, https://github.com/tidyverse/dplyr

269. Wickham, H., Averick, M., Bryan, J., Chang, W., McGowan, L.D., François, R., Grolemund, G., Hayes, A., Henry, L., Hester, J., Kuhn, M., Pedersen, T.L., Miller, E., Bache, S.M., Müller, K., Ooms, J., Robinson, D., Seidel, D.P., Spinu, V., Takahashi, K., Vaughan, D., Wilke, C., Woo, K. & Yutani, H. (2019). Welcome to the Tidyverse. The Journal of Open Source Software, 4(43), 1686. doi: 10.21105/joss.01686

270. Wiederholm, T. (ed.) (1989) Chironomidae of the Holarctic region. Keys and diagnoses. Part 3. Adult males. – Entomologica Scandinavica, Suppl. 34, 532.

271. Wiegmann, B.M., Trautwein, M.D., Winkler, I.S. & Yeates, D.K. (2011) Episodic radiations in the fly tree of life. Proceedings of the National Academy of Sciences of the USA, 108(14), 5690–5695. doi: 10.1073/pnas.1012675108

272. Wirta, H.K., Vesterinen, E.J., Hambäck, P.A., Weingartner, E., Rasmussen, C., Reneerkens, J., Schmidt, N.M., Gilg, O. & Roslin, T. (2015a). Exposing the structure of an Arctic food web. Ecology and Evolution, 5(17), 3842–3856. doi: 10.1002/ece3.1647

273. Wirta, H.K., Weingartner, E., Hambäck, P.A. & Roslin, T. (2015b). Extensive niche overlap among the dominant arthropod predators of the High Arctic. Basic and Applied Ecology, 16(1), 86–92. doi: 10.1016/j.baae.2014.11.003

274. Wirta, H., Várkonyi,, G., Rasmussen, C., Kaartinen, R., Schmidt, N.M., Hebert, P.D.N., Barták, M., Blagoev, G., Disney, H., Ertl, S., Gjelstrup, P., Gwiazdowicz, D.J., Huldén, L., Ilmonen, J., Jakovlev, J., Jaschhof, M., Kahanpää, J., Kankaanpää, T., Krogh, P.H., Labbee, R., Lettner, C., Michelsen, V., Nielsen, S.A., Nielsen, T.R., Paasivirta, L., Pedersen, S., Pohjoismäki, J., Salmela, J., Vilkamaa, P., Väre, H., von Tschirnhaus, M., Roslin, T. (2016). Establishing a community-wide DNA barcode library as a new tool for arctic research. Molecular Ecology Resources, 16(3), 809–822. doi: 10.1111/1755-0998.12489

275. Woodcock, T.S., Boyle, E.E., Roughley, R.E., Kevan, P.G., Labbee, R.N., Smith, A.B.T., Goulet, H., Steinke, D. & Adamowicz, S.J. (2013). The diversity and biogeography of the Coleoptera of Churchill: insights from DNA barcoding. BMC Ecology, 13(1), 40–40. doi: 10.1186/1472-6785-13-40

276. World Health Organization. (2016). Mosquito control: can it stop Zika at source? https://reliefweb.int/report/world/mosquito-control-can-it-stop-zika-source

277. World Health Organization (2020). World Malaria Report 2020: 20 Years of Global Progress and Challenges. https://www.who.int/publications/i/item/9789240015791

278. Wright, E.S. (2016). Using DECIPHER v2.0 to analyze big biological sequence data in R. The R Journal, 8(1), 352–359. Doi: 10.32614/RJ-2016-025

279. Wulder, M.A., Nelson, T.A. EOSD Land Cover Classification Legend Report. Version 2. (2003) Natural Resources Canada, Canadian Forest Service, Pacific Forestry Centre, Victoria, BC. https://cfs.nrcan.gc.ca/publications?id=33752

280. Wülker, W. (1956) Zur Kenntnis der Gattung Psectrocladius Kieff. (Dipt., Chironom.). Individuelle Variabilität, Grenzen und Möglichkeiten der Artentrennung, Ökologie und Verbreitung. Archiv für Hydrobiologie, Suppl., 24(1), 1–66.

281. Wülker, W., Ryser, H. M., & Scholl, A. (1981). Revision der Gattung Chironomus Meigen (Dipt.). VI. C. holomelas Keyl, C. saxatilis n. sp., C. melanescens Keyl. Revue Suisse De Zoologie, 88(4), 903–924. doi: 10.5962/bhl.part.82418

282. Young, M.R., Behan-Pelletier, V.M. & Hebert, P.D.N. (2012). Revealing the hyperdiverse mite fauna of Subarctic Canada through DNA barcoding. PLOS ONE, 7(11). doi: 10.1371/journal.pone.0048755

283. Zahiri, R., Lafontaine, J.D., Schmidt, B.C., deWaard, J.R., Zakharov, E.V. & Hebert, P.D.N. (2014). A transcontinental challenge — a test of DNA barcode performance for 1,541 species of Canadian Noctuoidea (Lepidoptera). PLOS ONE, 9(3). doi: 10.1371/journal.pone.0092797

284. Zelentsov, N. I. (1985). Tablitsy dlya opredeleniya podrod i vidov roda Psectrocladius Kieff. (Diptera, Chironomidae). [Keys for the identification of subgenera and species in the genus Psectrocladius (Diptera, Chironomidae)]. In: Vodnye soobshchestva i biologiya gidrobiontov [Aquatic communities and biology of aquatic organisms]. Akademija Nauk SSSR, Institut Biologii Vnutrennich Vod, Trudy, 52, 119–137. [in Russian].

285. Zhou, X., Adamowicz, S.J., Jacobus, L.M., DeWalt R.E. & Hebert, P.D.N. (2009). Towards a comprehensive barcode library for arctic life – Ephemeroptera, Plecoptera, and Trichoptera of Churchill, Manitoba, Canada. Frontiers in Zoology, 6(1), 30. doi: 10.1186/1742-9994-6-30

286. Zhou, X., Jacobus, L.M., DeWalt, R.E., Adamowicz, S.J. & Hebert, P.D.N. (2010). Ephemeroptera, Plecoptera, and Trichoptera fauna of Churchill (Manitoba, Canada): insights into biodiversity patterns from DNA barcoding. Freshwater Science, 29(3), 814–837. doi: 10.1899/09-121.1

287. Zlobin, V.V. (1995). Review of mining flies of the genus Metopomyza Enderlein (Diptera: Agromyzidae). International Journal of Dipterological Research 6(2), 143–168

288. Zoller, L., Bennett, J.M. & Knight, T.M. (2020). Diel-scale temporal dynamics in the abundance and composition of pollinators in the Arctic summer. Scientific Reports, 10, 21198. doi: 10.1038/s41598-020-78165-w

289. Zorina, O. V. (2004). Novye i maloizvestnye vidy komarov-zvontsov roda Demicryptochironomus Lenz (Diptera, Chironomidae) s ostrova Sakhalin. (New and little known species of the genus Demicryptochironomus Lenz (Diptera, Chironomidae) from Sakhalin Island). In: Bogatov, V. V., Barkalov, V. Yu., Lelej, A. S., Makarchenko, E. A., Storozhenko, S. Yu. (eds): Flora and fauna of Sakhalin Island (Materials of International Sakhalin Island Project), Pt 1: 224–237. Dal’nauka, Vladivostok.

