## Supplementary material for "A Collaborative DNA Barcode Library for the Diptera of Churchill, Canada: Exploration of Biodiversity and Biogeography": A phenogram showing the relationship between the taxa collected in this study shows that COI sequences within families generally group together for th

### BOLD TaxonID Tree

| | | |
|---|---|---|
| Title | : | Tree Result - DS-CDIPTERA |
| Date | : | 08-Jul-2026 |
| Data Type | : | Nucleotide |
| Distance Model | : | Kimura 2 Parameter |
| Marker | : | COI-5P |
| Colourization | : | Taxonomy: Family |
| Label | : | Process ID |
| Label | : | Taxon |
| Label | : | Barcode Cluster (BIN) |
| Filter | : | length > 400bp only |
| Filter | : | exclude records flagged as misidentifications |
| Filter | : | exclude records with stop codons |
| Filter | : | exclude contaminants |
| Sequence Count | : | 16616 |
| Species count | : | 1212 |
| Genus count | : | 443 |
| Family count | : | 67 |
| Unidentified | : | 3953 |
| BIN Count | : | 2225 |

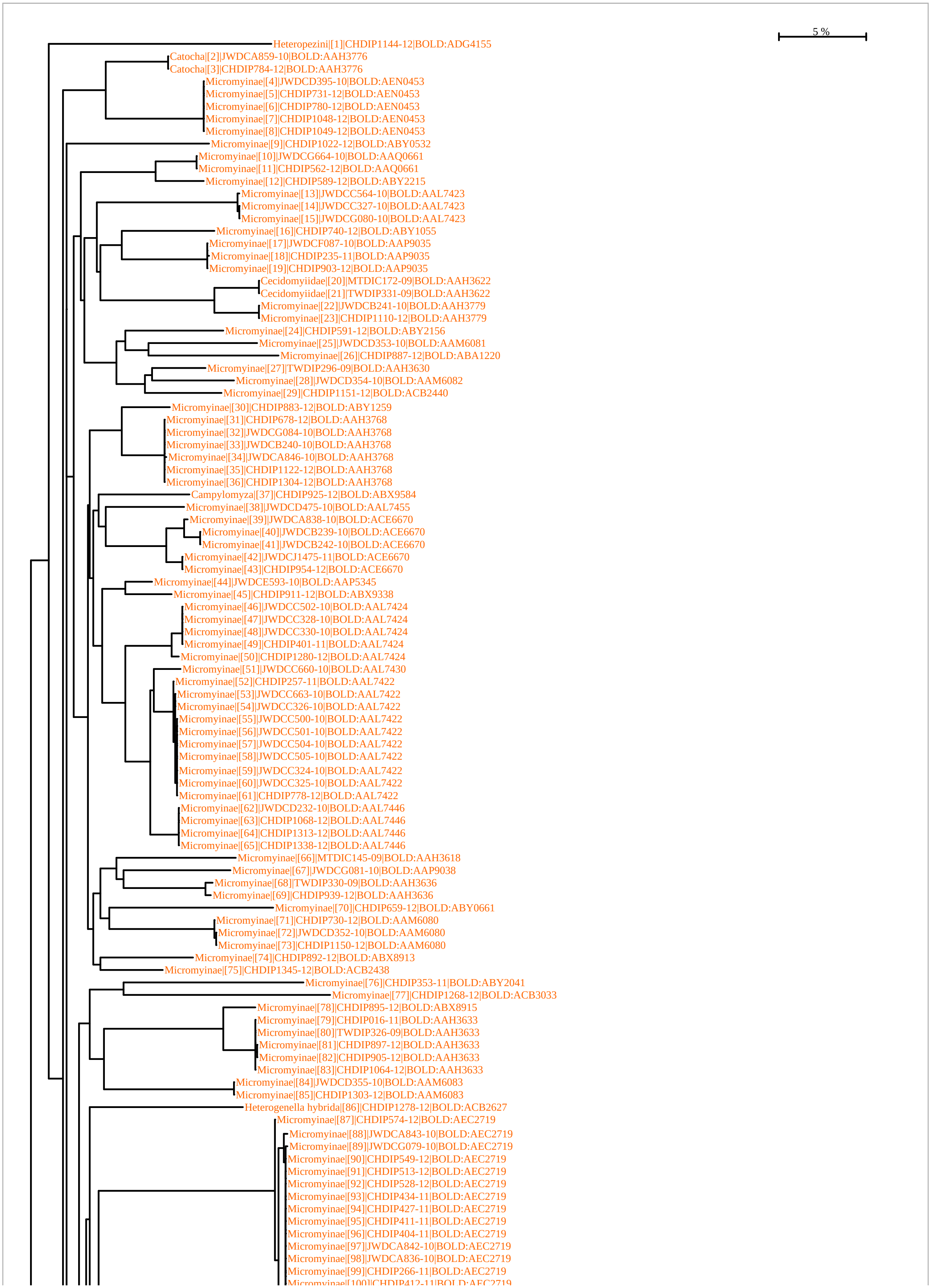
5 %
Heteropezinini|[1]|CHDIP1144-12|BOLD:ADG4155
Catocha|[2]|JWDCA859-10|BOLD:AAH3776
Catocha|[3]|CHDIP784-12|BOLD:AAH3776
Micromyinae|[4]|JWDCD395-10|BOLD:AEN0453
Micromyinae|[5]|CHDIP731-12|BOLD:AEN0453
Micromyinae|[6]|CHDIP780-12|BOLD:AEN0453
Micromyinae|[7]|CHDIP1048-12|BOLD:AEN0453
Micromyinae|[8]|CHDIP1049-12|BOLD:AEN0453
Micromyinae|[9]|CHDIP1022-12|BOLD:ABY0532
Micromyinae|[10]|JWDCG664-10|BOLD:AAQ0661
Micromyinae|[11]|CHDIP562-12|BOLD:AAQ0661
Micromyinae|[12]|CHDIP589-12|BOLD:ABY2215
Micromyinae|[13]|JWDCC564-10|BOLD:AAL7423
Micromyinae|[14]|JWDCC327-10|BOLD:AAL7423
Micromyinae|[15]|JWDCG080-10|BOLD:AAL7423
Micromyinae|[16]|CHDIP740-12|BOLD:ABY1055
Micromyinae|[17]|JWDCF087-10|BOLD:AAP9035
Micromyinae|[18]|CHDIP235-11|BOLD:AAP9035
Micromyinae|[19]|CHDIP903-12|BOLD:AAP9035
Cecidomyiidae|[20]|MTDIC172-09|BOLD:AAH3622
Cecidomyiidae|[21]|TWDIP331-09|BOLD:AAH3622
Micromyinae|[22]|JWDCB241-10|BOLD:AAH3779
Micromyinae|[23]|CHDIP1110-12|BOLD:AAH3779
Micromyinae|[24]|CHDIP591-12|BOLD:ABY2156
Micromyinae|[25]|JWDCD353-10|BOLD:AAM6081
Micromyinae|[26]|CHDIP887-12|BOLD:ABA1220
Micromyinae|[27]|TWDIP296-09|BOLD:AAH3630
Micromyinae|[28]|JWDCD354-10|BOLD:AAM6082
Micromyinae|[29]|CHDIP1151-12|BOLD:ACB2440
Micromyinae|[30]|CHDIP883-12|BOLD:ABY1259
Micromyinae|[31]|CHDIP678-12|BOLD:AAH3768
Micromyinae|[32]|JWDCG084-10|BOLD:AAH3768
Micromyinae|[33]|JWDCB240-10|BOLD:AAH3768
Micromyinae|[34]|JWDCA846-10|BOLD:AAH3768
Micromyinae|[35]|CHDIP1122-12|BOLD:AAH3768
Micromyinae|[36]|CHDIP1304-12|BOLD:AAH3768
Campylomyza|[37]|CHDIP925-12|BOLD:ABX9584
Micromyinae|[38]|JWDCD475-10|BOLD:AAL7455
Micromyinae|[39]|JWDCA838-10|BOLD:ACE6670
Micromyinae|[40]|JWDCB239-10|BOLD:ACE6670
Micromyinae|[41]|JWDCB242-10|BOLD:ACE6670
Micromyinae|[42]|JWDCJ1475-11|BOLD:ACE6670
Micromyinae|[43]|CHDIP954-12|BOLD:ACE6670
Micromyinae|[44]|JWDCE593-10|BOLD:AAP5345
Micromyinae|[45]|CHDIP911-12|BOLD:ABX9338
Micromyinae|[46]|JWDCC502-10|BOLD:AAL7424
Micromyinae|[47]|JWDCC328-10|BOLD:AAL7424
Micromyinae|[48]|JWDCC330-10|BOLD:AAL7424
Micromyinae|[49]|CHDIP401-11|BOLD:AAL7424
Micromyinae|[50]|CHDIP1280-12|BOLD:AAL7424
Micromyinae|[51]|JWDCC660-10|BOLD:AAL7430
Micromyinae|[52]|CHDIP257-11|BOLD:AAL7422
Micromyinae|[53]|JWDCC663-10|BOLD:AAL7422
Micromyinae|[54]|JWDCC326-10|BOLD:AAL7422
Micromyinae|[55]|JWDCC500-10|BOLD:AAL7422
Micromyinae|[56]|JWDCC501-10|BOLD:AAL7422
Micromyinae|[57]|JWDCC504-10|BOLD:AAL7422
Micromyinae|[58]|JWDCC505-10|BOLD:AAL7422
Micromyinae|[59]|JWDCC324-10|BOLD:AAL7422
Micromyinae|[60]|JWDCC325-10|BOLD:AAL7422
Micromyinae|[61]|CHDIP778-12|BOLD:AAL7422
Micromyinae|[62]|JWDCD232-10|BOLD:AAL7446
Micromyinae|[63]|CHDIP1068-12|BOLD:AAL7446
Micromyinae|[64]|CHDIP1313-12|BOLD:AAL7446
Micromyinae|[65]|CHDIP1338-12|BOLD:AAL7446
Micromyinae|[66]|MTDIC145-09|BOLD:AAH3618
Micromyinae|[67]|JWDCG081-10|BOLD:AAP9038
Micromyinae|[68]|TWDIP330-09|BOLD:AAH3636
Micromyinae|[69]|CHDIP939-12|BOLD:AAH3636
Micromyinae|[70]|CHDIP659-12|BOLD:ABY0661
Micromyinae|[71]|CHDIP730-12|BOLD:AAM6080
Micromyinae|[72]|JWDCD352-10|BOLD:AAM6080
Micromyinae|[73]|CHDIP1150-12|BOLD:AAM6080
Micromyinae|[74]|CHDIP892-12|BOLD:ABX8913
Micromyinae|[75]|CHDIP1345-12|BOLD:ACB2438
Micromyinae|[76]|CHDIP353-11|BOLD:ABY2041
Micromyinae|[77]|CHDIP1268-12|BOLD:ACB3033
Micromyinae|[78]|CHDIP895-12|BOLD:ABX8915
Micromyinae|[79]|CHDIP016-11|BOLD:AAH3633
Micromyinae|[80]|TWDIP326-09|BOLD:AAH3633
Micromyinae|[81]|CHDIP897-12|BOLD:AAH3633
Micromyinae|[82]|CHDIP905-12|BOLD:AAH3633
Micromyinae|[83]|CHDIP1064-12|BOLD:AAH3633
Micromyinae|[84]|JWDCD355-10|BOLD:AAM6083
Micromyinae|[85]|CHDIP1303-12|BOLD:AAM6083
Heterogenella hybrida|[86]|CHDIP1278-12|BOLD:ACB2627
Micromyinae|[87]|CHDIP574-12|BOLD:AEC2719
Micromyinae|[88]|JWDCA843-10|BOLD:AEC2719
Micromyinae|[89]|JWDCG079-10|BOLD:AEC2719
Micromyinae|[90]|CHDIP549-12|BOLD:AEC2719
Micromyinae|[91]|CHDIP513-12|BOLD:AEC2719
Micromyinae|[92]|CHDIP528-12|BOLD:AEC2719
Micromyinae|[93]|CHDIP434-11|BOLD:AEC2719
Micromyinae|[94]|CHDIP427-11|BOLD:AEC2719
Micromyinae|[95]|CHDIP411-11|BOLD:AEC2719
Micromyinae|[96]|CHDIP404-11|BOLD:AEC2719
Micromyinae|[97]|JWDCA842-10|BOLD:AEC2719
Micromyinae|[98]|JWDCA836-10|BOLD:AEC2719
Micromyinae|[99]|CHDIP266-11|BOLD:AEC2719

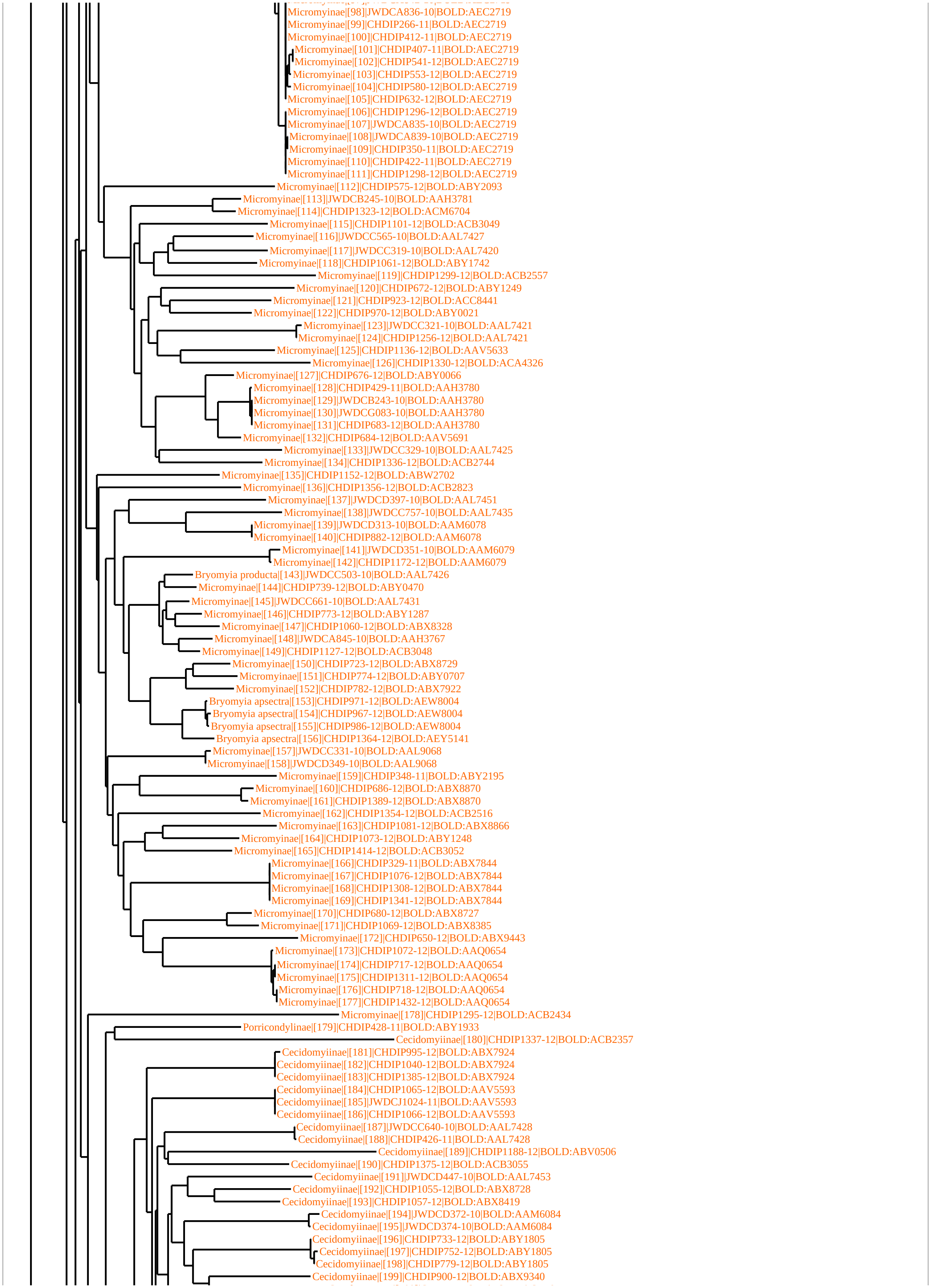
Micromyinae|[98]|JWDCA836-10|BOLD:AEC2719
Micromyinae|[99]|CHDIP266-11|BOLD:AEC2719
Micromyinae|[100]|CHDIP412-11|BOLD:AEC2719
Micromyinae|[101]|CHDIP407-11|BOLD:AEC2719
Micromyinae|[102]|CHDIP541-12|BOLD:AEC2719
Micromyinae|[103]|CHDIP553-12|BOLD:AEC2719
Micromyinae|[104]|CHDIP580-12|BOLD:AEC2719
Micromyinae|[105]|CHDIP632-12|BOLD:AEC2719
Micromyinae|[106]|CHDIP1296-12|BOLD:AEC2719
Micromyinae|[107]|JWDCA835-10|BOLD:AEC2719
Micromyinae|[108]|JWDCA839-10|BOLD:AEC2719
Micromyinae|[109]|CHDIP350-11|BOLD:AEC2719
Micromyinae|[110]|CHDIP422-11|BOLD:AEC2719
Micromyinae|[111]|CHDIP1298-12|BOLD:AEC2719
Micromyinae|[112]|CHDIP575-12|BOLD:ABY2093
Micromyinae|[113]|JWDCB245-10|BOLD:AAH3781
Micromyinae|[114]|CHDIP1323-12|BOLD:ACM6704
Micromyinae|[115]|CHDIP1101-12|BOLD:ACB3049
Micromyinae|[116]|JWDCC565-10|BOLD:AAL7427
Micromyinae|[117]|JWDCC319-10|BOLD:AAL7420
Micromyinae|[118]|CHDIP1061-12|BOLD:ABY1742
Micromyinae|[119]|CHDIP1299-12|BOLD:ACB2557
Micromyinae|[120]|CHDIP672-12|BOLD:ABY1249
Micromyinae|[121]|CHDIP923-12|BOLD:ACC8441
Micromyinae|[122]|CHDIP970-12|BOLD:ABY0021
Micromyinae|[123]|JWDCC321-10|BOLD:AAL7421
Micromyinae|[124]|CHDIP1256-12|BOLD:AAL7421
Micromyinae|[125]|CHDIP1136-12|BOLD:AAV5633
Micromyinae|[126]|CHDIP1330-12|BOLD:ACA4326
Micromyinae|[127]|CHDIP676-12|BOLD:ABY0066
Micromyinae|[128]|CHDIP429-11|BOLD:AAH3780
Micromyinae|[129]|JWDCB243-10|BOLD:AAH3780
Micromyinae|[130]|JWDCG083-10|BOLD:AAH3780
Micromyinae|[131]|CHDIP683-12|BOLD:AAH3780
Micromyinae|[132]|CHDIP684-12|BOLD:AAV5691
Micromyinae|[133]|JWDCC329-10|BOLD:AAL7425
Micromyinae|[134]|CHDIP1336-12|BOLD:ACB2744
Micromyinae|[135]|CHDIP1152-12|BOLD:ABW2702
Micromyinae|[136]|CHDIP1356-12|BOLD:ACB2823
Micromyinae|[137]|JWDCD397-10|BOLD:AAL7451
Micromyinae|[138]|JWDCC757-10|BOLD:AAL7435
Micromyinae|[139]|JWDCD313-10|BOLD:AAM6078
Micromyinae|[140]|CHDIP882-12|BOLD:AAM6078
Micromyinae|[141]|JWDCD351-10|BOLD:AAM6079
Micromyinae|[142]|CHDIP1172-12|BOLD:AAM6079
Bryomyia producta|[143]|JWDCC503-10|BOLD:AAL7426
Micromyinae|[144]|CHDIP739-12|BOLD:ABY0470
Micromyinae|[145]|JWDCC661-10|BOLD:AAL7431
Micromyinae|[146]|CHDIP773-12|BOLD:ABY1287
Micromyinae|[147]|CHDIP1060-12|BOLD:ABX8328
Micromyinae|[148]|JWDCA845-10|BOLD:AAH3767
Micromyinae|[149]|CHDIP1127-12|BOLD:ACB3048
Micromyinae|[150]|CHDIP723-12|BOLD:ABX8729
Micromyinae|[151]|CHDIP774-12|BOLD:ABY0707
Micromyinae|[152]|CHDIP782-12|BOLD:ABX7922
Bryomyia apsectra|[153]|CHDIP971-12|BOLD:AEW8004
Bryomyia apsectra|[154]|CHDIP967-12|BOLD:AEW8004
Bryomyia apsectra|[155]|CHDIP986-12|BOLD:AEW8004
Bryomyia apsectra|[156]|CHDIP1364-12|BOLD:AEY5141
Micromyinae|[157]|JWDCC331-10|BOLD:AAL9068
Micromyinae|[158]|JWDCD349-10|BOLD:AAL9068
Micromyinae|[159]|CHDIP348-11|BOLD:ABY2195
Micromyinae|[160]|CHDIP686-12|BOLD:ABX8870
Micromyinae|[161]|CHDIP1389-12|BOLD:ABX8870
Micromyinae|[162]|CHDIP1354-12|BOLD:ACB2516
Micromyinae|[163]|CHDIP1081-12|BOLD:ABX8866
Micromyinae|[164]|CHDIP1073-12|BOLD:ABY1248
Micromyinae|[165]|CHDIP1414-12|BOLD:ACB3052
Micromyinae|[166]|CHDIP329-11|BOLD:ABX7844
Micromyinae|[167]|CHDIP1076-12|BOLD:ABX7844
Micromyinae|[168]|CHDIP1308-12|BOLD:ABX7844
Micromyinae|[169]|CHDIP1341-12|BOLD:ABX7844
Micromyinae|[170]|CHDIP680-12|BOLD:ABX8727
Micromyinae|[171]|CHDIP1069-12|BOLD:ABX8385
Micromyinae|[172]|CHDIP650-12|BOLD:ABX9443
Micromyinae|[173]|CHDIP1072-12|BOLD:AAQ0654
Micromyinae|[174]|CHDIP717-12|BOLD:AAQ0654
Micromyinae|[175]|CHDIP1311-12|BOLD:AAQ0654
Micromyinae|[176]|CHDIP718-12|BOLD:AAQ0654
Micromyinae|[177]|CHDIP1432-12|BOLD:AAQ0654
Micromyinae|[178]|CHDIP1295-12|BOLD:ACB2434
Porricondylinae|[179]|CHDIP428-11|BOLD:ABY1933
Cecidomyiinae|[180]|CHDIP1337-12|BOLD:ACB2357
Cecidomyiinae|[181]|CHDIP995-12|BOLD:ABX7924
Cecidomyiinae|[182]|CHDIP1040-12|BOLD:ABX7924
Cecidomyiinae|[183]|CHDIP1385-12|BOLD:ABX7924
Cecidomyiinae|[184]|CHDIP1065-12|BOLD:AAV5593
Cecidomyiinae|[185]|JWDCJ1024-11|BOLD:AAV5593
Cecidomyiinae|[186]|CHDIP1066-12|BOLD:AAV5593
Cecidomyiinae|[187]|JWDCC640-10|BOLD:AAL7428
Cecidomyiinae|[188]|CHDIP426-11|BOLD:AAL7428
Cecidomyiinae|[189]|CHDIP1188-12|BOLD:ABV0506
Cecidomyiinae|[190]|CHDIP1375-12|BOLD:ACB3055
Cecidomyiinae|[191]|JWDCD447-10|BOLD:AAL7453
Cecidomyiinae|[192]|CHDIP1055-12|BOLD:ABX8728
Cecidomyiinae|[193]|CHDIP1057-12|BOLD:ABX8419
Cecidomyiinae|[194]|JWDCD372-10|BOLD:AAM6084
Cecidomyiinae|[195]|JWDCD374-10|BOLD:AAM6084
Cecidomyiinae|[196]|CHDIP733-12|BOLD:ABY1805
Cecidomyiinae|[197]|CHDIP752-12|BOLD:ABY1805
Cecidomyiinae|[198]|CHDIP779-12|BOLD:ABY1805
Cecidomyiinae|[199]|CHDIP900-12|BOLD:ABX9340

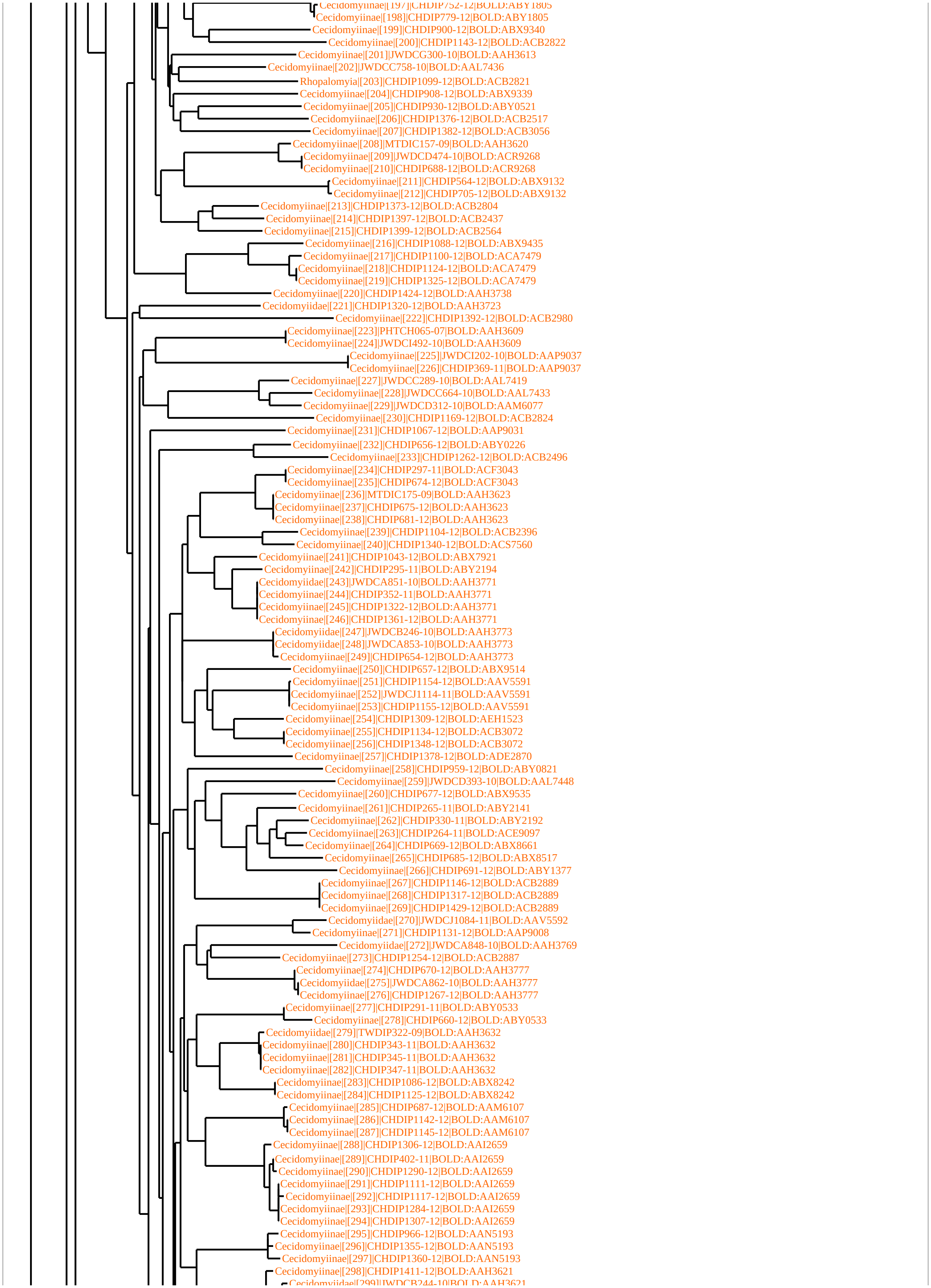
Cecidomyiinae|[197]|CHDIP752-12|BOLD:ABY1805
Cecidomyiinae|[198]|CHDIP779-12|BOLD:ABY1805
Cecidomyiinae|[199]|CHDIP900-12|BOLD:ABX9340
Cecidomyiinae|[200]|CHDIP1143-12|BOLD:ACB2822
Cecidomyiinae|[201]|JWDCG300-10|BOLD:AAH3613
Cecidomyiinae|[202]|JWDCC758-10|BOLD:AAL7436
Rhopalomyia|[203]|CHDIP1099-12|BOLD:ACB2821
Cecidomyiinae|[204]|CHDIP908-12|BOLD:ABX9339
Cecidomyiinae|[205]|CHDIP930-12|BOLD:ABY0521
Cecidomyiinae|[206]|CHDIP1376-12|BOLD:ACB2517
Cecidomyiinae|[207]|CHDIP1382-12|BOLD:ACB3056
Cecidomyiinae|[208]|MTDIC157-09|BOLD:AAH3620
Cecidomyiinae|[209]|JWDCD474-10|BOLD:ACR9268
Cecidomyiinae|[210]|CHDIP688-12|BOLD:ACR9268
Cecidomyiinae|[211]|CHDIP564-12|BOLD:ABX9132
Cecidomyiinae|[212]|CHDIP705-12|BOLD:ABX9132
Cecidomyiinae|[213]|CHDIP1373-12|BOLD:ACB2804
Cecidomyiinae|[214]|CHDIP1397-12|BOLD:ACB2437
Cecidomyiinae|[215]|CHDIP1399-12|BOLD:ACB2564
Cecidomyiinae|[216]|CHDIP1088-12|BOLD:ABX9435
Cecidomyiinae|[217]|CHDIP1100-12|BOLD:ACA7479
Cecidomyiinae|[218]|CHDIP1124-12|BOLD:ACA7479
Cecidomyiinae|[219]|CHDIP1325-12|BOLD:ACA7479
Cecidomyiinae|[220]|CHDIP1424-12|BOLD:AAH3738
Cecidomyiidae|[221]|CHDIP1320-12|BOLD:AAH3723
Cecidomyiinae|[222]|CHDIP1392-12|BOLD:ACB2980
Cecidomyiinae|[223]|PHTCH065-07|BOLD:AAH3609
Cecidomyiinae|[224]|JWDCI492-10|BOLD:AAH3609
Cecidomyiinae|[225]|JWDCI202-10|BOLD:AAP9037
Cecidomyiinae|[226]|CHDIP369-11|BOLD:AAP9037
Cecidomyiinae|[227]|JWDCC289-10|BOLD:AAL7419
Cecidomyiinae|[228]|JWDCC664-10|BOLD:AAL7433
Cecidomyiinae|[229]|JWDCD312-10|BOLD:AAM6077
Cecidomyiinae|[230]|CHDIP1169-12|BOLD:ACB2824
Cecidomyiinae|[231]|CHDIP1067-12|BOLD:AAP9031
Cecidomyiinae|[232]|CHDIP656-12|BOLD:ABY0226
Cecidomyiinae|[233]|CHDIP1262-12|BOLD:ACB2496
Cecidomyiinae|[234]|CHDIP297-11|BOLD:ACF3043
Cecidomyiinae|[235]|CHDIP674-12|BOLD:ACF3043
Cecidomyiinae|[236]|MTDIC175-09|BOLD:AAH3623
Cecidomyiinae|[237]|CHDIP675-12|BOLD:AAH3623
Cecidomyiinae|[238]|CHDIP681-12|BOLD:AAH3623
Cecidomyiinae|[239]|CHDIP1104-12|BOLD:ACB2396
Cecidomyiinae|[240]|CHDIP1340-12|BOLD:ACS7560
Cecidomyiinae|[241]|CHDIP1043-12|BOLD:ABX7921
Cecidomyiinae|[242]|CHDIP295-11|BOLD:ABY2194
Cecidomyiidae|[243]|JWDCA851-10|BOLD:AAH3771
Cecidomyiinae|[244]|CHDIP352-11|BOLD:AAH3771
Cecidomyiinae|[245]|CHDIP1322-12|BOLD:AAH3771
Cecidomyiinae|[246]|CHDIP1361-12|BOLD:AAH3771
Cecidomyiidae|[247]|JWDCB246-10|BOLD:AAH3773
Cecidomyiidae|[248]|JWDCA853-10|BOLD:AAH3773
Cecidomyiinae|[249]|CHDIP654-12|BOLD:AAH3773
Cecidomyiinae|[250]|CHDIP657-12|BOLD:ABX9514
Cecidomyiinae|[251]|CHDIP1154-12|BOLD:AAV5591
Cecidomyiinae|[252]|JWDCJ1114-11|BOLD:AAV5591
Cecidomyiinae|[253]|CHDIP1155-12|BOLD:AAV5591
Cecidomyiinae|[254]|CHDIP1309-12|BOLD:AEH1523
Cecidomyiinae|[255]|CHDIP1134-12|BOLD:ACB3072
Cecidomyiinae|[256]|CHDIP1348-12|BOLD:ACB3072
Cecidomyiinae|[257]|CHDIP1378-12|BOLD:ADE2870
Cecidomyiinae|[258]|CHDIP959-12|BOLD:ABY0821
Cecidomyiinae|[259]|JWDCD393-10|BOLD:AAL7448
Cecidomyiinae|[260]|CHDIP677-12|BOLD:ABX9535
Cecidomyiinae|[261]|CHDIP265-11|BOLD:ABY2141
Cecidomyiinae|[262]|CHDIP330-11|BOLD:ABY2192
Cecidomyiinae|[263]|CHDIP264-11|BOLD:ACE9097
Cecidomyiinae|[264]|CHDIP669-12|BOLD:ABX8661
Cecidomyiinae|[265]|CHDIP685-12|BOLD:ABX8517
Cecidomyiinae|[266]|CHDIP691-12|BOLD:ABY1377
Cecidomyiinae|[267]|CHDIP1146-12|BOLD:ACB2889
Cecidomyiinae|[268]|CHDIP1317-12|BOLD:ACB2889
Cecidomyiinae|[269]|CHDIP1429-12|BOLD:ACB2889
Cecidomyiidae|[270]|JWDCJ1084-11|BOLD:AAV5592
Cecidomyiinae|[271]|CHDIP1131-12|BOLD:AAP9008
Cecidomyiidae|[272]|JWDCA848-10|BOLD:AAH3769
Cecidomyiinae|[273]|CHDIP1254-12|BOLD:ACB2887
Cecidomyiinae|[274]|CHDIP670-12|BOLD:AAH3777
Cecidomyiidae|[275]|JWDCA862-10|BOLD:AAH3777
Cecidomyiinae|[276]|CHDIP1267-12|BOLD:AAH3777
Cecidomyiinae|[277]|CHDIP291-11|BOLD:ABY0533
Cecidomyiinae|[278]|CHDIP660-12|BOLD:ABY0533
Cecidomyiidae|[279]|TWDIP322-09|BOLD:AAH3632
Cecidomyiinae|[280]|CHDIP343-11|BOLD:AAH3632
Cecidomyiinae|[281]|CHDIP345-11|BOLD:AAH3632
Cecidomyiinae|[282]|CHDIP347-11|BOLD:AAH3632
Cecidomyiinae|[283]|CHDIP1086-12|BOLD:ABX8242
Cecidomyiinae|[284]|CHDIP1125-12|BOLD:ABX8242
Cecidomyiinae|[285]|CHDIP687-12|BOLD:AAM6107
Cecidomyiinae|[286]|CHDIP1142-12|BOLD:AAM6107
Cecidomyiinae|[287]|CHDIP1145-12|BOLD:AAM6107
Cecidomyiinae|[288]|CHDIP1306-12|BOLD:AAI2659
Cecidomyiinae|[289]|CHDIP402-11|BOLD:AAI2659
Cecidomyiinae|[290]|CHDIP1290-12|BOLD:AAI2659
Cecidomyiinae|[291]|CHDIP1111-12|BOLD:AAI2659
Cecidomyiinae|[292]|CHDIP1117-12|BOLD:AAI2659
Cecidomyiinae|[293]|CHDIP1284-12|BOLD:AAI2659
Cecidomyiinae|[294]|CHDIP1307-12|BOLD:AAI2659
Cecidomyiinae|[295]|CHDIP966-12|BOLD:AAN5193
Cecidomyiinae|[296]|CHDIP1355-12|BOLD:AAN5193
Cecidomyiinae|[297]|CHDIP1360-12|BOLD:AAN5193
Cecidomyiinae|[298]|CHDIP1411-12|BOLD:AAH3621
Cecidomyiidae|[299]|JWDCB244-10|BOLD:AAH3621

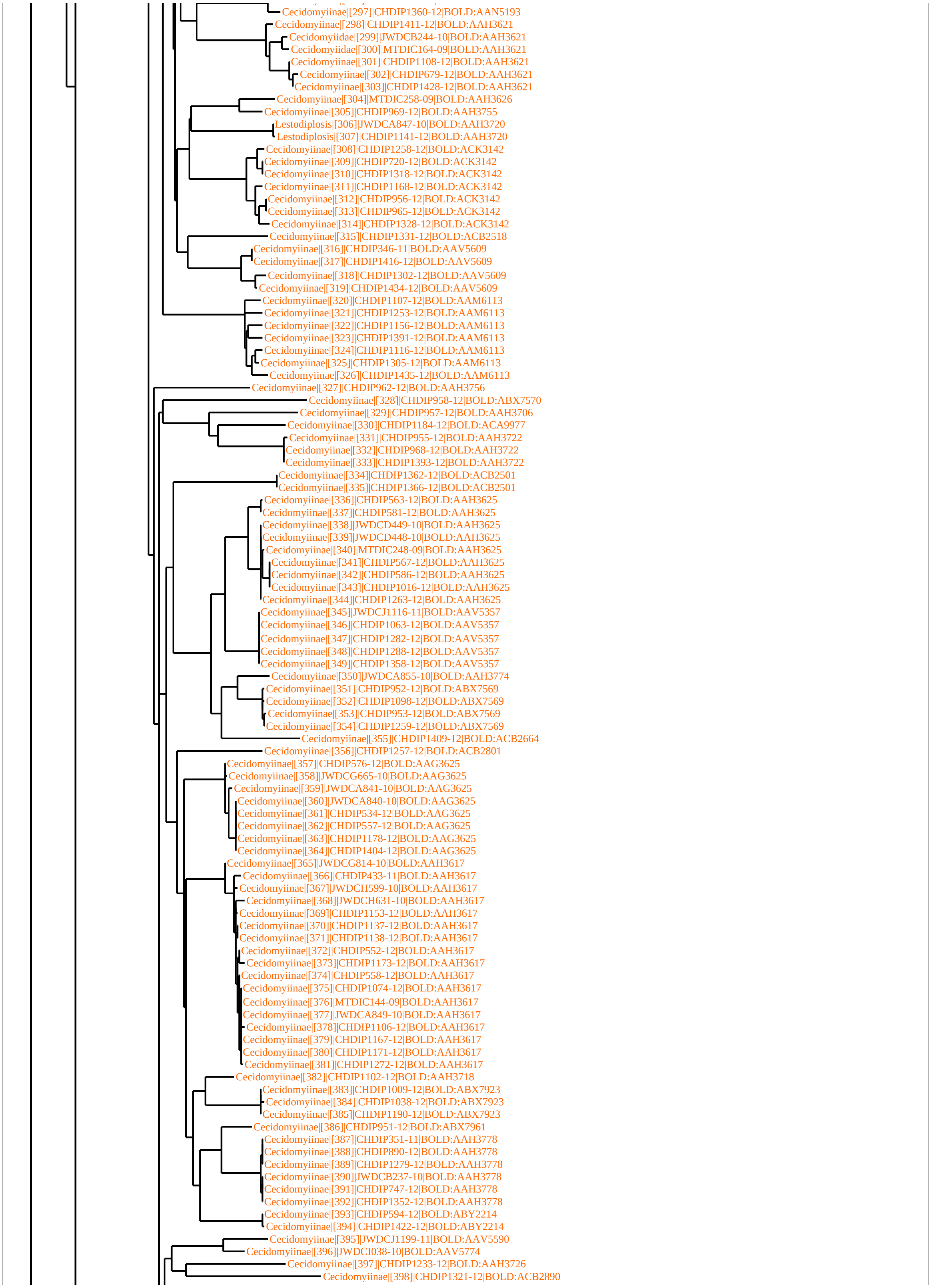
Cecidomyiinae|[297]|CHDIP1360-12|BOLD:AAN5193
Cecidomyiinae|[298]|CHDIP1411-12|BOLD:AAH3621
Cecidomyiidae|[299]|JWDCB244-10|BOLD:AAH3621
Cecidomyiidae|[300]|MTDIC164-09|BOLD:AAH3621
Cecidomyiinae|[301]|CHDIP1108-12|BOLD:AAH3621
Cecidomyiinae|[302]|CHDIP679-12|BOLD:AAH3621
Cecidomyiinae|[303]|CHDIP1428-12|BOLD:AAH3621
Cecidomyiinae|[304]|MTDIC258-09|BOLD:AAH3626
Cecidomyiinae|[305]|CHDIP969-12|BOLD:AAH3755
Lestodiplosis|[306]|JWDCA847-10|BOLD:AAH3720
Lestodiplosis|[307]|CHDIP1141-12|BOLD:AAH3720
Cecidomyiinae|[308]|CHDIP1258-12|BOLD:ACK3142
Cecidomyiinae|[309]|CHDIP720-12|BOLD:ACK3142
Cecidomyiinae|[310]|CHDIP1318-12|BOLD:ACK3142
Cecidomyiinae|[311]|CHDIP1168-12|BOLD:ACK3142
Cecidomyiinae|[312]|CHDIP956-12|BOLD:ACK3142
Cecidomyiinae|[313]|CHDIP965-12|BOLD:ACK3142
Cecidomyiinae|[314]|CHDIP1328-12|BOLD:ACK3142
Cecidomyiinae|[315]|CHDIP1331-12|BOLD:ACB2518
Cecidomyiinae|[316]|CHDIP346-11|BOLD:AAV5609
Cecidomyiinae|[317]|CHDIP1416-12|BOLD:AAV5609
Cecidomyiinae|[318]|CHDIP1302-12|BOLD:AAV5609
Cecidomyiinae|[319]|CHDIP1434-12|BOLD:AAV5609
Cecidomyiinae|[320]|CHDIP1107-12|BOLD:AAM6113
Cecidomyiinae|[321]|CHDIP1253-12|BOLD:AAM6113
Cecidomyiinae|[322]|CHDIP1156-12|BOLD:AAM6113
Cecidomyiinae|[323]|CHDIP1391-12|BOLD:AAM6113
Cecidomyiinae|[324]|CHDIP1116-12|BOLD:AAM6113
Cecidomyiinae|[325]|CHDIP1305-12|BOLD:AAM6113
Cecidomyiinae|[326]|CHDIP1435-12|BOLD:AAM6113
Cecidomyiinae|[327]|CHDIP962-12|BOLD:AAH3756
Cecidomyiinae|[328]|CHDIP958-12|BOLD:ABX7570
Cecidomyiinae|[329]|CHDIP957-12|BOLD:AAH3706
Cecidomyiinae|[330]|CHDIP1184-12|BOLD:ACA9977
Cecidomyiinae|[331]|CHDIP955-12|BOLD:AAH3722
Cecidomyiinae|[332]|CHDIP968-12|BOLD:AAH3722
Cecidomyiinae|[333]|CHDIP1393-12|BOLD:AAH3722
Cecidomyiinae|[334]|CHDIP1362-12|BOLD:ACB2501
Cecidomyiinae|[335]|CHDIP1366-12|BOLD:ACB2501
Cecidomyiinae|[336]|CHDIP563-12|BOLD:AAH3625
Cecidomyiinae|[337]|CHDIP581-12|BOLD:AAH3625
Cecidomyiinae|[338]|JWDCD449-10|BOLD:AAH3625
Cecidomyiinae|[339]|JWDCD448-10|BOLD:AAH3625
Cecidomyiinae|[340]|MTDIC248-09|BOLD:AAH3625
Cecidomyiinae|[341]|CHDIP567-12|BOLD:AAH3625
Cecidomyiinae|[342]|CHDIP586-12|BOLD:AAH3625
Cecidomyiinae|[343]|CHDIP1016-12|BOLD:AAH3625
Cecidomyiinae|[344]|CHDIP1263-12|BOLD:AAH3625
Cecidomyiinae|[345]|JWDCJ1116-11|BOLD:AAV5357
Cecidomyiinae|[346]|CHDIP1063-12|BOLD:AAV5357
Cecidomyiinae|[347]|CHDIP1282-12|BOLD:AAV5357
Cecidomyiinae|[348]|CHDIP1288-12|BOLD:AAV5357
Cecidomyiinae|[349]|CHDIP1358-12|BOLD:AAV5357
Cecidomyiinae|[350]|JWDCA855-10|BOLD:AAH3774
Cecidomyiinae|[351]|CHDIP952-12|BOLD:ABX7569
Cecidomyiinae|[352]|CHDIP1098-12|BOLD:ABX7569
Cecidomyiinae|[353]|CHDIP953-12|BOLD:ABX7569
Cecidomyiinae|[354]|CHDIP1259-12|BOLD:ABX7569
Cecidomyiinae|[355]|CHDIP1409-12|BOLD:ACB2664
Cecidomyiinae|[356]|CHDIP1257-12|BOLD:ACB2801
Cecidomyiinae|[357]|CHDIP576-12|BOLD:AAG3625
Cecidomyiinae|[358]|JWDCG665-10|BOLD:AAG3625
Cecidomyiinae|[359]|JWDCA841-10|BOLD:AAG3625
Cecidomyiinae|[360]|JWDCA840-10|BOLD:AAG3625
Cecidomyiinae|[361]|CHDIP534-12|BOLD:AAG3625
Cecidomyiinae|[362]|CHDIP557-12|BOLD:AAG3625
Cecidomyiinae|[363]|CHDIP1178-12|BOLD:AAG3625
Cecidomyiinae|[364]|CHDIP1404-12|BOLD:AAG3625
Cecidomyiinae|[365]|JWDCG814-10|BOLD:AAH3617
Cecidomyiinae|[366]|CHDIP433-11|BOLD:AAH3617
Cecidomyiinae|[367]|JWDCH599-10|BOLD:AAH3617
Cecidomyiinae|[368]|JWDCH631-10|BOLD:AAH3617
Cecidomyiinae|[369]|CHDIP1153-12|BOLD:AAH3617
Cecidomyiinae|[370]|CHDIP1137-12|BOLD:AAH3617
Cecidomyiinae|[371]|CHDIP1138-12|BOLD:AAH3617
Cecidomyiinae|[372]|CHDIP552-12|BOLD:AAH3617
Cecidomyiinae|[373]|CHDIP1173-12|BOLD:AAH3617
Cecidomyiinae|[374]|CHDIP558-12|BOLD:AAH3617
Cecidomyiinae|[375]|CHDIP1074-12|BOLD:AAH3617
Cecidomyiinae|[376]|MTDIC144-09|BOLD:AAH3617
Cecidomyiinae|[377]|JWDCA849-10|BOLD:AAH3617
Cecidomyiinae|[378]|CHDIP1106-12|BOLD:AAH3617
Cecidomyiinae|[379]|CHDIP1167-12|BOLD:AAH3617
Cecidomyiinae|[380]|CHDIP1171-12|BOLD:AAH3617
Cecidomyiinae|[381]|CHDIP1272-12|BOLD:AAH3617
Cecidomyiinae|[382]|CHDIP1102-12|BOLD:AAH3718
Cecidomyiinae|[383]|CHDIP1009-12|BOLD:ABX7923
Cecidomyiinae|[384]|CHDIP1038-12|BOLD:ABX7923
Cecidomyiinae|[385]|CHDIP1190-12|BOLD:ABX7923
Cecidomyiinae|[386]|CHDIP951-12|BOLD:ABX7961
Cecidomyiinae|[387]|CHDIP351-11|BOLD:AAH3778
Cecidomyiinae|[388]|CHDIP890-12|BOLD:AAH3778
Cecidomyiinae|[389]|CHDIP1279-12|BOLD:AAH3778
Cecidomyiinae|[390]|JWDCB237-10|BOLD:AAH3778
Cecidomyiinae|[391]|CHDIP747-12|BOLD:AAH3778
Cecidomyiinae|[392]|CHDIP1352-12|BOLD:AAH3778
Cecidomyiinae|[393]|CHDIP594-12|BOLD:ABY2214
Cecidomyiinae|[394]|CHDIP1422-12|BOLD:ABY2214
Cecidomyiinae|[395]|JWDCJ1199-11|BOLD:AAV5590
Cecidomyiinae|[396]|JWDCI038-10|BOLD:AAV5774
Cecidomyiinae|[397]|CHDIP1233-12|BOLD:AAH3726
Cecidomyiinae|[398]|CHDIP1321-12|BOLD:ACB2890

Cecidomyiinae|[396]|JWDCI038-10|BOLD:AAV5774
Cecidomyiinae|[397]|CHDIP1233-12|BOLD:AAH3726
Cecidomyiinae|[398]|CHDIP1321-12|BOLD:ACB2890
Cecidomyiinae|[399]|CHDIP640-12|BOLD:ABX8097
Cecidomyiinae|[400]|CHDIP413-11|BOLD:AAH3631
Cecidomyiinae|[401]|TWDIP321-09|BOLD:AAH3631
Cecidomyiinae|[402]|CHDIP716-12|BOLD:AAH3631
Cecidomyiinae|[403]|CHDIP1324-12|BOLD:ACB2888
Cecidomyiinae|[404]|CHDIP1181-12|BOLD:ACB2377
Cecidomyiinae|[405]|CHDIP1353-12|BOLD:ACB2377
Cecidomyiidae|[406]|JWDCA850-10|BOLD:AAH3770
Cecidomyiinae|[407]|CHDIP1161-12|BOLD:ABZ2405
Cecidomyiinae|[408]|CHDIP1095-12|BOLD:ABZ2405
Cecidomyiinae|[409]|CHDIP1182-12|BOLD:ABZ2405
Cecidomyiinae|[410]|CHDIP1191-12|BOLD:ABZ2405
Cecidomyiinae|[411]|CHDIP1319-12|BOLD:ABZ2405
Cecidomyiinae|[412]|CHDIP1344-12|BOLD:ABZ2405
Cecidomyiinae|[413]|CHDIP1347-12|BOLD:ABZ2405
Cecidomyiinae|[414]|CHDIP1349-12|BOLD:ABZ2405
Cecidomyiinae|[415]|CHDIP1359-12|BOLD:ABZ2405
Cecidomyiidae|[416]|CHDIP569-12|BOLD:AAN5190
Cecidomyiinae|[417]|CHDIP585-12|BOLD:ABY2094
Cecidomyiinae|[418]|CHDIP1427-12|BOLD:ACB2619
Cecidomyiinae|[419]|JWDCD473-10|BOLD:AAL7454
Cecidomyiinae|[420]|CHDIP1029-12|BOLD:ABX9583
Cecidomyiinae|[421]|CHDIP972-12|BOLD:ABX9218
Cecidomyiinae|[422]|MTDIC156-09|BOLD:AAH3619
Cecidomyiinae|[423]|CHDIP406-11|BOLD:AAH3619
Cecidomyiinae|[424]|CHDIP420-11|BOLD:AAH3619
Cecidomyiinae|[425]|CHDIP430-11|BOLD:AAH3619
Cecidomyiinae|[426]|CHDIP443-11|BOLD:AAH3619
Cecidomyiinae|[427]|CHDIP572-12|BOLD:AAH3619
Cecidomyiinae|[428]|CHDIP1008-12|BOLD:ACR8175
Cecidomyiinae|[429]|JWDCE026-10|BOLD:AAP1835
Cecidomyiinae|[430]|CHDIP1380-12|BOLD:AAH3752
Cecidomyiinae|[431]|CHDIP588-12|BOLD:ABY2157
Cecidomyiinae|[432]|CHDIP590-12|BOLD:ABY2157
Cecidomyiinae|[433]|CHDIP987-12|BOLD:AAY6420
Cecidomyiinae|[434]|JWDCB247-10|BOLD:AAH3782
Cecidomyiinae|[435]|CHDIP1400-12|BOLD:AAH3782
Cecidomyiinae|[436]|CHDIP1403-12|BOLD:ACB2982
Cecidomyiinae|[437]|JWDCJ1195-11|BOLD:AAV5589
Cecidomyiinae|[438]|CHDIP1085-12|BOLD:AAV5589
Cecidomyiinae|[439]|CHDIP1093-12|BOLD:AAV5589
Cecidomyiinae|[440]|CHDIP1094-12|BOLD:AAV5589
Cecidomyiinae|[441]|CHDIP1192-12|BOLD:AAV5589
Cecidomyiinae|[442]|CHDIP1398-12|BOLD:ACB2900
Cecidomyiinae|[443]|CHDIP571-12|BOLD:ACX5107
Cecidomyiinae|[444]|CHDIP1407-12|BOLD:ACX5107
Cecidomyiinae|[445]|CHDIP1316-12|BOLD:ACX5107
Cecidomyiinae|[446]|CHDIP1430-12|BOLD:ACX5107
Cecidomyiinae|[447]|CHDIP1438-12|BOLD:ACX5107
Cecidomyiinae|[448]|CHDIP1395-12|BOLD:AAN5281
Cecidomyiinae|[449]|CHDIP963-12|BOLD:AAH3713
Cecidomyiinae|[450]|CHDIP1426-12|BOLD:ACB2530
Cecidomyiinae|[451]|CHDIP1120-12|BOLD:ACO4990
Cecidomyiinae|[452]|CHDIP1441-12|BOLD:ACO4990
Winnertziinae|[453]|TWDIP328-09|BOLD:AAH3635
Winnertziinae|[454]|JWDCA852-10|BOLD:AAH3772
Winnertziinae|[455]|JWDCC641-10|BOLD:AAL7429
Winnertziinae|[456]|TWDIP250-09|BOLD:AAL7429
Winnertziinae|[457]|JWDCD373-10|BOLD:AAM6085
Winnertziinae|[458]|CHDIP1315-12|BOLD:ACB3032
Porricondylinae|[459]|JWDCA837-10|BOLD:AAH3624
Porricondylinae|[460]|JWDCA856-10|BOLD:AGA8658
Porricondylinae|[461]|MTDIC239-09|BOLD:AAH3624
Porricondylinae|[462]|JWDCH812-10|BOLD:AAH3624
Porricondylinae|[463]|CHDIP341-11|BOLD:AAH3627
Porricondylinae|[464]|CHDIP435-11|BOLD:AAH3627
Porricondylinae|[465]|JWDCA834-10|BOLD:AAH3627
Porricondylinae|[466]|JWDCA861-10|BOLD:AAH3627
Porricondylinae|[467]|CHDIP517-12|BOLD:AAH3627
Porricondylinae|[468]|CHDIP556-12|BOLD:AAH3627
Porricondylinae|[469]|CHDIP565-12|BOLD:AAH3627
Porricondylinae|[470]|TWDIP287-09|BOLD:AAH3627
Porricondylinae|[471]|CHDIP938-12|BOLD:AAH3627
Porricondylinae|[472]|TWDIP288-09|BOLD:AAH3628
Porricondylinae|[473]|CHDIP568-12|BOLD:AAH3628
Porricondylinae|[474]|CHDIP577-12|BOLD:AAH3628
Porricondylinae|[475]|CHDIP579-12|BOLD:AAH3628
Porricondylinae|[476]|CHDIP344-11|BOLD:AAH3628
Porricondylinae|[477]|CHDIP1261-12|BOLD:AAH3628
Porricondylinae|[478]|CHDIP1283-12|BOLD:ACB2911
Porricondylinae|[479]|JWDCF785-10|BOLD:AAQ0658
Porricondylinae|[480]|CHDIP1369-12|BOLD:ACB3014
Porricondylinae|[481]|TWDIP327-09|BOLD:AAH3634
Porricondylinae|[482]|CHDIP223-11|BOLD:ABY0116
Porricondylinae|[483]|CHDIP885-12|BOLD:ABX7729
Porricondylinae|[484]|JWDCF547-10|BOLD:AAQ0657
Porricondylinae|[485]|CHDIP1062-12|BOLD:AAQ0657
Porricondylinae|[486]|TWDIP289-09|BOLD:AAH3629
Porricondylinae|[487]|CHDIP1229-12|BOLD:AAH3629
Porricondylinae|[488]|CHDIP777-12|BOLD:ABX8329
Porricondylinae|[489]|JWDCD311-10|BOLD:AAM6076
Porricondylinae|[490]|CHDIP1285-12|BOLD:ACB2559
Porricondylinae|[491]|CHDIP1365-12|BOLD:ACB2524
Porricondylinae|[492]|JWDCD398-10|BOLD:AAL7452
Porricondylinae|[493]|CHDIP899-12|BOLD:AAL7452
Porricondylinae|[494]|CHDIP1312-12|BOLD:AAL7452
Porricondylinae|[495]|CHDIP1326-12|BOLD:AAP6867
Porricondylinae|[496]|JWDCI569-10|BOLD:AAQ0659
Porricondylinae|[497]|CHDIP312-11|BOLD:ADD2169
Porricondylinae|[498]|CHDIP1039-12|BOLD:ABX7925

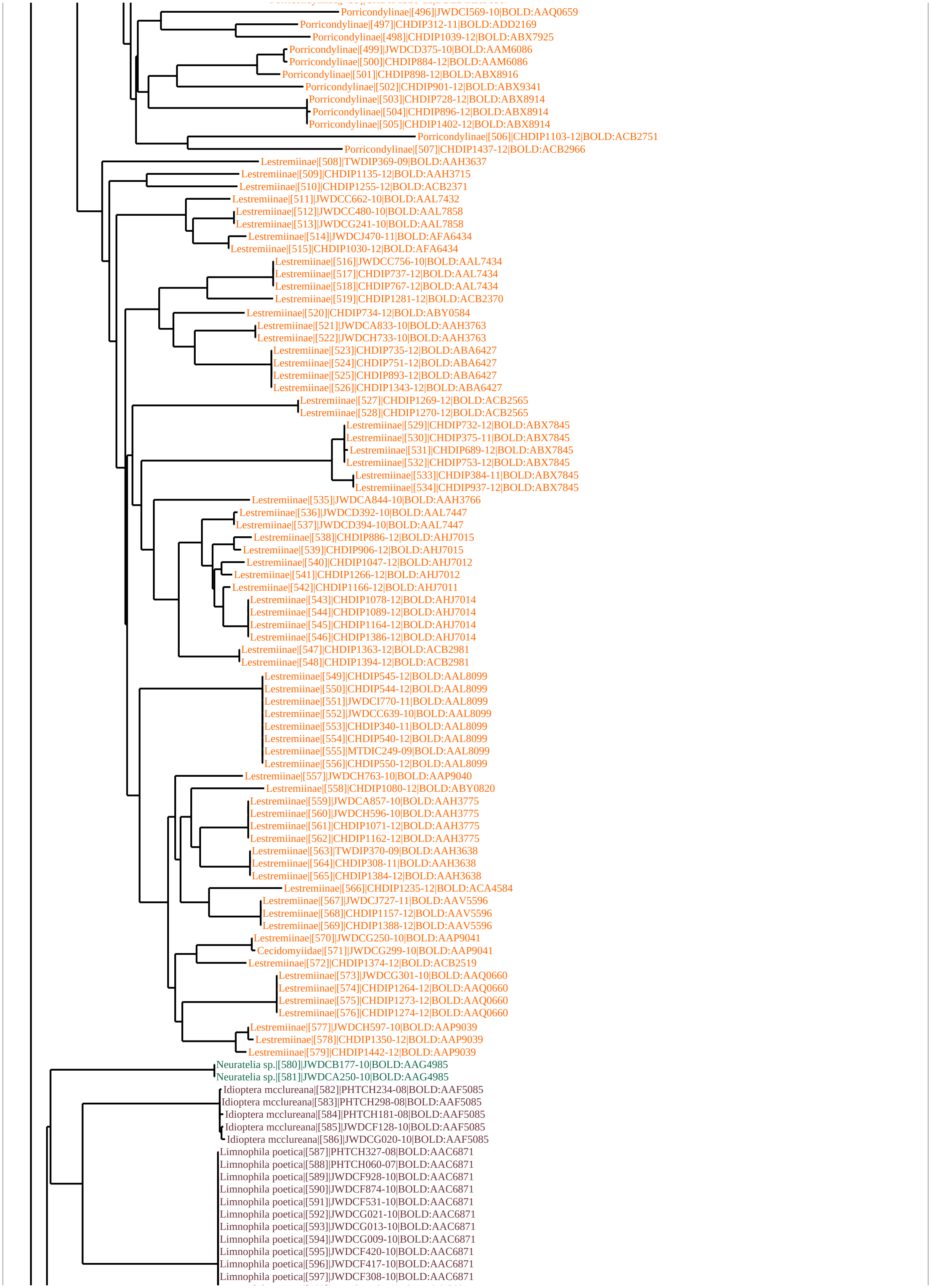
Porricondylinae|[496]|JWDCI569-10|BOLD:AAQ0659
Porricondylinae|[497]|CHDIP312-11|BOLD:ADD2169
Porricondylinae|[498]|CHDIP1039-12|BOLD:ABX7925
Porricondylinae|[499]|JWDCD375-10|BOLD:AAM6086
Porricondylinae|[500]|CHDIP884-12|BOLD:AAM6086
Porricondylinae|[501]|CHDIP898-12|BOLD:ABX8916
Porricondylinae|[502]|CHDIP901-12|BOLD:ABX9341
Porricondylinae|[503]|CHDIP728-12|BOLD:ABX8914
Porricondylinae|[504]|CHDIP896-12|BOLD:ABX8914
Porricondylinae|[505]|CHDIP1402-12|BOLD:ABX8914
Porricondylinae|[506]|CHDIP1103-12|BOLD:ACB2751
Porricondylinae|[507]|CHDIP1437-12|BOLD:ACB2966
Lestremiinae|[508]|TWDIP369-09|BOLD:AAH3637
Lestremiinae|[509]|CHDIP1135-12|BOLD:AAH3715
Lestremiinae|[510]|CHDIP1255-12|BOLD:ACB2371
Lestremiinae|[511]|JWDCC662-10|BOLD:AAL7432
Lestremiinae|[512]|JWDCC480-10|BOLD:AAL7858
Lestremiinae|[513]|JWDCG241-10|BOLD:AAL7858
Lestremiinae|[514]|JWDCJ470-11|BOLD:AFA6434
Lestremiinae|[515]|CHDIP1030-12|BOLD:AFA6434
Lestremiinae|[516]|JWDCC756-10|BOLD:AAL7434
Lestremiinae|[517]|CHDIP737-12|BOLD:AAL7434
Lestremiinae|[518]|CHDIP767-12|BOLD:AAL7434
Lestremiinae|[519]|CHDIP1281-12|BOLD:ACB2370
Lestremiinae|[520]|CHDIP734-12|BOLD:ABY0584
Lestremiinae|[521]|JWDCA833-10|BOLD:AAH3763
Lestremiinae|[522]|JWDCH733-10|BOLD:AAH3763
Lestremiinae|[523]|CHDIP735-12|BOLD:ABA6427
Lestremiinae|[524]|CHDIP751-12|BOLD:ABA6427
Lestremiinae|[525]|CHDIP893-12|BOLD:ABA6427
Lestremiinae|[526]|CHDIP1343-12|BOLD:ABA6427
Lestremiinae|[527]|CHDIP1269-12|BOLD:ACB2565
Lestremiinae|[528]|CHDIP1270-12|BOLD:ACB2565
Lestremiinae|[529]|CHDIP732-12|BOLD:ABX7845
Lestremiinae|[530]|CHDIP375-11|BOLD:ABX7845
Lestremiinae|[531]|CHDIP689-12|BOLD:ABX7845
Lestremiinae|[532]|CHDIP753-12|BOLD:ABX7845
Lestremiinae|[533]|CHDIP384-11|BOLD:ABX7845
Lestremiinae|[534]|CHDIP937-12|BOLD:ABX7845
Lestremiinae|[535]|JWDCA844-10|BOLD:AAH3766
Lestremiinae|[536]|JWDCD392-10|BOLD:AAL7447
Lestremiinae|[537]|JWDCD394-10|BOLD:AAL7447
Lestremiinae|[538]|CHDIP886-12|BOLD:AHJ7015
Lestremiinae|[539]|CHDIP906-12|BOLD:AHJ7015
Lestremiinae|[540]|CHDIP1047-12|BOLD:AHJ7012
Lestremiinae|[541]|CHDIP1266-12|BOLD:AHJ7012
Lestremiinae|[542]|CHDIP1166-12|BOLD:AHJ7011
Lestremiinae|[543]|CHDIP1078-12|BOLD:AHJ7014
Lestremiinae|[544]|CHDIP1089-12|BOLD:AHJ7014
Lestremiinae|[545]|CHDIP1164-12|BOLD:AHJ7014
Lestremiinae|[546]|CHDIP1386-12|BOLD:AHJ7014
Lestremiinae|[547]|CHDIP1363-12|BOLD:ACB2981
Lestremiinae|[548]|CHDIP1394-12|BOLD:ACB2981
Lestremiinae|[549]|CHDIP545-12|BOLD:AAL8099
Lestremiinae|[550]|CHDIP544-12|BOLD:AAL8099
Lestremiinae|[551]|JWDCI770-11|BOLD:AAL8099
Lestremiinae|[552]|JWDCC639-10|BOLD:AAL8099
Lestremiinae|[553]|CHDIP340-11|BOLD:AAL8099
Lestremiinae|[554]|CHDIP540-12|BOLD:AAL8099
Lestremiinae|[555]|MTDIC249-09|BOLD:AAL8099
Lestremiinae|[556]|CHDIP550-12|BOLD:AAL8099
Lestremiinae|[557]|JWDCH763-10|BOLD:AAP9040
Lestremiinae|[558]|CHDIP1080-12|BOLD:ABY0820
Lestremiinae|[559]|JWDCA857-10|BOLD:AAH3775
Lestremiinae|[560]|JWDCH596-10|BOLD:AAH3775
Lestremiinae|[561]|CHDIP1071-12|BOLD:AAH3775
Lestremiinae|[562]|CHDIP1162-12|BOLD:AAH3775
Lestremiinae|[563]|TWDIP370-09|BOLD:AAH3638
Lestremiinae|[564]|CHDIP308-11|BOLD:AAH3638
Lestremiinae|[565]|CHDIP1384-12|BOLD:AAH3638
Lestremiinae|[566]|CHDIP1235-12|BOLD:ACA4584
Lestremiinae|[567]|JWDCJ727-11|BOLD:AAV5596
Lestremiinae|[568]|CHDIP1157-12|BOLD:AAV5596
Lestremiinae|[569]|CHDIP1388-12|BOLD:AAV5596
Lestremiinae|[570]|JWDCG250-10|BOLD:AAP9041
Cecidomyiidae|[571]|JWDCG299-10|BOLD:AAP9041
Lestremiinae|[572]|CHDIP1374-12|BOLD:ACB2519
Lestremiinae|[573]|JWDCG301-10|BOLD:AAQ0660
Lestremiinae|[574]|CHDIP1264-12|BOLD:AAQ0660
Lestremiinae|[575]|CHDIP1273-12|BOLD:AAQ0660
Lestremiinae|[576]|CHDIP1274-12|BOLD:AAQ0660
Lestremiinae|[577]|JWDCH597-10|BOLD:AAP9039
Lestremiinae|[578]|CHDIP1350-12|BOLD:AAP9039
Lestremiinae|[579]|CHDIP1442-12|BOLD:AAP9039
Neuratelia sp.|[580]|JWDCB177-10|BOLD:AAG4985
Neuratelia sp.|[581]|JWDCA250-10|BOLD:AAG4985
Idioptera mcclureana|[582]|PHTCH234-08|BOLD:AAF5085
Idioptera mcclureana|[583]|PHTCH298-08|BOLD:AAF5085
Idioptera mcclureana|[584]|PHTCH181-08|BOLD:AAF5085
Idioptera mcclureana|[585]|JWDCF128-10|BOLD:AAF5085
Idioptera mcclureana|[586]|JWDCG020-10|BOLD:AAF5085
Limnophila poetica|[587]|PHTCH327-08|BOLD:AAC6871
Limnophila poetica|[588]|PHTCH060-07|BOLD:AAC6871
Limnophila poetica|[589]|JWDCF928-10|BOLD:AAC6871
Limnophila poetica|[590]|JWDCF874-10|BOLD:AAC6871
Limnophila poetica|[591]|JWDCF531-10|BOLD:AAC6871
Limnophila poetica|[592]|JWDCG021-10|BOLD:AAC6871
Limnophila poetica|[593]|JWDCG013-10|BOLD:AAC6871
Limnophila poetica|[594]|JWDCG009-10|BOLD:AAC6871
Limnophila poetica|[595]|JWDCF420-10|BOLD:AAC6871
Limnophila poetica|[596]|JWDCF417-10|BOLD:AAC6871
Limnophila poetica|[597]|JWDCF308-10|BOLD:AAC6871

Limnophila poetica|[595]|JWDCF420-10|BOLD:AAC6871
Limnophila poetica|[596]|JWDCF417-10|BOLD:AAC6871
Limnophila poetica|[597]|JWDCF308-10|BOLD:AAC6871
Limnophila poetica|[598]|JWDCF173-10|BOLD:AAC6871
Limnophila poetica|[599]|JWDCG587-10|BOLD:AAC6871
Limnophila poetica|[600]|JWDCG586-10|BOLD:AAC6871
Limnophila poetica|[601]|PHTCH010-07|BOLD:AAC6871
Limnophila poetica|[602]|PHTCH004-07|BOLD:AAC6871
Limnophila poetica|[603]|PHTCH003-07|BOLD:AAC6871
Limnophila poetica|[604]|PHTCH384-08|BOLD:AAC6871
Limnophila poetica|[605]|PHTCH204-08|BOLD:AAC6871
Limnophila poetica|[606]|PHTCH203-08|BOLD:AAC6871
Limnophila poetica|[607]|PHTCH185-08|BOLD:AAC6871
Limnophila poetica|[608]|PHTCH184-08|BOLD:AAC6871
Limnophila poetica|[609]|PHTCH180-08|BOLD:AAC6871
Limnophila poetica|[610]|PHTCH130-08|BOLD:AAC6871
Limnophila poetica|[611]|PHTCH108-08|BOLD:AAC6871
Limnophila poetica|[612]|PHTCH416-08|BOLD:AAC6871
Limnophila poetica|[613]|PHTCH101-08|BOLD:AAC6871
Limnophila poetica|[614]|PHTCH182-08|BOLD:AAC6871
Limnophila poetica|[615]|PHTCH128-08|BOLD:AAC6871
Limnophila poetica|[616]|PHTCH129-08|BOLD:AAC6871
Limnophila poetica|[617]|PHTCH131-08|BOLD:AAC6871
Limnophila poetica|[618]|JWDCK001-11|BOLD:AAC6871
Limnophila poetica|[619]|JWDCF931-10|BOLD:AAC6871
Limnophila poetica|[620]|JWDCI367-10|BOLD:AAC6871
Limnophila poetica|[621]|JWDCI368-10|BOLD:AAC6871
Limnophila poetica|[622]|JWDCI242-10|BOLD:AAC6871
Limnophila poetica|[623]|JWDCE945-10|BOLD:AAC6871
Limnophila poetica|[624]|JWDCE946-10|BOLD:AAC6871
Scathophaga furcata|[625]|JWDCK059-11|BOLD:AEU1602
Scathophaga furcata|[626]|JWDCC031-10|BOLD:AEU1602
Scathophaga furcata|[627]|JWDCE277-10|BOLD:AEU1602
Scathophaga furcata|[628]|JWDCF238-10|BOLD:AEU1602
Scathophaga furcata|[629]|JWDCH315-10|BOLD:AEU1602
Scathophaga furcata|[630]|JWDCH600-10|BOLD:AEU1602
Scathophaga furcata|[631]|JWDCL649-11|BOLD:AEU1602
Scathophaga furcata|[632]|JWDCJ498-11|BOLD:AEU1602
Scathophaga furcata|[633]|JWDCL195-11|BOLD:AEU1602
Scathophaga furcata|[634]|JWDCA293-10|BOLD:AEU1602
Scathophaga furcata|[635]|JWDCA284-10|BOLD:AEU1602
Scathophaga|[636]|MHFLY384-09|BOLD:AEU1602
Scathophaga furcata|[637]|MHFLY391-09|BOLD:AEU1602
Scathophaga furcata|[638]|JWDCA302-10|BOLD:AEU1602
Scathophaga furcata|[639]|JWDCJ041-11|BOLD:AEU1602
Scathophaga furcata|[640]|JWDCC193-10|BOLD:AEU1602
Scathophaga furcata|[641]|MHFLY061-06|BOLD:AEU1602
Scathophaga furcata|[642]|JWDCJ615-11|BOLD:AEU1602
Scathophaga furcata|[643]|JWDCL081-11|BOLD:AEU1602
Scathophaga furcata|[644]|JWDCG449-10|BOLD:AEU1602
Scathophaga furcata|[645]|JWDCE708-10|BOLD:AEU1602
Scathophaga furcata|[646]|JWDCE762-10|BOLD:AEU1602
Scathophaga furcata|[647]|MHFLY057-06|BOLD:AEU1602
Scathophaga furcata|[648]|JWDCC383-10|BOLD:AEU1602
Scathophaga furcata|[649]|MHFLI049-06|BOLD:AEU1602
Scathophaga furcata|[650]|JWDCJ493-11|BOLD:AEU1602
Scathophaga furcata|[651]|JWDCF140-10|BOLD:AEU1602
Scathophaga furcata|[652]|JWDCF628-10|BOLD:AEU1602
Scathophaga furcata|[653]|JWDCE476-10|BOLD:AEU1602
Scathophaga furcata|[654]|JWDCA280-10|BOLD:AEU1602
Scathophaga furcata|[655]|MHFLI225-06|BOLD:AEU1602
Scathophaga furcata|[656]|MHFLI223-06|BOLD:AEU1602
Scathophaga furcata|[657]|JWDCA277-10|BOLD:AEU1602
Scathophaga furcata|[658]|JWDCC259-10|BOLD:AEU1602
Scathophaga furcata|[659]|JWDCA259-10|BOLD:AEU1602
Scathophaga furcata|[660]|JWDCE713-10|BOLD:AEU1602
Scathophaga furcata|[661]|JWDCE521-10|BOLD:AEU1602
Scathophaga furcata|[662]|JWDCJ617-11|BOLD:AEU1602
Scathophaga furcata|[663]|MHFLY383-09|BOLD:AEU1602
Scathophaga furcata|[664]|JWDCA269-10|BOLD:AEU1602
Scathophaga furcata|[665]|JWDCC845-10|BOLD:AEU1602
Scathophaga furcata|[666]|JWDCH916-10|BOLD:AEU1602
Scathophaga furcata|[667]|JWDCJ616-11|BOLD:AEU1602
Scathophaga furcata|[668]|JWDCD130-10|BOLD:AEU1602
Scathophaga furcata|[669]|MHFLI052-06|BOLD:AEU1602
Scathophaga furcata|[670]|MHFLI229-06|BOLD:AEU1602
Scathophaga furcata|[671]|JWDCJ622-11|BOLD:AEU1602
Scathophaga furcata|[672]|JWDCE393-10|BOLD:AEU1602
Scathophaga furcata|[673]|JWDCF896-10|BOLD:AEU1602
Scathophaga furcata|[674]|MHFLI054-06|BOLD:AEU1602
Scathophaga furcata|[675]|MHFLI066-06|BOLD:AEU1602
Scathophaga furcata|[676]|MHFLI226-06|BOLD:AEU1602
Scathophaga furcata|[677]|JWDCA296-10|BOLD:AEU1602
Scathophaga furcata|[678]|JWDCJ1415-11|BOLD:AEU1602
Scathophaga furcata|[679]|MHFLI063-06|BOLD:AEU1602
Scathophaga furcata|[680]|JWDCC793-10|BOLD:AEU1602
Scathophaga furcata|[681]|JWDCJ620-11|BOLD:AEU1602
Scathophaga furcata|[682]|JWDCC078-10|BOLD:AEU1602
Scathophaga furcata|[683]|JWDCJ040-11|BOLD:AEU1602
Scathophaga furcata|[684]|JWDCJ567-11|BOLD:AEU1602
Scathophaga furcata|[685]|JWDCE394-10|BOLD:AEU1602
Scathophaga furcata|[686]|MHFLY056-06|BOLD:AEU1602
Scathophaga furcata|[687]|MHFLY402-09|BOLD:AEU1602
Scathophaga furcata|[688]|JWDCE707-10|BOLD:AEU1602
Scathophaga furcata|[689]|JWDCC008-10|BOLD:AEU1602
Scathophaga furcata|[690]|JWDCA294-10|BOLD:AEU1602
Scathophaga furcata|[691]|JWDCJ568-11|BOLD:AEU1602
Scathophaga furcata|[692]|JWDCI656-10|BOLD:AEU1602
Scathophaga furcata|[693]|JWDCC795-10|BOLD:AEU1602
Scathophaga furcata|[694]|JWDCF597-10|BOLD:AEU1602
Scathophaga furcata|[695]|JWDCF750-10|BOLD:AEU1602
Scathophaga furcata|[696]|JWDCF100-10|BOLD:AEU1602
Scathophaga furcata|[697]|JWDCA304-10|BOLD:AEU1602

Scathophaga furcata|[695]|JWDCF750-10|BOLD:AEU1602
Scathophaga furcata|[696]|JWDCF100-10|BOLD:AEU1602
Scathophaga furcata|[697]|JWDCA304-10|BOLD:AEU1602
Scathophaga furcata|[698]|JWDCD114-10|BOLD:AEU1602
Scathophaga furcata|[699]|MHFLI051-06|BOLD:AEU1602
Scathophaga furcata|[700]|JWDCJ497-11|BOLD:AEU1602
Scathophaga|[701]|JWDCF141-10|BOLD:AEU1602
Scathophaga furcata|[702]|JWDCE571-10|BOLD:AEU1602
Scathophaga furcata|[703]|JWDCJ495-11|BOLD:AEU1602
Scathophaga furcata|[704]|MHFLY373-09|BOLD:AEU1602
Scathophaga furcata|[705]|JWDCC258-10|BOLD:AEU1602
Scathophaga furcata|[706]|JWDCE209-10|BOLD:AEU1602
Scathophaga furcata|[707]|JWDCL266-11|BOLD:AEU1602
Scathophaga furcata|[708]|JWDCG628-10|BOLD:AEU1602
Scathophaga furcata|[709]|MHFLI227-06|BOLD:AEU1602
Scathophaga furcata|[710]|MHFLY375-09|BOLD:AEU1602
Scathophaga furcata|[711]|JWDCI788-11|BOLD:AEU1602
Scathophaga furcata|[712]|MHFLI228-06|BOLD:AEU1602
Scathophaga furcata|[713]|JWDCE410-10|BOLD:AEU1602
Scathophaga furcata|[714]|JWDCJ496-11|BOLD:AEU1602
Scathophaga furcata|[715]|JWDCA610-10|BOLD:AEU1602
Scathophaga furcata|[716]|JWDCE676-10|BOLD:AEU1602
Scathophaga furcata|[717]|JWDCE484-10|BOLD:AEU1602
Scathophaga furcata|[718]|MHFLY371-09|BOLD:AEU1602
Scathophaga furcata|[719]|JWDCJ042-11|BOLD:AEU1602
Scathophaga furcata|[720]|JWDCF440-10|BOLD:AEU1602
Scathophaga furcata|[721]|JWDCL161-11|BOLD:AEU1602
Scathophaga furcata|[722]|JWDCC797-10|BOLD:AEU1602
Scathophaga furcata|[723]|JWDCA275-10|BOLD:AEU1602
Scathophaga furcata|[724]|JWDCL268-11|BOLD:AEU1602
Scathophaga furcata|[725]|MHFLY052-06|BOLD:AEU1602
Scathophaga furcata|[726]|JWDCL650-11|BOLD:AEU1602
Scathophaga furcata|[727]|JWDCE395-10|BOLD:AEU1602
Scathophaga furcata|[728]|JWDCE246-10|BOLD:AEU1602
Scathophaga furcata|[729]|JWDCC030-10|BOLD:AEU1602
Scathophaga furcata|[730]|JWDCC082-10|BOLD:AEU1602
Scathophaga furcata|[731]|JWDCA264-10|BOLD:AEU1602
Scathophaga furcata|[732]|JWDCE572-10|BOLD:AEU1602
Scathophaga furcata|[733]|JWDCJ566-11|BOLD:AEU1602
Scathophaga furcata|[734]|JWDCC121-10|BOLD:AEU1602
Scathophaga furcata|[735]|JWDCC074-10|BOLD:AEU1602
Scathophaga furcata|[736]|JWDCC899-10|BOLD:AEU1602
Scathophaga furcata|[737]|MTDIC247-09|BOLD:AEU1602
Scathophaga furcata|[738]|JWDCF101-10|BOLD:AEU1602
Scathophaga furcata|[739]|MHFLI040-06|BOLD:AEU1602
Scathophaga furcata|[740]|MHFLI044-06|BOLD:AEU1602
Scathophaga furcata|[741]|JWDCC003-10|BOLD:AEU1602
Scathophaga furcata|[742]|JWDCC794-10|BOLD:AEU1602
Scathophaga furcata|[743]|JWDCI234-10|BOLD:AEU1602
Scathophaga furcata|[744]|JWDCJ1164-11|BOLD:AEU1602
Scathophaga furcata|[745]|JWDCE208-10|BOLD:AEU1602
Scathophaga furcata|[746]|JWDCE483-10|BOLD:AEU1602
Scathophaga furcata|[747]|MHFLI894-09|BOLD:AEU1602
Scathophaga furcata|[748]|MHFLI053-06|BOLD:AEU1602
Scathophaga furcata|[749]|JWDCE247-10|BOLD:AEU1602
Scathophaga furcata|[750]|JWDCE761-10|BOLD:AEU1602
Scathophaga furcata|[751]|JWDCC077-10|BOLD:AEU1602
Scathophaga furcata|[752]|JWDCJ613-11|BOLD:AEU1602
Scathophaga furcata|[753]|JWDCE034-10|BOLD:AEU1602
Scathophaga furcata|[754]|MHFLI233-06|BOLD:AEU1602
Scathophaga furcata|[755]|JWDCC898-10|BOLD:AEU1602
Scathophaga furcata|[756]|MHFLY386-09|BOLD:AEU1602
Scathophaga furcata|[757]|JWDCA292-10|BOLD:AEU1602
Scathophaga furcata|[758]|JWDCJ494-11|BOLD:AEU1602
Scathophaga furcata|[759]|JWDCE724-10|BOLD:AEU1602
Scathophaga furcata|[760]|JWDCF682-10|BOLD:AEU1602
Scathophaga furcata|[761]|MHFLI224-06|BOLD:AEU1602
Scathophaga furcata|[762]|JWDCC252-10|BOLD:AEU1602
Scathophaga furcata|[763]|JWDCC257-10|BOLD:AEU1602
Scathophaga furcata|[764]|MHFLI234-06|BOLD:AEU1602
Scathophaga furcata|[765]|MHFLY374-09|BOLD:AEU1602
Scathophaga furcata|[766]|JWDCE475-10|BOLD:AEU1602
Scathophaga furcata|[767]|MHFLY377-09|BOLD:AEU1602
Scathophaga furcata|[768]|JWDCD115-10|BOLD:AEU1602
Scathophaga furcata|[769]|JWDCL648-11|BOLD:AEU1602
Scathophaga furcata|[770]|JWDCJ246-11|BOLD:AEU1602
Scathophaga furcata|[771]|JWDCH731-10|BOLD:AEU1602
Scathophaga furcata|[772]|JWDCE035-10|BOLD:AEU1602
Scathophaga furcata|[773]|JWDCE763-10|BOLD:AEU1602
Scathophaga furcata|[774]|JWDCE059-10|BOLD:AEU1602
Scathophaga furcata|[775]|JWDCC894-10|BOLD:AEU1602
Scathophaga furcata|[776]|JWDCC081-10|BOLD:AEU1602
Scathophaga furcata|[777]|JWDCJ668-11|BOLD:AEU1602
Scathophaga furcata|[778]|JWDCE574-10|BOLD:AEU1602
Scathophaga furcata|[779]|JWDCC194-10|BOLD:AEU1602
Scathophaga furcata|[780]|MHFLI925-09|BOLD:AEU1602
Scathophaga furcata|[781]|JWDCJ188-11|BOLD:AEU1602
Scathophaga furcata|[782]|JWDCJ923-11|BOLD:AEU1602
Scathophaga furcata|[783]|JWDCG574-10|BOLD:AEU1602
Scathophaga furcata|[784]|MHFLI282-06|BOLD:AEU1602
Scathophaga furcata|[785]|JWDCE210-10|BOLD:AEU1602
Scathophaga furcata|[786]|MHFLI278-06|BOLD:AEU1602
Scathophaga furcata|[787]|MHFLI231-06|BOLD:AEU1602
Scathophaga furcata|[788]|MHFLI929-09|BOLD:AEU1602
Scathophaga furcata|[789]|JWDCC079-10|BOLD:AEU1602
Scathophaga furcata|[790]|JWDCE409-10|BOLD:AEU1602
Scathophaga furcata|[791]|MHFLI230-06|BOLD:AEU1602
Scathophaga furcata|[792]|MHFLY053-06|BOLD:AEU1602
Scathophaga furcata|[793]|JWDCC075-10|BOLD:AEU1602
Scathophaga furcata|[794]|MHFLI064-06|BOLD:AEU1602
Scathophaga furcata|[795]|JWDCE324-10|BOLD:AEU1602
Scathophaga furcata|[796]|JWDCD017-10|BOLD:AEU1602

Scathophaga furcata|[794]|MHFLI064-06|BOLD:AEU1602
Scathophaga furcata|[795]|JWDCE324-10|BOLD:AEU1602
Scathophaga furcata|[796]|JWDCD017-10|BOLD:AEU1602
Scathophaga furcata|[797]|MTDIC251-09|BOLD:AEU1602
Scathophaga furcata|[798]|JWDCC080-10|BOLD:AEU1602
Scathophaga furcata|[799]|JWDCI331-10|BOLD:AEU1602
Scathophaga furcata|[800]|JWDCF024-10|BOLD:AEU1602
Scathophaga furcata|[801]|JWDCE678-10|BOLD:AEU1602
Scathophaga furcata|[802]|JWDCL156-11|BOLD:AEU1602
Scathophaga furcata|[803]|JWDCC895-10|BOLD:AEU1602
Scathophaga furcata|[804]|JWDCJ922-11|BOLD:AEU1602
Scathophaga furcata|[805]|JWDCE055-10|BOLD:AEU1602
Scathophaga furcata|[806]|JWDCD055-10|BOLD:AEU1602
Scathophaga furcata|[807]|JWDCL324-11|BOLD:AEU1602
Scathophaga furcata|[808]|JWDCB075-10|BOLD:AEU1602
Scathophaga furcata|[809]|MHFLI238-06|BOLD:AEU1602
Scathophaga furcata|[810]|MHFLY366-09|BOLD:AEU1602
Scathophaga furcata|[811]|JWDCC896-10|BOLD:AEU1602
Scathophaga furcata|[812]|MHFLY400-09|BOLD:AEU1602
Scathophaga furcata|[813]|JWDCJ1006-11|BOLD:AEU1602
Scathophaga furcata|[814]|JWDCL651-11|BOLD:AEU1602
Scathophaga furcata|[815]|JWDCC076-10|BOLD:AEU1602
Scathophaga furcata|[816]|JWDCE857-10|BOLD:AEU1602
Scathophaga furcata|[817]|JWDCJ1418-11|BOLD:AEU1602
Scathophaga furcata|[818]|JWDCE482-10|BOLD:AEU1602
Scathophaga furcata|[819]|MHFLI055-06|BOLD:AEU1602
Scathophaga furcata|[820]|JWDCD070-10|BOLD:AEU1602
Scathophaga furcata|[821]|MHFLY398-09|BOLD:AEU1602
Scathophaga furcata|[822]|TWDIP174-09|BOLD:AEU1602
Scathophaga spurca|[823]|JWDCD069-10|BOLD:AEV5717
Scathophaga spurca|[824]|JWDCC084-10|BOLD:AEV5717
Scathophaga spurca|[825]|JWDCG307-10|BOLD:AEV5717
Scathophaga spurca|[826]|JWDCL411-11|BOLD:AEV5717
Scathophaga spurca|[827]|JWDCL409-11|BOLD:AEV5717
Scathophaga spurca|[828]|JWDCL006-11|BOLD:AEV5717
Scathophaga spurca|[829]|JWDCJ614-11|BOLD:AEV5717
Scathophaga spurca|[830]|JWDCJ564-11|BOLD:AEV5717
Scathophaga spurca|[831]|JWDCA268-10|BOLD:AEV5717
Scathophaga spurca|[832]|JWDCC083-10|BOLD:AEV5717
Scathophaga spurca|[833]|MTDIC029-09|BOLD:AEV5717
Scathophaga spurca|[834]|JWDCC164-10|BOLD:AEV5717
Scathophaga spurca|[835]|DSCNI050-07|BOLD:AEV5717
Scathophaga spurca|[836]|MHFLI258-06|BOLD:AEV5717
Scathophaga spurca|[837]|MHFLY130-07|BOLD:AEV5717
Scathophaga spurca|[838]|JWDCC148-10|BOLD:AEV5717
Scathophaga spurca|[839]|MHFLY382-09|BOLD:AEV5717
Scathophaga spurca|[840]|JWDCK249-11|BOLD:AEV5717
Scathophaga spurca|[841]|JWDCF098-10|BOLD:AEV5717
Orthacheta|[842]|TWDIP116-09|BOLD:AAH4223
Orthacheta|[843]|JWDCD015-10|BOLD:AAH4223
Orthacheta|[844]|JWDCD020-10|BOLD:AAH4223
Orthacheta|[845]|JWDCA285-10|BOLD:AAH4223
Orthacheta|[846]|JWDCK283-11|BOLD:AAH4223
Orthacheta|[847]|JWEMP036-10|BOLD:AAH4223
Orthacheta|[848]|JWDCG843-10|BOLD:AAH4223
Orthacheta|[849]|JWDCH375-10|BOLD:AAH4223
Orthacheta|[850]|JWDCI653-10|BOLD:AAH4223
Scathophagini|[851]|JWDCH771-10|BOLD:AAP8772
Scathophaginae|[852]|JWDCA260-10|BOLD:AAH4216
Scathophaginae|[853]|MHFLI895-09|BOLD:AAH4216
Scathophaginae|[854]|MHFLI893-09|BOLD:AAH4216
Scathophaginae|[855]|JWDCE677-10|BOLD:AAH4216
Scathophaginae|[856]|MHFLI056-06|BOLD:AAH4216
Scathophaginae|[857]|MHFLI142-06|BOLD:AAH4216
Scathophaginae|[858]|JWDCL541-11|BOLD:AAH4216
Scathophaginae|[859]|MHFLI148-06|BOLD:AAH4216
Scathophaginae|[860]|MHFLI968-09|BOLD:AAH4216
Scathophaginae|[861]|MHFLI124-06|BOLD:AAH4216
Scathophaginae|[862]|MHFLI143-06|BOLD:AAH4216
Scathophaginae|[863]|JWDCA263-10|BOLD:AAH4216
Scathophaginae|[864]|JWDCE391-10|BOLD:AAH4216
Scathophaginae|[865]|JWDCE392-10|BOLD:AAH4216
Scathophaginae|[866]|JWDCE628-10|BOLD:AAH4216
Scathophaginae|[867]|JWDCE766-10|BOLD:AAH4216
Scathophaga|[868]|MHFLY385-09|BOLD:AAH4215
Scathophaga|[869]|MHCOL349-07|BOLD:AAH4215
Scathophaga|[870]|MHFLY405-09|BOLD:AAH4215
Scathophaga|[871]|MHFLY389-09|BOLD:AAH4215
Scathophaga|[872]|MHFLY015-06|BOLD:AAH4215
Scathophaga|[873]|MHFLY020-06|BOLD:AAH4215
Scathophaga|[874]|TWDIP170-09|BOLD:AAH4215
Scathophaga|[875]|MHFLY044-06|BOLD:AAH4215
Scathophaga|[876]|MHFLY027-06|BOLD:AAH4215
Scathophaga|[877]|MHFLY013-06|BOLD:AAH4215
Scathophaga|[878]|MHFLI156-06|BOLD:AAH4215
Scathophaga|[879]|MHFLI144-06|BOLD:AAH4215
Scathophaga|[880]|JWDCE365-10|BOLD:AAH4215
Scathophaga|[881]|MHFLI641-07|BOLD:AAH4215
Scathophaga|[882]|JWDCA276-10|BOLD:AAH4215
Scathophaga|[883]|JWDCA279-10|BOLD:AAH4215
Scathophaga|[884]|JWDCI544-10|BOLD:AAH4215
Scathophaga|[885]|MHCOL355-07|BOLD:AAH4215
Scathophaga|[886]|MHCOL342-07|BOLD:AAH4215
Scathophaga|[887]|MHFLI645-07|BOLD:AAH4215
Scathophaga|[888]|MHCOL354-07|BOLD:AAH4215
Scathophaga|[889]|MHFLY381-09|BOLD:AAH4215
Scathophaga|[890]|MTDIC228-09|BOLD:AAH4215
Scathophaga|[891]|MHCOL358-07|BOLD:AAH4215
Scathophaga|[892]|MHCOL359-07|BOLD:AAH4215
Scathophaga|[893]|MHFLY403-09|BOLD:AAH4215
Scathophaga|[894]|MHCOL356-07|BOLD:AAH4215
Scathophaga|[895]|MHCOL352-07|BOLD:AAH4215
Scathophaga|[896]|MHCOL353-07|BOLD:AAH4215

Scathophaga|[894]|MHCOL356-07|BOLD:AAH4215
Scathophaga|[895]|MHCOL352-07|BOLD:AAH4215
Scathophaga|[896]|MHCOL353-07|BOLD:AAH4215
Scathophaga|[897]|JSFLA008-08|BOLD:AAH4215
Scathophaga|[898]|TWDIP171-09|BOLD:AAH4215
Scathophaga|[899]|TWDIP172-09|BOLD:AAH4215
Scathophaga|[900]|TWDIP173-09|BOLD:AAH4215
Scathophaga|[901]|JWDCC792-10|BOLD:AAH4215
Scathophaga|[902]|JWDCC835-10|BOLD:AAH4215
Scathophaga|[903]|JWDCC844-10|BOLD:AAH4215
Scathophaga|[904]|JWDCC846-10|BOLD:AAH4215
Scathophaga|[905]|JWDCC847-10|BOLD:AAH4215
Scathophaga|[906]|JWDCC860-10|BOLD:AAH4215
Scathophaga|[907]|JWDCC893-10|BOLD:AAH4215
Scathophaga|[908]|JWDCC897-10|BOLD:AAH4215
Scathophaga|[909]|JWDCC900-10|BOLD:AAH4215
Scathophaga|[910]|JWDCA265-10|BOLD:AAH4215
Scathophaga|[911]|JWDCA270-10|BOLD:AAH4215
Scathophaga|[912]|JWDCA271-10|BOLD:AAH4215
Scathophaga|[913]|JWDCA278-10|BOLD:AAH4215
Scathophaga|[914]|JWDCA295-10|BOLD:AAH4215
Scathophaga|[915]|MHFLI892-09|BOLD:AAH4215
Scathophaga|[916]|MHCOL357-07|BOLD:AAH4215
Scathophaga|[917]|JWDCA297-10|BOLD:AAH4215
Scathophaga|[918]|JWDCA467-10|BOLD:AAH4215
Scathophaga|[919]|MHFLI919-09|BOLD:AAH4215
Scathophaga|[920]|MHFLI944-09|BOLD:AAH4215
Scathophaga|[921]|JWDCA267-10|BOLD:AAH4215
Scathophaga|[922]|JWDCA298-10|BOLD:AAH4215
Scathophaga|[923]|MHCOL351-07|BOLD:AAH4215
Scathophaga|[924]|MHCOL350-07|BOLD:AAH4215
Scathophaga|[925]|MHCOL348-07|BOLD:AAH4215
Scathophaga|[926]|MHCOL346-07|BOLD:AAH4215
Scathophaga|[927]|MHCOL345-07|BOLD:AAH4215
Scathophaga|[928]|MHCOL344-07|BOLD:AAH4215
Scathophaga|[929]|MHCOL343-07|BOLD:AAH4215
Scathophaga|[930]|MHCOL347-07|BOLD:AAH4215
Scathophaga|[931]|MHCOL341-07|BOLD:AAH4215
Scathophaga|[932]|MHCOL340-07|BOLD:AAH4215
Scathophaga|[933]|MHCOL288-07|BOLD:AAH4215
Scathophaga|[934]|JWDCA266-10|BOLD:AAH4215
Scathophaga|[935]|MHFLY356-09|BOLD:AAH4215
Scathophaga|[936]|MHFLY369-09|BOLD:AAH4215
Scathophaga|[937]|JWDCH316-10|BOLD:AAH4215
Scathophaga|[938]|JWDCI111-10|BOLD:AAH4215
Scathophaga|[939]|MHFLY372-09|BOLD:AAH4215
Scathophaga|[940]|MHFLY390-09|BOLD:AAH4215
Scathophaga|[941]|JWDCI112-10|BOLD:AAH4215
Scathophaga|[942]|JWDCI516-10|BOLD:AAH4215
Scathophaga|[943]|JWDCJ669-11|BOLD:AAH4215
Scathophaga|[944]|JWDCJ949-11|BOLD:AAH4215
Scathophaga|[945]|JWDCJ1417-11|BOLD:AAH4215
Scathophaga|[946]|JWDCL078-11|BOLD:AAH4215
Scathophaga|[947]|JWDCL556-11|BOLD:AAH4215
Scathophaga|[948]|JWDCG579-10|BOLD:AAH4215
Scathophaga|[949]|JWDCH915-10|BOLD:AAH4215
Scathophaga|[950]|JWDCG001-10|BOLD:AAH4215
Scathophaga|[951]|JWDCH173-10|BOLD:AAH4215
Scathophaga|[952]|JWDCG277-10|BOLD:AAH4215
Scathophaga|[953]|JWDCF026-10|BOLD:AAH4215
Scathophaga|[954]|JWDCF027-10|BOLD:AAH4215
Scathophaga|[955]|JWDCE389-10|BOLD:AAH4215
Scathophaga|[956]|JWDCE390-10|BOLD:AAH4215
Scathophaga|[957]|JWDCF232-10|BOLD:AAH4215
Scathophaga|[958]|JWDCF243-10|BOLD:AAH4215
Scathophaga|[959]|JWDCE520-10|BOLD:AAH4215
Scathophaga|[960]|JWDCE539-10|BOLD:AAH4215
Scathophaga|[961]|JWDCF271-10|BOLD:AAH4215
Scathophaga|[962]|JWDCE573-10|BOLD:AAH4215
Scathophaga|[963]|JWDCE589-10|BOLD:AAH4215
Scathophaga|[964]|JWDCF455-10|BOLD:AAH4215
Scathophaga|[965]|JWDCE765-10|BOLD:AAH4215
Scathophaga|[966]|JWDCF681-10|BOLD:AAH4215
Scathophaga|[967]|JWDCE177-10|BOLD:AAH4215
Scathophaga|[968]|JWDCE259-10|BOLD:AAH4215
Scathophaginae|[969]|JWDCI249-10|BOLD:AAM9280
Scathophaginae|[970]|JWDCI250-10|BOLD:AAM9280
Trichopalpus palpalis|[971]|MHFLI125-06|BOLD:AAH4217
Trichopalpus punctipes|[972]|JWDCJ1008-11|BOLD:AAH4220
Trichopalpus punctipes|[973]|JWDCI837-11|BOLD:AAH4220
Trichopalpus punctipes|[974]|JWDCJ709-11|BOLD:AAH4220
Trichopalpus punctipes|[975]|JWDCJ101-11|BOLD:AAH4220
Trichopalpus punctipes|[976]|JWDCF241-10|BOLD:AAH4220
Trichopalpus punctipes|[977]|JWDCC719-10|BOLD:AAH4220
Trichopalpus punctipes|[978]|JWDCC421-10|BOLD:AAH4220
Trichopalpus punctipes|[979]|JWDCJ099-11|BOLD:AAH4220
Trichopalpus punctipes|[980]|JWDCI694-11|BOLD:AAH4220
Trichopalpus punctipes|[981]|MHFLI542-07|BOLD:AAH4220
Trichopalpus punctipes|[982]|JWDCD021-10|BOLD:AAH4220
Trichopalpus punctipes|[983]|JWDCE537-10|BOLD:AAH4220
Trichopalpus punctipes|[984]|MHFLI865-07|BOLD:AAH4220
Trichopalpus punctipes|[985]|JWDCF518-10|BOLD:AAH4220
Trichopalpus punctipes|[986]|JWDCK132-11|BOLD:AAH4220
Trichopalpus punctipes|[987]|JWDCL471-11|BOLD:AAH4220
Trichopalpus punctipes|[988]|JWDCI835-11|BOLD:AAH4220
Trichopalpus punctipes|[989]|JWDCH213-10|BOLD:AAH4220
Trichopalpus punctipes|[990]|JWDCJ791-11|BOLD:AAH4220
Trichopalpus punctipes|[991]|JWDCL160-11|BOLD:AAH4220
Trichopalpus punctipes|[992]|JWDCH374-10|BOLD:AAH4220
Trichopalpus punctipes|[993]|JWDCI836-11|BOLD:AAH4220
Trichopalpus punctipes|[994]|JWDCK056-11|BOLD:AAH4220
Trichopalpus punctipes|[995]|JWDCD072-10|BOLD:AAH4220

Trichopalpus punctipes|[993]|JWDCI836-11|BOLD:AAH4220
Trichopalpus punctipes|[994]|JWDCK056-11|BOLD:AAH4220
Trichopalpus punctipes|[995]|JWDCD072-10|BOLD:AAH4220
Trichopalpus punctipes|[996]|JWDCK058-11|BOLD:AAH4220
Trichopalpus punctipes|[997]|JWDCK280-11|BOLD:AAH4220
Trichopalpus punctipes|[998]|JWDCJ100-11|BOLD:AAH4220
Trichopalpus punctipes|[999]|JWDCF519-10|BOLD:AAH4220
Trichopalpus punctipes|[1000]|JWDCF658-10|BOLD:AAH4220
Trichopalpus punctipes|[1001]|JWDCD117-10|BOLD:AAH4220
Trichopalpus punctipes|[1002]|JWDCL001-11|BOLD:AAH4220
Trichopalpus punctipes|[1003]|JWDCF338-10|BOLD:AAH4220
Trichopalpus punctipes|[1004]|JWDCK285-11|BOLD:AAH4220
Trichopalpus punctipes|[1005]|MHFLI489-07|BOLD:AAH4220
Trichopalpus punctipes|[1006]|JWDCI834-11|BOLD:AAH4220
Trichopalpus punctipes|[1007]|JWDCC721-10|BOLD:AAH4220
Trichopalpus punctipes|[1008]|JWDCK281-11|BOLD:AAH4220
Trichopalpus punctipes|[1009]|JWDCH842-10|BOLD:AAH4220
Trichopalpus punctipes|[1010]|JWDCL473-11|BOLD:AAH4220
Trichopalpus punctipes|[1011]|JWDCC013-10|BOLD:AAH4220
Trichopalpus punctipes|[1012]|JWDCH373-10|BOLD:AAH4220
Trichopalpus punctipes|[1013]|JWDCL557-11|BOLD:AAH4220
Trichopalpus punctipes|[1014]|JWDCF661-10|BOLD:AAH4220
Scathophaginae|[1015]|JWDCL652-11|BOLD:AAP8771
Scathophaginae|[1016]|JWDCH746-10|BOLD:AAP8771
Pogonota barbata|[1017]|JWDCK053-11|BOLD:AAG9681
Pogonota barbata|[1018]|JWDCK055-11|BOLD:AAG9681
Pogonota barbata|[1019]|MHFLI272-06|BOLD:AAG9681
Pogonota barbata|[1020]|JWDCK282-11|BOLD:AAG9681
Pogonota barbata|[1021]|JWDCK278-11|BOLD:AAG9681
Pogonota barbata|[1022]|JWDCJ1541-11|BOLD:AAG9681
Pogonota barbata|[1023]|JWDCK054-11|BOLD:AAG9681
Pogonota barbata|[1024]|JWDCL540-11|BOLD:AAG9681
Pogonota barbata|[1025]|JWDCJ186-11|BOLD:AAG9681
Pogonota barbata|[1026]|JWDCJ243-11|BOLD:AAG9681
Pogonota barbata|[1027]|JWDCH363-10|BOLD:AAG9681
Pogonota barbata|[1028]|MHFLI905-09|BOLD:AAG9681
Pogonota barbata|[1029]|JWDCK279-11|BOLD:AAG9681
Pogonota barbata|[1030]|JWDCG842-10|BOLD:AAG9681
Scathophaginae|[1031]|JWDCB073-10|BOLD:AAH4238
Cosmetopus longus|[1032]|JWDCI659-10|BOLD:AAH4239
Cosmetopus longus|[1033]|JWDCG477-10|BOLD:AAH4239
Cosmetopus longus|[1034]|JWDCB074-10|BOLD:AAH4239
Cosmetopus longus|[1035]|JWDCC118-10|BOLD:AAH4239
Cosmetopus longus|[1036]|JWDCJ950-11|BOLD:AAH4239
Cosmetopus longus|[1037]|JWDCL267-11|BOLD:AAH4239
Cosmetopus longus|[1038]|JWDCI655-10|BOLD:AAH4239
Cosmetopus longus|[1039]|JWDCI658-10|BOLD:AAH4239
Cosmetopus longus|[1040]|JWDCI660-10|BOLD:AAH4239
Cosmetopus longus|[1041]|JWDCG278-10|BOLD:AAH4239
Cosmetopus longus|[1042]|JWDCI654-10|BOLD:AAH4239
Cosmetopus longus|[1043]|JWDCI921-11|BOLD:AAH4239
Cosmetopus longus|[1044]|JWDCI923-11|BOLD:AAH4239
Pogonota|[1045]|JWDCJ190-11|BOLD:AAH4222
Pogonota|[1046]|JWDCC264-10|BOLD:AAH4222
Pogonota|[1047]|JWDCJ102-11|BOLD:AAH4222
Pogonota|[1048]|JWDCH179-10|BOLD:AAH4222
Pogonota|[1049]|JWDCL410-11|BOLD:AAH4222
Pogonota|[1050]|JWDCL472-11|BOLD:AAH4222
Pogonota|[1051]|JWDCF048-10|BOLD:AAH4222
Pogonota|[1052]|JWDCE519-10|BOLD:AAH4222
Pogonota|[1053]|JWDCG835-10|BOLD:AAH4222
Pogonota|[1054]|JWDCJ1540-11|BOLD:AAH4222
Pogonota|[1055]|MHFLI957-09|BOLD:AAH4222
Pogonota|[1056]|JWDCH841-10|BOLD:AAH4222
Pogonota|[1057]|JWDCF025-10|BOLD:AAH4222
Pogonota|[1058]|JWDCE518-10|BOLD:AAH4222
Pogonota|[1059]|JWDCH214-10|BOLD:AAH4222
Pogonota|[1060]|JWDCI922-11|BOLD:AAH4222
Pogonota|[1061]|JWDCF047-10|BOLD:AAH4222
Pogonota|[1062]|JWDCF096-10|BOLD:AAH4222
Pogonota|[1063]|JWDCJ187-11|BOLD:AAH4222
Pogonota|[1064]|JWDCJ189-11|BOLD:AAH4222
Pogonota|[1065]|JWDCE764-10|BOLD:AAH4222
Microprosopa|[1066]|JWDCI695-11|BOLD:AAH4224
Microprosopa|[1067]|TWDIP117-09|BOLD:AAH4224
Microprosopa|[1068]|JWDCJ710-11|BOLD:AAH4224
Microprosopa|[1069]|JWDCJ242-11|BOLD:AAH4224
Microprosopa|[1070]|JWDCJ244-11|BOLD:AAH4224
Microprosopa|[1071]|JWDCI697-11|BOLD:AAH4224
Microprosopa|[1072]|JWDCI575-10|BOLD:AAH4224
Microprosopa|[1073]|JWDCI924-11|BOLD:AAH4224
Microprosopa|[1074]|JWDCF275-10|BOLD:AAH4224
Microprosopa|[1075]|JWDCF324-10|BOLD:AAH4224
Microprosopa|[1076]|JWDCJ191-11|BOLD:AAH4224
Microprosopa|[1077]|JWDCB072-10|BOLD:AAH4224
Microprosopa|[1078]|TWDIP119-09|BOLD:AAH4224
Microprosopa|[1079]|JWDCB071-10|BOLD:AAH4224
Microprosopa|[1080]|JWDCJ1416-11|BOLD:AAH4224
Microprosopa|[1081]|JWDCJ245-11|BOLD:AAH4224
Microprosopa|[1082]|JWDCH314-10|BOLD:AAH4224
Microprosopa|[1083]|JWDCF274-10|BOLD:AAH4224
Microprosopa|[1084]|JWDCF272-10|BOLD:AAH4224
Scathophaginae|[1085]|JWDCD093-10|BOLD:AEU1321
Scathophaginae|[1086]|JWDCD071-10|BOLD:AEU1321
Scathophaginae|[1087]|JWDCI251-10|BOLD:AEU1321
Scathophaginae|[1088]|JWDCF659-10|BOLD:AAP9874
Scathophaginae|[1089]|JWDCF660-10|BOLD:AAP9874
Cordilura confusa|[1090]|JWDCB063-10|BOLD:AAH4218
Cordilura confusa|[1091]|MHFLI206-06|BOLD:AAH4218
Cordilura confusa|[1092]|JWDCC191-10|BOLD:AAH4218
Cordilura confusa|[1093]|JWDCB065-10|BOLD:AAH4218
Cordilura confusa|[1094]|JWDCB066-10|BOLD:AAH4218
Cordilura confusa|[1095]|JWDCB068-10|BOLD:AAH4218

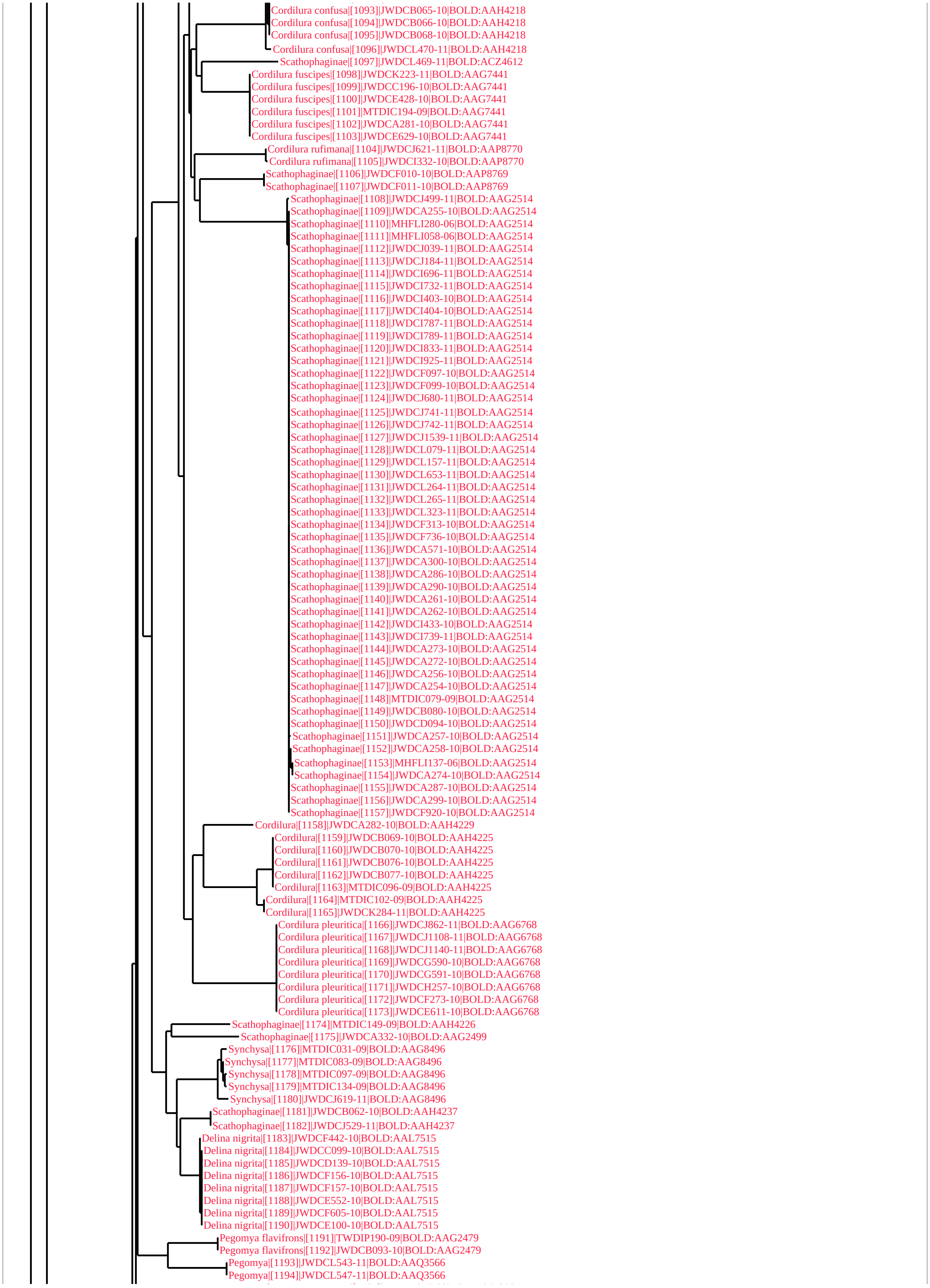
Cordilura confusa|[1093]|JWDCB065-10|BOLD:AAH4218
Cordilura confusa|[1094]|JWDCB066-10|BOLD:AAH4218
Cordilura confusa|[1095]|JWDCB068-10|BOLD:AAH4218
Cordilura confusa|[1096]|JWDCL470-11|BOLD:AAH4218
Scathophaginae|[1097]|JWDCL469-11|BOLD:ACZ4612
Cordilura fuscipes|[1098]|JWDCK223-11|BOLD:AAG7441
Cordilura fuscipes|[1099]|JWDCC196-10|BOLD:AAG7441
Cordilura fuscipes|[1100]|JWDCE428-10|BOLD:AAG7441
Cordilura fuscipes|[1101]|MTDIC194-09|BOLD:AAG7441
Cordilura fuscipes|[1102]|JWDCA281-10|BOLD:AAG7441
Cordilura fuscipes|[1103]|JWDCE629-10|BOLD:AAG7441
Cordilura rufimana|[1104]|JWDCJ621-11|BOLD:AAP8770
Cordilura rufimana|[1105]|JWDCI332-10|BOLD:AAP8770
Scathophaginae|[1106]|JWDCF010-10|BOLD:AAP8769
Scathophaginae|[1107]|JWDCF011-10|BOLD:AAP8769
Scathophaginae|[1108]|JWDCJ499-11|BOLD:AAG2514
Scathophaginae|[1109]|JWDCA255-10|BOLD:AAG2514
Scathophaginae|[1110]|MHFLI280-06|BOLD:AAG2514
Scathophaginae|[1111]|MHFLI058-06|BOLD:AAG2514
Scathophaginae|[1112]|JWDCJ039-11|BOLD:AAG2514
Scathophaginae|[1113]|JWDCJ184-11|BOLD:AAG2514
Scathophaginae|[1114]|JWDCI696-11|BOLD:AAG2514
Scathophaginae|[1115]|JWDCI732-11|BOLD:AAG2514
Scathophaginae|[1116]|JWDCI403-10|BOLD:AAG2514
Scathophaginae|[1117]|JWDCI404-10|BOLD:AAG2514
Scathophaginae|[1118]|JWDCI787-11|BOLD:AAG2514
Scathophaginae|[1119]|JWDCI789-11|BOLD:AAG2514
Scathophaginae|[1120]|JWDCI833-11|BOLD:AAG2514
Scathophaginae|[1121]|JWDCI925-11|BOLD:AAG2514
Scathophaginae|[1122]|JWDCF097-10|BOLD:AAG2514
Scathophaginae|[1123]|JWDCF099-10|BOLD:AAG2514
Scathophaginae|[1124]|JWDCJ680-11|BOLD:AAG2514
Scathophaginae|[1125]|JWDCJ741-11|BOLD:AAG2514
Scathophaginae|[1126]|JWDCJ742-11|BOLD:AAG2514
Scathophaginae|[1127]|JWDCJ1539-11|BOLD:AAG2514
Scathophaginae|[1128]|JWDCL079-11|BOLD:AAG2514
Scathophaginae|[1129]|JWDCL157-11|BOLD:AAG2514
Scathophaginae|[1130]|JWDCL653-11|BOLD:AAG2514
Scathophaginae|[1131]|JWDCL264-11|BOLD:AAG2514
Scathophaginae|[1132]|JWDCL265-11|BOLD:AAG2514
Scathophaginae|[1133]|JWDCL323-11|BOLD:AAG2514
Scathophaginae|[1134]|JWDCF313-10|BOLD:AAG2514
Scathophaginae|[1135]|JWDCF736-10|BOLD:AAG2514
Scathophaginae|[1136]|JWDCA571-10|BOLD:AAG2514
Scathophaginae|[1137]|JWDCA300-10|BOLD:AAG2514
Scathophaginae|[1138]|JWDCA286-10|BOLD:AAG2514
Scathophaginae|[1139]|JWDCA290-10|BOLD:AAG2514
Scathophaginae|[1140]|JWDCA261-10|BOLD:AAG2514
Scathophaginae|[1141]|JWDCA262-10|BOLD:AAG2514
Scathophaginae|[1142]|JWDCI433-10|BOLD:AAG2514
Scathophaginae|[1143]|JWDCI739-11|BOLD:AAG2514
Scathophaginae|[1144]|JWDCA273-10|BOLD:AAG2514
Scathophaginae|[1145]|JWDCA272-10|BOLD:AAG2514
Scathophaginae|[1146]|JWDCA256-10|BOLD:AAG2514
Scathophaginae|[1147]|JWDCA254-10|BOLD:AAG2514
Scathophaginae|[1148]|MTDIC079-09|BOLD:AAG2514
Scathophaginae|[1149]|JWDCB080-10|BOLD:AAG2514
Scathophaginae|[1150]|JWDCD094-10|BOLD:AAG2514
Scathophaginae|[1151]|JWDCA257-10|BOLD:AAG2514
Scathophaginae|[1152]|JWDCA258-10|BOLD:AAG2514
Scathophaginae|[1153]|MHFLI137-06|BOLD:AAG2514
Scathophaginae|[1154]|JWDCA274-10|BOLD:AAG2514
Scathophaginae|[1155]|JWDCA287-10|BOLD:AAG2514
Scathophaginae|[1156]|JWDCA299-10|BOLD:AAG2514
Scathophaginae|[1157]|JWDCF920-10|BOLD:AAG2514
Cordilura|[1158]|JWDCA282-10|BOLD:AAH4229
Cordilura|[1159]|JWDCB069-10|BOLD:AAH4225
Cordilura|[1160]|JWDCB070-10|BOLD:AAH4225
Cordilura|[1161]|JWDCB076-10|BOLD:AAH4225
Cordilura|[1162]|JWDCB077-10|BOLD:AAH4225
Cordilura|[1163]|MTDIC096-09|BOLD:AAH4225
Cordilura|[1164]|MTDIC102-09|BOLD:AAH4225
Cordilura|[1165]|JWDCK284-11|BOLD:AAH4225
Cordilura pleuritica|[1166]|JWDCJ862-11|BOLD:AAG6768
Cordilura pleuritica|[1167]|JWDCJ1108-11|BOLD:AAG6768
Cordilura pleuritica|[1168]|JWDCJ1140-11|BOLD:AAG6768
Cordilura pleuritica|[1169]|JWDCG590-10|BOLD:AAG6768
Cordilura pleuritica|[1170]|JWDCG591-10|BOLD:AAG6768
Cordilura pleuritica|[1171]|JWDCH257-10|BOLD:AAG6768
Cordilura pleuritica|[1172]|JWDCF273-10|BOLD:AAG6768
Cordilura pleuritica|[1173]|JWDCE611-10|BOLD:AAG6768
Scathophaginae|[1174]|MTDIC149-09|BOLD:AAH4226
Scathophaginae|[1175]|JWDCA332-10|BOLD:AAG2499
Synchysa|[1176]|MTDIC031-09|BOLD:AAG8496
Synchysa|[1177]|MTDIC083-09|BOLD:AAG8496
Synchysa|[1178]|MTDIC097-09|BOLD:AAG8496
Synchysa|[1179]|MTDIC134-09|BOLD:AAG8496
Synchysa|[1180]|JWDCJ619-11|BOLD:AAG8496
Scathophaginae|[1181]|JWDCB062-10|BOLD:AAH4237
Scathophaginae|[1182]|JWDCJ529-11|BOLD:AAH4237
Delina nigrita|[1183]|JWDCF442-10|BOLD:AAL7515
Delina nigrita|[1184]|JWDCC099-10|BOLD:AAL7515
Delina nigrita|[1185]|JWDCD139-10|BOLD:AAL7515
Delina nigrita|[1186]|JWDCF156-10|BOLD:AAL7515
Delina nigrita|[1187]|JWDCF157-10|BOLD:AAL7515
Delina nigrita|[1188]|JWDCE552-10|BOLD:AAL7515
Delina nigrita|[1189]|JWDCF605-10|BOLD:AAL7515
Delina nigrita|[1190]|JWDCE100-10|BOLD:AAL7515
Pegomya flavifrons|[1191]|TWDIP190-09|BOLD:AAG2479
Pegomya flavifrons|[1192]|JWDCB093-10|BOLD:AAG2479
Pegomya|[1193]|JWDCL543-11|BOLD:AAQ3566
Pegomya|[1194]|JWDCL547-11|BOLD:AAQ3566

Pegomya flavifrons|[1192]|JWDCB093-10|BOLD:AAG2479
Pegomya|[1193]|JWDCL543-11|BOLD:AAQ3566
Pegomya|[1194]|JWDCL547-11|BOLD:AAQ3566
Protophormia terraenovae|[1195]|TWDIP350-09|BOLD:AAC9614
Protophormia terraenovae|[1196]|JSFLA044-09|BOLD:AAC9614
Protophormia terraenovae|[1197]|MHFLI899-09|BOLD:AAC9614
Protophormia terraenovae|[1198]|TWDIP351-09|BOLD:AAC9614
Protophormia terraenovae|[1199]|TWDIP352-09|BOLD:AAC9614
Protocalliphora sapphira|[1200]|JWDCJ1260-11|BOLD:AAH3037
Protocalliphora sapphira|[1201]|JWDCJ1261-11|BOLD:AAH3037
Protocalliphora sapphira|[1202]|JWDCH178-10|BOLD:AAH3037
Protocalliphora spatulata|[1203]|JWDCE307-10|BOLD:AAH3037
Protocalliphora spatulata|[1204]|JWDCF162-10|BOLD:AAH3029
Protocalliphora spatulata|[1205]|JWDCK108-11|BOLD:AAH3029
Protocalliphora spatulata|[1206]|JWDCH555-10|BOLD:AAH3029
Protocalliphora spatulata|[1207]|DSCNI009-07|BOLD:AAH3029
Protocalliphora spatulata|[1208]|JWDCE613-10|BOLD:AAH3029
Protocalliphora spatulata|[1209]|JWDCE114-10|BOLD:AAH3029
Protocalliphora spatulata|[1210]|JWDCE460-10|BOLD:AAH3029
Protocalliphora spatulata|[1211]|JWDCE461-10|BOLD:AAH3029
Protocalliphora spatulata|[1212]|JWDCE557-10|BOLD:AAH3029
Protocalliphora spatulata|[1213]|JWDCE558-10|BOLD:AAH3029
Protocalliphora spatulata|[1214]|JWDCE559-10|BOLD:AAH3029
Protocalliphora spatulata|[1215]|JWDCE612-10|BOLD:AAH3029
Protocalliphora spatulata|[1216]|JWDCE306-10|BOLD:AAH3029
Protocalliphora spatulata|[1217]|JWDCE376-10|BOLD:AAH3029
Protocalliphora spatulata|[1218]|JWDCH557-10|BOLD:AAH3029
Protocalliphora spatulata|[1219]|JWDCH556-10|BOLD:AAH3029
Protocalliphora spatulata|[1220]|JWDCH044-10|BOLD:AAH3029
Protocalliphora spatulata|[1221]|JWDCI114-10|BOLD:AAH3029
Protocalliphora spatulata|[1222]|JWDCG839-10|BOLD:AAH3029
Protocalliphora spatulata|[1223]|JWDCJ089-11|BOLD:AAH3029
Protocalliphora spatulata|[1224]|JWDCJ1390-11|BOLD:AAH3029
Protocalliphora spatulata|[1225]|JWDCJ820-11|BOLD:AAH3029
Protocalliphora spatulata|[1226]|JWDCJ662-11|BOLD:AAH3029
Protocalliphora spatulata|[1227]|JWDCJ552-11|BOLD:AAH3029
Protocalliphora spatulata|[1228]|JWDCC015-10|BOLD:AAH3029
Protocalliphora metallica|[1229]|JWDCB057-10|BOLD:AAH3029
Protocalliphora spatulata|[1230]|MHFLY029-06|BOLD:AAH3029
Protocalliphora spatulata|[1231]|MTDIC197-09|BOLD:AAH3029
Protocalliphora metallica|[1232]|JWDCB058-10|BOLD:AAH3029
Protocalliphora spatulata|[1233]|JWDCH713-10|BOLD:AAH3029
Protocalliphora spatulata|[1234]|JWDCE305-10|BOLD:AAH3029
Protocalliphora spatulata|[1235]|JWDCE115-10|BOLD:AAH3029
Lucilia illustris|[1236]|MHFLY030-06|BOLD:AAA7470
Lucilia illustris|[1237]|MHFLI237-06|BOLD:AAA7470
Lucilia illustris|[1238]|MHFLY031-06|BOLD:AAA7470
Lucilia illustris|[1239]|MTDIC101-09|BOLD:AAA7470
Lucilia illustris|[1240]|MTDIC198-09|BOLD:AAA7470
Lucilia illustris|[1241]|JWDCJ1262-11|BOLD:AAA7470
Lucilia magnicornis|[1242]|JWDCI407-10|BOLD:AAA6618
Lucilia magnicornis|[1243]|MHFLI179-06|BOLD:AAA6618
Lucilia sericata|[1244]|MTDIC199-09|BOLD:AAA6618
Lucilia magnicornis|[1245]|JWDCJ538-11|BOLD:AAA6618
Lucilia magnicornis|[1246]|JWDCK201-11|BOLD:AAA6618
Lucilia magnicornis|[1247]|JWDCF555-10|BOLD:AAA6618
Calliphora genarum|[1248]|MHFLI188-06|BOLD:AAH3030
Calliphora genarum|[1249]|MHFLI257-06|BOLD:AAH3030
Calliphora genarum|[1250]|JWDCC236-10|BOLD:AAH3030
Calliphora genarum|[1251]|JWDCJ1160-11|BOLD:AAH3030
Calliphora genarum|[1252]|JWDCL176-11|BOLD:AAH3030
Calliphora genarum|[1253]|JWDCJ094-11|BOLD:AAH3030
Calliphora montana|[1254]|JWDCJ1453-11|BOLD:ABZ4059
Calliphora montana|[1255]|JWDCH043-10|BOLD:ABZ4059
Calliphora montana|[1256]|JWDCI791-11|BOLD:ABZ4059
Calliphora montana|[1257]|MHFLY379-09|BOLD:ABZ4059
Calliphora montana|[1258]|JWDCA535-10|BOLD:ABZ4059
Calliphora montana|[1259]|JWDCA536-10|BOLD:ABZ4059
Calliphora montana|[1260]|JWDCJ898-11|BOLD:ABZ4059
Calliphora montana|[1261]|JWDCJ968-11|BOLD:ABZ4059
Calliphora montana|[1262]|JWDCH714-10|BOLD:ABZ4059
Calliphora montana|[1263]|JWDCG800-10|BOLD:ABZ4059
Calliphora montana|[1264]|JWDCI113-10|BOLD:ABZ4059
Calliphora montana|[1265]|JWDCI155-10|BOLD:ABZ4059
Calliphora montana|[1266]|JWDCH545-10|BOLD:ABZ4059
Calliphora terraenovae|[1267]|MHFLI245-06|BOLD:AGO4847
Calliphora terraenovae|[1268]|MHFLI242-06|BOLD:AGO4847
Calliphora terraenovae|[1269]|MHFLI180-06|BOLD:AGO4847
Calliphora terraenovae|[1270]|JWDCE830-10|BOLD:AGO4847
Cynomya|[1271]|MHFLI243-06|BOLD:AAB0868
Cynomya|[1272]|JWDCF315-10|BOLD:AAB0868
Cynomya|[1273]|JWDCE643-10|BOLD:AAB0868
Cynomya|[1274]|JSFLA034-08|BOLD:AAB0868
Cynomya|[1275]|JWDCI633-10|BOLD:AAB0868
Cynomya|[1276]|JWDCE709-10|BOLD:AAB0868
Cynomya|[1277]|JWDCE897-10|BOLD:AAB0868
Cynomya|[1278]|JWDCF556-10|BOLD:AAB0868
Cynomya|[1279]|JWDCG486-10|BOLD:AAB0868
Cynomya|[1280]|JWDCE377-10|BOLD:AAB0868
Cynomya|[1281]|JWDCE133-10|BOLD:AAB0868
Cynomya|[1282]|MHFLI244-06|BOLD:AAB0868
Cynomya|[1283]|MTDIC235-09|BOLD:AAB0868
Cynomya|[1284]|MHFLI246-06|BOLD:AAB0868
Cynomya|[1285]|MHFLI240-06|BOLD:AAB0868
Cynomya|[1286]|MHFLI250-06|BOLD:AAB0868
Cynomya|[1287]|MHFLY035-06|BOLD:AAB0868
Cynomya|[1288]|MTDIC221-09|BOLD:AAB0868
Cynomya|[1289]|JWDCA521-10|BOLD:AAB0868
Cynomya|[1290]|JWDCG485-10|BOLD:AAB0868
Cynomya|[1291]|JWDCA513-10|BOLD:AAB0868
Cynomya|[1292]|JWDCJ450-11|BOLD:AAB0868
Cynomya|[1293]|HMCOL076-07|BOLD:AAB0868
Cynomya|[1294]|HMCOL078-07|BOLD:AAB0868

Cynomya|[1292]|JWDCJ450-11|BOLD:AAB0868
Cynomya|[1293]|HMCOL076-07|BOLD:AAB0868
Cynomya|[1294]|HMCOL078-07|BOLD:AAB0868
Cynomya|[1295]|HMCOL079-07|BOLD:AAB0868
Cynomya|[1296]|HMCOL081-07|BOLD:AAB0868
Cynomya|[1297]|HMCOL082-07|BOLD:AAB0868
Cynomya|[1298]|HMCOL083-07|BOLD:AAB0868
Cynomya|[1299]|HMCOL087-07|BOLD:AAB0868
Cynomya|[1300]|HMCOL077-07|BOLD:AAB0868
Cynomya|[1301]|MHFLI185-06|BOLD:AAB0868
Cynomya|[1302]|HMCOL084-07|BOLD:AAB0868
Cynomya|[1303]|HMCOL088-07|BOLD:AAB0868
Cynomya|[1304]|JWDCF680-10|BOLD:AAB0868
Cynomya|[1305]|JWDCF859-10|BOLD:AAB0868
Cynomya|[1306]|JWDCF206-10|BOLD:AAB0868
Cynomya|[1307]|JWDCF207-10|BOLD:AAB0868
Cynomya|[1308]|JWDCF029-10|BOLD:AAB0868
Cynomya|[1309]|JWDCF133-10|BOLD:AAB0868
Cynomya|[1310]|JWDCH117-10|BOLD:AAB0868
Cynomya|[1311]|JWDCH245-10|BOLD:AAB0868
Cynomya|[1312]|JWDCE685-10|BOLD:AAB0868
Cynomya|[1313]|JWDCF433-10|BOLD:AAB0868
Cynomya|[1314]|JWDCF314-10|BOLD:AAB0868
Cynomya|[1315]|JWDCE655-10|BOLD:AAB0868
Cynomya|[1316]|JWDCG487-10|BOLD:AAB0868
Cynomya|[1317]|JWDCF028-10|BOLD:AAB0868
Cynomya|[1318]|JWDCG393-10|BOLD:AAB0868
Cynomya|[1319]|JWDCH401-10|BOLD:AAB0868
Cynomya|[1320]|JWDCI152-10|BOLD:AAB0868
Cynomya|[1321]|JWDCI874-11|BOLD:AAB0868
Cynomya|[1322]|JWDCG571-10|BOLD:AAB0868
Cynomya|[1323]|JWDCH938-10|BOLD:AAB0868
Cynomya|[1324]|JWDCJ539-11|BOLD:AAB0868
Cynomya|[1325]|JWDCA520-10|BOLD:AAB0868
Cynomya|[1326]|JWDCA517-10|BOLD:AAB0868
Cynomya|[1327]|JWDCA516-10|BOLD:AAB0868
Cynomya|[1328]|JWDCB059-10|BOLD:AAB0868
Cynomya|[1329]|HMCOL080-07|BOLD:AAB0868
Cynomya|[1330]|MHFLI184-06|BOLD:AAB0868
Cynomya|[1331]|MHFLI241-06|BOLD:AAB0868
Cynomya|[1332]|MHFLI186-06|BOLD:AAB0868
Cynomya|[1333]|MHFLI181-06|BOLD:AAB0868
Cynomya|[1334]|MHFLI247-06|BOLD:AAB0868
Cynomya|[1335]|JSFLA028-08|BOLD:AAB0868
Cynomya|[1336]|JWDCF640-10|BOLD:AAB0868
Cynomya|[1337]|JSFLA039-08|BOLD:AAB0868
Cynomya|[1338]|JWDCF762-10|BOLD:AAB0868
Cynomya|[1339]|HMCOL086-07|BOLD:AAB0868
Cynomya|[1340]|JWDCE462-10|BOLD:AAB0868
Cynomya|[1341]|JWDCE134-10|BOLD:AAB0868
Cynomya|[1342]|JWDCE265-10|BOLD:AAB0868
Anthomyiidae|[1343]|JWDCA469-10|BOLD:AAM6382
Anthomyiidae|[1344]|JWDCA345-10|BOLD:AAM6378
Anthomyiidae|[1345]|JWDCA342-10|BOLD:AAM6378
Anthomyiidae|[1346]|JWDCA348-10|BOLD:AAM6378
Anthomyiidae|[1347]|JWDCL262-11|BOLD:AAM6378
Anthomyiidae|[1348]|JWDCI024-10|BOLD:AAP8835
Anthomyiidae|[1349]|JWDCJ1280-11|BOLD:AAP8840
Anthomyiidae|[1350]|JWDCG413-10|BOLD:AAP8840
Zaphne barbiventris|[1351]|JWDCE293-10|BOLD:AAG2501
Zaphne barbiventris|[1352]|JWDCE697-10|BOLD:AAG2501
Zaphne barbiventris|[1353]|JWDCE165-10|BOLD:AAG2501
Zaphne barbiventris|[1354]|JWDCE295-10|BOLD:AAG2501
Zaphne barbiventris|[1355]|JWDCF049-10|BOLD:AAG2501
Zaphne barbiventris|[1356]|JWDCE099-10|BOLD:AAG2501
Zaphne barbiventris|[1357]|JWDCE167-10|BOLD:AAG2501
Zaphne zetterstedtii|[1358]|TWDIP095-09|BOLD:AAG2443
Zaphne zetterstedtii|[1359]|MHFLI133-06|BOLD:AAG2443
Zaphne zetterstedtii|[1360]|JWDCI426-10|BOLD:AAG2443
Zaphne zetterstedtii|[1361]|JWDCJ918-11|BOLD:AAG2443
Zaphne zetterstedtii|[1362]|JWDCJ550-11|BOLD:AAG2443
Zaphne zetterstedtii|[1363]|JWDCJ115-11|BOLD:AAG2443
Zaphne zetterstedtii|[1364]|JWDCJ113-11|BOLD:AAG2443
Zaphne zetterstedtii|[1365]|JWDCJ107-11|BOLD:AAG2443
Zaphne zetterstedtii|[1366]|JWDCJ503-11|BOLD:AAG2443
Zaphne zetterstedtii|[1367]|JWDCC103-10|BOLD:AAG2443
Zaphne zetterstedtii|[1368]|JWDCD095-10|BOLD:AAG2443
Zaphne zetterstedtii|[1369]|JWDCJ1612-11|BOLD:AAG2443
Zaphne zetterstedtii|[1370]|JWDCE555-10|BOLD:AAG2443
Zaphne nearctica|[1371]|MHFLI920-09|BOLD:AAG2434
Zaphne nearctica|[1372]|MHFLI189-06|BOLD:AAG2434
Zaphne nearctica|[1373]|MHFLI248-06|BOLD:AAG2434
Zaphne nearctica|[1374]|MHFLI259-06|BOLD:AAG2434
Zaphne nearctica|[1375]|TWDIP187-09|BOLD:AAG2434
Zaphne subarctica|[1376]|JWDCE691-10|BOLD:AAG2434
Zaphne subarctica|[1377]|JWDCE485-10|BOLD:AAG2434
Zaphne subarctica|[1378]|JWDCL155-11|BOLD:AAG2434
Zaphne subarctica|[1379]|JWDCE837-10|BOLD:AAG2434
Zaphne subarctica|[1380]|JWDCE486-10|BOLD:AAG2434
Zaphne lineatocollis|[1381]|MHFLY001-06|BOLD:AAG2434
Zaphne lineatocollis|[1382]|JWDCI916-11|BOLD:AAG2434
Zaphne lineatocollis|[1383]|JWDCG115-10|BOLD:AAG2434
Zaphne lineatocollis|[1384]|JWDCG114-10|BOLD:AAG2434
Zaphne lineatocollis|[1385]|JWDCG113-10|BOLD:AAG2434
Zaphne lineatocollis|[1386]|JWDCI431-10|BOLD:AAG2434
Zaphne lineatocollis|[1387]|JWDCI430-10|BOLD:AAG2434
Zaphne lineatocollis|[1388]|JWDCI427-10|BOLD:AAG2434
Zaphne lineatocollis|[1389]|JWDCI211-10|BOLD:AAG2434
Zaphne lineatocollis|[1390]|JWDCJ1180-11|BOLD:AAG2434
Zaphne lineatocollis|[1391]|JWDCJ1163-11|BOLD:AAG2434
Zaphne lineatocollis|[1392]|JWDCJ666-11|BOLD:AAG2434
Zaphne lineatocollis|[1393]|JWDCJ569-11|BOLD:AAG2434

Zaphne lineatocollis|[1391]|JWDCJ1163-11|BOLD:AAG2434
Zaphne lineatocollis|[1392]|JWDCJ666-11|BOLD:AAG2434
Zaphne lineatocollis|[1393]|JWDCJ569-11|BOLD:AAG2434
Zaphne lineatocollis|[1394]|JWDCJ549-11|BOLD:AAG2434
Zaphne lineatocollis|[1395]|JWDCJ545-11|BOLD:AAG2434
Zaphne lineatocollis|[1396]|JWDCJ544-11|BOLD:AAG2434
Zaphne lineatocollis|[1397]|JWDCJ543-11|BOLD:AAG2434
Zaphne lineatocollis|[1398]|JWDCC422-10|BOLD:AAG2434
Zaphne lineatocollis|[1399]|JWDCC395-10|BOLD:AAG2434
Zaphne lineatocollis|[1400]|JWDCC381-10|BOLD:AAG2434
Zaphne lineatocollis|[1401]|JWDCC265-10|BOLD:AAG2434
Zaphne lineatocollis|[1402]|JWDCC261-10|BOLD:AAG2434
Zaphne lineatocollis|[1403]|JWDCC023-10|BOLD:AAG2434
Zaphne lineatocollis|[1404]|JWDCC251-10|BOLD:AAG2434
Zaphne lineatocollis|[1405]|JWDCC275-10|BOLD:AAG2434
Zaphne lineatocollis|[1406]|JWDCC184-10|BOLD:AAG2434
Zaphne lineatocollis|[1407]|JWDCB086-10|BOLD:AAG2434
Zaphne lineatocollis|[1408]|JWDCB082-10|BOLD:AAG2434
Zaphne lineatocollis|[1409]|JWDCF012-10|BOLD:AAG2434
Zaphne lineatocollis|[1410]|JWDCE620-10|BOLD:AAG2434
Zaphne lineatocollis|[1411]|JWDCF469-10|BOLD:AAG2434
Zaphne lineatocollis|[1412]|JWDCF569-10|BOLD:AAG2434
Zaphne lineatocollis|[1413]|JWDCF540-10|BOLD:AAG2434
Zaphne lineatocollis|[1414]|JWDCJ506-11|BOLD:AAG2434
Zaphne lineatocollis|[1415]|JWDCE459-10|BOLD:AAG2434
Zaphne lineatocollis|[1416]|MHFLI262-06|BOLD:AAG2434
Zaphne lineatocollis|[1417]|JWDCE470-10|BOLD:AAG2434
Zaphne lineatocollis|[1418]|JWDCE891-10|BOLD:AAG2434
Zaphne lineatocollis|[1419]|JWDCE234-10|BOLD:AAG2434
Zaphne lineatocollis|[1420]|JWDCF596-10|BOLD:AAG2434
Zaphne lineatocollis|[1421]|JWDCE712-10|BOLD:AAG2434
Zaphne lineatocollis|[1422]|JWDCE641-10|BOLD:AAG2434
Zaphne lineatocollis|[1423]|JWDCF457-10|BOLD:AAG2434
Zaphne lineatocollis|[1424]|JWDCE458-10|BOLD:AAG2434
Zaphne lineatocollis|[1425]|JWDCE599-10|BOLD:AAG2434
Zaphne lineatocollis|[1426]|JWDCE089-10|BOLD:AAG2434
Zaphne lineatocollis|[1427]|JWDCE241-10|BOLD:AAG2434
Zaphne lineatocollis|[1428]|JWDCE331-10|BOLD:AAG2434
Zaphne lineatocollis|[1429]|JWDCE457-10|BOLD:AAG2434
Zaphne lineatocollis|[1430]|JWDCG205-10|BOLD:AAG2434
Zaphne lineatocollis|[1431]|JWDCE554-10|BOLD:AAG2434
Zaphne lineatocollis|[1432]|JWDCJ1342-11|BOLD:AAG2434
Zaphne lineatocollis|[1433]|JWDCI400-10|BOLD:AAG2434
Zaphne lineatocollis|[1434]|JWDCK259-11|BOLD:AAG2434
Zaphne lineatocollis|[1435]|JWDCJ802-11|BOLD:AAG2434
Zaphne lineatocollis|[1436]|JWDCJ631-11|BOLD:AAG2434
Zaphne lineatocollis|[1437]|JWDCJ674-11|BOLD:AAG2434
Zaphne lineatocollis|[1438]|JWDCJ579-11|BOLD:AAG2434
Zaphne lineatocollis|[1439]|JWDCJ630-11|BOLD:AAG2434
Zaphne lineatocollis|[1440]|JWDCJ548-11|BOLD:AAG2434
Zaphne lineatocollis|[1441]|JWDCJ571-11|BOLD:AAG2434
Zaphne lineatocollis|[1442]|JWDCJ520-11|BOLD:AAG2434
Zaphne lineatocollis|[1443]|JWDCJ523-11|BOLD:AAG2434
Zaphne lineatocollis|[1444]|JWDCI432-10|BOLD:AAG2434
Zaphne lineatocollis|[1445]|JWDCG112-10|BOLD:AAG2434
Zaphne lineatocollis|[1446]|JWDCJ515-11|BOLD:AAG2434
Zaphne lineatocollis|[1447]|JWDCJ516-11|BOLD:AAG2434
Zaphne lineatocollis|[1448]|JWDCJ511-11|BOLD:AAG2434
Zaphne lineatocollis|[1449]|JWDCJ307-11|BOLD:AAG2434
Zaphne lineatocollis|[1450]|JWDCJ114-11|BOLD:AAG2434
Zaphne lineatocollis|[1451]|JWDCJ095-11|BOLD:AAG2434
Zaphne lineatocollis|[1452]|JWDCJ504-11|BOLD:AAG2434
Zaphne lineatocollis|[1453]|JWDCB078-10|BOLD:AAG2434
Zaphne lineatocollis|[1454]|JWDCC097-10|BOLD:AAG2434
Zaphne lineatocollis|[1455]|JWDCJ510-11|BOLD:AAG2434
Zaphne lineatocollis|[1456]|JWDCE264-10|BOLD:AAG2434
Zaphne ambigua|[1457]|MHFLI261-06|BOLD:AAG2447
Zaphne ambigua|[1458]|JWDCI542-10|BOLD:AAG2447
Zaphne implicata|[1459]|JWDCI026-10|BOLD:AAG2442
Zaphne implicata|[1460]|JWDCJ1488-11|BOLD:AAG2442
Zaphne implicata|[1461]|JWDCF539-10|BOLD:AAG2442
Zaphne implicata|[1462]|MHFLI123-06|BOLD:AAG2442
Zaphne implicata|[1463]|MHFLI254-06|BOLD:AAG2442
Zaphne implicata|[1464]|MHFLI268-06|BOLD:AAG2442
Zaphne implicata|[1465]|JWDCH513-10|BOLD:AAG2442
Zaphne implicata|[1466]|JWDCJ1344-11|BOLD:AAG2442
Zaphne divisa|[1467]|TWDIP186-09|BOLD:AAG2441
Zaphne divisa|[1468]|MHFLI122-06|BOLD:AAG2441
Zaphne divisa|[1469]|MHFLY357-09|BOLD:AAG2441
Zaphne divisa|[1470]|JWDCA369-10|BOLD:AAG2441
Zaphne divisa|[1471]|JWDCA354-10|BOLD:AAG2441
Zaphne divisa|[1472]|JWDCA366-10|BOLD:AAG2441
Zaphne divisa|[1473]|JWDCA367-10|BOLD:AAG2441
Zaphne divisa|[1474]|JWDCJ921-11|BOLD:AAG2441
Zaphne divisa|[1475]|MHFLY380-09|BOLD:AAG2441
Zaphne divisa|[1476]|JWDCA412-10|BOLD:AAG2441
Zaphne divisa|[1477]|JWDCK193-11|BOLD:AAG2441
Zaphne divisa|[1478]|JWDCH413-10|BOLD:AAG2441
Zaphne brunneifrons|[1479]|JWDCJ1343-11|BOLD:ACF0122
Zaphne brunneifrons|[1480]|JWDCI543-10|BOLD:ACF0122
Zaphne brunneifrons|[1481]|JWDCG375-10|BOLD:ACF0122
Zaphne brunneifrons|[1482]|JWDCG481-10|BOLD:ACF0122
Zaphne brunneifrons|[1483]|JWDCA329-10|BOLD:ACF0122
Zaphne brunneifrons|[1484]|TWDIP180-09|BOLD:ACF0122
Zaphne brunneifrons|[1485]|MHFLI908-09|BOLD:ACF0122
Zaphne brunneifrons|[1486]|JWDCH514-10|BOLD:ACF0122
Zaphne brunneifrons|[1487]|JWDCI803-11|BOLD:ACF0122
Zaphne brunneifrons|[1488]|JWDCH601-10|BOLD:ACF0122
Zaphne brunneifrons|[1489]|JWDCJ948-11|BOLD:ACF0122
Zaphne brunneifrons|[1490]|JWDCJ947-11|BOLD:ACF0122
Zaphne brunneifrons|[1491]|JWDCJ513-11|BOLD:ACF0122
Zaphne brunneifrons|[1492]|JWDCJ180-11|BOLD:ACF0122
Zaphne brunneifrons|[1493]|JWDCG786-10|BOLD:ACF0122

Zaphne brunneifrons|[1491]|JWDCJ513-11|BOLD:ACF0122
Zaphne brunneifrons|[1492]|JWDCJ180-11|BOLD:ACF0122
Zaphne brunneifrons|[1493]|JWDCG786-10|BOLD:ACF0122
Zaphne brunneifrons|[1494]|MHFLI255-06|BOLD:ACF0122
Zaphne brunneifrons|[1495]|MHFLI615-07|BOLD:ACF0122
Zaphne brunneifrons|[1496]|JWDCG530-10|BOLD:ACF0122
Zaphne brunneifrons|[1497]|JWDCE211-10|BOLD:ACF0122
Zaphne proxima|[1498]|MHFLI918-09|BOLD:AAG2470
Zaphne altilega|[1499]|MHFLI917-09|BOLD:ABX6111
Zaphne altilega|[1500]|JWDCG559-10|BOLD:ABX6111
Zaphne altilega|[1501]|JWDCE280-10|BOLD:ABX6111
Zaphne|[1502]|TWDIP179-09|BOLD:ABY6007
Zaphne|[1503]|JWDCG480-10|BOLD:AAG1723
Zaphne tundrica|[1504]|JWDCG379-10|BOLD:AAG1723
Zaphne tundrica|[1505]|JWDCG570-10|BOLD:AAG1723
Zaphne tundrica|[1506]|JWDCL063-11|BOLD:AAG1723
Zaphne tundrica|[1507]|MHFLI253-06|BOLD:AAG1723
Zaphne tundrica|[1508]|MHFLI190-06|BOLD:AAG1723
Zaphne tundrica|[1509]|JWDCI513-10|BOLD:AAG1723
Zaphne tundrica|[1510]|JWDCH099-10|BOLD:AAG1723
Zaphne tundrica|[1511]|JWDCG348-10|BOLD:AAG1723
Zaphne tundrica|[1512]|JWDCG349-10|BOLD:AAG1723
Zaphne tundrica|[1513]|JWDCG376-10|BOLD:AAG1723
Zaphne tundrica|[1514]|JWDCG426-10|BOLD:AAG1723
Zaphne tundrica|[1515]|JWDCG479-10|BOLD:AAG1723
Zaphne gymnura|[1516]|MHFLY378-09|BOLD:AAG1723
Zaphne gymnura|[1517]|MHFLY368-09|BOLD:AAG1723
Zaphne gymnura|[1518]|JWDCI097-10|BOLD:AAG1723
Zaphne gymnura|[1519]|JWDCH359-10|BOLD:AAG1723
Zaphne gymnura|[1520]|MHFLI252-06|BOLD:AAG1723
Zaphne gymnura|[1521]|JWDCH986-10|BOLD:AAG1723
Zaphne gymnura|[1522]|JWDCI551-10|BOLD:AAG1723
Zaphne gymnura|[1523]|JWDCH038-10|BOLD:AAG1723
Zaphne gymnura|[1524]|JWDCG005-10|BOLD:AAG1723
Zaphne gymnura|[1525]|JWDCG038-10|BOLD:AAG1723
Zaphne gymnura|[1526]|JWDCG681-10|BOLD:AAG1723
Zaphne gymnura|[1527]|JWDCJ512-11|BOLD:AAG1723
Zaphne gymnura|[1528]|JWDCC101-10|BOLD:AAG1723
Zaphne gymnura|[1529]|MHFLI210-06|BOLD:AAG1723
Zaphne gymnura|[1530]|MHFLI274-06|BOLD:AAG1723
Zaphne gymnura|[1531]|MHFLI207-06|BOLD:AAG1723
Zaphne gymnura|[1532]|MHFLI208-06|BOLD:AAG1723
Zaphne gymnura|[1533]|MHFLY019-06|BOLD:AAG1723
Zaphne gymnura|[1534]|MHFLI204-06|BOLD:AAG1723
Zaphne gymnura|[1535]|MHFLI202-06|BOLD:AAG1723
Zaphne gymnura|[1536]|MHFLI201-06|BOLD:AAG1723
Zaphne gymnura|[1537]|MHFLI200-06|BOLD:AAG1723
Zaphne gymnura|[1538]|MHFLI197-06|BOLD:AAG1723
Zaphne gymnura|[1539]|MHFLI196-06|BOLD:AAG1723
Zaphne gymnura|[1540]|MHFLI194-06|BOLD:AAG1723
Zaphne gymnura|[1541]|MHFLI263-06|BOLD:AAG1723
Zaphne gymnura|[1542]|JWDCG424-10|BOLD:AAG1723
Zaphne gymnura|[1543]|JWDCH548-10|BOLD:AAG1723
Zaphne gymnura|[1544]|JWDCG423-10|BOLD:AAG1723
Zaphne gymnura|[1545]|MHFLI912-09|BOLD:AAG1723
Zaphne gymnura|[1546]|JWDCH551-10|BOLD:AAG1723
Fannia|[1547]|MTDIC279-09|BOLD:AAD9983
Fannia|[1548]|MFDC653-09|BOLD:AAD9983
Fannia atripes|[1549]|JWDCJ800-11|BOLD:AAD9984
Fannia atripes|[1550]|MFDC318-09|BOLD:AAD9984
Fannia atripes|[1551]|MFDC655-09|BOLD:AAD9984
Fannia atripes|[1552]|MFDC654-09|BOLD:AAD9984
Fannia atripes|[1553]|JWDCG003-10|BOLD:AAD9984
Fannia atripes|[1554]|JWDCJ1181-11|BOLD:AAD9984
Fannia atripes|[1555]|MFDC319-09|BOLD:AAD9984
Fannia abrupta|[1556]|MFDC315-09|BOLD:AAJ0307
Fannia abrupta|[1557]|MFDC314-09|BOLD:AAJ0307
Fannia|[1558]|MFDC1090-11|BOLD:AAV6534
Fannia aethiops|[1559]|MFDC302-09|BOLD:AAM6399
Fannia aethiops|[1560]|JWDCA453-10|BOLD:AAM6399
Fannia|[1561]|JWDCJ1299-11|BOLD:AAM6399
Fannia|[1562]|JWDCA455-10|BOLD:AAM6399
Fannia|[1563]|JWDCL430-11|BOLD:AAM6399
Fannia aethiops|[1564]|MFDC303-09|BOLD:AAM6399
Fannia aethiops|[1565]|MFDC304-09|BOLD:AAM6399
Fannia aethiops|[1566]|MFDC305-09|BOLD:AAM6399
Fannia postica|[1567]|MFDC306-09|BOLD:AAF7093
Fannia postica|[1568]|MFDC307-09|BOLD:AAF7093
Fannia scyphocerca|[1569]|MFDC661-09|BOLD:AAJ0316
Fannia scyphocerca|[1570]|MFDC662-09|BOLD:AAJ0316
Fannia scyphocerca|[1571]|JWDCJ1294-11|BOLD:AAJ0316
Fannia scyphocerca|[1572]|JWDCA563-10|BOLD:AAJ0316
Fannia scyphocerca|[1573]|JWDCJ1284-11|BOLD:AAJ0316
Fannia scyphocerca|[1574]|MTDIC206-09|BOLD:AAJ0316
Fannia scyphocerca|[1575]|JWDCG850-10|BOLD:AAJ0316
Fannia scyphocerca|[1576]|MFDC312-09|BOLD:AAJ0316
Fannia tundrarum|[1577]|MFDC317-09|BOLD:ABA2387
Fannia|[1578]|MFDC658-09|BOLD:AAL5108
Fannia falcata|[1579]|JWDCF845-10|BOLD:AAF7100
Fannia falcata|[1580]|JWDCH408-10|BOLD:AAF7100
Fannia falcata|[1581]|MFDC299-09|BOLD:AAF7100
Fannia falcata|[1582]|JWDCK194-11|BOLD:AAF7100
Fannia falcata|[1583]|MHCOL462-07|BOLD:AAF7100
Fannia falcata|[1584]|JWDCH247-10|BOLD:AAF7100
Fannia falcata|[1585]|MFDC298-09|BOLD:AAF7100
Fannia falcata|[1586]|MFDC300-09|BOLD:AAF7100
Fannia lugubrina|[1587]|JWDCJ637-11|BOLD:AAJ0305
Fannia lugubrina|[1588]|JWDCC277-10|BOLD:AAJ0305
Fannia lugubrina|[1589]|MFDC308-09|BOLD:AAJ0305
Fannia lugubrina|[1590]|MFDC309-09|BOLD:AAJ0305
Fannia metallipennis|[1591]|MFDC650-09|BOLD:AAJ0306
Fannia metallipennis|[1592]|MFDC649-09|BOLD:AAJ0306
[illegible]

Fannia lugubrina|[1590]|MFDC309-09|BOLD:AAJ0305
Fannia metallipennis|[1591]|MFDC650-09|BOLD:AAJ0306
Fannia metallipennis|[1592]|MFDC649-09|BOLD:AAJ0306
Fannia metallipennis|[1593]|MFDC651-09|BOLD:AAJ0306
Fannia metallipennis|[1594]|MFDC652-09|BOLD:AAJ0306
Fannia metallipennis|[1595]|MFDC316-09|BOLD:AAJ0306
Fannia fuscitibia|[1596]|MFDC310-09|BOLD:AAG4621
Fannia fuscitibia|[1597]|MFDC311-09|BOLD:AAG4621
Fannia|[1598]|MTDIC246-09|BOLD:AAG4621
Fannia corvina|[1599]|JWDCG087-10|BOLD:AAG4632
Fannia corvina|[1600]|JWDCA608-10|BOLD:AAG4632
Fannia corvina|[1601]|JWDCJ1293-11|BOLD:AAG4632
Fannia corvina|[1602]|JWDCK262-11|BOLD:AAG4632
Fannia corvina|[1603]|MFDC322-09|BOLD:AAG4632
Alliopsis fractiseta|[1604]|JWDCE302-10|BOLD:AAL7529
Alliopsis fractiseta|[1605]|JWDCJ448-11|BOLD:AAL7529
Alliopsis fractiseta|[1606]|JWDCJ667-11|BOLD:AAL7529
Alliopsis fractiseta|[1607]|JWDCI892-11|BOLD:AAL7529
Alliopsis fractiseta|[1608]|JWDCE603-10|BOLD:AAL7529
Alliopsis fractiseta|[1609]|JWDCJ572-11|BOLD:AAL7529
Alliopsis fractiseta|[1610]|JWDCD022-10|BOLD:AAL7529
Alliopsis fractiseta|[1611]|JWDCJ664-11|BOLD:AAL7529
Alliopsis fractiseta|[1612]|JWDCE354-10|BOLD:AAL7529
Alliopsis fractiseta|[1613]|JWDCJ744-11|BOLD:AAL7529
Alliopsis fractiseta|[1614]|JWDCJ665-11|BOLD:AAL7529
Alliopsis fractiseta|[1615]|JWDCJ919-11|BOLD:AAL7529
Alliopsis fractiseta|[1616]|JWDCI641-10|BOLD:AAL7529
Alliopsis fractiseta|[1617]|JWDCI909-11|BOLD:AAL7529
Alliopsis fractiseta|[1618]|JWDCE303-10|BOLD:AAL7529
Alliopsis fractiseta|[1619]|JWDCE602-10|BOLD:AAL7529
Alliopsis fractiseta|[1620]|JWDCE604-10|BOLD:AAL7529
Alliopsis fractiseta|[1621]|JWDCE046-10|BOLD:AAL7529
Anthomyiidae|[1622]|JWDCI649-10|BOLD:AAG2435
Anthomyiidae|[1623]|JWDCG873-10|BOLD:AAG2435
Anthomyiidae|[1624]|JWDCL089-11|BOLD:AAG2435
Anthomyiidae|[1625]|JWDCJ1502-11|BOLD:AAG2435
Anthomyiidae|[1626]|JWDCJ181-11|BOLD:AAG2435
Anthomyiidae|[1627]|JWDCI248-10|BOLD:AAG2435
Anthomyiidae|[1628]|JWDCI802-11|BOLD:AAG2435
Anthomyiidae|[1629]|JWDCJ857-11|BOLD:AAG2435
Anthomyiidae|[1630]|JWDCI805-11|BOLD:AAG2435
Anthomyiidae|[1631]|JWDCH162-10|BOLD:AAG2435
Anthomyiidae|[1632]|MHFLY021-06|BOLD:AAG2435
Alliopsis constrictor|[1633]|JWDCC022-10|BOLD:AAN5493
Alliopsis constrictor|[1634]|JWDCE373-10|BOLD:AAN5493
Alliopsis constrictor|[1635]|JWDCE374-10|BOLD:AAN5493
Alliopsis constrictor|[1636]|JWDCE404-10|BOLD:AAN5493
Botanophila|[1637]|JWDCC037-10|BOLD:AAG2502
Botanophila|[1638]|JWDCC073-10|BOLD:AAG2502
Botanophila|[1639]|JWDCI401-10|BOLD:AAG2502
Botanophila|[1640]|JWDCC145-10|BOLD:AAG2502
Botanophila|[1641]|JWDCC043-10|BOLD:AAG2502
Botanophila|[1642]|JWDCC166-10|BOLD:AAG2502
Botanophila|[1643]|JWDCC165-10|BOLD:AAG2502
Botanophila|[1644]|JWDCC116-10|BOLD:AAG2502
Botanophila|[1645]|JWDCC036-10|BOLD:AAG2502
Botanophila|[1646]|JWDCC035-10|BOLD:AAG2502
Botanophila|[1647]|JWDCC170-10|BOLD:AAG2502
Botanophila|[1648]|JWDCC167-10|BOLD:AAG2502
Botanophila|[1649]|JWDCC211-10|BOLD:AAG2502
Botanophila|[1650]|JWDCE934-10|BOLD:AAG2502
Alliopsis silvestris|[1651]|JWDCC046-10|BOLD:AAG2500
Alliopsis silvestris|[1652]|JWDCF130-10|BOLD:AAG2500
Alliopsis silvestris|[1653]|JWDCE939-10|BOLD:AAG2500
Alliopsis silvestris|[1654]|JWDCE730-10|BOLD:AAG2500
Anthomyiidae|[1655]|JWDCE219-10|BOLD:AAP6806
Anthomyiidae|[1656]|JWDCJ509-11|BOLD:AAP6806
Anthomyiidae|[1657]|JWDCJ508-11|BOLD:AAP6806
Anthomyiidae|[1658]|JWDCJ514-11|BOLD:AAP6806
Anthomyiidae|[1659]|JWDCE235-10|BOLD:AAP6806
Anthomyiidae|[1660]|JWDCF449-10|BOLD:AAG6758
Anthomyiidae|[1661]|MHFLI935-09|BOLD:AAG6758
Anthomyiidae|[1662]|MHFLI205-06|BOLD:AAG6758
Anthomyiidae|[1663]|MHFLI276-06|BOLD:AAG6758
Anthomyiidae|[1664]|JWDCF460-10|BOLD:AAG6758
Anthomyiidae|[1665]|JWDCJ578-11|BOLD:AAL7519
Anthomyiidae|[1666]|JWDCJ574-11|BOLD:AAL7519
Anthomyiidae|[1667]|JWDCC276-10|BOLD:AAL7519
Anthomyiidae|[1668]|JWDCF609-10|BOLD:AAL7519
Anthomyiidae|[1669]|JWDCC384-10|BOLD:AAL7519
Anthomyiidae|[1670]|JWDCJ518-11|BOLD:AAL7519
Anthomyiidae|[1671]|JWDCJ573-11|BOLD:AAL7519
Anthomyiidae|[1672]|JWDCJ628-11|BOLD:AAL7519
Anthomyiidae|[1673]|JWDCE600-10|BOLD:AAL7519
Anthomyiidae|[1674]|JWDCE601-10|BOLD:AAL7519
Anthomyiidae|[1675]|JWDCE621-10|BOLD:AAL7519
Anthomyiidae|[1676]|JWDCJ1071-11|BOLD:AAV4969
Anthomyia mimetica|[1677]|JWDCC853-10|BOLD:AAL7527
Anthomyia mimetica|[1678]|JWDCC092-10|BOLD:AAL7527
Anthomyia|[1679]|JWDCC093-10|BOLD:ABY5744
Egle subarctica|[1680]|JWDCA360-10|BOLD:AAM6380
Egle subarctica|[1681]|JWDCA428-10|BOLD:AAM6380
Egle subarctica|[1682]|JWDCJ112-11|BOLD:AAM6380
Egle|[1683]|JWDCC407-10|BOLD:AAL7521
Egle|[1684]|JWDCC408-10|BOLD:AAL7521
Egle|[1685]|JWDCF456-10|BOLD:AAL7521
Egle atomaria|[1686]|JWDCA593-10|BOLD:AAI6225
Egle atomaria|[1687]|JWDCA592-10|BOLD:AAI6225
Egle atomaria|[1688]|JWDCC400-10|BOLD:AAI6225
Egle atomaria|[1689]|JWDCA422-10|BOLD:AAI6225
Egle atomaria|[1690]|JWDCA580-10|BOLD:AAI6225
Egle atomaria|[1691]|JWDCA558-10|BOLD:AAI6225
Egle atomaria|[1692]|JWDCA595-10|BOLD:AAI6225

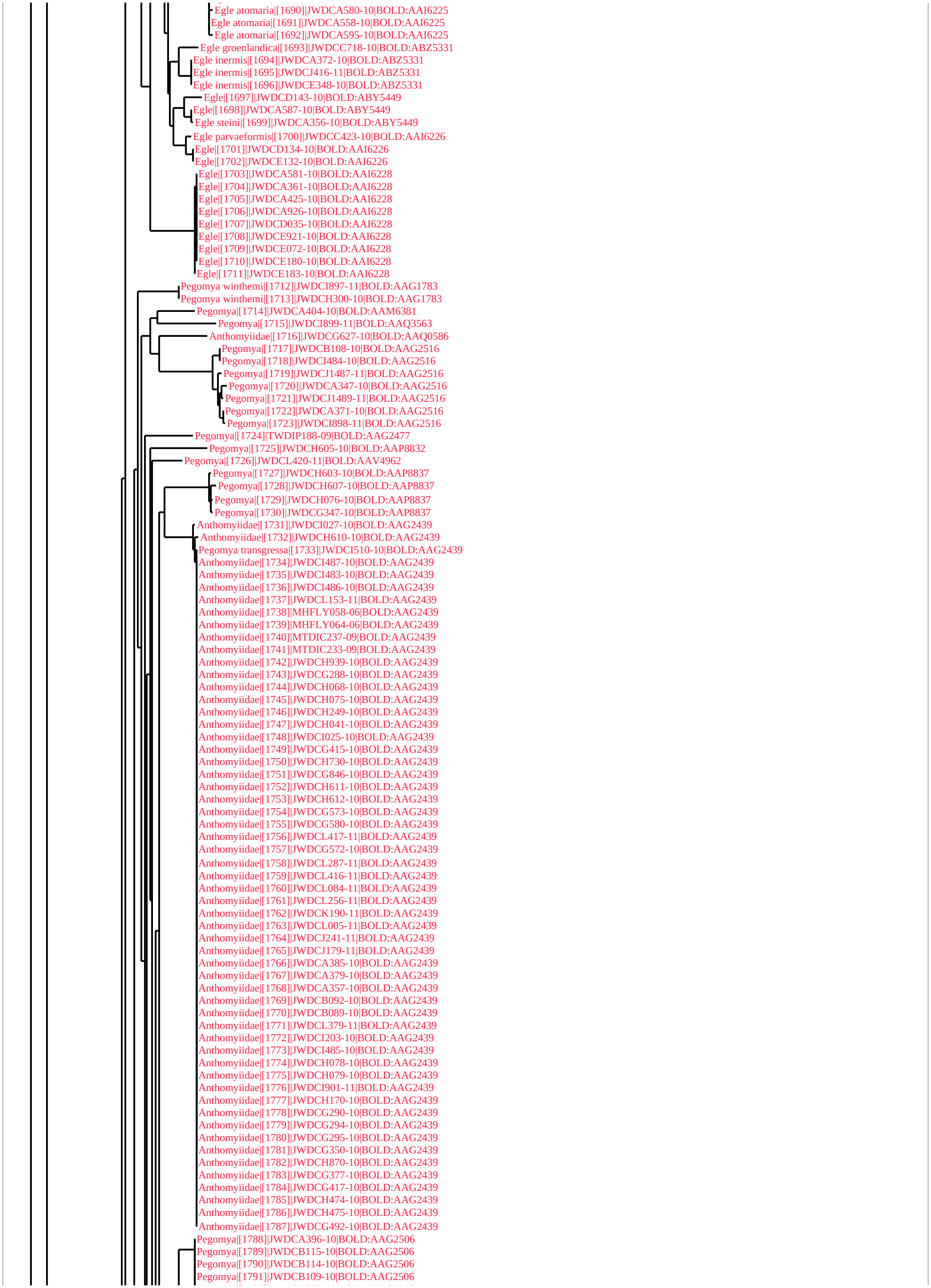
Egle atomaria|[1690]|JWDCA580-10|BOLD:AAI6225
Egle atomaria|[1691]|JWDCA558-10|BOLD:AAI6225
Egle atomaria|[1692]|JWDCA595-10|BOLD:AAI6225
Egle groenlandica|[1693]|JWDCC718-10|BOLD:ABZ5331
Egle inermis|[1694]|JWDCA372-10|BOLD:ABZ5331
Egle inermis|[1695]|JWDCJ416-11|BOLD:ABZ5331
Egle inermis|[1696]|JWDCE348-10|BOLD:ABZ5331
Egle|[1697]|JWDCD143-10|BOLD:ABY5449
Egle|[1698]|JWDCA587-10|BOLD:ABY5449
Egle steini|[1699]|JWDCA356-10|BOLD:ABY5449
Egle parvaeformis|[1700]|JWDCC423-10|BOLD:AAI6226
Egle|[1701]|JWDCD134-10|BOLD:AAI6226
Egle|[1702]|JWDCE132-10|BOLD:AAI6226
Egle|[1703]|JWDCA581-10|BOLD:AAI6228
Egle|[1704]|JWDCA361-10|BOLD:AAI6228
Egle|[1705]|JWDCA425-10|BOLD:AAI6228
Egle|[1706]|JWDCA926-10|BOLD:AAI6228
Egle|[1707]|JWDCD035-10|BOLD:AAI6228
Egle|[1708]|JWDCE921-10|BOLD:AAI6228
Egle|[1709]|JWDCE072-10|BOLD:AAI6228
Egle|[1710]|JWDCE180-10|BOLD:AAI6228
Egle|[1711]|JWDCE183-10|BOLD:AAI6228
Pegomya winthemi|[1712]|JWDCI897-11|BOLD:AAG1783
Pegomya winthemi|[1713]|JWDCH300-10|BOLD:AAG1783
Pegomya|[1714]|JWDCA404-10|BOLD:AAM6381
Pegomya|[1715]|JWDCI899-11|BOLD:AAQ3563
Anthomyiidae|[1716]|JWDCG627-10|BOLD:AAQ0586
Pegomya|[1717]|JWDCB108-10|BOLD:AAG2516
Pegomya|[1718]|JWDCI484-10|BOLD:AAG2516
Pegomya|[1719]|JWDCJ1487-11|BOLD:AAG2516
Pegomya|[1720]|JWDCA347-10|BOLD:AAG2516
Pegomya|[1721]|JWDCJ1489-11|BOLD:AAG2516
Pegomya|[1722]|JWDCA371-10|BOLD:AAG2516
Pegomya|[1723]|JWDCI898-11|BOLD:AAG2516
Pegomya|[1724]|TWDIP188-09|BOLD:AAG2477
Pegomya|[1725]|JWDCH605-10|BOLD:AAP8832
Pegomya|[1726]|JWDCL420-11|BOLD:AAV4962
Pegomya|[1727]|JWDCH603-10|BOLD:AAP8837
Pegomya|[1728]|JWDCH607-10|BOLD:AAP8837
Pegomya|[1729]|JWDCH076-10|BOLD:AAP8837
Pegomya|[1730]|JWDCG347-10|BOLD:AAP8837
Anthomyiidae|[1731]|JWDCI027-10|BOLD:AAG2439
Anthomyiidae|[1732]|JWDCH610-10|BOLD:AAG2439
Pegomya transgressa|[1733]|JWDCI510-10|BOLD:AAG2439
Anthomyiidae|[1734]|JWDCI487-10|BOLD:AAG2439
Anthomyiidae|[1735]|JWDCI483-10|BOLD:AAG2439
Anthomyiidae|[1736]|JWDCI486-10|BOLD:AAG2439
Anthomyiidae|[1737]|JWDCL153-11|BOLD:AAG2439
Anthomyiidae|[1738]|MHFLY058-06|BOLD:AAG2439
Anthomyiidae|[1739]|MHFLY064-06|BOLD:AAG2439
Anthomyiidae|[1740]|MTDIC237-09|BOLD:AAG2439
Anthomyiidae|[1741]|MTDIC233-09|BOLD:AAG2439
Anthomyiidae|[1742]|JWDCH939-10|BOLD:AAG2439
Anthomyiidae|[1743]|JWDCG288-10|BOLD:AAG2439
Anthomyiidae|[1744]|JWDCH068-10|BOLD:AAG2439
Anthomyiidae|[1745]|JWDCH075-10|BOLD:AAG2439
Anthomyiidae|[1746]|JWDCH249-10|BOLD:AAG2439
Anthomyiidae|[1747]|JWDCH041-10|BOLD:AAG2439
Anthomyiidae|[1748]|JWDCI025-10|BOLD:AAG2439
Anthomyiidae|[1749]|JWDCG415-10|BOLD:AAG2439
Anthomyiidae|[1750]|JWDCH730-10|BOLD:AAG2439
Anthomyiidae|[1751]|JWDCG846-10|BOLD:AAG2439
Anthomyiidae|[1752]|JWDCH611-10|BOLD:AAG2439
Anthomyiidae|[1753]|JWDCH612-10|BOLD:AAG2439
Anthomyiidae|[1754]|JWDCG573-10|BOLD:AAG2439
Anthomyiidae|[1755]|JWDCG580-10|BOLD:AAG2439
Anthomyiidae|[1756]|JWDCL417-11|BOLD:AAG2439
Anthomyiidae|[1757]|JWDCG572-10|BOLD:AAG2439
Anthomyiidae|[1758]|JWDCL287-11|BOLD:AAG2439
Anthomyiidae|[1759]|JWDCL416-11|BOLD:AAG2439
Anthomyiidae|[1760]|JWDCL084-11|BOLD:AAG2439
Anthomyiidae|[1761]|JWDCL256-11|BOLD:AAG2439
Anthomyiidae|[1762]|JWDCK190-11|BOLD:AAG2439
Anthomyiidae|[1763]|JWDCL005-11|BOLD:AAG2439
Anthomyiidae|[1764]|JWDCJ241-11|BOLD:AAG2439
Anthomyiidae|[1765]|JWDCJ179-11|BOLD:AAG2439
Anthomyiidae|[1766]|JWDCA385-10|BOLD:AAG2439
Anthomyiidae|[1767]|JWDCA379-10|BOLD:AAG2439
Anthomyiidae|[1768]|JWDCA357-10|BOLD:AAG2439
Anthomyiidae|[1769]|JWDCB092-10|BOLD:AAG2439
Anthomyiidae|[1770]|JWDCB089-10|BOLD:AAG2439
Anthomyiidae|[1771]|JWDCL379-11|BOLD:AAG2439
Anthomyiidae|[1772]|JWDCI203-10|BOLD:AAG2439
Anthomyiidae|[1773]|JWDCI485-10|BOLD:AAG2439
Anthomyiidae|[1774]|JWDCH078-10|BOLD:AAG2439
Anthomyiidae|[1775]|JWDCH079-10|BOLD:AAG2439
Anthomyiidae|[1776]|JWDCI901-11|BOLD:AAG2439
Anthomyiidae|[1777]|JWDCH170-10|BOLD:AAG2439
Anthomyiidae|[1778]|JWDCG290-10|BOLD:AAG2439
Anthomyiidae|[1779]|JWDCG294-10|BOLD:AAG2439
Anthomyiidae|[1780]|JWDCG295-10|BOLD:AAG2439
Anthomyiidae|[1781]|JWDCG350-10|BOLD:AAG2439
Anthomyiidae|[1782]|JWDCH870-10|BOLD:AAG2439
Anthomyiidae|[1783]|JWDCG377-10|BOLD:AAG2439
Anthomyiidae|[1784]|JWDCG417-10|BOLD:AAG2439
Anthomyiidae|[1785]|JWDCH474-10|BOLD:AAG2439
Anthomyiidae|[1786]|JWDCH475-10|BOLD:AAG2439
Anthomyiidae|[1787]|JWDCG492-10|BOLD:AAG2439
Pegomya|[1788]|JWDCA396-10|BOLD:AAG2506
Pegomya|[1789]|JWDCB115-10|BOLD:AAG2506
Pegomya|[1790]|JWDCB114-10|BOLD:AAG2506
Pegomya|[1791]|JWDCB109-10|BOLD:AAG2506

Pegomya|[1789]|JWDCB115-10|BOLD:AAG2506
Pegomya|[1790]|JWDCB114-10|BOLD:AAG2506
Pegomya|[1791]|JWDCB109-10|BOLD:AAG2506
Pegomya|[1792]|JWDCH868-10|BOLD:AAG2506
Pegomya|[1793]|JWDCH309-10|BOLD:AAG2506
Pegomya|[1794]|JWDCJ960-11|BOLD:AAG2506
Pegomya|[1795]|TWDIP182-09|BOLD:AAG2506
Pegomya|[1796]|JWDCH312-10|BOLD:AAG2506
Pegomya|[1797]|JWDCH311-10|BOLD:AAG2506
Pegomya|[1798]|JWDCH729-10|BOLD:AAG2506
Pegomya|[1799]|JWDCI895-11|BOLD:AAG2506
Pegomya|[1800]|JWDCH398-10|BOLD:AAG2506
Pegomya|[1801]|JWDCI652-10|BOLD:AAG2506
Pegomya|[1802]|JWDCI428-10|BOLD:AAG2506
Pegomya|[1803]|JWDCL328-11|BOLD:AAG2506
Pegomya|[1804]|JWDCG833-10|BOLD:AAG2506
Pegomya|[1805]|JWDCL254-11|BOLD:AAG2506
Pegomya|[1806]|JWDCL259-11|BOLD:AAG2506
Pegomya|[1807]|JWDCL082-11|BOLD:AAG2506
Pegomya|[1808]|JWDCL002-11|BOLD:AAG2506
Pegomya|[1809]|JWDCJ1282-11|BOLD:AAG2506
Pegomya|[1810]|JWDCJ1274-11|BOLD:AAG2506
Pegomya|[1811]|JWDCK153-11|BOLD:AAG2506
Pegomya|[1812]|JWDCJ304-11|BOLD:AAG2506
Pegomya|[1813]|JWDCH414-10|BOLD:AAG2506
Pegomya|[1814]|JWDCH169-10|BOLD:AAG2506
Pegomya|[1815]|JWDCL257-11|BOLD:AAG2506
Pegomya|[1816]|JWDCH071-10|BOLD:AAG2506
Pegomya|[1817]|JWDCH602-10|BOLD:AAG2506
Pegomya|[1818]|JWDCL263-11|BOLD:AAG2506
Pegomya|[1819]|JWDCL260-11|BOLD:AAG2506
Pegomya|[1820]|JWDCJ856-11|BOLD:AAG2506
Pegomya|[1821]|JWDCH172-10|BOLD:AAG2506
Pegomya|[1822]|JWDCG493-10|BOLD:AAG2506
Pegomya|[1823]|JWDCG494-10|BOLD:AAG2506
Pegomya|[1824]|JWDCG496-10|BOLD:AAG2506
Pegomya|[1825]|JWDCG558-10|BOLD:AAG2506
Anthomyiidae|[1826]|JWDCJ1273-11|BOLD:AAP8839
Anthomyiidae|[1827]|JWDCH613-10|BOLD:AAP8839
Pegomya|[1828]|TWDIP183-09|BOLD:AAG1782
Pegomya|[1829]|TWDIP185-09|BOLD:AAG1782
Pegomya|[1830]|JWDCG845-10|BOLD:AAG1782
Pegomya|[1831]|JWDCA327-10|BOLD:AAG1782
Pegomya flavoscutellata|[1832]|TWDIP093-09|BOLD:AAG1782
Pegomya|[1833]|JWDCL270-11|BOLD:AAG1782
Pegomya|[1834]|JWDCL326-11|BOLD:AAG1782
Pegomya|[1835]|JWDCA330-10|BOLD:AAG1782
Pegomya|[1836]|JWDCA340-10|BOLD:AAG1782
Pegomya|[1837]|JWDCJ108-11|BOLD:AAG1782
Pegomya|[1838]|JWDCJ535-11|BOLD:AAG1782
Pegomya|[1839]|JWDCL252-11|BOLD:AAG1782
Pegomya|[1840]|JWDCL253-11|BOLD:AAG1782
Pegomya|[1841]|JWDCL255-11|BOLD:AAG1782
Pegomya|[1842]|JWDCL258-11|BOLD:AAG1782
Pegomya|[1843]|JWDCL329-11|BOLD:AAG1782
Pegomya|[1844]|JWDCL412-11|BOLD:AAG1782
Pegomya|[1845]|JWDCL413-11|BOLD:AAG1782
Pegomya|[1846]|JWDCL438-11|BOLD:AAG1782
Pegomya|[1847]|JWDCI213-10|BOLD:AAG1782
Pegomya|[1848]|JWDCI919-11|BOLD:AAG1782
Pegomya|[1849]|JWDCG847-10|BOLD:AAG1782
Pegomya|[1850]|JWDCG293-10|BOLD:AAG1782
Pegomya|[1851]|JWDCB094-10|BOLD:AAG1782
Pegomya|[1852]|JWDCB112-10|BOLD:AAG1782
Pegomya|[1853]|JWDCF459-10|BOLD:AAG1782
Pegomya|[1854]|JWDCA326-10|BOLD:AAG1782
Pegomya|[1855]|JWDCB095-10|BOLD:AAG1782
Pegomya|[1856]|JWDCB091-10|BOLD:AAG1782
Pegomya|[1857]|TWDIP096-09|BOLD:AAG1782
Pegomya|[1858]|JWDCC801-10|BOLD:AAG1782
Pegomya|[1859]|JWDCA336-10|BOLD:AAG1782
Pegomya|[1860]|JWDCA338-10|BOLD:AAG1782
Pegomya|[1861]|JWDCA349-10|BOLD:AAG1782
Pegomya|[1862]|JWDCA380-10|BOLD:AAG1782
Pegomya|[1863]|JWDCJ103-11|BOLD:AAG1782
Pegomya|[1864]|JWDCJ105-11|BOLD:AAG1782
Pegomya|[1865]|JWDCJ116-11|BOLD:AAG1782
Pegomya|[1866]|JWDCJ860-11|BOLD:AAG1782
Pegomya|[1867]|JWDCL418-11|BOLD:AAG1782
Pegomya|[1868]|JWDCI210-10|BOLD:AAG1782
Pegomya|[1869]|JWDCH171-10|BOLD:AAG1782
Pegomya|[1870]|JWDCI902-11|BOLD:AAG1782
Pegomya|[1871]|JWDCE731-10|BOLD:AAG1782
Pegomya geniculata|[1872]|MTDIC174-09|BOLD:AAG2466
Pegomya flavifrons|[1873]|MHFLY054-06|BOLD:AAG6754
Anthomyiidae|[1874]|JWDCA352-10|BOLD:AAL8103
Anthomyiidae|[1875]|JWDCJ1063-11|BOLD:AAL8103
Anthomyiidae|[1876]|JWDCL325-11|BOLD:AAL8103
Eutrichota|[1877]|MHFLY063-06|BOLD:AAG2440
Eutrichota|[1878]|MHFLI933-09|BOLD:AAG2440
Eutrichota tunicata|[1879]|JWDCJ1005-11|BOLD:AAG2440
Eutrichota|[1880]|MTDIC222-09|BOLD:AAG2440
Eutrichota tunicata|[1881]|JWDCA343-10|BOLD:AAG2440
Eutrichota tunicata|[1882]|JWDCJ106-11|BOLD:AAG2440
Eutrichota tunicata|[1883]|JWDCK189-11|BOLD:AAG2440
Eutrichota|[1884]|MTDIC196-09|BOLD:AAG2440
Eutrichota tunicata|[1885]|JWDCL004-11|BOLD:AAG2440
Eutrichota|[1886]|MHFLI921-09|BOLD:AAG2440
Eutrichota|[1887]|MHFLY392-09|BOLD:AAG2440
Eutrichota|[1888]|MTDIC225-09|BOLD:AAG2440
Eutrichota tunicata|[1889]|JWDCK149-11|BOLD:AAG2440
Eutrichota tunicata|[1890]|JWDCJ1607-11|BOLD:AAG2440
Eutrichota tunicata|[1891]|JWDCJ1421-11|BOLD:AAG2440

Eutrichota tunicata|[1889]|JWDCK149-11|BOLD:AAG2440
Eutrichota tunicata|[1890]|JWDCJ1607-11|BOLD:AAG2440
Eutrichota tunicata|[1891]|JWDCJ1421-11|BOLD:AAG2440
Eutrichota tunicata|[1892]|JWDCI893-11|BOLD:AAG2440
Eutrichota tunicata|[1893]|JWDCB088-10|BOLD:AAG2440
Eutrichota tunicata|[1894]|JWDCK151-11|BOLD:AAG2440
Eutrichota tunicata|[1895]|JWDCI896-11|BOLD:AAG2440
Eutrichota tunicata|[1896]|JWDCC120-10|BOLD:AAG2440
Eutrichota tunicata|[1897]|JWDCC117-10|BOLD:AAG2440
Eutrichota|[1898]|MHFLI924-09|BOLD:AAG2440
Eutrichota tunicata|[1899]|JWDCJ1062-11|BOLD:AAG2440
Eutrichota tunicata|[1900]|JWDCA363-10|BOLD:AAG2440
Eutrichota tunicata|[1901]|JWDCC192-10|BOLD:AAG2440
Eutrichota tunicata|[1902]|JWDCA344-10|BOLD:AAG2440
Eutrichota tunicata|[1903]|JWDCJ1424-11|BOLD:AAG2440
Eutrichota tunicata|[1904]|JWDCL261-11|BOLD:AAG2440
Eutrichota tunicata|[1905]|JWDCL327-11|BOLD:AAG2440
Eutrichota tunicata|[1906]|JWDCI488-10|BOLD:AAG2440
Eutrichota tunicata|[1907]|JWDCI900-11|BOLD:AAG2440
Eutrichota tunicata|[1908]|JWDCH310-10|BOLD:AAG2440
Eutrichota tunicata|[1909]|JWDCG351-10|BOLD:AAG2440
Eutrichota|[1910]|JWDCC198-10|BOLD:AAL7517
Anthomyiidae|[1911]|JWDCE033-10|BOLD:AAL7517
Anthomyiidae|[1912]|JWDCC403-10|BOLD:AAL7517
Anthomyiidae|[1913]|JWDCE233-10|BOLD:AAL7517
Delia bucculenta|[1914]|JWDCE540-10|BOLD:AAP3547
Delia|[1915]|JWDCH524-10|BOLD:AAP8838
Delia longicauda|[1916]|JWDCC405-10|BOLD:AAG2475
Delia longicauda|[1917]|JWDCI247-10|BOLD:AAG2475
Delia longicauda|[1918]|TWDIP094-09|BOLD:AAG2475
Delia longicauda|[1919]|JWDCA355-10|BOLD:AAG2475
Delia longicauda|[1920]|JWDCE294-10|BOLD:AAG2475
Delia|[1921]|JWDCE353-10|BOLD:AAI5778
Delia platura|[1922]|MHFLI128-06|BOLD:AAA3453
Delia platura|[1923]|JSFLA032-08|BOLD:AAA3453
Delia platura|[1924]|JWDCE064-10|BOLD:AAA3453
Delia tarsata|[1925]|TWDIP097-09|BOLD:AAG2476
Delia|[1926]|MHFLI940-09|BOLD:AAG2471
Delia lineariventris|[1927]|TWDIP189-09|BOLD:AAG2478
Delia|[1928]|JWDCC045-10|BOLD:AAL9120
Delia|[1929]|JWDCC042-10|BOLD:AAL9120
Delia|[1930]|JWDCC213-10|BOLD:AAL9120
Delia albula|[1931]|JWDCG682-10|BOLD:AAP8834
Delia albula|[1932]|JWDCH879-10|BOLD:AAP8834
Delia albula|[1933]|JWDCH869-10|BOLD:AAP8834
Delia albula|[1934]|JWDCG425-10|BOLD:AAP8834
Delia albula|[1935]|JWDCF334-10|BOLD:AAP8834
Delia|[1936]|JWDCJ1420-11|BOLD:AAG2472
Delia|[1937]|JWDCH727-10|BOLD:AAG2472
Delia|[1938]|JWDCE538-10|BOLD:AAP3546
Delia|[1939]|JWDCE347-10|BOLD:AAP3546
Delia|[1940]|JWDCE777-10|BOLD:AAP3546
Delia|[1941]|JWDCL563-11|BOLD:AAQ3567
Delia planipalpis|[1942]|JWDCL542-11|BOLD:AAQ3565
Delia linearis|[1943]|JWDCC115-10|BOLD:AAL7516
Delia linearis|[1944]|JWDCA351-10|BOLD:AAL7516
Delia setigera|[1945]|MHFLY055-06|BOLD:AAG2438
Delia setigera|[1946]|JWDCA350-10|BOLD:AAG2438
Delia|[1947]|JWDCE252-10|BOLD:AAP6807
Anthomyiidae|[1948]|JWDCC837-10|BOLD:AAL7526
Botanophila varians|[1949]|TWDIP181-09|BOLD:AAP6568
Botanophila profuga|[1950]|JWDCA401-10|BOLD:ACP6334
Botanophila profuga|[1951]|JWDCA382-10|BOLD:ACP6334
Botanophila profuga|[1952]|JWDCA377-10|BOLD:ACP6334
Botanophila profuga|[1953]|JWDCA373-10|BOLD:ACP6334
Botanophila profuga|[1954]|JWDCB102-10|BOLD:ACP6334
Botanophila profuga|[1955]|JWDCA392-10|BOLD:ACP6334
Botanophila profuga|[1956]|JWDCA394-10|BOLD:ACP6334
Botanophila profuga|[1957]|JWDCA395-10|BOLD:ACP6334
Botanophila profuga|[1958]|JWDCK188-11|BOLD:ACP6334
Botanophila profuga|[1959]|JWDCK252-11|BOLD:ACP6334
Botanophila profuga|[1960]|JWDCJ1422-11|BOLD:ACP6334
Botanophila profuga|[1961]|JWDCJ1423-11|BOLD:ACP6334
Botanophila profuga|[1962]|JWDCJ1486-11|BOLD:ACP6334
Botanophila profuga|[1963]|JWDCG763-10|BOLD:ACP6334
Botanophila profuga|[1964]|JWDCH073-10|BOLD:ACP6334
Botanophila profuga|[1965]|JWDCG416-10|BOLD:ACP6334
Botanophila profuga|[1966]|JWDCI109-10|BOLD:ACP6334
Botanophila profuga|[1967]|JWDCI160-10|BOLD:ACP6334
Botanophila profuga|[1968]|MHFLY025-06|BOLD:ACP6334
Botanophila profuga|[1969]|JWDCL008-11|BOLD:ACP6334
Botanophila profuga|[1970]|MHFLI922-09|BOLD:ACP6334
Botanophila profuga|[1971]|JWDCA323-10|BOLD:ACP6334
Botanophila profuga|[1972]|JWDCB100-10|BOLD:ACP6334
Botanophila profuga|[1973]|JWDCA400-10|BOLD:ACP6334
Botanophila profuga|[1974]|JWDCA384-10|BOLD:ACP6334
Botanophila profuga|[1975]|JWDCA393-10|BOLD:ACP6334
Botanophila profuga|[1976]|JWDCJ1275-11|BOLD:ACP6334
Botanophila profuga|[1977]|JWDCH604-10|BOLD:ACP6334
Botanophila profuga|[1978]|JWDCH617-10|BOLD:ACP6334
Botanophila profuga|[1979]|JWDCH726-10|BOLD:ACP6334
Botanophila profuga|[1980]|JWDCG844-10|BOLD:ACP6334
Botanophila profuga|[1981]|JWDCI651-10|BOLD:ACP6334
Botanophila profuga|[1982]|JWDCH248-10|BOLD:ACP6334
Botanophila profuga|[1983]|JWDCG296-10|BOLD:ACP6334
Botanophila profuga|[1984]|JWDCG346-10|BOLD:ACP6334
Botanophila profuga|[1985]|JWDCG422-10|BOLD:ACP6334
Botanophila profuga|[1986]|JWDCG482-10|BOLD:ACP6334
Boreophorbia hirtipes|[1987]|JWDCJ1345-11|BOLD:AAG2448
Boreophorbia hirtipes|[1988]|JWDCJ1491-11|BOLD:AAG2448
Boreophorbia hirtipes|[1989]|JWDCH770-10|BOLD:AAG2448
Boreophorbia hirtipes|[1990]|JWDCF122-10|BOLD:AAG2448

Boreophorbia hirtipes|[1988]|JWDCJ1491-11|BOLD:AAG2448
Boreophorbia hirtipes|[1989]|JWDCH770-10|BOLD:AAG2448
Boreophorbia hirtipes|[1990]|JWDCF122-10|BOLD:AAG2448
Boreophorbia hirtipes|[1991]|JWDCE452-10|BOLD:AAG2448
Boreophorbia hirtipes|[1992]|JWDCI057-10|BOLD:AAG2448
Boreophorbia hirtipes|[1993]|JWDCJ789-11|BOLD:AAG2448
Boreophorbia hirtipes|[1994]|JWDCJ790-11|BOLD:AAG2448
Boreophorbia hirtipes|[1995]|JWDCJ1546-11|BOLD:AAG2448
Boreophorbia hirtipes|[1996]|JWDCJ1565-11|BOLD:AAG2448
Boreophorbia hirtipes|[1997]|JWDCJ397-11|BOLD:AAG2448
Boreophorbia hirtipes|[1998]|JWDCJ183-11|BOLD:AAG2448
Boreophorbia hirtipes|[1999]|JWDCI141-10|BOLD:AAG2448
Boreophorbia hirtipes|[2000]|JWDCF303-10|BOLD:AAG2448
Boreophorbia hirtipes|[2001]|MHFLI392-07|BOLD:AAG2448
Fucellia tergina|[2002]|JWDCH985-10|BOLD:AAG2497
Fucellia ariciiformis|[2003]|JWDCA390-10|BOLD:AAG2437
Fucellia ariciiformis|[2004]|MHFLY133-07|BOLD:AAG2437
Fucellia ariciiformis|[2005]|MHFLY123-07|BOLD:AAG2437
Fucellia ariciiformis|[2006]|JWDCK257-11|BOLD:AAG2437
Fucellia ariciiformis|[2007]|MHFLY135-07|BOLD:AAG2437
Fucellia ariciiformis|[2008]|JWDCI110-10|BOLD:AAG2437
Fucellia ariciiformis|[2009]|JWDCA388-10|BOLD:AAG2437
Fucellia ariciiformis|[2010]|JWDCA378-10|BOLD:AAG2437
Fucellia ariciiformis|[2011]|JWDCL606-11|BOLD:AAG2437
Fucellia ariciiformis|[2012]|JWDCI512-10|BOLD:AAG2437
Fucellia ariciiformis|[2013]|JWDCI515-10|BOLD:AAG2437
Fucellia ariciiformis|[2014]|MHFLY137-07|BOLD:AAG2437
Fucellia ariciiformis|[2015]|MHFLY028-06|BOLD:AAG2437
Fucellia ariciiformis|[2016]|MHFLI165-06|BOLD:AAG2437
Fucellia ariciiformis|[2017]|MHFLI132-06|BOLD:AAG2437
Fucellia ariciiformis|[2018]|MHFLI126-06|BOLD:AAG2437
Fucellia ariciiformis|[2019]|JWDCH808-10|BOLD:AAG2437
Fucellia ariciiformis|[2020]|JWDCI514-10|BOLD:AAG2437
Fucellia ariciiformis|[2021]|JWDCH806-10|BOLD:AAG2437
Fucellia ariciiformis|[2022]|JWDCH807-10|BOLD:AAG2437
Fucellia ariciiformis|[2023]|JWDCG683-10|BOLD:AAG2437
Fucellia ariciiformis|[2024]|JWDCH805-10|BOLD:AAG2437
Fucellia ariciiformis|[2025]|JWDCL433-11|BOLD:AAG2437
Fucellia ariciiformis|[2026]|JWDCG678-10|BOLD:AAG2437
Fucellia ariciiformis|[2027]|JWDCL087-11|BOLD:AAG2437
Fucellia ariciiformis|[2028]|JWDCL605-11|BOLD:AAG2437
Fucellia ariciiformis|[2029]|JWDCL083-11|BOLD:AAG2437
Fucellia ariciiformis|[2030]|JWDCL086-11|BOLD:AAG2437
Fucellia ariciiformis|[2031]|JWDCL562-11|BOLD:AAG2437
Fucellia ariciiformis|[2032]|JWDCL154-11|BOLD:AAG2437
Fucellia ariciiformis|[2033]|JWDCL559-11|BOLD:AAG2437
Fucellia ariciiformis|[2034]|JWDCL561-11|BOLD:AAG2437
Fucellia ariciiformis|[2035]|JWDCA402-10|BOLD:AAG2437
Fucellia ariciiformis|[2036]|JWDCA604-10|BOLD:AAG2437
Fucellia ariciiformis|[2037]|JWDCA386-10|BOLD:AAG2437
Fucellia ariciiformis|[2038]|JWDCA389-10|BOLD:AAG2437
Fucellia ariciiformis|[2039]|JWDCA376-10|BOLD:AAG2437
Fucellia ariciiformis|[2040]|JWDCA374-10|BOLD:AAG2437
Fucellia ariciiformis|[2041]|JWDCA375-10|BOLD:AAG2437
Fucellia ariciiformis|[2042]|MHFLI164-06|BOLD:AAG2437
Fucellia ariciiformis|[2043]|JWDCG334-10|BOLD:AAG2437
Fucellia ariciiformis|[2044]|JWDCG335-10|BOLD:AAG2437
Fucellia ariciiformis|[2045]|JWDCH029-10|BOLD:AAG2437
Fucellia ariciiformis|[2046]|JWDCH151-10|BOLD:AAG2437
Fucellia ariciiformis|[2047]|JWDCH411-10|BOLD:AAG2437
Fucellia ariciiformis|[2048]|JWDCH415-10|BOLD:AAG2437
Fucellia ariciiformis|[2049]|JWDCH525-10|BOLD:AAG2437
Fucellia ariciiformis|[2050]|JWDCG560-10|BOLD:AAG2437
Fucellia ariciiformis|[2051]|JWDCE375-10|BOLD:AAG2437
Fucellia ariciiformis|[2052]|JWDCE401-10|BOLD:AAG2437
Fucellia ariciiformis|[2053]|JWDCE402-10|BOLD:AAG2437
Fucellia ariciiformis|[2054]|JWDCF277-10|BOLD:AAG2437
Fucellia ariciiformis|[2055]|MHFLI891-09|BOLD:AAG2437
Fucellia ariciiformis|[2056]|MHFLI931-09|BOLD:AAG2437
Fucellia ariciiformis|[2057]|MHFLI932-09|BOLD:AAG2437
Fucellia ariciiformis|[2058]|MHFLY136-07|BOLD:AAG2437
Fucellia ariciiformis|[2059]|JWDCE044-10|BOLD:AAG2437
Fucellia ariciiformis|[2060]|JWDCE060-10|BOLD:AAG2437
Fucellia ariciiformis|[2061]|JWDCE063-10|BOLD:AAG2437
Fucellia ariciiformis|[2062]|JWDCE065-10|BOLD:AAG2437
Fucellia ariciiformis|[2063]|JWDCE066-10|BOLD:AAG2437
Fucellia ariciiformis|[2064]|JWDCE067-10|BOLD:AAG2437
Fucellia ariciiformis|[2065]|JWDCE068-10|BOLD:AAG2437
Fucellia ariciiformis|[2066]|JWDCE070-10|BOLD:AAG2437
Fucellia ariciiformis|[2067]|JWDCE130-10|BOLD:AAG2437
Fucellia ariciiformis|[2068]|JWDCE141-10|BOLD:AAG2437
Fucellia ariciiformis|[2069]|JWDCE166-10|BOLD:AAG2437
Fucellia ariciiformis|[2070]|JWDCE182-10|BOLD:AAG2437
Fucellia ariciiformis|[2071]|JWDCE251-10|BOLD:AAG2437
Fucellia ariciiformis|[2072]|JWDCE253-10|BOLD:AAG2437
Fucellia ariciiformis|[2073]|JWDCE254-10|BOLD:AAG2437
Fucellia ariciiformis|[2074]|JWDCE256-10|BOLD:AAG2437
Pegoplata nigroscutellata|[2075]|JWDCC044-10|BOLD:AAN5494
Pegoplata nigroscutellata|[2076]|JWDCI650-10|BOLD:AAN5494
Pegoplata tundrica|[2077]|JWDCH042-10|BOLD:AAG2449
Pegoplata tundrica|[2078]|MHFLI658-07|BOLD:AAG2449
Pegoplata tundrica|[2079]|JWDCH105-10|BOLD:AAG2449
Gymnocheta|[2080]|JWDCC162-10|BOLD:AAL7400
Gymnocheta|[2081]|JWDCC130-10|BOLD:AAL7400
Gymnocheta|[2082]|JWDCC160-10|BOLD:AAL7400
Gymnocheta|[2083]|JWDCC177-10|BOLD:AAL7400
Gymnocheta|[2084]|JWDCJ355-11|BOLD:AAL7400
Angioneura|[2085]|JWDCJ323-11|BOLD:AAQ0759
Angioneura|[2086]|JWDCL352-11|BOLD:AAQ0759
Angioneura|[2087]|JWDCL142-11|BOLD:AAQ0759
Angioneura|[2088]|JWDCL132-11|BOLD:AAQ0759
Angioneura|[2089]|JWDCK197-11|BOLD:AAQ0759
Angioneura|[2090]|JWDCK082-11|BOLD:AAQ0759

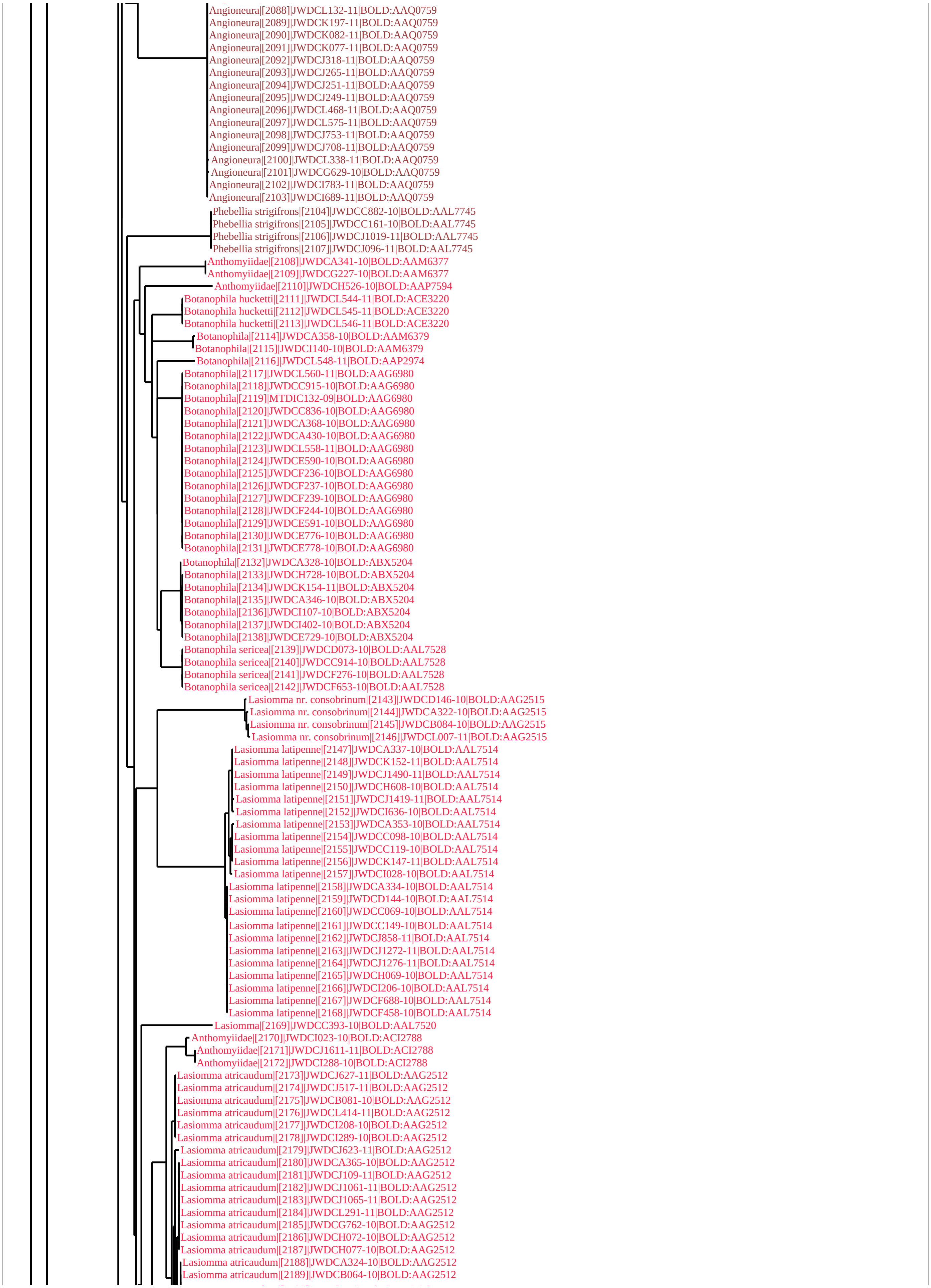
Angioneura|[2088]|JWDCL132-11|BOLD:AAQ0759
Angioneura|[2089]|JWDCK197-11|BOLD:AAQ0759
Angioneura|[2090]|JWDCK082-11|BOLD:AAQ0759
Angioneura|[2091]|JWDCK077-11|BOLD:AAQ0759
Angioneura|[2092]|JWDCJ318-11|BOLD:AAQ0759
Angioneura|[2093]|JWDCJ265-11|BOLD:AAQ0759
Angioneura|[2094]|JWDCJ251-11|BOLD:AAQ0759
Angioneura|[2095]|JWDCJ249-11|BOLD:AAQ0759
Angioneura|[2096]|JWDCL468-11|BOLD:AAQ0759
Angioneura|[2097]|JWDCL575-11|BOLD:AAQ0759
Angioneura|[2098]|JWDCJ753-11|BOLD:AAQ0759
Angioneura|[2099]|JWDCJ708-11|BOLD:AAQ0759
Angioneura|[2100]|JWDCL338-11|BOLD:AAQ0759
Angioneura|[2101]|JWDCG629-10|BOLD:AAQ0759
Angioneura|[2102]|JWDCI783-11|BOLD:AAQ0759
Angioneura|[2103]|JWDCI689-11|BOLD:AAQ0759
Phebellia strigifrons|[2104]|JWDCC882-10|BOLD:AAL7745
Phebellia strigifrons|[2105]|JWDCC161-10|BOLD:AAL7745
Phebellia strigifrons|[2106]|JWDCJ1019-11|BOLD:AAL7745
Phebellia strigifrons|[2107]|JWDCJ096-11|BOLD:AAL7745
Anthomyiidae|[2108]|JWDCA341-10|BOLD:AAM6377
Anthomyiidae|[2109]|JWDCG227-10|BOLD:AAM6377
Anthomyiidae|[2110]|JWDCH526-10|BOLD:AAP7594
Botanophila hucketti|[2111]|JWDCL544-11|BOLD:ACE3220
Botanophila hucketti|[2112]|JWDCL545-11|BOLD:ACE3220
Botanophila hucketti|[2113]|JWDCL546-11|BOLD:ACE3220
Botanophila|[2114]|JWDCA358-10|BOLD:AAM6379
Botanophila|[2115]|JWDCI140-10|BOLD:AAM6379
Botanophila|[2116]|JWDCL548-11|BOLD:AAP2974
Botanophila|[2117]|JWDCL560-11|BOLD:AAG6980
Botanophila|[2118]|JWDCC915-10|BOLD:AAG6980
Botanophila|[2119]|MTDIC132-09|BOLD:AAG6980
Botanophila|[2120]|JWDCC836-10|BOLD:AAG6980
Botanophila|[2121]|JWDCA368-10|BOLD:AAG6980
Botanophila|[2122]|JWDCA430-10|BOLD:AAG6980
Botanophila|[2123]|JWDCL558-11|BOLD:AAG6980
Botanophila|[2124]|JWDCE590-10|BOLD:AAG6980
Botanophila|[2125]|JWDCF236-10|BOLD:AAG6980
Botanophila|[2126]|JWDCF237-10|BOLD:AAG6980
Botanophila|[2127]|JWDCF239-10|BOLD:AAG6980
Botanophila|[2128]|JWDCF244-10|BOLD:AAG6980
Botanophila|[2129]|JWDCE591-10|BOLD:AAG6980
Botanophila|[2130]|JWDCE776-10|BOLD:AAG6980
Botanophila|[2131]|JWDCE778-10|BOLD:AAG6980
Botanophila|[2132]|JWDCA328-10|BOLD:ABX5204
Botanophila|[2133]|JWDCH728-10|BOLD:ABX5204
Botanophila|[2134]|JWDCK154-11|BOLD:ABX5204
Botanophila|[2135]|JWDCA346-10|BOLD:ABX5204
Botanophila|[2136]|JWDCI107-10|BOLD:ABX5204
Botanophila|[2137]|JWDCI402-10|BOLD:ABX5204
Botanophila|[2138]|JWDCE729-10|BOLD:ABX5204
Botanophila sericea|[2139]|JWDCD073-10|BOLD:AAL7528
Botanophila sericea|[2140]|JWDCC914-10|BOLD:AAL7528
Botanophila sericea|[2141]|JWDCF276-10|BOLD:AAL7528
Botanophila sericea|[2142]|JWDCF653-10|BOLD:AAL7528
Lasiomma nr. consobrinum|[2143]|JWDCD146-10|BOLD:AAG2515
Lasiomma nr. consobrinum|[2144]|JWDCA322-10|BOLD:AAG2515
Lasiomma nr. consobrinum|[2145]|JWDCB084-10|BOLD:AAG2515
Lasiomma nr. consobrinum|[2146]|JWDCL007-11|BOLD:AAG2515
Lasiomma latipenne|[2147]|JWDCA337-10|BOLD:AAL7514
Lasiomma latipenne|[2148]|JWDCK152-11|BOLD:AAL7514
Lasiomma latipenne|[2149]|JWDCJ1490-11|BOLD:AAL7514
Lasiomma latipenne|[2150]|JWDCH608-10|BOLD:AAL7514
Lasiomma latipenne|[2151]|JWDCJ1419-11|BOLD:AAL7514
Lasiomma latipenne|[2152]|JWDCI636-10|BOLD:AAL7514
Lasiomma latipenne|[2153]|JWDCA353-10|BOLD:AAL7514
Lasiomma latipenne|[2154]|JWDCC098-10|BOLD:AAL7514
Lasiomma latipenne|[2155]|JWDCC119-10|BOLD:AAL7514
Lasiomma latipenne|[2156]|JWDCK147-11|BOLD:AAL7514
Lasiomma latipenne|[2157]|JWDCI028-10|BOLD:AAL7514
Lasiomma latipenne|[2158]|JWDCA334-10|BOLD:AAL7514
Lasiomma latipenne|[2159]|JWDCD144-10|BOLD:AAL7514
Lasiomma latipenne|[2160]|JWDCC069-10|BOLD:AAL7514
Lasiomma latipenne|[2161]|JWDCC149-10|BOLD:AAL7514
Lasiomma latipenne|[2162]|JWDCJ858-11|BOLD:AAL7514
Lasiomma latipenne|[2163]|JWDCJ1272-11|BOLD:AAL7514
Lasiomma latipenne|[2164]|JWDCJ1276-11|BOLD:AAL7514
Lasiomma latipenne|[2165]|JWDCH069-10|BOLD:AAL7514
Lasiomma latipenne|[2166]|JWDCI206-10|BOLD:AAL7514
Lasiomma latipenne|[2167]|JWDCF688-10|BOLD:AAL7514
Lasiomma latipenne|[2168]|JWDCF458-10|BOLD:AAL7514
Lasiomma|[2169]|JWDCC393-10|BOLD:AAL7520
Anthomyiidae|[2170]|JWDCI023-10|BOLD:ACI2788
Anthomyiidae|[2171]|JWDCJ1611-11|BOLD:ACI2788
Anthomyiidae|[2172]|JWDCI288-10|BOLD:ACI2788
Lasiomma atricaudum|[2173]|JWDCJ627-11|BOLD:AAG2512
Lasiomma atricaudum|[2174]|JWDCJ517-11|BOLD:AAG2512
Lasiomma atricaudum|[2175]|JWDCB081-10|BOLD:AAG2512
Lasiomma atricaudum|[2176]|JWDCL414-11|BOLD:AAG2512
Lasiomma atricaudum|[2177]|JWDCI208-10|BOLD:AAG2512
Lasiomma atricaudum|[2178]|JWDCI289-10|BOLD:AAG2512
Lasiomma atricaudum|[2179]|JWDCJ623-11|BOLD:AAG2512
Lasiomma atricaudum|[2180]|JWDCA365-10|BOLD:AAG2512
Lasiomma atricaudum|[2181]|JWDCJ109-11|BOLD:AAG2512
Lasiomma atricaudum|[2182]|JWDCJ1061-11|BOLD:AAG2512
Lasiomma atricaudum|[2183]|JWDCJ1065-11|BOLD:AAG2512
Lasiomma atricaudum|[2184]|JWDCL291-11|BOLD:AAG2512
Lasiomma atricaudum|[2185]|JWDCG762-10|BOLD:AAG2512
Lasiomma atricaudum|[2186]|JWDCH072-10|BOLD:AAG2512
Lasiomma atricaudum|[2187]|JWDCH077-10|BOLD:AAG2512
Lasiomma atricaudum|[2188]|JWDCA324-10|BOLD:AAG2512
Lasiomma atricaudum|[2189]|JWDCB064-10|BOLD:AAG2512

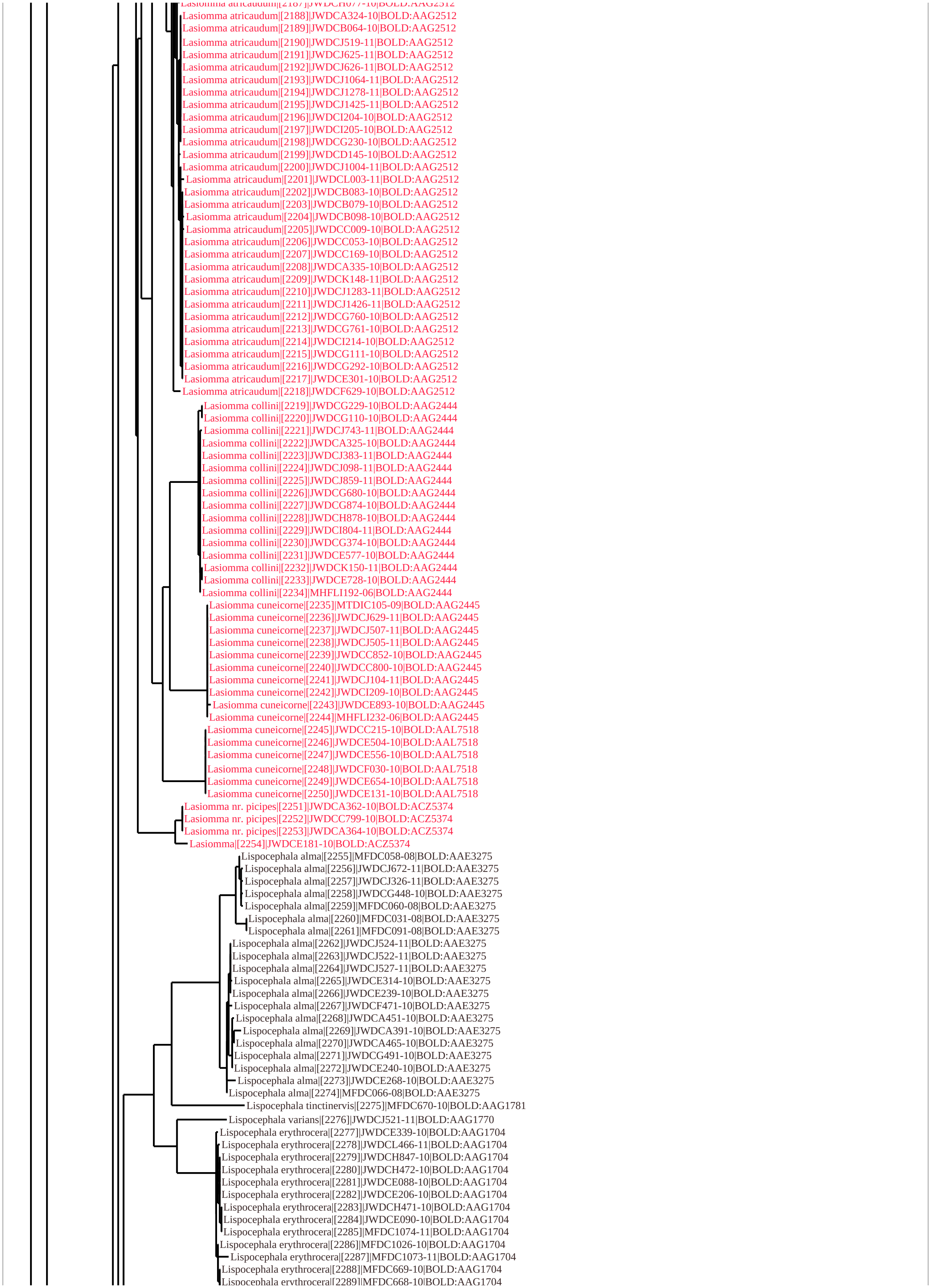
Lasiomma atricaudum|[2187]|JWDCH077-10|BOLD:AAG2512
Lasiomma atricaudum|[2188]|JWDCA324-10|BOLD:AAG2512
Lasiomma atricaudum|[2189]|JWDCB064-10|BOLD:AAG2512
Lasiomma atricaudum|[2190]|JWDCJ519-11|BOLD:AAG2512
Lasiomma atricaudum|[2191]|JWDCJ625-11|BOLD:AAG2512
Lasiomma atricaudum|[2192]|JWDCJ626-11|BOLD:AAG2512
Lasiomma atricaudum|[2193]|JWDCJ1064-11|BOLD:AAG2512
Lasiomma atricaudum|[2194]|JWDCJ1278-11|BOLD:AAG2512
Lasiomma atricaudum|[2195]|JWDCJ1425-11|BOLD:AAG2512
Lasiomma atricaudum|[2196]|JWDCI204-10|BOLD:AAG2512
Lasiomma atricaudum|[2197]|JWDCI205-10|BOLD:AAG2512
Lasiomma atricaudum|[2198]|JWDCG230-10|BOLD:AAG2512
Lasiomma atricaudum|[2199]|JWDCD145-10|BOLD:AAG2512
Lasiomma atricaudum|[2200]|JWDCJ1004-11|BOLD:AAG2512
Lasiomma atricaudum|[2201]|JWDCL003-11|BOLD:AAG2512
Lasiomma atricaudum|[2202]|JWDCB083-10|BOLD:AAG2512
Lasiomma atricaudum|[2203]|JWDCB079-10|BOLD:AAG2512
Lasiomma atricaudum|[2204]|JWDCB098-10|BOLD:AAG2512
Lasiomma atricaudum|[2205]|JWDCC009-10|BOLD:AAG2512
Lasiomma atricaudum|[2206]|JWDCC053-10|BOLD:AAG2512
Lasiomma atricaudum|[2207]|JWDCC169-10|BOLD:AAG2512
Lasiomma atricaudum|[2208]|JWDCA335-10|BOLD:AAG2512
Lasiomma atricaudum|[2209]|JWDCK148-11|BOLD:AAG2512
Lasiomma atricaudum|[2210]|JWDCJ1283-11|BOLD:AAG2512
Lasiomma atricaudum|[2211]|JWDCJ1426-11|BOLD:AAG2512
Lasiomma atricaudum|[2212]|JWDCG760-10|BOLD:AAG2512
Lasiomma atricaudum|[2213]|JWDCG761-10|BOLD:AAG2512
Lasiomma atricaudum|[2214]|JWDCI214-10|BOLD:AAG2512
Lasiomma atricaudum|[2215]|JWDCG111-10|BOLD:AAG2512
Lasiomma atricaudum|[2216]|JWDCG292-10|BOLD:AAG2512
Lasiomma atricaudum|[2217]|JWDCE301-10|BOLD:AAG2512
Lasiomma atricaudum|[2218]|JWDCF629-10|BOLD:AAG2512
Lasiomma collini|[2219]|JWDCG229-10|BOLD:AAG2444
Lasiomma collini|[2220]|JWDCG110-10|BOLD:AAG2444
Lasiomma collini|[2221]|JWDCJ743-11|BOLD:AAG2444
Lasiomma collini|[2222]|JWDCA325-10|BOLD:AAG2444
Lasiomma collini|[2223]|JWDCJ383-11|BOLD:AAG2444
Lasiomma collini|[2224]|JWDCJ098-11|BOLD:AAG2444
Lasiomma collini|[2225]|JWDCJ859-11|BOLD:AAG2444
Lasiomma collini|[2226]|JWDCG680-10|BOLD:AAG2444
Lasiomma collini|[2227]|JWDCG874-10|BOLD:AAG2444
Lasiomma collini|[2228]|JWDCH878-10|BOLD:AAG2444
Lasiomma collini|[2229]|JWDCI804-11|BOLD:AAG2444
Lasiomma collini|[2230]|JWDCG374-10|BOLD:AAG2444
Lasiomma collini|[2231]|JWDCE577-10|BOLD:AAG2444
Lasiomma collini|[2232]|JWDCK150-11|BOLD:AAG2444
Lasiomma collini|[2233]|JWDCE728-10|BOLD:AAG2444
Lasiomma collini|[2234]|MHFLI192-06|BOLD:AAG2444
Lasiomma cuneicorne|[2235]|MTDIC105-09|BOLD:AAG2445
Lasiomma cuneicorne|[2236]|JWDCJ629-11|BOLD:AAG2445
Lasiomma cuneicorne|[2237]|JWDCJ507-11|BOLD:AAG2445
Lasiomma cuneicorne|[2238]|JWDCJ505-11|BOLD:AAG2445
Lasiomma cuneicorne|[2239]|JWDCC852-10|BOLD:AAG2445
Lasiomma cuneicorne|[2240]|JWDCC800-10|BOLD:AAG2445
Lasiomma cuneicorne|[2241]|JWDCJ104-11|BOLD:AAG2445
Lasiomma cuneicorne|[2242]|JWDCI209-10|BOLD:AAG2445
Lasiomma cuneicorne|[2243]|JWDCE893-10|BOLD:AAG2445
Lasiomma cuneicorne|[2244]|MHFLI232-06|BOLD:AAG2445
Lasiomma cuneicorne|[2245]|JWDCC215-10|BOLD:AAL7518
Lasiomma cuneicorne|[2246]|JWDCE504-10|BOLD:AAL7518
Lasiomma cuneicorne|[2247]|JWDCE556-10|BOLD:AAL7518
Lasiomma cuneicorne|[2248]|JWDCF030-10|BOLD:AAL7518
Lasiomma cuneicorne|[2249]|JWDCE654-10|BOLD:AAL7518
Lasiomma cuneicorne|[2250]|JWDCE131-10|BOLD:AAL7518
Lasiomma nr. picipes|[2251]|JWDCA362-10|BOLD:ACZ5374
Lasiomma nr. picipes|[2252]|JWDCC799-10|BOLD:ACZ5374
Lasiomma nr. picipes|[2253]|JWDCA364-10|BOLD:ACZ5374
Lasiomma|[2254]|JWDCE181-10|BOLD:ACZ5374
Lispocephala alma|[2255]|MFDC058-08|BOLD:AAE3275
Lispocephala alma|[2256]|JWDCJ672-11|BOLD:AAE3275
Lispocephala alma|[2257]|JWDCJ326-11|BOLD:AAE3275
Lispocephala alma|[2258]|JWDCG448-10|BOLD:AAE3275
Lispocephala alma|[2259]|MFDC060-08|BOLD:AAE3275
Lispocephala alma|[2260]|MFDC031-08|BOLD:AAE3275
Lispocephala alma|[2261]|MFDC091-08|BOLD:AAE3275
Lispocephala alma|[2262]|JWDCJ524-11|BOLD:AAE3275
Lispocephala alma|[2263]|JWDCJ522-11|BOLD:AAE3275
Lispocephala alma|[2264]|JWDCJ527-11|BOLD:AAE3275
Lispocephala alma|[2265]|JWDCE314-10|BOLD:AAE3275
Lispocephala alma|[2266]|JWDCE239-10|BOLD:AAE3275
Lispocephala alma|[2267]|JWDCF471-10|BOLD:AAE3275
Lispocephala alma|[2268]|JWDCA451-10|BOLD:AAE3275
Lispocephala alma|[2269]|JWDCA391-10|BOLD:AAE3275
Lispocephala alma|[2270]|JWDCA465-10|BOLD:AAE3275
Lispocephala alma|[2271]|JWDCG491-10|BOLD:AAE3275
Lispocephala alma|[2272]|JWDCE240-10|BOLD:AAE3275
Lispocephala alma|[2273]|JWDCE268-10|BOLD:AAE3275
Lispocephala alma|[2274]|MFDC066-08|BOLD:AAE3275
Lispocephala tinctinervis|[2275]|MFDC670-10|BOLD:AAG1781
Lispocephala varians|[2276]|JWDCJ521-11|BOLD:AAG1770
Lispocephala erythrocera|[2277]|JWDCE339-10|BOLD:AAG1704
Lispocephala erythrocera|[2278]|JWDCL466-11|BOLD:AAG1704
Lispocephala erythrocera|[2279]|JWDCH847-10|BOLD:AAG1704
Lispocephala erythrocera|[2280]|JWDCH472-10|BOLD:AAG1704
Lispocephala erythrocera|[2281]|JWDCE088-10|BOLD:AAG1704
Lispocephala erythrocera|[2282]|JWDCE206-10|BOLD:AAG1704
Lispocephala erythrocera|[2283]|JWDCH471-10|BOLD:AAG1704
Lispocephala erythrocera|[2284]|JWDCE090-10|BOLD:AAG1704
Lispocephala erythrocera|[2285]|MFDC1074-11|BOLD:AAG1704
Lispocephala erythrocera|[2286]|MFDC1026-10|BOLD:AAG1704
Lispocephala erythrocera|[2287]|MFDC1073-11|BOLD:AAG1704
Lispocephala erythrocera|[2288]|MFDC669-10|BOLD:AAG1704
Lispocephala erythrocera|[2289]|MFDC668-10|BOLD:AAG1704

Lispocephala erythrocera|[2287]|MFDC1073-11|BOLD:AAG1704
Lispocephala erythrocera|[2288]|MFDC669-10|BOLD:AAG1704
Lispocephala erythrocera|[2289]|MFDC668-10|BOLD:AAG1704
Lispocephala erythrocera|[2290]|MFDC667-10|BOLD:AAG1704
Lispocephala erythrocera|[2291]|MFDC666-10|BOLD:AAG1704
Lispocephala erythrocera|[2292]|MFDC665-10|BOLD:AAG1704
Lispocephala erythrocera|[2293]|MFDC1025-10|BOLD:AAG1704
Lispocephala erythrocera|[2294]|MFDC671-10|BOLD:AAG1704
Lispocephala erythrocera|[2295]|JWDCE231-10|BOLD:AAG1704
Lispocephala erythrocera|[2296]|JWDCE205-10|BOLD:AAG1704
Lispocephala erythrocera|[2297]|JWDCE204-10|BOLD:AAG1704
Lispocephala erythrocera|[2298]|JWDCE091-10|BOLD:AAG1704
Lispocephala erythrocera|[2299]|JWDCE040-10|BOLD:AAG1704
Lispocephala erythrocera|[2300]|JWDCF522-10|BOLD:AAG1704
Lispocephala erythrocera|[2301]|JWDCE568-10|BOLD:AAG1704
Lispocephala erythrocera|[2302]|JWDCE567-10|BOLD:AAG1704
Lispocephala erythrocera|[2303]|JWDCE566-10|BOLD:AAG1704
Lispocephala erythrocera|[2304]|JWDCF216-10|BOLD:AAG1704
Lispocephala erythrocera|[2305]|JWDCF014-10|BOLD:AAG1704
Lispocephala erythrocera|[2306]|JWDCI911-11|BOLD:AAG1704
Lispocephala erythrocera|[2307]|JWDCI350-10|BOLD:AAG1704
Lispocephala erythrocera|[2308]|JWDCG537-10|BOLD:AAG1704
Lispocephala erythrocera|[2309]|JWDCG536-10|BOLD:AAG1704
Lispocephala erythrocera|[2310]|JWDCG534-10|BOLD:AAG1704
Lispocephala erythrocera|[2311]|JWDCG533-10|BOLD:AAG1704
Lispocephala erythrocera|[2312]|JWDCG532-10|BOLD:AAG1704
Lispocephala erythrocera|[2313]|JWDCG531-10|BOLD:AAG1704
Lispocephala erythrocera|[2314]|JWDCG519-10|BOLD:AAG1704
Lispocephala erythrocera|[2315]|JWDCG518-10|BOLD:AAG1704
Lispocephala erythrocera|[2316]|JWDCH512-10|BOLD:AAG1704
Lispocephala erythrocera|[2317]|JWDCG311-10|BOLD:AAG1704
Lispocephala erythrocera|[2318]|JWDCH849-10|BOLD:AAG1704
Lispocephala erythrocera|[2319]|JWDCH216-10|BOLD:AAG1704
Lispocephala erythrocera|[2320]|JWDCH096-10|BOLD:AAG1704
Lispocephala erythrocera|[2321]|JWDCH028-10|BOLD:AAG1704
Lispocephala erythrocera|[2322]|JWDCG726-10|BOLD:AAG1704
Lispocephala erythrocera|[2323]|JWDCH921-10|BOLD:AAG1704
Lispocephala erythrocera|[2324]|JWDCH919-10|BOLD:AAG1704
Lispocephala erythrocera|[2325]|JWDCH850-10|BOLD:AAG1704
Lispocephala erythrocera|[2326]|JWDCH848-10|BOLD:AAG1704
Lispocephala erythrocera|[2327]|JWDCJ1346-11|BOLD:AAG1704
Lispocephala erythrocera|[2328]|JWDCJ671-11|BOLD:AAG1704
Lispocephala erythrocera|[2329]|JWDCC420-10|BOLD:AAG1704
Lispocephala erythrocera|[2330]|MFDC882-10|BOLD:AAG1704
Lispocephala erythrocera|[2331]|JWDCE317-10|BOLD:AAG1704
Lispocephala erythrocera|[2332]|MFDC672-10|BOLD:AAG1704
Helina maculipennis|[2333]|MFDC035-08|BOLD:AAK4685
Helina maculipennis|[2334]|MFDC719-10|BOLD:AAK4685
Helina maculipennis|[2335]|MFDC720-10|BOLD:AAK4685
Helina humilis|[2336]|JWDCC094-10|BOLD:AAL5778
Helina humilis|[2337]|JWDCJ530-11|BOLD:AAL5778
Helina humilis|[2338]|MFDC967-10|BOLD:AAL5778
Helina humilis|[2339]|MFDC968-10|BOLD:AAL5778
Helina humilis|[2340]|MFDC969-10|BOLD:AAL5778
Helina longicornis|[2341]|MFDC714-10|BOLD:AAL9497
Helina longicornis|[2342]|MFDC715-10|BOLD:AAL9497
Helina annosa|[2343]|JWDCI021-10|BOLD:AAK4683
Helina annosa|[2344]|MFDC015-08|BOLD:AAK4683
Helina flavisquama|[2345]|MFDC100-09|BOLD:AAC7186
Helina flavisquama|[2346]|JWDCL061-11|BOLD:AAC7186
Helina flavisquama|[2347]|JWDCA477-10|BOLD:AAC7186
Helina flavisquama|[2348]|JWDCL348-11|BOLD:AAC7186
Helina flavisquama|[2349]|JWDCJ308-11|BOLD:AAC7186
Helina flavisquama|[2350]|JWDCA448-10|BOLD:AAC7186
Helina flavisquama|[2351]|JWDCA432-10|BOLD:AAC7186
Helina flavisquama|[2352]|JWDCA460-10|BOLD:AAC7186
Helina flavisquama|[2353]|JWDCL208-11|BOLD:AAC7186
Helina flavisquama|[2354]|JWDCI419-10|BOLD:AAC7186
Helina flavisquama|[2355]|JWDCI424-10|BOLD:AAC7186
Helina flavisquama|[2356]|MHFLI173-06|BOLD:AAC7186
Helina flavisquama|[2357]|JWDCL071-11|BOLD:AAC7186
Helina flavisquama|[2358]|MFDC099-09|BOLD:AAC7186
Helina flavisquama|[2359]|JWDCA445-10|BOLD:AAC7186
Helina flavisquama|[2360]|JWDCA523-10|BOLD:AAC7186
Helina flavisquama|[2361]|MFDC106-09|BOLD:AAC7186
Helina flavisquama|[2362]|MFDC103-09|BOLD:AAC7186
Helina flavisquama|[2363]|JWDCL068-11|BOLD:AAC7186
Helina flavisquama|[2364]|JWDCL053-11|BOLD:AAC7186
Helina flavisquama|[2365]|JWDCG737-10|BOLD:AAC7186
Helina flavisquama|[2366]|JWDCG037-10|BOLD:AAC7186
Helina flavisquama|[2367]|MHFLI167-06|BOLD:AAC7186
Helina flavisquama|[2368]|JWDCJ386-11|BOLD:AAC7186
Helina flavisquama|[2369]|JWDCJ1010-11|BOLD:AAC7186
Helina flavisquama|[2370]|JWDCJ025-11|BOLD:AAC7186
Helina flavisquama|[2371]|JWDCL337-11|BOLD:AAC7186
Helina flavisquama|[2372]|JWDCG117-10|BOLD:AAC7186
Helina flavisquama|[2373]|JWDCH303-10|BOLD:AAC7186
Helina flavisquama|[2374]|JWDCI429-10|BOLD:AAC7186
Helina flavisquama|[2375]|MFDC101-09|BOLD:AAC7186
Helina flavisquama|[2376]|MFDC102-09|BOLD:AAC7186
Helina flavisquama|[2377]|MFDC104-09|BOLD:AAC7186
Helina flavisquama|[2378]|MHFLI916-09|BOLD:AAC7186
Helina flavisquama|[2379]|MHFLY370-09|BOLD:AAC7186
Helina fulvisquama|[2380]|JWDCJ092-11|BOLD:ABX5991
Helina fulvisquama|[2381]|JWDCC919-10|BOLD:ABX5991
Helina fulvisquama|[2382]|MFDC109-09|BOLD:ABX5991
Helina fulvisquama|[2383]|JWDCC785-10|BOLD:ABX5991
Helina fulvisquama|[2384]|JWDCC038-10|BOLD:ABX5991
Helina fulvisquama|[2385]|JWDCK273-11|BOLD:ABX5991
Helina fulvisquama|[2386]|JWDCJ1067-11|BOLD:ABX5991
Helina fulvisquama|[2387]|MTDIC099-09|BOLD:ABX5991
Helina fulvisquama|[2388]|MFDC107-09|BOLD:ABX5991
[illegible]

Helina fulvisquama|[2386]|JWDCJ1067-11|BOLD:ABX5991
Helina fulvisquama|[2387]|MTDIC099-09|BOLD:ABX5991
Helina fulvisquama|[2388]|MFDC107-09|BOLD:ABX5991
Helina fulvisquama|[2389]|JWDCG787-10|BOLD:ABX5991
Helina fulvisquama|[2390]|MFDC108-09|BOLD:ABX5991
Helina fulvisquama|[2391]|JWDCC883-10|BOLD:ABX5991
Helina fulvisquama|[2392]|MFDC1099-11|BOLD:ABX5991
Helina fulvisquama|[2393]|MFDC074-08|BOLD:ABX5991
Helina fulvisquama|[2394]|MFDC716-10|BOLD:ABX5991
Helina fulvisquama|[2395]|MFDC717-10|BOLD:ABX5991
Helina fulvisquama|[2396]|MFDC718-10|BOLD:ABX5991
Helina evecta|[2397]|JWDCC885-10|BOLD:AAC2498
Helina evecta|[2398]|JWDCJ878-11|BOLD:AAC2498
Helina evecta|[2399]|MHFLI275-06|BOLD:AAC2498
Helina evecta|[2400]|JWDCI792-11|BOLD:AAC2498
Helina evecta|[2401]|JWDCI885-11|BOLD:AAC2498
Helina evecta|[2402]|MFDC105-09|BOLD:AAC2498
Helina evecta|[2403]|MFDC126-09|BOLD:AAC2498
Helina evecta|[2404]|MFDC119-09|BOLD:AAC2498
Helina evecta|[2405]|MHFLI169-06|BOLD:AAC2498
Helina evecta|[2406]|JWDCF404-10|BOLD:AAC2498
Helina evecta|[2407]|JWDCE747-10|BOLD:AAC2498
Helina evecta|[2408]|JWDCF453-10|BOLD:AAC2498
Helina evecta|[2409]|JWDCF685-10|BOLD:AAC2498
Helina evecta|[2410]|JWDCE710-10|BOLD:AAC2498
Helina evecta|[2411]|JWDCF630-10|BOLD:AAC2498
Helina evecta|[2412]|JWDCI637-10|BOLD:AAC2498
Helina evecta|[2413]|JWDCI638-10|BOLD:AAC2498
Helina evecta|[2414]|JWDCI292-10|BOLD:AAC2498
Helina evecta|[2415]|JWDCI801-11|BOLD:AAC2498
Helina evecta|[2416]|JWDCF217-10|BOLD:AAC2498
Helina evecta|[2417]|JWDCE664-10|BOLD:AAC2498
Helina evecta|[2418]|JWDCJ867-11|BOLD:AAC2498
Helina evecta|[2419]|JWDCJ876-11|BOLD:AAC2498
Helina evecta|[2420]|JWDCJ1172-11|BOLD:AAC2498
Helina evecta|[2421]|JWDCJ1178-11|BOLD:AAC2498
Helina evecta|[2422]|JWDCJ853-11|BOLD:AAC2498
Helina evecta|[2423]|JWDCJ1007-11|BOLD:AAC2498
Helina evecta|[2424]|JWDCJ1296-11|BOLD:AAC2498
Helina evecta|[2425]|JWDCJ324-11|BOLD:AAC2498
Helina evecta|[2426]|JWDCJ675-11|BOLD:AAC2498
Helina evecta|[2427]|JWDCJ677-11|BOLD:AAC2498
Helina evecta|[2428]|JWDCJ1341-11|BOLD:AAC2498
Helina evecta|[2429]|JWDCL059-11|BOLD:AAC2498
Helina evecta|[2430]|JWDCL271-11|BOLD:AAC2498
Helina evecta|[2431]|JWDCL280-11|BOLD:AAC2498
Helina evecta|[2432]|JWDCL421-11|BOLD:AAC2498
Helina evecta|[2433]|JWDCL422-11|BOLD:AAC2498
Helina evecta|[2434]|JWDCJ555-11|BOLD:AAC2498
Helina evecta|[2435]|JWDCJ570-11|BOLD:AAC2498
Helina evecta|[2436]|JWDCL432-11|BOLD:AAC2498
Helina evecta|[2437]|JWDCG592-10|BOLD:AAC2498
Helina evecta|[2438]|JWDCG594-10|BOLD:AAC2498
Helina evecta|[2439]|JWDCG596-10|BOLD:AAC2498
Helina evecta|[2440]|JWDCG849-10|BOLD:AAC2498
Helina evecta|[2441]|JWDCG868-10|BOLD:AAC2498
Helina evecta|[2442]|JWDCG872-10|BOLD:AAC2498
Helina evecta|[2443]|JWDCG016-10|BOLD:AAC2498
Helina evecta|[2444]|JWDCG017-10|BOLD:AAC2498
Helina evecta|[2445]|JWDCG018-10|BOLD:AAC2498
Helina evecta|[2446]|JWDCH164-10|BOLD:AAC2498
Helina evecta|[2447]|JWDCG283-10|BOLD:AAC2498
Helina evecta|[2448]|JWDCI886-11|BOLD:AAC2498
Helina evecta|[2449]|JWDCI888-11|BOLD:AAC2498
Helina evecta|[2450]|JWDCF135-10|BOLD:AAC2498
Helina evecta|[2451]|JWDCF212-10|BOLD:AAC2498
Helina evecta|[2452]|MFDC127-09|BOLD:AAC2498
Helina evecta|[2453]|MFDC128-09|BOLD:AAC2498
Helina evecta|[2454]|MFDC129-09|BOLD:AAC2498
Helina evecta|[2455]|MFDC118-09|BOLD:AAC2498
Helina evecta|[2456]|MFDC120-09|BOLD:AAC2498
Helina evecta|[2457]|MFDC034-08|BOLD:AAC2498
Helina evecta|[2458]|JWDCJ532-11|BOLD:AAC2498
Helina evecta|[2459]|JWDCJ554-11|BOLD:AAC2498
Helina evecta|[2460]|JWDCA525-10|BOLD:AAC2498
Helina evecta|[2461]|JWDCA531-10|BOLD:AAC2498
Helina evecta|[2462]|JWDCA515-10|BOLD:AAC2498
Helina evecta|[2463]|JWDCA518-10|BOLD:AAC2498
Helina evecta|[2464]|JWDCA461-10|BOLD:AAC2498
Helina evecta|[2465]|JWDCA514-10|BOLD:AAC2498
Helina evecta|[2466]|JWDCA435-10|BOLD:AAC2498
Helina evecta|[2467]|JWDCA459-10|BOLD:AAC2498
Helina evecta|[2468]|JWDCI890-11|BOLD:AAC2498
Helina evecta|[2469]|JWDCC266-10|BOLD:AAC2498
Helina evecta|[2470]|JWDCC100-10|BOLD:AAC2498
Helina evecta|[2471]|JWDCC047-10|BOLD:AAC2498
Helina evecta|[2472]|JWDCC040-10|BOLD:AAC2498
Helina evecta|[2473]|JWDCC917-10|BOLD:AAC2498
Helina evecta|[2474]|JWDCB120-10|BOLD:AAC2498
Helina evecta|[2475]|JWDCB118-10|BOLD:AAC2498
Helina evecta|[2476]|MTDIC098-09|BOLD:AAC2498
Helina evecta|[2477]|JWDCC096-10|BOLD:AAC2498
Helina evecta|[2478]|JWDCB117-10|BOLD:AAC2498
Helina evecta|[2479]|JWDCA526-10|BOLD:AAC2498
Helina evecta|[2480]|JWDCF161-10|BOLD:AAC2498
Helina evecta|[2481]|MFDC045-08|BOLD:AAC2498
Helina evecta|[2482]|MFDC125-09|BOLD:AAC2498
Helina evecta|[2483]|MFDC124-09|BOLD:AAC2498
Helina evecta|[2484]|MFDC123-09|BOLD:AAC2498
Helina evecta|[2485]|MFDC1098-11|BOLD:AAC2498
Helina evecta|[2486]|MFDC130-09|BOLD:AAC2498
Helina evecta|[2487]|MFDC121-09|BOLD:AAC2498
Helina evecta|[2488]|JWDCC787-10|BOLD:AAC2498

Helina evecta|[2486]|MFDC130-09|BOLD:AAC2498
Helina evecta|[2487]|MFDC121-09|BOLD:AAC2498
Helina evecta|[2488]|JWDCC787-10|BOLD:AAC2498
Helina evecta|[2489]|MFDC115-09|BOLD:AAC2498
Helina evecta|[2490]|JWDCC142-10|BOLD:AAC2498
Helina evecta|[2491]|MHFLI175-06|BOLD:AAC2498
Helina evecta|[2492]|JWDCE642-10|BOLD:AAC2498
Helina evecta|[2493]|JWDCF538-10|BOLD:AAC2498
Helina evecta|[2494]|JWDCE711-10|BOLD:AAC2498
Helina evecta|[2495]|JWDCF765-10|BOLD:AAC2498
Helina evecta|[2496]|JWDCE038-10|BOLD:AAC2498
Helina evecta|[2497]|MFDC019-08|BOLD:AAC2498
Helina evecta|[2498]|JWDCE640-10|BOLD:AAC2498
Helina evecta|[2499]|JWDCF240-10|BOLD:AAC2498
Helina evecta|[2500]|JWDCF159-10|BOLD:AAC2498
Helina evecta|[2501]|JWDCF033-10|BOLD:AAC2498
Helina evecta|[2502]|JWDCH138-10|BOLD:AAC2498
Helina evecta|[2503]|JWDCH791-10|BOLD:AAC2498
Helina evecta|[2504]|JWDCJ952-11|BOLD:AAC2498
Helina evecta|[2505]|JWDCJ795-11|BOLD:AAC2498
Helina evecta|[2506]|JWDCC158-10|BOLD:AAC2498
Helina evecta|[2507]|JWDCC107-10|BOLD:AAC2498
Helina evecta|[2508]|JWDCJ097-11|BOLD:AAC2498
Helina evecta|[2509]|JWDCC106-10|BOLD:AAC2498
Helina evecta|[2510]|JWDCC104-10|BOLD:AAC2498
Helina evecta|[2511]|JWDCF632-10|BOLD:AAC2498
Helina evecta|[2512]|MFDC020-08|BOLD:AAC2498
Helina evecta|[2513]|MFDC032-08|BOLD:AAC2498
Helina evecta|[2514]|MFDC116-09|BOLD:AAC2498
Helina evecta|[2515]|JWDCC282-10|BOLD:AAC2498
Helina evecta|[2516]|JWDCA431-10|BOLD:AAC2498
Helina evecta|[2517]|JWDCC186-10|BOLD:AAC2498
Helina evecta|[2518]|JWDCF402-10|BOLD:AAC2498
Helina evecta|[2519]|JWDCC918-10|BOLD:AAC2498
Helina evecta|[2520]|JWDCI793-11|BOLD:AAC2498
Helina evecta|[2521]|MHFLI260-06|BOLD:AAC2498
Helina evecta|[2522]|MHFLI170-06|BOLD:AAC2498
Helina evecta|[2523]|MHFLI166-06|BOLD:AAC2498
Helina evecta|[2524]|JWDCF608-10|BOLD:AAC2498
Helina evecta|[2525]|JWDCE363-10|BOLD:AAC2498
Helina evecta|[2526]|JWDCE746-10|BOLD:AAC2498
Helina evecta|[2527]|JWDCE383-10|BOLD:AAC2498
Helina evecta|[2528]|JWDCF035-10|BOLD:AAC2498
Helina evecta|[2529]|JWDCI784-11|BOLD:AAC2498
Helina evecta|[2530]|JWDCH137-10|BOLD:AAC2498
Helina evecta|[2531]|JWDCJ553-11|BOLD:AAC2498
Helina evecta|[2532]|JWDCC402-10|BOLD:AAC2498
Helina evecta|[2533]|JWDCC839-10|BOLD:AAC2498
Helina evecta|[2534]|JWDCC788-10|BOLD:AAC2498
Helina evecta|[2535]|JWDCC786-10|BOLD:AAC2498
Helina evecta|[2536]|JWDCF641-10|BOLD:AAC2498
Helina evecta|[2537]|JWDCF687-10|BOLD:AAC2498
Helina evecta|[2538]|JWDCE786-10|BOLD:AAC2498
Helina evecta|[2539]|JWDCF917-10|BOLD:AAC2498
Helina evecta|[2540]|MFDC117-09|BOLD:AAC2498
Helina evecta|[2541]|MFDC122-09|BOLD:AAC2498
Helina evecta|[2542]|MFDC050-08|BOLD:AAC2498
Helina evecta|[2543]|MFDC711-10|BOLD:AAC2498
Helina cinerella|[2544]|JWDCF205-10|BOLD:AAG1706
Helina cinerella|[2545]|JWDCF582-10|BOLD:AAG1706
Helina cinerella|[2546]|JWDCF580-10|BOLD:AAG1706
Helina cinerella|[2547]|JWDCF802-10|BOLD:AAG1706
Helina cinerella|[2548]|MFDC973-10|BOLD:AAG1706
Helina cinerella|[2549]|JWDCI237-10|BOLD:AAG1706
Helina cinerella|[2550]|MFDC705-10|BOLD:AAG1706
Helina cinerella|[2551]|MFDC706-10|BOLD:AAG1706
Helina cinerella|[2552]|MFDC707-10|BOLD:AAG1706
Helina cinerella|[2553]|MFDC708-10|BOLD:AAG1706
Helina cinerella|[2554]|MFDC725-10|BOLD:AAG1706
Helina obscurata|[2555]|MHFLI160-06|BOLD:AAE6931
Helina obscurata|[2556]|JWDCL343-11|BOLD:AAE6931
Helina obscurata|[2557]|JWDCG345-10|BOLD:AAE6931
Helina obscurata|[2558]|MFDC112-09|BOLD:AAE6931
Helina obscurata|[2559]|MFDC110-09|BOLD:AAE6931
Helina obscurata|[2560]|JWDCD024-10|BOLD:AAE6931
Helina obscurata|[2561]|JWDCJ261-11|BOLD:AAE6931
Helina obscurata|[2562]|JWDCK137-11|BOLD:AAE6931
Helina obscurata|[2563]|JWDCL336-11|BOLD:AAE6931
Helina obscurata|[2564]|JWDCH152-10|BOLD:AAE6931
Helina obscurata|[2565]|MFDC111-09|BOLD:AAE6931
Helina obscurata|[2566]|MFDC721-10|BOLD:AAE6931
Helina spinosa|[2567]|MHFLI117-06|BOLD:AAG1687
Helina spinosa|[2568]|MHFLI625-07|BOLD:AAG1687
Helina spinosa|[2569]|MFDC726-10|BOLD:AAG1687
Helina marguerita|[2570]|MFDC113-09|BOLD:AAF1480
Helina marguerita|[2571]|MFDC087-08|BOLD:AAF1480
Helina marguerita|[2572]|MFDC114-09|BOLD:AAF1480
Helina marguerita|[2573]|MFDC723-10|BOLD:AAF1480
Helina subvittata|[2574]|MFDC724-10|BOLD:AAG1694
Helina laxifrons|[2575]|MFDC709-10|BOLD:AAL6541
Helina laxifrons|[2576]|MFDC710-10|BOLD:AAL6541
Helina laxifrons|[2577]|MFDC712-10|BOLD:AAL6541
Helina squalens|[2578]|MFDC733-10|BOLD:AAG1695
Helina squalens|[2579]|JWDCE408-10|BOLD:AAG1695
Helina squalens|[2580]|JWDCC188-10|BOLD:AAG1695
Helina squalens|[2581]|JWDCC412-10|BOLD:AAG1695
Helina squalens|[2582]|JWDCE362-10|BOLD:AAG1695
Helina squalens|[2583]|JWDCE382-10|BOLD:AAG1695
Helina squalens|[2584]|JWDCF136-10|BOLD:AAG1695
Helina squalens|[2585]|JWDCE546-10|BOLD:AAG1695
Helina squalens|[2586]|JWDCE745-10|BOLD:AAG1695
Helina squalens|[2587]|MFDC734-10|BOLD:AAG1695
[illegible]

Helina squalens|[2585]|JWDCE546-10|BOLD:AAG1695
Helina squalens|[2586]|JWDCE745-10|BOLD:AAG1695
Helina squalens|[2587]|MFDC734-10|BOLD:AAG1695
Helina squalens|[2588]|MFDC735-10|BOLD:AAG1695
Helina squalens|[2589]|JWDCF134-10|BOLD:AAG1695
Helina squalens|[2590]|JWDCE381-10|BOLD:AAG1695
Helina squalens|[2591]|JSFLA031-08|BOLD:AAG1695
Helina squalens|[2592]|MFDC736-10|BOLD:AAG1695
Phaonia subfuscinervis|[2593]|JWDCF567-10|BOLD:AAF6792
Phaonia subfuscinervis|[2594]|JWDCC153-10|BOLD:AAF6792
Phaonia subfuscinervis|[2595]|JWDCJ534-11|BOLD:AAF6792
Phaonia subfuscinervis|[2596]|JWDCC183-10|BOLD:AAF6792
Phaonia subfuscinervis|[2597]|JWDCG187-10|BOLD:AAF6792
Phaonia subfuscinervis|[2598]|JWDCE895-10|BOLD:AAF6792
Phaonia subfuscinervis|[2599]|JWDCB119-10|BOLD:AAF6792
Phaonia subfuscinervis|[2600]|JWDCC238-10|BOLD:AAF6792
Phaonia subfuscinervis|[2601]|JWDCJ528-11|BOLD:AAF6792
Phaonia subfuscinervis|[2602]|JWDCG125-10|BOLD:AAF6792
Phaonia subfuscinervis|[2603]|JWDCI912-11|BOLD:AAF6792
Phaonia subfuscinervis|[2604]|JWDCF438-10|BOLD:AAF6792
Phaonia subfuscinervis|[2605]|JWDCE787-10|BOLD:AAF6792
Phaonia subfuscinervis|[2606]|JWDCF631-10|BOLD:AAF6792
Phaonia subfuscinervis|[2607]|JWDCF643-10|BOLD:AAF6792
Phaonia subfuscinervis|[2608]|JWDCE075-10|BOLD:AAF6792
Phaonia subfuscinervis|[2609]|JWDCE262-10|BOLD:AAF6792
Phaonia subfuscinervis|[2610]|MFDC013-08|BOLD:AAF6792
Phaonia subfuscinervis|[2611]|MFDC011-08|BOLD:AAF6792
Phaonia subfuscinervis|[2612]|MFDC713-10|BOLD:AAF6792
Phaonia serva|[2613]|MFDC329-09|BOLD:AAC0888
Phaonia|[2614]|JWDCJ705-11|BOLD:AAC0888
Phaonia serva|[2615]|MFDC769-10|BOLD:AAC0888
Phaonia serva|[2616]|MFDC379-09|BOLD:AAC0888
Phaonia serva|[2617]|MFDC770-10|BOLD:AAC0888
Phaonia|[2618]|JWDCG797-10|BOLD:AAC0888
Phaonia|[2619]|JWDCJ1614-11|BOLD:AAC0888
Phaonia|[2620]|JWDCK271-11|BOLD:AAC0888
Phaonia savonoskii|[2621]|MFDC340-09|BOLD:AAC0888
Phaonia savonoskii|[2622]|MFDC341-09|BOLD:AAC0888
Phaonia savonoskii|[2623]|MFDC334-09|BOLD:AAC0888
Phaonia savonoskii|[2624]|MFDC335-09|BOLD:AAC0888
Phaonia|[2625]|JWDCL206-11|BOLD:AAC0888
Phaonia|[2626]|JWDCC884-10|BOLD:AAC0888
Phaonia|[2627]|JWDCJ388-11|BOLD:AAC0888
Phaonia|[2628]|JWDCJ024-11|BOLD:AAC0888
Phaonia|[2629]|JWDCJ1545-11|BOLD:AAC0888
Phaonia|[2630]|JWDCJ1009-11|BOLD:AAC0888
Phaonia|[2631]|JWDCJ1011-11|BOLD:AAC0888
Phaonia|[2632]|JWDCJ1495-11|BOLD:AAC0888
Phaonia|[2633]|JWDCJ1503-11|BOLD:AAC0888
Phaonia|[2634]|JWDCJ022-11|BOLD:AAC0888
Phaonia|[2635]|JWDCG785-10|BOLD:AAC0888
Phaonia|[2636]|JWDCH104-10|BOLD:AAC0888
Phaonia|[2637]|JWDCH139-10|BOLD:AAC0888
Phaonia|[2638]|JWDCI236-10|BOLD:AAC0888
Phaonia|[2639]|JWDCF439-10|BOLD:AAC0888
Phaonia|[2640]|JWDCF888-10|BOLD:AAC0888
Phaonia|[2641]|JWDCF895-10|BOLD:AAC0888
Phaonia savonoskii|[2642]|MFDC330-09|BOLD:AAC0888
Phaonia savonoskii|[2643]|MFDC337-09|BOLD:AAC0888
Phaonia savonoskii|[2644]|MFDC349-09|BOLD:AAC0888
Phaonia savonoskii|[2645]|MFDC333-09|BOLD:AAC0888
Phaonia savonoskii|[2646]|MFDC336-09|BOLD:AAC0888
Phaonia savonoskii|[2647]|MFDC338-09|BOLD:AAC0888
Phaonia savonoskii|[2648]|MFDC339-09|BOLD:AAC0888
Phaonia savonoskii|[2649]|MFDC331-09|BOLD:AAC0888
Phaonia savonoskii|[2650]|MFDC332-09|BOLD:AAC0888
Phaonia savonoskii|[2651]|MFDC012-08|BOLD:AAC0888
Phaonia savonoskii|[2652]|MFDC067-08|BOLD:AAC0888
Phaonia savonoskii|[2653]|MFDC722-10|BOLD:AAC0888
Phaonia inenarrabilis|[2654]|MFDC346-09|BOLD:AAL2820
Phaonia luteva|[2655]|JWDCJ1435-11|BOLD:AAE6220
Phaonia luteva|[2656]|JWDCI643-10|BOLD:AAE6220
Phaonia luteva|[2657]|JWDCG593-10|BOLD:AAE6220
Phaonia luteva|[2658]|JWDCL331-11|BOLD:AAE6220
Phaonia luteva|[2659]|JWDCL276-11|BOLD:AAE6220
Phaonia luteva|[2660]|JWDCL060-11|BOLD:AAE6220
Phaonia luteva|[2661]|JWDCJ091-11|BOLD:AAE6220
Phaonia luteva|[2662]|JWDCJ854-11|BOLD:AAE6220
Phaonia luteva|[2663]|JWDCJ745-11|BOLD:AAE6220
Phaonia luteva|[2664]|JWDCC124-10|BOLD:AAE6220
Phaonia luteva|[2665]|JWDCC017-10|BOLD:AAE6220
Phaonia luteva|[2666]|JWDCJ090-11|BOLD:AAE6220
Phaonia luteva|[2667]|JWDCH308-10|BOLD:AAE6220
Phaonia luteva|[2668]|MFDC361-09|BOLD:AAE6220
Phaonia luteva|[2669]|JWDCA533-10|BOLD:AAE6220
Phaonia luteva|[2670]|JWDCG280-10|BOLD:AAE6220
Phaonia luteva|[2671]|MFDC353-09|BOLD:AAE6220
Phaonia luteva|[2672]|MFDC364-09|BOLD:AAE6220
Phaonia luteva|[2673]|MFDC350-09|BOLD:AAE6220
Phaonia luteva|[2674]|MFDC354-09|BOLD:AAE6220
Phaonia luteva|[2675]|MFDC039-08|BOLD:AAE6220
Phaonia errans|[2676]|MFDC325-09|BOLD:AAE6221
Phaonia errans|[2677]|MFDC328-09|BOLD:AAE6221
Phaonia errans|[2678]|MFDC360-09|BOLD:AAE6221
Phaonia errans|[2679]|JWDCC033-10|BOLD:AAE6221
Phaonia errans|[2680]|MFDC324-09|BOLD:AAE6221
Phaonia errans|[2681]|JWDCI889-11|BOLD:AAE6221
Phaonia errans|[2682]|JWDCJ1609-11|BOLD:AAE6221
Phaonia errans|[2683]|JWDCK272-11|BOLD:AAE6221
Phaonia errans|[2684]|JWDCK142-11|BOLD:AAE6221
Phaonia errans|[2685]|JWDCA458-10|BOLD:AAE6221
Phaonia errans|[2686]|JWDCC150-10|BOLD:AAE6221
Phaonia errans|[2687]|JWDCC091-10|BOLD:AAE6221

Phaonia errans|[2685]|JWDCA458-10|BOLD:AAE6221
Phaonia errans|[2686]|JWDCC150-10|BOLD:AAE6221
Phaonia errans|[2687]|JWDCC091-10|BOLD:AAE6221
Phaonia errans|[2688]|MFDC326-09|BOLD:AAE6221
Phaonia errans|[2689]|MFDC327-09|BOLD:AAE6221
Phaonia errans|[2690]|MFDC086-08|BOLD:AAE6221
Phaonia protuberans|[2691]|JWDCF568-10|BOLD:AAE6219
Phaonia protuberans|[2692]|JWDCG764-10|BOLD:AAE6219
Phaonia protuberans|[2693]|JWDCJ1613-11|BOLD:AAE6219
Phaonia protuberans|[2694]|JWDCJ643-11|BOLD:AAE6219
Phaonia protuberans|[2695]|JWDCJ641-11|BOLD:AAE6219
Phaonia protuberans|[2696]|JWDCJ531-11|BOLD:AAE6219
Phaonia protuberans|[2697]|JWDCC129-10|BOLD:AAE6219
Phaonia protuberans|[2698]|JWDCC789-10|BOLD:AAE6219
Phaonia protuberans|[2699]|JWDCG765-10|BOLD:AAE6219
Phaonia protuberans|[2700]|JWDCG116-10|BOLD:AAE6219
Phaonia protuberans|[2701]|JWDCF892-10|BOLD:AAE6219
Phaonia protuberans|[2702]|MFDC357-09|BOLD:AAE6219
Phaonia protuberans|[2703]|JWDCJ546-11|BOLD:AAE6219
Phaonia protuberans|[2704]|JWDCE422-10|BOLD:AAE6219
Phaonia protuberans|[2705]|JWDCF865-10|BOLD:AAE6219
Phaonia protuberans|[2706]|JWDCF861-10|BOLD:AAE6219
Phaonia protuberans|[2707]|MFDC356-09|BOLD:AAE6219
Phaonia protuberans|[2708]|MFDC382-09|BOLD:AAE6219
Phaonia protuberans|[2709]|MFDC075-08|BOLD:AAE6219
Phaonia alpicola|[2710]|JWDCI794-11|BOLD:AAG1764
Phaonia alpicola|[2711]|MFDC765-10|BOLD:AAG1764
Phaonia alpicola|[2712]|MFDC766-10|BOLD:AAG1764
Phaonia monticola|[2713]|MFDC345-09|BOLD:AAI2888
Phaonia rugia|[2714]|JWDCF358-10|BOLD:AAL2787
Phaonia rugia|[2715]|JWDCF870-10|BOLD:AAL2787
Phaonia rugia|[2716]|MFDC348-09|BOLD:AAL2787
Phaonia consobrina|[2717]|MFDC008-08|BOLD:AAF6699
Phaonia consobrina|[2718]|MFDC344-09|BOLD:AAF6699
Phaonia consobrina|[2719]|JWDCC102-10|BOLD:AAF6699
Phaonia consobrina|[2720]|JWDCC016-10|BOLD:AAF6699
Phaonia consobrina|[2721]|JWDCC881-10|BOLD:AAF6699
Phaonia consobrina|[2722]|JWDCI395-10|BOLD:AAF6699
Phaonia consobrina|[2723]|JWDCE896-10|BOLD:AAF6699
Phaonia consobrina|[2724]|JWDCF337-10|BOLD:AAF6699
Phaonia consobrina|[2725]|MFDC347-09|BOLD:AAF6699
Phaonia consobrina|[2726]|JWDCC382-10|BOLD:AAF6699
Phaonia consobrina|[2727]|JWDCF361-10|BOLD:AAF6699
Phaonia consobrina|[2728]|JWDCF841-10|BOLD:AAF6699
Phaonia consobrina|[2729]|JWDCF860-10|BOLD:AAF6699
Phaonia consobrina|[2730]|MFDC768-10|BOLD:AAF6699
Spilogona trigonata|[2731]|JWDCE704-10|BOLD:AAF3084
Spilogona trigonata|[2732]|JWDCC151-10|BOLD:AAF3084
Spilogona trigonata|[2733]|MFDC046-08|BOLD:AAF3084
Spilogona trigonata|[2734]|MFDC985-10|BOLD:AAF3084
Spilogona trigonata|[2735]|MFDC861-10|BOLD:AAF3084
Spilogona trigonata|[2736]|MFDC860-10|BOLD:AAF3084
Spilogona trigonata|[2737]|MFDC858-10|BOLD:AAF3084
Spilogona trigonata|[2738]|JWDCE230-10|BOLD:AAF3084
Spilogona trigonata|[2739]|JWDCE154-10|BOLD:AAF3084
Spilogona trigonata|[2740]|JWDCF743-10|BOLD:AAF3084
Spilogona trigonata|[2741]|JWDCE471-10|BOLD:AAF3084
Spilogona trigonata|[2742]|JWDCE469-10|BOLD:AAF3084
Spilogona trigonata|[2743]|JWDCF116-10|BOLD:AAF3084
Spilogona trigonata|[2744]|JWDCJ916-11|BOLD:AAF3084
Spilogona trigonata|[2745]|JWDCC791-10|BOLD:AAF3084
Spilogona trigonata|[2746]|JWDCD121-10|BOLD:AAF3084
Spilogona trigonata|[2747]|JWDCD120-10|BOLD:AAF3084
Spilogona trigonata|[2748]|JWDCE092-10|BOLD:AAF3084
Spilogona trigonata|[2749]|JWDCE796-10|BOLD:AAF3084
Spilogona trigonata|[2750]|MFDC864-10|BOLD:AAF3084
Spilogona trigonata|[2751]|MFDC986-10|BOLD:AAF3084
Spilogona trigonata|[2752]|MFDC006-08|BOLD:AAF3084
Spilogona trigonata|[2753]|MFDC987-10|BOLD:AAF3084
Spilogona trigonata|[2754]|MFDC042-08|BOLD:AAF3084
Spilogona opaca|[2755]|JWDCJ1170-11|BOLD:AAI0276
Spilogona opaca|[2756]|JWDCG036-10|BOLD:AAI0276
Spilogona opaca|[2757]|MFDC471-09|BOLD:AAI0276
Spilogona opaca|[2758]|MFDC470-09|BOLD:AAI0276
Spilogona opaca|[2759]|JWDCA438-10|BOLD:AAI0276
Spilogona opaca|[2760]|MFDC837-10|BOLD:AAI0276
Spilogona opaca|[2761]|MFDC1101-11|BOLD:AAI0276
Spilogona sp. 9 AKR|[2762]|JWDCF663-10|BOLD:AAL5981
Spilogona sp. 9 AKR|[2763]|MFDC908-10|BOLD:AAL5981
Spilogona sp. 9 AKR|[2764]|MFDC916-10|BOLD:AAL5981
Spilogona calcaria|[2765]|MFDC906-10|BOLD:ACC0942
Spilogona calcaria|[2766]|MFDC800-10|BOLD:ACA2089
Spilogona flavinervis|[2767]|MFDC819-10|BOLD:ACA2089
Spilogona surda|[2768]|JWDCL127-11|BOLD:AAG1689
Spilogona surda|[2769]|MHFLI193-06|BOLD:AAG1689
Spilogona surda|[2770]|MHFLY388-09|BOLD:AAG1689
Spilogona surda|[2771]|MFDC853-10|BOLD:AAG1689
Spilogona surda|[2772]|MFDC002-08|BOLD:AAG1689
Spilogona surda|[2773]|MFDC852-10|BOLD:AAG1689
Spilogona surda|[2774]|MFDC851-10|BOLD:AAG1689
Spilogona surda|[2775]|MFDC850-10|BOLD:AAG1689
Spilogona surda|[2776]|JWDCF325-10|BOLD:AAG1689
Spilogona surda|[2777]|JWDCI598-10|BOLD:AAG1689
Spilogona surda|[2778]|JWDCL137-11|BOLD:AAG1689
Spilogona surda|[2779]|JWDCL136-11|BOLD:AAG1689
Spilogona surda|[2780]|JWDCL134-11|BOLD:AAG1689
Spilogona surda|[2781]|JWDCK063-11|BOLD:AAG1689
Spilogona surda|[2782]|JWDCJ259-11|BOLD:AAG1689
Spilogona surda|[2783]|JWDCJ208-11|BOLD:AAG1689
Spilogona surda|[2784]|JWDCJ205-11|BOLD:AAG1689
Spilogona surda|[2785]|JWDCL607-11|BOLD:AAG1689
Spilogona surda|[2786]|JWDCL602-11|BOLD:AAG1689
[illegible]

Spilogona surda|[2784]|JWDCJ205-11|BOLD:AAG1689
Spilogona surda|[2785]|JWDCL607-11|BOLD:AAG1689
Spilogona surda|[2786]|JWDCL602-11|BOLD:AAG1689
Spilogona surda|[2787]|JWDCJ798-11|BOLD:AAG1689
Spilogona surda|[2788]|MHFLI632-07|BOLD:AAG1689
Spilogona tornensis|[2789]|MFDC855-10|BOLD:ABX6359
Spilogona depressula|[2790]|MFDC840-10|BOLD:AAL6514
Spilogona depressula|[2791]|MFDC842-10|BOLD:AAL6514
Spilogona depressula|[2792]|MFDC841-10|BOLD:AAL6514
Spilogona depressula|[2793]|MFDC1102-11|BOLD:AAL6514
Spilogona setipes|[2794]|MFDC849-10|BOLD:AAM0157
Spilogona bifimbriata|[2795]|MFDC798-10|BOLD:AAL9640
Spilogona bifimbriata|[2796]|MFDC799-10|BOLD:AAL9640
Spilogona micans|[2797]|MHFLY048-06|BOLD:AAG1686
Spilogona micans|[2798]|MFDC830-10|BOLD:AAG1686
Spilogona micans|[2799]|MFDC828-10|BOLD:AAG1686
Spilogona micans|[2800]|MFDC829-10|BOLD:AAG1686
Spilogona|[2801]|JWDCI346-10|BOLD:AAP9048
Spilogona|[2802]|JWDCH412-10|BOLD:AAP9048
Spilogona quinquelineata|[2803]|MFDC862-10|BOLD:AAL6933
Spilogona sp. 1 AKR|[2804]|MFDC896-10|BOLD:AAL5980
Spilogona sp. 1 AKR|[2805]|MFDC900-10|BOLD:AAL5980
Spilogona sp. 1 AKR|[2806]|MFDC899-10|BOLD:AAL5980
Spilogona sp. 1 AKR|[2807]|MFDC897-10|BOLD:AAL5980
Spilogona sp. 1 AKR|[2808]|MFDC898-10|BOLD:AAL5980
Spilogona sp. 1 AKR|[2809]|MFDC1138-11|BOLD:AAL5980
Spilogona sp. 1 AKR|[2810]|MFDC1137-11|BOLD:AAL5980
Spilogona pusilla|[2811]|JWDCL626-11|BOLD:AAE2792
Spilogona pusilla|[2812]|JWDCJ801-11|BOLD:AAE2792
Spilogona pusilla|[2813]|MFDC894-10|BOLD:AAE2792
Spilogona pusilla|[2814]|MFDC895-10|BOLD:AAE2792
Spilogona pusilla|[2815]|MFDC843-10|BOLD:AAE2792
Spilogona pusilla|[2816]|MFDC844-10|BOLD:AAE2792
Spilogona pusilla|[2817]|JWDCJ811-11|BOLD:AAE2792
Spilogona pusilla|[2818]|JWDCL572-11|BOLD:AAE2792
Spilogona pusilla|[2819]|JWDCJ320-11|BOLD:AAE2792
Spilogona pusilla|[2820]|JWDCJ1184-11|BOLD:AAE2792
Spilogona pusilla|[2821]|JWDCL056-11|BOLD:AAE2792
Spilogona pusilla|[2822]|JWDCL075-11|BOLD:AAE2792
Spilogona pusilla|[2823]|JWDCL151-11|BOLD:AAE2792
Spilogona pusilla|[2824]|JWDCL351-11|BOLD:AAE2792
Spilogona pusilla|[2825]|JWDCI684-11|BOLD:AAE2792
Spilogona pusilla|[2826]|JWDCF521-10|BOLD:AAE2792
Spilogona pusilla|[2827]|MFDC481-09|BOLD:AAE2792
Spilogona pusilla|[2828]|MFDC482-09|BOLD:AAE2792
Spilogona pusilla|[2829]|MFDC498-09|BOLD:AAE2792
Spilogona pusilla|[2830]|MFDC794-10|BOLD:AAE2792
Spilogona pusilla|[2831]|MFDC836-10|BOLD:AAE2792
Spilogona pusilla|[2832]|MFDC483-09|BOLD:AAE2792
Spilogona pusilla|[2833]|MFDC496-09|BOLD:AAE2792
Spilogona pusilla|[2834]|MFDC796-10|BOLD:AAE2792
Spilogona atrisquamula|[2835]|MFDC475-09|BOLD:AAC8383
Spilogona atrisquamula|[2836]|MFDC519-09|BOLD:AAC8383
Spilogona atrisquamula|[2837]|MFDC1089-11|BOLD:AAC8383
Spilogona atrisquamula|[2838]|MHFLI438-07|BOLD:AAC8382
Spilogona atrisquamula|[2839]|MFDC474-09|BOLD:AAC8382
Spilogona atrisquamula|[2840]|MFDC473-09|BOLD:AAC8382
Spilogona atrisquamula|[2841]|MFDC455-09|BOLD:AAC8382
Spilogona atrisquamula|[2842]|MFDC984-10|BOLD:AAC8382
Spilogona atrisquamula|[2843]|JWDCI736-11|BOLD:AAC8382
Spilogona atrisquamula|[2844]|JWDCH550-10|BOLD:AAC8382
Spilogona atrisquamula|[2845]|JWDCH549-10|BOLD:AAC8382
Spilogona atrisquamula|[2846]|JWDCH511-10|BOLD:AAC8382
Spilogona atrisquamula|[2847]|JWDCH510-10|BOLD:AAC8382
Spilogona atrisquamula|[2848]|JWDCH473-10|BOLD:AAC8382
Spilogona atrisquamula|[2849]|JWDCH467-10|BOLD:AAC8382
Spilogona atrisquamula|[2850]|JWDCH405-10|BOLD:AAC8382
Spilogona atrisquamula|[2851]|JWDCG380-10|BOLD:AAC8382
Spilogona atrisquamula|[2852]|JWDCI397-10|BOLD:AAC8382
Spilogona atrisquamula|[2853]|JWDCJ1355-11|BOLD:AAC8382
Spilogona atrisquamula|[2854]|JWDCL196-11|BOLD:AAC8382
Spilogona atrisquamula|[2855]|JWDCJ1347-11|BOLD:AAC8382
Spilogona atrisquamula|[2856]|JWDCJ1182-11|BOLD:AAC8382
Spilogona atrisquamula|[2857]|JWDCK062-11|BOLD:AAC8382
Spilogona atrisquamula|[2858]|JWDCJ210-11|BOLD:AAC8382
Spilogona atrisquamula|[2859]|JWDCJ030-11|BOLD:AAC8382
Spilogona atrisquamula|[2860]|JWDCJ1512-11|BOLD:AAC8382
Spilogona atrisquamula|[2861]|JWDCJ1494-11|BOLD:AAC8382
Spilogona atrisquamula|[2862]|JWDCJ1551-11|BOLD:AAC8382
Spilogona atrisquamula|[2863]|MTDIC207-09|BOLD:AAC8382
Spilogona atrisquamula|[2864]|MHFLY393-09|BOLD:AAC8382
Spilogona atrisquamula|[2865]|MFDC477-09|BOLD:AAC8382
Spilogona atrisquamula|[2866]|MFDC520-09|BOLD:AAC8382
Spilogona atrisquamula|[2867]|MFDC1088-11|BOLD:AAC8382
Spilogona atrisquamula|[2868]|MFDC1086-11|BOLD:AAC8382
Spilogona atrisquamula|[2869]|MFDC1087-11|BOLD:AAC8382
Spilogona atrisquamula|[2870]|MFDC521-09|BOLD:AAC8382
Spilogona atrisquamula|[2871]|MFDC459-09|BOLD:AAC8382
Spilogona atrisquamula|[2872]|MFDC788-10|BOLD:AAC8382
Spilogona atrisquamula|[2873]|MFDC789-10|BOLD:AAC8382
Spilogona atrisquamula|[2874]|MFDC792-10|BOLD:AAC8382
Spilogona atrisquamula|[2875]|MFDC793-10|BOLD:AAC8382
Spilogona atrisquamula|[2876]|MFDC795-10|BOLD:AAC8382
Spilogona confluens|[2877]|MFDC476-09|BOLD:ABY5454
Spilogona confluens|[2878]|MFDC797-10|BOLD:ABY5454
Spilogona sp. 4 AKR|[2879]|JWDCF283-10|BOLD:AAM0158
Spilogona sp. 4 AKR|[2880]|MFDC846-10|BOLD:AAM0158
Spilogona genualis|[2881]|MFDC821-10|BOLD:AAL9641
Spilogona genualis|[2882]|MFDC820-10|BOLD:AAL9641
Spilogona melanosoma|[2883]|MFDC901-10|BOLD:AAF3057
Spilogona melanosoma|[2884]|MFDC517-09|BOLD:AAF3057
Spilogona melanosoma|[2885]|MFDC518-09|BOLD:AAF3057
Spilogona melanosoma|[2886]|MFDC092-08|BOLD:AAF3057

Spilogona melanosoma|[2884]|MFDC517-09|BOLD:AAF3057
Spilogona melanosoma|[2885]|MFDC518-09|BOLD:AAF3057
Spilogona melanosoma|[2886]|MFDC092-08|BOLD:AAF3057
Spilogona malaisei|[2887]|JWDCF119-10|BOLD:AAL9573
Spilogona malaisei|[2888]|JWDCE705-10|BOLD:AAL9573
Spilogona malaisei|[2889]|JWDCI844-11|BOLD:AAL9573
Spilogona malaisei|[2890]|JWDCI849-11|BOLD:AAL9573
Spilogona malaisei|[2891]|JWDCI574-10|BOLD:AAL9573
Spilogona malaisei|[2892]|JWDCI843-11|BOLD:AAL9573
Spilogona malaisei|[2893]|JWDCH039-10|BOLD:AAL9573
Spilogona malaisei|[2894]|JWDCH407-10|BOLD:AAL9573
Spilogona malaisei|[2895]|JWDCE920-10|BOLD:AAL9573
Spilogona malaisei|[2896]|MFDC826-10|BOLD:AAL9573
Spilogona malaisei|[2897]|JWDCK078-11|BOLD:AAL9573
Spilogona malaisei|[2898]|JWDCL197-11|BOLD:AAL9573
Spilogona malaisei|[2899]|JWDCK070-11|BOLD:AAL9573
Spilogona malaisei|[2900]|JWDCJ212-11|BOLD:AAL9573
Spilogona malaisei|[2901]|JWDCJ192-11|BOLD:AAL9573
Spilogona malaisei|[2902]|JWDCJ636-11|BOLD:AAL9573
Spilogona malaisei|[2903]|JWDCJ036-11|BOLD:AAL9573
Spilogona malaisei|[2904]|JWDCL612-11|BOLD:AAL9573
Spilogona malaisei|[2905]|JWDCL601-11|BOLD:AAL9573
Spilogona malaisei|[2906]|JWDCL577-11|BOLD:AAL9573
Spilogona malaisei|[2907]|JWDCJ1552-11|BOLD:AAL9573
Spilogona malaisei|[2908]|JWDCJ547-11|BOLD:AAL9573
Spilogona malaisei|[2909]|JWDCJ182-11|BOLD:AAL9573
Spilogona malaisei|[2910]|MFDC827-10|BOLD:AAL9573
Spilogona sp. 6 AKR|[2911]|JWDCJ1281-11|BOLD:ABZ5879
Spilogona sp. 6 AKR|[2912]|MFDC940-10|BOLD:ABZ5879
Spilogona sp. 3 AKR|[2913]|JWDCA446-10|BOLD:ABZ5880
Spilogona sp. 3 AKR|[2914]|MFDC938-10|BOLD:ABZ5880
Spilogona sp. 3 AKR|[2915]|MFDC480-09|BOLD:ABZ5880
Spilogona griseola|[2916]|JWDCJ315-11|BOLD:AAC0135
Spilogona griseola|[2917]|MFDC923-10|BOLD:AAC0135
Spilogona griseola|[2918]|MFDC931-10|BOLD:AAC0135
Spilogona griseola|[2919]|MFDC960-10|BOLD:AAC0135
Spilogona griseola|[2920]|MFDC983-10|BOLD:AAC0135
Spilogona griseola|[2921]|MFDC465-09|BOLD:AAC0135
Spilogona griseola|[2922]|MFDC464-09|BOLD:AAC0135
Spilogona griseola|[2923]|MFDC463-09|BOLD:AAC0135
Spilogona griseola|[2924]|MFDC462-09|BOLD:AAC0135
Spilogona griseola|[2925]|MFDC458-09|BOLD:AAC0135
Spilogona griseola|[2926]|MFDC456-09|BOLD:AAC0135
Spilogona griseola|[2927]|MFDC055-08|BOLD:AAC0135
Spilogona griseola|[2928]|MFDC468-09|BOLD:AAC0135
Spilogona griseola|[2929]|MFDC467-09|BOLD:AAC0135
Spilogona griseola|[2930]|MFDC466-09|BOLD:AAC0135
Spilogona griseola|[2931]|MFDC461-09|BOLD:AAC0135
Spilogona griseola|[2932]|MFDC457-09|BOLD:AAC0135
Spilogona griseola|[2933]|MFDC963-10|BOLD:AAC0135
Spilogona griseola|[2934]|MFDC958-10|BOLD:AAC0135
Spilogona griseola|[2935]|MFDC957-10|BOLD:AAC0135
Spilogona griseola|[2936]|MFDC956-10|BOLD:AAC0135
Spilogona griseola|[2937]|MFDC955-10|BOLD:AAC0135
Spilogona griseola|[2938]|MFDC954-10|BOLD:AAC0135
Spilogona griseola|[2939]|MFDC953-10|BOLD:AAC0135
Spilogona griseola|[2940]|MFDC952-10|BOLD:AAC0135
Spilogona griseola|[2941]|MFDC951-10|BOLD:AAC0135
Spilogona griseola|[2942]|MFDC950-10|BOLD:AAC0135
Spilogona griseola|[2943]|MFDC943-10|BOLD:AAC0135
Spilogona griseola|[2944]|MFDC941-10|BOLD:AAC0135
Spilogona griseola|[2945]|MFDC939-10|BOLD:AAC0135
Spilogona griseola|[2946]|MFDC937-10|BOLD:AAC0135
Spilogona griseola|[2947]|MFDC936-10|BOLD:AAC0135
Spilogona griseola|[2948]|MFDC935-10|BOLD:AAC0135
Spilogona griseola|[2949]|MFDC934-10|BOLD:AAC0135
Spilogona griseola|[2950]|MFDC930-10|BOLD:AAC0135
Spilogona griseola|[2951]|MFDC927-10|BOLD:AAC0135
Spilogona griseola|[2952]|MFDC926-10|BOLD:AAC0135
Spilogona griseola|[2953]|MFDC925-10|BOLD:AAC0135
Spilogona griseola|[2954]|MFDC922-10|BOLD:AAC0135
Spilogona griseola|[2955]|MFDC921-10|BOLD:AAC0135
Spilogona griseola|[2956]|MFDC920-10|BOLD:AAC0135
Spilogona griseola|[2957]|MFDC919-10|BOLD:AAC0135
Spilogona griseola|[2958]|MFDC892-10|BOLD:AAC0135
Spilogona griseola|[2959]|JWDCF922-10|BOLD:AAC0135
Spilogona griseola|[2960]|JWDCH980-10|BOLD:AAC0135
Spilogona griseola|[2961]|JWDCL355-11|BOLD:AAC0135
Spilogona griseola|[2962]|JWDCL073-11|BOLD:AAC0135
Spilogona griseola|[2963]|JWDCA463-10|BOLD:AAC0135
Spilogona griseola|[2964]|MFDC928-10|BOLD:AAC0135
Spilogona griseola|[2965]|MFDC929-10|BOLD:AAC0135
Spilogona griseola|[2966]|MFDC949-10|BOLD:AAC0135
Spilogona griseola|[2967]|MFDC946-10|BOLD:AAC0135
Spilogona griseola|[2968]|MFDC948-10|BOLD:AAC0135
Spilogona griseola|[2969]|MFDC947-10|BOLD:AAC0135
Spilogona griseola|[2970]|MFDC942-10|BOLD:AAC0135
Spilogona griseola|[2971]|MFDC959-10|BOLD:AAC0135
Spilogona griseola|[2972]|MFDC961-10|BOLD:AAC0135
Spilogona griseola|[2973]|MFDC962-10|BOLD:AAC0135
Spilogona griseola|[2974]|MFDC469-09|BOLD:AAC0135
Spilogona griseola|[2975]|MFDC460-09|BOLD:AAC0135
Spilogona griseola|[2976]|MFDC822-10|BOLD:AAC0135
Spilogona sp. 10 AKR|[2977]|MFDC965-10|BOLD:AAL9121
Spilogona sp. 10 AKR|[2978]|MFDC966-10|BOLD:AAL9121
Spilogona sp. 10 AKR|[2979]|MFDC971-10|BOLD:AAL9121
Spilogona incerta|[2980]|MFDC823-10|BOLD:AAM0159
Spilogona setinervis|[2981]|JWDCB130-10|BOLD:AAG1785
Spilogona trianguligera|[2982]|MFDC847-10|BOLD:AAG1785
Spilogona trianguligera|[2983]|MFDC845-10|BOLD:AAG1785
Spilogona trianguligera|[2984]|MFDC838-10|BOLD:AAG1785
Spilogona trianguligera|[2985]|MFDC848-10|BOLD:AAG1785
[illegible]

Spilogona trianguligera|[2983]|MFDC843-10|BOLD:AAG1785
Spilogona trianguligera|[2984]|MFDC838-10|BOLD:AAG1785
Spilogona trianguligera|[2985]|MFDC848-10|BOLD:AAG1785
Spilogona zaitzevi|[2986]|MFDC902-10|BOLD:AAL9576
Spilogona zaitzevi|[2987]|MFDC825-10|BOLD:AAL9576
Spilogona imitatrix|[2988]|JWDCJ870-11|BOLD:AAD8451
Spilogona imitatrix|[2989]|JWDCJ387-11|BOLD:AAD8451
Spilogona imitatrix|[2990]|JWDCC185-10|BOLD:AAD8451
Spilogona imitatrix|[2991]|JWDCJ911-11|BOLD:AAD8451
Spilogona imitatrix|[2992]|JWDCE936-10|BOLD:AAD8451
Spilogona imitatrix|[2993]|MFDC488-09|BOLD:AAD8451
Spilogona imitatrix|[2994]|MFDC489-09|BOLD:AAD8451
Spilogona imitatrix|[2995]|MFDC499-09|BOLD:AAD8451
Spilogona imitatrix|[2996]|MFDC487-09|BOLD:AAD8451
Spilogona imitatrix|[2997]|MFDC500-09|BOLD:AAD8451
Spilogona sororcula|[2998]|MHFLI134-06|BOLD:AAD8449
Spilogona sororcula|[2999]|JWDCJ1291-11|BOLD:AAD8449
Spilogona sororcula|[3000]|MFDC029-08|BOLD:AAD8449
Spilogona sororcula|[3001]|MFDC492-09|BOLD:AAD8449
Spilogona sororcula|[3002]|JWDCF570-10|BOLD:AAD8449
Spilogona sororcula|[3003]|MFDC504-09|BOLD:AAD8449
Spilogona sororcula|[3004]|JWDCG094-10|BOLD:AAD8449
Spilogona sororcula|[3005]|JWDCI193-10|BOLD:AAD8449
Spilogona sororcula|[3006]|JWDCK131-11|BOLD:AAD8449
Spilogona sororcula|[3007]|JWDCG091-10|BOLD:AAD8449
Spilogona sororcula|[3008]|JWDCJ1297-11|BOLD:AAD8449
Spilogona sororcula|[3009]|JWDCJ1290-11|BOLD:AAD8449
Spilogona sororcula|[3010]|JWDCJ1287-11|BOLD:AAD8449
Spilogona sororcula|[3011]|JWDCJ1070-11|BOLD:AAD8449
Spilogona sororcula|[3012]|JWDCJ1069-11|BOLD:AAD8449
Spilogona sororcula|[3013]|JWDCJ1066-11|BOLD:AAD8449
Spilogona sororcula|[3014]|JWDCJ920-11|BOLD:AAD8449
Spilogona sororcula|[3015]|JWDCK144-11|BOLD:AAD8449
Spilogona sororcula|[3016]|JWDCK141-11|BOLD:AAD8449
Spilogona sororcula|[3017]|JWDCK138-11|BOLD:AAD8449
Spilogona sororcula|[3018]|JWDCJ1073-11|BOLD:AAD8449
Spilogona sororcula|[3019]|JWDCJ1289-11|BOLD:AAD8449
Spilogona sororcula|[3020]|JWDCE890-10|BOLD:AAD8449
Spilogona sororcula|[3021]|MFDC493-09|BOLD:AAD8449
Spilogona sororcula|[3022]|MFDC505-09|BOLD:AAD8449
Spilogona imitatrix|[3023]|JWDCD023-10|BOLD:AAG1688
Spilogona imitatrix|[3024]|JWDCL608-11|BOLD:AAG1688
Spilogona imitatrix|[3025]|MHFLI131-06|BOLD:AAG1688
Spilogona pacifica|[3026]|JWDCL076-11|BOLD:AAD8448
Spilogona pacifica|[3027]|JWDCB139-10|BOLD:AAD8448
Spilogona pacifica|[3028]|JWDCH877-10|BOLD:AAD8448
Spilogona pacifica|[3029]|JWDCH304-10|BOLD:AAD8448
Spilogona pacifica|[3030]|MFDC044-08|BOLD:AAD8448
Spilogona pacifica|[3031]|MFDC503-09|BOLD:AAD8448
Spilogona pacifica|[3032]|MFDC043-08|BOLD:AAD8448
Spilogona pacifica|[3033]|JWDCI103-10|BOLD:AAD8448
Spilogona pacifica|[3034]|MFDC911-10|BOLD:AAD8448
Spilogona pacifica|[3035]|MFDC910-10|BOLD:AAD8448
Spilogona pacifica|[3036]|MFDC909-10|BOLD:AAD8448
Spilogona pacifica|[3037]|JWDCF919-10|BOLD:AAD8448
Spilogona pacifica|[3038]|JWDCH722-10|BOLD:AAD8448
Spilogona pacifica|[3039]|JWDCK277-11|BOLD:AAD8448
Spilogona pacifica|[3040]|JWDCK274-11|BOLD:AAD8448
Spilogona pacifica|[3041]|MFDC912-10|BOLD:AAD8448
Spilogona pacifica|[3042]|MFDC078-08|BOLD:AAD8448
Spilogona pacifica|[3043]|MFDC497-09|BOLD:AAD8448
Pseudocoenosia fletcheri|[3044]|MFDC775-10|BOLD:AAF6899
Pseudocoenosia fletcheri|[3045]|MFDC376-09|BOLD:AAF6899
Pseudocoenosia fletcheri|[3046]|MFDC004-08|BOLD:AAF6899
Pseudocoenosia fletcheri|[3047]|MFDC005-08|BOLD:AAF6899
Pseudocoenosia fletcheri|[3048]|MFDC774-10|BOLD:AAF6899
Pseudocoenosia fletcheri|[3049]|MFDC776-10|BOLD:AAF6899
Spilogona aenea|[3050]|JWDCB141-10|BOLD:ACX5124
Spilogona aenea|[3051]|MFDC917-10|BOLD:ACX5124
Spilogona aenea|[3052]|JWDCB143-10|BOLD:ACX5124
Spilogona aenea|[3053]|JWDCB127-10|BOLD:ACX5124
Spilogona aenea|[3054]|JWDCH717-10|BOLD:ACX5124
Spilogona aenea|[3055]|MFDC037-08|BOLD:ACX5124
Spilogona aenea|[3056]|MFDC787-10|BOLD:ACX5124
Spilogona aenea|[3057]|MFDC784-10|BOLD:ACX5124
Spilogona aenea|[3058]|MFDC785-10|BOLD:ACX5124
Spilogona aenea|[3059]|MFDC786-10|BOLD:ACX5124
Spilogona|[3060]|JWDCG232-10|BOLD:ABZ0955
Spilogona sp. 12AKR|[3061]|MFDC944-10|BOLD:ABZ0955
Spilogona sp. 12AKR|[3062]|MFDC366-09|BOLD:ABZ0955
Spilogona sp. 12AKR|[3063]|MFDC449-09|BOLD:ABZ0955
Spilogona|[3064]|JWDCB110-10|BOLD:ABZ0955
Spilogona|[3065]|JWDCB103-10|BOLD:ABZ0955
Spilogona|[3066]|JWDCB136-10|BOLD:ABZ0955
Spilogona|[3067]|JWDCA411-10|BOLD:ABZ0955
Spilogona|[3068]|JWDCA403-10|BOLD:ABZ0955
Spilogona|[3069]|JWDCA491-10|BOLD:ABZ0955
Spilogona|[3070]|JWDCH619-10|BOLD:ABZ0955
Spilogona|[3071]|JWDCH161-10|BOLD:ABZ0955
Spilogona|[3072]|JWDCH168-10|BOLD:ABZ0955
Spilogona|[3073]|JWDCG284-10|BOLD:ABZ0955
Spilogona forticula|[3074]|MFDC903-10|BOLD:ABZ0955
Spilogona forticula|[3075]|MFDC905-10|BOLD:ABZ0955
Spilogona forticula|[3076]|MFDC369-09|BOLD:ABZ0955
Spilogona forticula|[3077]|MFDC372-09|BOLD:ABZ0955
Spilogona forticula|[3078]|MFDC479-09|BOLD:ABZ0955
Spilogona forticula|[3079]|MFDC051-08|BOLD:ABZ0955
Spilogona forticula|[3080]|MFDC988-10|BOLD:ABZ0955
Spilogona forticula|[3081]|MFDC023-08|BOLD:ABZ0955
Spilogona forticula|[3082]|MFDC478-09|BOLD:ABZ0955
Spilogona suspecta|[3083]|JWDCA468-10|BOLD:AAC4649
Spilogona suspecta|[3084]|MHFLY395-09|BOLD:AAC4649
Spilogona suspecta|[3085]|MHFLI136-06|BOLD:AAC4649

Spilogona suspecta|[3083]|JWDCA468-10|BOLD:AAC4649
Spilogona suspecta|[3084]|MHFLY395-09|BOLD:AAC4649
Spilogona suspecta|[3085]|MHFLI136-06|BOLD:AAC4649
Spilogona suspecta|[3086]|MFDC485-09|BOLD:AAC4649
Spilogona suspecta|[3087]|MFDC472-09|BOLD:AAC4649
Spilogona suspecta|[3088]|MFDC059-08|BOLD:AAC4649
Spilogona suspecta|[3089]|MHFLI927-09|BOLD:AAC4649
Spilogona suspecta|[3090]|JWDCH066-10|BOLD:AAC4649
Spilogona suspecta|[3091]|MFDC486-09|BOLD:AAC4649
Spilogona suspecta|[3092]|MFDC484-09|BOLD:AAC4649
Spilogona suspecta|[3093]|MHFLY355-09|BOLD:AAC4649
Spilogona suspecta|[3094]|MHFLY132-07|BOLD:AAC4649
Spilogona suspecta|[3095]|MHFLY394-09|BOLD:AAC4649
Spilogona arctica|[3096]|JWDCC386-10|BOLD:ACF0384
Spilogona arctica|[3097]|JWDCI540-10|BOLD:ACF0384
Spilogona arctica|[3098]|MFDC090-08|BOLD:ACF0384
Spilogona arctica|[3099]|MFDC072-08|BOLD:ACF0384
Spilogona arctica|[3100]|MFDC440-09|BOLD:ACF0384
Spilogona arctica|[3101]|MFDC373-09|BOLD:ACF0384
Spilogona arctica|[3102]|JWDCG289-10|BOLD:ACF0384
Spilogona arctica|[3103]|MFDC945-10|BOLD:ACF0384
Spilogona arctica|[3104]|MFDC367-09|BOLD:ACF0384
Spilogona arctica|[3105]|JWDCG867-10|BOLD:ACF0384
Spilogona arctica|[3106]|MFDC904-10|BOLD:ACF0384
Spilogona arctica|[3107]|MFDC914-10|BOLD:ACF0384
Spilogona arctica|[3108]|MFDC918-10|BOLD:ACF0384
Spilogona arctica|[3109]|MFDC924-10|BOLD:ACF0384
Spilogona arctica|[3110]|JWDCG495-10|BOLD:ACF0384
Spilogona arctica|[3111]|JWDCG728-10|BOLD:ACF0384
Spilogona arctica|[3112]|JWDCI742-11|BOLD:ACF0384
Spilogona arctica|[3113]|JWDCI917-11|BOLD:ACF0384
Spilogona arctica|[3114]|MFDC365-09|BOLD:ACF0384
Spilogona arctica|[3115]|MFDC017-08|BOLD:ACF0384
Spilogona arctica|[3116]|MFDC451-09|BOLD:ACF0384
Spilogona arctica|[3117]|MFDC452-09|BOLD:ACF0384
Spilogona arctica|[3118]|MFDC436-09|BOLD:ACF0384
Spilogona arctica|[3119]|MFDC025-08|BOLD:ACF0384
Spilogona|[3120]|JWDCF769-10|BOLD:AAB5278
Spilogona|[3121]|JWDCB128-10|BOLD:AAB5278
Spilogona|[3122]|JWDCI056-10|BOLD:AAB5278
Spilogona|[3123]|JWDCG282-10|BOLD:AAB5278
Spilogona|[3124]|JWDCK080-11|BOLD:AAB5278
Spilogona contractifrons|[3125]|MFDC915-10|BOLD:AAB5278
Spilogona|[3126]|JWDCG631-10|BOLD:AAB5278
Spilogona|[3127]|JWDCH983-10|BOLD:AAB5278
Spilogona|[3128]|JWDCL625-11|BOLD:AAB5278
Spilogona|[3129]|JWDCL214-11|BOLD:AAB5278
Spilogona|[3130]|JWDCL069-11|BOLD:AAB5278
Spilogona|[3131]|JWDCL618-11|BOLD:AAB5278
Spilogona|[3132]|JWDCJ202-11|BOLD:AAB5278
Spilogona|[3133]|JWDCJ266-11|BOLD:AAB5278
Spilogona contractifrons|[3134]|MFDC438-09|BOLD:AAB5278
Spilogona contractifrons|[3135]|MFDC442-09|BOLD:AAB5278
Spilogona contractifrons|[3136]|MFDC443-09|BOLD:AAB5278
Spilogona contractifrons|[3137]|MFDC446-09|BOLD:AAB5278
Spilogona contractifrons|[3138]|MFDC448-09|BOLD:AAB5278
Spilogona contractifrons|[3139]|MFDC450-09|BOLD:AAB5278
Spilogona contractifrons|[3140]|MFDC030-08|BOLD:AAB5278
Spilogona contractifrons|[3141]|MFDC089-08|BOLD:AAB5278
Spilogona contractifrons|[3142]|MFDC053-08|BOLD:AAB5278
Spilogona contractifrons|[3143]|MFDC439-09|BOLD:AAB5278
Spilogona contractifrons|[3144]|MFDC441-09|BOLD:AAB5278
Spilogona contractifrons|[3145]|MFDC010-08|BOLD:AAB5278
Spilogona contractifrons|[3146]|MFDC447-09|BOLD:AAB5278
Spilogona contractifrons|[3147]|MFDC027-08|BOLD:AAB5278
Spilogona|[3148]|JWDCJ032-11|BOLD:AAB5278
Spilogona|[3149]|JWDCJ200-11|BOLD:AAB5278
Spilogona contractifrons|[3150]|MFDC371-09|BOLD:AAB5278
Spilogona contractifrons|[3151]|MFDC437-09|BOLD:AAB5278
Spilogona contractifrons|[3152]|MFDC859-10|BOLD:AAB5278
Spilogona contractifrons|[3153]|MFDC932-10|BOLD:AAB5278
Spilogona|[3154]|JWDCH509-10|BOLD:AAB5278
Spilogona|[3155]|JWDCF281-10|BOLD:AAB5278
Spilogona|[3156]|JWDCG006-10|BOLD:AAB5278
Spilogona|[3157]|JWDCH357-10|BOLD:AAB5278
Spilogona|[3158]|JWDCA429-10|BOLD:AAB5278
Spilogona|[3159]|JWDCA423-10|BOLD:AAB5278
Spilogona|[3160]|JWDCA405-10|BOLD:AAB5278
Spilogona|[3161]|JWDCB129-10|BOLD:AAB5278
Spilogona|[3162]|JWDCA440-10|BOLD:AAB5278
Spilogona|[3163]|JWDCA452-10|BOLD:AAB5278
Spilogona|[3164]|JWDCJ1560-11|BOLD:AAB5278
Spilogona|[3165]|JWDCL576-11|BOLD:AAB5278
Spilogona|[3166]|JWDCL616-11|BOLD:AAB5278
Spilogona|[3167]|JWDCJ027-11|BOLD:AAB5278
Spilogona contractifrons|[3168]|MFDC933-10|BOLD:AAB5278
Spilogona contractifrons|[3169]|MFDC370-09|BOLD:AAB5278
Spilogona contractifrons|[3170]|MFDC022-08|BOLD:AAB5278
Spilogona contractifrons|[3171]|MFDC062-08|BOLD:AAB5278
Spilogona|[3172]|MHFLI118-06|BOLD:AAB5278
Spilogona sp. 7 AKR|[3173]|JWDCC214-10|BOLD:AAL5982
Spilogona sp. 7 AKR|[3174]|MFDC913-10|BOLD:AAL5982
Spilogona churchillensis|[3175]|MHFLI617-07|BOLD:AAB7605
Spilogona churchillensis|[3176]|MHFLI620-07|BOLD:AAB7605
Spilogona churchillensis|[3177]|MHFLI623-07|BOLD:AAB7605
Spilogona churchillensis|[3178]|MHFLI667-07|BOLD:AAB7605
Spilogona churchillensis|[3179]|MHFLI619-07|BOLD:AAB7605
Spilogona churchillensis|[3180]|MHFLI618-07|BOLD:AAB7605
Spilogona churchillensis|[3181]|MFDC515-09|BOLD:AAB7605
Spilogona churchillensis|[3182]|MHFLI969-09|BOLD:AAB7605
Spilogona churchillensis|[3183]|MHFLI954-09|BOLD:AAB7605
Spilogona churchillensis|[3184]|MHFLI951-09|BOLD:AAB7605
[illegible]

Spilogona churchillensis|[3182]|MHFLI969-09|BOLD:AAB7605
Spilogona churchillensis|[3183]|MHFLI954-09|BOLD:AAB7605
Spilogona churchillensis|[3184]|MHFLI951-09|BOLD:AAB7605
Spilogona churchillensis|[3185]|MHFLI949-09|BOLD:AAB7605
Spilogona churchillensis|[3186]|MHFLI948-09|BOLD:AAB7605
Spilogona churchillensis|[3187]|MHFLI945-09|BOLD:AAB7605
Spilogona churchillensis|[3188]|MHFLI896-09|BOLD:AAB7605
Spilogona churchillensis|[3189]|MFDC516-09|BOLD:AAB7605
Spilogona churchillensis|[3190]|MHFLI971-09|BOLD:AAB7605
Spilogona churchillensis|[3191]|MHFLI621-07|BOLD:AAB7605
Spilogona churchillensis|[3192]|MHFLI636-07|BOLD:AAB7605
Spilogona sp. 2 AKR|[3193]|JWDCG090-10|BOLD:AAJ3564
Spilogona sp. 2 AKR|[3194]|MFDC368-09|BOLD:AAJ3564
Spilogona trilineata|[3195]|JWDCH100-10|BOLD:AAG1685
Spilogona trilineata|[3196]|JWDCJ1492-11|BOLD:AAG1685
Spilogona trilineata|[3197]|MFDC865-10|BOLD:AAG1685
Spilogona trilineata|[3198]|MFDC863-10|BOLD:AAG1685
Spilogona trilineata|[3199]|MFDC866-10|BOLD:AAG1685
Spilogona trilineata|[3200]|MFDC502-09|BOLD:AAG1685
Spilogona trilineata|[3201]|MHFLY014-06|BOLD:AAG1685
Spilogona infuscata|[3202]|MFDC803-10|BOLD:AAL9574
Spilogona infuscata|[3203]|MFDC824-10|BOLD:AAL9574
Spilogona narina|[3204]|MFDC831-10|BOLD:AAG3155
Spilogona narina|[3205]|MFDC832-10|BOLD:AAG3155
Spilogona|[3206]|MHFLI648-07|BOLD:AAG1693
Spilogona leucogaster|[3207]|MHFLY023-06|BOLD:AAE2782
Spilogona leucogaster|[3208]|MFDC511-09|BOLD:AAE2782
Spilogona leucogaster|[3209]|MFDC513-09|BOLD:AAE2782
Spilogona leucogaster|[3210]|MFDC514-09|BOLD:AAE2782
Spilogona leucogaster|[3211]|MFDC512-09|BOLD:AAE2782
Spilogona deflorata|[3212]|MFDC893-10|BOLD:AAG1692
Spilogona deflorata|[3213]|MFDC808-10|BOLD:AAG1692
Spilogona deflorata|[3214]|MFDC809-10|BOLD:AAG1692
Spilogona deflorata|[3215]|MFDC989-10|BOLD:AAG1692
Spilogona deflorata|[3216]|MHFLI613-07|BOLD:AAG1692
Spilogona deflorata|[3217]|MHFLI453-07|BOLD:AAG1692
Spilogona deflorata|[3218]|MFDC806-10|BOLD:AAG1692
Spilogona deflorata|[3219]|MFDC807-10|BOLD:AAG1692
Spilogona obscuripennis|[3220]|MFDC833-10|BOLD:ACZ6405
Spilogona obscuripennis|[3221]|MFDC805-10|BOLD:ACZ6405
Spilogona obscuripennis|[3222]|MFDC804-10|BOLD:ACZ6405
Spilogona obscuripennis|[3223]|MHFLY401-09|BOLD:ACZ6405
Spilogona obscuripennis|[3224]|JWDCJ1498-11|BOLD:ACZ6405
Spilogona obscuripennis|[3225]|MFDC834-10|BOLD:ACZ6405
Spilogona obscuripennis|[3226]|MFDC835-10|BOLD:ACZ6405
Spilogona aerea|[3227]|MFDC507-09|BOLD:AAA7731
Spilogona aerea|[3228]|MHFLI676-07|BOLD:AAA7731
Spilogona aerea|[3229]|MHFLI656-07|BOLD:AAA7731
Spilogona aerea|[3230]|MHFLI685-07|BOLD:AAA7731
Spilogona aerea|[3231]|MHFLI391-07|BOLD:AAA7731
Spilogona aerea|[3232]|MHFLI605-07|BOLD:AAA7731
Spilogona aerea|[3233]|MHFLI660-07|BOLD:AAA7731
Spilogona aerea|[3234]|MHFLI595-07|BOLD:AAA7731
Spilogona aerea|[3235]|MHFLI611-07|BOLD:AAA7731
Spilogona aerea|[3236]|MHFLI606-07|BOLD:AAA7731
Spilogona aerea|[3237]|MHFLI597-07|BOLD:AAA7731
Spilogona aerea|[3238]|MHFLI609-07|BOLD:AAA7731
Spilogona aerea|[3239]|MHFLI684-07|BOLD:AAA7731
Spilogona aerea|[3240]|MHFLI662-07|BOLD:AAA7731
Spilogona aerea|[3241]|MFDC506-09|BOLD:AAA7731
Spilogona aerea|[3242]|MHFLI890-09|BOLD:AAA7731
Spilogona aerea|[3243]|MHFLI961-09|BOLD:AAA7731
Spilogona aerea|[3244]|MHFLI964-09|BOLD:AAA7731
Spilogona aerea|[3245]|MFDC052-08|BOLD:AAA7731
Spilogona aerea|[3246]|MFDC495-09|BOLD:AAA7731
Spilogona aerea|[3247]|MHFLI959-09|BOLD:AAA7731
Spilogona aerea|[3248]|MHFLI960-09|BOLD:AAA7731
Spilogona aerea|[3249]|MHFLI946-09|BOLD:AAA7731
Spilogona aerea|[3250]|MHFLI947-09|BOLD:AAA7731
Spilogona aerea|[3251]|MHFLI942-09|BOLD:AAA7731
Spilogona aerea|[3252]|MHFLI943-09|BOLD:AAA7731
Spilogona aerea|[3253]|MHFLI938-09|BOLD:AAA7731
Spilogona aerea|[3254]|MHFLI939-09|BOLD:AAA7731
Spilogona aerea|[3255]|JWDCH344-10|BOLD:AAA7731
Spilogona aerea|[3256]|JWDCL632-11|BOLD:AAA7731
Spilogona aerea|[3257]|JWDCL617-11|BOLD:AAA7731
Spilogona aerea|[3258]|MHFLI651-07|BOLD:AAA7731
Spilogona aerea|[3259]|MHFLI669-07|BOLD:AAA7731
Spilogona aerea|[3260]|MHFLI682-07|BOLD:AAA7731
Spilogona aerea|[3261]|MHFLI424-07|BOLD:AAA7731
Spilogona aerea|[3262]|MHFLI683-07|BOLD:AAA7731
Spilogona aerea|[3263]|MHFLI659-07|BOLD:AAA7731
Spilogona aerea|[3264]|MHFLI600-07|BOLD:AAA7731
Spilogona aerea|[3265]|MHFLI596-07|BOLD:AAA7731
Spilogona aerea|[3266]|MHFLI677-07|BOLD:AAA7731
Spilogona aerea|[3267]|MHFLI966-09|BOLD:AAA7731
Spilogona aerea|[3268]|MHFLI466-07|BOLD:AAA7731
Spilogona aerea|[3269]|MHFLI610-07|BOLD:AAA7731
Spilogona aerea|[3270]|MHFLI403-07|BOLD:AAA7731
Spilogona aerea|[3271]|MHFLI594-07|BOLD:AAA7731
Spilogona aerea|[3272]|MHFLI601-07|BOLD:AAA7731
Spilogona aerea|[3273]|MHFLI603-07|BOLD:AAA7731
Spilogona aerea|[3274]|MHFLI655-07|BOLD:AAA7731
Spilogona aerea|[3275]|MHFLI665-07|BOLD:AAA7731
Spilogona aerea|[3276]|MHFLI670-07|BOLD:AAA7731
Tachinidae|[3277]|TWDIP241-09|BOLD:AAG2109
Tachinidae|[3278]|TWDIP242-09|BOLD:AAG2109
Tachinidae|[3279]|JWDCJ1451-11|BOLD:AAG2109
Tachinidae|[3280]|JWDCJ1452-11|BOLD:AAG2109
Tachinidae|[3281]|JWDCI154-10|BOLD:AAG2109
Tachinidae|[3282]|MHFLI177-06|BOLD:AAG2109
Tachinidae|[3283]|MHFLI171-06|BOLD:AAG2109
Tachina marklini|[3284]|JWDCC178-10|BOLD:ABY5409

Tachinidae|[3282]|MHFLI177-06|BOLD:AAG2109
Tachinidae|[3283]|MHFLI171-06|BOLD:AAG2109
Tachina marklini|[3284]|JWDCC178-10|BOLD:ABY5409
Tachina ampliforceps|[3285]|MHFLI172-06|BOLD:AAG2110
Tachina ampliforceps|[3286]|JWDCG803-10|BOLD:AAG2110
Muscina sp. 1 AKR|[3287]|JWDCF203-10|BOLD:AAB8818
Muscina sp. 1 AKR|[3288]|MFDC1096-11|BOLD:AAB8818
Muscina flukei|[3289]|MHFLY362-09|BOLD:ABZ4354
Muscina levida|[3290]|MHFLI168-06|BOLD:AAB8817
Muscina levida|[3291]|MFDC633-09|BOLD:AAB8817
Muscina levida|[3292]|MFDC635-09|BOLD:AAB8817
Muscina levida|[3293]|MFDC048-08|BOLD:AAB8817
Muscina levida|[3294]|MFDC076-08|BOLD:AAB8817
Muscina levida|[3295]|JWDCE039-10|BOLD:AAB8817
Muscina levida|[3296]|JWDCK051-11|BOLD:AAB8817
Muscina levida|[3297]|MHFLI187-06|BOLD:AAB8817
Muscina levida|[3298]|MFDC630-09|BOLD:AAB8817
Muscina levida|[3299]|MFDC631-09|BOLD:AAB8817
Muscina levida|[3300]|MFDC634-09|BOLD:AAB8817
Muscina levida|[3301]|MHFLY363-09|BOLD:AAB8817
Muscina levida|[3302]|MFDC753-10|BOLD:AAB8817
Muscina levida|[3303]|MFDC754-10|BOLD:AAB8817
Anthomyiidae|[3304]|JWDCC798-10|BOLD:AAL7524
Actia interrupta|[3305]|JWDCH616-10|BOLD:AAB8437
Actia interrupta|[3306]|JWDCH614-10|BOLD:AAB8437
Actia interrupta|[3307]|TWDIP203-09|BOLD:AAB8437
Actia interrupta|[3308]|JWDCB104-10|BOLD:AAB8437
Actia interrupta|[3309]|TWDIP272-09|BOLD:AAB8437
Actia interrupta|[3310]|JWDCH618-10|BOLD:AAB8437
Actia interrupta|[3311]|JWDCH620-10|BOLD:AAB8437
Actia diffidens|[3312]|JWDCF185-10|BOLD:AAG2354
Blaesoxipha kyrtonidion|[3313]|JWDCJ1059-11|BOLD:AAV1091
Blaesoxipha kyrtonidion|[3314]|JWDCJ1060-11|BOLD:AAV1091
Blaesoxipha|[3315]|JWDCG218-10|BOLD:AAP7703
Sarcophaga aldrichi|[3316]|JWDCC803-10|BOLD:AAH0007
Sarcophagidae|[3317]|JWDCG387-10|BOLD:AAP8791
Sarcophaga sinuata|[3318]|JWDCA528-10|BOLD:AAG3629
Sarcophaga sinuata|[3319]|JWDCG738-10|BOLD:AAG3629
Sarcophaga sinuata|[3320]|JWDCF204-10|BOLD:AAG3629
Sarcophaga sinuata|[3321]|JWDCF317-10|BOLD:AAG3629
Sarcophaga|[3322]|JWDCG600-10|BOLD:AAH0008
Sarcophaga|[3323]|JWDCG840-10|BOLD:AAH0008
Sarcophaga|[3324]|JWDCG217-10|BOLD:AAH0008
Sarcophaga|[3325]|JWDCH135-10|BOLD:AAH0008
Sarcophaga sexpunctata|[3326]|MTDIC155-09|BOLD:AAH7150
Sarcophaga sexpunctata|[3327]|JWDCA497-10|BOLD:AAH7150
Sarcophaga sexpunctata|[3328]|JWDCG783-10|BOLD:AAH7150
Sarcophaga sexpunctata|[3329]|JWDCI644-10|BOLD:AAH7150
Sarcophaginae|[3330]|JWDCI153-10|BOLD:AAG2430
Sarcophaginae|[3331]|MTDIC103-09|BOLD:AAG2430
Sarcophaginae|[3332]|JWDCB096-10|BOLD:AAG2430
Sarcophaginae|[3333]|JWDCB067-10|BOLD:AAG2430
Sarcophaginae|[3334]|JWDCB061-10|BOLD:AAG2430
Sarcophaginae|[3335]|JWDCB060-10|BOLD:AAG2430
Sarcophaginae|[3336]|JWDCB054-10|BOLD:AAG2430
Sarcophaginae|[3337]|MTDIC191-09|BOLD:AAG2430
Sarcophaginae|[3338]|MTDIC192-09|BOLD:AAG2430
Sarcophaginae|[3339]|JWDCJ1409-11|BOLD:AAG2430
Sarcophaginae|[3340]|JWDCG784-10|BOLD:AAG2430
Sarcophaginae|[3341]|JWDCG869-10|BOLD:AAG2430
Sarcophaginae|[3342]|JWDCF160-10|BOLD:AAG2430
Sarcophaga libera|[3343]|JWDCI022-10|BOLD:AAP8790
Sarcophaga libera|[3344]|JWDCF844-10|BOLD:AAP8790
Strongygaster|[3345]|JWDCK052-11|BOLD:AAV0905
Strongygaster celer|[3346]|JWDCJ168-11|BOLD:AAV0905
Solieria piperi|[3347]|JWDCJ1413-11|BOLD:AAV0932
Besseria anthophila|[3348]|MTDIC265-09|BOLD:AAG6981
Besseria anthophila|[3349]|JWDCG788-10|BOLD:AAG6981
Voria|[3350]|JWDCA569-10|BOLD:AAG2358
Voria|[3351]|JWDCA508-10|BOLD:AAG2358
Voria|[3352]|JWDCJ1266-11|BOLD:AAG2358
Voria|[3353]|JWDCJ1410-11|BOLD:AAG2358
Voria|[3354]|JWDCJ1412-11|BOLD:AAG2358
Voria|[3355]|JWDCJ1414-11|BOLD:AAG2358
Voria|[3356]|JWDCG633-10|BOLD:AAG2358
Voria|[3357]|JWDCH883-10|BOLD:AAG2358
Voria|[3358]|JWDCG231-10|BOLD:AAG2358
Acemya oestriforme|[3359]|JWDCI345-10|BOLD:AAG2132
Eurithia|[3360]|MHFLI178-06|BOLD:AAG2113
Eurithia|[3361]|JWDCH790-10|BOLD:AAG2113
Eurithia|[3362]|JWDCG388-10|BOLD:AAG2113
Lydina americana|[3363]|JWDCB056-10|BOLD:AAG2432
Admontia|[3364]|MHFLI267-06|BOLD:AAG2114
Periscepsia laevigata|[3365]|JWDCC858-10|BOLD:AAG2175
Periscepsia laevigata|[3366]|TWDIP262-09|BOLD:AAG2175
Periscepsia laevigata|[3367]|JWDCA559-10|BOLD:AAG2175
Periscepsia laevigata|[3368]|JWDCA567-10|BOLD:AAG2175
Periscepsia|[3369]|JWDCJ1072-11|BOLD:AAV0917
Blondelia eufitchiae|[3370]|JWDCI434-10|BOLD:AAG2346
Platymya confusionis|[3371]|MHFLI909-09|BOLD:AAG2162
Platymya confusionis|[3372]|MHFLI911-09|BOLD:AAG2162
Platymya confusionis|[3373]|MHFLI913-09|BOLD:AAG2162
Platymya confusionis|[3374]|MHFLI914-09|BOLD:AAG2162
Platymya confusionis|[3375]|MHFLI930-09|BOLD:AAG2162
Platymya confusionis|[3376]|JWDCA495-10|BOLD:AAG2162
Platymya confusionis|[3377]|JWDCA498-10|BOLD:AAG2162
Platymya confusionis|[3378]|JWDCA499-10|BOLD:AAG2162
Platymya confusionis|[3379]|JWDCA502-10|BOLD:AAG2162
Platymya confusionis|[3380]|JWDCG390-10|BOLD:AAG2162
Chetogena|[3381]|JWDCA496-10|BOLD:AAG2431
Chetogena|[3382]|JWDCA504-10|BOLD:AAG2431
Chetogena|[3383]|JWDCA529-10|BOLD:AAG2431
[illegible]

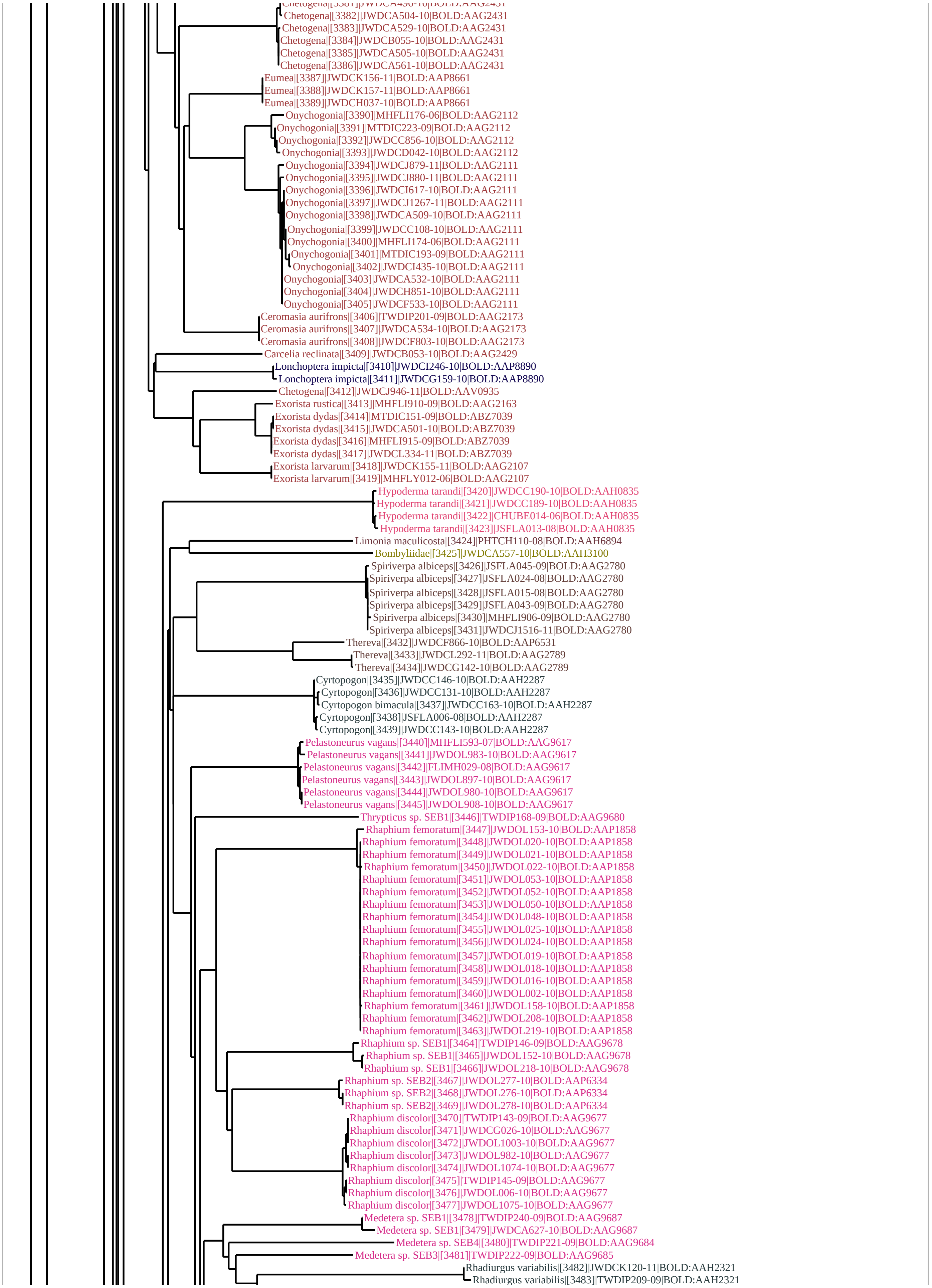
Chetogena|[3381]|JWDCA496-10|BOLD:AAG2431
Chetogena|[3382]|JWDCA504-10|BOLD:AAG2431
Chetogena|[3383]|JWDCA529-10|BOLD:AAG2431
Chetogena|[3384]|JWDCB055-10|BOLD:AAG2431
Chetogena|[3385]|JWDCA505-10|BOLD:AAG2431
Chetogena|[3386]|JWDCA561-10|BOLD:AAG2431
Eumea|[3387]|JWDCK156-11|BOLD:AAP8661
Eumea|[3388]|JWDCK157-11|BOLD:AAP8661
Eumea|[3389]|JWDCH037-10|BOLD:AAP8661
Onychogonia|[3390]|MHFLI176-06|BOLD:AAG2112
Onychogonia|[3391]|MTDIC223-09|BOLD:AAG2112
Onychogonia|[3392]|JWDCC856-10|BOLD:AAG2112
Onychogonia|[3393]|JWDCD042-10|BOLD:AAG2112
Onychogonia|[3394]|JWDCJ879-11|BOLD:AAG2111
Onychogonia|[3395]|JWDCJ880-11|BOLD:AAG2111
Onychogonia|[3396]|JWDCI617-10|BOLD:AAG2111
Onychogonia|[3397]|JWDCJ1267-11|BOLD:AAG2111
Onychogonia|[3398]|JWDCA509-10|BOLD:AAG2111
Onychogonia|[3399]|JWDCC108-10|BOLD:AAG2111
Onychogonia|[3400]|MHFLI174-06|BOLD:AAG2111
Onychogonia|[3401]|MTDIC193-09|BOLD:AAG2111
Onychogonia|[3402]|JWDCI435-10|BOLD:AAG2111
Onychogonia|[3403]|JWDCA532-10|BOLD:AAG2111
Onychogonia|[3404]|JWDCH851-10|BOLD:AAG2111
Onychogonia|[3405]|JWDCF533-10|BOLD:AAG2111
Ceromasia aurifrons|[3406]|TWDIP201-09|BOLD:AAG2173
Ceromasia aurifrons|[3407]|JWDCA534-10|BOLD:AAG2173
Ceromasia aurifrons|[3408]|JWDCF803-10|BOLD:AAG2173
Carcelia reclinata|[3409]|JWDCB053-10|BOLD:AAG2429
Lonchoptera impicta|[3410]|JWDCI246-10|BOLD:AAP8890
Lonchoptera impicta|[3411]|JWDCG159-10|BOLD:AAP8890
Chetogena|[3412]|JWDCJ946-11|BOLD:AAV0935
Exorista rustica|[3413]|MHFLI910-09|BOLD:AAG2163
Exorista dydas|[3414]|MTDIC151-09|BOLD:ABZ7039
Exorista dydas|[3415]|JWDCA501-10|BOLD:ABZ7039
Exorista dydas|[3416]|MHFLI915-09|BOLD:ABZ7039
Exorista dydas|[3417]|JWDCL334-11|BOLD:ABZ7039
Exorista larvarum|[3418]|JWDCK155-11|BOLD:AAG2107
Exorista larvarum|[3419]|MHFLY012-06|BOLD:AAG2107
Hypoderma tarandi|[3420]|JWDCC190-10|BOLD:AAH0835
Hypoderma tarandi|[3421]|JWDCC189-10|BOLD:AAH0835
Hypoderma tarandi|[3422]|CHUBE014-06|BOLD:AAH0835
Hypoderma tarandi|[3423]|JSFLA013-08|BOLD:AAH0835
Limonia maculicosta|[3424]|PHTCH110-08|BOLD:AAH6894
Bombyliidae|[3425]|JWDCA557-10|BOLD:AAH3100
Spiriverpa albiceps|[3426]|JSFLA045-09|BOLD:AAG2780
Spiriverpa albiceps|[3427]|JSFLA024-08|BOLD:AAG2780
Spiriverpa albiceps|[3428]|JSFLA015-08|BOLD:AAG2780
Spiriverpa albiceps|[3429]|JSFLA043-09|BOLD:AAG2780
Spiriverpa albiceps|[3430]|MHFLI906-09|BOLD:AAG2780
Spiriverpa albiceps|[3431]|JWDCJ1516-11|BOLD:AAG2780
Thereva|[3432]|JWDCF866-10|BOLD:AAP6531
Thereva|[3433]|JWDCL292-11|BOLD:AAG2789
Thereva|[3434]|JWDCG142-10|BOLD:AAG2789
Cyrtopogon|[3435]|JWDCC146-10|BOLD:AAH2287
Cyrtopogon|[3436]|JWDCC131-10|BOLD:AAH2287
Cyrtopogon bimacula|[3437]|JWDCC163-10|BOLD:AAH2287
Cyrtopogon|[3438]|JSFLA006-08|BOLD:AAH2287
Cyrtopogon|[3439]|JWDCC143-10|BOLD:AAH2287
Pelastoneurus vagans|[3440]|MHFLI593-07|BOLD:AAG9617
Pelastoneurus vagans|[3441]|JWDOL983-10|BOLD:AAG9617
Pelastoneurus vagans|[3442]|FLIMH029-08|BOLD:AAG9617
Pelastoneurus vagans|[3443]|JWDOL897-10|BOLD:AAG9617
Pelastoneurus vagans|[3444]|JWDOL980-10|BOLD:AAG9617
Pelastoneurus vagans|[3445]|JWDOL908-10|BOLD:AAG9617
Thrypticus sp. SEB1|[3446]|TWDIP168-09|BOLD:AAG9680
Rhaphium femoratum|[3447]|JWDOL153-10|BOLD:AAP1858
Rhaphium femoratum|[3448]|JWDOL020-10|BOLD:AAP1858
Rhaphium femoratum|[3449]|JWDOL021-10|BOLD:AAP1858
Rhaphium femoratum|[3450]|JWDOL022-10|BOLD:AAP1858
Rhaphium femoratum|[3451]|JWDOL053-10|BOLD:AAP1858
Rhaphium femoratum|[3452]|JWDOL052-10|BOLD:AAP1858
Rhaphium femoratum|[3453]|JWDOL050-10|BOLD:AAP1858
Rhaphium femoratum|[3454]|JWDOL048-10|BOLD:AAP1858
Rhaphium femoratum|[3455]|JWDOL025-10|BOLD:AAP1858
Rhaphium femoratum|[3456]|JWDOL024-10|BOLD:AAP1858
Rhaphium femoratum|[3457]|JWDOL019-10|BOLD:AAP1858
Rhaphium femoratum|[3458]|JWDOL018-10|BOLD:AAP1858
Rhaphium femoratum|[3459]|JWDOL016-10|BOLD:AAP1858
Rhaphium femoratum|[3460]|JWDOL002-10|BOLD:AAP1858
Rhaphium femoratum|[3461]|JWDOL158-10|BOLD:AAP1858
Rhaphium femoratum|[3462]|JWDOL208-10|BOLD:AAP1858
Rhaphium femoratum|[3463]|JWDOL219-10|BOLD:AAP1858
Rhaphium sp. SEB1|[3464]|TWDIP146-09|BOLD:AAG9678
Rhaphium sp. SEB1|[3465]|JWDOL152-10|BOLD:AAG9678
Rhaphium sp. SEB1|[3466]|JWDOL218-10|BOLD:AAG9678
Rhaphium sp. SEB2|[3467]|JWDOL277-10|BOLD:AAP6334
Rhaphium sp. SEB2|[3468]|JWDOL276-10|BOLD:AAP6334
Rhaphium sp. SEB2|[3469]|JWDOL278-10|BOLD:AAP6334
Rhaphium discolor|[3470]|TWDIP143-09|BOLD:AAG9677
Rhaphium discolor|[3471]|JWDCG026-10|BOLD:AAG9677
Rhaphium discolor|[3472]|JWDOL1003-10|BOLD:AAG9677
Rhaphium discolor|[3473]|JWDOL982-10|BOLD:AAG9677
Rhaphium discolor|[3474]|JWDOL1074-10|BOLD:AAG9677
Rhaphium discolor|[3475]|TWDIP145-09|BOLD:AAG9677
Rhaphium discolor|[3476]|JWDOL006-10|BOLD:AAG9677
Rhaphium discolor|[3477]|JWDOL1075-10|BOLD:AAG9677
Medetera sp. SEB1|[3478]|TWDIP240-09|BOLD:AAG9687
Medetera sp. SEB1|[3479]|JWDCA627-10|BOLD:AAG9687
Medetera sp. SEB4|[3480]|TWDIP221-09|BOLD:AAG9684
Medetera sp. SEB3|[3481]|TWDIP222-09|BOLD:AAG9685
Rhadiurgus variabilis|[3482]|JWDCK120-11|BOLD:AAH2321
Rhadiurgus variabilis|[3483]|TWDIP209-09|BOLD:AAH2321

Medetera sp. SEB3|[3481]|TWDIP222-09|BOLD:AAG9685
Rhadiurgus variabilis|[3482]|JWDCK120-11|BOLD:AAH2321
Rhadiurgus variabilis|[3483]|TWDIP209-09|BOLD:AAH2321
Medetera sp. SEB2|[3484]|JWDOL867-10|BOLD:AAP5021
Hercostomus unicolor|[3485]|JWDCB164-10|BOLD:AAG9651
Hercostomus unicolor|[3486]|JWDOL993-10|BOLD:AAG9651
Hercostomus unicolor|[3487]|JWDOL825-10|BOLD:AAG9651
Hercostomus unicolor|[3488]|JWDOL887-10|BOLD:AAG9651
Hercostomus unicolor|[3489]|JWDOL976-10|BOLD:AAG9651
Hercostomus unicolor|[3490]|JWDOL824-10|BOLD:AAG9651
Hercostomus unicolor|[3491]|JWDOL293-10|BOLD:AAG9651
Hercostomus unicolor|[3492]|JWDOL917-10|BOLD:AAG9651
Hercostomus unicolor|[3493]|JWDCB165-10|BOLD:AAG9651
Hercostomus unicolor|[3494]|MTDIC179-09|BOLD:AAG9651
Hercostomus unicolor|[3495]|JWDOL975-10|BOLD:AAG9651
Hercostomus unicolor|[3496]|JWDOL994-10|BOLD:AAG9651
Hercostomus unicolor|[3497]|JWDOL296-10|BOLD:AAG9651
Hercostomus unicolor|[3498]|JWDCB166-10|BOLD:AAG9651
Hercostomus unicolor|[3499]|JWDCB167-10|BOLD:AAG9651
Hercostomus unicolor|[3500]|JWDOL292-10|BOLD:AAG9651
Hercostomus unicolor|[3501]|JWDOL295-10|BOLD:AAG9651
Hercostomus unicolor|[3502]|JWDOL352-10|BOLD:AAG9651
Hercostomus unicolor|[3503]|JWDOL353-10|BOLD:AAG9651
Hercostomus unicolor|[3504]|JWDOL431-10|BOLD:AAG9651
Hercostomus unicolor|[3505]|JWDOL432-10|BOLD:AAG9651
Hercostomus unicolor|[3506]|JWDOL442-10|BOLD:AAG9651
Hercostomus unicolor|[3507]|JWDOL977-10|BOLD:AAG9651
Hercostomus unicolor|[3508]|JWDOL959-10|BOLD:AAG9651
Hercostomus unicolor|[3509]|JWDOL961-10|BOLD:AAG9651
Dolichopus sp. nr. discolor|[3510]|JWDOL223-10|BOLD:AAP1857
Dolichopus sp. nr. discolor|[3511]|JWDOL147-10|BOLD:AAP1857
Dolichopus sp. nr. discolor|[3512]|JWDOL221-10|BOLD:AAP1857
Dolichopus sp. nr. discolor|[3513]|JWDOL349-10|BOLD:AAP1857
Dolichopus sp. nr. discolor|[3514]|JWDOL350-10|BOLD:AAP1857
Dolichopus sp. nr. discolor|[3515]|JWDOL001-10|BOLD:AAP1857
Dolichopus sp. nr. discolor|[3516]|JWDOL148-10|BOLD:AAP1857
Dolichopus sp. nr. discolor|[3517]|JWDOL149-10|BOLD:AAP1857
Dolichopus sp. nr. discolor|[3518]|JWDOL005-10|BOLD:AAP1857
Dolichopus sp. nr. discolor|[3519]|JWDOL045-10|BOLD:AAP1857
Dolichopus sp. nr. discolor|[3520]|JWDOL283-10|BOLD:AAP1857
Dolichopus sp. nr. discolor|[3521]|JWDOL421-10|BOLD:AAP1857
Dolichopus sp. nr. discolor|[3522]|JWDOL222-10|BOLD:AAP1857
Dolichopus sp. nr. discolor|[3523]|JWDOL220-10|BOLD:AAP1857
Dolichopus sp. nr. discolor|[3524]|JWDOL163-10|BOLD:AAP1857
Dolichopus sp. nr. discolor|[3525]|JWDOL162-10|BOLD:AAP1857
Dolichopus sp. nr. discolor|[3526]|JWDOL146-10|BOLD:AAP1857
Dolichopus sp. nr. discolor|[3527]|JWDOL046-10|BOLD:AAP1857
Dolichopus sp. nr. discolor|[3528]|JWDOL043-10|BOLD:AAP1857
Dolichopus sp. nr. discolor|[3529]|JWDCF834-10|BOLD:AAP1857
Dolichopus sp. nr. discolor|[3530]|JWDCF340-10|BOLD:AAP1857
Dolichopus sp. nr. discolor|[3531]|JWDCF341-10|BOLD:AAP1857
Dolichopus sp. nr. discolor|[3532]|JWDOL229-10|BOLD:AAP1857
Dolichopus sp. nr. discolor|[3533]|JWDOL285-10|BOLD:AAP1857
Dolichopus sp. nr. discolor|[3534]|JWDOL345-10|BOLD:AAP1857
Dolichopus sp. nr. discolor|[3535]|JWDOL907-10|BOLD:AAP1857
Dolichopus latipennis|[3536]|MHFLI953-09|BOLD:AAG9611
Dolichopus latipennis|[3537]|JWDOL102-10|BOLD:AAG9611
Dolichopus latipennis|[3538]|JWDOL340-10|BOLD:AAG9611
Dolichopus latipennis|[3539]|TWDIP164-09
Dolichopus latipennis|[3540]|MHFLI657-07|BOLD:AAG9611
Dolichopus latipennis|[3541]|JWDOL028-10|BOLD:AAG9611
Dolichopus latipennis|[3542]|JWDOL023-10|BOLD:AAG9611
Dolichopus latipennis|[3543]|MHFLI650-07|BOLD:AAG9611
Dolichopus latipennis|[3544]|MHFLI635-07|BOLD:AAG9611
Dolichopus latipennis|[3545]|MHFLI614-07|BOLD:AAG9611
Dolichopus latipennis|[3546]|JWDOL338-10|BOLD:AAG9611
Dolichopus latipennis|[3547]|JWDOL320-10|BOLD:AAG9611
Dolichopus latipennis|[3548]|JWDOL319-10|BOLD:AAG9611
Dolichopus latipennis|[3549]|JWDOL318-10|BOLD:AAG9611
Dolichopus latipennis|[3550]|JWDOL240-10|BOLD:AAG9611
Dolichopus latipennis|[3551]|JWDOL192-10|BOLD:AAG9611
Dolichopus latipennis|[3552]|JWDOL191-10|BOLD:AAG9611
Dolichopus latipennis|[3553]|JWDOL186-10|BOLD:AAG9611
Dolichopus latipennis|[3554]|JWDOL185-10|BOLD:AAG9611
Dolichopus latipennis|[3555]|JWDOL182-10|BOLD:AAG9611
Dolichopus latipennis|[3556]|JWDOL170-10|BOLD:AAG9611
Dolichopus latipennis|[3557]|JWDOL169-10|BOLD:AAG9611
Dolichopus latipennis|[3558]|JWDOL081-10|BOLD:AAG9611
Dolichopus latipennis|[3559]|JWDOL080-10|BOLD:AAG9611
Dolichopus latipennis|[3560]|JWDOL073-10|BOLD:AAG9611
Dolichopus latipennis|[3561]|JWDOL072-10|BOLD:AAG9611
Dolichopus latipennis|[3562]|JWDOL027-10|BOLD:AAG9611
Dolichopus latipennis|[3563]|JWDCA035-10|BOLD:AAG9611
Dolichopus latipennis|[3564]|TWDIP150-09|BOLD:AAG9611
Dolichopus latipennis|[3565]|JWDOL082-10|BOLD:AAG9611
Dolichopus latipennis|[3566]|JWDOL074-10|BOLD:AAG9611
Dolichopus latipennis|[3567]|MHFLI088-06|BOLD:AAG9611
Dolichopus latipennis|[3568]|TWDIP162-09|BOLD:AAG9611
Dolichopus latipennis|[3569]|TWDIP166-09|BOLD:AAG9611
Dolichopus latipennis|[3570]|JWDOL183-10|BOLD:AAG9611
Dolichopus latipennis|[3571]|JWDOL1083-10|BOLD:AAG9611
Dolichopus latipennis|[3572]|JWDOL1084-10|BOLD:AAG9611
Dolichopus sp. nr. remus|[3573]|JWDOL055-10|BOLD:AAN5527
Dolichopus sp. nr. remus|[3574]|JWDOL128-10|BOLD:AAN5527
Dolichopus sp. nr. remus|[3575]|JWDOL135-10|BOLD:AAN5527
Dolichopus sp. nr. remus|[3576]|JWDOL136-10|BOLD:AAN5527
Dolichopus sp. nr. remus|[3577]|JWDOL137-10|BOLD:AAN5527
Dolichopus sp. nr. remus|[3578]|JWDOL228-10|BOLD:AAN5527
Dolichopus sp. nr. remus|[3579]|JWDOL289-10|BOLD:AAN5527
Dolichopus sp. nr. remus|[3580]|JWDOL347-10|BOLD:AAN5527
Dolichopus sp. nr. remus|[3581]|JWDOL372-10|BOLD:AAN5527
Dolichopus sp. nr. remus|[3582]|JWDCF600-10|BOLD:AAN5527
[illegible]

Dolichopus sp. nr. remus|[3580]|JWDOL347-10|BOLD:AAN5527
Dolichopus sp. nr. remus|[3581]|JWDOL372-10|BOLD:AAN5527
Dolichopus sp. nr. remus|[3582]|JWDCF600-10|BOLD:AAN5527
Dolichopus sp. nr. remus|[3583]|JWDOL057-10|BOLD:AAN5527
Dolichopus sp. nr. remus|[3584]|JWDOL362-10|BOLD:AAN5527
Dolichopus sp. nr. remus|[3585]|JWDOL127-10|BOLD:AAN5527
Dolichopus sp. nr. remus|[3586]|JWDOL065-10|BOLD:AAN5527
Dolichopus sp. nr. remus|[3587]|JWDOL058-10|BOLD:AAN5527
Dolichopus sp. nr. remus|[3588]|JWDCF599-10|BOLD:AAN5527
Dolichopus sp. nr. remus|[3589]|JWDCC090-10|BOLD:AAN5527
Dolichopus sp. nr. remus|[3590]|JWDCC088-10|BOLD:AAN5527
Dolichopus sp. nr. remus|[3591]|JWDOL079-10|BOLD:AAN5527
Dolichopus sp. nr. remus|[3592]|JWDCC089-10|BOLD:AAN5527
Dolichopus sp. nr. remus|[3593]|JWDOL290-10|BOLD:AAN5527
Dolichopus sp. nr. remus|[3594]|JWDOL306-10|BOLD:AAN5527
Dolichopus sp. nr. remus|[3595]|JWDOL433-10|BOLD:AAN5527
Dolichopus solidus|[3596]|JWDOL007-10|BOLD:AAP1859
Dolichopus solidus|[3597]|JWDOL396-10|BOLD:AAP1859
Dolichopus sp. nr. elegans|[3598]|MTDIC009-09|BOLD:ACZ6834
Dolichopus sp. nr. elegans|[3599]|JWDOL206-10|BOLD:ACZ6834
Dolichopus sp. nr. elegans|[3600]|JWDOL047-10|BOLD:ACZ6834
Dolichopus sp. nr. elegans|[3601]|JWDOL003-10|BOLD:ACZ6834
Dolichopus sp. nr. elegans|[3602]|JWDCG191-10|BOLD:ACZ6834
Dolichopus sp. nr. elegans|[3603]|JWDOL235-10|BOLD:ACZ6834
Dolichopus sp. nr. elegans|[3604]|JWDCF833-10|BOLD:ACZ6834
Dolichopus sp. nr. elegans|[3605]|JWDCH033-10|BOLD:ACZ6834
Dolichopus sp. nr. elegans|[3606]|JWDOL888-10|BOLD:ACZ6834
Dolichopus sp. nr. elegans|[3607]|JWDOL889-10|BOLD:ACZ6834
Dolichopus sp. nr. elegans|[3608]|JWDOL1016-10|BOLD:ACZ6834
Dolichopus sp. nr. elegans|[3609]|JWDOL424-10|BOLD:ACZ6834
Dolichopus sp. nr. elegans|[3610]|JWDOL378-10|BOLD:ACZ6834
Dolichopus sp. nr. elegans|[3611]|JWDOL357-10|BOLD:ACZ6834
Dolichopus sp. nr. elegans|[3612]|JWDOL288-10|BOLD:ACZ6834
Dolichopus sp. nr. elegans|[3613]|JWDOL257-10|BOLD:ACZ6834
Dolichopus sp. nr. elegans|[3614]|JWDOL237-10|BOLD:ACZ6834
Dolichopus sp. nr. elegans|[3615]|JWDOL236-10|BOLD:ACZ6834
Dolichopus sp. nr. elegans|[3616]|JWDOL198-10|BOLD:ACZ6834
Dolichopus sp. nr. elegans|[3617]|JWDOL040-10|BOLD:ACZ6834
Dolichopus sp. nr. elegans|[3618]|JWDCF722-10|BOLD:ACZ6834
Dolichopus sp. nr. elegans|[3619]|JWDCA032-10|BOLD:ACZ6834
Dolichopus sp. nr. elegans|[3620]|JWDOL154-10|BOLD:ACZ6834
Dolichopus sp. nr. elegans|[3621]|JWDOL224-10|BOLD:ACZ6834
Dolichopus sp. nr. elegans|[3622]|JWDOL826-10|BOLD:ACZ6834
Dolichopus sp. nr. elegans|[3623]|JWDOL890-10|BOLD:ACZ6834
Dolichopus sp. nr. elegans|[3624]|JWDOL200-10|BOLD:ACZ6834
Dolichopus sp. nr. elegans|[3625]|JWDOL923-10|BOLD:ACZ6834
Dolichopus sp. nr. elegans|[3626]|TWDIP234-09|BOLD:ACZ6834
Dolichopus sp. nr. elegans|[3627]|JWDOL064-10|BOLD:ACZ6834
Dolichopus sp. nr. elegans|[3628]|JWDOL403-10|BOLD:ACZ6834
Dolichopus sp. nr. elegans|[3629]|JWDOL1017-10|BOLD:ACZ6834
Dolichopus sp. nr. elegans|[3630]|JWDOL190-10|BOLD:ACZ6834
Dolichopus sp. nr. elegans|[3631]|JWDOL197-10|BOLD:ACZ6834
Dolichopus sp. nr. elegans|[3632]|JWDOL868-10|BOLD:ACZ6834
Dolichopus sp. nr. elegans|[3633]|JWDOL903-10|BOLD:ACZ6834
Dolichopus sp. nr. diversipennis|[3634]|JWDCA034-10|BOLD:AAG7065
Dolichopus sp. nr. diversipennis|[3635]|JWDOL049-10|BOLD:AAG7065
Dolichopus sp. nr. diversipennis|[3636]|JWDCD184-10|BOLD:AAG7065
Dolichopus sp. nr. diversipennis|[3637]|JWDCA038-10|BOLD:AAG7065
Dolichopus sp. nr. diversipennis|[3638]|JWDCF208-10|BOLD:AAG7065
Dolichopus sp. nr. diversipennis|[3639]|JWDOL042-10|BOLD:AAG7065
Dolichopus sp. nr. diversipennis|[3640]|JWDOL172-10|BOLD:AAG7065
Dolichopus sp. nr. diversipennis|[3641]|JWDOL173-10|BOLD:AAG7065
Dolichopus diversipennis|[3642]|JWDOL176-10|BOLD:AAG9614
Dolichopus diversipennis|[3643]|JWDCC007-10|BOLD:AAG9614
Dolichopus diversipennis|[3644]|JWDOL030-10|BOLD:AAG9614
Dolichopus diversipennis|[3645]|JWDOL175-10|BOLD:AAG9614
Dolichopus diversipennis|[3646]|JWDOL195-10|BOLD:AAG9614
Dolichopus diversipennis|[3647]|MHFLI265-06|BOLD:AAG9614
Dolichopus diversipennis|[3648]|JWDOL029-10|BOLD:AAG9614
Dolichopus diversipennis|[3649]|JWDOL174-10|BOLD:AAG9614
Dolichopus diversipennis|[3650]|JWDOL177-10|BOLD:AAG9614
Dolichopus diversipennis|[3651]|JWDOL199-10|BOLD:AAG9614
Dolichopus diversipennis|[3652]|JWDOL202-10|BOLD:AAG9614
Dolichopus diversipennis|[3653]|JWDOL201-10|BOLD:AAG9614
Dolichopus diversipennis|[3654]|JWDOL241-10|BOLD:AAG9614
Dolichopus detersus|[3655]|JWDCD085-10|BOLD:ACK3631
Dolichopus detersus|[3656]|JWDOL187-10|BOLD:ACK3631
Dolichopus detersus|[3657]|JWDOL467-10|BOLD:ACK3631
Dolichopus genualis|[3658]|JWDOL895-10|BOLD:AAP5022
Dolichopus sp. nr. obcordatus SEB1|[3659]|MHFLI078-06|BOLD:AAG9610
Dolichopus sp. nr. obcordatus SEB1|[3660]|MHFLI084-06|BOLD:AAG9610
Dolichopus sp. nr. obcordatus SEB2|[3661]|JWDOL216-10|BOLD:AAG9610
Dolichopus sp. nr. obcordatus SEB2|[3662]|JWDOL327-10|BOLD:AAG9610
Dolichopus tetricus|[3663]|JWDCD127-10|BOLD:AAG9688
Dolichopus tetricus|[3664]|TWDIP349-09|BOLD:AAG9688
Dolichopus tetricus|[3665]|JWDCA030-10|BOLD:AAG9688
Dolichopus tetricus|[3666]|JWDOL400-10|BOLD:AAG9688
Dolichopus sp. SEB03|[3667]|JWDOL427-10|BOLD:AAP6335
Dolichopus sp. SEB11|[3668]|JWDOL821-10|BOLD:AAG9634
Dolichopus nudus|[3669]|JWDOL813-10|BOLD:AAG9723
Dolichopus nudus|[3670]|JWDCH657-10|BOLD:AAG9723
Dolichopus nudus|[3671]|JWDOL822-10|BOLD:AAG9723
Dolichopus nudus|[3672]|JWDOL291-10|BOLD:AAG9723
Dolichopus nudus|[3673]|JWDOL428-10|BOLD:AAG9723
Dolichopus nudus|[3674]|JWDOL940-10|BOLD:AAG9723
Dolichopus nudus|[3675]|JWDOL863-10|BOLD:AAG9723
Dolichopus nudus|[3676]|JWDOL430-10|BOLD:AAG9723
Dolichopus nudus|[3677]|JWDOL941-10|BOLD:AAG9723
Dolichopus nudus|[3678]|JWDOL819-10|BOLD:AAG9723
Dolichopus nudus|[3679]|JWDOL878-10|BOLD:AAG9723
Dolichopus brevipennis|[3680]|JWDOL090-10|BOLD:AAG9609
Dolichopus brevipennis|[3681]|MHFLY049-06|BOLD:AAG9609
Dolichopus brevipennis|[3682]|JWDOL332-10|BOLD:AAG9609

Dolichopus brevipennis|[3680]|JWDOL090-10|BOLD:AAG9609
Dolichopus brevipennis|[3681]|MHFLY049-06|BOLD:AAG9609
Dolichopus brevipennis|[3682]|JWDOL332-10|BOLD:AAG9609
Dolichopus brevipennis|[3683]|JWDOL384-10|BOLD:AAG9609
Dolichopus brevipennis|[3684]|MHFLI887-09|BOLD:AAG9609
Dolichopus brevipennis|[3685]|MHFLI079-06|BOLD:AAG9609
Dolichopus brevipennis|[3686]|MHFLI886-09|BOLD:AAG9609
Dolichopus brevipennis|[3687]|MHFLI082-06|BOLD:AAG9609
Dolichopus brevipennis|[3688]|MHFLI081-06|BOLD:AAG9609
Dolichopus brevipennis|[3689]|MHFLY043-06|BOLD:AAG9609
Dolichopus brevipennis|[3690]|JWDCL539-11|BOLD:AAG9609
Dolichopus brevipennis|[3691]|JWDOL105-10|BOLD:AAG9609
Dolichopus brevipennis|[3692]|JWDOL385-10|BOLD:AAG9609
Dolichopus brevipennis|[3693]|JWDOL1014-10|BOLD:AAG9609
Dolichopus brevipennis|[3694]|JWDOL461-10|BOLD:AAG9609
Dolichopus brevipennis|[3695]|JWDOL933-10|BOLD:AAG9609
Dolichopus brevipennis|[3696]|JWDOL929-10|BOLD:AAG9609
Dolichopus brevipennis|[3697]|JWDOL382-10|BOLD:AAG9609
Dolichopus brevipennis|[3698]|JWDOL1012-10|BOLD:AAG9609
Dolichopus brevipennis|[3699]|JWDOL126-10|BOLD:AAG9609
Dolichopus brevipennis|[3700]|JWDOL1013-10|BOLD:AAG9609
Dolichopus brevipennis|[3701]|JWDOL875-10|BOLD:AAG9609
Dolichopus brevipennis|[3702]|JWDOL087-10|BOLD:AAG9609
Dolichopus brevipennis|[3703]|JWDOL013-10|BOLD:AAG9609
Dolichopus brevipennis|[3704]|JWDOL932-10|BOLD:AAG9609
Dolichopus brevipennis|[3705]|JWDOL934-10|BOLD:AAG9609
Dolichopus brevipennis|[3706]|JWDOL1015-10|BOLD:AAG9609
Dolichopus brevipennis|[3707]|JWDOL881-10|BOLD:AAG9609
Dolichopus brevipennis|[3708]|MHFLI076-06|BOLD:AAG9609
Dolichopus brevipennis|[3709]|MHFLI083-06|BOLD:AAG9609
Dolichopus brevipennis|[3710]|JWDOL1048-10|BOLD:AAG9609
Dolichopus brevipennis|[3711]|JWDOL1011-10|BOLD:AAG9609
Dolichopus brevipennis|[3712]|JWDOL892-10|BOLD:AAG9609
Dolichopus brevipennis|[3713]|MHFLI087-06|BOLD:AAG9609
Dolichopus brevipennis|[3714]|MHFLI080-06|BOLD:AAG9609
Dolichopus brevipennis|[3715]|JWDOL375-10|BOLD:AAG9609
Dolichopus brevipennis|[3716]|MHFLI075-06|BOLD:AAG9609
Dolichopus brevipennis|[3717]|JWDOL935-10|BOLD:AAG9609
Dolichopus brevipennis|[3718]|JWDOL883-10|BOLD:AAG9609
Dolichopus brevipennis|[3719]|JWDOL891-10|BOLD:AAG9609
Dolichopus brevipennis|[3720]|JWDOL460-10|BOLD:AAG9609
Dolichopus brevipennis|[3721]|JWDOL882-10|BOLD:AAG9609
Dolichopus brevipennis|[3722]|JWDOL1040-10|BOLD:AAG9609
Dolichopus brevipennis|[3723]|JWDOL1049-10|BOLD:AAG9609
Dolichopus brevipennis|[3724]|JWDOL930-10|BOLD:AAG9609
Dolichopus brevipennis|[3725]|JWDOL931-10|BOLD:AAG9609
Dolichopus brevipennis|[3726]|JWDOL336-10|BOLD:AAG9609
Dolichopus brevipennis|[3727]|JWDOL386-10|BOLD:AAG9609
Dolichopus brevipennis|[3728]|MHFLI085-06|BOLD:AAG9609
Dolichopus brevipennis|[3729]|JWDCA014-10|BOLD:AAG9609
Dolichopus brevipennis|[3730]|JWDCH942-10|BOLD:AAG9609
Dolichopus brevipennis|[3731]|JWDOL1008-10|BOLD:AAG9609
Dolichopus brevipennis|[3732]|JWDOL014-10|BOLD:AAG9609
Dolichopus brevipennis|[3733]|JWDOL015-10|BOLD:AAG9609
Dolichopus brevipennis|[3734]|JWDOL088-10|BOLD:AAG9609
Dolichopus brevipennis|[3735]|JWDOL089-10|BOLD:AAG9609
Dolichopus brevipennis|[3736]|JWDOL091-10|BOLD:AAG9609
Dolichopus brevipennis|[3737]|JWDOL244-10|BOLD:AAG9609
Dolichopus brevipennis|[3738]|JWDOL333-10|BOLD:AAG9609
Dolichopus brevipennis|[3739]|JWDOL334-10|BOLD:AAG9609
Dolichopus brevipennis|[3740]|JWDOL335-10|BOLD:AAG9609
Dolichopus brevipennis|[3741]|JWDOL893-10|BOLD:AAG9609
Dolichopus sp. SEB06|[3742]|TWDIP238-09|BOLD:AAG9686
Dolichopus sp. SEB06|[3743]|JWDOL860-10|BOLD:AAG9686
Dolichopus sp. SEB04|[3744]|JWDCA036-10|BOLD:ABY5310
Dolichopus sp. SEB05|[3745]|JWDCG792-10|BOLD:AAP3632
Dolichopus sp. SEB05|[3746]|JWDOL364-10|BOLD:AAP3632
Dolichopus stenhammari|[3747]|JWDCE434-10|BOLD:AAG9682
Dolichopus stenhammari|[3748]|JWDOL004-10|BOLD:AAG9682
Dolichopus stenhammari|[3749]|JWDOL253-10|BOLD:AAG9682
Dolichopus stenhammari|[3750]|JWDCE266-10|BOLD:AAG9682
Dolichopus stenhammari|[3751]|JWDOL251-10|BOLD:AAG9682
Dolichopus stenhammari|[3752]|JWDOL213-10|BOLD:AAG9682
Dolichopus stenhammari|[3753]|JWDOL212-10|BOLD:AAG9682
Dolichopus stenhammari|[3754]|JWDOL181-10|BOLD:AAG9682
Dolichopus stenhammari|[3755]|JWDCF863-10|BOLD:AAG9682
Dolichopus stenhammari|[3756]|JWDCB160-10|BOLD:AAG9682
Dolichopus stenhammari|[3757]|JWDOL252-10|BOLD:AAG9682
Dolichopus stenhammari|[3758]|JWDOL254-10|BOLD:AAG9682
Dolichopus stenhammari|[3759]|JWDOL281-10|BOLD:AAG9682
Dolichopus stenhammari|[3760]|JWDOL298-10|BOLD:AAG9682
Dolichopus stenhammari|[3761]|JWDOL321-10|BOLD:AAG9682
Dolichopus stenhammari|[3762]|JWDOL322-10|BOLD:AAG9682
Dolichopus stenhammari|[3763]|JWDOL1077-10|BOLD:AAG9682
Dolichopus stenhammari|[3764]|MTDIC100-09|BOLD:AAG9682
Dolichopus stenhammari|[3765]|MTDIC106-09|BOLD:AAG9682
Dolichopus stenhammari|[3766]|JWDOL078-10|BOLD:AAG9682
Dolichopus stenhammari|[3767]|JWDOL140-10|BOLD:AAG9682
Dolichopus stenhammari|[3768]|JWDOL210-10|BOLD:AAG9682
Dolichopus stenhammari|[3769]|JWDOL211-10|BOLD:AAG9682
Dolichopus stenhammari|[3770]|JWDOL282-10|BOLD:AAG9682
Dolichopus stenhammari|[3771]|JWDOL376-10|BOLD:AAG9682
Dolichopus stenhammari|[3772]|JWDOL962-10|BOLD:AAG9682
Dolichopus sp. nr. defectus|[3773]|JWDOL919-10|BOLD:AAP5023
Dolichopus sp. nr. defectus|[3774]|JWDOL910-10|BOLD:AAP5023
Dolichopus sp. SEB13 nr. boreus|[3775]|JWDOL161-10|BOLD:AAG9628
Dolichopus sp. SEB13 nr. boreus|[3776]|JWDCH866-10|BOLD:AAG9628
Dolichopus sp. SEB13 nr. boreus|[3777]|TWDIP151-09|BOLD:AAG9628
Dolichopus sp. SEB13 nr. boreus|[3778]|JWDOL323-10|BOLD:AAG9628
Dolichopus sp. SEB13 nr. boreus|[3779]|JWDOL324-10|BOLD:AAG9628
Dolichopus sp. SEB13 nr. boreus|[3780]|JWDOL373-10|BOLD:AAG9628
Dolichopus lamellipes|[3781]|JWDCF881-10|BOLD:ACZ5095
[illegible]

Dolichopus sp. SEB13 nr. boreus|[3779]|JWDOL324-10|BOLD:AAG9628
Dolichopus sp. SEB13 nr. boreus|[3780]|JWDOL373-10|BOLD:AAG9628
Dolichopus lamellipes|[3781]|JWDCF881-10|BOLD:ACZ5095
Dolichopus lamellipes|[3782]|JWDCG675-10|BOLD:ACZ5095
Dolichopus lamellipes|[3783]|JWDCF721-10|BOLD:ACZ5095
Dolichopus lamellipes|[3784]|JWDCF794-10|BOLD:ACZ5095
Dolichopus lamellipes|[3785]|JWDOL230-10|BOLD:ACZ5095
Dolichopus lamellipes|[3786]|JWDCI760-11|BOLD:ACZ5095
Dolichopus lamellipes|[3787]|JWDOL284-10|BOLD:ACZ5095
Dolichopus lamellipes|[3788]|TWDIP232-09|BOLD:ACZ5095
Dolichopus lamellipes|[3789]|JWDCF215-10|BOLD:ACZ5095
Dolichopus lamellipes|[3790]|JWDOL346-10|BOLD:ACZ5095
Dolichopus lamellipes|[3791]|JWDOL358-10|BOLD:ACZ5095
Dolichopus lamellipes|[3792]|JWDOL418-10|BOLD:ACZ5095
Dolichopus lamellipes|[3793]|JWDCA029-10|BOLD:ACZ5095
Dolichopus lamellipes|[3794]|JWDCI146-10|BOLD:ACZ5095
Dolichopus lamellipes|[3795]|JWDOL075-10|BOLD:ACZ5095
Dolichopus lamellipes|[3796]|JWDOL189-10|BOLD:ACZ5095
Dolichopus lamellipes|[3797]|JWDCF941-10|BOLD:ACZ5095
Dolichopus lamellipes|[3798]|JWDOL033-10|BOLD:ACZ5095
Dolichopus lamellipes|[3799]|JWDOL410-10|BOLD:ACZ5095
Dolichopus lamellipes|[3800]|JWDOL1082-10|BOLD:ACZ5095
Dolichopus lamellipes|[3801]|JWDOL233-10|BOLD:ACZ5095
Dolichopus lamellipes|[3802]|JWDOL420-10|BOLD:ACZ5095
Dolichopus lamellipes|[3803]|JWDCG015-10|BOLD:ACZ5095
Dolichopus lamellipes|[3804]|JWDCG790-10|BOLD:ACZ5095
Dolichopus lamellipes|[3805]|TWDIP233-09|BOLD:ACZ5095
Dolichopus lamellipes|[3806]|JWDOL273-10|BOLD:ACZ5095
Dolichopus lamellipes|[3807]|JWDOL308-10|BOLD:ACZ5095
Dolichopus lamellipes|[3808]|JWDOL077-10|BOLD:ACZ5095
Dolichopus lamellipes|[3809]|JWDCG637-10|BOLD:ACZ5095
Dolichopus lamellipes|[3810]|JWDOL054-10|BOLD:ACZ5095
Dolichopus lamellipes|[3811]|JWDOL388-10|BOLD:ACZ5095
Dolichopus lamellipes|[3812]|JWDOL188-10|BOLD:ACZ5095
Dolichopus lamellipes|[3813]|JWDOL041-10|BOLD:ACZ5095
Dolichopus lamellipes|[3814]|JWDCG861-10|BOLD:ACZ5095
Dolichopus lamellipes|[3815]|MTDIC209-09|BOLD:ACZ5095
Dolichopus lamellipes|[3816]|JWDOL262-10|BOLD:ACZ5095
Dolichopus lamellipes|[3817]|JWDOL286-10|BOLD:ACZ5095
Dolichopus lamellipes|[3818]|JWDOL979-10|BOLD:ACZ5095
Dolichopus lamellipes|[3819]|JWDOL1089-10|BOLD:ACZ5095
Dolichopus lamellipes|[3820]|JWDOL142-10|BOLD:ACZ5095
Dolichopus lamellipes|[3821]|JWDOL084-10|BOLD:ACZ5095
Dolichopus lamellipes|[3822]|JWDOL300-10|BOLD:ACZ5095
Dolichopus lamellipes|[3823]|JWDCA002-10|BOLD:ACZ5095
Dolichopus lamellipes|[3824]|JWDOL390-10|BOLD:ACZ5095
Dolichopus lamellipes|[3825]|JWDOL068-10|BOLD:ACZ5095
Dolichopus lamellipes|[3826]|JWDCI758-11|BOLD:ACZ5095
Dolichopus lamellipes|[3827]|JWDCF209-10|BOLD:ACZ5095
Dolichopus lamellipes|[3828]|JWDOL314-10|BOLD:ACZ5095
Dolichopus lamellipes|[3829]|JWDOL193-10|BOLD:ACZ5095
Dolichopus lamellipes|[3830]|TWDIP152-09|BOLD:ACZ5095
Dolichopus lamellipes|[3831]|JWDCI224-10|BOLD:ACZ5095
Dolichopus lamellipes|[3832]|JWDOL258-10|BOLD:ACZ5095
Dolichopus lamellipes|[3833]|JWDOL225-10|BOLD:ACZ5095
Dolichopus lamellipes|[3834]|JWDCI228-10|BOLD:ACZ5095
Dolichopus lamellipes|[3835]|JWDOL312-10|BOLD:ACZ5095
Dolichopus lamellipes|[3836]|JWDOL317-10|BOLD:ACZ5095
Dolichopus lamellipes|[3837]|JWDOL328-10|BOLD:ACZ5095
Dolichopus lamellipes|[3838]|JWDOL343-10|BOLD:ACZ5095
Dolichopus lamellipes|[3839]|JWDOL348-10|BOLD:ACZ5095
Dolichopus lamellipes|[3840]|JWDOL351-10|BOLD:ACZ5095
Dolichopus lamellipes|[3841]|JWDOL356-10|BOLD:ACZ5095
Dolichopus lamellipes|[3842]|JWDOL363-10|BOLD:ACZ5095
Dolichopus lamellipes|[3843]|JWDOL365-10|BOLD:ACZ5095
Dolichopus lamellipes|[3844]|JWDOL411-10|BOLD:ACZ5095
Dolichopus lamellipes|[3845]|JWDOL412-10|BOLD:ACZ5095
Dolichopus lamellipes|[3846]|JWDOL413-10|BOLD:ACZ5095
Dolichopus lamellipes|[3847]|JWDOL422-10|BOLD:ACZ5095
Dolichopus lamellipes|[3848]|JWDOL425-10|BOLD:ACZ5095
Dolichopus lamellipes|[3849]|JWDOL446-10|BOLD:ACZ5095
Dolichopus lamellipes|[3850]|JWDOL971-10|BOLD:ACZ5095
Dolichopus lamellipes|[3851]|JWDOL1050-10|BOLD:ACZ5095
Dolichopus lamellipes|[3852]|JWDOL1078-10|BOLD:ACZ5095
Dolichopus lamellipes|[3853]|JWDOL272-10|BOLD:ACZ5095
Dolichopus lamellipes|[3854]|JWDOL274-10|BOLD:ACZ5095
Dolichopus lamellipes|[3855]|JWDOL205-10|BOLD:ACZ5095
Dolichopus lamellipes|[3856]|JWDOL227-10|BOLD:ACZ5095
Dolichopus lamellipes|[3857]|JWDOL232-10|BOLD:ACZ5095
Dolichopus lamellipes|[3858]|JWDOL242-10|BOLD:ACZ5095
Dolichopus lamellipes|[3859]|JWDOL069-10|BOLD:ACZ5095
Dolichopus lamellipes|[3860]|JWDOL070-10|BOLD:ACZ5095
Dolichopus lamellipes|[3861]|JWDOL131-10|BOLD:ACZ5095
Dolichopus lamellipes|[3862]|JWDOL138-10|BOLD:ACZ5095
Dolichopus lamellipes|[3863]|JWDOL259-10|BOLD:ACZ5095
Dolichopus lamellipes|[3864]|JWDOL260-10|BOLD:ACZ5095
Dolichopus lamellipes|[3865]|JWDOL263-10|BOLD:ACZ5095
Dolichopus lamellipes|[3866]|JWDOL268-10|BOLD:ACZ5095
Dolichopus lamellipes|[3867]|JWDOL139-10|BOLD:ACZ5095
Dolichopus lamellipes|[3868]|JWDOL155-10|BOLD:ACZ5095
Dolichopus lamellipes|[3869]|JWDCI222-10|BOLD:ACZ5095
Dolichopus lamellipes|[3870]|JWDCF354-10|BOLD:ACZ5095
Dolichopus lamellipes|[3871]|JWDCI759-11|BOLD:ACZ5095
Dolichopus lamellipes|[3872]|JWDCH349-10|BOLD:ACZ5095
Dolichopus lamellipes|[3873]|JWDCF795-10|BOLD:ACZ5095
Dolichopus lamellipes|[3874]|JWDCF835-10|BOLD:ACZ5095
Dolichopus lamellipes|[3875]|JWDCF880-10|BOLD:ACZ5095
Dolichopus lamellipes|[3876]|JWDOL039-10|BOLD:ACZ5095
Dolichopus lamellipes|[3877]|JWDOL056-10|BOLD:ACZ5095
Dolichopus lamellipes|[3878]|JWDOL067-10|BOLD:ACZ5095
Dolichopus lamellipes|[3879]|JWDCF355-10|BOLD:ACZ5095
Dolichopus lamellipes|[3880]|JWDCF793-10|BOLD:ACZ5095
Dolichopus lamellipes|[3881]|JWDCI221-10|BOLD:ACZ5095

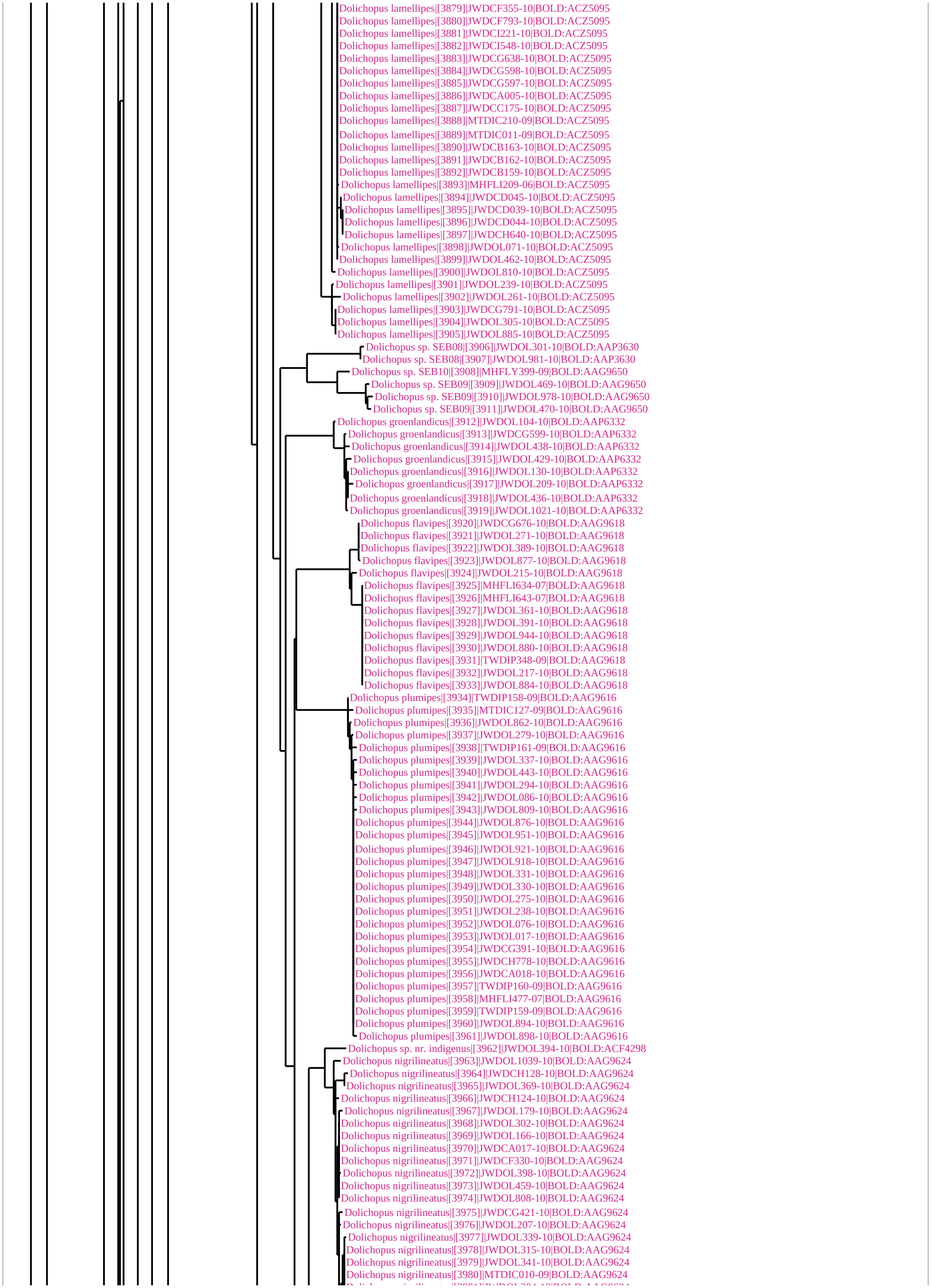
Dolichopus lamellipes|[3879]|JWDCF355-10|BOLD:ACZ5095
Dolichopus lamellipes|[3880]|JWDCF793-10|BOLD:ACZ5095
Dolichopus lamellipes|[3881]|JWDCI221-10|BOLD:ACZ5095
Dolichopus lamellipes|[3882]|JWDCI548-10|BOLD:ACZ5095
Dolichopus lamellipes|[3883]|JWDCG638-10|BOLD:ACZ5095
Dolichopus lamellipes|[3884]|JWDCG598-10|BOLD:ACZ5095
Dolichopus lamellipes|[3885]|JWDCG597-10|BOLD:ACZ5095
Dolichopus lamellipes|[3886]|JWDCA005-10|BOLD:ACZ5095
Dolichopus lamellipes|[3887]|JWDCC175-10|BOLD:ACZ5095
Dolichopus lamellipes|[3888]|MTDIC210-09|BOLD:ACZ5095
Dolichopus lamellipes|[3889]|MTDIC011-09|BOLD:ACZ5095
Dolichopus lamellipes|[3890]|JWDCB163-10|BOLD:ACZ5095
Dolichopus lamellipes|[3891]|JWDCB162-10|BOLD:ACZ5095
Dolichopus lamellipes|[3892]|JWDCB159-10|BOLD:ACZ5095
Dolichopus lamellipes|[3893]|MHFLI209-06|BOLD:ACZ5095
Dolichopus lamellipes|[3894]|JWDCD045-10|BOLD:ACZ5095
Dolichopus lamellipes|[3895]|JWDCD039-10|BOLD:ACZ5095
Dolichopus lamellipes|[3896]|JWDCD044-10|BOLD:ACZ5095
Dolichopus lamellipes|[3897]|JWDCH640-10|BOLD:ACZ5095
Dolichopus lamellipes|[3898]|JWDOL071-10|BOLD:ACZ5095
Dolichopus lamellipes|[3899]|JWDOL462-10|BOLD:ACZ5095
Dolichopus lamellipes|[3900]|JWDOL810-10|BOLD:ACZ5095
Dolichopus lamellipes|[3901]|JWDOL239-10|BOLD:ACZ5095
Dolichopus lamellipes|[3902]|JWDOL261-10|BOLD:ACZ5095
Dolichopus lamellipes|[3903]|JWDCG791-10|BOLD:ACZ5095
Dolichopus lamellipes|[3904]|JWDOL305-10|BOLD:ACZ5095
Dolichopus lamellipes|[3905]|JWDOL885-10|BOLD:ACZ5095
Dolichopus sp. SEB08|[3906]|JWDOL301-10|BOLD:AAP3630
Dolichopus sp. SEB08|[3907]|JWDOL981-10|BOLD:AAP3630
Dolichopus sp. SEB10|[3908]|MHFLY399-09|BOLD:AAG9650
Dolichopus sp. SEB09|[3909]|JWDOL469-10|BOLD:AAG9650
Dolichopus sp. SEB09|[3910]|JWDOL978-10|BOLD:AAG9650
Dolichopus sp. SEB09|[3911]|JWDOL470-10|BOLD:AAG9650
Dolichopus groenlandicus|[3912]|JWDOL104-10|BOLD:AAP6332
Dolichopus groenlandicus|[3913]|JWDCG599-10|BOLD:AAP6332
Dolichopus groenlandicus|[3914]|JWDOL438-10|BOLD:AAP6332
Dolichopus groenlandicus|[3915]|JWDOL429-10|BOLD:AAP6332
Dolichopus groenlandicus|[3916]|JWDOL130-10|BOLD:AAP6332
Dolichopus groenlandicus|[3917]|JWDOL209-10|BOLD:AAP6332
Dolichopus groenlandicus|[3918]|JWDOL436-10|BOLD:AAP6332
Dolichopus groenlandicus|[3919]|JWDOL1021-10|BOLD:AAP6332
Dolichopus flavipes|[3920]|JWDCG676-10|BOLD:AAG9618
Dolichopus flavipes|[3921]|JWDOL271-10|BOLD:AAG9618
Dolichopus flavipes|[3922]|JWDOL389-10|BOLD:AAG9618
Dolichopus flavipes|[3923]|JWDOL877-10|BOLD:AAG9618
Dolichopus flavipes|[3924]|JWDOL215-10|BOLD:AAG9618
Dolichopus flavipes|[3925]|MHFLI634-07|BOLD:AAG9618
Dolichopus flavipes|[3926]|MHFLI643-07|BOLD:AAG9618
Dolichopus flavipes|[3927]|JWDOL361-10|BOLD:AAG9618
Dolichopus flavipes|[3928]|JWDOL391-10|BOLD:AAG9618
Dolichopus flavipes|[3929]|JWDOL944-10|BOLD:AAG9618
Dolichopus flavipes|[3930]|JWDOL880-10|BOLD:AAG9618
Dolichopus flavipes|[3931]|TWDIP348-09|BOLD:AAG9618
Dolichopus flavipes|[3932]|JWDOL217-10|BOLD:AAG9618
Dolichopus flavipes|[3933]|JWDOL884-10|BOLD:AAG9618
Dolichopus plumipes|[3934]|TWDIP158-09|BOLD:AAG9616
Dolichopus plumipes|[3935]|MTDIC127-09|BOLD:AAG9616
Dolichopus plumipes|[3936]|JWDOL862-10|BOLD:AAG9616
Dolichopus plumipes|[3937]|JWDOL279-10|BOLD:AAG9616
Dolichopus plumipes|[3938]|TWDIP161-09|BOLD:AAG9616
Dolichopus plumipes|[3939]|JWDOL337-10|BOLD:AAG9616
Dolichopus plumipes|[3940]|JWDOL443-10|BOLD:AAG9616
Dolichopus plumipes|[3941]|JWDOL294-10|BOLD:AAG9616
Dolichopus plumipes|[3942]|JWDOL086-10|BOLD:AAG9616
Dolichopus plumipes|[3943]|JWDOL809-10|BOLD:AAG9616
Dolichopus plumipes|[3944]|JWDOL876-10|BOLD:AAG9616
Dolichopus plumipes|[3945]|JWDOL951-10|BOLD:AAG9616
Dolichopus plumipes|[3946]|JWDOL921-10|BOLD:AAG9616
Dolichopus plumipes|[3947]|JWDOL918-10|BOLD:AAG9616
Dolichopus plumipes|[3948]|JWDOL331-10|BOLD:AAG9616
Dolichopus plumipes|[3949]|JWDOL330-10|BOLD:AAG9616
Dolichopus plumipes|[3950]|JWDOL275-10|BOLD:AAG9616
Dolichopus plumipes|[3951]|JWDOL238-10|BOLD:AAG9616
Dolichopus plumipes|[3952]|JWDOL076-10|BOLD:AAG9616
Dolichopus plumipes|[3953]|JWDOL017-10|BOLD:AAG9616
Dolichopus plumipes|[3954]|JWDCG391-10|BOLD:AAG9616
Dolichopus plumipes|[3955]|JWDCH778-10|BOLD:AAG9616
Dolichopus plumipes|[3956]|JWDCA018-10|BOLD:AAG9616
Dolichopus plumipes|[3957]|TWDIP160-09|BOLD:AAG9616
Dolichopus plumipes|[3958]|MHFLI477-07|BOLD:AAG9616
Dolichopus plumipes|[3959]|TWDIP159-09|BOLD:AAG9616
Dolichopus plumipes|[3960]|JWDOL894-10|BOLD:AAG9616
Dolichopus plumipes|[3961]|JWDOL898-10|BOLD:AAG9616
Dolichopus sp. nr. indigenus|[3962]|JWDOL394-10|BOLD:ACF4298
Dolichopus nigrilineatus|[3963]|JWDOL1039-10|BOLD:AAG9624
Dolichopus nigrilineatus|[3964]|JWDCH128-10|BOLD:AAG9624
Dolichopus nigrilineatus|[3965]|JWDOL369-10|BOLD:AAG9624
Dolichopus nigrilineatus|[3966]|JWDCH124-10|BOLD:AAG9624
Dolichopus nigrilineatus|[3967]|JWDOL179-10|BOLD:AAG9624
Dolichopus nigrilineatus|[3968]|JWDOL302-10|BOLD:AAG9624
Dolichopus nigrilineatus|[3969]|JWDOL166-10|BOLD:AAG9624
Dolichopus nigrilineatus|[3970]|JWDCA017-10|BOLD:AAG9624
Dolichopus nigrilineatus|[3971]|JWDCF330-10|BOLD:AAG9624
Dolichopus nigrilineatus|[3972]|JWDOL398-10|BOLD:AAG9624
Dolichopus nigrilineatus|[3973]|JWDOL459-10|BOLD:AAG9624
Dolichopus nigrilineatus|[3974]|JWDOL808-10|BOLD:AAG9624
Dolichopus nigrilineatus|[3975]|JWDCG421-10|BOLD:AAG9624
Dolichopus nigrilineatus|[3976]|JWDOL207-10|BOLD:AAG9624
Dolichopus nigrilineatus|[3977]|JWDOL339-10|BOLD:AAG9624
Dolichopus nigrilineatus|[3978]|JWDOL315-10|BOLD:AAG9624
Dolichopus nigrilineatus|[3979]|JWDOL341-10|BOLD:AAG9624
Dolichopus nigrilineatus|[3980]|MTDIC010-09|BOLD:AAG9624

Dolichopus nigrilineatus|[3978]|JWDOL315-10|BOLD:AAG9624
Dolichopus nigrilineatus|[3979]|JWDOL341-10|BOLD:AAG9624
Dolichopus nigrilineatus|[3980]|MTDIC010-09|BOLD:AAG9624
Dolichopus nigrilineatus|[3981]|JWDOL304-10|BOLD:AAG9624
Dolichopus nigrilineatus|[3982]|JWDOL370-10|BOLD:AAG9624
Dolichopus nigrilineatus|[3983]|JWDOL414-10|BOLD:AAG9624
Dolichopus nigrilineatus|[3984]|JWDCG793-10|BOLD:AAG9624
Dolichopus nigrilineatus|[3985]|JWDOL167-10|BOLD:AAG9624
Dolichopus nigrilineatus|[3986]|JWDOL371-10|BOLD:AAG9624
Dolichopus nigrilineatus|[3987]|JWDOL401-10|BOLD:AAG9624
Dolichopus nigrilineatus|[3988]|JWDCH547-10|BOLD:AAG9624
Dolichopus nigrilineatus|[3989]|JWDOL310-10|BOLD:AAG9624
Dolichopus nigrilineatus|[3990]|JWDOL474-10|BOLD:AAG9624
Dolichopus nigrilineatus|[3991]|JWDOL440-10|BOLD:AAG9624
Dolichopus nigrilineatus|[3992]|JWDOL383-10|BOLD:AAG9624
Dolichopus nigrilineatus|[3993]|JWDCI549-10|BOLD:AAG9624
Dolichopus nigrilineatus|[3994]|JWDCG674-10|BOLD:AAG9624
Dolichopus nigrilineatus|[3995]|MHFLI958-09|BOLD:AAG9624
Dolichopus nigrilineatus|[3996]|JWDCA027-10|BOLD:AAG9624
Dolichopus nigrilineatus|[3997]|JWDCH347-10|BOLD:AAG9624
Dolichopus nigrilineatus|[3998]|JWDCH348-10|BOLD:AAG9624
Dolichopus nigrilineatus|[3999]|JWDCI225-10|BOLD:AAG9624
Dolichopus nigrilineatus|[4000]|JWDCF942-10|BOLD:AAG9624
Dolichopus nigrilineatus|[4001]|JWDOL1009-10|BOLD:AAG9624
Dolichopus nigrilineatus|[4002]|JWDOL132-10|BOLD:AAG9624
Dolichopus nigrilineatus|[4003]|JWDOL134-10|BOLD:AAG9624
Dolichopus nigrilineatus|[4004]|JWDOL165-10|BOLD:AAG9624
Dolichopus nigrilineatus|[4005]|JWDOL171-10|BOLD:AAG9624
Dolichopus nigrilineatus|[4006]|JWDOL247-10|BOLD:AAG9624
Dolichopus nigrilineatus|[4007]|JWDOL248-10|BOLD:AAG9624
Dolichopus nigrilineatus|[4008]|JWDOL249-10|BOLD:AAG9624
Dolichopus nigrilineatus|[4009]|JWDOL256-10|BOLD:AAG9624
Dolichopus nigrilineatus|[4010]|JWDOL270-10|BOLD:AAG9624
Dolichopus nigrilineatus|[4011]|JWDOL297-10|BOLD:AAG9624
Dolichopus nigrilineatus|[4012]|JWDOL329-10|BOLD:AAG9624
Dolichopus nigrilineatus|[4013]|JWDOL368-10|BOLD:AAG9624
Dolichopus nigrilineatus|[4014]|JWDOL397-10|BOLD:AAG9624
Dolichopus nigrilineatus|[4015]|JWDOL409-10|BOLD:AAG9624
Dolichopus nigrilineatus|[4016]|JWDOL415-10|BOLD:AAG9624
Dolichopus nigrilineatus|[4017]|JWDOL465-10|BOLD:AAG9624
Dolichopus nigrilineatus|[4018]|JWDOL805-10|BOLD:AAG9624
Dolichopus nigrilineatus|[4019]|JWDOL806-10|BOLD:AAG9624
Dolichopus nigrilineatus|[4020]|JWDOL902-10|BOLD:AAG9624
Dolichopus sp. SEB01|[4021]|JWDOL1010-10|BOLD:ABZ0681
Dolichopus shelfordi|[4022]|JWDOL812-10|BOLD:ABY8812
Dolichopus shelfordi|[4023]|JWDOL946-10|BOLD:ABY8812
Dolichopus shelfordi|[4024]|JWDOL129-10|BOLD:ABY8812
Dolichopus shelfordi|[4025]|JWDOL814-10|BOLD:ABY8812
Dolichopus shelfordi|[4026]|JWDCG744-10|BOLD:ABY8812
Dolichopus shelfordi|[4027]|JWDOL299-10|BOLD:ABY8812
Dolichopus shelfordi|[4028]|JWDOL896-10|BOLD:ABY8812
Dolichopus shelfordi|[4029]|JWDOL984-10|BOLD:ABY8812
Dolichopus shelfordi|[4030]|JWDOL143-10|BOLD:ABY8812
Dolichopus shelfordi|[4031]|JWDCA008-10|BOLD:ABY8812
Dolichopus shelfordi|[4032]|JWDOL1034-10|BOLD:ABY8812
Dolichopus shelfordi|[4033]|JWDOL280-10|BOLD:ABY8812
Dolichopus shelfordi|[4034]|JWDCH034-10|BOLD:ABY8812
Dolichopus shelfordi|[4035]|JWDOL970-10|BOLD:ABY8812
Dolichopus shelfordi|[4036]|JWDOL909-10|BOLD:ABY8812
Dolichopus shelfordi|[4037]|JWDOL034-10|BOLD:AFA6974
Dolichopus shelfordi|[4038]|JWDOL036-10|BOLD:AFA6974
Dolichopus shelfordi|[4039]|JWDOL037-10|BOLD:AFA6974
Dolichopus shelfordi|[4040]|JWDOL159-10|BOLD:AEX2014
Dolichopus shelfordi|[4041]|JWDOL231-10|BOLD:AEX2015
Dolichopus shelfordi|[4042]|JWDCF944-10
Dolichopus shelfordi|[4043]|JWDOL811-10|BOLD:AAM6770
Dolichopus shelfordi|[4044]|JWDOL945-10|BOLD:AAM6770
Dolichopus shelfordi|[4045]|JWDOL309-10|BOLD:AAM6770
Dolichopus shelfordi|[4046]|JWDCA009-10|BOLD:AAM6770
Dolichopus shelfordi|[4047]|JWDOL311-10|BOLD:AAM6770
Dolichopus shelfordi|[4048]|JWDOL366-10|BOLD:AAM6770
Dolichopus shelfordi|[4049]|JWDOL377-10|BOLD:AAM6770
Dolichopus shelfordi|[4050]|JWDOL344-10|BOLD:AAM6770
Dolichopus shelfordi|[4051]|JWDOL083-10|BOLD:AAM6770
Dolichopus shelfordi|[4052]|JWDOL374-10|BOLD:AAM6770
Dolichopus shelfordi|[4053]|JWDOL156-10|BOLD:AAM6770
Dolichopus shelfordi|[4054]|JWDCA007-10|BOLD:AAM6770
Dolichopus shelfordi|[4055]|JWDCH125-10|BOLD:AAM6770
Dolichopus shelfordi|[4056]|JWDCG385-10|BOLD:AAM6770
Dolichopus shelfordi|[4057]|JWDOL032-10|BOLD:AAM6770
Dolichopus shelfordi|[4058]|JWDOL441-10|BOLD:AAM6770
Dolichopus shelfordi|[4059]|JWDOL464-10|BOLD:AAM6770
Dolichopus shelfordi|[4060]|JWDOL168-10|BOLD:AAM6770
Dolichopus shelfordi|[4061]|JWDOL404-10|BOLD:AAM6770
Dolichopus shelfordi|[4062]|JWDCG024-10|BOLD:AAM6770
Dolichopus shelfordi|[4063]|JWDOL157-10|BOLD:AAM6770
Dolichopus shelfordi|[4064]|JWDOL419-10|BOLD:AAM6770
Dolichopus shelfordi|[4065]|JWDOL449-10|BOLD:AAM6770
Dolichopus shelfordi|[4066]|JWDOL455-10|BOLD:AAM6770
Dolichopus shelfordi|[4067]|JWDOL886-10|BOLD:AAM6770
Dolichopus shelfordi|[4068]|JWDOL899-10|BOLD:AAM6770
Dolichopus shelfordi|[4069]|JWDOL963-10|BOLD:AAM6770
Dolichopus sp. SEB02|[4070]|JWDOL287-10|BOLD:AAP3629
Dolichopus sp. SEB02|[4071]|JWDOL316-10|BOLD:AAP3629
Dolichopus sp. SEB02|[4072]|JWDOL417-10|BOLD:AAP3629
Dolichopus sp. SEB02|[4073]|JWDOL471-10|BOLD:AAP3629
Dolichopus sp. SEB02|[4074]|JWDOL901-10|BOLD:AAP3629
Dolichopus sp. nr. vanduzeei|[4075]|JWDOL360-10|BOLD:AAP3631
Dolichopus sp. nr. vanduzeei|[4076]|JWDOL392-10|BOLD:AAP3631
Dolichopus sp. nr. vanduzeei|[4077]|JWDOL406-10|BOLD:AAP3631
Dolichopus sp. nr. vanduzeei|[4078]|JWDOL1006-10|BOLD:AAP3631
Dolichopus sp. nr. vanduzeei|[4079]|JWDOL405-10|BOLD:AAP3631
Dolichopus sp. nr. vanduzeei|[4080]|JWDOL473-10|BOLD:AAP3631

Dolichopus sp. nr. vanduzeei|[4078]|JWDOL1006-10|BOLD:AAP3631
Dolichopus sp. nr. vanduzeei|[4079]|JWDOL405-10|BOLD:AAP3631
Dolichopus sp. nr. vanduzeei|[4080]|JWDOL473-10|BOLD:AAP3631
Dolichopus sp. nr. vanduzeei|[4081]|JWDOL359-10|BOLD:AAP3631
Dolichopus sp. nr. vanduzeei|[4082]|JWDCH252-10|BOLD:AAP3631
Dolichopus sp. nr. vanduzeei|[4083]|JWDOL1004-10|BOLD:AAP3631
Dolichopus sp. nr. vanduzeei|[4084]|JWDOL184-10|BOLD:AAP3631
Dolichopus sp. nr. vanduzeei|[4085]|JWDOL395-10|BOLD:AAP3631
Dolichopus sp. nr. vanduzeei|[4086]|JWDOL407-10|BOLD:AAP3631
Dolichopus sp. nr. vanduzeei|[4087]|JWDOL900-10|BOLD:AAP3631
Dolichopus sp. nr. vanduzeei|[4088]|JWDOL964-10|BOLD:AAP3631
Laphria|[4089]|MTDIC205-09|BOLD:AAI1265
Coelopa frigida|[4090]|TWDIP195-09|BOLD:AAG0822
Coelopa frigida|[4091]|TWDIP197-09|BOLD:AAG0822
Coelopa frigida|[4092]|TWDIP196-09|BOLD:AAG0822
Coelopa frigida|[4093]|SWSWP047-09|BOLD:AAG0822
Coelopa frigida|[4094]|MTDIC232-09|BOLD:AAG0822
Coelopa frigida|[4095]|MTDIC270-09|BOLD:AAG0822
Coelopa frigida|[4096]|MHFLY040-06|BOLD:AAG0822
Coelopa frigida|[4097]|MHFLY041-06|BOLD:AAG0822
Coelopa frigida|[4098]|MHFLY047-06|BOLD:AAG0822
Coelopa frigida|[4099]|JWDCA600-10|BOLD:AAG0822
Coelopa frigida|[4100]|JWDCA601-10|BOLD:AAG0822
Coelopa frigida|[4101]|JWDCA602-10|BOLD:AAG0822
Coelopa frigida|[4102]|JWDCA603-10|BOLD:AAG0822
Coelopa frigida|[4103]|MHFLY039-06|BOLD:AAG0822
Coelopa frigida|[4104]|MHFLY045-06|BOLD:AAG0822
Coelopa frigida|[4105]|JWDCJ1123-11|BOLD:AAG0822
Coelopa frigida|[4106]|JWDCE257-10|BOLD:AAG0822
Coelopa frigida|[4107]|JWDCA606-10|BOLD:AAG0822
Herina nigribasis|[4108]|JWDCJ977-11|BOLD:AAG0259
Herina nigribasis|[4109]|JWDCJ976-11|BOLD:AAG0259
Herina nigribasis|[4110]|JWDCJ1577-11|BOLD:AAG0259
Herina nigribasis|[4111]|JWDCJ1576-11|BOLD:AAG0259
Herina nigribasis|[4112]|JWDCJ1575-11|BOLD:AAG0259
Herina nigribasis|[4113]|JWDCJ901-11|BOLD:AAG0259
Herina nigribasis|[4114]|JWDCJ899-11|BOLD:AAG0259
Herina nigribasis|[4115]|JWDCJ075-11|BOLD:AAG0259
Herina nigribasis|[4116]|JWDCK235-11|BOLD:AAG0259
Herina nigribasis|[4117]|JWDCK234-11|BOLD:AAG0259
Herina nigribasis|[4118]|JWDCK233-11|BOLD:AAG0259
Herina nigribasis|[4119]|JWDCJ014-11|BOLD:AAG0259
Herina nigribasis|[4120]|JWDCA562-10|BOLD:AAG0259
Herina nigribasis|[4121]|JWDCJ978-11|BOLD:AAG0259
Herina nigribasis|[4122]|JWDCJ1368-11|BOLD:AAG0259
Herina nigribasis|[4123]|JWDCJ1369-11|BOLD:AAG0259
Herina nigribasis|[4124]|JWDCJ1449-11|BOLD:AAG0259
Herina nigribasis|[4125]|JWDCJ1450-11|BOLD:AAG0259
Herina nigribasis|[4126]|JWDCJ074-11|BOLD:AAG0259
Herina nigribasis|[4127]|JWDCF189-10|BOLD:AAG0259
Herina nigribasis|[4128]|JWDCF252-10|BOLD:AAG0259
Ulidiidae|[4129]|MTDIC278-09|BOLD:AAG6982
Graphomya|[4130]|MFDC1064-11|BOLD:AAG1735
Graphomya|[4131]|MFDC1061-11|BOLD:AAG1735
Graphomya minor|[4132]|TWDIP075-09|BOLD:AAG1735
Graphomya minor|[4133]|MFDC1065-11|BOLD:AAG1735
Graphomya minor|[4134]|MFDC1062-11|BOLD:AAG1735
Graphomya minor|[4135]|MFDC1058-11|BOLD:AAG1735
Graphomya minor|[4136]|MFDC1056-11|BOLD:AAG1735
Graphomya minor|[4137]|MFDC1052-11|BOLD:AAG1735
Graphomya minor|[4138]|MFDC1050-11|BOLD:AAG1735
Graphomya minor|[4139]|MFDC1048-11|BOLD:AAG1735
Graphomya minor|[4140]|MFDC1045-11|BOLD:AAG1735
Graphomya|[4141]|JWDCE892-10|BOLD:AAG1735
Graphomya|[4142]|JWDCF557-10|BOLD:AAG1735
Graphomya|[4143]|JWDCF604-10|BOLD:AAG1735
Graphomya|[4144]|JWDCF357-10|BOLD:AAG1735
Graphomya|[4145]|JWDCF316-10|BOLD:AAG1735
Graphomya|[4146]|JWDCI235-10|BOLD:AAG1735
Graphomya|[4147]|JWDCH400-10|BOLD:AAG1735
Graphomya|[4148]|JWDCG206-10|BOLD:AAG1735
Graphomya minor|[4149]|MFDC1046-11|BOLD:AAG1735
Graphomya|[4150]|JWDCJ905-11|BOLD:AAG1735
Graphomya|[4151]|JWDCJ1018-11|BOLD:AAG1735
Graphomya minor|[4152]|TWDIP204-09|BOLD:AAG1735
Graphomya|[4153]|MFDC1055-11|BOLD:AAG1735
Graphomya|[4154]|MFDC1066-11|BOLD:AAG1735
Graphomya minor|[4155]|MFDC1053-11|BOLD:AAG1735
Graphomya minuta|[4156]|JWDCG741-10|BOLD:ABY7767
Graphomya|[4157]|MFDC1063-11|BOLD:ABY7767
Graphomya cf. ungava|[4158]|JWDCA524-10|BOLD:ABZ7314
Graphomya cf. ungava|[4159]|JWDCG207-10|BOLD:ABZ7314
Graphomya cf. ungava|[4160]|JWDCG208-10|BOLD:ABZ7314
Graphomya cf. ungava|[4161]|JWDCF842-10|BOLD:ABZ7314
Graphomya cf. ungava|[4162]|MFDC1060-11|BOLD:ABZ7314
Graphomya cf. ungava|[4163]|MFDC1057-11|BOLD:ABZ7314
Hydrotaea aenescens|[4164]|MFDC159-09|BOLD:AAK3419
Hydrotaea aenescens|[4165]|MFDC160-09|BOLD:AAK3419
Hydrotaea aenescens|[4166]|MFDC737-10|BOLD:AAK3419
Hydrotaea aenescens|[4167]|MFDC738-10|BOLD:AAK3419
Brachicoma devia|[4168]|JWDCC048-10|BOLD:AAG2171
Brachicoma devia|[4169]|TWDIP126-09|BOLD:AAG2171
Brachicoma devia|[4170]|JWDCA506-10|BOLD:AAG2171
Brachicoma|[4171]|JWDCA507-10|BOLD:AAG2428
Brachicoma|[4172]|JWDCB052-10|BOLD:AAG2428
Brachicoma|[4173]|JWDCC854-10|BOLD:AAG2428
Brachicoma|[4174]|JWDCC802-10|BOLD:AAG2428
Brachicoma|[4175]|JWDCC855-10|BOLD:AAG2428
Brachicoma|[4176]|JWDCJ088-11|BOLD:AAG2428
Brachicoma|[4177]|JWDCC039-10|BOLD:AAG2428
Brachicoma|[4178]|JWDCI634-10|BOLD:AAG2428
Myospila meditabunda|[4179]|JWDCC272-10|BOLD:AAD7145
[illegible]

Brachicoma|[4177]|JWDCC039-10|BOLD:AAG2428
Brachicoma|[4178]|JWDCI634-10|BOLD:AAG2428
Myospila meditabunda|[4179]|JWDCC272-10|BOLD:AAD7145
Myospila meditabunda|[4180]|MFDC627-09|BOLD:AAD7145
Myospila meditabunda|[4181]|MFDC628-09|BOLD:AAD7145
Myospila meditabunda|[4182]|MFDC626-09|BOLD:AAD7145
Myospila meditabunda|[4183]|JWDCC159-10|BOLD:AAD7145
Myospila meditabunda|[4184]|MFDC629-09|BOLD:AAD7145
Myospila meditabunda|[4185]|MFDC764-10|BOLD:AAD7145
Ochthera mantis group|[4186]|JWDCK044-11|BOLD:AAI3530
Ochthera mantis group|[4187]|JWDCJ485-11|BOLD:AAI3530
Ochthera mantis group|[4188]|MHFLI962-09|BOLD:AAI3530
Ochthera mantis group|[4189]|MHFLI900-09|BOLD:AAI3530
Ochthera mantis group|[4190]|JWDCK043-11|BOLD:AAI3530
Ochthera mantis group|[4191]|JWDCG455-10|BOLD:AAI3530
Ochthera mantis group|[4192]|JWDCG454-10|BOLD:AAI3530
Ochthera mantis group|[4193]|JWDCG705-10|BOLD:AAI3530
Ochthera mantis group|[4194]|JWDCL312-11|BOLD:AAI3530
Ochthera mantis group|[4195]|JWDCI517-10|BOLD:AAI3530
Ochthera mantis group|[4196]|MHFLI212-06|BOLD:AAI3530
Ochthera mantis group|[4197]|JWDCL454-11|BOLD:AAI3530
Ochthera mantis group|[4198]|JWDCG581-10|BOLD:AAI3530
Ochthera mantis group|[4199]|JWDCE297-10|BOLD:AAI3530
Ochthera mantis group|[4200]|JWDCE359-10|BOLD:AAI3530
Ochthera mantis group|[4201]|JWDCE360-10|BOLD:AAI3530
Ochthera mantis group|[4202]|JWDCE021-10|BOLD:AAI3530
Lophosceles frenatus|[4203]|JWDCA433-10|BOLD:AAE1843
Lophosceles frenatus|[4204]|MFDC970-10|BOLD:AAE1843
Lophosceles frenatus|[4205]|MFDC637-09|BOLD:AAE1843
Lophosceles frenatus|[4206]|MFDC638-09|BOLD:AAE1843
Lophosceles frenatus|[4207]|MFDC639-09|BOLD:AAE1843
Lophosceles frenatus|[4208]|MFDC640-09|BOLD:AAE1843
Lophosceles frenatus|[4209]|MFDC644-09|BOLD:AAE1843
Lophosceles impar|[4210]|MFDC641-09|BOLD:AAK6862
Lophosceles impar|[4211]|MFDC751-10|BOLD:AAK6862
Lophosceles cinereiventris|[4212]|MFDC643-09|BOLD:AAI1152
Lophosceles cinereiventris|[4213]|MFDC642-09|BOLD:AAI1152
Lophosceles cinereiventris|[4214]|MFDC749-10|BOLD:AAI1152
Lophosceles cinereiventris|[4215]|MFDC750-10|BOLD:AAI1152
Lophosceles cinereiventris|[4216]|MFDC752-10|BOLD:AAI1152
Fannia|[4217]|DSANN027-08|BOLD:AAG6828
Fannia|[4218]|DSANN029-08|BOLD:AAG6828
Cephenemyia trompe|[4219]|JSFLA041-08|BOLD:AAH0838
Cephenemyia trompe|[4220]|JSFLA042-08|BOLD:AAH0838
Musca domestica|[4221]|MFDC645-09|BOLD:AAA6020
Musca domestica|[4222]|MFDC646-09|BOLD:AAA6020
Musca domestica|[4223]|MFDC647-09|BOLD:AAA6020
Musca domestica|[4224]|MFDC648-09|BOLD:AAA6020
Limnophora groenlandica|[4225]|MFDC609-09|BOLD:AAC6873
Limnophora groenlandica|[4226]|MFDC528-09|BOLD:AAC6873
Limnophora groenlandica|[4227]|MFDC608-09|BOLD:AAC6873
Limnophora groenlandica|[4228]|MFDC531-09|BOLD:AAC6873
Limnophora groenlandica|[4229]|MFDC529-09|BOLD:AAC6873
Limnophora groenlandica|[4230]|MFDC530-09|BOLD:AAC6873
Limnophora groenlandica|[4231]|MFDC556-09|BOLD:AAC6873
Limnophora groenlandica|[4232]|MFDC554-09|BOLD:AAC6873
Limnophora groenlandica|[4233]|MFDC605-09|BOLD:AAC6873
Limnophora groenlandica|[4234]|MFDC620-09|BOLD:AAC6873
Limnophora groenlandica|[4235]|MFDC093-08|BOLD:AAC6873
Limnophora groenlandica|[4236]|MFDC555-09|BOLD:AAC6873
Limnophora uniseta|[4237]|JWDCE496-10|BOLD:AAB9421
Limnophora uniseta|[4238]|JWDCC274-10|BOLD:AAB9421
Limnophora uniseta|[4239]|JWDCC385-10|BOLD:AAB9421
Limnophora uniseta|[4240]|JWDCC427-10|BOLD:AAB9421
Limnophora uniseta|[4241]|JWDCE666-10|BOLD:AAB9421
Limnophora uniseta|[4242]|MFDC552-09|BOLD:AAB9421
Limnophora uniseta|[4243]|JWDCI313-10|BOLD:AAB9421
Limnophora uniseta|[4244]|JWDCF523-10|BOLD:AAB9421
Limnophora uniseta|[4245]|JWDCJ913-11|BOLD:AAB9421
Limnophora uniseta|[4246]|SWSWP030-09|BOLD:AAB9421
Limnophora uniseta|[4247]|JWDCJ955-11|BOLD:AAB9421
Limnophora uniseta|[4248]|JWDCC391-10|BOLD:AAB9421
Limnophora uniseta|[4249]|JWDCC387-10|BOLD:AAB9421
Limnophora uniseta|[4250]|JWDCC025-10|BOLD:AAB9421
Limnophora uniseta|[4251]|JWDCC004-10|BOLD:AAB9421
Limnophora uniseta|[4252]|JWDCD183-10|BOLD:AAB9421
Limnophora uniseta|[4253]|JWDCC250-10|BOLD:AAB9421
Limnophora uniseta|[4254]|JWDCC392-10|BOLD:AAB9421
Limnophora uniseta|[4255]|JWDCC401-10|BOLD:AAB9421
Limnophora uniseta|[4256]|JWDCC410-10|BOLD:AAB9421
Limnophora uniseta|[4257]|JWDCC415-10|BOLD:AAB9421
Limnophora uniseta|[4258]|JWDCC430-10|BOLD:AAB9421
Limnophora uniseta|[4259]|JWDCJ810-11|BOLD:AAB9421
Limnophora uniseta|[4260]|JWDCJ1171-11|BOLD:AAB9421
Limnophora uniseta|[4261]|JWDCI293-10|BOLD:AAB9421
Limnophora uniseta|[4262]|JWDCI319-10|BOLD:AAB9421
Limnophora uniseta|[4263]|JWDCE497-10|BOLD:AAB9421
Limnophora uniseta|[4264]|JWDCE692-10|BOLD:AAB9421
Limnophora uniseta|[4265]|JWDCF497-10|BOLD:AAB9421
Limnophora uniseta|[4266]|JWDCF499-10|BOLD:AAB9421
Limnophora uniseta|[4267]|JWDCE834-10|BOLD:AAB9421
Limnophora uniseta|[4268]|JWDCE838-10|BOLD:AAB9421
Limnophora uniseta|[4269]|JWDCF692-10|BOLD:AAB9421
Limnophora uniseta|[4270]|JWDCE855-10|BOLD:AAB9421
Limnophora uniseta|[4271]|MFDC597-09|BOLD:AAB9421
Limnophora uniseta|[4272]|MFDC599-09|BOLD:AAB9421
Limnophora uniseta|[4273]|MFDC600-09|BOLD:AAB9421
Limnophora uniseta|[4274]|MFDC615-09|BOLD:AAB9421
Limnophora uniseta|[4275]|JWDCJ799-11|BOLD:AAB9421
Limnophora uniseta|[4276]|JWDCF013-10|BOLD:AAB9421
Limnophora uniseta|[4277]|JWDCC243-10|BOLD:AAB9421
Limnophora uniseta|[4278]|MFDC527-09|BOLD:AAB9421
Limnophora uniseta|[4279]|JWDCG188-10|BOLD:AAB9421

Limnophora uniseta|[4277]|JWDCC243-10|BOLD:AAB9421
Limnophora uniseta|[4278]|MFDC527-09|BOLD:AAB9421
Limnophora uniseta|[4279]|JWDCG188-10|BOLD:AAB9421
Limnophora uniseta|[4280]|JWDCD181-10|BOLD:AAB9421
Limnophora uniseta|[4281]|JWDCI291-10|BOLD:AAB9421
Limnophora uniseta|[4282]|MFDC616-09|BOLD:AAB9421
Limnophora uniseta|[4283]|JWDCC424-10|BOLD:AAB9421
Limnophora uniseta|[4284]|JWDCC426-10|BOLD:AAB9421
Limnophora uniseta|[4285]|JWDCC425-10|BOLD:AAB9421
Limnophora uniseta|[4286]|JWDCC260-10|BOLD:AAB9421
Limnophora uniseta|[4287]|JWDCC026-10|BOLD:AAB9421
Limnophora uniseta|[4288]|MFDC525-09|BOLD:AAB9421
Limnophora uniseta|[4289]|MFDC018-08|BOLD:AAB9421
Limnophora uniseta|[4290]|MFDC606-09|BOLD:AAB9421
Limnophora uniseta|[4291]|MFDC598-09|BOLD:AAB9421
Limnophora uniseta|[4292]|MFDC596-09|BOLD:AAB9421
Limnophora uniseta|[4293]|MFDC524-09|BOLD:AAB9421
Limnophora uniseta|[4294]|JWDCE175-10|BOLD:AAB9421
Limnophora uniseta|[4295]|JWDCE854-10|BOLD:AAB9421
Limnophora uniseta|[4296]|JWDCF843-10|BOLD:AAB9421
Limnophora uniseta|[4297]|JWDCF745-10|BOLD:AAB9421
Limnophora uniseta|[4298]|JWDCF741-10|BOLD:AAB9421
Limnophora uniseta|[4299]|JWDCF739-10|BOLD:AAB9421
Limnophora uniseta|[4300]|JWDCF642-10|BOLD:AAB9421
Limnophora uniseta|[4301]|JWDCE835-10|BOLD:AAB9421
Limnophora uniseta|[4302]|JWDCE812-10|BOLD:AAB9421
Limnophora uniseta|[4303]|JWDCF089-10|BOLD:AAB9421
Limnophora uniseta|[4304]|JWDCF495-10|BOLD:AAB9421
Limnophora uniseta|[4305]|JWDCF494-10|BOLD:AAB9421
Limnophora uniseta|[4306]|JWDCF493-10|BOLD:AAB9421
Limnophora uniseta|[4307]|JWDCF470-10|BOLD:AAB9421
Limnophora uniseta|[4308]|JWDCE706-10|BOLD:AAB9421
Limnophora uniseta|[4309]|JWDCE696-10|BOLD:AAB9421
Limnophora uniseta|[4310]|JWDCE687-10|BOLD:AAB9421
Limnophora uniseta|[4311]|JWDCE495-10|BOLD:AAB9421
Limnophora uniseta|[4312]|JWDCF117-10|BOLD:AAB9421
Limnophora uniseta|[4313]|JWDCF088-10|BOLD:AAB9421
Limnophora uniseta|[4314]|JWDCI425-10|BOLD:AAB9421
Limnophora uniseta|[4315]|JWDCI318-10|BOLD:AAB9421
Limnophora uniseta|[4316]|JWDCI316-10|BOLD:AAB9421
Limnophora uniseta|[4317]|JWDCI315-10|BOLD:AAB9421
Limnophora uniseta|[4318]|JWDCI314-10|BOLD:AAB9421
Limnophora uniseta|[4319]|JWDCI312-10|BOLD:AAB9421
Limnophora uniseta|[4320]|JWDCG189-10|BOLD:AAB9421
Limnophora uniseta|[4321]|JWDCI394-10|BOLD:AAB9421
Limnophora uniseta|[4322]|JWDCL139-11|BOLD:AAB9421
Limnophora uniseta|[4323]|JWDCJ312-11|BOLD:AAB9421
Limnophora uniseta|[4324]|JWDCL571-11|BOLD:AAB9421
Limnophora uniseta|[4325]|JWDCJ748-11|BOLD:AAB9421
Limnophora uniseta|[4326]|JWDCJ749-11|BOLD:AAB9421
Limnophora uniseta|[4327]|JWDCC396-10|BOLD:AAB9421
Limnophora uniseta|[4328]|JWDCC262-10|BOLD:AAB9421
Limnophora uniseta|[4329]|JWDCC254-10|BOLD:AAB9421
Limnophora uniseta|[4330]|JWDCC253-10|BOLD:AAB9421
Limnophora uniseta|[4331]|JWDCC248-10|BOLD:AAB9421
Limnophora uniseta|[4332]|JWDCC237-10|BOLD:AAB9421
Limnophora uniseta|[4333]|JWDCC126-10|BOLD:AAB9421
Limnophora uniseta|[4334]|JWDCC027-10|BOLD:AAB9421
Limnophora uniseta|[4335]|JWDCC018-10|BOLD:AAB9421
Limnophora uniseta|[4336]|JWDCC404-10|BOLD:AAB9421
Limnophora uniseta|[4337]|JWDCC406-10|BOLD:AAB9421
Limnophora uniseta|[4338]|JWDCC249-10|BOLD:AAB9421
Limnophora uniseta|[4339]|JWDCD025-10|BOLD:AAB9421
Limnophora uniseta|[4340]|JWDCD074-10|BOLD:AAB9421
Limnophora uniseta|[4341]|JWDCG185-10|BOLD:AAB9421
Limnophora uniseta|[4342]|JWDCE833-10|BOLD:AAB9421
Limnophora uniseta|[4343]|JWDCF115-10|BOLD:AAB9421
Limnophora uniseta|[4344]|MFDC622-09|BOLD:AAB9421
Limnophora uniseta|[4345]|MFDC523-09|BOLD:AAB9421
Limnophora sp. 1 AKR|[4346]|JWDCE517-10|BOLD:AAB2730
Limnophora sp. 1 AKR|[4347]|JWDCE839-10|BOLD:AAB2730
Limnophora sp. 1 AKR|[4348]|JWDCJ751-11|BOLD:AAB2730
Limnophora sp. 1 AKR|[4349]|JWDCJ1177-11|BOLD:AAB2730
Limnophora sp. 1 AKR|[4350]|JWDCE894-10|BOLD:AAB2730
Limnophora sp. 1 AKR|[4351]|MHFLY360-09|BOLD:AAB2730
Limnophora sp. 1 AKR|[4352]|JWDCI290-10|BOLD:AAB2730
Limnophora sp. 1 AKR|[4353]|MFDC546-09|BOLD:AAB2730
Limnophora sp. 1 AKR|[4354]|MFDC604-09|BOLD:AAB2730
Limnophora sp. 1 AKR|[4355]|MFDC547-09|BOLD:AAB2730
Limnophora sp. 1 AKR|[4356]|MFDC545-09|BOLD:AAB2730
Limnophora sp. 1 AKR|[4357]|MFDC084-08|BOLD:AAB2730
Limnophora sp. 1 AKR|[4358]|MFDC079-08|BOLD:AAB2730
Limnophora sp. 1 AKR|[4359]|MFDC619-09|BOLD:AAB2730
Limnophora sp. 1 AKR|[4360]|MFDC618-09|BOLD:AAB2730
Limnophora sp. 1 AKR|[4361]|MFDC617-09|BOLD:AAB2730
Limnophora sp. 1 AKR|[4362]|MFDC614-09|BOLD:AAB2730
Limnophora sp. 1 AKR|[4363]|MFDC612-09|BOLD:AAB2730
Limnophora sp. 1 AKR|[4364]|MFDC611-09|BOLD:AAB2730
Limnophora sp. 1 AKR|[4365]|MFDC610-09|BOLD:AAB2730
Limnophora sp. 1 AKR|[4366]|MFDC607-09|BOLD:AAB2730
Limnophora sp. 1 AKR|[4367]|MFDC603-09|BOLD:AAB2730
Limnophora sp. 1 AKR|[4368]|MFDC602-09|BOLD:AAB2730
Limnophora sp. 1 AKR|[4369]|MFDC551-09|BOLD:AAB2730
Limnophora sp. 1 AKR|[4370]|MFDC550-09|BOLD:AAB2730
Limnophora sp. 1 AKR|[4371]|MFDC549-09|BOLD:AAB2730
Limnophora sp. 1 AKR|[4372]|MFDC548-09|BOLD:AAB2730
Limnophora sp. 1 AKR|[4373]|MFDC544-09|BOLD:AAB2730
Limnophora sp. 1 AKR|[4374]|MFDC543-09|BOLD:AAB2730
Limnophora sp. 1 AKR|[4375]|JWDCE286-10|BOLD:AAB2730
Limnophora sp. 1 AKR|[4376]|JWDCF451-10|BOLD:AAB2730
Limnophora sp. 1 AKR|[4377]|JWDCF448-10|BOLD:AAB2730
Limnophora sp. 1 AKR|[4378]|JWDCF437-10|BOLD:AAB2730
[illegible]

Limnophora sp. 1 AKR|[4376]|JWDCF431-10|BOLD:AAB2730
Limnophora sp. 1 AKR|[4377]|JWDCF448-10|BOLD:AAB2730
Limnophora sp. 1 AKR|[4378]|JWDCF437-10|BOLD:AAB2730
Limnophora sp. 1 AKR|[4379]|JWDCF282-10|BOLD:AAB2730
Limnophora sp. 1 AKR|[4380]|JWDCG190-10|BOLD:AAB2730
Limnophora sp. 1 AKR|[4381]|JWDCJ1186-11|BOLD:AAB2730
Limnophora sp. 1 AKR|[4382]|JWDCJ305-11|BOLD:AAB2730
Limnophora sp. 1 AKR|[4383]|JWDCJ755-11|BOLD:AAB2730
Limnophora sp. 1 AKR|[4384]|JWDCJ029-11|BOLD:AAB2730
Limnophora sp. 1 AKR|[4385]|JWDCJ122-11|BOLD:AAB2730
Limnophora sp. 1 AKR|[4386]|JWDCJ746-11|BOLD:AAB2730
Limnophora sp. 1 AKR|[4387]|JWDCJ750-11|BOLD:AAB2730
Limnophora sp. 1 AKR|[4388]|JWDCC428-10|BOLD:AAB2730
Limnophora sp. 1 AKR|[4389]|JWDCC409-10|BOLD:AAB2730
Limnophora sp. 1 AKR|[4390]|JWDCC240-10|BOLD:AAB2730
Limnophora sp. 1 AKR|[4391]|JWDCC239-10|BOLD:AAB2730
Limnophora sp. 1 AKR|[4392]|JWDCC187-10|BOLD:AAB2730
Limnophora sp. 1 AKR|[4393]|JWDCC021-10|BOLD:AAB2730
Limnophora sp. 1 AKR|[4394]|JWDCC011-10|BOLD:AAB2730
Limnophora sp. 1 AKR|[4395]|JWDCC242-10|BOLD:AAB2730
Limnophora sp. 1 AKR|[4396]|JWDCB133-10|BOLD:AAB2730
Limnophora sp. 1 AKR|[4397]|MHFLI213-06|BOLD:AAB2730
Limnophora sp. 1 AKR|[4398]|JWDCF581-10|BOLD:AAB2730
Limnophora sp. 1 AKR|[4399]|JWDCC157-10|BOLD:AAB2730
Limnophora sp. 1 AKR|[4400]|MFDC613-09|BOLD:AAB2730
Limnophora sp. 1 AKR|[4401]|MFDC601-09|BOLD:AAB2730
Limnophora sp. 1 AKR|[4402]|MFDC522-09|BOLD:AAB2730
Limnophora sp. 1 AKR|[4403]|MFDC588-09|BOLD:AAB2730
Limnophora|[4404]|JWDCC783-10|BOLD:AAC2702
Limnophora|[4405]|JWDCC429-10|BOLD:AAC2702
Limnophora sp. 2 AKR|[4406]|JWDCJ391-11|BOLD:AAC2702
Limnophora sp. 2 AKR|[4407]|JWDCI423-10|BOLD:AAC2702
Limnophora sp. 2 AKR|[4408]|JWDCL290-11|BOLD:AAC2702
Limnophora sp. 2 AKR|[4409]|JWDCJ872-11|BOLD:AAC2702
Limnophora sp. 2 AKR|[4410]|JWDCK263-11|BOLD:AAC2702
Limnophora|[4411]|MFDC573-09|BOLD:AAC2702
Limnophora|[4412]|MFDC592-09|BOLD:AAC2702
Limnophora sp. 2 AKR|[4413]|JWDCL282-11|BOLD:AAC2702
Limnophora|[4414]|MFDC577-09|BOLD:AAC2702
Limnophora|[4415]|MFDC590-09|BOLD:AAC2702
Limnophora sp. 2 AKR|[4416]|JWDCH867-10|BOLD:AAC2702
Limnophora|[4417]|MFDC594-09|BOLD:AAC2702
Limnophora|[4418]|MFDC586-09|BOLD:AAC2702
Limnophora|[4419]|MFDC587-09|BOLD:AAC2702
Limnophora|[4420]|MFDC576-09|BOLD:AAC2702
Limnophora|[4421]|MFDC578-09|BOLD:AAC2702
Limnophora|[4422]|MFDC571-09|BOLD:AAC2702
Limnophora|[4423]|MFDC575-09|BOLD:AAC2702
Limnophora sp. 2 AKR|[4424]|JWDCI648-10|BOLD:AAC2702
Limnophora sp. 2 AKR|[4425]|JWDCF336-10|BOLD:AAC2702
Limnophora sp. 2 AKR|[4426]|JWDCI647-10|BOLD:AAC2702
Limnophora sp. 2 AKR|[4427]|JWDCI646-10|BOLD:AAC2702
Limnophora|[4428]|JWDCH027-10|BOLD:AAC2702
Limnophora|[4429]|JWDCH793-10|BOLD:AAC2702
Limnophora sp. 2 AKR|[4430]|JWDCG595-10|BOLD:AAC2702
Limnophora sp. 2 AKR|[4431]|JWDCJ866-11|BOLD:AAC2702
Limnophora sp. 2 AKR|[4432]|JWDCJ754-11|BOLD:AAC2702
Limnophora|[4433]|JWDCC199-10|BOLD:AAC2702
Limnophora|[4434]|MTDIC125-09|BOLD:AAC2702
Limnophora|[4435]|MFDC572-09|BOLD:AAC2702
Limnophora|[4436]|MFDC593-09|BOLD:AAC2702
Limnophora|[4437]|MFDC591-09|BOLD:AAC2702
Limnophora|[4438]|MFDC574-09|BOLD:AAC2702
Limnophora rotundata|[4439]|MFDC585-09|BOLD:AAB9451
Limnophora rotundata|[4440]|JWDCC263-10|BOLD:AAB9451
Limnophora rotundata|[4441]|MFDC579-09|BOLD:AAB9451
Limnophora rotundata|[4442]|JWDCJ268-11|BOLD:AAB9451
Limnophora rotundata|[4443]|MFDC065-08|BOLD:AAB9451
Limnophora rotundata|[4444]|MFDC068-08|BOLD:AAB9451
Limnophora rotundata|[4445]|MFDC558-09|BOLD:AAB9451
Limnophora rotundata|[4446]|MFDC560-09|BOLD:AAB9451
Limnophora rotundata|[4447]|MFDC580-09|BOLD:AAB9451
Limnophora rotundata|[4448]|MFDC061-08|BOLD:AAB9451
Limnophora rotundata|[4449]|MFDC583-09|BOLD:AAB9451
Limnophora rotundata|[4450]|MFDC562-09|BOLD:AAB9451
Limnophora rotundata|[4451]|JWDCH040-10|BOLD:AAB9451
Limnophora rotundata|[4452]|MFDC559-09|BOLD:AAB9451
Limnophora rotundata|[4453]|JWDCF889-10|BOLD:AAB9451
Limnophora rotundata|[4454]|JWDCC416-10|BOLD:AAB9451
Limnophora rotundata|[4455]|JWDCA427-10|BOLD:AAB9451
Limnophora rotundata|[4456]|MFDC581-09|BOLD:AAB9451
Limnophora rotundata|[4457]|MFDC584-09|BOLD:AAB9451
Limnophora rotundata|[4458]|MFDC595-09|BOLD:AAB9451
Limnophora rotundata|[4459]|MFDC621-09|BOLD:AAB9451
Limnophora rotundata|[4460]|MFDC057-08|BOLD:AAB9451
Limnophora rotundata|[4461]|MFDC561-09|BOLD:AAB9451
Limnophora rotundata|[4462]|MFDC589-09|BOLD:AAB9451
Lispe uliginosa|[4463]|MFDC210-09|BOLD:AAD3484
Lispe uliginosa|[4464]|MFDC211-09|BOLD:AAD3484
Lispe uliginosa|[4465]|MFDC212-09|BOLD:AAD3484
Lispe uliginosa|[4466]|MFDC213-09|BOLD:AAD3484
Lispe uliginosa|[4467]|MFDC214-09|BOLD:AAD3484
Lispe uliginosa|[4468]|MFDC215-09|BOLD:AAD3484
Lispe uliginosa|[4469]|JWDCJ678-11|BOLD:AAD3484
Lispe uliginosa|[4470]|JWDCH794-10|BOLD:AAD3484
Lispe uliginosa|[4471]|MHFLI907-09|BOLD:AAD3484
Lispe uliginosa|[4472]|MFDC080-08|BOLD:AAD3484
Lispe uliginosa|[4473]|MFDC209-09|BOLD:AAD3484
Lispe uliginosa|[4474]|MFDC208-09|BOLD:AAD3484
Lispe uliginosa|[4475]|MFDC207-09|BOLD:AAD3484
Lispe uliginosa|[4476]|MHFLI203-06|BOLD:AAD3484
Lispe uliginosa|[4477]|MHFLI220-06|BOLD:AAD3484
Lispe tentaculata|[4478]|MHFLI183-06|BOLD:AAB8430

Lispe uliginosa|[4476]|MHFLI203-06|BOLD:AAD3484
Lispe uliginosa|[4477]|MHFLI220-06|BOLD:AAD3484
Lispe tentaculata|[4478]|MHFLI183-06|BOLD:AAB8430
Lispe tentaculata|[4479]|MHFLY016-06|BOLD:AAB8430
Lispe tentaculata|[4480]|MHFLY129-07|BOLD:AAB8429
Lispe tentaculata|[4481]|JWDCG412-10|BOLD:AAB8429
Lispe tentaculata|[4482]|JWDCG228-10|BOLD:AAB8429
Lispe tentaculata|[4483]|MFDC206-09|BOLD:AAB8429
Lispe tentaculata|[4484]|MHFLY003-06|BOLD:AAB8429
Lispe tentaculata|[4485]|MFDC265-09|BOLD:AAB8429
Lispe tentaculata|[4486]|JWDCE619-10|BOLD:AAB8429
Lispe tentaculata|[4487]|MFDC205-09|BOLD:AAB8429
Lispe tentaculata|[4488]|JWDCC012-10|BOLD:AAB8429
Lispe tentaculata|[4489]|JWDCH507-10|BOLD:AAB8429
Lispe tentaculata|[4490]|JWDCI480-10|BOLD:AAB8429
Lispe tentaculata|[4491]|MHFLY354-09|BOLD:AAB8429
Lispe tentaculata|[4492]|MFDC047-08|BOLD:AAB8429
Lispe tentaculata|[4493]|JWDCH246-10|BOLD:AAB8429
Lispe tentaculata|[4494]|JWDCG870-10|BOLD:AAB8429
Lispe tentaculata|[4495]|JWDCG582-10|BOLD:AAB8429
Lispe tentaculata|[4496]|JWDCG279-10|BOLD:AAB8429
Lispe tentaculata|[4497]|MHFLY026-06|BOLD:AAB8429
Lispe frigida|[4498]|MFDC223-09|BOLD:AAD8875
Lispe frigida|[4499]|MFDC225-09|BOLD:AAD8875
Lispe frigida|[4500]|MFDC222-09|BOLD:AAD8875
Lispe frigida|[4501]|MFDC221-09|BOLD:AAD8875
Lispe frigida|[4502]|MFDC224-09|BOLD:AAD8875
Lispe frigida|[4503]|MHFLI129-06|BOLD:AAD8875
Lispe frigida|[4504]|MFDC746-10|BOLD:ACY4326
Lispe frigida|[4505]|MFDC747-10|BOLD:ACY4326
Lispe salina|[4506]|JWDCE037-10|BOLD:AAD8886
Lispe salina|[4507]|MFDC220-09|BOLD:AAD8886
Lispe salina|[4508]|JWDCH406-10|BOLD:AAD8886
Lispe salina|[4509]|MFDC218-09|BOLD:AAD8886
Lispe salina|[4510]|MFDC021-08|BOLD:AAD8886
Lispe salina|[4511]|MFDC219-09|BOLD:AAD8886
Lispe salina|[4512]|MFDC216-09|BOLD:AAD8886
Lispe salina|[4513]|MFDC217-09|BOLD:AAD8886
Lispe salina|[4514]|MFDC748-10|BOLD:AAD8886
Helina nigribasis|[4515]|MFDC883-10|BOLD:AAL5954
Helina nigribasis|[4516]|MFDC884-10|BOLD:AAL5954
Helina nigribasis|[4517]|MFDC885-10|BOLD:AAL5954
Helina nigribasis|[4518]|MFDC978-10|BOLD:AAL5954
Helina nigribasis|[4519]|JWDCK133-11|BOLD:AAL5954
Helina nigribasis|[4520]|JWDCF864-10|BOLD:AAL5954
Helina nigribasis|[4521]|MFDC979-10|BOLD:AAL5954
Helina nigribasis|[4522]|MFDC980-10|BOLD:AAL5954
Helina nigribasis|[4523]|MFDC981-10|BOLD:AAL5954
Helina nigribasis|[4524]|MFDC982-10|BOLD:AAL5954
Cylindromyia interrupta|[4525]|JWDCH021-10|BOLD:AAG2144
Helina reversio|[4526]|MFDC028-08|BOLD:AAD0642
Helina reversio|[4527]|JWDCC125-10|BOLD:AAD0642
Helina reversio|[4528]|JWDCJ1542-11|BOLD:AAD0642
Helina reversio|[4529]|JWDCE938-10|BOLD:AAD0642
Helina reversio|[4530]|JWDCC155-10|BOLD:AAD0642
Helina reversio|[4531]|JWDCC168-10|BOLD:AAD0642
Helina reversio|[4532]|JWDCC154-10|BOLD:AAD0642
Helina reversio|[4533]|JWDCC152-10|BOLD:AAD0642
Helina reversio|[4534]|JWDCC144-10|BOLD:AAD0642
Helina reversio|[4535]|JWDCC127-10|BOLD:AAD0642
Helina reversio|[4536]|JWDCC114-10|BOLD:AAD0642
Helina reversio|[4537]|JWDCC019-10|BOLD:AAD0642
Helina reversio|[4538]|JWDCC010-10|BOLD:AAD0642
Helina reversio|[4539]|MFDC096-09|BOLD:AAD0642
Helina reversio|[4540]|JWDCD122-10|BOLD:AAD0642
Helina reversio|[4541]|JWDCC877-10|BOLD:AAD0642
Helina reversio|[4542]|MFDC097-09|BOLD:AAD0642
Helina reversio|[4543]|MFDC095-09|BOLD:AAD0642
Helina reversio|[4544]|MFDC098-09|BOLD:AAD0642
Helina reversio|[4545]|JWDCC876-10|BOLD:AAD0642
Helina reversio|[4546]|MFDC009-08|BOLD:AAD0642
Phaonia apicalis|[4547]|MFDC358-09|BOLD:AAL2821
Phaonia apicalis|[4548]|MFDC359-09|BOLD:AAL2821
Phaonia|[4549]|JWDCJ1615-11|BOLD:AAQ3549
Phaonia atrocyanea|[4550]|MFDC342-09|BOLD:AAI4570
Phaonia atrocyanea|[4551]|MFDC343-09|BOLD:AAI4570
Phaonia atrocyanea|[4552]|MFDC767-10|BOLD:AAI4570
Mydaea occidentalis|[4553]|MFDC254-09|BOLD:AAE0058
Mydaea occidentalis|[4554]|MFDC253-09|BOLD:AAE0058
Mydaea occidentalis|[4555]|JWDCA473-10|BOLD:AAE0058
Mydaea occidentalis|[4556]|MFDC255-09|BOLD:AAE0058
Mydaea occidentalis|[4557]|MFDC256-09|BOLD:AAE0058
Mydaea obscurella|[4558]|JWDCC790-10|BOLD:AAC6709
Mydaea obscurella|[4559]|MFDC257-09|BOLD:AAC6709
Mydaea obscurella|[4560]|MFDC258-09|BOLD:AAC6709
Mydaea obscurella|[4561]|MFDC259-09|BOLD:AAC6709
Mydaea obscurella|[4562]|MFDC260-09|BOLD:AAC6709
Mydaea obscurella|[4563]|MFDC261-09|BOLD:AAC6709
Mydaea obscurella|[4564]|MFDC262-09|BOLD:AAC6709
Mydaea obscurella|[4565]|MFDC263-09|BOLD:AAC6709
Mydaea obscurella|[4566]|MFDC264-09|BOLD:AAC6709
Mydaea obscurella|[4567]|MFDC014-08|BOLD:AAC6709
Mydaea pseudonubila|[4568]|MFDC241-09|BOLD:AAE0077
Mydaea pseudonubila|[4569]|MFDC077-08|BOLD:AAE0077
Mydaea pseudonubila|[4570]|MFDC857-10|BOLD:AAE0077
Mydaea pseudonubila|[4571]|MFDC856-10|BOLD:AAE0077
Mydaea pseudonubila|[4572]|MFDC240-09|BOLD:AAE0077
Mydaea affinis|[4573]|JWDCG736-10|BOLD:AAD2891
Mydaea affinis|[4574]|JWDCF360-10|BOLD:AAD2891
Mydaea affinis|[4575]|MFDC251-09|BOLD:AAD2891
Mydaea affinis|[4576]|MFDC252-09|BOLD:AAD2891
Mydaea affinis|[4577]|MFDC755-10|BOLD:AAD2891
Mydaea affinis|[4578]|MFDC756-10|BOLD:AAD2891

Mydaea affinis|[4575]|MFDC251-09|BOLD:AAD2891
Mydaea affinis|[4576]|MFDC252-09|BOLD:AAD2891
Mydaea affinis|[4577]|MFDC755-10|BOLD:AAD2891
Mydaea affinis|[4578]|MFDC756-10|BOLD:AAD2891
Opsolasia orichalcea|[4579]|MFDC247-09|BOLD:AAE6151
Opsolasia orichalcea|[4580]|MFDC250-09|BOLD:AAE6151
Opsolasia orichalcea|[4581]|MFDC248-09|BOLD:AAE6151
Opsolasia orichalcea|[4582]|MFDC249-09|BOLD:AAE6151
Opsolasia orichalcea|[4583]|MFDC758-10|BOLD:AAE6151
Mydaea palpalis|[4584]|JWDCK266-11|BOLD:AAL7145
Mydaea palpalis|[4585]|MFDC757-10|BOLD:AAL7145
Mydaea palpalis|[4586]|MFDC759-10|BOLD:AAL7145
Mydaea furtiva|[4587]|JWDCJ1497-11|BOLD:AAG1779
Mydaea furtiva|[4588]|MFDC761-10|BOLD:AAG1779
Thricops diaphanus|[4589]|MFDC868-10|BOLD:AAG1710
Thricops diaphanus|[4590]|MFDC867-10|BOLD:AAG1710
Thricops diaphanus|[4591]|JWDCA462-10|BOLD:AAG1710
Thricops diaphanus|[4592]|MFDC869-10|BOLD:AAG1710
Thricops albibasalis|[4593]|MFDC285-09|BOLD:AAD2373
Thricops albibasalis|[4594]|MFDC026-08|BOLD:AAD2373
Thricops albibasalis|[4595]|JWDCJ121-11|BOLD:AAD2373
Thricops albibasalis|[4596]|MFDC870-10|BOLD:AAD2373
Thricops albibasalis|[4597]|MFDC069-08|BOLD:AAD2373
Thricops albibasalis|[4598]|JWDCJ1015-11|BOLD:AAD2373
Thricops albibasalis|[4599]|MFDC282-09|BOLD:AAD2373
Thricops albibasalis|[4600]|MFDC283-09|BOLD:AAD2373
Thricops albibasalis|[4601]|MFDC284-09|BOLD:AAD2373
Thricops albibasalis|[4602]|MFDC040-08|BOLD:AAD2373
Thricops albibasalis|[4603]|JWDCJ382-11|BOLD:AAD2373
Thricops albibasalis|[4604]|MFDC281-09|BOLD:AAD2373
Thricops hirtulus|[4605]|JWDCJ250-11|BOLD:ABY7837
Thricops hirtulus|[4606]|JWDCB131-10|BOLD:ABY7837
Thricops hirtulus|[4607]|JWDCA442-10|BOLD:ABY7837
Thricops hirtulus|[4608]|JWDCI913-11|BOLD:ABY7837
Thricops hirtulus|[4609]|JWDCC212-10|BOLD:ABY7837
Thricops hirtulus|[4610]|JWDCH313-10|BOLD:ABY7837
Thricops hirtulus|[4611]|JWDCJ264-11|BOLD:ABY7837
Thricops hirtulus|[4612]|JWDCJ1559-11|BOLD:ABY7837
Thricops hirtulus|[4613]|JWDCF280-10|BOLD:ABY7837
Thricops hirtulus|[4614]|JWDCI908-11|BOLD:ABY7837
Thricops hirtulus|[4615]|JWDCI904-11|BOLD:ABY7837
Thricops hirtulus|[4616]|JWDCI847-11|BOLD:ABY7837
Thricops hirtulus|[4617]|JWDCI839-11|BOLD:ABY7837
Thricops hirtulus|[4618]|JWDCI838-11|BOLD:ABY7837
Thricops hirtulus|[4619]|JWDCI688-11|BOLD:ABY7837
Thricops hirtulus|[4620]|JWDCI635-10|BOLD:ABY7837
Thricops hirtulus|[4621]|JWDCI573-10|BOLD:ABY7837
Thricops hirtulus|[4622]|JWDCH609-10|BOLD:ABY7837
Thricops hirtulus|[4623]|JWDCG848-10|BOLD:ABY7837
Thricops hirtulus|[4624]|JWDCG186-10|BOLD:ABY7837
Thricops hirtulus|[4625]|JWDCI393-10|BOLD:ABY7837
Thricops hirtulus|[4626]|JWDCH982-10|BOLD:ABY7837
Thricops hirtulus|[4627]|JWDCL428-11|BOLD:ABY7837
Thricops hirtulus|[4628]|JWDCL423-11|BOLD:ABY7837
Thricops hirtulus|[4629]|JWDCL346-11|BOLD:ABY7837
Thricops hirtulus|[4630]|JWDCL330-11|BOLD:ABY7837
Thricops hirtulus|[4631]|JWDCL437-11|BOLD:ABY7837
Thricops hirtulus|[4632]|JWDCL285-11|BOLD:ABY7837
Thricops hirtulus|[4633]|JWDCL283-11|BOLD:ABY7837
Thricops hirtulus|[4634]|JWDCL274-11|BOLD:ABY7837
Thricops hirtulus|[4635]|JWDCL072-11|BOLD:ABY7837
Thricops hirtulus|[4636]|JWDCJ1432-11|BOLD:ABY7837
Thricops hirtulus|[4637]|JWDCJ869-11|BOLD:ABY7837
Thricops hirtulus|[4638]|JWDCJ868-11|BOLD:ABY7837
Thricops hirtulus|[4639]|JWDCJ314-11|BOLD:ABY7837
Thricops hirtulus|[4640]|JWDCJ257-11|BOLD:ABY7837
Thricops hirtulus|[4641]|JWDCK261-11|BOLD:ABY7837
Thricops hirtulus|[4642]|JWDCK254-11|BOLD:ABY7837
Thricops hirtulus|[4643]|JWDCJ395-11|BOLD:ABY7837
Thricops hirtulus|[4644]|JWDCA443-10|BOLD:ABY7837
Thricops hirtulus|[4645]|JWDCL415-11|BOLD:ABY7837
Thricops hirtulus|[4646]|JWDCK264-11|BOLD:ABY7837
Thricops hirtulus|[4647]|JWDCJ1360-11|BOLD:ABY7837
Thricops hirtulus|[4648]|JWDCG832-10|BOLD:ABY7837
Thricops hirtulus|[4649]|JWDCI845-11|BOLD:ABY7837
Thricops hirtulus|[4650]|JWDCF359-10|BOLD:ABY7837
Thricops innocuus|[4651]|MFDC271-09|BOLD:ABY7837
Thricops hirtulus|[4652]|MFDC083-08|BOLD:ABY7837
Thricops hirtulus|[4653]|MFDC081-08|BOLD:ABY7837
Thricops hirtulus|[4654]|JWDCJ119-11|BOLD:ABY7837
Thricops hirtulus|[4655]|JWDCL289-11|BOLD:ABY7837
Thricops septentrionalis|[4656]|MFDC295-09|BOLD:ABY7837
Thricops innocuus|[4657]|MFDC270-09|BOLD:ABY7837
Thricops innocuus|[4658]|MFDC269-09|BOLD:ABY7837
Thricops innocuus|[4659]|MFDC268-09|BOLD:ABY7837
Thricops innocuus|[4660]|MFDC267-09|BOLD:ABY7837
Thricops hirtulus|[4661]|JWDCJ110-11|BOLD:ABY7837
Thricops hirtulus|[4662]|JWDCF891-10|BOLD:ABY7837
Thricops hirtulus|[4663]|JWDCF450-10|BOLD:ABY7837
Thricops hirtulus|[4664]|JWDCF405-10|BOLD:ABY7837
Thricops hirtulus|[4665]|JWDCH217-10|BOLD:ABY7837
Thricops hirtulus|[4666]|JWDCH215-10|BOLD:ABY7837
Thricops hirtulus|[4667]|JWDCI914-11|BOLD:ABY7837
Thricops hirtulus|[4668]|JWDCI907-11|BOLD:ABY7837
Thricops hirtulus|[4669]|JWDCI906-11|BOLD:ABY7837
Thricops hirtulus|[4670]|JWDCI905-11|BOLD:ABY7837
Thricops hirtulus|[4671]|JWDCI903-11|BOLD:ABY7837
Thricops hirtulus|[4672]|JWDCI841-11|BOLD:ABY7837
Thricops hirtulus|[4673]|JWDCI840-11|BOLD:ABY7837
Thricops hirtulus|[4674]|JWDCI740-11|BOLD:ABY7837
Thricops hirtulus|[4675]|JWDCI692-11|BOLD:ABY7837
Thricops hirtulus|[4676]|JWDCI687-11|BOLD:ABY7837
Thricops hirtulus|[4677]|JWDCI645-10|BOLD:ABY7837

Thricops hirtulus|[4675]|JWDCI692-11|BOLD:ABY7837
Thricops hirtulus|[4676]|JWDCI687-11|BOLD:ABY7837
Thricops hirtulus|[4677]|JWDCI645-10|BOLD:ABY7837
Thricops hirtulus|[4678]|JWDCI642-10|BOLD:ABY7837
Thricops hirtulus|[4679]|JWDCI640-10|BOLD:ABY7837
Thricops hirtulus|[4680]|JWDCI639-10|BOLD:ABY7837
Thricops hirtulus|[4681]|JWDCI571-10|BOLD:ABY7837
Thricops hirtulus|[4682]|JWDCH469-10|BOLD:ABY7837
Thricops hirtulus|[4683]|JWDCG285-10|BOLD:ABY7837
Thricops hirtulus|[4684]|JWDCH844-10|BOLD:ABY7837
Thricops hirtulus|[4685]|JWDCI392-10|BOLD:ABY7837
Thricops hirtulus|[4686]|JWDCI349-10|BOLD:ABY7837
Thricops hirtulus|[4687]|JWDCI348-10|BOLD:ABY7837
Thricops hirtulus|[4688]|JWDCI344-10|BOLD:ABY7837
Thricops hirtulus|[4689]|JWDCG093-10|BOLD:ABY7837
Thricops hirtulus|[4690]|JWDCG871-10|BOLD:ABY7837
Thricops hirtulus|[4691]|JWDCH843-10|BOLD:ABY7837
Thricops hirtulus|[4692]|JWDCL429-11|BOLD:ABY7837
Thricops hirtulus|[4693]|JWDCL427-11|BOLD:ABY7837
Thricops hirtulus|[4694]|JWDCL426-11|BOLD:ABY7837
Thricops hirtulus|[4695]|JWDCL424-11|BOLD:ABY7837
Thricops hirtulus|[4696]|JWDCL350-11|BOLD:ABY7837
Thricops hirtulus|[4697]|JWDCL349-11|BOLD:ABY7837
Thricops hirtulus|[4698]|JWDCL333-11|BOLD:ABY7837
Thricops hirtulus|[4699]|JWDCL332-11|BOLD:ABY7837
Thricops hirtulus|[4700]|JWDCL286-11|BOLD:ABY7837
Thricops hirtulus|[4701]|JWDCL284-11|BOLD:ABY7837
Thricops hirtulus|[4702]|JWDCL279-11|BOLD:ABY7837
Thricops hirtulus|[4703]|JWDCL277-11|BOLD:ABY7837
Thricops hirtulus|[4704]|JWDCL273-11|BOLD:ABY7837
Thricops hirtulus|[4705]|JWDCL272-11|BOLD:ABY7837
Thricops hirtulus|[4706]|JWDCL131-11|BOLD:ABY7837
Thricops hirtulus|[4707]|JWDCJ873-11|BOLD:ABY7837
Thricops hirtulus|[4708]|JWDCL335-11|BOLD:ABY7837
Thricops hirtulus|[4709]|JWDCJ1188-11|BOLD:ABY7837
Thricops hirtulus|[4710]|JWDCJ877-11|BOLD:ABY7837
Thricops hirtulus|[4711]|JWDCJ875-11|BOLD:ABY7837
Thricops hirtulus|[4712]|JWDCJ874-11|BOLD:ABY7837
Thricops hirtulus|[4713]|JWDCJ871-11|BOLD:ABY7837
Thricops hirtulus|[4714]|JWDCJ865-11|BOLD:ABY7837
Thricops hirtulus|[4715]|JWDCJ863-11|BOLD:ABY7837
Thricops hirtulus|[4716]|JWDCK134-11|BOLD:ABY7837
Thricops hirtulus|[4717]|JWDCJ325-11|BOLD:ABY7837
Thricops hirtulus|[4718]|JWDCJ322-11|BOLD:ABY7837
Thricops hirtulus|[4719]|JWDCJ321-11|BOLD:ABY7837
Thricops hirtulus|[4720]|JWDCJ317-11|BOLD:ABY7837
Thricops hirtulus|[4721]|JWDCJ313-11|BOLD:ABY7837
Thricops hirtulus|[4722]|JWDCJ311-11|BOLD:ABY7837
Thricops hirtulus|[4723]|JWDCJ309-11|BOLD:ABY7837
Thricops hirtulus|[4724]|JWDCJ306-11|BOLD:ABY7837
Thricops hirtulus|[4725]|JWDCJ270-11|BOLD:ABY7837
Thricops hirtulus|[4726]|JWDCJ269-11|BOLD:ABY7837
Thricops hirtulus|[4727]|JWDCJ254-11|BOLD:ABY7837
Thricops hirtulus|[4728]|JWDCJ128-11|BOLD:ABY7837
Thricops hirtulus|[4729]|JWDCJ120-11|BOLD:ABY7837
Thricops hirtulus|[4730]|JWDCK143-11|BOLD:ABY7837
Thricops hirtulus|[4731]|JWDCJ031-11|BOLD:ABY7837
Thricops hirtulus|[4732]|JWDCL467-11|BOLD:ABY7837
Thricops hirtulus|[4733]|JWDCL465-11|BOLD:ABY7837
Thricops hirtulus|[4734]|JWDCL340-11|BOLD:ABY7837
Thricops hirtulus|[4735]|JWDCJ1292-11|BOLD:ABY7837
Thricops hirtulus|[4736]|JWDCJ1068-11|BOLD:ABY7837
Thricops hirtulus|[4737]|JWDCJ951-11|BOLD:ABY7837
Thricops hirtulus|[4738]|JWDCJ914-11|BOLD:ABY7837
Thricops hirtulus|[4739]|JWDCK270-11|BOLD:ABY7837
Thricops hirtulus|[4740]|JWDCK256-11|BOLD:ABY7837
Thricops hirtulus|[4741]|JWDCK253-11|BOLD:ABY7837
Thricops hirtulus|[4742]|JWDCK251-11|BOLD:ABY7837
Thricops hirtulus|[4743]|JWDCJ634-11|BOLD:ABY7837
Thricops hirtulus|[4744]|JWDCJ632-11|BOLD:ABY7837
Thricops hirtulus|[4745]|JWDCJ584-11|BOLD:ABY7837
Thricops hirtulus|[4746]|JWDCJ390-11|BOLD:ABY7837
Thricops hirtulus|[4747]|JWDCJ389-11|BOLD:ABY7837
Thricops hirtulus|[4748]|JWDCJ385-11|BOLD:ABY7837
Thricops hirtulus|[4749]|JWDCJ384-11|BOLD:ABY7837
Thricops hirtulus|[4750]|JWDCA457-10|BOLD:ABY7837
Thricops hirtulus|[4751]|JWDCA437-10|BOLD:ABY7837
Thricops hirtulus|[4752]|JWDCA436-10|BOLD:ABY7837
Thricops hirtulus|[4753]|JWDCC218-10|BOLD:ABY7837
Thricops hirtulus|[4754]|JWDCC128-10|BOLD:ABY7837
Thricops hirtulus|[4755]|JWDCC095-10|BOLD:ABY7837
Thricops hirtulus|[4756]|JWDCC014-10|BOLD:ABY7837
Thricops hirtulus|[4757]|JWDCL431-11|BOLD:ABY7837
Thricops hirtulus|[4758]|JWDCA410-10|BOLD:ABY7837
Thricops hirtulus|[4759]|JWDCJ124-11|BOLD:ABY7837
Thricops innocuus|[4760]|MFDC273-09|BOLD:ABY7837
Thricops hirtulus|[4761]|JWDCJ123-11|BOLD:ABY7837
Thricops hirtulus|[4762]|JWDCF862-10|BOLD:ABY7837
Thricops hirtulus|[4763]|JWDCI920-11|BOLD:ABY7837
Thricops hirtulus|[4764]|JWDCF389-10|BOLD:ABY7837
Thricops hirtulus|[4765]|JWDCL275-11|BOLD:ABY7837
Thricops hirtulus|[4766]|JWDCH218-10|BOLD:ABY7837
Thricops hirtulus|[4767]|JWDCJ381-11|BOLD:ABY7837
Thricops hirtulus|[4768]|JWDCD182-10|BOLD:ABY7837
Thricops hirtulus|[4769]|JWDCK269-11|BOLD:ABY7837
Thricops innocuus|[4770]|MFDC886-10|BOLD:ABY7837
Thricops innocuus|[4771]|MFDC266-09|BOLD:ABY7837
Thricops innocuus|[4772]|MFDC272-09|BOLD:ABY7837
Thricops hirtulus|[4773]|MFDC024-08|BOLD:ABY7837
Thricops hirtulus|[4774]|MHFLI928-09|BOLD:ABY7837
Thricops hirtulus|[4775]|MFDC082-08|BOLD:ABY7837
Thricops hirtulus|[4776]|JWDCC840-10|BOLD:ABY7837
Thricops hirtulus|[4777]|JWDCC782-10|BOLD:ABY7837

Thricops hirtulus|[4774]|MHFLI528-09|BOLD:ABY7837
Thricops hirtulus|[4775]|MFDC082-08|BOLD:ABY7837
Thricops hirtulus|[4776]|JWDCC840-10|BOLD:ABY7837
Thricops hirtulus|[4777]|JWDCC782-10|BOLD:ABY7837
Thricops hirtulus|[4778]|JWDCJ803-11|BOLD:ABY7837
Thricops hirtulus|[4779]|JWDCH358-10|BOLD:ABY7837
Thricops hirtulus|[4780]|MFDC274-09|BOLD:ABY7837
Thricops hirtulus|[4781]|MFDC275-09|BOLD:ABY7837
Thricops hirtulus|[4782]|MFDC276-09|BOLD:ABY7837
Thricops hirtulus|[4783]|MFDC277-09|BOLD:ABY7837
Thricops hirtulus|[4784]|MFDC278-09|BOLD:ABY7837
Thricops hirtulus|[4785]|MFDC279-09|BOLD:ABY7837
Thricops hirtulus|[4786]|MFDC280-09|BOLD:ABY7837
Thricops hirtulus|[4787]|MHFLI198-06|BOLD:ABY7837
Thricops hirtulus|[4788]|MFDC741-10|BOLD:ABY7837
Thricops villicrus|[4789]|MHFLI150-06|BOLD:AAB6566
Thricops|[4790]|MHFLI149-06|BOLD:AAB6566
Thricops|[4791]|JWDCA512-10|BOLD:AAB6566
Thricops villicrus|[4792]|MHFLI147-06|BOLD:AAB6566
Thricops spiniger|[4793]|MFDC287-09|BOLD:AAB6566
Thricops spiniger|[4794]|MFDC290-09|BOLD:AAB6566
Thricops|[4795]|JWDCB097-10|BOLD:AAB6566
Thricops spiniger|[4796]|MFDC286-09|BOLD:AAB6566
Thricops septentrionalis|[4797]|MFDC297-09|BOLD:AAB6566
Thricops villicrus|[4798]|MHFLI155-06|BOLD:AAB6566
Thricops villicrus|[4799]|MHFLI151-06|BOLD:AAB6566
Thricops villicrus|[4800]|MFDC071-08|BOLD:AAB6566
Thricops villicrus|[4801]|MHFLI074-06|BOLD:AAB6566
Thricops septentrionalis|[4802]|MFDC292-09|BOLD:AAB6566
Thricops spiniger|[4803]|MFDC288-09|BOLD:AAB6566
Thricops villicrus|[4804]|MFDC007-08|BOLD:AAB6566
Thricops|[4805]|MTDIC231-09|BOLD:AAB6566
Thricops villicrus|[4806]|MHFLI157-06|BOLD:AAB6566
Thricops villicrus|[4807]|MHFLI140-06|BOLD:AAB6566
Thricops villicrus|[4808]|MHFLI139-06|BOLD:AAB6566
Thricops septentrionalis|[4809]|MFDC296-09|BOLD:AAB6566
Thricops villicrus|[4810]|MFDC073-08|BOLD:AAB6566
Thricops septentrionalis|[4811]|MFDC293-09|BOLD:AAB6566
Thricops spiniger|[4812]|MFDC289-09|BOLD:AAB6566
Thricops septentrionalis|[4813]|MFDC291-09|BOLD:AAB6566
Thricops septentrionalis|[4814]|MFDC294-09|BOLD:AAB6566
Thricops villicrus|[4815]|MHFLI146-06|BOLD:AAB6566
Thricops villicrus|[4816]|MHFLI158-06|BOLD:AAB6566
Thricops villicrus|[4817]|MHFLI642-07|BOLD:AAB6566
Haematobosca alcis|[4818]|MFDC624-09|BOLD:AAF8038
Haematobosca alcis|[4819]|MFDC625-09|BOLD:AAF8038
Stomoxys calcitrans|[4820]|MFDC636-09|BOLD:AAA3181
Stomoxys calcitrans|[4821]|MFDC854-10|BOLD:AAA3181
Morellia podagrica|[4822]|MFDC491-09|BOLD:AAC7840
Potamia littoralis|[4823]|MFDC964-10|BOLD:AEN0503
Potamia littoralis|[4824]|MFDC881-10|BOLD:AEN0503
Potamia littoralis|[4825]|MFDC355-09|BOLD:AEN0503
Hydrotaea floccosa|[4826]|MFDC888-10|BOLD:AAG1760
Hydrotaea scambus|[4827]|MFDC974-10|BOLD:AAG1768
Hydrotaea scambus|[4828]|MFDC975-10|BOLD:AAG1768
Hydrotaea scambus|[4829]|MFDC731-10|BOLD:AAG1768
Hydrotaea scambus|[4830]|MFDC732-10|BOLD:AAG1768
Drymeia quadrisetosa|[4831]|JWDCF213-10|BOLD:AAC1021
Drymeia quadrisetosa|[4832]|JWDCL128-11|BOLD:AAC1021
Drymeia quadrisetosa|[4833]|JWDCG004-10|BOLD:AAC1021
Drymeia quadrisetosa|[4834]|JWDCG007-10|BOLD:AAC1021
Drymeia quadrisetosa|[4835]|JWDCF090-10|BOLD:AAC1021
Drymeia quadrisetosa|[4836]|JWDCF091-10|BOLD:AAC1021
Drymeia quadrisetosa|[4837]|JWDCF714-10|BOLD:AAC1021
Drymeia quadrisetosa|[4838]|JWDCF715-10|BOLD:AAC1021
Drymeia quadrisetosa|[4839]|JWDCF716-10|BOLD:AAC1021
Drymeia quadrisetosa|[4840]|JWDCF767-10|BOLD:AAC1021
Drymeia quadrisetosa|[4841]|JWDCF768-10|BOLD:AAC1021
Drymeia quadrisetosa|[4842]|MFDC191-09|BOLD:AAC1021
Drymeia quadrisetosa|[4843]|MFDC192-09|BOLD:AAC1021
Drymeia quadrisetosa|[4844]|MFDC193-09|BOLD:AAC1021
Drymeia quadrisetosa|[4845]|MFDC194-09|BOLD:AAC1021
Drymeia quadrisetosa|[4846]|MFDC195-09|BOLD:AAC1021
Drymeia quadrisetosa|[4847]|MFDC196-09|BOLD:AAC1021
Drymeia quadrisetosa|[4848]|MFDC197-09|BOLD:AAC1021
Drymeia quadrisetosa|[4849]|MFDC199-09|BOLD:AAC1021
Drymeia quadrisetosa|[4850]|MFDC200-09|BOLD:AAC1021
Drymeia quadrisetosa|[4851]|MFDC201-09|BOLD:AAC1021
Drymeia quadrisetosa|[4852]|MFDC202-09|BOLD:AAC1021
Drymeia quadrisetosa|[4853]|MFDC203-09|BOLD:AAC1021
Drymeia quadrisetosa|[4854]|MFDC204-09|BOLD:AAC1021
Drymeia quadrisetosa|[4855]|MFDC198-09|BOLD:AAC1021
Drymeia groenlandica|[4856]|MFDC976-10|BOLD:AAL9801
Drymeia groenlandica|[4857]|MFDC977-10|BOLD:AAL9801
Drymeia pribilofensis|[4858]|MFDC187-09|BOLD:AAD7664
Drymeia pribilofensis|[4859]|MFDC185-09|BOLD:AAD7664
Drymeia pribilofensis|[4860]|MFDC188-09|BOLD:AAD7664
Drymeia pribilofensis|[4861]|MFDC189-09|BOLD:AAD7664
Drymeia pribilofensis|[4862]|MFDC190-09|BOLD:AAD7664
Drymeia pribilofensis|[4863]|MFDC186-09|BOLD:AAD7664
Drymeia pribilofensis|[4864]|MFDC704-10|BOLD:AAD7664
Hydrotaea armipes|[4865]|MFDC729-10|BOLD:AAG6908
Hydrotaea ringdahli|[4866]|MFDC730-10|BOLD:AAM0878
Hydrotaea houghi|[4867]|MFDC176-09|BOLD:AAI8770
Hydrotaea cristata|[4868]|MFDC889-10|BOLD:AAD0824
Hydrotaea cristata|[4869]|MFDC178-09|BOLD:AAD0824
Hydrotaea cristata|[4870]|MFDC181-09|BOLD:AAD0824
Hydrotaea cristata|[4871]|MFDC184-09|BOLD:AAD0824
Hydrotaea cristata|[4872]|MFDC890-10|BOLD:AAD0824
Hydrotaea cristata|[4873]|MFDC891-10|BOLD:AAD0824
Hydrotaea cristata|[4874]|MFDC183-09|BOLD:AAD0824
Hydrotaea cristata|[4875]|MFDC001-08|BOLD:AAD0824
Hydrotaea cristata|[4876]|MFDC887-10|BOLD:AAD0824

Hydrotaea cristata|[4874]|MFDC183-09|BOLD:AAD0824
Hydrotaea cristata|[4875]|MFDC001-08|BOLD:AAD0824
Hydrotaea cristata|[4876]|MFDC887-10|BOLD:AAD0824
Hydrotaea cristata|[4877]|MFDC175-09|BOLD:AAD0824
Hydrotaea cristata|[4878]|MFDC177-09|BOLD:AAD0824
Hydrotaea cristata|[4879]|MFDC182-09|BOLD:AAD0824
Hydrotaea cristata|[4880]|MFDC727-10|BOLD:AAD0824
Hydrotaea cristata|[4881]|MFDC728-10|BOLD:AAD0824
Hydrotaea anxia|[4882]|MFDC163-09|BOLD:AAC6497
Hydrotaea anxia|[4883]|MFDC169-09|BOLD:AAC6497
Hydrotaea anxia|[4884]|MFDC173-09|BOLD:AAC6497
Hydrotaea anxia|[4885]|MFDC174-09|BOLD:AAC6497
Hydrotaea anxia|[4886]|MFDC167-09|BOLD:AAC6497
Hydrotaea anxia|[4887]|MFDC168-09|BOLD:AAC6497
Hydrotaea anxia|[4888]|MFDC170-09|BOLD:AAC6497
Hydrotaea anxia|[4889]|MFDC164-09|BOLD:AAC6497
Hydrotaea anxia|[4890]|MFDC171-09|BOLD:AAC6497
Hydrotaea anxia|[4891]|MFDC172-09|BOLD:AAC6497
Hydrotaea anxia|[4892]|MFDC740-10|BOLD:AAC6497
Hydrotaea anxia|[4893]|MFDC739-10|BOLD:AAC6497
Sarcophagidae|[4894]|JWDCC235-10|BOLD:AAL7746
Sarcophagidae|[4895]|JWDCC394-10|BOLD:AAL7746
Chymomyza caudatula|[4896]|TWDIP082-09|BOLD:AAG8494
Chymomyza caudatula|[4897]|JWDCA909-10|BOLD:AAG8494
Chymomyza caudatula|[4898]|MTDIC208-09|BOLD:AAG8494
Chymomyza caudatula|[4899]|JWDCH572-10|BOLD:AAG8494
Metopia campestris|[4900]|JWDCB051-10|BOLD:AAG2427
Metopia campestris|[4901]|JWDCE937-10|BOLD:AAG2427
Hebecnema nigra|[4902]|JWDCE315-10|BOLD:AAG1726
Hebecnema nigra|[4903]|MHFLI952-09|BOLD:AAG1726
Leucophora marylandica|[4904]|JWDCE775-10|BOLD:AAG2496
Leucophora marylandica|[4905]|JWDCC147-10|BOLD:AAG2496
Leucophora marylandica|[4906]|JWDCE780-10|BOLD:AAG2496
Subhylemyia longula|[4907]|JWDCI192-10|BOLD:AAP8836
Subhylemyia longula|[4908]|JWDCE935-10|BOLD:AAP8836
Coenosia pumila|[4909]|JWDCJ038-11|BOLD:AAG6936
Coenosia pumila|[4910]|MFDC879-10|BOLD:AAG6936
Coenosia pumila|[4911]|MFDC878-10|BOLD:AAG6936
Coenosia pumila|[4912]|JWDCC760-10|BOLD:AAG6936
Coenosia pumila|[4913]|JWDCC759-10|BOLD:AAG6936
Coenosia pumila|[4914]|MFDC880-10|BOLD:AAG6936
Coenosia pumila|[4915]|MFDC700-10|BOLD:AAG6936
Coenosia pumila|[4916]|MFDC685-10|BOLD:AAG6936
Coenosia campestris|[4917]|MFDC388-09|BOLD:AAC1880
Coenosia campestris|[4918]|MFDC872-10|BOLD:AAC1880
Coenosia campestris|[4919]|JWDCL434-11|BOLD:AAC1880
Coenosia campestris|[4920]|JWDCJ125-11|BOLD:AAC1880
Coenosia campestris|[4921]|JWDCB142-10|BOLD:AAC1880
Coenosia campestris|[4922]|JWDCB134-10|BOLD:AAC1880
Coenosia campestris|[4923]|JWDCB124-10|BOLD:AAC1880
Coenosia campestris|[4924]|JWDCB121-10|BOLD:AAC1880
Coenosia campestris|[4925]|MFDC877-10|BOLD:AAC1880
Coenosia campestris|[4926]|MFDC687-10|BOLD:AAC1880
Coenosia campestris|[4927]|MFDC876-10|BOLD:AAC1880
Coenosia campestris|[4928]|JWDCJ316-11|BOLD:AAC1880
Coenosia campestris|[4929]|JWDCJ1298-11|BOLD:AAC1880
Coenosia campestris|[4930]|JWDCJ1288-11|BOLD:AAC1880
Coenosia campestris|[4931]|JWDCA407-10|BOLD:AAC1880
Coenosia campestris|[4932]|JWDCA408-10|BOLD:AAC1880
Coenosia campestris|[4933]|MFDC874-10|BOLD:AAC1880
Coenosia campestris|[4934]|MFDC875-10|BOLD:AAC1880
Coenosia campestris|[4935]|MFDC383-09|BOLD:AAC1880
Coenosia campestris|[4936]|MFDC384-09|BOLD:AAC1880
Coenosia campestris|[4937]|MFDC387-09|BOLD:AAC1880
Coenosia campestris|[4938]|MFDC389-09|BOLD:AAC1880
Coenosia campestris|[4939]|MFDC390-09|BOLD:AAC1880
Coenosia campestris|[4940]|MFDC391-09|BOLD:AAC1880
Coenosia campestris|[4941]|MFDC085-08|BOLD:AAC1880
Coenosia campestris|[4942]|MFDC088-08|BOLD:AAC1880
Coenosia campestris|[4943]|MFDC686-10|BOLD:AAC1880
Coenosia campestris|[4944]|MFDC688-10|BOLD:AAC1880
Coenosia campestris|[4945]|MFDC690-10|BOLD:AAC1880
Coenosia transiens|[4946]|JWDCL200-11|BOLD:AAC4500
Coenosia transiens|[4947]|JWDCJ1509-11|BOLD:AAC4500
Coenosia transiens|[4948]|JWDCH768-10|BOLD:AAC4500
Coenosia transiens|[4949]|JWDCI137-10|BOLD:AAC4500
Coenosia transiens|[4950]|JWDCI138-10|BOLD:AAC4500
Coenosia transiens|[4951]|MFDC432-09|BOLD:AAC4500
Coenosia transiens|[4952]|JWDCH765-10|BOLD:AAC4500
Coenosia transiens|[4953]|JWDCJ1566-11|BOLD:AAC4500
Coenosia transiens|[4954]|JWDCJ1568-11|BOLD:AAC4500
Coenosia transiens|[4955]|JWDCJ1350-11|BOLD:AAC4500
Coenosia transiens|[4956]|JWDCL201-11|BOLD:AAC4500
Coenosia transiens|[4957]|JWDCH724-10|BOLD:AAC4500
Coenosia transiens|[4958]|JWDCH766-10|BOLD:AAC4500
Coenosia transiens|[4959]|JWDCH767-10|BOLD:AAC4500
Coenosia transiens|[4960]|JWDCI399-10|BOLD:AAC4500
Coenosia transiens|[4961]|JWDCI136-10|BOLD:AAC4500
Coenosia transiens|[4962]|MFDC131-09|BOLD:AAC4500
Coenosia transiens|[4963]|JWDCL204-11|BOLD:AAC4500
Coenosia transiens|[4964]|JWDCL212-11|BOLD:AAC4500
Coenosia transiens|[4965]|JWDCI139-10|BOLD:AAC4500
Coenosia transiens|[4966]|MFDC132-09|BOLD:AAC4500
Coenosia transiens|[4967]|MFDC063-08|BOLD:AAC4500
Coenosia frisoni|[4968]|MFDC684-10|BOLD:AAL9581
Coenosia frisoni|[4969]|MFDC689-10|BOLD:AAL9581
Coenosia|[4970]|MHFLI324-07|BOLD:AAG1698
Coenosia|[4971]|JWDCL639-11|BOLD:AAG1698
Coenosia|[4972]|JWDCL143-11|BOLD:AAG1698
Coenosia|[4973]|JWDCL621-11|BOLD:AAG1698
Coenosia|[4974]|JWDCK072-11|BOLD:AAG1698
Coenosia|[4975]|JWDCJ255-11|BOLD:AAG1698
Coenosia|[4976]|[illegible]

Coenosia|[4973]|JWDCL621-11|BOLD:AAG1698
Coenosia|[4974]|JWDCK072-11|BOLD:AAG1698
Coenosia|[4975]|JWDCJ255-11|BOLD:AAG1698
Coenosia|[4976]|JWDCJ213-11|BOLD:AAG1698
Coenosia|[4977]|JWDCJ198-11|BOLD:AAG1698
Coenosia|[4978]|JWDCL611-11|BOLD:AAG1698
Coenosia|[4979]|JWDCJ809-11|BOLD:AAG1698
Coenosia|[4980]|JWDCJ806-11|BOLD:AAG1698
Coenosia|[4981]|JWDCJ796-11|BOLD:AAG1698
Coenosia|[4982]|JWDCJ794-11|BOLD:AAG1698
Coenosia|[4983]|JWDCJ193-11|BOLD:AAG1698
Coenosia|[4984]|JWDCL624-11|BOLD:AAG1698
Coenosia|[4985]|JWDCL627-11|BOLD:AAG1698
Coenosia|[4986]|JWDCL629-11|BOLD:AAG1698
Coenosia|[4987]|JWDCL630-11|BOLD:AAG1698
Coenosia|[4988]|JWDCL631-11|BOLD:AAG1698
Coenosia|[4989]|JWDCL643-11|BOLD:AAG1698
Coenosia|[4990]|JWDCL152-11|BOLD:AAG1698
Coenosia|[4991]|JWDCG634-10|BOLD:AAG1698
Coenosia|[4992]|JWDCG636-10|BOLD:AAG1698
Coenosia|[4993]|JWDCH924-10|BOLD:AAG1698
Coenosia|[4994]|JWDCH380-10|BOLD:AAG1698
Coenosia|[4995]|JWDCH156-10|BOLD:AAG1698
Coenosia|[4996]|JWDCI597-10|BOLD:AAG1698
Coenosia|[4997]|JWDCI599-10|BOLD:AAG1698
Coenosia|[4998]|JWDCF333-10|BOLD:AAG1698
Coenosia|[4999]|JWDCF747-10|BOLD:AAG1698
Coenosia|[5000]|JWDCF950-10|BOLD:AAG1698
Coenosia|[5001]|MHFLI787-07|BOLD:AAG1698
Coenosia verralli|[5002]|JWDCJ1493-11|BOLD:AAC8884
Coenosia verralli|[5003]|JWDCG118-10|BOLD:AAC8884
Coenosia verralli|[5004]|JWDCG119-10|BOLD:AAC8884
Coenosia verralli|[5005]|JWDCH720-10|BOLD:AAC8884
Coenosia verralli|[5006]|JWDCH725-10|BOLD:AAC8884
Coenosia verralli|[5007]|JWDCJ1511-11|BOLD:AAC8884
Coenosia verralli|[5008]|JWDCC431-10|BOLD:AAC8884
Coenosia verralli|[5009]|JWDCE933-10|BOLD:AAC8884
Coenosia verralli|[5010]|JWDCJ1183-11|BOLD:AAC8884
Coenosia verralli|[5011]|JWDCH103-10|BOLD:AAC8884
Coenosia verralli|[5012]|JWDCA474-10|BOLD:AAC8884
Coenosia verralli|[5013]|JWDCJ449-11|BOLD:AAC8884
Coenosia verralli|[5014]|JWDCL553-11|BOLD:AAC8884
Coenosia verralli|[5015]|JWDCL555-11|BOLD:AAC8884
Coenosia verralli|[5016]|JWDCL552-11|BOLD:AAC8884
Coenosia verralli|[5017]|JWDCK064-11|BOLD:AAC8884
Coenosia verralli|[5018]|JWDCH154-10|BOLD:AAC8884
Coenosia verralli|[5019]|JWDCH155-10|BOLD:AAC8884
Coenosia verralli|[5020]|JWDCH523-10|BOLD:AAC8884
Coenosia verralli|[5021]|MFDC385-09|BOLD:AAC8884
Coenosia verralli|[5022]|MFDC404-09|BOLD:AAC8884
Coenosia verralli|[5023]|MFDC406-09|BOLD:AAC8884
Coenosia verralli|[5024]|MFDC409-09|BOLD:AAC8884
Coenosia verralli|[5025]|MFDC410-09|BOLD:AAC8884
Coenosia verralli|[5026]|MFDC411-09|BOLD:AAC8884
Coenosia verralli|[5027]|MFDC412-09|BOLD:AAC8884
Coenosia verralli|[5028]|JWDCF610-10|BOLD:AAC8884
Coenosia verralli|[5029]|MFDC408-09|BOLD:AAC8884
Coenosia verralli|[5030]|JWDCE579-10|BOLD:AAC8884
Coenosia verralli|[5031]|JWDCF373-10|BOLD:AAC8884
Coenosia verralli|[5032]|JWDCH101-10|BOLD:AAC8884
Coenosia verralli|[5033]|JWDCF158-10|BOLD:AAC8884
Coenosia verralli|[5034]|JWDCH769-10|BOLD:AAC8884
Coenosia verralli|[5035]|JWDCG906-10|BOLD:AAC8884
Coenosia verralli|[5036]|JWDCH102-10|BOLD:AAC8884
Coenosia verralli|[5037]|JWDCH097-10|BOLD:AAC8884
Coenosia verralli|[5038]|JWDCH719-10|BOLD:AAC8884
Coenosia verralli|[5039]|JWDCL213-11|BOLD:AAC8884
Coenosia verralli|[5040]|JWDCL211-11|BOLD:AAC8884
Coenosia verralli|[5041]|JWDCJ1440-11|BOLD:AAC8884
Coenosia verralli|[5042]|JWDCJ1358-11|BOLD:AAC8884
Coenosia verralli|[5043]|JWDCJ129-11|BOLD:AAC8884
Coenosia verralli|[5044]|JWDCG898-10|BOLD:AAC8884
Coenosia verralli|[5045]|JWDCJ1510-11|BOLD:AAC8884
Coenosia verralli|[5046]|JWDCK267-11|BOLD:AAC8884
Coenosia verralli|[5047]|JWDCC068-10|BOLD:AAC8884
Coenosia verralli|[5048]|JWDCC589-10|BOLD:AAC8884
Coenosia verralli|[5049]|JWDCC414-10|BOLD:AAC8884
Coenosia verralli|[5050]|JWDCD119-10|BOLD:AAC8884
Coenosia verralli|[5051]|MFDC623-09|BOLD:AAC8884
Coenosia verralli|[5052]|MFDC701-10|BOLD:AAC8884
Coenosia|[5053]|JWDCC838-10|BOLD:AAL6947
Coenosia|[5054]|JWDCF121-10|BOLD:AAL6947
Coenosia atritibia|[5055]|MFDC673-10|BOLD:AAL6947
Coenosia atritibia|[5056]|MFDC674-10|BOLD:AAL6947
Coenosia atritibia|[5057]|MFDC676-10|BOLD:AAL6947
Coenosia nigricoxa|[5058]|JWDCK081-11|BOLD:AAB6353
Coenosia nigricoxa|[5059]|JWDCJ1363-11|BOLD:AAB6353
Coenosia nigricoxa|[5060]|JWDCL125-11|BOLD:AAB6353
Coenosia nigricoxa|[5061]|MHFLI127-06|BOLD:AAB6353
Coenosia nigricoxa|[5062]|JWDCL619-11|BOLD:AAB6353
Coenosia nigricoxa|[5063]|JWDCK061-11|BOLD:AAB6353
Coenosia nigricoxa|[5064]|JWDCH307-10|BOLD:AAB6353
Coenosia nigricoxa|[5065]|MFDC064-08|BOLD:AAB6353
Coenosia nigricoxa|[5066]|MFDC400-09|BOLD:AAB6353
Coenosia nigricoxa|[5067]|JWDCF893-10|BOLD:AAB6353
Coenosia nigricoxa|[5068]|JWDCL637-11|BOLD:AAB6353
Coenosia nigricoxa|[5069]|JWDCL148-11|BOLD:AAB6353
Coenosia nigricoxa|[5070]|JWDCL133-11|BOLD:AAB6353
Coenosia nigricoxa|[5071]|JWDCL122-11|BOLD:AAB6353
Coenosia nigricoxa|[5072]|JWDCK073-11|BOLD:AAB6353
Coenosia nigricoxa|[5073]|JWDCJ126-11|BOLD:AAB6353
Coenosia nigricoxa|[5074]|JWDCL614-11|BOLD:AAB6353
Coenosia nigricoxa|[5075]|JWDCJ1555-11|BOLD:AAB6353

Coenosia nigricoxa|[5073]|JWDCJ126-11|BOLD:AAB6353
Coenosia nigricoxa|[5074]|JWDCL614-11|BOLD:AAB6353
Coenosia nigricoxa|[5075]|JWDCJ1555-11|BOLD:AAB6353
Coenosia nigricoxa|[5076]|MHFLI465-07|BOLD:AAB6353
Coenosia nigricoxa|[5077]|MFDC402-09|BOLD:AAB6353
Coenosia nigricoxa|[5078]|MHFLI318-07|BOLD:AAB6353
Coenosia nigricoxa|[5079]|MHFLI422-07|BOLD:AAB6353
Coenosia nigricoxa|[5080]|MHFLI331-07|BOLD:AAB6353
Coenosia nigricoxa|[5081]|JWDCH153-10|BOLD:AAB6353
Coenosia nigricoxa|[5082]|MFDC435-09|BOLD:AAB6353
Coenosia nigricoxa|[5083]|MFDC407-09|BOLD:AAB6353
Coenosia nigricoxa|[5084]|MFDC405-09|BOLD:AAB6353
Coenosia nigricoxa|[5085]|MFDC403-09|BOLD:AAB6353
Coenosia nigricoxa|[5086]|MFDC399-09|BOLD:AAB6353
Coenosia nigricoxa|[5087]|MFDC396-09|BOLD:AAB6353
Coenosia nigricoxa|[5088]|MFDC395-09|BOLD:AAB6353
Coenosia nigricoxa|[5089]|MFDC394-09|BOLD:AAB6353
Coenosia nigricoxa|[5090]|JWDCF199-10|BOLD:AAB6353
Coenosia nigricoxa|[5091]|JWDCF890-10|BOLD:AAB6353
Coenosia nigricoxa|[5092]|JWDCF804-10|BOLD:AAB6353
Coenosia nigricoxa|[5093]|JWDCF766-10|BOLD:AAB6353
Coenosia nigricoxa|[5094]|JWDCH342-10|BOLD:AAB6353
Coenosia nigricoxa|[5095]|JWDCH302-10|BOLD:AAB6353
Coenosia nigricoxa|[5096]|JWDCH764-10|BOLD:AAB6353
Coenosia nigricoxa|[5097]|JWDCH341-10|BOLD:AAB6353
Coenosia nigricoxa|[5098]|JWDCH339-10|BOLD:AAB6353
Coenosia nigricoxa|[5099]|JWDCH918-10|BOLD:AAB6353
Coenosia nigricoxa|[5100]|JWDCK075-11|BOLD:AAB6353
Coenosia nigricoxa|[5101]|JWDCL633-11|BOLD:AAB6353
Coenosia nigricoxa|[5102]|JWDCL141-11|BOLD:AAB6353
Coenosia nigricoxa|[5103]|JWDCL138-11|BOLD:AAB6353
Coenosia nigricoxa|[5104]|JWDCL126-11|BOLD:AAB6353
Coenosia nigricoxa|[5105]|JWDCL121-11|BOLD:AAB6353
Coenosia nigricoxa|[5106]|JWDCJ1438-11|BOLD:AAB6353
Coenosia nigricoxa|[5107]|JWDCJ1361-11|BOLD:AAB6353
Coenosia nigricoxa|[5108]|JWDCK066-11|BOLD:AAB6353
Coenosia nigricoxa|[5109]|JWDCK065-11|BOLD:AAB6353
Coenosia nigricoxa|[5110]|JWDCJ204-11|BOLD:AAB6353
Coenosia nigricoxa|[5111]|JWDCJ117-11|BOLD:AAB6353
Coenosia nigricoxa|[5112]|JWDCJ021-11|BOLD:AAB6353
Coenosia nigricoxa|[5113]|JWDCL613-11|BOLD:AAB6353
Coenosia nigricoxa|[5114]|JWDCJ1504-11|BOLD:AAB6353
Coenosia nigricoxa|[5115]|JWDCJ1561-11|BOLD:AAB6353
Coenosia nigricoxa|[5116]|JWDCJ1554-11|BOLD:AAB6353
Coenosia nigricoxa|[5117]|JWDCJ1553-11|BOLD:AAB6353
Coenosia nigricoxa|[5118]|JWDCK268-11|BOLD:AAB6353
Coenosia nigricoxa|[5119]|JWDCL549-11|BOLD:AAB6353
Coenosia nigricoxa|[5120]|MFDC070-08|BOLD:AAB6353
Coenosia nigricoxa|[5121]|MFDC152-09|BOLD:AAB6353
Coenosia nigricoxa|[5122]|MFDC401-09|BOLD:AAB6353
Coenosia nigricoxa|[5123]|JWDCJ195-11|BOLD:AAB6353
Coenosia nigricoxa|[5124]|MFDC398-09|BOLD:AAB6353
Coenosia nigricoxa|[5125]|MHFLI336-07|BOLD:AAB6353
Coenosia nigricoxa|[5126]|MHFLI423-07|BOLD:AAB6353
Coenosia nigricoxa|[5127]|MFDC675-10|BOLD:AAB6353
Coenosia nigricoxa|[5128]|MFDC696-10|BOLD:AAB6353
Coenosia sp. 1 AKR|[5129]|MFDC699-10|BOLD:AAG1725
Coenosia flaviseta|[5130]|MFDC682-10|BOLD:ABY2665
Coenosia flaviseta|[5131]|MFDC683-10|BOLD:ABY2665
Coenosia remissa|[5132]|MFDC697-10|BOLD:AAM0327
Coenosia remissa|[5133]|MFDC698-10|BOLD:AAM0327
Coenosia tarsata|[5134]|JWDCL550-11|BOLD:AAJ7001
Coenosia tarsata|[5135]|MFDC434-09|BOLD:AAJ7001
Coenosia mollicula|[5136]|JWDCJ1506-11|BOLD:AAE8903
Coenosia mollicula|[5137]|JWDCI158-10|BOLD:AAE8903
Coenosia mollicula|[5138]|JWDCK139-11|BOLD:AAE8903
Coenosia mollicula|[5139]|JWDCJ1285-11|BOLD:AAE8903
Coenosia mollicula|[5140]|JWDCJ1362-11|BOLD:AAE8903
Coenosia mollicula|[5141]|JWDCJ1437-11|BOLD:AAE8903
Coenosia mollicula|[5142]|JWDCJ1507-11|BOLD:AAE8903
Coenosia mollicula|[5143]|JWDCJ1508-11|BOLD:AAE8903
Coenosia mollicula|[5144]|JWDCG344-10|BOLD:AAE8903
Coenosia mollicula|[5145]|MFDC139-09|BOLD:AAE8903
Coenosia mollicula|[5146]|JWDCJ1500-11|BOLD:AAE8903
Coenosia mollicula|[5147]|JWDCI099-10|BOLD:AAE8903
Coenosia mollicula|[5148]|JWDCI157-10|BOLD:AAE8903
Coenosia mollicula|[5149]|JWDCJ1496-11|BOLD:AAE8903
Coenosia mollicula|[5150]|JWDCK260-11|BOLD:AAE8903
Coenosia mollicula|[5151]|JWDCJ1501-11|BOLD:AAE8903
Coenosia mollicula|[5152]|JWDCK145-11|BOLD:AAE8903
Coenosia mollicula|[5153]|JWDCI101-10|BOLD:AAE8903
Coenosia mollicula|[5154]|MFDC140-09|BOLD:AAE8903
Coenosia mollicula|[5155]|JWDCK140-11|BOLD:AAE8903
Coenosia mollicula|[5156]|JWDCJ1295-11|BOLD:AAE8903
Coenosia mollicula|[5157]|JWDCJ1427-11|BOLD:AAE8903
Coenosia mollicula|[5158]|JWDCJ1433-11|BOLD:AAE8903
Coenosia mollicula|[5159]|JWDCJ1436-11|BOLD:AAE8903
Coenosia mollicula|[5160]|JWDCL198-11|BOLD:AAE8903
Coenosia mollicula|[5161]|JWDCG092-10|BOLD:AAE8903
Coenosia mollicula|[5162]|JWDCI100-10|BOLD:AAE8903
Coenosia mollicula|[5163]|MFDC137-09|BOLD:AAE8903
Coenosia conforma|[5164]|MFDC134-09|BOLD:AAE8917
Coenosia conforma|[5165]|MFDC677-10|BOLD:AAE8917
Coenosia conforma|[5166]|JWDCH721-10|BOLD:AAE8917
Coenosia conforma|[5167]|JWDCJ1558-11|BOLD:AAE8917
Coenosia conforma|[5168]|JWDCJ1353-11|BOLD:AAE8917
Coenosia conforma|[5169]|JWDCJ1356-11|BOLD:AAE8917
Coenosia conforma|[5170]|JWDCH723-10|BOLD:AAE8917
Coenosia conforma|[5171]|JWDCJ023-11|BOLD:AAE8917
Coenosia conforma|[5172]|JWDCH023-10|BOLD:AAE8917
Coenosia conforma|[5173]|MFDC429-09|BOLD:AAE8917
Coenosia conforma|[5174]|JWDCI098-10|BOLD:AAE8917
Coenosia conforma|[5175]|JWDCI481-10|BOLD:AAE8917

Coenosia conforma|[5172]|[illegible]|BOLD:AAE8917
Coenosia conforma|[5173]|MFDC429-09|BOLD:AAE8917
Coenosia conforma|[5174]|JWDCI098-10|BOLD:AAE8917
Coenosia conforma|[5175]|JWDCI481-10|BOLD:AAE8917
Coenosia conforma|[5176]|JWDCG382-10|BOLD:AAE8917
Coenosia conforma|[5177]|MFDC871-10|BOLD:AAE8917
Coenosia conforma|[5178]|JWDCG287-10|BOLD:AAE8917
Coenosia conforma|[5179]|JWDCG381-10|BOLD:AAE8917
Coenosia conforma|[5180]|JWDCL210-11|BOLD:AAE8917
Coenosia conforma|[5181]|JWDCL342-11|BOLD:AAE8917
Coenosia conforma|[5182]|JWDCL203-11|BOLD:AAE8917
Coenosia conforma|[5183]|JWDCL209-11|BOLD:AAE8917
Coenosia conforma|[5184]|JWDCJ1439-11|BOLD:AAE8917
Coenosia conforma|[5185]|JWDCL199-11|BOLD:AAE8917
Coenosia conforma|[5186]|JWDCJ1434-11|BOLD:AAE8917
Coenosia conforma|[5187]|JWDCJ1431-11|BOLD:AAE8917
Coenosia conforma|[5188]|JWDCJ1428-11|BOLD:AAE8917
Coenosia conforma|[5189]|JWDCJ1354-11|BOLD:AAE8917
Coenosia conforma|[5190]|JWDCK200-11|BOLD:AAE8917
Coenosia conforma|[5191]|JWDCJ247-11|BOLD:AAE8917
Coenosia conforma|[5192]|JWDCK265-11|BOLD:AAE8917
Coenosia conforma|[5193]|JWDCJ1357-11|BOLD:AAE8917
Coenosia conforma|[5194]|JWDCJ1556-11|BOLD:AAE8917
Coenosia conforma|[5195]|JWDCJ396-11|BOLD:AAE8917
Coenosia conforma|[5196]|JWDCJ392-11|BOLD:AAE8917
Coenosia conforma|[5197]|JWDCJ1499-11|BOLD:AAE8917
Coenosia conforma|[5198]|JWDCJ393-11|BOLD:AAE8917
Coenosia conforma|[5199]|MFDC133-09|BOLD:AAE8917
Coenosia conforma|[5200]|MFDC428-09|BOLD:AAE8917
Coenosia conforma|[5201]|MFDC054-08|BOLD:AAE8917
Coenosia conforma|[5202]|MFDC678-10|BOLD:AAE8917
Coenosia comita|[5203]|JWDCD185-10|BOLD:AAB7602
Coenosia comita|[5204]|JWDCA456-10|BOLD:AAB7602
Coenosia comita|[5205]|JWDCL341-11|BOLD:AAB7602
Coenosia comita|[5206]|JWDCF520-10|BOLD:AAB7602
Coenosia comita|[5207]|JWDCF472-10|BOLD:AAB7602
Coenosia comita|[5208]|MFDC972-10|BOLD:AAB7602
Coenosia comita|[5209]|MFDC414-09|BOLD:AAB7602
Coenosia comita|[5210]|MFDC420-09|BOLD:AAB7602
Coenosia comita|[5211]|JWDCL065-11|BOLD:AAB7602
Coenosia comita|[5212]|MFDC426-09|BOLD:AAB7602
Coenosia comita|[5213]|JWDCJ310-11|BOLD:AAB7602
Coenosia comita|[5214]|MFDC424-09|BOLD:AAB7602
Coenosia comita|[5215]|JWDCL064-11|BOLD:AAB7602
Coenosia comita|[5216]|MFDC419-09|BOLD:AAB7602
Coenosia comita|[5217]|JWDCA419-10|BOLD:AAB7602
Coenosia comita|[5218]|JWDCK060-11|BOLD:AAB7602
Coenosia comita|[5219]|JWDCJ956-11|BOLD:AAB7602
Coenosia comita|[5220]|JWDCH508-10|BOLD:AAB7602
Coenosia comita|[5221]|JWDCH468-10|BOLD:AAB7602
Coenosia comita|[5222]|JWDCI799-11|BOLD:AAB7602
Coenosia comita|[5223]|JWDCJ260-11|BOLD:AAB7602
Coenosia comita|[5224]|JWDCH163-10|BOLD:AAB7602
Coenosia comita|[5225]|JWDCA413-10|BOLD:AAB7602
Coenosia comita|[5226]|JWDCC105-10|BOLD:AAB7602
Coenosia comita|[5227]|JWDCA417-10|BOLD:AAB7602
Coenosia comita|[5228]|JWDCA450-10|BOLD:AAB7602
Coenosia comita|[5229]|JWDCA454-10|BOLD:AAB7602
Coenosia comita|[5230]|JWDCJ706-11|BOLD:AAB7602
Coenosia comita|[5231]|JWDCJ707-11|BOLD:AAB7602
Coenosia comita|[5232]|JWDCJ209-11|BOLD:AAB7602
Coenosia comita|[5233]|JWDCJ248-11|BOLD:AAB7602
Coenosia comita|[5234]|JWDCJ1185-11|BOLD:AAB7602
Coenosia comita|[5235]|JWDCL062-11|BOLD:AAB7602
Coenosia comita|[5236]|JWDCL077-11|BOLD:AAB7602
Coenosia comita|[5237]|JWDCL145-11|BOLD:AAB7602
Coenosia comita|[5238]|JWDCL353-11|BOLD:AAB7602
Coenosia comita|[5239]|JWDCI420-10|BOLD:AAB7602
Coenosia comita|[5240]|JWDCI421-10|BOLD:AAB7602
Coenosia comita|[5241]|JWDCI686-11|BOLD:AAB7602
Coenosia comita|[5242]|JWDCI738-11|BOLD:AAB7602
Coenosia comita|[5243]|JWDCF918-10|BOLD:AAB7602
Coenosia comita|[5244]|JWDCF921-10|BOLD:AAB7602
Coenosia comita|[5245]|MFDC416-09|BOLD:AAB7602
Coenosia comita|[5246]|JWDCH470-10|BOLD:AAB7602
Coenosia comita|[5247]|JWDCA449-10|BOLD:AAB7602
Coenosia comita|[5248]|JWDCL146-11|BOLD:AAB7602
Coenosia comita|[5249]|MFDC421-09|BOLD:AAB7602
Coenosia comita|[5250]|MFDC415-09|BOLD:AAB7602
Coenosia comita|[5251]|JWDCJ319-11|BOLD:AAB7602
Coenosia comita|[5252]|JWDCJ267-11|BOLD:AAB7602
Coenosia comita|[5253]|JWDCJ206-11|BOLD:AAB7602
Coenosia comita|[5254]|JWDCJ194-11|BOLD:AAB7602
Coenosia comita|[5255]|JWDCJ033-11|BOLD:AAB7602
Coenosia comita|[5256]|JWDCL339-11|BOLD:AAB7602
Coenosia comita|[5257]|JWDCJ954-11|BOLD:AAB7602
Coenosia comita|[5258]|JWDCJ704-11|BOLD:AAB7602
Coenosia comita|[5259]|JWDCJ633-11|BOLD:AAB7602
Coenosia comita|[5260]|JWDCA447-10|BOLD:AAB7602
Coenosia comita|[5261]|JWDCA418-10|BOLD:AAB7602
Coenosia comita|[5262]|JWDCA416-10|BOLD:AAB7602
Coenosia comita|[5263]|JWDCA415-10|BOLD:AAB7602
Coenosia comita|[5264]|JWDCA426-10|BOLD:AAB7602
Coenosia comita|[5265]|JWDCJ203-11|BOLD:AAB7602
Coenosia comita|[5266]|JWDCK196-11|BOLD:AAB7602
Coenosia comita|[5267]|JWDCK199-11|BOLD:AAB7602
Coenosia comita|[5268]|JWDCJ1349-11|BOLD:AAB7602
Coenosia comita|[5269]|JWDCL066-11|BOLD:AAB7602
Coenosia comita|[5270]|JWDCL067-11|BOLD:AAB7602
Coenosia comita|[5271]|JWDCL074-11|BOLD:AAB7602
Coenosia comita|[5272]|JWDCL124-11|BOLD:AAB7602
Coenosia comita|[5273]|JWDCH301-10|BOLD:AAB7602
Coenosia comita|[5274]|JWDCI055-10|BOLD:AAB7602

Coenosia comita|[5272]|JWDCL124-11|BOLD:AAB7602
Coenosia comita|[5273]|JWDCH301-10|BOLD:AAB7602
Coenosia comita|[5274]|JWDCI055-10|BOLD:AAB7602
Coenosia comita|[5275]|JWDCI786-11|BOLD:AAB7602
Coenosia comita|[5276]|JWDCI422-10|BOLD:AAB7602
Coenosia comita|[5277]|JWDCI347-10|BOLD:AAB7602
Coenosia comita|[5278]|JWDCI690-11|BOLD:AAB7602
Coenosia comita|[5279]|JWDCI691-11|BOLD:AAB7602
Coenosia comita|[5280]|JWDCI737-11|BOLD:AAB7602
Coenosia comita|[5281]|JWDCI790-11|BOLD:AAB7602
Coenosia comita|[5282]|JWDCI795-11|BOLD:AAB7602
Coenosia comita|[5283]|JWDCI796-11|BOLD:AAB7602
Coenosia comita|[5284]|JWDCF594-10|BOLD:AAB7602
Coenosia comita|[5285]|JWDCF923-10|BOLD:AAB7602
Coenosia comita|[5286]|MFDC144-09|BOLD:AAB7602
Coenosia comita|[5287]|MFDC145-09|BOLD:AAB7602
Coenosia comita|[5288]|MFDC397-09|BOLD:AAB7602
Coenosia comita|[5289]|MFDC417-09|BOLD:AAB7602
Coenosia comita|[5290]|MFDC418-09|BOLD:AAB7602
Coenosia comita|[5291]|MFDC422-09|BOLD:AAB7602
Coenosia comita|[5292]|MFDC423-09|BOLD:AAB7602
Coenosia comita|[5293]|MFDC425-09|BOLD:AAB7602
Coenosia comita|[5294]|MFDC041-08|BOLD:AAB7602
Coenosia octopunctata|[5295]|JWDCF496-10|BOLD:AAC6280
Coenosia octopunctata|[5296]|MFDC147-09|BOLD:AAC6280
Coenosia octopunctata|[5297]|JWDCF120-10|BOLD:AAC6280
Coenosia octopunctata|[5298]|JWDCJ958-11|BOLD:AAC6280
Coenosia octopunctata|[5299]|JWDCD075-10|BOLD:AAC6280
Coenosia octopunctata|[5300]|JWDCD056-10|BOLD:AAC6280
Coenosia octopunctata|[5301]|JWDCF034-10|BOLD:AAC6280
Coenosia octopunctata|[5302]|JWDCF691-10|BOLD:AAC6280
Coenosia octopunctata|[5303]|MFDC151-09|BOLD:AAC6280
Coenosia octopunctata|[5304]|JWDCF690-10|BOLD:AAC6280
Coenosia octopunctata|[5305]|JWDCF686-10|BOLD:AAC6280
Coenosia octopunctata|[5306]|JWDCE665-10|BOLD:AAC6280
Coenosia octopunctata|[5307]|JWDCF118-10|BOLD:AAC6280
Coenosia octopunctata|[5308]|JWDCF092-10|BOLD:AAC6280
Coenosia octopunctata|[5309]|JWDCF032-10|BOLD:AAC6280
Coenosia octopunctata|[5310]|JWDCF031-10|BOLD:AAC6280
Coenosia octopunctata|[5311]|JWDCI572-10|BOLD:AAC6280
Coenosia octopunctata|[5312]|JWDCJ959-11|BOLD:AAC6280
Coenosia octopunctata|[5313]|JWDCJ575-11|BOLD:AAC6280
Coenosia octopunctata|[5314]|JWDCA444-10|BOLD:AAC6280
Coenosia octopunctata|[5315]|JWDCF689-10|BOLD:AAC6280
Coenosia octopunctata|[5316]|JWDCF740-10|BOLD:AAC6280
Coenosia octopunctata|[5317]|JWDCF746-10|BOLD:AAC6280
Coenosia octopunctata|[5318]|JWDCF770-10|BOLD:AAC6280
Coenosia octopunctata|[5319]|MFDC427-09|BOLD:AAC6280
Coenosia octopunctata|[5320]|MFDC033-08|BOLD:AAC6280
Coenosia octopunctata|[5321]|MFDC663-09|BOLD:AAC6280
Coenosia octopunctata|[5322]|MFDC664-09|BOLD:AAC6280
Coenosia octopunctata|[5323]|MFDC056-08|BOLD:AAC6280
Coenosia octopunctata|[5324]|MFDC148-09|BOLD:AAC6280
Coenosia demoralis|[5325]|MFDC1068-11|BOLD:AAE1399
Coenosia|[5326]|JWDCB126-10|BOLD:AAE1399
Coenosia|[5327]|JWDCA565-10|BOLD:AAE1399
Coenosia demoralis|[5328]|MFDC094-08|BOLD:AAE1399
Coenosia|[5329]|JWDCA566-10|BOLD:AAE1399
Coenosia|[5330]|JWDCA564-10|BOLD:AAE1399
Coenosia|[5331]|JWDCA420-10|BOLD:AAE1399
Coenosia|[5332]|JWDCB123-10|BOLD:AAE1399
Coenosia|[5333]|JWDCB122-10|BOLD:AAE1399
Coenosia|[5334]|JWDCB125-10|BOLD:AAE1399
Coenosia demoralis|[5335]|MFDC142-09|BOLD:AAE1399
Coenosia demoralis|[5336]|MFDC1067-11|BOLD:AAE1399
Coenosia demoralis|[5337]|MFDC680-10|BOLD:AAE1399
Coenosia demoralis|[5338]|MFDC681-10|BOLD:AAE1399
Coenosia minor|[5339]|MFDC691-10|BOLD:AAE1399
Coenosia minor|[5340]|MFDC135-09|BOLD:AAE1399
Coenosia minor|[5341]|MFDC136-09|BOLD:AAE1399
Coenosia minor|[5342]|MFDC692-10|BOLD:AAE1399
Coenosia minor|[5343]|MFDC693-10|BOLD:AAE1399
Coenosia minor|[5344]|MFDC694-10|BOLD:AAE1399
Coenosia|[5345]|JWDCJ262-11|BOLD:AAC8885
Coenosia|[5346]|JWDCF894-10|BOLD:AAC8885
Coenosia|[5347]|JWDCB106-10|BOLD:AAC8885
Coenosia imperator|[5348]|MHFLI119-06|BOLD:AAC8885
Coenosia|[5349]|JWDCJ019-11|BOLD:AAC8885
Coenosia|[5350]|JWDCI741-11|BOLD:AAC8885
Coenosia|[5351]|JWDCG281-10|BOLD:AAC8885
Coenosia|[5352]|JWDCH409-10|BOLD:AAC8885
Coenosia|[5353]|JWDCH165-10|BOLD:AAC8885
Coenosia|[5354]|JWDCH167-10|BOLD:AAC8885
Coenosia|[5355]|JWDCH917-10|BOLD:AAC8885
Coenosia|[5356]|JWDCH981-10|BOLD:AAC8885
Coenosia|[5357]|JWDCG720-10|BOLD:AAC8885
Coenosia|[5358]|JWDCG722-10|BOLD:AAC8885
Coenosia|[5359]|JWDCL347-11|BOLD:AAC8885
Coenosia|[5360]|JWDCH716-10|BOLD:AAC8885
Coenosia|[5361]|JWDCL055-11|BOLD:AAC8885
Coenosia|[5362]|JWDCL436-11|BOLD:AAC8885
Coenosia|[5363]|JWDCJ1175-11|BOLD:AAC8885
Coenosia|[5364]|JWDCJ1352-11|BOLD:AAC8885
Coenosia|[5365]|JWDCI800-11|BOLD:AAC8885
Coenosia|[5366]|JWDCJ1169-11|BOLD:AAC8885
Coenosia|[5367]|JWDCJ201-11|BOLD:AAC8885
Coenosia|[5368]|JWDCL609-11|BOLD:AAC8885
Coenosia|[5369]|JWDCA424-10|BOLD:AAC8885
Coenosia|[5370]|JWDCA409-10|BOLD:AAC8885
Coenosia|[5371]|JWDCB087-10|BOLD:AAC8885
Coenosia|[5372]|JWDCH404-10|BOLD:AAC8885
Coenosia|[5373]|JWDCL054-11|BOLD:AAC8885
Coenosia imperator|[5374]|MFDC380-09|BOLD:AAC8885

Coenosia|[5371]|JWDCB087-10|BOLD:AAC8885
Coenosia|[5372]|JWDCH404-10|BOLD:AAC8885
Coenosia|[5373]|JWDCL054-11|BOLD:AAC8885
Coenosia imperator|[5374]|MFDC380-09|BOLD:AAC8885
Coenosia|[5375]|JWDCH305-10|BOLD:AAC8885
Coenosia|[5376]|JWDCL278-11|BOLD:AAC8885
Coenosia|[5377]|JWDCF606-10|BOLD:AAC8885
Coenosia imperator|[5378]|MHFLI138-06|BOLD:AAC8885
Coenosia|[5379]|JWDCF924-10|BOLD:AAC8885
Coenosia|[5380]|JWDCF607-10|BOLD:AAC8885
Coenosia|[5381]|JWDCI918-11|BOLD:AAC8885
Coenosia|[5382]|JWDCI910-11|BOLD:AAC8885
Coenosia|[5383]|JWDCI798-11|BOLD:AAC8885
Coenosia|[5384]|JWDCI797-11|BOLD:AAC8885
Coenosia|[5385]|JWDCI743-11|BOLD:AAC8885
Coenosia|[5386]|JWDCI685-11|BOLD:AAC8885
Coenosia|[5387]|JWDCI106-10|BOLD:AAC8885
Coenosia|[5388]|JWDCI104-10|BOLD:AAC8885
Coenosia|[5389]|JWDCH166-10|BOLD:AAC8885
Coenosia|[5390]|JWDCG725-10|BOLD:AAC8885
Coenosia|[5391]|JWDCG088-10|BOLD:AAC8885
Coenosia|[5392]|JWDCH984-10|BOLD:AAC8885
Coenosia|[5393]|JWDCG834-10|BOLD:AAC8885
Coenosia|[5394]|JWDCG831-10|BOLD:AAC8885
Coenosia|[5395]|JWDCG635-10|BOLD:AAC8885
Coenosia|[5396]|JWDCG632-10|BOLD:AAC8885
Coenosia|[5397]|JWDCG630-10|BOLD:AAC8885
Coenosia|[5398]|JWDCL435-11|BOLD:AAC8885
Coenosia|[5399]|JWDCL354-11|BOLD:AAC8885
Coenosia|[5400]|JWDCL620-11|BOLD:AAC8885
Coenosia|[5401]|JWDCL070-11|BOLD:AAC8885
Coenosia|[5402]|JWDCL058-11|BOLD:AAC8885
Coenosia|[5403]|JWDCJ1187-11|BOLD:AAC8885
Coenosia|[5404]|JWDCJ1179-11|BOLD:AAC8885
Coenosia|[5405]|JWDCJ1174-11|BOLD:AAC8885
Coenosia|[5406]|JWDCJ1173-11|BOLD:AAC8885
Coenosia|[5407]|JWDCJ912-11|BOLD:AAC8885
Coenosia|[5408]|JWDCK198-11|BOLD:AAC8885
Coenosia|[5409]|JWDCJ256-11|BOLD:AAC8885
Coenosia|[5410]|JWDCJ207-11|BOLD:AAC8885
Coenosia|[5411]|JWDCJ197-11|BOLD:AAC8885
Coenosia|[5412]|JWDCJ118-11|BOLD:AAC8885
Coenosia|[5413]|JWDCK255-11|BOLD:AAC8885
Coenosia|[5414]|JWDCL439-11|BOLD:AAC8885
Coenosia|[5415]|JWDCL615-11|BOLD:AAC8885
Coenosia|[5416]|JWDCL600-11|BOLD:AAC8885
Coenosia|[5417]|JWDCL574-11|BOLD:AAC8885
Coenosia|[5418]|JWDCJ961-11|BOLD:AAC8885
Coenosia|[5419]|JWDCJ805-11|BOLD:AAC8885
Coenosia|[5420]|JWDCK275-11|BOLD:AAC8885
Coenosia|[5421]|JWDCA470-10|BOLD:AAC8885
Coenosia|[5422]|JWDCA472-10|BOLD:AAC8885
Coenosia|[5423]|JWDCB140-10|BOLD:AAC8885
Coenosia|[5424]|JWDCJ1562-11|BOLD:AAC8885
Coenosia|[5425]|JWDCI511-10|BOLD:AAC8885
Coenosia|[5426]|JWDCJ263-11|BOLD:AAC8885
Coenosia imperator|[5427]|MFDC157-09|BOLD:AAC8885
Coenosia imperator|[5428]|MFDC158-09|BOLD:AAC8885
Coenosia imperator|[5429]|MFDC038-08|BOLD:AAC8885
Coenosia imperator|[5430]|MFDC702-10|BOLD:AAC8885
Coenosia imperator|[5431]|MHFLI120-06|BOLD:AAC8885
Schoenomyza|[5432]|JWDCH230-10|BOLD:AAP9049
Schoenomyza|[5433]|JWDCI846-11|BOLD:AAP9049
Schoenomyza|[5434]|JWDCF662-10|BOLD:AAP9049
Schoenomyza dorsalis|[5435]|MFDC235-09|BOLD:AAE9281
Schoenomyza dorsalis|[5436]|MFDC234-09|BOLD:AAE9281
Schoenomyza dorsalis|[5437]|JWDCF200-10|BOLD:AAE9281
Schoenomyza dorsalis|[5438]|JWDCF201-10|BOLD:AAE9281
Schoenomyza dorsalis|[5439]|JWDCF191-10|BOLD:AAE9281
Schoenomyza dorsalis|[5440]|JWDCI782-11|BOLD:AAE9281
Schoenomyza dorsalis|[5441]|JWDCG095-10|BOLD:AAE9281
Schoenomyza dorsalis|[5442]|JWDCL216-11|BOLD:AAE9281
Schoenomyza dorsalis|[5443]|JWDCL207-11|BOLD:AAE9281
Schoenomyza dorsalis|[5444]|JWDCJ214-11|BOLD:AAE9281
Schoenomyza dorsalis|[5445]|JWDCJ028-11|BOLD:AAE9281
Schoenomyza dorsalis|[5446]|JWDCJ1017-11|BOLD:AAE9281
Schoenomyza dorsalis|[5447]|MFDC236-09|BOLD:AAE9281
Schoenomyza dorsalis|[5448]|JWDCJ374-11|BOLD:AAE9281
Schoenomyza dorsalis|[5449]|JWDCH098-10|BOLD:AAE9281
Schoenomyza dorsalis|[5450]|MHCOL184-07|BOLD:AAE9281
Schoenomyza litorella|[5451]|JWDCF595-10|BOLD:AAB9109
Schoenomyza litorella|[5452]|JWDCI848-11|BOLD:AAB9109
Schoenomyza litorella|[5453]|MFDC783-10|BOLD:AAB9109
Schoenomyza litorella|[5454]|JWDCI601-10|BOLD:AAB9109
Schoenomyza litorella|[5455]|JWDCH379-10|BOLD:AAB9109
Schoenomyza litorella|[5456]|FLIMH016-08|BOLD:AAB9109
Schoenomyza litorella|[5457]|JWDCI596-10|BOLD:AAB9109
Schoenomyza litorella|[5458]|JWDCH718-10|BOLD:AAB9109
Schoenomyza litorella|[5459]|JWDCI600-10|BOLD:AAB9109
Schoenomyza litorella|[5460]|JWDCF748-10|BOLD:AAB9109
Schoenomyza litorella|[5461]|JWDCJ035-11|BOLD:AAB9109
Schoenomyza litorella|[5462]|MHFLI439-07|BOLD:AAB9109
Schoenomyza litorella|[5463]|MHFLI371-07|BOLD:AAB9109
Schoenomyza litorella|[5464]|MHFLI589-07|BOLD:AAB9109
Schoenomyza litorella|[5465]|JWDCJ026-11|BOLD:AAB9109
Schoenomyza litorella|[5466]|JWDCJ037-11|BOLD:AAB9109
Schoenomyza litorella|[5467]|JWDCF015-10|BOLD:AAB9109
Schoenomyza litorella|[5468]|JWDCF190-10|BOLD:AAB9109
Schoenomyza litorella|[5469]|JWDCF202-10|BOLD:AAB9109
Schoenomyza litorella|[5470]|JWDCF284-10|BOLD:AAB9109
Schoenomyza litorella|[5471]|JWDCF285-10|BOLD:AAB9109
Schoenomyza litorella|[5472]|JWDCF742-10|BOLD:AAB9109
Schoenomyza litorella|[5473]|JWDCF744-10|BOLD:AAB9109

Schoenomyza litorella|[5471]|JWDCF285-10|BOLD:AAB9109
Schoenomyza litorella|[5472]|JWDCF742-10|BOLD:AAB9109
Schoenomyza litorella|[5473]|JWDCF744-10|BOLD:AAB9109
Schoenomyza litorella|[5474]|MFDC228-09|BOLD:AAB9109
Schoenomyza litorella|[5475]|MFDC230-09|BOLD:AAB9109
Schoenomyza litorella|[5476]|MFDC231-09|BOLD:AAB9109
Schoenomyza litorella|[5477]|MFDC232-09|BOLD:AAB9109
Schoenomyza litorella|[5478]|MFDC233-09|BOLD:AAB9109
Schoenomyza litorella|[5479]|MFDC237-09|BOLD:AAB9109
Schoenomyza litorella|[5480]|MFDC238-09|BOLD:AAB9109
Schoenomyza litorella|[5481]|MFDC239-09|BOLD:AAB9109
Schoenomyza litorella|[5482]|MFDC781-10|BOLD:AAB9109
Schoenomyza litorella|[5483]|JWDCL634-11|BOLD:AAB9109
Schoenomyza litorella|[5484]|JWDCL636-11|BOLD:AAB9109
Schoenomyza litorella|[5485]|JWDCL164-11|BOLD:AAB9109
Schoenomyza litorella|[5486]|JWDCL623-11|BOLD:AAB9109
Schoenomyza litorella|[5487]|JWDCJ1348-11|BOLD:AAB9109
Schoenomyza litorella|[5488]|JWDCJ1364-11|BOLD:AAB9109
Schoenomyza litorella|[5489]|JWDCG089-10|BOLD:AAB9109
Schoenomyza litorella|[5490]|JWDCH224-10|BOLD:AAB9109
Schoenomyza litorella|[5491]|JWDCH229-10|BOLD:AAB9109
Schoenomyza litorella|[5492]|JWDCG535-10|BOLD:AAB9109
Schoenomyza litorella|[5493]|JWDCJ1604-11|BOLD:AAB9109
Schoenomyza litorella|[5494]|JWDCL440-11|BOLD:AAB9109
Schoenomyza litorella|[5495]|JWDCL638-11|BOLD:AAB9109
Schoenomyza litorella|[5496]|JWDCL657-11|BOLD:AAB9109
Schoenomyza litorella|[5497]|JWDCL205-11|BOLD:AAB9109
Schoenomyza litorella|[5498]|JWDCH928-10|BOLD:AAB9109
Schoenomyza litorella|[5499]|JWDCJ1557-11|BOLD:AAB9109
Schoenomyza litorella|[5500]|JWDCJ1564-11|BOLD:AAB9109
Schoenomyza litorella|[5501]|JWDCJ1549-11|BOLD:AAB9109
Schoenomyza litorella|[5502]|JWDCJ1548-11|BOLD:AAB9109
Schoenomyza litorella|[5503]|JWDCJ804-11|BOLD:AAB9109
Schoenomyza litorella|[5504]|JWDCJ797-11|BOLD:AAB9109
Schoenomyza litorella|[5505]|JWDCJ788-11|BOLD:AAB9109
Schoenomyza litorella|[5506]|JWDCL554-11|BOLD:AAB9109
Schoenomyza litorella|[5507]|JWDCJ020-11|BOLD:AAB9109
Schoenomyza litorella|[5508]|JWDCB137-10|BOLD:AAB9109
Schoenomyza litorella|[5509]|JWDCB135-10|BOLD:AAB9109
Schoenomyza litorella|[5510]|JWDCL167-11|BOLD:AAB9109
Schoenomyza litorella|[5511]|MHFLI432-07|BOLD:AAB9109
Schoenomyza litorella|[5512]|JWDCJ1563-11|BOLD:AAB9109
Schoenomyza litorella|[5513]|MHFLI451-07|BOLD:AAB9109
Schoenomyza litorella|[5514]|MHFLI577-07|BOLD:AAB9109
Schoenomyza litorella|[5515]|JWDCJ034-11|BOLD:AAB9109
Schoenomyza litorella|[5516]|MHFLI343-07|BOLD:AAB9109
Schoenomyza litorella|[5517]|FLIMH032-08|BOLD:AAB9109
Schoenomyza litorella|[5518]|MFDC782-10|BOLD:AAB9109
Spilogona sp. 5 AKR|[5519]|MFDC375-09|BOLD:AAE2783
Pseudocoenosia brevicauda|[5520]|MFDC771-10|BOLD:ACZ4665
Pseudocoenosia|[5521]|JWDCH022-10|BOLD:ACZ4665
Pseudocoenosia|[5522]|JWDCL573-11|BOLD:ACZ4665
Pseudocoenosia|[5523]|JWDCJ253-11|BOLD:ACZ4665
Pseudocoenosia|[5524]|JWDCK067-11|BOLD:ACZ4665
Pseudocoenosia|[5525]|JWDCL144-11|BOLD:ACZ4665
Pseudocoenosia|[5526]|JWDCL147-11|BOLD:ACZ4665
Pseudocoenosia|[5527]|JWDCH920-10|BOLD:ACZ4665
Pseudocoenosia|[5528]|JWDCH922-10|BOLD:ACZ4665
Pseudocoenosia|[5529]|JWDCG727-10|BOLD:ACZ4665
Pseudocoenosia brevicauda|[5530]|MFDC772-10|BOLD:ACZ4665
Pseudocoenosia brevicauda|[5531]|MFDC773-10|BOLD:ACZ4665
Pseudocoenosia solitaria|[5532]|MHFLI221-06|BOLD:AAG1690
Pseudocoenosia solitaria|[5533]|JWDCK068-11|BOLD:AAG1690
Pseudocoenosia solitaria|[5534]|JWDCL622-11|BOLD:AAG1690
Pseudocoenosia solitaria|[5535]|JWDCI398-10|BOLD:AAG1690
Pseudocoenosia solitaria|[5536]|JWDCH024-10|BOLD:AAG1690
Pseudocoenosia solitaria|[5537]|MFDC374-09|BOLD:AAG1690
Pseudocoenosia solitaria|[5538]|MFDC778-10|BOLD:AAG1690
Pseudocoenosia solitaria|[5539]|JWDCJ808-11|BOLD:AAG1690
Pseudocoenosia solitaria|[5540]|JWDCJ258-11|BOLD:AAG1690
Pseudocoenosia solitaria|[5541]|MFDC779-10|BOLD:AAG1690
Pseudocoenosia solitaria|[5542]|JWDCJ252-11|BOLD:AAG1690
Pseudocoenosia solitaria|[5543]|JWDCL610-11|BOLD:AAG1690
Pseudocoenosia solitaria|[5544]|JWDCL603-11|BOLD:AAG1690
Pseudocoenosia solitaria|[5545]|JWDCJ1505-11|BOLD:AAG1690
Pseudocoenosia solitaria|[5546]|JWDCL150-11|BOLD:AAG1690
Pseudocoenosia solitaria|[5547]|JWDCL281-11|BOLD:AAG1690
Pseudocoenosia solitaria|[5548]|JWDCI730-11|BOLD:AAG1690
Pseudocoenosia solitaria|[5549]|JWDCI842-11|BOLD:AAG1690
Pseudocoenosia solitaria|[5550]|MFDC777-10|BOLD:AAG1690
Pseudocoenosia solitaria|[5551]|MFDC780-10|BOLD:AAG1690
Spilogona|[5552]|JWDCF664-10|BOLD:AAE2784
Spilogona|[5553]|JWDCC413-10|BOLD:AAE2784
Spilogona|[5554]|JWDCC411-10|BOLD:AAE2784
Spilogona|[5555]|JWDCJ673-11|BOLD:AAE2784
Spilogona|[5556]|JWDCJ525-11|BOLD:AAE2784
Spilogona|[5557]|JWDCJ526-11|BOLD:AAE2784
Spilogona|[5558]|JWDCF452-10|BOLD:AAE2784
Spilogona novemmaculata|[5559]|MFDC508-09|BOLD:AAE2784
Spilogona|[5560]|JWDCB132-10|BOLD:AAE2784
Spilogona|[5561]|JWDCC241-10|BOLD:AAE2784
Spilogona novemmaculata|[5562]|MFDC812-10|BOLD:AAE2784
Spilogona fatima|[5563]|JWDCJ199-11|BOLD:ACF1455
Spilogona fatima|[5564]|JWDCJ957-11|BOLD:ACF1455
Spilogona fatima|[5565]|JWDCJ953-11|BOLD:ACF1455
Spilogona fatima|[5566]|JWDCF016-10|BOLD:ACF1455
Spilogona fatima|[5567]|JWDCE515-10|BOLD:ACF1455
Spilogona fatima|[5568]|MFDC509-09|BOLD:ACF1455
Spilogona fatima|[5569]|MFDC510-09|BOLD:ACF1455
Spilogona fatima|[5570]|MFDC810-10|BOLD:ACF1455
Spilogona fatima|[5571]|MFDC811-10|BOLD:ACF1455
Spilogona fatima|[5572]|MFDC813-10|BOLD:ACF1455
Limnospila albifrons|[5573]|MHFLI598-07|BOLD:AAG1691

Spilogona fatima|[5570]|MFDC810-10|BOLD:ACF1455
Spilogona fatima|[5571]|MFDC811-10|BOLD:ACF1455
Spilogona fatima|[5572]|MFDC813-10|BOLD:ACF1455
Limnospila albifrons|[5573]|MHFLI598-07|BOLD:AAG1691
Limnospila albifrons|[5574]|MHFLI937-09|BOLD:AAG1691
Limnospila albifrons|[5575]|MFDC745-10|BOLD:AAG1691
Limnospila albifrons|[5576]|MFDC744-10|BOLD:AAG1691
Limnospila albifrons|[5577]|MFDC743-10|BOLD:AAG1691
Limnospila albifrons|[5578]|MFDC742-10|BOLD:AAG1691
Limnospila albifrons|[5579]|JWDCJ962-11|BOLD:AAG1691
Limnospila albifrons|[5580]|MHFLI663-07|BOLD:AAG1691
Limnospila albifrons|[5581]|MHFLI419-07|BOLD:AAG1691
Limnospila albifrons|[5582]|MHFLI646-07|BOLD:AAG1691
Spilogona firmidisetosa|[5583]|MFDC992-10|BOLD:AAL5985
Spilogona firmidisetosa|[5584]|MFDC790-10|BOLD:AAL5985
Spilogona firmidisetosa|[5585]|MFDC791-10|BOLD:AAL5985
Spilogona firmidisetosa|[5586]|MFDC814-10|BOLD:AAL5985
Spilogona firmidisetosa|[5587]|MFDC990-10|BOLD:AAL5985
Spilogona firmidisetosa|[5588]|MFDC991-10|BOLD:AAL5985
Spilogona firmidisetosa|[5589]|MFDC815-10|BOLD:AAL5985
Spilogona firmidisetosa|[5590]|MFDC817-10|BOLD:AAL5985
Spilogona firmidisetosa|[5591]|MFDC816-10|BOLD:AAL5985
Pelomyiella mallochi|[5592]|MHFLI357-07|BOLD:AAG6759
Pelomyiella mallochi|[5593]|FLIMH010-08|BOLD:AAG6827
Pelomyiella mallochi|[5594]|FLIMH024-08|BOLD:AAG6827
Pelomyiella mallochi|[5595]|FLIMH025-08|BOLD:AAG6827
Pelomyiella mallochi|[5596]|FLIMH027-08|BOLD:AAG6827
Pelomyiella mallochi|[5597]|JWDCJ1514-11|BOLD:AAG3916
Chrysops frigidus|[5598]|JWDCB041-10|BOLD:AAD4375
Chrysops frigidus|[5599]|JWDCK122-11|BOLD:AAD4375
Chrysops frigidus|[5600]|JWDCJ166-11|BOLD:AAD4375
Chrysops frigidus|[5601]|JWDCG837-10|BOLD:AAD4375
Chrysops frigidus|[5602]|JWDCI233-10|BOLD:AAD4375
Chrysops frigidus|[5603]|JWDCG183-10|BOLD:AAD4375
Chrysops frigidus|[5604]|JWDCF356-10|BOLD:AAD4375
Chrysops frigidus|[5605]|JWDCF878-10|BOLD:AAD4375
Chrysops furcatus|[5606]|JWDCI148-10|BOLD:AAB1142
Chrysops furcatus|[5607]|JWDCB042-10|BOLD:AAB1142
Chrysops furcatus|[5608]|JWDCI560-10|BOLD:AAB1142
Chrysops furcatus|[5609]|JWDCG137-10|BOLD:AAB1142
Chrysops furcatus|[5610]|MHFLI875-09|BOLD:AAB1142
Chrysops furcatus|[5611]|JWDCI409-10|BOLD:AAB1142
Chrysops furcatus|[5612]|JWDCG138-10|BOLD:AAB1142
Chrysops furcatus|[5613]|JWDCG139-10|BOLD:AAB1142
Chrysops furcatus|[5614]|SAMOS095-09|BOLD:AAB1142
Chrysops furcatus|[5615]|JWDCI360-10|BOLD:AAB1142
Chrysops furcatus|[5616]|JWDCH735-10|BOLD:AAB1142
Chrysops furcatus|[5617]|JWDCL113-11|BOLD:AAB1142
Chrysops furcatus|[5618]|JWDCB046-10|BOLD:AAB1142
Chrysops furcatus|[5619]|JWDCB040-10|BOLD:AAB1142
Chrysops furcatus|[5620]|JWDCF368-10|BOLD:AAB1142
Chrysops mitis|[5621]|JWDCI820-11|BOLD:AAA8870
Chrysops ater|[5622]|JWDCB039-10|BOLD:ACE5640
Chrysops ater|[5623]|JWDCA538-10|BOLD:AAB4140
Chrysops ater|[5624]|JWDCI592-10|BOLD:AAB4140
Chrysops ater|[5625]|JWDCB031-10|BOLD:AAB4140
Chrysops ater|[5626]|JWDCJ1524-11|BOLD:AAB4140
Chrysops ater|[5627]|JWDCF857-10|BOLD:AAB4140
Chrysops ater|[5628]|JWDCJ774-11|BOLD:AAB4140
Chrysops ater|[5629]|JWDCJ776-11|BOLD:AAB4140
Chrysops ater|[5630]|JWDCJ078-11|BOLD:AAB4140
Chrysops ater|[5631]|JWDCI593-10|BOLD:AAB4140
Chrysops ater|[5632]|JWDCI561-10|BOLD:AAB4140
Chrysops ater|[5633]|JWDCI948-11|BOLD:AAB4140
Chrysops ater|[5634]|JWDCF169-10|BOLD:AAB4140
Chrysops ater|[5635]|JWDCF879-10|BOLD:AAB4140
Chrysops nigripes|[5636]|JWDCA541-10|BOLD:AAF2575
Chrysops nigripes|[5637]|JWDCH873-10|BOLD:AAF2575
Chrysops nigripes|[5638]|JWDCB037-10|BOLD:AAF2575
Chrysops nigripes|[5639]|JWDCA542-10|BOLD:AAF2575
Chrysops nigripes|[5640]|JWDCJ1190-11|BOLD:AAF2575
Chrysops nigripes|[5641]|JWDCI074-10|BOLD:AAF2575
Chrysops nigripes|[5642]|JWDCI410-10|BOLD:AAF2575
Chrysops nigripes|[5643]|JWDCB032-10|BOLD:AAF2575
Chrysops nigripes|[5644]|JWDCL318-11|BOLD:AAF2575
Chrysops nigripes|[5645]|JWDCI411-10|BOLD:AAF2575
Chrysops nigripes|[5646]|JWDCF167-10|BOLD:AAF2575
Chrysops nigripes|[5647]|JWDCJ155-11|BOLD:AAF2575
Chrysops nigripes|[5648]|JWDCB030-10|BOLD:AAF2575
Chrysops nigripes|[5649]|JWDCB044-10|BOLD:AAF2575
Chrysops nigripes|[5650]|JWDCK231-11|BOLD:AAF2575
Chrysops nigripes|[5651]|JWDCJ841-11|BOLD:AAF2575
Chrysops nigripes|[5652]|JWDCJ997-11|BOLD:AAF2575
Chrysops nigripes|[5653]|JWDCL052-11|BOLD:AAF2575
Chrysops nigripes|[5654]|JWDCL243-11|BOLD:AAF2575
Chrysops nigripes|[5655]|JWDCJ076-11|BOLD:AAF2575
Chrysops nigripes|[5656]|JWDCG645-10|BOLD:AAF2575
Chrysops nigripes|[5657]|JWDCG184-10|BOLD:AAF2575
Chrysops nigripes|[5658]|JWDCF914-10|BOLD:AAF2575
Chrysops nigripes|[5659]|JWDCJ703-11|BOLD:AAF2575
Chrysops nigripes|[5660]|JWDCB028-10|BOLD:AAF2575
Chrysops nigripes|[5661]|JWDCB027-10|BOLD:AAF2575
Chrysops nigripes|[5662]|JWDCA540-10|BOLD:AAF2575
Chrysops nigripes|[5663]|JWDCA537-10|BOLD:AAF2575
Chrysops nigripes|[5664]|JWDCB038-10|BOLD:AAF2575
Chrysops nigripes|[5665]|JWDCB043-10|BOLD:AAF2575
Chrysops nigripes|[5666]|JWDCB026-10|BOLD:AAF2575
Chrysops nigripes|[5667]|JWDCK184-11|BOLD:AAF2575
Chrysops nigripes|[5668]|JWDCJ701-11|BOLD:AAF2575
Chrysops nigripes|[5669]|JWDCK232-11|BOLD:AAF2575
Chrysops nigripes|[5670]|JWDCJ996-11|BOLD:AAF2575
Chrysops nigripes|[5671]|JWDCJ1038-11|BOLD:AAF2575
Chrysops nigripes|[5672]|JWDCJ1398-11|BOLD:AAF2575

Chrysops nigripes|[5670]|JWDCJ996-11|BOLD:AAF2575
Chrysops nigripes|[5671]|JWDCJ1038-11|BOLD:AAF2575
Chrysops nigripes|[5672]|JWDCJ1398-11|BOLD:AAF2575
Chrysops nigripes|[5673]|JWDCJ1399-11|BOLD:AAF2575
Chrysops nigripes|[5674]|JWDCL317-11|BOLD:AAF2575
Chrysops nigripes|[5675]|JWDCL366-11|BOLD:AAF2575
Chrysops nigripes|[5676]|JWDCL402-11|BOLD:AAF2575
Chrysops nigripes|[5677]|JWDCG801-10|BOLD:AAF2575
Chrysops nigripes|[5678]|JWDCI607-10|BOLD:AAF2575
Chrysops nigripes|[5679]|JWDCG056-10|BOLD:AAF2575
Chrysops nigripes|[5680]|JWDCI073-10|BOLD:AAF2575
Chrysops nigripes|[5681]|JWDCI767-11|BOLD:AAF2575
Chrysops nigripes|[5682]|JWDCI768-11|BOLD:AAF2575
Chrysops nigripes|[5683]|JWDCI769-11|BOLD:AAF2575
Chrysops nigripes|[5684]|JWDCI875-11|BOLD:AAF2575
Chrysops nigripes|[5685]|JWDCI876-11|BOLD:AAF2575
Chrysops nigripes|[5686]|JWDCG182-10|BOLD:AAF2575
Chrysops nigripes|[5687]|JWDCF915-10|BOLD:AAF2575
Chrysops nigripes|[5688]|JWDCL403-11|BOLD:AAF2575
Chrysops nigripes|[5689]|JWDCG646-10|BOLD:AAF2575
Chrysops nigripes|[5690]|JWDCA539-10|BOLD:AAF2575
Chrysops nigripes|[5691]|JWDCJ735-11|BOLD:AAF2575
Chrysops nigripes|[5692]|JWDCJ1189-11|BOLD:AAF2575
Chrysops nigripes|[5693]|JWDCL450-11|BOLD:AAF2575
Chrysops nigripes|[5694]|JWDCG838-10|BOLD:AAF2575
Chrysops nigripes|[5695]|JWDCF916-10|BOLD:AAF2575
Atylotus sublunaticornis|[5696]|JWDCJ758-11|BOLD:ABY8099
Atylotus sublunaticornis|[5697]|JWDCJ757-11|BOLD:ABY8099
Atylotus sublunaticornis|[5698]|JWDCJ759-11|BOLD:ABY8099
Atylotus sp.|[5699]|JWDCG136-10|BOLD:AAI4870
Hybomitra frontalis|[5700]|MHFLY007-06|BOLD:AAA3296
Hybomitra frontalis|[5701]|JWDCJ702-11|BOLD:AAA3296
Hybomitra frontalis|[5702]|JWDCI712-11|BOLD:AAA3296
Hybomitra frontalis|[5703]|JWDCH387-10|BOLD:AAA3296
Hybomitra frontalis|[5704]|JWDCG701-10|BOLD:AAA3296
Hybomitra frontalis|[5705]|JWDCB048-10|BOLD:AAA3296
Hybomitra frontalis|[5706]|JWDCI463-10|BOLD:AAA3296
Hybomitra frontalis|[5707]|JWDCI464-10|BOLD:AAA3296
Hybomitra frontalis|[5708]|JWDCF168-10|BOLD:AAA3296
Hybomitra frontalis|[5709]|JWDCL365-11|BOLD:AAA3296
Hybomitra frontalis|[5710]|JWDCB029-10|BOLD:AAA3296
Hybomitra frontalis|[5711]|JWDCK021-11|BOLD:AAA3296
Hybomitra frontalis|[5712]|JWDCJ1264-11|BOLD:AAA3296
Hybomitra frontalis|[5713]|JWDCL319-11|BOLD:AAA3296
Hybomitra frontalis|[5714]|JWDCJ217-11|BOLD:AAA3296
Hybomitra frontalis|[5715]|JWDCG857-10|BOLD:AAA3296
Hybomitra frontalis|[5716]|JWDCH001-10|BOLD:AAA3296
Hybomitra frontalis|[5717]|JWDCL367-11|BOLD:AAA3296
Hybomitra frontalis|[5718]|JWDCL589-11|BOLD:AAA3296
Hybomitra frontalis|[5719]|JWDCI947-11|BOLD:AAA3296
Hybomitra frontalis|[5720]|JWDCG841-10|BOLD:AAA3296
Hybomitra frontalis|[5721]|JWDCL316-11|BOLD:AAA3296
Hybomitra frontalis|[5722]|JWDCI818-11|BOLD:AAA3296
Hybomitra frontalis|[5723]|MHFLI009-06|BOLD:AAA3296
Hybomitra frontalis|[5724]|JWDCJ303-11|BOLD:AAA3296
Hybomitra frontalis|[5725]|JWDCJ1039-11|BOLD:AAA3296
Hybomitra frontalis|[5726]|JWDCJ998-11|BOLD:AAA3296
Hybomitra frontalis|[5727]|JWDCJ1602-11|BOLD:AAA3296
Hybomitra frontalis|[5728]|JWDCJ165-11|BOLD:AAA3296
Hybomitra frontalis|[5729]|JWDCB045-10|BOLD:AAA3296
Hybomitra frontalis|[5730]|MMADL061-09|BOLD:AAA3296
Hybomitra frontalis|[5731]|JWDCL590-11|BOLD:AAA3296
Hybomitra frontalis|[5732]|JWDCG769-10|BOLD:AAA3296
Hybomitra frontalis|[5733]|JWDCG858-10|BOLD:AAA3296
Hybomitra frontalis|[5734]|JWDCI819-11|BOLD:AAA3296
Hybomitra frontalis|[5735]|JWDCI877-11|BOLD:AAA3296
Hybomitra frontalis|[5736]|JWDCH243-10|BOLD:AAA3296
Hybomitra frontalis|[5737]|JWDCH199-10|BOLD:AAA3296
Hybomitra frontalis|[5738]|MHFLI005-06|BOLD:AAA3296
Hybomitra frontalis|[5739]|JWDCH175-10|BOLD:AAA3296
Hybomitra frontalis|[5740]|JWDCH318-10|BOLD:AAA3296
Hybomitra frontalis|[5741]|JWDCJ215-11|BOLD:AAA3296
Hybomitra frontalis|[5742]|JWDCI462-10|BOLD:AAA3296
Hybomitra frontalis|[5743]|JWDCK121-11|BOLD:AAA3296
Hybomitra frontalis|[5744]|MHFLI006-06|BOLD:AAA3296
Hybomitra frontalis|[5745]|JWDCH319-10|BOLD:AAA3296
Hybomitra frontalis|[5746]|JWDCF367-10|BOLD:AAA3296
Hybomitra frontalis|[5747]|JWDCK020-11|BOLD:AAA3296
Hybomitra frontalis|[5748]|JWDCG891-10|BOLD:AAA3296
Hybomitra frontalis|[5749]|JWDCJ1523-11|BOLD:AAA3296
Hybomitra frontalis|[5750]|JWDCK123-11|BOLD:AAA3296
Hybomitra frontalis|[5751]|JWDCJ302-11|BOLD:AAA3296
Hybomitra frontalis|[5752]|JWDCH002-10|BOLD:AAA3296
Hybomitra frontalis|[5753]|JWDCH365-10|BOLD:AAA3296
Hybomitra frontalis|[5754]|JWDCF855-10|BOLD:AAA3296
Hybomitra frontalis|[5755]|JWDCJ301-11|BOLD:AAA3296
Hybomitra frontalis|[5756]|JWDCJ1146-11|BOLD:AAA3296
Hybomitra frontalis|[5757]|JWDCL187-11|BOLD:AAA3296
Hybomitra frontalis|[5758]|JWDCG601-10|BOLD:AAA3296
Hybomitra frontalis|[5759]|JWDCG889-10|BOLD:AAA3296
Hybomitra frontalis|[5760]|JWDCJ164-11|BOLD:AAA3296
Hybomitra frontalis|[5761]|JWDCH244-10|BOLD:AAA3296
Hybomitra frontalis|[5762]|MHFLY008-06|BOLD:AAA3296
Hybomitra frontalis|[5763]|JWDCG644-10|BOLD:AAA3296
Hybomitra frontalis|[5764]|JWDCJ775-11|BOLD:AAA3296
Hybomitra frontalis|[5765]|JWDCG890-10|BOLD:AAA3296
Hybomitra frontalis|[5766]|JWDCF869-10|BOLD:AAA3296
Hybomitra frontalis|[5767]|JWDCI713-11|BOLD:AAA3296
Hybomitra frontalis|[5768]|JWDCH386-10|BOLD:AAA3296
Hybomitra frontalis|[5769]|JWDCF912-10|BOLD:AAA3296
Hybomitra itasca|[5770]|JWDCK022-11|BOLD:AAE1681
Hybomitra itasca|[5771]|JWDCJ1263-11|BOLD:AAE1681
Hybomitra itasca|[5772]|JWDCL051-11|BOLD:AAE1681

Hybomitra frontalis|[5769]|JWDCF912-10|BOLD:[illegible]
Hybomitra itasca|[5770]|JWDCK022-11|BOLD:AAE1681
Hybomitra itasca|[5771]|JWDCJ1263-11|BOLD:AAE1681
Hybomitra itasca|[5772]|JWDCL051-11|BOLD:AAE1681
Hybomitra itasca|[5773]|JWDCL245-11|BOLD:AAE1681
Hybomitra itasca|[5774]|JWDCL244-11|BOLD:AAE1681
Hybomitra itasca|[5775]|JWDCK185-11|BOLD:AAE1681
Hybomitra hearlei|[5776]|JWDCI664-10|BOLD:AAE1681
Hybomitra hearlei|[5777]|SAMOS074-09|BOLD:AAE1681
Hybomitra hearlei|[5778]|SAMOS073-09|BOLD:AAE1681
Hybomitra hearlei|[5779]|JWDCJ737-11|BOLD:AAE1681
Hybomitra hearlei|[5780]|JWDCI408-10|BOLD:AAE1681
Hybomitra zonalis|[5781]|JWDCC284-10|BOLD:AAD8656
Hybomitra zonalis|[5782]|JWDCC283-10|BOLD:AAD8656
Hybomitra zonalis|[5783]|SAMOS071-09|BOLD:AAD8656
Hybomitra zonalis|[5784]|SAMOS072-09|BOLD:AAD8656
Hybomitra zonalis|[5785]|JWDCC285-10|BOLD:AAD8656
Hybomitra affinis|[5786]|JWDCJ1145-11|BOLD:AAB1544
Hybomitra affinis|[5787]|JWDCB050-10|BOLD:AAB1544
Hybomitra affinis|[5788]|JWDCC281-10|BOLD:AAB1544
Hybomitra affinis|[5789]|JWDCC443-10|BOLD:AAB1544
Hybomitra affinis|[5790]|JWDCC457-10|BOLD:AAB1544
Hybomitra affinis|[5791]|JWDCC458-10|BOLD:AAB1544
Hybomitra affinis|[5792]|JWDCI363-10|BOLD:AAB1544
Hybomitra affinis|[5793]|SAMOS077-09|BOLD:AAB1544
Hybomitra affinis|[5794]|SAMOS076-09|BOLD:AAB1544
Hybomitra affinis|[5795]|JWDCC438-10|BOLD:AAB1544
Hybomitra affinis|[5796]|JWDCI412-10|BOLD:AAB1544
Hybomitra affinis|[5797]|JWDCI950-11|BOLD:AAB1544
Hybomitra affinis|[5798]|JWDCD174-10|BOLD:AAB1544
Hybomitra affinis|[5799]|JWDCD180-10|BOLD:AAB1544
Hybomitra affinis|[5800]|DSCNI011-07|BOLD:AAB1544
Hybomitra affinis|[5801]|JWDCB033-10|BOLD:AAB1544
Hybomitra affinis|[5802]|JWDCI949-11|BOLD:AAB1544
Hybomitra affinis|[5803]|JWDCL186-11|BOLD:AAB1544
Hybomitra affinis|[5804]|JWDCK186-11|BOLD:AAB1544
Hybomitra affinis|[5805]|MHFLI874-09|BOLD:AAB1544
Hybomitra affinis|[5806]|SAMOS081-09|BOLD:AAB1544
Hybomitra affinis|[5807]|JWDCC051-10|BOLD:AAB1544
Hybomitra affinis|[5808]|SAMOS084-09|BOLD:AAB1544
Hybomitra affinis|[5809]|JWDCH554-10|BOLD:AAB1544
Hybomitra arpadi|[5810]|JWDCD173-10|BOLD:AAB1544
Hybomitra arpadi|[5811]|JWDCC468-10|BOLD:AAB1544
Hybomitra arpadi|[5812]|JWDCJ842-11|BOLD:AAB1544
Hybomitra arpadi|[5813]|JWDCJ401-11|BOLD:AAB1544
Hybomitra arpadi|[5814]|JWDCC222-10|BOLD:AAB1544
Hybomitra arpadi|[5815]|JWDCC467-10|BOLD:AAB1544
Hybomitra arpadi|[5816]|JWDCC442-10|BOLD:AAB1544
Hybomitra arpadi|[5817]|JWDCC050-10|BOLD:AAB1544
Hybomitra arpadi|[5818]|SAMOS080-09|BOLD:AAB1544
Hybomitra arpadi|[5819]|SAMOS079-09|BOLD:AAB1544
Hybomitra arpadi|[5820]|SAMOS075-09|BOLD:AAB1544
Hybomitra arpadi|[5821]|JWDCB036-10|BOLD:AAB1544
Hybomitra arpadi|[5822]|JWDCC210-10|BOLD:AAB1544
Hybomitra arpadi|[5823]|JWDCC451-10|BOLD:AAB1544
Hybomitra arpadi|[5824]|JWDCJ400-11|BOLD:AAB1544
Hybomitra arpadi|[5825]|JWDCJ903-11|BOLD:AAB1544
Hybomitra arpadi|[5826]|JWDCJ1191-11|BOLD:AAB1544
Hybomitra arpadi|[5827]|JWDCJ1265-11|BOLD:AAB1544
Hybomitra arpadi|[5828]|JWDCJ077-11|BOLD:AAB1544
Hybomitra arpadi|[5829]|JWDCI608-10|BOLD:AAB1544
Hybomitra arpadi|[5830]|JWDCI605-10|BOLD:AAB1544
Hybomitra arpadi|[5831]|JWDCF867-10|BOLD:AAB1544
Hybomitra arpadi|[5832]|JWDCB035-10|BOLD:AAB1544
Hybomitra arpadi|[5833]|SAMOS082-09|BOLD:AAB1544
Hybomitra arpadi|[5834]|JWDCC219-10|BOLD:AAB1544
Hybomitra arpadi|[5835]|JWDCC440-10|BOLD:AAB1544
Hybomitra arpadi|[5836]|JWDCC441-10|BOLD:AAB1544
Hybomitra arpadi|[5837]|JWDCC460-10|BOLD:AAB1544
Hybomitra arpadi|[5838]|JWDCC465-10|BOLD:AAB1544
Hybomitra arpadi|[5839]|JWDCJ904-11|BOLD:AAB1544
Hybomitra arpadi|[5840]|JWDCL401-11|BOLD:AAB1544
Hybomitra arpadi|[5841]|SAMOS083-09|BOLD:AAB1544
Hybomitra arpadi|[5842]|JWDCB049-10|BOLD:AAB1544
Hybomitra arpadi|[5843]|JWDCB047-10|BOLD:AAB1544
Hybomitra arpadi|[5844]|JWDCB034-10|BOLD:AAB1544
Hybomitra arpadi|[5845]|SAMOS078-09|BOLD:AAB1544
Hybomitra arpadi|[5846]|JWDCC280-10|BOLD:AAB1544
Hybomitra arpadi|[5847]|JWDCK124-11|BOLD:AAB1544
Hybomitra arpadi|[5848]|JWDCJ843-11|BOLD:AAB1544
Hybomitra arpadi|[5849]|JWDCJ902-11|BOLD:AAB1544
Hybomitra arpadi|[5850]|JWDCC209-10|BOLD:AAB1544
Hybomitra arpadi|[5851]|JWDCC459-10|BOLD:AAB1544
Hybomitra arpadi|[5852]|JWDCI606-10|BOLD:AAB1544
Hybomitra arpadi|[5853]|JWDCI878-11|BOLD:AAB1544
Hybomitra arpadi|[5854]|JWDCC049-10|BOLD:AAB1544
Hybomitra arpadi|[5855]|JWDCC220-10|BOLD:AAB1544
Hybomitra arpadi|[5856]|JWDCJ399-11|BOLD:AAB1544
Hybomitra arpadi|[5857]|JWDCF552-10|BOLD:AAB1544
Hybomitra arpadi|[5858]|JWDCC223-10|BOLD:AAB1544
Hybomitra arpadi|[5859]|JWDCD175-10|BOLD:AAB1544
Hybomitra arpadi|[5860]|JWDCJ1040-11|BOLD:AAB1544
Hybomitra arpadi|[5861]|JWDCJ1144-11|BOLD:AAB1544
Hybomitra arpadi|[5862]|JWDCF868-10|BOLD:AAB1544
Hybomitra illota|[5863]|SAMOS093-09|BOLD:AAD1086
Hybomitra lurida|[5864]|JWDCF508-10|BOLD:AAB0648
Hybomitra lurida|[5865]|JWDCF369-10|BOLD:AAB0648
Hybomitra lurida|[5866]|SAMOS094-09|BOLD:AAB0648
Hybomitra lurida|[5867]|JWDCI562-10|BOLD:AAB0648
Hybomitra lurida|[5868]|SAMOS088-09|BOLD:AAB0648
Hybomitra lurida|[5869]|JWDCC206-10|BOLD:AAB0648
Hybomitra lurida|[5870]|JWDCC208-10|BOLD:AAB0648
Hybomitra lurida|[5871]|JWDCI663-10|BOLD:AAB0648

Hybomitra lurida|[5869]|JWDCC206-10|BOLD:AAB0648
Hybomitra lurida|[5870]|JWDCC208-10|BOLD:AAB0648
Hybomitra lurida|[5871]|JWDCI663-10|BOLD:AAB0648
Hybomitra lurida|[5872]|JWDCF856-10|BOLD:AAB0648
Hybomitra lurida|[5873]|JWDCF537-10|BOLD:AAB0648
Hybomitra lurida|[5874]|JWDCF509-10|BOLD:AAB0648
Hybomitra lurida|[5875]|JWDCE480-10|BOLD:AAB0648
Hybomitra lurida|[5876]|JWDCE479-10|BOLD:AAB0648
Hybomitra lurida|[5877]|JWDCI662-10|BOLD:AAB0648
Hybomitra lurida|[5878]|JWDCJ582-11|BOLD:AAB0648
Hybomitra lurida|[5879]|JWDCC452-10|BOLD:AAB0648
Hybomitra lurida|[5880]|JWDCC221-10|BOLD:AAB0648
Hybomitra lurida|[5881]|JWDCC207-10|BOLD:AAB0648
Hybomitra lurida|[5882]|SAMOS086-09|BOLD:AAB0648
Hybomitra lurida|[5883]|SAMOS091-09|BOLD:AAB0648
Hybomitra lurida|[5884]|SAMOS089-09|BOLD:AAB0648
Hybomitra lurida|[5885]|JWDCE550-10|BOLD:AAB0648
Hybomitra lurida|[5886]|JWDCE481-10|BOLD:AAB0648
Hybomitra lurida|[5887]|JWDCE887-10|BOLD:AAB0648
Hybomitra lurida|[5888]|JWDCE889-10|BOLD:AAB0648
Hybomitra lurida|[5889]|JWDCJ581-11|BOLD:AAB0648
Hybomitra lurida|[5890]|JWDCG120-10|BOLD:AAB0648
Hybomitra lurida|[5891]|SAMOS085-09|BOLD:AAB0648
Hybomitra lurida|[5892]|SAMOS090-09|BOLD:AAB0648
Hybomitra lurida|[5893]|SAMOS092-09|BOLD:AAB0648
Hybomitra lurida|[5894]|JWDCF510-10|BOLD:AAB0648
Hybomitra lurida|[5895]|JWDCG122-10|BOLD:AAB0648
Hybomitra lurida|[5896]|JWDCC439-10|BOLD:AAB0648
Hybomitra lurida|[5897]|JWDCC466-10|BOLD:AAB0648
Hybomitra lurida|[5898]|JWDCJ736-11|BOLD:AAB0648
Hybomitra lurida|[5899]|JWDCI361-10|BOLD:AAB0648
Hybomitra lurida|[5900]|JWDCE820-10|BOLD:AAB0648
Hybomitra lurida|[5901]|JWDCC230-10|BOLD:AAB0648
Hybomitra lurida|[5902]|SAMOS087-09|BOLD:AAB0648
Hybomitra lurida|[5903]|JWDCC052-10|BOLD:AAB0648
Hybomitra lurida|[5904]|JWDCJ583-11|BOLD:AAB0648
Hybomitra lurida|[5905]|JWDCI362-10|BOLD:AAB0648
Hybomitra lurida|[5906]|JWDCG121-10|BOLD:AAB0648
Hybomitra lurida|[5907]|JWDCE818-10|BOLD:AAB0648
Hybomitra lurida|[5908]|JWDCE819-10|BOLD:AAB0648
Hybomitra lurida|[5909]|JWDCE888-10|BOLD:AAB0648
Hybomitra lurida|[5910]|JWDCE551-10|BOLD:AAB0648
Ephydridae|[5911]|FLIMH003-08|BOLD:AAG2731
Bolitophila cinerea|[5912]|MHMOS094-07|BOLD:AAG4863
Bolitophila cinerea|[5913]|PHTCH170-08|BOLD:AAG4863
Euhybus triplex|[5914]|JWDOL651-10|BOLD:AAF9778
Euhybus triplex|[5915]|JWDOL749-10|BOLD:AAF9778
Euhybus triplex|[5916]|JWDOL567-10|BOLD:AAF9778
Euhybus triplex|[5917]|JWDCH256-10|BOLD:AAF9778
Euhybus triplex|[5918]|JWDOL624-10|BOLD:AAF9778
Euhybus triplex|[5919]|JWDOL636-10|BOLD:AAF9778
Euhybus triplex|[5920]|JWEMP232-10|BOLD:AAF9778
Euhybus triplex|[5921]|JWDOL673-10|BOLD:AAF9778
Euhybus triplex|[5922]|JWEMP261-10|BOLD:AAF9778
Euhybus triplex|[5923]|JWDOL707-10|BOLD:AAF9778
Euhybus triplex|[5924]|JWEMP184-10|BOLD:AAF9778
Euhybus triplex|[5925]|JWEMP182-10|BOLD:AAF9778
Euhybus triplex|[5926]|JWEMP159-10|BOLD:AAF9778
Euhybus triplex|[5927]|JWEMP093-10|BOLD:AAF9778
Euhybus triplex|[5928]|JWEMP009-10|BOLD:AAF9778
Euhybus triplex|[5929]|JWDOL756-10|BOLD:AAF9778
Euhybus triplex|[5930]|JWDOL623-10|BOLD:AAF9778
Euhybus triplex|[5931]|JWDOL834-10|BOLD:AAF9778
Euhybus triplex|[5932]|JWDOL831-10|BOLD:AAF9778
Euhybus triplex|[5933]|JWDCA085-10|BOLD:AAF9778
Euhybus triplex|[5934]|JWEMP264-10|BOLD:AAF9778
Euhybus triplex|[5935]|JWDOL509-10|BOLD:AAF9778
Euhybus triplex|[5936]|JWDOL577-10|BOLD:AAF9778
Euhybus triplex|[5937]|JWDOL580-10|BOLD:AAF9778
Euhybus triplex|[5938]|JWDOL700-10|BOLD:AAF9778
Euhybus triplex|[5939]|JWDOL660-10|BOLD:AAF9778
Euhybus triplex|[5940]|JWEMP258-10|BOLD:AAF9778
Euhybus triplex|[5941]|JWEMP259-10|BOLD:AAF9778
Euhybus triplex|[5942]|JWDOL764-10|BOLD:AAF9778
Euhybus triplex|[5943]|JWDOL592-10|BOLD:AAF9778
Euhybus triplex|[5944]|JWDOL771-10|BOLD:AAF9778
Euhybus triplex|[5945]|JWEMP272-10|BOLD:AAF9778
Euhybus triplex|[5946]|JWDOL762-10|BOLD:AAF9778
Euhybus triplex|[5947]|JWDOL758-10|BOLD:AAF9778
Euhybus triplex|[5948]|JWDOL752-10|BOLD:AAF9778
Euhybus triplex|[5949]|JWDOL800-10|BOLD:AAF9778
Euhybus triplex|[5950]|JWEMP012-10|BOLD:AAF9778
Euhybus triplex|[5951]|JWDOL697-10|BOLD:AAF9778
Euhybus triplex|[5952]|JWEMP004-10|BOLD:AAF9778
Euhybus triplex|[5953]|JWEMP079-10|BOLD:AAF9778
Euhybus triplex|[5954]|JWDOL799-10|BOLD:AAF9778
Euhybus triplex|[5955]|JWDOL566-10|BOLD:AAF9778
Euhybus triplex|[5956]|JWEMP008-10|BOLD:AAF9778
Euhybus triplex|[5957]|JWEMP284-10|BOLD:AAF9778
Euhybus triplex|[5958]|TWDIP312-09|BOLD:AAF9778
Euhybus triplex|[5959]|JWEMP124-10|BOLD:AAF9778
Euhybus triplex|[5960]|JWEMP125-10|BOLD:AAF9778
Euhybus triplex|[5961]|JWEMP183-10|BOLD:AAF9778
Euhybus triplex|[5962]|JWEMP234-10|BOLD:AAF9778
Euhybus triplex|[5963]|JWDOL775-10|BOLD:AAF9778
Euhybus triplex|[5964]|JWDOL776-10|BOLD:AAF9778
Euhybus triplex|[5965]|JWDOL698-10|BOLD:AAF9778
Euhybus triplex|[5966]|JWDOL699-10|BOLD:AAF9778
Euhybus triplex|[5967]|JWEMP254-10|BOLD:AAF9778
Euhybus triplex|[5968]|JWEMP292-10|BOLD:AAF9778
Euhybus triplex|[5969]|JWEMP005-10|BOLD:AAF9778
Euhybus triplex|[5970]|JWEMP090-10|BOLD:AAF9778
Euhybus triplex|[5971]|JWDOL477-10|BOLD:AAF9778

[illegible]
Euhybus triplex|[5969]|JWEMP005-10|BOLD:AAF9778
Euhybus triplex|[5970]|JWEMP090-10|BOLD:AAF9778
Euhybus triplex|[5971]|JWDOL477-10|BOLD:AAF9778
Euhybus triplex|[5972]|JWDOL547-10|BOLD:AAF9778
Euhybus triplex|[5973]|JWDOL565-10|BOLD:AAF9778
Euhybus triplex|[5974]|JWDOL568-10|BOLD:AAF9778
Euhybus triplex|[5975]|JWDOL578-10|BOLD:AAF9778
Euhybus triplex|[5976]|JWDOL579-10|BOLD:AAF9778
Euhybus triplex|[5977]|JWDOL581-10|BOLD:AAF9778
Euhybus triplex|[5978]|JWDOL661-10|BOLD:AAF9778
Euhybus triplex|[5979]|JWDOL701-10|BOLD:AAF9778
Euhybus triplex|[5980]|JWDOL702-10|BOLD:AAF9778
Euhybus triplex|[5981]|JWEMP091-10|BOLD:AAF9778
Euhybus triplex|[5982]|JWEMP092-10|BOLD:AAF9778
Euhybus triplex|[5983]|JWDOL774-10|BOLD:AAF9778
Euhybus triplex|[5984]|JWDOL773-10|BOLD:AAF9778
Euhybus triplex|[5985]|JWDOL769-10|BOLD:AAF9778
Euhybus triplex|[5986]|JWDOL759-10|BOLD:AAF9778
Euhybus triplex|[5987]|JWDOL751-10|BOLD:AAF9778
Euhybus triplex|[5988]|JWDOL725-10|BOLD:AAF9778
Euhybus triplex|[5989]|JWDOL721-10|BOLD:AAF9778
Euhybus triplex|[5990]|JWDOL625-10|BOLD:AAF9778
Euhybus triplex|[5991]|JWDOL621-10|BOLD:AAF9778
Euhybus triplex|[5992]|JWDOL845-10|BOLD:AAF9778
Euhybus triplex|[5993]|JWDCI935-11|BOLD:AAF9778
Euhybus triplex|[5994]|JWDCA087-10|BOLD:AAF9778
Euhybus triplex|[5995]|JWDCA060-10|BOLD:AAF9778
Euhybus triplex|[5996]|JWDOL720-10|BOLD:AAF9778
Euhybus triplex|[5997]|JWDOL496-10|BOLD:AAF9778
Euhybus triplex|[5998]|JWDOL792-10|BOLD:AAF9778
Euhybus triplex|[5999]|JWDOL653-10|BOLD:AAF9778
Euhybus triplex|[6000]|JWEMP233-10|BOLD:AAF9778
Euhybus triplex|[6001]|JWDCA059-10|BOLD:AAF9778
Euhybus triplex|[6002]|JWDOL761-10|BOLD:AAF9778
Euhybus triplex|[6003]|JWEMP011-10|BOLD:AAF9778
Euhybus triplex|[6004]|JWEMP044-10|BOLD:AAF9778
Euhybus triplex|[6005]|JWEMP214-10|BOLD:AAF9778
Euhybus triplex|[6006]|JWDOL541-10|BOLD:AAF9778
Euhybus triplex|[6007]|JWDOL777-10|BOLD:AAF9778
Euhybus triplex|[6008]|JWDOL659-10|BOLD:AAF9778
Euhybus triplex|[6009]|JWDOL836-10|BOLD:AAF9778
Euhybus triplex|[6010]|JWDCI936-11|BOLD:AAF9778
Euhybus triplex|[6011]|MTDIC204-09|BOLD:AAF9778
Euhybus triplex|[6012]|JWDOL843-10|BOLD:AAF9778
Euhybus triplex|[6013]|JWDOL715-10|BOLD:AAF9778
Euhybus triplex|[6014]|JWDOL719-10|BOLD:AAF9778
Euhybus triplex|[6015]|JWEMP010-10|BOLD:AAF9778
Euhybus triplex|[6016]|JWDOL703-10|BOLD:AAF9778
Euhybus triplex|[6017]|JWEMP260-10|BOLD:AAF9778
Euhybus triplex|[6018]|JWDOL546-10|BOLD:AAF9778
Euhybus triplex|[6019]|JWDOL677-10|BOLD:AAF9778
Euhybus triplex|[6020]|JWDOL705-10|BOLD:AAF9778
Euhybus triplex|[6021]|JWDOL783-10|BOLD:AAF9778
Euhybus triplex|[6022]|JWDOL674-10|BOLD:AAF9778
Euhybus triplex|[6023]|JWDOL704-10|BOLD:AAF9778
Euhybus triplex|[6024]|JWDOL784-10|BOLD:AAF9778
Euhybus triplex|[6025]|JWDOL654-10|BOLD:AAF9778
Euhybus triplex|[6026]|JWDOL708-10|BOLD:AAF9778
Euhybus triplex|[6027]|JWDOL801-10|BOLD:AAF9778
Euhybus triplex|[6028]|JWDOL676-10|BOLD:AAF9778
Axysta|[6029]|JWDCE003-10|BOLD:AAP1854
Lamproscatella|[6030]|FLIMH009-08|BOLD:AAG2726
Lamproscatella|[6031]|MHFLI859-07|BOLD:AAG2726
Lamproscatella brunnipennis|[6032]|JWDCJ1307-11|BOLD:AAG2726
Lamproscatella|[6033]|MHFLI864-07|BOLD:AAG2726
Lamproscatella|[6034]|FLIMH030-08|BOLD:AAG2726
Lamproscatella|[6035]|MHFLI806-07|BOLD:AAG2726
Lamproscatella|[6036]|FLIMH008-08|BOLD:AAG2726
Lamproscatella|[6037]|MHFLI846-07|BOLD:AAG2726
Lamproscatella|[6038]|FLIMH012-08|BOLD:AAG2726
Lamproscatella brunnipennis|[6039]|JWDCJ1306-11|BOLD:AAG2726
Lamproscatella|[6040]|MHFLI372-07|BOLD:AAG2726
Lamproscatella|[6041]|MHFLI483-07|BOLD:AAG2726
Lamproscatella|[6042]|FLIMH031-08|BOLD:AAG2726
Lamproscatella|[6043]|FLIMH006-08|BOLD:AAG2726
Lamproscatella brunnipennis|[6044]|JWDCL182-11|BOLD:AAG2726
Lamproscatella brunnipennis|[6045]|JWDCG897-10|BOLD:AAG2726
Ephydra sp. TAW1|[6046]|MHFLI607-07|BOLD:AAG2729
Ephydra sp. TAW1|[6047]|JWDCF077-10|BOLD:AAG2729
Ephydra sp. TAW1|[6048]|JWDCE407-10|BOLD:AAG2729
Ephydra sp. TAW1|[6049]|JWDCE684-10|BOLD:AAG2729
Lamproscatella quadrisetosa|[6050]|JWDCE420-10|BOLD:AAP5196
Lamproscatella quadrisetosa|[6051]|MHFLI463-07|BOLD:AAG2728
Lamproscatella quadrisetosa|[6052]|JWDCI535-10
Lamproscatella quadrisetosa|[6053]|JWDCG314-10|BOLD:AAG2725
Lamproscatella quadrisetosa|[6054]|JWDCG451-10|BOLD:AAG2725
Lamproscatella quadrisetosa|[6055]|JWDCL655-11|BOLD:AAG2725
Lamproscatella quadrisetosa|[6056]|JWDCI083-10|BOLD:AAG2725
Lamproscatella quadrisetosa|[6057]|JWDCH506-10|BOLD:AAG2725
Lamproscatella quadrisetosa|[6058]|JWDCH110-10|BOLD:AAG2725
Lamproscatella quadrisetosa|[6059]|JWDCI081-10|BOLD:AAG2725
Lamproscatella quadrisetosa|[6060]|JWDCH752-10|BOLD:AAG2725
Lamproscatella quadrisetosa|[6061]|FLIMH014-08|BOLD:AAG2725
Lamproscatella quadrisetosa|[6062]|JWDCH702-10|BOLD:AAG2725
Lamproscatella quadrisetosa|[6063]|MHFLI370-07|BOLD:AAG2725
Lamproscatella quadrisetosa|[6064]|JWDCH902-10|BOLD:AAG2725
Lamproscatella quadrisetosa|[6065]|JWDCI128-10|BOLD:AAG2725
Lamproscatella quadrisetosa|[6066]|JWDCE544-10|BOLD:AAG2725
Lamproscatella quadrisetosa|[6067]|JWDCE081-10|BOLD:AAG2725
Lamproscatella quadrisetosa|[6068]|JWDCE190-10|BOLD:AAG2725
Parydra|[6069]|FLIMH026-08|BOLD:AAG2733
Parydra|[6070]|JWDCE369-10|BOLD:AAG2733

Lamproscatella quadrisetosa|[6068]|JWDCE190-10|BOLD:AAG2725
Parydra|[6069]|FLIMH026-08|BOLD:AAG2733
Parydra|[6070]|JWDCE369-10|BOLD:AAG2733
Ephydridae|[6071]|JWDCH527-10|BOLD:AAP1855
Ephydridae|[6072]|JWDCH493-10|BOLD:AAP1855
Ephydridae|[6073]|JWDCH369-10|BOLD:AAP1855
Ephydridae|[6074]|JWDCH750-10|BOLD:AAP1855
Ephydridae|[6075]|JWDCH366-10|BOLD:AAP1855
Ephydridae|[6076]|JWDCH368-10|BOLD:AAP1855
Ephydridae|[6077]|JWDCE022-10|BOLD:AAP1855
Ephydridae|[6078]|JWDCE023-10|BOLD:AAP1855
Paracoenia bisetosa|[6079]|JWDCF040-10|BOLD:AAG2734
Ephydridae|[6080]|JWDCA866-10|BOLD:AAG2769
Lamproscatella muria|[6081]|JWDCJ1481-11|BOLD:AAG2741
Lamproscatella muria|[6082]|JWDCD217-10|BOLD:AAG2741
Lamproscatella muria|[6083]|JWDCG236-10|BOLD:AAG2741
Lamproscatella muria|[6084]|JWDCE447-10|BOLD:AAG2741
Coenia|[6085]|JWDCI438-10|BOLD:AAP8986
Coenia|[6086]|JWDCG702-10|BOLD:AAP8986
Coenia|[6087]|JWDCH457-10|BOLD:AAP8986
Coenia|[6088]|JWDCH459-10|BOLD:AAP8986
Coenia|[6089]|JWDCH012-10|BOLD:AAP8986
Philotelma alaskense|[6090]|JWDCC736-10|BOLD:AAL8953
Philotelma alaskense|[6091]|JWDCE512-10|BOLD:AAL8953
Philotelma alaskense|[6092]|JWDCE513-10|BOLD:AAL8953
Philotelma alaskense|[6093]|JWDCE151-10|BOLD:AAL8953
Scatella stagnalis|[6094]|TWDIP076-09|BOLD:ACZ6583
Scatella stagnalis|[6095]|JWDCE592-10|BOLD:ACZ6583
Scatella stagnalis|[6096]|JWDCA863-10|BOLD:AAG2736
Scatella stagnalis|[6097]|JWDCA864-10|BOLD:AAG2736
Scatella stagnalis|[6098]|JWDCA945-10|BOLD:AAG2736
Scatella stagnalis|[6099]|JWDCA916-10|BOLD:AAG2736
Scatella stagnalis|[6100]|JWDCF186-10|BOLD:AAG2736
Scatella stagnalis|[6101]|JWDCF461-10|BOLD:AAG2736
Scatella crassicosta|[6102]|MHFLI479-07|BOLD:AAG2760
Scatella crassicosta|[6103]|MHFLI734-07|BOLD:AAG2760
Scatella crassicosta|[6104]|JWDCG864-10|BOLD:AAG2760
Scatella crassicosta|[6105]|JWDCE397-10|BOLD:AAG2760
Scatella stagnalis|[6106]|MHFLI867-07|BOLD:ACG1868
Scatella stagnalis|[6107]|MHFLI536-07|BOLD:ACG1868
Scatella stagnalis|[6108]|MHFLI822-07|BOLD:ACG1868
Scatella stagnalis|[6109]|MHFLI345-07|BOLD:ACG1868
Scatella stagnalis|[6110]|MHFLI832-07|BOLD:ACG1868
Scatella stagnalis|[6111]|MHFLI430-07|BOLD:ACG1868
Scatella stagnalis|[6112]|MHFLI725-07|BOLD:ACG1868
Scatella stagnalis|[6113]|MHFLI337-07|BOLD:ACG1868
Scatella stagnalis|[6114]|MHFLI558-07|BOLD:ACG1868
Scatella stagnalis|[6115]|MHFLI319-07|BOLD:ACG1868
Scatella stagnalis|[6116]|MHFLI406-07|BOLD:ACG1868
Scatella stagnalis|[6117]|MHFLI485-07|BOLD:ACG1868
Scatella stagnalis|[6118]|JWDCH330-10|BOLD:ACG1868
Scatella stagnalis|[6119]|MHFLI761-07|BOLD:ACG1868
Scatella stagnalis|[6120]|MHFLI785-07|BOLD:ACG1868
Scatella stagnalis|[6121]|MHFLI764-07|BOLD:ACG1868
Scatella stagnalis|[6122]|MHFLI444-07|BOLD:ACG1868
Scatella stagnalis|[6123]|MHFLI449-07|BOLD:ACG1868
Scatella stagnalis|[6124]|MHFLI788-07|BOLD:ACG1868
Scatella stagnalis|[6125]|MHFLI469-07|BOLD:ACG1868
Scatella stagnalis|[6126]|MHFLI580-07|BOLD:ACG1868
Scatella stagnalis|[6127]|MHFLI585-07|BOLD:ACG1868
Scatella stagnalis|[6128]|MHFLI707-07
Scatella stagnalis|[6129]|MHFLI526-07|BOLD:ACG1868
Scatella stagnalis|[6130]|MHFLI796-07|BOLD:ACG1868
Scatella stagnalis|[6131]|MHFLI803-07|BOLD:ACG1868
Scatella stagnalis|[6132]|MHFLI800-07|BOLD:ACG1868
Scatella stagnalis|[6133]|MHFLI373-07|BOLD:ACG1868
Scatella stagnalis|[6134]|MHFLI468-07|BOLD:ACG1868
Scatella stagnalis|[6135]|MHFLI687-07|BOLD:ACG1868
Scatella stagnalis|[6136]|MHFLI369-07|BOLD:ACG1868
Scatella stagnalis|[6137]|MHFLI491-07|BOLD:ACG1868
Scatella stagnalis|[6138]|MHFLI586-07|BOLD:ACG1868
Scatella stagnalis|[6139]|MHFLI588-07|BOLD:ACG1868
Scatella stagnalis|[6140]|MHFLI452-07|BOLD:ACG1868
Scatella stagnalis|[6141]|FLIMH034-08|BOLD:ACG1868
Scatella stagnalis|[6142]|MHFLI861-07
Scatella stagnalis|[6143]|MHFLI700-07|BOLD:ACG1868
Scatella stagnalis|[6144]|MHFLI873-07|BOLD:ACG1868
Scatella stagnalis|[6145]|MHFLI845-07|BOLD:ACG1868
Scatella stagnalis|[6146]|MHFLI812-07|BOLD:ACG1868
Scatella stagnalis|[6147]|MHFLI849-07|BOLD:ACG1868
Scatella stagnalis|[6148]|MHFLI698-07|BOLD:ACG1868
Scatella stagnalis|[6149]|MHFLI710-07|BOLD:ACG1868
Scatella stagnalis|[6150]|MHFLI590-07|BOLD:ACG1868
Scatella stagnalis|[6151]|MHFLI821-07
Scatella stagnalis|[6152]|MHFLI332-07|BOLD:ACG1868
Scatella stagnalis|[6153]|JWDCH903-10|BOLD:ACG1868
Scatella stagnalis|[6154]|MHFLI705-07
Scatella stagnalis|[6155]|MHFLI819-07|BOLD:ACG1868
Scatella stagnalis|[6156]|MHFLI763-07|BOLD:ACG1868
Scatella stagnalis|[6157]|MHFLI325-07|BOLD:ACG1868
Scatella stagnalis|[6158]|MHFLI416-07|BOLD:ACG1868
Scatella stagnalis|[6159]|MHFLI476-07|BOLD:ACG1868
Scatella stagnalis|[6160]|JWDCH332-10|BOLD:ACG1868
Scatella stagnalis|[6161]|MHFLI769-07|BOLD:ACG1868
Scatella stagnalis|[6162]|MHFLI333-07|BOLD:ACG1868
Scatella stagnalis|[6163]|MHFLI330-07|BOLD:ACG1868
Scatella stagnalis|[6164]|MHFLI870-07|BOLD:ACG1868
Scatella stagnalis|[6165]|MHFLI805-07|BOLD:ACG1868
Scatella stagnalis|[6166]|MHFLI472-07|BOLD:ACG1868
Scatella stagnalis|[6167]|MHFLI853-07|BOLD:ACG1868
Scatella stagnalis|[6168]|MHFLI346-07|BOLD:ACG1868
Scatella stagnalis|[6169]|MHFLI381-07|BOLD:ACG1868
Scatella stagnalis|[6170]|MHFLI385-07|BOLD:ACG1868

Scatella stagnalis|[6167]|MHFLI[illegible]-07|BOLD:ACG1868
Scatella stagnalis|[6168]|MHFLI346-07|BOLD:ACG1868
Scatella stagnalis|[6169]|MHFLI381-07|BOLD:ACG1868
Scatella stagnalis|[6170]|MHFLI385-07|BOLD:ACG1868
Scatella stagnalis|[6171]|MHFLI863-07|BOLD:ACG1868
Scatella stagnalis|[6172]|MHFLI757-07|BOLD:ACG1868
Scatella stagnalis|[6173]|MHFLI414-07|BOLD:ACG1868
Scatella stagnalis|[6174]|MHFLI437-07|BOLD:ACG1868
Scatella stagnalis|[6175]|MHFLI347-07|BOLD:ACG1868
Scatella stagnalis|[6176]|MHFLI334-07|BOLD:ACG1868
Scatella stagnalis|[6177]|MHFLI411-07|BOLD:ACG1868
Scatella stagnalis|[6178]|MHFLI404-07|BOLD:ACG1868
Scatella stagnalis|[6179]|MHFLI379-07|BOLD:ACG1868
Scatella stagnalis|[6180]|MHFLI312-07|BOLD:ACG1868
Scatella stagnalis|[6181]|MHFLI457-07|BOLD:ACG1868
Scatella stagnalis|[6182]|MHFLI461-07|BOLD:ACG1868
Scatella stagnalis|[6183]|MHFLI574-07|BOLD:ACG1868
Scatella stagnalis|[6184]|MHFLI360-07|BOLD:ACG1868
Scatella stagnalis|[6185]|FLIMH002-08|BOLD:ACG1868
Scatella stagnalis|[6186]|JWDCE419-10|BOLD:ACG1868
Scatella stagnalis|[6187]|MHFLI801-07|BOLD:ACG1868
Scatella stagnalis|[6188]|MHFLI493-07|BOLD:ACG1868
Scatella stagnalis|[6189]|MHFLI779-07|BOLD:ACG1868
Scatella stagnalis|[6190]|JWDCH950-10|BOLD:ACG1868
Scatella stagnalis|[6191]|JWDCH456-10|BOLD:ACG1868
Scatella stagnalis|[6192]|JWDCL013-11|BOLD:ACG1868
Scatella stagnalis|[6193]|JWDCL184-11|BOLD:ACG1868
Scatella stagnalis|[6194]|JWDCA865-10|BOLD:ACG1868
Scatella stagnalis|[6195]|MHFLI353-07|BOLD:ACG1868
Scatella stagnalis|[6196]|MHFLI322-07|BOLD:ACG1868
Scatella stagnalis|[6197]|MHFLI488-07|BOLD:ACG1868
Scatella stagnalis|[6198]|MHFLI474-07|BOLD:ACG1868
Scatella stagnalis|[6199]|MHFLI446-07|BOLD:ACG1868
Scatella stagnalis|[6200]|MHFLI431-07|BOLD:ACG1868
Scatella stagnalis|[6201]|MHFLI417-07|BOLD:ACG1868
Scatella stagnalis|[6202]|MHFLI781-07|BOLD:ACG1868
Scatella stagnalis|[6203]|MHFLI549-07|BOLD:ACG1868
Scatella stagnalis|[6204]|MHFLI464-07|BOLD:ACG1868
Scatella stagnalis|[6205]|MHFLI809-07|BOLD:ACG1868
Scatella stagnalis|[6206]|MHFLI554-07|BOLD:ACG1868
Scatella stagnalis|[6207]|JWDCF288-10|BOLD:ACG1868
Scatella stagnalis|[6208]|JWDCE174-10|BOLD:ACG1868
Scatella stagnalis|[6209]|JWDCH531-10|BOLD:ACG1868
Scatella stagnalis|[6210]|JWDCE215-10|BOLD:ACG1868
Scatella stagnalis|[6211]|MHFLI702-07|BOLD:ACG1868
Scatella stagnalis|[6212]|MHFLI354-07|BOLD:ACG1868
Scatella stagnalis|[6213]|MHFLI341-07|BOLD:ACG1868
Scatella stagnalis|[6214]|MHFLI335-07|BOLD:ACG1868
Scatella stagnalis|[6215]|MHFLI386-07|BOLD:ACG1868
Scatella stagnalis|[6216]|MHFLI367-07|BOLD:ACG1868
Scatella stagnalis|[6217]|MHFLI344-07|BOLD:ACG1868
Scatella stagnalis|[6218]|MHFLI358-07|BOLD:ACG1868
Scatella stagnalis|[6219]|MHFLI363-07|BOLD:ACG1868
Scatella stagnalis|[6220]|MHFLI686-07|BOLD:ACG1868
Scatella stagnalis|[6221]|MHFLI314-07|BOLD:ACG1868
Scatella stagnalis|[6222]|MHFLI482-07|BOLD:ACG1868
Scatella stagnalis|[6223]|MHFLI352-07|BOLD:ACG1868
Scatella stagnalis|[6224]|MHFLI548-07|BOLD:ACG1868
Scatella stagnalis|[6225]|JWDCG529-10|BOLD:ACG1868
Scatella stagnalis|[6226]|MHFLI847-07|BOLD:ACG1868
Scatella stagnalis|[6227]|MHFLI448-07|BOLD:ACG1868
Scatella stagnalis|[6228]|MHFLI694-07|BOLD:ACG1868
Scatella stagnalis|[6229]|JWDCJ1308-11|BOLD:ACG1868
Scatella stagnalis|[6230]|MHFLI340-07|BOLD:ACG1868
Scatella stagnalis|[6231]|MHFLI361-07|BOLD:ACG1868
Scatella stagnalis|[6232]|MHFLI342-07|BOLD:ACG1868
Scatella stagnalis|[6233]|JWDCE095-10|BOLD:ACG1868
Scatella stagnalis|[6234]|MHFLI426-07|BOLD:ACG1868
Scatella stagnalis|[6235]|MHFLI454-07|BOLD:ACG1868
Scatella stagnalis|[6236]|MHFLI434-07|BOLD:ACG1868
Scatella stagnalis|[6237]|MHFLI429-07|BOLD:ACG1868
Scatella stagnalis|[6238]|MHFLI425-07|BOLD:ACG1868
Scatella stagnalis|[6239]|MHFLI415-07|BOLD:ACG1868
Scatella stagnalis|[6240]|MHFLI770-07|BOLD:ACG1868
Scatella stagnalis|[6241]|MHFLI848-07|BOLD:ACG1868
Scatella stagnalis|[6242]|MHFLI497-07|BOLD:ACG1868
Scatella stagnalis|[6243]|MHFLI467-07|BOLD:ACG1868
Scatella stagnalis|[6244]|MHFLI436-07|BOLD:ACG1868
Scatella stagnalis|[6245]|JWDCH225-10|BOLD:ACG1868
Scatella stagnalis|[6246]|MHFLI377-07|BOLD:ACG1868
Scatella stagnalis|[6247]|MHFLI460-07|BOLD:ACG1868
Scatella stagnalis|[6248]|MHFLI850-07|BOLD:ACG1868
Scatella stagnalis|[6249]|MHFLI752-07|BOLD:ACG1868
Scatella stagnalis|[6250]|MHFLI828-07|BOLD:ACG1868
Scatella stagnalis|[6251]|MHFLI513-07|BOLD:ACG1868
Scatella stagnalis|[6252]|MHFLI772-07|BOLD:ACG1868
Scatella stagnalis|[6253]|MHFLI844-07|BOLD:ACG1868
Scatella stagnalis|[6254]|MHFLI826-07|BOLD:ACG1868
Scatella stagnalis|[6255]|MHFLI462-07|BOLD:ACG1868
Scatella stagnalis|[6256]|MHFLI445-07|BOLD:ACG1868
Scatella stagnalis|[6257]|MHFLI442-07|BOLD:ACG1868
Scatella stagnalis|[6258]|MHFLI808-07|BOLD:ACG1868
Scatella stagnalis|[6259]|MHFLI487-07|BOLD:ACG1868
Scatella stagnalis|[6260]|MHFLI475-07|BOLD:ACG1868
Scatella stagnalis|[6261]|MHFLI471-07|BOLD:ACG1868
Scatella stagnalis|[6262]|MHFLI470-07|BOLD:ACG1868
Scatella stagnalis|[6263]|MHFLI455-07|BOLD:ACG1868
Scatella stagnalis|[6264]|MHFLI440-07|BOLD:ACG1868
Scatella stagnalis|[6265]|MHFLI420-07|BOLD:ACG1868
Scatella stagnalis|[6266]|MHFLI413-07|BOLD:ACG1868
Scatella stagnalis|[6267]|MHFLI405-07|BOLD:ACG1868
Scatella stagnalis|[6268]|MHFLI351-07|BOLD:ACG1868
Scatella stagnalis|[6269]|MHFLI807-07|BOLD:ACG1868

Scatella stagnalis|[6267]|MHFLI405-07|BOLD:ACG1868
Scatella stagnalis|[6268]|MHFLI351-07|BOLD:ACG1868
Scatella stagnalis|[6269]|MHFLI807-07|BOLD:ACG1868
Scatella stagnalis|[6270]|MHFLI786-07|BOLD:ACG1868
Scatella stagnalis|[6271]|MHFLI412-07|BOLD:ACG1868
Scatella stagnalis|[6272]|MHFLI459-07|BOLD:ACG1868
Scatella stagnalis|[6273]|MHFLI428-07|BOLD:ACG1868
Scatella stagnalis|[6274]|MHFLI833-07|BOLD:ACG1868
Scatella stagnalis|[6275]|MHFLI366-07|BOLD:ACG1868
Scatella stagnalis|[6276]|MHFLI868-07|BOLD:ACG1868
Scatella stagnalis|[6277]|MHFLI811-07|BOLD:ACG1868
Scatella stagnalis|[6278]|JWDCI391-10|BOLD:ACG1868
Scatella stagnalis|[6279]|MHFLI711-07|BOLD:ACG1868
Scatella stagnalis|[6280]|MHFLI862-07|BOLD:ACG1868
Scatella stagnalis|[6281]|MHFLI380-07|BOLD:ACG1868
Scatella stagnalis|[6282]|MHFLI350-07|BOLD:ACG1868
Scatella stagnalis|[6283]|MHFLI458-07|BOLD:ACG1868
Scatella stagnalis|[6284]|MHFLI349-07|BOLD:ACG1868
Scatella stagnalis|[6285]|MHFLI327-07|BOLD:ACG1868
Scatella stagnalis|[6286]|MHFLI326-07|BOLD:ACG1868
Scatella stagnalis|[6287]|MHFLI450-07|BOLD:ACG1868
Scatella stagnalis|[6288]|MHFLI782-07|BOLD:ACG1868
Scatella stagnalis|[6289]|MHFLI810-07|BOLD:ACG1868
Scatella stagnalis|[6290]|MHFLI799-07|BOLD:ACG1868
Scatella stagnalis|[6291]|MHFLI716-07|BOLD:ACG1868
Scatella stagnalis|[6292]|JWDCK046-11|BOLD:ACG1868
Scatella stagnalis|[6293]|JWDCG528-10|BOLD:ACG1868
Scatella stagnalis|[6294]|JWDCH458-10|BOLD:ACG1868
Scatella stagnalis|[6295]|JWDCG452-10|BOLD:ACG1868
Scatella stagnalis|[6296]|JWDCH371-10|BOLD:ACG1868
Scatella stagnalis|[6297]|JWDCH370-10|BOLD:ACG1868
Scatella stagnalis|[6298]|JWDCH367-10|BOLD:ACG1868
Scatella stagnalis|[6299]|JWDCG315-10|BOLD:ACG1868
Scatella stagnalis|[6300]|JWDCG313-10|BOLD:ACG1868
Scatella stagnalis|[6301]|JWDCI390-10|BOLD:ACG1868
Scatella stagnalis|[6302]|JWDCH901-10|BOLD:ACG1868
Scatella stagnalis|[6303]|JWDCH528-10|BOLD:ACG1868
Scatella stagnalis|[6304]|JWDCH830-10|BOLD:ACG1868
Scatella stagnalis|[6305]|JWDCH829-10|BOLD:ACG1868
Scatella stagnalis|[6306]|JWDCH753-10|BOLD:ACG1868
Scatella stagnalis|[6307]|JWDCG730-10|BOLD:ACG1868
Scatella stagnalis|[6308]|JWDCJ149-11|BOLD:ACG1868
Scatella stagnalis|[6309]|JWDCJ148-11|BOLD:ACG1868
Scatella stagnalis|[6310]|JWDCJ147-11|BOLD:ACG1868
Scatella stagnalis|[6311]|JWDCL183-11|BOLD:ACG1868
Scatella stagnalis|[6312]|JWDCL181-11|BOLD:ACG1868
Scatella stagnalis|[6313]|JWDCL168-11|BOLD:ACG1868
Scatella stagnalis|[6314]|JWDCL163-11|BOLD:ACG1868
Scatella stagnalis|[6315]|JWDCL014-11|BOLD:ACG1868
Scatella stagnalis|[6316]|JWDCL012-11|BOLD:ACG1868
Scatella stagnalis|[6317]|MHFLI758-07|BOLD:ACG1868
Scatella stagnalis|[6318]|FLIMH013-08|BOLD:ACG1868
Scatella stagnalis|[6319]|FLIMH001-08|BOLD:ACG1868
Scatella stagnalis|[6320]|MHFLI486-07|BOLD:ACG1868
Scatella stagnalis|[6321]|MHFLI362-07|BOLD:ACG1868
Scatella stagnalis|[6322]|MHFLI329-07|BOLD:ACG1868
Scatella stagnalis|[6323]|MHFLI480-07|BOLD:ACG1868
Scatella stagnalis|[6324]|MHFLI478-07|BOLD:ACG1868
Scatella stagnalis|[6325]|MHFLI447-07|BOLD:ACG1868
Scatella stagnalis|[6326]|MHFLI418-07|BOLD:ACG1868
Scatella stagnalis|[6327]|MHFLI409-07|BOLD:ACG1868
Scatella stagnalis|[6328]|MHFLI378-07|BOLD:ACG1868
Scatella stagnalis|[6329]|MHFLI376-07|BOLD:ACG1868
Scatella stagnalis|[6330]|MHFLI387-07|BOLD:ACG1868
Scatella stagnalis|[6331]|MHFLI496-07|BOLD:ACG1868
Scatella stagnalis|[6332]|JWDCL442-11|BOLD:ACG1868
Scatella stagnalis|[6333]|JWDCL441-11|BOLD:ACG1868
Scatella stagnalis|[6334]|MHFLI750-07|BOLD:ACG1868
Scatella stagnalis|[6335]|MHFLI338-07|BOLD:ACG1868
Scatella stagnalis|[6336]|MHFLI375-07|BOLD:ACG1868
Scatella stagnalis|[6337]|MHFLI695-07|BOLD:ACG1868
Scatella stagnalis|[6338]|MHFLI825-07|BOLD:ACG1868
Scatella stagnalis|[6339]|MHFLI573-07|BOLD:ACG1868
Scatella stagnalis|[6340]|MHFLI311-07|BOLD:ACG1868
Scatella stagnalis|[6341]|MHFLI315-07|BOLD:ACG1868
Scatella stagnalis|[6342]|MHFLI368-07|BOLD:ACG1868
Scatella stagnalis|[6343]|MHFLI356-07|BOLD:ACG1868
Scatella stagnalis|[6344]|MHFLI410-07|BOLD:ACG1868
Scatella stagnalis|[6345]|JWDCI533-10|BOLD:ACG1868
Scatella stagnalis|[6346]|JWDCI826-11|BOLD:ACG1868
Scatella stagnalis|[6347]|JWDCH223-10|BOLD:ACG1868
Scatella stagnalis|[6348]|JWDCI534-10|BOLD:ACG1868
Scatella stagnalis|[6349]|JWDCH331-10|BOLD:ACG1868
Scatella stagnalis|[6350]|JWDCE445-10|BOLD:ACG1868
Scatella stagnalis|[6351]|JWDCE216-10|BOLD:ACG1868
Scatella stagnalis|[6352]|JWDCE217-10|BOLD:ACG1868
Philygria debilis|[6353]|JWDCA910-10|BOLD:AAG2740
Philygria debilis|[6354]|JWDCA915-10|BOLD:AAG2740
Philygria debilis|[6355]|JWDCA919-10|BOLD:AAG2740
Philygria debilis|[6356]|JWDCA922-10|BOLD:AAG2740
Ephydridae|[6357]|JWCHL295-10|BOLD:AAV4070
Sepsis flavimana|[6358]|JWDCA621-10|BOLD:AAJ7591
Sepsis flavimana|[6359]|JWDCJ967-11|BOLD:AAJ7591
Meromyza sp. TAW1|[6360]|JWCHL351-10|BOLD:AAV4318
Hydrellia notata|[6361]|JWDCE446-10|BOLD:AAG9656
Hydrellia notata|[6362]|JWDCE543-10|BOLD:AAG9656
Hydrellia notata|[6363]|JWDCE545-10|BOLD:AAG9656
Ephydridae|[6364]|MHFLI374-07|BOLD:AAG2727
Ephydridae|[6365]|MHFLI495-07|BOLD:AAG2727
Ephydridae|[6366]|MHFLI435-07|BOLD:AAG2727
Ephydridae|[6367]|FLIMH007-08|BOLD:AAG2727
Ephydridae|[6368]|JWDCH535-10|BOLD:AAP8988
Hydrellia|[6369]|JWDCH460-10|BOLD:ADE5061

Ephydridae|[illegible]
Ephydridae|[6367]|FLIMH007-08|BOLD:AAG2727
Ephydridae|[6368]|JWDCH535-10|BOLD:AAP8988
Hydrellia|[6369]|JWDCH460-10|BOLD:ADE5061
Hydrellia|[6370]|FLIMH004-08|BOLD:AAG2732
Ephydridae|[6371]|MHFLI780-07|BOLD:AAG2730
Paramyia silvestris|[6372]|JWDCB355-10|BOLD:ABX4879
Paramyia silvestris|[6373]|JWDCA747-10|BOLD:ABX4879
Paramyia silvestris|[6374]|JWDCJ618-11|BOLD:ABX4879
Leptometopa latipes|[6375]|JWDCG333-10|BOLD:AAP8985
Leptometopa latipes|[6376]|JWDCJ1101-11|BOLD:AAP8985
Leptometopa latipes|[6377]|JWDCJ1082-11|BOLD:AAP8985
Leptometopa latipes|[6378]|JWDCJ1102-11|BOLD:AAP8985
Leptometopa latipes|[6379]|JWDCJ1103-11|BOLD:AAP8985
Leptometopa latipes|[6380]|JWDCL492-11|BOLD:AAP8985
Leptometopa latipes|[6381]|JWDCG330-10|BOLD:AAP8985
Leptometopa latipes|[6382]|JWDCG331-10|BOLD:AAP8985
Leptometopa latipes|[6383]|JWDCG394-10|BOLD:AAP8985
Discocerina obscurella|[6384]|JWDCG160-10|BOLD:AAG2739
Campiglossa|[6385]|JWDCH662-10|BOLD:AAI6491
Campiglossa|[6386]|JWDCH661-10|BOLD:AAI6491
Campiglossa|[6387]|JWDCC041-10|BOLD:AAI6491
Campiglossa|[6388]|JWDCH660-10|BOLD:AAI6491
Campiglossa|[6389]|JWDCE947-10|BOLD:AAI6491
Campiglossa|[6390]|JWDCE948-10|BOLD:AAI6491
Campiglossa|[6391]|JWDCE949-10|BOLD:AAI6491
Campiglossa|[6392]|JWDCE950-10|BOLD:AAI6491
Chamaemyia sp. TAW1|[6393]|JWDCL640-11|BOLD:AAG9063
Cephalops pallidivittipes|[6394]|TWDIP276-09|BOLD:AAG1658
Cephalops grootaerti|[6395]|JWDCJ1242-11|BOLD:AAI3760
Cephalops grootaerti|[6396]|JWPIP093-10|BOLD:AAI3760
Semicephalops varius|[6397]|JWPIP088-10|BOLD:AEE1018
Tomosvaryella sp. MB1|[6398]|JWDCJ1118-11|BOLD:AAU5983
Tomosvaryella sp. AK4|[6399]|JWPIP085-10|BOLD:AAO3574
Tomosvaryella sp. AK4|[6400]|JWDCJ1193-11|BOLD:AAO3574
Tomosvaryella sp. AK4|[6401]|JWPIP060-10|BOLD:AAO3574
Tomosvaryella sp. AK4|[6402]|JWPIP067-10|BOLD:AAO3574
Tomosvaryella sp. AK4|[6403]|JWPIP084-10|BOLD:AAO3574
Tomosvaryella sp. AK4|[6404]|JWPIP091-10|BOLD:AAO3574
Tomosvaryella sp. AK4|[6405]|JWDCH649-10|BOLD:AAO3574
Tomosvaryella sylvatica|[6406]|TWDIP278-09|BOLD:AAG1660
Tomosvaryella sylvatica|[6407]|JWPIP055-10|BOLD:AAG1660
Tomosvaryella sylvatica|[6408]|JWPIP056-10|BOLD:AAG1660
Tomosvaryella sylvatica|[6409]|JWPIP050-10|BOLD:AAG1660
Tomosvaryella sp. ON16|[6410]|JWDCI258-10|BOLD:AAG1659
Tomosvaryella sp. ON16|[6411]|JWPIP014-10|BOLD:AAG1659
Tomosvaryella sp. ON16|[6412]|JWPIP015-10|BOLD:AAG1659
Tomosvaryella sp. ON16|[6413]|JWPIP009-10|BOLD:AAG1659
Tomosvaryella sp. ON16|[6414]|JWPIP034-10|BOLD:AAG1659
Tomosvaryella sp. ON16|[6415]|JWPIP036-10|BOLD:AAG1659
Tomosvaryella sp. ON16|[6416]|JWPIP025-10|BOLD:AAG1659
Tomosvaryella sp. ON16|[6417]|JWPIP019-10|BOLD:AAG1659
Tomosvaryella sp. ON16|[6418]|JWPIP013-10|BOLD:AAG1659
Tomosvaryella sp. ON16|[6419]|JWPIP012-10|BOLD:AAG1659
Tomosvaryella sp. ON16|[6420]|JWPIP011-10|BOLD:AAG1659
Tomosvaryella sp. ON16|[6421]|JWPIP010-10|BOLD:AAG1659
Tomosvaryella sp. ON16|[6422]|JWPIP008-10|BOLD:AAG1659
Tomosvaryella sp. ON16|[6423]|JWPIP007-10|BOLD:AAG1659
Tomosvaryella sp. ON16|[6424]|JWPIP006-10|BOLD:AAG1659
Tomosvaryella sp. ON16|[6425]|JWDCC726-10|BOLD:AAG1659
Tomosvaryella sp. ON16|[6426]|JWPIP027-10|BOLD:AAG1659
Tomosvaryella sp. ON16|[6427]|JWPIP045-10|BOLD:AAG1659
Tomosvaryella sp. ON16|[6428]|JWPIP044-10|BOLD:AAG1659
Tomosvaryella sp. ON16|[6429]|JWPIP046-10|BOLD:AAG1659
Tomosvaryella sp. ON16|[6430]|JWPIP047-10|BOLD:AAG1659
Tomosvaryella sp. ON16|[6431]|JWPIP048-10|BOLD:AAG1659
Tomosvaryella sp. ON16|[6432]|JWPIP049-10|BOLD:AAG1659
Tomosvaryella sp. ON16|[6433]|JWPIP005-10|BOLD:AAG1659
Tomosvaryella sp. ON16|[6434]|JWDCG127-10|BOLD:AAG1659
Tomosvaryella sp. ON16|[6435]|JWDCG130-10|BOLD:AAG1659
Tomosvaryella sp. ON16|[6436]|JWPIP033-10|BOLD:AAG1659
Tomosvaryella sp. ON16|[6437]|JWPIP068-10|BOLD:AAG1659
Tomosvaryella sp. ON16|[6438]|TWDIP277-09|BOLD:AAG1659
Tomosvaryella sp. ON16|[6439]|JWPIP037-10|BOLD:AAG1659
Tomosvaryella sp. ON16|[6440]|JWPIP016-10|BOLD:AAG1659
Tomosvaryella sp. ON16|[6441]|JWPIP035-10|BOLD:AAG1659
Tomosvaryella sp. ON16|[6442]|JWPIP039-10|BOLD:AAG1659
Tomosvaryella sp. ON16|[6443]|JWPIP021-10|BOLD:AAG1659
Tomosvaryella sp. ON16|[6444]|JWPIP023-10|BOLD:AAG1659
Tomosvaryella sp. ON16|[6445]|JWPIP022-10|BOLD:AAG1659
Tomosvaryella sp. ON16|[6446]|JWPIP020-10|BOLD:AAG1659
Tomosvaryella sp. ON16|[6447]|JWPIP017-10|BOLD:AAG1659
Tomosvaryella sp. ON16|[6448]|JWDCJ1025-11|BOLD:AAG1659
Tomosvaryella sp. ON16|[6449]|JWPIP024-10|BOLD:AAG1659
Tomosvaryella sp. ON16|[6450]|JWDCG129-10|BOLD:AAG1659
Tomosvaryella sp. ON16|[6451]|JWDCG128-10|BOLD:AAG1659
Pipunculus hertzogi|[6452]|JWPIP028-10|BOLD:AAE4793
Pipunculus hertzogi|[6453]|JWPIP029-10|BOLD:AAE4793
Pipunculus hertzogi|[6454]|JWPIP052-10|BOLD:AAE4793
Pipunculus hertzogi|[6455]|JWPIP041-10|BOLD:AAE4793
Pipunculus hertzogi|[6456]|JWPIP082-10|BOLD:AAE4793
Pipunculus hertzogi|[6457]|JWPIP065-10|BOLD:AAE4793
Pipunculus hertzogi|[6458]|JWPIP038-10|BOLD:AAE4793
Pipunculus hertzogi|[6459]|JWPIP086-10|BOLD:AAE4793
Pipunculus hertzogi|[6460]|JWPIP087-10|BOLD:AAE4793
Pipunculus hertzogi|[6461]|JWPIP062-10|BOLD:AAE4793
Pipunculus hertzogi|[6462]|JWPIP043-10|BOLD:AAE4793
Pipunculus hertzogi|[6463]|JWPIP030-10|BOLD:AAE4793
Pipunculus hertzogi|[6464]|JWPIP071-10|BOLD:AAE4793
Pipunculus hertzogi|[6465]|JWPIP032-10|BOLD:AAE4793
Pipunculus hertzogi|[6466]|JWPIP040-10|BOLD:AAE4793
Pipunculus hertzogi|[6467]|JWPIP061-10|BOLD:AAE4793
Dorylomorpha sp. ABY5401|[6468]|JWDCJ232-11|BOLD:ABY5401

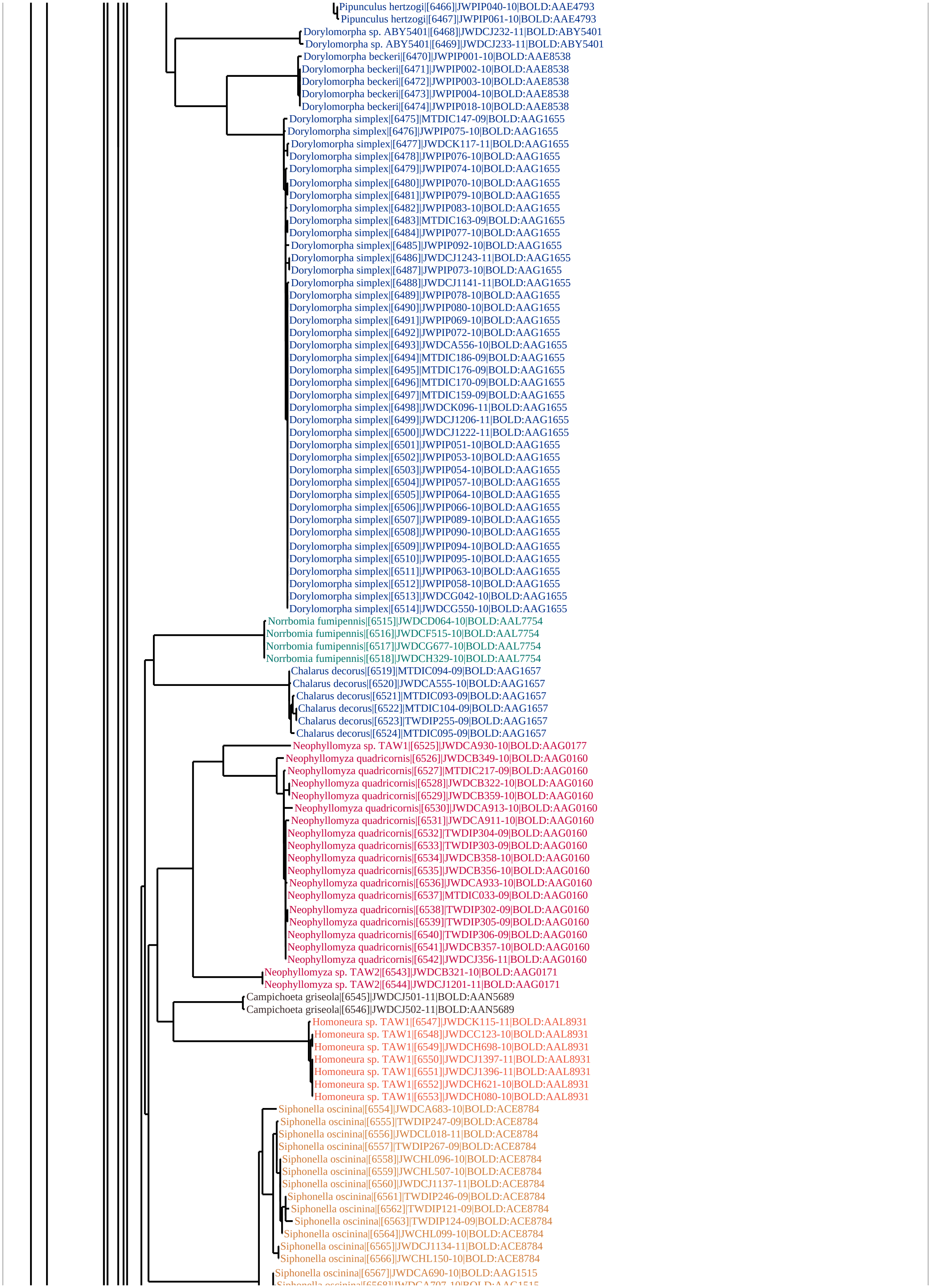
Pipunculus hertzogi|[6466]|JWPIP040-10|BOLD:AAE4793
Pipunculus hertzogi|[6467]|JWPIP061-10|BOLD:AAE4793
Dorylomorpha sp. ABY5401|[6468]|JWDCJ232-11|BOLD:ABY5401
Dorylomorpha sp. ABY5401|[6469]|JWDCJ233-11|BOLD:ABY5401
Dorylomorpha beckeri|[6470]|JWPIP001-10|BOLD:AAE8538
Dorylomorpha beckeri|[6471]|JWPIP002-10|BOLD:AAE8538
Dorylomorpha beckeri|[6472]|JWPIP003-10|BOLD:AAE8538
Dorylomorpha beckeri|[6473]|JWPIP004-10|BOLD:AAE8538
Dorylomorpha beckeri|[6474]|JWPIP018-10|BOLD:AAE8538
Dorylomorpha simplex|[6475]|MTDIC147-09|BOLD:AAG1655
Dorylomorpha simplex|[6476]|JWPIP075-10|BOLD:AAG1655
Dorylomorpha simplex|[6477]|JWDCK117-11|BOLD:AAG1655
Dorylomorpha simplex|[6478]|JWPIP076-10|BOLD:AAG1655
Dorylomorpha simplex|[6479]|JWPIP074-10|BOLD:AAG1655
Dorylomorpha simplex|[6480]|JWPIP070-10|BOLD:AAG1655
Dorylomorpha simplex|[6481]|JWPIP079-10|BOLD:AAG1655
Dorylomorpha simplex|[6482]|JWPIP083-10|BOLD:AAG1655
Dorylomorpha simplex|[6483]|MTDIC163-09|BOLD:AAG1655
Dorylomorpha simplex|[6484]|JWPIP077-10|BOLD:AAG1655
Dorylomorpha simplex|[6485]|JWPIP092-10|BOLD:AAG1655
Dorylomorpha simplex|[6486]|JWDCJ1243-11|BOLD:AAG1655
Dorylomorpha simplex|[6487]|JWPIP073-10|BOLD:AAG1655
Dorylomorpha simplex|[6488]|JWDCJ1141-11|BOLD:AAG1655
Dorylomorpha simplex|[6489]|JWPIP078-10|BOLD:AAG1655
Dorylomorpha simplex|[6490]|JWPIP080-10|BOLD:AAG1655
Dorylomorpha simplex|[6491]|JWPIP069-10|BOLD:AAG1655
Dorylomorpha simplex|[6492]|JWPIP072-10|BOLD:AAG1655
Dorylomorpha simplex|[6493]|JWDCA556-10|BOLD:AAG1655
Dorylomorpha simplex|[6494]|MTDIC186-09|BOLD:AAG1655
Dorylomorpha simplex|[6495]|MTDIC176-09|BOLD:AAG1655
Dorylomorpha simplex|[6496]|MTDIC170-09|BOLD:AAG1655
Dorylomorpha simplex|[6497]|MTDIC159-09|BOLD:AAG1655
Dorylomorpha simplex|[6498]|JWDCK096-11|BOLD:AAG1655
Dorylomorpha simplex|[6499]|JWDCJ1206-11|BOLD:AAG1655
Dorylomorpha simplex|[6500]|JWDCJ1222-11|BOLD:AAG1655
Dorylomorpha simplex|[6501]|JWPIP051-10|BOLD:AAG1655
Dorylomorpha simplex|[6502]|JWPIP053-10|BOLD:AAG1655
Dorylomorpha simplex|[6503]|JWPIP054-10|BOLD:AAG1655
Dorylomorpha simplex|[6504]|JWPIP057-10|BOLD:AAG1655
Dorylomorpha simplex|[6505]|JWPIP064-10|BOLD:AAG1655
Dorylomorpha simplex|[6506]|JWPIP066-10|BOLD:AAG1655
Dorylomorpha simplex|[6507]|JWPIP089-10|BOLD:AAG1655
Dorylomorpha simplex|[6508]|JWPIP090-10|BOLD:AAG1655
Dorylomorpha simplex|[6509]|JWPIP094-10|BOLD:AAG1655
Dorylomorpha simplex|[6510]|JWPIP095-10|BOLD:AAG1655
Dorylomorpha simplex|[6511]|JWPIP063-10|BOLD:AAG1655
Dorylomorpha simplex|[6512]|JWPIP058-10|BOLD:AAG1655
Dorylomorpha simplex|[6513]|JWDCG042-10|BOLD:AAG1655
Dorylomorpha simplex|[6514]|JWDCG550-10|BOLD:AAG1655
Norrbomia fumipennis|[6515]|JWDCD064-10|BOLD:AAL7754
Norrbomia fumipennis|[6516]|JWDCF515-10|BOLD:AAL7754
Norrbomia fumipennis|[6517]|JWDCG677-10|BOLD:AAL7754
Norrbomia fumipennis|[6518]|JWDCH329-10|BOLD:AAL7754
Chalarus decorus|[6519]|MTDIC094-09|BOLD:AAG1657
Chalarus decorus|[6520]|JWDCA555-10|BOLD:AAG1657
Chalarus decorus|[6521]|MTDIC093-09|BOLD:AAG1657
Chalarus decorus|[6522]|MTDIC104-09|BOLD:AAG1657
Chalarus decorus|[6523]|TWDIP255-09|BOLD:AAG1657
Chalarus decorus|[6524]|MTDIC095-09|BOLD:AAG1657
Neophyllomyza sp. TAW1|[6525]|JWDCA930-10|BOLD:AAG0177
Neophyllomyza quadricornis|[6526]|JWDCB349-10|BOLD:AAG0160
Neophyllomyza quadricornis|[6527]|MTDIC217-09|BOLD:AAG0160
Neophyllomyza quadricornis|[6528]|JWDCB322-10|BOLD:AAG0160
Neophyllomyza quadricornis|[6529]|JWDCB359-10|BOLD:AAG0160
Neophyllomyza quadricornis|[6530]|JWDCA913-10|BOLD:AAG0160
Neophyllomyza quadricornis|[6531]|JWDCA911-10|BOLD:AAG0160
Neophyllomyza quadricornis|[6532]|TWDIP304-09|BOLD:AAG0160
Neophyllomyza quadricornis|[6533]|TWDIP303-09|BOLD:AAG0160
Neophyllomyza quadricornis|[6534]|JWDCB358-10|BOLD:AAG0160
Neophyllomyza quadricornis|[6535]|JWDCB356-10|BOLD:AAG0160
Neophyllomyza quadricornis|[6536]|JWDCA933-10|BOLD:AAG0160
Neophyllomyza quadricornis|[6537]|MTDIC033-09|BOLD:AAG0160
Neophyllomyza quadricornis|[6538]|TWDIP302-09|BOLD:AAG0160
Neophyllomyza quadricornis|[6539]|TWDIP305-09|BOLD:AAG0160
Neophyllomyza quadricornis|[6540]|TWDIP306-09|BOLD:AAG0160
Neophyllomyza quadricornis|[6541]|JWDCB357-10|BOLD:AAG0160
Neophyllomyza quadricornis|[6542]|JWDCJ356-11|BOLD:AAG0160
Neophyllomyza sp. TAW2|[6543]|JWDCB321-10|BOLD:AAG0171
Neophyllomyza sp. TAW2|[6544]|JWDCJ1201-11|BOLD:AAG0171
Campichoeta griseola|[6545]|JWDCJ501-11|BOLD:AAN5689
Campichoeta griseola|[6546]|JWDCJ502-11|BOLD:AAN5689
Homoneura sp. TAW1|[6547]|JWDCK115-11|BOLD:AAL8931
Homoneura sp. TAW1|[6548]|JWDCC123-10|BOLD:AAL8931
Homoneura sp. TAW1|[6549]|JWDCH698-10|BOLD:AAL8931
Homoneura sp. TAW1|[6550]|JWDCJ1397-11|BOLD:AAL8931
Homoneura sp. TAW1|[6551]|JWDCJ1396-11|BOLD:AAL8931
Homoneura sp. TAW1|[6552]|JWDCH621-10|BOLD:AAL8931
Homoneura sp. TAW1|[6553]|JWDCH080-10|BOLD:AAL8931
Siphonella oscinina|[6554]|JWDCA683-10|BOLD:ACE8784
Siphonella oscinina|[6555]|TWDIP247-09|BOLD:ACE8784
Siphonella oscinina|[6556]|JWDCL018-11|BOLD:ACE8784
Siphonella oscinina|[6557]|TWDIP267-09|BOLD:ACE8784
Siphonella oscinina|[6558]|JWCHL096-10|BOLD:ACE8784
Siphonella oscinina|[6559]|JWCHL507-10|BOLD:ACE8784
Siphonella oscinina|[6560]|JWDCJ1137-11|BOLD:ACE8784
Siphonella oscinina|[6561]|TWDIP246-09|BOLD:ACE8784
Siphonella oscinina|[6562]|TWDIP121-09|BOLD:ACE8784
Siphonella oscinina|[6563]|TWDIP124-09|BOLD:ACE8784
Siphonella oscinina|[6564]|JWCHL099-10|BOLD:ACE8784
Siphonella oscinina|[6565]|JWDCJ1134-11|BOLD:ACE8784
Siphonella oscinina|[6566]|JWCHL150-10|BOLD:ACE8784
Siphonella oscinina|[6567]|JWDCA690-10|BOLD:AAG1515

[illegible]
Siphonella oscinina|[6566]|JWCHL150-10|BOLD:ACE8784
Siphonella oscinina|[6567]|JWDCA690-10|BOLD:AAG1515
Siphonella oscinina|[6568]|JWDCA707-10|BOLD:AAG1515
Siphonella oscinina|[6569]|TWDIP125-09|BOLD:AAG1515
Siphonella oscinina|[6570]|MTDIC189-09|BOLD:AAG1515
Siphonella oscinina|[6571]|TWDIP123-09|BOLD:AAG1515
Siphonella oscinina|[6572]|JWDCB204-10|BOLD:AAG1515
Siphonella oscinina|[6573]|JWDCJ398-11|BOLD:AAG1515
Siphonella oscinina|[6574]|JWDCJ1213-11|BOLD:AAG1515
Siphonella oscinina|[6575]|JWCHL032-10|BOLD:AAG1515
Siphonella oscinina|[6576]|JWCHL436-10|BOLD:AAG1515
Siphonella oscinina|[6577]|JWCHL509-10|BOLD:AAG1515
Siphonella oscinina|[6578]|JWCHL164-10|BOLD:AAG1515
Siphonella oscinina|[6579]|JWCHL394-10|BOLD:AAG1515
Siphonella oscinina|[6580]|JWDCA706-10|BOLD:AAG1515
Siphonella oscinina|[6581]|JWDCA686-10|BOLD:AAG1515
Siphonella oscinina|[6582]|JWCHL095-10|BOLD:AAG1515
Siphonella oscinina|[6583]|JWCHL313-10|BOLD:AAG1515
Siphonella oscinina|[6584]|JWCHL167-10|BOLD:AAG1515
Siphonella oscinina|[6585]|JWCHL165-10|BOLD:AAG1515
Siphonella oscinina|[6586]|JWCHL147-10|BOLD:AAG1515
Siphonella oscinina|[6587]|JWCHL097-10|BOLD:AAG1515
Siphonella oscinina|[6588]|JWDCH811-10|BOLD:AAG1515
Siphonella oscinina|[6589]|JWDCJ1136-11|BOLD:AAG1515
Siphonella oscinina|[6590]|JWDCJ1128-11|BOLD:AAG1515
Siphonella oscinina|[6591]|JWDCJ357-11|BOLD:AAG1515
Siphonella oscinina|[6592]|JWDCB223-10|BOLD:AAG1515
Siphonella oscinina|[6593]|JWDCA681-10|BOLD:AAG1515
Siphonella oscinina|[6594]|TWDIP268-09|BOLD:AAG1515
Siphonella oscinina|[6595]|TWDIP249-09|BOLD:AAG1515
Siphonella oscinina|[6596]|JWDCA709-10|BOLD:AAG1515
Siphonella oscinina|[6597]|TWDIP245-09|BOLD:AAG1515
Siphonella oscinina|[6598]|JWCHL395-10|BOLD:AAG1515
Siphonella oscinina|[6599]|JWCHL434-10|BOLD:AAG1515
Chlorops sp. TAW1|[6600]|JWCHL338-10|BOLD:AAH4154
Chlorops sp. TAW2|[6601]|JWCHL256-10|BOLD:AAP2737
Chlorops sp. TAW2|[6602]|JWCHL411-10|BOLD:AAP2737
Chlorops sp. TAW2|[6603]|JWCHL413-10|BOLD:AAP2737
Chlorops sp. TAW2|[6604]|JWDCI931-11|BOLD:AAP2737
Chlorops sp. TAW2|[6605]|JWCHL414-10|BOLD:AAP2737
Epichlorops puncticollis|[6606]|MHFLI100-06|BOLD:AAI5374
Epichlorops puncticollis|[6607]|MHFLI099-06|BOLD:AAI5374
Epichlorops puncticollis|[6608]|MHFLI098-06|BOLD:AAI5374
Epichlorops puncticollis|[6609]|JWCHL020-10|BOLD:AAI5374
Epichlorops puncticollis|[6610]|JWCHL064-10|BOLD:AAI5374
Epichlorops puncticollis|[6611]|JWCHL065-10|BOLD:AAI5374
Epichlorops puncticollis|[6612]|JWCHL076-10|BOLD:AAI5374
Epichlorops puncticollis|[6613]|JWDCL024-11|BOLD:AAI5374
Epichlorops puncticollis|[6614]|JWCHL011-10|BOLD:AAI5374
Epichlorops puncticollis|[6615]|JWCHL105-10|BOLD:AAI5374
Epichlorops puncticollis|[6616]|JWCHL075-10|BOLD:AAI5374
Epichlorops puncticollis|[6617]|MHFLI097-06|BOLD:AAI5374
Epichlorops puncticollis|[6618]|JWCHL106-10|BOLD:AAI5374
Epichlorops puncticollis|[6619]|JWCHL063-10|BOLD:AAI5374
Epichlorops puncticollis|[6620]|JWCHL107-10|BOLD:AAI5374
Epichlorops puncticollis|[6621]|JWCHL410-10|BOLD:AAI5374
Epichlorops puncticollis|[6622]|MHFLI096-06|BOLD:AAI5374
Epichlorops puncticollis|[6623]|MHFLI095-06|BOLD:AAI5374
Epichlorops puncticollis|[6624]|JWCHL044-10|BOLD:AAI5374
Epichlorops puncticollis|[6625]|JWCHL043-10|BOLD:AAI5374
Epichlorops puncticollis|[6626]|JWCHL042-10|BOLD:AAI5374
Epichlorops puncticollis|[6627]|JWCHL019-10|BOLD:AAI5374
Epichlorops puncticollis|[6628]|JWCHL018-10|BOLD:AAI5374
Epichlorops puncticollis|[6629]|JWCHL017-10|BOLD:AAI5374
Epichlorops puncticollis|[6630]|JWCHL014-10|BOLD:AAI5374
Epichlorops puncticollis|[6631]|JWCHL013-10|BOLD:AAI5374
Epichlorops puncticollis|[6632]|JWCHL012-10|BOLD:AAI5374
Epichlorops puncticollis|[6633]|JWCHL010-10|BOLD:AAI5374
Epichlorops puncticollis|[6634]|JWCHL116-10|BOLD:AAI5374
Epichlorops puncticollis|[6635]|JWDCL475-11|BOLD:AAI5374
Epichlorops puncticollis|[6636]|JWDCL169-11|BOLD:AAI5374
Epichlorops puncticollis|[6637]|JWDCL585-11|BOLD:AAI5374
Epichlorops puncticollis|[6638]|JWDCL023-11|BOLD:AAI5374
Epichlorops puncticollis|[6639]|JWDCK041-11|BOLD:AAI5374
Epichlorops puncticollis|[6640]|JWCHL041-10|BOLD:AAI5374
Epichlorops puncticollis|[6641]|JWCHL046-10|BOLD:AAI5374
Epichlorops puncticollis|[6642]|JWCHL062-10|BOLD:AAI5374
Epichlorops puncticollis|[6643]|JWCHL066-10|BOLD:AAI5374
Epichlorops puncticollis|[6644]|JWCHL074-10|BOLD:AAI5374
Epichlorops puncticollis|[6645]|JWCHL079-10|BOLD:AAI5374
Epichlorops puncticollis|[6646]|JWCHL103-10|BOLD:AAI5374
Epichlorops puncticollis|[6647]|JWCHL104-10|BOLD:AAI5374
Epichlorops puncticollis|[6648]|JWCHL114-10|BOLD:AAI5374
Epichlorops puncticollis|[6649]|JWCHL115-10|BOLD:AAI5374
Epichlorops puncticollis|[6650]|JWCHL117-10|BOLD:AAI5374
Epichlorops puncticollis|[6651]|JWCHL118-10|BOLD:AAI5374
Epichlorops puncticollis|[6652]|JWCHL132-10|BOLD:AAI5374
Epichlorops puncticollis|[6653]|JWCHL140-10|BOLD:AAI5374
Epichlorops puncticollis|[6654]|JWCHL153-10|BOLD:AAI5374
Epichlorops puncticollis|[6655]|JWCHL160-10|BOLD:AAI5374
Epichlorops puncticollis|[6656]|JWCHL161-10|BOLD:AAI5374
Epichlorops puncticollis|[6657]|JWCHL387-10|BOLD:AAI5374
Epichlorops puncticollis|[6658]|JWCHL408-10|BOLD:AAI5374
Epichlorops puncticollis|[6659]|JWCHL409-10|BOLD:AAI5374
Epichlorops puncticollis|[6660]|JWCHL499-10|BOLD:AAI5374
Chlorops subniger|[6661]|JWCHL471-10|BOLD:AAP3664
Chlorops subniger|[6662]|JWCHL133-10|BOLD:AAP3664
Chlorops subniger|[6663]|JWCHL479-10|BOLD:AAP3664
Epichlorops scaber|[6664]|JWCHL506-10|BOLD:AAH4206
Epichlorops scaber|[6665]|JWCHL505-10|BOLD:AAH4206
Epichlorops scaber|[6666]|JWCHL470-10|BOLD:AAH4206
Epichlorops scaber|[6667]|JWCHL467-10|BOLD:AAH4206

Epichlorops scaber|[6665]|JWCHL505-10|BOLD:AAH4206
Epichlorops scaber|[6666]|JWCHL470-10|BOLD:AAH4206
Epichlorops scaber|[6667]|JWCHL467-10|BOLD:AAH4206
Epichlorops scaber|[6668]|JWCHL458-10|BOLD:AAH4206
Epichlorops scaber|[6669]|JWCHL341-10|BOLD:AAH4206
Epichlorops scaber|[6670]|JWCHL170-10|BOLD:AAH4206
Epichlorops scaber|[6671]|JWCHL050-10|BOLD:AAH4206
Epichlorops scaber|[6672]|JWCHL047-10|BOLD:AAH4206
Epichlorops puncticollis|[6673]|MTDIC012-09|BOLD:AAH4206
Epichlorops scaber|[6674]|JWCHL511-10|BOLD:AAH4206
Chlorops sp. TAW5|[6675]|JWDCK040-11|BOLD:AAP5941
Chlorops sp. TAW5|[6676]|JWCHL081-10|BOLD:AAP5941
Chlorops sp. TAW3|[6677]|JWCHL514-10|BOLD:AAQ0882
Chlorops sp. TAW4|[6678]|JWCHL152-10|BOLD:AAH4137
Chlorops sp. TAW4|[6679]|JWCHL162-10|BOLD:AAH4137
Chlorops sp. TAW4|[6680]|JWCHL468-10|BOLD:AAH4137
Chlorops sp. TAW4|[6681]|JWCHL396-10|BOLD:AAH4137
Chlorops sp. TAW4|[6682]|JWCHL144-10|BOLD:AAH4137
Chlorops sp. TAW4|[6683]|JWCHL001-10|BOLD:AAH4137
Chlorops sp. TAW4|[6684]|JWCHL397-10|BOLD:AAH4137
Chlorops sp. TAW4|[6685]|JWCHL431-10|BOLD:AAH4137
Chlorops sp. TAW4|[6686]|JWCHL469-10|BOLD:AAH4137
Chlorops sp. TAW4|[6687]|JWCHL516-10|BOLD:AAH4137
Chlorops sp. TAW4|[6688]|JWCHL504-10|BOLD:AAH4137
Chlorops sp. TAW4|[6689]|JWCHL481-10|BOLD:AAH4137
Chlorops sp. TAW4|[6690]|JWCHL500-10|BOLD:AAH4137
Chlorops sp. TAW4|[6691]|JWCHL021-10|BOLD:AAH4137
Chlorops sp. TAW4|[6692]|JWCHL049-10|BOLD:AAH4137
Chlorops sp. TAW4|[6693]|JWCHL077-10|BOLD:AAH4137
Chlorops sp. TAW4|[6694]|JWCHL078-10|BOLD:AAH4137
Chlorops sp. TAW4|[6695]|JWCHL080-10|BOLD:AAH4137
Chlorops sp. TAW4|[6696]|JWCHL130-10|BOLD:AAH4137
Chlorops sp. TAW4|[6697]|JWCHL131-10|BOLD:AAH4137
Chlorops sp. TAW4|[6698]|JWCHL142-10|BOLD:AAH4137
Chlorops sp. TAW4|[6699]|JWCHL145-10|BOLD:AAH4137
Chlorops sp. TAW4|[6700]|JWCHL155-10|BOLD:AAH4137
Chlorops sp. TAW4|[6701]|JWCHL156-10|BOLD:AAH4137
Chlorops sp. TAW4|[6702]|JWCHL163-10|BOLD:AAH4137
Chlorops sp. TAW4|[6703]|JWCHL421-10|BOLD:AAH4137
Chlorops sp. TAW4|[6704]|JWCHL484-10|BOLD:AAH4137
Chlorops sp. TAW4|[6705]|JWCHL522-10|BOLD:AAH4137
Notiphila atripes|[6706]|JWDCJ176-11|BOLD:AAP6351
Notiphila atripes|[6707]|JWDCL185-11|BOLD:AAP6351
Notiphila atripes|[6708]|JWDOL164-10|BOLD:AAP6351
Notiphila atripes|[6709]|JWDCJ787-11|BOLD:AAP6351
Notiphila atripes|[6710]|JWDCJ786-11|BOLD:AAP6351
Notiphila atripes|[6711]|JWDCK050-11|BOLD:AAP6351
Notiphila atripes|[6712]|JWDCL661-11|BOLD:AAP6351
Notiphila atripes|[6713]|JWDCL660-11|BOLD:AAP6351
Notiphila atripes|[6714]|JWDCL659-11|BOLD:AAP6351
Notiphila atripes|[6715]|JWDCL658-11|BOLD:AAP6351
Notiphila atripes|[6716]|JWDCJ178-11|BOLD:AAP6351
Notiphila atripes|[6717]|JWDCJ238-11|BOLD:AAP6351
Notiphila atripes|[6718]|JWDCH900-10|BOLD:AAP6351
Notiphila atripes|[6719]|JWDCI576-10|BOLD:AAP6351
Notiphila atripes|[6720]|JWDCI744-11|BOLD:AAP6351
Notiphila atripes|[6721]|JWDCI745-11|BOLD:AAP6351
Notiphila atripes|[6722]|JWDCF279-10|BOLD:AAP6351
Notiphila atripes|[6723]|JWDCF287-10|BOLD:AAP6351
Speccafrons mallochi|[6724]|JWDCC879-10|BOLD:ABZ9888
Speccafrons mallochi|[6725]|JWCHL143-10|BOLD:ABZ9888
Thaumatomyia pulla|[6726]|JWDCA743-10|BOLD:AAG9241
Thaumatomyia pulla|[6727]|JWDCB199-10|BOLD:AAG9241
Thaumatomyia pulla|[6728]|JWCHL435-10|BOLD:AAG9241
Thaumatomyia pulla|[6729]|JWDCA691-10|BOLD:ABZ3900
Thaumatomyia pulla|[6730]|JWDCA698-10|BOLD:ABZ3900
Thaumatomyia pulla|[6731]|JWDCB222-10|BOLD:ABZ3900
Thaumatomyia pulla|[6732]|JWCHL068-10|BOLD:ABZ3900
Thaumatomyia pulla|[6733]|JWCHL269-10|BOLD:AAG9240
Thaumatomyia pulla|[6734]|JWCHL271-10|BOLD:AAG9240
Thaumatomyia pulla|[6735]|JWCHL344-10|BOLD:AAG9240
Thaumatomyia pulla|[6736]|JWCHL416-10|BOLD:AAG9240
Thaumatomyia pulla|[6737]|JWDCA732-10|BOLD:AAG9240
Thaumatomyia pulla|[6738]|JWDCD061-10|BOLD:AAG9240
Thaumatomyia pulla|[6739]|JWDCA682-10|BOLD:AAG9240
Thaumatomyia pulla|[6740]|JWDCJ1027-11|BOLD:AAG9240
Thaumatomyia pulla|[6741]|JWCHL263-10|BOLD:AAG9240
Thaumatomyia pulla|[6742]|JWCHL264-10|BOLD:AAG9240
Thaumatomyia pulla|[6743]|JWCHL439-10|BOLD:AAG9240
Thaumatomyia pulla|[6744]|JWCHL440-10|BOLD:AAG9240
Thaumatomyia trifasciata|[6745]|JWDCA666-10|BOLD:AAG4705
Thaumatomyia trifasciata|[6746]|JWDCA671-10|BOLD:AAG4705
Thaumatomyia trifasciata|[6747]|JWCHL188-10|BOLD:AAG4705
Thaumatomyia trifasciata|[6748]|JWCHL318-10|BOLD:AAG4705
Thaumatomyia trifasciata|[6749]|JWCHL187-10|BOLD:AAG4705
Thaumatomyia trifasciata|[6750]|JWCHL186-10|BOLD:AAG4705
Thaumatomyia trifasciata|[6751]|JWDCA672-10|BOLD:AAG4705
Thaumatomyia trifasciata|[6752]|JWDCA667-10|BOLD:AAG4705
Thaumatomyia trifasciata|[6753]|JWCHL456-10|BOLD:AAG4705
Thaumatomyia trifasciata|[6754]|JWCHL502-10|BOLD:AAG4705
Thaumatomyia trifasciata|[6755]|TWDIP049-09|BOLD:AAG4705
Thaumatomyia trifasciata|[6756]|JWDCA670-10|BOLD:AAG4705
Thaumatomyia trifasciata|[6757]|TWDIP050-09|BOLD:AAG4705
Thaumatomyia trifasciata|[6758]|JWCHL321-10|BOLD:AAG4705
Thaumatomyia trifasciata|[6759]|MTDIC092-09|BOLD:AAG4705
Thaumatomyia trifasciata|[6760]|JWDCB193-10|BOLD:AAG4705
Thaumatomyia trifasciata|[6761]|MTDIC263-09|BOLD:AAG4705
Thaumatomyia trifasciata|[6762]|TWDIP048-09|BOLD:AAG4705
Thaumatomyia trifasciata|[6763]|JWDCJ1026-11|BOLD:AAG4705
Thaumatomyia trifasciata|[6764]|JWCHL139-10|BOLD:AAG4705
Thaumatomyia trifasciata|[6765]|JWCHL348-10|BOLD:AAG4705
Thaumatomyia trifasciata|[6766]|JWCHL350-10|BOLD:AAG4705
Thaumatomyia trifasciata|[6767]|JWCHL247-10|BOLD:AAG4705

[illegible]
Thaumatomyia trifasciata|[6765]|JWCHL348-10|BOLD:AAG4705
Thaumatomyia trifasciata|[6766]|JWCHL350-10|BOLD:AAG4705
Thaumatomyia trifasciata|[6767]|JWCHL247-10|BOLD:AAG4705
Thaumatomyia trifasciata|[6768]|TWDIP047-09|BOLD:AAG4705
Thaumatomyia trifasciata|[6769]|JWCHL255-10|BOLD:AAG4705
Thaumatomyia trifasciata|[6770]|JWCHL525-10|BOLD:AAG4705
Thaumatomyia trifasciata|[6771]|MTDIC055-09|BOLD:AAG4705
Thaumatomyia trifasciata|[6772]|JWDCA677-10|BOLD:AAG4705
Thaumatomyia trifasciata|[6773]|JWDCA724-10|BOLD:AAG4705
Thaumatomyia trifasciata|[6774]|JWDCA675-10|BOLD:AAG4705
Thaumatomyia trifasciata|[6775]|JWDCA719-10|BOLD:AAG4705
Thaumatomyia trifasciata|[6776]|JWDCA723-10|BOLD:AAG4705
Thaumatomyia trifasciata|[6777]|JWDCA722-10|BOLD:AAG4705
Thaumatomyia trifasciata|[6778]|JWCHL346-10|BOLD:AAG4705
Thaumatomyia trifasciata|[6779]|JWCHL359-10|BOLD:AAG4705
Thaumatomyia trifasciata|[6780]|JWDCA720-10|BOLD:AAG4705
Thaumatomyia trifasciata|[6781]|TWDIP208-09|BOLD:AAG4705
Thaumatomyia trifasciata|[6782]|JWCHL347-10|BOLD:AAG4705
Thaumatomyia trifasciata|[6783]|JWCHL189-10|BOLD:AAG4705
Thaumatomyia trifasciata|[6784]|JWDCA674-10|BOLD:AAG4705
Thaumatomyia trifasciata|[6785]|JWDCA668-10|BOLD:AAG4705
Thaumatomyia trifasciata|[6786]|TWDIP207-09|BOLD:AAG4705
Thaumatomyia trifasciata|[6787]|TWDIP206-09|BOLD:AAG4705
Thaumatomyia trifasciata|[6788]|MTDIC256-09|BOLD:AAG4705
Thaumatomyia trifasciata|[6789]|JWCHL285-10|BOLD:AAG4705
Thaumatomyia trifasciata|[6790]|JWCHL317-10|BOLD:AAG4705
Thaumatomyia trifasciata|[6791]|JWDCJ1130-11|BOLD:AAG4705
Thaumatomyia trifasciata|[6792]|JWDCL523-11|BOLD:AAG4705
Thaumatomyia trifasciata|[6793]|JWCHL190-10|BOLD:AAG4705
Thaumatomyia trifasciata|[6794]|JWCHL349-10|BOLD:AAG4705
Thaumatomyia trifasciata|[6795]|TWDIP046-09|BOLD:AAG4705
Thaumatomyia trifasciata|[6796]|JWDCA673-10|BOLD:AAG4705
Thaumatomyia trifasciata|[6797]|JWDCA676-10|BOLD:AAG4705
Thaumatomyia trifasciata|[6798]|JWDCA680-10|BOLD:AAG4705
Thaumatomyia trifasciata|[6799]|JWCHL455-10|BOLD:AAG4705
Thaumatomyia trifasciata|[6800]|JWCHL534-10|BOLD:AAG4705
Thaumatomyia glabra|[6801]|JWCHL345-10|BOLD:AAH4135
Thaumatomyia glabra|[6802]|JWCHL262-10|BOLD:AAH4135
Thaumatomyia glabra|[6803]|JWCHL365-10|BOLD:AAH4135
Thaumatomyia glabra|[6804]|JWCHL185-10|BOLD:AAH4135
Thaumatomyia glabra|[6805]|JWCHL039-10|BOLD:AAH4135
Thaumatomyia glabra|[6806]|JWCHL037-10|BOLD:AAH4135
Thaumatomyia glabra|[6807]|JWCHL362-10|BOLD:AAH4135
Thaumatomyia glabra|[6808]|JWCHL444-10|BOLD:AAH4135
Thaumatomyia glabra|[6809]|JWCHL036-10|BOLD:AAH4135
Thaumatomyia glabra|[6810]|JWCHL445-10|BOLD:AAH4135
Thaumatomyia glabra|[6811]|JWCHL363-10|BOLD:AAH4135
Thaumatomyia glabra|[6812]|JWCHL443-10|BOLD:AAH4135
Thaumatomyia glabra|[6813]|JWCHL366-10|BOLD:AAH4135
Thaumatomyia glabra|[6814]|JWCHL368-10|BOLD:AAH4135
Thaumatomyia glabra|[6815]|JWCHL441-10|BOLD:AAH4135
Thaumatomyia glabra|[6816]|JWDCL485-11|BOLD:AAH4135
Thaumatomyia glabra|[6817]|JWCHL360-10|BOLD:AAH4135
Thaumatomyia glabra|[6818]|JWCHL038-10|BOLD:AAH4135
Thaumatomyia bistriata|[6819]|JWDCA679-10|BOLD:ABZ1503
Thaumatomyia bistriata|[6820]|JWDCA678-10|BOLD:AAG9238
Thaumatomyia bistriata|[6821]|JWDCJ1097-11|BOLD:AAG9238
Thaumatomyia bistriata|[6822]|JWDCJ1098-11|BOLD:AAG9238
Thaumatomyia bistriata|[6823]|JWDCJ137-11|BOLD:AAG9238
Thaumatomyia bistriata|[6824]|JWCHL040-10|BOLD:AAG9238
Thaumatomyia bistriata|[6825]|JWCHL108-10|BOLD:AAG9238
Thaumatomyia bistriata|[6826]|JWCHL361-10|BOLD:AAG9238
Thaumatomyia bistriata|[6827]|JWCHL364-10|BOLD:AAG9238
Thaumatomyia bistriata|[6828]|JWCHL415-10|BOLD:AAG9238
Thaumatomyia bistriata|[6829]|JWCHL442-10|BOLD:AAG9238
Thaumatomyia bistriata|[6830]|JWCHL533-10|BOLD:AAG9238
Thaumatomyia bistriata|[6831]|JWCHL532-10|BOLD:AAG9238
Thaumatomyia bistriata|[6832]|JWCHL536-10|BOLD:AAG9238
Fiebrigella sp. TAW1|[6833]|JWDCA688-10|BOLD:AAH4212
Fiebrigella sp. TAW1|[6834]|JWDCH800-10|BOLD:AAH4212
Fiebrigella sp. TAW1|[6835]|JWDCH881-10|BOLD:AAH4212
Fiebrigella sp. TAW1|[6836]|JWDCL090-11|BOLD:AAH4212
Fiebrigella sp. TAW1|[6837]|JWDCJ1135-11|BOLD:AAH4212
Fiebrigella sp. TAW1|[6838]|JWDCJ1133-11|BOLD:AAH4212
Fiebrigella sp. TAW1|[6839]|JWDCJ1132-11|BOLD:AAH4212
Fiebrigella sp. TAW1|[6840]|JWDCL091-11|BOLD:AAH4212
Fiebrigella sp. TAW1|[6841]|JWDCH943-10|BOLD:AAH4212
Fiebrigella sp. TAW1|[6842]|JWDCI080-10|BOLD:AAH4212
Fiebrigella sp. TAW1|[6843]|JWDCH274-10|BOLD:AAH4212
Gaurax pallidipes|[6844]|JWDCA693-10|BOLD:AAG1517
Gaurax sp. TAW3|[6845]|MTDIC141-09|BOLD:AAH4165
Gaurax sp. TAW4|[6846]|JWCHL129-10|BOLD:AAP3903
Gaurax sp. TAW2|[6847]|JWDCC554-10|BOLD:AAL7387
Gaurax sp. TAW2|[6848]|JWDCC567-10|BOLD:AAL7387
Gaurax sp. TAW1|[6849]|JWDCA708-10|BOLD:AAG1520
Gaurax sp. TAW1|[6850]|JWDCH570-10|BOLD:AAG1520
Gaurax sp. TAW1|[6851]|JWDCH571-10|BOLD:AAG1520
Gaurax sp. TAW1|[6852]|JWCHL530-10|BOLD:AAG1520
Rhopalopterum soror|[6853]|JWCHL023-10|BOLD:ACZ6867
Rhopalopterum soror|[6854]|JWCHL135-10|BOLD:ACZ6867
Rhopalopterum soror|[6855]|JWDCL474-11|BOLD:ACZ6867
Rhopalopterum soror|[6856]|JWCHL035-10|BOLD:ACZ6867
Rhopalopterum soror|[6857]|JWCHL433-10|BOLD:ACZ6867
Rhopalopterum soror|[6858]|JWCHL400-10|BOLD:ACZ6867
Rhopalopterum soror|[6859]|JWCHL459-10|BOLD:ACZ6867
Rhopalopterum soror|[6860]|JWCHL254-10|BOLD:ACZ6867
Rhopalopterum soror|[6861]|JWCHL026-10|BOLD:ACZ6867
Rhopalopterum soror|[6862]|JWCHL022-10|BOLD:ACZ6867
Rhopalopterum soror|[6863]|JWDCJ1447-11|BOLD:ACZ6867
Rhopalopterum soror|[6864]|JWCHL475-10|BOLD:ACZ6867
Rhopalopterum soror|[6865]|JWCHL029-10|BOLD:ACZ6867
Rhopalopterum soror|[6866]|JWCHL052-10|BOLD:ACZ6867

Rhopalopterum soror|[6864]|JWCHL475-10|BOLD:ACZ6867
Rhopalopterum soror|[6865]|JWCHL029-10|BOLD:ACZ6867
Rhopalopterum soror|[6866]|JWCHL052-10|BOLD:ACZ6867
Rhopalopterum soror|[6867]|JWCHL059-10|BOLD:ACZ6867
Rhopalopterum soror|[6868]|JWCHL405-10|BOLD:ACZ6867
Rhopalopterum soror|[6869]|JWCHL399-10|BOLD:ACZ6867
Rhopalopterum soror|[6870]|JWCHL388-10|BOLD:ACZ6867
Rhopalopterum soror|[6871]|JWCHL384-10|BOLD:ACZ6867
Rhopalopterum soror|[6872]|JWCHL378-10|BOLD:ACZ6867
Rhopalopterum soror|[6873]|JWCHL373-10|BOLD:ACZ6867
Rhopalopterum soror|[6874]|JWCHL372-10|BOLD:ACZ6867
Rhopalopterum soror|[6875]|JWCHL371-10|BOLD:ACZ6867
Rhopalopterum soror|[6876]|JWCHL370-10|BOLD:ACZ6867
Rhopalopterum soror|[6877]|JWCHL335-10|BOLD:ACZ6867
Rhopalopterum soror|[6878]|JWCHL253-10|BOLD:ACZ6867
Rhopalopterum soror|[6879]|JWCHL137-10|BOLD:ACZ6867
Rhopalopterum soror|[6880]|JWCHL136-10|BOLD:ACZ6867
Rhopalopterum soror|[6881]|JWCHL124-10|BOLD:ACZ6867
Rhopalopterum soror|[6882]|JWCHL109-10|BOLD:ACZ6867
Rhopalopterum soror|[6883]|JWCHL090-10|BOLD:ACZ6867
Rhopalopterum soror|[6884]|JWCHL084-10|BOLD:ACZ6867
Rhopalopterum soror|[6885]|JWCHL051-10|BOLD:ACZ6867
Rhopalopterum soror|[6886]|JWCHL030-10|BOLD:ACZ6867
Rhopalopterum soror|[6887]|JWCHL027-10|BOLD:ACZ6867
Rhopalopterum soror|[6888]|JWCHL024-10|BOLD:ACZ6867
Rhopalopterum soror|[6889]|JWCHL088-10|BOLD:ACZ6867
Rhopalopterum soror|[6890]|JWCHL083-10|BOLD:ACZ6867
Rhopalopterum soror|[6891]|JWDCI387-10|BOLD:ACZ6867
Rhopalopterum soror|[6892]|JWDCG879-10|BOLD:ACZ6867
Rhopalopterum soror|[6893]|JWDCL477-11|BOLD:ACZ6867
Rhopalopterum soror|[6894]|JWDCJ1448-11|BOLD:ACZ6867
Rhopalopterum soror|[6895]|JWDCJ1446-11|BOLD:ACZ6867
Rhopalopterum soror|[6896]|JWDCJ350-11|BOLD:ACZ6867
Rhopalopterum soror|[6897]|JWCHL053-10|BOLD:ACZ6867
Rhopalopterum soror|[6898]|JWCHL055-10|BOLD:ACZ6867
Rhopalopterum soror|[6899]|JWCHL015-10|BOLD:ACZ6867
Rhopalopterum soror|[6900]|JWCHL418-10|BOLD:ACZ6867
Rhopalopterum soror|[6901]|JWCHL432-10|BOLD:ACZ6867
Rhopalopterum soror|[6902]|JWCHL473-10|BOLD:ACZ6867
Rhopalopterum soror|[6903]|JWCHL482-10|BOLD:ACZ6867
Rhopalopterum soror|[6904]|JWCHL488-10|BOLD:ACZ6867
Rhopalopterum soror|[6905]|JWCHL501-10|BOLD:ACZ6867
Rhopalopterum soror|[6906]|JWCHL524-10|BOLD:ACZ6867
Oscinella sp. TAW7|[6907]|JWCHL111-10|BOLD:AAG1522
Oscinella sp. TAW7|[6908]|JWCHL377-10|BOLD:AAG1522
Oscinella sp. TAW7|[6909]|JWCHL060-10|BOLD:AAG1522
Oscinella sp. TAW7|[6910]|JWCHL320-10|BOLD:AAG1522
Oscinella sp. TAW7|[6911]|JWCHL169-10|BOLD:AAG1522
Oscinella sp. TAW7|[6912]|JWCHL113-10|BOLD:AAG1522
Oscinella sp. TAW7|[6913]|JWCHL056-10|BOLD:AAG1522
Oscinella sp. TAW7|[6914]|JWCHL054-10|BOLD:AAG1522
Oscinella sp. TAW7|[6915]|JWCHL016-10|BOLD:AAG1522
Oscinella sp. TAW7|[6916]|JWDCG818-10|BOLD:AAG1522
Oscinella sp. TAW7|[6917]|JWCHL563-10|BOLD:AAG1522
Oscinella sp. TAW7|[6918]|JWDCA734-10|BOLD:AAG1522
Oscinella sp. TAW7|[6919]|JWCHL407-10|BOLD:AAG1522
Oscinella|[6920]|JWCHL087-10|BOLD:AAP5974
Conioscinella sp. TAW3|[6921]|JWCHL281-10|BOLD:AAP2545
Conioscinella sp. TAW3|[6922]|JWCHL376-10|BOLD:AAP2545
Conioscinella sp. TAW3|[6923]|JWCHL252-10|BOLD:ACG8143
Conioscinella sp. TAW3|[6924]|JWCHL288-10|BOLD:ACG8143
Conioscinella sp. TAW3|[6925]|JWDCI451-10|BOLD:ACG8143
Conioscinella sp. TAW3|[6926]|JWCHL309-10|BOLD:ACG8143
Conioscinella sp. TAW3|[6927]|JWCHL425-10|BOLD:ACG8143
Aphanotrigonum scabrum|[6928]|JWCHL477-10|BOLD:AAQ0868
Aphanotrigonum scabrum|[6929]|JWCHL478-10|BOLD:AAQ0868
Conioscinella sp. TAW1|[6930]|JWCHL003-10|BOLD:AAP6928
Conioscinella sp. TAW2|[6931]|JWDCB191-10|BOLD:AAG6803
Conioscinella sp. TAW2|[6932]|MTDIC087-09|BOLD:AAG6803
Conioscinella sp. TAW2|[6933]|JWDCC669-10|BOLD:AAG6803
Conioscinella sp. TAW2|[6934]|JWDCB192-10|BOLD:AAG6803
Conioscinella sp. TAW2|[6935]|JWDCB215-10|BOLD:AAG6803
Conioscinella sp. TAW2|[6936]|JWDCB226-10|BOLD:AAG6803
Conioscinella sp. TAW2|[6937]|JWCHL315-10|BOLD:AAG6803
Conioscinella sp. TAW2|[6938]|JWDCL521-11|BOLD:AAG6803
Conioscinella sp. TAW2|[6939]|JWDCH944-10|BOLD:AAG6803
Conioscinella sp. TAW2|[6940]|JWCHL337-10|BOLD:AAG6803
Conioscinella sp. TAW2|[6941]|JWCHL404-10|BOLD:AAG6803
Eribolus nana|[6942]|JWDCJ679-11|BOLD:AAL7392
Eribolus nana|[6943]|JWDCD138-10|BOLD:AAL7392
Eribolus nana|[6944]|JWCHL319-10|BOLD:AAL7392
Eribolus nana|[6945]|JWCHL299-10|BOLD:AAL7392
Eribolus nana|[6946]|JWCHL283-10|BOLD:AAL7392
Eribolus nana|[6947]|JWCHL220-10|BOLD:AAL7392
Eribolus nana|[6948]|JWCHL172-10|BOLD:AAL7392
Eribolus nana|[6949]|JWCHL141-10|BOLD:AAL7392
Eribolus nana|[6950]|JWCHL134-10|BOLD:AAL7392
Eribolus nana|[6951]|JWCHL556-10|BOLD:AAL7392
Eribolus nana|[6952]|JWDCD124-10|BOLD:AAL7392
Eribolus nana|[6953]|JWCHL287-10|BOLD:AAL7392
Eribolus nana|[6954]|JWCHL325-10|BOLD:AAL7392
Eribolus nana|[6955]|JWCHL327-10|BOLD:AAL7392
Eribolus nana|[6956]|JWCHL336-10|BOLD:AAL7392
Eribolus nana|[6957]|JWCHL367-10|BOLD:AAL7392
Eribolus nana|[6958]|JWCHL374-10|BOLD:AAL7392
Eribolus nana|[6959]|JWCHL385-10|BOLD:AAL7392
Eribolus nana|[6960]|JWCHL390-10|BOLD:AAL7392
Eribolus nana|[6961]|JWCHL474-10|BOLD:AAL7392
Eribolus nana|[6962]|JWCHL490-10|BOLD:AAL7392
Eribolus nana|[6963]|JWCHL497-10|BOLD:AAL7392
Eribolus nana|[6964]|JWCHL513-10|BOLD:AAL7392
Elachiptera decipiens|[6965]|JWDCJ489-11|BOLD:AAP3561
Elachiptera decipiens|[6966]|JWDCJ491-11|BOLD:AAP3561

[illegible]
Eribolus nana|[6964]|JWCHL513-10|BOLD:AAL7392
Elachiptera decipiens|[6965]|JWDCJ489-11|BOLD:AAP3561
Elachiptera decipiens|[6966]|JWDCJ491-11|BOLD:AAP3561
Elachiptera decipiens|[6967]|JWDCJ1142-11|BOLD:AAP3561
Elachiptera decipiens|[6968]|JWDCJ490-11|BOLD:AAP3561
Elachiptera decipiens|[6969]|JWCHL422-10|BOLD:AAP3561
Elachiptera decipiens|[6970]|JWCHL192-10|BOLD:AAH4164
Elachiptera decipiens|[6971]|JWDCC612-10|BOLD:AAH4164
Elachiptera decipiens|[6972]|JWCHL491-10|BOLD:AAH4164
Elachiptera decipiens|[6973]|JWCHL403-10|BOLD:AAH4164
Elachiptera decipiens|[6974]|JWCHL369-10|BOLD:AAH4164
Elachiptera decipiens|[6975]|JWCHL227-10|BOLD:AAH4164
Elachiptera decipiens|[6976]|JWCHL224-10|BOLD:AAH4164
Elachiptera decipiens|[6977]|JWDCJ056-11|BOLD:AAH4164
Elachiptera decipiens|[6978]|JWCHL541-10|BOLD:AAH4164
Elachiptera decipiens|[6979]|JWCHL546-10|BOLD:AAH4164
Elachiptera decipiens|[6980]|JWCHL515-10|BOLD:AAH4164
Elachiptera decipiens|[6981]|JWCHL125-10|BOLD:AAH4164
Elachiptera decipiens|[6982]|JWCHL483-10|BOLD:AAH4164
Elachiptera decipiens|[6983]|JWCHL193-10|BOLD:AAH4164
Elachiptera decipiens|[6984]|JWCHL215-10|BOLD:AAH4164
Elachiptera decipiens|[6985]|JWCHL419-10|BOLD:AAH4164
Elachiptera decipiens|[6986]|TWDIP153-09|BOLD:AAH4164
Elachiptera decipiens|[6987]|JWCHL531-10|BOLD:AAH4164
Elachiptera decipiens|[6988]|JWCHL221-10|BOLD:AAH4164
Elachiptera decipiens|[6989]|JWCHL545-10|BOLD:AAH4164
Elachiptera decipiens|[6990]|JWCHL182-10|BOLD:AAH4164
Elachiptera decipiens|[6991]|JWDCI030-10|BOLD:AAH4164
Elachiptera decipiens|[6992]|JWDCC587-10|BOLD:AAH4164
Elachiptera decipiens|[6993]|JWCHL543-10|BOLD:AAH4164
Elachiptera decipiens|[6994]|JWCHL542-10|BOLD:AAH4164
Elachiptera decipiens|[6995]|JWCHL529-10|BOLD:AAH4164
Elachiptera decipiens|[6996]|JWCHL528-10|BOLD:AAH4164
Elachiptera decipiens|[6997]|JWCHL526-10|BOLD:AAH4164
Elachiptera decipiens|[6998]|JWCHL520-10|BOLD:AAH4164
Elachiptera decipiens|[6999]|JWCHL512-10|BOLD:AAH4164
Elachiptera decipiens|[7000]|JWCHL496-10|BOLD:AAH4164
Elachiptera decipiens|[7001]|JWCHL480-10|BOLD:AAH4164
Elachiptera decipiens|[7002]|JWCHL539-10|BOLD:AAH4164
Elachiptera decipiens|[7003]|JWCHL523-10|BOLD:AAH4164
Elachiptera decipiens|[7004]|JWCHL465-10|BOLD:AAH4164
Elachiptera decipiens|[7005]|JWCHL426-10|BOLD:AAH4164
Elachiptera decipiens|[7006]|JWCHL402-10|BOLD:AAH4164
Elachiptera decipiens|[7007]|JWCHL392-10|BOLD:AAH4164
Elachiptera decipiens|[7008]|JWCHL391-10|BOLD:AAH4164
Elachiptera decipiens|[7009]|JWCHL379-10|BOLD:AAH4164
Elachiptera decipiens|[7010]|JWCHL356-10|BOLD:AAH4164
Elachiptera decipiens|[7011]|JWCHL355-10|BOLD:AAH4164
Elachiptera decipiens|[7012]|JWCHL340-10|BOLD:AAH4164
Elachiptera decipiens|[7013]|JWCHL324-10|BOLD:AAH4164
Elachiptera decipiens|[7014]|JWCHL323-10|BOLD:AAH4164
Elachiptera decipiens|[7015]|JWCHL259-10|BOLD:AAH4164
Elachiptera decipiens|[7016]|JWCHL248-10|BOLD:AAH4164
Elachiptera decipiens|[7017]|JWCHL246-10|BOLD:AAH4164
Elachiptera decipiens|[7018]|JWCHL237-10|BOLD:AAH4164
Elachiptera decipiens|[7019]|JWCHL236-10|BOLD:AAH4164
Elachiptera decipiens|[7020]|JWCHL235-10|BOLD:AAH4164
Elachiptera decipiens|[7021]|JWCHL234-10|BOLD:AAH4164
Elachiptera decipiens|[7022]|JWCHL232-10|BOLD:AAH4164
Elachiptera decipiens|[7023]|JWCHL223-10|BOLD:AAH4164
Elachiptera decipiens|[7024]|JWCHL222-10|BOLD:AAH4164
Elachiptera decipiens|[7025]|JWCHL219-10|BOLD:AAH4164
Elachiptera decipiens|[7026]|JWCHL217-10|BOLD:AAH4164
Elachiptera decipiens|[7027]|JWCHL209-10|BOLD:AAH4164
Elachiptera decipiens|[7028]|JWCHL204-10|BOLD:AAH4164
Elachiptera decipiens|[7029]|JWCHL199-10|BOLD:AAH4164
Elachiptera decipiens|[7030]|JWCHL194-10|BOLD:AAH4164
Elachiptera decipiens|[7031]|JWCHL191-10|BOLD:AAH4164
Elachiptera decipiens|[7032]|JWCHL157-10|BOLD:AAH4164
Elachiptera decipiens|[7033]|JWCHL148-10|BOLD:AAH4164
Elachiptera decipiens|[7034]|JWCHL128-10|BOLD:AAH4164
Elachiptera decipiens|[7035]|JWCHL091-10|BOLD:AAH4164
Elachiptera decipiens|[7036]|JWCHL004-10|BOLD:AAH4164
Elachiptera decipiens|[7037]|JWCHL086-10|BOLD:AAH4164
Elachiptera decipiens|[7038]|JWDCD140-10|BOLD:AAH4164
Elachiptera decipiens|[7039]|JWDCC707-10|BOLD:AAH4164
Elachiptera decipiens|[7040]|JWDCC588-10|BOLD:AAH4164
Elachiptera decipiens|[7041]|JWCHL231-10|BOLD:AAH4164
Elachiptera decipiens|[7042]|JWCHL195-10|BOLD:AAH4164
Elachiptera decipiens|[7043]|JWCHL286-10|BOLD:AAH4164
Elachiptera decipiens|[7044]|JWCHL561-10|BOLD:AAH4164
Elachiptera decipiens|[7045]|JWCHL218-10|BOLD:AAH4164
Elachiptera decipiens|[7046]|JWDCI388-10|BOLD:AAH4164
Elachiptera decipiens|[7047]|JWCHL492-10|BOLD:AAH4164
Elachiptera decipiens|[7048]|JWCHL547-10|BOLD:AAH4164
Tricimba melancholica|[7049]|MTDIC128-09|BOLD:AAG1503
Tricimba melancholica|[7050]|MTDIC181-09|BOLD:AAG1503
Tricimba melancholica|[7051]|JWDCC673-10|BOLD:AAG1503
Tricimba melancholica|[7052]|JWDCB232-10|BOLD:AAG1503
Tricimba melancholica|[7053]|JWDCA730-10|BOLD:AAG1503
Tricimba melancholica|[7054]|JWDCC670-10|BOLD:AAG1503
Tricimba melancholica|[7055]|JWDCB211-10|BOLD:AAG1503
Tricimba melancholica|[7056]|JWDCB213-10|BOLD:AAG1503
Tricimba melancholica|[7057]|JWDCA712-10|BOLD:AAG1503
Tricimba melancholica|[7058]|JWDCA702-10|BOLD:AAG1503
Tricimba melancholica|[7059]|JWDCB194-10|BOLD:AAG1503
Tricimba melancholica|[7060]|JWDCA749-10|BOLD:AAG1503
Tricimba melancholica|[7061]|JWDCH185-10|BOLD:AAG1503
Tricimba melancholica|[7062]|JWDCH863-10|BOLD:AAG1503
Tricimba melancholica|[7063]|JWDCJ1202-11|BOLD:AAG1503
Tricimba melancholica|[7064]|JWDCJ1200-11|BOLD:AAG1503
Tricimba melancholica|[7065]|JWDCB210-10|BOLD:AAG1503

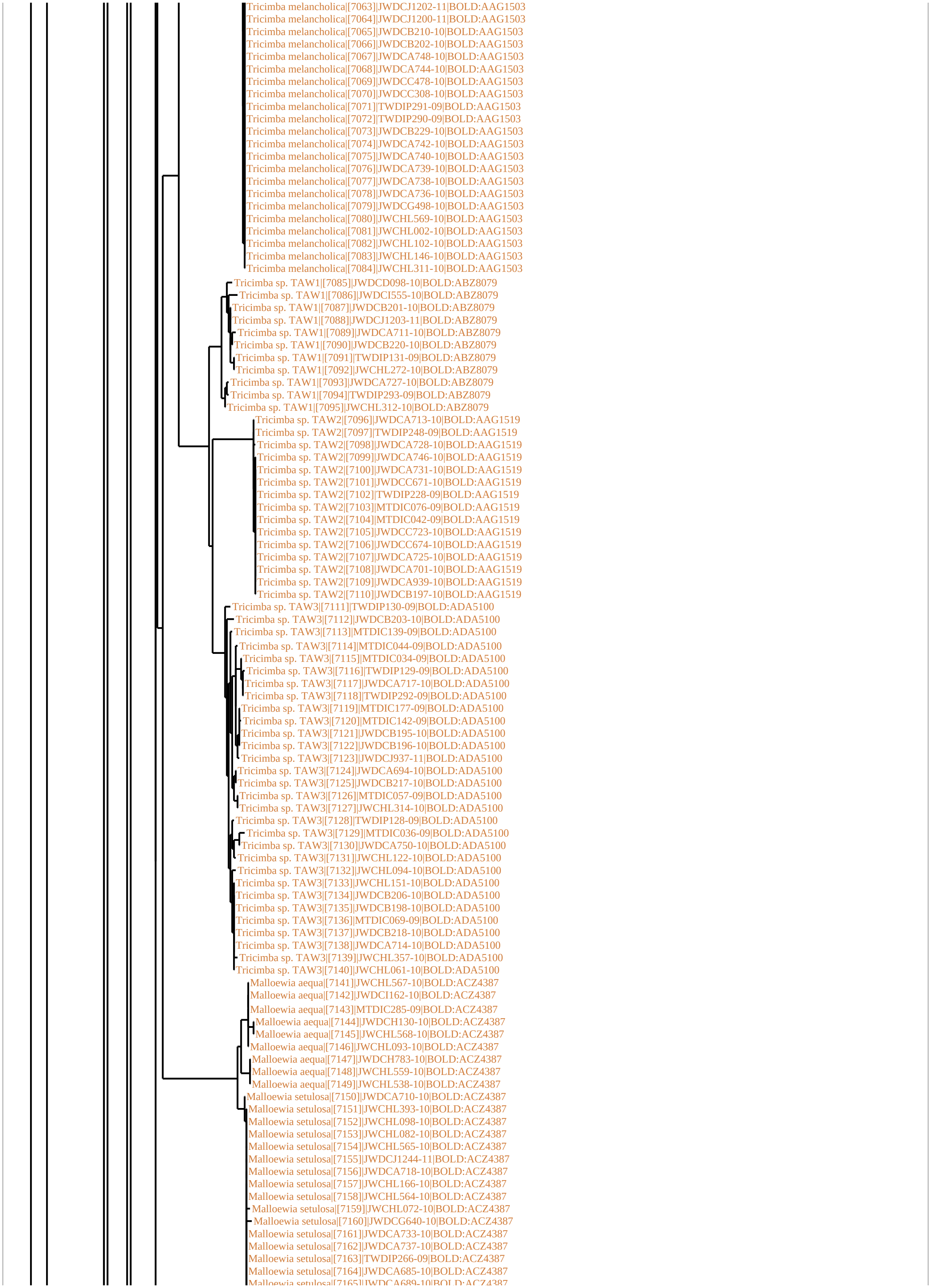
Tricimba melancholica|[7063]|JWDCJ1202-11|BOLD:AAG1503
Tricimba melancholica|[7064]|JWDCJ1200-11|BOLD:AAG1503
Tricimba melancholica|[7065]|JWDCB210-10|BOLD:AAG1503
Tricimba melancholica|[7066]|JWDCB202-10|BOLD:AAG1503
Tricimba melancholica|[7067]|JWDCA748-10|BOLD:AAG1503
Tricimba melancholica|[7068]|JWDCA744-10|BOLD:AAG1503
Tricimba melancholica|[7069]|JWDCC478-10|BOLD:AAG1503
Tricimba melancholica|[7070]|JWDCC308-10|BOLD:AAG1503
Tricimba melancholica|[7071]|TWDIP291-09|BOLD:AAG1503
Tricimba melancholica|[7072]|TWDIP290-09|BOLD:AAG1503
Tricimba melancholica|[7073]|JWDCB229-10|BOLD:AAG1503
Tricimba melancholica|[7074]|JWDCA742-10|BOLD:AAG1503
Tricimba melancholica|[7075]|JWDCA740-10|BOLD:AAG1503
Tricimba melancholica|[7076]|JWDCA739-10|BOLD:AAG1503
Tricimba melancholica|[7077]|JWDCA738-10|BOLD:AAG1503
Tricimba melancholica|[7078]|JWDCA736-10|BOLD:AAG1503
Tricimba melancholica|[7079]|JWDCG498-10|BOLD:AAG1503
Tricimba melancholica|[7080]|JWCHL569-10|BOLD:AAG1503
Tricimba melancholica|[7081]|JWCHL002-10|BOLD:AAG1503
Tricimba melancholica|[7082]|JWCHL102-10|BOLD:AAG1503
Tricimba melancholica|[7083]|JWCHL146-10|BOLD:AAG1503
Tricimba melancholica|[7084]|JWCHL311-10|BOLD:AAG1503
Tricimba sp. TAW1|[7085]|JWDCD098-10|BOLD:ABZ8079
Tricimba sp. TAW1|[7086]|JWDCI555-10|BOLD:ABZ8079
Tricimba sp. TAW1|[7087]|JWDCB201-10|BOLD:ABZ8079
Tricimba sp. TAW1|[7088]|JWDCJ1203-11|BOLD:ABZ8079
Tricimba sp. TAW1|[7089]|JWDCA711-10|BOLD:ABZ8079
Tricimba sp. TAW1|[7090]|JWDCB220-10|BOLD:ABZ8079
Tricimba sp. TAW1|[7091]|TWDIP131-09|BOLD:ABZ8079
Tricimba sp. TAW1|[7092]|JWCHL272-10|BOLD:ABZ8079
Tricimba sp. TAW1|[7093]|JWDCA727-10|BOLD:ABZ8079
Tricimba sp. TAW1|[7094]|TWDIP293-09|BOLD:ABZ8079
Tricimba sp. TAW1|[7095]|JWCHL312-10|BOLD:ABZ8079
Tricimba sp. TAW2|[7096]|JWDCA713-10|BOLD:AAG1519
Tricimba sp. TAW2|[7097]|TWDIP248-09|BOLD:AAG1519
Tricimba sp. TAW2|[7098]|JWDCA728-10|BOLD:AAG1519
Tricimba sp. TAW2|[7099]|JWDCA746-10|BOLD:AAG1519
Tricimba sp. TAW2|[7100]|JWDCA731-10|BOLD:AAG1519
Tricimba sp. TAW2|[7101]|JWDCC671-10|BOLD:AAG1519
Tricimba sp. TAW2|[7102]|TWDIP228-09|BOLD:AAG1519
Tricimba sp. TAW2|[7103]|MTDIC076-09|BOLD:AAG1519
Tricimba sp. TAW2|[7104]|MTDIC042-09|BOLD:AAG1519
Tricimba sp. TAW2|[7105]|JWDCC723-10|BOLD:AAG1519
Tricimba sp. TAW2|[7106]|JWDCC674-10|BOLD:AAG1519
Tricimba sp. TAW2|[7107]|JWDCA725-10|BOLD:AAG1519
Tricimba sp. TAW2|[7108]|JWDCA701-10|BOLD:AAG1519
Tricimba sp. TAW2|[7109]|JWDCA939-10|BOLD:AAG1519
Tricimba sp. TAW2|[7110]|JWDCB197-10|BOLD:AAG1519
Tricimba sp. TAW3|[7111]|TWDIP130-09|BOLD:ADA5100
Tricimba sp. TAW3|[7112]|JWDCB203-10|BOLD:ADA5100
Tricimba sp. TAW3|[7113]|MTDIC139-09|BOLD:ADA5100
Tricimba sp. TAW3|[7114]|MTDIC044-09|BOLD:ADA5100
Tricimba sp. TAW3|[7115]|MTDIC034-09|BOLD:ADA5100
Tricimba sp. TAW3|[7116]|TWDIP129-09|BOLD:ADA5100
Tricimba sp. TAW3|[7117]|JWDCA717-10|BOLD:ADA5100
Tricimba sp. TAW3|[7118]|TWDIP292-09|BOLD:ADA5100
Tricimba sp. TAW3|[7119]|MTDIC177-09|BOLD:ADA5100
Tricimba sp. TAW3|[7120]|MTDIC142-09|BOLD:ADA5100
Tricimba sp. TAW3|[7121]|JWDCB195-10|BOLD:ADA5100
Tricimba sp. TAW3|[7122]|JWDCB196-10|BOLD:ADA5100
Tricimba sp. TAW3|[7123]|JWDCJ937-11|BOLD:ADA5100
Tricimba sp. TAW3|[7124]|JWDCA694-10|BOLD:ADA5100
Tricimba sp. TAW3|[7125]|JWDCB217-10|BOLD:ADA5100
Tricimba sp. TAW3|[7126]|MTDIC057-09|BOLD:ADA5100
Tricimba sp. TAW3|[7127]|JWCHL314-10|BOLD:ADA5100
Tricimba sp. TAW3|[7128]|TWDIP128-09|BOLD:ADA5100
Tricimba sp. TAW3|[7129]|MTDIC036-09|BOLD:ADA5100
Tricimba sp. TAW3|[7130]|JWDCA750-10|BOLD:ADA5100
Tricimba sp. TAW3|[7131]|JWCHL122-10|BOLD:ADA5100
Tricimba sp. TAW3|[7132]|JWCHL094-10|BOLD:ADA5100
Tricimba sp. TAW3|[7133]|JWCHL151-10|BOLD:ADA5100
Tricimba sp. TAW3|[7134]|JWDCB206-10|BOLD:ADA5100
Tricimba sp. TAW3|[7135]|JWDCB198-10|BOLD:ADA5100
Tricimba sp. TAW3|[7136]|MTDIC069-09|BOLD:ADA5100
Tricimba sp. TAW3|[7137]|JWDCB218-10|BOLD:ADA5100
Tricimba sp. TAW3|[7138]|JWDCA714-10|BOLD:ADA5100
Tricimba sp. TAW3|[7139]|JWCHL357-10|BOLD:ADA5100
Tricimba sp. TAW3|[7140]|JWCHL061-10|BOLD:ADA5100
Malloewia aequa|[7141]|JWCHL567-10|BOLD:ACZ4387
Malloewia aequa|[7142]|JWDCI162-10|BOLD:ACZ4387
Malloewia aequa|[7143]|MTDIC285-09|BOLD:ACZ4387
Malloewia aequa|[7144]|JWDCH130-10|BOLD:ACZ4387
Malloewia aequa|[7145]|JWCHL568-10|BOLD:ACZ4387
Malloewia aequa|[7146]|JWCHL093-10|BOLD:ACZ4387
Malloewia aequa|[7147]|JWDCH783-10|BOLD:ACZ4387
Malloewia aequa|[7148]|JWCHL559-10|BOLD:ACZ4387
Malloewia aequa|[7149]|JWCHL538-10|BOLD:ACZ4387
Malloewia setulosa|[7150]|JWDCA710-10|BOLD:ACZ4387
Malloewia setulosa|[7151]|JWCHL393-10|BOLD:ACZ4387
Malloewia setulosa|[7152]|JWCHL098-10|BOLD:ACZ4387
Malloewia setulosa|[7153]|JWCHL082-10|BOLD:ACZ4387
Malloewia setulosa|[7154]|JWCHL565-10|BOLD:ACZ4387
Malloewia setulosa|[7155]|JWDCJ1244-11|BOLD:ACZ4387
Malloewia setulosa|[7156]|JWDCA718-10|BOLD:ACZ4387
Malloewia setulosa|[7157]|JWCHL166-10|BOLD:ACZ4387
Malloewia setulosa|[7158]|JWCHL564-10|BOLD:ACZ4387
Malloewia setulosa|[7159]|JWCHL072-10|BOLD:ACZ4387
Malloewia setulosa|[7160]|JWDCG640-10|BOLD:ACZ4387
Malloewia setulosa|[7161]|JWDCA733-10|BOLD:ACZ4387
Malloewia setulosa|[7162]|JWDCA737-10|BOLD:ACZ4387
Malloewia setulosa|[7163]|TWDIP266-09|BOLD:ACZ4387
Malloewia setulosa|[7164]|JWDCA685-10|BOLD:ACZ4387
Malloewia setulosa|[7165]|JWDCA689-10|BOLD:ACZ4387

[illegible]
Malloewia setulosa|[7163]|TWDIP266-09|BOLD:ACZ4387
Malloewia setulosa|[7164]|JWDCA685-10|BOLD:ACZ4387
Malloewia setulosa|[7165]|JWDCA689-10|BOLD:ACZ4387
Malloewia setulosa|[7166]|JWDCB214-10|BOLD:ACZ4387
Malloewia setulosa|[7167]|JWDCB212-10|BOLD:ACZ4387
Malloewia setulosa|[7168]|JWDCB235-10|BOLD:ACZ4387
Malloewia setulosa|[7169]|JWDCJ1205-11|BOLD:ACZ4387
Malloewia setulosa|[7170]|JWDCJ1204-11|BOLD:ACZ4387
Malloewia setulosa|[7171]|JWDCL093-11|BOLD:ACZ4387
Malloewia setulosa|[7172]|JWDCL094-11|BOLD:ACZ4387
Malloewia setulosa|[7173]|JWDCG639-10|BOLD:ACZ4387
Malloewia setulosa|[7174]|JWDCG686-10|BOLD:ACZ4387
Malloewia setulosa|[7175]|JWDCH945-10|BOLD:ACZ4387
Malloewia setulosa|[7176]|JWDCI452-10|BOLD:ACZ4387
Malloewia setulosa|[7177]|JWDCH266-10|BOLD:ACZ4387
Malloewia setulosa|[7178]|JWDCH328-10|BOLD:ACZ4387
Malloewia setulosa|[7179]|JWDCH275-10|BOLD:ACZ4387
Malloewia setulosa|[7180]|JWCHL558-10|BOLD:ACZ4387
Malloewia setulosa|[7181]|JWCHL570-10|BOLD:ACZ4387
Malloewia setulosa|[7182]|JWCHL048-10|BOLD:ACZ4387
Malloewia setulosa|[7183]|JWCHL067-10|BOLD:ACZ4387
Malloewia setulosa|[7184]|JWCHL069-10|BOLD:ACZ4387
Malloewia setulosa|[7185]|JWCHL070-10|BOLD:ACZ4387
Malloewia setulosa|[7186]|JWCHL071-10|BOLD:ACZ4387
Malloewia setulosa|[7187]|JWCHL085-10|BOLD:ACZ4387
Malloewia setulosa|[7188]|JWCHL092-10|BOLD:ACZ4387
Malloewia setulosa|[7189]|JWCHL100-10|BOLD:ACZ4387
Malloewia setulosa|[7190]|JWCHL119-10|BOLD:ACZ4387
Malloewia setulosa|[7191]|JWCHL120-10|BOLD:ACZ4387
Malloewia setulosa|[7192]|JWCHL121-10|BOLD:ACZ4387
Malloewia setulosa|[7193]|JWCHL123-10|BOLD:ACZ4387
Malloewia setulosa|[7194]|JWCHL184-10|BOLD:ACZ4387
Malloewia setulosa|[7195]|JWCHL429-10|BOLD:ACZ4387
Malloewia setulosa|[7196]|JWCHL437-10|BOLD:ACZ4387
Malloewia setulosa|[7197]|JWCHL031-10|BOLD:ACZ4387
Malloewia setulosa|[7198]|JWCHL476-10|BOLD:ACZ4387
Malloewia setulosa|[7199]|JWCHL551-10|BOLD:ACZ4387
Malloewia setulosa|[7200]|JWCHL552-10|BOLD:ACZ4387
Dasyopa sp. TAW1|[7201]|MTDIC183-09|BOLD:AAG1504
Dasyopa sp. TAW1|[7202]|JWDCA729-10|BOLD:AAG1504
Dasyopa sp. TAW1|[7203]|JWDCA726-10|BOLD:AAG1504
Dasyopa sp. TAW1|[7204]|MTDIC089-09|BOLD:AAG1504
Dasyopa sp. TAW1|[7205]|MTDIC109-09|BOLD:AAG1504
Dasyopa sp. TAW1|[7206]|JWDCB224-10|BOLD:AAG1504
Dasyopa sp. TAW1|[7207]|JWDCB228-10|BOLD:AAG1504
Dasyopa sp. TAW1|[7208]|JWCHL550-10|BOLD:AAG1504
Dasyopa triangulata|[7209]|JWDCB209-10|BOLD:AAG1518
Dasyopa triangulata|[7210]|JWDCA705-10|BOLD:AAG1518
Dasyopa triangulata|[7211]|JWDCA704-10|BOLD:AAG1518
Dasyopa triangulata|[7212]|JWDCA703-10|BOLD:AAG1518
Dasyopa triangulata|[7213]|JWDCB219-10|BOLD:AAG1518
Dasyopa triangulata|[7214]|JWDCB233-10|BOLD:AAG1518
Dasyopa triangulata|[7215]|JWDCB230-10|BOLD:AAG1518
Dasyopa triangulata|[7216]|JWDCB227-10|BOLD:AAG1518
Dasyopa triangulata|[7217]|JWDCB208-10|BOLD:AAG1518
Dasyopa triangulata|[7218]|JWDCA944-10|BOLD:AAG1518
Dasyopa triangulata|[7219]|JWDCA751-10|BOLD:AAG1518
Dasyopa triangulata|[7220]|JWDCA745-10|BOLD:AAG1518
Dasyopa triangulata|[7221]|MTDIC162-09|BOLD:AAG1518
Dasyopa triangulata|[7222]|JWDCA697-10|BOLD:AAG1518
Dasyopa triangulata|[7223]|JWDCA735-10|BOLD:AAG1518
Dasyopa triangulata|[7224]|JWDCA716-10|BOLD:AAG1518
Dasyopa triangulata|[7225]|JWDCA715-10|BOLD:AAG1518
Dasyopa triangulata|[7226]|JWDCA700-10|BOLD:AAG1518
Dasyopa triangulata|[7227]|JWDCA696-10|BOLD:AAG1518
Dasyopa triangulata|[7228]|JWDCB231-10|BOLD:AAG1518
Dasyopa triangulata|[7229]|TWDIP227-09|BOLD:AAG1518
Dasyopa triangulata|[7230]|JWDCB207-10|BOLD:AAG1518
Dasyopa triangulata|[7231]|JWDCB236-10|BOLD:AAG1518
Dasyopa triangulata|[7232]|JWDCB221-10|BOLD:AAG1518
Dasyopa triangulata|[7233]|JWDCB225-10|BOLD:AAG1518
Dasyopa triangulata|[7234]|JWDCJ1138-11|BOLD:AAG1518
Dasyopa triangulata|[7235]|JWCHL168-10|BOLD:AAG1518
Dasyopa triangulata|[7236]|JWCHL503-10|BOLD:AAG1518
Rhopalopterum soror|[7237]|JWDCA741-10|BOLD:AAG1523
Rhopalopterum carbonarium|[7238]|JWCHL386-10|BOLD:AAP3813
Rhopalopterum carbonarium|[7239]|JWCHL057-10|BOLD:AAP3813
Rhopalopterum carbonarium|[7240]|JWCHL489-10|BOLD:AAP3813
Rhopalopterum carbonarium|[7241]|JWCHL089-10|BOLD:AAP3813
Rhopalopterum carbonarium|[7242]|JWCHL112-10|BOLD:AAP3813
Rhopalopterum carbonarium|[7243]|JWCHL138-10|BOLD:AAP3813
Rhopalopterum carbonarium|[7244]|JWCHL158-10|BOLD:AAP3813
Rhopalopterum carbonarium|[7245]|JWCHL334-10|BOLD:AAP3813
Rhopalopterum carbonarium|[7246]|JWCHL381-10|BOLD:AAP3813
Rhopalopterum carbonarium|[7247]|JWCHL423-10|BOLD:AAP3813
Rhopalopterum carbonarium|[7248]|JWCHL427-10|BOLD:AAP3813
Rhopalopterum carbonarium|[7249]|JWCHL028-10|BOLD:AAP3813
Rhopalopterum carbonarium|[7250]|JWCHL058-10|BOLD:AAP3813
Rhopalopterum carbonarium|[7251]|JWCHL485-10|BOLD:AAP3813
Rhopalopterum carbonarium|[7252]|JWCHL498-10|BOLD:AAP3813
Rhopalopterum carbonarium|[7253]|JWCHL544-10|BOLD:AAP3813
Microcercis albipalpis|[7254]|JWCHL034-10|BOLD:AAP6790
Microcercis albipalpis|[7255]|JWCHL449-10|BOLD:AAP6790
Microcercis albipalpis|[7256]|JWCHL417-10|BOLD:AAP6790
Microcercis albipalpis|[7257]|JWCHL290-10|BOLD:AAP6790
Microcercis albipalpis|[7258]|JWCHL450-10|BOLD:AAP6790
Rhopalopterum soror|[7259]|JWCHL033-10|BOLD:AGA8316
Rhopalopterum soror|[7260]|JWCHL358-10|BOLD:AGA8316
Microcercis bispina|[7261]|JWCHL494-10|BOLD:AGA8316
Microcercis bispina|[7262]|JWCHL495-10|BOLD:AGA8316
Oscinella sp. TAW1|[7263]|JWCHL278-10|BOLD:AAL7389
Oscinella sp. TAW1|[7264]|JWCHL294-10|BOLD:AAL7389

Microcercis bispina|[7262]|JWCHL495-10|BOLD:AGA8316
Oscinella sp. TAW1|[7263]|JWCHL278-10|BOLD:AAL7389
Oscinella sp. TAW1|[7264]|JWCHL294-10|BOLD:AAL7389
Oscinella sp. TAW1|[7265]|JWCHL493-10|BOLD:AAL7389
Oscinella sp. TAW1|[7266]|JWDCC672-10|BOLD:AAL7389
Oscinella sp. TAW1|[7267]|JWDCC667-10|BOLD:AAL7389
Oscinella sp. TAW1|[7268]|JWCHL005-10|BOLD:AAL7389
Oscinella sp. TAW1|[7269]|JWCHL007-10|BOLD:AAL7389
Oscinella sp. TAW1|[7270]|JWCHL292-10|BOLD:AAL7389
Oscinella sp. TAW1|[7271]|MTDIC178-09|BOLD:AAL7389
Oscinella sp. TAW1|[7272]|JWCHL307-10|BOLD:AAL7389
Oscinella sp. TAW1|[7273]|JWCHL275-10|BOLD:AAL7389
Oscinella sp. TAW1|[7274]|JWCHL276-10|BOLD:AAL7389
Oscinella sp. TAW1|[7275]|JWCHL553-10|BOLD:AAL7389
Oscinella sp. TAW5|[7276]|JWCHL310-10|BOLD:AAB1152
Oscinella sp. TAW2|[7277]|JWCHL177-10|BOLD:AAI3425
Oscinella sp. TAW2|[7278]|JWCHL178-10|BOLD:AAI3425
Oscinella sp. TAW2|[7279]|JWCHL179-10|BOLD:AAI3425
Oscinella sp. TAW2|[7280]|JWCHL181-10|BOLD:AAI3425
Oscinella sp. TAW6|[7281]|JWCHL267-10
Oscinella sp. TAW6|[7282]|JWDCC571-10|BOLD:AAL7388
Oscinella sp. TAW6|[7283]|JWCHL342-10|BOLD:AAL7388
Oscinella sp. TAW6|[7284]|JWCHL266-10|BOLD:AAL7388
Oscinella sp. TAW6|[7285]|JWCHL265-10|BOLD:AAL7388
Oscinella sp. TAW6|[7286]|JWCHL270-10|BOLD:AAL7388
Oscinella sp. TAW6|[7287]|JWCHL343-10|BOLD:AAL7388
Oscinella sp. TAW6|[7288]|JWCHL420-10|BOLD:AAL7388
Oscinella sp. TAW6|[7289]|JWCHL424-10|BOLD:AAL7388
Oscinella sp. TAW4|[7290]|JWCHL289-10|BOLD:AAB1150
Oscinella sp. TAW3|[7291]|JWCHL009-10
Oscinella sp. TAW3|[7292]|JWDCJ351-11|BOLD:AAB1148
Oscinella sp. TAW3|[7293]|JWCHL460-10|BOLD:AAB1148
Oscinella sp. TAW3|[7294]|JWCHL462-10|BOLD:AAB1148
Oscinella sp. TAW3|[7295]|JWCHL461-10|BOLD:AAB1148
Oscinella sp. TAW3|[7296]|JWCHL463-10|BOLD:AAB1148
Oscinella sp. TAW3|[7297]|JWCHL464-10|BOLD:AAB1148
Oscinella sp. TAW3|[7298]|JWCHL180-10|BOLD:AAB1148
Oscinella sp. TAW3|[7299]|JWDCJ1028-11|BOLD:AAB1148
Oscinella sp. TAW3|[7300]|JWCHL305-10|BOLD:AAB1148
Oscinella sp. TAW3|[7301]|JWCHL008-10|BOLD:AAB1148
Oscinella sp. TAW3|[7302]|JWCHL279-10|BOLD:AAB1148
Oscinella sp. TAW3|[7303]|JWCHL291-10|BOLD:AAB1148
Oscinella sp. TAW3|[7304]|JWCHL293-10|BOLD:AAB1148
Oscinella sp. TAW3|[7305]|JWDCC668-10|BOLD:AAB1148
Oscinella sp. TAW3|[7306]|JWCHL006-10|BOLD:AAB1148
Oscinella sp. TAW3|[7307]|JWCHL554-10|BOLD:AAB1148
Oscinella sp. TAW3|[7308]|JWCHL555-10|BOLD:AAB1148
Crumomyia annulus|[7309]|JWDCF399-10|BOLD:AAP8767
Suillia nemorum|[7310]|JWDCB099-10|BOLD:ACX5303
Suillia nemorum|[7311]|JWDCA253-10|BOLD:ACX5303
Suillia nemorum|[7312]|JWDCJ1271-11|BOLD:ACX5303
Lonchaea sp. TAW1|[7313]|TWDIP360-09|BOLD:AAG7068
Lonchaea sp. TAW9|[7314]|MTDIC216-09|BOLD:AAG7067
Lonchaea sp. TAW6|[7315]|JWDCA626-10|BOLD:AAG7079
Lonchaea sp. TAW7|[7316]|JWDCJ359-11|BOLD:AAG7076
Lonchaea sp. TAW8|[7317]|JWDCA550-10|BOLD:AAG6844
Lonchaea sp. TAW8|[7318]|MTDIC049-09|BOLD:AAG6844
Lonchaea sp. TAW8|[7319]|TWDIP009-09|BOLD:AAG6844
Lonchaea sp. TAW8|[7320]|JWDCA551-10|BOLD:AAG6844
Lonchaea sp. TAW8|[7321]|JWDCJ361-11|BOLD:AAG6844
Lonchaea sp. TAW8|[7322]|JWDCJ1111-11|BOLD:AAG6844
Lonchaea sp. TAW8|[7323]|JWDCJ1110-11|BOLD:AAG6844
Lonchaea sp. TAW8|[7324]|JWDCJ1197-11|BOLD:AAG6844
Lonchaea sp. TAW8|[7325]|JWDCJ1105-11|BOLD:AAG6844
Lonchaea sp. TAW8|[7326]|JWDCA552-10|BOLD:AAG6844
Lonchaea sp. TAW8|[7327]|TWDIP008-09|BOLD:AAG6844
Lonchaea sp. TAW8|[7328]|MTDIC047-09|BOLD:AAG6844
Lonchaea sp. TAW8|[7329]|MTDIC072-09|BOLD:AAG6844
Lonchaea sp. TAW8|[7330]|JWDCA549-10|BOLD:AAG6844
Lonchaea sp. TAW8|[7331]|JWDCA553-10|BOLD:AAG6844
Lonchaea sp. TAW8|[7332]|JWDCA554-10|BOLD:AAG6844
Lonchaea sp. TAW8|[7333]|JWDCA572-10|BOLD:AAG6844
Lonchaea sp. TAW8|[7334]|JWDCJ360-11|BOLD:AAG6844
Lonchaea sp. TAW8|[7335]|JWDCJ536-11|BOLD:AAG6844
Lonchaea sp. TAW8|[7336]|JWDCJ738-11|BOLD:AAG6844
Lonchaea sp. TAW8|[7337]|JWDCJ1143-11|BOLD:AAG6844
Lonchaea sp. TAW8|[7338]|JWDCJ1192-11|BOLD:AAG6844
Lonchaea sp. TAW8|[7339]|JWDCJ1198-11|BOLD:AAG6844
Lonchaea sp. TAW4|[7340]|JWDCG054-10|BOLD:AAP8895
Lonchaea sp. TAW5|[7341]|JWDCA624-10|BOLD:AAG7078
Lonchaea sp. TAW5|[7342]|JWDCK191-11|BOLD:AAG7078
Lonchaea sp. TAW5|[7343]|JWDCH416-10|BOLD:AAG7078
Lonchaea sp. TAW3|[7344]|JWDCE618-10|BOLD:AAP4836
Lonchaea sp. TAW2|[7345]|JWDCE516-10|BOLD:AAG6956
Mycetaulus bipunctatus group|[7346]|JWDCJ1139-11|BOLD:AAU5480
Allopiophila luteata|[7347]|JWDCH177-10|BOLD:AAP8546
Allopiophila luteata|[7348]|JWDCI460-10|BOLD:AAP8546
Parapiophila sp. TAW3|[7349]|TWDIP105-09|BOLD:AAG1786
Parapiophila sp. TAW3|[7350]|TWDIP106-09|BOLD:AAG1786
Parapiophila sp. TAW3|[7351]|MTDIC212-09|BOLD:AAG1786
Parapiophila sp. TAW3|[7352]|JWDCA560-10|BOLD:AAG1786
Parapiophila sp. TAW2|[7353]|TWDIP108-09|BOLD:AAG1787
Parapiophila sp. TAW2|[7354]|JWDCA721-10|BOLD:AAG1787
Parapiophila sp. TAW2|[7355]|JWDCA588-10|BOLD:AAG1787
Parapiophila sp. TAW1|[7356]|JWDCI926-11|BOLD:AAI2623
Parapiophila sp. TAW4|[7357]|JWDCA618-10|BOLD:AAG1788
Parapiophila sp. TAW4|[7358]|MTDIC271-09|BOLD:AAG1788
Parapiophila sp. TAW4|[7359]|JWDCG852-10|BOLD:AAG1788
Parapiophila sp. TAW4|[7360]|JWDCA611-10|BOLD:AAG1788
Parapiophila sp. TAW4|[7361]|MTDIC137-09|BOLD:AAG1788
Parapiophila sp. TAW4|[7362]|JWDCA616-10|BOLD:AAG1788
Parapiophila sp. TAW4|[7363]|JWDCK085-11|BOLD:AAG1788
Poecilolycia sp.|[7364]|MTDIC245-09|BOLD:AAH3532

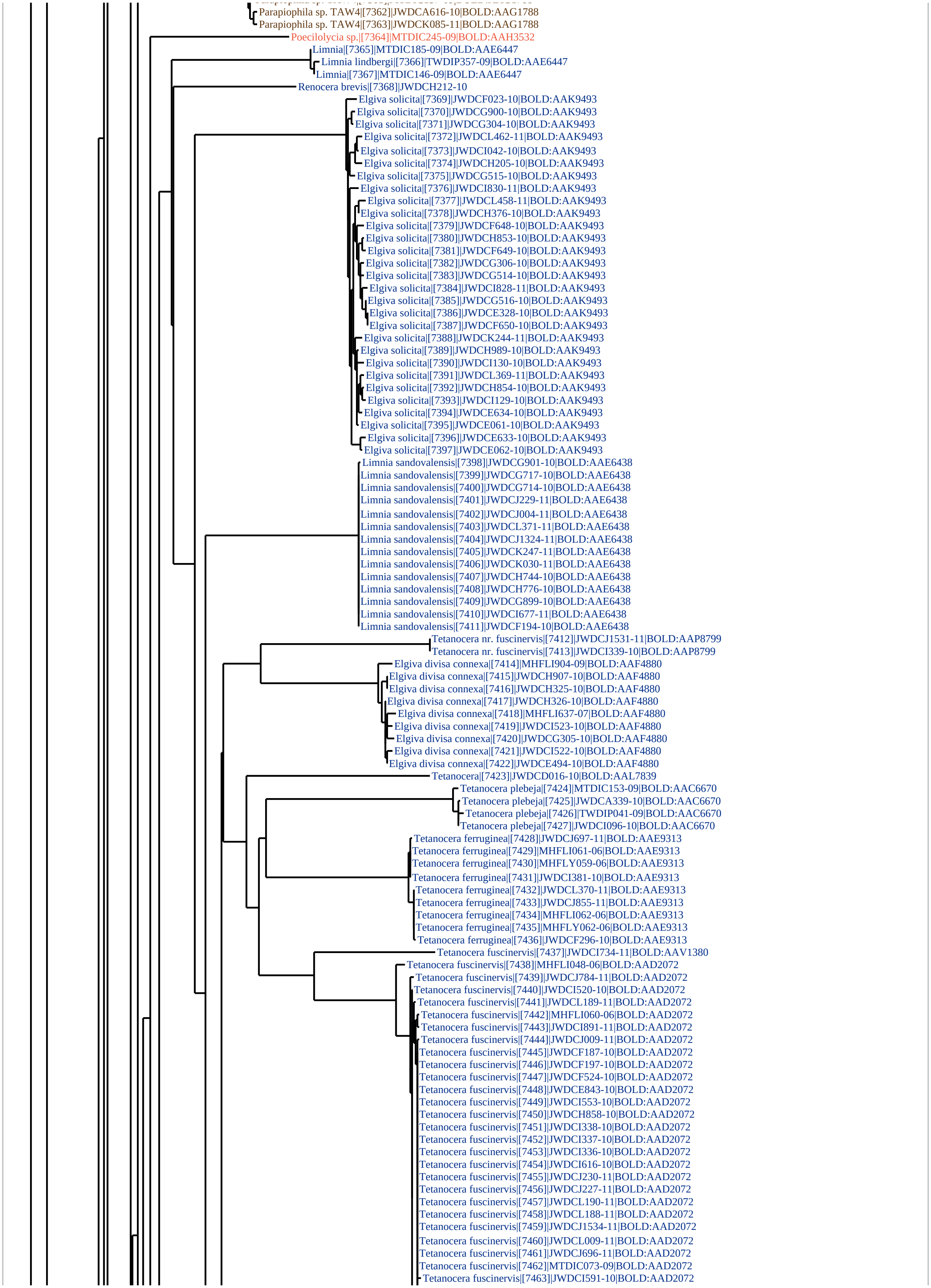

Parapiophila sp. TAW4|[7362]|JWDCA616-10|BOLD:AAG1788
Parapiophila sp. TAW4|[7363]|JWDCK085-11|BOLD:AAG1788
Poecilolycia sp.|[7364]|MTDIC245-09|BOLD:AAH3532
Limnia|[7365]|MTDIC185-09|BOLD:AAE6447
Limnia lindbergi|[7366]|TWDIP357-09|BOLD:AAE6447
Limnia|[7367]|MTDIC146-09|BOLD:AAE6447
Renocera brevis|[7368]|JWDCH212-10
Elgiva solicita|[7369]|JWDCF023-10|BOLD:AAK9493
Elgiva solicita|[7370]|JWDCG900-10|BOLD:AAK9493
Elgiva solicita|[7371]|JWDCG304-10|BOLD:AAK9493
Elgiva solicita|[7372]|JWDCL462-11|BOLD:AAK9493
Elgiva solicita|[7373]|JWDCI042-10|BOLD:AAK9493
Elgiva solicita|[7374]|JWDCH205-10|BOLD:AAK9493
Elgiva solicita|[7375]|JWDCG515-10|BOLD:AAK9493
Elgiva solicita|[7376]|JWDCI830-11|BOLD:AAK9493
Elgiva solicita|[7377]|JWDCL458-11|BOLD:AAK9493
Elgiva solicita|[7378]|JWDCH376-10|BOLD:AAK9493
Elgiva solicita|[7379]|JWDCF648-10|BOLD:AAK9493
Elgiva solicita|[7380]|JWDCH853-10|BOLD:AAK9493
Elgiva solicita|[7381]|JWDCF649-10|BOLD:AAK9493
Elgiva solicita|[7382]|JWDCG306-10|BOLD:AAK9493
Elgiva solicita|[7383]|JWDCG514-10|BOLD:AAK9493
Elgiva solicita|[7384]|JWDCI828-11|BOLD:AAK9493
Elgiva solicita|[7385]|JWDCG516-10|BOLD:AAK9493
Elgiva solicita|[7386]|JWDCE328-10|BOLD:AAK9493
Elgiva solicita|[7387]|JWDCF650-10|BOLD:AAK9493
Elgiva solicita|[7388]|JWDCK244-11|BOLD:AAK9493
Elgiva solicita|[7389]|JWDCH989-10|BOLD:AAK9493
Elgiva solicita|[7390]|JWDCI130-10|BOLD:AAK9493
Elgiva solicita|[7391]|JWDCL369-11|BOLD:AAK9493
Elgiva solicita|[7392]|JWDCH854-10|BOLD:AAK9493
Elgiva solicita|[7393]|JWDCI129-10|BOLD:AAK9493
Elgiva solicita|[7394]|JWDCE634-10|BOLD:AAK9493
Elgiva solicita|[7395]|JWDCE061-10|BOLD:AAK9493
Elgiva solicita|[7396]|JWDCE633-10|BOLD:AAK9493
Elgiva solicita|[7397]|JWDCE062-10|BOLD:AAK9493
Limnia sandovalensis|[7398]|JWDCG901-10|BOLD:AAE6438
Limnia sandovalensis|[7399]|JWDCG717-10|BOLD:AAE6438
Limnia sandovalensis|[7400]|JWDCG714-10|BOLD:AAE6438
Limnia sandovalensis|[7401]|JWDCJ229-11|BOLD:AAE6438
Limnia sandovalensis|[7402]|JWDCJ004-11|BOLD:AAE6438
Limnia sandovalensis|[7403]|JWDCL371-11|BOLD:AAE6438
Limnia sandovalensis|[7404]|JWDCJ1324-11|BOLD:AAE6438
Limnia sandovalensis|[7405]|JWDCK247-11|BOLD:AAE6438
Limnia sandovalensis|[7406]|JWDCK030-11|BOLD:AAE6438
Limnia sandovalensis|[7407]|JWDCH744-10|BOLD:AAE6438
Limnia sandovalensis|[7408]|JWDCH776-10|BOLD:AAE6438
Limnia sandovalensis|[7409]|JWDCG899-10|BOLD:AAE6438
Limnia sandovalensis|[7410]|JWDCI677-11|BOLD:AAE6438
Limnia sandovalensis|[7411]|JWDCF194-10|BOLD:AAE6438
Tetanocera nr. fuscinervis|[7412]|JWDCJ1531-11|BOLD:AAP8799
Tetanocera nr. fuscinervis|[7413]|JWDCI339-10|BOLD:AAP8799
Elgiva divisa connexa|[7414]|MHFLI904-09|BOLD:AAF4880
Elgiva divisa connexa|[7415]|JWDCH907-10|BOLD:AAF4880
Elgiva divisa connexa|[7416]|JWDCH325-10|BOLD:AAF4880
Elgiva divisa connexa|[7417]|JWDCH326-10|BOLD:AAF4880
Elgiva divisa connexa|[7418]|MHFLI637-07|BOLD:AAF4880
Elgiva divisa connexa|[7419]|JWDCI523-10|BOLD:AAF4880
Elgiva divisa connexa|[7420]|JWDCG305-10|BOLD:AAF4880
Elgiva divisa connexa|[7421]|JWDCI522-10|BOLD:AAF4880
Elgiva divisa connexa|[7422]|JWDCE494-10|BOLD:AAF4880
Tetanocera|[7423]|JWDCD016-10|BOLD:AAL7839
Tetanocera plebeja|[7424]|MTDIC153-09|BOLD:AAC6670
Tetanocera plebeja|[7425]|JWDCA339-10|BOLD:AAC6670
Tetanocera plebeja|[7426]|TWDIP041-09|BOLD:AAC6670
Tetanocera plebeja|[7427]|JWDCI096-10|BOLD:AAC6670
Tetanocera ferruginea|[7428]|JWDCJ697-11|BOLD:AAE9313
Tetanocera ferruginea|[7429]|MHFLI061-06|BOLD:AAE9313
Tetanocera ferruginea|[7430]|MHFLY059-06|BOLD:AAE9313
Tetanocera ferruginea|[7431]|JWDCI381-10|BOLD:AAE9313
Tetanocera ferruginea|[7432]|JWDCL370-11|BOLD:AAE9313
Tetanocera ferruginea|[7433]|JWDCJ855-11|BOLD:AAE9313
Tetanocera ferruginea|[7434]|MHFLI062-06|BOLD:AAE9313
Tetanocera ferruginea|[7435]|MHFLY062-06|BOLD:AAE9313
Tetanocera ferruginea|[7436]|JWDCF296-10|BOLD:AAE9313
Tetanocera fuscinervis|[7437]|JWDCI734-11|BOLD:AAV1380
Tetanocera fuscinervis|[7438]|MHFLI048-06|BOLD:AAD2072
Tetanocera fuscinervis|[7439]|JWDCJ784-11|BOLD:AAD2072
Tetanocera fuscinervis|[7440]|JWDCI520-10|BOLD:AAD2072
Tetanocera fuscinervis|[7441]|JWDCL189-11|BOLD:AAD2072
Tetanocera fuscinervis|[7442]|MHFLI060-06|BOLD:AAD2072
Tetanocera fuscinervis|[7443]|JWDCI891-11|BOLD:AAD2072
Tetanocera fuscinervis|[7444]|JWDCJ009-11|BOLD:AAD2072
Tetanocera fuscinervis|[7445]|JWDCF187-10|BOLD:AAD2072
Tetanocera fuscinervis|[7446]|JWDCF197-10|BOLD:AAD2072
Tetanocera fuscinervis|[7447]|JWDCF524-10|BOLD:AAD2072
Tetanocera fuscinervis|[7448]|JWDCE843-10|BOLD:AAD2072
Tetanocera fuscinervis|[7449]|JWDCI553-10|BOLD:AAD2072
Tetanocera fuscinervis|[7450]|JWDCH858-10|BOLD:AAD2072
Tetanocera fuscinervis|[7451]|JWDCI338-10|BOLD:AAD2072
Tetanocera fuscinervis|[7452]|JWDCI337-10|BOLD:AAD2072
Tetanocera fuscinervis|[7453]|JWDCI336-10|BOLD:AAD2072
Tetanocera fuscinervis|[7454]|JWDCI616-10|BOLD:AAD2072
Tetanocera fuscinervis|[7455]|JWDCJ230-11|BOLD:AAD2072
Tetanocera fuscinervis|[7456]|JWDCJ227-11|BOLD:AAD2072
Tetanocera fuscinervis|[7457]|JWDCL190-11|BOLD:AAD2072
Tetanocera fuscinervis|[7458]|JWDCL188-11|BOLD:AAD2072
Tetanocera fuscinervis|[7459]|JWDCJ1534-11|BOLD:AAD2072
Tetanocera fuscinervis|[7460]|JWDCL009-11|BOLD:AAD2072
Tetanocera fuscinervis|[7461]|JWDCJ696-11|BOLD:AAD2072
Tetanocera fuscinervis|[7462]|MTDIC073-09|BOLD:AAD2072
Tetanocera fuscinervis|[7463]|JWDCI591-10|BOLD:AAD2072

Tetanocera fuscinervis|[7461]|JWDCJ696-11|BOLD:AAD2072
Tetanocera fuscinervis|[7462]|MTDIC073-09|BOLD:AAD2072
Tetanocera fuscinervis|[7463]|JWDCI591-10|BOLD:AAD2072
Tetanocera fuscinervis|[7464]|JWDCI382-10|BOLD:AAD2072
Tetanocera fuscinervis|[7465]|JWDCF726-10|BOLD:AAD2072
Tetanocera fuscinervis|[7466]|JWDCJ008-11|BOLD:AAD2072
Tetanocera fuscinervis|[7467]|JWDCI589-10|BOLD:AAD2072
Tetanocera fuscinervis|[7468]|JWDCJ782-11|BOLD:AAD2072
Tetanocera fuscinervis|[7469]|JWDCJ1327-11|BOLD:AAD2072
Tetanocera fuscinervis|[7470]|JWDCG719-10|BOLD:AAD2072
Tetanocera fuscinervis|[7471]|JWDCI131-10|BOLD:AAD2072
Tetanocera fuscinervis|[7472]|JWDCI735-11|BOLD:AAD2072
Tetanocera fuscinervis|[7473]|JWDCI733-11|BOLD:AAD2072
Tetanocera fuscinervis|[7474]|JWDCI590-10|BOLD:AAD2072
Tetanocera fuscinervis|[7475]|JWDCF593-10|BOLD:AAD2072
Tetanocera fuscinervis|[7476]|JWDCF103-10|BOLD:AAD2072
Tetanocera fuscinervis|[7477]|JWDCF756-10|BOLD:AAD2072
Tetanocera fuscinervis|[7478]|JWDCJ695-11|BOLD:AAD2072
Tetanocera fuscinervis|[7479]|JWDCK026-11|BOLD:AAD2072
Tetanocera fuscinervis|[7480]|JWDCK033-11|BOLD:AAD2072
Tetanocera fuscinervis|[7481]|JWDCJ1322-11|BOLD:AAD2072
Tetanocera fuscinervis|[7482]|JWDCJ1325-11|BOLD:AAD2072
Tetanocera fuscinervis|[7483]|JWDCJ1326-11|BOLD:AAD2072
Tetanocera fuscinervis|[7484]|JWDCL460-11|BOLD:AAD2072
Tetanocera fuscinervis|[7485]|JWDCL567-11|BOLD:AAD2072
Tetanocera fuscinervis|[7486]|JWDCJ158-11|BOLD:AAD2072
Tetanocera fuscinervis|[7487]|JWDCI044-10|BOLD:AAD2072
Tetanocera fuscinervis|[7488]|JWDCI667-11|BOLD:AAD2072
Tetanocera fuscinervis|[7489]|JWDCH210-10|BOLD:AAD2072
Tetanocera fuscinervis|[7490]|JWDCE506-10|BOLD:AAD2072
Tetanocera fuscinervis|[7491]|JWDCF294-10|BOLD:AAD2072
Tetanocera fuscinervis|[7492]|JWDCF312-10|BOLD:AAD2072
Tetanocera fuscinervis|[7493]|JWDCF447-10|BOLD:AAD2072
Tetanocera fuscinervis|[7494]|JWDCF749-10|BOLD:AAD2072
Tetanocera fuscinervis|[7495]|JWDCF764-10|BOLD:AAD2072
Tetanocera fuscinervis|[7496]|JWDCF102-10|BOLD:AAD2072
Hedria mixta|[7497]|JWDCI132-10|BOLD:AAI2043
Tetanocera robusta|[7498]|JWDCH951-10|BOLD:AAG7434
Tetanocera robusta|[7499]|JWDCK246-11|BOLD:AAG7434
Tetanocera robusta|[7500]|JWDCH446-10|BOLD:AAG7434
Tetanocera robusta|[7501]|JWDCC886-10|BOLD:AAG7434
Tetanocera robusta|[7502]|MHFLI191-06|BOLD:AAG7434
Tetanocera robusta|[7503]|JWDCJ085-11|BOLD:AAG7434
Tetanocera robusta|[7504]|JWDCH953-10|BOLD:AAG7434
Tetanocera robusta|[7505]|JWDCG716-10|BOLD:AAG7434
Tetanocera robusta|[7506]|JWDCK183-11|BOLD:AAG7434
Tetanocera robusta|[7507]|JWDCA289-10|BOLD:AAG7434
Tetanocera robusta|[7508]|JWDCJ1571-11|BOLD:AAG7434
Tetanocera robusta|[7509]|JWDCG669-10|BOLD:AAG7434
Tetanocera robusta|[7510]|JWDCH952-10|BOLD:AAG7434
Tetanocera robusta|[7511]|JWDCH987-10|BOLD:AAG7434
Tetanocera robusta|[7512]|JWDCI169-10|BOLD:AAG7434
Tetanocera robusta|[7513]|JWDCI341-10|BOLD:AAG7434
Tetanocera robusta|[7514]|JWDCI552-10|BOLD:AAG7434
Tetanocera robusta|[7515]|JWDCH482-10|BOLD:AAG7434
Tetanocera robusta|[7516]|JWDCF297-10|BOLD:AAG7434
Tetanocera robusta|[7517]|JWDCH396-10|BOLD:AAG7434
Tetanocera robusta|[7518]|JWDCF298-10|BOLD:AAG7434
Tetanocera robusta|[7519]|JWDCH395-10|BOLD:AAG7434
Tetanocera robusta|[7520]|JWDCK248-11|BOLD:AAG7434
Tetanocera robusta|[7521]|TWDIP043-09|BOLD:AAG7434
Tetanocera robusta|[7522]|JWDCI665-10|BOLD:AAG7434
Tetanocera robusta|[7523]|JWDCI171-10|BOLD:AAG7434
Tetanocera robusta|[7524]|JWDCH394-10|BOLD:AAG7434
Tetanocera robusta|[7525]|JWDCF295-10|BOLD:AAG7434
Tetanocera robusta|[7526]|JWDCF738-10|BOLD:AAG7434
Tetanocera robusta|[7527]|JWDCH481-10|BOLD:AAG7434
Tetanocera robusta|[7528]|JWDCF796-10|BOLD:AAG7434
Tetanocera robusta|[7529]|JWDCJ1478-11|BOLD:AAG7435
Tetanocera robusta|[7530]|MHFLI271-06|BOLD:AAG7435
Tetanocera robusta|[7531]|JWDCG668-10|BOLD:AAG7435
Tetanocera robusta|[7532]|JWDCF832-10|BOLD:AAG7435
Tetanocera robusta|[7533]|JWDCH852-10|BOLD:AAG7435
Tetanocera robusta|[7534]|JWDCF885-10|BOLD:AAG7435
Tetanocera|[7535]|MHFLI038-06|BOLD:AAB7398
Tetanocera|[7536]|MHFLI045-06|BOLD:AAB7398
Tetanocera|[7537]|MHFLI041-06|BOLD:AAB7398
Tetanocera|[7538]|MHFLI039-06|BOLD:AAB7398
Tetanocera|[7539]|MHFLI037-06|BOLD:AAB7398
Tetanocera|[7540]|MHFLI218-06|BOLD:AAB7398
Tetanocera nr. annae|[7541]|MHFLI216-06|BOLD:AAB7398
Tetanocera|[7542]|MHFLI043-06|BOLD:AAB7398
Tetanocera|[7543]|MHFLI047-06|BOLD:AAB7398
Tetanocera|[7544]|MHFLI215-06|BOLD:AAB7398
Tetanocera|[7545]|MHFLI046-06|BOLD:AAB7398
Tetanocera|[7546]|MHFLI036-06|BOLD:AAB7398
Tetanocera nr. annae|[7547]|MHFLI217-06|BOLD:AAB7398
Tetanocera|[7548]|MHFLI222-06|BOLD:AAB7398
Tetanocera|[7549]|MHFLI034-06|BOLD:AAB7398
Tetanocera|[7550]|MHFLI035-06|BOLD:AAB7398
Tetanocera|[7551]|MHFLI219-06|BOLD:AAB7398
Tetanocera|[7552]|MMADL076-09|BOLD:AAB7398
Tetanocera montana|[7553]|DSCNI049-07|BOLD:AAF2065
Tetanocera montana|[7554]|JWDCK027-11|BOLD:AAF2065
Tetanocera montana|[7555]|JWDCK031-11|BOLD:AAF2065
Tetanocera montana|[7556]|JWDCK034-11|BOLD:AAF2065
Tetanocera montana|[7557]|JWDCJ1392-11|BOLD:AAF2065
Tetanocera montana|[7558]|JWDCL117-11|BOLD:AAF2065
Tetanocera montana|[7559]|JWDCL118-11|BOLD:AAF2065
Tetanocera montana|[7560]|JWDCL596-11|BOLD:AAF2065
Tetanocera montana|[7561]|JWDCL368-11|BOLD:AAF2065
Tetanocera montana|[7562]|JWDCH743-10|BOLD:AAF2065
Tetanocera montana|[7563]|JWDCH988-10|BOLD:AAF2065

Tetanocera montana|[7561]|JWDCL368-11|BOLD:AAF2065
Tetanocera montana|[7562]|JWDCH743-10|BOLD:AAF2065
Tetanocera montana|[7563]|JWDCH988-10|BOLD:AAF2065
Tetanocera montana|[7564]|JWDCI133-10|BOLD:AAF2065
Tetanocera montana|[7565]|JWDCI525-10|BOLD:AAF2065
Tetanocera montana|[7566]|JWDCI731-11|BOLD:AAF2065
Tetanocera montana|[7567]|JWDCH447-10|BOLD:AAF2065
Tetanocera montana|[7568]|JWDCH448-10|BOLD:AAF2065
Tetanocera montana|[7569]|JWDCH480-10|BOLD:AAF2065
Tetanocera sp. TAW13|[7570]|MTDIC266-09|BOLD:AAD2073
Tetanocera sp. TAW13|[7571]|MTDIC021-09|BOLD:AAD2073
Tetanocera sp. TAW13|[7572]|JWDCI170-10|BOLD:AAD2073
Tetanocera sp. TAW13|[7573]|MHFLI923-09|BOLD:AAD2073
Tetanocera sp. TAW13|[7574]|JWDCH174-10|BOLD:AAD2073
Tetanocera sp. TAW13|[7575]|MHFLI050-06|BOLD:AAD2073
Tetanocera sp. TAW13|[7576]|JWDCJ943-11|BOLD:AAD2073
Tetanocera sp. TAW13|[7577]|JWDCF763-10|BOLD:AAD2073
Tetanocera silvatica|[7578]|JWDCJ1162-11|BOLD:AAL7840
Tetanocera silvatica|[7579]|JWDCD054-10|BOLD:AAL7840
Tetanocera silvatica|[7580]|JWDCL313-11|BOLD:AAL7840
Tetanocera silvatica|[7581]|JWDCJ159-11|BOLD:AAL7840
Tetanocera silvatica|[7582]|JWDCH906-10|BOLD:ACR6973
Tetanocera silvatica|[7583]|JWDCA331-10|BOLD:AAD2077
Tetanocera silvatica|[7584]|JWDCJ1569-11|BOLD:AAD2077
Tetanocera silvatica|[7585]|TWDIP040-09|BOLD:AAD2077
Tetanocera silvatica|[7586]|JWDCA288-10|BOLD:AAD2077
Tetanocera silvatica|[7587]|JWDCL050-11|BOLD:AAD2077
Tetanocera silvatica|[7588]|JWDCJ007-11|BOLD:AAD2077
Tetanocera silvatica|[7589]|JWDCG905-10|BOLD:AAD2077
Tetanocera silvatica|[7590]|JWDCG670-10|BOLD:AAD2077
Tetanocera silvatica|[7591]|JWDCB090-10|BOLD:AAD2077
Tetanocera silvatica|[7592]|JWDCK245-11|BOLD:AAD2077
Tetanocera silvatica|[7593]|JWDCG642-10|BOLD:AAD2077
Tetanocera silvatica|[7594]|JWDCJ783-11|BOLD:AAD2077
Tetanocera silvatica|[7595]|JWDCK243-11|BOLD:AAD2077
Tetanocera silvatica|[7596]|JSFLA007-08|BOLD:AAD2077
Tetanocera silvatica|[7597]|JWDCI335-10|BOLD:AAD2077
Tetanocera silvatica|[7598]|JWDCL315-11|BOLD:AAD2077
Tetanocera silvatica|[7599]|JWDCJ084-11|BOLD:AAD2077
Tetanocera silvatica|[7600]|JWDCG641-10|BOLD:AAD2077
Tetanocera silvatica|[7601]|JWDCH745-10|BOLD:AAD2077
Tetanocera silvatica|[7602]|JWDCJ1003-11|BOLD:AAD2077
Tetanocera silvatica|[7603]|JWDCL250-11|BOLD:AAD2077
Tetanocera silvatica|[7604]|JWDCJ910-11|BOLD:AAD2077
Tetanocera silvatica|[7605]|JWDCC195-10|BOLD:AAD2077
Tetanocera silvatica|[7606]|JWDCD116-10|BOLD:AAD2077
Tetanocera silvatica|[7607]|JWDCA333-10|BOLD:AAD2077
Tetanocera silvatica|[7608]|JWDCA303-10|BOLD:AAD2077
Tetanocera silvatica|[7609]|JWDCA291-10|BOLD:AAD2077
Tetanocera silvatica|[7610]|JSFLA009-08|BOLD:AAD2077
Tetanocera silvatica|[7611]|JWDCJ1529-11|BOLD:AAD2077
Tetanocera silvatica|[7612]|JWDCD019-10|BOLD:AAD2077
Tetanocera silvatica|[7613]|JWDCD068-10|BOLD:AAD2077
Tetanocera silvatica|[7614]|JWDCJ942-11|BOLD:AAD2077
Tetanocera silvatica|[7615]|JWDCJ944-11|BOLD:AAD2077
Tetanocera silvatica|[7616]|JWDCJ1570-11|BOLD:AAD2077
Tetanocera silvatica|[7617]|JWDCG903-10|BOLD:AAD2077
Tetanocera silvatica|[7618]|JWDCE505-10|BOLD:AAD2077
Dictya sp. TAW1|[7619]|JWDCJ781-11|BOLD:ACZ5284
Dictya sp. TAW1|[7620]|JWDCF193-10|BOLD:ACZ5284
Dictya sp. TAW1|[7621]|JWDCI230-10|BOLD:ACZ5284
Dictya sp. TAW1|[7622]|JWDCH445-10|BOLD:ACZ5284
Dictya sp. TAW1|[7623]|MTDIC264-09|BOLD:ACZ5284
Dictya sp. TAW1|[7624]|JWDCI041-10|BOLD:ACZ5284
Dictya sp. TAW1|[7625]|JWDCF125-10|BOLD:ACZ5284
Dictya sp. TAW1|[7626]|JWDCE514-10|BOLD:ACZ5284
Dictya sp. TAW1|[7627]|JWDCF301-10|BOLD:ACZ5284
Dictya sp. TAW1|[7628]|JWDCG902-10|BOLD:ACZ5284
Dictya sp. TAW1|[7629]|JWDCH992-10|BOLD:ACZ5284
Dictya sp. TAW1|[7630]|JWDCH908-10|BOLD:ACZ5284
Dictya sp. TAW1|[7631]|JWDCF196-10|BOLD:ACZ5284
Dictya sp. TAW1|[7632]|JWDCG904-10|BOLD:ACZ5284
Dictya sp. TAW1|[7633]|JWDCJ017-11|BOLD:ACZ5284
Dictya sp. TAW1|[7634]|JWDCH990-10|BOLD:ACZ5284
Dictya sp. TAW1|[7635]|JWDCH035-10|BOLD:ACZ5284
Dictya sp. TAW1|[7636]|JWDCI827-11|BOLD:ACZ5284
Dictya sp. TAW1|[7637]|JWDCF300-10|BOLD:ACZ5284
Dictya sp. TAW1|[7638]|JWDCF797-10|BOLD:ACZ5284
Dictya sp. TAW1|[7639]|JWDCI701-11|BOLD:ACZ5284
Dictya sp. TAW1|[7640]|JWDCF651-10|BOLD:ACZ5284
Dictya sp. TAW1|[7641]|JWDCF652-10|BOLD:ACZ5284
Dictya sp. TAW1|[7642]|JWDCJ006-11|BOLD:ACZ5284
Dictya sp. TAW1|[7643]|JWDCG802-10|BOLD:ACZ5284
Dictya sp. TAW1|[7644]|JWDCI587-10|BOLD:ACZ5284
Dictya sp. TAW1|[7645]|JWDCH991-10|BOLD:ACZ5284
Dictya sp. TAW1|[7646]|JWDCI831-11|BOLD:ACZ5284
Dictya sp. TAW1|[7647]|JWDCF299-10|BOLD:ACZ5284
Dictya sp. TAW1|[7648]|JWDCH909-10|BOLD:ACZ5284
Dictya sp. TAW1|[7649]|JWDCL372-11|BOLD:ACZ5284
Dictya sp. TAW1|[7650]|JWDCF221-10|BOLD:ACZ5284
Dictya sp. TAW1|[7651]|JWDCG718-10|BOLD:ACZ5284
Dictya sp. TAW1|[7652]|JWDCH954-10|BOLD:ACZ5284
Dictya sp. TAW1|[7653]|JWDCF235-10|BOLD:ACZ5284
Tetanocera|[7654]|JWDCI436-10|BOLD:AAV1381
Tetanocera|[7655]|JWDCI437-10|BOLD:AAV1381
Anticheta melanosoma|[7656]|TWDIP042-09|BOLD:AAG6971
Sepedon borealis|[7657]|JWDCJ492-11|BOLD:AAP7698
Sepedon borealis|[7658]|JWDCJ005-11|BOLD:AAP7698
Sepedon borealis|[7659]|JWDCJ082-11|BOLD:AAP7698
Sepedon borealis|[7660]|JWDCF192-10|BOLD:AAP7698
Minettia lupulina|[7661]|JWDCH665-10|BOLD:AAB8787
Minettia lupulina|[7662]|TWDIP210-09|BOLD:AAB8787

Sepedon borealis|[7660]|JWDCF192-10|BOLD:AAP7698
Minettia lupulina|[7661]|JWDCH665-10|BOLD:AAB8787
Minettia lupulina|[7662]|TWDIP210-09|BOLD:AAB8787
Minettia lupulina|[7663]|JWDCB116-10|BOLD:AAB8787
Minettia lupulina|[7664]|JWDCA625-10|BOLD:AAB8787
Minettia lupulina|[7665]|TWDIP212-09|BOLD:AAB8787
Minettia lupulina|[7666]|JWDCG757-10|BOLD:AAB8787
Minettia lupulina|[7667]|JWDCG758-10|BOLD:AAB8787
Minettia lupulina|[7668]|JWDCA613-10|BOLD:AAB8787
Minettia lupulina|[7669]|JWDCH567-10|BOLD:AAB8787
Minettia lupulina|[7670]|JWDCJ1312-11|BOLD:AAB8787
Minettia lupulina|[7671]|MTDIC118-09|BOLD:AAB8787
Minettia lupulina|[7672]|JWDCG055-10|BOLD:AAB8787
Minettia lupulina|[7673]|JWDCH057-10|BOLD:AAB8787
Minettia lupulina|[7674]|JWDCH058-10|BOLD:AAB8787
Minettia lupulina|[7675]|JWDCH059-10|BOLD:AAB8787
Minettia lupulina|[7676]|JWDCH094-10|BOLD:AAB8787
Minettia lupulina|[7677]|JWDCH095-10|BOLD:AAB8787
Minettia lupulina|[7678]|JWDCJ369-11|BOLD:AAB8787
Minettia lupulina|[7679]|JWDCF251-10|BOLD:AAB8787
Heteromyza tarsalis|[7680]|JWDCE454-10|BOLD:AAP4578
Pseudocollinella abhorrens|[7681]|JWDCI385-10|BOLD:AAP4725
Pseudocollinella abhorrens|[7682]|JWDCE337-10|BOLD:AAP4725
Pseudocollinella abhorrens|[7683]|JWDCE575-10|BOLD:AAP4725
Pseudocollinella abhorrens|[7684]|JWDCK226-11|BOLD:AAP4725
Pseudocollinella abhorrens|[7685]|JWDCI357-10|BOLD:AAP4725
Pseudocollinella abhorrens|[7686]|JWDCI386-10|BOLD:AAP4725
Pseudocollinella abhorrens|[7687]|JWDCL483-11|BOLD:AAP4725
Pseudocollinella abhorrens|[7688]|JWDCJ329-11|BOLD:AAP4725
Pseudocollinella caelobata|[7689]|JWDCG161-10|BOLD:AAQ0530
Pseudocollinella arctopellucida|[7690]|JWDCJ341-11|BOLD:AAQ2033
Heteromyza oculata|[7691]|JWDCE455-10|BOLD:AAG0366
Heteromyza oculata|[7692]|JWDCE232-10|BOLD:AAG0366
Borboropsis puberula|[7693]|JWDCJ1090-11|BOLD:AAU6650
Leptocera neofinalis|[7694]|JWDCE260-10|BOLD:AAP6443
Leptocera neofinalis|[7695]|JWDCJ337-11|BOLD:AAP6443
Leptocera neofinalis|[7696]|JWDCG152-10|BOLD:AAF9268
Leptocera neofinalis|[7697]|JWDCH240-10|BOLD:AAF9268
Leptocera neofinalis|[7698]|JWDCJ338-11|BOLD:AAF9268
Leptocera neofinalis|[7699]|JWDCE179-10|BOLD:AAF9268
Leptocera neofinalis|[7700]|JWDCJ330-11|BOLD:AAF9268
Leptocera neofinalis|[7701]|JWDCL528-11|BOLD:AAF9268
Leptocera neofinalis|[7702]|JWDCJ333-11|BOLD:AAF9268
Leptocera neofinalis|[7703]|JWDCE291-10|BOLD:AAF9268
Leptocera neofinalis|[7704]|JWDCJ331-11|BOLD:AAF9268
Leptocera neofinalis|[7705]|JWDCJ334-11|BOLD:AAF9268
Leptocera neofinalis|[7706]|JWDCE279-10|BOLD:AAF9268
Themira minor|[7707]|JWDCH451-10|BOLD:AAD2272
Themira lucida|[7708]|JWDCJ349-11|BOLD:AAD7140
Themira lucida|[7709]|JWDCJ348-11|BOLD:AAD7140
Themira lucida|[7710]|TWDIP066-09|BOLD:AAD7140
Themira lucida|[7711]|MTDIC013-09|BOLD:AAD7140
Themira lucida|[7712]|JWDCA591-10|BOLD:AAD7140
Themira lucida|[7713]|JWDCJ1076-11|BOLD:AAD7140
Themira lucida|[7714]|JWDCJ347-11|BOLD:AAD7140
Themira lucida|[7715]|JWDCJ1099-11|BOLD:AAD7140
Themira lucida|[7716]|JWDCG544-10|BOLD:AAD7140
Themira germanica|[7717]|JWDCJ1442-11|BOLD:AAP8796
Themira germanica|[7718]|JWDCH899-10|BOLD:AAP8796
Themira germanica|[7719]|JWDCH831-10|BOLD:AAP8796
Themira germanica|[7720]|JWDCH220-10|BOLD:AAP8796
Themira germanica|[7721]|JWDCG545-10|BOLD:AAP8796
Themira pusilla|[7722]|JWDCD005-10|BOLD:AAG5641
Themira pusilla|[7723]|JWDCA924-10|BOLD:AAG5641
Themira pusilla|[7724]|JWDCD218-10|BOLD:AAG5641
Themira pusilla|[7725]|JWDCJ170-11|BOLD:AAG5641
Themira pusilla|[7726]|JWDCJ169-11|BOLD:AAG5641
Themira pusilla|[7727]|JWDCJ813-11|BOLD:AAI1006
Themira pusilla|[7728]|MTDIC272-09|BOLD:AAI1006
Themira pusilla|[7729]|JWDCJ1340-11|BOLD:AAI1006
Themira pusilla|[7730]|JWDCH897-10|BOLD:AAI1006
Themira malformans|[7731]|JWDCH484-10|BOLD:AAG2671
Themira malformans|[7732]|MTDIC273-09|BOLD:AAG2671
Themira malformans|[7733]|JWDCH221-10|BOLD:AAG2671
Themira malformans|[7734]|MTDIC067-09|BOLD:AAG2671
Themira malformans|[7735]|JWDCL025-11|BOLD:AAG2671
Themira malformans|[7736]|MTDIC253-09|BOLD:AAG2671
Themira malformans|[7737]|TWDIP062-09|BOLD:AAG2671
Themira malformans|[7738]|TWDIP063-09|BOLD:AAG2671
Themira malformans|[7739]|JWDCA612-10|BOLD:AAG2671
Themira malformans|[7740]|JWDCH898-10|BOLD:AAG2671
Themira malformans|[7741]|MTDIC242-09|BOLD:AAG2671
Themira malformans|[7742]|TWDIP065-09|BOLD:AAG2671
Themira malformans|[7743]|JWDCG565-10|BOLD:AAG2671
Themira malformans|[7744]|MTDIC070-09|BOLD:AAG2671
Themira malformans|[7745]|MTDIC071-09|BOLD:AAG2671
Themira malformans|[7746]|JWDCI661-10|BOLD:AAG2671
Themira malformans|[7747]|JWDCH483-10|BOLD:AAG2671
Themira malformans|[7748]|JWDCH450-10|BOLD:AAG2671
Themira malformans|[7749]|MTDIC058-09|BOLD:AAG2671
Themira malformans|[7750]|MTDIC086-09|BOLD:AAG2671
Themira malformans|[7751]|TWDIP064-09|BOLD:AAG2671
Themira malformans|[7752]|JWDCG566-10|BOLD:AAG2671
Themira malformans|[7753]|JWDCA576-10|BOLD:AAG2671
Themira malformans|[7754]|JWDCH222-10|BOLD:AAG2671
Themira malformans|[7755]|JWDCH449-10|BOLD:AAG2671
Themira malformans|[7756]|JWDCH485-10|BOLD:AAG2671
Themira malformans|[7757]|JWDCF229-10|BOLD:AAG2671
Themira malformans|[7758]|JWDCF751-10|BOLD:AAG2671
Heleomyzidae|[7759]|JWDCA614-10|BOLD:AAG7060
Neoleria prominens|[7760]|JWDCE255-10|BOLD:AAP6808
Neoleria inscripta|[7761]|JWDCJ1196-11|BOLD:AAU6649
Opacifrons convexa|[7762]|JWDCJ332-11|BOLD:AAQ2032

Neoleria prominens|[7760]|JWDCE255-10|BOLD:AAP6808
Neoleria inscripta|[7761]|JWDCJ1196-11|BOLD:AAU6649
Opacifrons convexa|[7762]|JWDCJ332-11|BOLD:AAQ2032
Phthitia quadricercus|[7763]|JWDCH112-10|BOLD:AAQ0529
Sapromyza sp. TAW1|[7764]|JWDCA617-10|BOLD:AAG6931
Lauxania shewelli|[7765]|JWDCD371-10|BOLD:ABZ2891
Lauxania shewelli|[7766]|JWDCB187-10|BOLD:ABZ2891
Lauxania shewelli|[7767]|JWDCF468-10|BOLD:ABZ2891
Lauxania shewelli|[7768]|JWDCD336-10|BOLD:ACF2909
Lauxania shewelli|[7769]|JWDCD219-10|BOLD:ACM8283
Lauxania shewelli|[7770]|MTDIC038-09|BOLD:ACM8283
Lauxania shewelli|[7771]|JWDCD370-10|BOLD:ACM8283
Lauxania shewelli|[7772]|JWDCF467-10|BOLD:ACM8283
Lauxania shewelli|[7773]|MTDIC107-09|BOLD:ACM8283
Lauxania shewelli|[7774]|TWDIP220-09|BOLD:ACM8283
Lauxania shewelli|[7775]|JWDCB189-10|BOLD:ACM8283
Lauxania shewelli|[7776]|JWDCA577-10|BOLD:ACM8283
Lauxania shewelli|[7777]|JWDCA544-10|BOLD:ACM8283
Lauxania shewelli|[7778]|MTDIC229-09|BOLD:ACM8283
Lauxania shewelli|[7779]|MTDIC040-09|BOLD:ACM8283
Lauxania shewelli|[7780]|JWDCD413-10|BOLD:ACM8283
Lauxania shewelli|[7781]|MTDIC050-09|BOLD:ACM8283
Lauxania shewelli|[7782]|JWDCA547-10|BOLD:ACM8283
Lauxania shewelli|[7783]|JWDCG150-10|BOLD:ACM8283
Lauxania shewelli|[7784]|MTDIC039-09|BOLD:ACM8283
Lauxania shewelli|[7785]|JWDCC722-10|BOLD:ACM8283
Lauxania shewelli|[7786]|MTDIC084-09|BOLD:ACM8283
Lauxania shewelli|[7787]|MTDIC158-09|BOLD:ACM8283
Lauxania shewelli|[7788]|MTDIC161-09|BOLD:ACM8283
Lauxania shewelli|[7789]|MTDIC187-09|BOLD:ACM8283
Lauxania shewelli|[7790]|JWDCC029-10|BOLD:ACM8283
Lauxania shewelli|[7791]|JWDCD245-10|BOLD:ACM8283
Lauxania shewelli|[7792]|JWDCA543-10|BOLD:ACM8283
Lauxania shewelli|[7793]|JWDCA545-10|BOLD:ACM8283
Lauxania shewelli|[7794]|JWDCA546-10|BOLD:ACM8283
Lauxania shewelli|[7795]|JWDCD369-10|BOLD:ACM8283
Lauxania shewelli|[7796]|JWDCC477-10|BOLD:ACM8283
Lauxania shewelli|[7797]|JWDCL293-11|BOLD:ACM8283
Lauxania shewelli|[7798]|JWDCD229-10|BOLD:ACM8283
Lauxania shewelli|[7799]|TWDIP219-09|BOLD:ACM8283
Lauxania shewelli|[7800]|MTDIC230-09|BOLD:ACM8283
Lauxania shewelli|[7801]|MTDIC108-09|BOLD:ACM8283
Lauxania shewelli|[7802]|JWDCB190-10|BOLD:ACM8283
Lauxania shewelli|[7803]|JWDCB186-10|BOLD:ACM8283
Lauxania shewelli|[7804]|TWDIP333-09|BOLD:ACM8283
Lauxania shewelli|[7805]|JWDCF574-10|BOLD:ACM8283
Nearcticorpus canadense|[7806]|JWDCD381-10
Spelobia tufta|[7807]|JWDCD141-10|BOLD:AAL7755
Spelobia tufta|[7808]|JWDCF155-10|BOLD:AAL7755
Spelobia tufta|[7809]|JWDCE163-10|BOLD:AAL7755
Spelobia luteilabris|[7810]|JWDCF435-10|BOLD:AAL7752
Spelobia luteilabris|[7811]|JWDCF514-10|BOLD:AAL7752
Spelobia luteilabris|[7812]|JWDCH402-10|BOLD:AAL7752
Spelobia luteilabris|[7813]|JWDCH403-10|BOLD:AAL7752
Spelobia luteilabris|[7814]|JWDCH648-10|BOLD:AAG7305
Spelobia luteilabris|[7815]|JWDCF949-10|BOLD:AAG7305
Spelobia luteilabris|[7816]|JWDCG399-10|BOLD:AAG7305
Spelobia luteilabris|[7817]|CHDIP286-11|BOLD:AAG7305
Leucopis sp. TAW1|[7818]|JWDCA912-10|BOLD:AAG9064
Leucopis sp. TAW1|[7819]|JWDCA932-10|BOLD:AAG9064
Leucopis sp. TAW4|[7820]|TWDIP087-09|BOLD:AAG6973
Leucopis sp. TAW3|[7821]|TWDIP213-09|BOLD:AAG6983
Leucopis sp. TAW3|[7822]|TWDIP214-09|BOLD:AAG6983
Leucopis sp. TAW2|[7823]|JWDCC313-10|BOLD:AAL8098
Pteromicra leucothrix|[7824]|JWDCF854-10|BOLD:AAP1862
Pteromicra leucothrix|[7825]|JWDCJ816-11|BOLD:AAP1862
Pteromicra leucothrix|[7826]|JWDCI852-11|BOLD:AAP1862
Pteromicra leucothrix|[7827]|JWDCH836-10|BOLD:AAP1862
Pteromicra leucothrix|[7828]|JWDCH835-10|BOLD:AAP1862
Pteromicra leucothrix|[7829]|JWDOL949-10|BOLD:AAP1862
Pteromicra leucothrix|[7830]|JWDCI752-11|BOLD:AAP1862
Pteromicra leucothrix|[7831]|JWDCG712-10|BOLD:AAP1862
Pteromicra leucothrix|[7832]|JWDCG723-10|BOLD:AAP1862
Pteromicra leucothrix|[7833]|JWDCG729-10|BOLD:AAP1862
Pteromicra leucothrix|[7834]|JWDOL092-10|BOLD:AAP1862
Pteromicra leucothrix|[7835]|JWDCJ740-11|BOLD:AAP1862
Pteromicra leucothrix|[7836]|JWDCJ273-11|BOLD:AAP1862
Pherbellia argyra|[7837]|JWDCH377-10|BOLD:AAP2681
Pherbellia argyra|[7838]|JWDCE686-10|BOLD:AAP2681
Pherbellia vitalis|[7839]|JWDCJ157-11|BOLD:AAG6845
Pherbellia nana|[7840]|JWDCJ711-11|BOLD:AHC5717
Pherbellia nana|[7841]|JWDCL598-11|BOLD:AHC5717
Pherbellia nana|[7842]|JWDCH747-10|BOLD:AHC5717
Pherbellia nana|[7843]|JWDCI045-10|BOLD:AHC5717
Pherbellia nana|[7844]|JWDCH454-10|BOLD:AHC5717
Pherbellia nana|[7845]|JWDCI702-11|BOLD:AHC5717
Pherbellia nana|[7846]|JWDCI678-11|BOLD:AHC5717
Pherbellia nana|[7847]|JWDCH004-10|BOLD:AHC5717
Pherbellia nana|[7848]|JWDCI135-10|BOLD:AHC5717
Pherbellia nana|[7849]|JWDCH993-10|BOLD:AHC5717
Pherbellia nana|[7850]|JWDCG711-10|BOLD:AHC5717
Pherbellia nana|[7851]|JWDCJ046-11|BOLD:AHC5717
Pherbellia nana|[7852]|JWDCI746-11|BOLD:AHC5717
Pherbellia nana|[7853]|JWDCH113-10|BOLD:AHC5717
Pherbellia nana|[7854]|JWDCH452-10|BOLD:AHC5717
Pherbellia nana|[7855]|JWDCH455-10|BOLD:AHC5717
Pherbellia nana|[7856]|JWDCH476-10|BOLD:AHC5717
Pherbellia nana|[7857]|JWDCF305-10|BOLD:AHC5717
Pherbellia nana|[7858]|JWDCF526-10|BOLD:AHC5717
Pherbellia nana|[7859]|JWDCF304-10|BOLD:AHC5717
Pherbellia schoenherri|[7860]|JWDCH748-10|BOLD:AAG7442
Pherbellia schoenherri|[7861]|JWDCH913-10|BOLD:AAG7442

Pherbellia nana|[7859]|JWDCF304-10|BOLD:AHC5717
Pherbellia schoenherri|[7860]|JWDCH748-10|BOLD:AAG7442
Pherbellia schoenherri|[7861]|JWDCH913-10|BOLD:AAG7442
Pherbellia schoenherri|[7862]|JWDCI334-10|BOLD:AAG7442
Pherbellia schoenherri|[7863]|JWDCE298-10|BOLD:AAG7442
Pherbellia schoenherri|[7864]|JWDCG469-10|BOLD:AAG7442
Pherbellia schoenherri|[7865]|JWDCE508-10|BOLD:AAG7442
Pherbellia schoenherri|[7866]|JWDCE082-10|BOLD:AAG7442
Pherbellia schoenherri|[7867]|JWDCI519-10|BOLD:AAG7442
Pherbellia schoenherri|[7868]|JWDCF124-10|BOLD:AAG7442
Pherbellia schoenherri|[7869]|JWDCE002-10|BOLD:AAG7442
Pherbellia schoenherri|[7870]|JWDCH334-10|BOLD:AAG7442
Pherbellia schoenherri|[7871]|JWDCI521-10|BOLD:AAG7442
Pherbellia schoenherri maculata|[7872]|JWDCH912-10|BOLD:AAG7442
Pherbellia schoenherri maculata|[7873]|JWDCH911-10|BOLD:AAG7442
Pherbellia schoenherri|[7874]|JWDCI043-10|BOLD:AAG7442
Pherbellia schoenherri|[7875]|JWDCH333-10|BOLD:AAG7442
Pherbellia schoenherri|[7876]|JWDCE507-10|BOLD:AAG7442
Pherbellia schoenherri|[7877]|JWDCE226-10|BOLD:AAG7442
Pherbellia|[7878]|JWDCI850-11|BOLD:AAP8802
Pherbellia|[7879]|JWDCI680-11|BOLD:AAP8802
Pherbellia|[7880]|JWDCH857-10|BOLD:AAP8802
Pherbellia sp. TAW04|[7881]|JWDCH856-10|BOLD:AAP8797
Pherbellia sp. TAW04|[7882]|JWDCG724-10|BOLD:AAP8797
Pherbellia sp. TAW04|[7883]|JWDCH937-10|BOLD:AAP8797
Pherbellia sp. TAW04|[7884]|JWDCH378-10|BOLD:AAP8797
Pherbellia sp. TAW04|[7885]|JWDCF017-10|BOLD:AAP8797
Pherbellia sp. TAW04|[7886]|JWDCH936-10|BOLD:AAP8797
Pherbellia|[7887]|JWDCL376-11|BOLD:AAP8797
Pherbellia|[7888]|JWDCL378-11|BOLD:AAP8797
Pherbellia sp. TAW04|[7889]|JWDCL464-11|BOLD:AAP8797
Pherbellia sp. TAW05|[7890]|JWDCJ818-11|BOLD:AAL8105
Pherbellia sp. TAW05|[7891]|JWDCL015-11|BOLD:AAL8105
Pherbellia sp. TAW05|[7892]|JWDCE340-10|BOLD:AAL8105
Pherbellia sp. TAW05|[7893]|JWDCD190-10|BOLD:AAL8105
Sciomyza simplex|[7894]|MHFLI067-06|BOLD:AAH0830
Sciomyza simplex|[7895]|MTDIC213-09|BOLD:AAH0830
Sciomyza simplex|[7896]|MHFLI879-09|BOLD:AAH0830
Sciomyza dryomyzina|[7897]|TWDIP044-09|BOLD:AAH0833
Sciomyza dryomyzina|[7898]|JWDCJ739-11|BOLD:AAH0833
Sciomyza dryomyzina|[7899]|JWDCJ779-11|BOLD:AAH0833
Sciomyza dryomyzina|[7900]|JWDCL595-11|BOLD:AAH0833
Pherbellia phela|[7901]|JWDCH534-10|BOLD:AAP8801
Pherbellia sp. TAW06|[7902]|JWDCE732-10|BOLD:AEH9782
Pherbellia obscura|[7903]|JWDCE036-10|BOLD:AAP1967
Pherbellia anubis|[7904]|JWDCE045-10|BOLD:AAN6420
Pherbellia griseola|[7905]|JWDCI470-10|BOLD:AAP4749
Pherbellia griseola|[7906]|JWDCH453-10|BOLD:AAP4749
Pherbellia griseola|[7907]|JWDCL251-11|BOLD:AAP4749
Pherbellia griseola|[7908]|JWDCI747-11|BOLD:AAP4749
Pherbellia griseola|[7909]|JWDCH399-10|BOLD:AAP4749
Pherbellia griseola|[7910]|JWDCH211-10|BOLD:AAP4749
Pherbellia griseola|[7911]|JWDCG512-10|BOLD:AAP4749
Pherbellia griseola|[7912]|JWDCG513-10|BOLD:AAP4749
Pherbellia fisheri|[7913]|JWDCL408-11|BOLD:AAP1971
Pherbellia fisheri|[7914]|JWDCE078-10|BOLD:AAP1971
Pherbellia fisheri|[7915]|JWDCE343-10|BOLD:AAP1971
Pherbellia fisheri|[7916]|JWDCE093-10|BOLD:AAP1971
Pherbellia fisheri|[7917]|JWDCE076-10|BOLD:AAP1971
Pherbellia fisheri|[7918]|JWDCE372-10|BOLD:AAP1971
Pherbellia fisheri|[7919]|JWDCI340-10|BOLD:AAP1971
Pherbellia fisheri|[7920]|JWDCE553-10|BOLD:AAP1971
Pherbellia fisheri|[7921]|JWDCE932-10|BOLD:AAP1971
Pherbellia fisheri|[7922]|JWDCE094-10|BOLD:AAP1971
Pherbellia fisheri|[7923]|JWDCE218-10|BOLD:AAP1971
Pherbellia fisheri|[7924]|JWDCF198-10|BOLD:AAP1971
Pherbellia fisheri|[7925]|JWDCE077-10|BOLD:AAP1971
Pherbellia fisheri|[7926]|JWDCE207-10|BOLD:AAP1971
Pherbellia fisheri|[7927]|JWDCE456-10|BOLD:AAP1971
Pherbellia fisheri|[7928]|MTDIC082-09|BOLD:AAP1971
Pherbellia fisheri|[7929]|JWDCJ173-11|BOLD:AAP1971
Pteromicra sp. TAW1|[7930]|JWDCJ1606-11|BOLD:AHJ8243
Pteromicra sp. TAW1|[7931]|JWDCL242-11|BOLD:AHJ8243
Pteromicra sp. TAW1|[7932]|JWDCL016-11|BOLD:AHJ8243
Pteromicra sp. TAW1|[7933]|JWDCE453-10|BOLD:AHJ8243
Pteromicra sp. TAW1|[7934]|JWDCJ044-11|BOLD:AHJ8243
Pteromicra sp. TAW1|[7935]|JWDCI751-11|BOLD:AHJ8243
Pteromicra sp. TAW1|[7936]|JWDCJ093-11|BOLD:AHJ8243
Pteromicra glabricula|[7937]|JWDCL193-11|BOLD:AAP4579
Pteromicra|[7938]|JWDCI342-10|BOLD:AAP4579
Pteromicra nr. angustipennis|[7939]|JWDCJ817-11|BOLD:AAP4579
Pteromicra|[7940]|JWDCE605-10|BOLD:AAP4579
Pteromicra nr. angustipennis|[7941]|JWDCJ815-11|BOLD:AAP4579
Pteromicra|[7942]|JWDCJ1605-11|BOLD:AAP4579
Pteromicra|[7943]|JWDCI333-10|BOLD:AAP4579
Pteromicra|[7944]|JWDCH994-10|BOLD:AAP4579
Pteromicra|[7945]|JWDCI693-11|BOLD:AAP4579
Pteromicra|[7946]|JWDCI343-10|BOLD:AAP4579
Pteromicra nr. angustipennis|[7947]|JWDCL215-11|BOLD:AAP4579
Pteromicra|[7948]|JWDCJ274-11|BOLD:AAP4579
Pteromicra|[7949]|JWDCK084-11|BOLD:AAP4579
Pteromicra|[7950]|JWDCJ275-11|BOLD:AAP4579
Pteromicra angustipennis|[7951]|JWDCL511-11|BOLD:AAG0358
Pteromicra angustipennis|[7952]|JWDCL516-11|BOLD:AAG0358
Pteromicra angustipennis|[7953]|JWDCA594-10|BOLD:AAG0358
Drosophila sp. TAW1|[7954]|JWDCA615-10|BOLD:ACF0720
Drosophila sp. TAW1|[7955]|MTDIC171-09|BOLD:AAA2674
Drosophila sp. TAW1|[7956]|JWDCJ537-11|BOLD:AAA2674
Crossopalpus|[7957]|TWDIP178-09
Crossopalpus|[7958]|JWEMP287-10|BOLD:AAF9854
Crossopalpus curvipes|[7959]|JWEMP251-10|BOLD:ABZ0555
Crossopalpus curvipes|[7960]|JWEMP293-10|BOLD:ABZ0555
Crossopalpus setiger|[7961]|JWEMP249-10|BOLD:ACF6928

[illegible]
Crossopalpus curvipes|[7959]|JWEMP251-10|BOLD:ABZ0555
Crossopalpus curvipes|[7960]|JWEMP293-10|BOLD:ABZ0555
Crossopalpus setiger|[7961]|JWEMP249-10|BOLD:ACE6928
Crossopalpus|[7962]|JWDCD157-10|BOLD:AAF9799
Crossopalpus|[7963]|JWDCD159-10|BOLD:AAF9799
Crossopalpus|[7964]|JWDCG400-10|BOLD:AAF9799
Crossopalpus|[7965]|JWDOL757-10|BOLD:AAF9799
Crossopalpus|[7966]|JWDCD156-10|BOLD:AAF9799
Crossopalpus|[7967]|JWDCE565-10|BOLD:AAF9799
Crossopalpus setiger|[7968]|JWDOL772-10|BOLD:AAF9799
Crossopalpus|[7969]|JWEMP262-10|BOLD:AAF9799
Crossopalpus|[7970]|JWEMP289-10|BOLD:AAF9799
Crossopalpus|[7971]|JWDCE032-10|BOLD:AAF9799
Crossopalpus|[7972]|JWDCC572-10|BOLD:AAF9799
Crossopalpus|[7973]|JWDCD086-10|BOLD:AAF9799
Crossopalpus|[7974]|JWEMP250-10|BOLD:AAF9799
Crossopalpus|[7975]|JWEMP320-10|BOLD:AAF9799
Crossopalpus|[7976]|JWDCD154-10|BOLD:AAF9799
Crossopalpus|[7977]|JWDCE030-10|BOLD:AAF9799
Crossopalpus|[7978]|JWDCE578-10|BOLD:AAF9799
Crossopalpus|[7979]|JWDCE296-10|BOLD:AAF9799
Crossopalpus|[7980]|JWDCE311-10|BOLD:AAF9799
Crossopalpus|[7981]|JWDCE584-10|BOLD:AAF9799
Crossopalpus|[7982]|JWDCE312-10|BOLD:AAF9799
Crossopalpus|[7983]|JWDCE281-10|BOLD:AAF9799
Crossopalpus|[7984]|JWDCG267-10|BOLD:AAF9799
Crossopalpus|[7985]|JWDCI550-10|BOLD:AAF9799
Crossopalpus|[7986]|JWDCD155-10|BOLD:AAF9799
Crossopalpus|[7987]|JWDCD153-10|BOLD:AAF9799
Crossopalpus|[7988]|JWDCD152-10|BOLD:AAF9799
Crossopalpus|[7989]|JWDCD158-10|BOLD:AAF9799
Crossopalpus|[7990]|JWDCD003-10|BOLD:AAF9799
Crossopalpus|[7991]|TWDIP177-09|BOLD:AAF9799
Crossopalpus|[7992]|JWDCE564-10|BOLD:AAF9799
Crossopalpus|[7993]|JWDCE112-10|BOLD:AAF9799
Crossopalpus|[7994]|JWDCE213-10|BOLD:AAF9799
Crossopalpus|[7995]|JWDOL1061-10|BOLD:AAF9799
Crossopalpus|[7996]|JWDCE214-10|BOLD:AAF9799
Crossopalpus|[7997]|JWDCE220-10|BOLD:AAF9799
Crossopalpus|[7998]|JWDCE110-10|BOLD:AAF9799
Crossopalpus|[7999]|JWDCE111-10|BOLD:AAF9799
Crossopalpus|[8000]|JWDCE031-10|BOLD:AAF9799
Crossopalpus|[8001]|JWDCE043-10|BOLD:AAF9799
Crossopalpus|[8002]|JWDCE238-10|BOLD:AAF9799
Crossopalpus|[8003]|JWDCE267-10|BOLD:AAF9799
Crossopalpus setiger|[8004]|JWEMP148-10|BOLD:AAF9799
Crossopalpus|[8005]|JWEMP150-10|BOLD:AAF9799
Crossopalpus|[8006]|JWEMP247-10|BOLD:AAF9799
Crossopalpus|[8007]|JWEMP248-10|BOLD:AAF9799
Crossopalpus|[8008]|JWEMP077-10|BOLD:AAF9799
Crossopalpus|[8009]|JWEMP080-10|BOLD:AAF9799
Crossopalpus|[8010]|JWEMP040-10|BOLD:AAF9799
Crossopalpus|[8011]|JWEMP041-10|BOLD:AAF9799
Crossopalpus|[8012]|JWDOL1086-10|BOLD:AAF9799
Crossopalpus|[8013]|JWDOL1087-10|BOLD:AAF9799
Crossopalpus|[8014]|JWDOL1072-10|BOLD:AAF9799
Crossopalpus|[8015]|JWDOL1085-10|BOLD:AAF9799
Crossopalpus|[8016]|JWEMP255-10|BOLD:AAF9799
Crossopalpus|[8017]|JWEMP263-10|BOLD:AAF9799
Crossopalpus|[8018]|JWEMP271-10|BOLD:AAF9799
Crossopalpus|[8019]|JWEMP290-10|BOLD:AAF9799
Crossopalpus|[8020]|JWEMP291-10|BOLD:AAF9799
Crossopalpus|[8021]|JWDCA098-10|BOLD:AAF9799
Carexomyza marshalli|[8022]|JWDCL665-11|BOLD:AAQ3561
Carexomyza marshalli|[8023]|JWDCL481-11|BOLD:AAQ3561
Arganthomyza duplex|[8024]|JWDCH668-10|BOLD:AAG6938
Arganthomyza duplex|[8025]|JWDCH670-10|BOLD:AAG6938
Arganthomyza duplex|[8026]|JWDCH671-10|BOLD:AAG6938
Arganthomyza disjuncta|[8027]|JWDCJ139-11|BOLD:AAP8841
Arganthomyza disjuncta|[8028]|JWDCL533-11|BOLD:AAP8841
Arganthomyza disjuncta|[8029]|JWDCL534-11|BOLD:AAP8841
Arganthomyza disjuncta|[8030]|JWDCL535-11|BOLD:AAP8841
Arganthomyza disjuncta|[8031]|JWDCH810-10|BOLD:AAP8841
Anthomyza gilviventris|[8032]|JWDCJ172-11|BOLD:AAG4829
Anthomyza gilviventris|[8033]|JWDCL664-11|BOLD:AAG4829
Anthomyza gilviventris|[8034]|JWDCH020-10|BOLD:AAG4829
Anthomyza gilviventris|[8035]|JWDCJ1444-11|BOLD:AAG4829
Anthomyza gilviventris|[8036]|JWDCL662-11|BOLD:AAG4829
Anthomyza gilviventris|[8037]|JWDCL663-11|BOLD:AAG4829
Anthomyza gilviventris|[8038]|JWDCK227-11|BOLD:AAG4829
Anthomyza gilviventris|[8039]|JWDCK228-11|BOLD:AAG4829
Anthomyza gilviventris|[8040]|JWDCJ1443-11|BOLD:AAG4829
Anthomyza gilviventris|[8041]|JWDCL241-11|BOLD:AAG4829
Anthomyza gilviventris|[8042]|JWDCH839-10|BOLD:AAG4829
Anthomyza gilviventris|[8043]|JWDCI851-11|BOLD:AAG4829
Anthomyza gilviventris|[8044]|JWDCH466-10|BOLD:AAG4829
Anthomyza gilviventris|[8045]|JWDCI854-11|BOLD:AAG4829
Anthomyza gilviventris|[8046]|JWDCI855-11|BOLD:AAG4829
Anthomyza gilviventris|[8047]|JWDCH837-10|BOLD:AAG4829
Anthomyza gilviventris|[8048]|JWDCH242-10|BOLD:AAG4829
Anthomyza pengellyi|[8049]|JWDCH669-10|BOLD:AAL8100
Anthomyza pengellyi|[8050]|JWDCJ353-11|BOLD:AAL8100
Anthomyza pengellyi|[8051]|JWDCC720-10|BOLD:AAL8100
Cnodacophora nasoni|[8052]|JWDCJ1079-11|BOLD:AAP8873
Cnodacophora nasoni|[8053]|JWDCJ1092-11|BOLD:AAP8873
Cnodacophora nasoni|[8054]|JWDCJ1077-11|BOLD:AAP8873
Cnodacophora nasoni|[8055]|JWDCJ1078-11|BOLD:AAP8873
Cnodacophora nasoni|[8056]|JWDCG337-10|BOLD:AAP8873
Cnodacophora nasoni|[8057]|JWDCJ1093-11|BOLD:AAP8873
Cnodacophora nasoni|[8058]|JWDCJ1091-11|BOLD:AAP8873
Cnodacophora nasoni|[8059]|JWDCL486-11|BOLD:AAP8873
Cnodacophora nasoni|[8060]|JWDCL536-11|BOLD:AAP8873
[illegible]

Cnodacophora nasoni|[8058]|JWDCJ1091-11|BOLD:AAP8873
Cnodacophora nasoni|[8059]|JWDCL486-11|BOLD:AAP8873
Cnodacophora nasoni|[8060]|JWDCL536-11|BOLD:AAP8873
Cnodacophora nasoni|[8061]|JWDCG338-10|BOLD:AAP8873
Compsobata pallipes|[8062]|CHDIP835-12|BOLD:AAF8557
Pseudopachychaeta ruficeps|[8063]|JWCHL331-10|BOLD:AAL7391
Pseudopachychaeta ruficeps|[8064]|JWCHL352-10|BOLD:AAL7391
Pseudopachychaeta ruficeps|[8065]|JWCHL258-10|BOLD:AAL7391
Pseudopachychaeta ruficeps|[8066]|JWCHL242-10|BOLD:AAL7391
Pseudopachychaeta ruficeps|[8067]|JWCHL241-10|BOLD:AAL7391
Pseudopachychaeta ruficeps|[8068]|JWCHL211-10|BOLD:AAL7391
Pseudopachychaeta ruficeps|[8069]|JWCHL207-10|BOLD:AAL7391
Pseudopachychaeta ruficeps|[8070]|JWCHL202-10|BOLD:AAL7391
Pseudopachychaeta ruficeps|[8071]|JWCHL196-10|BOLD:AAL7391
Pseudopachychaeta ruficeps|[8072]|JWCHL174-10|BOLD:AAL7391
Pseudopachychaeta ruficeps|[8073]|JWCHL171-10|BOLD:AAL7391
Pseudopachychaeta ruficeps|[8074]|JWDCH464-10|BOLD:AAL7391
Pseudopachychaeta ruficeps|[8075]|JWCHL260-10|BOLD:AAL7391
Pseudopachychaeta ruficeps|[8076]|JWCHL273-10|BOLD:AAL7391
Pseudopachychaeta ruficeps|[8077]|JWCHL282-10|BOLD:AAL7391
Pseudopachychaeta ruficeps|[8078]|JWCHL301-10|BOLD:AAL7391
Pseudopachychaeta ruficeps|[8079]|JWCHL303-10|BOLD:AAL7391
Pseudopachychaeta ruficeps|[8080]|JWCHL375-10|BOLD:AAL7391
Pseudopachychaeta ruficeps|[8081]|JWCHL508-10|BOLD:AAL7391
Pseudopachychaeta ruficeps|[8082]|JWCHL518-10|BOLD:AAL7391
Pseudopachychaeta ruficeps|[8083]|JWCHL175-10|BOLD:AAL7391
Pseudopachychaeta ruficeps|[8084]|JWCHL210-10|BOLD:AAL7391
Pseudopachychaeta ruficeps|[8085]|JWCHL261-10|BOLD:AAL7391
Pseudopachychaeta ruficeps|[8086]|JWCHL332-10|BOLD:AAL7391
Pseudopachychaeta ruficeps|[8087]|JWCHL354-10|BOLD:AAL7391
Pseudopachychaeta ruficeps|[8088]|JWCHL401-10|BOLD:AAL7391
Pseudopachychaeta ruficeps|[8089]|JWCHL521-10|BOLD:AAL7391
Pseudopachychaeta ruficeps|[8090]|JWDCL479-11|BOLD:AAL7391
Pseudopachychaeta ruficeps|[8091]|JWCHL208-10|BOLD:AAL7391
Pseudopachychaeta ruficeps|[8092]|JWDCH228-10|BOLD:AAL7391
Pseudopachychaeta ruficeps|[8093]|JWCHL353-10|BOLD:AAL7391
Pseudopachychaeta ruficeps|[8094]|JWCHL173-10|BOLD:AAL7391
Pseudopachychaeta ruficeps|[8095]|JWCHL298-10|BOLD:AAL7391
Pseudopachychaeta ruficeps|[8096]|JWCHL297-10|BOLD:AAL7391
Pseudopachychaeta ruficeps|[8097]|JWCHL257-10|BOLD:AAL7391
Pseudopachychaeta ruficeps|[8098]|JWCHL250-10|BOLD:AAL7391
Pseudopachychaeta ruficeps|[8099]|JWCHL203-10|BOLD:AAL7391
Pseudopachychaeta ruficeps|[8100]|JWCHL201-10|BOLD:AAL7391
Pseudopachychaeta ruficeps|[8101]|JWCHL198-10|BOLD:AAL7391
Pseudopachychaeta ruficeps|[8102]|JWCHL176-10|BOLD:AAL7391
Pseudopachychaeta ruficeps|[8103]|JWCHL557-10|BOLD:AAL7391
Pseudopachychaeta ruficeps|[8104]|JWDCH018-10|BOLD:AAL7391
Pseudopachychaeta ruficeps|[8105]|JWCHL244-10|BOLD:AAL7391
Pseudopachychaeta ruficeps|[8106]|JWCHL205-10|BOLD:AAL7391
Pseudopachychaeta ruficeps|[8107]|JWCHL243-10|BOLD:AAL7391
Pseudopachychaeta ruficeps|[8108]|JWCHL328-10|BOLD:AAL7391
Pseudopachychaeta ruficeps|[8109]|JWCHL339-10|BOLD:AAL7391
Pseudopachychaeta ruficeps|[8110]|JWCHL382-10|BOLD:AAL7391
Pseudopachychaeta ruficeps|[8111]|JWCHL510-10|BOLD:AAL7391
Pseudopachychaeta ruficeps|[8112]|JWCHL517-10|BOLD:AAL7391
Pseudopachychaeta ruficeps|[8113]|JWCHL519-10|BOLD:AAL7391
Pseudopachychaeta ruficeps|[8114]|JWCHL487-10|BOLD:AAL7391
Pseudopachychaeta ruficeps|[8115]|JWCHL383-10|BOLD:AAL7391
Pseudopachychaeta ruficeps|[8116]|JWCHL230-10|BOLD:AAL7391
Pseudopachychaeta ruficeps|[8117]|JWDCI052-10|BOLD:AAL7391
Pseudopachychaeta ruficeps|[8118]|JWCHL562-10|BOLD:AAL7391
Pseudopachychaeta ruficeps|[8119]|JWCHL159-10|BOLD:AAL7391
Pseudopachychaeta ruficeps|[8120]|JWCHL197-10|BOLD:AAL7391
Pseudopachychaeta ruficeps|[8121]|JWCHL200-10|BOLD:AAL7391
Pseudopachychaeta ruficeps|[8122]|JWCHL229-10|BOLD:AAL7391
Pseudopachychaeta ruficeps|[8123]|JWCHL238-10|BOLD:AAL7391
Pseudopachychaeta ruficeps|[8124]|JWCHL274-10|BOLD:AAL7391
Pseudopachychaeta ruficeps|[8125]|JWCHL280-10|BOLD:AAL7391
Pseudopachychaeta ruficeps|[8126]|JWCHL300-10|BOLD:AAL7391
Pseudopachychaeta ruficeps|[8127]|JWDCD062-10|BOLD:AAL7391
Pseudopachychaeta ruficeps|[8128]|JWDCL478-11|BOLD:AAL7391
Pseudopachychaeta ruficeps|[8129]|JWCHL302-10|BOLD:AAL7391
Pseudopachychaeta ruficeps|[8130]|JWCHL304-10|BOLD:AAL7391
Pseudopachychaeta ruficeps|[8131]|JWCHL329-10|BOLD:AAL7391
Pseudopachychaeta ruficeps|[8132]|JWCHL330-10|BOLD:AAL7391
Pseudopachychaeta ruficeps|[8133]|JWCHL380-10|BOLD:AAL7391
Pseudopachychaeta ruficeps|[8134]|JWCHL389-10|BOLD:AAL7391
Pseudopachychaeta ruficeps|[8135]|JWCHL486-10|BOLD:AAL7391
Pseudopachychaeta ruficeps|[8136]|JWCHL548-10|BOLD:AAL7391
Meoneura triangularis|[8137]|CHDIP436-11|BOLD:AAH6446
Meoneura triangularis|[8138]|JWDCA943-10|BOLD:AAH6446
Meoneura triangularis|[8139]|JWDCA948-10|BOLD:AAH6446
Meoneura triangularis|[8140]|CHDIP510-12|BOLD:AAH6446
Meoneura triangularis|[8141]|JWDCA941-10|BOLD:AAH6446
Meoneura triangularis|[8142]|JWDCC666-10|BOLD:AAH6446
Meoneura triangularis|[8143]|JWDCA923-10|BOLD:AAH6446
Meoneura triangularis|[8144]|JWDCA940-10|BOLD:AAH6446
Meoneura triangularis|[8145]|CHDIP748-12|BOLD:AAH6446
Siphona multifaria|[8146]|TWDIP252-09|BOLD:AAG2174
Siphona multifaria|[8147]|TWDIP253-09|BOLD:AAG2174
Siphona multifaria|[8148]|TWDIP251-09|BOLD:AAG2174
Siphona multifaria|[8149]|TWDIP254-09|BOLD:AAG2174
Siphona hokkaidensis|[8150]|JWDCB101-10|BOLD:AAG2172
Siphona hokkaidensis|[8151]|JWDCB111-10|BOLD:AAG2172
Siphona hokkaidensis|[8152]|JWDCB113-10|BOLD:AAG2172
Siphona hokkaidensis|[8153]|MTDIC160-09|BOLD:AAG2172
Siphona geniculata|[8154]|MHFLI963-09|BOLD:AAG2108
Siphona geniculata|[8155]|MHFLI673-07|BOLD:AAG2108
Siphona geniculata|[8156]|MHFLI639-07|BOLD:AAG2108
Siphona geniculata|[8157]|MHFLI644-07|BOLD:AAG2108
Siphona geniculata|[8158]|MHFLI936-09|BOLD:AAG2108
Siphona geniculata|[8159]|MHFLI161-06|BOLD:AAG2108
Siphona geniculata|[8160]|MHFLI965-09|BOLD:AAG2108

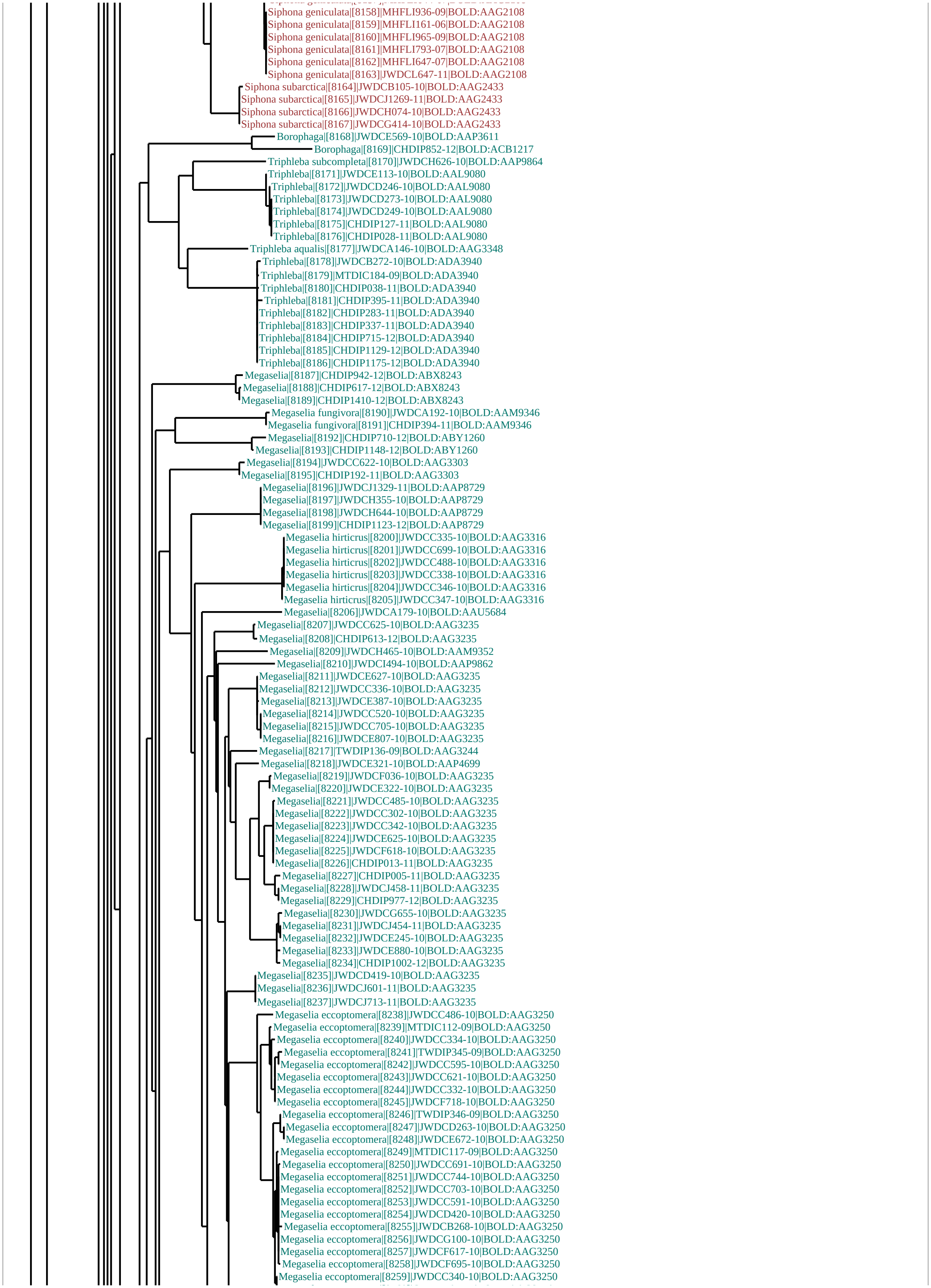
Siphona geniculata|[8158]|MHFLI936-09|BOLD:AAG2108
Siphona geniculata|[8159]|MHFLI161-06|BOLD:AAG2108
Siphona geniculata|[8160]|MHFLI965-09|BOLD:AAG2108
Siphona geniculata|[8161]|MHFLI793-07|BOLD:AAG2108
Siphona geniculata|[8162]|MHFLI647-07|BOLD:AAG2108
Siphona geniculata|[8163]|JWDCL647-11|BOLD:AAG2108
Siphona subarctica|[8164]|JWDCB105-10|BOLD:AAG2433
Siphona subarctica|[8165]|JWDCJ1269-11|BOLD:AAG2433
Siphona subarctica|[8166]|JWDCH074-10|BOLD:AAG2433
Siphona subarctica|[8167]|JWDCG414-10|BOLD:AAG2433
Borophaga|[8168]|JWDCE569-10|BOLD:AAP3611
Borophaga|[8169]|CHDIP852-12|BOLD:ACB1217
Triphleba subcompleta|[8170]|JWDCH626-10|BOLD:AAP9864
Triphleba|[8171]|JWDCE113-10|BOLD:AAL9080
Triphleba|[8172]|JWDCD246-10|BOLD:AAL9080
Triphleba|[8173]|JWDCD273-10|BOLD:AAL9080
Triphleba|[8174]|JWDCD249-10|BOLD:AAL9080
Triphleba|[8175]|CHDIP127-11|BOLD:AAL9080
Triphleba|[8176]|CHDIP028-11|BOLD:AAL9080
Triphleba aqualis|[8177]|JWDCA146-10|BOLD:AAG3348
Triphleba|[8178]|JWDCB272-10|BOLD:ADA3940
Triphleba|[8179]|MTDIC184-09|BOLD:ADA3940
Triphleba|[8180]|CHDIP038-11|BOLD:ADA3940
Triphleba|[8181]|CHDIP395-11|BOLD:ADA3940
Triphleba|[8182]|CHDIP283-11|BOLD:ADA3940
Triphleba|[8183]|CHDIP337-11|BOLD:ADA3940
Triphleba|[8184]|CHDIP715-12|BOLD:ADA3940
Triphleba|[8185]|CHDIP1129-12|BOLD:ADA3940
Triphleba|[8186]|CHDIP1175-12|BOLD:ADA3940
Megaselia|[8187]|CHDIP942-12|BOLD:ABX8243
Megaselia|[8188]|CHDIP617-12|BOLD:ABX8243
Megaselia|[8189]|CHDIP1410-12|BOLD:ABX8243
Megaselia fungivora|[8190]|JWDCA192-10|BOLD:AAM9346
Megaselia fungivora|[8191]|CHDIP394-11|BOLD:AAM9346
Megaselia|[8192]|CHDIP710-12|BOLD:ABY1260
Megaselia|[8193]|CHDIP1148-12|BOLD:ABY1260
Megaselia|[8194]|JWDCC622-10|BOLD:AAG3303
Megaselia|[8195]|CHDIP192-11|BOLD:AAG3303
Megaselia|[8196]|JWDCJ1329-11|BOLD:AAP8729
Megaselia|[8197]|JWDCH355-10|BOLD:AAP8729
Megaselia|[8198]|JWDCH644-10|BOLD:AAP8729
Megaselia|[8199]|CHDIP1123-12|BOLD:AAP8729
Megaselia hirticrus|[8200]|JWDCC335-10|BOLD:AAG3316
Megaselia hirticrus|[8201]|JWDCC699-10|BOLD:AAG3316
Megaselia hirticrus|[8202]|JWDCC488-10|BOLD:AAG3316
Megaselia hirticrus|[8203]|JWDCC338-10|BOLD:AAG3316
Megaselia hirticrus|[8204]|JWDCC346-10|BOLD:AAG3316
Megaselia hirticrus|[8205]|JWDCC347-10|BOLD:AAG3316
Megaselia|[8206]|JWDCA179-10|BOLD:AAU5684
Megaselia|[8207]|JWDCC625-10|BOLD:AAG3235
Megaselia|[8208]|CHDIP613-12|BOLD:AAG3235
Megaselia|[8209]|JWDCH465-10|BOLD:AAM9352
Megaselia|[8210]|JWDCI494-10|BOLD:AAP9862
Megaselia|[8211]|JWDCE627-10|BOLD:AAG3235
Megaselia|[8212]|JWDCC336-10|BOLD:AAG3235
Megaselia|[8213]|JWDCE387-10|BOLD:AAG3235
Megaselia|[8214]|JWDCC520-10|BOLD:AAG3235
Megaselia|[8215]|JWDCC705-10|BOLD:AAG3235
Megaselia|[8216]|JWDCE807-10|BOLD:AAG3235
Megaselia|[8217]|TWDIP136-09|BOLD:AAG3244
Megaselia|[8218]|JWDCE321-10|BOLD:AAP4699
Megaselia|[8219]|JWDCF036-10|BOLD:AAG3235
Megaselia|[8220]|JWDCE322-10|BOLD:AAG3235
Megaselia|[8221]|JWDCC485-10|BOLD:AAG3235
Megaselia|[8222]|JWDCC302-10|BOLD:AAG3235
Megaselia|[8223]|JWDCC342-10|BOLD:AAG3235
Megaselia|[8224]|JWDCE625-10|BOLD:AAG3235
Megaselia|[8225]|JWDCF618-10|BOLD:AAG3235
Megaselia|[8226]|CHDIP013-11|BOLD:AAG3235
Megaselia|[8227]|CHDIP005-11|BOLD:AAG3235
Megaselia|[8228]|JWDCJ458-11|BOLD:AAG3235
Megaselia|[8229]|CHDIP977-12|BOLD:AAG3235
Megaselia|[8230]|JWDCG655-10|BOLD:AAG3235
Megaselia|[8231]|JWDCJ454-11|BOLD:AAG3235
Megaselia|[8232]|JWDCE245-10|BOLD:AAG3235
Megaselia|[8233]|JWDCE880-10|BOLD:AAG3235
Megaselia|[8234]|CHDIP1002-12|BOLD:AAG3235
Megaselia|[8235]|JWDCD419-10|BOLD:AAG3235
Megaselia|[8236]|JWDCJ601-11|BOLD:AAG3235
Megaselia|[8237]|JWDCJ713-11|BOLD:AAG3235
Megaselia eccoptomera|[8238]|JWDCC486-10|BOLD:AAG3250
Megaselia eccoptomera|[8239]|MTDIC112-09|BOLD:AAG3250
Megaselia eccoptomera|[8240]|JWDCC334-10|BOLD:AAG3250
Megaselia eccoptomera|[8241]|TWDIP345-09|BOLD:AAG3250
Megaselia eccoptomera|[8242]|JWDCC595-10|BOLD:AAG3250
Megaselia eccoptomera|[8243]|JWDCC621-10|BOLD:AAG3250
Megaselia eccoptomera|[8244]|JWDCC332-10|BOLD:AAG3250
Megaselia eccoptomera|[8245]|JWDCF718-10|BOLD:AAG3250
Megaselia eccoptomera|[8246]|TWDIP346-09|BOLD:AAG3250
Megaselia eccoptomera|[8247]|JWDCD263-10|BOLD:AAG3250
Megaselia eccoptomera|[8248]|JWDCE672-10|BOLD:AAG3250
Megaselia eccoptomera|[8249]|MTDIC117-09|BOLD:AAG3250
Megaselia eccoptomera|[8250]|JWDCC691-10|BOLD:AAG3250
Megaselia eccoptomera|[8251]|JWDCC744-10|BOLD:AAG3250
Megaselia eccoptomera|[8252]|JWDCC703-10|BOLD:AAG3250
Megaselia eccoptomera|[8253]|JWDCC591-10|BOLD:AAG3250
Megaselia eccoptomera|[8254]|JWDCD420-10|BOLD:AAG3250
Megaselia eccoptomera|[8255]|JWDCB268-10|BOLD:AAG3250
Megaselia eccoptomera|[8256]|JWDCG100-10|BOLD:AAG3250
Megaselia eccoptomera|[8257]|JWDCF617-10|BOLD:AAG3250
Megaselia eccoptomera|[8258]|JWDCF695-10|BOLD:AAG3250
Megaselia eccoptomera|[8259]|JWDCC340-10|BOLD:AAG3250

Megaselia eccoptomera|[8257]|JWDCF617-10|BOLD:AAG3250
Megaselia eccoptomera|[8258]|JWDCF695-10|BOLD:AAG3250
Megaselia eccoptomera|[8259]|JWDCC340-10|BOLD:AAG3250
Megaselia eccoptomera|[8260]|CHDIP204-11|BOLD:AAG3250
Megaselia|[8261]|MTDIC025-09|BOLD:AAG3234
Megaselia|[8262]|JWDCJ285-11|BOLD:AAG3234
Megaselia|[8263]|JWDCH435-10|BOLD:AAG3234
Megaselia|[8264]|JWDCJ1468-11|BOLD:AAG3234
Megaselia|[8265]|JWDCG576-10|BOLD:AAG3234
Megaselia|[8266]|JWDCG359-10|BOLD:AAG3234
Megaselia|[8267]|JWDCH498-10|BOLD:AAG3234
Megaselia|[8268]|CHDIP912-12|BOLD:AAG3234
Megaselia|[8269]|CHDIP913-12|BOLD:AAG3234
Megaselia|[8270]|JWDCB274-10|BOLD:AAG3234
Megaselia|[8271]|JWDCI086-10|BOLD:AAG3234
Megaselia|[8272]|CHDIP391-11|BOLD:AAG3234
Megaselia|[8273]|JWDCH497-10|BOLD:AAG3234
Megaselia|[8274]|JWDCG028-10|BOLD:AAG3234
Megaselia|[8275]|JWDCH505-10|BOLD:AAG3234
Megaselia|[8276]|CHDIP646-12|BOLD:AAG3234
Megaselia|[8277]|JWDCB282-10|BOLD:AAG3234
Megaselia|[8278]|CHDIP621-12|BOLD:AAG3234
Megaselia|[8279]|JWDCF351-10|BOLD:AAG3234
Megaselia|[8280]|JWDCI240-10|BOLD:AAG3234
Megaselia|[8281]|JWDCG255-10|BOLD:AAG3234
Megaselia|[8282]|JWDCG656-10|BOLD:AAG3234
Megaselia|[8283]|JWDCL119-11|BOLD:AAG3234
Megaselia|[8284]|JWDCL032-11|BOLD:AAG3234
Megaselia|[8285]|JWDCL031-11|BOLD:AAG3234
Megaselia|[8286]|JWDCJ935-11|BOLD:AAG3234
Megaselia|[8287]|JWDCK209-11|BOLD:AAG3234
Megaselia|[8288]|CHDIP655-12|BOLD:AAG3234
Megaselia|[8289]|CHDIP1128-12|BOLD:AAG3234
Megaselia|[8290]|JWDCK101-11|BOLD:ADA3078
Megaselia|[8291]|JWDCL221-11|BOLD:AAU5579
Megaselia|[8292]|TWDIP265-09|BOLD:ACB9833
Megaselia|[8293]|JWDCA177-10|BOLD:ACB9833
Megaselia|[8294]|JWDCA182-10|BOLD:ACB9833
Megaselia|[8295]|JWDCD343-10|BOLD:ACB9833
Megaselia|[8296]|JWDCI604-10|BOLD:ACB9833
Megaselia|[8297]|CHDIP417-11|BOLD:ACB9833
Megaselia|[8298]|CHDIP509-12|BOLD:ACB9833
Megaselia|[8299]|JWDCJ1149-11|BOLD:AAG3235
Megaselia|[8300]|JWDCH888-10|BOLD:AAG3235
Megaselia|[8301]|JWDCG866-10|BOLD:AAG3235
Megaselia|[8302]|JWDCI444-10|BOLD:AAG3235
Megaselia|[8303]|JWDCG575-10|BOLD:AAG3235
Megaselia|[8304]|JWDCH959-10|BOLD:AAG3235
Megaselia|[8305]|JWDCG312-10|BOLD:AAG3235
Megaselia|[8306]|JWDCG562-10|BOLD:AAG3235
Megaselia|[8307]|JWDCG561-10|BOLD:AAG3235
Megaselia|[8308]|JWDCH904-10|BOLD:AAG3235
Megaselia|[8309]|JWDCH645-10|BOLD:AAG3235
Megaselia|[8310]|FLIMH011-08|BOLD:AAG3235
Megaselia|[8311]|JWDCA193-10|BOLD:AAG3235
Megaselia|[8312]|JWDCJ1150-11|BOLD:AAG3235
Megaselia|[8313]|JWDCG253-10|BOLD:AAG3235
Megaselia|[8314]|JWDCA171-10|BOLD:AAG3235
Megaselia|[8315]|JWDCA172-10|BOLD:AAG3235
Megaselia|[8316]|JWDCA174-10|BOLD:AAG3235
Megaselia|[8317]|JWDCA180-10|BOLD:AAG3235
Megaselia|[8318]|JWDCA178-10|BOLD:AAG3235
Megaselia|[8319]|CHDIP371-11|BOLD:AAG3235
Megaselia|[8320]|TWDIP140-09|BOLD:AAG3235
Megaselia|[8321]|TWDIP344-09|BOLD:AAG3235
Megaselia|[8322]|JWDCL222-11|BOLD:AAG3235
Megaselia|[8323]|CHDIP374-11|BOLD:AAG3235
Megaselia|[8324]|CHDIP390-11|BOLD:AAG3235
Megaselia|[8325]|CHDIP278-11|BOLD:AAG3235
Megaselia|[8326]|CHDIP554-12|BOLD:AAG3235
Megaselia|[8327]|CHDIP947-12|BOLD:AAG3235
Megaselia|[8328]|JWDCH056-10|BOLD:AAG3235
Megaselia|[8329]|TWDIP141-09|BOLD:AAG3235
Megaselia|[8330]|JWDCJ1228-11|BOLD:AAG3235
Megaselia|[8331]|JWDCA173-10|BOLD:AAG3235
Megaselia|[8332]|JWDCJ1470-11|BOLD:AAG3235
Megaselia|[8333]|JWDCH696-10|BOLD:AAG3235
Megaselia|[8334]|JWDCG460-10|BOLD:AAG3235
Megaselia|[8335]|JWDCH533-10|BOLD:AAG3235
Megaselia|[8336]|JWDCI004-10|BOLD:AAG3235
Megaselia|[8337]|JWDCI071-10|BOLD:AAG3235
Megaselia|[8338]|JWDCI085-10|BOLD:AAG3235
Megaselia|[8339]|JWDCH055-10|BOLD:AAG3235
Megaselia|[8340]|CHDIP593-12|BOLD:AAG3235
Megaselia|[8341]|CHDIP919-12|BOLD:AAG3235
Megaselia|[8342]|JWDCD342-10|BOLD:AAG3235
Megaselia|[8343]|TWDIP133-09|BOLD:AAG3235
Megaselia|[8344]|CHDIP003-11|BOLD:AAG3235
Megaselia|[8345]|CHDIP040-11|BOLD:AAG3235
Megaselia|[8346]|CHDIP052-11|BOLD:AAG3235
Megaselia|[8347]|CHDIP255-11|BOLD:AAG3235
Megaselia|[8348]|CHDIP252-11|BOLD:AAG3235
Megaselia|[8349]|CHDIP245-11|BOLD:AAG3235
Megaselia|[8350]|CHDIP238-11|BOLD:AAG3235
Megaselia|[8351]|JWDCI777-11|BOLD:AAG3235
Megaselia|[8352]|JWDCD341-10|BOLD:AAG3235
Megaselia|[8353]|JWDCD196-10|BOLD:AAG3235
Megaselia|[8354]|JWDCA151-10|BOLD:AAG3235
Megaselia|[8355]|JWDCA149-10|BOLD:AAG3235
Megaselia|[8356]|JWDCA145-10|BOLD:AAG3235
Megaselia|[8357]|TWDIP263-09|BOLD:AAG3235
Megaselia|[8358]|CHDIP948-12|BOLD:AAG3235
Megaselia|[8359]|JWDCK208-11|BOLD:AAG3235

[illegible]
Megaselia|[8357]|TWDIP263-09|BOLD:AAG3235
Megaselia|[8358]|CHDIP948-12|BOLD:AAG3235
Megaselia|[8359]|JWDCK208-11|BOLD:AAG3235
Megaselia|[8360]|JWDCA570-10|BOLD:AAG3235
Megaselia|[8361]|JWDCA195-10|BOLD:AAG3235
Megaselia|[8362]|JWDCA191-10|BOLD:AAG3235
Megaselia|[8363]|TWDIP347-09|BOLD:AAG3235
Megaselia|[8364]|TWDIP132-09|BOLD:AAG3235
Megaselia|[8365]|TWDIP139-09|BOLD:AAG3235
Megaselia|[8366]|MTDIC202-09|BOLD:AAG3235
Megaselia|[8367]|JWDCJ1587-11|BOLD:AAG3235
Megaselia|[8368]|JWDCH695-10|BOLD:AAG3235
Megaselia|[8369]|JWDCI084-10|BOLD:AAG3235
Megaselia|[8370]|CHDIP388-11|BOLD:AAG3235
Megaselia|[8371]|CHDIP333-11|BOLD:AAG3235
Megaselia|[8372]|JWDCB283-10|BOLD:AAG3235
Megaselia|[8373]|CHDIP002-11|BOLD:AAG3235
Megaselia|[8374]|CHDIP164-11|BOLD:AAG3235
Megaselia|[8375]|JWDCI389-10|BOLD:AAG3235
Megaselia|[8376]|JWDCA155-10|BOLD:AAG3235
Megaselia|[8377]|TWDIP225-09|BOLD:AAG3235
Megaselia|[8378]|JWDCA154-10|BOLD:AAG3235
Megaselia|[8379]|CHDIP168-11|BOLD:AAG3235
Megaselia|[8380]|CHDIP1387-12|BOLD:AAG3235
Megaselia|[8381]|CHDIP625-12|BOLD:AAD7101
Megaselia|[8382]|TWDIP224-09|BOLD:AAD7101
Megaselia|[8383]|JWDCJ1471-11|BOLD:AAD7101
Megaselia|[8384]|CHDIP708-12|BOLD:AAD7101
Megaselia brejchaorum|[8385]|JWDCC695-10|BOLD:AAG3313
Megaselia|[8386]|JWDCI305-10|BOLD:AAP8723
Megaselia|[8387]|CHDIP377-11|BOLD:AAP8723
Megaselia|[8388]|CHDIP918-12|BOLD:ABY0262
Megaselia|[8389]|CHDIP699-12|BOLD:ABY0932
Megaselia|[8390]|JWDCC483-10|BOLD:AAL9072
Megaselia wiegmanae|[8391]|CHDIP043-11|BOLD:AAG3312
Megaselia|[8392]|JWDCE274-10|BOLD:AAG3232
Megaselia|[8393]|JWDCI557-10|BOLD:AAG3232
Megaselia|[8394]|JWDCI556-10|BOLD:AAG3232
Megaselia|[8395]|JWDCF320-10|BOLD:AAG3232
Megaselia|[8396]|JWDCE155-10|BOLD:AAG3232
Megaselia|[8397]|JWDCE156-10|BOLD:AAG3232
Megaselia|[8398]|JWDCE157-10|BOLD:AAG3232
Megaselia|[8399]|CHDIP981-12|BOLD:AAG3232
Megaselia|[8400]|CHDIP1133-12|BOLD:AAG3232
Megaselia|[8401]|CHDIP964-12|BOLD:ABU5538
Megaselia|[8402]|JWDCF321-10|BOLD:AAP8720
Megaselia|[8403]|JWDCJ712-11|BOLD:AAU5616
Megaselia|[8404]|JWDCF362-10|BOLD:AAP8721
Megaselia|[8405]|JWDCA181-10|BOLD:AAG3353
Megaselia|[8406]|JWDCA187-10|BOLD:AAG3353
Megaselia|[8407]|JWDCG027-10|BOLD:AAU5673
Megaselia|[8408]|TWDIP223-09|BOLD:AAG3253
Megaselia|[8409]|MTDIC236-09|BOLD:AAG3253
Megaselia|[8410]|CHDIP597-12|BOLD:AAG3253
Megaselia atrox|[8411]|CHDIP985-12|BOLD:ABX8427
Megaselia shatesae|[8412]|JWDCF776-10|BOLD:AAP8722
Megaselia shatesae|[8413]|JWDCJ1372-11|BOLD:AAP8722
Megaselia shatesae|[8414]|JWDCG689-10|BOLD:AAP8722
Megaselia shatesae|[8415]|JWDCG691-10|BOLD:AAP8722
Megaselia shatesae|[8416]|JWDCH940-10|BOLD:AAP8722
Megaselia shatesae|[8417]|JWDCH356-10|BOLD:AAP8722
Megaselia shatesae|[8418]|JWDCI493-10|BOLD:AAP8722
Megaselia shatesae|[8419]|JWDCF394-10|BOLD:AAP8722
Megaselia shatesae|[8420]|JWDCF947-10|BOLD:AAP8722
Megaselia shatesae|[8421]|JWDCF948-10|BOLD:AAP8722
Megaselia|[8422]|JWDCG102-10|BOLD:AAP6424
Megaselia|[8423]|CHDIP961-12|BOLD:AAP6424
Megaselia|[8424]|CHDIP1003-12|BOLD:AAP6424
Megaselia|[8425]|JWDCE229-10|BOLD:AAP6424
Megaselia|[8426]|JWDCE228-10|BOLD:AAP6424
Megaselia|[8427]|JWDCF696-10|BOLD:AAP6424
Megaselia|[8428]|JWDCF551-10|BOLD:AAP6424
Megaselia|[8429]|JWDCF550-10|BOLD:AAP6424
Megaselia|[8430]|CHDIP982-12|BOLD:AAP6424
Megaselia|[8431]|CHDIP992-12|BOLD:AAP6424
Megaselia|[8432]|CHDIP994-12|BOLD:AAP6424
Megaselia|[8433]|CHDIP1023-12|BOLD:AAP6424
Megaselia|[8434]|JWDCB281-10|BOLD:AAG3361
Megaselia|[8435]|JWDCA186-10|BOLD:AAG3355
Megaselia|[8436]|JWDCG397-10|BOLD:AAP8733
Megaselia|[8437]|CHDIP1179-12|BOLD:ACI6669
Megaselia|[8438]|JWDCC521-10|BOLD:AAG3266
Megaselia|[8439]|JWDCC344-10|BOLD:AAG3266
Megaselia|[8440]|JWDCG398-10|BOLD:AAG3266
Megaselia|[8441]|JWDCC597-10|BOLD:AAG3266
Megaselia|[8442]|JWDCF507-10|BOLD:AAG3266
Megaselia|[8443]|JWDCH840-10|BOLD:AAG3266
Megaselia|[8444]|JWDCE198-10|BOLD:AAG3266
Megaselia|[8445]|JWDCE199-10|BOLD:AAG3266
Megaselia|[8446]|JWDCE200-10|BOLD:AAG3266
Megaselia|[8447]|CHDIP1294-12|BOLD:AAG3266
Megaselia|[8448]|JWDCE287-10|BOLD:AAG3266
Megaselia|[8449]|JWDCE876-10|BOLD:AAG3266
Megaselia|[8450]|JWDCA189-10|BOLD:AAG3266
Megaselia|[8451]|CHDIP537-12|BOLD:AAG3266
Megaselia|[8452]|CHDIP1367-12|BOLD:AAG3266
Megaselia|[8453]|CHDIP1371-12|BOLD:AAG3266
Megaselia hardingorum|[8454]|JWDCC690-10|BOLD:AAG3274
Megaselia hardingorum|[8455]|JWDCC577-10|BOLD:AAG3274
Megaselia hardingorum|[8456]|JWDCD409-10|BOLD:AAG3274
Megaselia hardingorum|[8457]|CHDIP320-11|BOLD:AAG3274
Megaselia|[8458]|JWDCE126-10|BOLD:AAP4698
[illegible]

Megaselia hardingorum|[8456]|JWDCD409-10|BOLD:AAG3274
Megaselia hardingorum|[8457]|CHDIP320-11|BOLD:AAG3274
Megaselia|[8458]|JWDCE126-10|BOLD:AAP4698
Megaselia|[8459]|CHDIP976-12|BOLD:AAP4698
Megaselia|[8460]|JWDCB264-10|BOLD:AAG3356
Megaselia pulicaria complex|[8461]|JWDCC522-10|BOLD:AAL9073
Megaselia pulicaria complex|[8462]|CHDIP323-11|BOLD:AAL9073
Megaselia|[8463]|CHDIP979-12|BOLD:ABY9105
Megaselia|[8464]|CHDIP1084-12|BOLD:ABY9105
Megaselia mallochi|[8465]|JWDCD441-10|BOLD:AAL9084
Megaselia mallochi|[8466]|JWDCF152-10|BOLD:AAL9084
Megaselia mallochi|[8467]|JWDCE127-10|BOLD:AAL9084
Megaselia mallochi|[8468]|CHDIP004-11|BOLD:AAL9084
Megaselia mallochi|[8469]|CHDIP978-12|BOLD:AAL9084
Megaselia mallochi|[8470]|CHDIP1351-12|BOLD:AAL9084
Megaselia|[8471]|JWDCL389-11|BOLD:AAL9076
Megaselia|[8472]|JWDCF616-10|BOLD:AAL9076
Megaselia|[8473]|CHDIP008-11|BOLD:AAL9076
Megaselia|[8474]|JWDCC745-10|BOLD:AAL9076
Megaselia|[8475]|JWDCD265-10|BOLD:AAL9076
Megaselia|[8476]|CHDIP035-11|BOLD:AAL9076
Megaselia giraudii|[8477]|CHDIP197-11|BOLD:ADA4621
Megaselia giraudii|[8478]|CHDIP209-11|BOLD:ADA4621
Megaselia giraudii|[8479]|JWDCD403-10|BOLD:ADA4621
Megaselia giraudii|[8480]|JWDCA194-10|BOLD:ADA4621
Megaselia giraudii|[8481]|CHDIP247-11|BOLD:ADA4621
Megaselia giraudii|[8482]|CHDIP246-11|BOLD:ADA4621
Megaselia giraudii|[8483]|JWDCF154-10|BOLD:ADA4621
Megaselia giraudii|[8484]|JWDCD344-10|BOLD:ADA4621
Megaselia giraudii|[8485]|JWDCD220-10|BOLD:ADA4621
Megaselia giraudii|[8486]|JWDCA157-10|BOLD:ADA4621
Megaselia giraudii|[8487]|JWDCC623-10|BOLD:ADA4621
Megaselia lucifrons|[8488]|TWDIP374-09|BOLD:ADA4621
Megaselia giraudii|[8489]|CHDIP250-11|BOLD:ADA4621
Megaselia giraudii|[8490]|CHDIP051-11|BOLD:ADA4621
Megaselia giraudii|[8491]|CHDIP009-11|BOLD:ADA4621
Megaselia giraudii|[8492]|CHDIP033-11|BOLD:ADA4621
Megaselia giraudii|[8493]|CHDIP034-11|BOLD:ADA4621
Megaselia giraudii|[8494]|CHDIP037-11|BOLD:ADA4621
Megaselia giraudii|[8495]|CHDIP999-12|BOLD:ADA4621
Megaselia|[8496]|JWDCC590-10|BOLD:AAL9074
Megaselia|[8497]|JWDCC286-10|BOLD:AAL9066
Megaselia|[8498]|TWDIP373-09|BOLD:AAG3259
Megaselia|[8499]|CHDIP373-11|BOLD:AAG3259
Megaselia|[8500]|CHDIP1189-12|BOLD:AAG3259
Megaselia arcticae|[8501]|JWDCC339-10|BOLD:ACS7008
Megaselia arcticae|[8502]|TWDIP264-09|BOLD:ACS7008
Megaselia arcticae|[8503]|JWDCI445-10|BOLD:ACS7008
Megaselia arcticae|[8504]|JWDCF775-10|BOLD:ACS7008
Megaselia arcticae|[8505]|CHDIP652-12|BOLD:ACS7008
Megaselia arcticae|[8506]|JWDCF393-10|BOLD:ACS7008
Megaselia arcticae|[8507]|JWDCA190-10|BOLD:ACS7008
Megaselia arcticae|[8508]|CHDIP372-11|BOLD:ACS7008
Megaselia arcticae|[8509]|CHDIP011-11|BOLD:ACS7008
Megaselia arcticae|[8510]|CHDIP031-11|BOLD:ACS7008
Megaselia arcticae|[8511]|CHDIP620-12|BOLD:ACS7008
Megaselia arcticae|[8512]|CHDIP1097-12|BOLD:ACS7008
Megaselia arcticae|[8513]|CHDIP622-12|BOLD:ACS7008
Megaselia arcticae|[8514]|CHDIP029-11|BOLD:ACS7008
Megaselia arcticae|[8515]|CHDIP211-11|BOLD:ACS7008
Megaselia arcticae|[8516]|CHDIP1140-12|BOLD:ACS7008
Megaselia arcticae|[8517]|CHDIP392-11|BOLD:ACS7008
Megaselia arcticae|[8518]|CHDIP712-12|BOLD:ACS7008
Megaselia arcticae|[8519]|CHDIP440-11|BOLD:ACS7008
Megaselia arcticae|[8520]|MTDIC122-09|BOLD:ACS7008
Megaselia arcticae|[8521]|JWDCA197-10|BOLD:ACS7008
Megaselia arcticae|[8522]|MTDIC060-09|BOLD:ACS7008
Megaselia arcticae|[8523]|JWDCB273-10|BOLD:ACS7008
Megaselia arcticae|[8524]|MTDIC061-09|BOLD:ACS7008
Megaselia arcticae|[8525]|CHDIP019-11|BOLD:ACS7008
Megaselia arcticae|[8526]|JWDCB267-10|BOLD:ACS7008
Megaselia arcticae|[8527]|JWDCC698-10|BOLD:ACS7008
Megaselia arcticae|[8528]|JWDCC576-10|BOLD:ACS7008
Megaselia arcticae|[8529]|CHDIP176-11|BOLD:ACS7008
Megaselia arcticae|[8530]|CHDIP389-11|BOLD:ACS7008
Megaselia arcticae|[8531]|CHDIP284-11|BOLD:ACS7008
Megaselia arcticae|[8532]|CHDIP627-12|BOLD:ACS7008
Megaselia arcticae|[8533]|CHDIP643-12|BOLD:ACS7008
Megaselia arcticae|[8534]|CHDIP053-11|BOLD:ACS7008
Megaselia arcticae|[8535]|CHDIP001-11|BOLD:ACS7008
Megaselia arcticae|[8536]|CHDIP162-11|BOLD:ACS7008
Megaselia arcticae|[8537]|CHDIP017-11|BOLD:ACS7008
Megaselia arcticae|[8538]|CHDIP027-11|BOLD:ACS7008
Megaselia arcticae|[8539]|CHDIP603-12|BOLD:ACS7008
Megaselia arcticae|[8540]|CHDIP628-12|BOLD:ACS7008
Megaselia arcticae|[8541]|CHDIP030-11|BOLD:ACS7008
Megaselia arcticae|[8542]|CHDIP598-12|BOLD:ACS7008
Megaselia arcticae|[8543]|CHDIP610-12|BOLD:ACS7008
Megaselia arcticae|[8544]|CHDIP626-12|BOLD:ACS7008
Megaselia arcticae|[8545]|CHDIP630-12|BOLD:ACS7008
Megaselia arcticae|[8546]|CHDIP698-12|BOLD:ACS7008
Megaselia arcticae|[8547]|CHDIP642-12|BOLD:ACS7008
Megaselia arcticae|[8548]|CHDIP697-12|BOLD:ACS7008
Megaselia arcticae|[8549]|CHDIP703-12|BOLD:ACS7008
Megaselia arcticae|[8550]|CHDIP922-12|BOLD:ACS7008
Megaselia arcticae|[8551]|CHDIP1121-12|BOLD:ACS7008
Megaselia arcticae|[8552]|CHDIP1170-12|BOLD:ACS7008
Megaselia arcticae|[8553]|CHDIP1327-12|BOLD:ACS7008
Megaselia arcticae|[8554]|CHDIP602-12|BOLD:ACS7008
Megaselia arcticae|[8555]|CHDIP707-12|BOLD:ACS7008
Megaselia arcticae|[8556]|CHDIP1431-12|BOLD:ACS7008
Megaselia plebeia|[8557]|JWDCD266-10|BOLD:AAL9081
Megaselia|[8558]|JWDCH134-10|BOLD:AEA0066

[illegible]
Megaselia arcticae|[8556]|CHDIP1431-12|BOLD:ACS7008
Megaselia plebeia|[8557]|JWDCD266-10|BOLD:AAL9081
Megaselia|[8558]|JWDCH134-10|BOLD:AEA0066
Megaselia|[8559]|JWDCD250-10|BOLD:AEA0066
Megaselia|[8560]|JWDCD198-10|BOLD:AEA0066
Megaselia|[8561]|JWDCG563-10|BOLD:AEA0066
Megaselia|[8562]|JWDCF774-10|BOLD:AEA0066
Megaselia|[8563]|JWDCE808-10|BOLD:AEA0066
Megaselia|[8564]|CHDIP975-12|BOLD:AEA0066
Megaselia|[8565]|CHDIP980-12|BOLD:AEA0066
Phora|[8566]|JWDCD399-10|BOLD:AAL9082
Phora|[8567]|CHDIP015-11|BOLD:AAL9082
Phora viridinota|[8568]|TWDIP098-09|BOLD:AAG3240
Phora viridinota|[8569]|JWDCF575-10|BOLD:AAG3240
Phora|[8570]|JWDCE042-10|BOLD:AAP1929
Phora|[8571]|JWDCC341-10|BOLD:AAL9071
Phora|[8572]|JWDCA152-10|BOLD:AAG3249
Phora|[8573]|JWDCE760-10|BOLD:AAG3249
Phora|[8574]|JWDCD221-10|BOLD:AAG3249
Phora|[8575]|CHDIP125-11|BOLD:AAG3249
Phora|[8576]|MTDIC090-09|BOLD:AAG3249
Phora|[8577]|JWDCE875-10|BOLD:AAG3249
Phora|[8578]|JWDCA053-10|BOLD:AAG3249
Phora|[8579]|CHDIP126-11|BOLD:AAG3249
Phora|[8580]|JWDCA150-10|BOLD:AAG3249
Phora|[8581]|JWDCA147-10|BOLD:AAG3249
Phora|[8582]|JWDCA144-10|BOLD:AAG3249
Phora|[8583]|JWDCA140-10|BOLD:AAG3249
Phora|[8584]|JWDCD416-10|BOLD:AAG3249
Phora|[8585]|JWDCD287-10|BOLD:AAG3249
Phora|[8586]|JWDCD241-10|BOLD:AAG3249
Phora|[8587]|JWDCC700-10|BOLD:AAG3249
Phora|[8588]|JWDCI196-10|BOLD:AAG3249
Phora|[8589]|MTDIC091-09|BOLD:AAG3249
Phora|[8590]|JWDCA141-10|BOLD:AAG3249
Phora|[8591]|CHDIP046-11|BOLD:AAG3249
Phora carlina|[8592]|JWDCD199-10|BOLD:AAG3239
Phora|[8593]|MTDIC022-09|BOLD:AAG3247
Phora|[8594]|CHDIP651-12|BOLD:AAG3247
Megaselia|[8595]|JWDCC704-10|BOLD:AAL9070
Megaselia|[8596]|JWDCC624-10|BOLD:AAL9070
Megaselia|[8597]|CHDIP012-11|BOLD:AAL9070
Megaselia|[8598]|JWDCF153-10|BOLD:AAL9070
Megaselia|[8599]|JWDCJ936-11|BOLD:AAL9070
Megaselia|[8600]|JWDCJ421-11|BOLD:AAL9070
Megaselia|[8601]|JWDCC337-10|BOLD:AAL9070
Megaselia|[8602]|JWDCD418-10|BOLD:AAL9070
Megaselia|[8603]|JWDCC484-10|BOLD:AAL9070
Megaselia|[8604]|JWDCD417-10|BOLD:AAL9070
Megaselia|[8605]|CHDIP990-12|BOLD:AAL9070
Megaselia|[8606]|CHDIP1005-12|BOLD:AAL9070
Megaselia|[8607]|CHDIP1381-12|BOLD:AAL9070
Megaselia|[8608]|CHDIP508-12|BOLD:ABY0968
Megaselia|[8609]|CHDIP941-12|BOLD:ABY0968
Megaselia|[8610]|TWDIP100-09|BOLD:AAG3241
Megaselia|[8611]|CHDIP1015-12|BOLD:AAG3241
Megaselia|[8612]|CHDIP050-11|BOLD:AAG3241
Megaselia|[8613]|TWDIP103-09|BOLD:AAG3241
Megaselia|[8614]|JWDCA156-10|BOLD:AAG3241
Megaselia|[8615]|CHDIP023-11|BOLD:AAG3241
Megaselia|[8616]|CHDIP251-11|BOLD:AAG3241
Megaselia|[8617]|CHDIP018-11|BOLD:AAG3241
Megaselia|[8618]|CHDIP1013-12|BOLD:AAG3241
Megaselia|[8619]|CHDIP1014-12|BOLD:AAG3241
Megaselia|[8620]|JWDCD443-10|BOLD:AAG3241
Megaselia|[8621]|CHDIP1020-12|BOLD:AAG3241
Megaselia|[8622]|CHDIP025-11|BOLD:AAG3358
Megaselia|[8623]|CHDIP042-11|BOLD:AAG3358
Megaselia|[8624]|JWDCB269-10|BOLD:AAG3358
Megaselia|[8625]|CHDIP010-11|BOLD:AAG3358
Megaselia|[8626]|CHDIP943-12|BOLD:AAG3358
Megaselia|[8627]|CHDIP946-12|BOLD:AAG3358
Megaselia|[8628]|CHDIP1396-12|BOLD:AAG3358
Megaselia|[8629]|JWDCE874-10|BOLD:AAP9863
Megaselia|[8630]|TWDIP135-09|BOLD:AAG3243
Megaselia|[8631]|JWDCD197-10|BOLD:AAG3243
Megaselia tecticauda|[8632]|JWDCG358-10|BOLD:AAG3311
Megaselia tecticauda|[8633]|JWDCC539-10|BOLD:AAG3311
Megaselia tecticauda|[8634]|CHDIP032-11|BOLD:AAG3311
Megaselia|[8635]|JWDCE041-10|BOLD:AAP1928
Megaselia|[8636]|CHDIP960-12|BOLD:AAP1928
Megaselia|[8637]|JWDCJ1589-11|BOLD:AAQ3659
Megaselia|[8638]|JWDCJ1229-11|BOLD:AAQ3659
Megaselia|[8639]|CHDIP645-12|BOLD:ADA3077
Megaselia|[8640]|CHDIP1126-12|BOLD:ADA3077
Megaselia|[8641]|CHDIP386-11|BOLD:AAG3341
Megaselia|[8642]|JWDCG506-10|BOLD:AAP8734
Megaselia arizonensis|[8643]|CHDIP253-11|BOLD:AAP4684
Megaselia arizonensis|[8644]|CHDIP653-12|BOLD:AAP4684
Megaselia|[8645]|CHDIP916-12|BOLD:ABY1806
Megaselia|[8646]|CHDIP432-11|BOLD:ABY1806
Megaselia|[8647]|CHDIP920-12|BOLD:ABY1806
Megaselia|[8648]|CHDIP926-12|BOLD:ABY1806
Megaselia|[8649]|CHDIP700-12|BOLD:ACZ3800
Megaselia|[8650]|CHDIP1113-12|BOLD:ACZ3800
Megaselia|[8651]|CHDIP599-12|BOLD:ACZ3800
Megaselia|[8652]|CHDIP1287-12|BOLD:ACZ3800
Megaselia|[8653]|CHDIP719-12|BOLD:ACZ3800
Megaselia|[8654]|CHDIP644-12|BOLD:ACZ3800
Megaselia|[8655]|JWDCL388-11|BOLD:ACZ3800
Megaselia|[8656]|JWDCH697-10|BOLD:ACZ3800
Megaselia|[8657]|JWDCI627-10|BOLD:ACZ3800
[illegible]

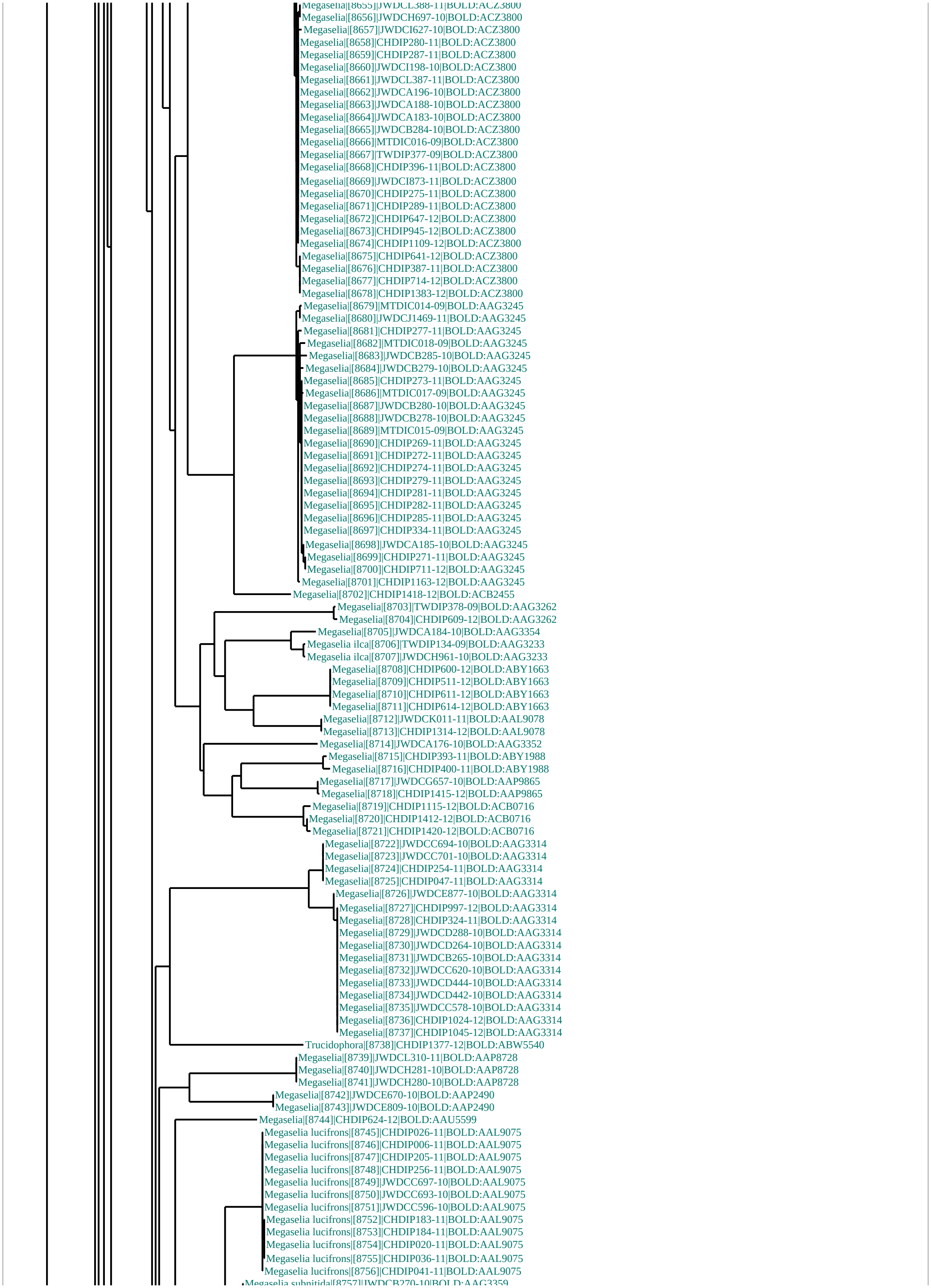
Megaselia|[8655]|JWDCL388-11|BOLD:ACZ3800
Megaselia|[8656]|JWDCH697-10|BOLD:ACZ3800
Megaselia|[8657]|JWDCI627-10|BOLD:ACZ3800
Megaselia|[8658]|CHDIP280-11|BOLD:ACZ3800
Megaselia|[8659]|CHDIP287-11|BOLD:ACZ3800
Megaselia|[8660]|JWDCI198-10|BOLD:ACZ3800
Megaselia|[8661]|JWDCL387-11|BOLD:ACZ3800
Megaselia|[8662]|JWDCA196-10|BOLD:ACZ3800
Megaselia|[8663]|JWDCA188-10|BOLD:ACZ3800
Megaselia|[8664]|JWDCA183-10|BOLD:ACZ3800
Megaselia|[8665]|JWDCB284-10|BOLD:ACZ3800
Megaselia|[8666]|MTDIC016-09|BOLD:ACZ3800
Megaselia|[8667]|TWDIP377-09|BOLD:ACZ3800
Megaselia|[8668]|CHDIP396-11|BOLD:ACZ3800
Megaselia|[8669]|JWDCI873-11|BOLD:ACZ3800
Megaselia|[8670]|CHDIP275-11|BOLD:ACZ3800
Megaselia|[8671]|CHDIP289-11|BOLD:ACZ3800
Megaselia|[8672]|CHDIP647-12|BOLD:ACZ3800
Megaselia|[8673]|CHDIP945-12|BOLD:ACZ3800
Megaselia|[8674]|CHDIP1109-12|BOLD:ACZ3800
Megaselia|[8675]|CHDIP641-12|BOLD:ACZ3800
Megaselia|[8676]|CHDIP387-11|BOLD:ACZ3800
Megaselia|[8677]|CHDIP714-12|BOLD:ACZ3800
Megaselia|[8678]|CHDIP1383-12|BOLD:ACZ3800
Megaselia|[8679]|MTDIC014-09|BOLD:AAG3245
Megaselia|[8680]|JWDCJ1469-11|BOLD:AAG3245
Megaselia|[8681]|CHDIP277-11|BOLD:AAG3245
Megaselia|[8682]|MTDIC018-09|BOLD:AAG3245
Megaselia|[8683]|JWDCB285-10|BOLD:AAG3245
Megaselia|[8684]|JWDCB279-10|BOLD:AAG3245
Megaselia|[8685]|CHDIP273-11|BOLD:AAG3245
Megaselia|[8686]|MTDIC017-09|BOLD:AAG3245
Megaselia|[8687]|JWDCB280-10|BOLD:AAG3245
Megaselia|[8688]|JWDCB278-10|BOLD:AAG3245
Megaselia|[8689]|MTDIC015-09|BOLD:AAG3245
Megaselia|[8690]|CHDIP269-11|BOLD:AAG3245
Megaselia|[8691]|CHDIP272-11|BOLD:AAG3245
Megaselia|[8692]|CHDIP274-11|BOLD:AAG3245
Megaselia|[8693]|CHDIP279-11|BOLD:AAG3245
Megaselia|[8694]|CHDIP281-11|BOLD:AAG3245
Megaselia|[8695]|CHDIP282-11|BOLD:AAG3245
Megaselia|[8696]|CHDIP285-11|BOLD:AAG3245
Megaselia|[8697]|CHDIP334-11|BOLD:AAG3245
Megaselia|[8698]|JWDCA185-10|BOLD:AAG3245
Megaselia|[8699]|CHDIP271-11|BOLD:AAG3245
Megaselia|[8700]|CHDIP711-12|BOLD:AAG3245
Megaselia|[8701]|CHDIP1163-12|BOLD:AAG3245
Megaselia|[8702]|CHDIP1418-12|BOLD:ACB2455
Megaselia|[8703]|TWDIP378-09|BOLD:AAG3262
Megaselia|[8704]|CHDIP609-12|BOLD:AAG3262
Megaselia|[8705]|JWDCA184-10|BOLD:AAG3354
Megaselia ilca|[8706]|TWDIP134-09|BOLD:AAG3233
Megaselia ilca|[8707]|JWDCH961-10|BOLD:AAG3233
Megaselia|[8708]|CHDIP600-12|BOLD:ABY1663
Megaselia|[8709]|CHDIP511-12|BOLD:ABY1663
Megaselia|[8710]|CHDIP611-12|BOLD:ABY1663
Megaselia|[8711]|CHDIP614-12|BOLD:ABY1663
Megaselia|[8712]|JWDCK011-11|BOLD:AAL9078
Megaselia|[8713]|CHDIP1314-12|BOLD:AAL9078
Megaselia|[8714]|JWDCA176-10|BOLD:AAG3352
Megaselia|[8715]|CHDIP393-11|BOLD:ABY1988
Megaselia|[8716]|CHDIP400-11|BOLD:ABY1988
Megaselia|[8717]|JWDCG657-10|BOLD:AAP9865
Megaselia|[8718]|CHDIP1415-12|BOLD:AAP9865
Megaselia|[8719]|CHDIP1115-12|BOLD:ACB0716
Megaselia|[8720]|CHDIP1412-12|BOLD:ACB0716
Megaselia|[8721]|CHDIP1420-12|BOLD:ACB0716
Megaselia|[8722]|JWDCC694-10|BOLD:AAG3314
Megaselia|[8723]|JWDCC701-10|BOLD:AAG3314
Megaselia|[8724]|CHDIP254-11|BOLD:AAG3314
Megaselia|[8725]|CHDIP047-11|BOLD:AAG3314
Megaselia|[8726]|JWDCE877-10|BOLD:AAG3314
Megaselia|[8727]|CHDIP997-12|BOLD:AAG3314
Megaselia|[8728]|CHDIP324-11|BOLD:AAG3314
Megaselia|[8729]|JWDCD288-10|BOLD:AAG3314
Megaselia|[8730]|JWDCD264-10|BOLD:AAG3314
Megaselia|[8731]|JWDCB265-10|BOLD:AAG3314
Megaselia|[8732]|JWDCC620-10|BOLD:AAG3314
Megaselia|[8733]|JWDCD444-10|BOLD:AAG3314
Megaselia|[8734]|JWDCD442-10|BOLD:AAG3314
Megaselia|[8735]|JWDCC578-10|BOLD:AAG3314
Megaselia|[8736]|CHDIP1024-12|BOLD:AAG3314
Megaselia|[8737]|CHDIP1045-12|BOLD:AAG3314
Trucidophora|[8738]|CHDIP1377-12|BOLD:ABW5540
Megaselia|[8739]|JWDCL310-11|BOLD:AAP8728
Megaselia|[8740]|JWDCH281-10|BOLD:AAP8728
Megaselia|[8741]|JWDCH280-10|BOLD:AAP8728
Megaselia|[8742]|JWDCE670-10|BOLD:AAP2490
Megaselia|[8743]|JWDCE809-10|BOLD:AAP2490
Megaselia|[8744]|CHDIP624-12|BOLD:AAU5599
Megaselia lucifrons|[8745]|CHDIP026-11|BOLD:AAL9075
Megaselia lucifrons|[8746]|CHDIP006-11|BOLD:AAL9075
Megaselia lucifrons|[8747]|CHDIP205-11|BOLD:AAL9075
Megaselia lucifrons|[8748]|CHDIP256-11|BOLD:AAL9075
Megaselia lucifrons|[8749]|JWDCC697-10|BOLD:AAL9075
Megaselia lucifrons|[8750]|JWDCC693-10|BOLD:AAL9075
Megaselia lucifrons|[8751]|JWDCC596-10|BOLD:AAL9075
Megaselia lucifrons|[8752]|CHDIP183-11|BOLD:AAL9075
Megaselia lucifrons|[8753]|CHDIP184-11|BOLD:AAL9075
Megaselia lucifrons|[8754]|CHDIP020-11|BOLD:AAL9075
Megaselia lucifrons|[8755]|CHDIP036-11|BOLD:AAL9075
Megaselia lucifrons|[8756]|CHDIP041-11|BOLD:AAL9075
Megaselia subnitida|[8757]|JWDCB270-10|BOLD:AAG3359

Megaselia lucifrons|[8755]|CHDIP036-11|BOLD:AAL9075
Megaselia lucifrons|[8756]|CHDIP041-11|BOLD:AAL9075
Megaselia subnitida|[8757]|JWDCB270-10|BOLD:AAG3359
Megaselia subnitida|[8758]|JWDCC300-10|BOLD:AAG3359
Megaselia subnitida|[8759]|JWDCD242-10|BOLD:AAG3359
Megaselia subnitida|[8760]|JWDCD247-10|BOLD:AAG3359
Megaselia subnitida|[8761]|JWDCJ989-11|BOLD:AAG3359
Megaselia subnitida|[8762]|JWDCE386-10|BOLD:AAG3359
Megaselia subnitida|[8763]|CHDIP044-11|BOLD:AAG3359
Megaselia subnitida|[8764]|JWDCD402-10|BOLD:AAG3359
Megaselia subnitida|[8765]|JWDCC746-10|BOLD:AAG3359
Megaselia subnitida|[8766]|JWDCC743-10|BOLD:AAG3359
Megaselia subnitida|[8767]|JWDCC626-10|BOLD:AAG3359
Megaselia subnitida|[8768]|JWDCB277-10|BOLD:AAG3359
Megaselia subnitida|[8769]|JWDCJ420-11|BOLD:AAG3359
Megaselia subnitida|[8770]|JWDCE388-10|BOLD:AAG3359
Megaselia subnitida|[8771]|JWDCE758-10|BOLD:AAG3359
Megaselia subnitida|[8772]|JWDCF548-10|BOLD:AAG3359
Megaselia subnitida|[8773]|JWDCE878-10|BOLD:AAG3359
Megaselia subnitida|[8774]|CHDIP222-11|BOLD:AAG3359
Megaselia subnitida|[8775]|CHDIP048-11|BOLD:AAG3359
Megaselia subnitida|[8776]|CHDIP1000-12|BOLD:AAG3359
Megaselia|[8777]|CHDIP1083-12|BOLD:AAN8683
Megaselia|[8778]|JWDCH584-10|BOLD:AAP8725
Megaselia|[8779]|JWDCH585-10|BOLD:AAP8725
Megaselia|[8780]|JWDCH060-10|BOLD:AAP8725
Megaselia|[8781]|JWDCA160-10|BOLD:ACZ4279
Megaselia|[8782]|JWDCA163-10|BOLD:ACZ4279
Megaselia|[8783]|JWDCA169-10|BOLD:ACZ4279
Megaselia|[8784]|JWDCE547-10|BOLD:AAP3610
Megaselia|[8785]|CHDIP983-12|BOLD:AAP3610
Megaselia|[8786]|CHDIP623-12|BOLD:AAG3255
Megaselia|[8787]|MTDIC267-09|BOLD:AAG3255
Megaselia|[8788]|JWDCG062-10|BOLD:AAG3255
Megaselia|[8789]|CHDIP605-12|BOLD:AAG3255
Megaselia|[8790]|CHDIP1096-12|BOLD:AAG3255
Megaselia|[8791]|CHDIP322-11|BOLD:AAG3251
Megaselia|[8792]|CHDIP321-11|BOLD:AAG3251
Megaselia|[8793]|JWDCA167-10|BOLD:AAG3251
Megaselia|[8794]|JWDCK102-11|BOLD:AAG3251
Megaselia|[8795]|MTDIC114-09|BOLD:AAG3251
Megaselia|[8796]|CHDIP398-11|BOLD:AAG3251
Megaselia|[8797]|TWDIP375-09|BOLD:AAG3251
Megaselia|[8798]|JWDCA164-10|BOLD:AAG3251
Megaselia|[8799]|JWDCA165-10|BOLD:AAG3251
Megaselia|[8800]|JWDCA170-10|BOLD:AAG3251
Megaselia|[8801]|JWDCA166-10|BOLD:AAG3251
Megaselia|[8802]|CHDIP424-11|BOLD:AAG3251
Megaselia|[8803]|CHDIP276-11|BOLD:AAG3251
Megaselia|[8804]|CHDIP596-12|BOLD:AAG3251
Megaselia lutea|[8805]|JWDCA161-10|BOLD:AAG3351
Megaselia lutea|[8806]|CHDIP615-12|BOLD:AAG3351
Megaselia|[8807]|CHDIP619-12|BOLD:ACZ4314
Megaselia|[8808]|CHDIP455-12|BOLD:ACZ4314
Megaselia|[8809]|JWDCA168-10|BOLD:ACZ4314
Megaselia|[8810]|JWDCA158-10|BOLD:ACZ4314
Megaselia|[8811]|JWDCA162-10|BOLD:ACZ4314
Megaselia|[8812]|JWDCG254-10|BOLD:ACZ4314
Megaselia|[8813]|JWDCG268-10|BOLD:ACZ4314
Megaselia|[8814]|CHDIP612-12|BOLD:ACZ4314
Megaselia|[8815]|CHDIP618-12|BOLD:ACZ4314
Megaselia|[8816]|CHDIP702-12|BOLD:ACZ4314
Megaselia|[8817]|MTDIC255-09|BOLD:ACZ4314
Megaselia|[8818]|CHDIP629-12|BOLD:ACZ4314
Megaselia|[8819]|CHDIP1183-12|BOLD:ACZ4314
Megaselia|[8820]|JWDCA175-10|BOLD:AAG3349
Megaselia|[8821]|JWDCD201-10|BOLD:AAG3349
Megaselia|[8822]|JWDCA148-10|BOLD:AAG3349
Megaselia|[8823]|JWDCD407-10|BOLD:AAG3349
Megaselia|[8824]|JWDCD200-10|BOLD:AAG3349
Megaselia|[8825]|JWDCC696-10|BOLD:AAG3349
Megaselia|[8826]|JWDCB266-10|BOLD:AAG3349
Megaselia|[8827]|JWDCC692-10|BOLD:AAG3349
Megaselia|[8828]|JWDCC575-10|BOLD:AAG3349
Megaselia|[8829]|JWDCC538-10|BOLD:AAG3349
Megaselia|[8830]|JWDCD406-10|BOLD:AAG3349
Megaselia|[8831]|JWDCD248-10|BOLD:AAG3349
Megaselia|[8832]|JWDCD408-10|BOLD:AAG3349
Megaselia|[8833]|CHDIP249-11|BOLD:AAG3349
Megaselia|[8834]|CHDIP175-11|BOLD:AAG3349
Megaselia|[8835]|CHDIP007-11|BOLD:AAG3349
Megaselia|[8836]|CHDIP248-11|BOLD:AAG3349
Megaselia|[8837]|CHDIP024-11|BOLD:AAG3349
Megaselia pulicaria complex|[8838]|JWDCJ1230-11|BOLD:AAU8534
Megaselia pulicaria complex|[8839]|CHDIP915-12|BOLD:AAU8534
Megaselia pulicaria complex|[8840]|JWDCH192-10|BOLD:AAU8534
Megaselia pulicaria complex|[8841]|JWDCH189-10|BOLD:AAU8534
Megaselia pulicaria complex|[8842]|JWDCB276-10|BOLD:AAU8534
Megaselia pulicaria complex|[8843]|CHDIP1185-12|BOLD:AAU8534
Megaselia|[8844]|TWDIP376-09|BOLD:AAG3261
Megaselia|[8845]|JWDCB271-10|BOLD:AAG3360
Megaselia|[8846]|JWDCH190-10|BOLD:AAP8727
Megaselia|[8847]|JWDCG252-10|BOLD:AAP8731
Megaselia|[8848]|JWDCI495-10|BOLD:AAP8731
Megaselia|[8849]|CHDIP804-12|BOLD:ABZ0111
Megaselia|[8850]|CHDIP917-12|BOLD:ABZ0111
Megaselia nigriceps|[8851]|CHDIP608-12|BOLD:AAG3304
Megaselia nigriceps|[8852]|JWDCG505-10|BOLD:AAG3304
Megaselia nigriceps|[8853]|JWDCJ456-11|BOLD:AAG3304
Megaselia nigriceps|[8854]|JWDCA159-10|BOLD:AAG3304
Megaselia nigriceps|[8855]|JWDCF363-10|BOLD:AAG3304
Megaselia nigriceps|[8856]|CHDIP288-11|BOLD:AAG3304

Megaselia nigriceps|[8854]|JWDCA159-10|BOLD:AAG3304
Megaselia nigriceps|[8855]|JWDCF363-10|BOLD:AAG3304
Megaselia nigriceps|[8856]|CHDIP288-11|BOLD:AAG3304
Megaselia nigriceps|[8857]|CHDIP130-11|BOLD:AAG3304
Megaselia nigriceps|[8858]|CHDIP453-12|BOLD:AAG3304
Megaselia nigriceps|[8859]|CHDIP974-12|BOLD:AAG3304
Megaselia nigriceps|[8860]|CHDIP1206-12|BOLD:AAG3304
Phalacrotophora epeirae|[8861]|CHDIP704-12|BOLD:AAM9371
Phalacrotophora epeirae|[8862]|CHDIP1091-12|BOLD:AAM9371
Phalacrotophora epeirae|[8863]|CHDIP1436-12|BOLD:AAM9371
Phalacrotophora epeirae|[8864]|CHDIP1439-12|BOLD:AAM9371
Phalacrotophora epeirae|[8865]|JWDCK103-11|BOLD:AAM9371
Phalacrotophora epeirae|[8866]|CHDIP1230-12|BOLD:AAM9371
Phalacrotophora epeirae|[8867]|CHDIP1440-12|BOLD:AAM9371
Hammerschmidtia ferruginea|[8868]|MHSYR375-07|BOLD:AAI8336
Ferdinandea buccata|[8869]|JWDCA490-10|BOLD:AAE0948
Neoascia tenur|[8870]|JWDCH200-10|BOLD:ACE9059
Neoascia geniculata|[8871]|JWDCH818-10|BOLD:AAI3040
Neoascia geniculata|[8872]|JWDCL457-11|BOLD:AAI3040
Neoascia geniculata|[8873]|JWDCG881-10|BOLD:AAI3040
Neoascia geniculata|[8874]|JWDCG611-10|BOLD:AAI3040
Sphegina|[8875]|MHSYR326-07|BOLD:AAF2591
Sericomyia sexfasciata|[8876]|JWDCJ845-11|BOLD:AAD0792
Sericomyia sexfasciata|[8877]|JWDCG604-10|BOLD:AAD0792
Sericomyia sexfasciata|[8878]|JWDCG766-10|BOLD:AAD0792
Sericomyia sexfasciata|[8879]|JWDCG602-10|BOLD:AAD0792
Sericomyia sexfasciata|[8880]|JWDCK130-11|BOLD:AAD0792
Sericomyia sexfasciata|[8881]|JWDCF473-10|BOLD:AAD0792
Sericomyia sexfasciata|[8882]|JWDCJ1047-11|BOLD:AAD0792
Sericomyia sexfasciata|[8883]|JWDCJ849-11|BOLD:AAD0792
Sericomyia sexfasciata|[8884]|JWDCI416-10|BOLD:AAD0792
Sericomyia sexfasciata|[8885]|JWDCL249-11|BOLD:AAD0792
Sericomyia sexfasciata|[8886]|JWDCJ608-11|BOLD:AAD0792
Sericomyia sexfasciata|[8887]|JWDCG181-10|BOLD:AAD0792
Sericomyia sexfasciata|[8888]|JWDCG859-10|BOLD:AAD0792
Melanostoma mellinum|[8889]|TWDIP072-09|BOLD:AAB2866
Melanostoma mellinum|[8890]|JWDCJ300-11|BOLD:AAB2866
Melanostoma mellinum|[8891]|JWDCF591-10|BOLD:AAB2866
Melanostoma mellinum|[8892]|JWDCF592-10|BOLD:AAB2866
Melanostoma mellinum|[8893]|JWDCJ003-11|BOLD:AAB2866
Melanostoma mellinum|[8894]|JWDCB001-10|BOLD:AAB2866
Melanostoma mellinum|[8895]|JWDCE653-10|BOLD:AAB2866
Melanostoma mellinum|[8896]|JWDCL364-11|BOLD:AAB2866
Melanostoma mellinum|[8897]|JWDCF270-10|BOLD:AAB2866
Melanostoma mellinum|[8898]|JWDCI465-10|BOLD:AAB2866
Melanostoma mellinum|[8899]|JWDCI330-10|BOLD:AAB2866
Melanostoma mellinum|[8900]|JWDCI328-10|BOLD:AAB2866
Melanostoma mellinum|[8901]|JWDCF754-10|BOLD:AAB2866
Melanostoma mellinum|[8902]|JWDCF525-10|BOLD:AAB2866
Melanostoma mellinum|[8903]|JWDCJ692-11|BOLD:AAB2866
Melanostoma mellinum|[8904]|JWDCF269-10|BOLD:AAB2866
Melanostoma mellinum|[8905]|JWDCI466-10|BOLD:AAB2866
Melanostoma mellinum|[8906]|JWDCH119-10|BOLD:AAB2866
Melanostoma mellinum|[8907]|JWDCL455-11|BOLD:AAB2866
Melanostoma mellinum|[8908]|JWDCH317-10|BOLD:AAB2866
Melanostoma mellinum|[8909]|JWDCF656-10|BOLD:AAB2866
Melanostoma mellinum|[8910]|JWDCG693-10|BOLD:AAB2866
Melanostoma mellinum|[8911]|JWDCA503-10|BOLD:AAB2866
Melanostoma mellinum|[8912]|JWDCH201-10|BOLD:AAB2866
Melanostoma mellinum|[8913]|JWDCF020-10|BOLD:AAB2866
Melanostoma mellinum|[8914]|JWDCC002-10|BOLD:AAB2866
Melanostoma mellinum|[8915]|JWDCF753-10|BOLD:AAB2866
Melanostoma mellinum|[8916]|JWDCD099-10|BOLD:AAB2866
Melanostoma mellinum|[8917]|JWDCA479-10|BOLD:AAB2866
Melanostoma mellinum|[8918]|JWDCJ694-11|BOLD:AAB2866
Melanostoma mellinum|[8919]|JWDCF019-10|BOLD:AAB2866
Melanostoma mellinum|[8920]|JWDCF900-10|BOLD:AAB2866
Melanostoma mellinum|[8921]|JWDCJ1333-11|BOLD:AAB2866
Melanostoma mellinum|[8922]|JWDCI287-10|BOLD:AAB2866
Melanostoma mellinum|[8923]|JWDCI329-10|BOLD:AAB2866
Melanostoma mellinum|[8924]|JWDCL593-11|BOLD:AAB2866
Melanostoma mellinum|[8925]|JWDCE856-10|BOLD:AAB2866
Melanostoma mellinum|[8926]|JWDCF268-10|BOLD:AAB2866
Melanostoma mellinum|[8927]|JWDCJ693-11|BOLD:AAB2866
Melanostoma mellinum|[8928]|JWDCJ223-11|BOLD:AAB2866
Cheilosia|[8929]|MHSYR390-07|BOLD:AAC8002
Cheilosia|[8930]|MTDIC124-09|BOLD:AAC8002
Cheilosia latrans|[8931]|JWDCA485-10|BOLD:AAA3293
Cheilosia latrans|[8932]|JWDCH272-10|BOLD:AAA3293
Cheilosia latrans|[8933]|JWDCA486-10|BOLD:AAA3293
Cheilosia latrans|[8934]|JWDCJ1256-11|BOLD:AAA3293
Cheilosia latrans|[8935]|JWDCG497-10|BOLD:AAA3293
Cheilosia latrans|[8936]|JWDCG276-10|BOLD:AAA3293
Cheilosia latrans|[8937]|JWDCG438-10|BOLD:AAA3293
Cheilosia hunteri|[8938]|MHSYR320-07|BOLD:AAC1338
Cheilosia hunteri|[8939]|JWDCJ486-11|BOLD:AAC1338
Cheilosia hunteri|[8940]|JWDCJ500-11|BOLD:AAC1338
Cheilosia hunteri|[8941]|JSFLA040-08|BOLD:AAC1338
Cheilosia sp. nov.|[8942]|MHSYR323-07|BOLD:AAC8001
Cheilosia sp. nov.|[8943]|MHSYR328-07|BOLD:AAC8001
Cheilosia sp. nov.|[8944]|MHSYR350-07|BOLD:AAC8001
Cheilosia sp. nov.|[8945]|JWDCF737-10|BOLD:AAC8001
Cheilosia sp. nov.|[8946]|JWDCB024-10|BOLD:AAC8001
Cheilosia sp. nov.|[8947]|JWDCI700-11|BOLD:AAC8001
Cheilosia sp. nov.|[8948]|JWDCC200-10|BOLD:AAC8001
Cheilosia sp. nov.|[8949]|MHSYR327-07|BOLD:AAC8001
Cheilosia sp. nov.|[8950]|MHSYR325-07|BOLD:AAC8001
Cheilosia sp. nov.|[8951]|MHSYR333-07|BOLD:AAC8001
Cheilosia sp. nov.|[8952]|MHSYR357-07|BOLD:AAC8001
Cheilosia sp. nov.|[8953]|JWDCJ540-11|BOLD:AAC8001
Cheilosia sp. nov.|[8954]|JWDCF712-10|BOLD:AAC8001
Cheilosia sp. nov.|[8955]|JWDCE164-10|BOLD:AAC8001
Cheilosia sp. nov.|[8956]|MHSYR319-07|BOLD:AAC8001

Cheilosia sp. nov.|[8954]|JWDCF712-10|BOLD:AAC8001
Cheilosia sp. nov.|[8955]|JWDCE164-10|BOLD:AAC8001
Cheilosia sp. nov.|[8956]|MHSYR319-07|BOLD:AAC8001
Cheilosia sp. nov.|[8957]|JWDCB023-10|BOLD:AAC8001
Cheilosia sp. nov.|[8958]|MHSYR324-07|BOLD:AAC8001
Orthonevra robusta|[8959]|JWDCF498-10|BOLD:AAP8766
Orthonevra robusta|[8960]|JWDCF492-10|BOLD:AAP8766
Orthonevra robusta|[8961]|JWDCI245-10|BOLD:AAP8766
Orthonevra robusta|[8962]|JWDCI309-10|BOLD:AAP8766
Chrysosyrphus latus|[8963]|MHSYR355-07|BOLD:AAF2592
Chrysosyrphus latus|[8964]|MHSYR356-07|BOLD:AAF2592
Chrysosyrphus latus|[8965]|JWDCC182-10|BOLD:AAF2592
Chrysosyrphus latus|[8966]|JWDCF131-10|BOLD:AAF2592
Chrysosyrphus latus|[8967]|JWDCI285-10|BOLD:AAF2592
Chrysosyrphus latus|[8968]|JWDCB025-10|BOLD:AAF2592
Chrysosyrphus latus|[8969]|JWDCC006-10|BOLD:AAF2592
Chrysosyrphus latus|[8970]|JWDCL320-11|BOLD:AAF2592
Chrysosyrphus latus|[8971]|JWDCI564-10|BOLD:AAF2592
Chrysosyrphus latus|[8972]|JWDCE836-10|BOLD:AAF2592
Chrysosyrphus latus|[8973]|JWDCD040-10|BOLD:AAF2592
Chrysosyrphus latus|[8974]|JSFLA038-08|BOLD:AAF2592
Brachyopa notata|[8975]|MHSYR321-07|BOLD:AAI8335
Chalcosyrphus anomalus|[8976]|JWDCJ487-11|BOLD:AAQ2056
Chalcosyrphus anomalus|[8977]|JWDCJ541-11|BOLD:AAQ2056
Xylota quadrimaculata|[8978]|JWDCL321-11|BOLD:AAG4661
Xylota flavifrons|[8979]|MHSYR318-07|BOLD:AAC2125
Xylota flavifrons|[8980]|JWDCF553-10|BOLD:AAC2125
Xylota flavifrons|[8981]|MHSYR308-07|BOLD:AAC2125
Xylota flavifrons|[8982]|MHSYR309-07|BOLD:AAC2125
Xylota flavifrons|[8983]|MHSYR307-07|BOLD:AAC2125
Xylota flavifrons|[8984]|MHSYR315-07|BOLD:AAC2125
Xylota flavifrons|[8985]|JWDCH118-10|BOLD:AAC2125
Xylota flavifrons|[8986]|JWDCF871-10|BOLD:AAC2125
Xylota flavifrons|[8987]|JWDCI007-10|BOLD:AAC2125
Xylota flavifrons|[8988]|JWDCI006-10|BOLD:AAC2125
Xylota flavifrons|[8989]|MHSYR314-07|BOLD:AAC2125
Xylota flavifrons|[8990]|MHSYR313-07|BOLD:AAC2125
Xylota flavifrons|[8991]|MHSYR311-07|BOLD:AAC2125
Xylota flavifrons|[8992]|JWDCJ609-11|BOLD:AAC2125
Xylota flavifrons|[8993]|JWDCG767-10|BOLD:AAC2125
Xylota flavifrons|[8994]|MHSYR312-07|BOLD:AAC2125
Xylota flavifrons|[8995]|JWDCI005-10|BOLD:AAC2125
Xylota flavifrons|[8996]|MHFLY037-06|BOLD:AAC2125
Xylota flavifrons|[8997]|MHSYR316-07|BOLD:AAC2125
Xylota flavifrons|[8998]|MHSYR310-07|BOLD:AAC2125
Xylota flavifrons|[8999]|LRSYR003-07|BOLD:AAC2125
Xylota ouelleti|[9000]|MHFLI199-06|BOLD:ABZ0281
Xylota ouelleti|[9001]|JWDCF872-10|BOLD:ABZ0281
Xylota ouelleti|[9002]|JWDCG223-10|BOLD:ABZ0281
Xylota ouelleti|[9003]|JWDCB018-10|BOLD:ABZ0281
Xylota ouelleti|[9004]|JWDCG603-10|BOLD:ABZ0281
Xylota ouelleti|[9005]|JWDCG014-10|BOLD:ABZ0281
Xylota ouelleti|[9006]|JWDCG034-10|BOLD:ABZ0281
Xylota ouelleti|[9007]|JWDCG225-10|BOLD:ABZ0281
Xylota ouelleti|[9008]|JWDCG226-10|BOLD:ABZ0281
Xylota ouelleti|[9009]|JWDCA493-10|BOLD:ABZ0281
Xylota ouelleti|[9010]|MHSYR317-07|BOLD:ABZ0281
Xylota ouelleti|[9011]|JWDCH546-10|BOLD:ABZ0281
Volucella|[9012]|CHDIP1444-12|BOLD:AAB8627
Neocnemodon rita|[9013]|MHFLY065-06|BOLD:AAG4653
Paragus haemorrhous|[9014]|JWDCB013-10|BOLD:AAC2439
Paragus haemorrhous|[9015]|MHSYR258-07|BOLD:AAC2439
Paragus haemorrhous|[9016]|MRACC002-07|BOLD:AAC2439
Toxomerus marginatus|[9017]|MHSYR099-07|BOLD:AAA4277
Toxomerus marginatus|[9018]|LRSYR046-07|BOLD:AAA4277
Sphaerophoria|[9019]|JWDCA483-10|BOLD:AAA7374
Sphaerophoria|[9020]|TWDIP022-09|BOLD:AAA7374
Sphaerophoria philanthus|[9021]|MHSYR028-06|BOLD:AAA7374
Sphaerophoria philanthus|[9022]|JWDCK240-11|BOLD:AAA7374
Sphaerophoria philanthus|[9023]|JWDCJ1404-11|BOLD:AAA7374
Sphaerophoria philanthus|[9024]|MHSYR302-07|BOLD:AAA7374
Sphaerophoria philanthus|[9025]|JWDCI418-10|BOLD:AAA7374
Sphaerophoria philanthus|[9026]|MHSYR034-06|BOLD:AAA7374
Sphaerophoria philanthus|[9027]|MHSYR102-07|BOLD:AAA7374
Sphaerophoria philanthus|[9028]|JWDCJ1406-11|BOLD:AAA7374
Sphaerophoria philanthus|[9029]|JWDCJ850-11|BOLD:AAA7374
Sphaerophoria philanthus|[9030]|JWDCG052-10|BOLD:AAA7374
Sphaerophoria philanthus|[9031]|JWDCK239-11|BOLD:AAA7374
Sphaerophoria philanthus|[9032]|JWDCK125-11|BOLD:AAA7374
Sphaerophoria philanthus|[9033]|JWDCH819-10|BOLD:AAA7374
Sphaerophoria|[9034]|JWDCB016-10|BOLD:AAA7374
Sphaerophoria philanthus|[9035]|MHSYR101-07|BOLD:AAA7374
Sphaerophoria philanthus|[9036]|MHSYR108-07|BOLD:AAA7374
Sphaerophoria philanthus|[9037]|MHSYR111-07|BOLD:AAA7374
Sphaerophoria philanthus|[9038]|JWDCE919-10|BOLD:AAA7374
Sphaerophoria philanthus|[9039]|JWDCL407-11|BOLD:AAA7374
Sphaerophoria|[9040]|JWDCG224-10|BOLD:AAA7374
Sphaerophoria|[9041]|JWDCH203-10|BOLD:AAA7374
Sphaerophoria philanthus|[9042]|LRSYR004-07|BOLD:AAA7374
Sphaerophoria philanthus|[9043]|LRSYR015-07|BOLD:AAA7374
Sphaerophoria philanthus|[9044]|MHSYR297-07|BOLD:AAA7374
Sphaerophoria philanthus|[9045]|MHSYR298-07|BOLD:AAA7374
Sphaerophoria philanthus|[9046]|MHSYR060-07|BOLD:AAA7374
Sphaerophoria philanthus|[9047]|LRSYR036-07|BOLD:AAA7374
Sphaerophoria|[9048]|TWDIP283-09|BOLD:AAA7374
Sphaerophoria|[9049]|JWDCE596-10|BOLD:AAA7374
Sphaerophoria philanthus|[9050]|MHSYR301-07|BOLD:AAA7374
Sphaerophoria philanthus|[9051]|MHSYR303-07|BOLD:AAA7374
Sphaerophoria philanthus|[9052]|MHFLI239-06|BOLD:AAA7374
Sphaerophoria philanthus|[9053]|MHSYR098-07|BOLD:AAA7374
Sphaerophoria|[9054]|JWDCD066-10|BOLD:AAA7374
Sphaerophoria philanthus|[9055]|JWDCE918-10|BOLD:AAA7374
[illegible]

Sphaerophoria philanthus|[9053]|MHSYR098-07|BOLD:AAA7374
Sphaerophoria philanthus|[9054]|JWDCD066-10|BOLD:AAA7374
Sphaerophoria philanthus|[9055]|JWDCE918-10|BOLD:AAA7374
Sphaerophoria|[9056]|TWDIP023-09|BOLD:AAA7374
Sphaerophoria philanthus|[9057]|JWDCH675-10|BOLD:AAA7374
Sphaerophoria|[9058]|JWDCE371-10|BOLD:AAA7374
Sphaerophoria philanthus|[9059]|JWDCJ733-11|BOLD:AAA7374
Sphaerophoria|[9060]|JWDCB017-10|BOLD:AAA7374
Sphaerophoria|[9061]|JWDCE598-10|BOLD:AAA7374
Sphaerophoria|[9062]|TWDIP067-09|BOLD:AAA7374
Sphaerophoria|[9063]|TWDIP024-09|BOLD:AAA7374
Sphaerophoria|[9064]|JWDCE370-10|BOLD:AAA7374
Sphaerophoria philanthus|[9065]|MHSYR300-07|BOLD:AAA7374
Sphaerophoria philanthus|[9066]|MHFLI094-06|BOLD:AAA7374
Sphaerophoria philanthus|[9067]|MHFLI214-06|BOLD:AAA7374
Sphaerophoria|[9068]|JWDCA489-10|BOLD:AAA7374
Sphaerophoria philanthus|[9069]|LRSYR035-07|BOLD:AAA7374
Sphaerophoria philanthus|[9070]|LRSYR038-07|BOLD:AAA7374
Sphaerophoria philanthus|[9071]|LRSYR039-07|BOLD:AAA7374
Sphaerophoria philanthus|[9072]|MHSYR095-07|BOLD:AAA7374
Sphaerophoria philanthus|[9073]|MHFLY127-07|BOLD:AAA7374
Sphaerophoria philanthus|[9074]|MHSYR296-07|BOLD:AAA7374
Sphaerophoria philanthus|[9075]|JWDCL247-11|BOLD:AAA7374
Sphaerophoria philanthus|[9076]|MHSYR105-07|BOLD:AAA7374
Sphaerophoria philanthus|[9077]|MHSYR222-07|BOLD:AAA7374
Sphaerophoria philanthus|[9078]|JSFLA011-08|BOLD:AAA7374
Sphaerophoria philanthus|[9079]|JWDCK236-11|BOLD:AAA7374
Sphaerophoria philanthus|[9080]|JWDCG053-10|BOLD:AAA7374
Sphaerophoria|[9081]|MHFLI888-09|BOLD:AAA7374
Sphaerophoria|[9082]|JWDCI008-10|BOLD:AAA7374
Sphaerophoria philanthus|[9083]|JWDCJ1402-11|BOLD:AAA7374
Sphaerophoria philanthus|[9084]|JWDCJ1484-11|BOLD:AAA7374
Sphaerophoria philanthus|[9085]|MHSYR100-07|BOLD:AAA7374
Sphaerophoria philanthus|[9086]|MHSYR059-07|BOLD:AAA7374
Sphaerophoria|[9087]|JWDCD136-10|BOLD:AAA7374
Sphaerophoria philanthus|[9088]|LRSYR030-07|BOLD:AAA7374
Sphaerophoria philanthus|[9089]|LRSYR045-07|BOLD:AAA7374
Chrysotoxum|[9090]|JWDCJ1257-11|BOLD:ABZ5562
Chrysotoxum|[9091]|JWDCJ1258-11|BOLD:ABZ5562
Chrysotoxum|[9092]|JWDCH553-10|BOLD:ABZ5562
Epistrophe terminalis|[9093]|JWDCC085-10|BOLD:AAN6464
Scaeva affinis|[9094]|JWDCJ402-11|BOLD:AAF2373
Lapposyrphus lapponicus|[9095]|MHFLI090-06|BOLD:AAB5561
Lapposyrphus lapponicus|[9096]|MHFLI266-06|BOLD:AAB5561
Lapposyrphus lapponicus|[9097]|MHSYR113-07|BOLD:AAB5561
Lapposyrphus lapponicus|[9098]|MHSYR158-07|BOLD:AAB5561
Lapposyrphus lapponicus|[9099]|MHSYR177-07|BOLD:AAB5561
Lapposyrphus lapponicus|[9100]|MHSYR210-07|BOLD:AAB5561
Lapposyrphus lapponicus|[9101]|JWDCE330-10|BOLD:AAB5561
Lapposyrphus lapponicus|[9102]|MHSYR120-07|BOLD:AAB5561
Lapposyrphus lapponicus|[9103]|JWDCK127-11|BOLD:AAB5561
Lapposyrphus lapponicus|[9104]|MHSYR121-07|BOLD:AAB5561
Lapposyrphus lapponicus|[9105]|LRSYR016-07|BOLD:AAB5561
Lapposyrphus lapponicus|[9106]|MHSYR103-07|BOLD:AAB5561
Lapposyrphus lapponicus|[9107]|JWDCK129-11|BOLD:AAB5561
Lapposyrphus lapponicus|[9108]|MHSYR123-07|BOLD:AAB5561
Lapposyrphus lapponicus|[9109]|JWDCE398-10|BOLD:AAB5561
Lapposyrphus lapponicus|[9110]|JWDCJ1000-11|BOLD:AAB5561
Lapposyrphus lapponicus|[9111]|LRSYR013-07|BOLD:AAB5561
Lapposyrphus|[9112]|JWDCB006-10|BOLD:AAB5561
Lapposyrphus|[9113]|JSFLA020-08|BOLD:AAB5561
Lapposyrphus lapponicus|[9114]|LRSYR026-07|BOLD:AAB5561
Lapposyrphus lapponicus|[9115]|LRSYR027-07|BOLD:AAB5561
Lapposyrphus lapponicus|[9116]|LRSYR031-07|BOLD:AAB5561
Lapposyrphus lapponicus|[9117]|LRSYR032-07|BOLD:AAB5561
Lapposyrphus lapponicus|[9118]|LRSYR044-07|BOLD:AAB5561
Melangyna labiatarum|[9119]|MHSYR106-07|BOLD:AAB5562
Melangyna labiatarum|[9120]|JWDCA488-10|BOLD:AAB5562
Melangyna arctica|[9121]|TWDIP038-09|BOLD:AAG4664
Melangyna arctica|[9122]|MHSYR209-07|BOLD:AAG4664
Melangyna arctica|[9123]|JWDCJ851-11|BOLD:AAG4664
Melangyna arctica|[9124]|LRSYR007-07|BOLD:AAG4664
Melangyna arctica|[9125]|TWDIP035-09|BOLD:AAG4664
Melangyna arctica|[9126]|JWDCH674-10|BOLD:AAG4664
Melangyna arctica|[9127]|MHSYR287-07|BOLD:AAG4664
Melangyna arctica|[9128]|TWDIP036-09|BOLD:AAG4664
Melangyna arctica|[9129]|JWDCI881-11|BOLD:AAG4664
Melangyna arctica|[9130]|LRSYR006-07|BOLD:AAG4664
Melangyna arctica|[9131]|LRSYR009-07|BOLD:AAG4664
Eupeodes latifasciatus|[9132]|MHSYR243-07|BOLD:ADJ8356
Eupeodes latifasciatus|[9133]|MHSYR241-07|BOLD:ADJ8356
Eupeodes latifasciatus|[9134]|MHSYR242-07|BOLD:ADJ8356
Eupeodes latifasciatus|[9135]|MHSYR240-07|BOLD:ADJ8356
Eupeodes|[9136]|JWDCB007-10|BOLD:AAB2384
Eupeodes luniger|[9137]|MHSYR124-07|BOLD:AAB2384
Eupeodes luniger|[9138]|MHSYR134-07|BOLD:AAB2384
Eupeodes|[9139]|JWDCL246-11|BOLD:AAB2384
Eupeodes|[9140]|TWDIP027-09|BOLD:AAB2384
Eupeodes|[9141]|MHSYR234-07|BOLD:AAB2384
Eupeodes curtus|[9142]|MHSYR201-07|BOLD:AAB2384
Eupeodes|[9143]|JSFLA025-08|BOLD:AAB2384
Eupeodes|[9144]|JWDCE478-10|BOLD:AAB2384
Eupeodes curtus|[9145]|MHSYR031-06|BOLD:AAB2384
Eupeodes|[9146]|JWDCG310-10|BOLD:AAB2384
Eupeodes americanus|[9147]|JWDCI815-11|BOLD:AAB2384
Eupeodes|[9148]|JWDCC173-10|BOLD:AAB2384
Eupeodes curtus|[9149]|MHFLI092-06|BOLD:AAB2384
Eupeodes curtus|[9150]|MHSYR169-07|BOLD:AAB2384
Eupeodes|[9151]|JWDCC256-10|BOLD:AAB2384
Eupeodes|[9152]|JWDCD041-10|BOLD:AAB2384
Eupeodes latifasciatus|[9153]|MRACC001-07|BOLD:AAB2384
Eupeodes|[9154]|MRACC004-07|BOLD:AAB2384
Eupeodes|[9155]|MRACC003-07|BOLD:AAB2384

Eupeodes latifasciatus|[9153]|MRACC001-07|BOLD:AAB2384
Eupeodes|[9154]|MRACC004-07|BOLD:AAB2384
Eupeodes|[9155]|MRACC003-07|BOLD:AAB2384
Eupeodes|[9156]|MRACC009-07|BOLD:AAB2384
Eupeodes|[9157]|JWDCK126-11|BOLD:AAB2384
Eupeodes|[9158]|JWDCH672-10|BOLD:AAB2384
Eupeodes luniger|[9159]|MHSYR236-07|BOLD:AAB2384
Eupeodes flukei|[9160]|MHSYR235-07|BOLD:AAB2384
Eupeodes curtus|[9161]|MHSYR030-06|BOLD:AAB2384
Eupeodes latifasciatus|[9162]|MHSYR118-07|BOLD:AAB2384
Eupeodes luniger|[9163]|MHSYR116-07|BOLD:AAB2384
Eupeodes|[9164]|MRACC007-07|BOLD:AAB2384
Eupeodes|[9165]|MRACC006-07|BOLD:AAB2384
Eupeodes|[9166]|JSFLA022-08|BOLD:AAB2384
Eupeodes luniger|[9167]|MHSYR126-07|BOLD:AAB2384
Eupeodes luniger|[9168]|MHSYR125-07|BOLD:AAB2384
Eupeodes luniger|[9169]|MHSYR096-07|BOLD:AAB2384
Eupeodes|[9170]|JWDCJ1048-11|BOLD:AAB2384
Eupeodes|[9171]|TWDIP028-09|BOLD:AAB2384
Eupeodes luniger|[9172]|MHSYR119-07|BOLD:AAB2384
Eupeodes|[9173]|JWDCH141-10|BOLD:AAB2384
Eupeodes luniger|[9174]|MHFLI093-06|BOLD:AAB2384
Eupeodes|[9175]|LRSYR014-07|BOLD:AAB2384
Eupeodes|[9176]|LRSYR043-07|BOLD:AAB2384
Eupeodes curtus|[9177]|MHSYR114-07|BOLD:AAB2384
Eupeodes|[9178]|JWDCI880-11|BOLD:AAB2384
Eupeodes luniger|[9179]|MHSYR208-07|BOLD:AAB2384
Eupeodes luniger|[9180]|MHSYR237-07|BOLD:AAB2384
Eupeodes|[9181]|JWDCC834-10|BOLD:AAB2384
Eupeodes|[9182]|JWDCH258-10|BOLD:AAB2384
Eupeodes montivagus|[9183]|MHSYR238-07|BOLD:AAB2384
Eupeodes|[9184]|JWDCJ1485-11|BOLD:AAB2384
Eupeodes|[9185]|JWDCJ1407-11|BOLD:AAB2384
Eupeodes|[9186]|JWDCJ1400-11|BOLD:AAB2384
Eupeodes|[9187]|JWDCF755-10|BOLD:AAB2384
Eupeodes|[9188]|JWDCF679-10|BOLD:AAB2384
Eupeodes|[9189]|JWDCJ1401-11|BOLD:AAB2384
Eupeodes|[9190]|JWDCC171-10|BOLD:AAB2384
Eupeodes flukei|[9191]|MHSYR239-07|BOLD:AAB2384
Eupeodes latifasciatus|[9192]|MHSYR244-07|BOLD:AAB2384
Parasyrphus genualis|[9193]|LRSYR008-07|BOLD:AAG4652
Parasyrphus relictus|[9194]|JWDCE441-10|BOLD:AAL3233
Parasyrphus relictus|[9195]|MHSYR007-06|BOLD:AAL3233
Parasyrphus tarsatus|[9196]|LSCCH025-07|BOLD:AAC1834
Parasyrphus tarsatus|[9197]|LRSYR088-07|BOLD:AAC1834
Parasyrphus|[9198]|TWDIP282-09|BOLD:AAC1834
Parasyrphus|[9199]|TWDIP029-09|BOLD:AAC1834
Parasyrphus|[9200]|TWDIP026-09|BOLD:AAC1834
Parasyrphus tarsatus|[9201]|JWDCB010-10|BOLD:AAC1834
Parasyrphus tarsatus|[9202]|JSFLA026-08|BOLD:AAC1834
Parasyrphus tarsatus|[9203]|JWDCC005-10|BOLD:AAC1834
Parasyrphus|[9204]|JSFLA004-08|BOLD:AAC1834
Parasyrphus tarsatus|[9205]|MHSYR247-07|BOLD:AAC1834
Parasyrphus tarsatus|[9206]|MHSYR262-07|BOLD:AAC1834
Parasyrphus tarsatus|[9207]|MHSYR260-07|BOLD:AAC1834
Parasyrphus tarsatus|[9208]|JWDCB009-10|BOLD:AAC1834
Parasyrphus tarsatus|[9209]|JWDCE329-10|BOLD:AAC1834
Parasyrphus tarsatus|[9210]|MHSYR261-07|BOLD:AAC1834
Parasyrphus tarsatus|[9211]|MHSYR263-07|BOLD:AAC1834
Parasyrphus tarsatus|[9212]|MHSYR225-07|BOLD:AAC1834
Parasyrphus nigritarsis|[9213]|MHSYR259-07|BOLD:AAL3230
Syrphus vitripennis|[9214]|LRSYR055-07|BOLD:AAB5577
Syrphus vitripennis|[9215]|LRSYR066-07|BOLD:AAB5577
Syrphus vitripennis|[9216]|LRSYR073-07|BOLD:AAB5577
Syrphus vitripennis|[9217]|MHSYR159-07|BOLD:AAB5577
Syrphus vitripennis|[9218]|JWDCJ1049-11|BOLD:AAB5577
Syrphus vitripennis|[9219]|MHSYR112-07|BOLD:AAB5577
Syrphus vitripennis|[9220]|MHSYR097-07|BOLD:AAB5577
Syrphus vitripennis|[9221]|JWDCI609-10|BOLD:AAB5577
Syrphus vitripennis|[9222]|MHSYR032-06|BOLD:AAB5577
Syrphus vitripennis|[9223]|MHSYR127-07|BOLD:AAB5577
Syrphus vitripennis|[9224]|MHSYR171-07|BOLD:AAB5577
Syrphus vitripennis|[9225]|JWDCE477-10|BOLD:AAB5577
Syrphus vitripennis|[9226]|MHSYR184-07|BOLD:AAB5577
Syrphus vitripennis|[9227]|JWDCH522-10|BOLD:AAB5577
Syrphus vitripennis|[9228]|MHSYR175-07|BOLD:AAB5577
Syrphus vitripennis|[9229]|MHSYR198-07|BOLD:AAB5577
Syrphus vitripennis|[9230]|LRSYR010-07|BOLD:AAB5577
Syrphus vitripennis|[9231]|LRSYR041-07|BOLD:AAB5577
Syrphus vitripennis|[9232]|LRSYR069-07|BOLD:AAB5577
Syrphus vitripennis|[9233]|LRSYR087-07|BOLD:AAB5577
Syrphus attenuatus|[9234]|JWDCJ1482-11|BOLD:AAD7605
Syrphus attenuatus|[9235]|MHSYR135-07|BOLD:AAD7605
Syrphus attenuatus|[9236]|MHSYR036-06|BOLD:AAD7605
Syrphus attenuatus|[9237]|JWDCE442-10|BOLD:AAD7605
Syrphus attenuatus|[9238]|MHSYR164-07|BOLD:AAD7605
Syrphus attenuatus|[9239]|MHSYR166-07|BOLD:AAD7605
Syrphus attenuatus|[9240]|MHSYR189-07|BOLD:AAD7605
Syrphus torvus|[9241]|LRSYR033-07|BOLD:AAC6088
Syrphus ribesii|[9242]|LRSYR077-07|BOLD:AAA4570
Syrphus ribesii|[9243]|LRSYR075-07|BOLD:AAA4570
Syrphus ribesii|[9244]|LRSYR085-07|BOLD:AAA4570
Syrphus ribesii|[9245]|DSSAW002-06|BOLD:AAA4570
Syrphus ribesii|[9246]|DSSAW070-06|BOLD:AAA4570
Syrphus ribesii|[9247]|DSSAW006-06|BOLD:AAA4570
Syrphus ribesii|[9248]|DSSAW005-06|BOLD:AAA4570
Syrphus ribesii|[9249]|DSSAW004-06|BOLD:AAA4570
Syrphus ribesii|[9250]|DSSAW003-06|BOLD:AAA4570
Syrphus ribesii|[9251]|DSSAW001-06|BOLD:AAA4570
Syrphus ribesii|[9252]|MHSYR037-06|BOLD:AAA4570
Syrphus ribesii|[9253]|LRSYR042-07|BOLD:AAA4570
Syrphus ribesii|[9254]|JWDCJ1483-11|BOLD:AAA4570

Syrphus ribesii|[9252]|MHSYR037-06|BOLD:AAA4570
Syrphus ribesii|[9253]|LRSYR042-07|BOLD:AAA4570
Syrphus ribesii|[9254]|JWDCJ1483-11|BOLD:AAA4570
Syrphus ribesii|[9255]|LRSYR068-07|BOLD:AAA4570
Syrphus ribesii|[9256]|LRSYR058-07|BOLD:AAA4570
Syrphus ribesii|[9257]|LRSYR071-07|BOLD:AAA4570
Syrphus ribesii|[9258]|LRSYR076-07|BOLD:AAA4570
Syrphus ribesii|[9259]|MHSYR304-07|BOLD:AAA4570
Syrphus ribesii|[9260]|LRSYR065-07|BOLD:AAA4570
Syrphus ribesii|[9261]|JWDCK238-11|BOLD:AAA4570
Syrphus ribesii|[9262]|MHSYR136-07|BOLD:AAA4570
Syrphus ribesii|[9263]|MHSYR004-06|BOLD:AAA4570
Syrphus ribesii|[9264]|MHSYR192-07|BOLD:AAA4570
Syrphus ribesii|[9265]|MHSYR305-07|BOLD:AAA4570
Syrphus ribesii|[9266]|MRACC008-07|BOLD:AAA4570
Syrphus ribesii|[9267]|MHSYR130-07|BOLD:AAA4570
Syrphus ribesii|[9268]|MHSYR006-06|BOLD:AAA4570
Syrphus ribesii|[9269]|MHSYR115-07|BOLD:AAA4570
Syrphus ribesii|[9270]|MHSYR178-07|BOLD:AAA4570
Syrphus ribesii|[9271]|MHSYR183-07|BOLD:AAA4570
Syrphus ribesii|[9272]|MHSYR205-07|BOLD:AAA4570
Syrphus ribesii|[9273]|MHSYR035-06|BOLD:AAA4570
Syrphus ribesii|[9274]|MHSYR199-07|BOLD:AAA4570
Syrphus ribesii|[9275]|LRSYR082-07|BOLD:AAA4570
Syrphus ribesii|[9276]|MHSYR122-07|BOLD:AAA4570
Syrphus ribesii|[9277]|MHSYR128-07|BOLD:AAA4570
Syrphus ribesii|[9278]|MHSYR172-07|BOLD:AAA4570
Syrphus ribesii|[9279]|LRSYR062-07|BOLD:AAA4570
Syrphus ribesii|[9280]|MHSYR204-07|BOLD:AAA4570
Syrphus ribesii|[9281]|MHSYR187-07|BOLD:AAA4570
Syrphus ribesii|[9282]|MHSYR132-07|BOLD:AAA4570
Syrphus ribesii|[9283]|LRSYR072-07|BOLD:AAA4570
Syrphus ribesii|[9284]|LRSYR086-07|BOLD:AAA4570
Syrphus ribesii|[9285]|LRSYR078-07|BOLD:AAA4570
Syrphus ribesii|[9286]|JWDCI817-11|BOLD:AAA4570
Syrphus ribesii|[9287]|MHSYR133-07|BOLD:AAA4570
Syrphus ribesii|[9288]|MHSYR131-07|BOLD:AAA4570
Syrphus ribesii|[9289]|LRSYR079-07|BOLD:AAA4570
Syrphus ribesii|[9290]|JWDCF446-10|BOLD:AAA4570
Syrphus ribesii|[9291]|MHSYR005-06|BOLD:AAA4570
Syrphus ribesii|[9292]|MHSYR003-06|BOLD:AAA4570
Syrphus ribesii|[9293]|MHSYR002-06|BOLD:AAA4570
Syrphus ribesii|[9294]|MHFLI089-06|BOLD:AAA4570
Syrphus ribesii|[9295]|MHSYR001-06|BOLD:AAA4570
Syrphus ribesii|[9296]|MHSYR160-07|BOLD:AAA4570
Syrphus ribesii|[9297]|MHSYR170-07|BOLD:AAA4570
Syrphus ribesii|[9298]|MHSYR188-07|BOLD:AAA4570
Syrphus ribesii|[9299]|MHSYR193-07|BOLD:AAA4570
Syrphus ribesii|[9300]|MHSYR202-07|BOLD:AAA4570
Syrphus ribesii|[9301]|MHSYR206-07|BOLD:AAA4570
Syrphus ribesii|[9302]|MHSYR207-07|BOLD:AAA4570
Syrphus ribesii|[9303]|LRSYR070-07|BOLD:AAA4570
Syrphus ribesii|[9304]|LRSYR081-07|BOLD:AAA4570
Syrphus ribesii|[9305]|LRSYR083-07|BOLD:AAA4570
Syrphus ribesii|[9306]|LRSYR064-07|BOLD:AAA4570
Syrphus ribesii|[9307]|LRSYR074-07|BOLD:AAA4570
Syrphus ribesii|[9308]|LRSYR080-07|BOLD:AAA4570
Syrphus ribesii|[9309]|LRSYR089-07|BOLD:AAA4570
Syrphus ribesii|[9310]|MHSYR110-07|BOLD:AAA4570
Syrphus ribesii|[9311]|MHSYR117-07|BOLD:AAA4570
Syrphus ribesii|[9312]|MHSYR168-07|BOLD:AAA4570
Syrphus ribesii|[9313]|LRSYR063-07|BOLD:AAA4570
Syrphus ribesii|[9314]|LRSYR067-07|BOLD:AAA4570
Syrphus ribesii|[9315]|MHSYR129-07|BOLD:AAA4570
Syrphus ribesii|[9316]|MHSYR033-06|BOLD:AAA4570
Syrphus ribesii|[9317]|MHSYR190-07|BOLD:AAA4570
Syrphus ribesii|[9318]|JWDCB020-10|BOLD:AAA4570
Syrphus ribesii|[9319]|MHSYR194-07|BOLD:AAA4570
Syrphus ribesii|[9320]|MHSYR197-07|BOLD:AAA4570
Syrphus ribesii|[9321]|MHSYR182-07|BOLD:AAA4570
Eristalis brousii|[9322]|MHSYR407-07|BOLD:AAA8223
Eristalis brousii|[9323]|MHSYR334-07|BOLD:AAA8223
Eristalis brousii|[9324]|MHSYR338-07|BOLD:AAA8223
Eristalis brousii|[9325]|MHSYR335-07|BOLD:AAA8223
Eristalis brousii|[9326]|MHSYR400-07|BOLD:AAA8223
Eristalis brousii|[9327]|MHSYR399-07|BOLD:AAA8223
Eristalis brousii|[9328]|MHSYR398-07|BOLD:AAA8223
Eristalis brousii|[9329]|MHSYR397-07|BOLD:AAA8223
Eristalis brousii|[9330]|MHSYR396-07|BOLD:AAA8223
Eristalis brousii|[9331]|MHSYR341-07|BOLD:AAA8223
Eristalis brousii|[9332]|MHSYR339-07|BOLD:AAA8223
Eristalis brousii|[9333]|MHSYR332-07|BOLD:AAA8223
Eristalis brousii|[9334]|MHSYR337-07|BOLD:AAA8223
Eristalis brousii|[9335]|MHSYR403-07|BOLD:AAA8223
Eristalis brousii|[9336]|MHSYR406-07|BOLD:AAA8223
Eristalis brousii|[9337]|MHSYR024-06|BOLD:AAA8223
Eristalis brousii|[9338]|MHSYR401-07|BOLD:AAA8223
Eristalis brousii|[9339]|MHSYR402-07|BOLD:AAA8223
Eristalis brousii|[9340]|MHSYR409-07|BOLD:AAA8223
Eristalis brousii|[9341]|MHSYR410-07|BOLD:AAA8223
Eristalis brousii|[9342]|MHSYR217-07|BOLD:AAA8223
Eristalis brousii|[9343]|JWDCF332-10|BOLD:AAA8223
Eristalis brousii|[9344]|LRSYR025-07|BOLD:AAA8223
Eristalis brousii|[9345]|LRSYR040-07|BOLD:AAA8223
Eristalis dimidiata|[9346]|MHSYR329-07|BOLD:AAB1409
Eristalis dimidiata|[9347]|MHSYR027-06|BOLD:AAB1409
Eristalis dimidiata|[9348]|MHSYR143-07|BOLD:AAB1409
Eristalis dimidiata|[9349]|JWDCG569-10|BOLD:AAB1409
Eristalis dimidiata|[9350]|JWDCG568-10|BOLD:AAB1409
Eristalis dimidiata|[9351]|JWDCG567-10|BOLD:AAB1409
Eristalis dimidiata|[9352]|MHSYR146-07|BOLD:AAB1409
Eristalis dimidiata|[9353]|MHSYR218-07|BOLD:AAB1409
Eristalis dimidiata|[9354]|LRSYR051-07|BOLD:AAB1409

Eristalis dimidiata|[9352]|MHSYR146-07|BOLD:AAB1409
Eristalis dimidiata|[9353]|MHSYR218-07|BOLD:AAB1409
Eristalis dimidiata|[9354]|LRSYR051-07|BOLD:AAB1409
Eristalis dimidiata|[9355]|JSFLA016-08|BOLD:AAB1409
Eristalis dimidiata|[9356]|JSFLA018-08|BOLD:AAB1409
Eristalis dimidiata|[9357]|JWDCG483-10|BOLD:AAB1409
Eristalis cryptarum|[9358]|JWDCG484-10|BOLD:AAC9280
Eristalis obscura|[9359]|MHSYR358-07|BOLD:AAA6459
Eristalis obscura|[9360]|MHSYR386-07|BOLD:AAA6459
Eristalis obscura|[9361]|MHSYR371-07|BOLD:AAA6459
Eristalis obscura|[9362]|MHSYR388-07|BOLD:AAA6459
Eristalis obscura|[9363]|MHSYR374-07|BOLD:AAA6459
Eristalis obscura|[9364]|LRSYR052-07|BOLD:AAA6459
Eristalis obscura|[9365]|MHSYR144-07|BOLD:AAA6459
Eristalis obscura|[9366]|MHSYR366-07|BOLD:AAA6459
Eristalis obscura|[9367]|MHSYR370-07|BOLD:AAA6459
Eristalis obscura|[9368]|MHSYR376-07|BOLD:AAA6459
Eristalis obscura|[9369]|MHSYR359-07|BOLD:AAA6459
Eristalis obscura|[9370]|MHSYR372-07|BOLD:AAA6459
Eristalis obscura|[9371]|MHSYR360-07|BOLD:AAA6459
Eristalis obscura|[9372]|MHSYR383-07|BOLD:AAA6459
Eristalis obscura|[9373]|MHSYR351-07|BOLD:AAA6459
Eristalis obscura|[9374]|MHSYR186-07|BOLD:AAA6459
Eristalis obscura|[9375]|MHSYR352-07|BOLD:AAA6459
Eristalis obscura|[9376]|MHSYR353-07|BOLD:AAA6459
Eristalis obscura|[9377]|MHSYR362-07|BOLD:AAA6459
Eristalis obscura|[9378]|MHSYR363-07|BOLD:AAA6459
Eristalis obscura|[9379]|MHSYR364-07|BOLD:AAA6459
Eristalis obscura|[9380]|MHSYR365-07|BOLD:AAA6459
Eristalis obscura|[9381]|MHSYR368-07|BOLD:AAA6459
Eristalis obscura|[9382]|MHSYR369-07|BOLD:AAA6459
Eristalis obscura|[9383]|MHSYR373-07|BOLD:AAA6459
Eristalis obscura|[9384]|MHSYR377-07|BOLD:AAA6459
Eristalis obscura|[9385]|MHSYR379-07|BOLD:AAA6459
Eristalis obscura|[9386]|MHSYR381-07|BOLD:AAA6459
Eristalis obscura|[9387]|MHSYR382-07|BOLD:AAA6459
Eristalis obscura|[9388]|MHSYR384-07|BOLD:AAA6459
Eristalis obscura|[9389]|MHSYR385-07|BOLD:AAA6459
Eristalis obscura|[9390]|MHSYR387-07|BOLD:AAA6459
Eristalis obscura|[9391]|MHSYR389-07|BOLD:AAA6459
Eristalis obscura|[9392]|MHSYR391-07|BOLD:AAA6459
Eristalis obscura|[9393]|MHSYR393-07|BOLD:AAA6459
Eristalis obscura|[9394]|MHSYR394-07|BOLD:AAA6459
Eristalis obscura|[9395]|MHSYR404-07|BOLD:AAA6459
Eristalis obscura|[9396]|MHSYR408-07|BOLD:AAA6459
Eristalis obscura|[9397]|MHSYR142-07|BOLD:AAA6459
Eristalis obscura|[9398]|LRSYR019-07|BOLD:AAA6459
Eristalis obscura|[9399]|MHSYR180-07|BOLD:AAA6459
Eristalis obscura|[9400]|MHSYR185-07|BOLD:AAA6459
Eristalis obscura|[9401]|LRSYR020-07|BOLD:AAA6459
Eristalis obscura|[9402]|LRSYR028-07|BOLD:AAA6459
Eristalis fraterculus|[9403]|MHSYR330-07|BOLD:AAD7912
Eristalis fraterculus|[9404]|MHSYR322-07|BOLD:AAD7912
Eristalis fraterculus|[9405]|JSFLA027-08|BOLD:AAD7912
Eristalis fraterculus|[9406]|LRSYR049-07|BOLD:AAD7912
Eristalis flavipes|[9407]|MHSYR140-07|BOLD:AAB9658
Eristalis flavipes|[9408]|JSFLA001-08|BOLD:AAB9658
Eristalis anthophorina|[9409]|MHSYR072-07|BOLD:AAB6090
Eristalis anthophorina|[9410]|JWDCH120-10|BOLD:AAB6090
Eristalis anthophorina|[9411]|MHSYR062-07|BOLD:AAB6090
Eristalis anthophorina|[9412]|TWDIP068-09|BOLD:AAB6090
Eristalis anthophorina|[9413]|MHSYR066-07|BOLD:AAB6090
Eristalis anthophorina|[9414]|MHSYR137-07|BOLD:AAB6090
Eristalis anthophorina|[9415]|MHSYR138-07|BOLD:AAB6090
Eristalis anthophorina|[9416]|JWDCH673-10|BOLD:AAB6090
Eristalis anthophorina|[9417]|MHSYR191-07|BOLD:AAB6090
Eristalis anthophorina|[9418]|LRSYR050-07|BOLD:AAB6090
Eristalis tenax|[9419]|MHSYR077-07|BOLD:AAB0391
Eristalis tenax|[9420]|MHSYR093-07|BOLD:AAB0391
Eristalis tenax|[9421]|MHSYR026-06|BOLD:AAB0391
Eristalis tenax|[9422]|MHSYR054-07|BOLD:AAB0391
Helophilus affinis|[9423]|MHSYR013-06|BOLD:AAA3292
Helophilus affinis|[9424]|MHSYR165-07|BOLD:AAA3292
Helophilus affinis|[9425]|MHSYR196-07|BOLD:AAA3292
Helophilus affinis|[9426]|MHSYR203-07|BOLD:AAA3292
Helophilus affinis|[9427]|MHSYR162-07|BOLD:AAA3292
Helophilus groenlandicus|[9428]|MHCOL066-07|BOLD:AAB1983
Helophilus lapponicus|[9429]|JWDCD125-10|BOLD:ACE4226
Helophilus lapponicus|[9430]|JWDCI150-10|BOLD:ACE4226
Helophilus lapponicus|[9431]|JWDCJ612-11|BOLD:ACE4226
Helophilus lapponicus|[9432]|JWDCI151-10|BOLD:ACE4226
Helophilus lapponicus|[9433]|MHSYR023-06|BOLD:ACE4226
Helophilus lapponicus|[9434]|MHSYR380-07|BOLD:ACE4226
Helophilus lapponicus|[9435]|JWDCJ610-11|BOLD:ACE4226
Helophilus lapponicus|[9436]|JWDCJ611-11|BOLD:ACE4226
Helophilus lapponicus|[9437]|JWDCB004-10|BOLD:ACE4226
Helophilus lapponicus|[9438]|JWDCI149-10|BOLD:ACE4226
Helophilus lapponicus|[9439]|JWDCJ542-11|BOLD:ACE4226
Helophilus lapponicus|[9440]|JWDCC857-10|BOLD:ACE4226
Helophilus lapponicus|[9441]|JSFLA019-08|BOLD:ACE4226
Helophilus lapponicus|[9442]|MHSYR405-07|BOLD:ACE4226
Helophilus lapponicus|[9443]|LRSYR029-07|BOLD:ACE4226
Helophilus groenlandicus|[9444]|MHSYR336-07|BOLD:AAB1982
Helophilus groenlandicus|[9445]|HMCOL004-07|BOLD:AAB1982
Helophilus groenlandicus|[9446]|HMCOL010-07|BOLD:AAB1982
Helophilus groenlandicus|[9447]|HMCOL007-07|BOLD:AAB1982
Helophilus groenlandicus|[9448]|HMCOL005-07|BOLD:AAB1982
Helophilus groenlandicus|[9449]|MHSYR411-07|BOLD:AAB1982
Helophilus groenlandicus|[9450]|MHSYR079-07|BOLD:AAB1982
Helophilus groenlandicus|[9451]|HMCOL009-07|BOLD:AAB1982
Helophilus groenlandicus|[9452]|HMCOL003-07|BOLD:AAB1982
Helophilus groenlandicus|[9453]|MHSYR055-07|BOLD:AAB1982
[illegible]

Helophilus groenlandicus|[9451]|HMCOL009-07|BOLD:AAB1982
Helophilus groenlandicus|[9452]|HMCOL003-07|BOLD:AAB1982
Helophilus groenlandicus|[9453]|MHSYR055-07|BOLD:AAB1982
Helophilus groenlandicus|[9454]|MHCOL067-07|BOLD:AAB1982
Helophilus groenlandicus|[9455]|MHCOL068-07|BOLD:AAB1982
Helophilus groenlandicus|[9456]|MHCOL070-07|BOLD:AAB1982
Helophilus groenlandicus|[9457]|MHSYR020-06|BOLD:AAB1982
Helophilus groenlandicus|[9458]|MHSYR145-07|BOLD:AAB1982
Helophilus groenlandicus|[9459]|HMCOL006-07|BOLD:AAB1982
Helophilus groenlandicus|[9460]|MHCOL065-07|BOLD:AAB1982
Helophilus groenlandicus|[9461]|MHCOL069-07|BOLD:AAB1982
Helophilus groenlandicus|[9462]|MHSYR057-07|BOLD:AAB1982
Helophilus groenlandicus|[9463]|MHSYR025-06|BOLD:AAB1982
Helophilus groenlandicus|[9464]|HMCOL002-07|BOLD:AAB1982
Helophilus groenlandicus|[9465]|HMCOL008-07|BOLD:AAB1982
Helophilus groenlandicus|[9466]|LRSYR021-07|BOLD:AAB1982
Helophilus groenlandicus|[9467]|HMCOL001-07|BOLD:AAB1982
Helophilus groenlandicus|[9468]|JSFLA017-08|BOLD:AAB1982
Helophilus groenlandicus|[9469]|MHSYR056-07|BOLD:AAB1982
Helophilus obscurus|[9470]|MHSYR016-06|BOLD:AAA3291
Helophilus obscurus|[9471]|MHSYR008-06|BOLD:AAA3291
Helophilus obscurus|[9472]|MHSYR014-06|BOLD:AAA3291
Helophilus obscurus|[9473]|MHSYR221-07|BOLD:AAA3291
Helophilus obscurus|[9474]|MHSYR149-07|BOLD:AAA3291
Helophilus obscurus|[9475]|MHSYR063-07|BOLD:AAA3291
Helophilus obscurus|[9476]|MHSYR075-07|BOLD:AAA3291
Helophilus obscurus|[9477]|MHSYR195-07|BOLD:AAA3291
Helophilus obscurus|[9478]|MHSYR010-06|BOLD:AAA3291
Helophilus obscurus|[9479]|MHSYR048-07|BOLD:AAA3291
Helophilus obscurus|[9480]|MHSYR045-07|BOLD:AAA3291
Helophilus obscurus|[9481]|MHSYR040-07|BOLD:AAA3291
Helophilus obscurus|[9482]|MHSYR156-07|BOLD:AAA3291
Helophilus obscurus|[9483]|MHSYR153-07|BOLD:AAA3291
Helophilus obscurus|[9484]|MHSYR139-07|BOLD:AAA3291
Helophilus obscurus|[9485]|MHSYR094-07|BOLD:AAA3291
Helophilus obscurus|[9486]|MHSYR082-07|BOLD:AAA3291
Helophilus obscurus|[9487]|MHSYR078-07|BOLD:AAA3291
Helophilus obscurus|[9488]|MHSYR015-06|BOLD:AAA3291
Helophilus obscurus|[9489]|MHSYR012-06|BOLD:AAA3291
Helophilus obscurus|[9490]|MHSYR009-06|BOLD:AAA3291
Helophilus obscurus|[9491]|MHSYR021-06|BOLD:AAA3291
Helophilus obscurus|[9492]|MHSYR223-07|BOLD:AAA3291
Helophilus obscurus|[9493]|MHSYR157-07|BOLD:AAA3291
Helophilus obscurus|[9494]|MHSYR167-07|BOLD:AAA3291
Helophilus obscurus|[9495]|MHSYR053-07|BOLD:AAA3291
Helophilus obscurus|[9496]|MHSYR083-07|BOLD:AAA3291
Helophilus obscurus|[9497]|MHSYR092-07|BOLD:AAA3291
Helophilus obscurus|[9498]|MHSYR046-07|BOLD:AAA3291
Helophilus obscurus|[9499]|MHSYR200-07|BOLD:AAA3291
Helophilus obscurus|[9500]|MHSYR085-07|BOLD:AAA3291
Helophilus obscurus|[9501]|MHSYR081-07|BOLD:AAA3291
Helophilus obscurus|[9502]|MHSYR214-07|BOLD:AAA3291
Helophilus obscurus|[9503]|MHSYR043-07|BOLD:AAA3291
Helophilus obscurus|[9504]|MHSYR042-07|BOLD:AAA3291
Helophilus obscurus|[9505]|MHSYR039-07|BOLD:AAA3291
Helophilus obscurus|[9506]|LRSYR024-07|BOLD:AAA3291
Helophilus obscurus|[9507]|MHSYR061-07|BOLD:AAA3291
Helophilus obscurus|[9508]|MHSYR050-07|BOLD:AAA3291
Helophilus obscurus|[9509]|MHSYR049-07|BOLD:AAA3291
Helophilus obscurus|[9510]|MHSYR044-07|BOLD:AAA3291
Helophilus obscurus|[9511]|MHSYR038-07|BOLD:AAA3291
Helophilus obscurus|[9512]|LRSYR056-07|BOLD:AAA3291
Helophilus obscurus|[9513]|MHSYR220-07|BOLD:AAA3291
Helophilus obscurus|[9514]|MHSYR216-07|BOLD:AAA3291
Helophilus obscurus|[9515]|MHSYR215-07|BOLD:AAA3291
Helophilus obscurus|[9516]|MHSYR213-07|BOLD:AAA3291
Helophilus obscurus|[9517]|MHSYR212-07|BOLD:AAA3291
Helophilus obscurus|[9518]|MHSYR211-07|BOLD:AAA3291
Helophilus obscurus|[9519]|MHSYR181-07|BOLD:AAA3291
Helophilus obscurus|[9520]|MHSYR179-07|BOLD:AAA3291
Helophilus obscurus|[9521]|MHSYR174-07|BOLD:AAA3291
Helophilus obscurus|[9522]|MHSYR163-07|BOLD:AAA3291
Helophilus obscurus|[9523]|MHSYR161-07|BOLD:AAA3291
Helophilus obscurus|[9524]|MHSYR041-07|BOLD:AAA3291
Helophilus obscurus|[9525]|MHSYR091-07|BOLD:AAA3291
Helophilus obscurus|[9526]|MHSYR090-07|BOLD:AAA3291
Helophilus obscurus|[9527]|MHSYR155-07|BOLD:AAA3291
Helophilus obscurus|[9528]|MHSYR154-07|BOLD:AAA3291
Helophilus obscurus|[9529]|MHSYR150-07|BOLD:AAA3291
Helophilus obscurus|[9530]|MHSYR147-07|BOLD:AAA3291
Helophilus obscurus|[9531]|MHSYR089-07|BOLD:AAA3291
Helophilus obscurus|[9532]|MHSYR088-07|BOLD:AAA3291
Helophilus obscurus|[9533]|MHSYR087-07|BOLD:AAA3291
Helophilus obscurus|[9534]|MHSYR086-07|BOLD:AAA3291
Helophilus obscurus|[9535]|MHSYR084-07|BOLD:AAA3291
Helophilus obscurus|[9536]|MHSYR080-07|BOLD:AAA3291
Helophilus obscurus|[9537]|MHSYR076-07|BOLD:AAA3291
Helophilus obscurus|[9538]|MHSYR073-07|BOLD:AAA3291
Helophilus obscurus|[9539]|MHSYR071-07|BOLD:AAA3291
Helophilus obscurus|[9540]|MHSYR070-07|BOLD:AAA3291
Helophilus obscurus|[9541]|MHSYR069-07|BOLD:AAA3291
Helophilus obscurus|[9542]|MHSYR068-07|BOLD:AAA3291
Helophilus obscurus|[9543]|MHSYR067-07|BOLD:AAA3291
Helophilus obscurus|[9544]|MHSYR065-07|BOLD:AAA3291
Helophilus obscurus|[9545]|MHSYR064-07|BOLD:AAA3291
Helophilus obscurus|[9546]|LRSYR053-07|BOLD:AAA3291
Helophilus obscurus|[9547]|MHSYR051-07|BOLD:AAA3291
Helophilus obscurus|[9548]|MHSYR173-07|BOLD:AAA3291
Helophilus obscurus|[9549]|MHSYR022-06|BOLD:AAA3291
Helophilus obscurus|[9550]|MHSYR019-06|BOLD:AAA3291
Helophilus obscurus|[9551]|MHSYR018-06|BOLD:AAA3291
Helophilus obscurus|[9552]|MHSYR017-06|BOLD:AAA3291
Helophilus obscurus|[9553]|MHSYR011-06|BOLD:AAA3291

Helophilus obscurus|[9551]|MHSYR018-06|BOLD:AAA3291
Helophilus obscurus|[9552]|MHSYR017-06|BOLD:AAA3291
Helophilus obscurus|[9553]|MHSYR011-06|BOLD:AAA3291
Helophilus obscurus|[9554]|MHSYR151-07|BOLD:AAA3291
Helophilus obscurus|[9555]|MHSYR152-07|BOLD:AAA3291
Helophilus obscurus|[9556]|LRSYR022-07|BOLD:AAA3291
Helophilus obscurus|[9557]|MHSYR047-07|BOLD:AAA3291
Helophilus obscurus|[9558]|MHSYR052-07|BOLD:AAA3291
Helophilus obscurus|[9559]|MHSYR058-07|BOLD:AAA3291
Polydontomyia curvipes|[9560]|MHSYR412-07|BOLD:AAC0442
Polydontomyia curvipes|[9561]|JWDCB019-10|BOLD:AAC0442
Polydontomyia curvipes|[9562]|LRSYR061-07|BOLD:AAC0442
Polydontomyia curvipes|[9563]|LRSYR059-07|BOLD:AAC0442
Polydontomyia curvipes|[9564]|LRSYR060-07|BOLD:AAC0442
Arctosyrphus willingi|[9565]|JSFLA010-08|BOLD:AAC0441
Arctosyrphus willingi|[9566]|MHSYR346-07|BOLD:AAC0441
Arctosyrphus willingi|[9567]|MHSYR246-07|BOLD:AAC0441
Arctosyrphus willingi|[9568]|MHSYR340-07|BOLD:AAC0441
Arctosyrphus willingi|[9569]|MHSYR331-07|BOLD:AAC0441
Arctosyrphus willingi|[9570]|MHSYR343-07|BOLD:AAC0441
Arctosyrphus willingi|[9571]|MHSYR344-07|BOLD:AAC0441
Arctosyrphus willingi|[9572]|MHSYR345-07|BOLD:AAC0441
Arctosyrphus willingi|[9573]|MHSYR347-07|BOLD:AAC0441
Arctosyrphus willingi|[9574]|MHSYR348-07|BOLD:AAC0441
Arctosyrphus willingi|[9575]|TWDIP069-09|BOLD:AAC0441
Arctosyrphus willingi|[9576]|MTDIC195-09|BOLD:AAC0441
Arctosyrphus willingi|[9577]|JWDCF132-10|BOLD:AAC0441
Arctosyrphus willingi|[9578]|JWDCC278-10|BOLD:AAC0441
Arctosyrphus willingi|[9579]|JWDCC087-10|BOLD:AAC0441
Arctosyrphus willingi|[9580]|JSFLA036-08|BOLD:AAC0441
Arctosyrphus willingi|[9581]|JWDCA478-10|BOLD:AAC0441
Arctosyrphus willingi|[9582]|JWDCA492-10|BOLD:AAC0441
Arctosyrphus willingi|[9583]|JWDCD067-10|BOLD:AAC0441
Arctosyrphus willingi|[9584]|MHSYR245-07|BOLD:AAC0441
Eurimyia stipata|[9585]|JWDCI075-10|BOLD:AAD8951
Eurimyia stipata|[9586]|JWDCF897-10|BOLD:AAD8951
Anasimyia diffusus|[9587]|JWDCI286-10|BOLD:AAD8954
Anasimyia anausis|[9588]|JWDCF899-10|BOLD:AAD8954
Anasimyia anausis|[9589]|JWDCG033-10|BOLD:AAD8954
Anasimyia anausis|[9590]|MHSYR367-07|BOLD:AAD8954
Anasimyia anausis|[9591]|JWDCF639-10|BOLD:AAD8954
Anasimyia anausis|[9592]|JWDCG860-10|BOLD:AAD8954
Anasimyia anausis|[9593]|JWDCJ299-11|BOLD:AAD8954
Anasimyia anausis|[9594]|JWDCF806-10|BOLD:AAD8954
Anasimyia anausis|[9595]|JWDCF838-10|BOLD:AAD8954
Anasimyia anausis|[9596]|JWDCI413-10|BOLD:AAD8954
Anasimyia anausis|[9597]|JWDCG740-10|BOLD:AAD8954
Anasimyia anausis|[9598]|JWDCC001-10|BOLD:AAD8954
Anasimyia anausis|[9599]|JWDCC086-10|BOLD:AAD8954
Anasimyia anausis|[9600]|JWDCJ846-11|BOLD:AAD8954
Anasimyia anausis|[9601]|JWDCG035-10|BOLD:AAD8954
Anasimyia anausis|[9602]|JWDCG799-10|BOLD:AAD8954
Anasimyia anausis|[9603]|MHFLI115-06|BOLD:AAD8954
Anasimyia anausis|[9604]|MHFLI195-06|BOLD:AAD8954
Anasimyia anausis|[9605]|JWDCI417-10|BOLD:AAD8954
Anasimyia anausis|[9606]|JWDCH036-10|BOLD:AAD8954
Anasimyia anausis|[9607]|JSFLA005-08|BOLD:AAD8954
Anasimyia bilinearis|[9608]|JWDCK237-11|BOLD:ACF0911
Anasimyia bilinearis|[9609]|JWDCC279-10|BOLD:ACF0911
Anasimyia bilinearis|[9610]|TWDIP020-09|BOLD:ACF0911
Anasimyia bilinearis|[9611]|TWDIP021-09|BOLD:ACF0911
Anasimyia bilinearis|[9612]|JWDCB021-10|BOLD:ACF0911
Anasimyia bilinearis|[9613]|JWDCH777-10|BOLD:ACF0911
Anasimyia bilinearis|[9614]|JWDCF805-10|BOLD:ACF0911
Anasimyia bilinearis|[9615]|JWDCD046-10|BOLD:ACF0911
Anasimyia bilinearis|[9616]|MHSYR354-07|BOLD:ACF0911
Anasimyia bilinearis|[9617]|JWDCF898-10|BOLD:ACF0911
Anasimyia bilinearis|[9618]|JWDCJ1405-11|BOLD:ACF0911
Anasimyia bilinearis|[9619]|JWDCE816-10|BOLD:ACF0911
Anasimyia bilinearis|[9620]|JWDCF655-10|BOLD:ACF0911
Anasimyia bilinearis|[9621]|JWDCF401-10|BOLD:ACF0911
Anasimyia bilinearis|[9622]|JWDCJ847-11|BOLD:ACF0911
Anasimyia bilinearis|[9623]|JWDCA480-10|BOLD:ACF0911
Parhelophilus obsoletus|[9624]|MHSYR266-07|BOLD:AAC0056
Parhelophilus obsoletus|[9625]|JWDCH204-10|BOLD:AAC0056
Parhelophilus obsoletus|[9626]|MHSYR271-07|BOLD:AAC0056
Parhelophilus obsoletus|[9627]|MHSYR264-07|BOLD:AAC0056
Parhelophilus obsoletus|[9628]|MHSYR270-07|BOLD:AAC0056
Parhelophilus obsoletus|[9629]|MHSYR265-07|BOLD:AAC0056
Parhelophilus obsoletus|[9630]|MHSYR269-07|BOLD:AAC0056
Parhelophilus obsoletus|[9631]|JWDCG789-10|BOLD:AAC0056
Parhelophilus obsoletus|[9632]|JWDCH202-10|BOLD:AAC0056
Parhelophilus obsoletus|[9633]|JWDCI610-10|BOLD:AAC0056
Parhelophilus obsoletus|[9634]|LRSYR011-07|BOLD:AAC0056
Parhelophilus obsoletus|[9635]|LRSYR057-07|BOLD:AAC0056
Parhelophilus obsoletus|[9636]|MHSYR268-07|BOLD:AAC0056
Parhelophilus obsoletus|[9637]|JWDCI461-10|BOLD:AAC0056
Parhelophilus obsoletus|[9638]|MHSYR272-07|BOLD:AAC0056
Parhelophilus obsoletus|[9639]|JWDCI884-11|BOLD:AAC0056
Parhelophilus obsoletus|[9640]|MHSYR267-07|BOLD:AAC0056
Parhelophilus obsoletus|[9641]|MHSYR273-07|BOLD:AAC0056
Parhelophilus porcus|[9642]|MHSYR277-07|BOLD:AAC0057
Parhelophilus porcus|[9643]|MHSYR276-07|BOLD:AAC0057
Parhelophilus porcus|[9644]|MHSYR275-07|BOLD:AAC0057
Parhelophilus porcus|[9645]|MHSYR282-07|BOLD:AAC0057
Parhelophilus porcus|[9646]|JWDCL322-11|BOLD:AAC0057
Parhelophilus porcus|[9647]|LRSYR023-07|BOLD:AAC0057
Parhelophilus porcus|[9648]|JWDCA494-10|BOLD:AAC0057
Parhelophilus porcus|[9649]|JWDCK241-11|BOLD:AAC0057
Parhelophilus porcus|[9650]|JWDCF839-10|BOLD:AAC0057
Parhelophilus porcus|[9651]|JWDCF840-10|BOLD:AAC0057
Parhelophilus porcus|[9652]|JWDCH646-10|BOLD:AAC0057
[illegible]

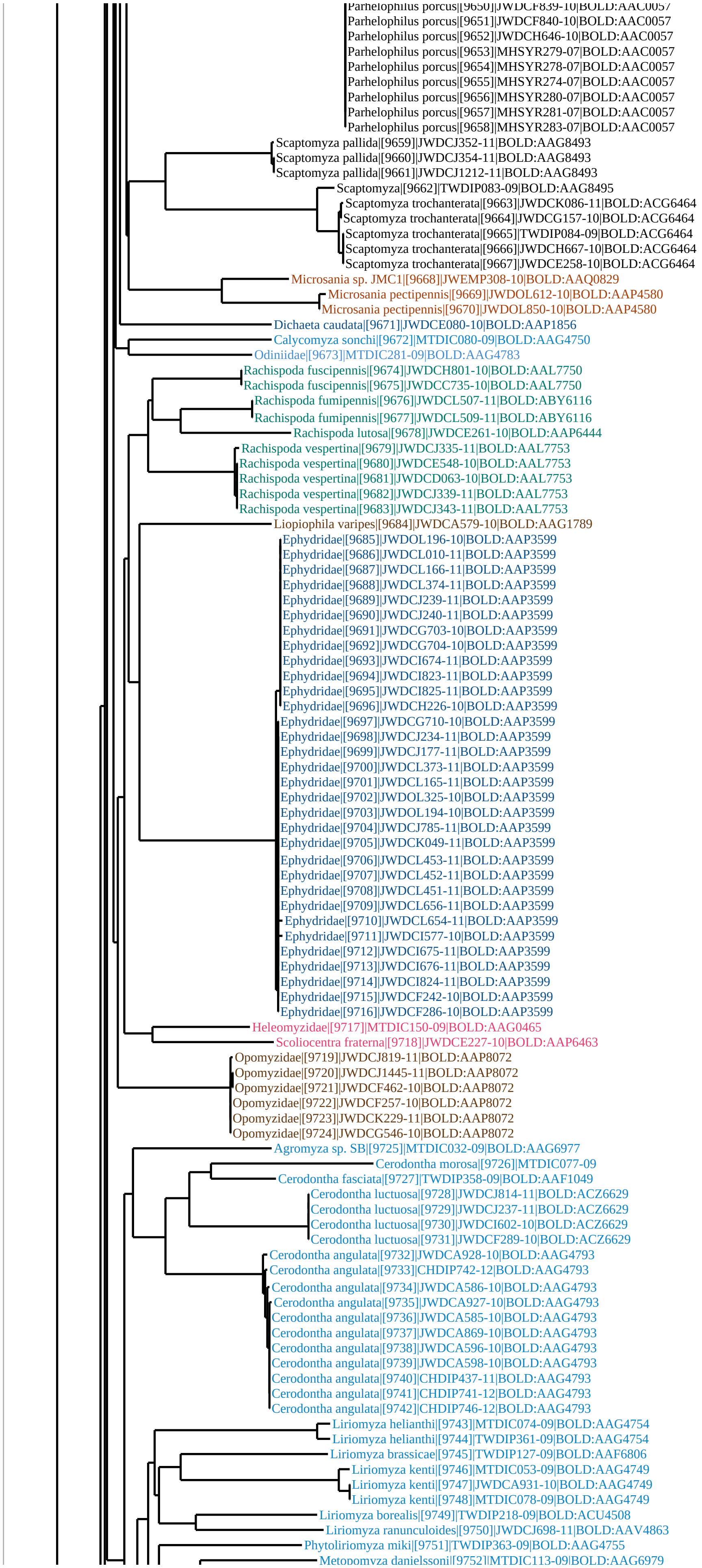
Parhelophilus porcus|[9650]|JWDCF839-10|BOLD:AAC0057
Parhelophilus porcus|[9651]|JWDCF840-10|BOLD:AAC0057
Parhelophilus porcus|[9652]|JWDCH646-10|BOLD:AAC0057
Parhelophilus porcus|[9653]|MHSYR279-07|BOLD:AAC0057
Parhelophilus porcus|[9654]|MHSYR278-07|BOLD:AAC0057
Parhelophilus porcus|[9655]|MHSYR274-07|BOLD:AAC0057
Parhelophilus porcus|[9656]|MHSYR280-07|BOLD:AAC0057
Parhelophilus porcus|[9657]|MHSYR281-07|BOLD:AAC0057
Parhelophilus porcus|[9658]|MHSYR283-07|BOLD:AAC0057
Scaptomyza pallida|[9659]|JWDCJ352-11|BOLD:AAG8493
Scaptomyza pallida|[9660]|JWDCJ354-11|BOLD:AAG8493
Scaptomyza pallida|[9661]|JWDCJ1212-11|BOLD:AAG8493
Scaptomyza|[9662]|TWDIP083-09|BOLD:AAG8495
Scaptomyza trochanterata|[9663]|JWDCK086-11|BOLD:ACG6464
Scaptomyza trochanterata|[9664]|JWDCG157-10|BOLD:ACG6464
Scaptomyza trochanterata|[9665]|TWDIP084-09|BOLD:ACG6464
Scaptomyza trochanterata|[9666]|JWDCH667-10|BOLD:ACG6464
Scaptomyza trochanterata|[9667]|JWDCE258-10|BOLD:ACG6464
Microsania sp. JMC1|[9668]|JWEMP308-10|BOLD:AAQ0829
Microsania pectipennis|[9669]|JWDOL612-10|BOLD:AAP4580
Microsania pectipennis|[9670]|JWDOL850-10|BOLD:AAP4580
Dichaeta caudata|[9671]|JWDCE080-10|BOLD:AAP1856
Calycomyza sonchi|[9672]|MTDIC080-09|BOLD:AAG4750
Odiniidae|[9673]|MTDIC281-09|BOLD:AAG4783
Rachispoda fuscipennis|[9674]|JWDCH801-10|BOLD:AAL7750
Rachispoda fuscipennis|[9675]|JWDCC735-10|BOLD:AAL7750
Rachispoda fumipennis|[9676]|JWDCL507-11|BOLD:ABY6116
Rachispoda fumipennis|[9677]|JWDCL509-11|BOLD:ABY6116
Rachispoda lutosa|[9678]|JWDCE261-10|BOLD:AAP6444
Rachispoda vespertina|[9679]|JWDCJ335-11|BOLD:AAL7753
Rachispoda vespertina|[9680]|JWDCE548-10|BOLD:AAL7753
Rachispoda vespertina|[9681]|JWDCD063-10|BOLD:AAL7753
Rachispoda vespertina|[9682]|JWDCJ339-11|BOLD:AAL7753
Rachispoda vespertina|[9683]|JWDCJ343-11|BOLD:AAL7753
Liopiophila varipes|[9684]|JWDCA579-10|BOLD:AAG1789
Ephydridae|[9685]|JWDOL196-10|BOLD:AAP3599
Ephydridae|[9686]|JWDCL010-11|BOLD:AAP3599
Ephydridae|[9687]|JWDCL166-11|BOLD:AAP3599
Ephydridae|[9688]|JWDCL374-11|BOLD:AAP3599
Ephydridae|[9689]|JWDCJ239-11|BOLD:AAP3599
Ephydridae|[9690]|JWDCJ240-11|BOLD:AAP3599
Ephydridae|[9691]|JWDCG703-10|BOLD:AAP3599
Ephydridae|[9692]|JWDCG704-10|BOLD:AAP3599
Ephydridae|[9693]|JWDCI674-11|BOLD:AAP3599
Ephydridae|[9694]|JWDCI823-11|BOLD:AAP3599
Ephydridae|[9695]|JWDCI825-11|BOLD:AAP3599
Ephydridae|[9696]|JWDCH226-10|BOLD:AAP3599
Ephydridae|[9697]|JWDCG710-10|BOLD:AAP3599
Ephydridae|[9698]|JWDCJ234-11|BOLD:AAP3599
Ephydridae|[9699]|JWDCJ177-11|BOLD:AAP3599
Ephydridae|[9700]|JWDCL373-11|BOLD:AAP3599
Ephydridae|[9701]|JWDCL165-11|BOLD:AAP3599
Ephydridae|[9702]|JWDOL325-10|BOLD:AAP3599
Ephydridae|[9703]|JWDOL194-10|BOLD:AAP3599
Ephydridae|[9704]|JWDCJ785-11|BOLD:AAP3599
Ephydridae|[9705]|JWDCK049-11|BOLD:AAP3599
Ephydridae|[9706]|JWDCL453-11|BOLD:AAP3599
Ephydridae|[9707]|JWDCL452-11|BOLD:AAP3599
Ephydridae|[9708]|JWDCL451-11|BOLD:AAP3599
Ephydridae|[9709]|JWDCL656-11|BOLD:AAP3599
Ephydridae|[9710]|JWDCL654-11|BOLD:AAP3599
Ephydridae|[9711]|JWDCI577-10|BOLD:AAP3599
Ephydridae|[9712]|JWDCI675-11|BOLD:AAP3599
Ephydridae|[9713]|JWDCI676-11|BOLD:AAP3599
Ephydridae|[9714]|JWDCI824-11|BOLD:AAP3599
Ephydridae|[9715]|JWDCF242-10|BOLD:AAP3599
Ephydridae|[9716]|JWDCF286-10|BOLD:AAP3599
Heleomyzidae|[9717]|MTDIC150-09|BOLD:AAG0465
Scoliocentra fraterna|[9718]|JWDCE227-10|BOLD:AAP6463
Opomyzidae|[9719]|JWDCJ819-11|BOLD:AAP8072
Opomyzidae|[9720]|JWDCJ1445-11|BOLD:AAP8072
Opomyzidae|[9721]|JWDCF462-10|BOLD:AAP8072
Opomyzidae|[9722]|JWDCF257-10|BOLD:AAP8072
Opomyzidae|[9723]|JWDCK229-11|BOLD:AAP8072
Opomyzidae|[9724]|JWDCG546-10|BOLD:AAP8072
Agromyza sp. SB|[9725]|MTDIC032-09|BOLD:AAG6977
Cerodontha morosa|[9726]|MTDIC077-09
Cerodontha fasciata|[9727]|TWDIP358-09|BOLD:AAF1049
Cerodontha luctuosa|[9728]|JWDCJ814-11|BOLD:ACZ6629
Cerodontha luctuosa|[9729]|JWDCJ237-11|BOLD:ACZ6629
Cerodontha luctuosa|[9730]|JWDCI602-10|BOLD:ACZ6629
Cerodontha luctuosa|[9731]|JWDCF289-10|BOLD:ACZ6629
Cerodontha angulata|[9732]|JWDCA928-10|BOLD:AAG4793
Cerodontha angulata|[9733]|CHDIP742-12|BOLD:AAG4793
Cerodontha angulata|[9734]|JWDCA586-10|BOLD:AAG4793
Cerodontha angulata|[9735]|JWDCA927-10|BOLD:AAG4793
Cerodontha angulata|[9736]|JWDCA585-10|BOLD:AAG4793
Cerodontha angulata|[9737]|JWDCA869-10|BOLD:AAG4793
Cerodontha angulata|[9738]|JWDCA596-10|BOLD:AAG4793
Cerodontha angulata|[9739]|JWDCA598-10|BOLD:AAG4793
Cerodontha angulata|[9740]|CHDIP437-11|BOLD:AAG4793
Cerodontha angulata|[9741]|CHDIP741-12|BOLD:AAG4793
Cerodontha angulata|[9742]|CHDIP746-12|BOLD:AAG4793
Liriomyza helianthi|[9743]|MTDIC074-09|BOLD:AAG4754
Liriomyza helianthi|[9744]|TWDIP361-09|BOLD:AAG4754
Liriomyza brassicae|[9745]|TWDIP127-09|BOLD:AAF6806
Liriomyza kenti|[9746]|MTDIC053-09|BOLD:AAG4749
Liriomyza kenti|[9747]|JWDCA931-10|BOLD:AAG4749
Liriomyza kenti|[9748]|MTDIC078-09|BOLD:AAG4749
Liriomyza borealis|[9749]|TWDIP218-09|BOLD:ACU4508
Liriomyza ranunculoides|[9750]|JWDCJ698-11|BOLD:AAV4863
Phytoliriomyza miki|[9751]|TWDIP363-09|BOLD:AAG4755
Metopomyza danielssoni|[9752]|MTDIC113-09|BOLD:AAG6979

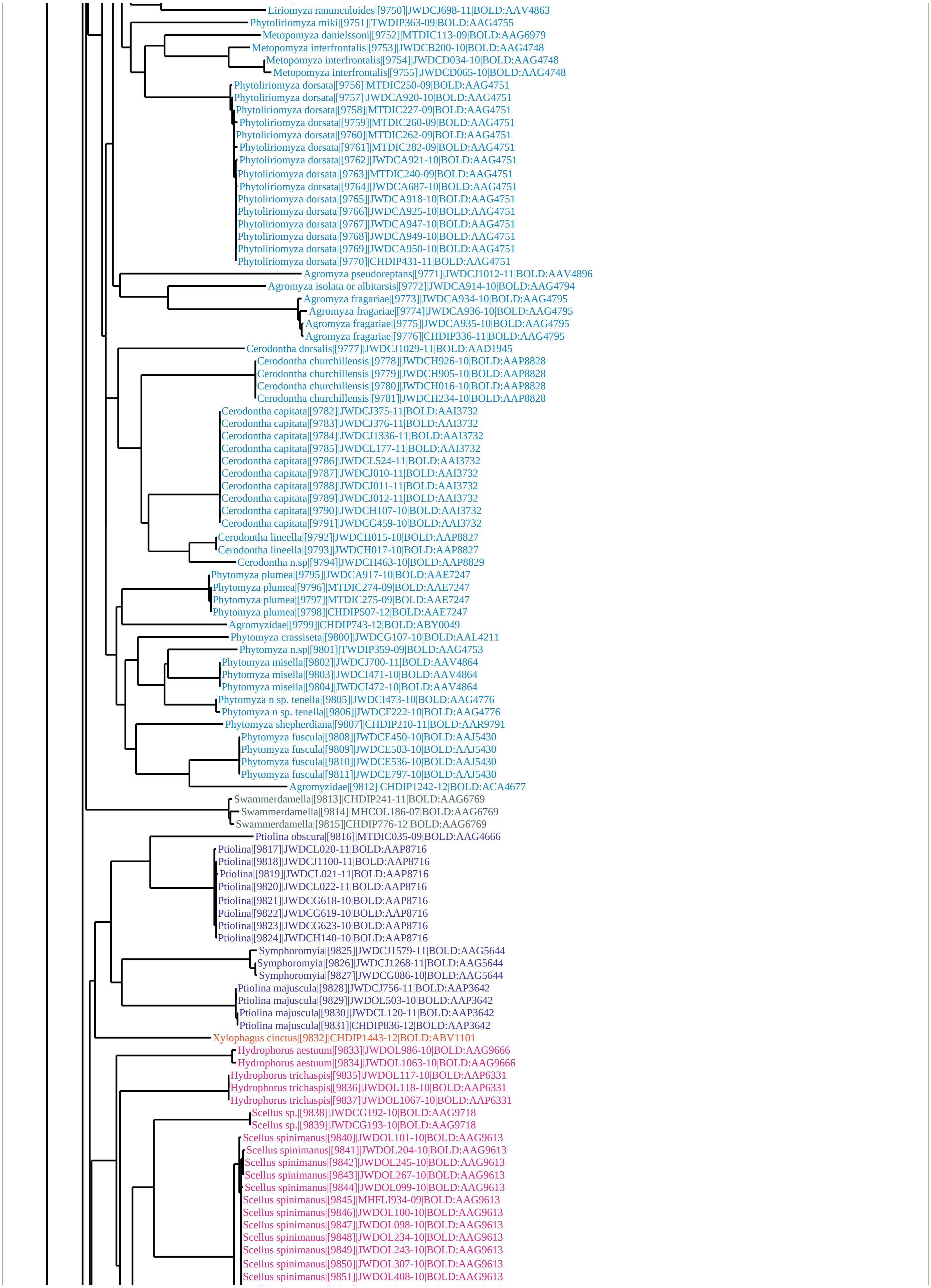
Liriomyza ranunculoides|[9750]|JWDCJ698-11|BOLD:AAV4863
Phytoliriomyza miki|[9751]|TWDIP363-09|BOLD:AAG4755
Metopomyza danielssoni|[9752]|MTDIC113-09|BOLD:AAG6979
Metopomyza interfrontalis|[9753]|JWDCB200-10|BOLD:AAG4748
Metopomyza interfrontalis|[9754]|JWDCD034-10|BOLD:AAG4748
Metopomyza interfrontalis|[9755]|JWDCD065-10|BOLD:AAG4748
Phytoliriomyza dorsata|[9756]|MTDIC250-09|BOLD:AAG4751
Phytoliriomyza dorsata|[9757]|JWDCA920-10|BOLD:AAG4751
Phytoliriomyza dorsata|[9758]|MTDIC227-09|BOLD:AAG4751
Phytoliriomyza dorsata|[9759]|MTDIC260-09|BOLD:AAG4751
Phytoliriomyza dorsata|[9760]|MTDIC262-09|BOLD:AAG4751
Phytoliriomyza dorsata|[9761]|MTDIC282-09|BOLD:AAG4751
Phytoliriomyza dorsata|[9762]|JWDCA921-10|BOLD:AAG4751
Phytoliriomyza dorsata|[9763]|MTDIC240-09|BOLD:AAG4751
Phytoliriomyza dorsata|[9764]|JWDCA687-10|BOLD:AAG4751
Phytoliriomyza dorsata|[9765]|JWDCA918-10|BOLD:AAG4751
Phytoliriomyza dorsata|[9766]|JWDCA925-10|BOLD:AAG4751
Phytoliriomyza dorsata|[9767]|JWDCA947-10|BOLD:AAG4751
Phytoliriomyza dorsata|[9768]|JWDCA949-10|BOLD:AAG4751
Phytoliriomyza dorsata|[9769]|JWDCA950-10|BOLD:AAG4751
Phytoliriomyza dorsata|[9770]|CHDIP431-11|BOLD:AAG4751
Agromyza pseudoreptans|[9771]|JWDCJ1012-11|BOLD:AAV4896
Agromyza isolata or albitarsis|[9772]|JWDCA914-10|BOLD:AAG4794
Agromyza fragariae|[9773]|JWDCA934-10|BOLD:AAG4795
Agromyza fragariae|[9774]|JWDCA936-10|BOLD:AAG4795
Agromyza fragariae|[9775]|JWDCA935-10|BOLD:AAG4795
Agromyza fragariae|[9776]|CHDIP336-11|BOLD:AAG4795
Cerodontha dorsalis|[9777]|JWDCJ1029-11|BOLD:AAD1945
Cerodontha churchillensis|[9778]|JWDCH926-10|BOLD:AAP8828
Cerodontha churchillensis|[9779]|JWDCH905-10|BOLD:AAP8828
Cerodontha churchillensis|[9780]|JWDCH016-10|BOLD:AAP8828
Cerodontha churchillensis|[9781]|JWDCH234-10|BOLD:AAP8828
Cerodontha capitata|[9782]|JWDCJ375-11|BOLD:AAI3732
Cerodontha capitata|[9783]|JWDCJ376-11|BOLD:AAI3732
Cerodontha capitata|[9784]|JWDCJ1336-11|BOLD:AAI3732
Cerodontha capitata|[9785]|JWDCL177-11|BOLD:AAI3732
Cerodontha capitata|[9786]|JWDCL524-11|BOLD:AAI3732
Cerodontha capitata|[9787]|JWDCJ010-11|BOLD:AAI3732
Cerodontha capitata|[9788]|JWDCJ011-11|BOLD:AAI3732
Cerodontha capitata|[9789]|JWDCJ012-11|BOLD:AAI3732
Cerodontha capitata|[9790]|JWDCH107-10|BOLD:AAI3732
Cerodontha capitata|[9791]|JWDCG459-10|BOLD:AAI3732
Cerodontha lineella|[9792]|JWDCH015-10|BOLD:AAP8827
Cerodontha lineella|[9793]|JWDCH017-10|BOLD:AAP8827
Cerodontha n.sp|[9794]|JWDCH463-10|BOLD:AAP8829
Phytomyza plumea|[9795]|JWDCA917-10|BOLD:AAE7247
Phytomyza plumea|[9796]|MTDIC274-09|BOLD:AAE7247
Phytomyza plumea|[9797]|MTDIC275-09|BOLD:AAE7247
Phytomyza plumea|[9798]|CHDIP507-12|BOLD:AAE7247
Agromyzidae|[9799]|CHDIP743-12|BOLD:ABY0049
Phytomyza crassiseta|[9800]|JWDCG107-10|BOLD:AAL4211
Phytomyza n.sp|[9801]|TWDIP359-09|BOLD:AAG4753
Phytomyza misella|[9802]|JWDCJ700-11|BOLD:AAV4864
Phytomyza misella|[9803]|JWDCI471-10|BOLD:AAV4864
Phytomyza misella|[9804]|JWDCI472-10|BOLD:AAV4864
Phytomyza n sp. tenella|[9805]|JWDCI473-10|BOLD:AAG4776
Phytomyza n sp. tenella|[9806]|JWDCF222-10|BOLD:AAG4776
Phytomyza shepherdiana|[9807]|CHDIP210-11|BOLD:AAR9791
Phytomyza fuscula|[9808]|JWDCE450-10|BOLD:AAJ5430
Phytomyza fuscula|[9809]|JWDCE503-10|BOLD:AAJ5430
Phytomyza fuscula|[9810]|JWDCE536-10|BOLD:AAJ5430
Phytomyza fuscula|[9811]|JWDCE797-10|BOLD:AAJ5430
Agromyzidae|[9812]|CHDIP1242-12|BOLD:ACA4677
Swammerdamella|[9813]|CHDIP241-11|BOLD:AAG6769
Swammerdamella|[9814]|MHCOL186-07|BOLD:AAG6769
Swammerdamella|[9815]|CHDIP776-12|BOLD:AAG6769
Ptiolina obscura|[9816]|MTDIC035-09|BOLD:AAG4666
Ptiolina|[9817]|JWDCL020-11|BOLD:AAP8716
Ptiolina|[9818]|JWDCJ1100-11|BOLD:AAP8716
Ptiolina|[9819]|JWDCL021-11|BOLD:AAP8716
Ptiolina|[9820]|JWDCL022-11|BOLD:AAP8716
Ptiolina|[9821]|JWDCG618-10|BOLD:AAP8716
Ptiolina|[9822]|JWDCG619-10|BOLD:AAP8716
Ptiolina|[9823]|JWDCG623-10|BOLD:AAP8716
Ptiolina|[9824]|JWDCH140-10|BOLD:AAP8716
Symphoromyia|[9825]|JWDCJ1579-11|BOLD:AAG5644
Symphoromyia|[9826]|JWDCJ1268-11|BOLD:AAG5644
Symphoromyia|[9827]|JWDCG086-10|BOLD:AAG5644
Ptiolina majuscula|[9828]|JWDCJ756-11|BOLD:AAP3642
Ptiolina majuscula|[9829]|JWDOL503-10|BOLD:AAP3642
Ptiolina majuscula|[9830]|JWDCL120-11|BOLD:AAP3642
Ptiolina majuscula|[9831]|CHDIP836-12|BOLD:AAP3642
Xylophagus cinctus|[9832]|CHDIP1443-12|BOLD:ABV1101
Hydrophorus aestuum|[9833]|JWDOL986-10|BOLD:AAG9666
Hydrophorus aestuum|[9834]|JWDOL1063-10|BOLD:AAG9666
Hydrophorus trichaspis|[9835]|JWDOL117-10|BOLD:AAP6331
Hydrophorus trichaspis|[9836]|JWDOL118-10|BOLD:AAP6331
Hydrophorus trichaspis|[9837]|JWDOL1067-10|BOLD:AAP6331
Scellus sp.|[9838]|JWDCG192-10|BOLD:AAG9718
Scellus sp.|[9839]|JWDCG193-10|BOLD:AAG9718
Scellus spinimanus|[9840]|JWDOL101-10|BOLD:AAG9613
Scellus spinimanus|[9841]|JWDOL204-10|BOLD:AAG9613
Scellus spinimanus|[9842]|JWDOL245-10|BOLD:AAG9613
Scellus spinimanus|[9843]|JWDOL267-10|BOLD:AAG9613
Scellus spinimanus|[9844]|JWDOL099-10|BOLD:AAG9613
Scellus spinimanus|[9845]|MHFLI934-09|BOLD:AAG9613
Scellus spinimanus|[9846]|JWDOL100-10|BOLD:AAG9613
Scellus spinimanus|[9847]|JWDOL098-10|BOLD:AAG9613
Scellus spinimanus|[9848]|JWDOL234-10|BOLD:AAG9613
Scellus spinimanus|[9849]|JWDOL243-10|BOLD:AAG9613
Scellus spinimanus|[9850]|JWDOL307-10|BOLD:AAG9613
Scellus spinimanus|[9851]|JWDOL408-10|BOLD:AAG9613

Scellus spinimanus|[9849]|JWDOL245-10|BOLD:AAG9613
Scellus spinimanus|[9850]|JWDOL307-10|BOLD:AAG9613
Scellus spinimanus|[9851]|JWDOL408-10|BOLD:AAG9613
Scellus spinimanus|[9852]|JWDOL985-10|BOLD:AAG9613
Scellus spinimanus|[9853]|JWDOL303-10|BOLD:AAG9613
Scellus spinimanus|[9854]|JWDOL095-10|BOLD:AAG9613
Scellus spinimanus|[9855]|MHFLI235-06|BOLD:AAG9613
Scellus spinimanus|[9856]|JWDOL246-10|BOLD:AAG9613
Scellus spinimanus|[9857]|JWDOL203-10|BOLD:AAG9613
Scellus spinimanus|[9858]|JWDOL097-10|BOLD:AAG9613
Scellus spinimanus|[9859]|JWDOL096-10|BOLD:AAG9613
Scellus spinimanus|[9860]|JWDOL012-10|BOLD:AAG9613
Scellus spinimanus|[9861]|JWDOL011-10|BOLD:AAG9613
Scellus spinimanus|[9862]|JWDOL010-10|BOLD:AAG9613
Scellus spinimanus|[9863]|JWDOL009-10|BOLD:AAG9613
Scellus spinimanus|[9864]|JWDOL008-10|BOLD:AAG9613
Scellus spinimanus|[9865]|MHFLI236-06|BOLD:AAG9613
Scellus spinimanus|[9866]|TWDIP147-09|BOLD:AAG9613
Scellus spinimanus|[9867]|JWDOL094-10|BOLD:AAG9613
Scellus spinimanus|[9868]|JWDCI226-10|BOLD:AAG9613
Scellus spinimanus|[9869]|JWDOL264-10|BOLD:AAG9613
Scellus spinimanus|[9870]|JWDOL265-10|BOLD:AAG9613
Scellus spinimanus|[9871]|JWDOL266-10|BOLD:AAG9613
Scellus spinimanus|[9872]|JWDOL326-10|BOLD:AAG9613
Scellus spinimanus|[9873]|JWDOL380-10|BOLD:AAG9613
Scellus spinimanus|[9874]|JWDOL381-10|BOLD:AAG9613
Scellus spinimanus|[9875]|JWDOL387-10|BOLD:AAG9613
Scellus spinimanus|[9876]|JWDOL399-10|BOLD:AAG9613
Scellus spinimanus|[9877]|JWDOL402-10|BOLD:AAG9613
Scellus spinimanus|[9878]|JWDOL416-10|BOLD:AAG9613
Scellus spinimanus|[9879]|JWDOL1038-10|BOLD:AAG9613
Scellus spinimanus|[9880]|JWDOL874-10|BOLD:AAG9613
Scellus spinimanus|[9881]|JWDOL904-10|BOLD:AAG9613
Hydrophorus brevicauda|[9882]|JWDOL109-10|BOLD:AAP6328
Hydrophorus flavipennis|[9883]|MHFLY125-07|BOLD:AAG9615
Hydrophorus flavipennis|[9884]|MHFLY126-07|BOLD:AAG9615
Hydrophorus chrysologus|[9885]|JWDCG517-10|BOLD:AAP6330
Hydrophorus chrysologus|[9886]|JWDOL113-10|BOLD:AAP6330
Hydrophorus manicatus|[9887]|JWDOL110-10|BOLD:AAP6329
Hydrophorus manicatus|[9888]|JWDOL111-10|BOLD:AAP6329
Hydrophorus manicatus|[9889]|JWDOL112-10|BOLD:AAP6329
Hydrophorus flavihirtus|[9890]|JWDOL1060-10|BOLD:AAQ0835
Hydrophorus flavihirtus|[9891]|JWDOL1018-10|BOLD:AAQ0835
Hydrophorus flavihirtus|[9892]|JWDOL1064-10|BOLD:AAQ0835
Hydrophorus glaber|[9893]|JWDOL913-10|BOLD:ACE7656
Hydrophorus hirpicifer|[9894]|JWDOL973-10|BOLD:AAP5024
Hydrophorus glaber|[9895]|JWDOL827-10|BOLD:AAG7066
Hydrophorus glaber|[9896]|JWDOL914-10|BOLD:AAG7066
Hydrophorus glaber|[9897]|MHFLI281-06|BOLD:AAG7066
Hydrophorus glaber|[9898]|JWDCE355-10|BOLD:AAG7066
Hydrophorus glaber|[9899]|JWDCE001-10|BOLD:AAG7066
Hydrophorus glaber|[9900]|JWDOL972-10|BOLD:AAG7066
Hydrophorus glaber|[9901]|JWDOL1065-10|BOLD:AAG7066
Hydrophorus glaber|[9902]|JWDOL472-10|BOLD:AAG7066
Hydrophorus glaber|[9903]|JWDOL915-10|BOLD:AAG7066
Campsicnemus|[9904]|JWDOL051-10|BOLD:AAP1861
Campsicnemus americanus|[9905]|JWDOL151-10|BOLD:AAP1861
Campsicnemus|[9906]|JWDOL180-10|BOLD:AAP1861
Campsicnemus sp. SEB1|[9907]|JWDOL1062-10|BOLD:AAG9637
Campsicnemus sp. SEB1|[9908]|JWDOL1059-10|BOLD:AAG9637
Campsicnemus sp. SEB1|[9909]|JWEMP266-10|BOLD:AAG9637
Campsicnemus sp. SEB1|[9910]|JWDOL966-10|BOLD:AAG9637
Campsicnemus sp. SEB1|[9911]|JWDOL1020-10|BOLD:AAG9637
Campsicnemus sp. SEB1|[9912]|JWDOL912-10|BOLD:AAG9637
Campsicnemus sp. SEB1|[9913]|JWDOL957-10|BOLD:AAG9637
Campsicnemus vanduzeei|[9914]|JWDOL1035-10|BOLD:ABY9817
Campsicnemus vanduzeei|[9915]|JWDOL1043-10|BOLD:ABY9817
Campsicnemus vanduzeei|[9916]|JWDOL1031-10|BOLD:ABY9817
Campsicnemus vanduzeei|[9917]|JWDOL1045-10|BOLD:ABY9817
Campsicnemus vanduzeei|[9918]|JWDOL1027-10|BOLD:ABY9817
Campsicnemus vanduzeei|[9919]|JWDOL828-10|BOLD:ABY9817
Campsicnemus vanduzeei|[9920]|JWEMP265-10|BOLD:ABY9817
Campsicnemus vanduzeei|[9921]|JWDOL1070-10|BOLD:ABY9817
Campsicnemus vanduzeei|[9922]|JWDOL1056-10|BOLD:ABY9817
Campsicnemus vanduzeei|[9923]|JWDOL1055-10|BOLD:ABY9817
Campsicnemus vanduzeei|[9924]|JWDOL1054-10|BOLD:ABY9817
Campsicnemus vanduzeei|[9925]|JWDOL1053-10|BOLD:ABY9817
Campsicnemus vanduzeei|[9926]|JWDOL1052-10|BOLD:ABY9817
Campsicnemus vanduzeei|[9927]|JWDOL1051-10|BOLD:ABY9817
Campsicnemus vanduzeei|[9928]|JWDOL1047-10|BOLD:ABY9817
Campsicnemus vanduzeei|[9929]|JWDOL1046-10|BOLD:ABY9817
Campsicnemus vanduzeei|[9930]|JWDOL1044-10|BOLD:ABY9817
Campsicnemus vanduzeei|[9931]|JWDOL1042-10|BOLD:ABY9817
Campsicnemus vanduzeei|[9932]|JWDOL1036-10|BOLD:ABY9817
Campsicnemus vanduzeei|[9933]|JWDOL1033-10|BOLD:ABY9817
Campsicnemus vanduzeei|[9934]|JWDOL1032-10|BOLD:ABY9817
Campsicnemus vanduzeei|[9935]|JWDOL1030-10|BOLD:ABY9817
Campsicnemus vanduzeei|[9936]|JWDOL1028-10|BOLD:ABY9817
Campsicnemus vanduzeei|[9937]|JWDOL1026-10|BOLD:ABY9817
Campsicnemus vanduzeei|[9938]|JWDOL1025-10|BOLD:ABY9817
Campsicnemus vanduzeei|[9939]|JWDOL1024-10|BOLD:ABY9817
Campsicnemus vanduzeei|[9940]|JWDOL987-10|BOLD:ABY9817
Campsicnemus vanduzeei|[9941]|JWDOL952-10|BOLD:ABY9817
Campsicnemus vanduzeei|[9942]|JWDOL938-10|BOLD:ABY9817
Campsicnemus vanduzeei|[9943]|JWDOL928-10|BOLD:ABY9817
Campsicnemus vanduzeei|[9944]|JWDOL927-10|BOLD:ABY9817
Campsicnemus vanduzeei|[9945]|JWDOL926-10|BOLD:ABY9817
Campsicnemus vanduzeei|[9946]|JWDOL925-10|BOLD:ABY9817
Campsicnemus vanduzeei|[9947]|JWDOL924-10|BOLD:ABY9817
Campsicnemus vanduzeei|[9948]|JWDOL457-10|BOLD:ABY9817
Campsicnemus vanduzeei|[9949]|JWDOL451-10|BOLD:ABY9817
Campsicnemus vanduzeei|[9950]|JWDOL439-10|BOLD:ABY9817
Campsicnemus vanduzeei|[9951]|JWDOL124-10|BOLD:ABY9817

Campsicnemus vanduzeei|[9949]|JWDOL451-10|BOLD:ABY9817
Campsicnemus vanduzeei|[9950]|JWDOL439-10|BOLD:ABY9817
Campsicnemus vanduzeei|[9951]|JWDOL124-10|BOLD:ABY9817
Campsicnemus vanduzeei|[9952]|JWDOL123-10|BOLD:ABY9817
Campsicnemus vanduzeei|[9953]|JWDOL122-10|BOLD:ABY9817
Campsicnemus vanduzeei|[9954]|JWDOL119-10|BOLD:ABY9817
Campsicnemus vanduzeei|[9955]|JWDCE196-10|BOLD:ABY9817
Campsicnemus vanduzeei|[9956]|JWDCE146-10|BOLD:ABY9817
Campsicnemus vanduzeei|[9957]|JWDCE120-10|BOLD:ABY9817
Campsicnemus vanduzeei|[9958]|JWDCE109-10|BOLD:ABY9817
Campsicnemus vanduzeei|[9959]|JWDCE108-10|BOLD:ABY9817
Campsicnemus vanduzeei|[9960]|JWDCE804-10|BOLD:ABY9817
Campsicnemus vanduzeei|[9961]|JWDCE405-10|BOLD:ABY9817
Campsicnemus vanduzeei|[9962]|JWDCE366-10|BOLD:ABY9817
Campsicnemus vanduzeei|[9963]|JWDCE336-10|BOLD:ABY9817
Campsicnemus vanduzeei|[9964]|JWDCE285-10|BOLD:ABY9817
Campsicnemus vanduzeei|[9965]|JWDCE284-10|BOLD:ABY9817
Campsicnemus vanduzeei|[9966]|JWDCE283-10|BOLD:ABY9817
Campsicnemus vanduzeei|[9967]|JWDCG520-10|BOLD:ABY9817
Campsicnemus vanduzeei|[9968]|JWDOL115-10|BOLD:ABY9817
Campsicnemus vanduzeei|[9969]|JWDCG450-10|BOLD:ABY9817
Campsicnemus vanduzeei|[9970]|JWDOL905-10|BOLD:ABY9817
Campsicnemus vanduzeei|[9971]|JWDOL906-10|BOLD:ABY9817
Campsicnemus vanduzeei|[9972]|JWDOL956-10|BOLD:ABY9817
Campsicnemus alexanderi|[9973]|JWDCE282-10|BOLD:AAG9638
Campsicnemus alexanderi|[9974]|JWDCE335-10|BOLD:AAG9638
Campsicnemus alexanderi|[9975]|JWDOL108-10|BOLD:AAG9638
Campsicnemus alexanderi|[9976]|JWDOL936-10|BOLD:AAG9638
Campsicnemus alexanderi|[9977]|JWDOL967-10|BOLD:AAG9638
Campsicnemus alexanderi|[9978]|JWDOL969-10|BOLD:AAG9638
Campsicnemus alexanderi|[9979]|JWDOL1041-10|BOLD:AAG9638
Campsicnemus alexanderi|[9980]|JWDOL107-10|BOLD:AAG9638
Campsicnemus alexanderi|[9981]|JWDOL1069-10|BOLD:AAG9638
Campsicnemus alexanderi|[9982]|JWDCE342-10|BOLD:AAG9638
Campsicnemus alexanderi|[9983]|JWDCE107-10|BOLD:AAG9638
Campsicnemus alexanderi|[9984]|JWDCE275-10|BOLD:AAG9638
Campsicnemus alexanderi|[9985]|JWDOL106-10|BOLD:AAG9638
Campsicnemus alexanderi|[9986]|JWDOL114-10|BOLD:AAG9638
Campsicnemus alexanderi|[9987]|JWDOL116-10|BOLD:AAG9638
Campsicnemus alexanderi|[9988]|JWDOL120-10|BOLD:AAG9638
Campsicnemus alexanderi|[9989]|JWDOL125-10|BOLD:AAG9638
Campsicnemus alexanderi|[9990]|JWDOL937-10|BOLD:AAG9638
Campsicnemus alexanderi|[9991]|JWDOL968-10|BOLD:AAG9638
Campsicnemus alexanderi|[9992]|JWDOL989-10|BOLD:AAG9638
Campsicnemus alexanderi|[9993]|JWDOL990-10|BOLD:AAG9638
Campsicnemus alexanderi|[9994]|JWDOL1068-10|BOLD:AAG9638
Campsicnemus alexanderi|[9995]|JWDOL1071-10|BOLD:AAG9638
Campsicnemus alexanderi|[9996]|JWEMP283-10|BOLD:AAG9638
Campsicnemus alexanderi|[9997]|JWDOL965-10|BOLD:AAG9638
Sympycnus sp. SEB2|[9998]|MTDIC180-09|BOLD:AAG9683
Sympycnus sp. SEB1|[9999]|JWDOL450-10|BOLD:ACW3856
Sympycnus sp. SEB1|[10000]|JWDOL1022-10|BOLD:ACW3856
Sympycnus sp. SEB1|[10001]|JWDOL866-10|BOLD:ACW3856
Sympycnus|[10002]|JWDOL872-10|BOLD:ACW3856
Sympycnus sp. SEB1|[10003]|JWDOL991-10|BOLD:ACW3856
Sympycnus sp. SEB1|[10004]|JWDCH129-10|BOLD:ACW3856
Sympycnus sp. SEB1|[10005]|JWDOL869-10|BOLD:ACW3856
Sympycnus sp. SEB1|[10006]|JWDOL954-10|BOLD:ACW3856
Sympycnus sp. SEB1|[10007]|JWDOL955-10|BOLD:ACW3856
Sympycnus sp. SEB1|[10008]|JWDOL865-10|BOLD:ACW3856
Sympycnus sp. SEB1|[10009]|JWDOL458-10|BOLD:ACW3856
Sympycnus sp. SEB1|[10010]|JWDOL1023-10|BOLD:ACW3856
Sympycnus sp. SEB1|[10011]|JWDOL454-10|BOLD:ACW3856
Sympycnus sp. SEB1|[10012]|JWDOL453-10|BOLD:ACW3856
Sympycnus sp. SEB1|[10013]|JWDOL452-10|BOLD:ACW3856
Sympycnus sp. SEB1|[10014]|JWDCI144-10|BOLD:ACW3856
Sympycnus sp. SEB1|[10015]|JWDOL444-10|BOLD:ACW3856
Sympycnus sp. SEB1|[10016]|JWDOL870-10|BOLD:ACW3856
Sympycnus sp. SEB1|[10017]|JWDOL871-10|BOLD:ACW3856
Sympycnus sp. SEB1|[10018]|JWDOL873-10|BOLD:ACW3856
Sympycnus sp. SEB1|[10019]|JWDOL953-10|BOLD:ACW3856
Sympycnus sp. SEB1|[10020]|JWDOL958-10|BOLD:ACW3856
Argyra sp. nr. nigricoxa|[10021]|JWDCF901-10|BOLD:AAP1860
Argyra sp. nr. nigricoxa|[10022]|JWDOL035-10|BOLD:AAP1860
Argyra sp. nr. nigricoxa|[10023]|JWDOL038-10|BOLD:AAP1860
Argyra sp. nr. nigricoxa|[10024]|JWDOL044-10|BOLD:AAP1860
Argyra sp. nr. nigricoxa|[10025]|JWDOL423-10|BOLD:AAP1860
Argyra sp. nr. nigricoxa|[10026]|JWEMP002-10|BOLD:AAP1860
Allanthalia sp.|[10027]|JWDCC307-10|BOLD:AAL8961
Swammerdamella|[10028]|JWDCJ1022-11|BOLD:AAV1137
Coboldia fuscipes|[10029]|JWDCJ1094-11|BOLD:AAN8523
Coboldia fuscipes|[10030]|JWDCJ1095-11|BOLD:AAN8523
Scatopse|[10031]|TWDIP073-09|BOLD:AAH4119
Scatopse|[10032]|TWDIP074-09|BOLD:AAH4119
Scatopse|[10033]|MTDIC019-09|BOLD:AAH4119
Scatopse|[10034]|MTDIC020-09|BOLD:AAH4119
Scatopse|[10035]|MTDIC120-09|BOLD:AAH4119
Scatopse|[10036]|MTDIC129-09|BOLD:AAH4119
Scatopse|[10037]|MTDIC130-09|BOLD:AAH4119
Scatopse|[10038]|JWDCB350-10|BOLD:AAH4119
Scatopse|[10039]|JWDCB351-10|BOLD:AAH4119
Scatopse|[10040]|JWDCB352-10|BOLD:AAH4119
Scatopse|[10041]|JWDCB354-10|BOLD:AAH4119
Scatopse|[10042]|JWDCB360-10|BOLD:AAH4119
Scatopse|[10043]|JWDCB353-10|BOLD:AAH4119
Scatopse|[10044]|JWDCJ1021-11|BOLD:AAH4119
Scatopse|[10045]|JWDCJ1023-11|BOLD:AAH4119
Scatopse|[10046]|JWDCG339-10|BOLD:AAH4119
Leptopeza flavipes|[10047]|JWEMP239-10|BOLD:AAF9766
Leptopeza flavipes|[10048]|JWEMP244-10|BOLD:AAF9766
Leptopeza flavipes|[10049]|JWDOL712-10|BOLD:AAF9766
Leptopeza flavipes|[10050]|JWDOL790-10|BOLD:AAF9766

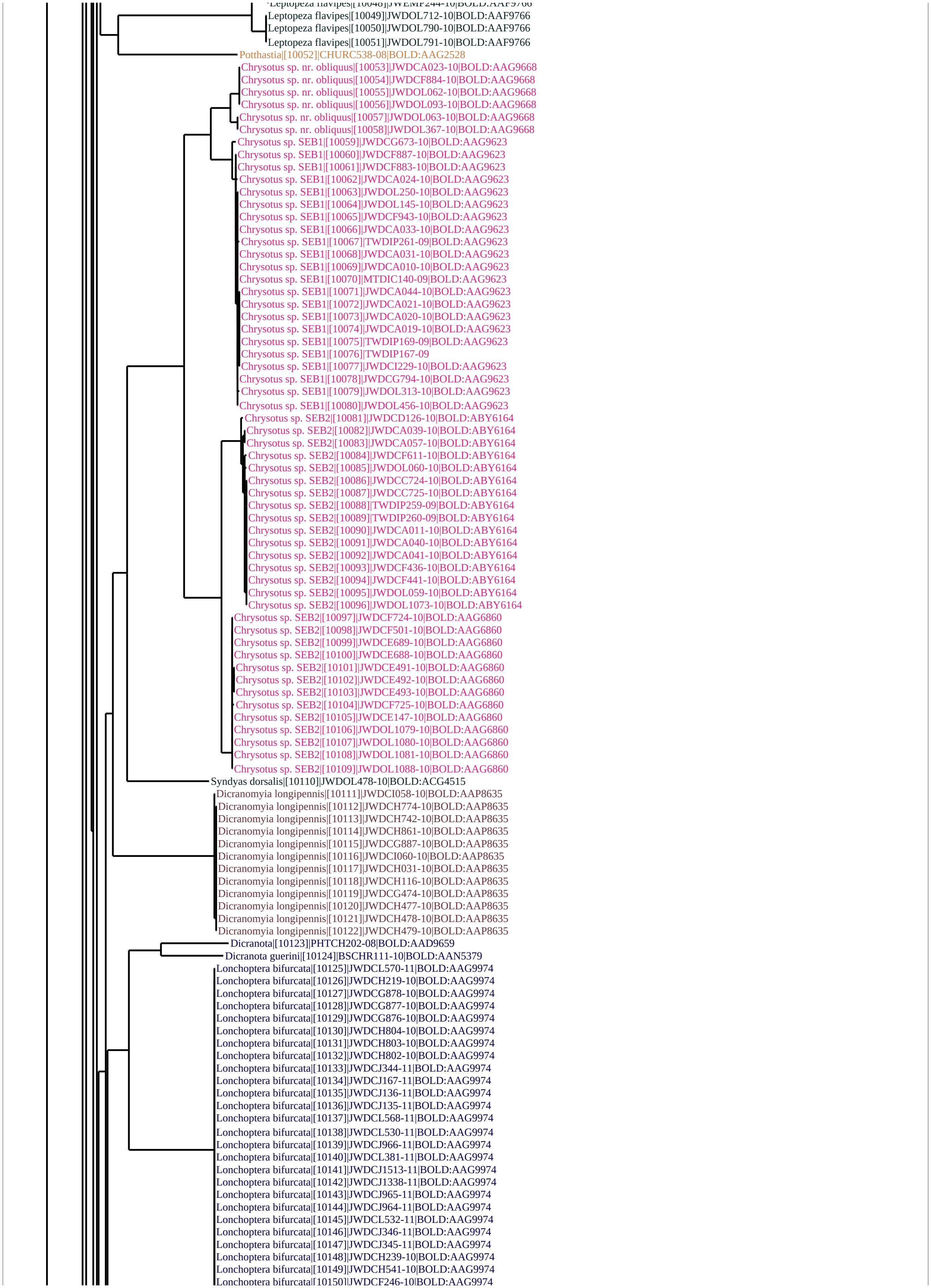
Leptopeza flavipes|[10048]|JWEMP244-10|BOLD:AAF9766
Leptopeza flavipes|[10049]|JWDOL712-10|BOLD:AAF9766
Leptopeza flavipes|[10050]|JWDOL790-10|BOLD:AAF9766
Leptopeza flavipes|[10051]|JWDOL791-10|BOLD:AAF9766
Potthastia|[10052]|CHURC538-08|BOLD:AAG2528
Chrysotus sp. nr. obliquus|[10053]|JWDCA023-10|BOLD:AAG9668
Chrysotus sp. nr. obliquus|[10054]|JWDCF884-10|BOLD:AAG9668
Chrysotus sp. nr. obliquus|[10055]|JWDOL062-10|BOLD:AAG9668
Chrysotus sp. nr. obliquus|[10056]|JWDOL093-10|BOLD:AAG9668
Chrysotus sp. nr. obliquus|[10057]|JWDOL063-10|BOLD:AAG9668
Chrysotus sp. nr. obliquus|[10058]|JWDOL367-10|BOLD:AAG9668
Chrysotus sp. SEB1|[10059]|JWDCG673-10|BOLD:AAG9623
Chrysotus sp. SEB1|[10060]|JWDCF887-10|BOLD:AAG9623
Chrysotus sp. SEB1|[10061]|JWDCF883-10|BOLD:AAG9623
Chrysotus sp. SEB1|[10062]|JWDCA024-10|BOLD:AAG9623
Chrysotus sp. SEB1|[10063]|JWDOL250-10|BOLD:AAG9623
Chrysotus sp. SEB1|[10064]|JWDOL145-10|BOLD:AAG9623
Chrysotus sp. SEB1|[10065]|JWDCF943-10|BOLD:AAG9623
Chrysotus sp. SEB1|[10066]|JWDCA033-10|BOLD:AAG9623
Chrysotus sp. SEB1|[10067]|TWDIP261-09|BOLD:AAG9623
Chrysotus sp. SEB1|[10068]|JWDCA031-10|BOLD:AAG9623
Chrysotus sp. SEB1|[10069]|JWDCA010-10|BOLD:AAG9623
Chrysotus sp. SEB1|[10070]|MTDIC140-09|BOLD:AAG9623
Chrysotus sp. SEB1|[10071]|JWDCA044-10|BOLD:AAG9623
Chrysotus sp. SEB1|[10072]|JWDCA021-10|BOLD:AAG9623
Chrysotus sp. SEB1|[10073]|JWDCA020-10|BOLD:AAG9623
Chrysotus sp. SEB1|[10074]|JWDCA019-10|BOLD:AAG9623
Chrysotus sp. SEB1|[10075]|TWDIP169-09|BOLD:AAG9623
Chrysotus sp. SEB1|[10076]|TWDIP167-09
Chrysotus sp. SEB1|[10077]|JWDCI229-10|BOLD:AAG9623
Chrysotus sp. SEB1|[10078]|JWDCG794-10|BOLD:AAG9623
Chrysotus sp. SEB1|[10079]|JWDOL313-10|BOLD:AAG9623
Chrysotus sp. SEB1|[10080]|JWDOL456-10|BOLD:AAG9623
Chrysotus sp. SEB2|[10081]|JWDCD126-10|BOLD:ABY6164
Chrysotus sp. SEB2|[10082]|JWDCA039-10|BOLD:ABY6164
Chrysotus sp. SEB2|[10083]|JWDCA057-10|BOLD:ABY6164
Chrysotus sp. SEB2|[10084]|JWDCF611-10|BOLD:ABY6164
Chrysotus sp. SEB2|[10085]|JWDOL060-10|BOLD:ABY6164
Chrysotus sp. SEB2|[10086]|JWDCC724-10|BOLD:ABY6164
Chrysotus sp. SEB2|[10087]|JWDCC725-10|BOLD:ABY6164
Chrysotus sp. SEB2|[10088]|TWDIP259-09|BOLD:ABY6164
Chrysotus sp. SEB2|[10089]|TWDIP260-09|BOLD:ABY6164
Chrysotus sp. SEB2|[10090]|JWDCA011-10|BOLD:ABY6164
Chrysotus sp. SEB2|[10091]|JWDCA040-10|BOLD:ABY6164
Chrysotus sp. SEB2|[10092]|JWDCA041-10|BOLD:ABY6164
Chrysotus sp. SEB2|[10093]|JWDCF436-10|BOLD:ABY6164
Chrysotus sp. SEB2|[10094]|JWDCF441-10|BOLD:ABY6164
Chrysotus sp. SEB2|[10095]|JWDOL059-10|BOLD:ABY6164
Chrysotus sp. SEB2|[10096]|JWDOL1073-10|BOLD:ABY6164
Chrysotus sp. SEB2|[10097]|JWDCF724-10|BOLD:AAG6860
Chrysotus sp. SEB2|[10098]|JWDCF501-10|BOLD:AAG6860
Chrysotus sp. SEB2|[10099]|JWDCE689-10|BOLD:AAG6860
Chrysotus sp. SEB2|[10100]|JWDCE688-10|BOLD:AAG6860
Chrysotus sp. SEB2|[10101]|JWDCE491-10|BOLD:AAG6860
Chrysotus sp. SEB2|[10102]|JWDCE492-10|BOLD:AAG6860
Chrysotus sp. SEB2|[10103]|JWDCE493-10|BOLD:AAG6860
Chrysotus sp. SEB2|[10104]|JWDCF725-10|BOLD:AAG6860
Chrysotus sp. SEB2|[10105]|JWDCE147-10|BOLD:AAG6860
Chrysotus sp. SEB2|[10106]|JWDOL1079-10|BOLD:AAG6860
Chrysotus sp. SEB2|[10107]|JWDOL1080-10|BOLD:AAG6860
Chrysotus sp. SEB2|[10108]|JWDOL1081-10|BOLD:AAG6860
Chrysotus sp. SEB2|[10109]|JWDOL1088-10|BOLD:AAG6860
Syndyas dorsalis|[10110]|JWDOL478-10|BOLD:ACG4515
Dicranomyia longipennis|[10111]|JWDCI058-10|BOLD:AAP8635
Dicranomyia longipennis|[10112]|JWDCH774-10|BOLD:AAP8635
Dicranomyia longipennis|[10113]|JWDCH742-10|BOLD:AAP8635
Dicranomyia longipennis|[10114]|JWDCH861-10|BOLD:AAP8635
Dicranomyia longipennis|[10115]|JWDCG887-10|BOLD:AAP8635
Dicranomyia longipennis|[10116]|JWDCI060-10|BOLD:AAP8635
Dicranomyia longipennis|[10117]|JWDCH031-10|BOLD:AAP8635
Dicranomyia longipennis|[10118]|JWDCH116-10|BOLD:AAP8635
Dicranomyia longipennis|[10119]|JWDCG474-10|BOLD:AAP8635
Dicranomyia longipennis|[10120]|JWDCH477-10|BOLD:AAP8635
Dicranomyia longipennis|[10121]|JWDCH478-10|BOLD:AAP8635
Dicranomyia longipennis|[10122]|JWDCH479-10|BOLD:AAP8635
Dicranota|[10123]|PHTCH202-08|BOLD:AAD9659
Dicranota guerini|[10124]|BSCHR111-10|BOLD:AAN5379
Lonchoptera bifurcata|[10125]|JWDCL570-11|BOLD:AAG9974
Lonchoptera bifurcata|[10126]|JWDCH219-10|BOLD:AAG9974
Lonchoptera bifurcata|[10127]|JWDCG878-10|BOLD:AAG9974
Lonchoptera bifurcata|[10128]|JWDCG877-10|BOLD:AAG9974
Lonchoptera bifurcata|[10129]|JWDCG876-10|BOLD:AAG9974
Lonchoptera bifurcata|[10130]|JWDCH804-10|BOLD:AAG9974
Lonchoptera bifurcata|[10131]|JWDCH803-10|BOLD:AAG9974
Lonchoptera bifurcata|[10132]|JWDCH802-10|BOLD:AAG9974
Lonchoptera bifurcata|[10133]|JWDCJ344-11|BOLD:AAG9974
Lonchoptera bifurcata|[10134]|JWDCJ167-11|BOLD:AAG9974
Lonchoptera bifurcata|[10135]|JWDCJ136-11|BOLD:AAG9974
Lonchoptera bifurcata|[10136]|JWDCJ135-11|BOLD:AAG9974
Lonchoptera bifurcata|[10137]|JWDCL568-11|BOLD:AAG9974
Lonchoptera bifurcata|[10138]|JWDCL530-11|BOLD:AAG9974
Lonchoptera bifurcata|[10139]|JWDCJ966-11|BOLD:AAG9974
Lonchoptera bifurcata|[10140]|JWDCL381-11|BOLD:AAG9974
Lonchoptera bifurcata|[10141]|JWDCJ1513-11|BOLD:AAG9974
Lonchoptera bifurcata|[10142]|JWDCJ1338-11|BOLD:AAG9974
Lonchoptera bifurcata|[10143]|JWDCJ965-11|BOLD:AAG9974
Lonchoptera bifurcata|[10144]|JWDCJ964-11|BOLD:AAG9974
Lonchoptera bifurcata|[10145]|JWDCL532-11|BOLD:AAG9974
Lonchoptera bifurcata|[10146]|JWDCJ346-11|BOLD:AAG9974
Lonchoptera bifurcata|[10147]|JWDCJ345-11|BOLD:AAG9974
Lonchoptera bifurcata|[10148]|JWDCH239-10|BOLD:AAG9974
Lonchoptera bifurcata|[10149]|JWDCH541-10|BOLD:AAG9974
Lonchoptera bifurcata|[10150]|JWDCF246-10|BOLD:AAG9974

Lonchoptera bifurcata|[10148]|JWDCH239-10|BOLD:AAG9974
Lonchoptera bifurcata|[10149]|JWDCH541-10|BOLD:AAG9974
Lonchoptera bifurcata|[10150]|JWDCF246-10|BOLD:AAG9974
Lonchoptera bifurcata|[10151]|JWDCF247-10|BOLD:AAG9974
Lonchoptera bifurcata|[10152]|JWDCF256-10|BOLD:AAG9974
Lonchoptera bifurcata|[10153]|JWDCF278-10|BOLD:AAG9974
Lonchoptera bifurcata|[10154]|JWDCF248-10|BOLD:AAG9974
Lonchoptera bifurcata|[10155]|JWDCF249-10|BOLD:AAG9974
Chrysopilus flavibarbis|[10156]|JWDCJ1031-11|BOLD:AAG5645
Chrysopilus flavibarbis|[10157]|JWDCG155-10|BOLD:AAG5645
Chrysopilus flavibarbis|[10158]|JWDCJ1032-11|BOLD:AAG5645
Chrysopilus flavibarbis|[10159]|JWDCJ1033-11|BOLD:AAG5645
Chrysopilus flavibarbis|[10160]|JWDCG153-10|BOLD:AAG5645
Chrysopilus flavibarbis|[10161]|JWDCG154-10|BOLD:AAG5645
Chrysopilus flavibarbis|[10162]|JWDCG156-10|BOLD:AAG5645
Chrysopilus flavibarbis|[10163]|JWDCF398-10|BOLD:AAG5645
Thripomorpha|[10164]|JWDCI928-11|BOLD:AAP8768
Thripomorpha|[10165]|JWDCF326-10|BOLD:AAP8768
Thripomorpha|[10166]|JWDCF777-10|BOLD:ACY5612
Thripomorpha|[10167]|JWDCF293-10|BOLD:ACY5612
Thripomorpha|[10168]|JWDCF327-10|BOLD:ACY5611
Thripomorpha|[10169]|JWDCI929-11|BOLD:ACY5611
Thripomorpha|[10170]|JWDCF328-10|BOLD:ACY5611
Pediciidae|[10171]|BSCHR157-10|BOLD:AAN5372
Pediciidae|[10172]|BSCHR056-10|BOLD:AAN5372
Pediciidae|[10173]|BSCHR093-10|BOLD:AAN5372
Pediciidae|[10174]|BSCHR115-10|BOLD:AAN5372
Odontomyia|[10175]|TWDIP226-09|BOLD:AAI3613
Odontomyia nigerrima|[10176]|JWDCD059-10|BOLD:AAI3613
Odontomyia|[10177]|JSFLA029-08|BOLD:AAI3613
Stratiomys laticeps|[10178]|HMCOL064-07|BOLD:AAG9066
Stratiomys laticeps|[10179]|MTDIC001-09|BOLD:AAG9066
Stratiomys laticeps|[10180]|MTDIC002-09|BOLD:AAG9066
Stratiomys laticeps|[10181]|JSFLA014-08|BOLD:AAG9066
Stratiomys laticeps|[10182]|JWDCL493-11|BOLD:AAG9066
Stratiomys laticeps|[10183]|JWDCL494-11|BOLD:AAG9066
Stratiomys laticeps|[10184]|JWDCL495-11|BOLD:AAG9066
Stratiomys laticeps|[10185]|JWDCG856-10|BOLD:AAG9066
Stratiomys laticeps|[10186]|JWDCI231-10|BOLD:AAG9066
Stratiomys laticeps|[10187]|JWDCF932-10|BOLD:AAG9066
Stratiomys|[10188]|JWDCG780-10|BOLD:AAE9052
Stratiomys|[10189]|JWDCF309-10|BOLD:AAE9052
Stratiomys|[10190]|JWDCI232-10|BOLD:AAE9052
Stratiomys|[10191]|JWDCF310-10|BOLD:AAE9052
Stratiomys|[10192]|JWDCF311-10|BOLD:AAE9052
Stratiomys|[10193]|JWDCG180-10|BOLD:AAE9052
Stratiomys|[10194]|JWDCG179-10|BOLD:AAE9052
Stratiomys|[10195]|JWDCH775-10|BOLD:AAE9052
Stratiomys|[10196]|JWDCG684-10|BOLD:AAE9052
Stratiomys|[10197]|JWDCH659-10|BOLD:AAE9052
Stratiomys|[10198]|JWDCL599-11|BOLD:AAE9052
Stratiomys|[10199]|JWDCG781-10|BOLD:AAE9052
Stratiomys|[10200]|JWDCH123-10|BOLD:AAE9052
Stratiomys|[10201]|JWDCH345-10|BOLD:AAE9052
Stratiomys|[10202]|JWDCF211-10|BOLD:AAE9052
Stratiomys|[10203]|JWDCF253-10|BOLD:AAE9052
Stratiomys|[10204]|JWDCF255-10|BOLD:AAE9052
Stratiomys|[10205]|JWDCF761-10|BOLD:AAE9052
Stratiomys barbata|[10206]|JWDCH121-10|BOLD:AAE9052
Stratiomys barbata|[10207]|JWDCH122-10|BOLD:AAE9052
Stratiomys barbata|[10208]|JWDCG782-10|BOLD:AAE9052
Stratiomys barbata|[10209]|JWDCF210-10|BOLD:AAE9052
Stratiomys barbata|[10210]|JWDCF254-10|BOLD:AAE9052
Oxycera centralis|[10211]|MTDIC081-09|BOLD:AAI3016
Nemotelus bonnarius|[10212]|JWDCI257-10|BOLD:ABU5578
Nemotelus bonnarius|[10213]|JWDCI256-10|BOLD:ABU5578
Nemotelus bonnarius|[10214]|JWDCI259-10|BOLD:ABU5578
Nemotelus bonnarius|[10215]|JWDCI260-10|BOLD:ABU5578
Nemotelus bonnarius|[10216]|JWDCG134-10|BOLD:ABU5578
Nemotelus bonnarius|[10217]|JWDCG135-10|BOLD:ABU5578
Nemotelus bonnarius|[10218]|JWDCG148-10|BOLD:AAP8753
Nemotelus bonnarius|[10219]|JSFLA021-08|BOLD:AAP8753
Nemotelus bonnarius|[10220]|JWDCJ1020-11|BOLD:AAP8753
Nemotelus bonnarius|[10221]|JWDCL496-11|BOLD:AAP8753
Nemotelus bonnarius|[10222]|JWDCI927-11|BOLD:AAP8753
Nemotelus canadensis|[10223]|JWDCJ963-11|BOLD:AAI3067
Nemotelus canadensis|[10224]|JWDCI244-10|BOLD:AAI3067
Nemotelus|[10225]|MHCOL468-07|BOLD:AAI3065
Nemotelus|[10226]|MHCOL459-07|BOLD:AAI3065
Nemotelus|[10227]|TWDIP056-09|BOLD:AAI3065
Nemotelus|[10228]|MTDIC056-09|BOLD:AAI3065
Nemotelus|[10229]|JWDCE523-10|BOLD:AAI3065
Nemotelus|[10230]|JWDCF107-10|BOLD:AAI3065
Nemotelus|[10231]|TWDIP057-09|BOLD:AAI3065
Nemotelus|[10232]|JWDCF078-10|BOLD:AAI3065
Nemotelus|[10233]|MTDIC065-09|BOLD:AAI3065
Nemotelus|[10234]|JWDCE522-10|BOLD:AAI3065
Nemotelus|[10235]|MHCOL457-07|BOLD:AAI3065
Nemotelus|[10236]|MHCOL461-07|BOLD:AAI3065
Nemotelus|[10237]|TWDIP055-09|BOLD:AAI3065
Nemotelus|[10238]|TWDIP058-09|BOLD:AAI3065
Nemotelus|[10239]|MTDIC037-09|BOLD:AAI3065
Nemotelus|[10240]|MTDIC051-09|BOLD:AAI3065
Nemotelus|[10241]|JWDCG149-10|BOLD:AAI3065
Nemotelus|[10242]|JWDCI930-11|BOLD:AAI3065
Nemotelus|[10243]|JWDCE524-10|BOLD:AAI3065
Tipula breviligula|[10244]|PHTCH442-10|BOLD:AAO3962
Metalimnobia immatura|[10245]|CNCTI257-12|BOLD:ACA9696
Metalimnobia solitaria|[10246]|PHTCH090-07|BOLD:ABZ1300
Metalimnobia solitaria|[10247]|CNCTI255-12|BOLD:ABZ1300
Metalimnobia hudsonica|[10248]|PHTCH424-08|BOLD:AAB3508
Metalimnobia hudsonica|[10249]|PHTCH423-08|BOLD:AAB3508
[illegible]

Metalimnobia solitaria|[10247]|CNCTI253-12|BOLD:ABZ1300
Metalimnobia hudsonica|[10248]|PHTCH424-08|BOLD:AAB3508
Metalimnobia hudsonica|[10249]|PHTCH423-08|BOLD:AAB3508
Metalimnobia hudsonica|[10250]|PHTCH106-08|BOLD:AAB3508
Metalimnobia solitaria|[10251]|JWDCI269-10|BOLD:AAB3508
Metalimnobia hudsonica|[10252]|TICHU099-09|BOLD:AAB3508
Metalimnobia hudsonica|[10253]|CNCTI254-12|BOLD:AAB3508
Metalimnobia solitaria|[10254]|JWDCG667-10|BOLD:AAB3507
Metalimnobia solitaria|[10255]|PHTCH024-07|BOLD:AAB3507
Metalimnobia solitaria|[10256]|PHTCH035-07|BOLD:AAB3507
Metalimnobia solitaria|[10257]|PHTCH086-07|BOLD:AAB3507
Metalimnobia solitaria|[10258]|JWDCF424-10|BOLD:AAB3507
Metalimnobia solitaria|[10259]|JWDCG008-10|BOLD:AAB3507
Metalimnobia solitaria|[10260]|PHTCH070-07|BOLD:AAB3507
Metalimnobia solitaria|[10261]|JWDCG666-10|BOLD:AAB3507
Metalimnobia solitaria|[10262]|PHTCH036-07|BOLD:AAB3507
Metalimnobia solitaria|[10263]|PHTCH067-07|BOLD:AAB3507
Metalimnobia solitaria|[10264]|JWDCF930-10|BOLD:AAB3507
Metalimnobia solitaria|[10265]|PHTCH400-08|BOLD:AAB3507
Metalimnobia solitaria|[10266]|PHTCH104-08|BOLD:AAB3507
Metalimnobia solitaria|[10267]|PHTCH015-07|BOLD:AAB3507
Metalimnobia solitaria|[10268]|JWDCF788-10|BOLD:AAB3507
Metalimnobia solitaria|[10269]|JWDCI216-10|BOLD:AAB3507
Metalimnobia solitaria|[10270]|JWDCI219-10|BOLD:AAB3507
Metalimnobia solitaria|[10271]|PHTCH135-08|BOLD:AAB3507
Metalimnobia solitaria|[10272]|PHTCH333-08|BOLD:AAB3507
Metalimnobia solitaria|[10273]|PHTCH341-08|BOLD:AAB3507
Metalimnobia solitaria|[10274]|PHTCH046-07|BOLD:AAB3507
Metalimnobia solitaria|[10275]|PHTCH062-07|BOLD:AAB3507
Metalimnobia solitaria|[10276]|JWDCF426-10|BOLD:AAB3507
Metalimnobia solitaria|[10277]|JWDCG022-10|BOLD:AAB3507
Metalimnobia solitaria|[10278]|JWDCI215-10|BOLD:AAB3507
Metalimnobia solitaria|[10279]|JWDCH346-10|BOLD:AAB3507
Metalimnobia solitaria|[10280]|JWDCF926-10|BOLD:AAB3507
Metalimnobia solitaria|[10281]|PHTCH343-08|BOLD:AAB3507
Metalimnobia solitaria|[10282]|PHTCH132-08|BOLD:AAB3507
Metalimnobia solitaria|[10283]|PHTCH342-08|BOLD:AAB3507
Metalimnobia solitaria|[10284]|PHTCH105-08|BOLD:AAB3507
Metalimnobia solitaria|[10285]|PHTCH064-07|BOLD:AAB3507
Metalimnobia solitaria|[10286]|PHTCH048-07|BOLD:AAB3507
Metalimnobia solitaria|[10287]|JWDCI217-10|BOLD:AAB3507
Metalimnobia solitaria|[10288]|PHTCH074-07|BOLD:AAB3507
Metalimnobia solitaria|[10289]|JWDCI243-10|BOLD:AAB3507
Metalimnobia solitaria|[10290]|JWDCI220-10|BOLD:AAB3507
Metalimnobia solitaria|[10291]|JWDCF925-10|BOLD:AAB3507
Metalimnobia solitaria|[10292]|PHTCH063-07|BOLD:AAB3507
Metalimnobia solitaria|[10293]|CNCTI259-12|BOLD:AAB3507
Simulium vittatum complex|[10294]|MHFLY184-07|BOLD:AAA4121
Simulium vittatum complex|[10295]|JWDCI270-10|BOLD:AAA4121
Simulium vittatum complex|[10296]|MHFLI398-07|BOLD:AAA4121
Simulium vittatum complex|[10297]|JWDCI272-10|BOLD:AAA4121
Simulium vittatum complex|[10298]|JWDCJ769-11|BOLD:AAA4121
Simulium vittatum complex|[10299]|JWDCJ886-11|BOLD:AAA4121
Simulium vittatum complex|[10300]|MHFLY244-07|BOLD:AAA4121
Simulium vittatum complex|[10301]|MHFLY193-07|BOLD:AAA4121
Simulium vittatum complex|[10302]|MHFLY192-07|BOLD:AAA4121
Simulium vittatum complex|[10303]|MHFLY253-07|BOLD:AAA4121
Simulium vittatum complex|[10304]|MHFLY231-07|BOLD:AAA4121
Simulium vittatum complex|[10305]|MHFLI494-07|BOLD:AAA4121
Simulium vittatum complex|[10306]|MHFLY111-07
Simulium vittatum complex|[10307]|MHFLY255-07|BOLD:AAA4121
Simulium vittatum complex|[10308]|MHFLY302-07|BOLD:AAA4121
Simulium vittatum complex|[10309]|MHFLY269-07|BOLD:AAA4121
Simulium vittatum complex|[10310]|JWDCG166-10|BOLD:AAA4121
Simulium vittatum complex|[10311]|JWDCI370-10|BOLD:AAA4121
Simulium vittatum complex|[10312]|JWDCF647-10|BOLD:AAA4121
Simulium vittatum complex|[10313]|JWDCF810-10|BOLD:AAA4121
Simulium vittatum complex|[10314]|JWDCG131-10|BOLD:AAA4121
Simulium vittatum complex|[10315]|MHFLY224-07|BOLD:AAA4121
Simulium vittatum complex|[10316]|JWDCG171-10|BOLD:AAA4121
Simulium vittatum complex|[10317]|JWDCF554-10|BOLD:AAA4121
Simulium vittatum complex|[10318]|MHFLI400-07|BOLD:AAA4121
Simulium vittatum complex|[10319]|MHFLY251-07|BOLD:AAA4121
Simulium vittatum complex|[10320]|JWDCF638-10|BOLD:AAA4121
Simulium vittatum complex|[10321]|MHFLY124-07|BOLD:AAA4121
Simulium vittatum complex|[10322]|MHFLY236-07|BOLD:AAA4121
Simulium vittatum complex|[10323]|MHFLY314-07|BOLD:AAA4121
Simulium vittatum complex|[10324]|JWDCB254-10|BOLD:AAA4121
Simulium vittatum complex|[10325]|JWDCF815-10|BOLD:AAA4121
Simulium vittatum complex|[10326]|JWDCF823-10|BOLD:AAA4121
Simulium vittatum complex|[10327]|JWDCB261-10|BOLD:AAA4121
Simulium vittatum complex|[10328]|JWDCG173-10|BOLD:AAA4121
Simulium vittatum complex|[10329]|JWDCF825-10|BOLD:AAA4121
Simulium vittatum complex|[10330]|TWDIP001-09|BOLD:AAA4121
Simulium vittatum complex|[10331]|JWDCJ403-11|BOLD:AAA4121
Simulium vittatum complex|[10332]|JWDCH322-10|BOLD:AAA4121
Simulium vittatum complex|[10333]|MHFLI321-07|BOLD:AAA4121
Simulium vittatum complex|[10334]|MHFLI401-07|BOLD:AAA4121
Simulium vittatum complex|[10335]|JWDCF828-10|BOLD:AAA4121
Simulium vittatum complex|[10336]|JWDCF829-10|BOLD:AAA4121
Simulium vittatum complex|[10337]|MHFLY245-07|BOLD:AAA4121
Simulium vittatum complex|[10338]|MHFLY191-07|BOLD:AAA4121
Simulium vittatum complex|[10339]|MHFLI791-07|BOLD:AAA4121
Simulium vittatum complex|[10340]|MHFLY182-07|BOLD:AAA4121
Simulium vittatum complex|[10341]|MHFLY222-07|BOLD:AAA4121
Simulium vittatum complex|[10342]|MHFLY227-07|BOLD:AAA4121
Simulium vittatum complex|[10343]|MHFLY183-07|BOLD:AAA4121
Simulium vittatum complex|[10344]|MHFLI364-07|BOLD:AAA4121
Simulium vittatum complex|[10345]|MHFLY197-07|BOLD:AAA4121
Simulium vittatum complex|[10346]|MHFLY203-07|BOLD:AAA4121
Simulium vittatum complex|[10347]|MHFLY273-07|BOLD:AAA4121
Simulium vittatum complex|[10348]|MHFLY312-07|BOLD:AAA4121
Simulium vittatum complex|[10349]|JWDCL029-11|BOLD:AAA4121

Simulium vittatum complex|[10347]|MHFLY273-07|BOLD:AAA4121
Simulium vittatum complex|[10348]|MHFLY312-07|BOLD:AAA4121
Simulium vittatum complex|[10349]|JWDCL029-11|BOLD:AAA4121
Simulium vittatum complex|[10350]|JWDCG172-10|BOLD:AAA4121
Simulium vittatum complex|[10351]|JWDCI265-10|BOLD:AAA4121
Simulium vittatum complex|[10352]|JWDCI371-10|BOLD:AAA4121
Simulium vittatum complex|[10353]|JWDCG132-10|BOLD:AAA4121
Simulium vittatum complex|[10354]|JWDCE772-10|BOLD:AAA4121
Simulium vittatum complex|[10355]|JWDCE790-10|BOLD:AAA4121
Simulium vittatum complex|[10356]|JWDCF814-10|BOLD:AAA4121
Simulium vittatum complex|[10357]|JWDCF824-10|BOLD:AAA4121
Simulium vittatum complex|[10358]|JWDCE771-10|BOLD:AAA4121
Helodon gibsoni|[10359]|JWDCD230-10|BOLD:AAB0333
Helodon gibsoni|[10360]|JWDCD227-10|BOLD:AAB0333
Helodon gibsoni|[10361]|JWDCD228-10|BOLD:AAB0333
Helodon gibsoni|[10362]|JWDCD231-10|BOLD:AAB0333
Helodon gibsoni|[10363]|JWDCJ991-11|BOLD:AAB0333
Helodon gibsoni|[10364]|JWDCD226-10|BOLD:AAB0333
Helodon gibsoni|[10365]|JWDCF598-10|BOLD:AAB0333
Simulium excisum|[10366]|JWDCD410-10|BOLD:AAE8832
Simulium excisum|[10367]|JWDCJ426-11|BOLD:AAE8832
Simulium silvestre|[10368]|JWDCL384-11|BOLD:AAA8323
Simulium silvestre|[10369]|JWDCL301-11|BOLD:AAA8323
Simulium furculatum|[10370]|JWDCG176-10|BOLD:AAB1117
Simulium furculatum|[10371]|JWDCI178-10|BOLD:AAB1117
Simulium furculatum|[10372]|JWDCJ834-11|BOLD:AAB1117
Simulium furculatum|[10373]|JWDCH391-10|BOLD:AAB1117
Simulium furculatum|[10374]|JWDCJ990-11|BOLD:AAB1117
Simulium furculatum|[10375]|JWDCG178-10|BOLD:AAB1117
Simulium furculatum|[10376]|JWDCG145-10|BOLD:AAB1117
Simulium furculatum|[10377]|JWDCK089-11|BOLD:AAB1117
Simulium furculatum|[10378]|JWDCA136-10|BOLD:AAB1117
Simulium furculatum|[10379]|JWDCJ885-11|BOLD:AAB1117
Simulium furculatum|[10380]|JWDCJ832-11|BOLD:AAB1117
Simulium furculatum|[10381]|JWDCJ833-11|BOLD:AAB1117
Simulium furculatum|[10382]|JWDCI631-10|BOLD:AAB1117
Simulium furculatum|[10383]|JWDCI632-10|BOLD:AAB1117
Simulium furculatum|[10384]|JWDCI859-11|BOLD:AAB1117
Simulium furculatum|[10385]|JWDCG177-10|BOLD:AAB1117
Simulium furculatum|[10386]|JWDCI179-10|BOLD:AAB1117
Simulium furculatum|[10387]|JWDCG143-10|BOLD:AAB1117
Simulium furculatum|[10388]|JWDCG144-10|BOLD:AAB1117
Simulium furculatum|[10389]|JWDCG146-10|BOLD:AAB1117
Simulium anatinum/randalense|[10390]|JWDCD274-10|BOLD:AAE8842
Simulium anatinum/randalense|[10391]|JWDCD411-10|BOLD:AAE8842
Simulium anatinum/randalense|[10392]|JWDCD257-10|BOLD:AAE8842
Simulium anatinum/randalense|[10393]|JWDCD338-10|BOLD:AAE8842
Simulium anatinum/randalense|[10394]|JWDCD256-10|BOLD:AAE8842
Simulium anatinum/randalense|[10395]|JWDCD390-10|BOLD:AAE8842
Simulium congareenarum|[10396]|JWDCH828-10|BOLD:AAL8152
Metacnephia saskatchewana|[10397]|JWDCF165-10|BOLD:AAB7193
Metacnephia saskatchewana|[10398]|JWDCG165-10|BOLD:AAB7193
Metacnephia saskatchewana|[10399]|JWDCF816-10|BOLD:AAB7193
Metacnephia saskatchewana|[10400]|JWDCJ404-11|BOLD:AAB7193
Metacnephia saskatchewana|[10401]|JWDCI308-10|BOLD:AAB7193
Metacnephia saskatchewana|[10402]|JWDCF809-10|BOLD:AAB7193
Metacnephia saskatchewana|[10403]|JWDCF812-10|BOLD:AAB7193
Metacnephia saskatchewana|[10404]|JWDCF820-10|BOLD:AAB7193
Metacnephia saskatchewana|[10405]|JWDCF813-10|BOLD:AAB7193
Metacnephia saskatchewana|[10406]|JWDCF811-10|BOLD:AAB7193
Metacnephia saskatchewana|[10407]|JWDCG163-10|BOLD:AAB7193
Metacnephia saskatchewana|[10408]|JWDCG162-10|BOLD:AAB7193
Metacnephia saskatchewana|[10409]|JWDCG174-10|BOLD:AAB7193
Metacnephia saskatchewana|[10410]|JWDCG886-10|BOLD:AAB7193
Metacnephia saskatchewana|[10411]|JWDCJ645-11|BOLD:AAB7193
Metacnephia saskatchewana|[10412]|JWDCJ644-11|BOLD:AAB7193
Metacnephia saskatchewana|[10413]|JWDCJ551-11|BOLD:AAB7193
Metacnephia saskatchewana|[10414]|JWDCF817-10|BOLD:AAB7193
Metacnephia saskatchewana|[10415]|JWDCF821-10|BOLD:AAB7193
Metacnephia saskatchewana|[10416]|JWDCF822-10|BOLD:AAB7193
Simulium decimatum|[10417]|MHFLY258-07|BOLD:AAC0365
Simulium decimatum|[10418]|JWDCH895-10|BOLD:AAC0365
Simulium arcticum complex|[10419]|JWDCA128-10|BOLD:AAA8954
Simulium noelleri|[10420]|JWDCJ1479-11|BOLD:AAB0056
Simulium noelleri|[10421]|JWDCJ1480-11|BOLD:AAB0056
Simulium noelleri|[10422]|JWDCH700-10|BOLD:AAB0056
Simulium noelleri|[10423]|JWDCH701-10|BOLD:AAB0056
Simulium decorum|[10424]|MHFLY110-07|BOLD:ACZ4859
Simulium decorum|[10425]|MHFLY050-06|BOLD:ACZ4859
Simulium decorum|[10426]|JWDCJ1578-11|BOLD:ACZ4859
Simulium decorum|[10427]|MHFLY229-07|BOLD:ACZ4859
Simulium decorum|[10428]|JWDCH062-10|BOLD:ACZ4859
Simulium decorum|[10429]|JWDCG057-10|BOLD:ACZ4859
Simulium tuberosum complex|[10430]|JWDCB250-10|BOLD:AAA8326
Simulium tuberosum complex|[10431]|JWDCA125-10|BOLD:AAA8326
Simulium tuberosum complex|[10432]|JWDCA129-10|BOLD:AAA8326
Simulium tuberosum complex|[10433]|JWDCC557-10|BOLD:AAA8326
Simulium tuberosum complex|[10434]|JWDCA120-10|BOLD:AAA8326
Simulium tuberosum complex|[10435]|JWDCA052-10|BOLD:AAA8326
Simulium tuberosum complex|[10436]|JWDCA590-10|BOLD:AAA8326
Simulium tuberosum complex|[10437]|JWDCK174-11|BOLD:AAA8326
Simulium tuberosum complex|[10438]|JWDCB249-10|BOLD:AAA9710
Simulium tuberosum complex|[10439]|MTDIC269-09|BOLD:AAA9710
Simulium tuberosum complex|[10440]|JWDCA127-10|BOLD:AAA9710
Simulium tuberosum complex|[10441]|JWDCF819-10|BOLD:AAA9710
Simulium venustum complex|[10442]|JWDCC617-10|BOLD:AAA4264
Simulium venustum complex|[10443]|SWSWP069-09|BOLD:AAA4264
Simulium venustum complex|[10444]|JWDCF407-10|BOLD:AAA4264
Simulium venustum complex|[10445]|JWDCE789-10|BOLD:AAA4264
Simulium venustum complex|[10446]|JWDCH231-10|BOLD:AGB0586
Simulium venustum complex|[10447]|MHFLY219-07|BOLD:AGB0586
Simulium venustum complex|[10448]|FLIMH036-08|BOLD:AGB0586
Simulium venustum complex|[10449]|JWDCC754-10|BOLD:AGB0586

Simulium venustum complex|[10446]|JWDCH231-10|BOLD:AGB0586
Simulium venustum complex|[10447]|MHFLY219-07|BOLD:AGB0586
Simulium venustum complex|[10448]|FLIMH036-08|BOLD:AGB0586
Simulium venustum complex|[10449]|JWDCG754-10|BOLD:AGB0586
Simulium venustum complex|[10450]|MHFLY233-07|BOLD:AGB0586
Simulium venustum complex|[10451]|MTDIC046-09|BOLD:AGB0586
Simulium venustum complex|[10452]|MHFLI851-07|BOLD:AGB0586
Simulium venustum complex|[10453]|MHFLY200-07|BOLD:AGB0586
Simulium venustum complex|[10454]|MHFLY301-07|BOLD:AGB0586
Simulium venustum complex|[10455]|JWDCL358-11|BOLD:AGB0586
Simulium venustum complex|[10456]|MHFLY189-07|BOLD:AGB0586
Simulium venustum complex|[10457]|MHFLY324-07|BOLD:AGB0586
Simulium venustum complex|[10458]|JWDCK038-11|BOLD:AGB0586
Simulium venustum complex|[10459]|JWDCH896-10|BOLD:AGB0586
Simulium venustum complex|[10460]|JWDCI051-10|BOLD:AGB0586
Simulium venustum complex|[10461]|MHFLI323-07|BOLD:AGB0586
Simulium venustum complex|[10462]|SWSWP025-09|BOLD:AGB0586
Simulium venustum complex|[10463]|JWDCI001-10|BOLD:AGB0586
Simulium venustum complex|[10464]|MHFLY051-06|BOLD:AGB0586
Simulium venustum complex|[10465]|MHFLY173-07|BOLD:AGB0586
Simulium venustum complex|[10466]|MHFLY225-07|BOLD:AGB0586
Simulium venustum complex|[10467]|MHFLY243-07|BOLD:AGB0586
Simulium venustum complex|[10468]|JWDCC556-10|BOLD:AGB0586
Simulium venustum complex|[10469]|JWDCJ683-11|BOLD:AGB0586
Simulium venustum complex|[10470]|JWDCF039-10|BOLD:AGB0586
Simulium venustum complex|[10471]|JWDCF853-10|BOLD:AGB0586
Simulium venustum complex|[10472]|MHFLY352-07|BOLD:AGB0586
Simulium venustum complex|[10473]|JWDCI568-10|BOLD:AGB0586
Simulium venustum complex|[10474]|JWDCG458-10|BOLD:AGB0586
Simulium venustum complex|[10475]|SWSWP022-09|BOLD:AGB0586
Simulium venustum complex|[10476]|JWDCI252-10|BOLD:AGB0586
Simulium venustum complex|[10477]|JWDCI566-10|BOLD:AGB0586
Simulium venustum complex|[10478]|JWDCC616-10|BOLD:AGB0586
Simulium venustum complex|[10479]|MHFLY249-07|BOLD:AGB0586
Simulium venustum complex|[10480]|JWDCF038-10|BOLD:AGB0586
Simulium venustum complex|[10481]|JWDCI254-10|BOLD:AGB0586
Simulium venustum complex|[10482]|JWDCF184-10|BOLD:AGB0586
Simulium venustum complex|[10483]|JWDCC654-10|BOLD:AGB0586
Simulium venustum complex|[10484]|JWDCC525-10|BOLD:AGB0586
Simulium venustum complex|[10485]|SWSWP015-09|BOLD:AGB0586
Simulium venustum complex|[10486]|SWSWP062-09|BOLD:AGB0586
Simulium venustum complex|[10487]|JWDCC652-10|BOLD:AGB0586
Simulium venustum complex|[10488]|JWDCJ770-11|BOLD:AGB0586
Simulium venustum complex|[10489]|JWDCJ771-11|BOLD:AGB0586
Simulium venustum complex|[10490]|JWDCC555-10|BOLD:AGB0586
Simulium venustum complex|[10491]|SWSWP007-09|BOLD:AGB0586
Simulium venustum complex|[10492]|JWDCG884-10|BOLD:AGB0586
Simulium venustum complex|[10493]|JWDCI933-11|BOLD:AGB0586
Simulium venustum complex|[10494]|SWSWP018-09|BOLD:AGB0586
Simulium venustum complex|[10495]|SWSWP020-09|BOLD:AGB0586
Simulium venustum complex|[10496]|SWSWP014-09|BOLD:AGB0586
Simulium venustum complex|[10497]|SWSWP012-09|BOLD:AGB0586
Simulium venustum complex|[10498]|JWDCJ292-11|BOLD:AGB0586
Simulium venustum complex|[10499]|JWDCG175-10|BOLD:AGB0586
Simulium venustum complex|[10500]|JWDCG199-10|BOLD:AGB0586
Simulium venustum complex|[10501]|JWDCE915-10|BOLD:AGB0586
Simulium venustum complex|[10502]|JWDCI595-10|BOLD:AGB0586
Simulium venustum complex|[10503]|JWDCJ405-11|BOLD:AGB0586
Simulium venustum complex|[10504]|MHFLY226-07|BOLD:AGB0586
Simulium venustum complex|[10505]|MHFLY317-07|BOLD:AGB0586
Simulium venustum complex|[10506]|JWDCF182-10|BOLD:AGB0586
Simulium venustum complex|[10507]|SWSWP038-09|BOLD:AGB0586
Simulium venustum complex|[10508]|MTDIC041-09|BOLD:AGB0586
Simulium venustum complex|[10509]|JWDCJ143-11|BOLD:AGB0586
Simulium venustum complex|[10510]|SWSWP092-09|BOLD:AGB0586
Simulium venustum complex|[10511]|JWDCF852-10|BOLD:AGB0586
Simulium venustum complex|[10512]|JWDCG885-10|BOLD:AGB0586
Simulium venustum complex|[10513]|JWDCC651-10|BOLD:AGB0586
Simulium venustum complex|[10514]|SWSWP021-09|BOLD:AGB0586
Simulium venustum complex|[10515]|MHFLY237-07|BOLD:AGB0586
Simulium venustum complex|[10516]|JWDCF037-10|BOLD:AGB0586
Simulium venustum complex|[10517]|JWDCG168-10|BOLD:AGB0586
Simulium venustum complex|[10518]|JWDCD334-10|BOLD:AGB0586
Simulium venustum complex|[10519]|JWDCC524-10|BOLD:AGB0586
Simulium venustum complex|[10520]|JWDCH496-10|BOLD:AGB0586
Simulium venustum complex|[10521]|SWSWP066-09|BOLD:AGB0586
Simulium venustum complex|[10522]|MHFLY259-07|BOLD:AGB0586
Simulium venustum complex|[10523]|MHFLY328-07|BOLD:AGB0586
Simulium venustum complex|[10524]|MHFLY353-07|BOLD:AGB0586
Simulium venustum complex|[10525]|JWDCI372-10|BOLD:AGB0586
Simulium venustum complex|[10526]|JWDCG883-10|BOLD:AGB0586
Simulium venustum complex|[10527]|JWDCI567-10|BOLD:AGB0586
Simulium venustum complex|[10528]|JWDCE788-10|BOLD:AGB0586
Simulium venustum complex|[10529]|JWDCI683-11|BOLD:AGB0586
Simulium venustum complex|[10530]|JWDCD335-10|BOLD:AGB0586
Simulium venustum complex|[10531]|MHFLI317-07|BOLD:AGB0586
Simulium venustum complex|[10532]|SWSWP064-09|BOLD:AGB0586
Simulium venustum complex|[10533]|JWDCD244-10|BOLD:AGB0586
Simulium venustum complex|[10534]|JWDCA589-10|BOLD:AGB0586
Simulium venustum complex|[10535]|JWDCF516-10|BOLD:AGB0586
Simulium venustum complex|[10536]|JWDCF831-10|BOLD:AGB0586
Simulium venustum complex|[10537]|JWDCD440-10|BOLD:AGB0586
Simulium venustum complex|[10538]|SWSWP017-09|BOLD:AGB0586
Simulium venustum complex|[10539]|JWDCF227-10|BOLD:AGB0586
Simulium venustum complex|[10540]|JWDCF183-10|BOLD:AGB0586
Simulium venustum complex|[10541]|SWSWP024-09|BOLD:AGB0586
Simulium venustum complex|[10542]|SWSWP008-09|BOLD:AGB0586
Simulium venustum complex|[10543]|JWDCF346-10|BOLD:AGB0586
Simulium venustum complex|[10544]|JWDCE916-10|BOLD:AGB0586
Simulium venustum complex|[10545]|JWDCB256-10|BOLD:AGB0586
Simulium venustum complex|[10546]|JWDCJ682-11|BOLD:AGB0586
Simulium venustum complex|[10547]|JWDCF517-10|BOLD:AGB0586
Simulium venustum complex|[10548]|JWDCE917-10|BOLD:AGB0586

Simulium venustum complex|[10546]|JWDCJ682-11|BOLD:AGB0586
Simulium venustum complex|[10547]|JWDCF517-10|BOLD:AGB0586
Simulium venustum complex|[10548]|JWDCE917-10|BOLD:AGB0586
Simulium verecundum complex|[10549]|MHFLY299-07
Simulium verecundum complex|[10550]|MHFLY178-07|BOLD:AAA1697
Simulium verecundum complex|[10551]|MHFLY319-07|BOLD:AAA1697
Simulium verecundum complex|[10552]|JWDCH320-10|BOLD:AAA1697
Simulium verecundum complex|[10553]|MHFLY268-07|BOLD:AAA1697
Simulium verecundum complex|[10554]|MHFLY337-07|BOLD:AAA1697
Simulium verecundum complex|[10555]|JWDCH625-10|BOLD:AAA1697
Simulium verecundum complex|[10556]|MHFLY265-07|BOLD:AAA1697
Simulium verecundum complex|[10557]|JWDCH563-10|BOLD:AAA1697
Simulium verecundum complex|[10558]|JWDCJ1227-11|BOLD:AAA1697
Simulium verecundum complex|[10559]|JWDCK088-11|BOLD:AAA1697
Simulium verecundum complex|[10560]|JWDCB253-10|BOLD:AAA1697
Simulium verecundum complex|[10561]|MTDIC268-09|BOLD:AAA1697
Simulium verecundum complex|[10562]|MTDIC261-09|BOLD:AAA1697
Simulium verecundum complex|[10563]|JWDCB158-10|BOLD:AAA1697
Simulium verecundum complex|[10564]|MHCOL334-07|BOLD:AAA1697
Simulium verecundum complex|[10565]|MHFLY326-07|BOLD:AAA1697
Simulium verecundum complex|[10566]|JWDCJ1224-11|BOLD:AAA1697
Simulium verecundum complex|[10567]|JWDCJ1393-11|BOLD:AAA1697
Simulium verecundum complex|[10568]|JWDCG234-10|BOLD:AAA1697
Simulium verecundum complex|[10569]|JWDCH653-10|BOLD:AAA1697
Simulium verecundum complex|[10570]|JWDCG707-10|BOLD:AAA1697
Simulium verecundum complex|[10571]|JWDCG774-10|BOLD:AAA1697
Simulium verecundum complex|[10572]|JWDCG795-10|BOLD:AAA1697
Simulium verecundum complex|[10573]|JWDCI586-10|BOLD:AAA1697
Simulium verecundum complex|[10574]|JWDCH063-10|BOLD:AAA1697
Simulium verecundum complex|[10575]|JWDCI681-11|BOLD:AAA1697
Simulium verecundum complex|[10576]|JWDCI724-11|BOLD:AAA1697
Simulium verecundum complex|[10577]|JWDCI013-10|BOLD:AAA1697
Simulium verecundum complex|[10578]|JWDCI014-10|BOLD:AAA1697
Simulium verecundum complex|[10579]|JWDCG257-10|BOLD:AAA1697
Simulium verecundum complex|[10580]|JWDCL359-11|BOLD:AAA1697
Simulium verecundum complex|[10581]|MHFLY181-07|BOLD:AAA1697
Simulium verecundum complex|[10582]|FLIMH023-08|BOLD:AAA1697
Simulium verecundum complex|[10583]|JWDCH561-10|BOLD:AAA1697
Simulium verecundum complex|[10584]|JWDCJ142-11|BOLD:AAA1697
Simulium verecundum complex|[10585]|JWDCA139-10|BOLD:AAA1697
Simulium verecundum complex|[10586]|MHCOL210-07|BOLD:AAA1697
Simulium verecundum complex|[10587]|JWDCL385-11|BOLD:AAA1697
Simulium verecundum complex|[10588]|MHFLY347-07|BOLD:AAA1697
Simulium verecundum complex|[10589]|JWDCB258-10|BOLD:AAA1697
Simulium verecundum complex|[10590]|MHCOL335-07|BOLD:AAA1697
Simulium verecundum complex|[10591]|MHCOL327-07|BOLD:AAA1697
Simulium verecundum complex|[10592]|JWDCL170-11|BOLD:AAA1697
Simulium verecundum complex|[10593]|JWDCI806-11|BOLD:AAA1697
Simulium verecundum complex|[10594]|FLIMH017-08|BOLD:AAA1697
Simulium verecundum complex|[10595]|MHFLY176-07|BOLD:AAA1697
Simulium verecundum complex|[10596]|MHFLI397-07|BOLD:AAA1697
Simulium verecundum complex|[10597]|MHFLY277-07|BOLD:AAA1697
Simulium verecundum complex|[10598]|MHFLY303-07|BOLD:AAA1697
Simulium verecundum complex|[10599]|MHFLY171-07|BOLD:AAA1697
Simulium verecundum complex|[10600]|MHFLY177-07|BOLD:AAA1697
Simulium verecundum complex|[10601]|MHCOL209-07|BOLD:AAA1697
Simulium verecundum complex|[10602]|JWDCH181-10|BOLD:AAA1697
Simulium verecundum complex|[10603]|JWDCI003-10|BOLD:AAA1697
Simulium verecundum complex|[10604]|JWDCH180-10|BOLD:AAA1697
Simulium verecundum complex|[10605]|MHFLY180-07|BOLD:AAA1697
Simulium verecundum complex|[10606]|MHCOL328-07|BOLD:AAA1697
Simulium verecundum complex|[10607]|JWDCL171-11|BOLD:AAA1697
Simulium verecundum complex|[10608]|JWDCG258-10|BOLD:AAA1697
Simulium verecundum complex|[10609]|MHFLY188-07|BOLD:AAA1697
Simulium verecundum complex|[10610]|MHFLY275-07|BOLD:AAA1697
Simulium verecundum complex|[10611]|MHFLY232-07|BOLD:AAA1697
Simulium verecundum complex|[10612]|JWDCH827-10|BOLD:AAA1697
Simulium verecundum complex|[10613]|JWDCI002-10|BOLD:AAA1697
Simulium verecundum complex|[10614]|JWDCF164-10|BOLD:AAA1697
Simulium verecundum complex|[10615]|FLIMH033-08|BOLD:AAA1697
Simulium verecundum complex|[10616]|JWDCF416-10|BOLD:AAA1697
Simulium verecundum complex|[10617]|MHFLY264-07|BOLD:AAA1697
Simulium verecundum complex|[10618]|JWDCL172-11|BOLD:AAA1697
Simulium verecundum complex|[10619]|MHFLY262-07|BOLD:AAA1697
Simulium verecundum complex|[10620]|MHFLY290-07|BOLD:AAA1697
Simulium verecundum complex|[10621]|MHFLY282-07|BOLD:AAA1697
Simulium verecundum complex|[10622]|MHFLY172-07|BOLD:AAA1697
Simulium verecundum complex|[10623]|MHFLY246-07|BOLD:AAA1697
Simulium verecundum complex|[10624]|MHCOL332-07|BOLD:AAA1697
Simulium verecundum complex|[10625]|MHCOL111-07|BOLD:AAA1697
Simulium verecundum complex|[10626]|JWDCA133-10|BOLD:AAA1697
Simulium verecundum complex|[10627]|FLIMH022-08|BOLD:AAA1697
Simulium verecundum complex|[10628]|MHFLY241-07|BOLD:AAA1697
Simulium verecundum complex|[10629]|MHFLI395-07|BOLD:AAA1697
Simulium verecundum complex|[10630]|MHFLY174-07|BOLD:AAA1697
Simulium verecundum complex|[10631]|JWDCH442-10|BOLD:AAA1697
Simulium verecundum complex|[10632]|MHFLY311-07|BOLD:AAA1697
Simulium verecundum complex|[10633]|MHFLY284-07|BOLD:AAA1697
Simulium verecundum complex|[10634]|JWDCJ1305-11|BOLD:AAA1697
Simulium verecundum complex|[10635]|JWDCF837-10|BOLD:AAA1697
Simulium verecundum complex|[10636]|MHFLI388-07|BOLD:AAA1697
Simulium verecundum complex|[10637]|MHFLY286-07|BOLD:AAA1697
Simulium verecundum complex|[10638]|MHFLI393-07|BOLD:AAA1697
Simulium verecundum complex|[10639]|MHFLI390-07|BOLD:AAA1697
Simulium verecundum complex|[10640]|MHFLY280-07|BOLD:AAA1697
Simulium verecundum complex|[10641]|JWDCI016-10|BOLD:AAA1697
Simulium verecundum complex|[10642]|MHFLY291-07|BOLD:AAA1697
Simulium verecundum complex|[10643]|JWDCJ1304-11|BOLD:AAA1697
Simulium verecundum complex|[10644]|JWDCB262-10|BOLD:AAA1697
Simulium verecundum complex|[10645]|MHFLY331-07|BOLD:AAA1697
Simulium verecundum complex|[10646]|MHFLY320-07|BOLD:AAA1697
Simulium verecundum complex|[10647]|MHCOL339-07|BOLD:AAA1697
Simulium verecundum complex|[10648]|[illegible]

Simulium verecundum complex|[10645]|MHFLY331-07|BOLD:AAA1697
Simulium verecundum complex|[10646]|MHFLY320-07|BOLD:AAA1697
Simulium verecundum complex|[10647]|MHCOL339-07|BOLD:AAA1697
Simulium verecundum complex|[10648]|JWDCG456-10|BOLD:AAA1697
Simulium verecundum complex|[10649]|JWDCK173-11|BOLD:AAA1697
Simulium verecundum complex|[10650]|JWDCJ1226-11|BOLD:AAA1697
Simulium verecundum complex|[10651]|MHFLI310-07|BOLD:AAA1697
Simulium verecundum complex|[10652]|MHFLY247-07|BOLD:AAA1697
Simulium verecundum complex|[10653]|MHFLY190-07|BOLD:AAA1697
Simulium verecundum complex|[10654]|JWDCG708-10|BOLD:AAA1697
Simulium verecundum complex|[10655]|JWDCH444-10|BOLD:AAA1697
Simulium verecundum complex|[10656]|JWDCI049-10|BOLD:AAA1697
Simulium verecundum complex|[10657]|JWDCH232-10|BOLD:AAA1697
Simulium verecundum complex|[10658]|JWDCG059-10|BOLD:AAA1697
Simulium verecundum complex|[10659]|JWDCG457-10|BOLD:AAA1697
Simulium verecundum complex|[10660]|JWDCH233-10|BOLD:AAA1697
Simulium verecundum complex|[10661]|JWDCJ1303-11|BOLD:AAA1697
Simulium verecundum complex|[10662]|JWDCJ1035-11|BOLD:AAA1697
Simulium verecundum complex|[10663]|JWDCA130-10|BOLD:AAA1697
Simulium verecundum complex|[10664]|JWDCA123-10|BOLD:AAA1697
Simulium verecundum complex|[10665]|MHFLY207-07|BOLD:AAA1697
Simulium verecundum complex|[10666]|MHFLY330-07|BOLD:AAA1697
Simulium verecundum complex|[10667]|MHFLY329-07|BOLD:AAA1697
Simulium verecundum complex|[10668]|JWDCI050-10|BOLD:AAA1697
Simulium verecundum complex|[10669]|MHFLY204-07|BOLD:AAA1697
Simulium verecundum complex|[10670]|MHFLY348-07|BOLD:AAA1697
Simulium verecundum complex|[10671]|MHFLY274-07|BOLD:AAA1697
Simulium verecundum complex|[10672]|MHFLY339-07|BOLD:AAA1697
Simulium verecundum complex|[10673]|MHFLY285-07|BOLD:AAA1697
Simulium verecundum complex|[10674]|MHFLI396-07|BOLD:AAA1697
Simulium verecundum complex|[10675]|MHFLY168-07|BOLD:AAA1697
Simulium verecundum complex|[10676]|JWDCI934-11|BOLD:AAA1697
Simulium verecundum complex|[10677]|JWDCG521-10|BOLD:AAA1697
Simulium verecundum complex|[10678]|JWDCG259-10|BOLD:AAA1697
Simulium verecundum complex|[10679]|JWDCF851-10|BOLD:AAA1697
Simulium verecundum complex|[10680]|MHFLY266-07|BOLD:AAA1697
Simulium verecundum complex|[10681]|MHFLY279-07|BOLD:AAA1697
Simulium verecundum complex|[10682]|JWDCJ1036-11|BOLD:AAA1697
Simulium verecundum complex|[10683]|MHFLI783-07|BOLD:AAA1697
Simulium verecundum complex|[10684]|MHFLY351-07|BOLD:AAA1697
Simulium verecundum complex|[10685]|MHFLY338-07|BOLD:AAA1697
Simulium verecundum complex|[10686]|JWDCJ141-11|BOLD:AAA1697
Simulium verecundum complex|[10687]|MHFLY212-07|BOLD:AAA1697
Simulium verecundum complex|[10688]|JWDCH962-10|BOLD:AAA1697
Simulium verecundum complex|[10689]|MHFLY166-07|BOLD:AAA1697
Simulium verecundum complex|[10690]|MHFLY201-07|BOLD:AAA1697
Simulium verecundum complex|[10691]|MHFLY205-07|BOLD:AAA1697
Simulium verecundum complex|[10692]|MHFLY208-07|BOLD:AAA1697
Simulium verecundum complex|[10693]|JWDCI015-10|BOLD:AAA1697
Simulium verecundum complex|[10694]|JWDCG709-10|BOLD:AAA1697
Simulium verecundum complex|[10695]|MHCOL330-07|BOLD:AAA1697
Simulium verecundum complex|[10696]|MHCOL333-07|BOLD:AAA1697
Simulium verecundum complex|[10697]|JWDCA121-10|BOLD:AAA1697
Simulium verecundum complex|[10698]|JWDCI528-10|BOLD:AAA1697
Simulium verecundum complex|[10699]|JWDCI725-11|BOLD:AAA1697
Simulium|[10700]|JWDCA126-10|BOLD:AAA1697
Simulium verecundum complex|[10701]|JWDCL101-11|BOLD:AAA1697
Simulium verecundum complex|[10702]|MHFLY217-07|BOLD:AAA1697
Simulium verecundum complex|[10703]|JWDCG746-10|BOLD:AAA1697
Simulium verecundum complex|[10704]|MHFLY209-07|BOLD:AAA1697
Simulium verecundum complex|[10705]|JWDCL360-11|BOLD:AAA1697
Simulium verecundum complex|[10706]|JWDCI630-10|BOLD:AAA1697
Simulium verecundum complex|[10707]|JWDCI807-11|BOLD:AAA1697
Simulium verecundum complex|[10708]|MHFLY261-07|BOLD:AAA1697
Simulium verecundum complex|[10709]|MHFLY263-07|BOLD:AAA1697
Simulium verecundum complex|[10710]|JWDCJ1223-11|BOLD:AAA1697
Simulium verecundum complex|[10711]|JWDCG523-10|BOLD:AAA1697
Simulium verecundum complex|[10712]|MHFLY298-07|BOLD:AAA1697
Simulium|[10713]|JWDCA131-10|BOLD:AAA1697
Simulium verecundum complex|[10714]|JWDCB252-10|BOLD:AAA1697
Simulium verecundum complex|[10715]|JWDCB257-10|BOLD:AAA1697
Simulium verecundum complex|[10716]|MHFLY175-07|BOLD:AAA1697
Simulium verecundum complex|[10717]|MHFLY344-07|BOLD:AAA1697
Simulium verecundum complex|[10718]|JWDCG811-10|BOLD:AAA1697
Simulium verecundum complex|[10719]|MHFLY214-07|BOLD:AAA1697
Simulium verecundum complex|[10720]|JWDCJ225-11|BOLD:AAA1697
Simulium verecundum complex|[10721]|JWDCH011-10|BOLD:AAA1697
Simulium verecundum complex|[10722]|JWDCH880-10|BOLD:AAA1697
Simulium verecundum complex|[10723]|MHFLY287-07|BOLD:AAA1697
Simulium verecundum complex|[10724]|JWDCH064-10|BOLD:AAA1697
Simulium verecundum complex|[10725]|MHFLI384-07|BOLD:AAA1697
Simulium verecundum complex|[10726]|MHFLY272-07|BOLD:AAA1697
Simulium verecundum complex|[10727]|MHFLY194-07|BOLD:AAA1697
Simulium verecundum complex|[10728]|JWDCK091-11|BOLD:AAA1697
Simulium verecundum complex|[10729]|MHFLY335-07|BOLD:AAA1697
Simulium verecundum complex|[10730]|MHFLY254-07|BOLD:AAA1697
Simulium verecundum complex|[10731]|JWDCK036-11|BOLD:AAA1697
Simulium verecundum complex|[10732]|MHFLI394-07|BOLD:AAA1697
Simulium verecundum complex|[10733]|JWDCB255-10|BOLD:AAA1697
Simulium verecundum complex|[10734]|MHFLY293-07|BOLD:AAA1697
Simulium verecundum complex|[10735]|MHFLY313-07|BOLD:AAA1697
Simulium verecundum complex|[10736]|JWDCH755-10|BOLD:AAA1697
Simulium verecundum complex|[10737]|MHFLY211-07|BOLD:AAA1697
Simulium verecundum complex|[10738]|MHFLY288-07|BOLD:AAA1697
Simulium verecundum complex|[10739]|MHFLY250-07|BOLD:AAA1697
Simulium verecundum complex|[10740]|MHFLY213-07|BOLD:AAA1697
Simulium verecundum complex|[10741]|JWDCB259-10|BOLD:AAA1697
Simulium verecundum complex|[10742]|JWDCL028-11|BOLD:AAA1697
Simulium verecundum complex|[10743]|JWDCH065-10|BOLD:AAA1697
Simulium verecundum complex|[10744]|MHFLY289-07|BOLD:AAA1697
Simulium verecundum complex|[10745]|MHFLY300-07|BOLD:AAA1697
Simulium verecundum complex|[10746]|MHFLY239-07|BOLD:AAA1697
Simulium verecundum complex|[10747]|FLIMH019-08|BOLD:AAA1697

Simulium verecundum complex|[10745]|MHFLY300-07|BOLD:AAA1697
Simulium verecundum complex|[10746]|MHFLY239-07|BOLD:AAA1697
Simulium verecundum complex|[10747]|FLIMH019-08|BOLD:AAA1697
Simulium verecundum complex|[10748]|FLIMH021-08|BOLD:AAA1697
Simulium verecundum complex|[10749]|MHFLI320-07|BOLD:AAA1697
Simulium verecundum complex|[10750]|MHFLY114-07|BOLD:AAA1697
Simulium verecundum complex|[10751]|JWDCJ365-11|BOLD:AAA1697
Simulium verecundum complex|[10752]|JWDCL578-11|BOLD:AAA1697
Simulium verecundum complex|[10753]|JWDCL444-11|BOLD:AAA1697
Simulium verecundum complex|[10754]|JWDCL445-11|BOLD:AAA1697
Simulium verecundum complex|[10755]|JWDCH566-10|BOLD:AAA1697
Simulium verecundum complex|[10756]|JWDCH654-10|BOLD:AAA1697
Simulium verecundum complex|[10757]|JWDCG753-10|BOLD:AAA1697
Simulium verecundum complex|[10758]|JWDCG755-10|BOLD:AAA1697
Simulium verecundum complex|[10759]|JWDCH756-10|BOLD:AAA1697
Simulium verecundum complex|[10760]|JWDCG895-10|BOLD:AAA1697
Simulium verecundum complex|[10761]|JWDCH255-10|BOLD:AAA1697
Simulium verecundum complex|[10762]|JWDCH562-10|BOLD:AAA1697
Simulium verecundum complex|[10763]|JWDCF383-10|BOLD:AAA1697
Simulium verecundum complex|[10764]|JWDCF384-10|BOLD:AAA1697
Simulium verecundum complex|[10765]|MHCOL338-07|BOLD:AAA1697
Simulium verecundum complex|[10766]|JWDCK037-11|BOLD:AAA1697
Simulium verecundum complex|[10767]|JWDCF385-10|BOLD:AAA1697
Simulium verecundum complex|[10768]|TWDIP002-09|BOLD:AAA1697
Simulium verecundum complex|[10769]|JWDCH894-10|BOLD:AAA1697
Simulium verecundum complex|[10770]|JWDCI858-11|BOLD:AAA1697
Simulium verecundum complex|[10771]|JWDCF406-10|BOLD:AAA1697
Simulium verecundum complex|[10772]|MHFLY306-07|BOLD:AAA1697
Simulium verecundum complex|[10773]|JWDCH443-10|BOLD:AAA1697
Simulium verecundum complex|[10774]|JWDCG896-10|BOLD:AAA1697
Simulium verecundum complex|[10775]|MHFLY235-07|BOLD:AAA1697
Simulium verecundum complex|[10776]|MHFLY307-07|BOLD:AAA1697
Simulium verecundum complex|[10777]|MHFLY295-07|BOLD:AAA1697
Simulium verecundum complex|[10778]|MHFLY215-07|BOLD:AAA1697
Simulium verecundum complex|[10779]|MHFLY220-07|BOLD:AAA1697
Simulium verecundum complex|[10780]|MHCOL329-07|BOLD:AAA1697
Simulium verecundum complex|[10781]|JWDCH826-10|BOLD:AAA1697
Simulium verecundum complex|[10782]|MHFLY228-07|BOLD:AAA1697
Simulium verecundum complex|[10783]|JWDCK008-11|BOLD:AAA1697
Simulium verecundum complex|[10784]|JWDCL217-11|BOLD:AAA1697
Simulium verecundum complex|[10785]|JWDCG812-10|BOLD:AAA1697
Simulium verecundum complex|[10786]|JWDCL446-11|BOLD:AAA1697
Simulium verecundum complex|[10787]|MHFLY202-07|BOLD:AAA1697
Simulium verecundum complex|[10788]|MHFLY260-07|BOLD:AAA1697
Simulium verecundum complex|[10789]|MHFLY341-07|BOLD:AAA1697
Simulium verecundum complex|[10790]|MHFLY252-07|BOLD:AAA1697
Simulium verecundum complex|[10791]|MHFLY327-07|BOLD:AAA1697
Simulium verecundum complex|[10792]|MHFLY345-07|BOLD:AAA1697
Simulium verecundum complex|[10793]|JWDCK009-11|BOLD:AAA1697
Simulium verecundum complex|[10794]|JWDCH010-10|BOLD:AAA1697
Simulium verecundum complex|[10795]|MHFLY271-07|BOLD:AAA1697
Simulium verecundum complex|[10796]|MHFLY210-07|BOLD:AAA1697
Simulium verecundum complex|[10797]|MHFLY278-07|BOLD:AAA1697
Simulium verecundum complex|[10798]|MHFLY216-07|BOLD:AAA1697
Simulium verecundum complex|[10799]|JWDCI682-11|BOLD:AAA1697
Simulium verecundum complex|[10800]|JWDCH150-10|BOLD:AAA1697
Simulium verecundum complex|[10801]|JWDCL383-11|BOLD:AAA1697
Simulium verecundum complex|[10802]|MHFLY267-07|BOLD:AAA1697
Simulium verecundum complex|[10803]|FLIMH018-08|BOLD:AAA1697
Simulium verecundum complex|[10804]|MHFLY333-07|BOLD:AAA1697
Simulium verecundum complex|[10805]|JWDCK175-11|BOLD:AAA1697
Simulium verecundum complex|[10806]|JWDCJ1225-11|BOLD:AAA1697
Simulium verecundum complex|[10807]|MHFLY185-07|BOLD:AAA1697
Simulium verecundum complex|[10808]|JWDCL100-11|BOLD:AAA1697
Simulium verecundum complex|[10809]|JWDCG810-10|BOLD:AAA1697
Simulium verecundum complex|[10810]|MHFLI389-07|BOLD:AAA1697
Simulium verecundum complex|[10811]|JWDCK087-11|BOLD:AAA1697
Simulium verecundum complex|[10812]|MHFLY113-07|BOLD:AAA1697
Simulium verecundum complex|[10813]|MHFLI473-07|BOLD:AAA1697
Simulium verecundum complex|[10814]|MHFLY340-07|BOLD:AAA1697
Simulium verecundum complex|[10815]|MHFLI399-07|BOLD:AAA1697
Simulium verecundum complex|[10816]|MHFLY198-07|BOLD:AAA1697
Simulium verecundum complex|[10817]|JWDCJ367-11|BOLD:AAA1697
Simulium verecundum complex|[10818]|JWDCL579-11|BOLD:AAA1697
Simulium verecundum complex|[10819]|MHFLY223-07|BOLD:AAA1697
Simulium verecundum complex|[10820]|MHFLY234-07|BOLD:AAA1697
Simulium verecundum complex|[10821]|MHFLY346-07|BOLD:AAA1697
Simulium verecundum complex|[10822]|JWDCA134-10|BOLD:AAA1697
Simulium verecundum complex|[10823]|JWDCH321-10|BOLD:AAA1697
Simulium verecundum complex|[10824]|MHFLY325-07|BOLD:AAA1697
Simulium verecundum complex|[10825]|JWDCF415-10|BOLD:AAA1697
Simulium verecundum complex|[10826]|FLIMH020-08|BOLD:AAA1697
Simulium verecundum complex|[10827]|MHFLY167-07|BOLD:AAA1697
Simulium verecundum complex|[10828]|MHFLY256-07|BOLD:AAA1697
Simulium verecundum complex|[10829]|MHFLI456-07|BOLD:AAA1697
Simulium verecundum complex|[10830]|MHFLI383-07|BOLD:AAA1697
Simulium verecundum complex|[10831]|MHFLY257-07|BOLD:AAA1697
Simulium verecundum complex|[10832]|JWDCH964-10|BOLD:AAA1697
Simulium verecundum complex|[10833]|MHFLY109-07|BOLD:AAA1697
Simulium verecundum complex|[10834]|JWDCB260-10|BOLD:AAA1697
Simulium verecundum complex|[10835]|MHFLY242-07|BOLD:AAA1697
Simulium verecundum complex|[10836]|JWDCA122-10|BOLD:AAA1697
Simulium verecundum complex|[10837]|MHFLY318-07|BOLD:AAA1697
Simulium verecundum complex|[10838]|JWDCH754-10|BOLD:AAA1697
Simulium verecundum complex|[10839]|MHFLY221-07|BOLD:AAA1697
Simulium verecundum complex|[10840]|JWDCL218-11|BOLD:AAA1697
Simulium verecundum complex|[10841]|JWDCG058-10|BOLD:AAA1697
Simulium verecundum complex|[10842]|JWDCI529-10|BOLD:AAA1697
Simulium verecundum complex|[10843]|JWDCI124-10|BOLD:AAA1697
Simulium verecundum complex|[10844]|JWDCJ1037-11|BOLD:AAA1697
Simulium verecundum complex|[10845]|MHCOL331-07|BOLD:AAA1697
Simulium verecundum complex|[10846]|MHCOL337-07|BOLD:AAA1697
[illegible]

Simulium verecundum complex|[10844]|JWDCJ1037-11|BOLD:AAA1697
Simulium verecundum complex|[10845]|MHCOL331-07|BOLD:AAA1697
Simulium verecundum complex|[10846]|MHCOL337-07|BOLD:AAA1697
Simulium verecundum complex|[10847]|MHCOL326-07|BOLD:AAA1697
Simulium|[10848]|JWDCA137-10|BOLD:AAA1697
Simulium verecundum complex|[10849]|MHFLI365-07|BOLD:AAA1697
Simulium verecundum complex|[10850]|MHFLY323-07|BOLD:AAA1697
Simulium verecundum complex|[10851]|MHFLY336-07|BOLD:AAA1697
Simulium verecundum complex|[10852]|MHFLY195-07|BOLD:AAA1697
Simulium verecundum complex|[10853]|MHFLY170-07
Simulium verecundum complex|[10854]|MHFLY186-07
Simulium verecundum complex|[10855]|MHFLY321-07|BOLD:AAA1697
Simulium verecundum complex|[10856]|MHFLY196-07|BOLD:AAA1697
Simulium verecundum complex|[10857]|MHFLY248-07|BOLD:AAA1697
Simulium verecundum complex|[10858]|MHFLY218-07|BOLD:AAA1697
Simulium verecundum complex|[10859]|JWDCI723-11|BOLD:AAA1697
Simulium|[10860]|JWDCA132-10|BOLD:AAA1697
Bicellaria bisetosa|[10861]|JWDOL526-10|BOLD:AAF9807
Bicellaria bisetosa|[10862]|JWDCA070-10|BOLD:AAF9807
Bicellaria bisetosa|[10863]|JWDCA071-10|BOLD:AAF9807
Bicellaria bisetosa|[10864]|JWDOL535-10|BOLD:AAF9807
Bicellaria bisetosa|[10865]|JWEMP230-10|BOLD:AAF9807
Bicellaria bisetosa|[10866]|JWDOL735-10|BOLD:AAF9807
Bicellaria bisetosa|[10867]|JWEMP126-10|BOLD:AAF9807
Bicellaria bisetosa|[10868]|JWDOL548-10|BOLD:AAF9807
Bicellaria bisetosa|[10869]|JWDOL549-10|BOLD:AAF9807
Bicellaria bisetosa|[10870]|JWDOL538-10|BOLD:AAF9807
Bicellaria bisetosa|[10871]|JWDOL539-10|BOLD:AAF9807
Bicellaria bisetosa|[10872]|JWDOL524-10|BOLD:AAF9807
Bicellaria bisetosa|[10873]|JWDOL525-10|BOLD:AAF9807
Bicellaria bisetosa|[10874]|JWDOL522-10|BOLD:AAF9807
Bicellaria bisetosa|[10875]|JWDOL523-10|BOLD:AAF9807
Bicellaria bisetosa|[10876]|JWDOL695-10|BOLD:AAF9807
Bicellaria bisetosa|[10877]|JWDOL782-10|BOLD:AAF9807
Bicellaria bisetosa|[10878]|JWDOL785-10|BOLD:AAF9807
Bicellaria bisetosa|[10879]|JWDOL788-10|BOLD:AAF9807
Bicellaria bisetosa|[10880]|JWDOL506-10|BOLD:AAF9807
Bicellaria bisetosa|[10881]|JWDOL508-10|BOLD:AAF9807
Bicellaria bisetosa|[10882]|JWDOL575-10|BOLD:AAF9807
Bicellaria bisetosa|[10883]|JWDOL693-10|BOLD:AAF9807
Bicellaria bisetosa|[10884]|JWDOL504-10|BOLD:AAF9807
Bicellaria bisetosa|[10885]|JWDOL505-10|BOLD:AAF9807
Bicellaria bisetosa|[10886]|JWDOL573-10|BOLD:AAF9807
Bicellaria bisetosa|[10887]|JWDOL574-10|BOLD:AAF9807
Bicellaria bisetosa|[10888]|JWDOL552-10|BOLD:AAF9807
Bicellaria bisetosa|[10889]|JWDOL571-10|BOLD:AAF9807
Bicellaria bisetosa|[10890]|JWEMP256-10|BOLD:AAF9807
Bicellaria bisetosa|[10891]|JWEMP286-10|BOLD:AAF9807
Bicellaria bisetosa|[10892]|JWDOL490-10|BOLD:AAF9807
Bicellaria bisetosa|[10893]|JWDOL491-10|BOLD:AAF9807
Bicellaria bisetosa|[10894]|JWDOL536-10|BOLD:AAF9807
Bicellaria bisetosa|[10895]|JWDOL537-10|BOLD:AAF9807
Bicellaria bisetosa|[10896]|JWEMP210-10|BOLD:AAF9807
Bicellaria bisetosa|[10897]|JWEMP228-10|BOLD:AAF9807
Bicellaria bisetosa|[10898]|JWEMP229-10|BOLD:AAF9807
Bicellaria bisetosa|[10899]|JWEMP243-10|BOLD:AAF9807
Bicellaria bisetosa|[10900]|JWEMP200-10|BOLD:AAF9807
Bicellaria bisetosa|[10901]|JWEMP206-10|BOLD:AAF9807
Bicellaria bisetosa|[10902]|JWEMP208-10|BOLD:AAF9807
Bicellaria bisetosa|[10903]|JWEMP209-10|BOLD:AAF9807
Bicellaria bisetosa|[10904]|JWEMP193-10|BOLD:AAF9807
Bicellaria bisetosa|[10905]|JWEMP197-10|BOLD:AAF9807
Bicellaria bisetosa|[10906]|JWEMP131-10|BOLD:AAF9807
Bicellaria bisetosa|[10907]|JWEMP132-10|BOLD:AAF9807
Bicellaria bisetosa|[10908]|JWEMP133-10|BOLD:AAF9807
Bicellaria bisetosa|[10909]|JWEMP158-10|BOLD:AAF9807
Bicellaria bisetosa|[10910]|JWEMP191-10|BOLD:AAF9807
Bicellaria bisetosa|[10911]|JWEMP192-10|BOLD:AAF9807
Bicellaria bisetosa|[10912]|JWEMP127-10|BOLD:AAF9807
Bicellaria bisetosa|[10913]|JWEMP128-10|BOLD:AAF9807
Bicellaria bisetosa|[10914]|JWEMP129-10|BOLD:AAF9807
Bicellaria bisetosa|[10915]|JWEMP130-10|BOLD:AAF9807
Bicellaria bisetosa|[10916]|JWEMP081-10|BOLD:AAF9807
Bicellaria bisetosa|[10917]|JWEMP088-10|BOLD:AAF9807
Bicellaria bisetosa|[10918]|JWDOL607-10|BOLD:AAF9807
Bicellaria bisetosa|[10919]|JWDOL611-10|BOLD:AAF9807
Bicellaria bisetosa|[10920]|JWDOL614-10|BOLD:AAF9807
Bicellaria bisetosa|[10921]|JWDOL736-10|BOLD:AAF9807
Bicellaria bisetosa|[10922]|JWDOL744-10|BOLD:AAF9807
Bicellaria bisetosa|[10923]|JWDOL753-10|BOLD:AAF9807
Bicellaria bisetosa|[10924]|JWDOL765-10|BOLD:AAF9807
Bicellaria bisetosa|[10925]|JWDOL778-10|BOLD:AAF9807
Bicellaria bisetosa|[10926]|JWEMP060-10|BOLD:AAF9807
Bicellaria bisetosa|[10927]|JWEMP063-10|BOLD:AAF9807
Bicellaria bisetosa|[10928]|JWDOL841-10|BOLD:AAF9807
Bicellaria bisetosa|[10929]|JWDOL584-10|BOLD:AAF9807
Bicellaria bisetosa|[10930]|JWDOL585-10|BOLD:AAF9807
Bicellaria bisetosa|[10931]|JWDOL586-10|BOLD:AAF9807
Bicellaria bisetosa|[10932]|JWDOL587-10|BOLD:AAF9807
Bicellaria bisetosa|[10933]|JWDOL591-10|BOLD:AAF9807
Bicellaria bisetosa|[10934]|JWDOL601-10|BOLD:AAF9807
Bicellaria bisetosa|[10935]|JWDOL605-10|BOLD:AAF9807
Bicellaria bisetosa|[10936]|JWDOL779-10|BOLD:AAF9807
Bicellaria bisetosa|[10937]|JWEMP048-10|BOLD:AAF9807
Bicellaria bisetosa|[10938]|JWEMP097-10|BOLD:AAF9807
Bicellaria bisetosa|[10939]|JWEMP119-10|BOLD:AAF9807
Bicellaria bisetosa|[10940]|JWDOL837-10|BOLD:AAF9807
Bicellaria bisetosa|[10941]|JWDOL838-10|BOLD:AAF9807
Bicellaria bisetosa|[10942]|JWDCI757-11|BOLD:AAF9807
Bicellaria bisetosa|[10943]|JWDOL835-10|BOLD:AAF9807
Bicellaria bisetosa|[10944]|JWDCI719-11|BOLD:AAF9807
Bicellaria bisetosa|[10945]|JWDCI718-11|BOLD:AAF9807
Bicellaria bisetosa|[10946]|JWDCL566-11|BOLD:AAF9807

Bicellaria bisetosa|[10944]|JWDCI719-11|BOLD:AAF9807
Bicellaria bisetosa|[10945]|JWDCI718-11|BOLD:AAF9807
Bicellaria bisetosa|[10946]|JWDCL566-11|BOLD:AAF9807
Bicellaria bisetosa|[10947]|JWDCL564-11|BOLD:AAF9807
Bicellaria bisetosa|[10948]|JWDCL538-11|BOLD:AAF9807
Bicellaria bisetosa|[10949]|JWDCD161-10|BOLD:AAF9807
Bicellaria bisetosa|[10950]|JWDCD129-10|BOLD:AAF9807
Bicellaria bisetosa|[10951]|JWDCA089-10|BOLD:AAF9807
Bicellaria bisetosa|[10952]|JWDCA065-10|BOLD:AAF9807
Bicellaria bisetosa|[10953]|TWDIP274-09|BOLD:AAF9807
Bicellaria bisetosa|[10954]|MTDIC244-09|BOLD:AAF9807
Bicellaria bisetosa|[10955]|JWDOL608-10|BOLD:AAF9807
Bicellaria bisetosa|[10956]|JWDCA064-10|BOLD:AAF9807
Bicellaria bisetosa|[10957]|JWEMP231-10|BOLD:AAF9807
Bicellaria bisetosa|[10958]|JWDOL637-10|BOLD:AAF9807
Bicellaria bisetosa|[10959]|JWDOL798-10|BOLD:AAF9807
Bicellaria bisetosa|[10960]|JWDOL646-10|BOLD:AAF9807
Bicellaria bisetosa|[10961]|JWDOL711-10|BOLD:AAF9807
Bicellaria bisetosa|[10962]|JWDOL709-10|BOLD:AAF9807
Bicellaria bisetosa|[10963]|JWDOL652-10|BOLD:AAF9807
Bicellaria bisetosa|[10964]|JWDOL640-10|BOLD:AAF9807
Bicellaria bisetosa|[10965]|JWDOL572-10|BOLD:AAF9807
Bicellaria bisetosa|[10966]|JWDOL507-10|BOLD:AAF9807
Bicellaria bisetosa|[10967]|JWDOL499-10|BOLD:AAF9807
Bicellaria bisetosa|[10968]|JWDOL492-10|BOLD:AAF9807
Bicellaria bisetosa|[10969]|JWDOL781-10|BOLD:AAF9807
Bicellaria bisetosa|[10970]|JWDOL794-10|BOLD:AAF9807
Bicellaria bisetosa|[10971]|JWEMP253-10|BOLD:AAF9807
Bicellaria bisetosa|[10972]|JWEMP195-10|BOLD:AAF9807
Bicellaria bisetosa|[10973]|JWEMP194-10|BOLD:AAF9807
Bicellaria bisetosa|[10974]|JWEMP171-10|BOLD:AAF9807
Bicellaria bisetosa|[10975]|JWEMP169-10|BOLD:AAF9807
Bicellaria bisetosa|[10976]|JWEMP163-10|BOLD:AAF9807
Bicellaria bisetosa|[10977]|JWEMP161-10|BOLD:AAF9807
Bicellaria bisetosa|[10978]|JWEMP160-10|BOLD:AAF9807
Bicellaria bisetosa|[10979]|JWEMP156-10|BOLD:AAF9807
Bicellaria bisetosa|[10980]|JWEMP100-10|BOLD:AAF9807
Bicellaria bisetosa|[10981]|JWEMP096-10|BOLD:AAF9807
Bicellaria bisetosa|[10982]|JWEMP087-10|BOLD:AAF9807
Bicellaria bisetosa|[10983]|JWEMP086-10|BOLD:AAF9807
Bicellaria bisetosa|[10984]|JWEMP085-10|BOLD:AAF9807
Bicellaria bisetosa|[10985]|JWEMP064-10|BOLD:AAF9807
Bicellaria bisetosa|[10986]|JWEMP061-10|BOLD:AAF9807
Bicellaria bisetosa|[10987]|JWDOL770-10|BOLD:AAF9807
Bicellaria bisetosa|[10988]|JWDOL755-10|BOLD:AAF9807
Bicellaria bisetosa|[10989]|JWDOL733-10|BOLD:AAF9807
Bicellaria bisetosa|[10990]|JWDOL631-10|BOLD:AAF9807
Bicellaria bisetosa|[10991]|JWDOL622-10|BOLD:AAF9807
Bicellaria bisetosa|[10992]|JWDOL609-10|BOLD:AAF9807
Bicellaria bisetosa|[10993]|JWDOL606-10|BOLD:AAF9807
Bicellaria bisetosa|[10994]|JWDOL598-10|BOLD:AAF9807
Bicellaria bisetosa|[10995]|JWDOL594-10|BOLD:AAF9807
Bicellaria bisetosa|[10996]|JWDOL593-10|BOLD:AAF9807
Bicellaria bisetosa|[10997]|JWDOL583-10|BOLD:AAF9807
Bicellaria bisetosa|[10998]|JWDCH269-10|BOLD:AAF9807
Bicellaria bisetosa|[10999]|JWDCI756-11|BOLD:AAF9807
Bicellaria bisetosa|[11000]|JWDCA069-10|BOLD:AAF9807
Bicellaria bisetosa|[11001]|JWDCA068-10|BOLD:AAF9807
Bicellaria bisetosa|[11002]|JWDCA066-10|BOLD:AAF9807
Bicellaria bisetosa|[11003]|MTDIC182-09|BOLD:AAF9807
Bicellaria bisetosa|[11004]|JWDOL644-10|BOLD:AAF9807
Bicellaria bisetosa|[11005]|JWEMP246-10|BOLD:AAF9807
Bicellaria bisetosa|[11006]|JWDOL599-10|BOLD:AAF9807
Bicellaria bisetosa|[11007]|JWDOL786-10|BOLD:AAF9807
Bicellaria bisetosa|[11008]|JWDOL787-10|BOLD:AAF9807
Bicellaria bisetosa|[11009]|JWDOL793-10|BOLD:AAF9807
Bicellaria bisetosa|[11010]|JWDOL795-10|BOLD:AAF9807
Bicellaria bisetosa|[11011]|JWDOL796-10|BOLD:AAF9807
Bicellaria bisetosa|[11012]|JWDOL645-10|BOLD:AAF9807
Bicellaria bisetosa|[11013]|JWDOL647-10|BOLD:AAF9807
Bicellaria lugubris|[11014]|JWEMP164-10|BOLD:AAF9812
Bicellaria lugubris|[11015]|JWDCB324-10|BOLD:AAF9812
Bicellaria lugubris|[11016]|TWDIP275-09|BOLD:AAF9812
Bicellaria lugubris|[11017]|JWDOL495-10|BOLD:AAF9812
Bicellaria lugubris|[11018]|JWDCC729-10|BOLD:AAF9812
Bicellaria lugubris|[11019]|JWDCB338-10|BOLD:AAF9812
Bicellaria lugubris|[11020]|JWDCB346-10|BOLD:AAF9812
Bicellaria lugubris|[11021]|JWDCA103-10|BOLD:AAF9812
Bicellaria lugubris|[11022]|JWDCA063-10|BOLD:AAF9812
Bicellaria lugubris|[11023]|JWDCA067-10|BOLD:AAF9812
Bicellaria lugubris|[11024]|JWDCA092-10|BOLD:AAF9812
Bicellaria lugubris|[11025]|JWDCB342-10|BOLD:AAF9812
Bicellaria lugubris|[11026]|JWDCB343-10|BOLD:AAF9812
Bicellaria lugubris|[11027]|JWDCD107-10|BOLD:AAF9812
Bicellaria lugubris|[11028]|JWEMP084-10|BOLD:AAF9812
Bicellaria lugubris|[11029]|JWEMP157-10|BOLD:AAF9812
Bicellaria lugubris|[11030]|JWEMP165-10|BOLD:AAF9812
Bicellaria lugubris|[11031]|JWEMP321-10|BOLD:AAF9812
Bicellaria lugubris|[11032]|JWDOL544-10|BOLD:AAF9812
Bicellaria lugubris|[11033]|JWDOL576-10|BOLD:AAF9812
Bicellaria furcifer|[11034]|JWDOL859-10|BOLD:AAF9752
Bicellaria furcifer|[11035]|JWDCB326-10|BOLD:AAF9752
Bicellaria furcifer|[11036]|JWDOL656-10|BOLD:AAF9752
Bicellaria furcifer|[11037]|JWDCA091-10|BOLD:AAF9752
Bicellaria furcifer|[11038]|JWDOL754-10|BOLD:AAF9752
Bicellaria furcifer|[11039]|MHMOS098-07|BOLD:AAF9752
Bicellaria furcifer|[11040]|JWDCB347-10|BOLD:AAF9752
Bicellaria furcifer|[11041]|TWDIP273-09|BOLD:AAF9752
Bicellaria furcifer|[11042]|JWDOL832-10|BOLD:AAF9752
Bicellaria furcifer|[11043]|JWDOL858-10|BOLD:AAF9752
Bicellaria furcifer|[11044]|JWDOL634-10|BOLD:AAF9752
Bicellaria furcifer|[11045]|JWDOL714-10|BOLD:AAF9752
[illegible]

Bicellaria furcifer|[11043]|JWDOL838-10|BOLD:AAF9752
Bicellaria furcifer|[11044]|JWDOL634-10|BOLD:AAF9752
Bicellaria furcifer|[11045]|JWDOL714-10|BOLD:AAF9752
Bicellaria furcifer|[11046]|JWDOL722-10|BOLD:AAF9752
Bicellaria furcifer|[11047]|JWDOL726-10|BOLD:AAF9752
Bicellaria furcifer|[11048]|JWDOL737-10|BOLD:AAF9752
Bicellaria furcifer|[11049]|JWDOL760-10|BOLD:AAF9752
Bicellaria furcifer|[11050]|JWDOL767-10|BOLD:AAF9752
Bicellaria furcifer|[11051]|JWEMP252-10|BOLD:AAF9752
Bicellaria furcifer|[11052]|JWEMP267-10|BOLD:AAF9752
Bicellaria furcifer|[11053]|JWEMP281-10|BOLD:AAF9752
Bicellaria furcifer|[11054]|JWEMP288-10|BOLD:AAF9752
Bicellaria furcifer|[11055]|JWDOL550-10|BOLD:AAF9752
Bicellaria furcifer|[11056]|JWDOL665-10|BOLD:AAF9752
Bicellaria furcifer|[11057]|JWDOL780-10|BOLD:AAF9752
Bicellaria furcifer|[11058]|JWDOL789-10|BOLD:AAF9752
Bicellaria furcifer|[11059]|JWDOL662-10|BOLD:AAF9752
Bicellaria furcifer|[11060]|JWDOL663-10|BOLD:AAF9752
Platypalpus juvenis complex sp. JMC3|[11061]|TWDIP006-09|BOLD:AAF9793
Platypalpus xanthopodus|[11062]|JWDCA075-10|BOLD:AAM6647
Platypalpus xanthopodus|[11063]|JWDCA078-10|BOLD:AAM6647
Platypalpus xanthopodus|[11064]|JWDOL857-10|BOLD:AAM6647
Platypalpus xanthopodus|[11065]|JWDOL615-10|BOLD:AAM6647
Platypalpus xanthopodus|[11066]|JWDOL738-10|BOLD:AAM6647
Platypalpus xanthopodus|[11067]|JWDOL747-10|BOLD:AAM6647
Platypalpus xanthopodus|[11068]|JWEMP034-10|BOLD:AAM6647
Platypalpus xanthopodus|[11069]|JWEMP335-10|BOLD:AAM6647
Platypalpus xanthopodus|[11070]|JWDOL649-10|BOLD:AAM6647
Platypalpus nr. sahlbergi|[11071]|JWDCA074-10|BOLD:AAM6646
Platypalpus nr. sahlbergi|[11072]|JWEMP275-10|BOLD:AAM6646
Platypalpus nr. sahlbergi|[11073]|JWDOL731-10|BOLD:AAM6646
Platypalpus nr. sahlbergi|[11074]|JWDOL729-10|BOLD:AAM6646
Platypalpus nr. sahlbergi|[11075]|JWDOL853-10|BOLD:AAM6646
Platypalpus nr. sahlbergi|[11076]|JWDOL852-10|BOLD:AAM6646
Platypalpus nr. sahlbergi|[11077]|JWEMP279-10|BOLD:AAM6646
Platypalpus sp. JMC3-4|[11078]|JWDOL766-10|BOLD:AAF9801
Platypalpus sp. JMC3-4|[11079]|JWDCB345-10|BOLD:AAF9801
Platypalpus sp. JMC3-4|[11080]|JWDCB348-10|BOLD:AAF9801
Platypalpus sp. JMC3-4|[11081]|JWDOL727-10|BOLD:AAF9801
Platypalpus sp. JMC3-4|[11082]|MTDIC006-09|BOLD:AAF9801
Platypalpus sp. JMC3-4|[11083]|JWDCB334-10|BOLD:AAF9801
Platypalpus sp. JMC3-4|[11084]|MTDIC135-09|BOLD:AAF9801
Platypalpus sp. JMC3-4|[11085]|TWDIP314-09|BOLD:AAF9801
Platypalpus sp. JMC3-4|[11086]|MTDIC004-09|BOLD:AAF9801
Platypalpus sp. JMC3-4|[11087]|MTDIC026-09|BOLD:AAF9801
Platypalpus sp. JMC3-4|[11088]|JWDOL844-10|BOLD:AAF9801
Platypalpus sp. JMC3-4|[11089]|JWDOL846-10|BOLD:AAF9801
Platypalpus sp. JMC3-4|[11090]|JWDOL856-10|BOLD:AAF9801
Platypalpus sp. JMC3-4|[11091]|JWDOL588-10|BOLD:AAF9801
Platypalpus sp. JMC3-4|[11092]|JWDOL717-10|BOLD:AAF9801
Platypalpus sp. JMC3-4|[11093]|JWDOL718-10|BOLD:AAF9801
Platypalpus sp. JMC3-4|[11094]|JWEMP315-10|BOLD:AAF9801
Platypalpus sp. JMC3-4|[11095]|JWDOL639-10|BOLD:AAF9801
Platypalpus|[11096]|JWDCA096-10|BOLD:AAF9801
Iteaphila nitidula|[11097]|JWDCB328-10|BOLD:AAF9884
Iteaphila nitidula|[11098]|JWEMP235-10|BOLD:AAF9884
Iteaphila macquarti|[11099]|JWEMP300-10|BOLD:AAL8966
Iteaphila macquarti|[11100]|JWEMP296-10|BOLD:AAL8966
Iteaphila macquarti|[11101]|JWEMP299-10|BOLD:AAL8966
Iteaphila macquarti|[11102]|JWEMP301-10|BOLD:AAL8966
Iteaphila macquarti|[11103]|JWDCE560-10|BOLD:AAL8966
Iteaphila macquarti|[11104]|JWEMP006-10|BOLD:AAL8966
Iteaphila macquarti|[11105]|JWDCC833-10|BOLD:AAL8966
Iteaphila macquarti|[11106]|JWDCE436-10|BOLD:AAL8966
Iteaphila macquarti|[11107]|JWEMP318-10|BOLD:AAL8966
Rhamphomyia polita group sp. BJS16|[11108]|JWDOL629-10|BOLD:AAM7328
Rhamphomyia polita group sp. BJS16|[11109]|JWEMP245-10|BOLD:AAM7328
Rhamphomyia latilobata|[11110]|MTDIC054-09|BOLD:AAF9802
Rhamphomyia latilobata|[11111]|JWDCB332-10|BOLD:AAF9802
Rhamphomyia latilobata|[11112]|JWDCC848-10|BOLD:AAF9802
Rhamphomyia latilobata|[11113]|JWDCC936-10|BOLD:AAF9802
Rhamphomyia latilobata|[11114]|JWDCC929-10|BOLD:AAF9802
Rhamphomyia latilobata|[11115]|JWDCC875-10|BOLD:AAF9802
Rhamphomyia latilobata|[11116]|JWDCC824-10|BOLD:AAF9802
Rhamphomyia latilobata|[11117]|JWDCA055-10|BOLD:AAF9802
Rhamphomyia latilobata|[11118]|JWDCA051-10|BOLD:AAF9802
Rhamphomyia latilobata|[11119]|JWDCB323-10|BOLD:AAF9802
Rhamphomyia latilobata|[11120]|MTDIC007-09|BOLD:AAF9802
Rhamphomyia latilobata|[11121]|JWDCA047-10|BOLD:AAF9802
Rhamphomyia latilobata|[11122]|JWDCC822-10|BOLD:AAF9802
Rhamphomyia latilobata|[11123]|JWDCC927-10|BOLD:AAF9802
Rhamphomyia latilobata|[11124]|JWDCA048-10|BOLD:AAF9802
Rhamphomyia latilobata|[11125]|JWDCA105-10|BOLD:AAF9802
Rhamphomyia latilobata|[11126]|JWDCC825-10|BOLD:AAF9802
Rhamphomyia latilobata|[11127]|JWDCD076-10|BOLD:AAF9802
Rhamphomyia latilobata|[11128]|JWEMP304-10|BOLD:AAF9802
Rhamphomyia latilobata|[11129]|JWEMP035-10|BOLD:AAF9802
Rhamphomyia latilobata|[11130]|JWDOL528-10|BOLD:AAF9802
Rhamphomyia pendens|[11131]|JWDCD160-10|BOLD:AAF9811
Rhamphomyia pendens|[11132]|JWEMP180-10|BOLD:AAF9811
Rhamphomyia pendens|[11133]|JWDCA101-10|BOLD:AAF9811
Rhamphomyia pendens|[11134]|TWDIP217-09|BOLD:AAF9811
Rhamphomyia pendens|[11135]|JWDOL839-10|BOLD:AAF9811
Rhamphomyia pendens|[11136]|JWEMP026-10|BOLD:AAF9811
Rhamphomyia pendens|[11137]|JWEMP067-10|BOLD:AAF9811
Rhamphomyia pendens|[11138]|JWEMP145-10|BOLD:AAF9811
Rhamphomyia pendens|[11139]|JWDCA086-10|BOLD:AAF9811
Rhamphomyia pendens|[11140]|JWDOL534-10|BOLD:AAF9811
Rhamphomyia pendens|[11141]|JWEMP143-10|BOLD:AAF9811
Rhamphomyia pendens|[11142]|JWEMP070-10|BOLD:AAF9811
Rhamphomyia pendens|[11143]|JWDOL679-10|BOLD:AAF9811
Rhamphomyia pendens|[11144]|JWEMP021-10|BOLD:AAF9811
Rhamphomyia pendens|[11145]|JWEMP153-10|BOLD:AAF9811

Rhamphomyia pendens|[11143]|JWDOL679-10|BOLD:AAF9811
Rhamphomyia pendens|[11144]|JWEMP021-10|BOLD:AAF9811
Rhamphomyia pendens|[11145]|JWEMP153-10|BOLD:AAF9811
Rhamphomyia pendens|[11146]|JWDCC821-10|BOLD:AAF9811
Rhamphomyia pendens|[11147]|JWDCA106-10|BOLD:AAF9811
Rhamphomyia pendens|[11148]|JWDOL696-10|BOLD:AAF9811
Rhamphomyia pendens|[11149]|TWDIP380-09|BOLD:AAF9811
Rhamphomyia pendens|[11150]|JWEMP074-10|BOLD:AAF9811
Rhamphomyia pendens|[11151]|JWDCC863-10|BOLD:AAF9811
Rhamphomyia pendens|[11152]|JWDOL517-10|BOLD:AAF9811
Rhamphomyia pendens|[11153]|JWDOL516-10|BOLD:AAF9811
Rhamphomyia pendens|[11154]|JWDOL493-10|BOLD:AAF9811
Rhamphomyia pendens|[11155]|JWDOL482-10|BOLD:AAF9811
Rhamphomyia pendens|[11156]|JWEMP146-10|BOLD:AAF9811
Rhamphomyia pendens|[11157]|JWEMP144-10|BOLD:AAF9811
Rhamphomyia pendens|[11158]|JWEMP076-10|BOLD:AAF9811
Rhamphomyia pendens|[11159]|JWEMP075-10|BOLD:AAF9811
Rhamphomyia pendens|[11160]|JWEMP073-10|BOLD:AAF9811
Rhamphomyia pendens|[11161]|JWEMP071-10|BOLD:AAF9811
Rhamphomyia pendens|[11162]|JWDOL602-10|BOLD:AAF9811
Rhamphomyia pendens|[11163]|JWEMP031-10|BOLD:AAF9811
Rhamphomyia pendens|[11164]|JWEMP068-10|BOLD:AAF9811
Rhamphomyia pendens|[11165]|JWEMP050-10|BOLD:AAF9811
Rhamphomyia pendens|[11166]|JWEMP032-10|BOLD:AAF9811
Rhamphomyia pendens|[11167]|JWEMP023-10|BOLD:AAF9811
Rhamphomyia pendens|[11168]|JWDOL618-10|BOLD:AAF9811
Rhamphomyia pendens|[11169]|JWDCC870-10|BOLD:AAF9811
Rhamphomyia pendens|[11170]|TWDIP216-09|BOLD:AAF9811
Rhamphomyia pendens|[11171]|JWDCD038-10|BOLD:AAF9811
Rhamphomyia pendens|[11172]|JWDOL763-10|BOLD:AAF9811
Rhamphomyia pendens|[11173]|JWEMP033-10|BOLD:AAF9811
Rhamphomyia pendens|[11174]|JWEMP147-10|BOLD:AAF9811
Rhamphomyia pendens|[11175]|JWDOL603-10|BOLD:AAF9811
Rhamphomyia pendens|[11176]|JWEMP022-10|BOLD:AAF9811
Rhamphomyia pendens|[11177]|JWEMP024-10|BOLD:AAF9811
Rhamphomyia pendens|[11178]|JWEMP027-10|BOLD:AAF9811
Rhamphomyia pendens|[11179]|JWDOL689-10|BOLD:AAF9811
Rhamphomyia pendens|[11180]|JWDOL518-10|BOLD:AAF9811
Rhamphomyia pendens|[11181]|JWDOL690-10|BOLD:AAF9811
Rhamphomyia pendens|[11182]|JWDOL692-10|BOLD:AAF9811
Rhamphomyia pendens|[11183]|JWEMP176-10|BOLD:AAF9811
Rhamphomyia pendens|[11184]|JWEMP072-10|BOLD:AAF9811
Rhamphomyia pendens|[11185]|JWDOL694-10|BOLD:AAF9811
Rhamphomyia pendens|[11186]|JWDOL619-10|BOLD:AAF9811
Rhamphomyia pendens|[11187]|JWDOL485-10|BOLD:AAF9811
Rhamphomyia pendens|[11188]|JWDCC865-10|BOLD:AAF9811
Rhamphomyia pendens|[11189]|JWDCC867-10|BOLD:AAF9811
Rhamphomyia pendens|[11190]|JWDOL617-10|BOLD:AAF9811
Rhamphomyia pendens|[11191]|JWEMP025-10|BOLD:AAF9811
Rhamphomyia pendens|[11192]|JWEMP028-10|BOLD:AAF9811
Rhamphomyia pendens|[11193]|JWEMP030-10|BOLD:AAF9811
Rhamphomyia pendens|[11194]|JWEMP155-10|BOLD:AAF9811
Rhamphomyia pendens|[11195]|JWDOL519-10|BOLD:AAF9811
Rhamphomyia pendens|[11196]|JWDOL688-10|BOLD:AAF9811
Rhamphomyia pendens|[11197]|JWDOL680-10|BOLD:AAF9811
Rhamphomyia pendens|[11198]|JWDOL691-10|BOLD:AAF9811
Rhamphomyia setosa|[11199]|JWEMP058-10|BOLD:AAP1851
Rhamphomyia setosa|[11200]|JWEMP055-10|BOLD:AAP1851
Rhamphomyia setosa|[11201]|JWEMP056-10|BOLD:AAP1851
Rhamphomyia setosa|[11202]|JWEMP057-10|BOLD:AAP1851
Rhamphomyia setosa|[11203]|JWEMP059-10|BOLD:AAP1851
Rhamphomyia setosa|[11204]|JWEMP062-10|BOLD:AAP1851
Rhamphomyia clavigera grp. sp. BJS1A|[11205]|JWEMP039-10|BOLD:AAP1849
Rhamphomyia clavigera grp. sp. BJS1A|[11206]|JWEMP302-10|BOLD:AAP1849
Rhamphomyia dispar|[11207]|JWDCE541-10|BOLD:AAP1847
Rhamphomyia dispar|[11208]|JWEMP017-10|BOLD:AAP1847
Rhamphomyia clavigera grp. sp. BJS1|[11209]|JWDCC944-10|BOLD:AAL7821
Rhamphomyia clavigera grp. sp. BJS1|[11210]|JWDCC817-10|BOLD:AAL7821
Rhamphomyia clavigera grp. sp. BJS1|[11211]|JWDCC934-10|BOLD:AAL7821
Rhamphomyia clavigera grp. sp. BJS1|[11212]|JWDCD162-10|BOLD:AAL7821
Rhamphomyia clavigera grp. sp. BJS1|[11213]|JWDCE542-10|BOLD:AAL7821
Rhamphomyia clavigera grp. sp. BJS1|[11214]|JWDCC939-10|BOLD:AAL7821
Rhamphomyia clavigera grp. sp. BJS1|[11215]|JWDCC943-10|BOLD:AAL7821
Rhamphomyia clavigera grp. sp. BJS1|[11216]|JWDCC935-10|BOLD:AAL7821
Rhamphomyia clavigera grp. sp. BJS1|[11217]|JWDCC937-10|BOLD:AAL7821
Rhamphomyia clavigera grp. sp. BJS1|[11218]|JWDCC931-10|BOLD:AAL7821
Rhamphomyia clavigera grp. sp. BJS1|[11219]|JWDCC932-10|BOLD:AAL7821
Rhamphomyia clavigera grp. sp. BJS1|[11220]|JWDCC930-10|BOLD:AAL7821
Rhamphomyia clavigera grp. sp. BJS1|[11221]|JWDCC923-10|BOLD:AAL7821
Rhamphomyia clavigera grp. sp. BJS1|[11222]|JWDCC921-10|BOLD:AAL7821
Rhamphomyia clavigera grp. sp. BJS1|[11223]|JWDCC920-10|BOLD:AAL7821
Rhamphomyia clavigera grp. sp. BJS1|[11224]|JWDCC864-10|BOLD:AAL7821
Rhamphomyia clavigera grp. sp. BJS1|[11225]|JWDCC830-10|BOLD:AAL7821
Rhamphomyia clavigera grp. sp. BJS1|[11226]|JWDCC828-10|BOLD:AAL7821
Rhamphomyia clavigera grp. sp. BJS1|[11227]|JWDCC850-10|BOLD:AAL7821
Rhamphomyia clavigera grp. sp. BJS1|[11228]|JWDCC946-10|BOLD:AAL7821
Rhamphomyia clavigera grp. sp. BJS1|[11229]|JWDCC831-10|BOLD:AAL7821
Rhamphomyia clavigera grp. sp. BJS1|[11230]|JWDCC926-10|BOLD:AAL7821
Rhamphomyia clavigera grp. sp. BJS1|[11231]|JWEMP018-10|BOLD:AAL7821
Rhamphomyia clavigera grp. sp. BJS1|[11232]|JWDCE663-10|BOLD:AAL7821
Rhamphomyia clavigera grp. sp. BJS1|[11233]|JWEMP020-10|BOLD:AAL7821
Rhamphomyia clavigera grp. sp. BJS1|[11234]|JWDCC942-10|BOLD:AAL7821
Rhamphomyia clavigera grp. sp. BJS1|[11235]|JWDCC947-10|BOLD:AAL7821
Rhamphomyia clavigera grp. sp. BJS1|[11236]|JWDCC948-10|BOLD:AAL7821
Rhamphomyia clavigera grp. sp. BJS1|[11237]|JWDCC949-10|BOLD:AAL7821
Rhamphomyia clavigera grp. sp. BJS1|[11238]|JWDCC940-10|BOLD:AAL7821
Rhamphomyia clavigera grp. sp. BJS1|[11239]|JWDCC941-10|BOLD:AAL7821
Rhamphomyia clavigera grp. sp. BJS1|[11240]|JWDCC933-10|BOLD:AAL7821
Rhamphomyia clavigera grp. sp. BJS1|[11241]|JWDCC928-10|BOLD:AAL7821
Rhamphomyia clavigera grp. sp. BJS1|[11242]|JWDCC924-10|BOLD:AAL7821
Rhamphomyia clavigera grp. sp. BJS1|[11243]|JWDCC922-10|BOLD:AAL7821
Rhamphomyia clavigera grp. sp. BJS1|[11244]|JWDCC871-10|BOLD:AAL7821
[illegible]

Rhamphomyia clavigera grp. sp. BJS1|[11242]|JWDCC924-10|BOLD:AAL7821
Rhamphomyia clavigera grp. sp. BJS1|[11243]|JWDCC922-10|BOLD:AAL7821
Rhamphomyia clavigera grp. sp. BJS1|[11244]|JWDCC871-10|BOLD:AAL7821
Rhamphomyia clavigera grp. sp. BJS1|[11245]|JWDCC862-10|BOLD:AAL7821
Rhamphomyia clavigera grp. sp. BJS1|[11246]|JWDCC861-10|BOLD:AAL7821
Rhamphomyia clavigera grp. sp. BJS1|[11247]|JWDCC851-10|BOLD:AAL7821
Rhamphomyia clavigera grp. sp. BJS1|[11248]|JWDCC849-10|BOLD:AAL7821
Rhamphomyia clavigera grp. sp. BJS1|[11249]|JWDCC819-10|BOLD:AAL7821
Rhamphomyia clavigera grp. sp. BJS1|[11250]|JWDCC816-10|BOLD:AAL7821
Rhamphomyia clavigera grp. sp. BJS1|[11251]|JWDCC829-10|BOLD:AAL7821
Rhamphomyia clavigera grp. sp. BJS1|[11252]|JWDCC827-10|BOLD:AAL7821
Rhamphomyia clavigera grp. sp. BJS1|[11253]|JWDCC938-10|BOLD:AAL7821
Rhamphomyia clavigera grp. sp. BJS1|[11254]|JWDCC925-10|BOLD:AAL7821
Rhamphomyia clavigera grp. sp. BJS1|[11255]|JWDCC945-10|BOLD:AAL7821
Rhamphomyia clavigera grp. sp. BJS1|[11256]|JWDCC950-10|BOLD:AAL7821
Rhamphomyia clavigera grp. sp. BJS1|[11257]|JWDCE358-10|BOLD:AAL7821
Rhamphomyia clavigera grp. sp. BJS1|[11258]|JWEMP019-10|BOLD:AAL7821
Rhamphomyia clavigera grp. sp. BJS1|[11259]|JWEMP297-10|BOLD:AAL7821
Rhamphomyia clavigera grp. sp. BJS1|[11260]|JWEMP298-10|BOLD:AAL7821
Rhamphomyia clavigera grp. sp. BJS1|[11261]|JWDCA100-10|BOLD:AAL7821
Rhamphomyia plumigera grp. sp. BJS27|[11262]|JWDCA062-10|BOLD:AAM6645
Rhamphomyia plumigera grp. sp. BJS29|[11263]|TWDIP019-09|BOLD:AAF9798
Empis lucida|[11264]|JWEMP102-10|BOLD:AAL8968
Empis lucida|[11265]|JWDCD047-10|BOLD:AAL8968
Empis lucida|[11266]|JWEMP110-10|BOLD:AAL8968
Empis lucida|[11267]|JWEMP211-10|BOLD:AAL8968
Empis lucida|[11268]|JWDOL589-10|BOLD:AAL8968
Empis lucida|[11269]|JWEMP001-10|BOLD:AAL8968
Empis lucida|[11270]|JWEMP045-10|BOLD:AAL8968
Empis lucida|[11271]|JWEMP046-10|BOLD:AAL8968
Empis lucida|[11272]|JWEMP095-10|BOLD:AAL8968
Empis lucida|[11273]|JWEMP101-10|BOLD:AAL8968
Empis lucida|[11274]|JWEMP109-10|BOLD:AAL8968
Empis lucida|[11275]|JWEMP112-10|BOLD:AAL8968
Empis lucida|[11276]|JWEMP113-10|BOLD:AAL8968
Empis lucida|[11277]|JWEMP185-10|BOLD:AAL8968
Empis lucida|[11278]|JWEMP186-10|BOLD:AAL8968
Empis lucida|[11279]|JWDOL531-10|BOLD:AAL8968
Empis lucida|[11280]|JWDOL553-10|BOLD:AAL8968
Empis lucida|[11281]|JWDOL642-10|BOLD:AAL8968
Empis lucida|[11282]|JWDOL643-10|BOLD:AAL8968
Empis lucida|[11283]|JWDOL668-10|BOLD:AAL8968
Rhamphomyia pusilla|[11284]|JWDCA056-10|BOLD:AAF9880
Rhamphomyia pusilla|[11285]|JWDCA107-10|BOLD:AAF9880
Rhamphomyia pusilla|[11286]|JWEMP137-10|BOLD:AAF9880
Rhamphomyia pusilla|[11287]|JWEMP120-10|BOLD:AAF9880
Rhamphomyia pusilla|[11288]|JWDCA049-10|BOLD:AAF9880
Rhamphomyia pusilla|[11289]|JWDCC866-10|BOLD:AAF9880
Rhamphomyia pusilla|[11290]|JWDCA102-10|BOLD:AAF9880
Rhamphomyia pusilla|[11291]|JWDCC823-10|BOLD:AAF9880
Rhamphomyia pusilla|[11292]|JWDCC869-10|BOLD:AAF9880
Rhamphomyia pusilla|[11293]|JWEMP172-10|BOLD:AAF9880
Rhamphomyia pusilla|[11294]|JWDCA115-10|BOLD:AAF9880
Rhamphomyia pusilla|[11295]|JWEMP167-10|BOLD:AAF9880
Rhamphomyia pusilla|[11296]|JWEMP173-10|BOLD:AAF9880
Rhamphomyia pusilla|[11297]|JWEMP136-10|BOLD:AAF9880
Rhamphomyia pusilla|[11298]|JWEMP135-10|BOLD:AAF9880
Rhamphomyia pusilla|[11299]|JWDCC826-10|BOLD:AAF9880
Rhamphomyia pusilla|[11300]|JWDOL487-10|BOLD:AAF9880
Rhamphomyia pusilla|[11301]|JWDOL488-10|BOLD:AAF9880
Rhamphomyia pusilla|[11302]|JWEMP227-10|BOLD:AAF9880
Rhamphomyia pusilla|[11303]|JWEMP168-10|BOLD:AAF9880
Rhamphomyia pusilla|[11304]|JWDOL489-10|BOLD:AAF9880
Rhamphomyia pusilla|[11305]|JWDOL486-10|BOLD:AAF9880
Rhamphomyia pusilla|[11306]|JWEMP166-10|BOLD:AAF9880
Rhamphomyia pusilla|[11307]|JWEMP121-10|BOLD:AAF9880
Rhamphomyia pusilla|[11308]|JWDCA118-10|BOLD:AAF9880
Rhamphomyia pusilla|[11309]|JWDCA050-10|BOLD:AAF9880
Rhamphomyia pusilla|[11310]|JWDCC873-10|BOLD:AAF9880
Rhamphomyia pusilla|[11311]|JWDCB339-10|BOLD:AAF9880
Rhamphomyia pusilla|[11312]|JWDCB329-10|BOLD:AAF9880
Rhamphomyia pusilla|[11313]|JWDCC490-10|BOLD:AAF9880
Rhamphomyia pusilla|[11314]|JWDOL551-10|BOLD:AAF9880
Rhamphomyia pusilla grp. sp. BJS2|[11315]|JWDCC650-10|BOLD:AAL8963
Rhamphomyia pusilla grp. sp. BJS2|[11316]|JWDCC675-10|BOLD:AAL8963
Rhamphomyia pusilla grp. sp. BJS2|[11317]|JWDCC676-10|BOLD:AAL8963
Rhamphomyia|[11318]|JWDCA104-10|BOLD:AAL8963
Nephrotoma altissima|[11319]|PHTCH337-08|BOLD:AAB4544
Nephrotoma altissima|[11320]|PHTCH344-08|BOLD:AAB4544
Nephrotoma altissima|[11321]|PHTCH345-08|BOLD:AAB4544
Nephrotoma altissima|[11322]|PHTCH346-08|BOLD:AAB4544
Nephrotoma altissima|[11323]|PHTCH347-08|BOLD:AAB4544
Nephrotoma altissima|[11324]|PHTCH348-08|BOLD:AAB4544
Nephrotoma lundbecki|[11325]|PHTCH016-07|BOLD:AAC8434
Nephrotoma lundbecki|[11326]|PHTCH134-08|BOLD:AAC8434
Nephrotoma lundbecki|[11327]|PHTCH082-07|BOLD:AAC8434
Nephrotoma lundbecki|[11328]|PHTCH008-07|BOLD:AAC8434
Nephrotoma lundbecki|[11329]|PHTCH026-07|BOLD:AAC8434
Nephrotoma lundbecki|[11330]|PHTCH027-07|BOLD:AAC8434
Nephrotoma lundbecki|[11331]|PHTCH037-07|BOLD:AAC8434
Nephrotoma lundbecki|[11332]|PHTCH051-07|BOLD:AAC8434
Nephrotoma lundbecki|[11333]|PHTCH068-07|BOLD:AAC8434
Nephrotoma penumbra|[11334]|PHTCH054-07|BOLD:ACE4443
Nephrotoma penumbra|[11335]|PHTCH042-07|BOLD:ACE4443
Nephrotoma penumbra|[11336]|PHTCH077-07|BOLD:ACE4443
Nephrotoma ferruginea|[11337]|JWDCF791-10|BOLD:ABX6186
Angarotipula illustris|[11338]|PHTCH265-08|BOLD:AAD6051
Angarotipula illustris|[11339]|PHTCH417-08|BOLD:AAD6051
Angarotipula illustris|[11340]|PHTCH236-08|BOLD:AAD6051
Angarotipula illustris|[11341]|PHTCH239-08|BOLD:AAD6051
Angarotipula illustris|[11342]|PHTCH270-08|BOLD:AAD6051
Angarotipula illustris|[11343]|PHTCH324-08|BOLD:AAD6051
Prionocera pubescens|[11344]|JWDCF463-10|BOLD:AAP3789

Angarotipula illustris|[11342]|PHTCH270-08|BOLD:AAD6051
Angarotipula illustris|[11343]|PHTCH324-08|BOLD:AAD6051
Prionocera pubescens|[11344]|JWDCF463-10|BOLD:AAP3789
Prionocera pubescens|[11345]|JWDCE527-10|BOLD:AAP3789
Prionocera pubescens|[11346]|JWDCF464-10|BOLD:AAP3789
Prionocera subserricornis|[11347]|PHTCH230-08|BOLD:AAA7256
Prionocera subserricornis|[11348]|PHTCH364-08|BOLD:AAA7256
Prionocera subserricornis|[11349]|PHTCH371-08|BOLD:AAA7256
Prionocera subserricornis|[11350]|PHTCH374-08|BOLD:AAA7256
Prionocera subserricornis|[11351]|JWDCG589-10|BOLD:AAA7256
Prionocera subserricornis|[11352]|JWDCF786-10|BOLD:AAA7256
Prionocera subserricornis|[11353]|PHTCH200-08|BOLD:AAA7256
Prionocera subserricornis|[11354]|PHTCH229-08|BOLD:AAA7256
Prionocera subserricornis|[11355]|JWDCJ773-11|BOLD:AAA7256
Prionocera subserricornis|[11356]|JWDCG010-10|BOLD:AAA7256
Prionocera subserricornis|[11357]|JWDCF418-10|BOLD:AAA7256
Prionocera subserricornis|[11358]|JWDCG588-10|BOLD:AAA7256
Prionocera subserricornis|[11359]|PHTCH381-08|BOLD:AAA7256
Prionocera subserricornis|[11360]|PHTCH380-08|BOLD:AAA7256
Prionocera subserricornis|[11361]|PHTCH378-08|BOLD:AAA7256
Prionocera subserricornis|[11362]|PHTCH376-08|BOLD:AAA7256
Prionocera subserricornis|[11363]|PHTCH217-08|BOLD:AAA7256
Prionocera subserricornis|[11364]|PHTCH213-08|BOLD:AAA7256
Prionocera subserricornis|[11365]|PHTCH206-08|BOLD:AAA7256
Prionocera subserricornis|[11366]|PHTCH198-08|BOLD:AAA7256
Prionocera subserricornis|[11367]|PHTCH190-08|BOLD:AAA7256
Prionocera subserricornis|[11368]|PHTCH189-08|BOLD:AAA7256
Prionocera subserricornis|[11369]|PHTCH372-08|BOLD:AAA7256
Prionocera subserricornis|[11370]|PHTCH370-08|BOLD:AAA7256
Prionocera subserricornis|[11371]|PHTCH363-08|BOLD:AAA7256
Prionocera subserricornis|[11372]|PHTCH351-08|BOLD:AAA7256
Prionocera subserricornis|[11373]|PHTCH272-08|BOLD:AAA7256
Prionocera subserricornis|[11374]|PHTCH263-08|BOLD:AAA7256
Prionocera subserricornis|[11375]|PHTCH250-08|BOLD:AAA7256
Prionocera subserricornis|[11376]|PHTCH247-08|BOLD:AAA7256
Prionocera subserricornis|[11377]|PHTCH244-08|BOLD:AAA7256
Prionocera subserricornis|[11378]|PHTCH242-08|BOLD:AAA7256
Prionocera subserricornis|[11379]|PHTCH240-08|BOLD:AAA7256
Prionocera subserricornis|[11380]|PHTCH235-08|BOLD:AAA7256
Prionocera subserricornis|[11381]|PHTCH232-08|BOLD:AAA7256
Prionocera subserricornis|[11382]|PHTCH226-08|BOLD:AAA7256
Prionocera subserricornis|[11383]|PHTCH225-08|BOLD:AAA7256
Prionocera subserricornis|[11384]|PHTCH401-08|BOLD:AAA7256
Prionocera subserricornis|[11385]|PHTCH402-08|BOLD:AAA7256
Prionocera subserricornis|[11386]|PHTCH373-08|BOLD:AAA7256
Prionocera subserricornis|[11387]|JWDCF422-10|BOLD:AAA7256
Prionocera subserricornis|[11388]|JWDCG011-10|BOLD:AAA7256
Prionocera subserricornis|[11389]|JWDCJ940-11|BOLD:AAA7256
Prionocera subserricornis|[11390]|JWDCF673-10|BOLD:AAA7256
Prionocera subserricornis|[11391]|JWDCF307-10|BOLD:AAA7256
Prionocera subserricornis|[11392]|JWDCI241-10|BOLD:AAA7256
Prionocera subserricornis|[11393]|JWDCF425-10|BOLD:AAA7256
Prionocera subserricornis|[11394]|PHTCH224-08|BOLD:AAA7256
Prionocera subserricornis|[11395]|PHTCH248-08|BOLD:AAA7256
Prionocera subserricornis|[11396]|PHTCH375-08|BOLD:AAA7256
Prionocera subserricornis|[11397]|PHTCH171-08|BOLD:AAA7256
Prionocera subserricornis|[11398]|PHTCH178-08|BOLD:AAA7256
Prionocera subserricornis|[11399]|PHTCH379-08|BOLD:AAA7256
Prionocera subserricornis|[11400]|TICHU105-09|BOLD:AAA7256
Prionocera subserricornis|[11401]|PHTCH205-08|BOLD:AAA7256
Prionocera subserricornis|[11402]|PHTCH233-08|BOLD:AAA7256
Prionocera subserricornis|[11403]|JWDCF674-10|BOLD:AAA7256
Prionocera subserricornis|[11404]|JWDCF465-10|BOLD:AAA7256
Prionocera subserricornis|[11405]|PHTCH249-08|BOLD:AAA7256
Prionocera subserricornis|[11406]|PHTCH246-08|BOLD:AAA7256
Prionocera subserricornis|[11407]|PHTCH245-08|BOLD:AAA7256
Prionocera subserricornis|[11408]|PHTCH241-08|BOLD:AAA7256
Prionocera subserricornis|[11409]|PHTCH231-08|BOLD:AAA7256
Prionocera subserricornis|[11410]|PHTCH228-08|BOLD:AAA7256
Prionocera subserricornis|[11411]|PHTCH227-08|BOLD:AAA7256
Prionocera subserricornis|[11412]|PHTCH403-08|BOLD:AAA7256
Prionocera subserricornis|[11413]|TICHU106-09|BOLD:AAA7256
Prionocera turcica|[11414]|PHTCH193-08|BOLD:ACE4050
Prionocera turcica|[11415]|JWDCI268-10|BOLD:ACE4050
Prionocera turcica|[11416]|PHTCH287-08|BOLD:ACE4050
Prionocera turcica|[11417]|PHTCH208-08|BOLD:ACE4050
Prionocera turcica|[11418]|PHTCH219-08|BOLD:ACE4050
Prionocera turcica|[11419]|JWDCG012-10|BOLD:ACE4050
Prionocera turcica|[11420]|TICHU114-09|BOLD:ACE4050
Prionocera turcica|[11421]|TICHU112-09|BOLD:ACE4050
Prionocera turcica|[11422]|TICHU110-09|BOLD:ACE4050
Prionocera turcica|[11423]|TICHU101-09|BOLD:ACE4050
Prionocera turcica|[11424]|TICHU097-09|BOLD:ACE4050
Prionocera turcica|[11425]|JWDCJ844-11|BOLD:ACE4050
Prionocera turcica|[11426]|JWDCE639-10|BOLD:ACE4050
Prionocera turcica|[11427]|JWDCG735-10|BOLD:ACE4050
Prionocera turcica|[11428]|JWDCF046-10|BOLD:ACE4050
Prionocera turcica|[11429]|PHTCH220-08|BOLD:ACE4050
Prionocera turcica|[11430]|PHTCH218-08|BOLD:ACE4050
Prionocera turcica|[11431]|PHTCH216-08|BOLD:ACE4050
Prionocera turcica|[11432]|PHTCH215-08|BOLD:ACE4050
Prionocera turcica|[11433]|PHTCH214-08|BOLD:ACE4050
Prionocera turcica|[11434]|PHTCH211-08|BOLD:ACE4050
Prionocera turcica|[11435]|PHTCH210-08|BOLD:ACE4050
Prionocera turcica|[11436]|PHTCH209-08|BOLD:ACE4050
Prionocera turcica|[11437]|PHTCH191-08|BOLD:ACE4050
Prionocera turcica|[11438]|PHTCH187-08|BOLD:ACE4050
Prionocera turcica|[11439]|PHTCH186-08|BOLD:ACE4050
Prionocera turcica|[11440]|PHTCH352-08|BOLD:ACE4050
Prionocera turcica|[11441]|PHTCH336-08|BOLD:ACE4050
Prionocera turcica|[11442]|PHTCH335-08|BOLD:ACE4050
Prionocera turcica|[11443]|PHTCH292-08|BOLD:ACE4050
[illegible]

Prionocera turcica|[11441]|PHTCH330-08|BOLD:ACE4050
Prionocera turcica|[11442]|PHTCH335-08|BOLD:ACE4050
Prionocera turcica|[11443]|PHTCH292-08|BOLD:ACE4050
Prionocera turcica|[11444]|PHTCH290-08|BOLD:ACE4050
Prionocera turcica|[11445]|PHTCH288-08|BOLD:ACE4050
Prionocera turcica|[11446]|PHTCH266-08|BOLD:ACE4050
Prionocera turcica|[11447]|PHTCH264-08|BOLD:ACE4050
Prionocera turcica|[11448]|PHTCH418-08|BOLD:ACE4050
Prionocera turcica|[11449]|PHTCH404-08|BOLD:ACE4050
Prionocera turcica|[11450]|PHTCH410-08|BOLD:ACE4050
Prionocera turcica|[11451]|PHTCH267-08|BOLD:ACE4050
Prionocera turcica|[11452]|PHTCH334-08|BOLD:ACE4050
Prionocera turcica|[11453]|PHTCH221-08|BOLD:ACE4050
Prionocera turcica|[11454]|PHTCH398-08|BOLD:ACE4050
Prionocera turcica|[11455]|PHTCH212-08|BOLD:ACE4050
Prionocera turcica|[11456]|PHTCH399-08|BOLD:ACE4050
Prionocera turcica|[11457]|TICHU116-09|BOLD:ACE4050
Prionocera dimidiata|[11458]|PHTCH366-08|BOLD:AAB5747
Prionocera dimidiata|[11459]|PHTCH243-08|BOLD:AAB5747
Prionocera dimidiata|[11460]|JWDCF129-10|BOLD:AAB5747
Prionocera dimidiata|[11461]|PHTCH222-08|BOLD:AAB5747
Prionocera dimidiata|[11462]|JWDCE361-10|BOLD:AAB5747
Prionocera dimidiata|[11463]|PHTCH238-08|BOLD:AAB5747
Prionocera dimidiata|[11464]|PHTCH223-08|BOLD:AAB5747
Prionocera dimidiata|[11465]|PHTCH273-08|BOLD:AAB5747
Prionocera dimidiata|[11466]|PHTCH350-08|BOLD:AAB5747
Prionocera dimidiata|[11467]|PHTCH360-08|BOLD:AAB5747
Prionocera dimidiata|[11468]|PHTCH361-08|BOLD:AAB5747
Prionocera dimidiata|[11469]|PHTCH365-08|BOLD:AAB5747
Prionocera dimidiata|[11470]|PHTCH367-08|BOLD:AAB5747
Prionocera dimidiata|[11471]|PHTCH368-08|BOLD:AAB5747
Prionocera dimidiata|[11472]|PHTCH369-08|BOLD:AAB5747
Prionocera dimidiata|[11473]|PHTCH349-08|BOLD:AAB5747
Prionocera dimidiata|[11474]|PHTCH177-08|BOLD:AAB5747
Prionocera dimidiata|[11475]|PHTCH194-08|BOLD:AAB5747
Prionocera dimidiata|[11476]|PHTCH199-08|BOLD:AAB5747
Prionocera dimidiata|[11477]|PHTCH201-08|BOLD:AAB5747
Prionocera dimidiata|[11478]|PHTCH377-08|BOLD:AAB5747
Prionocera dimidiata|[11479]|PHTCH030-07|BOLD:AAB5747
Prionocera dimidiata|[11480]|JWDCF022-10|BOLD:AAB5747
Prionocera dimidiata|[11481]|JWDCK002-11|BOLD:AAB5747
Prionocera dimidiata|[11482]|JWDCJ772-11|BOLD:AAB5747
Prionocera dimidiata|[11483]|JWDCG023-10|BOLD:AAB5747
Prionocera dimidiata|[11484]|JWDCF787-10|BOLD:AAB5747
Prionocera dimidiata|[11485]|JWDCF808-10|BOLD:AAB5747
Prionocera dimidiata|[11486]|JWDCF927-10|BOLD:AAB5747
Prionocera dimidiata|[11487]|JWDCF929-10|BOLD:AAB5747
Prionocera dimidiata|[11488]|TICHU122-09|BOLD:AAB5747
Prionocera unimicra|[11489]|JWDCE525-10|BOLD:AAK9530
Prionocera unimicra|[11490]|PHTCH176-08|BOLD:AAK9530
Prionocera unimicra|[11491]|JWDCE444-10|BOLD:AAK9530
Prionocera unimicra|[11492]|JWDCE526-10|BOLD:AAK9530
Prionocera ominosa|[11493]|JWDCE443-10|BOLD:AAP4846
Prionocera ominosa|[11494]|JWDCE300-10|BOLD:AAP4846
Prionocera ominosa|[11495]|JWDCF045-10|BOLD:AAP4846
Prionocera ominosa|[11496]|JWDCE580-10|BOLD:AAP4846
Prionocera pubescens|[11497]|JWDCE576-10|BOLD:AAP4846
Prionocera ominosa|[11498]|JWDCE299-10|BOLD:AAP4846
Prionocera ominosa|[11499]|CNCTI008-12|BOLD:AAP4846
Tipula coloradensis|[11500]|PHTCH276-08|BOLD:AAA8248
Tipula coloradensis|[11501]|PHTCH391-08|BOLD:AAA8248
Tipula coloradensis|[11502]|PHTCH409-08|BOLD:AAA8248
Tipula coloradensis|[11503]|PHTCH414-08|BOLD:AAA8248
Tipula coloradensis|[11504]|PHTCH274-08|BOLD:AAA8248
Tipula coloradensis|[11505]|PHTCH275-08|BOLD:AAA8248
Tipula coloradensis|[11506]|PHTCH278-08|BOLD:AAA8248
Tipula coloradensis|[11507]|PHTCH279-08|BOLD:AAA8248
Tipula coloradensis|[11508]|PHTCH280-08|BOLD:AAA8248
Tipula coloradensis|[11509]|PHTCH281-08|BOLD:AAA8248
Tipula coloradensis|[11510]|PHTCH282-08|BOLD:AAA8248
Tipula coloradensis|[11511]|PHTCH325-08|BOLD:AAA8248
Tipula coloradensis|[11512]|PHTCH124-08|BOLD:AAA8248
Tipula coloradensis|[11513]|PHTCH047-07|BOLD:AAA8248
Tipula resurgens|[11514]|PHTCH050-07|BOLD:AAA8248
Tipula coloradensis|[11515]|PHTCH085-07|BOLD:AAA8248
Tipula coloradensis|[11516]|PHTCH330-08|BOLD:AAA8248
Tipula resurgens|[11517]|PHTCH038-07|BOLD:AAA8248
Tipula coloradensis|[11518]|PHTCH087-07|BOLD:AAA8248
Tipula coloradensis|[11519]|PHTCH022-07|BOLD:AAA8248
Tipula coloradensis|[11520]|PHTCH044-07|BOLD:AAA8248
Tipula coloradensis|[11521]|PHTCH012-07|BOLD:AAA8248
Tipula coloradensis|[11522]|PHTCH413-08|BOLD:AAA8248
Tipula coloradensis|[11523]|PHTCH412-08|BOLD:AAA8248
Tipula coloradensis|[11524]|PHTCH041-07|BOLD:AAA8248
Tipula coloradensis|[11525]|PHTCH383-08|BOLD:AAA8248
Tipula coloradensis|[11526]|PHTCH045-07|BOLD:AAA8248
Tipula coloradensis|[11527]|PHTCH406-08|BOLD:AAA8248
Tipula coloradensis|[11528]|PHTCH277-08|BOLD:AAA8248
Tipula coloradensis|[11529]|PHTCH013-07|BOLD:AAA8248
Tipula coloradensis|[11530]|PHTCH396-08|BOLD:AAA8248
Tipula coloradensis|[11531]|PHTCH415-08|BOLD:AAA8248
Tipula coloradensis|[11532]|PHTCH411-08|BOLD:AAA8248
Tipula coloradensis|[11533]|PHTCH283-08|BOLD:AAA8248
Tipula coloradensis|[11534]|PHTCH286-08|BOLD:AAA8248
Tipula coloradensis|[11535]|PHTCH291-08|BOLD:AAA8248
Tipula coloradensis|[11536]|PHTCH339-08|BOLD:AAA8248
Tipula coloradensis|[11537]|PHTCH002-07|BOLD:AAA8248
Tipula coloradensis|[11538]|PHTCH032-07|BOLD:AAA8248
Tipula coloradensis|[11539]|PHTCH040-07|BOLD:AAA8248
Tipula coloradensis|[11540]|JWDCF792-10|BOLD:AAA8248
Tipula sulphurea|[11541]|PHTCH329-08|BOLD:AAD2627
Tipula sulphurea|[11542]|PHTCH326-08|BOLD:ACE9065
Tipula sulphurea|[11543]|PHTCH354-08|BOLD:ACE9065

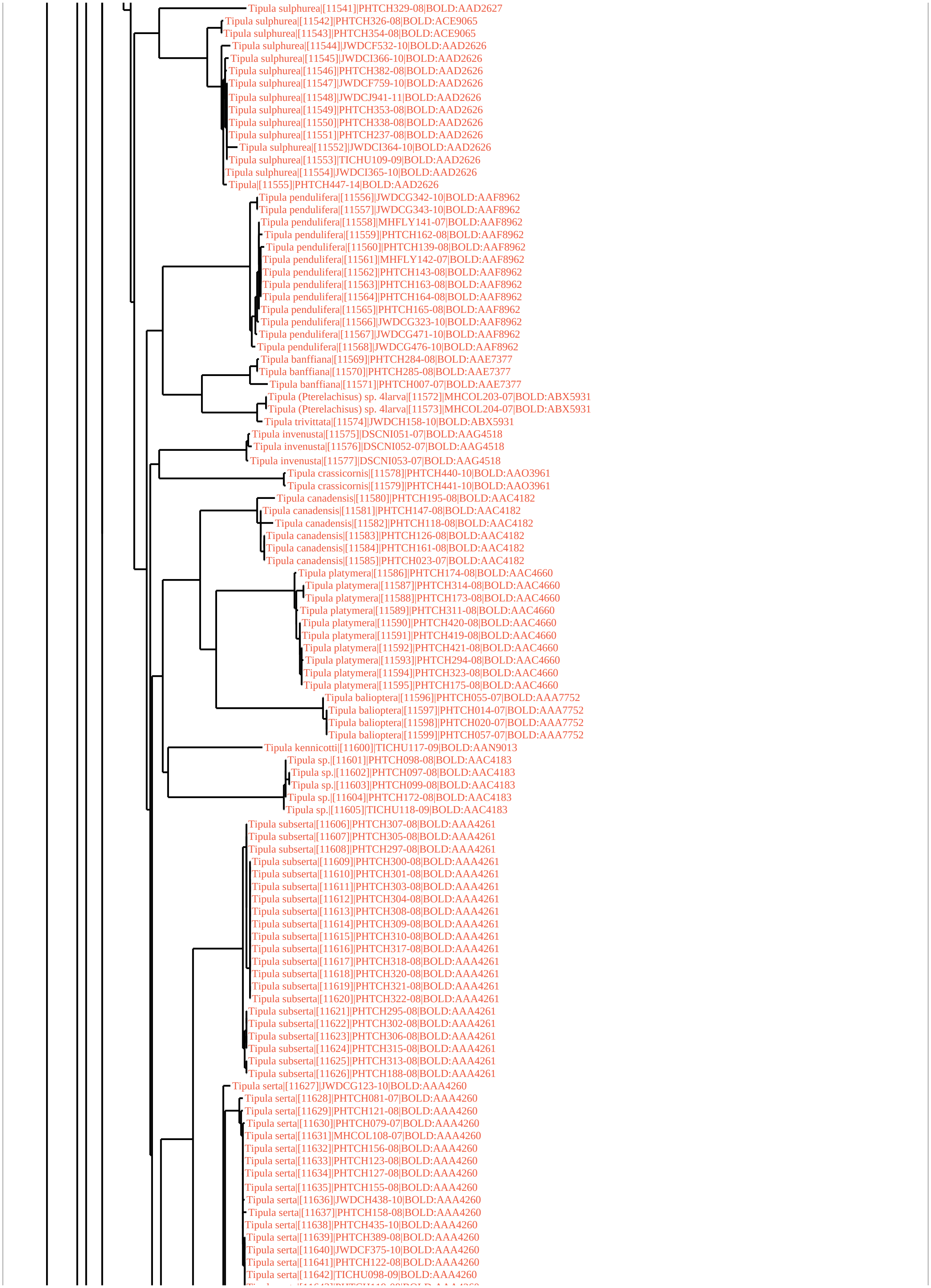
Tipula sulphurea|[11541]|PHTCH329-08|BOLD:AAD2627
Tipula sulphurea|[11542]|PHTCH326-08|BOLD:ACE9065
Tipula sulphurea|[11543]|PHTCH354-08|BOLD:ACE9065
Tipula sulphurea|[11544]|JWDCF532-10|BOLD:AAD2626
Tipula sulphurea|[11545]|JWDCI366-10|BOLD:AAD2626
Tipula sulphurea|[11546]|PHTCH382-08|BOLD:AAD2626
Tipula sulphurea|[11547]|JWDCF759-10|BOLD:AAD2626
Tipula sulphurea|[11548]|JWDCJ941-11|BOLD:AAD2626
Tipula sulphurea|[11549]|PHTCH353-08|BOLD:AAD2626
Tipula sulphurea|[11550]|PHTCH338-08|BOLD:AAD2626
Tipula sulphurea|[11551]|PHTCH237-08|BOLD:AAD2626
Tipula sulphurea|[11552]|JWDCI364-10|BOLD:AAD2626
Tipula sulphurea|[11553]|TICHU109-09|BOLD:AAD2626
Tipula sulphurea|[11554]|JWDCI365-10|BOLD:AAD2626
Tipula|[11555]|PHTCH447-14|BOLD:AAD2626
Tipula pendulifera|[11556]|JWDCG342-10|BOLD:AAF8962
Tipula pendulifera|[11557]|JWDCG343-10|BOLD:AAF8962
Tipula pendulifera|[11558]|MHFLY141-07|BOLD:AAF8962
Tipula pendulifera|[11559]|PHTCH162-08|BOLD:AAF8962
Tipula pendulifera|[11560]|PHTCH139-08|BOLD:AAF8962
Tipula pendulifera|[11561]|MHFLY142-07|BOLD:AAF8962
Tipula pendulifera|[11562]|PHTCH143-08|BOLD:AAF8962
Tipula pendulifera|[11563]|PHTCH163-08|BOLD:AAF8962
Tipula pendulifera|[11564]|PHTCH164-08|BOLD:AAF8962
Tipula pendulifera|[11565]|PHTCH165-08|BOLD:AAF8962
Tipula pendulifera|[11566]|JWDCG323-10|BOLD:AAF8962
Tipula pendulifera|[11567]|JWDCG471-10|BOLD:AAF8962
Tipula pendulifera|[11568]|JWDCG476-10|BOLD:AAF8962
Tipula banffiana|[11569]|PHTCH284-08|BOLD:AAE7377
Tipula banffiana|[11570]|PHTCH285-08|BOLD:AAE7377
Tipula banffiana|[11571]|PHTCH007-07|BOLD:AAE7377
Tipula (Pterelachisus) sp. 4larva|[11572]|MHCOL203-07|BOLD:ABX5931
Tipula (Pterelachisus) sp. 4larva|[11573]|MHCOL204-07|BOLD:ABX5931
Tipula trivittata|[11574]|JWDCH158-10|BOLD:ABX5931
Tipula invenusta|[11575]|DSCNI051-07|BOLD:AAG4518
Tipula invenusta|[11576]|DSCNI052-07|BOLD:AAG4518
Tipula invenusta|[11577]|DSCNI053-07|BOLD:AAG4518
Tipula crassicornis|[11578]|PHTCH440-10|BOLD:AAO3961
Tipula crassicornis|[11579]|PHTCH441-10|BOLD:AAO3961
Tipula canadensis|[11580]|PHTCH195-08|BOLD:AAC4182
Tipula canadensis|[11581]|PHTCH147-08|BOLD:AAC4182
Tipula canadensis|[11582]|PHTCH118-08|BOLD:AAC4182
Tipula canadensis|[11583]|PHTCH126-08|BOLD:AAC4182
Tipula canadensis|[11584]|PHTCH161-08|BOLD:AAC4182
Tipula canadensis|[11585]|PHTCH023-07|BOLD:AAC4182
Tipula platymera|[11586]|PHTCH174-08|BOLD:AAC4660
Tipula platymera|[11587]|PHTCH314-08|BOLD:AAC4660
Tipula platymera|[11588]|PHTCH173-08|BOLD:AAC4660
Tipula platymera|[11589]|PHTCH311-08|BOLD:AAC4660
Tipula platymera|[11590]|PHTCH420-08|BOLD:AAC4660
Tipula platymera|[11591]|PHTCH419-08|BOLD:AAC4660
Tipula platymera|[11592]|PHTCH421-08|BOLD:AAC4660
Tipula platymera|[11593]|PHTCH294-08|BOLD:AAC4660
Tipula platymera|[11594]|PHTCH323-08|BOLD:AAC4660
Tipula platymera|[11595]|PHTCH175-08|BOLD:AAC4660
Tipula balioptera|[11596]|PHTCH055-07|BOLD:AAA7752
Tipula balioptera|[11597]|PHTCH014-07|BOLD:AAA7752
Tipula balioptera|[11598]|PHTCH020-07|BOLD:AAA7752
Tipula balioptera|[11599]|PHTCH057-07|BOLD:AAA7752
Tipula kennicotti|[11600]|TICHU117-09|BOLD:AAN9013
Tipula sp.|[11601]|PHTCH098-08|BOLD:AAC4183
Tipula sp.|[11602]|PHTCH097-08|BOLD:AAC4183
Tipula sp.|[11603]|PHTCH099-08|BOLD:AAC4183
Tipula sp.|[11604]|PHTCH172-08|BOLD:AAC4183
Tipula sp.|[11605]|TICHU118-09|BOLD:AAC4183
Tipula subserta|[11606]|PHTCH307-08|BOLD:AAA4261
Tipula subserta|[11607]|PHTCH305-08|BOLD:AAA4261
Tipula subserta|[11608]|PHTCH297-08|BOLD:AAA4261
Tipula subserta|[11609]|PHTCH300-08|BOLD:AAA4261
Tipula subserta|[11610]|PHTCH301-08|BOLD:AAA4261
Tipula subserta|[11611]|PHTCH303-08|BOLD:AAA4261
Tipula subserta|[11612]|PHTCH304-08|BOLD:AAA4261
Tipula subserta|[11613]|PHTCH308-08|BOLD:AAA4261
Tipula subserta|[11614]|PHTCH309-08|BOLD:AAA4261
Tipula subserta|[11615]|PHTCH310-08|BOLD:AAA4261
Tipula subserta|[11616]|PHTCH317-08|BOLD:AAA4261
Tipula subserta|[11617]|PHTCH318-08|BOLD:AAA4261
Tipula subserta|[11618]|PHTCH320-08|BOLD:AAA4261
Tipula subserta|[11619]|PHTCH321-08|BOLD:AAA4261
Tipula subserta|[11620]|PHTCH322-08|BOLD:AAA4261
Tipula subserta|[11621]|PHTCH295-08|BOLD:AAA4261
Tipula subserta|[11622]|PHTCH302-08|BOLD:AAA4261
Tipula subserta|[11623]|PHTCH306-08|BOLD:AAA4261
Tipula subserta|[11624]|PHTCH315-08|BOLD:AAA4261
Tipula subserta|[11625]|PHTCH313-08|BOLD:AAA4261
Tipula subserta|[11626]|PHTCH188-08|BOLD:AAA4261
Tipula serta|[11627]|JWDCG123-10|BOLD:AAA4260
Tipula serta|[11628]|PHTCH081-07|BOLD:AAA4260
Tipula serta|[11629]|PHTCH121-08|BOLD:AAA4260
Tipula serta|[11630]|PHTCH079-07|BOLD:AAA4260
Tipula serta|[11631]|MHCOL108-07|BOLD:AAA4260
Tipula serta|[11632]|PHTCH156-08|BOLD:AAA4260
Tipula serta|[11633]|PHTCH123-08|BOLD:AAA4260
Tipula serta|[11634]|PHTCH127-08|BOLD:AAA4260
Tipula serta|[11635]|PHTCH155-08|BOLD:AAA4260
Tipula serta|[11636]|JWDCH438-10|BOLD:AAA4260
Tipula serta|[11637]|PHTCH158-08|BOLD:AAA4260
Tipula serta|[11638]|PHTCH435-10|BOLD:AAA4260
Tipula serta|[11639]|PHTCH389-08|BOLD:AAA4260
Tipula serta|[11640]|JWDCF375-10|BOLD:AAA4260
Tipula serta|[11641]|PHTCH122-08|BOLD:AAA4260
Tipula serta|[11642]|TICHU098-09|BOLD:AAA4260

Tipula serta|[11640]|JWDCF373-10|BOLD:AAA4260
Tipula serta|[11641]|PHTCH122-08|BOLD:AAA4260
Tipula serta|[11642]|TICHU098-09|BOLD:AAA4260
Tipula serta|[11643]|PHTCH119-08|BOLD:AAA4260
Tipula serta|[11644]|PHTCH434-10|BOLD:AAA4260
Tipula serta|[11645]|PHTCH153-08|BOLD:AAA4260
Tipula serta|[11646]|PHTCH001-07|BOLD:AAA4260
Tipula serta|[11647]|PHTCH083-07|BOLD:AAA4260
Tipula serta|[11648]|PHTCH073-07|BOLD:AAA4260
Tipula serta|[11649]|JWDCJ1034-11|BOLD:AAA4260
Tipula serta|[11650]|JWDCG019-10|BOLD:AAA4260
Tipula serta|[11651]|JWDCH088-10|BOLD:AAA4260
Tipula serta|[11652]|JWDCF379-10|BOLD:AAA4260
Tipula serta|[11653]|JWDCG779-10|BOLD:AAA4260
Tipula serta|[11654]|JWDCG747-10|BOLD:AAA4260
Tipula serta|[11655]|PHTCH437-10|BOLD:AAA4260
Tipula serta|[11656]|PHTCH053-07|BOLD:AAA4260
Tipula serta|[11657]|PHTCH049-07|BOLD:AAA4260
Tipula serta|[11658]|PHTCH018-07|BOLD:AAA4260
Tipula serta|[11659]|PHTCH390-08|BOLD:AAA4260
Tipula serta|[11660]|PHTCH387-08|BOLD:AAA4260
Tipula serta|[11661]|PHTCH192-08|BOLD:AAA4260
Tipula serta|[11662]|PHTCH089-07|BOLD:AAA4260
Tipula serta|[11663]|PHTCH157-08|BOLD:AAA4260
Tipula serta|[11664]|PHTCH125-08|BOLD:AAA4260
Tipula serta|[11665]|PHTCH117-08|BOLD:AAA4260
Tipula serta|[11666]|PHTCH100-08|BOLD:AAA4260
Tipula serta|[11667]|PHTCH355-08|BOLD:AAA4260
Tipula serta|[11668]|PHTCH331-08|BOLD:AAA4260
Tipula serta|[11669]|PHTCH328-08|BOLD:AAA4260
Tipula serta|[11670]|PHTCH319-08|BOLD:AAA4260
Tipula serta|[11671]|PHTCH316-08|BOLD:AAA4260
Tipula serta|[11672]|PHTCH252-08|BOLD:AAA4260
Tipula serta|[11673]|PHTCH269-08|BOLD:AAA4260
Tipula serta|[11674]|PHTCH268-08|BOLD:AAA4260
Tipula serta|[11675]|PHTCH262-08|BOLD:AAA4260
Tipula serta|[11676]|PHTCH261-08|BOLD:AAA4260
Tipula serta|[11677]|PHTCH260-08|BOLD:AAA4260
Tipula serta|[11678]|PHTCH259-08|BOLD:AAA4260
Tipula serta|[11679]|PHTCH258-08|BOLD:AAA4260
Tipula serta|[11680]|PHTCH257-08|BOLD:AAA4260
Tipula serta|[11681]|PHTCH256-08|BOLD:AAA4260
Tipula serta|[11682]|PHTCH255-08|BOLD:AAA4260
Tipula serta|[11683]|PHTCH253-08|BOLD:AAA4260
Tipula serta|[11684]|PHTCH251-08|BOLD:AAA4260
Tipula serta|[11685]|PHTCH393-08|BOLD:AAA4260
Tipula serta|[11686]|PHTCH392-08|BOLD:AAA4260
Tipula serta|[11687]|PHTCH056-07|BOLD:AAA4260
Tipula serta|[11688]|JWDCG749-10|BOLD:AAA4260
Tipula serta|[11689]|JWDCF427-10|BOLD:AAA4260
Tipula serta|[11690]|PHTCH386-08|BOLD:AAA4260
Tipula serta|[11691]|PHTCH271-08|BOLD:AAA4260
Tipula serta|[11692]|PHTCH433-10|BOLD:AAA4260
Tipula serta|[11693]|PHTCH432-10|BOLD:AAA4260
Tipula serta|[11694]|PHTCH152-08|BOLD:AAA4260
Tipula serta|[11695]|PHTCH254-08|BOLD:AAA4260
Tipula serta|[11696]|PHTCH084-07|BOLD:AAA4260
Tipula serta|[11697]|TICHU107-09|BOLD:AAA4260
Tipula serta|[11698]|SATIP220-09|BOLD:AAA4260
Tipula serta|[11699]|SATIP221-09|BOLD:AAA4260
Tipula serta|[11700]|SATIP223-09|BOLD:AAA4260
Tipula serta|[11701]|CNCTI071-12|BOLD:AAA4260
Tipula serta|[11702]|PHTCH031-07|BOLD:AAA4260
Tipula serta|[11703]|PHTCH439-10|BOLD:AAA4260
Tipula serta|[11704]|CNCTI072-12|BOLD:AAA4260
Tipula parshleyi|[11705]|PHTCH405-08|BOLD:AAF8980
Tipula macrolabis|[11706]|PHTCH151-08|BOLD:AAG4520
Tipula macrolabis|[11707]|PHTCH137-08
Tipula saxemontana|[11708]|PHTCH332-08|BOLD:AAA4262
Tipula saxemontana|[11709]|PHTCH076-07|BOLD:AAA4262
Tipula saxemontana|[11710]|PHTCH116-08|BOLD:AAA4262
Tipula saxemontana|[11711]|PHTCH395-08|BOLD:AAA4262
Tipula saxemontana|[11712]|PHTCH394-08|BOLD:AAA4262
Tipula saxemontana|[11713]|PHTCH388-08|BOLD:AAA4262
Tipula saxemontana|[11714]|PHTCH028-07|BOLD:AAA4262
Tipula saxemontana|[11715]|PHTCH061-07|BOLD:AAA4262
Tipula saxemontana|[11716]|PHTCH066-07|BOLD:AAA4262
Tipula saxemontana|[11717]|PHTCH069-07|BOLD:AAA4262
Tipula saxemontana|[11718]|PHTCH071-07|BOLD:AAA4262
Tipula saxemontana|[11719]|PHTCH072-07|BOLD:AAA4262
Tipula saxemontana|[11720]|SATIP222-09|BOLD:AAA4262
Tipula saxemontana|[11721]|CNCTI084-12|BOLD:AAA4262
Tipula saxemontana|[11722]|CNCDI107-11|BOLD:AAA4262
Euthyneura|[11723]|JWDCA114-10|BOLD:AAF9859
Sylvicola punctatus|[11724]|JWDCJ1408-11|BOLD:AAG1999
Sylvicola punctatus|[11725]|JWDCF166-10|BOLD:AAG1999
Villa|[11726]|MHFLY066-06|BOLD:AAH0541
Saetheria tylus|[11727]|CHURC464-07|BOLD:AAJ7404
Euphylidorea platyphallus|[11728]|PHTCH407-08|BOLD:AAD6847
Euphylidorea platyphallus|[11729]|JWDCF873-10|BOLD:AAD6847
Euphylidorea platyphallus|[11730]|PHTCH340-08|BOLD:AAD6847
Euphylidorea platyphallus|[11731]|PHTCH075-07|BOLD:AAD6847
Euphylidorea platyphallus|[11732]|JWDCI306-10|BOLD:AAD6847
Euphylidorea platyphallus|[11733]|JWDCG124-10|BOLD:AAD6847
Euphylidorea platyphallus|[11734]|JWDCF807-10|BOLD:AAD6847
Euphylidorea platyphallus|[11735]|PHTCH034-07|BOLD:AAD6847
Euphylidorea platyphallus|[11736]|PHTCH408-08|BOLD:AAD6847
Euphylidorea platyphallus|[11737]|JWDCH159-10|BOLD:AAD6847
Euphylidorea platyphallus|[11738]|JWDCH157-10|BOLD:AAD6847
Euphylidorea platyphallus|[11739]|TICHU096-09|BOLD:AAD6847
Dixella|[11740]|MHCOL443-07|BOLD:AAI1321
Dixella|[11741]|MHCOL444-07|BOLD:AAI1321
Dixella|[11742]|JWDCK004-11|BOLD:AAI1321

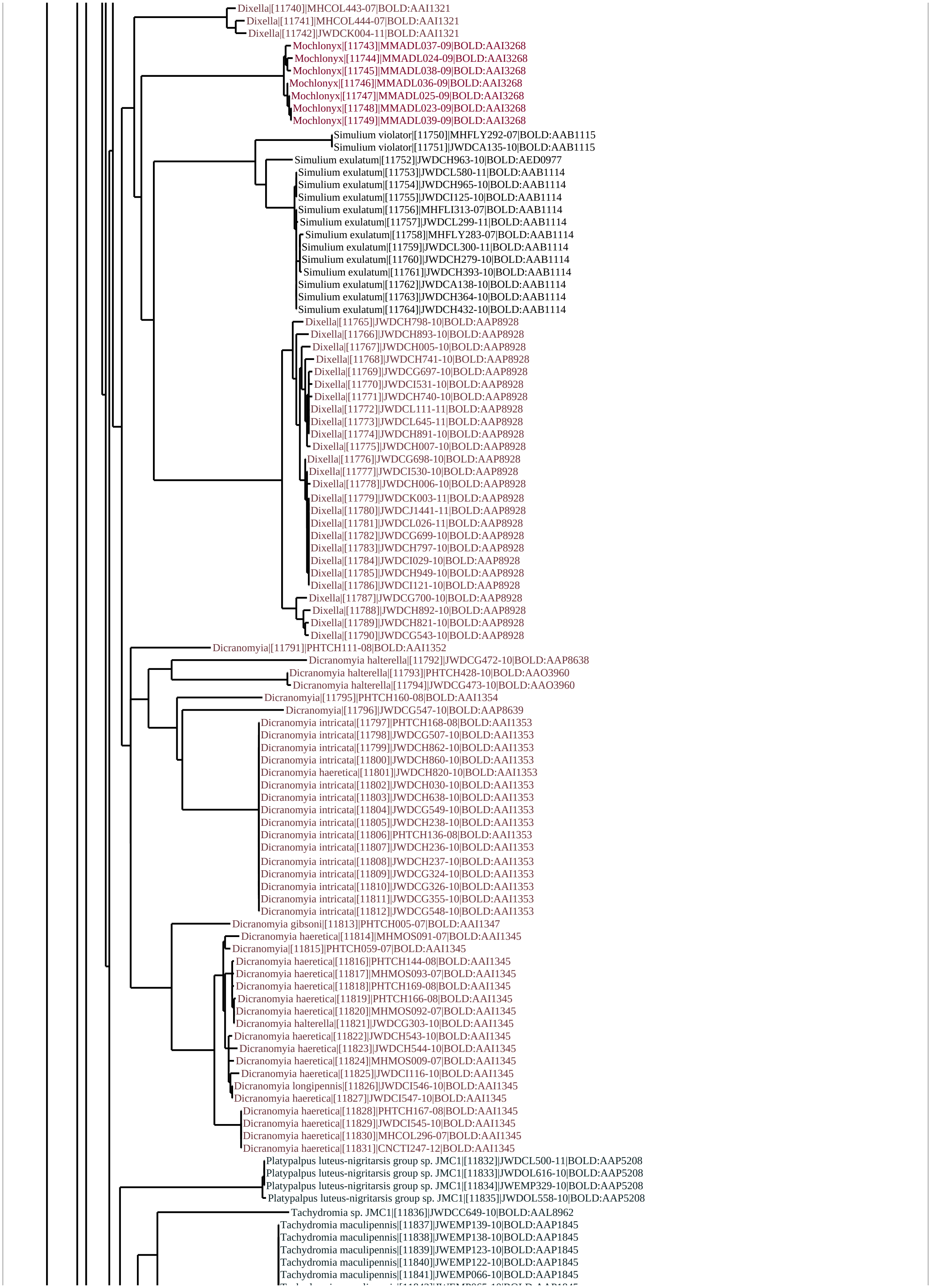
Dixella|[11740]|MHCOL443-07|BOLD:AAI1321
Dixella|[11741]|MHCOL444-07|BOLD:AAI1321
Dixella|[11742]|JWDCK004-11|BOLD:AAI1321
Mochlonyx|[11743]|MMADL037-09|BOLD:AAI3268
Mochlonyx|[11744]|MMADL024-09|BOLD:AAI3268
Mochlonyx|[11745]|MMADL038-09|BOLD:AAI3268
Mochlonyx|[11746]|MMADL036-09|BOLD:AAI3268
Mochlonyx|[11747]|MMADL025-09|BOLD:AAI3268
Mochlonyx|[11748]|MMADL023-09|BOLD:AAI3268
Mochlonyx|[11749]|MMADL039-09|BOLD:AAI3268
Simulium violator|[11750]|MHFLY292-07|BOLD:AAB1115
Simulium violator|[11751]|JWDCA135-10|BOLD:AAB1115
Simulium exulatum|[11752]|JWDCH963-10|BOLD:AED0977
Simulium exulatum|[11753]|JWDCL580-11|BOLD:AAB1114
Simulium exulatum|[11754]|JWDCH965-10|BOLD:AAB1114
Simulium exulatum|[11755]|JWDCI125-10|BOLD:AAB1114
Simulium exulatum|[11756]|MHFLI313-07|BOLD:AAB1114
Simulium exulatum|[11757]|JWDCL299-11|BOLD:AAB1114
Simulium exulatum|[11758]|MHFLY283-07|BOLD:AAB1114
Simulium exulatum|[11759]|JWDCL300-11|BOLD:AAB1114
Simulium exulatum|[11760]|JWDCH279-10|BOLD:AAB1114
Simulium exulatum|[11761]|JWDCH393-10|BOLD:AAB1114
Simulium exulatum|[11762]|JWDCA138-10|BOLD:AAB1114
Simulium exulatum|[11763]|JWDCH364-10|BOLD:AAB1114
Simulium exulatum|[11764]|JWDCH432-10|BOLD:AAB1114
Dixella|[11765]|JWDCH798-10|BOLD:AAP8928
Dixella|[11766]|JWDCH893-10|BOLD:AAP8928
Dixella|[11767]|JWDCH005-10|BOLD:AAP8928
Dixella|[11768]|JWDCH741-10|BOLD:AAP8928
Dixella|[11769]|JWDCG697-10|BOLD:AAP8928
Dixella|[11770]|JWDCI531-10|BOLD:AAP8928
Dixella|[11771]|JWDCH740-10|BOLD:AAP8928
Dixella|[11772]|JWDCL111-11|BOLD:AAP8928
Dixella|[11773]|JWDCL645-11|BOLD:AAP8928
Dixella|[11774]|JWDCH891-10|BOLD:AAP8928
Dixella|[11775]|JWDCH007-10|BOLD:AAP8928
Dixella|[11776]|JWDCG698-10|BOLD:AAP8928
Dixella|[11777]|JWDCI530-10|BOLD:AAP8928
Dixella|[11778]|JWDCH006-10|BOLD:AAP8928
Dixella|[11779]|JWDCK003-11|BOLD:AAP8928
Dixella|[11780]|JWDCJ1441-11|BOLD:AAP8928
Dixella|[11781]|JWDCL026-11|BOLD:AAP8928
Dixella|[11782]|JWDCG699-10|BOLD:AAP8928
Dixella|[11783]|JWDCH797-10|BOLD:AAP8928
Dixella|[11784]|JWDCI029-10|BOLD:AAP8928
Dixella|[11785]|JWDCH949-10|BOLD:AAP8928
Dixella|[11786]|JWDCI121-10|BOLD:AAP8928
Dixella|[11787]|JWDCG700-10|BOLD:AAP8928
Dixella|[11788]|JWDCH892-10|BOLD:AAP8928
Dixella|[11789]|JWDCH821-10|BOLD:AAP8928
Dixella|[11790]|JWDCG543-10|BOLD:AAP8928
Dicranomyia|[11791]|PHTCH111-08|BOLD:AAI1352
Dicranomyia halterella|[11792]|JWDCG472-10|BOLD:AAP8638
Dicranomyia halterella|[11793]|PHTCH428-10|BOLD:AAO3960
Dicranomyia halterella|[11794]|JWDCG473-10|BOLD:AAO3960
Dicranomyia|[11795]|PHTCH160-08|BOLD:AAI1354
Dicranomyia|[11796]|JWDCG547-10|BOLD:AAP8639
Dicranomyia intricata|[11797]|PHTCH168-08|BOLD:AAI1353
Dicranomyia intricata|[11798]|JWDCG507-10|BOLD:AAI1353
Dicranomyia intricata|[11799]|JWDCH862-10|BOLD:AAI1353
Dicranomyia intricata|[11800]|JWDCH860-10|BOLD:AAI1353
Dicranomyia haeretica|[11801]|JWDCH820-10|BOLD:AAI1353
Dicranomyia intricata|[11802]|JWDCH030-10|BOLD:AAI1353
Dicranomyia intricata|[11803]|JWDCH638-10|BOLD:AAI1353
Dicranomyia intricata|[11804]|JWDCG549-10|BOLD:AAI1353
Dicranomyia intricata|[11805]|JWDCH238-10|BOLD:AAI1353
Dicranomyia intricata|[11806]|PHTCH136-08|BOLD:AAI1353
Dicranomyia intricata|[11807]|JWDCH236-10|BOLD:AAI1353
Dicranomyia intricata|[11808]|JWDCH237-10|BOLD:AAI1353
Dicranomyia intricata|[11809]|JWDCG324-10|BOLD:AAI1353
Dicranomyia intricata|[11810]|JWDCG326-10|BOLD:AAI1353
Dicranomyia intricata|[11811]|JWDCG355-10|BOLD:AAI1353
Dicranomyia intricata|[11812]|JWDCG548-10|BOLD:AAI1353
Dicranomyia gibsoni|[11813]|PHTCH005-07|BOLD:AAI1347
Dicranomyia haeretica|[11814]|MHMOS091-07|BOLD:AAI1345
Dicranomyia|[11815]|PHTCH059-07|BOLD:AAI1345
Dicranomyia haeretica|[11816]|PHTCH144-08|BOLD:AAI1345
Dicranomyia haeretica|[11817]|MHMOS093-07|BOLD:AAI1345
Dicranomyia haeretica|[11818]|PHTCH169-08|BOLD:AAI1345
Dicranomyia haeretica|[11819]|PHTCH166-08|BOLD:AAI1345
Dicranomyia haeretica|[11820]|MHMOS092-07|BOLD:AAI1345
Dicranomyia halterella|[11821]|JWDCG303-10|BOLD:AAI1345
Dicranomyia haeretica|[11822]|JWDCH543-10|BOLD:AAI1345
Dicranomyia haeretica|[11823]|JWDCH544-10|BOLD:AAI1345
Dicranomyia haeretica|[11824]|MHMOS009-07|BOLD:AAI1345
Dicranomyia haeretica|[11825]|JWDCI116-10|BOLD:AAI1345
Dicranomyia longipennis|[11826]|JWDCI546-10|BOLD:AAI1345
Dicranomyia haeretica|[11827]|JWDCI547-10|BOLD:AAI1345
Dicranomyia haeretica|[11828]|PHTCH167-08|BOLD:AAI1345
Dicranomyia haeretica|[11829]|JWDCI545-10|BOLD:AAI1345
Dicranomyia haeretica|[11830]|MHCOL296-07|BOLD:AAI1345
Dicranomyia haeretica|[11831]|CNCTI247-12|BOLD:AAI1345
Platypalpus luteus-nigritarsis group sp. JMC1|[11832]|JWDCL500-11|BOLD:AAP5208
Platypalpus luteus-nigritarsis group sp. JMC1|[11833]|JWDOL616-10|BOLD:AAP5208
Platypalpus luteus-nigritarsis group sp. JMC1|[11834]|JWEMP329-10|BOLD:AAP5208
Platypalpus luteus-nigritarsis group sp. JMC1|[11835]|JWDOL558-10|BOLD:AAP5208
Tachydromia sp. JMC1|[11836]|JWDCC649-10|BOLD:AAL8962
Tachydromia maculipennis|[11837]|JWEMP139-10|BOLD:AAP1845
Tachydromia maculipennis|[11838]|JWEMP138-10|BOLD:AAP1845
Tachydromia maculipennis|[11839]|JWEMP123-10|BOLD:AAP1845
Tachydromia maculipennis|[11840]|JWEMP122-10|BOLD:AAP1845
Tachydromia maculipennis|[11841]|JWEMP066-10|BOLD:AAP1845

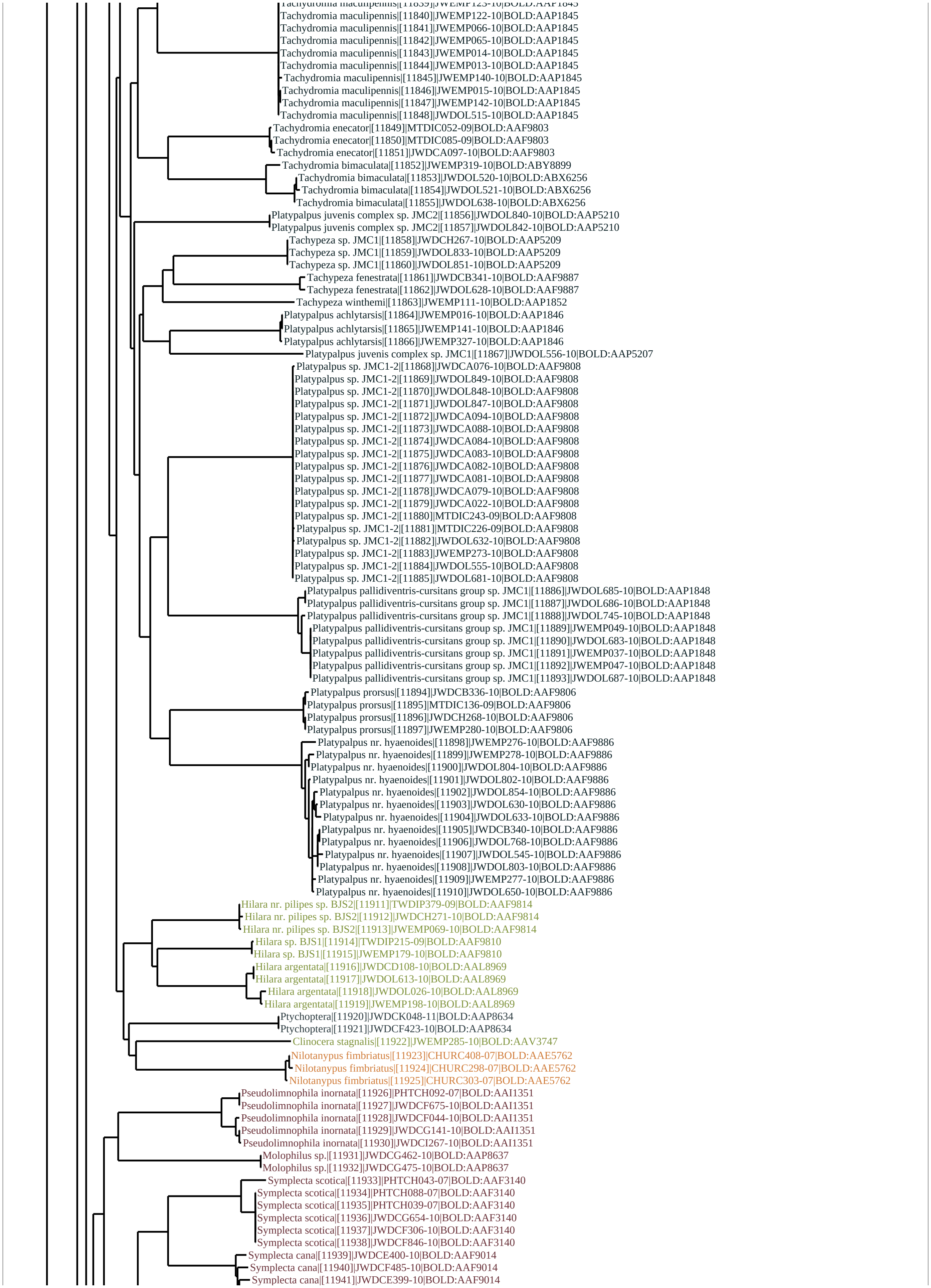
Tachydromia maculipennis|[11839]|JWEMP123-10|BOLD:AAP1845
Tachydromia maculipennis|[11840]|JWEMP122-10|BOLD:AAP1845
Tachydromia maculipennis|[11841]|JWEMP066-10|BOLD:AAP1845
Tachydromia maculipennis|[11842]|JWEMP065-10|BOLD:AAP1845
Tachydromia maculipennis|[11843]|JWEMP014-10|BOLD:AAP1845
Tachydromia maculipennis|[11844]|JWEMP013-10|BOLD:AAP1845
Tachydromia maculipennis|[11845]|JWEMP140-10|BOLD:AAP1845
Tachydromia maculipennis|[11846]|JWEMP015-10|BOLD:AAP1845
Tachydromia maculipennis|[11847]|JWEMP142-10|BOLD:AAP1845
Tachydromia maculipennis|[11848]|JWDOL515-10|BOLD:AAP1845
Tachydromia enecator|[11849]|MTDIC052-09|BOLD:AAF9803
Tachydromia enecator|[11850]|MTDIC085-09|BOLD:AAF9803
Tachydromia enecator|[11851]|JWDCA097-10|BOLD:AAF9803
Tachydromia bimaculata|[11852]|JWEMP319-10|BOLD:ABY8899
Tachydromia bimaculata|[11853]|JWDOL520-10|BOLD:ABX6256
Tachydromia bimaculata|[11854]|JWDOL521-10|BOLD:ABX6256
Tachydromia bimaculata|[11855]|JWDOL638-10|BOLD:ABX6256
Platypalpus juvenis complex sp. JMC2|[11856]|JWDOL840-10|BOLD:AAP5210
Platypalpus juvenis complex sp. JMC2|[11857]|JWDOL842-10|BOLD:AAP5210
Tachypeza sp. JMC1|[11858]|JWDCH267-10|BOLD:AAP5209
Tachypeza sp. JMC1|[11859]|JWDOL833-10|BOLD:AAP5209
Tachypeza sp. JMC1|[11860]|JWDOL851-10|BOLD:AAP5209
Tachypeza fenestrata|[11861]|JWDCB341-10|BOLD:AAF9887
Tachypeza fenestrata|[11862]|JWDOL628-10|BOLD:AAF9887
Tachypeza winthemi|[11863]|JWEMP111-10|BOLD:AAP1852
Platypalpus achlytarsis|[11864]|JWEMP016-10|BOLD:AAP1846
Platypalpus achlytarsis|[11865]|JWEMP141-10|BOLD:AAP1846
Platypalpus achlytarsis|[11866]|JWEMP327-10|BOLD:AAP1846
Platypalpus juvenis complex sp. JMC1|[11867]|JWDOL556-10|BOLD:AAP5207
Platypalpus sp. JMC1-2|[11868]|JWDCA076-10|BOLD:AAF9808
Platypalpus sp. JMC1-2|[11869]|JWDOL849-10|BOLD:AAF9808
Platypalpus sp. JMC1-2|[11870]|JWDOL848-10|BOLD:AAF9808
Platypalpus sp. JMC1-2|[11871]|JWDOL847-10|BOLD:AAF9808
Platypalpus sp. JMC1-2|[11872]|JWDCA094-10|BOLD:AAF9808
Platypalpus sp. JMC1-2|[11873]|JWDCA088-10|BOLD:AAF9808
Platypalpus sp. JMC1-2|[11874]|JWDCA084-10|BOLD:AAF9808
Platypalpus sp. JMC1-2|[11875]|JWDCA083-10|BOLD:AAF9808
Platypalpus sp. JMC1-2|[11876]|JWDCA082-10|BOLD:AAF9808
Platypalpus sp. JMC1-2|[11877]|JWDCA081-10|BOLD:AAF9808
Platypalpus sp. JMC1-2|[11878]|JWDCA079-10|BOLD:AAF9808
Platypalpus sp. JMC1-2|[11879]|JWDCA022-10|BOLD:AAF9808
Platypalpus sp. JMC1-2|[11880]|MTDIC243-09|BOLD:AAF9808
Platypalpus sp. JMC1-2|[11881]|MTDIC226-09|BOLD:AAF9808
Platypalpus sp. JMC1-2|[11882]|JWDOL632-10|BOLD:AAF9808
Platypalpus sp. JMC1-2|[11883]|JWEMP273-10|BOLD:AAF9808
Platypalpus sp. JMC1-2|[11884]|JWDOL555-10|BOLD:AAF9808
Platypalpus sp. JMC1-2|[11885]|JWDOL681-10|BOLD:AAF9808
Platypalpus pallidiventris-cursitans group sp. JMC1|[11886]|JWDOL685-10|BOLD:AAP1848
Platypalpus pallidiventris-cursitans group sp. JMC1|[11887]|JWDOL686-10|BOLD:AAP1848
Platypalpus pallidiventris-cursitans group sp. JMC1|[11888]|JWDOL745-10|BOLD:AAP1848
Platypalpus pallidiventris-cursitans group sp. JMC1|[11889]|JWEMP049-10|BOLD:AAP1848
Platypalpus pallidiventris-cursitans group sp. JMC1|[11890]|JWDOL683-10|BOLD:AAP1848
Platypalpus pallidiventris-cursitans group sp. JMC1|[11891]|JWEMP037-10|BOLD:AAP1848
Platypalpus pallidiventris-cursitans group sp. JMC1|[11892]|JWEMP047-10|BOLD:AAP1848
Platypalpus pallidiventris-cursitans group sp. JMC1|[11893]|JWDOL687-10|BOLD:AAP1848
Platypalpus prorsus|[11894]|JWDCB336-10|BOLD:AAF9806
Platypalpus prorsus|[11895]|MTDIC136-09|BOLD:AAF9806
Platypalpus prorsus|[11896]|JWDCH268-10|BOLD:AAF9806
Platypalpus prorsus|[11897]|JWEMP280-10|BOLD:AAF9806
Platypalpus nr. hyaenoides|[11898]|JWEMP276-10|BOLD:AAF9886
Platypalpus nr. hyaenoides|[11899]|JWEMP278-10|BOLD:AAF9886
Platypalpus nr. hyaenoides|[11900]|JWDOL804-10|BOLD:AAF9886
Platypalpus nr. hyaenoides|[11901]|JWDOL802-10|BOLD:AAF9886
Platypalpus nr. hyaenoides|[11902]|JWDOL854-10|BOLD:AAF9886
Platypalpus nr. hyaenoides|[11903]|JWDOL630-10|BOLD:AAF9886
Platypalpus nr. hyaenoides|[11904]|JWDOL633-10|BOLD:AAF9886
Platypalpus nr. hyaenoides|[11905]|JWDCB340-10|BOLD:AAF9886
Platypalpus nr. hyaenoides|[11906]|JWDOL768-10|BOLD:AAF9886
Platypalpus nr. hyaenoides|[11907]|JWDOL545-10|BOLD:AAF9886
Platypalpus nr. hyaenoides|[11908]|JWDOL803-10|BOLD:AAF9886
Platypalpus nr. hyaenoides|[11909]|JWEMP277-10|BOLD:AAF9886
Platypalpus nr. hyaenoides|[11910]|JWDOL650-10|BOLD:AAF9886
Hilara nr. pilipes sp. BJS2|[11911]|TWDIP379-09|BOLD:AAF9814
Hilara nr. pilipes sp. BJS2|[11912]|JWDCH271-10|BOLD:AAF9814
Hilara nr. pilipes sp. BJS2|[11913]|JWEMP069-10|BOLD:AAF9814
Hilara sp. BJS1|[11914]|TWDIP215-09|BOLD:AAF9810
Hilara sp. BJS1|[11915]|JWEMP179-10|BOLD:AAF9810
Hilara argentata|[11916]|JWDCD108-10|BOLD:AAL8969
Hilara argentata|[11917]|JWDOL613-10|BOLD:AAL8969
Hilara argentata|[11918]|JWDOL026-10|BOLD:AAL8969
Hilara argentata|[11919]|JWEMP198-10|BOLD:AAL8969
Ptychoptera|[11920]|JWDCK048-11|BOLD:AAP8634
Ptychoptera|[11921]|JWDCF423-10|BOLD:AAP8634
Clinocera stagnalis|[11922]|JWEMP285-10|BOLD:AAV3747
Nilotanypus fimbriatus|[11923]|CHURC408-07|BOLD:AAE5762
Nilotanypus fimbriatus|[11924]|CHURC298-07|BOLD:AAE5762
Nilotanypus fimbriatus|[11925]|CHURC303-07|BOLD:AAE5762
Pseudolimnophila inornata|[11926]|PHTCH092-07|BOLD:AAI1351
Pseudolimnophila inornata|[11927]|JWDCF675-10|BOLD:AAI1351
Pseudolimnophila inornata|[11928]|JWDCF044-10|BOLD:AAI1351
Pseudolimnophila inornata|[11929]|JWDCG141-10|BOLD:AAI1351
Pseudolimnophila inornata|[11930]|JWDCI267-10|BOLD:AAI1351
Molophilus sp.|[11931]|JWDCG462-10|BOLD:AAP8637
Molophilus sp.|[11932]|JWDCG475-10|BOLD:AAP8637
Symplecta scotica|[11933]|PHTCH043-07|BOLD:AAF3140
Symplecta scotica|[11934]|PHTCH088-07|BOLD:AAF3140
Symplecta scotica|[11935]|PHTCH039-07|BOLD:AAF3140
Symplecta scotica|[11936]|JWDCG654-10|BOLD:AAF3140
Symplecta scotica|[11937]|JWDCF306-10|BOLD:AAF3140
Symplecta scotica|[11938]|JWDCF846-10|BOLD:AAF3140
Symplecta cana|[11939]|JWDCE400-10|BOLD:AAF9014
Symplecta cana|[11940]|JWDCF485-10|BOLD:AAF9014
Symplecta cana|[11941]|JWDCE399-10|BOLD:AAF9014

Symplecta cana|[11939]|JWDCE400-10|BOLD:AAF9014
Symplecta cana|[11940]|JWDCF485-10|BOLD:AAF9014
Symplecta cana|[11941]|JWDCE399-10|BOLD:AAF9014
Symplecta cana|[11942]|JWDCG151-10|BOLD:AAF9014
Symplecta stictica angularis|[11943]|JWDCF075-10|BOLD:AAP2736
Symplecta stictica angularis|[11944]|JWDCE638-10|BOLD:AAP2736
Symplecta stictica angularis|[11945]|JWDCJ440-11|BOLD:AAP2736
Symplecta stictica angularis|[11946]|JWDCF073-10|BOLD:AAP2736
Symplecta stictica angularis|[11947]|JWDCF074-10|BOLD:AAP2736
Symplecta stictica angularis|[11948]|JWDCI218-10|BOLD:AAP2736
Symplecta stictica angularis|[11949]|JWDCE693-10|BOLD:AAP2736
Dicranomyia spinifera|[11950]|JWDCG734-10|BOLD:AAP8640
Rhipidia maculata|[11951]|PHTCH017-07|BOLD:AAI1348
Rhipidia maculata|[11952]|PHTCH021-07|BOLD:AAI1348
Dicranomyia|[11953]|PHTCH080-07|BOLD:AAI1350
Dicranomyia humidicola|[11954]|PHTCH299-08|BOLD:AAJ2209
Dicranomyia humidicola|[11955]|PHTCH312-08|BOLD:AAJ2209
Dicranomyia chorea|[11956]|PHTCH431-10|BOLD:AAI1349
Dicranomyia chorea|[11957]|PHTCH058-07|BOLD:AAI1349
Dicranomyia chorea|[11958]|PHTCH444-14|BOLD:AAI1349
Dicranomyia chorea|[11959]|CNCTI245-12|BOLD:AAI1349
Dicranomyia chillcotti|[11960]|JWDCE925-10|BOLD:AAF4859
Dicranomyia chillcotti|[11961]|PHTCH133-08|BOLD:AAF4859
Dicranomyia chillcotti|[11962]|JWDCF711-10|BOLD:AAF4859
Dicranomyia chillcotti|[11963]|JWDCF709-10|BOLD:AAF4859
Dicranomyia chillcotti|[11964]|JWDCF095-10|BOLD:AAF4859
Dicranomyia chillcotti|[11965]|PHTCH183-08|BOLD:AAF4859
Dicranomyia chillcotti|[11966]|PHTCH179-08|BOLD:AAF4859
Dicranomyia chillcotti|[11967]|PHTCH149-08|BOLD:AAF4859
Dicranomyia chillcotti|[11968]|PHTCH148-08|BOLD:AAF4859
Dicranomyia chillcotti|[11969]|PHTCH293-08|BOLD:AAF4859
Dicranomyia chillcotti|[11970]|JWDCF094-10|BOLD:AAF4859
Dicranomyia chillcotti|[11971]|JWDCF710-10|BOLD:AAF4859
Dicranomyia chillcotti|[11972]|JWDCE926-10|BOLD:AAF4859
Dicranomyia chillcotti|[11973]|JWDCF760-10|BOLD:AAF4859
Dicranomyia chillcotti|[11974]|TICHU123-09|BOLD:AAF4859
Dicranomyia chillcotti|[11975]|JWDCF093-10|BOLD:AAF4859
Dicranomyia chillcotti|[11976]|CNCTI246-12|BOLD:AAF4859
Dicranomyia brevivena|[11977]|JWDCH734-10|BOLD:AAP8636
Dicranomyia brevivena|[11978]|JWDCH958-10|BOLD:AAP8636
Dicranomyia brevivena|[11979]|JWDCI143-10|BOLD:AAP8636
Dicranomyia brevivena|[11980]|JWDCH032-10|BOLD:AAP8636
Dicranomyia brevivena|[11981]|JWDCG356-10|BOLD:AAP8636
Dicranomyia venusta|[11982]|JWDCF043-10|BOLD:AAP8633
Dicranomyia venusta|[11983]|JWDCI266-10|BOLD:AAP8633
Dicranomyia venusta|[11984]|JWDCI307-10|BOLD:AAP8633
Dicranomyia terraenovae|[11985]|CNCTI248-12|BOLD:AAP8633
Dicranomyia terraenovae|[11986]|CNCTI249-12|BOLD:AAP8633
Dicranomyia terraenovae|[11987]|CNCTI250-12|BOLD:AAP8633
Phalacrocera replicata|[11988]|PHTCH358-08|BOLD:AAD9776
Phalacrocera replicata|[11989]|PHTCH356-08|BOLD:AAD9776
Phalacrocera replicata|[11990]|PHTCH357-08|BOLD:AAD9776
Phalacrocera replicata|[11991]|PHTCH359-08|BOLD:AAD9776
Phalacrocera replicata|[11992]|PHTCH385-08|BOLD:AAD9776
Meoneura vagans|[11993]|TWDIP061-09|BOLD:AAG6972
Meoneura vagans|[11994]|TWDIP086-09|BOLD:AAG6972
Meoneura vagans|[11995]|JWDCD415-10|BOLD:AAG6972
Meoneura|[11996]|JWDCD337-10|BOLD:AAM7329
Meoneura tinkerbellae|[11997]|CHDIP516-12|BOLD:ABY2112
Amauromyza shepherdiae|[11998]|TWDIP341-09|BOLD:AAG4752
Amauromyza shepherdiae|[11999]|TWDIP342-09|BOLD:AAG4752
Amauromyza shepherdiae|[12000]|TWDIP343-09|BOLD:AAG4752
Amauromyza shepherdiae|[12001]|CHDIP311-11|BOLD:AAG4752
Amauromyza shepherdiae|[12002]|TWDIP339-09|BOLD:AAG4752
Amauromyza shepherdiae|[12003]|TWDIP340-09|BOLD:AAG4752
Amauromyza shepherdiae|[12004]|CHDIP745-12|BOLD:AAG4752
Culiseta inornata|[12005]|JWDCA308-10|BOLD:AAC9132
Culiseta|[12006]|MHMOS031-07|BOLD:AAD3254
Culiseta morsitans|[12007]|JWDCG243-10|BOLD:AAE3210
Culiseta morsitans|[12008]|JWDCI166-10|BOLD:AAE3210
Culiseta morsitans|[12009]|JWDCG244-10|BOLD:AAE3210
Culiseta morsitans|[12010]|JWDCH560-10|BOLD:AAE3210
Aedes cinereus|[12011]|JWDCL098-11|BOLD:AAP8896
Aedes cinereus|[12012]|JWDCH208-10|BOLD:AAP8896
Aedes cinereus|[12013]|JWDCL449-11|BOLD:AAP8897
Aedes cinereus|[12014]|JWDCH209-10|BOLD:AAP8897
Aedes vexans|[12015]|MHMOS051-07|BOLD:AAA7067
Aedes fitchii|[12016]|JWDCJ1219-11|BOLD:AAC9062
Aedes campestris|[12017]|SAMOS009-09|BOLD:AAD7982
Aedes campestris|[12018]|SAMOS006-09|BOLD:AAD7982
Aedes campestris|[12019]|SAMOS005-09|BOLD:AAD7982
Aedes campestris|[12020]|SAMOS003-09|BOLD:AAD7982
Aedes campestris|[12021]|SAMOS008-09|BOLD:AAD7982
Aedes campestris|[12022]|SAMOS004-09|BOLD:AAD7982
Aedes|[12023]|JWDCJ717-11|BOLD:AAD7982
Aedes rempeli|[12024]|SAMOS027-09|BOLD:AAA6147
Aedes rempeli|[12025]|SAMOS039-09|BOLD:AAA6147
Aedes rempeli|[12026]|SAMOS037-09|BOLD:AAA6147
Aedes rempeli|[12027]|SAMOS038-09|BOLD:AAA6147
Aedes rempeli|[12028]|MHMOS033-07|BOLD:AAA6147
Aedes nigripes|[12029]|JWDCF657-10|BOLD:AAA3750
Aedes nigripes|[12030]|JWDCE725-10|BOLD:AAA3750
Aedes nigripes|[12031]|JWDCF677-10|BOLD:AAA3750
Aedes nigripes|[12032]|JWDCJ445-11|BOLD:AAA3750
Aedes nigripes|[12033]|JWDCF729-10|BOLD:AAA3750
Aedes nigripes|[12034]|JWDCF104-10|BOLD:AAA3750
Aedes nigripes|[12035]|SAMOS026-09|BOLD:AAA3750
Aedes nigripes|[12036]|SAMOS029-09|BOLD:AAA3750
Aedes nigripes|[12037]|JWDCC841-10|BOLD:AAA3750
Aedes nigripes|[12038]|JWDCJ761-11|BOLD:AAA3750
Aedes nigripes|[12039]|JWDCF105-10|BOLD:AAA3750
Aedes nigripes|[12040]|SAMOS028-09|BOLD:AAA3750
[illegible]

Aedes nigripes|[12038]|JWDCJ761-11|BOLD:AAA3750
Aedes nigripes|[12039]|JWDCF105-10|BOLD:AAA3750
Aedes nigripes|[12040]|SAMOS028-09|BOLD:AAA3750
Aedes nigripes|[12041]|SAMOS044-09|BOLD:AAA3750
Aedes nigripes|[12042]|SAMOS020-09|BOLD:AAA3750
Aedes nigripes|[12043]|SAMOS047-09|BOLD:AAA3750
Aedes nigripes|[12044]|SAMOS045-09|BOLD:AAA3750
Aedes nigripes|[12045]|SAMOS046-09|BOLD:AAA3750
Aedes nigripes|[12046]|SAMOS030-09|BOLD:AAA3750
Aedes nigripes|[12047]|JWDCJ446-11|BOLD:AAA3750
Aedes nigripes|[12048]|JWDCJ760-11|BOLD:AAA3750
Aedes nigripes|[12049]|JWDCI765-11|BOLD:AAA3750
Aedes nigripes|[12050]|JWDCF076-10|BOLD:AAA3750
Aedes nigripes|[12051]|JWDCE659-10|BOLD:AAA3750
Aedes nigripes|[12052]|JWDCE702-10|BOLD:AAA3750
Aedes nigripes|[12053]|JWDCF683-10|BOLD:AAA3750
Aedes nigripes|[12054]|JWDCF757-10|BOLD:AAA3750
Aedes nigripes|[12055]|JWDCE171-10|BOLD:AAA3750
Aedes nigripes|[12056]|JWDCE172-10|BOLD:AAA3750
Aedes nigripes|[12057]|SAMOS025-09|BOLD:AAA3750
Aedes nigripes|[12058]|JWDCE701-10|BOLD:AAA3750
Aedes nigripes|[12059]|JWDCJ932-11|BOLD:AAA3750
Aedes nigripes|[12060]|JWDCJ971-11|BOLD:AAA3750
Aedes nigripes|[12061]|JWDCE145-10|BOLD:AAA3750
Aedes nigripes|[12062]|SAMOS022-09|BOLD:AAA3750
Aedes nigripes|[12063]|JWDCE173-10|BOLD:AAA3750
Aedes impiger|[12064]|JWDCJ762-11|BOLD:AAA3750
Aedes impiger|[12065]|JWDCJ151-11|BOLD:AAA3750
Aedes impiger|[12066]|JWDCJ447-11|BOLD:AAA3750
Aedes impiger|[12067]|JWDCE106-10|BOLD:AAA3750
Aedes impiger|[12068]|JWDCE332-10|BOLD:AAA3750
Aedes impiger|[12069]|JWDCE417-10|BOLD:AAA3750
Aedes impiger|[12070]|JWDCE104-10|BOLD:AAA3750
Aedes impiger|[12071]|JWDCJ418-11|BOLD:AAA3750
Aedes impiger|[12072]|JWDCE463-10|BOLD:AAA3750
Aedes impiger|[12073]|JWDCJ406-11|BOLD:AAA3750
Aedes impiger|[12074]|JWDCI263-10|BOLD:AAA3750
Aedes impiger|[12075]|JWDCE119-10|BOLD:AAA3750
Aedes impiger|[12076]|JWDCJ419-11|BOLD:AAA3750
Aedes impiger|[12077]|JWDCE187-10|BOLD:AAA3750
Aedes impiger|[12078]|JWDCE333-10|BOLD:AAA3750
Aedes impiger|[12079]|JWDCE334-10|BOLD:AAA3750
Aedes impiger|[12080]|JWDCE188-10|BOLD:AAA3750
Aedes impiger|[12081]|JWDCB145-10|BOLD:AAA3750
Aedes pionips|[12082]|JWDCJ1221-11|BOLD:AAB6338
Aedes pionips|[12083]|JWDCG750-10|BOLD:AAB6338
Aedes pionips|[12084]|JWDCK158-11|BOLD:AAB6338
Aedes pionips|[12085]|MMADL012-09|BOLD:AAB6338
Aedes pionips|[12086]|JWDCH559-10|BOLD:AAB6338
Aedes pionips|[12087]|JWDCK098-11|BOLD:AAB6338
Aedes pionips|[12088]|JWDCI047-10|BOLD:AAB6338
Aedes pionips|[12089]|JWDCH045-10|BOLD:AAB6338
Aedes pionips|[12090]|JWDCH092-10|BOLD:AAB6338
Aedes pionips|[12091]|JWDCL174-11|BOLD:AAB6338
Aedes pionips|[12092]|JWDCI046-10|BOLD:AAB6338
Aedes pionips|[12093]|JWDCL362-11|BOLD:AAB6338
Aedes pionips|[12094]|JWDCK159-11|BOLD:AAB6338
Aedes pionips|[12095]|JWDCJ364-11|BOLD:AAB6338
Aedes pionips|[12096]|MMADL015-09|BOLD:AAB6338
Aedes pionips|[12097]|MMADL014-09|BOLD:AAB6338
Aedes pionips|[12098]|MMADL013-09|BOLD:AAB6338
Aedes pionips|[12099]|MMADL011-09|BOLD:AAB6338
Aedes pionips|[12100]|MMADL010-09|BOLD:AAB6338
Aedes pionips|[12101]|MMADL009-09|BOLD:AAB6338
Aedes pionips|[12102]|JWDCH090-10|BOLD:AAB6338
Aedes pionips|[12103]|JWDCG214-10|BOLD:AAB6338
Aedes pionips|[12104]|JWDCB151-10|BOLD:AAB6338
Aedes pionips|[12105]|JWDCB154-10|BOLD:AAB6338
Aedes communis complex|[12106]|MHMOS029-07|BOLD:AAA3751
Aedes communis complex|[12107]|SAMOS040-09|BOLD:AAA3751
Aedes communis complex|[12108]|SAMOS062-09|BOLD:AAA3751
Aedes communis complex|[12109]|SAMOS042-09|BOLD:AAA3751
Aedes communis complex|[12110]|JWDCJ600-11|BOLD:AAA3751
Aedes communis complex|[12111]|MHMOS032-07|BOLD:AAA3751
Aedes communis complex|[12112]|JWDCI262-10|BOLD:AAA3751
Aedes communis complex|[12113]|JWDCJ451-11|BOLD:AAA3751
Aedes communis complex|[12114]|SAMOS069-09|BOLD:AAA3751
Aedes communis complex|[12115]|SAMOS041-09|BOLD:AAA3751
Aedes communis complex|[12116]|MHMOS022-07|BOLD:AAA3751
Aedes communis complex|[12117]|MHMOS028-07|BOLD:AAA3751
Aedes communis complex|[12118]|JWDCJ453-11|BOLD:AAA3751
Aedes communis complex|[12119]|JWDCJ599-11|BOLD:AAA3751
Aedes communis complex|[12120]|SAMOS061-09|BOLD:AAA3751
Aedes communis complex|[12121]|JWDCI264-10|BOLD:AAA3751
Aedes communis complex|[12122]|JWDCE418-10|BOLD:AAA3751
Aedes communis complex|[12123]|JWDCE432-10|BOLD:AAA3751
Aedes communis complex|[12124]|JWDCE433-10|BOLD:AAA3751
Aedes communis|[12125]|JWDCI940-11|BOLD:AAA6148
Aedes communis|[12126]|MHMOS083-07|BOLD:AAA6148
Aedes communis|[12127]|SWSWP043-09|BOLD:AAA6148
Aedes communis|[12128]|SWSWP042-09|BOLD:AAA6148
Aedes communis|[12129]|JWDCJ452-11|BOLD:AAA6148
Aedes communis|[12130]|JWDCF849-10|BOLD:AAA6148
Aedes communis|[12131]|JWDCE924-10|BOLD:AAA6148
Aedes|[12132]|JWDCA317-10|BOLD:AAA3748
Aedes|[12133]|JWDCF139-10|BOLD:AAA3748
Aedes|[12134]|SAMOS034-09|BOLD:AAA3748
Aedes|[12135]|JWDCF370-10|BOLD:AAA3748
Aedes|[12136]|JWDCC137-10|BOLD:AAA3748
Aedes|[12137]|SAMOS059-09|BOLD:AAA3748
Aedes abserratus|[12138]|JWDCC054-10|BOLD:AAA3748
Aedes abserratus|[12139]|MHMOS016-07|BOLD:AAA3748
Aedes abserratus|[12140]|JWDCG511-10|BOLD:AAA3748

Aedes abserratus|[12138]|JWDCC054-10|BOLD:AAA3748
Aedes abserratus|[12139]|MHMOS016-07|BOLD:AAA3748
Aedes abserratus|[12140]|JWDCG511-10|BOLD:AAA3748
Aedes abserratus|[12141]|AMMC013-07|BOLD:AAA3748
Aedes abserratus|[12142]|JWDCI468-10|BOLD:AAA3748
Aedes abserratus|[12143]|JWDCG752-10|BOLD:AAA3748
Aedes abserratus|[12144]|MHMOS080-07|BOLD:AAA3748
Aedes abserratus|[12145]|JWDCA313-10|BOLD:AAA3748
Aedes abserratus|[12146]|MHMOS036-07|BOLD:AAA3748
Aedes abserratus|[12147]|JWDCC204-10|BOLD:AAA3748
Aedes abserratus|[12148]|JWDCF137-10|BOLD:AAA3748
Aedes abserratus|[12149]|JWDCF511-10|BOLD:AAA3748
Aedes abserratus|[12150]|MHMOS084-07|BOLD:AAA3748
Aedes abserratus|[12151]|JWDCI668-11|BOLD:AAA3748
Aedes abserratus|[12152]|JWDCH009-10|BOLD:AAA3748
Aedes abserratus|[12153]|JWDCI373-10|BOLD:AAA3748
Aedes abserratus|[12154]|JWDCI282-10|BOLD:AAA3748
Aedes abserratus|[12155]|JWDCI273-10|BOLD:AAA3748
Aedes abserratus|[12156]|JWDCH889-10|BOLD:AAA3748
Aedes abserratus|[12157]|JWDCH864-10|BOLD:AAA3748
Aedes abserratus|[12158]|JWDCG265-10|BOLD:AAA3748
Aedes abserratus|[12159]|JWDCG751-10|BOLD:AAA3748
Aedes abserratus|[12160]|JWDCH736-10|BOLD:AAA3748
Aedes abserratus|[12161]|JWDCG696-10|BOLD:AAA3748
Aedes abserratus|[12162]|JWDCJ048-11|BOLD:AAA3748
Aedes abserratus|[12163]|JWDCJ1519-11|BOLD:AAA3748
Aedes abserratus|[12164]|JWDCJ1367-11|BOLD:AAA3748
Aedes abserratus|[12165]|JWDCC473-10|BOLD:AAA3748
Aedes abserratus|[12166]|JWDCC472-10|BOLD:AAA3748
Aedes abserratus|[12167]|JWDCC270-10|BOLD:AAA3748
Aedes abserratus|[12168]|JWDCC269-10|BOLD:AAA3748
Aedes abserratus|[12169]|SWSWP077-09|BOLD:AAA3748
Aedes abserratus|[12170]|JWDCD101-10|BOLD:AAA3748
Aedes abserratus|[12171]|MHMOS027-07|BOLD:AAA3748
Aedes abserratus|[12172]|MHMOS021-07|BOLD:AAA3748
Aedes abserratus|[12173]|MHMOS086-07|BOLD:AAA3748
Aedes abserratus|[12174]|MHMOS085-07|BOLD:AAA3748
Aedes abserratus|[12175]|MHMOS072-07|BOLD:AAA3748
Aedes abserratus|[12176]|MHMOS070-07|BOLD:AAA3748
Aedes abserratus|[12177]|JWDCA307-10|BOLD:AAA3748
Aedes abserratus|[12178]|JWDCC072-10|BOLD:AAA3748
Aedes abserratus|[12179]|SWSWP049-09|BOLD:AAA3748
Aedes abserratus|[12180]|AMMC016-07|BOLD:AAA3748
Aedes abserratus|[12181]|AMMC031-07|BOLD:AAA3748
Aedes abserratus|[12182]|SAMOS054-09|BOLD:AAA3748
Aedes abserratus|[12183]|SAMOS067-09|BOLD:AAA3748
Aedes abserratus|[12184]|SAMOS051-09|BOLD:AAA3748
Aedes abserratus|[12185]|AMMC077-07|BOLD:AAA3748
Aedes abserratus|[12186]|AMMC070-07|BOLD:AAA3748
Aedes abserratus|[12187]|AMMC003-07|BOLD:AAA3748
Aedes abserratus|[12188]|SAMOS032-09|BOLD:AAA3748
Aedes abserratus|[12189]|JWDCC180-10|BOLD:AAA3748
Aedes abserratus|[12190]|AMMC084-07|BOLD:AAA3748
Aedes abserratus|[12191]|SAMOS052-09|BOLD:AAA3748
Aedes abserratus|[12192]|JWDCC055-10|BOLD:AAA3748
Aedes abserratus|[12193]|MHMOS082-07|BOLD:AAA3748
Aedes abserratus|[12194]|SWSWP010-09|BOLD:AAA3748
Aedes abserratus|[12195]|AMMC094-07|BOLD:AAA3748
Aedes abserratus|[12196]|JWDCC903-10|BOLD:AAA3748
Aedes abserratus|[12197]|JWDCJ585-11|BOLD:AAA3748
Aedes abserratus|[12198]|JWDCI009-10|BOLD:AAA3748
Aedes abserratus|[12199]|JWDCG308-10|BOLD:AAA3748
Aedes abserratus|[12200]|JWDCG309-10|BOLD:AAA3748
Aedes abserratus|[12201]|JWDCH439-10|BOLD:AAA3748
Aedes abserratus|[12202]|JWDCG463-10|BOLD:AAA3748
Aedes abserratus|[12203]|JWDCG510-10|BOLD:AAA3748
Aedes abserratus|[12204]|JWDCF138-10|BOLD:AAA3748
Aedes abserratus|[12205]|JWDCF171-10|BOLD:AAA3748
Aedes abserratus|[12206]|JWDCE609-10|BOLD:AAA3748
Aedes abserratus|[12207]|JWDCE717-10|BOLD:AAA3748
Aedes abserratus|[12208]|JWDCE719-10|BOLD:AAA3748
Aedes abserratus|[12209]|JWDCE727-10|BOLD:AAA3748
Aedes abserratus|[12210]|JWDCF513-10|BOLD:AAA3748
Aedes abserratus|[12211]|JWDCE852-10|BOLD:AAA3748
Aedes abserratus|[12212]|JWDCF645-10|BOLD:AAA3748
Aedes abserratus|[12213]|JWDCE608-10|BOLD:AAA3748
Aedes abserratus|[12214]|JWDCB146-10|BOLD:AAA3748
Aedes|[12215]|JWDCC434-10|BOLD:AAA3748
Aedes|[12216]|JWDCJ1002-11|BOLD:AAA3748
Aedes|[12217]|JWDCH966-10|BOLD:AAA3748
Aedes|[12218]|AMMC088-07|BOLD:AAA3748
Aedes|[12219]|AMMC014-07|BOLD:AAA3748
Aedes|[12220]|AMMC054-07|BOLD:AAA3748
Aedes|[12221]|JWDCA312-10|BOLD:AAA3748
Aedes|[12222]|SWSWP082-09|BOLD:AAA3748
Aedes|[12223]|AMMC025-07|BOLD:AAA3748
Aedes|[12224]|JWDCG695-10|BOLD:AAA3748
Aedes|[12225]|JWDCH008-10|BOLD:AAA3748
Aedes|[12226]|SWSWP063-09|BOLD:AAA3748
Aedes hexodontus|[12227]|JWDCD178-10|BOLD:AAA3748
Aedes|[12228]|AMMC093-07|BOLD:AAA3748
Aedes|[12229]|AMMC081-07|BOLD:AAA3748
Aedes|[12230]|SAMOS068-09|BOLD:AAA3748
Aedes|[12231]|SWSWP011-09|BOLD:AAA3748
Aedes|[12232]|MHMOS067-07|BOLD:AAA3748
Aedes|[12233]|JWDCC475-10|BOLD:AAA3748
Aedes|[12234]|SAMOS023-09|BOLD:AAA3748
Aedes|[12235]|JWDCF060-10|BOLD:AAA3748
Aedes|[12236]|JWDCI856-11|BOLD:AAA3748
Aedes|[12237]|AMMC076-07|BOLD:AAA3748
Aedes|[12238]|JWDCE742-10|BOLD:AAA3748
Aedes|[12239]|JWDCF602-10|BOLD:AAA3748
[illegible]

Aedes|[12237]|AMMC070-07|BOLD:AAA3748
Aedes|[12238]|JWDCE742-10|BOLD:AAA3748
Aedes|[12239]|JWDCF602-10|BOLD:AAA3748
Aedes|[12240]|JWDCF603-10|BOLD:AAA3748
Aedes hexodontus|[12241]|JWDCC228-10|BOLD:AAA3748
Aedes hexodontus|[12242]|SWSWP019-09|BOLD:AAA3748
Aedes hexodontus|[12243]|JWDCC433-10|BOLD:AAA3748
Aedes hexodontus|[12244]|JWDCI383-10|BOLD:AAA3748
Aedes hexodontus|[12245]|SAMOS050-09|BOLD:AAA3748
Aedes hexodontus|[12246]|JWDCC474-10|BOLD:AAA3748
Aedes hexodontus|[12247]|AMMC001-07|BOLD:AAA3748
Aedes hexodontus|[12248]|JWDCI384-10|BOLD:AAA3748
Aedes hexodontus|[12249]|AMMC087-07|BOLD:AAA3748
Aedes hexodontus|[12250]|JWDCC225-10|BOLD:AAA3748
Aedes hexodontus|[12251]|MHMOS089-07|BOLD:AAA3748
Aedes hexodontus|[12252]|JWDCF432-10|BOLD:AAA3748
Aedes hexodontus|[12253]|SAMOS057-09|BOLD:AAA3748
Aedes hexodontus|[12254]|JWDCC135-10|BOLD:AAA3748
Aedes hexodontus|[12255]|JWDCJ417-11|BOLD:AAA3748
Aedes hexodontus|[12256]|AMMC085-07|BOLD:AAA3748
Aedes hexodontus|[12257]|AMMC092-07|BOLD:AAA3748
Aedes hexodontus|[12258]|AMMC022-07|BOLD:AAA3748
Aedes hexodontus|[12259]|JWDCH890-10|BOLD:AAA3748
Aedes hexodontus|[12260]|MHMOS075-07|BOLD:AAA3748
Aedes hexodontus|[12261]|JWDCF444-10|BOLD:AAA3748
Aedes hexodontus|[12262]|SAMOS036-09|BOLD:AAA3748
Aedes hexodontus|[12263]|JWDCE904-10|BOLD:AAA3748
Aedes hexodontus|[12264]|JWDCI810-11|BOLD:AAA3748
Aedes hexodontus|[12265]|MHFLY115-07|BOLD:AAA3748
Aedes hexodontus|[12266]|SWSWP040-09|BOLD:AAA3748
Aedes hexodontus|[12267]|SWSWP041-09|BOLD:AAA3748
Aedes hexodontus|[12268]|JWDCF573-10|BOLD:AAA3748
Aedes hexodontus|[12269]|JWDCD096-10|BOLD:AAA3748
Aedes hexodontus|[12270]|SAMOS049-09|BOLD:AAA3748
Aedes hexodontus|[12271]|JWDCC445-10|BOLD:AAA3748
Aedes hexodontus|[12272]|JWDCC109-10|BOLD:AAA3748
Aedes hexodontus|[12273]|JWDCF001-10|BOLD:AAA3748
Aedes hexodontus|[12274]|JWDCI311-10|BOLD:AAA3748
Aedes hexodontus|[12275]|JWDCF486-10|BOLD:AAA3748
Aedes hexodontus|[12276]|JWDCC902-10|BOLD:AAA3748
Aedes hexodontus|[12277]|AMMC069-07|BOLD:AAA3748
Aedes hexodontus|[12278]|AMMC047-07|BOLD:AAA3748
Aedes hexodontus|[12279]|JWDCA314-10|BOLD:AAA3748
Aedes hexodontus|[12280]|SWSWP029-09|BOLD:AAA3748
Aedes hexodontus|[12281]|JWDCA311-10|BOLD:AAA3748
Aedes hexodontus|[12282]|AMMC049-07|BOLD:AAA3748
Aedes hexodontus|[12283]|AMMC015-07|BOLD:AAA3748
Aedes hexodontus|[12284]|AMMC052-07|BOLD:AAA3748
Aedes hexodontus|[12285]|AMMC041-07|BOLD:AAA3748
Aedes|[12286]|JWDCF571-10|BOLD:AAA3748
Aedes hexodontus|[12287]|AMMC020-07|BOLD:AAA3748
Aedes hexodontus|[12288]|SAMOS053-09|BOLD:AAA3748
Aedes hexodontus|[12289]|AMMC027-07|BOLD:AAA3748
Aedes hexodontus|[12290]|AMMC075-07|BOLD:AAA3748
Aedes hexodontus|[12291]|AMMC064-07|BOLD:AAA3748
Aedes hexodontus|[12292]|JWDCC132-10|BOLD:AAA3748
Aedes hexodontus|[12293]|JWDCJ219-11|BOLD:AAA3748
Aedes hexodontus|[12294]|JWDCI584-10|BOLD:AAA3748
Aedes hexodontus|[12295]|JWDCC842-10|BOLD:AAA3748
Aedes hexodontus|[12296]|JWDCC205-10|BOLD:AAA3748
Aedes hexodontus|[12297]|SWSWP054-09|BOLD:AAA3748
Aedes hexodontus|[12298]|AMMC073-07|BOLD:AAA3748
Aedes hexodontus|[12299]|AMMC023-07|BOLD:AAA3748
Aedes hexodontus|[12300]|JWDCC450-10|BOLD:AAA3748
Aedes hexodontus|[12301]|JWDCF170-10|BOLD:AAA3748
Aedes hexodontus|[12302]|JWDCF601-10|BOLD:AAA3748
Aedes hexodontus|[12303]|JWDCA318-10|BOLD:AAA3748
Aedes hexodontus|[12304]|MHMOS074-07|BOLD:AAA3748
Aedes hexodontus|[12305]|JWDCC136-10|BOLD:AAA3748
Aedes hexodontus|[12306]|JWDCD103-10|BOLD:AAA3748
Aedes hexodontus|[12307]|JWDCD105-10|BOLD:AAA3748
Aedes hexodontus|[12308]|JWDCC203-10|BOLD:AAA3748
Aedes hexodontus|[12309]|JWDCC267-10|BOLD:AAA3748
Aedes hexodontus|[12310]|JWDCJ686-11|BOLD:AAA3748
Aedes hexodontus|[12311]|JWDCJ715-11|BOLD:AAA3748
Aedes hexodontus|[12312]|JWDCJ822-11|BOLD:AAA3748
Aedes hexodontus|[12313]|JWDCJ1574-11|BOLD:AAA3748
Aedes hexodontus|[12314]|JWDCG671-10|BOLD:AAA3748
Aedes hexodontus|[12315]|JWDCI670-11|BOLD:AAA3748
Aedes hexodontus|[12316]|JWDCI325-10|BOLD:AAA3748
Aedes hexodontus|[12317]|JWDCI351-10|BOLD:AAA3748
Aedes hexodontus|[12318]|JWDCI714-11|BOLD:AAA3748
Aedes hexodontus|[12319]|JWDCI715-11|BOLD:AAA3748
Aedes hexodontus|[12320]|JWDCF224-10|BOLD:AAA3748
Aedes hexodontus|[12321]|JWDCE610-10|BOLD:AAA3748
Aedes hexodontus|[12322]|JWDCE744-10|BOLD:AAA3748
Aedes hexodontus|[12323]|JWDCE782-10|BOLD:AAA3748
Aedes hexodontus|[12324]|JWDCF535-10|BOLD:AAA3748
Aedes hexodontus|[12325]|JWDCF558-10|BOLD:AAA3748
Aedes hexodontus|[12326]|JWDCF627-10|BOLD:AAA3748
Aedes hexodontus|[12327]|JWDCF646-10|BOLD:AAA3748
Aedes hexodontus|[12328]|JWDCF875-10|BOLD:AAA3748
Aedes hexodontus|[12329]|JWDCE905-10|BOLD:AAA3748
Aedes hexodontus|[12330]|JWDCF002-10|BOLD:AAA3748
Aedes hexodontus|[12331]|MHFLY118-07|BOLD:AAA3748
Aedes hexodontus|[12332]|MHFLY117-07|BOLD:AAA3748
Aedes|[12333]|JWDCI613-10|BOLD:AAA3748
Aedes|[12334]|JWDCC456-10|BOLD:AAA3748
Aedes|[12335]|JWDCC113-10|BOLD:AAA3748
Aedes|[12336]|SAMOS021-09|BOLD:AAA3748
Aedes|[12337]|JWDCH047-10|BOLD:AAA3748
Aedes|[12338]|AMMC050-07|BOLD:AAA3748
Aedes|[12339]|SWSWP055-09|BOLD:AAA3748

Aedes|[12337]|JWDCH047-10|BOLD:AAA3748
Aedes|[12338]|AMMC050-07|BOLD:AAA3748
Aedes|[12339]|SWSWP055-09|BOLD:AAA3748
Aedes|[12340]|JWDCF062-10|BOLD:AAA3748
Aedes|[12341]|JWDCE781-10|BOLD:AAA3748
Aedes|[12342]|JWDCE903-10|BOLD:AAA3748
Aedes|[12343]|AMMC078-07|BOLD:AAA3748
Aedes|[12344]|SWSWP036-09|BOLD:AAA3748
Aedes|[12345]|JWDCC226-10|BOLD:AAA3748
Aedes|[12346]|JWDCI048-10|BOLD:AAA3748
Aedes|[12347]|JWDCI375-10|BOLD:AAA3748
Aedes|[12348]|SWSWP048-09|BOLD:AAA3748
Aedes|[12349]|JWDCC454-10|BOLD:AAA3748
Aedes|[12350]|JWDCC471-10|BOLD:AAA3748
Aedes|[12351]|AMMC011-07|BOLD:AAA3748
Aedes|[12352]|SWSWP027-09|BOLD:AAA3748
Aedes|[12353]|JWDCF534-10|BOLD:AAA3748
Aedes|[12354]|JWDCF466-10|BOLD:AAA3748
Aedes|[12355]|AMMC071-07|BOLD:AAA3748
Aedes|[12356]|JWDCC112-10|BOLD:AAA3748
Aedes|[12357]|JWDCH967-10|BOLD:AAA3748
Aedes|[12358]|JWDCJ687-11|BOLD:AAA3748
Aedes|[12359]|JWDCJ823-11|BOLD:AAA3748
Aedes|[12360]|SAMOS024-09|BOLD:AAA3748
Aedes|[12361]|AMMC044-07|BOLD:AAA3748
Aedes|[12362]|SWSWP028-09|BOLD:AAA3748
Aedes|[12363]|JWDCF488-10|BOLD:AAA3748
Aedes|[12364]|AMMC017-07|BOLD:AAA3748
Aedes|[12365]|JWDCC110-10|BOLD:AAA3748
Aedes|[12366]|JWDCF382-10|BOLD:AAA3748
Aedes|[12367]|JWDCF583-10|BOLD:AAA3748
Aedes|[12368]|JWDCE853-10|BOLD:AAA3748
Aedes|[12369]|JWDCF172-10|BOLD:AAA3748
Aedes|[12370]|AMMC032-07|BOLD:AAA3748
Aedes|[12371]|SWSWP076-09|BOLD:AAA3748
Aedes|[12372]|JWDCC469-10|BOLD:AAA3748
Aedes|[12373]|AMMC057-07|BOLD:AAA3748
Aedes|[12374]|JWDCC111-10|BOLD:AAA3748
Aedes|[12375]|MHMOS015-07|BOLD:AAA3748
Aedes|[12376]|SWSWP026-09|BOLD:AAA3748
Aedes|[12377]|AMMC010-07|BOLD:AAA3748
Aedes|[12378]|SAMOS048-09|BOLD:AAA3748
Aedes|[12379]|AMMC063-07|BOLD:AAA3748
Aedes|[12380]|AMMC079-07|BOLD:AAA3748
Aedes|[12381]|JWDCE928-10|BOLD:AAA3748
Aedes|[12382]|SAMOS060-09|BOLD:AAA3748
Aedes|[12383]|JWDCC462-10|BOLD:AAA3748
Aedes|[12384]|JWDCJ884-11|BOLD:AAA3748
Aedes|[12385]|JWDCE794-10|BOLD:AAA3748
Aedes|[12386]|MHMOS026-07|BOLD:AAA3748
Aedes|[12387]|JWDCI469-10|BOLD:AAA3748
Aedes|[12388]|JWDCJ150-11|BOLD:AAA3748
Aedes|[12389]|JWDCJ408-11|BOLD:AAA3748
Aedes|[12390]|JWDCE922-10|BOLD:AAA3748
Aedes|[12391]|JWDCF713-10|BOLD:AAA3748
Aedes|[12392]|JWDCF678-10|BOLD:AAA3748
Aedes|[12393]|JWDCF445-10|BOLD:AAA3748
Aedes|[12394]|JWDCF226-10|BOLD:AAA3748
Aedes|[12395]|JWDCI310-10|BOLD:AAA3748
Aedes|[12396]|JWDCJ821-11|BOLD:AAA3748
Aedes|[12397]|JWDCJ716-11|BOLD:AAA3748
Aedes|[12398]|JWDCC464-10|BOLD:AAA3748
Aedes|[12399]|JWDCC444-10|BOLD:AAA3748
Aedes|[12400]|SWSWP072-09|BOLD:AAA3748
Aedes|[12401]|SWSWP067-09|BOLD:AAA3748
Aedes|[12402]|MHMOS081-07|BOLD:AAA3748
Aedes|[12403]|SWSWP013-09|BOLD:AAA3748
Aedes|[12404]|JWDCC056-10|BOLD:AAA3748
Aedes|[12405]|AMMC009-07|BOLD:AAA3748
Aedes|[12406]|SAMOS031-09|BOLD:AAA3748
Aedes|[12407]|AMMC053-07|BOLD:AAA3748
Aedes|[12408]|AMMC021-07|BOLD:AAA3748
Aedes|[12409]|AMMC061-07|BOLD:AAA3748
Aedes|[12410]|AMMC039-07|BOLD:AAA3748
Aedes|[12411]|AMMC035-07|BOLD:AAA3748
Aedes|[12412]|SAMOS055-09|BOLD:AAA3748
Aedes|[12413]|AMMC030-07|BOLD:AAA3748
Aedes|[12414]|AMMC074-07|BOLD:AAA3748
Aedes|[12415]|AMMC033-07|BOLD:AAA3748
Aedes|[12416]|MHMOS014-07|BOLD:AAA3748
Aedes|[12417]|JWDCI716-11|BOLD:AAA3748
Aedes|[12418]|AMMC060-07|BOLD:AAA3748
Aedes|[12419]|JWDCI326-10|BOLD:AAA3748
Aedes|[12420]|SWSWP065-09|BOLD:AAA3748
Aedes|[12421]|SWSWP059-09|BOLD:AAA3748
Aedes|[12422]|JWDCI327-10|BOLD:AAA3748
Aedes|[12423]|JWDCD001-10|BOLD:AAA3748
Aedes|[12424]|JWDCE528-10|BOLD:AAA3748
Aedes|[12425]|JWDCF335-10|BOLD:AAA3748
Aedes|[12426]|JWDCF644-10|BOLD:AAA3748
Aedes|[12427]|JWDCB144-10|BOLD:AAA3748
Aedes|[12428]|AMMC080-07|BOLD:AAA3748
Aedes|[12429]|SWSWP058-09|BOLD:AAA3748
Aedes|[12430]|SWSWP056-09|BOLD:AAA3748
Aedes|[12431]|JWDCJ363-11|BOLD:AAA3748
Aedes|[12432]|JWDCJ362-11|BOLD:AAA3748
Aedes|[12433]|JWDCB148-10|BOLD:AAA3748
Aedes|[12434]|MHFLY116-07|BOLD:AAA3748
Aedes|[12435]|MHMOS088-07|BOLD:AAA3748
Aedes|[12436]|JWDCF572-10|BOLD:AAA3748
Aedes|[12437]|JWDCI812-11|BOLD:AAA3748
Aedes|[12438]|JWDCJ1520-11|BOLD:AAA3748
Aedes|[12439]|JWDCC446-10|BOLD:AAA3748

Aedes|[12436]|JWDCF372-10|BOLD:AAA3748
Aedes|[12437]|JWDCI812-11|BOLD:AAA3748
Aedes|[12438]|JWDCJ1520-11|BOLD:AAA3748
Aedes|[12439]|JWDCC446-10|BOLD:AAA3748
Aedes|[12440]|JWDCH738-10|BOLD:AAA3748
Aedes|[12441]|JWDCC181-10|BOLD:AAA3748
Aedes|[12442]|JWDCE795-10|BOLD:AAA3748
Aedes|[12443]|JWDCL363-11|BOLD:AAA3748
Aedes|[12444]|JWDCC224-10|BOLD:AAA3748
Aedes|[12445]|JWDCE649-10|BOLD:AAA3748
Aedes|[12446]|JWDCJ688-11|BOLD:AAA3748
Aedes|[12447]|JWDCF676-10|BOLD:AAA3748
Aedes|[12448]|JWDCD097-10|BOLD:AAA3748
Aedes|[12449]|JWDCI583-10|BOLD:AAA3748
Aedes|[12450]|JWDCF489-10|BOLD:AAA3748
Aedes|[12451]|JWDCF585-10|BOLD:AAA3748
Aedes|[12452]|AMMC034-07|BOLD:AAA3748
Aedes|[12453]|JWDCJ648-11|BOLD:AAA3748
Aedes|[12454]|JWDCE651-10|BOLD:AAA3748
Aedes|[12455]|SAMOS018-09|BOLD:AAA3748
Aedes|[12456]|JWDCC268-10|BOLD:AAA3748
Aedes|[12457]|JWDCL361-11|BOLD:AAA3748
Aedes|[12458]|JWDCG126-10|BOLD:AAA3748
Aedes|[12459]|JWDCF536-10|BOLD:AAA3748
Aedes|[12460]|JWDCJ970-11|BOLD:AAA3748
Aedes|[12461]|JWDCH263-10|BOLD:AAA3748
Aedes|[12462]|JWDCE799-10|BOLD:AAA3748
Aedes|[12463]|JWDCC179-10|BOLD:AAA3748
Aedes|[12464]|JWDCI275-10|BOLD:AAA3748
Aedes|[12465]|JWDCC901-10|BOLD:AAA3748
Aedes|[12466]|JWDCC432-10|BOLD:AAA3748
Aedes|[12467]|AMMC046-07|BOLD:AAA3748
Aedes|[12468]|AMMC062-07|BOLD:AAA3748
Aedes|[12469]|JWDCF512-10|BOLD:AAA3748
Aedes|[12470]|JWDCE831-10|BOLD:AAA3748
Aedes|[12471]|MHMOS071-07|BOLD:AAA3748
Aedes|[12472]|MHMOS037-07|BOLD:AAA3748
Aedes|[12473]|JWDCH815-10|BOLD:AAA3748
Aedes|[12474]|JWDCF626-10|BOLD:AAA3748
Aedes|[12475]|AMMC065-07|BOLD:AAA3748
Aedes|[12476]|AMMC086-07|BOLD:AAA3748
Aedes|[12477]|JWDCC455-10|BOLD:AAA3748
Aedes|[12478]|JWDCD177-10|BOLD:AAA3748
Aedes|[12479]|JWDCE832-10|BOLD:AAA3748
Aedes|[12480]|SWSWP075-09|BOLD:AAA3748
Aedes|[12481]|JWDCE866-10|BOLD:AAA3748
Aedes|[12482]|AMMC059-07|BOLD:AAA3748
Aedes|[12483]|JWDCC780-10|BOLD:AAA3748
Aedes|[12484]|JWDCJ276-11|BOLD:AAA3748
Aedes|[12485]|AMMC008-07|BOLD:AAA3748
Aedes|[12486]|JWDCD106-10|BOLD:AAA3748
Aedes|[12487]|JWDCJ1301-11|BOLD:AAA3748
Aedes|[12488]|AMMC004-07|BOLD:AAA3748
Aedes|[12489]|AMMC012-07|BOLD:AAA3748
Aedes|[12490]|JWDCH390-10|BOLD:AAA3748
Aedes|[12491]|JWDCE793-10|BOLD:AAA3748
Aedes|[12492]|AMMC018-07|BOLD:AAA3748
Aedes|[12493]|AMMC040-07|BOLD:AAA3748
Aedes|[12494]|JWDCC463-10|BOLD:AAA3748
Aedes|[12495]|JWDCF061-10|BOLD:AAA3748
Aedes|[12496]|JWDCF431-10|BOLD:AAA3748
Aedes|[12497]|JWDCC134-10|BOLD:AAA3748
Aedes|[12498]|JWDCF625-10|BOLD:AAA3748
Aedes|[12499]|JWDCJ277-11|BOLD:AAA3748
Aedes|[12500]|JWDCF487-10|BOLD:AAA3748
Aedes|[12501]|JWDCF491-10|BOLD:AAA3748
Aedes|[12502]|SAMOS058-09|BOLD:AAA3748
Aedes|[12503]|SAMOS056-09|BOLD:AAA3748
Aedes|[12504]|JWDCF225-10|BOLD:AAA3748
Aedes|[12505]|JWDCE798-10|BOLD:AAA3748
Aedes|[12506]|JWDCF584-10|BOLD:AAA3748
Aedes|[12507]|AMMC056-07|BOLD:AAA3748
Aedes|[12508]|SAMOS017-09|BOLD:AAA3748
Aedes|[12509]|AMMC024-07|BOLD:AAA3748
Aedes|[12510]|JWDCC070-10|BOLD:AAA3748
Aedes|[12511]|MHMOS077-07|BOLD:AAA3748
Aedes|[12512]|JWDCE743-10|BOLD:AAA3748
Aedes|[12513]|JWDCE800-10|BOLD:AAA3748
Aedes|[12514]|JWDCE851-10|BOLD:AAA3748
Aedes|[12515]|JWDCE865-10|BOLD:AAA3748
Aedes|[12516]|JWDCC449-10|BOLD:AAA3748
Aedes|[12517]|JWDCC271-10|BOLD:AAA3748
Aedes|[12518]|JWDCD100-10|BOLD:AAA3748
Aedes|[12519]|SWSWP074-09|BOLD:AAA3748
Aedes|[12520]|JWDCC202-10|BOLD:AAA3748
Aedes|[12521]|JWDCD135-10|BOLD:AAA3748
Aedes|[12522]|SWSWP016-09|BOLD:AAA3748
Aedes|[12523]|SWSWP061-09|BOLD:AAA3748
Aedes|[12524]|JWDCD104-10|BOLD:AAA3748
Aedes|[12525]|MHMOS017-07|BOLD:AAA3748
Aedes|[12526]|JWDCC437-10|BOLD:AAA3748
Aedes|[12527]|JWDCC453-10|BOLD:AAA3748
Aedes|[12528]|JWDCJ407-11|BOLD:AAA3748
Aedes|[12529]|JWDCJ931-11|BOLD:AAA3748
Aedes|[12530]|JWDCG626-10|BOLD:AAA3748
Aedes|[12531]|JWDCI669-11|BOLD:AAA3748
Aedes|[12532]|JWDCC435-10|BOLD:AAA3748
Aedes|[12533]|JWDCC436-10|BOLD:AAA3748
Aedes|[12534]|JWDCL448-11|BOLD:AAA3748
Aedes|[12535]|JWDCJ218-11|BOLD:AAA3748
Aedes|[12536]|JWDCG210-10|BOLD:AAA3748
Aedes|[12537]|JWDCG211-10|BOLD:AAA3748
Aedes|[12538]|JWDCH389-10|BOLD:AAA3748

Aedes|[12536]|JWDCG210-10|BOLD:AAA3748
Aedes|[12537]|JWDCG211-10|BOLD:AAA3748
Aedes|[12538]|JWDCH389-10|BOLD:AAA3748
Aedes|[12539]|JWDCH441-10|BOLD:AAA3748
Aedes|[12540]|JWDCE529-10|BOLD:AAA3748
Aedes|[12541]|JWDCE650-10|BOLD:AAA3748
Aedes|[12542]|JWDCE703-10|BOLD:AAA3748
Aedes|[12543]|JWDCF443-10|BOLD:AAA3748
Aedes|[12544]|JWDCE923-10|BOLD:AAA3748
Aedes|[12545]|JWDCE927-10|BOLD:AAA3748
Aedes|[12546]|MHMOS076-07|BOLD:AAA3748
Aedes|[12547]|MHMOS078-07|BOLD:AAA3748
Aedes|[12548]|MHMOS046-07|BOLD:AAA3748
Aedes|[12549]|MHMOS062-07|BOLD:AAA3748
Aedes|[12550]|JWDCC057-10|BOLD:AAA3748
Aedes|[12551]|JWDCC071-10|BOLD:AAA3748
Aedes|[12552]|JWDCC461-10|BOLD:AAA3748
Aedes|[12553]|JWDCC133-10|BOLD:AAA3748
Aedes|[12554]|AMMC055-07|BOLD:AAA3748
Aedes|[12555]|AMMC019-07|BOLD:AAA3748
Aedes|[12556]|SAMOS019-09|BOLD:AAA3748
Aedes|[12557]|SAMOS035-09|BOLD:AAA3748
Aedes|[12558]|AMMC043-07|BOLD:AAA3748
Aedes|[12559]|AMMC045-07|BOLD:AAA3748
Aedes|[12560]|AMMC042-07|BOLD:AAA3748
Aedes|[12561]|AMMC037-07|BOLD:AAA3748
Aedes|[12562]|AMMC036-07|BOLD:AAA3748
Aedes|[12563]|AMMC029-07|BOLD:AAA3748
Aedes|[12564]|AMMC005-07|BOLD:AAA3748
Aedes|[12565]|SAMOS011-09|BOLD:AAA3748
Aedes|[12566]|AMMC082-07|BOLD:AAA3748
Aedes|[12567]|AMMC048-07|BOLD:AAA3748
Aedes|[12568]|JWDCG624-10|BOLD:AAA3748
Aedes|[12569]|JWDCC058-10|BOLD:AAA3748
Aedes|[12570]|AMMC007-07|BOLD:AAA3748
Aedes|[12571]|MHMOS073-07|BOLD:AAA3748
Aedes|[12572]|JWDCE929-10|BOLD:AAA3748
Aedes|[12573]|MHMOS047-07|BOLD:AAA3748
Aedes|[12574]|JWDCC777-10|BOLD:AAA3748
Aedes|[12575]|JWDCC878-10|BOLD:AAA3748
Aedes|[12576]|AMMC028-07|BOLD:AAA3748
Aedes|[12577]|SWSWP023-09|BOLD:AAA3748
Aedes|[12578]|JWDCC778-10|BOLD:AAA3748
Aedes|[12579]|AMMC090-07|BOLD:AAA3748
Aedes|[12580]|JWDCF380-10|BOLD:AAA3748
Aedes|[12581]|JWDCD123-10|BOLD:AAA3748
Aedes|[12582]|JWDCE530-10|BOLD:AAA3748
Aedes|[12583]|JWDCF372-10|BOLD:AAA3748
Aedes|[12584]|AMMC083-07|BOLD:AAA3748
Aedes|[12585]|JWDCC779-10|BOLD:AAA3748
Aedes|[12586]|JWDCE549-10|BOLD:AAA3748
Aedes|[12587]|MHFLY119-07|BOLD:AAA3748
Aedes|[12588]|MHFLY121-07|BOLD:AAA3748
Aedes diantaeus|[12589]|JWDCH091-10|BOLD:AAF2904
Aedes|[12590]|JWDCK160-11|BOLD:AAD4326
Aedes|[12591]|JWDCH642-10|BOLD:AAD4326
Aedes flavescens|[12592]|MHMOS005-07|BOLD:AAA6145
Aedes flavescens|[12593]|MHMOS063-07|BOLD:AAA6145
Aedes flavescens|[12594]|MHMOS056-07|BOLD:AAA6145
Aedes flavescens|[12595]|MHMOS055-07|BOLD:AAA6145
Aedes flavescens|[12596]|MHMOS012-07|BOLD:AAA6145
Aedes flavescens|[12597]|MHMOS011-07|BOLD:AAA6145
Aedes flavescens|[12598]|MHMOS010-07|BOLD:AAA6145
Aedes flavescens|[12599]|MHMOS008-07|BOLD:AAA6145
Aedes flavescens|[12600]|MHMOS007-07|BOLD:AAA6145
Aedes flavescens|[12601]|MHMOS004-07|BOLD:AAA6145
Aedes flavescens|[12602]|MHMOS002-07|BOLD:AAA6145
Aedes flavescens|[12603]|MHMOS001-07|BOLD:AAA6145
Aedes flavescens|[12604]|SAMOS065-09|BOLD:AAA6145
Aedes flavescens|[12605]|SAMOS064-09|BOLD:AAA6145
Aedes flavescens|[12606]|MHMOS057-07|BOLD:AAA6145
Aedes flavescens|[12607]|MHMOS003-07|BOLD:AAA6145
Aedes flavescens|[12608]|MHMOS059-07|BOLD:AAA6145
Aedes flavescens|[12609]|MHMOS061-07|BOLD:AAA6145
Aedes flavescens|[12610]|JWDCL447-11|BOLD:AAA6145
Aedes flavescens|[12611]|JWDCI766-11|BOLD:AAA6145
Aedes flavescens|[12612]|JWDCI811-11|BOLD:AAA6145
Aedes flavescens|[12613]|JWDCI857-11|BOLD:AAA6145
Aedes flavescens|[12614]|JWDCF258-10|BOLD:AAA6145
Aedes excrucians|[12615]|JWDCJ1365-11|BOLD:AAB1098
Aedes excrucians|[12616]|SAMOS012-09|BOLD:AAB1098
Aedes excrucians|[12617]|JWDCG478-10|BOLD:AAB1098
Aedes excrucians|[12618]|MMADL030-09|BOLD:AAB1098
Aedes excrucians|[12619]|MMADL002-09|BOLD:AAB1098
Aedes excrucians|[12620]|MMADL003-09|BOLD:AAB1098
Aedes excrucians|[12621]|MMADL004-09|BOLD:AAB1098
Aedes excrucians|[12622]|MHMOS079-07|BOLD:AAB1098
Aedes excrucians|[12623]|MMADL001-09|BOLD:AAB1098
Aedes excrucians|[12624]|JWDCJ049-11|BOLD:AAB1098
Aedes excrucians|[12625]|JWDCG694-10|BOLD:AAB1098
Aedes excrucians|[12626]|MMADL068-09|BOLD:AAB1098
Aedes excrucians|[12627]|MMADL026-09|BOLD:AAB1098
Aedes excrucians|[12628]|MHMOS023-07|BOLD:AAB1098
Aedes excrucians|[12629]|MMADL017-09|BOLD:AAB1098
Aedes excrucians|[12630]|AMMC072-07|BOLD:AAB1098
Aedes excrucians|[12631]|MMADL031-09|BOLD:AAB1098
Aedes excrucians|[12632]|JWDCI615-10|BOLD:AAB1098
Aedes excrucians|[12633]|JWDCB149-10|BOLD:AAB1098
Aedes excrucians|[12634]|JWDCF758-10|BOLD:AAB1098
Aedes excrucians|[12635]|JWDCG353-10|BOLD:AAB1098
Aedes excrucians|[12636]|JWDCI119-10|BOLD:AAB1098
Aedes excrucians|[12637]|JWDCB152-10|BOLD:AAB1098
Aedes excrucians|[12638]|JWDCL175-11|BOLD:AAB1098

Aedes excrucians|[illegible]|[illegible]-10|BOLD:AAB1098
Aedes excrucians|[12636]|JWDCI119-10|BOLD:AAB1098
Aedes excrucians|[12637]|JWDCB152-10|BOLD:AAB1098
Aedes excrucians|[12638]|JWDCL175-11|BOLD:AAB1098
Aedes excrucians|[12639]|JWDCF381-10|BOLD:AAB1098
Aedes excrucians|[12640]|MHMOS069-07|BOLD:AAB1098
Aedes excrucians|[12641]|MHMOS040-07|BOLD:AAB1098
Aedes excrucians|[12642]|AMMC089-07|BOLD:AAB1098
Aedes excrucians|[12643]|MMADL057-09|BOLD:AAB1098
Aedes excrucians|[12644]|JWDCJ1220-11|BOLD:AAB1098
Aedes excrucians|[12645]|JWDCI168-10|BOLD:AAB1098
Aedes excrucians|[12646]|JWDCF259-10|BOLD:AAB1098
Aedes excrucians|[12647]|JWDCF886-10|BOLD:AAB1098
Aedes excrucians|[12648]|SAMOS015-09|BOLD:AAB1098
Aedes excrucians|[12649]|JWDCI167-10|BOLD:AAB1098
Aedes excrucians|[12650]|JWDCL219-11|BOLD:AAB1098
Aedes excrucians|[12651]|JWDCA309-10|BOLD:AAB1098
Aedes excrucians|[12652]|JWDCA305-10|BOLD:AAB1098
Aedes excrucians|[12653]|MMADL007-09|BOLD:AAB1098
Aedes excrucians|[12654]|MMADL008-09|BOLD:AAB1098
Aedes excrucians|[12655]|MMADL065-09|BOLD:AAB1098
Aedes excrucians|[12656]|JWDCJ1302-11|BOLD:AAB1098
Aedes excrucians|[12657]|JWDCL173-11|BOLD:AAB1098
Aedes excrucians|[12658]|JWDCH046-10|BOLD:AAB1098
Aedes excrucians|[12659]|JWDCH265-10|BOLD:AAB1098
Aedes excrucians|[12660]|JWDCF106-10|BOLD:AAB1098
Aedes excrucians|[12661]|AMMC026-07|BOLD:AAB1098
Aedes excrucians|[12662]|JWDCI120-10|BOLD:AAB1098
Aedes excrucians|[12663]|JWDCJ1366-11|BOLD:AAB1098
Aedes excrucians|[12664]|JWDCA319-10|BOLD:AAB1098
Aedes excrucians|[12665]|JWDCG863-10|BOLD:AAB1098
Aedes excrucians|[12666]|JWDCJ278-11|BOLD:AAB1098
Aedes excrucians|[12667]|JWDCJ1517-11|BOLD:AAB1098
Aedes excrucians|[12668]|JWDCJ1454-11|BOLD:AAB1098
Aedes excrucians|[12669]|JWDCJ1041-11|BOLD:AAB1098
Aedes excrucians|[12670]|JWDCK213-11|BOLD:AAB1098
Aedes excrucians|[12671]|JWDCL220-11|BOLD:AAB1098
Aedes excrucians|[12672]|JWDCH865-10|BOLD:AAB1098
Aedes excrucians|[12673]|JWDCG039-10|BOLD:AAB1098
Aedes excrucians|[12674]|JWDCG216-10|BOLD:AAB1098
Aedes excrucians|[12675]|JWDCH250-10|BOLD:AAB1098
Aedes excrucians|[12676]|MMADL032-09|BOLD:AAB1098
Aedes excrucians|[12677]|JWDCJ1043-11|BOLD:AAB1098
Aedes excrucians|[12678]|JWDCG862-10|BOLD:AAB1098
Aedes excrucians|[12679]|JWDCF850-10|BOLD:AAB1098
Aedes excrucians|[12680]|SAMOS016-09|BOLD:AAB1098
Aedes excrucians|[12681]|JWDCL399-11|BOLD:AAB1098
Aedes excrucians|[12682]|JWDCE680-10|BOLD:AAB1098
Aedes excrucians|[12683]|MMADL021-09|BOLD:AAB1098
Aedes excrucians|[12684]|SAMOS063-09|BOLD:AAB1098
Aedes excrucians|[12685]|MHMOS042-07|BOLD:AAB1098
Aedes excrucians|[12686]|JWDCG625-10|BOLD:AAB1098
Aedes excrucians|[12687]|JWDCB155-10|BOLD:AAB1098
Aedes excrucians|[12688]|MMADL027-09|BOLD:AAB1098
Aedes excrucians|[12689]|JWDCI614-10|BOLD:AAB1098
Aedes excrucians|[12690]|JWDCG215-10|BOLD:AAB1098
Aedes excrucians|[12691]|MMADL005-09|BOLD:AAB1098
Aedes excrucians|[12692]|JWDCG804-10|BOLD:AAB1098
Aedes excrucians|[12693]|JWDCH694-10|BOLD:AAB1098
Aedes excrucians|[12694]|JWDCL099-11|BOLD:AAB1098
Aedes excrucians|[12695]|MMADL067-09|BOLD:AAB1098
Aedes excrucians|[12696]|JWDCA320-10|BOLD:AAB1098
Aedes excrucians|[12697]|AMMC091-07|BOLD:AAB1098
Aedes excrucians|[12698]|MHMOS034-07|BOLD:AAB1098
Aedes excrucians|[12699]|MMADL069-09|BOLD:AAB1098
Aedes excrucians|[12700]|JWDCJ152-11|BOLD:AAB1098
Aedes excrucians|[12701]|MMADL063-09|BOLD:AAB1098
Aedes excrucians|[12702]|JWDCC201-10|BOLD:AAB1098
Aedes excrucians|[12703]|JWDCH693-10|BOLD:AAB1098
Aedes excrucians|[12704]|JWDCH737-10|BOLD:AAB1098
Aedes excrucians|[12705]|JWDCG894-10|BOLD:AAB1098
Aedes excrucians|[12706]|JWDCG041-10|BOLD:AAB1098
Aedes excrucians|[12707]|JWDCI072-10|BOLD:AAB1098
Aedes excrucians|[12708]|JWDCE431-10|BOLD:AAB1098
Aedes excrucians|[12709]|JWDCA306-10|BOLD:AAB1098
Aedes excrucians|[12710]|AMMC058-07|BOLD:AAB1098
Aedes excrucians|[12711]|MMADL033-09|BOLD:AAB1098
Aedes excrucians|[12712]|JWDCJ969-11|BOLD:AAB1098
Aedes excrucians|[12713]|JWDCH486-10|BOLD:AAB1098
Aedes excrucians|[12714]|AMMC051-07|BOLD:AAB1098
Aedes excrucians|[12715]|JWDCC470-10|BOLD:AAB1098
Aedes excrucians|[12716]|JWDCL398-11|BOLD:AAB1098
Aedes excrucians|[12717]|JWDCC227-10|BOLD:AAB1098
Aedes excrucians|[12718]|JWDCF848-10|BOLD:AAB1098
Aedes excrucians|[12719]|JWDCJ047-11|BOLD:AAB1098
Aedes excrucians|[12720]|JWDCL027-11|BOLD:AAB1098
Aedes excrucians|[12721]|MMADL064-09|BOLD:AAB1098
Aedes excrucians|[12722]|JWDCG040-10|BOLD:AAB1098
Aedes excrucians|[12723]|JWDCI938-11|BOLD:AAB1098
Aedes excrucians|[12724]|AMMC068-07|BOLD:AAB1098
Aedes excrucians|[12725]|JWDCJ1300-11|BOLD:AAB1098
Aedes excrucians|[12726]|JWDCJ1455-11|BOLD:AAB1098
Aedes excrucians|[12727]|JWDCG352-10|BOLD:AAB1098
Aedes excrucians|[12728]|JWDCH440-10|BOLD:AAB1098
Aedes excrucians|[12729]|JWDCG213-10|BOLD:AAB1098
Aedes excrucians|[12730]|JWDCH558-10|BOLD:AAB1098
Aedes excrucians|[12731]|JWDCF318-10|BOLD:AAB1098
Aedes excrucians|[12732]|JWDCB153-10|BOLD:AAB1098
Aedes excrucians|[12733]|JWDCK221-11|BOLD:AAB1098
Aedes excrucians|[12734]|JWDCJ1573-11|BOLD:AAB1098
Aedes excrucians|[12735]|JWDCI467-10|BOLD:AAB1098
Aedes excrucians|[12736]|JWDCH264-10|BOLD:AAB1098
Aedes excrucians|[12737]|JWDCC776-10|BOLD:AAB1098

Aedes excrucians|[12735]|JWDCI467-10|BOLD:AAB1098
Aedes excrucians|[12736]|JWDCH264-10|BOLD:AAB1098
Aedes excrucians|[12737]|JWDCC776-10|BOLD:AAB1098
Aedes excrucians|[12738]|JWDCK097-11|BOLD:AAB1098
Aedes excrucians|[12739]|JWDCI118-10|BOLD:AAB1098
Aedes excrucians|[12740]|JWDCI274-10|BOLD:AAB1098
Aedes excrucians|[12741]|JWDCL294-11|BOLD:AAB1098
Aedes excrucians|[12742]|JWDCG892-10|BOLD:AAB1098
Aedes excrucians|[12743]|MMADL070-09|BOLD:AAB1098
Aedes excrucians|[12744]|MMADL090-09|BOLD:AAB1098
Aedes excrucians|[12745]|MMADL034-09|BOLD:AAB1098
Aedes excrucians|[12746]|MMADL066-09|BOLD:AAB1098
Aedes excrucians|[12747]|JWDCA315-10|BOLD:AAB1098
Aedes excrucians|[12748]|JWDCA321-10|BOLD:AAB1098
Aedes excrucians|[12749]|MMADL022-09|BOLD:AAB1098
Aedes excrucians|[12750]|MMADL020-09|BOLD:AAB1098
Aedes excrucians|[12751]|MMADL016-09|BOLD:AAB1098
Aedes excrucians|[12752]|MHMOS030-07|BOLD:AAB1098
Aedes excrucians|[12753]|SAMOS014-09|BOLD:AAB1098
Aedes excrucians|[12754]|AMMC067-07|BOLD:AAB1098
Aedes excrucians|[12755]|AMMC038-07|BOLD:AAB1098
Aedes excrucians|[12756]|AMMC006-07|BOLD:AAB1098
Aedes excrucians|[12757]|JWDCA316-10|BOLD:AAB1098
Aedes excrucians|[12758]|AMMC066-07|BOLD:AAB1098
Aedes excrucians|[12759]|SAMOS010-09|BOLD:AAB1098
Aedes excrucians|[12760]|MHMOS090-07|BOLD:AAB1098
Aedes excrucians|[12761]|MMADL006-09|BOLD:AAB1098
Aedes excrucians|[12762]|MMADL018-09|BOLD:AAB1098
Aedes excrucians|[12763]|MMADL019-09|BOLD:AAB1098
Aedes excrucians|[12764]|MMADL035-09|BOLD:AAB1098
Aedes excrucians|[12765]|MMADL052-09|BOLD:AAB1098
Aedes excrucians|[12766]|MMADL071-09|BOLD:AAB1098
Aedes excrucians|[12767]|JWDCL295-11|BOLD:AAB1098
Aedes excrucians|[12768]|JWDCI939-11|BOLD:AAB1098
Aedes excrucians|[12769]|SAMOS066-09|BOLD:AAB1098
Aedes excrucians|[12770]|JWDCJ1042-11|BOLD:AAB1098
Aedes excrucians|[12771]|JWDCF371-10|BOLD:AAB1098
Aedes excrucians|[12772]|JWDCG893-10|BOLD:AAB1098
Aedes excrucians|[12773]|MHMOS041-07|BOLD:AAB1098
Aedes excrucians|[12774]|SAMOS013-09|BOLD:AAB1098
Aedes excrucians|[12775]|MHFLY122-07|BOLD:AAB1098
Paratanytarsus cf. tenuis|[12776]|JWDCE310-10|BOLD:AAP5145
Paratanytarsus cf. tenuis|[12777]|CHURC900-09|BOLD:AAE3685
Paratanytarsus cf. tenuis|[12778]|CHURC545-08|BOLD:AAE3685
Paratanytarsus sp. TE08|[12779]|CHURC673-09|BOLD:AAI2608
Paratanytarsus sp. TE08|[12780]|CHURC677-09|BOLD:AAI2608
Paratanytarsus sp. TE08|[12781]|CHURC683-09|BOLD:AAI2608
Paratanytarsus sp. TE03|[12782]|CHURC034-06|BOLD:AAE3675
Paratanytarsus|[12783]|JWDCD298-10|BOLD:AAM6259
Paratanytarsus setosimanus|[12784]|MHMOS154-07|BOLD:AAC4205
Paratanytarsus setosimanus|[12785]|CHURC320-07|BOLD:AAC4206
Paratanytarsus setosimanus|[12786]|CHURC319-07|BOLD:AAC4206
Paratanytarsus setosimanus|[12787]|CHURC316-07|BOLD:AAC4206
Paratanytarsus setosimanus|[12788]|CHURC323-07|BOLD:AAC4206
Paratanytarsus setosimanus|[12789]|JWDCF733-10|BOLD:AAC4206
Paratanytarsus setosimanus|[12790]|JWDCJ883-11|BOLD:AAC4206
Paratanytarsus setosimanus|[12791]|JWDCH708-10|BOLD:AAC4206
Paratanytarsus setosimanus|[12792]|CHURC310-07|BOLD:AAC4206
Paratanytarsus sp. 1ES|[12793]|JWDCE118-10|BOLD:ADY7766
Paratanytarsus sp. 1ES|[12794]|JWDCE006-10|BOLD:ADY7766
Paratanytarsus sp. 1ES|[12795]|CHURC295-07|BOLD:ADY7766
Paratanytarsus sp. TE04|[12796]|JWDCE009-10|BOLD:AAI2607
Paratanytarsus sp. TE04|[12797]|JWDCE005-10|BOLD:AAI2607
Paratanytarsus sp. TE04|[12798]|CHURC636-09|BOLD:AAI2607
Paratanytarsus|[12799]|JWDCK107-11|BOLD:AAV5946
Paratanytarsus austriacus|[12800]|CHURC477-08|BOLD:AAA7954
Paratanytarsus austriacus|[12801]|CHURC426-07|BOLD:AAA7954
Paratanytarsus austriacus|[12802]|CHURC1028-09|BOLD:AAA7954
Paratanytarsus austriacus|[12803]|CHURC554-08|BOLD:AAA7954
Paratanytarsus abiskoensis|[12804]|JWDCD223-10|BOLD:AAE3677
Paratanytarsus abiskoensis|[12805]|JWDCE695-10|BOLD:AAE3677
Paratanytarsus abiskoensis|[12806]|CHURC930-09|BOLD:AAE3677
Paratanytarsus abiskoensis|[12807]|CHURC557-08|BOLD:AAE3677
Paratanytarsus dissimilis|[12808]|CHURC196-07|BOLD:AAE3698
Paratanytarsus dissimilis|[12809]|CHURC262-07|BOLD:AAE3698
Paratanytarsus dissimilis|[12810]|CHURC265-07|BOLD:AAE3698
Paratanytarsus laccophilus|[12811]|JWDCD306-10|BOLD:AAC8842
Paratanytarsus laccophilus|[12812]|JWDCD358-10|BOLD:AAC8842
Paratanytarsus laccophilus|[12813]|JWDCD357-10|BOLD:AAC8842
Paratanytarsus laccophilus|[12814]|CHURC322-07|BOLD:AAC8842
Paratanytarsus laccophilus|[12815]|CHURC318-07|BOLD:AAC8842
Paratanytarsus laccophilus|[12816]|CHURC415-07|BOLD:AAC8842
Paratanytarsus laccophilus|[12817]|JWDCF072-10|BOLD:AAC8842
Paratanytarsus laccophilus|[12818]|CHURC1033-09|BOLD:AAC8842
Paratanytarsus laccophilus|[12819]|CHURC1000-09|BOLD:AAC8842
Paratanytarsus laccophilus|[12820]|JWDCC357-10|BOLD:AAC8842
Paratanytarsus laccophilus|[12821]|JWDCC363-10|BOLD:AAC8842
Paratanytarsus laccophilus|[12822]|CHURC782-09|BOLD:AAC8842
Paratanytarsus laccophilus|[12823]|CHURC327-07|BOLD:AAC8842
Paratanytarsus laccophilus|[12824]|JWDCC377-10|BOLD:AAC8842
Paratanytarsus laccophilus|[12825]|JWDCC368-10|BOLD:AAC8842
Paratanytarsus laccophilus|[12826]|JWDCC362-10|BOLD:AAC8842
Paratanytarsus laccophilus|[12827]|JWDCC356-10|BOLD:AAC8842
Paratanytarsus laccophilus|[12828]|JWDCC496-10|BOLD:AAC8842
Paratanytarsus laccophilus|[12829]|JWDCJ1155-11|BOLD:AAC8842
Paratanytarsus laccophilus|[12830]|JWDCC495-10|BOLD:AAC8842
Paratanytarsus laccophilus|[12831]|CHURC1054-09|BOLD:AAC8842
Paratanytarsus laccophilus|[12832]|JWDCC366-10|BOLD:AAC8842
Paratanytarsus laccophilus|[12833]|JWDCD360-10|BOLD:AAC8842
Paratanytarsus laccophilus|[12834]|JWDCC374-10|BOLD:AAC8842
Paratanytarsus laccophilus|[12835]|CHURC1061-09|BOLD:AAC8842
Paratanytarsus laccophilus|[12836]|CHURC790-09|BOLD:AAC8842
Paratanytarsus laccophilus|[12837]|CHURC997-09|BOLD:AAC8842

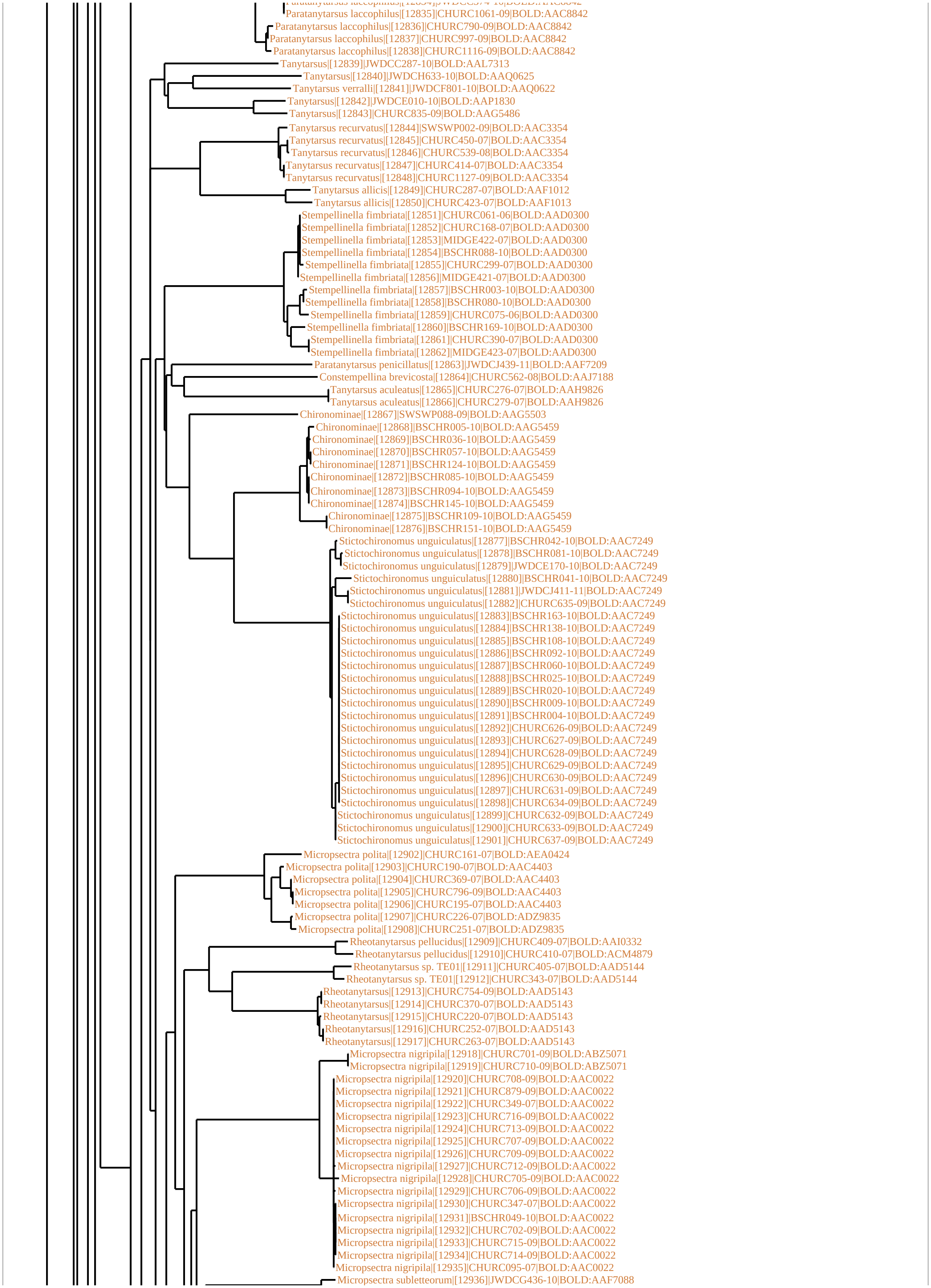
Paratanytarsus laccophilus|[12835]|CHURC1061-09|BOLD:AAC8842
Paratanytarsus laccophilus|[12836]|CHURC790-09|BOLD:AAC8842
Paratanytarsus laccophilus|[12837]|CHURC997-09|BOLD:AAC8842
Paratanytarsus laccophilus|[12838]|CHURC1116-09|BOLD:AAC8842
Tanytarsus|[12839]|JWDCC287-10|BOLD:AAL7313
Tanytarsus|[12840]|JWDCH633-10|BOLD:AAQ0625
Tanytarsus verralli|[12841]|JWDCF801-10|BOLD:AAQ0622
Tanytarsus|[12842]|JWDCE010-10|BOLD:AAP1830
Tanytarsus|[12843]|CHURC835-09|BOLD:AAG5486
Tanytarsus recurvatus|[12844]|SWSWP002-09|BOLD:AAC3354
Tanytarsus recurvatus|[12845]|CHURC450-07|BOLD:AAC3354
Tanytarsus recurvatus|[12846]|CHURC539-08|BOLD:AAC3354
Tanytarsus recurvatus|[12847]|CHURC414-07|BOLD:AAC3354
Tanytarsus recurvatus|[12848]|CHURC1127-09|BOLD:AAC3354
Tanytarsus allicis|[12849]|CHURC287-07|BOLD:AAF1012
Tanytarsus allicis|[12850]|CHURC423-07|BOLD:AAF1013
Stempellinella fimbriata|[12851]|CHURC061-06|BOLD:AAD0300
Stempellinella fimbriata|[12852]|CHURC168-07|BOLD:AAD0300
Stempellinella fimbriata|[12853]|MIDGE422-07|BOLD:AAD0300
Stempellinella fimbriata|[12854]|BSCHR088-10|BOLD:AAD0300
Stempellinella fimbriata|[12855]|CHURC299-07|BOLD:AAD0300
Stempellinella fimbriata|[12856]|MIDGE421-07|BOLD:AAD0300
Stempellinella fimbriata|[12857]|BSCHR003-10|BOLD:AAD0300
Stempellinella fimbriata|[12858]|BSCHR080-10|BOLD:AAD0300
Stempellinella fimbriata|[12859]|CHURC075-06|BOLD:AAD0300
Stempellinella fimbriata|[12860]|BSCHR169-10|BOLD:AAD0300
Stempellinella fimbriata|[12861]|CHURC390-07|BOLD:AAD0300
Stempellinella fimbriata|[12862]|MIDGE423-07|BOLD:AAD0300
Paratanytarsus penicillatus|[12863]|JWDCJ439-11|BOLD:AAF7209
Constempellina brevicosta|[12864]|CHURC562-08|BOLD:AAJ7188
Tanytarsus aculeatus|[12865]|CHURC276-07|BOLD:AAH9826
Tanytarsus aculeatus|[12866]|CHURC279-07|BOLD:AAH9826
Chironominae|[12867]|SWSWP088-09|BOLD:AAG5503
Chironominae|[12868]|BSCHR005-10|BOLD:AAG5459
Chironominae|[12869]|BSCHR036-10|BOLD:AAG5459
Chironominae|[12870]|BSCHR057-10|BOLD:AAG5459
Chironominae|[12871]|BSCHR124-10|BOLD:AAG5459
Chironominae|[12872]|BSCHR085-10|BOLD:AAG5459
Chironominae|[12873]|BSCHR094-10|BOLD:AAG5459
Chironominae|[12874]|BSCHR145-10|BOLD:AAG5459
Chironominae|[12875]|BSCHR109-10|BOLD:AAG5459
Chironominae|[12876]|BSCHR151-10|BOLD:AAG5459
Stictochironomus unguiculatus|[12877]|BSCHR042-10|BOLD:AAC7249
Stictochironomus unguiculatus|[12878]|BSCHR081-10|BOLD:AAC7249
Stictochironomus unguiculatus|[12879]|JWDCE170-10|BOLD:AAC7249
Stictochironomus unguiculatus|[12880]|BSCHR041-10|BOLD:AAC7249
Stictochironomus unguiculatus|[12881]|JWDCJ411-11|BOLD:AAC7249
Stictochironomus unguiculatus|[12882]|CHURC635-09|BOLD:AAC7249
Stictochironomus unguiculatus|[12883]|BSCHR163-10|BOLD:AAC7249
Stictochironomus unguiculatus|[12884]|BSCHR138-10|BOLD:AAC7249
Stictochironomus unguiculatus|[12885]|BSCHR108-10|BOLD:AAC7249
Stictochironomus unguiculatus|[12886]|BSCHR092-10|BOLD:AAC7249
Stictochironomus unguiculatus|[12887]|BSCHR060-10|BOLD:AAC7249
Stictochironomus unguiculatus|[12888]|BSCHR025-10|BOLD:AAC7249
Stictochironomus unguiculatus|[12889]|BSCHR020-10|BOLD:AAC7249
Stictochironomus unguiculatus|[12890]|BSCHR009-10|BOLD:AAC7249
Stictochironomus unguiculatus|[12891]|BSCHR004-10|BOLD:AAC7249
Stictochironomus unguiculatus|[12892]|CHURC626-09|BOLD:AAC7249
Stictochironomus unguiculatus|[12893]|CHURC627-09|BOLD:AAC7249
Stictochironomus unguiculatus|[12894]|CHURC628-09|BOLD:AAC7249
Stictochironomus unguiculatus|[12895]|CHURC629-09|BOLD:AAC7249
Stictochironomus unguiculatus|[12896]|CHURC630-09|BOLD:AAC7249
Stictochironomus unguiculatus|[12897]|CHURC631-09|BOLD:AAC7249
Stictochironomus unguiculatus|[12898]|CHURC634-09|BOLD:AAC7249
Stictochironomus unguiculatus|[12899]|CHURC632-09|BOLD:AAC7249
Stictochironomus unguiculatus|[12900]|CHURC633-09|BOLD:AAC7249
Stictochironomus unguiculatus|[12901]|CHURC637-09|BOLD:AAC7249
Micropsectra polita|[12902]|CHURC161-07|BOLD:AEA0424
Micropsectra polita|[12903]|CHURC190-07|BOLD:AAC4403
Micropsectra polita|[12904]|CHURC369-07|BOLD:AAC4403
Micropsectra polita|[12905]|CHURC796-09|BOLD:AAC4403
Micropsectra polita|[12906]|CHURC195-07|BOLD:AAC4403
Micropsectra polita|[12907]|CHURC226-07|BOLD:ADZ9835
Micropsectra polita|[12908]|CHURC251-07|BOLD:ADZ9835
Rheotanytarsus pellucidus|[12909]|CHURC409-07|BOLD:AAI0332
Rheotanytarsus pellucidus|[12910]|CHURC410-07|BOLD:ACM4879
Rheotanytarsus sp. TE01|[12911]|CHURC405-07|BOLD:AAD5144
Rheotanytarsus sp. TE01|[12912]|CHURC343-07|BOLD:AAD5144
Rheotanytarsus|[12913]|CHURC754-09|BOLD:AAD5143
Rheotanytarsus|[12914]|CHURC370-07|BOLD:AAD5143
Rheotanytarsus|[12915]|CHURC220-07|BOLD:AAD5143
Rheotanytarsus|[12916]|CHURC252-07|BOLD:AAD5143
Rheotanytarsus|[12917]|CHURC263-07|BOLD:AAD5143
Micropsectra nigripila|[12918]|CHURC701-09|BOLD:ABZ5071
Micropsectra nigripila|[12919]|CHURC710-09|BOLD:ABZ5071
Micropsectra nigripila|[12920]|CHURC708-09|BOLD:AAC0022
Micropsectra nigripila|[12921]|CHURC879-09|BOLD:AAC0022
Micropsectra nigripila|[12922]|CHURC349-07|BOLD:AAC0022
Micropsectra nigripila|[12923]|CHURC716-09|BOLD:AAC0022
Micropsectra nigripila|[12924]|CHURC713-09|BOLD:AAC0022
Micropsectra nigripila|[12925]|CHURC707-09|BOLD:AAC0022
Micropsectra nigripila|[12926]|CHURC709-09|BOLD:AAC0022
Micropsectra nigripila|[12927]|CHURC712-09|BOLD:AAC0022
Micropsectra nigripila|[12928]|CHURC705-09|BOLD:AAC0022
Micropsectra nigripila|[12929]|CHURC706-09|BOLD:AAC0022
Micropsectra nigripila|[12930]|CHURC347-07|BOLD:AAC0022
Micropsectra nigripila|[12931]|BSCHR049-10|BOLD:AAC0022
Micropsectra nigripila|[12932]|CHURC702-09|BOLD:AAC0022
Micropsectra nigripila|[12933]|CHURC715-09|BOLD:AAC0022
Micropsectra nigripila|[12934]|CHURC714-09|BOLD:AAC0022
Micropsectra nigripila|[12935]|CHURC095-07|BOLD:AAC0022
Micropsectra subletteorum|[12936]|JWDCG436-10|BOLD:AAF7088

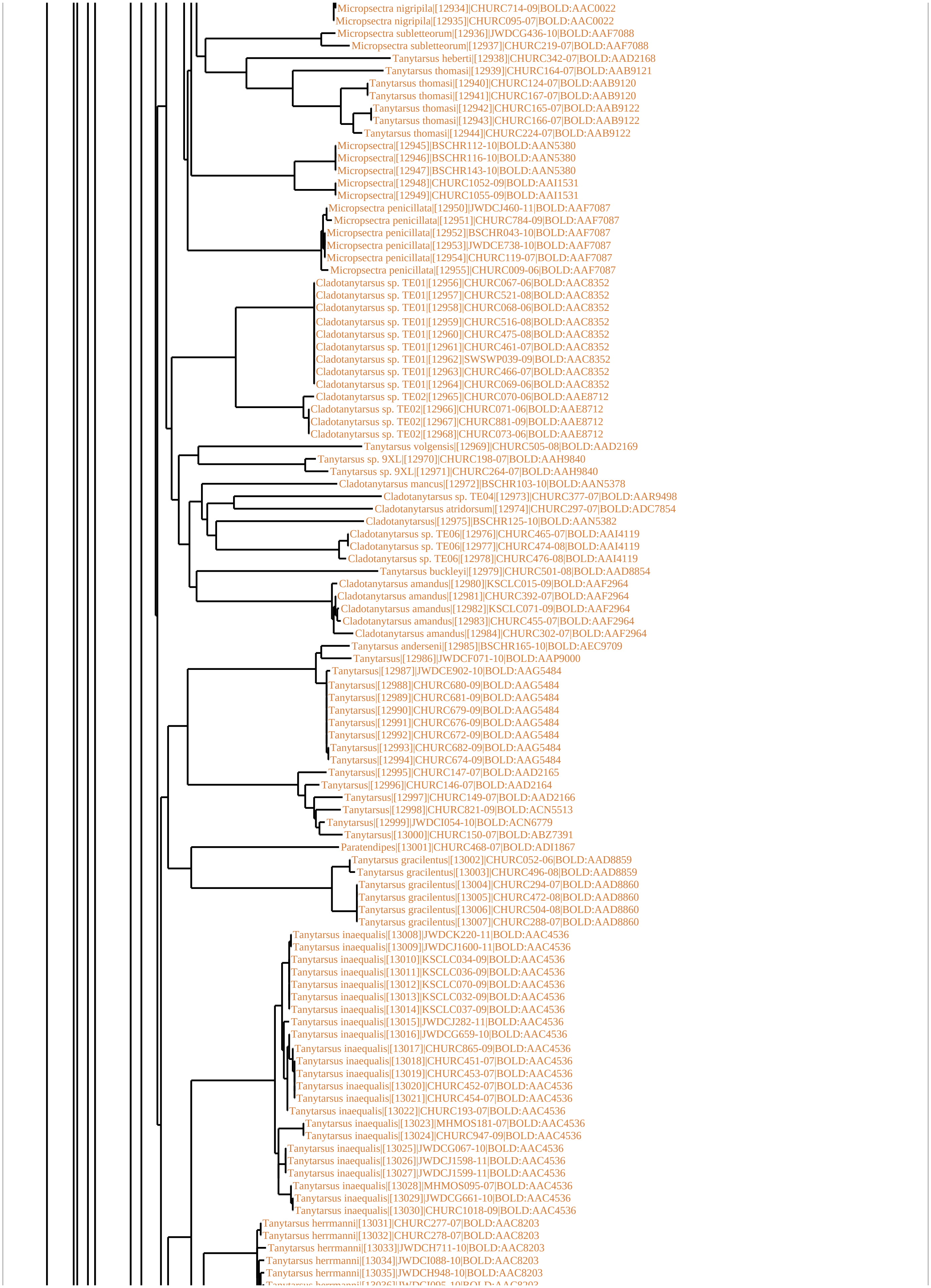
Micropsectra nigripila|[12934]|CHURC714-09|BOLD:AAC0022
Micropsectra nigripila|[12935]|CHURC095-07|BOLD:AAC0022
Micropsectra subletteorum|[12936]|JWDCG436-10|BOLD:AAF7088
Micropsectra subletteorum|[12937]|CHURC219-07|BOLD:AAF7088
Tanytarsus heberti|[12938]|CHURC342-07|BOLD:AAD2168
Tanytarsus thomasi|[12939]|CHURC164-07|BOLD:AAB9121
Tanytarsus thomasi|[12940]|CHURC124-07|BOLD:AAB9120
Tanytarsus thomasi|[12941]|CHURC167-07|BOLD:AAB9120
Tanytarsus thomasi|[12942]|CHURC165-07|BOLD:AAB9122
Tanytarsus thomasi|[12943]|CHURC166-07|BOLD:AAB9122
Tanytarsus thomasi|[12944]|CHURC224-07|BOLD:AAB9122
Micropsectra|[12945]|BSCHR112-10|BOLD:AAN5380
Micropsectra|[12946]|BSCHR116-10|BOLD:AAN5380
Micropsectra|[12947]|BSCHR143-10|BOLD:AAN5380
Micropsectra|[12948]|CHURC1052-09|BOLD:AAI1531
Micropsectra|[12949]|CHURC1055-09|BOLD:AAI1531
Micropsectra penicillata|[12950]|JWDCJ460-11|BOLD:AAF7087
Micropsectra penicillata|[12951]|CHURC784-09|BOLD:AAF7087
Micropsectra penicillata|[12952]|BSCHR043-10|BOLD:AAF7087
Micropsectra penicillata|[12953]|JWDCE738-10|BOLD:AAF7087
Micropsectra penicillata|[12954]|CHURC119-07|BOLD:AAF7087
Micropsectra penicillata|[12955]|CHURC009-06|BOLD:AAF7087
Cladotanytarsus sp. TE01|[12956]|CHURC067-06|BOLD:AAC8352
Cladotanytarsus sp. TE01|[12957]|CHURC521-08|BOLD:AAC8352
Cladotanytarsus sp. TE01|[12958]|CHURC068-06|BOLD:AAC8352
Cladotanytarsus sp. TE01|[12959]|CHURC516-08|BOLD:AAC8352
Cladotanytarsus sp. TE01|[12960]|CHURC475-08|BOLD:AAC8352
Cladotanytarsus sp. TE01|[12961]|CHURC461-07|BOLD:AAC8352
Cladotanytarsus sp. TE01|[12962]|SWSWP039-09|BOLD:AAC8352
Cladotanytarsus sp. TE01|[12963]|CHURC466-07|BOLD:AAC8352
Cladotanytarsus sp. TE01|[12964]|CHURC069-06|BOLD:AAC8352
Cladotanytarsus sp. TE02|[12965]|CHURC070-06|BOLD:AAE8712
Cladotanytarsus sp. TE02|[12966]|CHURC071-06|BOLD:AAE8712
Cladotanytarsus sp. TE02|[12967]|CHURC881-09|BOLD:AAE8712
Cladotanytarsus sp. TE02|[12968]|CHURC073-06|BOLD:AAE8712
Tanytarsus volgensis|[12969]|CHURC505-08|BOLD:AAD2169
Tanytarsus sp. 9XL|[12970]|CHURC198-07|BOLD:AAH9840
Tanytarsus sp. 9XL|[12971]|CHURC264-07|BOLD:AAH9840
Cladotanytarsus mancus|[12972]|BSCHR103-10|BOLD:AAN5378
Cladotanytarsus sp. TE04|[12973]|CHURC377-07|BOLD:AAR9498
Cladotanytarsus atridorsum|[12974]|CHURC297-07|BOLD:ADC7854
Cladotanytarsus|[12975]|BSCHR125-10|BOLD:AAN5382
Cladotanytarsus sp. TE06|[12976]|CHURC465-07|BOLD:AAI4119
Cladotanytarsus sp. TE06|[12977]|CHURC474-08|BOLD:AAI4119
Cladotanytarsus sp. TE06|[12978]|CHURC476-08|BOLD:AAI4119
Tanytarsus buckleyi|[12979]|CHURC501-08|BOLD:AAD8854
Cladotanytarsus amandus|[12980]|KSCLC015-09|BOLD:AAF2964
Cladotanytarsus amandus|[12981]|CHURC392-07|BOLD:AAF2964
Cladotanytarsus amandus|[12982]|KSCLC071-09|BOLD:AAF2964
Cladotanytarsus amandus|[12983]|CHURC455-07|BOLD:AAF2964
Cladotanytarsus amandus|[12984]|CHURC302-07|BOLD:AAF2964
Tanytarsus anderseni|[12985]|BSCHR165-10|BOLD:AEC9709
Tanytarsus|[12986]|JWDCF071-10|BOLD:AAP9000
Tanytarsus|[12987]|JWDCE902-10|BOLD:AAG5484
Tanytarsus|[12988]|CHURC680-09|BOLD:AAG5484
Tanytarsus|[12989]|CHURC681-09|BOLD:AAG5484
Tanytarsus|[12990]|CHURC679-09|BOLD:AAG5484
Tanytarsus|[12991]|CHURC676-09|BOLD:AAG5484
Tanytarsus|[12992]|CHURC672-09|BOLD:AAG5484
Tanytarsus|[12993]|CHURC682-09|BOLD:AAG5484
Tanytarsus|[12994]|CHURC674-09|BOLD:AAG5484
Tanytarsus|[12995]|CHURC147-07|BOLD:AAD2165
Tanytarsus|[12996]|CHURC146-07|BOLD:AAD2164
Tanytarsus|[12997]|CHURC149-07|BOLD:AAD2166
Tanytarsus|[12998]|CHURC821-09|BOLD:ACN5513
Tanytarsus|[12999]|JWDCI054-10|BOLD:ACN6779
Tanytarsus|[13000]|CHURC150-07|BOLD:ABZ7391
Paratendipes|[13001]|CHURC468-07|BOLD:ADI1867
Tanytarsus gracilentus|[13002]|CHURC052-06|BOLD:AAD8859
Tanytarsus gracilentus|[13003]|CHURC496-08|BOLD:AAD8859
Tanytarsus gracilentus|[13004]|CHURC294-07|BOLD:AAD8860
Tanytarsus gracilentus|[13005]|CHURC472-08|BOLD:AAD8860
Tanytarsus gracilentus|[13006]|CHURC504-08|BOLD:AAD8860
Tanytarsus gracilentus|[13007]|CHURC288-07|BOLD:AAD8860
Tanytarsus inaequalis|[13008]|JWDCK220-11|BOLD:AAC4536
Tanytarsus inaequalis|[13009]|JWDCJ1600-11|BOLD:AAC4536
Tanytarsus inaequalis|[13010]|KSCLC034-09|BOLD:AAC4536
Tanytarsus inaequalis|[13011]|KSCLC036-09|BOLD:AAC4536
Tanytarsus inaequalis|[13012]|KSCLC070-09|BOLD:AAC4536
Tanytarsus inaequalis|[13013]|KSCLC032-09|BOLD:AAC4536
Tanytarsus inaequalis|[13014]|KSCLC037-09|BOLD:AAC4536
Tanytarsus inaequalis|[13015]|JWDCJ282-11|BOLD:AAC4536
Tanytarsus inaequalis|[13016]|JWDCG659-10|BOLD:AAC4536
Tanytarsus inaequalis|[13017]|CHURC865-09|BOLD:AAC4536
Tanytarsus inaequalis|[13018]|CHURC451-07|BOLD:AAC4536
Tanytarsus inaequalis|[13019]|CHURC453-07|BOLD:AAC4536
Tanytarsus inaequalis|[13020]|CHURC452-07|BOLD:AAC4536
Tanytarsus inaequalis|[13021]|CHURC454-07|BOLD:AAC4536
Tanytarsus inaequalis|[13022]|CHURC193-07|BOLD:AAC4536
Tanytarsus inaequalis|[13023]|MHMOS181-07|BOLD:AAC4536
Tanytarsus inaequalis|[13024]|CHURC947-09|BOLD:AAC4536
Tanytarsus inaequalis|[13025]|JWDCG067-10|BOLD:AAC4536
Tanytarsus inaequalis|[13026]|JWDCJ1598-11|BOLD:AAC4536
Tanytarsus inaequalis|[13027]|JWDCJ1599-11|BOLD:AAC4536
Tanytarsus inaequalis|[13028]|MHMOS095-07|BOLD:AAC4536
Tanytarsus inaequalis|[13029]|JWDCG661-10|BOLD:AAC4536
Tanytarsus inaequalis|[13030]|CHURC1018-09|BOLD:AAC4536
Tanytarsus herrmanni|[13031]|CHURC277-07|BOLD:AAC8203
Tanytarsus herrmanni|[13032]|CHURC278-07|BOLD:AAC8203
Tanytarsus herrmanni|[13033]|JWDCH711-10|BOLD:AAC8203
Tanytarsus herrmanni|[13034]|JWDCI088-10|BOLD:AAC8203
Tanytarsus herrmanni|[13035]|JWDCH948-10|BOLD:AAC8203

[illegible]
Tanytarsus herrmanni|[13034]|JWDCI088-10|BOLD:AAC8203
Tanytarsus herrmanni|[13035]|JWDCH948-10|BOLD:AAC8203
Tanytarsus herrmanni|[13036]|JWDCI095-10|BOLD:AAC8203
Tanytarsus herrmanni|[13037]|JWDCH086-10|BOLD:AAC8203
Tanytarsus herrmanni|[13038]|JWDCK105-11|BOLD:AAC8203
Tanytarsus herrmanni|[13039]|JWDCH935-10|BOLD:AAC8203
Tanytarsus herrmanni|[13040]|JWDCH931-10|BOLD:AAC8203
Tanytarsus herrmanni|[13041]|BSCHR082-10|BOLD:AAC8203
Tanytarsus herrmanni|[13042]|BSCHR076-10|BOLD:AAC8203
Tanytarsus herrmanni|[13043]|JWDCH235-10|BOLD:AAC8203
Tanytarsus herrmanni|[13044]|JWDCJ1379-11|BOLD:AAC8203
Tanytarsus herrmanni|[13045]|JWDCI185-10|BOLD:AAC8203
Tanytarsus herrmanni|[13046]|JWDCH296-10|BOLD:AAC8203
Tanytarsus herrmanni|[13047]|JWDCG443-10|BOLD:AAC8203
Tanytarsus herrmanni|[13048]|CHURC695-09|BOLD:AAC8203
Tanytarsus herrmanni|[13049]|CHURC162-07|BOLD:AAC8203
Tanytarsus herrmanni|[13050]|CHURC1073-09|BOLD:AAC8203
Tanytarsus herrmanni|[13051]|CHURC015-06|BOLD:AAC8203
Tanytarsus herrmanni|[13052]|CHURC044-06|BOLD:AAC8203
Tanytarsus herrmanni|[13053]|JWDCG663-10|BOLD:AAC8203
Tanytarsus herrmanni|[13054]|JWDCI477-10|BOLD:AAC8203
Tanytarsus herrmanni|[13055]|CHURC088-06|BOLD:AAC8203
Tanytarsus bathophilus|[13056]|CHURC624-09|BOLD:AAH2253
Tanytarsus aigos|[13057]|CHURC422-07|BOLD:AAK0228
Tanytarsus aigos|[13058]|CHURC913-09
Tanytarsus aigos|[13059]|CHURC917-09|BOLD:AAK0228
Tanytarsus|[13060]|CHURC1102-09|BOLD:AAG5492
Stempellinella cf. leptocelloides|[13061]|CHURC386-07|BOLD:AAJ4951
Tanytarsus cf. dispar|[13062]|CHURC385-07|BOLD:AAH9825
Tanytarsus cf. dispar|[13063]|CHURC391-07|BOLD:AAH9825
Tanytarsus cf. dispar|[13064]|CHURC780-09|BOLD:AAH9825
Tanytarsus|[13065]|KSCLC019-09|BOLD:AAZ2451
Tanytarsus telmaticus|[13066]|CHURC1048-09|BOLD:AAF0835
Tanytarsus mendax|[13067]|MHMOS138-07|BOLD:AAA3776
Tanytarsus mendax|[13068]|MHMOS141-07|BOLD:AAA3776
Tanytarsus mendax|[13069]|MHMOS188-07|BOLD:AAA3776
Tanytarsus mendax|[13070]|MHMOS152-07|BOLD:AAA3776
Tanytarsus mendax|[13071]|MHMOS185-07|BOLD:AAA3776
Tanytarsus mendax|[13072]|CHURC686-09|BOLD:AAA3776
Tanytarsus mendax|[13073]|CHURC699-09|BOLD:AAA3776
Tanytarsus mendax|[13074]|MHMOS142-07|BOLD:AAA3776
Tanytarsus mendax|[13075]|CHURC691-09|BOLD:AAA3776
Tanytarsus mendax|[13076]|MHMOS126-07|BOLD:AAA3776
Tanytarsus mendax|[13077]|MHMOS128-07|BOLD:AAA3776
Tanytarsus mendax|[13078]|JWDCH712-10|BOLD:AAA3776
Tanytarsus mendax|[13079]|CHURC690-09|BOLD:AAA3776
Tanytarsus mendax|[13080]|CHURC191-07|BOLD:AAA3776
Tanytarsus mendax|[13081]|MHMOS120-07|BOLD:AAA3776
Tanytarsus mendax|[13082]|JWDCG822-10|BOLD:AAA3776
Tanytarsus mendax|[13083]|CHURC259-07|BOLD:AAA3776
Tanytarsus mendax|[13084]|MHMOS139-07|BOLD:AAA3776
Tanytarsus mendax|[13085]|MHMOS176-07|BOLD:AAA3776
Tanytarsus mendax|[13086]|MHMOS179-07|BOLD:AAA3776
Tanytarsus mendax|[13087]|MHMOS173-07|BOLD:AAA3776
Tanytarsus mendax|[13088]|CHURC726-09|BOLD:AAA3776
Tanytarsus mendax|[13089]|CHURC144-07|BOLD:AAA3776
Tanytarsus mendax|[13090]|CHURC148-07|BOLD:AAA3776
Tanytarsus mendax|[13091]|JWDCK015-11|BOLD:AAA3776
Tanytarsus mendax|[13092]|JWDCH295-10|BOLD:AAA3776
Tanytarsus mendax|[13093]|CHURC1041-09|BOLD:AAA3776
Tanytarsus mendax|[13094]|MHMOS149-07|BOLD:AAA3776
Tanytarsus mendax|[13095]|MHMOS143-07|BOLD:AAA3776
Tanytarsus mendax|[13096]|JWDCH294-10|BOLD:AAA3776
Tanytarsus mendax|[13097]|JWDCH196-10|BOLD:AAA3776
Tanytarsus mendax|[13098]|JWDCG824-10|BOLD:AAA3776
Tanytarsus mendax|[13099]|JWDCI089-10|BOLD:AAA3776
Tanytarsus mendax|[13100]|MHMOS125-07|BOLD:AAA3776
Tanytarsus mendax|[13101]|JWDCI092-10|BOLD:AAA3776
Tanytarsus mendax|[13102]|CHURC160-07|BOLD:AAA3776
Tanytarsus mendax|[13103]|CHURC1049-09|BOLD:AAA3776
Tanytarsus mendax|[13104]|MHMOS119-07|BOLD:AAA3776
Tanytarsus mendax|[13105]|MHMOS137-07|BOLD:AAA3776
Tanytarsus mendax|[13106]|MHMOS172-07|BOLD:AAA3776
Tanytarsus mendax|[13107]|MHMOS175-07|BOLD:AAA3776
Tanytarsus mendax|[13108]|MHMOS121-07|BOLD:AAA3776
Tanytarsus mendax|[13109]|JWDCH132-10|BOLD:AAA3776
Tanytarsus mendax|[13110]|JWDCI087-10|BOLD:AAA3776
Tanytarsus mendax|[13111]|CHURC698-09|BOLD:AAA3776
Tanytarsus mendax|[13112]|MHMOS174-07|BOLD:AAA3776
Tanytarsus mendax|[13113]|CHURC118-07|BOLD:AAA3776
Tanytarsus mendax|[13114]|MHMOS147-07|BOLD:AAA3776
Tanytarsus mendax|[13115]|CHURC685-09|BOLD:AAA3776
Tanytarsus mendax|[13116]|CHURC258-07|BOLD:AAA3776
Tanytarsus mendax|[13117]|MHMOS187-07|BOLD:AAA3776
Tanytarsus mendax|[13118]|JWDCG373-10|BOLD:AAA3776
Tanytarsus mendax|[13119]|MHMOS134-07|BOLD:AAA3776
Tanytarsus mendax|[13120]|MHMOS131-07|BOLD:AAA3776
Tanytarsus mendax|[13121]|CHURC696-09|BOLD:AAA3776
Tanytarsus mendax|[13122]|CHURC824-09|BOLD:AAA3776
Tanytarsus mendax|[13123]|CHURC012-06|BOLD:AAA3776
Tanytarsus mendax|[13124]|CHURC688-09|BOLD:AAA3777
Tanytarsus mendax|[13125]|JWDCI507-10|BOLD:AAQ0623
Tanytarsus mendax|[13126]|JWDCI093-10|BOLD:AAA3774
Tanytarsus mendax|[13127]|JWDCG658-10|BOLD:AAA3774
Tanytarsus mendax|[13128]|CHURC1043-09|BOLD:AAA3774
Tanytarsus mendax|[13129]|MHMOS132-07|BOLD:AAA3774
Tanytarsus mendax|[13130]|MHMOS180-07|BOLD:AAA3774
Tanytarsus mendax|[13131]|MHMOS113-07|BOLD:AAA3774
Tanytarsus mendax|[13132]|MHMOS127-07|BOLD:AAA3774
Tanytarsus mendax|[13133]|CHURC700-09|BOLD:AAA3774
Tanytarsus mendax|[13134]|JWDCJ1461-11|BOLD:AAA3774
Tanytarsus mendax|[13135]|CHURC189-07|BOLD:AAA3774

Tanytarsus mendax|[13133]|CHURC700-09|BOLD:AAA3774
Tanytarsus mendax|[13134]|JWDCJ1461-11|BOLD:AAA3774
Tanytarsus mendax|[13135]|CHURC189-07|BOLD:AAA3774
Tanytarsus mendax|[13136]|CHURC762-09|BOLD:AAA3774
Tanytarsus mendax|[13137]|JWDCH709-10|BOLD:AAA3774
Tanytarsus mendax|[13138]|CHURC222-07|BOLD:AAA3774
Tanytarsus mendax|[13139]|CHURC720-09|BOLD:AAA3774
Tanytarsus mendax|[13140]|CHURC798-09|BOLD:AAA3774
Tanytarsus mendax|[13141]|CHURC803-09|BOLD:AAA3774
Tanytarsus mendax|[13142]|CHURC746-09|BOLD:AAA3774
Tanytarsus mendax|[13143]|CHURC039-06|BOLD:AAA3774
Tanytarsus mendax|[13144]|JWDCG446-10|BOLD:AAA3774
Tanytarsus mendax|[13145]|CHURC751-09|BOLD:AAA3774
Tanytarsus mendax|[13146]|CHURC250-07|BOLD:AAA3774
Tanytarsus mendax|[13147]|CHURC722-09|BOLD:AAA3774
Tanytarsus mendax|[13148]|CHURC795-09|BOLD:AAA3774
Tanytarsus mendax|[13149]|CHURC737-09|BOLD:AAA3774
Tanytarsus mendax|[13150]|CHURC792-09|BOLD:AAA3774
Tanytarsus mendax|[13151]|CHURC099-07|BOLD:AAA3774
Tanytarsus mendax|[13152]|CHURC120-07|BOLD:AAA3774
Tanytarsus mendax|[13153]|MHMOS182-07|BOLD:AAA3774
Tanytarsus mendax|[13154]|CHURC221-07|BOLD:AAA3774
Tanytarsus mendax|[13155]|CHURC721-09|BOLD:AAA3774
Tanytarsus mendax|[13156]|JWDCH788-10|BOLD:AAA3774
Tanytarsus mendax|[13157]|CHURC249-07|BOLD:AAA3774
Tanytarsus mendax|[13158]|CHURC743-09|BOLD:AAA3774
Tanytarsus mendax|[13159]|CHURC038-06|BOLD:AAA3774
Tanytarsus mendax|[13160]|CHURC804-09|BOLD:AAA3774
Tanytarsus mendax|[13161]|CHURC773-09|BOLD:AAA3774
Tanytarsus mendax|[13162]|CHURC192-07|BOLD:AAA3774
Tanytarsus mendax|[13163]|CHURC794-09|BOLD:AAA3774
Tanytarsus mendax|[13164]|CHURC755-09|BOLD:AAA3774
Tanytarsus mendax|[13165]|CHURC819-09|BOLD:AAA3774
Tanytarsus mendax|[13166]|CHURC199-07|BOLD:AAA3774
Tanytarsus mendax|[13167]|CHURC223-07|BOLD:AAA3774
Tanytarsus mendax|[13168]|CHURC478-08|BOLD:AAA3774
Tanytarsus mendax|[13169]|CHURC1047-09|BOLD:AAA3774
Tanytarsus mendax|[13170]|CHURC815-09|BOLD:AAA3774
Tanytarsus mendax|[13171]|CHURC194-07|BOLD:AAA3774
Tanytarsus mendax|[13172]|JWDCI508-10|BOLD:AAA3774
Tanytarsus mendax|[13173]|CHURC261-07|BOLD:AAA3774
Tanytarsus mendax|[13174]|CHURC006-06|BOLD:AAA3774
Tanytarsus mendax|[13175]|JWDCG371-10|BOLD:AAA3774
Tanytarsus mendax|[13176]|CHURC007-06|BOLD:AAA3774
Tanytarsus mendax|[13177]|CHURC758-09|BOLD:AAA3774
Tanytarsus mendax|[13178]|CHURC740-09|BOLD:AAA3774
Tanytarsus mendax|[13179]|CHURC800-09|BOLD:AAA3774
Tanytarsus mendax|[13180]|CHURC008-06|BOLD:AAA3774
Tanytarsus mendax|[13181]|CHURC014-06|BOLD:AAA3774
Tanytarsus|[13182]|CHURC727-09|BOLD:AAH2254
Tanytarsus wirthi|[13183]|CHURC197-07|BOLD:AAD2144
Tanytarsus wirthi|[13184]|CHURC411-07|BOLD:AAD2144
Tanytarsus wirthi|[13185]|CHURC801-09|BOLD:AAD2144
Tanytarsus wirthi|[13186]|CHURC831-09|BOLD:AAD2144
Tanytarsus wirthi|[13187]|CHURC101-07|BOLD:AAD2144
Tanytarsus wirthi|[13188]|CHURC1044-09|BOLD:AAD2144
Tanytarsus wirthi|[13189]|CHURC820-09|BOLD:AAD2144
Tanytarsus wirthi|[13190]|CHURC849-09|BOLD:AAD2144
Tanytarsus dendyi|[13191]|CHURC748-09|BOLD:ADF0632
Tanytarsus dendyi|[13192]|CHURC260-07|BOLD:ADF0632
Tanytarsus dendyi|[13193]|CHURC814-09|BOLD:ADF0632
Tanytarsus dendyi|[13194]|CHURC513-08|BOLD:AAC6577
Tanytarsus dendyi|[13195]|CHURC518-08|BOLD:AAC6577
Tanytarsus dendyi|[13196]|CHURC725-09|BOLD:AAC6577
Tanytarsus dendyi|[13197]|CHURC200-07|BOLD:AAC6577
Tanytarsus dendyi|[13198]|CHURC515-08|BOLD:AAC6577
Tanytarsus dendyi|[13199]|CHURC035-06|BOLD:AAC6577
Tanytarsus dendyi|[13200]|CHURC047-06|BOLD:ADF0634
Phaenopsectra punctipes|[13201]|CHURC929-09|BOLD:AAD9171
Phaenopsectra punctipes|[13202]|CHURC225-07|BOLD:AAD9171
Phaenopsectra punctipes|[13203]|CHURC309-07|BOLD:AAD9171
Phaenopsectra punctipes|[13204]|CHURC326-07|BOLD:AAD9171
Phaenopsectra punctipes|[13205]|CHURC032-06|BOLD:AAD9171
Phaenopsectra punctipes|[13206]|CHURC033-06|BOLD:AAD9171
Paralauterborniella nigrohalteralis|[13207]|CHURC510-08|BOLD:AAI6341
Paralauterborniella nigrohalteralis|[13208]|CHURC301-07|BOLD:AAI6341
Microtendipes lugubris|[13209]|JWDCE740-10|BOLD:AAG5479
Microtendipes lugubris|[13210]|BSCHR154-10|BOLD:AAG5479
Microtendipes lugubris|[13211]|JWDCE379-10|BOLD:AAG5479
Microtendipes lugubris|[13212]|BSCHR147-10|BOLD:AAG5479
Microtendipes lugubris|[13213]|BSCHR133-10|BOLD:AAG5479
Microtendipes lugubris|[13214]|BSCHR117-10|BOLD:AAG5479
Microtendipes lugubris|[13215]|JWDCE615-10|BOLD:AAG5479
Microtendipes lugubris|[13216]|JWDCE309-10|BOLD:AAG5479
Microtendipes lugubris|[13217]|BSCHR184-10|BOLD:AAG5479
Microtendipes lugubris|[13218]|BSCHR176-10|BOLD:AAG5479
Microtendipes lugubris|[13219]|BSCHR178-10|BOLD:AAG5479
Microtendipes lugubris|[13220]|BSCHR181-10|BOLD:AAG5479
Microtendipes lugubris|[13221]|BSCHR171-10|BOLD:AAG5479
Microtendipes lugubris|[13222]|BSCHR175-10|BOLD:AAG5479
Microtendipes lugubris|[13223]|BSCHR166-10|BOLD:AAG5479
Microtendipes lugubris|[13224]|BSCHR167-10|BOLD:AAG5479
Microtendipes lugubris|[13225]|BSCHR161-10|BOLD:AAG5479
Microtendipes lugubris|[13226]|BSCHR162-10|BOLD:AAG5479
Microtendipes lugubris|[13227]|BSCHR142-10|BOLD:AAG5479
Microtendipes lugubris|[13228]|BSCHR160-10|BOLD:AAG5479
Microtendipes lugubris|[13229]|JWDCE588-10|BOLD:AAG5479
Microtendipes lugubris|[13230]|BSCHR140-10|BOLD:AAG5479
Microtendipes lugubris|[13231]|BSCHR137-10|BOLD:AAG5479
Microtendipes lugubris|[13232]|BSCHR136-10|BOLD:AAG5479
Microtendipes lugubris|[13233]|BSCHR123-10|BOLD:AAG5479
Microtendipes lugubris|[13234]|BSCHR119-10|BOLD:AAG5479
Microtendipes lugubris|[13235]|BSCHR118-10|BOLD:AAG5479

Microtendipes lugubris|[13232]|BSCHR130-10|BOLD:AAG5479
Microtendipes lugubris|[13233]|BSCHR123-10|BOLD:AAG5479
Microtendipes lugubris|[13234]|BSCHR119-10|BOLD:AAG5479
Microtendipes lugubris|[13235]|BSCHR118-10|BOLD:AAG5479
Microtendipes lugubris|[13236]|BSCHR100-10|BOLD:AAG5479
Microtendipes lugubris|[13237]|BSCHR099-10|BOLD:AAG5479
Microtendipes lugubris|[13238]|BSCHR096-10|BOLD:AAG5479
Microtendipes lugubris|[13239]|JWDCE468-10|BOLD:AAG5479
Microtendipes lugubris|[13240]|BSCHR097-10|BOLD:AAG5479
Microtendipes lugubris|[13241]|JWDCE859-10|BOLD:AAG5479
Microtendipes|[13242]|CHURC566-09|BOLD:AAG5479
Microtendipes|[13243]|CHURC567-09|BOLD:AAG5479
Microtendipes chloris|[13244]|BSCHR051-10|BOLD:AAN5371
Microtendipes chloris|[13245]|BSCHR131-10|BOLD:AAN5371
Microtendipes chloris|[13246]|BSCHR070-10|BOLD:AAN5371
Microtendipes chloris|[13247]|BSCHR050-10|BOLD:AAN5371
Microtendipes chloris|[13248]|BSCHR048-10|BOLD:AAN5371
Polypedilum sp. TE01|[13249]|CHURC444-07|BOLD:AAG5511
Polypedilum sp. TE01|[13250]|CHURC489-08|BOLD:AAG5511
Pseudochironomus crassus|[13251]|JWDCF670-10|BOLD:AAQ0620
Pseudochironomus crassus|[13252]|JWDCJ1156-11|BOLD:AAV5952
Pseudochironomus prasinatus|[13253]|JWDCI406-10|BOLD:AAV5962
Polypedilum sordens|[13254]|BSCHR177-10|BOLD:ACY3854
Polypedilum|[13255]|BSCHR188-10|BOLD:AAC4537
Polypedilum|[13256]|BSCHR189-10|BOLD:AAC4537
Polypedilum|[13257]|KSCLC005-09|BOLD:AAC4537
Polypedilum|[13258]|BSCHR134-10|BOLD:AAN5384
Polypedilum quadriguttatum|[13259]|CHURC304-07|BOLD:AAE4099
Polypedilum|[13260]|SWSWP037-09|BOLD:AAE4099
Polypedilum quadriguttatum|[13261]|CHURC500-08|BOLD:AAE4099
Polypedilum quadriguttatum|[13262]|CHURC447-07|BOLD:AAE4100
Polypedilum quadriguttatum|[13263]|CHURC448-07|BOLD:AAE4100
Polypedilum sp. TE03|[13264]|CHURC556-08|BOLD:AAY3393
Polypedilum sp. TE02|[13265]|CHURC100-07|BOLD:AAY3407
Chironominae|[13266]|BSCHR084-10|BOLD:AAG5457
Chironominae|[13267]|JWDCL226-11|BOLD:AAG5457
Chironominae|[13268]|JWDCA660-10|BOLD:AAG5457
Chironominae|[13269]|BSCHR001-10|BOLD:AAG5457
Chironominae|[13270]|JWDCL391-11|BOLD:AAG5457
Chironominae|[13271]|JWDCC913-10|BOLD:AAG5457
Chironominae|[13272]|JWDCC906-10|BOLD:AAG5457
Chironominae|[13273]|JWDCF065-10|BOLD:AAG5457
Chironominae|[13274]|JWDCJ658-11|BOLD:AAG5457
Chironominae|[13275]|JWDCI861-11|BOLD:AAG5457
Chironominae|[13276]|BSCHR016-10|BOLD:AAG5457
Chironominae|[13277]|JWDCE646-10|BOLD:AAG5457
Chironominae|[13278]|BSCHR026-10|BOLD:AAG5457
Chironominae|[13279]|JWDCA631-10|BOLD:AAG5457
Chironominae|[13280]|JWDCE308-10|BOLD:AAG5457
Chironominae|[13281]|KSCLC003-09|BOLD:AAG5457
Chironominae|[13282]|JWDCF637-10|BOLD:AAG5457
Chironominae|[13283]|JWDCF634-10|BOLD:AAG5457
Chironominae|[13284]|BSCHR055-10|BOLD:AAG5457
Chironominae|[13285]|JWDCJ656-11|BOLD:AAG5457
Endochironomus nigricans|[13286]|CHURC397-07|BOLD:AAC9540
Endochironomus nigricans|[13287]|CHURC906-09|BOLD:AAC9540
Endochironomus nigricans|[13288]|CHURC380-07|BOLD:AAC9540
Endochironomus nigricans|[13289]|JWDCJ974-11|BOLD:AAC9540
Endochironomus nigricans|[13290]|BSCHR002-10|BOLD:AAC9540
Endochironomus nigricans|[13291]|BSCHR072-10|BOLD:AAC9540
Endochironomus nigricans|[13292]|JWDCA665-10|BOLD:AAC9540
Endochironomus nigricans|[13293]|JWDCA662-10|BOLD:AAC9540
Endochironomus nigricans|[13294]|JWDCA657-10|BOLD:AAC9540
Endochironomus nigricans|[13295]|CHURC967-09|BOLD:AAC9540
Endochironomus nigricans|[13296]|CHURC975-09|BOLD:AAC9540
Polypedilum tritum|[13297]|JWDCF396-10|BOLD:AAE4119
Polypedilum tritum|[13298]|CHURC328-07|BOLD:AAE4119
Polypedilum tritum|[13299]|CHURC400-07|BOLD:AAE4119
Polypedilum tritum|[13300]|CHURC457-07|BOLD:AAE4119
Polypedilum tritum|[13301]|CHURC583-09|BOLD:AAE4119
Polypedilum tritum|[13302]|CHURC432-07|BOLD:AAE4119
Cyphomella cornea|[13303]|CHURC668-09|BOLD:AAI9470
Cyphomella cornea|[13304]|CHURC529-08|BOLD:AAI9469
Chironominae|[13305]|JWDCF481-10|BOLD:AAP5149
Chironominae|[13306]|JWDCF530-10|BOLD:AAP5149
Chironominae|[13307]|JWDCJ724-11|BOLD:AAP5149
Chironominae|[13308]|JWDCE841-10|BOLD:AAP5149
Chironominae|[13309]|JWDCE842-10|BOLD:AAP5149
Polypedilum cultellatum|[13310]|JWDCK106-11|BOLD:AAH7761
Polypedilum cultellatum|[13311]|CHURC395-07|BOLD:AAH7761
Polypedilum cultellatum|[13312]|CHURC413-07|BOLD:AAH7761
Demicryptochironomus cf. evgenii|[13313]|CHURC876-09|BOLD:AAL1205
Xenochironomus xenolabis|[13314]|CHURC1042-09|BOLD:AAG8483
Dicrotendipes|[13315]|SWSWP050-09|BOLD:AAG5502
Cladopelma sp. TE02|[13316]|JWDCF070-10|BOLD:AAI4126
Cladopelma sp. TE02|[13317]|JWDCF732-10|BOLD:AAI4126
Cladopelma sp. TE02|[13318]|JWDCF735-10|BOLD:AAI4126
Cladopelma sp. TE02|[13319]|CHURC339-07|BOLD:AAI4126
Cladopelma viridulum|[13320]|SWSWP031-09|BOLD:AAD7364
Cladopelma viridulum|[13321]|CHURC331-07|BOLD:AAD7363
Cladopelma viridulum|[13322]|CHURC420-07|BOLD:AAD7363
Cladopelma viridulum|[13323]|CHURC394-07|BOLD:AAD7363
Cladopelma viridulum|[13324]|CHURC330-07|BOLD:AAD7363
Cladopelma viridulum|[13325]|CHURC905-09|BOLD:AAD7363
Chironominae|[13326]|BSCHR174-10|BOLD:AAH0044
Chironominae|[13327]|CHURC1005-09|BOLD:AAH0044
Dicrotendipes modestus|[13328]|BSCHR065-10|BOLD:AAN5374
Dicrotendipes modestus|[13329]|KSCLC046-09|BOLD:AAC5069
Dicrotendipes modestus|[13330]|BSCHR130-10|BOLD:AAC5069
Dicrotendipes modestus|[13331]|JWDCF179-10|BOLD:AAC5069
Dicrotendipes modestus|[13332]|BSCHR152-10|BOLD:AAC5069
Dicrotendipes modestus|[13333]|CHURC284-07|BOLD:AAC5069
Dicrotendipes modestus|[13334]|CHURC087-06|BOLD:AAC5069

Dicrotendipes modestus|[13332]|BSCHR152-10|BOLD:AAC5069
Dicrotendipes modestus|[13333]|CHURC284-07|BOLD:AAC5069
Dicrotendipes modestus|[13334]|CHURC087-06|BOLD:AAC5069
Dicrotendipes modestus|[13335]|JWDCF230-10|BOLD:AAC5069
Dicrotendipes modestus|[13336]|JWDCF783-10|BOLD:AAC5069
Dicrotendipes modestus|[13337]|CHURC098-07|BOLD:AAC5069
Dicrotendipes modestus|[13338]|CHURC283-07|BOLD:AAC5069
Chironominae|[13339]|BSCHR069-10|BOLD:AAN5375
Dicrotendipes nervosus|[13340]|CHURC534-08|BOLD:AAI9518
Dicrotendipes nervosus|[13341]|CHURC503-08|BOLD:AAI9518
Dicrotendipes|[13342]|JWDCC064-10|BOLD:AAN5308
Dicrotendipes|[13343]|JWDCC658-10|BOLD:AAN5308
Dicrotendipes modestus|[13344]|JWDCC349-10|BOLD:ACE8918
Dicrotendipes modestus|[13345]|JWDCF707-10|BOLD:ACE8918
Dicrotendipes modestus|[13346]|JWDCJ410-11|BOLD:ACE8918
Dicrotendipes modestus|[13347]|BSCHR008-10|BOLD:ACE8918
Dicrotendipes modestus|[13348]|JWDCC497-10|BOLD:ACE8918
Dicrotendipes modestus|[13349]|JWDCC379-10|BOLD:ACE8918
Dicrotendipes modestus|[13350]|CHURC893-09|BOLD:ACE8918
Dicrotendipes modestus|[13351]|JWDCF266-10|BOLD:AAC5070
Dicrotendipes modestus|[13352]|JWDCJ765-11|BOLD:AAC5070
Dicrotendipes modestus|[13353]|CHURC321-07|BOLD:AAC5070
Dicrotendipes modestus|[13354]|JWDCK219-11|BOLD:AAC5070
Dicrotendipes modestus|[13355]|CHURC484-08|BOLD:AAC5070
Dicrotendipes modestus|[13356]|JWDCC348-10|BOLD:AAC5070
Dicrotendipes modestus|[13357]|CHURC544-08|BOLD:AAC5070
Dicrotendipes modestus|[13358]|JWDCC295-10|BOLD:AAC5070
Dicrotendipes modestus|[13359]|JWDCJ409-11|BOLD:AAC5070
Dicrotendipes modestus|[13360]|BSCHR120-10|BOLD:AAC5070
Dicrotendipes modestus|[13361]|BSCHR101-10|BOLD:AAC5070
Dicrotendipes modestus|[13362]|BSCHR098-10|BOLD:AAC5070
Dicrotendipes modestus|[13363]|BSCHR029-10|BOLD:AAC5070
Dicrotendipes modestus|[13364]|BSCHR027-10|BOLD:AAC5070
Dicrotendipes modestus|[13365]|BSCHR013-10|BOLD:AAC5070
Dicrotendipes modestus|[13366]|JWDCC659-10|BOLD:AAC5070
Dicrotendipes modestus|[13367]|JWDCC297-10|BOLD:AAC5070
Dicrotendipes modestus|[13368]|JWDCC655-10|BOLD:AAC5070
Dicrotendipes modestus|[13369]|CHURC1085-09|BOLD:AAC5070
Dicrotendipes modestus|[13370]|JWDCE720-10|BOLD:AAC5070
Dicrotendipes modestus|[13371]|CHURC894-09|BOLD:AAC5070
Dicrotendipes modestus|[13372]|JWDCJ435-11|BOLD:AAC5070
Dicrotendipes modestus|[13373]|JWDCC656-10|BOLD:AAC5070
Dicrotendipes modestus|[13374]|BSCHR018-10|BOLD:AAC5070
Dicrotendipes modestus|[13375]|JWDCC353-10|BOLD:AAC5070
Dicrotendipes modestus|[13376]|JWDCC380-10|BOLD:AAC5070
Dicrotendipes modestus|[13377]|SWSWP035-09|BOLD:AAC5070
Dicrotendipes modestus|[13378]|JWDCC351-10|BOLD:AAC5070
Dicrotendipes modestus|[13379]|JWDCC293-10|BOLD:AAC5070
Dicrotendipes modestus|[13380]|JWDCC292-10|BOLD:AAC5070
Dicrotendipes modestus|[13381]|JWDCC291-10|BOLD:AAC5070
Dicrotendipes modestus|[13382]|JWDCD110-10|BOLD:AAC5070
Dicrotendipes modestus|[13383]|JWDCC294-10|BOLD:AAC5070
Dicrotendipes modestus|[13384]|BSCHR139-10|BOLD:AAC5070
Dicrotendipes modestus|[13385]|JWDCC657-10|BOLD:AAC5070
Dicrotendipes modestus|[13386]|JWDCC499-10|BOLD:AAC5070
Dicrotendipes modestus|[13387]|JWDCC492-10|BOLD:AAC5070
Dicrotendipes modestus|[13388]|JWDCC491-10|BOLD:AAC5070
Dicrotendipes modestus|[13389]|SWSWP078-09|BOLD:AAC5070
Dicrotendipes modestus|[13390]|JWDCE142-10|BOLD:AAC5070
Dicrotendipes modestus|[13391]|CHURC678-09|BOLD:ACE8918
Dicrotendipes modestus|[13392]|CHURC571-09|BOLD:ACE8918
Dicrotendipes modestus|[13393]|CHURC586-09|BOLD:AAC5070
Dicrotendipes modestus|[13394]|SWSWP052-09|BOLD:AAC5070
Dicrotendipes modestus|[13395]|JWDCF704-10|BOLD:AAC5070
Dicrotendipes modestus|[13396]|JWDCA636-10|BOLD:AAC5070
Dicrotendipes modestus|[13397]|JWDCD267-10|BOLD:AAC5070
Dicrotendipes modestus|[13398]|JWDCE912-10|BOLD:AAC5070
Dicrotendipes modestus|[13399]|BSCHR172-10|BOLD:AAC5070
Dicrotendipes modestus|[13400]|CHURC1100-09|BOLD:AAC5070
Dicrotendipes sp. 2TE|[13401]|BSCHR135-10|BOLD:AAN5385
Dicrotendipes|[13402]|BSCHR129-10|BOLD:AAN5383
Dicrotendipes|[13403]|JWDCF114-10|BOLD:AAP6882
Dicrotendipes|[13404]|JWDCF730-10|BOLD:AAP6882
Dicrotendipes tritomus|[13405]|CHURC1003-09|BOLD:AAC0706
Dicrotendipes tritomus|[13406]|CHURC988-09|BOLD:AAC0706
Dicrotendipes tritomus|[13407]|CHURC1015-09|BOLD:AAC0706
Dicrotendipes tritomus|[13408]|CHURC1120-09|BOLD:AAC0706
Dicrotendipes tritomus|[13409]|CHURC1014-09|BOLD:AAC0706
Dicrotendipes tritomus|[13410]|CHURC1011-09|BOLD:AAC0706
Dicrotendipes tritomus|[13411]|CHURC983-09|BOLD:AAC0706
Dicrotendipes tritomus|[13412]|CHURC976-09|BOLD:AAC0706
Dicrotendipes tritomus|[13413]|CHURC966-09|BOLD:AAC0706
Dicrotendipes tritomus|[13414]|CHURC954-09|BOLD:AAC0706
Dicrotendipes tritomus|[13415]|CHURC1123-09|BOLD:AAC0706
Dicrotendipes tritomus|[13416]|CHURC949-09|BOLD:AAC0706
Dicrotendipes tritomus|[13417]|CHURC955-09|BOLD:AAC0706
Dicrotendipes tritomus|[13418]|CHURC1124-09|BOLD:AAC0706
Paracladopelma winnelli|[13419]|MHMOS100-07|BOLD:AAK4963
Paracladopelma winnelli|[13420]|BSCHR127-10|BOLD:AAK4963
Paracladopelma winnelli|[13421]|BSCHR170-10|BOLD:AAK4963
Paracladopelma winnelli|[13422]|CHURC173-07|BOLD:AAK4963
Cryptochironomus|[13423]|JWDCF876-10|BOLD:ABX6314
Cryptochironomus sp. TE02|[13424]|JWDCI622-10|BOLD:AAF1504
Cryptochironomus digitatus|[13425]|CHURC379-07|BOLD:AAF1504
Cryptochironomus digitatus|[13426]|CHURC378-07|BOLD:AAF1504
Cryptochironomus digitatus|[13427]|CHURC870-09|BOLD:AAF1504
Cryptochironomus|[13428]|BSCHR122-10|BOLD:AAN5381
Cryptochironomus psittacinus|[13429]|BSCHR045-10|BOLD:AAN5370
Cryptochironomus psittacinus|[13430]|JWDCE699-10|BOLD:AAN5370
Cryptochironomus psittacinus|[13431]|JWDCI582-10|BOLD:AAN5370
Cryptochironomus psittacinus|[13432]|CHURC433-07|BOLD:AAE9464
Cryptochironomus psittacinus|[13433]|CHURC502-08|BOLD:AAE9464
Cryptochironomus psittacinus|[13434]|BSCHR083-10|BOLD:[illegible]

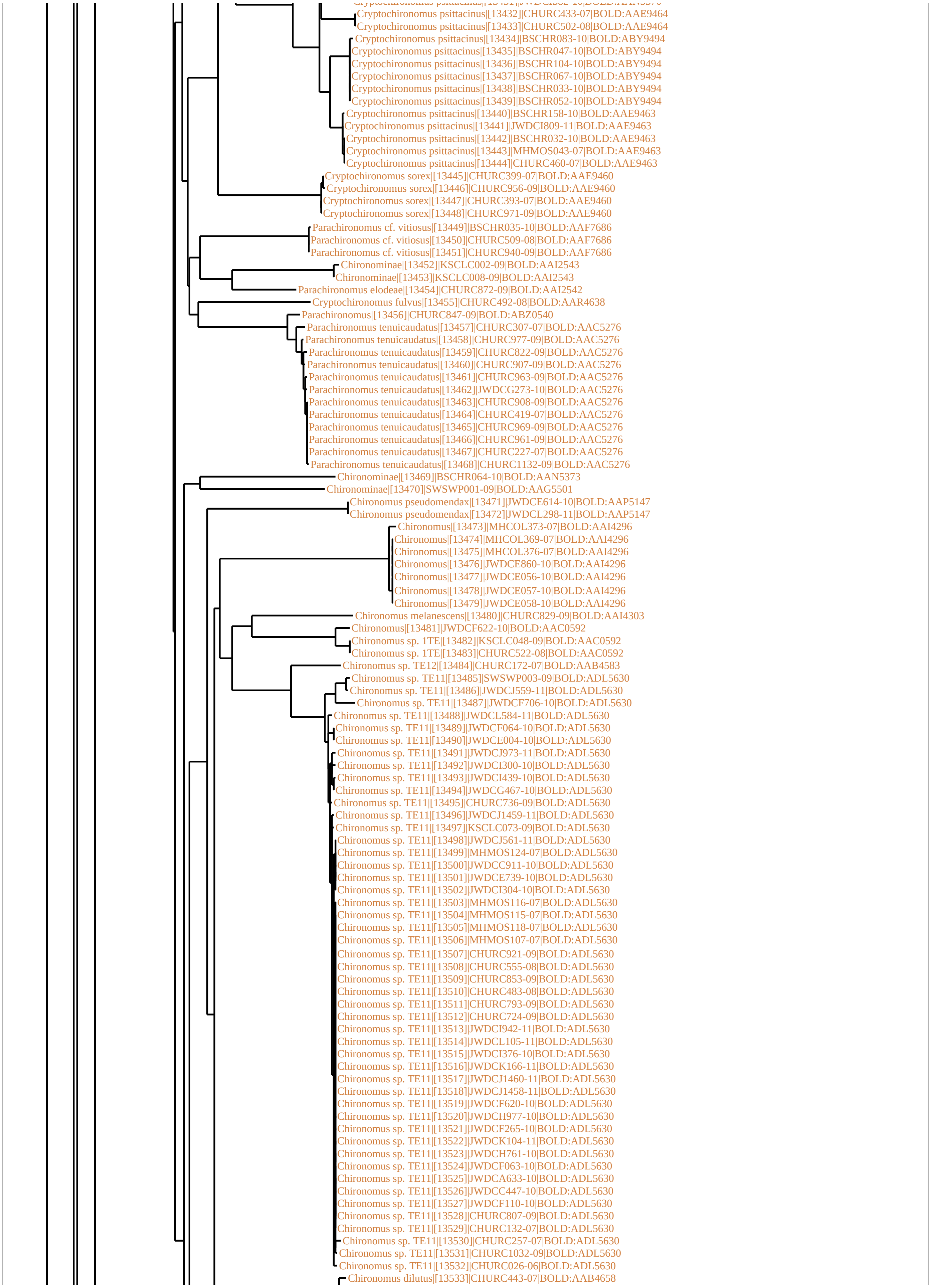
Cryptochironomus psittacinus|[illegible]
Cryptochironomus psittacinus|[13432]|CHURC433-07|BOLD:AAE9464
Cryptochironomus psittacinus|[13433]|CHURC502-08|BOLD:AAE9464
Cryptochironomus psittacinus|[13434]|BSCHR083-10|BOLD:ABY9494
Cryptochironomus psittacinus|[13435]|BSCHR047-10|BOLD:ABY9494
Cryptochironomus psittacinus|[13436]|BSCHR104-10|BOLD:ABY9494
Cryptochironomus psittacinus|[13437]|BSCHR067-10|BOLD:ABY9494
Cryptochironomus psittacinus|[13438]|BSCHR033-10|BOLD:ABY9494
Cryptochironomus psittacinus|[13439]|BSCHR052-10|BOLD:ABY9494
Cryptochironomus psittacinus|[13440]|BSCHR158-10|BOLD:AAE9463
Cryptochironomus psittacinus|[13441]|JWDCI809-11|BOLD:AAE9463
Cryptochironomus psittacinus|[13442]|BSCHR032-10|BOLD:AAE9463
Cryptochironomus psittacinus|[13443]|MHMOS043-07|BOLD:AAE9463
Cryptochironomus psittacinus|[13444]|CHURC460-07|BOLD:AAE9463
Cryptochironomus sorex|[13445]|CHURC399-07|BOLD:AAE9460
Cryptochironomus sorex|[13446]|CHURC956-09|BOLD:AAE9460
Cryptochironomus sorex|[13447]|CHURC393-07|BOLD:AAE9460
Cryptochironomus sorex|[13448]|CHURC971-09|BOLD:AAE9460
Parachironomus cf. vitiosus|[13449]|BSCHR035-10|BOLD:AAF7686
Parachironomus cf. vitiosus|[13450]|CHURC509-08|BOLD:AAF7686
Parachironomus cf. vitiosus|[13451]|CHURC940-09|BOLD:AAF7686
Chironominae|[13452]|KSCLC002-09|BOLD:AAI2543
Chironominae|[13453]|KSCLC008-09|BOLD:AAI2543
Parachironomus elodeae|[13454]|CHURC872-09|BOLD:AAI2542
Cryptochironomus fulvus|[13455]|CHURC492-08|BOLD:AAR4638
Parachironomus|[13456]|CHURC847-09|BOLD:ABZ0540
Parachironomus tenuicaudatus|[13457]|CHURC307-07|BOLD:AAC5276
Parachironomus tenuicaudatus|[13458]|CHURC977-09|BOLD:AAC5276
Parachironomus tenuicaudatus|[13459]|CHURC822-09|BOLD:AAC5276
Parachironomus tenuicaudatus|[13460]|CHURC907-09|BOLD:AAC5276
Parachironomus tenuicaudatus|[13461]|CHURC963-09|BOLD:AAC5276
Parachironomus tenuicaudatus|[13462]|JWDCG273-10|BOLD:AAC5276
Parachironomus tenuicaudatus|[13463]|CHURC908-09|BOLD:AAC5276
Parachironomus tenuicaudatus|[13464]|CHURC419-07|BOLD:AAC5276
Parachironomus tenuicaudatus|[13465]|CHURC969-09|BOLD:AAC5276
Parachironomus tenuicaudatus|[13466]|CHURC961-09|BOLD:AAC5276
Parachironomus tenuicaudatus|[13467]|CHURC227-07|BOLD:AAC5276
Parachironomus tenuicaudatus|[13468]|CHURC1132-09|BOLD:AAC5276
Chironominae|[13469]|BSCHR064-10|BOLD:AAN5373
Chironominae|[13470]|SWSWP001-09|BOLD:AAG5501
Chironomus pseudomendax|[13471]|JWDCE614-10|BOLD:AAP5147
Chironomus pseudomendax|[13472]|JWDCL298-11|BOLD:AAP5147
Chironomus|[13473]|MHCOL373-07|BOLD:AAI4296
Chironomus|[13474]|MHCOL369-07|BOLD:AAI4296
Chironomus|[13475]|MHCOL376-07|BOLD:AAI4296
Chironomus|[13476]|JWDCE860-10|BOLD:AAI4296
Chironomus|[13477]|JWDCE056-10|BOLD:AAI4296
Chironomus|[13478]|JWDCE057-10|BOLD:AAI4296
Chironomus|[13479]|JWDCE058-10|BOLD:AAI4296
Chironomus melanescens|[13480]|CHURC829-09|BOLD:AAI4303
Chironomus|[13481]|JWDCF622-10|BOLD:AAC0592
Chironomus sp. 1TE|[13482]|KSCLC048-09|BOLD:AAC0592
Chironomus sp. 1TE|[13483]|CHURC522-08|BOLD:AAC0592
Chironomus sp. TE12|[13484]|CHURC172-07|BOLD:AAB4583
Chironomus sp. TE11|[13485]|SWSWP003-09|BOLD:ADL5630
Chironomus sp. TE11|[13486]|JWDCJ559-11|BOLD:ADL5630
Chironomus sp. TE11|[13487]|JWDCF706-10|BOLD:ADL5630
Chironomus sp. TE11|[13488]|JWDCL584-11|BOLD:ADL5630
Chironomus sp. TE11|[13489]|JWDCF064-10|BOLD:ADL5630
Chironomus sp. TE11|[13490]|JWDCE004-10|BOLD:ADL5630
Chironomus sp. TE11|[13491]|JWDCJ973-11|BOLD:ADL5630
Chironomus sp. TE11|[13492]|JWDCI300-10|BOLD:ADL5630
Chironomus sp. TE11|[13493]|JWDCI439-10|BOLD:ADL5630
Chironomus sp. TE11|[13494]|JWDCG467-10|BOLD:ADL5630
Chironomus sp. TE11|[13495]|CHURC736-09|BOLD:ADL5630
Chironomus sp. TE11|[13496]|JWDCJ1459-11|BOLD:ADL5630
Chironomus sp. TE11|[13497]|KSCLC073-09|BOLD:ADL5630
Chironomus sp. TE11|[13498]|JWDCJ561-11|BOLD:ADL5630
Chironomus sp. TE11|[13499]|MHMOS124-07|BOLD:ADL5630
Chironomus sp. TE11|[13500]|JWDCC911-10|BOLD:ADL5630
Chironomus sp. TE11|[13501]|JWDCE739-10|BOLD:ADL5630
Chironomus sp. TE11|[13502]|JWDCI304-10|BOLD:ADL5630
Chironomus sp. TE11|[13503]|MHMOS116-07|BOLD:ADL5630
Chironomus sp. TE11|[13504]|MHMOS115-07|BOLD:ADL5630
Chironomus sp. TE11|[13505]|MHMOS118-07|BOLD:ADL5630
Chironomus sp. TE11|[13506]|MHMOS107-07|BOLD:ADL5630
Chironomus sp. TE11|[13507]|CHURC921-09|BOLD:ADL5630
Chironomus sp. TE11|[13508]|CHURC555-08|BOLD:ADL5630
Chironomus sp. TE11|[13509]|CHURC853-09|BOLD:ADL5630
Chironomus sp. TE11|[13510]|CHURC483-08|BOLD:ADL5630
Chironomus sp. TE11|[13511]|CHURC793-09|BOLD:ADL5630
Chironomus sp. TE11|[13512]|CHURC724-09|BOLD:ADL5630
Chironomus sp. TE11|[13513]|JWDCI942-11|BOLD:ADL5630
Chironomus sp. TE11|[13514]|JWDCL105-11|BOLD:ADL5630
Chironomus sp. TE11|[13515]|JWDCI376-10|BOLD:ADL5630
Chironomus sp. TE11|[13516]|JWDCK166-11|BOLD:ADL5630
Chironomus sp. TE11|[13517]|JWDCJ1460-11|BOLD:ADL5630
Chironomus sp. TE11|[13518]|JWDCJ1458-11|BOLD:ADL5630
Chironomus sp. TE11|[13519]|JWDCF620-10|BOLD:ADL5630
Chironomus sp. TE11|[13520]|JWDCH977-10|BOLD:ADL5630
Chironomus sp. TE11|[13521]|JWDCF265-10|BOLD:ADL5630
Chironomus sp. TE11|[13522]|JWDCK104-11|BOLD:ADL5630
Chironomus sp. TE11|[13523]|JWDCH761-10|BOLD:ADL5630
Chironomus sp. TE11|[13524]|JWDCF063-10|BOLD:ADL5630
Chironomus sp. TE11|[13525]|JWDCA633-10|BOLD:ADL5630
Chironomus sp. TE11|[13526]|JWDCC447-10|BOLD:ADL5630
Chironomus sp. TE11|[13527]|JWDCF110-10|BOLD:ADL5630
Chironomus sp. TE11|[13528]|CHURC807-09|BOLD:ADL5630
Chironomus sp. TE11|[13529]|CHURC132-07|BOLD:ADL5630
Chironomus sp. TE11|[13530]|CHURC257-07|BOLD:ADL5630
Chironomus sp. TE11|[13531]|CHURC1032-09|BOLD:ADL5630
Chironomus sp. TE11|[13532]|CHURC026-06|BOLD:ADL5630
Chironomus dilutus|[13533]|CHURC443-07|BOLD:AAB4658

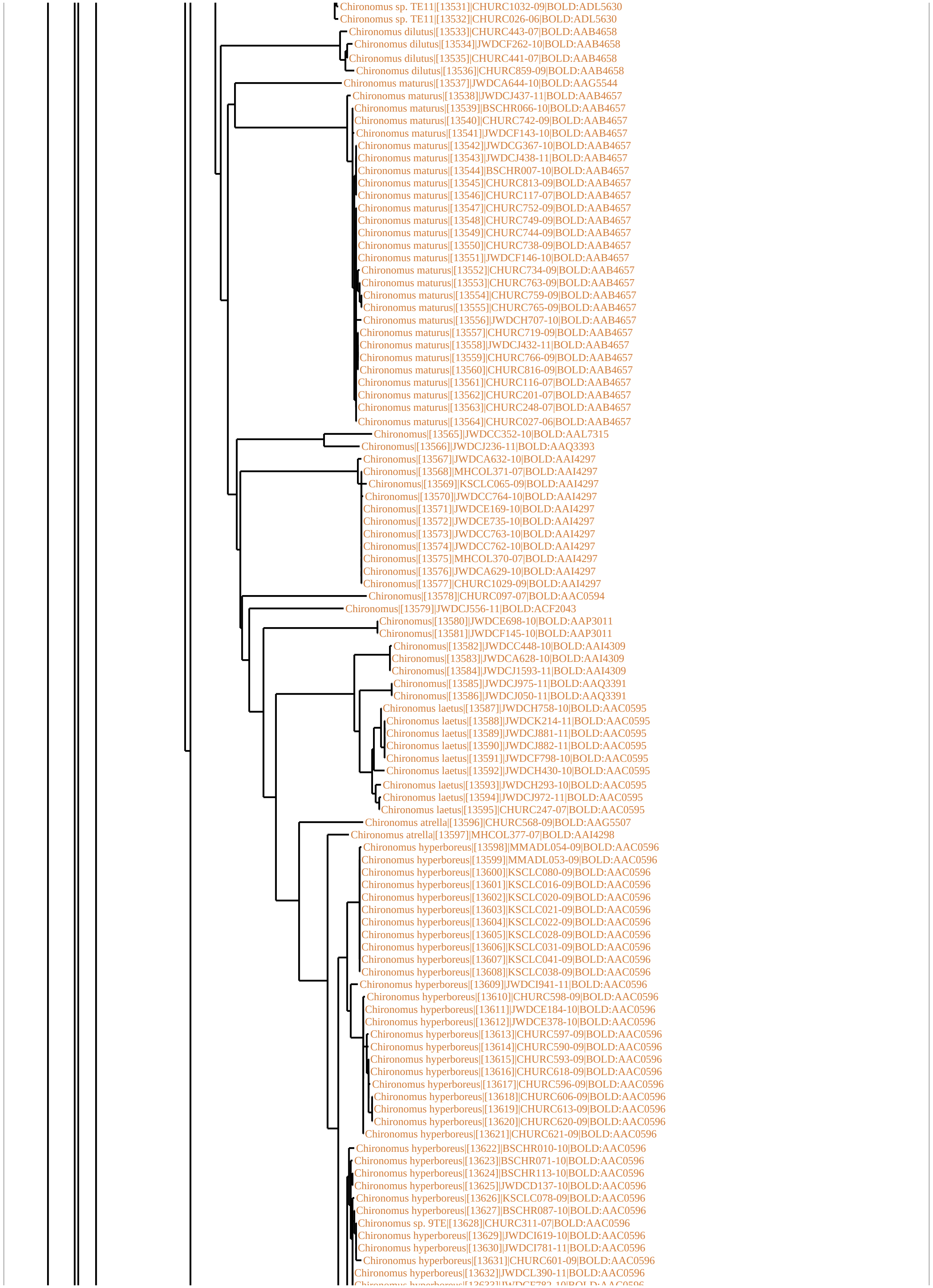
Chironomus sp. TE11|[13531]|CHURC1032-09|BOLD:ADL5630
Chironomus sp. TE11|[13532]|CHURC026-06|BOLD:ADL5630
Chironomus dilutus|[13533]|CHURC443-07|BOLD:AAB4658
Chironomus dilutus|[13534]|JWDCF262-10|BOLD:AAB4658
Chironomus dilutus|[13535]|CHURC441-07|BOLD:AAB4658
Chironomus dilutus|[13536]|CHURC859-09|BOLD:AAB4658
Chironomus maturus|[13537]|JWDCA644-10|BOLD:AAG5544
Chironomus maturus|[13538]|JWDCJ437-11|BOLD:AAB4657
Chironomus maturus|[13539]|BSCHR066-10|BOLD:AAB4657
Chironomus maturus|[13540]|CHURC742-09|BOLD:AAB4657
Chironomus maturus|[13541]|JWDCF143-10|BOLD:AAB4657
Chironomus maturus|[13542]|JWDCG367-10|BOLD:AAB4657
Chironomus maturus|[13543]|JWDCJ438-11|BOLD:AAB4657
Chironomus maturus|[13544]|BSCHR007-10|BOLD:AAB4657
Chironomus maturus|[13545]|CHURC813-09|BOLD:AAB4657
Chironomus maturus|[13546]|CHURC117-07|BOLD:AAB4657
Chironomus maturus|[13547]|CHURC752-09|BOLD:AAB4657
Chironomus maturus|[13548]|CHURC749-09|BOLD:AAB4657
Chironomus maturus|[13549]|CHURC744-09|BOLD:AAB4657
Chironomus maturus|[13550]|CHURC738-09|BOLD:AAB4657
Chironomus maturus|[13551]|JWDCF146-10|BOLD:AAB4657
Chironomus maturus|[13552]|CHURC734-09|BOLD:AAB4657
Chironomus maturus|[13553]|CHURC763-09|BOLD:AAB4657
Chironomus maturus|[13554]|CHURC759-09|BOLD:AAB4657
Chironomus maturus|[13555]|CHURC765-09|BOLD:AAB4657
Chironomus maturus|[13556]|JWDCH707-10|BOLD:AAB4657
Chironomus maturus|[13557]|CHURC719-09|BOLD:AAB4657
Chironomus maturus|[13558]|JWDCJ432-11|BOLD:AAB4657
Chironomus maturus|[13559]|CHURC766-09|BOLD:AAB4657
Chironomus maturus|[13560]|CHURC816-09|BOLD:AAB4657
Chironomus maturus|[13561]|CHURC116-07|BOLD:AAB4657
Chironomus maturus|[13562]|CHURC201-07|BOLD:AAB4657
Chironomus maturus|[13563]|CHURC248-07|BOLD:AAB4657
Chironomus maturus|[13564]|CHURC027-06|BOLD:AAB4657
Chironomus|[13565]|JWDCC352-10|BOLD:AAL7315
Chironomus|[13566]|JWDCJ236-11|BOLD:AAQ3393
Chironomus|[13567]|JWDCA632-10|BOLD:AAI4297
Chironomus|[13568]|MHCOL371-07|BOLD:AAI4297
Chironomus|[13569]|KSCLC065-09|BOLD:AAI4297
Chironomus|[13570]|JWDCC764-10|BOLD:AAI4297
Chironomus|[13571]|JWDCE169-10|BOLD:AAI4297
Chironomus|[13572]|JWDCE735-10|BOLD:AAI4297
Chironomus|[13573]|JWDCC763-10|BOLD:AAI4297
Chironomus|[13574]|JWDCC762-10|BOLD:AAI4297
Chironomus|[13575]|MHCOL370-07|BOLD:AAI4297
Chironomus|[13576]|JWDCA629-10|BOLD:AAI4297
Chironomus|[13577]|CHURC1029-09|BOLD:AAI4297
Chironomus|[13578]|CHURC097-07|BOLD:AAC0594
Chironomus|[13579]|JWDCJ556-11|BOLD:ACF2043
Chironomus|[13580]|JWDCE698-10|BOLD:AAP3011
Chironomus|[13581]|JWDCF145-10|BOLD:AAP3011
Chironomus|[13582]|JWDCC448-10|BOLD:AAI4309
Chironomus|[13583]|JWDCA628-10|BOLD:AAI4309
Chironomus|[13584]|JWDCJ1593-11|BOLD:AAI4309
Chironomus|[13585]|JWDCJ975-11|BOLD:AAQ3391
Chironomus|[13586]|JWDCJ050-11|BOLD:AAQ3391
Chironomus laetus|[13587]|JWDCH758-10|BOLD:AAC0595
Chironomus laetus|[13588]|JWDCK214-11|BOLD:AAC0595
Chironomus laetus|[13589]|JWDCJ881-11|BOLD:AAC0595
Chironomus laetus|[13590]|JWDCJ882-11|BOLD:AAC0595
Chironomus laetus|[13591]|JWDCF798-10|BOLD:AAC0595
Chironomus laetus|[13592]|JWDCH430-10|BOLD:AAC0595
Chironomus laetus|[13593]|JWDCH293-10|BOLD:AAC0595
Chironomus laetus|[13594]|JWDCJ972-11|BOLD:AAC0595
Chironomus laetus|[13595]|CHURC247-07|BOLD:AAC0595
Chironomus atrella|[13596]|CHURC568-09|BOLD:AAG5507
Chironomus atrella|[13597]|MHCOL377-07|BOLD:AAI4298
Chironomus hyperboreus|[13598]|MMADL054-09|BOLD:AAC0596
Chironomus hyperboreus|[13599]|MMADL053-09|BOLD:AAC0596
Chironomus hyperboreus|[13600]|KSCLC080-09|BOLD:AAC0596
Chironomus hyperboreus|[13601]|KSCLC016-09|BOLD:AAC0596
Chironomus hyperboreus|[13602]|KSCLC020-09|BOLD:AAC0596
Chironomus hyperboreus|[13603]|KSCLC021-09|BOLD:AAC0596
Chironomus hyperboreus|[13604]|KSCLC022-09|BOLD:AAC0596
Chironomus hyperboreus|[13605]|KSCLC028-09|BOLD:AAC0596
Chironomus hyperboreus|[13606]|KSCLC031-09|BOLD:AAC0596
Chironomus hyperboreus|[13607]|KSCLC041-09|BOLD:AAC0596
Chironomus hyperboreus|[13608]|KSCLC038-09|BOLD:AAC0596
Chironomus hyperboreus|[13609]|JWDCI941-11|BOLD:AAC0596
Chironomus hyperboreus|[13610]|CHURC598-09|BOLD:AAC0596
Chironomus hyperboreus|[13611]|JWDCE184-10|BOLD:AAC0596
Chironomus hyperboreus|[13612]|JWDCE378-10|BOLD:AAC0596
Chironomus hyperboreus|[13613]|CHURC597-09|BOLD:AAC0596
Chironomus hyperboreus|[13614]|CHURC590-09|BOLD:AAC0596
Chironomus hyperboreus|[13615]|CHURC593-09|BOLD:AAC0596
Chironomus hyperboreus|[13616]|CHURC618-09|BOLD:AAC0596
Chironomus hyperboreus|[13617]|CHURC596-09|BOLD:AAC0596
Chironomus hyperboreus|[13618]|CHURC606-09|BOLD:AAC0596
Chironomus hyperboreus|[13619]|CHURC613-09|BOLD:AAC0596
Chironomus hyperboreus|[13620]|CHURC620-09|BOLD:AAC0596
Chironomus hyperboreus|[13621]|CHURC621-09|BOLD:AAC0596
Chironomus hyperboreus|[13622]|BSCHR010-10|BOLD:AAC0596
Chironomus hyperboreus|[13623]|BSCHR071-10|BOLD:AAC0596
Chironomus hyperboreus|[13624]|BSCHR113-10|BOLD:AAC0596
Chironomus hyperboreus|[13625]|JWDCD137-10|BOLD:AAC0596
Chironomus hyperboreus|[13626]|KSCLC078-09|BOLD:AAC0596
Chironomus hyperboreus|[13627]|BSCHR087-10|BOLD:AAC0596
Chironomus sp. 9TE|[13628]|CHURC311-07|BOLD:AAC0596
Chironomus hyperboreus|[13629]|JWDCI619-10|BOLD:AAC0596
Chironomus hyperboreus|[13630]|JWDCI781-11|BOLD:AAC0596
Chironomus hyperboreus|[13631]|CHURC601-09|BOLD:AAC0596
Chironomus hyperboreus|[13632]|JWDCL390-11|BOLD:AAC0596

Chironomus hyperboreus|[13630]|JWDCI701-11|BOLD:AAC0596
Chironomus hyperboreus|[13631]|CHURC601-09|BOLD:AAC0596
Chironomus hyperboreus|[13632]|JWDCL390-11|BOLD:AAC0596
Chironomus hyperboreus|[13633]|JWDCF782-10|BOLD:AAC0596
Chironomus hyperboreus|[13634]|JWDCI623-10|BOLD:AAC0596
Chironomus hyperboreus|[13635]|KSCLC014-09|BOLD:AAC0596
Chironomus hyperboreus|[13636]|JWDCF261-10|BOLD:AAC0596
Chironomus hyperboreus|[13637]|KSCLC088-09|BOLD:AAC0596
Chironomus hyperboreus|[13638]|JWDCI862-11|BOLD:AAC0596
Chironomus hyperboreus|[13639]|JWDCA648-10|BOLD:AAC0596
Chironomus hyperboreus|[13640]|JWDCI579-10|BOLD:AAC0596
Chironomus sp. 9TE|[13641]|CHURC317-07|BOLD:AAC0596
Chironomus sp. 9TE|[13642]|CHURC315-07|BOLD:AAC0596
Chironomus hyperboreus|[13643]|JWDCI863-11|BOLD:AAC0596
Chironomus hyperboreus|[13644]|CHURC607-09|BOLD:AAC0596
Chironomus hyperboreus|[13645]|CHURC602-09|BOLD:AAC0596
Chironomus hyperboreus|[13646]|CHURC605-09|BOLD:AAC0596
Chironomus hyperboreus|[13647]|CHURC576-09|BOLD:AAC0596
Chironomus hyperboreus|[13648]|CHURC619-09|BOLD:AAC0596
Chironomus hyperboreus|[13649]|CHURC675-09|BOLD:AAC0596
Chironomus hyperboreus|[13650]|JWDCE186-10|BOLD:AAC0596
Chironomus hyperboreus|[13651]|JWDCE185-10|BOLD:AAC0596
Chironomus hyperboreus|[13652]|JWDCE137-10|BOLD:AAC0596
Chironomus hyperboreus|[13653]|JWDCE721-10|BOLD:AAC0596
Chironomus hyperboreus|[13654]|JWDCE380-10|BOLD:AAC0596
Chironomus hyperboreus|[13655]|JWDCF066-10|BOLD:AAC0596
Chironomus hyperboreus|[13656]|CHURC612-09|BOLD:AAC0596
Chironomus hyperboreus|[13657]|CHURC617-09|BOLD:AAC0596
Chironomus hyperboreus|[13658]|CHURC589-09|BOLD:AAC0596
Chironomus hyperboreus|[13659]|CHURC608-09|BOLD:AAC0596
Chironomus hyperboreus|[13660]|CHURC609-09|BOLD:AAC0596
Chironomus hyperboreus|[13661]|CHURC585-09|BOLD:AAC0596
Chironomus hyperboreus|[13662]|CHURC577-09|BOLD:AAC0596
Chironomus hyperboreus|[13663]|CHURC610-09|BOLD:AAC0596
Chironomus hyperboreus|[13664]|CHURC594-09|BOLD:AAC0596
Chironomus hyperboreus|[13665]|CHURC622-09|BOLD:AAC0596
Chironomus hyperboreus|[13666]|CHURC599-09|BOLD:AAC0596
Chironomus hyperboreus|[13667]|CHURC578-09|BOLD:AAC0596
Glyptotendipes lobiferus|[13668]|CHURC506-08|BOLD:AAF8348
Glyptotendipes lobiferus|[13669]|BSCHR179-10|BOLD:AAF8348
Glyptotendipes lobiferus|[13670]|BSCHR075-10|BOLD:AAF8348
Glyptotendipes lobiferus|[13671]|BSCHR126-10|BOLD:AAF8348
Glyptotendipes lobiferus|[13672]|BSCHR164-10|BOLD:AAF8348
Glyptotendipes lobiferus|[13673]|BSCHR183-10|BOLD:AAF8348
Glyptotendipes lobiferus|[13674]|BSCHR186-10|BOLD:AAF8348
Glyptotendipes lobiferus|[13675]|JWDCI621-10|BOLD:AAF8348
Glyptotendipes lobiferus|[13676]|JWDCJ824-11|BOLD:AAF8348
Glyptotendipes lobiferus|[13677]|JWDCJ829-11|BOLD:AAF8348
Glyptotendipes lobiferus|[13678]|BSCHR011-10|BOLD:AAF8348
Glyptotendipes lobiferus|[13679]|BSCHR038-10|BOLD:AAF8348
Glyptotendipes lobiferus|[13680]|CHURC871-09|BOLD:AAF8348
Cryptotendipes darbyi|[13681]|CHURC351-07|BOLD:AAC3085
Cryptotendipes darbyi|[13682]|CHURC353-07|BOLD:AAC3085
Cryptotendipes darbyi|[13683]|CHURC354-07|BOLD:AAC3085
Cryptotendipes darbyi|[13684]|CHURC355-07|BOLD:AAC3085
Cryptotendipes darbyi|[13685]|CHURC356-07|BOLD:AAC3085
Cryptotendipes darbyi|[13686]|CHURC437-07|BOLD:AAC3085
Cryptotendipes darbyi|[13687]|CHURC436-07|BOLD:AAC3084
Cryptotendipes darbyi|[13688]|CHURC350-07|BOLD:AAC3084
Cryptotendipes darbyi|[13689]|CHURC348-07|BOLD:AAC3084
Cryptotendipes darbyi|[13690]|CHURC344-07|BOLD:AAC3084
Cryptotendipes darbyi|[13691]|CHURC352-07|BOLD:AAC3084
Cryptotendipes darbyi|[13692]|CHURC357-07|BOLD:AAC3084
Cryptotendipes darbyi|[13693]|CHURC528-08|BOLD:AAC3084
Chironomus sp. TE3|[13694]|MHMOS068-07|BOLD:AAD4858
Chironomus sp. 3TE|[13695]|MHMOS064-07|BOLD:AAD4858
Chironomus sp. TE3|[13696]|MHMOS060-07|BOLD:AAD4858
Chironomus sp. 3TE|[13697]|CHURC493-08|BOLD:AAD4858
Chironomus sp. 3TE|[13698]|MHMOS065-07|BOLD:AAD4858
Chironomus sp. 3TE|[13699]|MHMOS006-07|BOLD:AAD4858
Chironomus sp. 3TE|[13700]|CHURC533-08|BOLD:AAD4858
Parachironomus frequens|[13701]|BSCHR053-10|BOLD:AAG5461
Parachironomus frequens|[13702]|CHURC936-09|BOLD:AAI4127
Dicrotendipes sp. TE01|[13703]|CHURC584-09|BOLD:AAL1066
Chironomus|[13704]|MHCOL374-07|BOLD:AAC0593
Chironomus|[13705]|MHCOL372-07|BOLD:AAC0593
Chironomus|[13706]|MHCOL378-07|BOLD:AAC0593
Chironomus|[13707]|MHCOL380-07|BOLD:AAC0593
Chironomus|[13708]|MHMOS066-07|BOLD:AAC0593
Glyptotendipes paripes|[13709]|JWDCE846-10|BOLD:AHJ7930
Glyptotendipes paripes|[13710]|JWDCI710-11|BOLD:AHJ7930
Glyptotendipes paripes|[13711]|BSCHR073-10|BOLD:AAC0597
Glyptotendipes paripes|[13712]|JWDCC907-10|BOLD:AAC0597
Glyptotendipes paripes|[13713]|JWDCF588-10|BOLD:AAC0597
Glyptotendipes paripes|[13714]|JWDCF587-10|BOLD:AAC0597
Glyptotendipes paripes|[13715]|JWDCF111-10|BOLD:AAC0597
Glyptotendipes paripes|[13716]|MHCOL379-07|BOLD:AAC0597
Glyptotendipes paripes|[13717]|JWDCA630-10|BOLD:AAC0597
Glyptotendipes paripes|[13718]|CHURC614-09|BOLD:AAC0597
Glyptotendipes paripes|[13719]|BSCHR014-10|BOLD:AAC0597
Glyptotendipes paripes|[13720]|CHURC591-09|BOLD:AAC0597
Glyptotendipes paripes|[13721]|CHURC611-09|BOLD:AAC0597
Glyptotendipes paripes|[13722]|JWDCF586-10|BOLD:AAC0597
Glyptotendipes paripes|[13723]|JWDCJ371-11|BOLD:AAC0597
Glyptotendipes paripes|[13724]|SWSWP085-09|BOLD:AAC0597
Glyptotendipes paripes|[13725]|JWDCC909-10|BOLD:AAC0597
Glyptotendipes paripes|[13726]|JWDCC908-10|BOLD:AAC0597
Glyptotendipes paripes|[13727]|JWDCC905-10|BOLD:AAC0597
Glyptotendipes paripes|[13728]|JWDCC843-10|BOLD:AAC0597
Glyptotendipes paripes|[13729]|BSCHR155-10|BOLD:AAC0597
Glyptotendipes paripes|[13730]|BSCHR089-10|BOLD:AAC0597
Glyptotendipes paripes|[13731]|BSCHR063-10|BOLD:AAC0597
Glyptotendipes paripes|[13732]|BSCHR058-10|BOLD:AAC0597

Glyptotendipes paripes|[13730]|BSCHR089-10|BOLD:AAC0597
Glyptotendipes paripes|[13731]|BSCHR063-10|BOLD:AAC0597
Glyptotendipes paripes|[13732]|BSCHR058-10|BOLD:AAC0597
Glyptotendipes paripes|[13733]|BSCHR046-10|BOLD:AAC0597
Glyptotendipes paripes|[13734]|BSCHR023-10|BOLD:AAC0597
Glyptotendipes paripes|[13735]|BSCHR019-10|BOLD:AAC0597
Glyptotendipes paripes|[13736]|JWDCB147-10|BOLD:AAC0597
Glyptotendipes paripes|[13737]|BSCHR061-10|BOLD:AAC0597
Glyptotendipes paripes|[13738]|JWDCF671-10|BOLD:AAC0597
Glyptotendipes paripes|[13739]|JWDCC065-10|BOLD:AAC0597
Glyptotendipes paripes|[13740]|JWDCC910-10|BOLD:AAC0597
Glyptotendipes paripes|[13741]|CHURC376-07|BOLD:AAC0597
Glyptotendipes paripes|[13742]|JWDCC066-10|BOLD:AAC0597
Glyptotendipes paripes|[13743]|CHURC600-09|BOLD:AAC0597
Glyptotendipes paripes|[13744]|BSCHR031-10|BOLD:AAC0597
Glyptotendipes paripes|[13745]|JWDCC063-10|BOLD:AAC0597
Glyptotendipes paripes|[13746]|CHURC429-07|BOLD:AAC0597
Glyptotendipes paripes|[13747]|CHURC581-09|BOLD:AAC0597
Cricotopus sp. 18ES|[13748]|CHURC692-09|BOLD:AAG0996
Cricotopus tricinctus|[13749]|CHURC511-08|BOLD:AAA5314
Cricotopus tricinctus|[13750]|JWDCH825-10|BOLD:AAA5305
Cricotopus tricinctus|[13751]|CHURC102-07|BOLD:AAA5305
Cricotopus sp. 4ES|[13752]|CHURC229-07|BOLD:AAA5303
Cricotopus sp. 4ES|[13753]|CHURC079-06|BOLD:AAA5303
Cricotopus sp. 4ES|[13754]|CHURC083-06|BOLD:AAA5303
Cricotopus sp. 4ES|[13755]|CHURC081-06|BOLD:AAA5303
Cricotopus sp. 4ES|[13756]|CHURC078-06|BOLD:AAA5303
Cricotopus sp. 4ES|[13757]|CHURC080-06|BOLD:AAA5303
Cricotopus sp. 4ES|[13758]|CHURC082-06|BOLD:AAA5303
Cricotopus sp. 6ES|[13759]|CHURC054-06|BOLD:AAA5301
Cricotopus sp. 6ES|[13760]|CHURC136-07|BOLD:AAA5301
Cricotopus sp. 6ES|[13761]|CHURC135-07|BOLD:AAA5301
Cricotopus sp. 6ES|[13762]|CHURC133-07|BOLD:AAA5301
Cricotopus sp. 6ES|[13763]|KSCLC061-09|BOLD:AAA5301
Cricotopus sp. 6ES|[13764]|CHURC134-07|BOLD:AAA5301
Cricotopus sp. 6ES|[13765]|CHURC094-06|BOLD:AAA5301
Cricotopus sp. 6ES|[13766]|CHURC292-07|BOLD:AAA5301
Cricotopus sp. 7ES|[13767]|KSCLC060-09|BOLD:AAA5308
Cricotopus sp. 7ES|[13768]|CHURC654-09|BOLD:AAA5308
Cricotopus sp. 7ES|[13769]|JWDCH539-10|BOLD:AAA5308
Cricotopus sp. 7ES|[13770]|CHURC291-07|BOLD:AAA5308
Cricotopus sp. 7ES|[13771]|CHURC639-09|BOLD:AAA5308
Cricotopus sp. 7ES|[13772]|CHURC640-09|BOLD:AAA5308
Cricotopus sp. 7ES|[13773]|CHURC525-08|BOLD:AAA5308
Cricotopus sp. 7ES|[13774]|JWDCH109-10|BOLD:AAA5308
Cricotopus sp. 7ES|[13775]|JWDCH148-10|BOLD:AAA5308
Cricotopus sp. 7ES|[13776]|KSCLC062-09|BOLD:AAA5308
Cricotopus sp. 7ES|[13777]|CHURC659-09|BOLD:AAA5308
Cricotopus|[13778]|CHURC731-09|BOLD:AAA5299
Cricotopus|[13779]|CHURC723-09|BOLD:AAA5299
Cricotopus sylvestris|[13780]|CHURC205-07|BOLD:AAA5299
Cricotopus sylvestris|[13781]|CHURC861-09|BOLD:AAA5299
Cricotopus sylvestris|[13782]|CHURC152-07|BOLD:AAA5299
Cricotopus|[13783]|JWDCJ1317-11|BOLD:AAA5299
Cricotopus|[13784]|JWDCJ1315-11|BOLD:AAA5299
Cricotopus sylvestris|[13785]|CHURC232-07|BOLD:AAA5299
Cricotopus|[13786]|JWDCH759-10|BOLD:AAA5299
Cricotopus sylvestris|[13787]|CHURC494-08|BOLD:AAA5299
Cricotopus sylvestris|[13788]|CHURC523-08|BOLD:AAA5299
Cricotopus sylvestris|[13789]|CHURC013-06|BOLD:AAA5299
Cricotopus sylvestris|[13790]|CHURC488-08|BOLD:AAA5299
Orthocladius knuthi|[13791]|CHURC895-09|BOLD:AAI3485
Procladius lugens|[13792]|BSCHR159-10|BOLD:AAN5387
Lasiodiamesa|[13793]|JWDCJ594-11|BOLD:AAQ0621
Lasiodiamesa|[13794]|JWDCF799-10|BOLD:AAQ0621
Tanypodinae|[13795]|BSCHR128-10|BOLD:AAG2646
Procladius|[13796]|BSCHR185-10|BOLD:AAN5390
Procladius|[13797]|BSCHR106-10|BOLD:AAL7334
Procladius sp. ES05|[13798]|CHURC469-07|BOLD:AAX7792
Procladius|[13799]|JWDCF108-10|BOLD:AAP9002
Procladius|[13800]|JWDCF731-10|BOLD:AAP9002
Procladius sp. ES06|[13801]|JWDCE656-10|BOLD:AAF6804
Procladius sp. ES06|[13802]|JWDCF067-10|BOLD:AAF6804
Procladius sp. ES06|[13803]|JWDCF069-10|BOLD:AAF6804
Procladius sp. ES06|[13804]|CHURC543-08|BOLD:AAF6804
Procladius appropinquatus|[13805]|JWDCH518-10|BOLD:AAE0538
Procladius appropinquatus|[13806]|CHURC487-08|BOLD:AAE0538
Procladius appropinquatus|[13807]|JWDCL033-11|BOLD:AAE0538
Procladius appropinquatus|[13808]|JWDCI190-10|BOLD:AAE0538
Procladius appropinquatus|[13809]|JWDCA635-10|BOLD:AAE0538
Procladius appropinquatus|[13810]|MMADL092-09|BOLD:AAE0538
Procladius appropinquatus|[13811]|KSCLC074-09|BOLD:AAE0538
Procladius appropinquatus|[13812]|BSCHR182-10|BOLD:AAE0538
Procladius appropinquatus|[13813]|BSCHR180-10|BOLD:AAE0538
Procladius appropinquatus|[13814]|JWDCA637-10|BOLD:AAE0538
Procladius appropinquatus|[13815]|CHURC507-08|BOLD:AAE0538
Procladius dentus|[13816]|JWDCI578-10|BOLD:AAD7251
Procladius dentus|[13817]|JWDCA638-10|BOLD:AAD7251
Procladius dentus|[13818]|KSCLC039-09|BOLD:AAD7251
Procladius dentus|[13819]|KSCLC086-09|BOLD:AAD7251
Procladius dentus|[13820]|MMADL078-09|BOLD:AAD7251
Procladius dentus|[13821]|KSCLC053-09|BOLD:AAD7251
Procladius dentus|[13822]|KSCLC054-09|BOLD:AAD7251
Procladius dentus|[13823]|JWDCE616-10|BOLD:AAD7251
Procladius dentus|[13824]|JWDCE648-10|BOLD:AAD7251
Procladius dentus|[13825]|BSCHR086-10|BOLD:AAD7251
Procladius dentus|[13826]|JWDCE658-10|BOLD:AAD7251
Procladius dentus|[13827]|BSCHR095-10|BOLD:AAD7251
Procladius dentus|[13828]|BSCHR107-10|BOLD:AAD7251
Procladius dentus|[13829]|BSCHR054-10|BOLD:AAD7251
Procladius dentus|[13830]|BSCHR030-10|BOLD:AAD7251
Procladius dentus|[13831]|JWDCC296-10|BOLD:AAD7251
Procladius dentus|[13832]|JWDCD058-10|BOLD:AAD7251

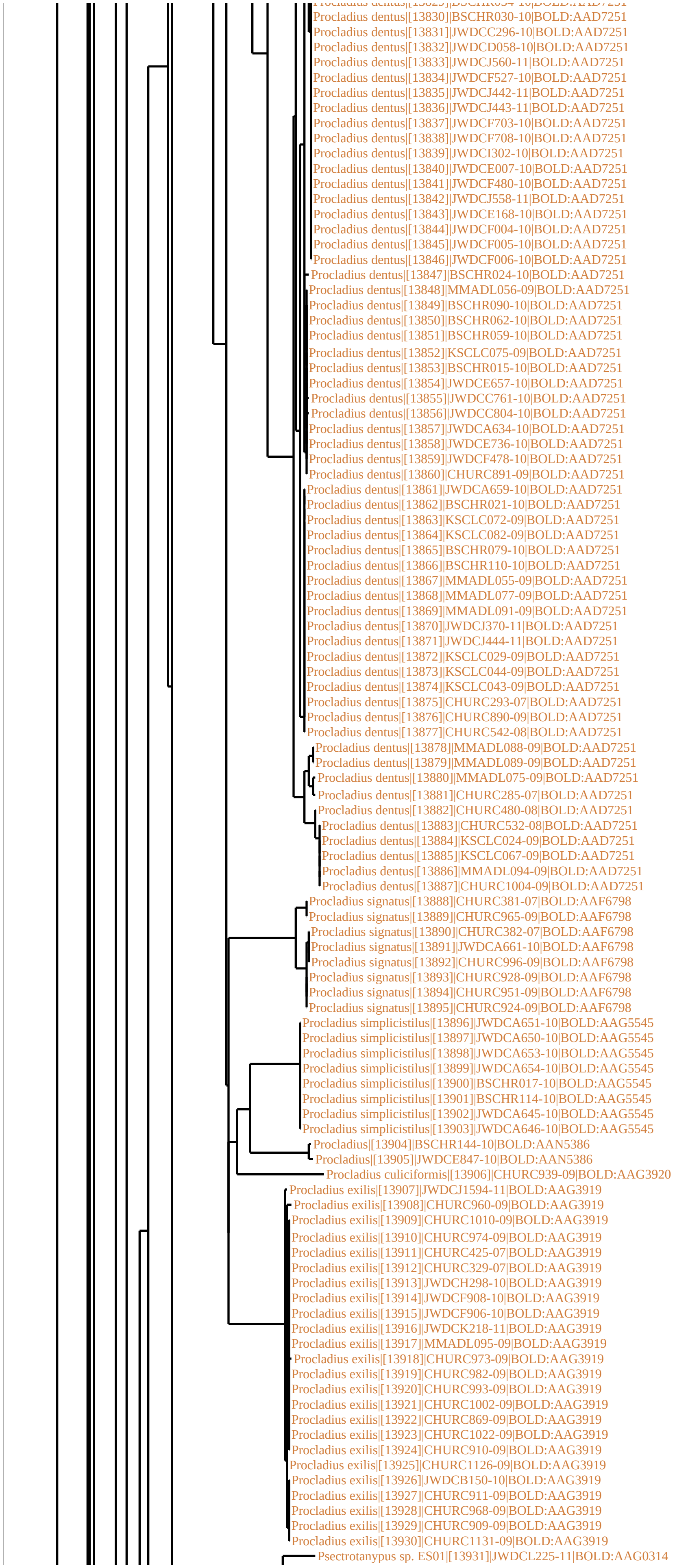
Procladius dentus|[13829]|BSCHR034-10|BOLD:AAD7251
Procladius dentus|[13830]|BSCHR030-10|BOLD:AAD7251
Procladius dentus|[13831]|JWDCC296-10|BOLD:AAD7251
Procladius dentus|[13832]|JWDCD058-10|BOLD:AAD7251
Procladius dentus|[13833]|JWDCJ560-11|BOLD:AAD7251
Procladius dentus|[13834]|JWDCF527-10|BOLD:AAD7251
Procladius dentus|[13835]|JWDCJ442-11|BOLD:AAD7251
Procladius dentus|[13836]|JWDCJ443-11|BOLD:AAD7251
Procladius dentus|[13837]|JWDCF703-10|BOLD:AAD7251
Procladius dentus|[13838]|JWDCF708-10|BOLD:AAD7251
Procladius dentus|[13839]|JWDCI302-10|BOLD:AAD7251
Procladius dentus|[13840]|JWDCE007-10|BOLD:AAD7251
Procladius dentus|[13841]|JWDCF480-10|BOLD:AAD7251
Procladius dentus|[13842]|JWDCJ558-11|BOLD:AAD7251
Procladius dentus|[13843]|JWDCE168-10|BOLD:AAD7251
Procladius dentus|[13844]|JWDCF004-10|BOLD:AAD7251
Procladius dentus|[13845]|JWDCF005-10|BOLD:AAD7251
Procladius dentus|[13846]|JWDCF006-10|BOLD:AAD7251
Procladius dentus|[13847]|BSCHR024-10|BOLD:AAD7251
Procladius dentus|[13848]|MMADL056-09|BOLD:AAD7251
Procladius dentus|[13849]|BSCHR090-10|BOLD:AAD7251
Procladius dentus|[13850]|BSCHR062-10|BOLD:AAD7251
Procladius dentus|[13851]|BSCHR059-10|BOLD:AAD7251
Procladius dentus|[13852]|KSCLC075-09|BOLD:AAD7251
Procladius dentus|[13853]|BSCHR015-10|BOLD:AAD7251
Procladius dentus|[13854]|JWDCE657-10|BOLD:AAD7251
Procladius dentus|[13855]|JWDCC761-10|BOLD:AAD7251
Procladius dentus|[13856]|JWDCC804-10|BOLD:AAD7251
Procladius dentus|[13857]|JWDCA634-10|BOLD:AAD7251
Procladius dentus|[13858]|JWDCE736-10|BOLD:AAD7251
Procladius dentus|[13859]|JWDCF478-10|BOLD:AAD7251
Procladius dentus|[13860]|CHURC891-09|BOLD:AAD7251
Procladius dentus|[13861]|JWDCA659-10|BOLD:AAD7251
Procladius dentus|[13862]|BSCHR021-10|BOLD:AAD7251
Procladius dentus|[13863]|KSCLC072-09|BOLD:AAD7251
Procladius dentus|[13864]|KSCLC082-09|BOLD:AAD7251
Procladius dentus|[13865]|BSCHR079-10|BOLD:AAD7251
Procladius dentus|[13866]|BSCHR110-10|BOLD:AAD7251
Procladius dentus|[13867]|MMADL055-09|BOLD:AAD7251
Procladius dentus|[13868]|MMADL077-09|BOLD:AAD7251
Procladius dentus|[13869]|MMADL091-09|BOLD:AAD7251
Procladius dentus|[13870]|JWDCJ370-11|BOLD:AAD7251
Procladius dentus|[13871]|JWDCJ444-11|BOLD:AAD7251
Procladius dentus|[13872]|KSCLC029-09|BOLD:AAD7251
Procladius dentus|[13873]|KSCLC044-09|BOLD:AAD7251
Procladius dentus|[13874]|KSCLC043-09|BOLD:AAD7251
Procladius dentus|[13875]|CHURC293-07|BOLD:AAD7251
Procladius dentus|[13876]|CHURC890-09|BOLD:AAD7251
Procladius dentus|[13877]|CHURC542-08|BOLD:AAD7251
Procladius dentus|[13878]|MMADL088-09|BOLD:AAD7251
Procladius dentus|[13879]|MMADL089-09|BOLD:AAD7251
Procladius dentus|[13880]|MMADL075-09|BOLD:AAD7251
Procladius dentus|[13881]|CHURC285-07|BOLD:AAD7251
Procladius dentus|[13882]|CHURC480-08|BOLD:AAD7251
Procladius dentus|[13883]|CHURC532-08|BOLD:AAD7251
Procladius dentus|[13884]|KSCLC024-09|BOLD:AAD7251
Procladius dentus|[13885]|KSCLC067-09|BOLD:AAD7251
Procladius dentus|[13886]|MMADL094-09|BOLD:AAD7251
Procladius dentus|[13887]|CHURC1004-09|BOLD:AAD7251
Procladius signatus|[13888]|CHURC381-07|BOLD:AAF6798
Procladius signatus|[13889]|CHURC965-09|BOLD:AAF6798
Procladius signatus|[13890]|CHURC382-07|BOLD:AAF6798
Procladius signatus|[13891]|JWDCA661-10|BOLD:AAF6798
Procladius signatus|[13892]|CHURC996-09|BOLD:AAF6798
Procladius signatus|[13893]|CHURC928-09|BOLD:AAF6798
Procladius signatus|[13894]|CHURC951-09|BOLD:AAF6798
Procladius signatus|[13895]|CHURC924-09|BOLD:AAF6798
Procladius simplicistilus|[13896]|JWDCA651-10|BOLD:AAG5545
Procladius simplicistilus|[13897]|JWDCA650-10|BOLD:AAG5545
Procladius simplicistilus|[13898]|JWDCA653-10|BOLD:AAG5545
Procladius simplicistilus|[13899]|JWDCA654-10|BOLD:AAG5545
Procladius simplicistilus|[13900]|BSCHR017-10|BOLD:AAG5545
Procladius simplicistilus|[13901]|BSCHR114-10|BOLD:AAG5545
Procladius simplicistilus|[13902]|JWDCA645-10|BOLD:AAG5545
Procladius simplicistilus|[13903]|JWDCA646-10|BOLD:AAG5545
Procladius|[13904]|BSCHR144-10|BOLD:AAN5386
Procladius|[13905]|JWDCE847-10|BOLD:AAN5386
Procladius culiciformis|[13906]|CHURC939-09|BOLD:AAG3920
Procladius exilis|[13907]|JWDCJ1594-11|BOLD:AAG3919
Procladius exilis|[13908]|CHURC960-09|BOLD:AAG3919
Procladius exilis|[13909]|CHURC1010-09|BOLD:AAG3919
Procladius exilis|[13910]|CHURC974-09|BOLD:AAG3919
Procladius exilis|[13911]|CHURC425-07|BOLD:AAG3919
Procladius exilis|[13912]|CHURC329-07|BOLD:AAG3919
Procladius exilis|[13913]|JWDCH298-10|BOLD:AAG3919
Procladius exilis|[13914]|JWDCF908-10|BOLD:AAG3919
Procladius exilis|[13915]|JWDCF906-10|BOLD:AAG3919
Procladius exilis|[13916]|JWDCK218-11|BOLD:AAG3919
Procladius exilis|[13917]|MMADL095-09|BOLD:AAG3919
Procladius exilis|[13918]|CHURC973-09|BOLD:AAG3919
Procladius exilis|[13919]|CHURC982-09|BOLD:AAG3919
Procladius exilis|[13920]|CHURC993-09|BOLD:AAG3919
Procladius exilis|[13921]|CHURC1002-09|BOLD:AAG3919
Procladius exilis|[13922]|CHURC869-09|BOLD:AAG3919
Procladius exilis|[13923]|CHURC1022-09|BOLD:AAG3919
Procladius exilis|[13924]|CHURC910-09|BOLD:AAG3919
Procladius exilis|[13925]|CHURC1126-09|BOLD:AAG3919
Procladius exilis|[13926]|JWDCB150-10|BOLD:AAG3919
Procladius exilis|[13927]|CHURC911-09|BOLD:AAG3919
Procladius exilis|[13928]|CHURC968-09|BOLD:AAG3919
Procladius exilis|[13929]|CHURC909-09|BOLD:AAG3919
Procladius exilis|[13930]|CHURC1131-09|BOLD:AAG3919
Psectrotanypus sp. ES01|[13931]|JWDCL225-11|BOLD:AAG0314

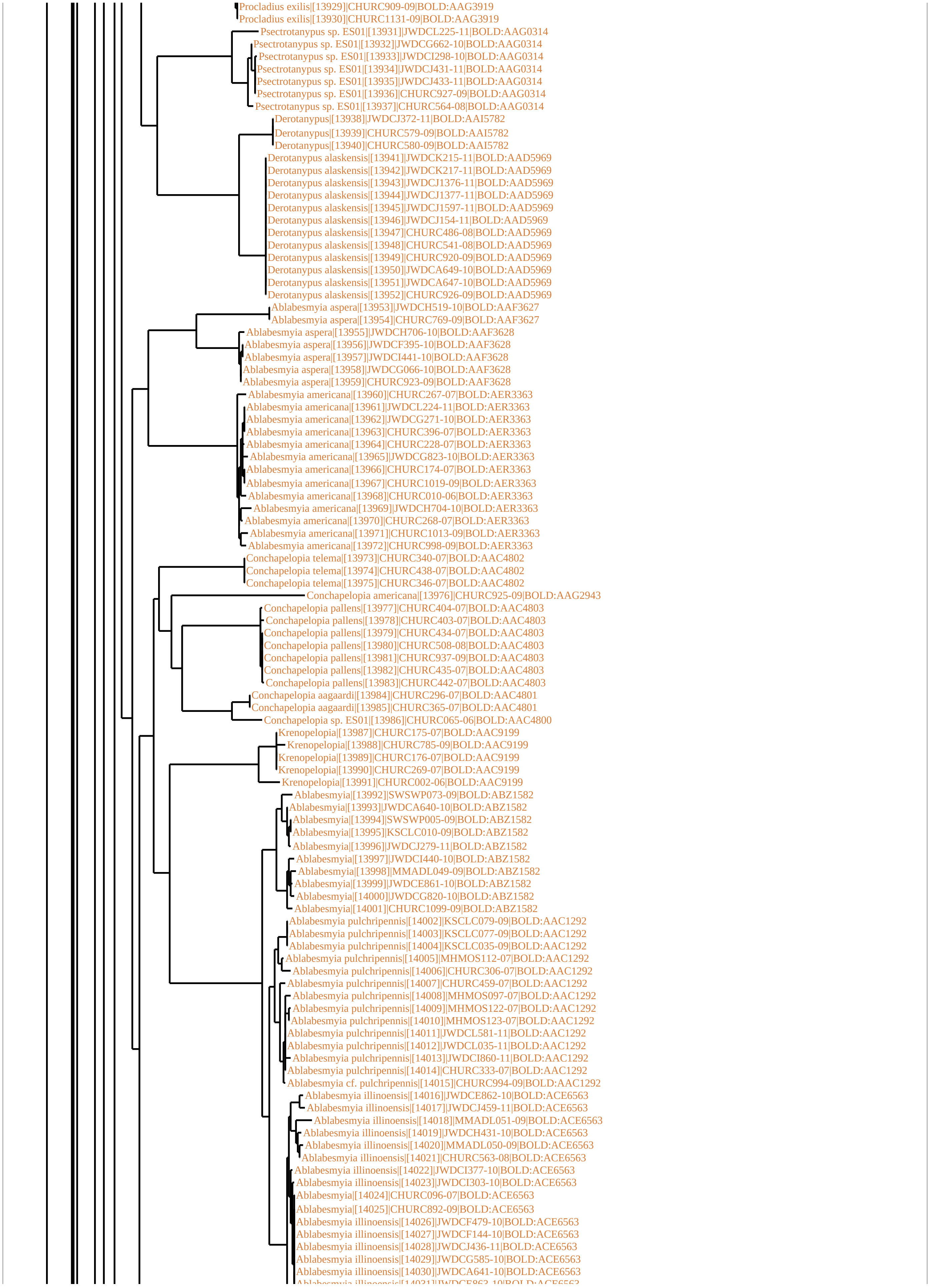
Procladius exilis|[13929]|CHURC909-09|BOLD:AAG3919
Procladius exilis|[13930]|CHURC1131-09|BOLD:AAG3919
Psectrotanypus sp. ES01|[13931]|JWDCL225-11|BOLD:AAG0314
Psectrotanypus sp. ES01|[13932]|JWDCG662-10|BOLD:AAG0314
Psectrotanypus sp. ES01|[13933]|JWDCI298-10|BOLD:AAG0314
Psectrotanypus sp. ES01|[13934]|JWDCJ431-11|BOLD:AAG0314
Psectrotanypus sp. ES01|[13935]|JWDCJ433-11|BOLD:AAG0314
Psectrotanypus sp. ES01|[13936]|CHURC927-09|BOLD:AAG0314
Psectrotanypus sp. ES01|[13937]|CHURC564-08|BOLD:AAG0314
Derotanypus|[13938]|JWDCJ372-11|BOLD:AAI5782
Derotanypus|[13939]|CHURC579-09|BOLD:AAI5782
Derotanypus|[13940]|CHURC580-09|BOLD:AAI5782
Derotanypus alaskensis|[13941]|JWDCK215-11|BOLD:AAD5969
Derotanypus alaskensis|[13942]|JWDCK217-11|BOLD:AAD5969
Derotanypus alaskensis|[13943]|JWDCJ1376-11|BOLD:AAD5969
Derotanypus alaskensis|[13944]|JWDCJ1377-11|BOLD:AAD5969
Derotanypus alaskensis|[13945]|JWDCJ1597-11|BOLD:AAD5969
Derotanypus alaskensis|[13946]|JWDCJ154-11|BOLD:AAD5969
Derotanypus alaskensis|[13947]|CHURC486-08|BOLD:AAD5969
Derotanypus alaskensis|[13948]|CHURC541-08|BOLD:AAD5969
Derotanypus alaskensis|[13949]|CHURC920-09|BOLD:AAD5969
Derotanypus alaskensis|[13950]|JWDCA649-10|BOLD:AAD5969
Derotanypus alaskensis|[13951]|JWDCA647-10|BOLD:AAD5969
Derotanypus alaskensis|[13952]|CHURC926-09|BOLD:AAD5969
Ablabesmyia aspera|[13953]|JWDCH519-10|BOLD:AAF3627
Ablabesmyia aspera|[13954]|CHURC769-09|BOLD:AAF3627
Ablabesmyia aspera|[13955]|JWDCH706-10|BOLD:AAF3628
Ablabesmyia aspera|[13956]|JWDCF395-10|BOLD:AAF3628
Ablabesmyia aspera|[13957]|JWDCI441-10|BOLD:AAF3628
Ablabesmyia aspera|[13958]|JWDCG066-10|BOLD:AAF3628
Ablabesmyia aspera|[13959]|CHURC923-09|BOLD:AAF3628
Ablabesmyia americana|[13960]|CHURC267-07|BOLD:AER3363
Ablabesmyia americana|[13961]|JWDCL224-11|BOLD:AER3363
Ablabesmyia americana|[13962]|JWDCG271-10|BOLD:AER3363
Ablabesmyia americana|[13963]|CHURC396-07|BOLD:AER3363
Ablabesmyia americana|[13964]|CHURC228-07|BOLD:AER3363
Ablabesmyia americana|[13965]|JWDCG823-10|BOLD:AER3363
Ablabesmyia americana|[13966]|CHURC174-07|BOLD:AER3363
Ablabesmyia americana|[13967]|CHURC1019-09|BOLD:AER3363
Ablabesmyia americana|[13968]|CHURC010-06|BOLD:AER3363
Ablabesmyia americana|[13969]|JWDCH704-10|BOLD:AER3363
Ablabesmyia americana|[13970]|CHURC268-07|BOLD:AER3363
Ablabesmyia americana|[13971]|CHURC1013-09|BOLD:AER3363
Ablabesmyia americana|[13972]|CHURC998-09|BOLD:AER3363
Conchapelopia telema|[13973]|CHURC340-07|BOLD:AAC4802
Conchapelopia telema|[13974]|CHURC438-07|BOLD:AAC4802
Conchapelopia telema|[13975]|CHURC346-07|BOLD:AAC4802
Conchapelopia americana|[13976]|CHURC925-09|BOLD:AAG2943
Conchapelopia pallens|[13977]|CHURC404-07|BOLD:AAC4803
Conchapelopia pallens|[13978]|CHURC403-07|BOLD:AAC4803
Conchapelopia pallens|[13979]|CHURC434-07|BOLD:AAC4803
Conchapelopia pallens|[13980]|CHURC508-08|BOLD:AAC4803
Conchapelopia pallens|[13981]|CHURC937-09|BOLD:AAC4803
Conchapelopia pallens|[13982]|CHURC435-07|BOLD:AAC4803
Conchapelopia pallens|[13983]|CHURC442-07|BOLD:AAC4803
Conchapelopia aagaardi|[13984]|CHURC296-07|BOLD:AAC4801
Conchapelopia aagaardi|[13985]|CHURC365-07|BOLD:AAC4801
Conchapelopia sp. ES01|[13986]|CHURC065-06|BOLD:AAC4800
Krenopelopia|[13987]|CHURC175-07|BOLD:AAC9199
Krenopelopia|[13988]|CHURC785-09|BOLD:AAC9199
Krenopelopia|[13989]|CHURC176-07|BOLD:AAC9199
Krenopelopia|[13990]|CHURC269-07|BOLD:AAC9199
Krenopelopia|[13991]|CHURC002-06|BOLD:AAC9199
Ablabesmyia|[13992]|SWSWP073-09|BOLD:ABZ1582
Ablabesmyia|[13993]|JWDCA640-10|BOLD:ABZ1582
Ablabesmyia|[13994]|SWSWP005-09|BOLD:ABZ1582
Ablabesmyia|[13995]|KSCLC010-09|BOLD:ABZ1582
Ablabesmyia|[13996]|JWDCJ279-11|BOLD:ABZ1582
Ablabesmyia|[13997]|JWDCI440-10|BOLD:ABZ1582
Ablabesmyia|[13998]|MMADL049-09|BOLD:ABZ1582
Ablabesmyia|[13999]|JWDCE861-10|BOLD:ABZ1582
Ablabesmyia|[14000]|JWDCG820-10|BOLD:ABZ1582
Ablabesmyia|[14001]|CHURC1099-09|BOLD:ABZ1582
Ablabesmyia pulchripennis|[14002]|KSCLC079-09|BOLD:AAC1292
Ablabesmyia pulchripennis|[14003]|KSCLC077-09|BOLD:AAC1292
Ablabesmyia pulchripennis|[14004]|KSCLC035-09|BOLD:AAC1292
Ablabesmyia pulchripennis|[14005]|MHMOS112-07|BOLD:AAC1292
Ablabesmyia pulchripennis|[14006]|CHURC306-07|BOLD:AAC1292
Ablabesmyia pulchripennis|[14007]|CHURC459-07|BOLD:AAC1292
Ablabesmyia pulchripennis|[14008]|MHMOS097-07|BOLD:AAC1292
Ablabesmyia pulchripennis|[14009]|MHMOS122-07|BOLD:AAC1292
Ablabesmyia pulchripennis|[14010]|MHMOS123-07|BOLD:AAC1292
Ablabesmyia pulchripennis|[14011]|JWDCL581-11|BOLD:AAC1292
Ablabesmyia pulchripennis|[14012]|JWDCL035-11|BOLD:AAC1292
Ablabesmyia pulchripennis|[14013]|JWDCI860-11|BOLD:AAC1292
Ablabesmyia pulchripennis|[14014]|CHURC333-07|BOLD:AAC1292
Ablabesmyia cf. pulchripennis|[14015]|CHURC994-09|BOLD:AAC1292
Ablabesmyia illinoensis|[14016]|JWDCE862-10|BOLD:ACE6563
Ablabesmyia illinoensis|[14017]|JWDCJ459-11|BOLD:ACE6563
Ablabesmyia illinoensis|[14018]|MMADL051-09|BOLD:ACE6563
Ablabesmyia illinoensis|[14019]|JWDCH431-10|BOLD:ACE6563
Ablabesmyia illinoensis|[14020]|MMADL050-09|BOLD:ACE6563
Ablabesmyia illinoensis|[14021]|CHURC563-08|BOLD:ACE6563
Ablabesmyia illinoensis|[14022]|JWDCI377-10|BOLD:ACE6563
Ablabesmyia illinoensis|[14023]|JWDCI303-10|BOLD:ACE6563
Ablabesmyia|[14024]|CHURC096-07|BOLD:ACE6563
Ablabesmyia|[14025]|CHURC892-09|BOLD:ACE6563
Ablabesmyia illinoensis|[14026]|JWDCF479-10|BOLD:ACE6563
Ablabesmyia illinoensis|[14027]|JWDCF144-10|BOLD:ACE6563
Ablabesmyia illinoensis|[14028]|JWDCJ436-11|BOLD:ACE6563
Ablabesmyia illinoensis|[14029]|JWDCG585-10|BOLD:ACE6563
Ablabesmyia illinoensis|[14030]|JWDCA641-10|BOLD:ACE6563

Ablabesmyia illinoensis|[14028]|[illegible]
Ablabesmyia illinoensis|[14029]|JWDCG585-10|BOLD:ACE6563
Ablabesmyia illinoensis|[14030]|JWDCA641-10|BOLD:ACE6563
Ablabesmyia illinoensis|[14031]|JWDCE863-10|BOLD:ACE6563
Ablabesmyia illinoensis|[14032]|JWDCJ434-11|BOLD:ACE6563
Ablabesmyia illinoensis|[14033]|CHURC1082-09|BOLD:ACE6563
Ablabesmyia|[14034]|CHURC011-06|BOLD:ACE6563
Ablabesmyia|[14035]|CHURC922-09|BOLD:ACE6563
Rheopelopia|[14036]|CHURC449-07|BOLD:AAF7638
Rheopelopia|[14037]|BSCHR037-10|BOLD:AAF7638
Rheopelopia|[14038]|CHURC667-09|BOLD:AAF7638
Rheopelopia|[14039]|CHURC491-08|BOLD:AAF7638
Smittia sp. ES11|[14040]|CHURC181-07|BOLD:AAB0378
Limnophyes sp. 12ES|[14041]|JWDCH583-10|BOLD:AAB1811
Limnophyes sp. 12ES|[14042]|CHURC171-07|BOLD:AAB1811
Limnophyes sp. 12ES|[14043]|CHURC904-09|BOLD:AAB1811
Limnophyes sp. 12ES|[14044]|CHURC183-07|BOLD:AAB1811
Limnophyes schnelli|[14045]|JWDCE203-10|BOLD:AAC9278
Limnophyes schnelli|[14046]|CHURC236-07|BOLD:AAC9278
Limnophyes schnelli|[14047]|CHURC255-07|BOLD:AAC9278
Limnophyes schnelli|[14048]|JWDCE202-10|BOLD:AAC9278
Limnophyes schnelli|[14049]|CHURC750-09|BOLD:AAC9278
Limnophyes schnelli|[14050]|JWDCH521-10|BOLD:AAC9278
Limnophyes schnelli|[14051]|JWDCD252-10|BOLD:AAC9278
Limnophyes schnelli|[14052]|CHURC745-09|BOLD:AAC9278
Limnophyes schnelli|[14053]|CHURC897-09|BOLD:AAC9278
Limnophyes schnelli|[14054]|CHURC139-07|BOLD:AAC9278
Limnophyes schnelli|[14055]|CHURC208-07|BOLD:AAC9278
Limnophyes schnelli|[14056]|CHURC049-06|BOLD:AAC9278
Limnophyes schnelli|[14057]|CHURC060-06|BOLD:AAC9278
Orthocladiinae|[14058]|SWSWP079-09|BOLD:AAG6992
Orthocladiinae|[14059]|SWSWP086-09|BOLD:AAG6992
Orthocladiinae|[14060]|JWDCF083-10|BOLD:AAP9001
Limnophyes ninae|[14061]|BSCHR078-10|BOLD:AAB1813
Limnophyes ninae|[14062]|BSCHR077-10|BOLD:AAB1813
Limnophyes ninae|[14063]|CHURC887-09|BOLD:AAB1813
Limnophyes ninae|[14064]|CHURC237-07|BOLD:AAB1813
Limnophyes minimus|[14065]|JWDCD435-10|BOLD:AAA8201
Limnophyes minimus|[14066]|CHURC050-06|BOLD:AAA8201
Limnophyes minimus|[14067]|JWDCC367-10|BOLD:AAA8201
Limnophyes minimus|[14068]|CHURC214-07|BOLD:AAA8201
Limnophyes minimus|[14069]|JWDCC355-10|BOLD:AAA8201
Limnophyes minimus|[14070]|CHURC028-06|BOLD:AAA8201
Limnophyes minimus|[14071]|CHURC043-06|BOLD:AAA8201
Limnophyes minimus|[14072]|CHURC1115-09|BOLD:AAA8201
Limnophyes minimus|[14073]|CHURC1095-09|BOLD:AAA8201
Limnophyes minimus|[14074]|CHURC108-07|BOLD:AAA8201
Limnophyes minimus|[14075]|CHURC243-07|BOLD:AAA8201
Limnophyes minimus|[14076]|CHURC274-07|BOLD:AAA8201
Limnophyes minimus|[14077]|CHURC854-09|BOLD:AAA8201
Limnophyes minimus|[14078]|CHURC1035-09|BOLD:AAA8201
Limnophyes minimus|[14079]|CHURC1059-09|BOLD:AAA8201
Limnophyes minimus|[14080]|CHURC1062-09|BOLD:AAA8201
Limnophyes minimus|[14081]|CHURC1071-09|BOLD:AAA8201
Limnophyes minimus|[14082]|CHURC1101-09|BOLD:AAA8201
Limnophyes minimus|[14083]|CHURC024-06|BOLD:AAA8201
Limnophyes minimus|[14084]|CHURC062-06|BOLD:AAA8201
Limnophyes minimus|[14085]|CHURC215-07|BOLD:AAA8201
Limnophyes minimus|[14086]|CHURC1106-09|BOLD:AAA8201
Limnophyes minimus|[14087]|CHURC241-07|BOLD:AAA8201
Limnophyes minimus|[14088]|CHURC216-07|BOLD:AAA8201
Limnophyes minimus|[14089]|CHURC142-07|BOLD:AAA8201
Limnophyes minimus|[14090]|CHURC786-09|BOLD:AAA8201
Limnophyes minimus|[14091]|JWDCG404-10|BOLD:AAA8201
Limnophyes minimus|[14092]|JWDCG402-10|BOLD:AAA8201
Limnophyes minimus|[14093]|JWDCD310-10|BOLD:AAA8201
Limnophyes minimus|[14094]|JWDCC372-10|BOLD:AAA8201
Limnophyes minimus|[14095]|JWDCC365-10|BOLD:AAA8201
Limnophyes minimus|[14096]|SWSWP068-09|BOLD:AAA8201
Limnophyes minimus|[14097]|JWDCC378-10|BOLD:AAA8201
Limnophyes minimus|[14098]|JWDCJ661-11|BOLD:AAA8201
Limnophyes minimus|[14099]|JWDCC369-10|BOLD:AAA8201
Limnophyes minimus|[14100]|JWDCD431-10|BOLD:AAA8201
Limnophyes minimus|[14101]|JWDCD472-10|BOLD:AAA8201
Limnophyes minimus|[14102]|CHURC1097-09|BOLD:AAA8201
Limnophyes minimus|[14103]|CHURC209-07|BOLD:AAA8201
Limnophyes minimus|[14104]|CHURC048-06|BOLD:AAA8201
Limnophyes minimus|[14105]|CHURC186-07|BOLD:AAA8201
Limnophyes minimus|[14106]|JWDCD272-10|BOLD:AAA8201
Limnophyes minimus|[14107]|CHURC1098-09|BOLD:AAA8201
Limnophyes minimus|[14108]|CHURC1109-09|BOLD:AAA8201
Limnophyes minimus|[14109]|CHURC1056-09|BOLD:AAA8202
Limnophyes minimus|[14110]|CHURC1067-09|BOLD:AAA8202
Limnophyes minimus|[14111]|CHURC157-07|BOLD:AAA8202
Limnophyes minimus|[14112]|CHURC159-07|BOLD:AAA8202
Limnophyes minimus|[14113]|CHURC158-07|BOLD:AAA8202
Limnophyes minimus|[14114]|CHURC789-09|BOLD:AAA8202
Limnophyes minimus|[14115]|CHURC1064-09|BOLD:AAA8202
Limnophyes minimus|[14116]|SWSWP009-09|BOLD:AAA8203
Limnophyes minimus|[14117]|CHURC1113-09|BOLD:AAA8203
Limnophyes sp. ES13|[14118]|CHURC170-07|BOLD:ACE8732
Limnophyes sp. ES13|[14119]|CHURC757-09|BOLD:ACE8732
Limnophyes sp. ES13|[14120]|CHURC812-09|BOLD:ACE8732
Limnophyes sp. ES13|[14121]|CHURC235-07|BOLD:ACE8732
Limnophyes sp. ES13|[14122]|CHURC254-07|BOLD:ACE8732
Limnophyes sp. ES13|[14123]|CHURC238-07|BOLD:AAB1808
Limnophyes sp. ES13|[14124]|CHURC123-07|BOLD:AAB1808
Limnophyes sp. ES13|[14125]|CHURC074-06|BOLD:AAB1808
Limnophyes sp. ES13|[14126]|CHURC022-06|BOLD:AAB1808
Orthocladiinae|[14127]|JWDCE028-10|BOLD:AAP1832
Smittia|[14128]|JWDCD383-10|BOLD:AAL5086
Smittia|[14129]|JWDCJ063-11|BOLD:AAQ3392
Smittia|[14130]|JWDCD433-10|BOLD:AAL7386

Smittia|[14128]|JWDCD383-10|BOLD:AAL5086
Smittia|[14129]|JWDCJ063-11|BOLD:AAQ3392
Smittia|[14130]|JWDCD433-10|BOLD:AAL7386
Smittia|[14131]|JWDCD382-10|BOLD:AAL7384
Smittia|[14132]|JWDCC359-10|BOLD:AAL7316
Smittia|[14133]|JWDCD254-10|BOLD:AAG6994
Smittia|[14134]|JWDCD384-10|BOLD:AAG6994
Smittia|[14135]|JWDCD271-10|BOLD:AAG6994
Smittia|[14136]|JWDCD276-10|BOLD:AAG6994
Smittia|[14137]|SWSWP094-09|BOLD:AAG6994
Smittia|[14138]|JWDCE101-10|BOLD:AAG6994
Smittia|[14139]|JWDCE102-10|BOLD:AAG6994
Smittia|[14140]|JWDCE103-10|BOLD:AAG6994
Orthocladiinae|[14141]|JWDCF345-10|BOLD:AAP9003
Limnophyes|[14142]|JWDCG819-10|BOLD:AAQ0626
Eukiefferiella claripennis|[14143]|CHURC817-09|BOLD:AHC6527
Eukiefferiella claripennis|[14144]|CHURC169-07|BOLD:AHC6527
Metriocnemus sp. 4ES|[14145]|CHURC145-07|BOLD:AAA9435
Metriocnemus sp. 4ES|[14146]|CHURC280-07|BOLD:AAA9431
Metriocnemus sp. 4ES|[14147]|CHURC036-06|BOLD:AAA9431
Metriocnemus sp. 4ES|[14148]|CHURC037-06|BOLD:ABX5809
Euryhapsis|[14149]|BSCHR102-10|BOLD:AAN5377
Euryhapsis cilium|[14150]|BSCHR149-10|BOLD:ACE3103
Euryhapsis cilium|[14151]|BSCHR150-10|BOLD:ACE3103
Brillia flavifrons|[14152]|CHURC485-08|BOLD:AAI3832
Orthocladiinae|[14153]|JWDCD270-10|BOLD:AAL7383
Cricotopus intersectus|[14154]|KSCLC087-09|BOLD:AAA5302
Cricotopus intersectus|[14155]|CHURC519-08|BOLD:AAA5302
Cricotopus intersectus|[14156]|CHURC056-06|BOLD:AAA5302
Cricotopus intersectus|[14157]|CHURC059-06|BOLD:AAA5302
Cricotopus intersectus|[14158]|CHURC364-07|BOLD:AAA5302
Cricotopus intersectus|[14159]|CHURC057-06|BOLD:AAA5302
Cricotopus perniger|[14160]|CHURC416-07|BOLD:AAA5313
Orthocladiinae|[14161]|JWDCE144-10|BOLD:AAP5144
Limnophyes pr. madeirae|[14162]|JWDCC376-10|BOLD:AAB1812
Limnophyes pr. madeirae|[14163]|CHURC760-09|BOLD:AAB1812
Limnophyes pr. madeirae|[14164]|CHURC753-09|BOLD:AAB1812
Limnophyes pr. madeirae|[14165]|JWDCC358-10|BOLD:AAB1812
Limnophyes pr. madeirae|[14166]|CHURC733-09|BOLD:AAB1812
Limnophyes pr. madeirae|[14167]|CHURC772-09|BOLD:AAB1812
Limnophyes pr. madeirae|[14168]|CHURC806-09|BOLD:AAB1812
Limnophyes pr. madeirae|[14169]|CHURC212-07|BOLD:AAB1812
Limnophyes pr. madeirae|[14170]|CHURC240-07|BOLD:AAB1812
Limnophyes pr. madeirae|[14171]|CHURC857-09|BOLD:AAB1812
Orthocladius sp. 9TE|[14172]|SWSWP051-09|BOLD:AAD4029
Orthocladius sp. 9TE|[14173]|JWDCE845-10|BOLD:AAD4029
Orthocladius subletteorum|[14174]|CHURC177-07|BOLD:AAF0487
Orthocladius subletteorum|[14175]|JWDCG029-10|BOLD:AAF0488
Orthocladius subletteorum|[14176]|CHURC1068-09|BOLD:AAF0488
Orthocladius subletteorum|[14177]|CHURC530-08|BOLD:AAF0488
Orthocladius sp. TE05|[14178]|CHURC206-07|BOLD:AAC0068
Orthocladius sp. TE05|[14179]|CHURC233-07|BOLD:AAC0068
Orthocladius sp. TE05|[14180]|CHURC046-06|BOLD:AAC0068
Orthocladius sp. TE05|[14181]|CHURC658-09|BOLD:AAC0068
Orthocladius sp. TE14|[14182]|CHURC1079-09|BOLD:AAG1001
Orthocladius oblidens|[14183]|JWDCF577-10|BOLD:AAD8971
Orthocladius oblidens|[14184]|CHURC272-07|BOLD:AAD8971
Orthocladius oblidens|[14185]|CHURC756-09|BOLD:AAD8971
Orthocladius oblidens|[14186]|CHURC884-09|BOLD:AAD8971
Orthocladius oblidens|[14187]|BSCHR068-10|BOLD:AAD8971
Orthocladius oblidens|[14188]|CHURC728-09|BOLD:AAD8971
Orthocladius oblidens|[14189]|CHURC550-08|BOLD:AAD8971
Orthocladius sp. TE13|[14190]|CHURC642-09|BOLD:AAC0069
Orthocladius sp. TE13|[14191]|CHURC646-09|BOLD:AAC0069
Orthocladius sp. TE13|[14192]|CHURC531-08|BOLD:AAC0069
Orthocladius sp. TE13|[14193]|CHURC664-09|BOLD:AAC0069
Orthocladius sp. TE13|[14194]|CHURC663-09|BOLD:AAC0069
Orthocladius sp. TE13|[14195]|CHURC656-09|BOLD:AAC0069
Orthocladius sp. TE13|[14196]|CHURC537-08|BOLD:AAC0069
Orthocladius sp. TE13|[14197]|CHURC655-09|BOLD:AAC0069
Orthocladius sp. TE13|[14198]|CHURC647-09|BOLD:AAC0069
Orthocladius sp. TE13|[14199]|CHURC653-09|BOLD:AAC0069
Orthocladius sp. TE13|[14200]|CHURC641-09|BOLD:AAC0069
Metriocnemus|[14201]|JWDCE737-10|BOLD:AAP1831
Metriocnemus|[14202]|JWDCE027-10|BOLD:AAP1831
Metriocnemus|[14203]|JWDCE050-10|BOLD:AAP1831
Metriocnemus|[14204]|JWDCE051-10|BOLD:AAP1831
Metriocnemus|[14205]|JWDCI356-10|BOLD:AAV5956
Metriocnemus|[14206]|JWDCJ723-11|BOLD:AAV5956
Metriocnemus|[14207]|JWDCG541-10|BOLD:AAV5956
Metriocnemus sp. 7ES|[14208]|MHMOS167-07|BOLD:AAA9432
Metriocnemus sp. 7ES|[14209]|JWDCG542-10|BOLD:AAA9432
Metriocnemus sp. 7ES|[14210]|JWDCI239-10|BOLD:AAA9432
Metriocnemus sp. 7ES|[14211]|JWDCG540-10|BOLD:AAA9432
Metriocnemus sp. 7ES|[14212]|JWDCH760-10|BOLD:AAA9432
Metriocnemus sp. 7ES|[14213]|CHURC085-06|BOLD:AAA9432
Metriocnemus sp. 7ES|[14214]|CHURC086-06|BOLD:AAA9432
Smittia sp. ES12|[14215]|CHURC779-09|BOLD:AAB0377
Smittia sp. ES12|[14216]|CHURC836-09|BOLD:AAB0377
Smittia sp. ES12|[14217]|CHURC818-09|BOLD:AAB0377
Smittia sp. ES12|[14218]|CHURC783-09|BOLD:AAB0377
Smittia sp. ES12|[14219]|CHURC830-09|BOLD:AAB0377
Smittia sp. ES12|[14220]|CHURC155-07|BOLD:AAB0377
Smittia sp. ES12|[14221]|CHURC834-09|BOLD:AAB0377
Smittia sp. ES12|[14222]|CHURC823-09|BOLD:AAB0377
Smittia sp. ES12|[14223]|JWDCC354-10|BOLD:AAB0377
Smittia sp. ES12|[14224]|CHURC826-09|BOLD:AAB0377
Smittia sp. ES12|[14225]|CHURC838-09|BOLD:AAB0377
Smittia sp. ES12|[14226]|CHURC1069-09|BOLD:AAB0377
Smittia sp. ES12|[14227]|CHURC1045-09|BOLD:AAB0377
Smittia sp. ES12|[14228]|CHURC832-09|BOLD:AAB0377
Smittia sp. ES12|[14229]|CHURC1057-09|BOLD:AAB0377
Smittia sp. ES12|[14230]|CHURC781-09|BOLD:AAB0377

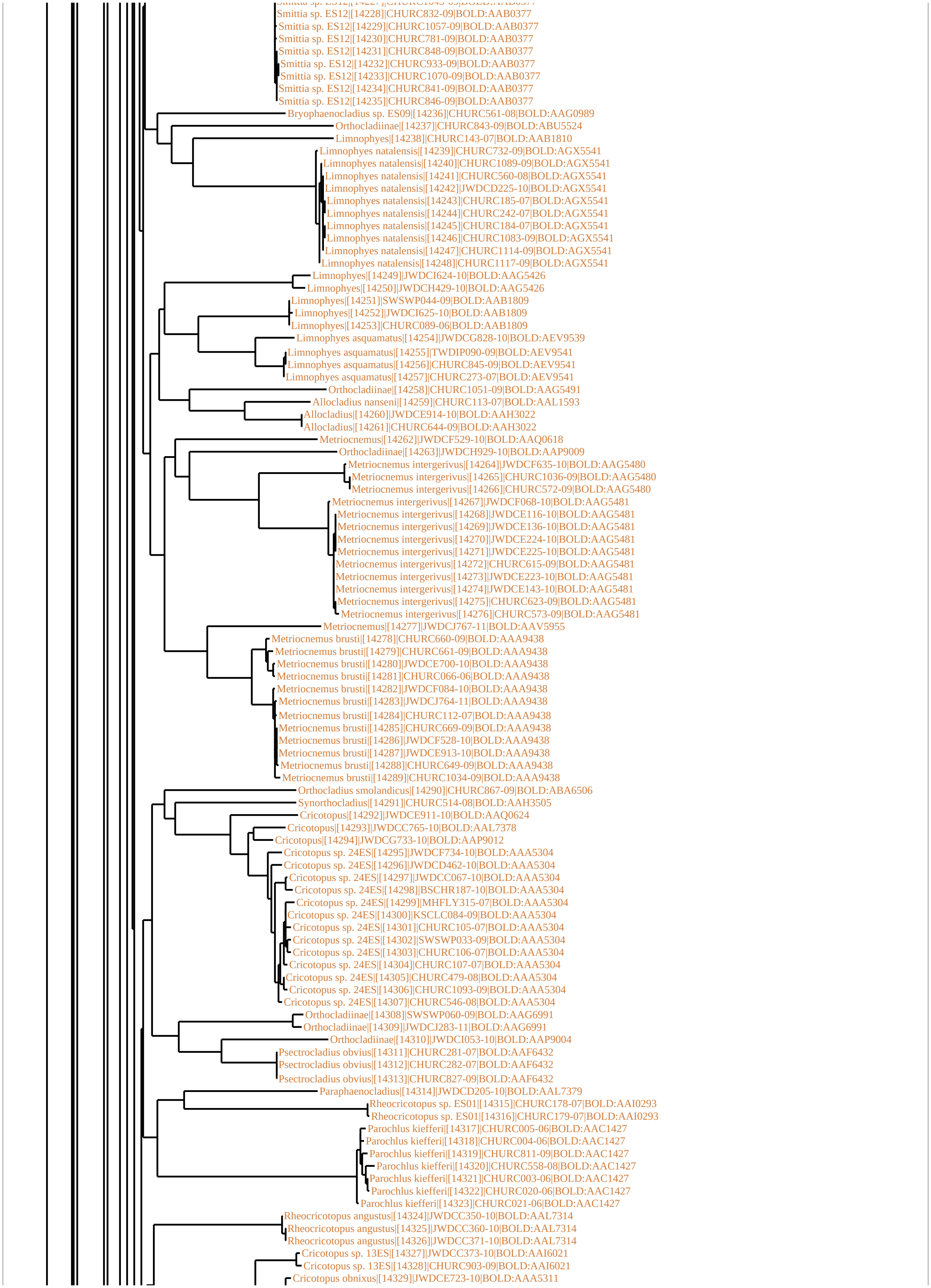
Smittia sp. ES12|[14228]|CHURC832-09|BOLD:AAB0377
Smittia sp. ES12|[14229]|CHURC1057-09|BOLD:AAB0377
Smittia sp. ES12|[14230]|CHURC781-09|BOLD:AAB0377
Smittia sp. ES12|[14231]|CHURC848-09|BOLD:AAB0377
Smittia sp. ES12|[14232]|CHURC933-09|BOLD:AAB0377
Smittia sp. ES12|[14233]|CHURC1070-09|BOLD:AAB0377
Smittia sp. ES12|[14234]|CHURC841-09|BOLD:AAB0377
Smittia sp. ES12|[14235]|CHURC846-09|BOLD:AAB0377
Bryophaenocladius sp. ES09|[14236]|CHURC561-08|BOLD:AAG0989
Orthocladiinae|[14237]|CHURC843-09|BOLD:ABU5524
Limnophyes|[14238]|CHURC143-07|BOLD:AAB1810
Limnophyes natalensis|[14239]|CHURC732-09|BOLD:AGX5541
Limnophyes natalensis|[14240]|CHURC1089-09|BOLD:AGX5541
Limnophyes natalensis|[14241]|CHURC560-08|BOLD:AGX5541
Limnophyes natalensis|[14242]|JWDCD225-10|BOLD:AGX5541
Limnophyes natalensis|[14243]|CHURC185-07|BOLD:AGX5541
Limnophyes natalensis|[14244]|CHURC242-07|BOLD:AGX5541
Limnophyes natalensis|[14245]|CHURC184-07|BOLD:AGX5541
Limnophyes natalensis|[14246]|CHURC1083-09|BOLD:AGX5541
Limnophyes natalensis|[14247]|CHURC1114-09|BOLD:AGX5541
Limnophyes natalensis|[14248]|CHURC1117-09|BOLD:AGX5541
Limnophyes|[14249]|JWDCI624-10|BOLD:AAG5426
Limnophyes|[14250]|JWDCH429-10|BOLD:AAG5426
Limnophyes|[14251]|SWSWP044-09|BOLD:AAB1809
Limnophyes|[14252]|JWDCI625-10|BOLD:AAB1809
Limnophyes|[14253]|CHURC089-06|BOLD:AAB1809
Limnophyes asquamatus|[14254]|JWDCG828-10|BOLD:AEV9539
Limnophyes asquamatus|[14255]|TWDIP090-09|BOLD:AEV9541
Limnophyes asquamatus|[14256]|CHURC845-09|BOLD:AEV9541
Limnophyes asquamatus|[14257]|CHURC273-07|BOLD:AEV9541
Orthocladiinae|[14258]|CHURC1051-09|BOLD:AAG5491
Allocladius nanseni|[14259]|CHURC113-07|BOLD:AAL1593
Allocladius|[14260]|JWDCE914-10|BOLD:AAH3022
Allocladius|[14261]|CHURC644-09|BOLD:AAH3022
Metriocnemus|[14262]|JWDCF529-10|BOLD:AAQ0618
Orthocladiinae|[14263]|JWDCH929-10|BOLD:AAP9009
Metriocnemus intergerivus|[14264]|JWDCF635-10|BOLD:AAG5480
Metriocnemus intergerivus|[14265]|CHURC1036-09|BOLD:AAG5480
Metriocnemus intergerivus|[14266]|CHURC572-09|BOLD:AAG5480
Metriocnemus intergerivus|[14267]|JWDCF068-10|BOLD:AAG5481
Metriocnemus intergerivus|[14268]|JWDCE116-10|BOLD:AAG5481
Metriocnemus intergerivus|[14269]|JWDCE136-10|BOLD:AAG5481
Metriocnemus intergerivus|[14270]|JWDCE224-10|BOLD:AAG5481
Metriocnemus intergerivus|[14271]|JWDCE225-10|BOLD:AAG5481
Metriocnemus intergerivus|[14272]|CHURC615-09|BOLD:AAG5481
Metriocnemus intergerivus|[14273]|JWDCE223-10|BOLD:AAG5481
Metriocnemus intergerivus|[14274]|JWDCE143-10|BOLD:AAG5481
Metriocnemus intergerivus|[14275]|CHURC623-09|BOLD:AAG5481
Metriocnemus intergerivus|[14276]|CHURC573-09|BOLD:AAG5481
Metriocnemus|[14277]|JWDCJ767-11|BOLD:AAV5955
Metriocnemus brusti|[14278]|CHURC660-09|BOLD:AAA9438
Metriocnemus brusti|[14279]|CHURC661-09|BOLD:AAA9438
Metriocnemus brusti|[14280]|JWDCE700-10|BOLD:AAA9438
Metriocnemus brusti|[14281]|CHURC066-06|BOLD:AAA9438
Metriocnemus brusti|[14282]|JWDCF084-10|BOLD:AAA9438
Metriocnemus brusti|[14283]|JWDCJ764-11|BOLD:AAA9438
Metriocnemus brusti|[14284]|CHURC112-07|BOLD:AAA9438
Metriocnemus brusti|[14285]|CHURC669-09|BOLD:AAA9438
Metriocnemus brusti|[14286]|JWDCF528-10|BOLD:AAA9438
Metriocnemus brusti|[14287]|JWDCE913-10|BOLD:AAA9438
Metriocnemus brusti|[14288]|CHURC649-09|BOLD:AAA9438
Metriocnemus brusti|[14289]|CHURC1034-09|BOLD:AAA9438
Orthocladius smolandicus|[14290]|CHURC867-09|BOLD:ABA6506
Synorthocladius|[14291]|CHURC514-08|BOLD:AAH3505
Cricotopus|[14292]|JWDCE911-10|BOLD:AAQ0624
Cricotopus|[14293]|JWDCC765-10|BOLD:AAL7378
Cricotopus|[14294]|JWDCG733-10|BOLD:AAP9012
Cricotopus sp. 24ES|[14295]|JWDCF734-10|BOLD:AAA5304
Cricotopus sp. 24ES|[14296]|JWDCD462-10|BOLD:AAA5304
Cricotopus sp. 24ES|[14297]|JWDCC067-10|BOLD:AAA5304
Cricotopus sp. 24ES|[14298]|BSCHR187-10|BOLD:AAA5304
Cricotopus sp. 24ES|[14299]|MHFLY315-07|BOLD:AAA5304
Cricotopus sp. 24ES|[14300]|KSCLC084-09|BOLD:AAA5304
Cricotopus sp. 24ES|[14301]|CHURC105-07|BOLD:AAA5304
Cricotopus sp. 24ES|[14302]|SWSWP033-09|BOLD:AAA5304
Cricotopus sp. 24ES|[14303]|CHURC106-07|BOLD:AAA5304
Cricotopus sp. 24ES|[14304]|CHURC107-07|BOLD:AAA5304
Cricotopus sp. 24ES|[14305]|CHURC479-08|BOLD:AAA5304
Cricotopus sp. 24ES|[14306]|CHURC1093-09|BOLD:AAA5304
Cricotopus sp. 24ES|[14307]|CHURC546-08|BOLD:AAA5304
Orthocladiinae|[14308]|SWSWP060-09|BOLD:AAG6991
Orthocladiinae|[14309]|JWDCJ283-11|BOLD:AAG6991
Orthocladiinae|[14310]|JWDCI053-10|BOLD:AAP9004
Psectrocladius obvius|[14311]|CHURC281-07|BOLD:AAF6432
Psectrocladius obvius|[14312]|CHURC282-07|BOLD:AAF6432
Psectrocladius obvius|[14313]|CHURC827-09|BOLD:AAF6432
Paraphaenocladius|[14314]|JWDCD205-10|BOLD:AAL7379
Rheocricotopus sp. ES01|[14315]|CHURC178-07|BOLD:AAI0293
Rheocricotopus sp. ES01|[14316]|CHURC179-07|BOLD:AAI0293
Parochlus kiefferi|[14317]|CHURC005-06|BOLD:AAC1427
Parochlus kiefferi|[14318]|CHURC004-06|BOLD:AAC1427
Parochlus kiefferi|[14319]|CHURC811-09|BOLD:AAC1427
Parochlus kiefferi|[14320]|CHURC558-08|BOLD:AAC1427
Parochlus kiefferi|[14321]|CHURC003-06|BOLD:AAC1427
Parochlus kiefferi|[14322]|CHURC020-06|BOLD:AAC1427
Parochlus kiefferi|[14323]|CHURC021-06|BOLD:AAC1427
Rheocricotopus angustus|[14324]|JWDCC350-10|BOLD:AAL7314
Rheocricotopus angustus|[14325]|JWDCC360-10|BOLD:AAL7314
Rheocricotopus angustus|[14326]|JWDCC371-10|BOLD:AAL7314
Cricotopus sp. 13ES|[14327]|JWDCC373-10|BOLD:AAI6021
Cricotopus sp. 13ES|[14328]|CHURC903-09|BOLD:AAI6021
Cricotopus obnixus|[14329]|JWDCE723-10|BOLD:AAA5311

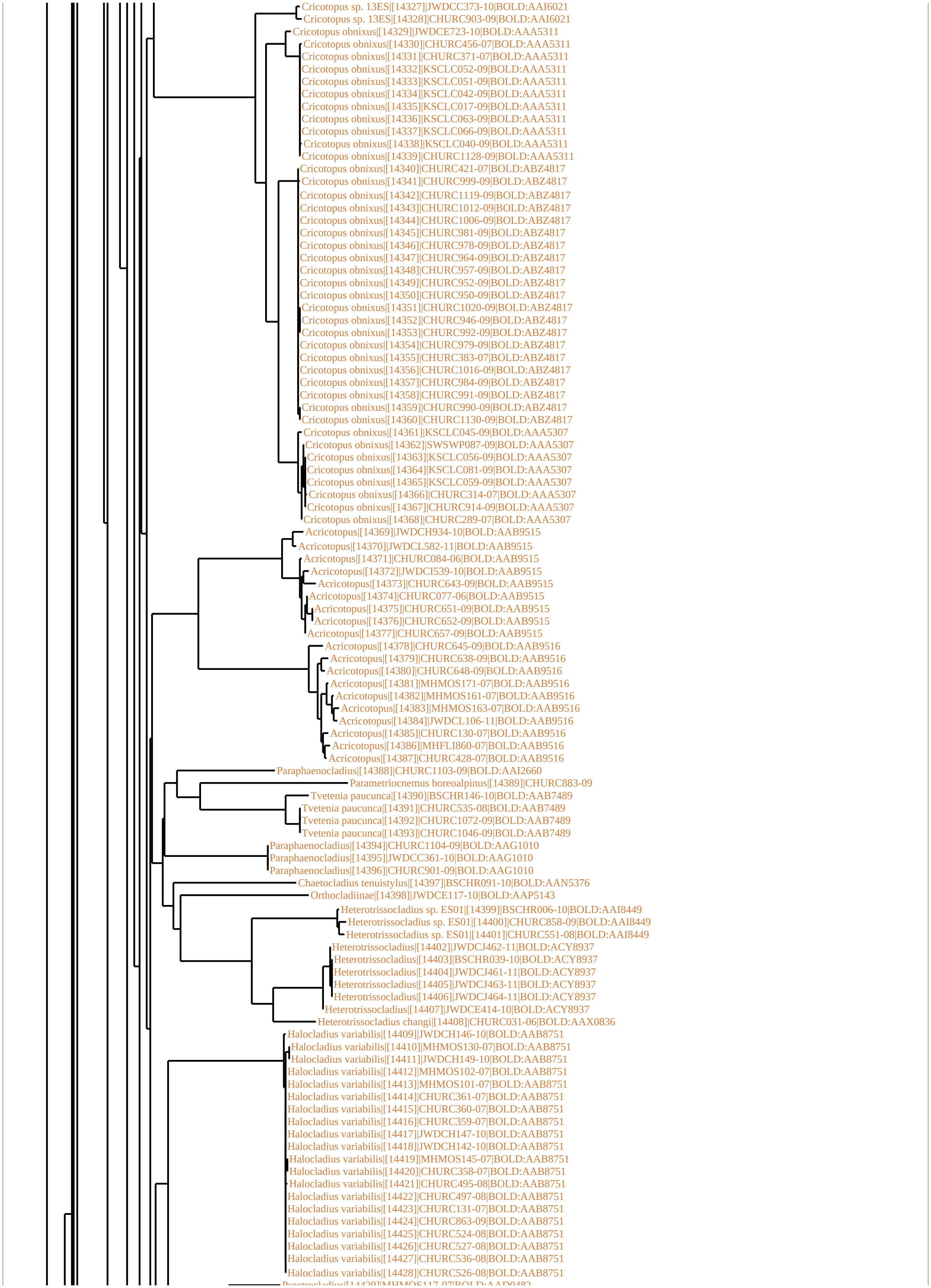
Cricotopus sp. 13ES|[14327]|JWDCC373-10|BOLD:AAI6021
Cricotopus sp. 13ES|[14328]|CHURC903-09|BOLD:AAI6021
Cricotopus obnixus|[14329]|JWDCE723-10|BOLD:AAA5311
Cricotopus obnixus|[14330]|CHURC456-07|BOLD:AAA5311
Cricotopus obnixus|[14331]|CHURC371-07|BOLD:AAA5311
Cricotopus obnixus|[14332]|KSCLC052-09|BOLD:AAA5311
Cricotopus obnixus|[14333]|KSCLC051-09|BOLD:AAA5311
Cricotopus obnixus|[14334]|KSCLC042-09|BOLD:AAA5311
Cricotopus obnixus|[14335]|KSCLC017-09|BOLD:AAA5311
Cricotopus obnixus|[14336]|KSCLC063-09|BOLD:AAA5311
Cricotopus obnixus|[14337]|KSCLC066-09|BOLD:AAA5311
Cricotopus obnixus|[14338]|KSCLC040-09|BOLD:AAA5311
Cricotopus obnixus|[14339]|CHURC1128-09|BOLD:AAA5311
Cricotopus obnixus|[14340]|CHURC421-07|BOLD:ABZ4817
Cricotopus obnixus|[14341]|CHURC999-09|BOLD:ABZ4817
Cricotopus obnixus|[14342]|CHURC1119-09|BOLD:ABZ4817
Cricotopus obnixus|[14343]|CHURC1012-09|BOLD:ABZ4817
Cricotopus obnixus|[14344]|CHURC1006-09|BOLD:ABZ4817
Cricotopus obnixus|[14345]|CHURC981-09|BOLD:ABZ4817
Cricotopus obnixus|[14346]|CHURC978-09|BOLD:ABZ4817
Cricotopus obnixus|[14347]|CHURC964-09|BOLD:ABZ4817
Cricotopus obnixus|[14348]|CHURC957-09|BOLD:ABZ4817
Cricotopus obnixus|[14349]|CHURC952-09|BOLD:ABZ4817
Cricotopus obnixus|[14350]|CHURC950-09|BOLD:ABZ4817
Cricotopus obnixus|[14351]|CHURC1020-09|BOLD:ABZ4817
Cricotopus obnixus|[14352]|CHURC946-09|BOLD:ABZ4817
Cricotopus obnixus|[14353]|CHURC992-09|BOLD:ABZ4817
Cricotopus obnixus|[14354]|CHURC979-09|BOLD:ABZ4817
Cricotopus obnixus|[14355]|CHURC383-07|BOLD:ABZ4817
Cricotopus obnixus|[14356]|CHURC1016-09|BOLD:ABZ4817
Cricotopus obnixus|[14357]|CHURC984-09|BOLD:ABZ4817
Cricotopus obnixus|[14358]|CHURC991-09|BOLD:ABZ4817
Cricotopus obnixus|[14359]|CHURC990-09|BOLD:ABZ4817
Cricotopus obnixus|[14360]|CHURC1130-09|BOLD:ABZ4817
Cricotopus obnixus|[14361]|KSCLC045-09|BOLD:AAA5307
Cricotopus obnixus|[14362]|SWSWP087-09|BOLD:AAA5307
Cricotopus obnixus|[14363]|KSCLC056-09|BOLD:AAA5307
Cricotopus obnixus|[14364]|KSCLC081-09|BOLD:AAA5307
Cricotopus obnixus|[14365]|KSCLC059-09|BOLD:AAA5307
Cricotopus obnixus|[14366]|CHURC314-07|BOLD:AAA5307
Cricotopus obnixus|[14367]|CHURC914-09|BOLD:AAA5307
Cricotopus obnixus|[14368]|CHURC289-07|BOLD:AAA5307
Acricotopus|[14369]|JWDCH934-10|BOLD:AAB9515
Acricotopus|[14370]|JWDCL582-11|BOLD:AAB9515
Acricotopus|[14371]|CHURC084-06|BOLD:AAB9515
Acricotopus|[14372]|JWDCI539-10|BOLD:AAB9515
Acricotopus|[14373]|CHURC643-09|BOLD:AAB9515
Acricotopus|[14374]|CHURC077-06|BOLD:AAB9515
Acricotopus|[14375]|CHURC651-09|BOLD:AAB9515
Acricotopus|[14376]|CHURC652-09|BOLD:AAB9515
Acricotopus|[14377]|CHURC657-09|BOLD:AAB9515
Acricotopus|[14378]|CHURC645-09|BOLD:AAB9516
Acricotopus|[14379]|CHURC638-09|BOLD:AAB9516
Acricotopus|[14380]|CHURC648-09|BOLD:AAB9516
Acricotopus|[14381]|MHMOS171-07|BOLD:AAB9516
Acricotopus|[14382]|MHMOS161-07|BOLD:AAB9516
Acricotopus|[14383]|MHMOS163-07|BOLD:AAB9516
Acricotopus|[14384]|JWDCL106-11|BOLD:AAB9516
Acricotopus|[14385]|CHURC130-07|BOLD:AAB9516
Acricotopus|[14386]|MHFLI860-07|BOLD:AAB9516
Acricotopus|[14387]|CHURC428-07|BOLD:AAB9516
Paraphaenocladius|[14388]|CHURC1103-09|BOLD:AAI2660
Parametriocnemus boreoalpinus|[14389]|CHURC883-09
Tvetenia paucunca|[14390]|BSCHR146-10|BOLD:AAB7489
Tvetenia paucunca|[14391]|CHURC535-08|BOLD:AAB7489
Tvetenia paucunca|[14392]|CHURC1072-09|BOLD:AAB7489
Tvetenia paucunca|[14393]|CHURC1046-09|BOLD:AAB7489
Paraphaenocladius|[14394]|CHURC1104-09|BOLD:AAG1010
Paraphaenocladius|[14395]|JWDCC361-10|BOLD:AAG1010
Paraphaenocladius|[14396]|CHURC901-09|BOLD:AAG1010
Chaetocladius tenuistylus|[14397]|BSCHR091-10|BOLD:AAN5376
Orthocladiinae|[14398]|JWDCE117-10|BOLD:AAP5143
Heterotrissocladius sp. ES01|[14399]|BSCHR006-10|BOLD:AAI8449
Heterotrissocladius sp. ES01|[14400]|CHURC858-09|BOLD:AAI8449
Heterotrissocladius sp. ES01|[14401]|CHURC551-08|BOLD:AAI8449
Heterotrissocladius|[14402]|JWDCJ462-11|BOLD:ACY8937
Heterotrissocladius|[14403]|BSCHR039-10|BOLD:ACY8937
Heterotrissocladius|[14404]|JWDCJ461-11|BOLD:ACY8937
Heterotrissocladius|[14405]|JWDCJ463-11|BOLD:ACY8937
Heterotrissocladius|[14406]|JWDCJ464-11|BOLD:ACY8937
Heterotrissocladius|[14407]|JWDCE414-10|BOLD:ACY8937
Heterotrissocladius changi|[14408]|CHURC031-06|BOLD:AAX0836
Halocladius variabilis|[14409]|JWDCH146-10|BOLD:AAB8751
Halocladius variabilis|[14410]|MHMOS130-07|BOLD:AAB8751
Halocladius variabilis|[14411]|JWDCH149-10|BOLD:AAB8751
Halocladius variabilis|[14412]|MHMOS102-07|BOLD:AAB8751
Halocladius variabilis|[14413]|MHMOS101-07|BOLD:AAB8751
Halocladius variabilis|[14414]|CHURC361-07|BOLD:AAB8751
Halocladius variabilis|[14415]|CHURC360-07|BOLD:AAB8751
Halocladius variabilis|[14416]|CHURC359-07|BOLD:AAB8751
Halocladius variabilis|[14417]|JWDCH147-10|BOLD:AAB8751
Halocladius variabilis|[14418]|JWDCH142-10|BOLD:AAB8751
Halocladius variabilis|[14419]|MHMOS145-07|BOLD:AAB8751
Halocladius variabilis|[14420]|CHURC358-07|BOLD:AAB8751
Halocladius variabilis|[14421]|CHURC495-08|BOLD:AAB8751
Halocladius variabilis|[14422]|CHURC497-08|BOLD:AAB8751
Halocladius variabilis|[14423]|CHURC131-07|BOLD:AAB8751
Halocladius variabilis|[14424]|CHURC863-09|BOLD:AAB8751
Halocladius variabilis|[14425]|CHURC524-08|BOLD:AAB8751
Halocladius variabilis|[14426]|CHURC527-08|BOLD:AAB8751
Halocladius variabilis|[14427]|CHURC536-08|BOLD:AAB8751
Halocladius variabilis|[14428]|CHURC526-08|BOLD:AAB8751

[illegible]
Halocladius variabilis|[14427]|CHURC536-08|BOLD:AAB8751
Halocladius variabilis|[14428]|CHURC526-08|BOLD:AAB8751
Psectrocladius|[14429]|MHMOS117-07|BOLD:AAD0482
Psectrocladius barbimanus|[14430]|CHURC547-08|BOLD:AAD0484
Psectrocladius barbimanus|[14431]|CHURC896-09|BOLD:AAD0484
Psectrocladius barbimanus|[14432]|JWDCE647-10|BOLD:AAD0483
Psectrocladius barbimanus|[14433]|CHURC548-08|BOLD:AAD0483
Psectrocladius barbimanus|[14434]|JWDCG821-10|BOLD:AAD0483
Psectrocladius barbimanus|[14435]|CHURC481-08|BOLD:AAD0483
Psectrocladius barbimanus|[14436]|CHURC873-09|BOLD:AAD0483
Psectrocladius barbimanus|[14437]|CHURC625-09|BOLD:AAD0483
Psectrocladius|[14438]|SWSWP095-09|BOLD:AAG6993
Psectrocladius|[14439]|SWSWP093-09|BOLD:AAG6993
Psectrocladius|[14440]|SWSWP089-09|BOLD:AAG6993
Psectrocladius|[14441]|SWSWP090-09|BOLD:AAG6993
Psectrocladius|[14442]|JWDCI580-10|BOLD:AAG6993
Smittia|[14443]|CHURC574-09|BOLD:AAG5482
Psectrocladius sp. 4TE|[14444]|CHURC512-08|BOLD:AAK9986
Paraphaenocladius sp. 4ES|[14445]|JWDCD253-10|BOLD:AAL7381
Paraphaenocladius impensus|[14446]|JWDCD434-10|BOLD:ACX4638
Paraphaenocladius impensus|[14447]|JWDCC494-10|BOLD:ACX4638
Paraphaenocladius impensus|[14448]|CHURC931-09|BOLD:ACX4638
Paraphaenocladius impensus|[14449]|JWDCD388-10|BOLD:AAC4201
Paraphaenocladius impensus|[14450]|JWDCD269-10|BOLD:AAC4201
Paraphaenocladius impensus|[14451]|CHURC1027-09|BOLD:AAC4201
Paraphaenocladius impensus|[14452]|CHURC1096-09|BOLD:AAC4201
Paraphaenocladius impensus|[14453]|JWDCD387-10|BOLD:AAC4201
Paraphaenocladius impensus|[14454]|CHURC1105-09|BOLD:AAC4201
Paraphaenocladius cf. nasthecus|[14455]|CHURC559-08|BOLD:AAX1168
Paraphaenocladius impensus|[14456]|JWDCD359-10|BOLD:AAI2661
Paraphaenocladius impensus|[14457]|JWDCD204-10|BOLD:AAI2661
Paraphaenocladius impensus|[14458]|CHURC1107-09|BOLD:AAI2661
Paraphaenocladius impensus|[14459]|CHURC1108-09|BOLD:AAI2661
Paraphaenocladius exagitans|[14460]|CHURC275-07|BOLD:AAI2658
Paraphaenocladius impensus|[14461]|CHURC336-07|BOLD:AAC4197
Paraphaenocladius impensus|[14462]|CHURC665-09|BOLD:AAC4197
Paraphaenocladius impensus|[14463]|CHURC040-06|BOLD:AAC4197
Paraphaenocladius impensus|[14464]|CHURC041-06|BOLD:AAC4197
Orthocladiinae|[14465]|JWDCD193-10|BOLD:AAL7890
Nanocladius dichromus|[14466]|CHURC156-07|BOLD:AAK9265
Nanocladius|[14467]|CHURC729-09|BOLD:AAF7613
Nanocladius|[14468]|CHURC771-09|BOLD:AAF7613
Nanocladius|[14469]|CHURC042-06|BOLD:AAF7613
Orthocladius mallochi|[14470]|CHURC1076-09|BOLD:AAB2644
Orthocladius mallochi|[14471]|CHURC490-08|BOLD:AAB2644
Orthocladius mallochi|[14472]|CHURC498-08|BOLD:AAB2644
Orthocladius mallochi|[14473]|CHURC864-09|BOLD:AAB2644
Orthocladius mallochi|[14474]|CHURC540-08|BOLD:AAB2644
Orthocladius mallochi|[14475]|JWDCH762-10|BOLD:AAB2644
Orthocladius mallochi|[14476]|CHURC739-09|BOLD:AAB2644
Orthocladius mallochi|[14477]|CHURC768-09|BOLD:AAB2644
Orthocladius mallochi|[14478]|CHURC778-09|BOLD:AAB2644
Orthocladius mallochi|[14479]|CHURC270-07|BOLD:AAB2644
Orthocladius mallochi|[14480]|CHURC029-06|BOLD:AAB2644
Orthocladius mallochi|[14481]|CHURC045-06|BOLD:AAB2644
Gymnometriocnemus|[14482]|JWDCD361-10|BOLD:AAM6261
Gymnometriocnemus|[14483]|JWDCD308-10|BOLD:AAM6261
Gymnometriocnemus|[14484]|JWDCD309-10|BOLD:AAI1981
Gymnometriocnemus|[14485]|JWDCD307-10|BOLD:AAI1981
Cricotopus sp. 14ES|[14486]|CHURC862-09|BOLD:ACV2627
Cricotopus sp. 8ES|[14487]|BSCHR105-10|BOLD:AAA5309
Cricotopus sp. 8ES|[14488]|JWDCJ766-11|BOLD:AAA5309
Cricotopus sp. 8ES|[14489]|JWDCG403-10|BOLD:AAA5309
Cricotopus sp. 8ES|[14490]|CHURC362-07|BOLD:AAA5309
Cricotopus sp. 21ES|[14491]|JWDCF147-10|BOLD:AAA5306
Cricotopus sp. 21ES|[14492]|JWDCE587-10|BOLD:AAA5306
Cricotopus sp. 21ES|[14493]|JWDCF181-10|BOLD:AAA5306
Cricotopus sp. 21ES|[14494]|JWDCE586-10|BOLD:AAA5306
Cricotopus sp. 21ES|[14495]|BSCHR156-10|BOLD:AAA5306
Cricotopus sp. 21ES|[14496]|CHURC693-09|BOLD:AAA5306
Cricotopus sp. 21ES|[14497]|CHURC204-07|BOLD:AAA5306
Cricotopus sp. 21ES|[14498]|CHURC234-07|BOLD:AAA5306
Cricotopus sp. 21ES|[14499]|CHURC1078-09|BOLD:AAA5306
Smittia sp. 16ES|[14500]|CHURC115-07|BOLD:AAB0376
Smittia sp. 16ES|[14501]|JWDCC318-10|BOLD:AAB0375
Smittia sp. 16ES|[14502]|JWDCD385-10|BOLD:AAB0375
Smittia sp. 16ES|[14503]|JWDCJ1521-11|BOLD:AAB0375
Smittia sp. 16ES|[14504]|CHURC1065-09|BOLD:AAB0375
Smittia sp. 16ES|[14505]|CHURC114-07|BOLD:AAB0375
Smittia sp. 16ES|[14506]|CHURC934-09|BOLD:AAB0375
Smittia sp. 16ES|[14507]|CHURC239-07|BOLD:AAB0375
Smittia sp. 16ES|[14508]|CHURC153-07|BOLD:AAB0375
Smittia sp. 16ES|[14509]|CHURC182-07|BOLD:AAB0375
Smittia sp. 16ES|[14510]|CHURC154-07|BOLD:AAB0375
Smittia sp. 16ES|[14511]|CHURC055-06|BOLD:AAB0375
Smittia sp. 16ES|[14512]|CHURC058-06|BOLD:AAB0375
Bryophaenocladius|[14513]|JWDCG272-10|BOLD:AAG5485
Bryophaenocladius|[14514]|CHURC787-09|BOLD:AAG5485
Metriocnemus ursinus|[14515]|CHURC290-07|BOLD:AAA9434
Metriocnemus ursinus|[14516]|CHURC137-07|BOLD:AAA9434
Metriocnemus sp. 1ES|[14517]|JWDCE221-10|BOLD:AAA9429
Metriocnemus sp. 1ES|[14518]|CHURC103-07|BOLD:AAA9429
Metriocnemus sp. 1ES|[14519]|CHURC552-08|BOLD:AAA9429
Metriocnemus sp. 1ES|[14520]|CHURC880-09|BOLD:AAA9429
Metriocnemus sp. 1ES|[14521]|CHURC207-07|BOLD:AAA9429
Metriocnemus sp. 1ES|[14522]|CHURC549-08|BOLD:AAA9429
Metriocnemus sp. 1ES|[14523]|CHURC138-07|BOLD:AAA9429
Metriocnemus sp. 1ES|[14524]|CHURC122-07|BOLD:AAA9429
Metriocnemus sp. 1ES|[14525]|CHURC104-07|BOLD:AAA9429
Metriocnemus sp. 1ES|[14526]|CHURC882-09|BOLD:AAA9429
Metriocnemus sp. 1ES|[14527]|CHURC694-09|BOLD:AAA9429
Metriocnemus sp. 1ES|[14528]|JWDCE222-10|BOLD:AAA9429

Metriocnemus sp. 1ES|[14526]|CHURC882-09|BOLD:AAA9429
Metriocnemus sp. 1ES|[14527]|CHURC694-09|BOLD:AAA9429
Metriocnemus sp. 1ES|[14528]|JWDCE222-10|BOLD:AAA9429
Metriocnemus sp. 1ES|[14529]|JWDCE073-10|BOLD:AAA9429
Metriocnemus sp. 1ES|[14530]|JWDCI538-10|BOLD:AAA9429
Metriocnemus sp. 1ES|[14531]|JWDCE049-10|BOLD:AAA9429
Metriocnemus sp. 1ES|[14532]|CHURC844-09|BOLD:AAA9429
Metriocnemus sp. 1ES|[14533]|CHURC885-09|BOLD:AAA9429
Metriocnemus sp. 1ES|[14534]|CHURC569-09|BOLD:AAA9429
Cricotopus sp. 22ES|[14535]|CHURC412-07|BOLD:AAA5310
Cricotopus sp. 22ES|[14536]|CHURC687-09|BOLD:AAA5310
Cricotopus sp. 22ES|[14537]|CHURC366-07|BOLD:AAA5310
Cricotopus sp. 23ES|[14538]|CHURC972-09|BOLD:AAI6022
Cricotopus sp. 23ES|[14539]|CHURC959-09|BOLD:AAI6022
Cricotopus sp. 23ES|[14540]|CHURC1017-09|BOLD:AAI6022
Cricotopus sp. 23ES|[14541]|CHURC1121-09|BOLD:AAI6022
Cricotopus|[14542]|JWDCH932-10|BOLD:AAP9005
Cricotopus|[14543]|JWDCI091-10|BOLD:AAP9005
Cricotopus sp. 25ES|[14544]|JWDCF667-10|BOLD:AAI6017
Cricotopus sp. 25ES|[14545]|CHURC180-07|BOLD:AAI6017
Cricotopus coronatus|[14546]|SWSWP032-09|BOLD:AAI6023
Cricotopus coronatus|[14547]|SWSWP053-09|BOLD:AAI6023
Cricotopus coronatus|[14548]|SWSWP080-09|BOLD:AAI6023
Cricotopus coronatus|[14549]|JWDCF668-10|BOLD:AAI6023
Cricotopus coronatus|[14550]|JWDCF408-10|BOLD:AAI6023
Cricotopus coronatus|[14551]|JWDCF264-10|BOLD:AAI6023
Cricotopus coronatus|[14552]|CHURC970-09|BOLD:AAI6023
Cricotopus coronatus|[14553]|CHURC1129-09|BOLD:AAI6023
Parakiefferiella bathophila|[14554]|CHURC286-07|BOLD:AAE6765
Parakiefferiella bathophila|[14555]|CHURC417-07|BOLD:AAE6766
Parakiefferiella bathophila|[14556]|CHURC995-09|BOLD:AAE6766
Parakiefferiella bathophila|[14557]|CHURC986-09|BOLD:AAE6766
Parakiefferiella bathophila|[14558]|CHURC958-09|BOLD:AAE6766
Parakiefferiella bathophila|[14559]|CHURC945-09|BOLD:AAE6766
Parakiefferiella bathophila|[14560]|CHURC398-07|BOLD:AAE6766
Parakiefferiella bathophila|[14561]|CHURC387-07|BOLD:AAE6766
Parakiefferiella bathophila|[14562]|CHURC1122-09|BOLD:AAE6766
Parakiefferiella bathophila|[14563]|CHURC1001-09|BOLD:AAE6766
Parakiefferiella bathophila|[14564]|CHURC1133-09|BOLD:AAE6766
Orthocladius|[14565]|JWDCE416-10|BOLD:AAP5146
Psectrocladius platypus|[14566]|KSCLC025-09|BOLD:AAG5499
Psectrocladius platypus|[14567]|KSCLC026-09|BOLD:AAG5499
Psectrocladius sp. TE03|[14568]|KSCLC068-09|BOLD:AAX8198
Psectrocladius sp. TE03|[14569]|KSCLC085-09|BOLD:AAX8198
Psectrocladius sp. TE03|[14570]|CHURC076-06|BOLD:AAX8198
Psectrocladius conjungens|[14571]|CHURC231-07|BOLD:AAF6405
Psectrocladius|[14572]|JWDCC493-10|BOLD:AAL7318
Psectrocladius sordidellus|[14573]|JWDCD268-10|BOLD:AAL7382
Psectrocladius|[14574]|JWDCC498-10|BOLD:AAL7319
Psectrocladius fennicus|[14575]|JWDCJ153-11|BOLD:AAD4712
Psectrocladius fennicus|[14576]|CHURC372-07|BOLD:AAD4712
Psectrocladius fennicus|[14577]|JWDCI581-10|BOLD:AAD4712
Psectrocladius fennicus|[14578]|JWDCF784-10|BOLD:AAD4712
Psectrocladius fennicus|[14579]|CHURC944-09|BOLD:AAD4712
Psectrocladius fennicus|[14580]|CHURC471-08|BOLD:AAD4712
Psectrocladius fennicus|[14581]|CHURC373-07|BOLD:AAD4712
Psectrocladius fennicus|[14582]|CHURC582-09|BOLD:AAD4712
Psectrocladius fennicus|[14583]|MHMOS103-07|BOLD:ADR7450
Psectrocladius fennicus|[14584]|KSCLC018-09|BOLD:AAD4713
Psectrocladius fennicus|[14585]|KSCLC069-09|BOLD:AAD4713
Psectrocladius fennicus|[14586]|KSCLC058-09|BOLD:AAD4713
Psectrocladius fennicus|[14587]|KSCLC076-09|BOLD:AAD4713
Psectrocladius fennicus|[14588]|KSCLC092-09|BOLD:AAD4713
Psectrocladius fennicus|[14589]|KSCLC012-09|BOLD:AAD4713
Psectrocladius fennicus|[14590]|KSCLC033-09|BOLD:AAD4713
Psectrocladius fennicus|[14591]|KSCLC055-09|BOLD:AAD4713
Psectrocladius fennicus|[14592]|CHURC430-07|BOLD:AAD4713
Orthocladius|[14593]|JWDCE791-10|BOLD:AAP5148
Cricotopus|[14594]|JWDCF636-10|BOLD:AAQ0619
Cricotopus|[14595]|JWDCF669-10|BOLD:AAQ0619
Cricotopus|[14596]|CHURC851-09|BOLD:ACH4204
Cricotopus pilosellus|[14597]|CHURC989-09|BOLD:AAA5315
Cricotopus pilosellus|[14598]|CHURC1021-09|BOLD:AAA5315
Cricotopus tibialis|[14599]|CHURC776-09|BOLD:AAA5300
Cricotopus tibialis|[14600]|CHURC202-07|BOLD:AAA5300
Cricotopus tibialis|[14601]|CHURC203-07|BOLD:AAA5300
Cricotopus tibialis|[14602]|CHURC805-09|BOLD:AAA5300
Cricotopus tibialis|[14603]|CHURC019-06|BOLD:AAA5300
Cricotopus tibialis|[14604]|CHURC016-06|BOLD:AAA5300
Cricotopus tibialis|[14605]|CHURC774-09|BOLD:AAA5300
Cricotopus tibialis|[14606]|CHURC797-09|BOLD:AAA5300
Cricotopus tibialis|[14607]|CHURC163-07|BOLD:AAA5300
Cricotopus tibialis|[14608]|CHURC777-09|BOLD:AAA5300
Cricotopus tibialis|[14609]|CHURC230-07|BOLD:AAA5300
Cricotopus tibialis|[14610]|CHURC121-07|BOLD:AAA5300
Cricotopus tibialis|[14611]|CHURC717-09|BOLD:AAA5300
Cricotopus tibialis|[14612]|JWDCF621-10|BOLD:AAA5300
Cricotopus tibialis|[14613]|JWDCH705-10|BOLD:AAA5300
Cricotopus tibialis|[14614]|CHURC802-09|BOLD:AAA5300
Cricotopus tibialis|[14615]|CHURC427-07|BOLD:AAA5300
Cricotopus tibialis|[14616]|CHURC266-07|BOLD:AAA5300
Cricotopus tibialis|[14617]|JWDCC479-10|BOLD:AAA5300
Cricotopus tibialis|[14618]|JWDCG368-10|BOLD:AAA5300
Cricotopus tibialis|[14619]|CHURC855-09|BOLD:AAA5300
Cricotopus tibialis|[14620]|CHURC030-06|BOLD:AAA5300
Cricotopus tibialis|[14621]|CHURC018-06|BOLD:AAA5300
Psectrocladius sordidellus|[14622]|MMADL058-09|BOLD:AAF8903
Psectrocladius sordidellus|[14623]|KSCLC093-09|BOLD:AAF8903
Psectrocladius sordidellus|[14624]|CHURC431-07|BOLD:AAF8903
Psectrocladius cf. limbatellus|[14625]|KSCLC050-09|BOLD:AAG5500
Psectrocladius cf. limbatellus|[14626]|KSCLC064-09|BOLD:AAG5500
Psectrocladius cf. limbatellus|[14627]|KSCLC057-09|BOLD:AAG5500
Psectrocladius cf. limbatellus|[14628]|JWDCH974-10|BOLD:AAG5500

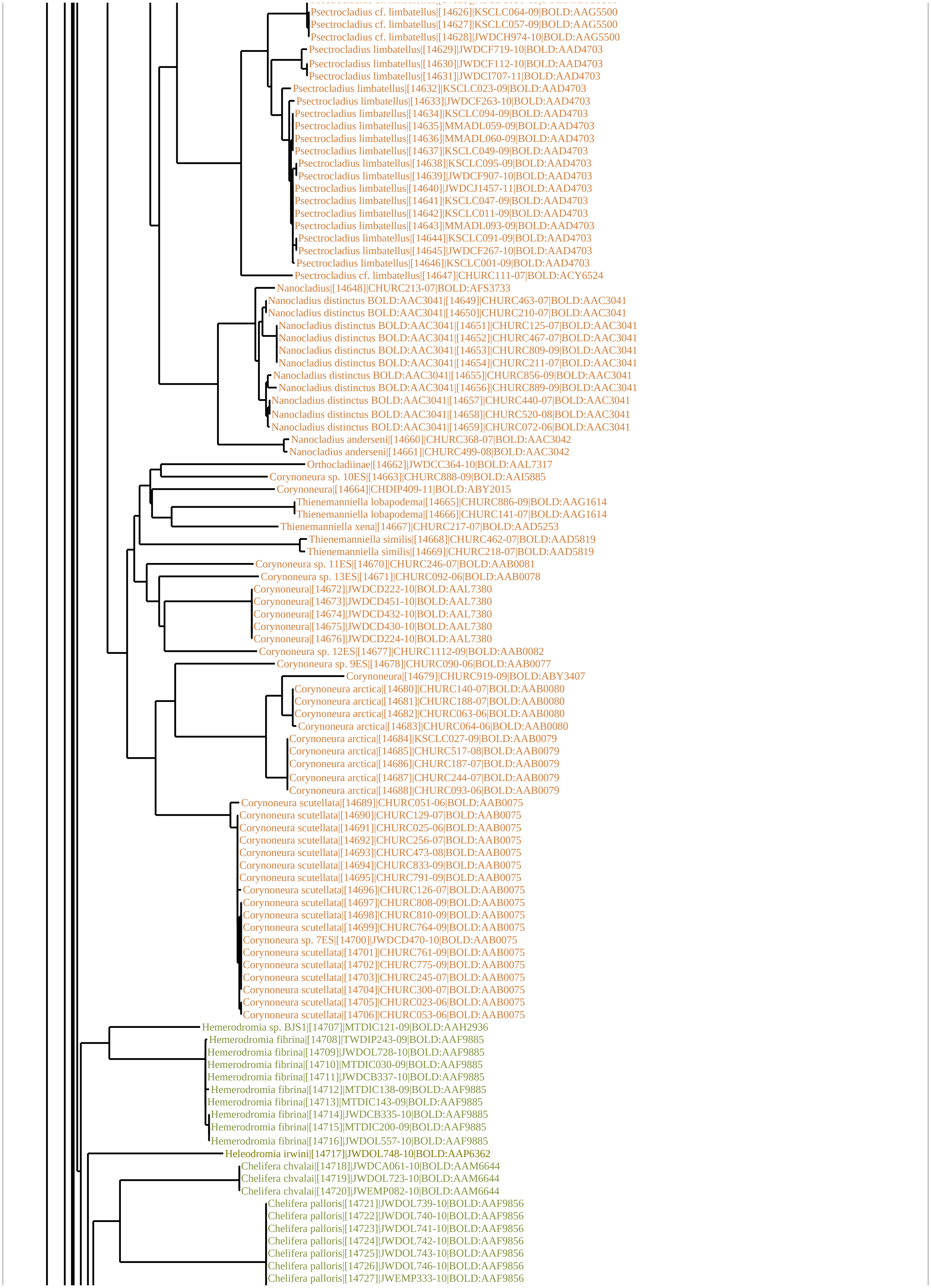
Psectrocladius cf. limbatellus|[14626]|KSCLC064-09|BOLD:AAG5500
Psectrocladius cf. limbatellus|[14627]|KSCLC057-09|BOLD:AAG5500
Psectrocladius cf. limbatellus|[14628]|JWDCH974-10|BOLD:AAG5500
Psectrocladius limbatellus|[14629]|JWDCF719-10|BOLD:AAD4703
Psectrocladius limbatellus|[14630]|JWDCF112-10|BOLD:AAD4703
Psectrocladius limbatellus|[14631]|JWDCI707-11|BOLD:AAD4703
Psectrocladius limbatellus|[14632]|KSCLC023-09|BOLD:AAD4703
Psectrocladius limbatellus|[14633]|JWDCF263-10|BOLD:AAD4703
Psectrocladius limbatellus|[14634]|KSCLC094-09|BOLD:AAD4703
Psectrocladius limbatellus|[14635]|MMADL059-09|BOLD:AAD4703
Psectrocladius limbatellus|[14636]|MMADL060-09|BOLD:AAD4703
Psectrocladius limbatellus|[14637]|KSCLC049-09|BOLD:AAD4703
Psectrocladius limbatellus|[14638]|KSCLC095-09|BOLD:AAD4703
Psectrocladius limbatellus|[14639]|JWDCF907-10|BOLD:AAD4703
Psectrocladius limbatellus|[14640]|JWDCJ1457-11|BOLD:AAD4703
Psectrocladius limbatellus|[14641]|KSCLC047-09|BOLD:AAD4703
Psectrocladius limbatellus|[14642]|KSCLC011-09|BOLD:AAD4703
Psectrocladius limbatellus|[14643]|MMADL093-09|BOLD:AAD4703
Psectrocladius limbatellus|[14644]|KSCLC091-09|BOLD:AAD4703
Psectrocladius limbatellus|[14645]|JWDCF267-10|BOLD:AAD4703
Psectrocladius limbatellus|[14646]|KSCLC001-09|BOLD:AAD4703
Psectrocladius cf. limbatellus|[14647]|CHURC111-07|BOLD:ACY6524
Nanocladius|[14648]|CHURC213-07|BOLD:AFS3733
Nanocladius distinctus BOLD:AAC3041|[14649]|CHURC463-07|BOLD:AAC3041
Nanocladius distinctus BOLD:AAC3041|[14650]|CHURC210-07|BOLD:AAC3041
Nanocladius distinctus BOLD:AAC3041|[14651]|CHURC125-07|BOLD:AAC3041
Nanocladius distinctus BOLD:AAC3041|[14652]|CHURC467-07|BOLD:AAC3041
Nanocladius distinctus BOLD:AAC3041|[14653]|CHURC809-09|BOLD:AAC3041
Nanocladius distinctus BOLD:AAC3041|[14654]|CHURC211-07|BOLD:AAC3041
Nanocladius distinctus BOLD:AAC3041|[14655]|CHURC856-09|BOLD:AAC3041
Nanocladius distinctus BOLD:AAC3041|[14656]|CHURC889-09|BOLD:AAC3041
Nanocladius distinctus BOLD:AAC3041|[14657]|CHURC440-07|BOLD:AAC3041
Nanocladius distinctus BOLD:AAC3041|[14658]|CHURC520-08|BOLD:AAC3041
Nanocladius distinctus BOLD:AAC3041|[14659]|CHURC072-06|BOLD:AAC3041
Nanocladius anderseni|[14660]|CHURC368-07|BOLD:AAC3042
Nanocladius anderseni|[14661]|CHURC499-08|BOLD:AAC3042
Orthocladiinae|[14662]|JWDCC364-10|BOLD:AAL7317
Corynoneura sp. 10ES|[14663]|CHURC888-09|BOLD:AAI5885
Corynoneura|[14664]|CHDIP409-11|BOLD:ABY2015
Thienemanniella lobapodema|[14665]|CHURC886-09|BOLD:AAG1614
Thienemanniella lobapodema|[14666]|CHURC141-07|BOLD:AAG1614
Thienemanniella xena|[14667]|CHURC217-07|BOLD:AAD5253
Thienemanniella similis|[14668]|CHURC462-07|BOLD:AAD5819
Thienemanniella similis|[14669]|CHURC218-07|BOLD:AAD5819
Corynoneura sp. 11ES|[14670]|CHURC246-07|BOLD:AAB0081
Corynoneura sp. 13ES|[14671]|CHURC092-06|BOLD:AAB0078
Corynoneura|[14672]|JWDCD222-10|BOLD:AAL7380
Corynoneura|[14673]|JWDCD451-10|BOLD:AAL7380
Corynoneura|[14674]|JWDCD432-10|BOLD:AAL7380
Corynoneura|[14675]|JWDCD430-10|BOLD:AAL7380
Corynoneura|[14676]|JWDCD224-10|BOLD:AAL7380
Corynoneura sp. 12ES|[14677]|CHURC1112-09|BOLD:AAB0082
Corynoneura sp. 9ES|[14678]|CHURC090-06|BOLD:AAB0077
Corynoneura|[14679]|CHURC919-09|BOLD:ABY3407
Corynoneura arctica|[14680]|CHURC140-07|BOLD:AAB0080
Corynoneura arctica|[14681]|CHURC188-07|BOLD:AAB0080
Corynoneura arctica|[14682]|CHURC063-06|BOLD:AAB0080
Corynoneura arctica|[14683]|CHURC064-06|BOLD:AAB0080
Corynoneura arctica|[14684]|KSCLC027-09|BOLD:AAB0079
Corynoneura arctica|[14685]|CHURC517-08|BOLD:AAB0079
Corynoneura arctica|[14686]|CHURC187-07|BOLD:AAB0079
Corynoneura arctica|[14687]|CHURC244-07|BOLD:AAB0079
Corynoneura arctica|[14688]|CHURC093-06|BOLD:AAB0079
Corynoneura scutellata|[14689]|CHURC051-06|BOLD:AAB0075
Corynoneura scutellata|[14690]|CHURC129-07|BOLD:AAB0075
Corynoneura scutellata|[14691]|CHURC025-06|BOLD:AAB0075
Corynoneura scutellata|[14692]|CHURC256-07|BOLD:AAB0075
Corynoneura scutellata|[14693]|CHURC473-08|BOLD:AAB0075
Corynoneura scutellata|[14694]|CHURC833-09|BOLD:AAB0075
Corynoneura scutellata|[14695]|CHURC791-09|BOLD:AAB0075
Corynoneura scutellata|[14696]|CHURC126-07|BOLD:AAB0075
Corynoneura scutellata|[14697]|CHURC808-09|BOLD:AAB0075
Corynoneura scutellata|[14698]|CHURC810-09|BOLD:AAB0075
Corynoneura scutellata|[14699]|CHURC764-09|BOLD:AAB0075
Corynoneura sp. 7ES|[14700]|JWDCD470-10|BOLD:AAB0075
Corynoneura scutellata|[14701]|CHURC761-09|BOLD:AAB0075
Corynoneura scutellata|[14702]|CHURC775-09|BOLD:AAB0075
Corynoneura scutellata|[14703]|CHURC245-07|BOLD:AAB0075
Corynoneura scutellata|[14704]|CHURC300-07|BOLD:AAB0075
Corynoneura scutellata|[14705]|CHURC023-06|BOLD:AAB0075
Corynoneura scutellata|[14706]|CHURC053-06|BOLD:AAB0075
Hemerodromia sp. BJS1|[14707]|MTDIC121-09|BOLD:AAH2936
Hemerodromia fibrina|[14708]|TWDIP243-09|BOLD:AAF9885
Hemerodromia fibrina|[14709]|JWDOL728-10|BOLD:AAF9885
Hemerodromia fibrina|[14710]|MTDIC030-09|BOLD:AAF9885
Hemerodromia fibrina|[14711]|JWDCB337-10|BOLD:AAF9885
Hemerodromia fibrina|[14712]|MTDIC138-09|BOLD:AAF9885
Hemerodromia fibrina|[14713]|MTDIC143-09|BOLD:AAF9885
Hemerodromia fibrina|[14714]|JWDCB335-10|BOLD:AAF9885
Hemerodromia fibrina|[14715]|MTDIC200-09|BOLD:AAF9885
Hemerodromia fibrina|[14716]|JWDOL557-10|BOLD:AAF9885
Heleodromia irwini|[14717]|JWDOL748-10|BOLD:AAP6362
Chelifera chvalai|[14718]|JWDCA061-10|BOLD:AAM6644
Chelifera chvalai|[14719]|JWDOL723-10|BOLD:AAM6644
Chelifera chvalai|[14720]|JWEMP082-10|BOLD:AAM6644
Chelifera palloris|[14721]|JWDOL739-10|BOLD:AAF9856
Chelifera palloris|[14722]|JWDOL740-10|BOLD:AAF9856
Chelifera palloris|[14723]|JWDOL741-10|BOLD:AAF9856
Chelifera palloris|[14724]|JWDOL742-10|BOLD:AAF9856
Chelifera palloris|[14725]|JWDOL743-10|BOLD:AAF9856
Chelifera palloris|[14726]|JWDOL746-10|BOLD:AAF9856
Chelifera palloris|[14727]|JWEMP333-10|BOLD:AAF9856

Chelifera palloris|[14725]|JWDOL743-10|BOLD:AAF9856
Chelifera palloris|[14726]|JWDOL746-10|BOLD:AAF9856
Chelifera palloris|[14727]|JWEMP333-10|BOLD:AAF9856
Chelifera palloris|[14728]|JWEMP336-10|BOLD:AAF9856
Rhamphomyia virgata grp. sp. BJS2|[14729]|TWDIP011-09|BOLD:AAF9795
Rhamphomyia virgata grp. sp. BJS2|[14730]|TWDIP013-09
Rhamphomyia virgata grp. sp. BJS2|[14731]|TWDIP010-09|BOLD:AAF9794
Rhamphomyia virgata grp. sp. BJS2|[14732]|JWDCA058-10|BOLD:AAF9794
Rhamphomyia virgata grp. sp. BJS2|[14733]|JWEMP282-10|BOLD:AAF9794
Rhamphomyia sp. BJS6|[14734]|JWDCE511-10|BOLD:AAP3640
Rhamphomyia sp. BJS1|[14735]|JWDCD031-10|BOLD:AAL8967
Rhamphomyia sp. BJS1|[14736]|JWDCD029-10|BOLD:AAL8967
Rhamphomyia sp. BJS1|[14737]|JWDCD028-10|BOLD:AAL8967
Rhamphomyia sp. BJS1|[14738]|JWDCD080-10|BOLD:AAL8967
Rhamphomyia sp. BJS1|[14739]|JWDCD032-10|BOLD:AAL8967
Rhamphomyia sp. BJS1|[14740]|JWDCD084-10|BOLD:AAL8967
Rhamphomyia sp. BJS1|[14741]|JWDCD026-10|BOLD:AAL8967
Rhamphomyia sp. BJS1|[14742]|JWDCD033-10|BOLD:AAL8967
Rhamphomyia sp. BJS1|[14743]|JWDCD077-10|BOLD:AAL8967
Rhamphomyia sp. BJS1|[14744]|JWDCD083-10|BOLD:AAL8967
Rhamphomyia sp. BJS1|[14745]|JWDCD027-10|BOLD:AAL8967
Rhamphomyia sp. BJS1|[14746]|JWEMP094-10|BOLD:AAL8967
Rhamphomyia sp. BJS1|[14747]|JWEMP305-10|BOLD:AAL8967
Rhamphomyia sp. BJS1|[14748]|JWEMP042-10|BOLD:AAL8967
Rhamphomyia sp. BJS1|[14749]|JWEMP003-10|BOLD:AAL8967
Rhamphomyia sp. BJS1|[14750]|JWDCE509-10|BOLD:AAL8967
Rhamphomyia sp. BJS1|[14751]|JWDCD030-10|BOLD:AAL8967
Rhamphomyia sp. BJS1|[14752]|JWDCD079-10|BOLD:AAL8967
Rhamphomyia sp. BJS1|[14753]|JWDCD078-10|BOLD:AAL8967
Rhamphomyia sp. BJS1|[14754]|JWDCD048-10|BOLD:AAL8967
Rhamphomyia sp. BJS1|[14755]|JWEMP309-10|BOLD:AAL8967
Rhamphomyia sp. BJS1|[14756]|JWEMP223-10|BOLD:AAL8967
Rhamphomyia sp. BJS1|[14757]|JWEMP310-10|BOLD:AAL8967
Rhamphomyia sp. BJS1|[14758]|JWEMP043-10|BOLD:AAL8967
Rhamphomyia sp. BJS1|[14759]|JWEMP103-10|BOLD:AAL8967
Rhamphomyia sp. BJS1|[14760]|JWEMP311-10|BOLD:AAL8967
Rhamphomyia sp. BJS1|[14761]|JWEMP312-10|BOLD:AAL8967
Rhamphomyia sp. BJS1|[14762]|JWEMP307-10|BOLD:AAQ0828
Rhamphomyia nigrita|[14763]|JWDCA042-10|BOLD:AAF9804
Rhamphomyia nigrita|[14764]|JWDCA043-10|BOLD:AAF9804
Rhamphomyia nigrita|[14765]|JWDCC231-10|BOLD:AAF9804
Rhamphomyia nigrita|[14766]|MTDIC066-09|BOLD:AAF9804
Rhamphomyia nigrita|[14767]|JWDCA045-10|BOLD:AAF9804
Rhamphomyia nigrita|[14768]|JWDCD049-10|BOLD:AAF9804
Rhamphomyia nigrita|[14769]|JWDCE561-10|BOLD:AAF9804
Rhamphomyia nigrita|[14770]|JWDCC020-10|BOLD:AAF9804
Rhamphomyia nigrita|[14771]|MTDIC115-09|BOLD:AAF9804
Rhamphomyia nigrita|[14772]|JWDCD037-10|BOLD:AAF9804
Rhamphomyia nigrita|[14773]|JWDCD036-10|BOLD:AAF9804
Rhamphomyia nigrita|[14774]|JWEMP314-10|BOLD:AAF9804
Rhamphomyia diversipennis|[14775]|JWDOL510-10|BOLD:AAP1853
Rhamphomyia diversipennis|[14776]|JWDOL604-10|BOLD:AAP1853
Rhamphomyia diversipennis|[14777]|JWEMP151-10|BOLD:AAP1853
Rhamphomyia diversipennis|[14778]|JWEMP152-10|BOLD:AAP1853
Rhamphomyia diversipennis|[14779]|JWEMP154-10|BOLD:AAP1853
Rhamphomyia diversipennis|[14780]|JWDOL511-10|BOLD:AAP1853
Rhamphomyia diversipennis|[14781]|JWDOL514-10|BOLD:AAP1853
Rhamphomyia diversipennis|[14782]|JWDOL530-10|BOLD:AAP1853
Rhamphomyia valga|[14783]|JWEMP241-10|BOLD:AAF9757
Rhamphomyia poplitea|[14784]|JWEMP270-10|BOLD:AAF9757
Rhamphomyia poplitea|[14785]|JWEMP238-10|BOLD:AAF9757
Rhamphomyia poplitea|[14786]|JWEMP269-10|BOLD:AAF9757
Rhamphomyia poplitea|[14787]|JWEMP268-10|BOLD:AAF9757
Rhamphomyia poplitea|[14788]|JWDOL675-10|BOLD:AAF9757
Rhamphomyia pusio grp. sp. BJS10|[14789]|JWEMP007-10|BOLD:AAF9883
Rhamphomyia pusio grp. sp. BJS10|[14790]|JWEMP237-10|BOLD:AAF9883
Rhamphomyia pusio grp. sp. BJS10|[14791]|JWEMP240-10|BOLD:AAF9883
Rhamphomyia pusio grp. sp. BJS10|[14792]|JWDCB325-10|BOLD:AAF9883
Rhamphomyia pusio grp. sp. BJS10|[14793]|JWDOL664-10|BOLD:AAF9883
Rhamphomyia phemius|[14794]|JWDCE338-10|BOLD:AAF9879
Rhamphomyia phemius|[14795]|JWDCE617-10|BOLD:AAF9879
Rhamphomyia phemius|[14796]|JWDCE149-10|BOLD:AAF9879
Rhamphomyia phemius|[14797]|JWDCE563-10|BOLD:AAF9879
Rhamphomyia phemius|[14798]|JWDCE356-10|BOLD:AAF9879
Rhamphomyia phemius|[14799]|JWDCD082-10|BOLD:AAF9879
Rhamphomyia phemius|[14800]|JWDCE801-10|BOLD:AAF9879
Rhamphomyia phemius|[14801]|JWDCE435-10|BOLD:AAF9879
Rhamphomyia phemius|[14802]|JWDCE357-10|BOLD:AAF9879
Rhamphomyia phemius|[14803]|JWDCD081-10|BOLD:AAF9879
Rhamphomyia phemius|[14804]|JWDCA099-10|BOLD:AAF9879
Rhamphomyia phemius|[14805]|JWDCD087-10|BOLD:AAF9879
Rhamphomyia phemius|[14806]|JWDCE802-10|BOLD:AAF9879
Rhamphomyia phemius|[14807]|JWDCE803-10|BOLD:AAF9879
Rhamphomyia phemius|[14808]|JWEMP306-10|BOLD:AAF9879
Rhamphomyia phemius|[14809]|JWEMP313-10|BOLD:AAF9879
Rhamphomyia lapponica grp. sp. BJS2|[14810]|MTDIC119-09|BOLD:AAF9805
Rhamphomyia nr. anaxo grp. sp. BJS4|[14811]|TWDIP017-09|BOLD:AAF9796
Rhamphomyia nr. anaxo grp. sp. BJS4|[14812]|TWDIP015-09|BOLD:AAF9796
Rhamphomyia nr. anaxo grp. sp. BJS4|[14813]|JWEMP053-10|BOLD:AAF9796
Rhamphomyia anaxo grp. sp. BJS4|[14814]|JWEMP224-10|BOLD:ABZ8252
Rhamphomyia anaxo grp. sp. BJS4|[14815]|TWDIP016-09|BOLD:AAF9797
Rhamphomyia anaxo grp. sp. BJS4|[14816]|TWDIP018-09|BOLD:AAF9797
Rhamphomyia anaxo grp. sp. BJS4|[14817]|JWDOL595-10|BOLD:AAF9797
Rhamphomyia anaxo grp. sp. BJS4|[14818]|JWDOL529-10|BOLD:AAF9797
Rhamphomyia anaxo grp. sp. BJS4|[14819]|JWDOL554-10|BOLD:AAF9797
Rhamphomyia anaxo grp. sp. BJS4|[14820]|JWEMP331-10|BOLD:AAP3643
Rhamphomyia anaxo grp. sp. BJS4|[14821]|JWEMP196-10|BOLD:AAP3643
Rhamphomyia anaxo grp. sp. BJS4|[14822]|JWDOL527-10|BOLD:AAP3643
Rhamphomyia anaxo grp. sp. BJS4|[14823]|JWDOL670-10|BOLD:AAP3643
Rhamphomyia fuscula grp. sp. BJS2|[14824]|JWEMP105-10|BOLD:AAP1850
Rhamphomyia fuscula grp. sp. BJS2|[14825]|JWEMP221-10|BOLD:AAP1850
Rhamphomyia fuscula grp. sp. BJS2|[14826]|JWEMP303-10|BOLD:AAP1850
Rhamphomyia fuscula grp. sp. BJS2|[14827]|JWDOL484-10|BOLD:AAP1850

[illegible]
Rhamphomyia fuscula grp. sp. BJS2|[14825]|JWEMP221-10|BOLD:AAP1850
Rhamphomyia fuscula grp. sp. BJS2|[14826]|JWEMP303-10|BOLD:AAP1850
Rhamphomyia fuscula grp. sp. BJS2|[14827]|JWDOL484-10|BOLD:AAP1850
Rhamphomyia fuscula grp. sp. BJS2|[14828]|JWDOL483-10|BOLD:AAP1850
Rhamphomyia fuscula grp. sp. BJS2|[14829]|JWDOL533-10|BOLD:AAP1850
Rhamphomyia fuscula grp. sp. BJS2|[14830]|JWDOL590-10|BOLD:AAP1850
Rhamphomyia fuscula grp. sp. BJS2|[14831]|JWDCE510-10|BOLD:AAP1850
Rhamphomyia fuscula grp. sp. BJS2|[14832]|JWEMP052-10|BOLD:AAP1850
Rhamphomyia fuscula grp. sp. BJS2|[14833]|JWEMP054-10|BOLD:AAP1850
Rhamphomyia fuscula grp. sp. BJS2|[14834]|JWEMP104-10|BOLD:AAP1850
Rhamphomyia fuscula grp. sp. BJS2|[14835]|JWEMP106-10|BOLD:AAP1850
Rhamphomyia fuscula grp. sp. BJS2|[14836]|JWEMP115-10|BOLD:AAP1850
Rhamphomyia fuscula grp. sp. BJS2|[14837]|JWEMP189-10|BOLD:AAP1850
Rhamphomyia fuscula grp. sp. BJS2|[14838]|JWEMP201-10|BOLD:AAP1850
Rhamphomyia fuscula grp. sp. BJS2|[14839]|JWEMP202-10|BOLD:AAP1850
Rhamphomyia fuscula grp. sp. BJS2|[14840]|JWEMP204-10|BOLD:AAP1850
Rhamphomyia fuscula grp. sp. BJS2|[14841]|JWEMP205-10|BOLD:AAP1850
Rhamphomyia fuscula grp. sp. BJS2|[14842]|JWEMP216-10|BOLD:AAP1850
Rhamphomyia fuscula grp. sp. BJS2|[14843]|JWEMP217-10|BOLD:AAP1850
Rhamphomyia fuscula grp. sp. BJS2|[14844]|JWEMP218-10|BOLD:AAP1850
Rhamphomyia fuscula grp. sp. BJS2|[14845]|JWEMP219-10|BOLD:AAP1850
Rhamphomyia fuscula grp. sp. BJS2|[14846]|JWEMP225-10|BOLD:AAP1850
Rhamphomyia fuscula grp. sp. BJS2|[14847]|JWDOL502-10|BOLD:AAP1850
Rhamphomyia fuscula grp. sp. BJS2|[14848]|JWDOL512-10|BOLD:AAP1850
Rhamphomyia fuscula grp. sp. BJS2|[14849]|JWDOL532-10|BOLD:AAP1850
Rhamphomyia fuscula grp. sp. BJS2|[14850]|JWDOL564-10|BOLD:AAP1850
Rhamphomyia fuscula grp. sp. BJS2|[14851]|JWDOL682-10|BOLD:AAP1850
Rhamphomyia fuscula grp. sp. BJS2|[14852]|JWEMP051-10|BOLD:AAP1850
Rhamphomyia fuscula grp. sp. BJS2|[14853]|JWEMP107-10|BOLD:AAP1850
Rhamphomyia fuscula grp. sp. BJS2|[14854]|JWEMP108-10|BOLD:AAP1850
Rhamphomyia fuscula grp. sp. BJS2|[14855]|JWEMP226-10|BOLD:AAP1850
Rhamphomyia fuscula grp. sp. BJS2|[14856]|JWDOL684-10|BOLD:AAP1850
Rhamphomyia fuscula grp. sp. BJS2|[14857]|JWDOL513-10|BOLD:AAP1850
Rhamphomyia fuscula grp. sp. BJS2|[14858]|JWDOL626-10|BOLD:AAP1850
Rhamphomyia fuscula grp. sp. BJS2|[14859]|JWDOL635-10|BOLD:AAP1850
Rhamphomyia fuscula grp. sp. BJS2|[14860]|JWEMP116-10|BOLD:AAP1850
Rhamphomyia fuscula grp. sp. BJS2|[14861]|JWEMP117-10|BOLD:AAP1850
Rhamphomyia fuscula grp. sp. BJS2|[14862]|JWEMP118-10|BOLD:AAP1850
Rhamphomyia fuscula grp. sp. BJS2|[14863]|JWEMP187-10|BOLD:AAP1850
Rhamphomyia fuscula grp. sp. BJS2|[14864]|JWEMP188-10|BOLD:AAP1850
Rhamphomyia fuscula grp. sp. BJS2|[14865]|JWEMP190-10|BOLD:AAP1850
Rhamphomyia fuscula grp. sp. BJS2|[14866]|JWEMP203-10|BOLD:AAP1850
Rhamphomyia fuscula grp. sp. BJS2|[14867]|JWEMP215-10|BOLD:AAP1850
Rhamphomyia fuscula grp. sp. BJS2|[14868]|JWEMP222-10|BOLD:AAP1850
Rhamphomyia fuscula grp. sp. BJS2|[14869]|JWDOL498-10|BOLD:AAP1850
Rhamphomyia fuscula grp. sp. BJS2|[14870]|JWDOL501-10|BOLD:AAP1850
Rhamphomyia fuscula grp. sp. BJS2|[14871]|JWDOL569-10|BOLD:AAP1850
Rhamphomyia fuscula grp. sp. BJS2|[14872]|JWDOL655-10|BOLD:AAP1850
Rhamphomyia tipularia grp. sp. BJS1|[14873]|JWDCA108-10|BOLD:AAF9881
Rhamphomyia tipularia grp. sp. BJS1|[14874]|JWDCE437-10|BOLD:AAF9881
Rhamphomyia tipularia grp. sp. BJS1|[14875]|JWDOL671-10|BOLD:AAF9881
Rhamphomyia tipularia grp. sp. BJS1|[14876]|JWDOL481-10|BOLD:AAF9881
Rhamphomyia tipularia grp. sp. BJS1|[14877]|JWDCC139-10|BOLD:AAF9881
Rhamphomyia tipularia grp. sp. BJS1|[14878]|JWDOL480-10|BOLD:AAF9881
Rhamphomyia tipularia grp. sp. BJS1|[14879]|JWDOL540-10|BOLD:AAF9881
Rhamphomyia tipularia grp. sp. BJS1|[14880]|JWDOL710-10|BOLD:AAF9881
Rhamphomyia nr. depilis grp. sp. BJS8|[14881]|JWDOL542-10|BOLD:AAF9813
Rhamphomyia nr. depilis grp. sp. BJS8|[14882]|JWDOL479-10|BOLD:AAF9813
Rhamphomyia nr. depilis grp. sp. BJS8|[14883]|JWDOL627-10|BOLD:AAF9813
Rhamphomyia nr. depilis grp. sp. BJS8|[14884]|JWDOL543-10|BOLD:AAF9813
Rhamphomyia nr. depilis grp. sp. BJS8|[14885]|JWDCA109-10|BOLD:AAF9813
Rhamphomyia nr. depilis grp. sp. BJS8|[14886]|TWDIP313-09|BOLD:AAF9813
Rhamphomyia nr. depilis grp. sp. BJS8|[14887]|JWDOL672-10|BOLD:AAF9813
Rhamphomyia sp. BJS2|[14888]|JWDCA112-10|BOLD:AAF9882
Rhamphomyia sp. BJS2|[14889]|JWDCA113-10|BOLD:AAF9882
Rhamphomyia sp. BJS2|[14890]|JWDCA116-10|BOLD:AAF9882
Rhamphomyia sp. BJS2|[14891]|JWDCA117-10|BOLD:AAF9882
Rhamphomyia sp. BJS2|[14892]|JWDCA119-10|BOLD:AAF9882
Rhamphomyia debilis grp. sp. BJS7|[14893]|JWDCC728-10|BOLD:AAF9800
Rhamphomyia debilis grp. sp. BJS7|[14894]|JWDOL657-10|BOLD:AAF9800
Rhamphomyia debilis grp. sp. BJS7|[14895]|JWDCC832-10|BOLD:AAF9800
Rhamphomyia debilis grp. sp. BJS7|[14896]|JWDCC734-10|BOLD:AAF9800
Rhamphomyia debilis grp. sp. BJS7|[14897]|JWDCC732-10|BOLD:AAF9800
Rhamphomyia debilis grp. sp. BJS7|[14898]|JWEMP083-10|BOLD:AAF9800
Rhamphomyia debilis grp. sp. BJS7|[14899]|JWEMP175-10|BOLD:AAF9800
Rhamphomyia debilis grp. sp. BJS7|[14900]|JWEMP177-10|BOLD:AAF9800
Rhamphomyia debilis grp. sp. BJS7|[14901]|JWEMP181-10|BOLD:AAF9800
Rhamphomyia debilis grp. sp. BJS7|[14902]|JWEMP213-10|BOLD:AAF9800
Rhamphomyia debilis grp. sp. BJS7|[14903]|JWDOL494-10|BOLD:AAF9800
Rhamphomyia debilis grp. sp. BJS7|[14904]|JWEMP178-10|BOLD:AAF9800
Rhamphomyia debilis grp. sp. BJS7|[14905]|JWDOL497-10|BOLD:AAF9800
Rhamphomyia debilis grp. sp. BJS7|[14906]|JWDOL724-10|BOLD:AAF9800
Rhamphomyia debilis grp. sp. BJS7|[14907]|JWDOL734-10|BOLD:AAF9800
Rhamphomyia debilis grp. sp. BJS7|[14908]|JWDOL620-10|BOLD:AAF9800
Rhamphomyia debilis grp. sp. BJS7|[14909]|JWDOL713-10|BOLD:AAF9800
Rhamphomyia debilis grp. sp. BJS7|[14910]|MTDIC003-09|BOLD:AAF9800
Rhamphomyia debilis grp. sp. BJS7|[14911]|JWDCA110-10|BOLD:AAF9800
Rhamphomyia debilis grp. sp. BJS7|[14912]|JWDCA095-10|BOLD:AAF9800
Rhamphomyia debilis grp. sp. BJS7|[14913]|JWDCA046-10|BOLD:AAF9800
Rhamphomyia debilis grp. sp. BJS7|[14914]|MTDIC008-09|BOLD:AAF9800
Rhamphomyia debilis grp. sp. BJS7|[14915]|JWDCC731-10|BOLD:AAF9800
Rhamphomyia debilis grp. sp. BJS7|[14916]|JWDCC730-10|BOLD:AAF9800
Rhamphomyia debilis grp. sp. BJS7|[14917]|JWDCC727-10|BOLD:AAF9800
Rhamphomyia debilis grp. sp. BJS7|[14918]|JWDOL559-10|BOLD:AAF9800
Rhamphomyia debilis grp. sp. BJS7|[14919]|JWDOL560-10|BOLD:AAF9800
Rhamphomyia debilis grp. sp. BJS7|[14920]|JWDOL561-10|BOLD:AAF9800
Rhamphomyia debilis grp. sp. BJS7|[14921]|JWDOL562-10|BOLD:AAF9800
Rhamphomyia debilis grp. sp. BJS7|[14922]|JWDOL563-10|BOLD:AAF9800
Rhamphomyia debilis grp. sp. BJS7|[14923]|JWDOL706-10|BOLD:AAF9800
Rhamphomyia debilis grp. sp. BJS7|[14924]|JWDOL678-10|BOLD:AAF9800
Dolichocephala woodi|[14925]|MTDIC059-09|BOLD:AAI6335
Bibio|[14926]|JWDCH874-10|BOLD:AAP8848

Rhamphomyia debilis grp. sp. BJS7|[14924]|JWDOL678-10|BOLD:AAF9800
Dolichocephala woodi|[14925]|MTDIC059-09|BOLD:AAI6335
Bibio|[14926]|JWDCH874-10|BOLD:AAP8848
Bibio labradorensis|[14927]|TWDIP355-09|BOLD:AAG6653
Bibio labradorensis|[14928]|TWDIP354-09|BOLD:AAG6653
Bibio labradorensis|[14929]|TWDIP353-09|BOLD:AAG6653
Bibio labradorensis|[14930]|MHFLI130-06|BOLD:AAG6653
Bibio inaequalis|[14931]|JWDCG328-10|BOLD:AAP8847
Bibio inaequalis|[14932]|JWDCJ1120-11|BOLD:AAP8847
Bibio inaequalis|[14933]|JWDCG617-10|BOLD:AAP8847
Bibio inaequalis|[14934]|JWDCG612-10|BOLD:AAP8847
Bibio inaequalis|[14935]|JWDCJ1121-11|BOLD:AAP8847
Bibio inaequalis|[14936]|JWDCJ1088-11|BOLD:AAP8847
Bibio inaequalis|[14937]|JWDCJ130-11|BOLD:AAP8847
Bibio inaequalis|[14938]|JWDCG616-10|BOLD:AAP8847
Bibio inaequalis|[14939]|JWDCG620-10|BOLD:AAP8847
Bibio inaequalis|[14940]|JWDCG622-10|BOLD:AAP8847
Bibio inaequalis|[14941]|JWDCH176-10|BOLD:AAP8847
Bibio inaequalis|[14942]|JWDCH260-10|BOLD:AAP8847
Bibio inaequalis|[14943]|JWDCI035-10|BOLD:AAP8847
Bibio inaequalis|[14944]|JWDCG621-10|BOLD:AAP8847
Bibio inaequalis|[14945]|JWDCG614-10|BOLD:AAP8847
Bibio inaequalis|[14946]|JWDCG613-10|BOLD:AAP8847
Bibio inaequalis|[14947]|JWDCJ133-11|BOLD:AAP8847
Bibio inaequalis|[14948]|JWDCJ132-11|BOLD:AAP8847
Bibio inaequalis|[14949]|JWDCL356-11|BOLD:AAP8847
Bibio inaequalis|[14950]|JWDCJ1119-11|BOLD:AAP8847
Bibio inaequalis|[14951]|JWDCJ1087-11|BOLD:AAP8847
Bibio inaequalis|[14952]|JWDCG615-10|BOLD:AAP8847
Bibio inaequalis|[14953]|JWDCI034-10|BOLD:AAP8847
Bibio inaequalis|[14954]|JWDCI036-10|BOLD:AAP8847
Bibio inaequalis|[14955]|JWDCH360-10|BOLD:AAP8847
Bibio inaequalis|[14956]|JWDCH361-10|BOLD:AAP8847
Bibio inaequalis|[14957]|JWDCH362-10|BOLD:AAP8847
Bibio fumipennis|[14958]|JWDCJ1044-11|BOLD:AAG6652
Bibio fumipennis|[14959]|JWDCK230-11|BOLD:AAG6652
Bibio fumipennis|[14960]|MTDIC152-09|BOLD:AAG6652
Bibio fumipennis|[14961]|MTDIC169-09|BOLD:AAG6652
Bibio fumipennis|[14962]|MTDIC166-09|BOLD:AAG6652
Bibio fumipennis|[14963]|JWDCB156-10|BOLD:AAG6652
Bibio fumipennis|[14964]|JWDCJ1045-11|BOLD:AAG6652
Bibio fumipennis|[14965]|JWDCB157-10|BOLD:AAG6652
Bibio fumipennis|[14966]|MTDIC168-09|BOLD:AAG6652
Bibio fumipennis|[14967]|JWDCH796-10|BOLD:AAG6652
Bibio fumipennis|[14968]|JWDCH048-10|BOLD:AAG6652
Bibio fumipennis|[14969]|JWDCH552-10|BOLD:AAG6652
Serromyia|[14970]|JWDCF176-10|BOLD:AAP9061
Serromyia|[14971]|JWDCF576-10|BOLD:AAP9061
Palpomyia lineata|[14972]|CHDIP788-12|BOLD:AAP9060
Palpomyia lineata|[14973]|JWDCF109-10|BOLD:AAP9060
Palpomyia lineata|[14974]|CHDIP829-12|BOLD:AAP9060
Palpomyia lineata|[14975]|CHDIP830-12|BOLD:AAP9060
Ceratopogonidae|[14976]|JWDCI355-10|BOLD:AAP9066
Ceratopogonidae|[14977]|JWDCG609-10|BOLD:AHE6322
Ceratopogonidae|[14978]|CHDIP1225-12|BOLD:AHE6322
Ceratopogoninae|[14979]|JWDCJ763-11|BOLD:AAV5157
Ceratopogonidae|[14980]|JWDCI301-10|BOLD:AAP9064
Bezzia sp. 8ES|[14981]|JWDCI299-10|BOLD:AAG6456
Bezzia sp. 8ES|[14982]|JWDCJ690-11|BOLD:AAG6456
Bezzia sp. 8ES|[14983]|SWSWP083-09|BOLD:AAG6456
Bezzia sp. 8ES|[14984]|CHDIP1056-12|BOLD:AAG6456
Bezzia sp. 8ES|[14985]|CHDIP1275-12|BOLD:AAG6456
Forcipomyia titillans|[14986]|JWDCA890-10|BOLD:AAG6501
Forcipomyia|[14987]|JWDCD437-10|BOLD:AAL7407
Forcipomyia|[14988]|JWDCC563-10|BOLD:AAL7407
Forcipomyia|[14989]|JWDCC562-10|BOLD:AAL7407
Forcipomyia|[14990]|JWDCC481-10|BOLD:AAL7407
Forcipomyia|[14991]|JWDCD294-10|BOLD:AAL7407
Ceratopogonidae|[14992]|JWDCC482-10|BOLD:AAL7408
Forcipomyia sp. 14ES|[14993]|CHURC839-09|BOLD:AEN8718
Forcipomyia sp. 14ES|[14994]|CHURC1058-09|BOLD:AEN8718
Forcipomyia sp. 14ES|[14995]|JWDCG826-10|BOLD:AEN8718
Forcipomyia sp. 14ES|[14996]|CHURC828-09|BOLD:AEN8718
Forcipomyia sp. 14ES|[14997]|JWDCI079-10|BOLD:AEN8718
Forcipomyia sp. 14ES|[14998]|JWDCH081-10|BOLD:AEN8718
Forcipomyia sp. 14ES|[14999]|JWDCH082-10|BOLD:AEN8718
Ceratopogonidae|[15000]|JWDCA901-10|BOLD:AAG6541
Ceratopogonidae|[15001]|CHDIP529-12|BOLD:ABY0685
Ceratopogonidae|[15002]|CHDIP770-12|BOLD:ABY0685
Atrichopogon|[15003]|JWDCH195-10|BOLD:AAG6519
Atrichopogon|[15004]|JWDCB364-10|BOLD:AAG6519
Atrichopogon|[15005]|JWDCA905-10|BOLD:AAG6519
Atrichopogon|[15006]|JWDCH677-10|BOLD:AAG6519
Atrichopogon|[15007]|JWDCA900-10|BOLD:AAG6519
Atrichopogon|[15008]|JWDCG262-10|BOLD:AAG6519
Atrichopogon|[15009]|JWDCA899-10|BOLD:AAG6519
Atrichopogon|[15010]|CHDIP055-11|BOLD:AAG6519
Atrichopogon|[15011]|CHDIP059-11|BOLD:AAG6519
Atrichopogon|[15012]|JWDCH503-10|BOLD:AAG6519
Atrichopogon|[15013]|JWDCL178-11|BOLD:AAG6519
Atrichopogon|[15014]|CHDIP446-11|BOLD:AAG6519
Atrichopogon|[15015]|JWDCH193-10|BOLD:AAG6519
Atrichopogon|[15016]|CHDIP060-11|BOLD:AAG6519
Atrichopogon|[15017]|CHDIP215-11|BOLD:AAG6519
Atrichopogon|[15018]|JWDCB366-10|BOLD:AAG6519
Atrichopogon|[15019]|JWDCH676-10|BOLD:AAG6519
Atrichopogon|[15020]|JWDCL228-11|BOLD:AAG6519
Atrichopogon|[15021]|JWDCL227-11|BOLD:AAG6519
Atrichopogon|[15022]|JWDCC740-10|BOLD:AAG6519
Atrichopogon|[15023]|JWDCB362-10|BOLD:AAG6519
Atrichopogon|[15024]|JWDCA898-10|BOLD:AAG6519
Atrichopogon|[15025]|JWDCB367-10|BOLD:AAG6519
Atrichopogon|[15026]|JWDCH422-10|BOLD:AAG6519

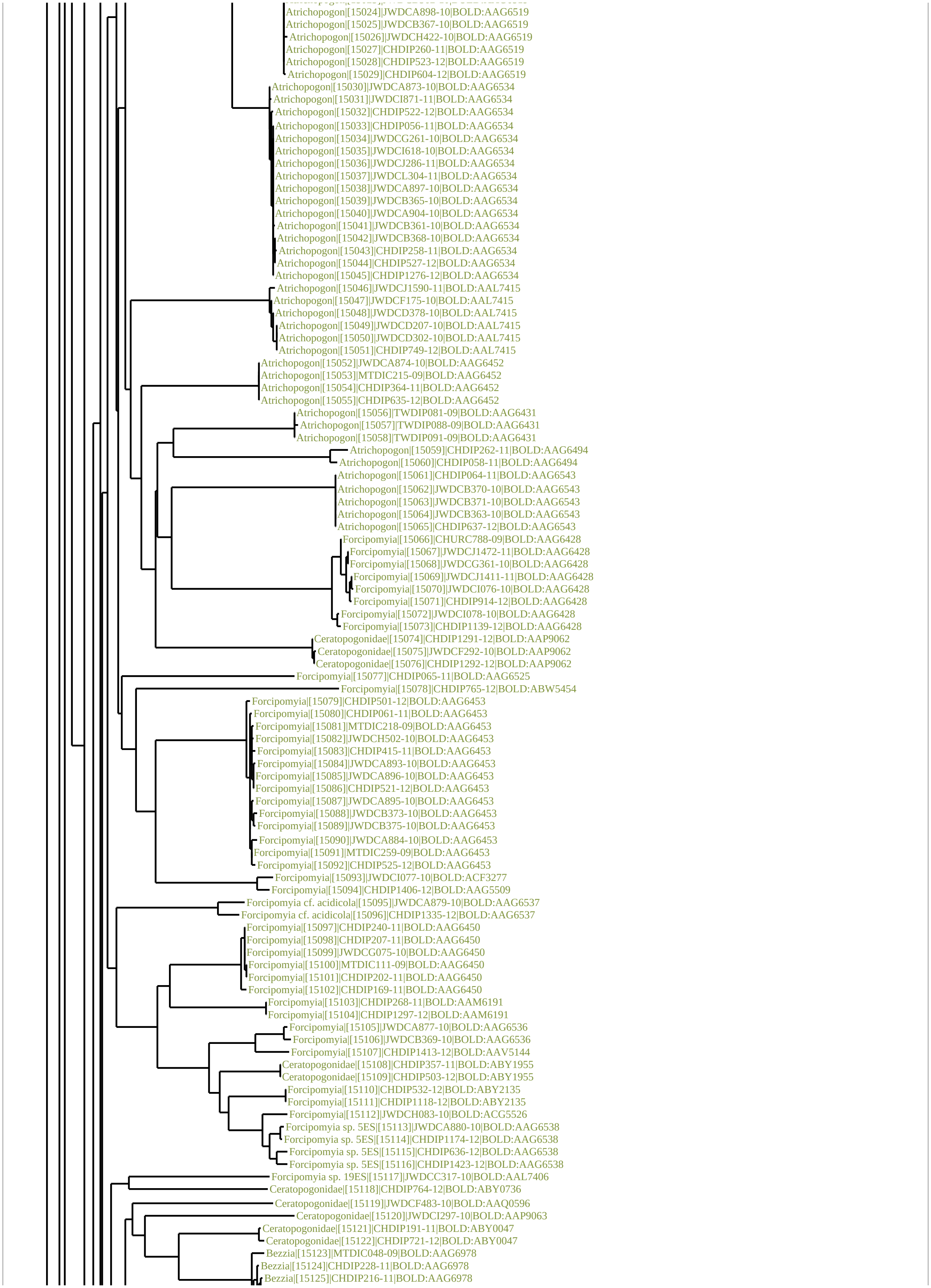
Atrichopogon|[15024]|JWDCA898-10|BOLD:AAG6519
Atrichopogon|[15025]|JWDCB367-10|BOLD:AAG6519
Atrichopogon|[15026]|JWDCH422-10|BOLD:AAG6519
Atrichopogon|[15027]|CHDIP260-11|BOLD:AAG6519
Atrichopogon|[15028]|CHDIP523-12|BOLD:AAG6519
Atrichopogon|[15029]|CHDIP604-12|BOLD:AAG6519
Atrichopogon|[15030]|JWDCA873-10|BOLD:AAG6534
Atrichopogon|[15031]|JWDCI871-11|BOLD:AAG6534
Atrichopogon|[15032]|CHDIP522-12|BOLD:AAG6534
Atrichopogon|[15033]|CHDIP056-11|BOLD:AAG6534
Atrichopogon|[15034]|JWDCG261-10|BOLD:AAG6534
Atrichopogon|[15035]|JWDCI618-10|BOLD:AAG6534
Atrichopogon|[15036]|JWDCJ286-11|BOLD:AAG6534
Atrichopogon|[15037]|JWDCL304-11|BOLD:AAG6534
Atrichopogon|[15038]|JWDCA897-10|BOLD:AAG6534
Atrichopogon|[15039]|JWDCB365-10|BOLD:AAG6534
Atrichopogon|[15040]|JWDCA904-10|BOLD:AAG6534
Atrichopogon|[15041]|JWDCB361-10|BOLD:AAG6534
Atrichopogon|[15042]|JWDCB368-10|BOLD:AAG6534
Atrichopogon|[15043]|CHDIP258-11|BOLD:AAG6534
Atrichopogon|[15044]|CHDIP527-12|BOLD:AAG6534
Atrichopogon|[15045]|CHDIP1276-12|BOLD:AAG6534
Atrichopogon|[15046]|JWDCJ1590-11|BOLD:AAL7415
Atrichopogon|[15047]|JWDCF175-10|BOLD:AAL7415
Atrichopogon|[15048]|JWDCD378-10|BOLD:AAL7415
Atrichopogon|[15049]|JWDCD207-10|BOLD:AAL7415
Atrichopogon|[15050]|JWDCD302-10|BOLD:AAL7415
Atrichopogon|[15051]|CHDIP749-12|BOLD:AAL7415
Atrichopogon|[15052]|JWDCA874-10|BOLD:AAG6452
Atrichopogon|[15053]|MTDIC215-09|BOLD:AAG6452
Atrichopogon|[15054]|CHDIP364-11|BOLD:AAG6452
Atrichopogon|[15055]|CHDIP635-12|BOLD:AAG6452
Atrichopogon|[15056]|TWDIP081-09|BOLD:AAG6431
Atrichopogon|[15057]|TWDIP088-09|BOLD:AAG6431
Atrichopogon|[15058]|TWDIP091-09|BOLD:AAG6431
Atrichopogon|[15059]|CHDIP262-11|BOLD:AAG6494
Atrichopogon|[15060]|CHDIP058-11|BOLD:AAG6494
Atrichopogon|[15061]|CHDIP064-11|BOLD:AAG6543
Atrichopogon|[15062]|JWDCB370-10|BOLD:AAG6543
Atrichopogon|[15063]|JWDCB371-10|BOLD:AAG6543
Atrichopogon|[15064]|JWDCB363-10|BOLD:AAG6543
Atrichopogon|[15065]|CHDIP637-12|BOLD:AAG6543
Forcipomyia|[15066]|CHURC788-09|BOLD:AAG6428
Forcipomyia|[15067]|JWDCJ1472-11|BOLD:AAG6428
Forcipomyia|[15068]|JWDCG361-10|BOLD:AAG6428
Forcipomyia|[15069]|JWDCJ1411-11|BOLD:AAG6428
Forcipomyia|[15070]|JWDCI076-10|BOLD:AAG6428
Forcipomyia|[15071]|CHDIP914-12|BOLD:AAG6428
Forcipomyia|[15072]|JWDCI078-10|BOLD:AAG6428
Forcipomyia|[15073]|CHDIP1139-12|BOLD:AAG6428
Ceratopogonidae|[15074]|CHDIP1291-12|BOLD:AAP9062
Ceratopogonidae|[15075]|JWDCF292-10|BOLD:AAP9062
Ceratopogonidae|[15076]|CHDIP1292-12|BOLD:AAP9062
Forcipomyia|[15077]|CHDIP065-11|BOLD:AAG6525
Forcipomyia|[15078]|CHDIP765-12|BOLD:ABW5454
Forcipomyia|[15079]|CHDIP501-12|BOLD:AAG6453
Forcipomyia|[15080]|CHDIP061-11|BOLD:AAG6453
Forcipomyia|[15081]|MTDIC218-09|BOLD:AAG6453
Forcipomyia|[15082]|JWDCH502-10|BOLD:AAG6453
Forcipomyia|[15083]|CHDIP415-11|BOLD:AAG6453
Forcipomyia|[15084]|JWDCA893-10|BOLD:AAG6453
Forcipomyia|[15085]|JWDCA896-10|BOLD:AAG6453
Forcipomyia|[15086]|CHDIP521-12|BOLD:AAG6453
Forcipomyia|[15087]|JWDCA895-10|BOLD:AAG6453
Forcipomyia|[15088]|JWDCB373-10|BOLD:AAG6453
Forcipomyia|[15089]|JWDCB375-10|BOLD:AAG6453
Forcipomyia|[15090]|JWDCA884-10|BOLD:AAG6453
Forcipomyia|[15091]|MTDIC259-09|BOLD:AAG6453
Forcipomyia|[15092]|CHDIP525-12|BOLD:AAG6453
Forcipomyia|[15093]|JWDCI077-10|BOLD:ACF3277
Forcipomyia|[15094]|CHDIP1406-12|BOLD:AAG5509
Forcipomyia cf. acidicola|[15095]|JWDCA879-10|BOLD:AAG6537
Forcipomyia cf. acidicola|[15096]|CHDIP1335-12|BOLD:AAG6537
Forcipomyia|[15097]|CHDIP240-11|BOLD:AAG6450
Forcipomyia|[15098]|CHDIP207-11|BOLD:AAG6450
Forcipomyia|[15099]|JWDCG075-10|BOLD:AAG6450
Forcipomyia|[15100]|MTDIC111-09|BOLD:AAG6450
Forcipomyia|[15101]|CHDIP202-11|BOLD:AAG6450
Forcipomyia|[15102]|CHDIP169-11|BOLD:AAG6450
Forcipomyia|[15103]|CHDIP268-11|BOLD:AAM6191
Forcipomyia|[15104]|CHDIP1297-12|BOLD:AAM6191
Forcipomyia|[15105]|JWDCA877-10|BOLD:AAG6536
Forcipomyia|[15106]|JWDCB369-10|BOLD:AAG6536
Forcipomyia|[15107]|CHDIP1413-12|BOLD:AAV5144
Ceratopogonidae|[15108]|CHDIP357-11|BOLD:ABY1955
Ceratopogonidae|[15109]|CHDIP503-12|BOLD:ABY1955
Forcipomyia|[15110]|CHDIP532-12|BOLD:ABY2135
Forcipomyia|[15111]|CHDIP1118-12|BOLD:ABY2135
Forcipomyia|[15112]|JWDCH083-10|BOLD:ACG5526
Forcipomyia sp. 5ES|[15113]|JWDCA880-10|BOLD:AAG6538
Forcipomyia sp. 5ES|[15114]|CHDIP1174-12|BOLD:AAG6538
Forcipomyia sp. 5ES|[15115]|CHDIP636-12|BOLD:AAG6538
Forcipomyia sp. 5ES|[15116]|CHDIP1423-12|BOLD:AAG6538
Forcipomyia sp. 19ES|[15117]|JWDCC317-10|BOLD:AAL7406
Ceratopogonidae|[15118]|CHDIP764-12|BOLD:ABY0736
Ceratopogonidae|[15119]|JWDCF483-10|BOLD:AAQ0596
Ceratopogonidae|[15120]|JWDCI297-10|BOLD:AAP9063
Ceratopogonidae|[15121]|CHDIP191-11|BOLD:ABY0047
Ceratopogonidae|[15122]|CHDIP721-12|BOLD:ABY0047
Bezzia|[15123]|MTDIC048-09|BOLD:AAG6978
Bezzia|[15124]|CHDIP228-11|BOLD:AAG6978
Bezzia|[15125]|CHDIP216-11|BOLD:AAG6978

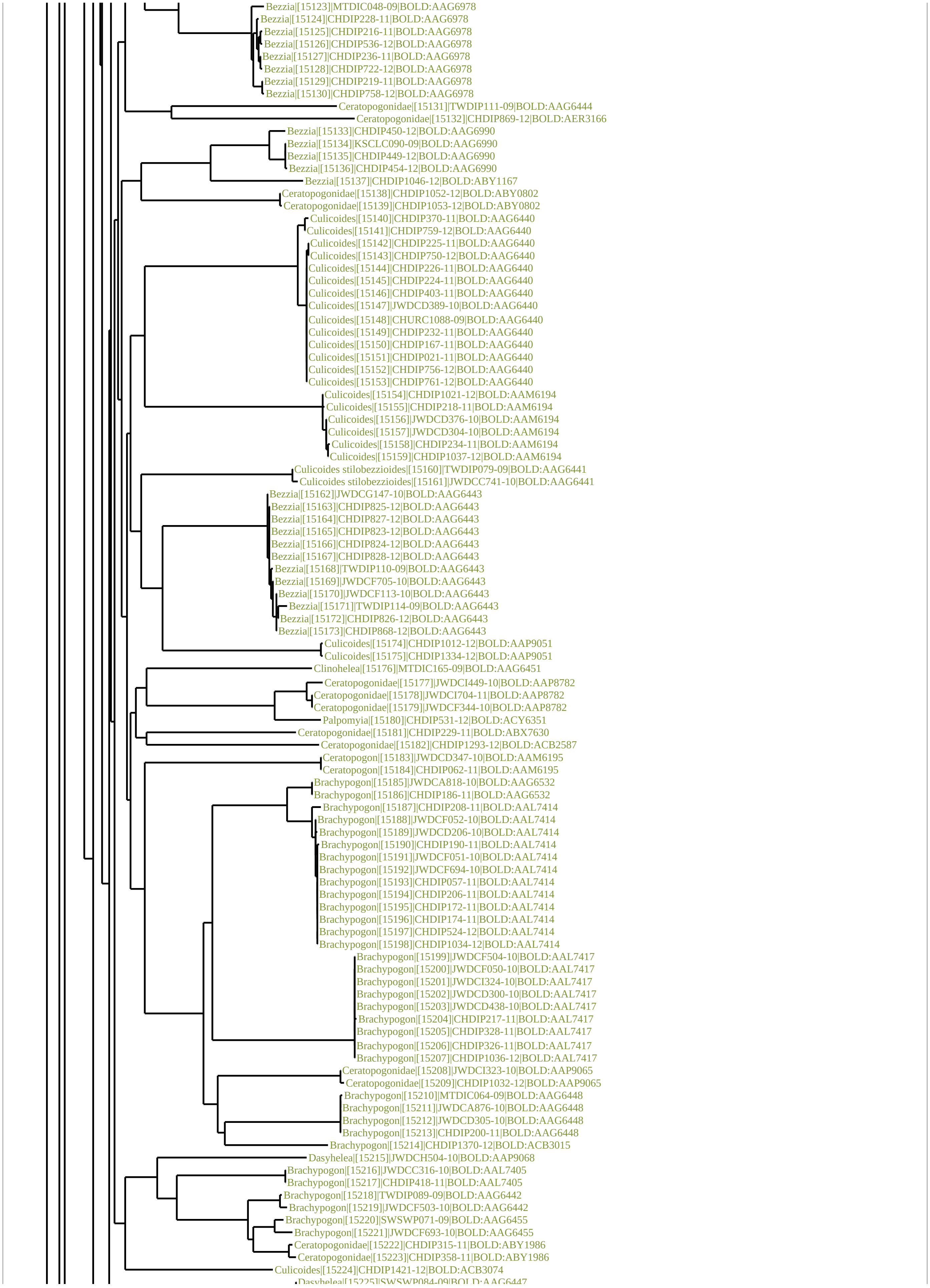
Bezzia|[15123]|MTDIC048-09|BOLD:AAG6978
Bezzia|[15124]|CHDIP228-11|BOLD:AAG6978
Bezzia|[15125]|CHDIP216-11|BOLD:AAG6978
Bezzia|[15126]|CHDIP536-12|BOLD:AAG6978
Bezzia|[15127]|CHDIP236-11|BOLD:AAG6978
Bezzia|[15128]|CHDIP722-12|BOLD:AAG6978
Bezzia|[15129]|CHDIP219-11|BOLD:AAG6978
Bezzia|[15130]|CHDIP758-12|BOLD:AAG6978
Ceratopogonidae|[15131]|TWDIP111-09|BOLD:AAG6444
Ceratopogonidae|[15132]|CHDIP869-12|BOLD:AER3166
Bezzia|[15133]|CHDIP450-12|BOLD:AAG6990
Bezzia|[15134]|KSCLC090-09|BOLD:AAG6990
Bezzia|[15135]|CHDIP449-12|BOLD:AAG6990
Bezzia|[15136]|CHDIP454-12|BOLD:AAG6990
Bezzia|[15137]|CHDIP1046-12|BOLD:ABY1167
Ceratopogonidae|[15138]|CHDIP1052-12|BOLD:ABY0802
Ceratopogonidae|[15139]|CHDIP1053-12|BOLD:ABY0802
Culicoides|[15140]|CHDIP370-11|BOLD:AAG6440
Culicoides|[15141]|CHDIP759-12|BOLD:AAG6440
Culicoides|[15142]|CHDIP225-11|BOLD:AAG6440
Culicoides|[15143]|CHDIP750-12|BOLD:AAG6440
Culicoides|[15144]|CHDIP226-11|BOLD:AAG6440
Culicoides|[15145]|CHDIP224-11|BOLD:AAG6440
Culicoides|[15146]|CHDIP403-11|BOLD:AAG6440
Culicoides|[15147]|JWDCD389-10|BOLD:AAG6440
Culicoides|[15148]|CHURC1088-09|BOLD:AAG6440
Culicoides|[15149]|CHDIP232-11|BOLD:AAG6440
Culicoides|[15150]|CHDIP167-11|BOLD:AAG6440
Culicoides|[15151]|CHDIP021-11|BOLD:AAG6440
Culicoides|[15152]|CHDIP756-12|BOLD:AAG6440
Culicoides|[15153]|CHDIP761-12|BOLD:AAG6440
Culicoides|[15154]|CHDIP1021-12|BOLD:AAM6194
Culicoides|[15155]|CHDIP218-11|BOLD:AAM6194
Culicoides|[15156]|JWDCD376-10|BOLD:AAM6194
Culicoides|[15157]|JWDCD304-10|BOLD:AAM6194
Culicoides|[15158]|CHDIP234-11|BOLD:AAM6194
Culicoides|[15159]|CHDIP1037-12|BOLD:AAM6194
Culicoides stilobezzioides|[15160]|TWDIP079-09|BOLD:AAG6441
Culicoides stilobezzioides|[15161]|JWDCC741-10|BOLD:AAG6441
Bezzia|[15162]|JWDCG147-10|BOLD:AAG6443
Bezzia|[15163]|CHDIP825-12|BOLD:AAG6443
Bezzia|[15164]|CHDIP827-12|BOLD:AAG6443
Bezzia|[15165]|CHDIP823-12|BOLD:AAG6443
Bezzia|[15166]|CHDIP824-12|BOLD:AAG6443
Bezzia|[15167]|CHDIP828-12|BOLD:AAG6443
Bezzia|[15168]|TWDIP110-09|BOLD:AAG6443
Bezzia|[15169]|JWDCF705-10|BOLD:AAG6443
Bezzia|[15170]|JWDCF113-10|BOLD:AAG6443
Bezzia|[15171]|TWDIP114-09|BOLD:AAG6443
Bezzia|[15172]|CHDIP826-12|BOLD:AAG6443
Bezzia|[15173]|CHDIP868-12|BOLD:AAG6443
Culicoides|[15174]|CHDIP1012-12|BOLD:AAP9051
Culicoides|[15175]|CHDIP1334-12|BOLD:AAP9051
Clinohelea|[15176]|MTDIC165-09|BOLD:AAG6451
Ceratopogonidae|[15177]|JWDCI449-10|BOLD:AAP8782
Ceratopogonidae|[15178]|JWDCI704-11|BOLD:AAP8782
Ceratopogonidae|[15179]|JWDCF344-10|BOLD:AAP8782
Palpomyia|[15180]|CHDIP531-12|BOLD:ACY6351
Ceratopogonidae|[15181]|CHDIP229-11|BOLD:ABX7630
Ceratopogonidae|[15182]|CHDIP1293-12|BOLD:ACB2587
Ceratopogon|[15183]|JWDCD347-10|BOLD:AAM6195
Ceratopogon|[15184]|CHDIP062-11|BOLD:AAM6195
Brachypogon|[15185]|JWDCA818-10|BOLD:AAG6532
Brachypogon|[15186]|CHDIP186-11|BOLD:AAG6532
Brachypogon|[15187]|CHDIP208-11|BOLD:AAL7414
Brachypogon|[15188]|JWDCF052-10|BOLD:AAL7414
Brachypogon|[15189]|JWDCD206-10|BOLD:AAL7414
Brachypogon|[15190]|CHDIP190-11|BOLD:AAL7414
Brachypogon|[15191]|JWDCF051-10|BOLD:AAL7414
Brachypogon|[15192]|JWDCF694-10|BOLD:AAL7414
Brachypogon|[15193]|CHDIP057-11|BOLD:AAL7414
Brachypogon|[15194]|CHDIP206-11|BOLD:AAL7414
Brachypogon|[15195]|CHDIP172-11|BOLD:AAL7414
Brachypogon|[15196]|CHDIP174-11|BOLD:AAL7414
Brachypogon|[15197]|CHDIP524-12|BOLD:AAL7414
Brachypogon|[15198]|CHDIP1034-12|BOLD:AAL7414
Brachypogon|[15199]|JWDCF504-10|BOLD:AAL7417
Brachypogon|[15200]|JWDCF050-10|BOLD:AAL7417
Brachypogon|[15201]|JWDCI324-10|BOLD:AAL7417
Brachypogon|[15202]|JWDCD300-10|BOLD:AAL7417
Brachypogon|[15203]|JWDCD438-10|BOLD:AAL7417
Brachypogon|[15204]|CHDIP217-11|BOLD:AAL7417
Brachypogon|[15205]|CHDIP328-11|BOLD:AAL7417
Brachypogon|[15206]|CHDIP326-11|BOLD:AAL7417
Brachypogon|[15207]|CHDIP1036-12|BOLD:AAL7417
Ceratopogonidae|[15208]|JWDCI323-10|BOLD:AAP9065
Ceratopogonidae|[15209]|CHDIP1032-12|BOLD:AAP9065
Brachypogon|[15210]|MTDIC064-09|BOLD:AAG6448
Brachypogon|[15211]|JWDCA876-10|BOLD:AAG6448
Brachypogon|[15212]|JWDCD305-10|BOLD:AAG6448
Brachypogon|[15213]|CHDIP200-11|BOLD:AAG6448
Brachypogon|[15214]|CHDIP1370-12|BOLD:ACB3015
Dasyhelea|[15215]|JWDCH504-10|BOLD:AAP9068
Brachypogon|[15216]|JWDCC316-10|BOLD:AAL7405
Brachypogon|[15217]|CHDIP418-11|BOLD:AAL7405
Brachypogon|[15218]|TWDIP089-09|BOLD:AAG6442
Brachypogon|[15219]|JWDCF503-10|BOLD:AAG6442
Brachypogon|[15220]|SWSWP071-09|BOLD:AAG6455
Brachypogon|[15221]|JWDCF693-10|BOLD:AAG6455
Ceratopogonidae|[15222]|CHDIP315-11|BOLD:ABY1986
Ceratopogonidae|[15223]|CHDIP358-11|BOLD:ABY1986
Culicoides|[15224]|CHDIP1421-12|BOLD:ACB3074
Dasyhelea|[15225]|SWSWP084-09|BOLD:AAG6447

[illegible]
Ceratopogonidae|[15223]|CHDIP358-11|BOLD:ABY1986
Culicoides|[15224]|CHDIP1421-12|BOLD:ACB3074
Dasyhelea|[15225]|SWSWP084-09|BOLD:AAG6447
Dasyhelea|[15226]|CHDIP189-11|BOLD:AAG6447
Dasyhelea|[15227]|MTDIC062-09|BOLD:AAG6447
Dasyhelea|[15228]|JWDCJ928-11|BOLD:AAG6447
Dasyhelea|[15229]|CHDIP198-11|BOLD:AAG6447
Dasyhelea|[15230]|CHDIP180-11|BOLD:AAG6447
Dasyhelea|[15231]|CHDIP182-11|BOLD:AAG5446
Dasyhelea|[15232]|CHDIP444-11|BOLD:AAG5446
Dasyhelea|[15233]|CHDIP195-11|BOLD:AAG5446
Dasyhelea|[15234]|CHDIP045-11|BOLD:AAG5446
Dasyhelea|[15235]|CHDIP530-12|BOLD:AAG5446
Dasyhelea|[15236]|CHDIP290-11|BOLD:ABY2126
Dasyhelea|[15237]|JWDCJ986-11|BOLD:AAG6544
Dasyhelea|[15238]|JWDCB379-10|BOLD:AAG6544
Dasyhelea|[15239]|JWDCB380-10|BOLD:AAG6544
Dasyhelea|[15240]|JWDCB374-10|BOLD:AAG6544
Dasyhelea|[15241]|CHDIP690-12|BOLD:AAG6544
Dasyhelea|[15242]|CHDIP1092-12|BOLD:AAG6544
Dasyhelea|[15243]|JWDCB376-10|BOLD:AAG6544
Dasyhelea|[15244]|CHDIP1329-12|BOLD:AAG6544
Dasyhelea|[15245]|JWDCC629-10|BOLD:AAL7409
Dasyhelea|[15246]|JWDCC633-10|BOLD:AAL7409
Dasyhelea|[15247]|JWDCC634-10|BOLD:AAL7409
Dasyhelea|[15248]|JWDCD301-10|BOLD:AAL7409
Dasyhelea|[15249]|JWDCD348-10|BOLD:AAL7409
Dasyhelea|[15250]|CHDIP201-11|BOLD:AAL7409
Dasyhelea|[15251]|JWDCF366-10|BOLD:AAG6454
Dasyhelea|[15252]|JWDCF377-10|BOLD:AAG6454
Dasyhelea|[15253]|SWSWP057-09|BOLD:AAG6454
Dasyhelea|[15254]|JWDCL644-11|BOLD:AAG6454
Dasyhelea|[15255]|JWDCI813-11|BOLD:AAG6454
Dasyhelea|[15256]|JWDCF233-10|BOLD:AAG6454
Dasyhelea|[15257]|JWDCF413-10|BOLD:AAG6454
Dasyhelea|[15258]|JWDCF672-10|BOLD:AAG6454
Dasyhelea|[15259]|JWDCF177-10|BOLD:AAG6454
Dasyhelea|[15260]|CHDIP1425-12|BOLD:AAG6454
Dasyhelea|[15261]|SWSWP091-09|BOLD:AAG6454
Dasyhelea|[15262]|JWDCL229-11|BOLD:AAG6454
Dasyhelea|[15263]|CHDIP1433-12|BOLD:AAG6454
Dasyhelea|[15264]|CHDIP356-11|BOLD:AAG6376
Ceratopogonidae|[15265]|CHDIP408-11|BOLD:ABY2049
Dasyhelea|[15266]|CHURC1053-09|BOLD:AAG6437
Dasyhelea|[15267]|JWDCA892-10|BOLD:AAG6437
Dasyhelea|[15268]|JWDCJ1473-11|BOLD:AAG6437
Dasyhelea|[15269]|CHDIP682-12|BOLD:AAN5161
Dasyhelea|[15270]|JWDCF414-10|BOLD:AAG6535
Dasyhelea|[15271]|CHDIP633-12|BOLD:AAG6535
Dasyhelea|[15272]|CHDIP054-11|BOLD:AAG6535
Dasyhelea|[15273]|CHDIP360-11|BOLD:AAG6535
Dasyhelea|[15274]|CHDIP359-11|BOLD:AAG6535
Dasyhelea|[15275]|JWDCA875-10|BOLD:AAG6535
Dasyhelea|[15276]|CHDIP649-12|BOLD:AAG6535
Dasyhelea|[15277]|CHDIP757-12|BOLD:AAG6535
Dasyhelea|[15278]|JWDCA878-10|BOLD:AAG6535
Dasyhelea|[15279]|CHDIP910-12|BOLD:AAG6535
Ceratopogonidae|[15280]|JWDCA871-10|BOLD:AAG6533
Ceratopogonidae|[15281]|JWDCA872-10|BOLD:AAG6533
Ceratopogonidae|[15282]|CHDIP639-12|BOLD:AAG6533
Ceratopogonidae|[15283]|CHDIP634-12|BOLD:AAG6533
Ceratopogonidae|[15284]|CHDIP1051-12|BOLD:AAG6533
Ceratopogonidae|[15285]|CHDIP760-12|BOLD:ABX7894
Ceratopogonidae|[15286]|CHDIP1054-12|BOLD:ABX7894
Ceratopogonidae|[15287]|JWDCJ606-11|BOLD:AAV5158
Ceratopogonidae|[15288]|JWDCD295-10|BOLD:AAM6193
Forcipomyia|[15289]|CHDIP338-11|BOLD:AAE0970
Forcipomyia|[15290]|JWDCF482-10|BOLD:AAE0970
Forcipomyia|[15291]|CHURC1086-09|BOLD:AAE0970
Forcipomyia|[15292]|JWDCC632-10|BOLD:AAE0970
Forcipomyia|[15293]|CHDIP193-11|BOLD:AAE0970
Forcipomyia|[15294]|JWDCC290-10|BOLD:AAE0970
Forcipomyia|[15295]|CHDIP775-12|BOLD:AAE0970
Forcipomyia|[15296]|CHDIP520-12|BOLD:AAE0970
Forcipomyia|[15297]|CHDIP768-12|BOLD:AAE0970
Forcipomyia|[15298]|CHDIP327-11|BOLD:AAE0970
Forcipomyia|[15299]|CHDIP177-11|BOLD:AAE0970
Forcipomyia|[15300]|CHDIP909-12|BOLD:AAE0970
Forcipomyia|[15301]|CHDIP230-11|BOLD:AAE0970
Forcipomyia|[15302]|CHDIP533-12|BOLD:AAE0970
Forcipomyia|[15303]|CHDIP950-12|BOLD:AAE0970
Forcipomyia|[15304]|JWDCC635-10|BOLD:AAE0970
Forcipomyia|[15305]|CHDIP1339-12|BOLD:AAE0970
Ceratopogonidae|[15306]|JWDCA881-10|BOLD:AAG6539
Ceratopogonidae|[15307]|JWDCD391-10|BOLD:AAG6539
Ceratopogonidae|[15308]|CHDIP1050-12|BOLD:ABY1515
Dasyhelea|[15309]|CHDIP1357-12|BOLD:ACB2575
Dasyhelea|[15310]|CHDIP1372-12|BOLD:ACB2575
Dasyhelea|[15311]|CHDIP1130-12|BOLD:ACX4903
Dasyhelea|[15312]|CHDIP1132-12|BOLD:ACX4903
Ceratopogonidae|[15313]|JWDCF086-10|BOLD:AAP9059
Ceratopogonidae|[15314]|CHDIP212-11|BOLD:AAP9059
Ceratopogonidae|[15315]|JWDCD208-10|BOLD:AAL7416
Dasyhelea|[15316]|CHDIP014-11|BOLD:ABW3988
Dasyhelea|[15317]|CHDIP771-12|BOLD:ABW3988
Dasyhelea|[15318]|JWDCA891-10|BOLD:AAG6540
Dasyhelea|[15319]|JWDCF085-10|BOLD:AAG6540
Dasyhelea|[15320]|CHDIP259-11|BOLD:AAG6540
Dasyhelea|[15321]|CHDIP362-11|BOLD:ACF3738
Ceratopogonidae|[15322]|JWDCI294-10|BOLD:AAG6438
Ceratopogonidae|[15323]|JWDCF564-10|BOLD:ACF3738
Ceratopogonidae|[15324]|JWDCA887-10|BOLD:ACF3738

Ceratopogonidae|[15322]|JWDCI294-10|BOLD:AAG6438
Ceratopogonidae|[15323]|JWDCF564-10|BOLD:ACF3738
Ceratopogonidae|[15324]|JWDCA887-10|BOLD:ACF3738
Ceratopogonidae|[15325]|MTDIC063-09|BOLD:ACF3738
Ceratopogonidae|[15326]|JWDCF566-10|BOLD:ACF3738
Ceratopogonidae|[15327]|CHURC1084-09|BOLD:AAG6438
Ceratopogonidae|[15328]|CHDIP763-12|BOLD:AAG6438
Ceratopogonidae|[15329]|CHDIP203-11|BOLD:ABZ0324
Dasyhelea|[15330]|JWDCF412-10|BOLD:AAL7404
Dasyhelea|[15331]|JWDCC288-10|BOLD:AAL7404
Dasyhelea|[15332]|JWDCF779-10|BOLD:AAL7404
Dasyhelea|[15333]|CHDIP196-11|BOLD:AAL7404
Dasyhelea|[15334]|CHDIP519-12|BOLD:AAL7404
Dasyhelea|[15335]|JWDCD445-10|BOLD:AAL7404
Dasyhelea|[15336]|JWDCF352-10|BOLD:AAL7404
Dasyhelea|[15337]|CHDIP949-12|BOLD:AAL7404
Dasyhelea sp. 20ES|[15338]|JWDCJ730-11|BOLD:AAG6439
Dasyhelea sp. 20ES|[15339]|JWDCF563-10|BOLD:AAG6439
Dasyhelea sp. 20ES|[15340]|CHDIP441-11|BOLD:AAG6439
Dasyhelea sp. 20ES|[15341]|CHDIP1033-12|BOLD:AAG6439
Dasyhelea sp. 20ES|[15342]|CHURC1087-09|BOLD:AAG6439
Dasyhelea sp. 20ES|[15343]|JWDCA889-10|BOLD:AAG6439
Dasyhelea sp. 20ES|[15344]|SWSWP070-09|BOLD:AAG6439
Dasyhelea sp. 20ES|[15345]|JWDCF565-10|BOLD:AAG6439
Dasyhelea sp. 20ES|[15346]|CHDIP242-11|BOLD:AAG6439
Dasyhelea sp. 20ES|[15347]|JWDCF780-10|BOLD:AAG6439
Dasyhelea sp. 20ES|[15348]|JWDCF353-10|BOLD:AAG6439
Dasyhelea sp. 20ES|[15349]|JWDCI296-10|BOLD:AAG6439
Dasyhelea sp. 20ES|[15350]|JWDCF234-10|BOLD:AAG6439
Dasyhelea sp. 20ES|[15351]|CHDIP231-11|BOLD:AAG6439
Dasyhelea sp. 20ES|[15352]|CHDIP314-11|BOLD:AAG6439
Dasyhelea sp. 20ES|[15353]|CHDIP376-11|BOLD:AAG6439
Dasyhelea sp. 20ES|[15354]|JWDCF800-10|BOLD:AAG6439
Dasyhelea sp. 20ES|[15355]|JWDCF562-10|BOLD:AAG6439
Dasyhelea sp. 20ES|[15356]|JWDCD303-10|BOLD:AAG6439
Dasyhelea sp. 20ES|[15357]|JWDCC638-10|BOLD:AAG6439
Dasyhelea sp. 20ES|[15358]|JWDCC636-10|BOLD:AAG6439
Dasyhelea sp. 20ES|[15359]|JWDCC630-10|BOLD:AAG6439
Dasyhelea sp. 20ES|[15360]|SWSWP004-09|BOLD:AAG6439
Dasyhelea sp. 20ES|[15361]|CHDIP1017-12|BOLD:AAG6439
Dasyhelea sp. 20ES|[15362]|CHDIP1018-12|BOLD:AAG6439
Dasyhelea sp. 20ES|[15363]|CHDIP1035-12|BOLD:AAG6439
Dasyhelea|[15364]|CHDIP1379-12|BOLD:ACB2687
Dasyhelea bifida|[15365]|CHDIP1342-12|BOLD:ABW3944
Dasyhelea bifida|[15366]|CHDIP1368-12|BOLD:ABW3944
Dasyhelea|[15367]|JWDCB377-10|BOLD:AAG6545
Dasyhelea|[15368]|JWDCI703-11|BOLD:AAG6545
Dasyhelea modesta|[15369]|JWDCC710-10|BOLD:AAG6542
Dasyhelea modesta|[15370]|JWDCA908-10|BOLD:AAG6542
Dasyhelea modesta|[15371]|JWDCF624-10|BOLD:AAG6542
Dasyhelea modesta|[15372]|CHDIP772-12|BOLD:AAG6542
Dasyhelea|[15373]|CHDIP928-12|BOLD:AAY6124
Ceratopogonidae|[15374]|CHDIP1332-12|BOLD:ACB2914
Dasyhelea|[15375]|JWDCJ605-11|BOLD:AAP8825
Dasyhelea|[15376]|JWDCJ731-11|BOLD:AAP8825
Dasyhelea|[15377]|JWDCJ992-11|BOLD:AAP8825
Dasyhelea|[15378]|JWDCG074-10|BOLD:AAP8825
Dasyhelea|[15379]|JWDCI458-10|BOLD:AAP8825
Dasyhelea|[15380]|JWDCG108-10|BOLD:AAP8825
Dasyhelea|[15381]|JWDCG109-10|BOLD:AAP8825
Dasyhelea|[15382]|CHDIP1031-12|BOLD:AAP8825
Dasyhelea|[15383]|JWDCI456-10|BOLD:AAP8825
Dasyhelea|[15384]|CHDIP1090-12|BOLD:AAP8825
Dasyhelea|[15385]|CHDIP1333-12|BOLD:AAP8825
Dasyhelea|[15386]|CHDIP1417-12|BOLD:AAP8825
Dasyhelea bensoni|[15387]|JWDCD439-10|BOLD:AAG6449
Dasyhelea bensoni|[15388]|JWDCD346-10|BOLD:AAG6449
Dasyhelea bensoni|[15389]|CHDIP239-11|BOLD:AAG6449
Dasyhelea bensoni|[15390]|JWDCC742-10|BOLD:AAG6449
Dasyhelea bensoni|[15391]|SWSWP006-09|BOLD:AAG6449
Dasyhelea bensoni|[15392]|CHDIP220-11|BOLD:AAG6449
Dasyhelea bensoni|[15393]|JWDCC665-10|BOLD:AAG6449
Dasyhelea bensoni|[15394]|CHDIP188-11|BOLD:AAG6449
Dasyhelea bensoni|[15395]|JWDCD292-10|BOLD:AAG6449
Dasyhelea bensoni|[15396]|JWDCC315-10|BOLD:AAG6449
Dasyhelea bensoni|[15397]|JWDCA907-10|BOLD:AAG6449
Dasyhelea bensoni|[15398]|JWDCA882-10|BOLD:AAG6449
Dasyhelea bensoni|[15399]|JWDCD290-10|BOLD:AAG6449
Dasyhelea bensoni|[15400]|CHDIP233-11|BOLD:AAG6449
Dasyhelea bensoni|[15401]|CHDIP165-11|BOLD:AAG6449
Dasyhelea bensoni|[15402]|CHDIP166-11|BOLD:AAG6449
Dasyhelea bensoni|[15403]|CHDIP181-11|BOLD:AAG6449
Dasyhelea bensoni|[15404]|JWDCD289-10|BOLD:AAG6449
Dasyhelea bensoni|[15405]|CHDIP179-11|BOLD:AAG6449
Dasyhelea bensoni|[15406]|JWDCD243-10|BOLD:AAG6449
Dasyhelea bensoni|[15407]|JWDCE931-10|BOLD:AAG6449
Dasyhelea bensoni|[15408]|CHDIP185-11|BOLD:AAG6449
Dasyhelea bensoni|[15409]|JWDCA906-10|BOLD:AAG6449
Dasyhelea bensoni|[15410]|JWDCD291-10|BOLD:AAG6449
Dasyhelea bensoni|[15411]|JWDCC628-10|BOLD:AAG6449
Dasyhelea bensoni|[15412]|JWDCA885-10|BOLD:AAG6449
Dasyhelea bensoni|[15413]|JWDCI570-10|BOLD:AAG6449
Dasyhelea bensoni|[15414]|JWDCA883-10|BOLD:AAG6449
Dasyhelea bensoni|[15415]|JWDCF623-10|BOLD:AAG6449
Dasyhelea bensoni|[15416]|CHDIP325-11|BOLD:AAG6449
Dasyhelea bensoni|[15417]|CHDIP170-11|BOLD:AAG6449
Dasyhelea bensoni|[15418]|MTDIC110-09|BOLD:AAG6449
Dasyhelea bensoni|[15419]|JWDCD293-10|BOLD:AAG6449
Dasyhelea bensoni|[15420]|CHDIP171-11|BOLD:AAG6449
Dasyhelea bensoni|[15421]|CHDIP766-12|BOLD:AAG6449
Dasyhelea bensoni|[15422]|CHDIP931-12|BOLD:AAG6449
Stenochironomus|[15423]|CHURC305-07|BOLD:AEZ7622
Sciophila|[15424]|CHDIP1195-12|BOLD:ACA4286

[illegible]
Dasyhelea bensoni|[15422]|CHDIP931-12|BOLD:AAG6449
Stenochironomus|[15423]|CHURC305-07|BOLD:AEZ7622
Sciophila|[15424]|CHDIP1195-12|BOLD:ACA4286
Sciophila|[15425]|CHDIP1200-12|BOLD:ACA4286
Sciophila|[15426]|CHDIP1244-12|BOLD:ACA4416
Sciophila|[15427]|CHDIP482-12|BOLD:ABW9766
Sciophila karelica|[15428]|CHDIP1231-12|BOLD:ABU6113
Sciophila sp. CJB1|[15429]|MTDIC028-09|BOLD:AAG4876
Sciophila|[15430]|CHDIP1243-12|BOLD:ACA4407
Sciophila sp. CJB1|[15431]|JWDCC216-10|BOLD:AAL9124
Sciophila sp. CJB1|[15432]|JWDCH290-10|BOLD:AAG4944
Sciophila lutea|[15433]|JWDCC060-10|BOLD:AEK6081
Sciophila lutea|[15434]|JWDCH289-10|BOLD:AEK6081
Sciophila ansluta|[15435]|CHDIP1210-12|BOLD:ACA4329
Sciophila hirta|[15436]|TWDIP319-09|BOLD:AAG4892
Sciophila|[15437]|CHDIP1228-12|BOLD:ACA4219
Sciophila|[15438]|CHDIP093-11|BOLD:ADQ1830
Sciophila|[15439]|CHDIP1251-12|BOLD:ACA4219
Neoplatyura noorae|[15440]|JWDCJ1586-11|BOLD:AAQ3542
Urytalpa atriceps|[15441]|JWDCA214-10|BOLD:AAM8955
Urytalpa atriceps|[15442]|JWDCA218-10|BOLD:AAM8955
Urytalpa atriceps|[15443]|JWDCA221-10|BOLD:AAM8955
Keroplatidae|[15444]|CHDIP494-12|BOLD:ABW9769
Keroplatidae|[15445]|CHDIP1218-12|BOLD:ACA4285
Macrocera|[15446]|JWDCK161-11|BOLD:AAU4909
Macrocera|[15447]|CHDIP1234-12|BOLD:ACA4302
Sciaridae|[15448]|CHDIP022-11|BOLD:ABW3421
Lycoriella|[15449]|JWDCB315-10|BOLD:AAH4031
Lycoriella|[15450]|JWDCJ722-11|BOLD:AAV1303
Cratyna|[15451]|JWDCC526-10|BOLD:AAL7870
Cratyna|[15452]|JWDCB297-10|BOLD:AAH3993
Corynoptera|[15453]|CHDIP988-12|BOLD:AAU6564
Corynoptera|[15454]|CHDIP1346-12|BOLD:AAU6564
Sciaridae|[15455]|JWDCC312-10|BOLD:AAL7862
Sciaridae|[15456]|JWDCC513-10|BOLD:AAL7862
Sciaridae|[15457]|JWDCC509-10|BOLD:AAL7862
Sciaridae|[15458]|JWDCC304-10|BOLD:AAL7862
Falakia modesta|[15459]|JWDCH537-10|BOLD:AAM9253
Trichosiopsis|[15460]|JWDCC642-10|BOLD:AAL7877
Trichosiopsis|[15461]|JWDCC512-10|BOLD:AAL7861
Trichosiopsis|[15462]|JWDCC511-10|BOLD:AAL7861
Trichosiopsis|[15463]|JWDCC530-10|BOLD:AAL7861
Trichosiopsis|[15464]|JWDCC301-10|BOLD:AAL7861
Sciaridae|[15465]|JWDCE025-10|BOLD:AAP1834
Sciaridae|[15466]|JWDCC689-10|BOLD:AAL7880
Dolichosciara saetosa|[15467]|JWDCG650-10|BOLD:AAH3908
Dolichosciara saetosa|[15468]|JWDCL307-11|BOLD:AAH3908
Dolichosciara saetosa|[15469]|JWDCG687-10|BOLD:AAH3908
Dolichosciara saetosa|[15470]|JWDCH287-10|BOLD:AAH3908
Dolichosciara saetosa|[15471]|CHDIP874-12|BOLD:AAH3908
Sciaridae|[15472]|JWDCC687-10|BOLD:AAL7879
Corynoptera trepida|[15473]|JWDCC306-10|BOLD:AAL7864
Corynoptera trepida|[15474]|JWDCC607-10|BOLD:AAL7864
Corynoptera trepida|[15475]|JWDCD327-10|BOLD:AAL7864
Corynoptera trepida|[15476]|JWDCC309-10|BOLD:AAL7864
Corynoptera trepida|[15477]|JWDCC528-10|BOLD:AAL7864
Corynoptera trepida|[15478]|JWDCC537-10|BOLD:AAL7864
Corynoptera trepida|[15479]|JWDCF429-10|BOLD:AAL7864
Corynoptera trepida|[15480]|JWDCE899-10|BOLD:AAL7864
Corynoptera|[15481]|CHDIP989-12|BOLD:AAH3612
Corynoptera|[15482]|JWDCD279-10|BOLD:AAH3612
Corynoptera|[15483]|JWDCF430-10|BOLD:AAH3612
Corynoptera|[15484]|JWDCF081-10|BOLD:AAH3612
Corynoptera|[15485]|CHDIP1019-12|BOLD:AAH3612
Alpinosciara subtrivialis|[15486]|JWDCG239-10|BOLD:AAP6422
Alpinosciara subtrivialis|[15487]|JWDCG363-10|BOLD:AAP6422
Alpinosciara subtrivialis|[15488]|JWDCG409-10|BOLD:AAP6422
Alpinosciara subtrivialis|[15489]|JWDCH499-10|BOLD:AAP6422
Dichopygina bernhardi|[15490]|JWDCF773-10|BOLD:AAP9901
Dichopygina aculeata|[15491]|JWDCI069-10|BOLD:AAH4002
Dichopygina aculeata|[15492]|JWDCG584-10|BOLD:AAH4002
Dichopygina aculeata|[15493]|JWEMP274-10|BOLD:AAH4002
Dichopygina aculeata|[15494]|JWDCB317-10|BOLD:AAH4002
Dichopygina aculeata|[15495]|JWDCA777-10|BOLD:AAH4002
Dichopygina aculeata|[15496]|JWDCD424-10|BOLD:AAH4002
Dichopygina aculeata|[15497]|JWDCL234-11|BOLD:AAH4002
Dichopygina aculeata|[15498]|JWDCJ1465-11|BOLD:AAH4002
Dichopygina aculeata|[15499]|JWDCH690-10|BOLD:AAH4002
Dichopygina aculeata|[15500]|JWDCI017-10|BOLD:AAH4002
Dichopygina aculeata|[15501]|JWDCI065-10|BOLD:AAH4002
Dichopygina aculeata|[15502]|JWDCI070-10|BOLD:AAH4002
Dichopygina duplicis|[15503]|JWDCA790-10|BOLD:AAH4007
Dichopygina duplicis|[15504]|CHDIP294-11|BOLD:AAH4007
Sciaridae|[15505]|JWDCD422-10|BOLD:AAL7895
Sciaridae|[15506]|JWDCC534-10|BOLD:AAL7872
Trichosiopsis|[15507]|TWDIP229-09|BOLD:AAH3930
Corynoptera blanda|[15508]|JWDCF772-10|BOLD:AAP9900
Corynoptera blanda|[15509]|CHDIP546-12|BOLD:AAP9900
Claustropyga acanthostyla|[15510]|JWDCC535-10|BOLD:AAL7873
Claustropyga|[15511]|JWDCC518-10|BOLD:AAL7868
Claustropyga corticis|[15512]|JWDCK164-11|BOLD:AAQ2140
Claustropyga corticis|[15513]|JWDCJ1583-11|BOLD:AAQ2140
Trichosia|[15514]|TWDIP269-09|BOLD:AAH3932
Corynoptera|[15515]|JWDCF546-10|BOLD:AAH3937
Corynoptera|[15516]|TWDIP372-09|BOLD:AAH3937
Corynoptera|[15517]|JWDCA829-10|BOLD:AAH3937
Corynoptera|[15518]|JWDCA831-10|BOLD:AAH3937
Corynoptera|[15519]|JWDCC515-10|BOLD:AAH3937
Corynoptera|[15520]|JWDCG072-10|BOLD:AAH3937
Corynoptera|[15521]|CHDIP385-11|BOLD:AAH3937
Corynoptera|[15522]|CHDIP307-11|BOLD:AAH3937
Scatopsciara subciliata|[15523]|JWDCA781-10|BOLD:AAH4004

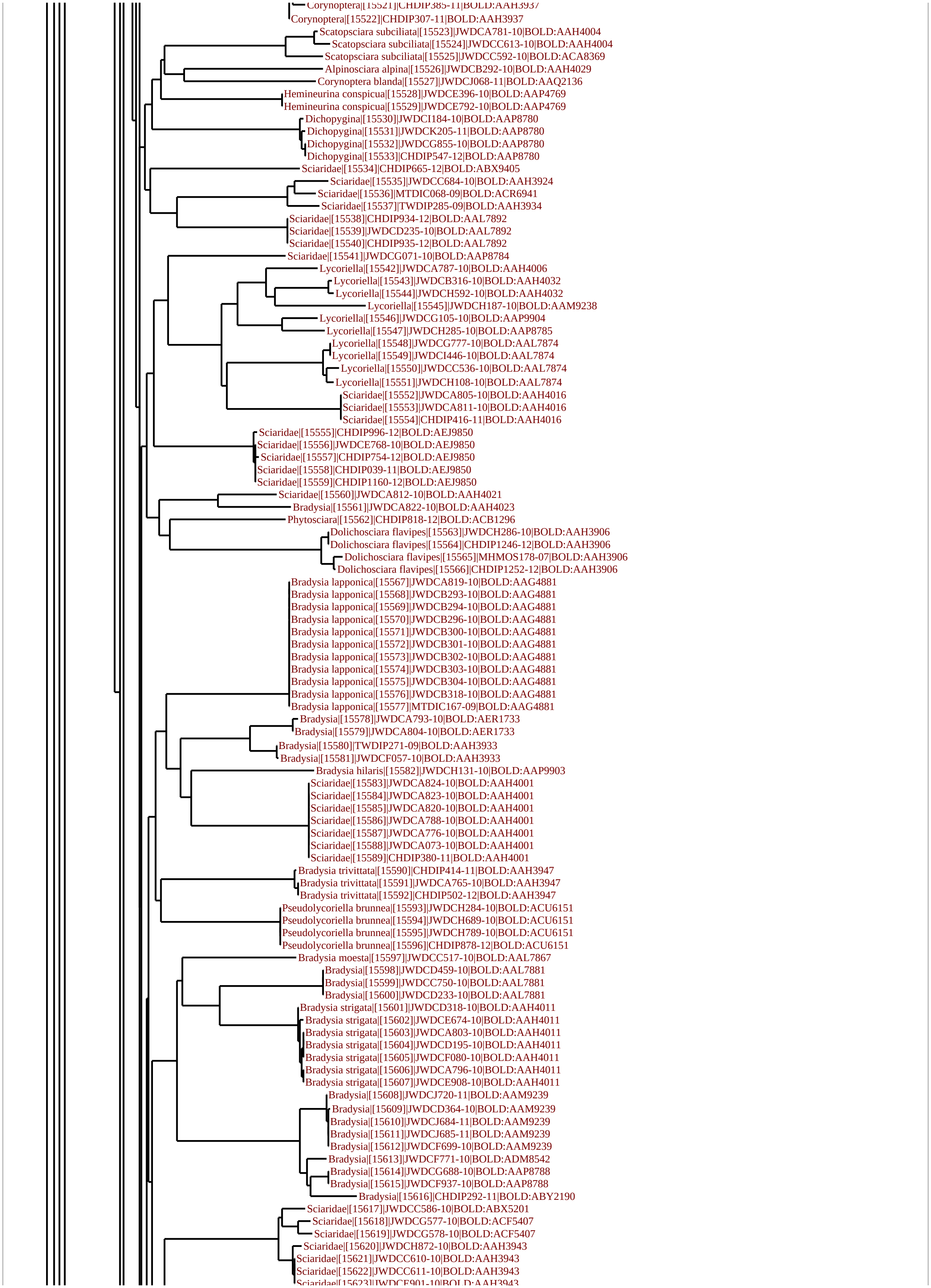
Corynoptera|[15521]|CHDIP385-11|BOLD:AAH3937
Corynoptera|[15522]|CHDIP307-11|BOLD:AAH3937
Scatopsciara subciliata|[15523]|JWDCA781-10|BOLD:AAH4004
Scatopsciara subciliata|[15524]|JWDCC613-10|BOLD:AAH4004
Scatopsciara subciliata|[15525]|JWDCC592-10|BOLD:ACA8369
Alpinosciara alpina|[15526]|JWDCB292-10|BOLD:AAH4029
Corynoptera blanda|[15527]|JWDCJ068-11|BOLD:AAQ2136
Hemineurina conspicua|[15528]|JWDCE396-10|BOLD:AAP4769
Hemineurina conspicua|[15529]|JWDCE792-10|BOLD:AAP4769
Dichopygina|[15530]|JWDCI184-10|BOLD:AAP8780
Dichopygina|[15531]|JWDCK205-11|BOLD:AAP8780
Dichopygina|[15532]|JWDCG855-10|BOLD:AAP8780
Dichopygina|[15533]|CHDIP547-12|BOLD:AAP8780
Sciaridae|[15534]|CHDIP665-12|BOLD:ABX9405
Sciaridae|[15535]|JWDCC684-10|BOLD:AAH3924
Sciaridae|[15536]|MTDIC068-09|BOLD:ACR6941
Sciaridae|[15537]|TWDIP285-09|BOLD:AAH3934
Sciaridae|[15538]|CHDIP934-12|BOLD:AAL7892
Sciaridae|[15539]|JWDCD235-10|BOLD:AAL7892
Sciaridae|[15540]|CHDIP935-12|BOLD:AAL7892
Sciaridae|[15541]|JWDCG071-10|BOLD:AAP8784
Lycoriella|[15542]|JWDCA787-10|BOLD:AAH4006
Lycoriella|[15543]|JWDCB316-10|BOLD:AAH4032
Lycoriella|[15544]|JWDCH592-10|BOLD:AAH4032
Lycoriella|[15545]|JWDCH187-10|BOLD:AAM9238
Lycoriella|[15546]|JWDCG105-10|BOLD:AAP9904
Lycoriella|[15547]|JWDCH285-10|BOLD:AAP8785
Lycoriella|[15548]|JWDCG777-10|BOLD:AAL7874
Lycoriella|[15549]|JWDCI446-10|BOLD:AAL7874
Lycoriella|[15550]|JWDCC536-10|BOLD:AAL7874
Lycoriella|[15551]|JWDCH108-10|BOLD:AAL7874
Sciaridae|[15552]|JWDCA805-10|BOLD:AAH4016
Sciaridae|[15553]|JWDCA811-10|BOLD:AAH4016
Sciaridae|[15554]|CHDIP416-11|BOLD:AAH4016
Sciaridae|[15555]|CHDIP996-12|BOLD:AEJ9850
Sciaridae|[15556]|JWDCE768-10|BOLD:AEJ9850
Sciaridae|[15557]|CHDIP754-12|BOLD:AEJ9850
Sciaridae|[15558]|CHDIP039-11|BOLD:AEJ9850
Sciaridae|[15559]|CHDIP1160-12|BOLD:AEJ9850
Sciaridae|[15560]|JWDCA812-10|BOLD:AAH4021
Bradysia|[15561]|JWDCA822-10|BOLD:AAH4023
Phytosciara|[15562]|CHDIP818-12|BOLD:ACB1296
Dolichosciara flavipes|[15563]|JWDCH286-10|BOLD:AAH3906
Dolichosciara flavipes|[15564]|CHDIP1246-12|BOLD:AAH3906
Dolichosciara flavipes|[15565]|MHMOS178-07|BOLD:AAH3906
Dolichosciara flavipes|[15566]|CHDIP1252-12|BOLD:AAH3906
Bradysia lapponica|[15567]|JWDCA819-10|BOLD:AAG4881
Bradysia lapponica|[15568]|JWDCB293-10|BOLD:AAG4881
Bradysia lapponica|[15569]|JWDCB294-10|BOLD:AAG4881
Bradysia lapponica|[15570]|JWDCB296-10|BOLD:AAG4881
Bradysia lapponica|[15571]|JWDCB300-10|BOLD:AAG4881
Bradysia lapponica|[15572]|JWDCB301-10|BOLD:AAG4881
Bradysia lapponica|[15573]|JWDCB302-10|BOLD:AAG4881
Bradysia lapponica|[15574]|JWDCB303-10|BOLD:AAG4881
Bradysia lapponica|[15575]|JWDCB304-10|BOLD:AAG4881
Bradysia lapponica|[15576]|JWDCB318-10|BOLD:AAG4881
Bradysia lapponica|[15577]|MTDIC167-09|BOLD:AAG4881
Bradysia|[15578]|JWDCA793-10|BOLD:AER1733
Bradysia|[15579]|JWDCA804-10|BOLD:AER1733
Bradysia|[15580]|TWDIP271-09|BOLD:AAH3933
Bradysia|[15581]|JWDCF057-10|BOLD:AAH3933
Bradysia hilaris|[15582]|JWDCH131-10|BOLD:AAP9903
Sciaridae|[15583]|JWDCA824-10|BOLD:AAH4001
Sciaridae|[15584]|JWDCA823-10|BOLD:AAH4001
Sciaridae|[15585]|JWDCA820-10|BOLD:AAH4001
Sciaridae|[15586]|JWDCA788-10|BOLD:AAH4001
Sciaridae|[15587]|JWDCA776-10|BOLD:AAH4001
Sciaridae|[15588]|JWDCA073-10|BOLD:AAH4001
Sciaridae|[15589]|CHDIP380-11|BOLD:AAH4001
Bradysia trivittata|[15590]|CHDIP414-11|BOLD:AAH3947
Bradysia trivittata|[15591]|JWDCA765-10|BOLD:AAH3947
Bradysia trivittata|[15592]|CHDIP502-12|BOLD:AAH3947
Pseudolycoriella brunnea|[15593]|JWDCH284-10|BOLD:ACU6151
Pseudolycoriella brunnea|[15594]|JWDCH689-10|BOLD:ACU6151
Pseudolycoriella brunnea|[15595]|JWDCH789-10|BOLD:ACU6151
Pseudolycoriella brunnea|[15596]|CHDIP878-12|BOLD:ACU6151
Bradysia moesta|[15597]|JWDCC517-10|BOLD:AAL7867
Bradysia|[15598]|JWDCD459-10|BOLD:AAL7881
Bradysia|[15599]|JWDCC750-10|BOLD:AAL7881
Bradysia|[15600]|JWDCD233-10|BOLD:AAL7881
Bradysia strigata|[15601]|JWDCD318-10|BOLD:AAH4011
Bradysia strigata|[15602]|JWDCE674-10|BOLD:AAH4011
Bradysia strigata|[15603]|JWDCA803-10|BOLD:AAH4011
Bradysia strigata|[15604]|JWDCD195-10|BOLD:AAH4011
Bradysia strigata|[15605]|JWDCF080-10|BOLD:AAH4011
Bradysia strigata|[15606]|JWDCA796-10|BOLD:AAH4011
Bradysia strigata|[15607]|JWDCE908-10|BOLD:AAH4011
Bradysia|[15608]|JWDCJ720-11|BOLD:AAM9239
Bradysia|[15609]|JWDCD364-10|BOLD:AAM9239
Bradysia|[15610]|JWDCJ684-11|BOLD:AAM9239
Bradysia|[15611]|JWDCJ685-11|BOLD:AAM9239
Bradysia|[15612]|JWDCF699-10|BOLD:AAM9239
Bradysia|[15613]|JWDCF771-10|BOLD:ADM8542
Bradysia|[15614]|JWDCG688-10|BOLD:AAP8788
Bradysia|[15615]|JWDCF937-10|BOLD:AAP8788
Bradysia|[15616]|CHDIP292-11|BOLD:ABY2190
Sciaridae|[15617]|JWDCC586-10|BOLD:ABX5201
Sciaridae|[15618]|JWDCG577-10|BOLD:ACF5407
Sciaridae|[15619]|JWDCG578-10|BOLD:ACF5407
Sciaridae|[15620]|JWDCH872-10|BOLD:AAH3943
Sciaridae|[15621]|JWDCC610-10|BOLD:AAH3943
Sciaridae|[15622]|JWDCC611-10|BOLD:AAH3943
Sciaridae|[15623]|JWDCE901-10|BOLD:AAH3943

Sciaridae|[15621]|JWDCC610-10|BOLD:AAH3943
Sciaridae|[15622]|JWDCC611-10|BOLD:AAH3943
Sciaridae|[15623]|JWDCE901-10|BOLD:AAH3943
Corynoptera|[15624]|JWDCI447-10|BOLD:AAV1308
Corynoptera|[15625]|JWDCI448-10|BOLD:AAV1308
Sciaridae|[15626]|JWDCC806-10|BOLD:AAH3909
Sciaridae|[15627]|JWDCH423-10|BOLD:AAH3909
Bradysia|[15628]|JWDCD453-10|BOLD:AAL7896
Bradysia|[15629]|JWDCC516-10|BOLD:AAH4026
Bradysia|[15630]|JWDCA830-10|BOLD:AAH4026
Bradysia|[15631]|JWDCD165-10|BOLD:AAH4026
Bradysia|[15632]|JWDCD421-10|BOLD:AAL7894
Bradysia|[15633]|JWDCF612-10|BOLD:AAL7894
Bradysia|[15634]|JWDCA774-10|BOLD:AAH3927
Bradysia|[15635]|JWDCA821-10|BOLD:AAH3927
Bradysia|[15636]|MTDIC234-09|BOLD:AAH3927
Bradysia|[15637]|CHDIP451-12|BOLD:AAH3927
Bradysia|[15638]|JWDCG103-10|BOLD:AAP9899
Bradysia|[15639]|JWDCF559-10|BOLD:ACR7948
Bradysia|[15640]|CHDIP316-11|BOLD:ACR7948
Bradysia|[15641]|CHDIP1041-12|BOLD:ABX9898
Bradysia|[15642]|JWDCA799-10|BOLD:AAH4012
Bradysia|[15643]|JWDCD365-10|BOLD:AAH4012
Bradysia|[15644]|JWDCA752-10|BOLD:ACF0037
Bradysia|[15645]|MTDIC252-09|BOLD:ACF0037
Bradysia iridipennis|[15646]|TWDIP192-09|BOLD:AAH3923
Bradysia iridipennis|[15647]|MTDIC027-09|BOLD:AAH3923
Bradysia iridipennis|[15648]|TWDIP315-09|BOLD:AAH3923
Bradysia iridipennis|[15649]|TWDIP316-09|BOLD:AAH3923
Bradysia iridipennis|[15650]|TWDIP318-09|BOLD:AAH3923
Bradysia iridipennis|[15651]|JWDCD239-10|BOLD:AAH3923
Bradysia iridipennis|[15652]|JWDCJ768-11|BOLD:AAH3923
Bradysia iridipennis|[15653]|JWDCI774-11|BOLD:AAH3923
Bradysia iridipennis|[15654]|JWDCI937-11|BOLD:AAH3923
Bradysia iridipennis|[15655]|JWDCF323-10|BOLD:AAH3923
Bradysia iridipennis|[15656]|TWDIP317-09|BOLD:AAH3923
Bradysia iridipennis|[15657]|TWDIP298-09|BOLD:AAH3923
Bradysia iridipennis|[15658]|JWDCL309-11|BOLD:AAH3923
Bradysia iridipennis|[15659]|JWDCI699-11|BOLD:AAH3923
Bradysia iridipennis|[15660]|JWDCF938-10|BOLD:AAH3923
Bradysia iridipennis|[15661]|CHDIP142-11|BOLD:AAH3923
Bradysia|[15662]|JWDCI946-11|BOLD:AAQ2135
Bradysia|[15663]|CHDIP151-11|BOLD:AAQ2135
Bradysia|[15664]|JWDCD258-10|BOLD:AAL7888
Bradysia|[15665]|JWDCD429-10|BOLD:AAL7888
Bradysia|[15666]|JWDCC805-10|BOLD:AAL7888
Bradysia|[15667]|JWDCD240-10|BOLD:AAL7888
Bradysia|[15668]|JWDCD259-10|BOLD:AAL7888
Bradysia|[15669]|JWDCD261-10|BOLD:AAL7888
Bradysia|[15670]|JWDCD280-10|BOLD:AAL7888
Bradysia|[15671]|JWDCD281-10|BOLD:AAL7888
Bradysia|[15672]|JWDCD283-10|BOLD:AAL7888
Bradysia|[15673]|JWDCD284-10|BOLD:AAL7888
Bradysia|[15674]|JWDCF150-10|BOLD:AAL7888
Sciaridae|[15675]|JWDCC685-10|BOLD:AAH4018
Sciaridae|[15676]|SAHYM320-10|BOLD:AAH4018
Sciaridae|[15677]|JWDCD368-10|BOLD:AAH4018
Sciaridae|[15678]|JWDCD234-10|BOLD:AAH4018
Sciaridae|[15679]|JWDCC298-10|BOLD:AAH4018
Sciaridae|[15680]|JWDCD172-10|BOLD:AAH4018
Sciaridae|[15681]|JWDCD170-10|BOLD:AAH4018
Sciaridae|[15682]|JWDCD168-10|BOLD:AAH4018
Sciaridae|[15683]|JWDCD166-10|BOLD:AAH4018
Sciaridae|[15684]|JWDCC813-10|BOLD:AAH4018
Sciaridae|[15685]|JWDCC810-10|BOLD:AAH4018
Sciaridae|[15686]|JWDCC531-10|BOLD:AAH4018
Sciaridae|[15687]|JWDCD457-10|BOLD:AAH4018
Sciaridae|[15688]|JWDCD456-10|BOLD:AAH4018
Sciaridae|[15689]|JWDCA827-10|BOLD:AAH4018
Sciaridae|[15690]|JWDCA826-10|BOLD:AAH4018
Sciaridae|[15691]|JWDCA807-10|BOLD:AAH4018
Sciaridae|[15692]|JWDCD460-10|BOLD:AAH4018
Sciaridae|[15693]|JWDCD427-10|BOLD:AAH4018
Sciaridae|[15694]|JWDCD425-10|BOLD:AAH4018
Sciaridae|[15695]|JWDCD423-10|BOLD:AAH4018
Sciaridae|[15696]|JWDCD282-10|BOLD:AAH4018
Sciaridae|[15697]|JWDCE942-10|BOLD:AAH4018
Sciaridae|[15698]|JWDCE162-10|BOLD:AAP3648
Sciaridae|[15699]|JWDCE500-10|BOLD:AAP3648
Bradysia|[15700]|JWDCE811-10|BOLD:AAP4772
Bradysia vernalis|[15701]|JWDCE290-10|BOLD:AAP4768
Bradysia vernalis|[15702]|JWDCF614-10|BOLD:AAP4768
Bradysia vernalis|[15703]|CHDIP932-12|BOLD:AAP3649
Bradysia vernalis|[15704]|JWDCE160-10|BOLD:AAP3649
Bradysia vernalis|[15705]|JWDCJ428-11|BOLD:AAP3649
Bradysia vernalis|[15706]|JWDCE288-10|BOLD:AAP3649
Bradysia vernalis|[15707]|JWDCE502-10|BOLD:AAP3649
Bradysia vernalis|[15708]|CHDIP936-12|BOLD:AAP3649
Sciaridae|[15709]|JWDCE885-10|BOLD:AAP9902
Sciaridae|[15710]|JWDCE884-10|BOLD:AAP9902
Sciaridae|[15711]|JWDCE882-10|BOLD:AAP9902
Sciaridae|[15712]|JWDCE886-10|BOLD:AAP9902
Sciaridae|[15713]|CHDIP1001-12|BOLD:AAP9902
Sciaridae|[15714]|CHDIP1025-12|BOLD:AAP9902
Sciaridae|[15715]|CHDIP1026-12|BOLD:AAP9902
Scatopsciara atomaria|[15716]|JWDCF390-10|BOLD:AAH3920
Scatopsciara atomaria|[15717]|TWDIP077-09|BOLD:AAH3920
Scatopsciara atomaria|[15718]|TWDIP078-09|BOLD:AAH3920
Scatopsciara atomaria|[15719]|TWDIP092-09|BOLD:AAH3920
Scatopsciara atomaria|[15720]|JWDCJ295-11|BOLD:AAH3920
Scatopsciara atomaria|[15721]|JWDCE909-10|BOLD:AAH3920
Scatopsciara atomaria|[15722]|JWDCB291-10|BOLD:AAH3920

Scatopsciara atomaria|[15720]|JWDCJ295-11|BOLD:AAH3920
Scatopsciara atomaria|[15721]|JWDCE909-10|BOLD:AAH3920
Scatopsciara atomaria|[15722]|JWDCB291-10|BOLD:AAH3920
Scatopsciara atomaria|[15723]|CHDIP305-11|BOLD:AAH3920
Sciaridae|[15724]|JWDCA785-10|BOLD:AAH4005
Scatopsciara vitripennis|[15725]|TWDIP200-09|BOLD:AAU6764
Bradysia flavipila|[15726]|JWDCA755-10|BOLD:AAH3929
Bradysia flavipila|[15727]|JWDCA760-10|BOLD:AAH3929
Bradysia flavipila|[15728]|CHDIP539-12|BOLD:AAH3929
Bradysia flavipila|[15729]|CHDIP452-12|BOLD:AAH3929
Bradysia flavipila|[15730]|JWDCA764-10|BOLD:AAH3929
Bradysia flavipila|[15731]|JWDCA753-10|BOLD:AAH3929
Bradysia flavipila|[15732]|JWDCA759-10|BOLD:AAH3929
Bradysia flavipila|[15733]|JWDCA758-10|BOLD:AAH3929
Bradysia flavipila|[15734]|JWDCA757-10|BOLD:AAH3929
Bradysia flavipila|[15735]|JWDCA754-10|BOLD:AAH3929
Bradysia flavipila|[15736]|TWDIP205-09|BOLD:AAH3929
Bradysia flavipila|[15737]|TWDIP270-09|BOLD:AAH3929
Bradysia flavipila|[15738]|CHDIP548-12|BOLD:AAH3929
Sciaridae|[15739]|CHDIP1158-12|BOLD:AAV1324
Lycoriella|[15740]|TWDIP080-09|BOLD:AAH3922
Trichocoelina dividua|[15741]|JWDCD455-10|BOLD:AAL7893
Trichocoelina dividua|[15742]|TWDIP155-09|BOLD:AAL7893
Trichocoelina dividua|[15743]|JWDCD237-10|BOLD:AAL7893
Trichocoelina dividua|[15744]|JWDCI775-11|BOLD:AAL7893
Trichocoelina dividua|[15745]|CHDIP309-11|BOLD:AAL7893
Lycoriella|[15746]|JWDCA802-10|BOLD:AAH4015
Lycoriella|[15747]|JWDCC809-10|BOLD:AAH4015
Lycoriella|[15748]|CHDIP163-11|BOLD:AAH4015
Hemineurina flavipeda|[15749]|JWDCC603-10|BOLD:AAL7869
Hemineurina flavipeda|[15750]|JWDCC583-10|BOLD:AAL7869
Hemineurina flavipeda|[15751]|JWDCC519-10|BOLD:AAL7869
Hemineurina flavipeda|[15752]|JWDCC686-10|BOLD:AAL7869
Hemineurina flavipeda|[15753]|JWDCC584-10|BOLD:AAL7869
Hemineurina flavipeda|[15754]|JWDCC605-10|BOLD:AAL7869
Hemineurina flavipeda|[15755]|JWDCC712-10|BOLD:AAL7869
Hemineurina flavipeda|[15756]|JWDCC606-10|BOLD:AAL7869
Hemineurina ventrosa|[15757]|JWDCE784-10|BOLD:AAP4771
Hemineurina ventrosa|[15758]|JWDCE783-10|BOLD:AAP4771
Lycoriella|[15759]|JWDCA767-10|BOLD:AAH3998
Lycoriella|[15760]|CHDIP538-12|BOLD:AAH3998
Bradysiopsis vittigera|[15761]|JWDCD215-10|BOLD:AAH4010
Bradysiopsis vittigera|[15762]|JWDCD214-10|BOLD:AAH4010
Bradysiopsis vittigera|[15763]|JWDCD163-10|BOLD:AAH4010
Bradysiopsis vittigera|[15764]|JWDCC508-10|BOLD:AAH4010
Bradysiopsis vittigera|[15765]|JWDCD216-10|BOLD:AAH4010
Bradysiopsis vittigera|[15766]|JWDCD320-10|BOLD:AAH4010
Bradysiopsis vittigera|[15767]|JWDCD324-10|BOLD:AAH4010
Bradysiopsis vittigera|[15768]|JWDCD329-10|BOLD:AAH4010
Bradysiopsis vittigera|[15769]|JWDCD332-10|BOLD:AAH4010
Bradysiopsis vittigera|[15770]|JWDCE910-10|BOLD:AAH4010
Bradysiopsis vittigera|[15771]|JWDCB286-10|BOLD:AAH4010
Bradysiopsis vittigera|[15772]|JWDCE881-10|BOLD:AAH4010
Bradysiopsis vittigera|[15773]|JWDCE087-10|BOLD:AAH4010
Bradysiopsis vittigera|[15774]|JWDCJ469-11|BOLD:AAH4010
Bradysiopsis vittigera|[15775]|JWDCJ468-11|BOLD:AAH4010
Bradysiopsis vittigera|[15776]|JWDCJ467-11|BOLD:AAH4010
Bradysiopsis vittigera|[15777]|JWDCJ466-11|BOLD:AAH4010
Bradysiopsis vittigera|[15778]|JWDCJ427-11|BOLD:AAH4010
Bradysiopsis vittigera|[15779]|JWDCJ415-11|BOLD:AAH4010
Bradysiopsis vittigera|[15780]|JWDCJ414-11|BOLD:AAH4010
Bradysiopsis vittigera|[15781]|JWDCJ413-11|BOLD:AAH4010
Bradysiopsis vittigera|[15782]|JWDCJ412-11|BOLD:AAH4010
Bradysiopsis vittigera|[15783]|JWDCD167-10|BOLD:AAH4010
Bradysiopsis vittigera|[15784]|JWDCA795-10|BOLD:AAH4010
Bradysiopsis vittigera|[15785]|JWDCC814-10|BOLD:AAH4010
Bradysiopsis vittigera|[15786]|JWDCE325-10|BOLD:AAH4010
Bradysiopsis vittigera|[15787]|JWDCJ465-11|BOLD:AAH4010
Bradysiopsis vittigera|[15788]|JWDCE326-10|BOLD:AAH4010
Bradysiopsis vittigera|[15789]|JWDCE327-10|BOLD:AAH4010
Bradysiopsis vittigera|[15790]|JWDCE767-10|BOLD:AAH4010
Bradysiopsis vittigera|[15791]|JWDCE810-10|BOLD:AAH4010
Bradysiopsis vittigera|[15792]|JWDCE011-10|BOLD:AAH4010
Bradysiopsis vittigera|[15793]|JWDCE012-10|BOLD:AAH4010
Bradysiopsis vittigera|[15794]|JWDCE013-10|BOLD:AAH4010
Bradysiopsis vittigera|[15795]|JWDCE014-10|BOLD:AAH4010
Bradysiopsis vittigera|[15796]|JWDCE015-10|BOLD:AAH4010
Bradysiopsis vittigera|[15797]|JWDCE128-10|BOLD:AAH4010
Bradysiopsis vittigera|[15798]|JWDCE197-10|BOLD:AAH4010
Bradysiopsis vittigera|[15799]|JWDCE248-10|BOLD:AAH4010
Bradysiopsis vittigera|[15800]|JWDCE250-10|BOLD:AAH4010
Bradysiopsis vittigera|[15801]|JWDCE263-10|BOLD:AAH4010
Bradysiopsis vittigera|[15802]|JWDCE273-10|BOLD:AAH4010
Bradysiopsis vittigera|[15803]|JWDCE278-10|BOLD:AAH4010
Bradysiopsis vittigera|[15804]|JWDCE630-10|BOLD:AAH4010
Bradysiopsis vittigera|[15805]|JWDCE631-10|BOLD:AAH4010
Bradysiopsis vittigera|[15806]|JWDCE632-10|BOLD:AAH4010
Bradysiopsis vittigera|[15807]|JWDCF545-10|BOLD:AAH4010
Bradysiopsis vittigera|[15808]|JWDCF613-10|BOLD:AAH4010
Bradysiopsis vittigera|[15809]|CHDIP1027-12|BOLD:AAH4010
Bradysiopsis vittigera|[15810]|JWDCG316-10|BOLD:AAP8777
Bradysiopsis vittigera|[15811]|JWDCF059-10|BOLD:AAP8777
Bradysiopsis vittigera|[15812]|CHDIP1271-12|BOLD:AAP8777
Bradysia|[15813]|JWDCD454-10|BOLD:AAL7897
Bradysia|[15814]|JWDCJ721-11|BOLD:AAV1302
Bradysia|[15815]|JWDCA792-10|BOLD:AAH4008
Bradysia excelsa|[15816]|JWDCA775-10|BOLD:AAH3907
Bradysia excelsa|[15817]|JWDCA782-10|BOLD:AAH3907
Bradysia excelsa|[15818]|JWDCB305-10|BOLD:AAH3907
Bradysia excelsa|[15819]|JWDCC527-10|BOLD:AAH3907
Bradysia excelsa|[15820]|JWDCL397-11|BOLD:AAH3907
Bradysia excelsa|[15821]|JWDCA780-10|BOLD:AAH3907
Bradysia excelsa|[15822]|JWDCG032-10|BOLD:AAH3907

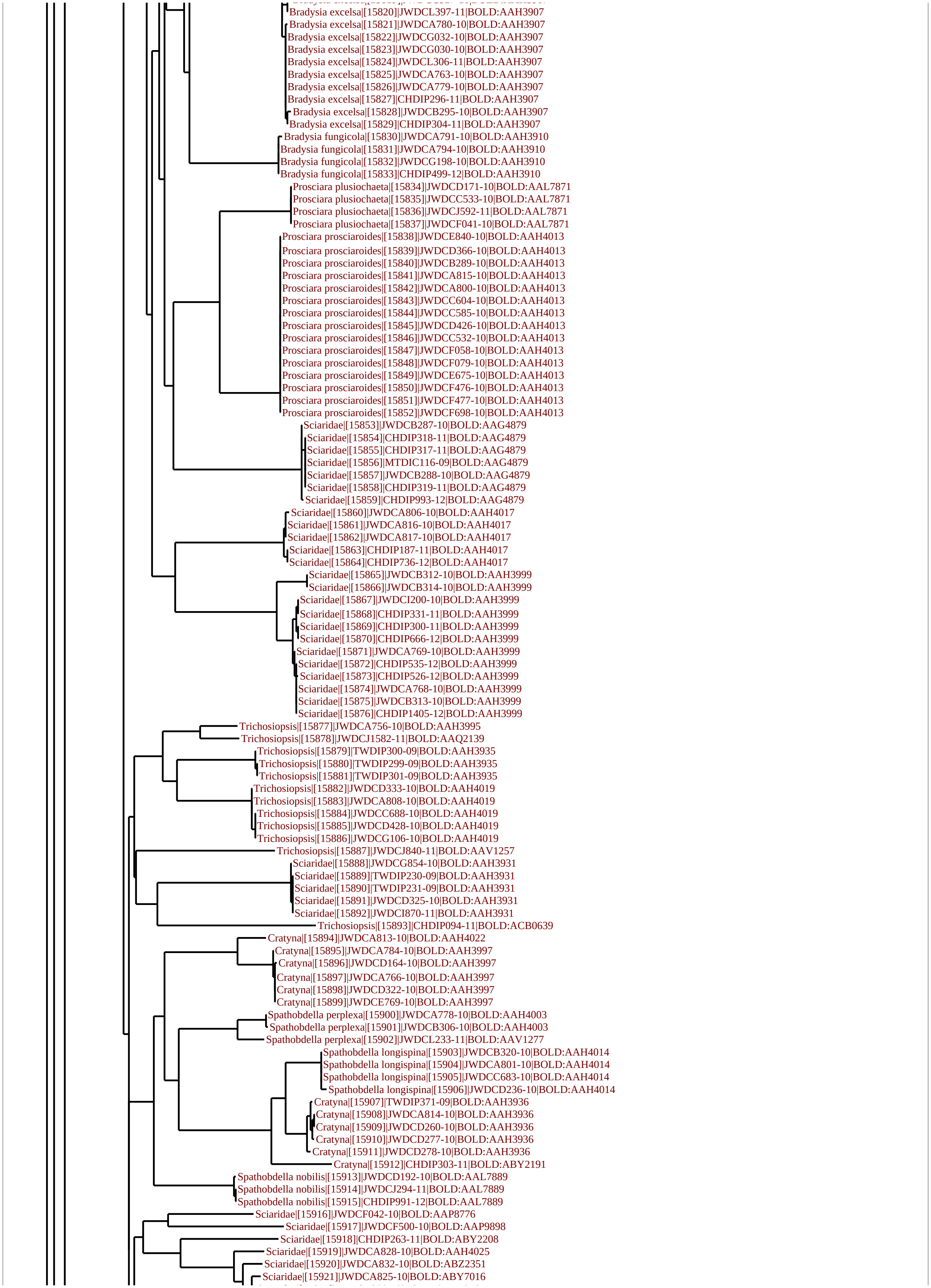

Bradysia excelsa|[15820]|JWDCL397-11|BOLD:AAH3907
Bradysia excelsa|[15821]|JWDCA780-10|BOLD:AAH3907
Bradysia excelsa|[15822]|JWDCG032-10|BOLD:AAH3907
Bradysia excelsa|[15823]|JWDCG030-10|BOLD:AAH3907
Bradysia excelsa|[15824]|JWDCL306-11|BOLD:AAH3907
Bradysia excelsa|[15825]|JWDCA763-10|BOLD:AAH3907
Bradysia excelsa|[15826]|JWDCA779-10|BOLD:AAH3907
Bradysia excelsa|[15827]|CHDIP296-11|BOLD:AAH3907
Bradysia excelsa|[15828]|JWDCB295-10|BOLD:AAH3907
Bradysia excelsa|[15829]|CHDIP304-11|BOLD:AAH3907
Bradysia fungicola|[15830]|JWDCA791-10|BOLD:AAH3910
Bradysia fungicola|[15831]|JWDCA794-10|BOLD:AAH3910
Bradysia fungicola|[15832]|JWDCG198-10|BOLD:AAH3910
Bradysia fungicola|[15833]|CHDIP499-12|BOLD:AAH3910
Prosciara plusiochaeta|[15834]|JWDCD171-10|BOLD:AAL7871
Prosciara plusiochaeta|[15835]|JWDCC533-10|BOLD:AAL7871
Prosciara plusiochaeta|[15836]|JWDCJ592-11|BOLD:AAL7871
Prosciara plusiochaeta|[15837]|JWDCF041-10|BOLD:AAL7871
Prosciara prosciaroides|[15838]|JWDCE840-10|BOLD:AAH4013
Prosciara prosciaroides|[15839]|JWDCD366-10|BOLD:AAH4013
Prosciara prosciaroides|[15840]|JWDCB289-10|BOLD:AAH4013
Prosciara prosciaroides|[15841]|JWDCA815-10|BOLD:AAH4013
Prosciara prosciaroides|[15842]|JWDCA800-10|BOLD:AAH4013
Prosciara prosciaroides|[15843]|JWDCC604-10|BOLD:AAH4013
Prosciara prosciaroides|[15844]|JWDCC585-10|BOLD:AAH4013
Prosciara prosciaroides|[15845]|JWDCD426-10|BOLD:AAH4013
Prosciara prosciaroides|[15846]|JWDCC532-10|BOLD:AAH4013
Prosciara prosciaroides|[15847]|JWDCF058-10|BOLD:AAH4013
Prosciara prosciaroides|[15848]|JWDCF079-10|BOLD:AAH4013
Prosciara prosciaroides|[15849]|JWDCE675-10|BOLD:AAH4013
Prosciara prosciaroides|[15850]|JWDCF476-10|BOLD:AAH4013
Prosciara prosciaroides|[15851]|JWDCF477-10|BOLD:AAH4013
Prosciara prosciaroides|[15852]|JWDCF698-10|BOLD:AAH4013
Sciaridae|[15853]|JWDCB287-10|BOLD:AAG4879
Sciaridae|[15854]|CHDIP318-11|BOLD:AAG4879
Sciaridae|[15855]|CHDIP317-11|BOLD:AAG4879
Sciaridae|[15856]|MTDIC116-09|BOLD:AAG4879
Sciaridae|[15857]|JWDCB288-10|BOLD:AAG4879
Sciaridae|[15858]|CHDIP319-11|BOLD:AAG4879
Sciaridae|[15859]|CHDIP993-12|BOLD:AAG4879
Sciaridae|[15860]|JWDCA806-10|BOLD:AAH4017
Sciaridae|[15861]|JWDCA816-10|BOLD:AAH4017
Sciaridae|[15862]|JWDCA817-10|BOLD:AAH4017
Sciaridae|[15863]|CHDIP187-11|BOLD:AAH4017
Sciaridae|[15864]|CHDIP736-12|BOLD:AAH4017
Sciaridae|[15865]|JWDCB312-10|BOLD:AAH3999
Sciaridae|[15866]|JWDCB314-10|BOLD:AAH3999
Sciaridae|[15867]|JWDCI200-10|BOLD:AAH3999
Sciaridae|[15868]|CHDIP331-11|BOLD:AAH3999
Sciaridae|[15869]|CHDIP300-11|BOLD:AAH3999
Sciaridae|[15870]|CHDIP666-12|BOLD:AAH3999
Sciaridae|[15871]|JWDCA769-10|BOLD:AAH3999
Sciaridae|[15872]|CHDIP535-12|BOLD:AAH3999
Sciaridae|[15873]|CHDIP526-12|BOLD:AAH3999
Sciaridae|[15874]|JWDCA768-10|BOLD:AAH3999
Sciaridae|[15875]|JWDCB313-10|BOLD:AAH3999
Sciaridae|[15876]|CHDIP1405-12|BOLD:AAH3999
Trichosiopsis|[15877]|JWDCA756-10|BOLD:AAH3995
Trichosiopsis|[15878]|JWDCJ1582-11|BOLD:AAQ2139
Trichosiopsis|[15879]|TWDIP300-09|BOLD:AAH3935
Trichosiopsis|[15880]|TWDIP299-09|BOLD:AAH3935
Trichosiopsis|[15881]|TWDIP301-09|BOLD:AAH3935
Trichosiopsis|[15882]|JWDCD333-10|BOLD:AAH4019
Trichosiopsis|[15883]|JWDCA808-10|BOLD:AAH4019
Trichosiopsis|[15884]|JWDCC688-10|BOLD:AAH4019
Trichosiopsis|[15885]|JWDCD428-10|BOLD:AAH4019
Trichosiopsis|[15886]|JWDCG106-10|BOLD:AAH4019
Trichosiopsis|[15887]|JWDCJ840-11|BOLD:AAV1257
Sciaridae|[15888]|JWDCG854-10|BOLD:AAH3931
Sciaridae|[15889]|TWDIP230-09|BOLD:AAH3931
Sciaridae|[15890]|TWDIP231-09|BOLD:AAH3931
Sciaridae|[15891]|JWDCD325-10|BOLD:AAH3931
Sciaridae|[15892]|JWDCI870-11|BOLD:AAH3931
Trichosiopsis|[15893]|CHDIP094-11|BOLD:ACB0639
Cratyna|[15894]|JWDCA813-10|BOLD:AAH4022
Cratyna|[15895]|JWDCA784-10|BOLD:AAH3997
Cratyna|[15896]|JWDCD164-10|BOLD:AAH3997
Cratyna|[15897]|JWDCA766-10|BOLD:AAH3997
Cratyna|[15898]|JWDCD322-10|BOLD:AAH3997
Cratyna|[15899]|JWDCE769-10|BOLD:AAH3997
Spathobdella perplexa|[15900]|JWDCA778-10|BOLD:AAH4003
Spathobdella perplexa|[15901]|JWDCB306-10|BOLD:AAH4003
Spathobdella perplexa|[15902]|JWDCL233-11|BOLD:AAV1277
Spathobdella longispina|[15903]|JWDCB320-10|BOLD:AAH4014
Spathobdella longispina|[15904]|JWDCA801-10|BOLD:AAH4014
Spathobdella longispina|[15905]|JWDCC683-10|BOLD:AAH4014
Spathobdella longispina|[15906]|JWDCD236-10|BOLD:AAH4014
Cratyna|[15907]|TWDIP371-09|BOLD:AAH3936
Cratyna|[15908]|JWDCA814-10|BOLD:AAH3936
Cratyna|[15909]|JWDCD260-10|BOLD:AAH3936
Cratyna|[15910]|JWDCD277-10|BOLD:AAH3936
Cratyna|[15911]|JWDCD278-10|BOLD:AAH3936
Cratyna|[15912]|CHDIP303-11|BOLD:ABY2191
Spathobdella nobilis|[15913]|JWDCD192-10|BOLD:AAL7889
Spathobdella nobilis|[15914]|JWDCJ294-11|BOLD:AAL7889
Spathobdella nobilis|[15915]|CHDIP991-12|BOLD:AAL7889
Sciaridae|[15916]|JWDCF042-10|BOLD:AAP8776
Sciaridae|[15917]|JWDCF500-10|BOLD:AAP9898
Sciaridae|[15918]|CHDIP263-11|BOLD:ABY2208
Sciaridae|[15919]|JWDCA828-10|BOLD:AAH4025
Sciaridae|[15920]|JWDCA832-10|BOLD:ABZ2351
Sciaridae|[15921]|JWDCA825-10|BOLD:ABY7016

Sciaridae|[15919]|JWDCA828-10|BOLD:AAH4025
Sciaridae|[15920]|JWDCA832-10|BOLD:ABZ2351
Sciaridae|[15921]|JWDCA825-10|BOLD:ABY7016
Sciaridae|[15922]|JWDCB308-10|BOLD:ABY7016
Sciaridae|[15923]|JWDCB310-10|BOLD:ABY7016
Sciaridae|[15924]|CHDIP302-11|BOLD:ABY7016
Sciaridae|[15925]|JWDCD396-10|BOLD:AAL7450
Sciaridae|[15926]|CHDIP382-11|BOLD:ABY2036
Sciaridae|[15927]|CHDIP542-12|BOLD:ABY2199
Corynoptera|[15928]|JWDCC510-10|BOLD:AAL7865
Corynoptera|[15929]|JWDCD194-10|BOLD:AAL7891
Corynoptera|[15930]|JWDCD285-10|BOLD:AHA4920
Sciaridae|[15931]|JWDCC305-10|BOLD:AAL7863
Sciaridae|[15932]|JWDCL308-11|BOLD:AAQ2138
Sciaridae|[15933]|JWDCG251-10|BOLD:AAP8787
Bradysia|[15934]|JWDCC614-10|BOLD:AAL7876
Bradysia|[15935]|JWDCF936-10|BOLD:AAL7876
Sciaridae|[15936]|MTDIC148-09|BOLD:AAH3925
Sciaridae|[15937]|CHDIP694-12|BOLD:AAH3925
Sciaridae|[15938]|CHDIP695-12|BOLD:AAH3925
Sciaridae|[15939]|CHDIP696-12|BOLD:AAH3925
Sciaridae|[15940]|CHDIP663-12|BOLD:AAH3928
Sciaridae|[15941]|CHDIP1119-12|BOLD:AAH3928
Sciaridae|[15942]|JWDCA789-10|BOLD:AAH3928
Sciaridae|[15943]|CHDIP1300-12|BOLD:AAH3928
Sciaridae|[15944]|MTDIC283-09|BOLD:AAH3928
Sciaridae|[15945]|CHDIP1105-12|BOLD:AAH3928
Sciaridae|[15946]|CHDIP379-11|BOLD:AAH3928
Sciaridae|[15947]|CHDIP381-11|BOLD:AAH3928
Sciaridae|[15948]|CHDIP1310-12|BOLD:AAH3928
Sciaridae|[15949]|CHDIP1390-12|BOLD:AAH3928
Xylosciara|[15950]|JWDCE770-10|BOLD:AAV1362
Sciaridae|[15951]|JWDCJ1147-11|BOLD:AAP8783
Sciaridae|[15952]|JWDCJ1168-11|BOLD:AAP8783
Sciaridae|[15953]|JWDCJ1328-11|BOLD:AAP8783
Sciaridae|[15954]|JWDCG031-10|BOLD:AAP8783
Sciaridae|[15955]|JWDCC514-10|BOLD:AAL7866
Sciaridae|[15956]|JWDCA773-10|BOLD:AAH4000
Sciaridae|[15957]|JWDCB298-10|BOLD:AAH4000
Sciaridae|[15958]|MTDIC220-09|BOLD:ACZ3939
Camptochaeta consimilis|[15959]|JWDCC751-10|BOLD:AAL7882
Camptochaeta consimilis|[15960]|JWDCF697-10|BOLD:AAL7882
Camptochaeta|[15961]|JWDCA762-10|BOLD:AAH3996
Camptochaeta flagellifera|[15962]|JWDCJ1467-11|BOLD:ACF1472
Camptochaeta flagellifera|[15963]|JWDCH688-10|BOLD:ACF1472
Camptochaeta|[15964]|JWDCH590-10|BOLD:AAP8778
Camptochaeta|[15965]|JWDCJ1466-11|BOLD:AAP8778
Camptochaeta|[15966]|JWDCI019-10|BOLD:AAP8778
Camptochaeta delicata|[15967]|JWDCI067-10|BOLD:AAP8779
Camptochaeta|[15968]|CHDIP490-12|BOLD:ABW9770
Camptochaeta|[15969]|JWDCE595-10|BOLD:AAH4020
Camptochaeta|[15970]|JWDCJ1581-11|BOLD:AAH4020
Camptochaeta|[15971]|JWDCK111-11|BOLD:AAH4020
Camptochaeta|[15972]|JWDCC615-10|BOLD:AAH4020
Camptochaeta|[15973]|JWDCC609-10|BOLD:AAH4020
Camptochaeta|[15974]|JWDCA810-10|BOLD:AAH4020
Camptochaeta|[15975]|JWDCD458-10|BOLD:AAH4020
Camptochaeta|[15976]|JWDCI194-10|BOLD:AAH4020
Camptochaeta|[15977]|JWDCI201-10|BOLD:AAH4020
Camptochaeta|[15978]|CHDIP1408-12|BOLD:AAH4020
Ectrepesthoneura|[15979]|CHDIP365-11|BOLD:ABW5550
Hadroneura|[15980]|JWDCF142-10|BOLD:AAG4984
Hadroneura|[15981]|JWDCE872-10|BOLD:AAG4984
Hadroneura|[15982]|JWDCC767-10|BOLD:AAG4984
Hadroneura|[15983]|JWDCJ654-11|BOLD:AAG4984
Hadroneura|[15984]|JWDCI866-11|BOLD:AAG4984
Hadroneura|[15985]|JWDCA246-10|BOLD:AAG4984
Hadroneura|[15986]|JWDCB175-10|BOLD:AAG4984
Hadroneura|[15987]|JWDCA252-10|BOLD:AAG4984
Hadroneura|[15988]|JWDCB174-10|BOLD:AAG4984
Hadroneura|[15989]|JWDCC771-10|BOLD:AAG4984
Hadroneura|[15990]|JWDCJ839-11|BOLD:AAG4984
Hadroneura|[15991]|JWDCE873-10|BOLD:AAG4984
Chaoborus|[15992]|MMADL073-09|BOLD:AAI4180
Chaoborus|[15993]|MMADL041-09|BOLD:AAI4180
Chaoborus|[15994]|MMADL044-09|BOLD:AAI4180
Chaoborus|[15995]|MMADL046-09|BOLD:AAI4180
Chaoborus|[15996]|MMADL048-09|BOLD:AAI4180
Chaoborus|[15997]|MMADL047-09|BOLD:AAI4180
Chaoborus|[15998]|KSCLC007-09|BOLD:AAI4180
Chaoborus|[15999]|MMADL040-09|BOLD:AAI4180
Chaoborus|[16000]|MMADL042-09|BOLD:AAI4180
Chaoborus|[16001]|MMADL043-09|BOLD:AAI4180
Chaoborus|[16002]|MMADL045-09|BOLD:AAI4180
Chaoborus|[16003]|MMADL062-09|BOLD:AAI4180
Chaoborus|[16004]|MMADL072-09|BOLD:AAI4180
Chaoborus|[16005]|MMADL074-09|BOLD:AAI4180
Chaoborus|[16006]|MMADL087-09|BOLD:AAI4179
Chaoborus|[16007]|CHURC1074-09|BOLD:AAI4179
Chaoborus|[16008]|MMADL086-09|BOLD:AAI4179
Chaoborus|[16009]|MMADL079-09|BOLD:AAI4179
Chaoborus|[16010]|MMADL084-09|BOLD:AAI4179
Chaoborus|[16011]|MHMOS053-07|BOLD:AAI4179
Chaoborus|[16012]|JWDCL036-11|BOLD:AAI4179
Chaoborus|[16013]|JWDCG468-10|BOLD:AAI4179
Chaoborus|[16014]|MHFLY134-07|BOLD:AAI4179
Chaoborus|[16015]|MHCOL293-07|BOLD:AAI4179
Chaoborus|[16016]|MMADL085-09|BOLD:AAI4179
Chaoborus|[16017]|MMADL083-09|BOLD:AAI4179
Chaoborus|[16018]|MMADL081-09|BOLD:AAI4179
Chaoborus|[16019]|MHCOL219-07|BOLD:AAI4179
Chaoborus|[16020]|KSCLC089-09|BOLD:AAI4179
Chaoborus|[16021]|KSCLC006-09|BOLD:AAI4179

[illegible]
Chaoborus|[16019]|MHCOL219-07|BOLD:AAI4179
Chaoborus|[16020]|KSCLC089-09|BOLD:AAI4179
Chaoborus|[16021]|KSCLC006-09|BOLD:AAI4179
Chaoborus|[16022]|CHURC1060-09|BOLD:AAI4179
Chaoborus|[16023]|MMADL080-09|BOLD:AAI4179
Chaoborus|[16024]|MMADL082-09|BOLD:AAI4179
Chaoborus|[16025]|BSCHR153-10|BOLD:AAI4179
Mycomya sp. CJB5|[16026]|JWDCF302-10|BOLD:AAP9081
Mycomya|[16027]|CHDIP871-12|BOLD:AAP9081
Docosia|[16028]|CHDIP477-12|BOLD:AAG4878
Docosia|[16029]|CHDIP458-12|BOLD:AAG4878
Docosia sp. CJB1|[16030]|MTDIC045-09|BOLD:AAG4878
Docosia|[16031]|CHDIP929-12|BOLD:AAG4878
Boletina hyperborea|[16032]|JWDCJ1053-11|BOLD:AAU4902
Boletina sp. CJB02|[16033]|JWDCI199-10|BOLD:AAG4889
Boletina sp. CJB02|[16034]|JWDCK113-11|BOLD:AAG4889
Boletina sp. CJB02|[16035]|JWDCG048-10|BOLD:AAG4889
Boletina sp. CJB02|[16036]|JWDCJ837-11|BOLD:AAG4889
Boletina sp. CJB02|[16037]|JWDCJ838-11|BOLD:AAG4889
Boletina sp. CJB02|[16038]|JWDCC544-10|BOLD:AAG4889
Boletina sp. CJB02|[16039]|JWDCI612-10|BOLD:AAG4889
Boletina sp. CJB02|[16040]|JWDCJ064-11|BOLD:AAG4889
Boletina sp. CJB02|[16041]|JWDCC542-10|BOLD:AAG4889
Boletina sp. CJB02|[16042]|JWDCB176-10|BOLD:AAG4889
Boletina sp. CJB02|[16043]|JWDCF386-10|BOLD:AAG4889
Boletina sp. CJB02|[16044]|JWDCF388-10|BOLD:AAG4889
Boletina sp. CJB02|[16045]|JWDCL393-11|BOLD:AAG4889
Boletina sp. CJB02|[16046]|JWDCJ906-11|BOLD:AAG4889
Boletina sp. CJB02|[16047]|TWDIP309-09|BOLD:AAG4889
Boletina sp. CJB02|[16048]|JWDCD149-10|BOLD:AAG4889
Boletina sp. CJB02|[16049]|JWDCG047-10|BOLD:AAG4889
Boletina sp. CJB02|[16050]|JWDCI869-11|BOLD:AAG4889
Boletina sp. CJB02|[16051]|JWDCG046-10|BOLD:AAG4889
Boletina sp. CJB02|[16052]|JWDCJ1054-11|BOLD:AAG4889
Boletina sp. CJB02|[16053]|JWDCJ981-11|BOLD:AAG4889
Boletina sp. CJB02|[16054]|JWDCB181-10|BOLD:AAG4889
Boletina sp. CJB02|[16055]|JWDCG049-10|BOLD:AAG4889
Boletina sp. CJB02|[16056]|JWDCF387-10|BOLD:AAG4889
Boletina sp. CJB01|[16057]|JWDCC891-10|BOLD:AAG4877
Boletina sp. CJB01|[16058]|JWDCC247-10|BOLD:AAG4877
Boletina sp. CJB01|[16059]|JWDCJ980-11|BOLD:AAG4877
Boletina sp. CJB01|[16060]|MTDIC043-09|BOLD:AAG4877
Boletina|[16061]|CHDIP070-11|BOLD:AAG4877
Boletina sp. CJB01|[16062]|JWDCC246-10|BOLD:AAG4877
Boletina sp. CJB01|[16063]|JWDCA235-10|BOLD:AAG4877
Boletina|[16064]|CHDIP075-11|BOLD:AAG4877
Boletina|[16065]|CHDIP076-11|BOLD:AAG4877
Boletina sp. CJB01|[16066]|JWDCC768-10|BOLD:AAG4877
Boletina sp. CJB01|[16067]|JWDCC888-10|BOLD:AAG4877
Boletina|[16068]|CHDIP071-11|BOLD:AAG4877
Boletina|[16069]|CHDIP091-11|BOLD:AAG4877
Boletina|[16070]|CHDIP069-11|BOLD:AAG4877
Boletina|[16071]|CHDIP081-11|BOLD:AAG4877
Boletina|[16072]|CHDIP074-11|BOLD:AAG4877
Boletina sp. CJB01|[16073]|JWDCJ924-11|BOLD:AAG4877
Boletina sp. CJB01|[16074]|JWDCJ889-11|BOLD:AAG4877
Boletina sp. CJB01|[16075]|JWDCJ836-11|BOLD:AAG4877
Boletina sp. CJB01|[16076]|JWDCC418-10|BOLD:AAG4877
Boletina sp. CJB01|[16077]|JWDCC388-10|BOLD:AAG4877
Boletina sp. CJB01|[16078]|JWDCD189-10|BOLD:AAG4877
Boletina sp. CJB01|[16079]|JWDCD186-10|BOLD:AAG4877
Boletina sp. CJB01|[16080]|JWDCC245-10|BOLD:AAG4877
Boletina sp. CJB01|[16081]|JWDCC244-10|BOLD:AAG4877
Boletina sp. CJB01|[16082]|JWDCC232-10|BOLD:AAG4877
Boletina sp. CJB01|[16083]|JWDCC229-10|BOLD:AAG4877
Boletina sp. CJB01|[16084]|JWDCA245-10|BOLD:AAG4877
Boletina sp. CJB01|[16085]|JWDCA244-10|BOLD:AAG4877
Boletina sp. CJB01|[16086]|JWDCA242-10|BOLD:AAG4877
Boletina sp. CJB01|[16087]|JWDCA240-10|BOLD:AAG4877
Boletina sp. CJB01|[16088]|JWDCA239-10|BOLD:AAG4877
Boletina sp. CJB01|[16089]|JWDCA241-10|BOLD:AAG4877
Boletina sp. CJB01|[16090]|JWDCA238-10|BOLD:AAG4877
Boletina sp. CJB01|[16091]|JWDCA236-10|BOLD:AAG4877
Boletina sp. CJB01|[16092]|JWDCA234-10|BOLD:AAG4877
Boletina sp. CJB01|[16093]|JWDCA233-10|BOLD:AAG4877
Boletina sp. CJB01|[16094]|JWDCA231-10|BOLD:AAG4877
Boletina sp. CJB01|[16095]|JWDCC890-10|BOLD:AAG4877
Boletina sp. CJB01|[16096]|JWDCC062-10|BOLD:AAG4877
Boletina sp. CJB01|[16097]|MTDIC088-09|BOLD:AAG4877
Boletina sp. CJB01|[16098]|JWDCC398-10|BOLD:AAG4877
Boletina sp. CJB01|[16099]|JWDCC773-10|BOLD:AAG4877
Boletina sp. CJB01|[16100]|JWDCA243-10|BOLD:AAG4877
Boletina sp. CJB01|[16101]|JWDCA237-10|BOLD:AAG4877
Boletina sp. CJB01|[16102]|MTDIC075-09|BOLD:AAG4877
Boletina sp. CJB01|[16103]|JWDCC233-10|BOLD:AAG4877
Boletina sp. CJB01|[16104]|JWDCD060-10|BOLD:AAG4877
Boletina sp. CJB01|[16105]|JWDCD008-10|BOLD:AAG4877
Boletina|[16106]|CHDIP078-11|BOLD:AAG4877
Boletina|[16107]|CHDIP073-11|BOLD:AAG4877
Boletina|[16108]|CHDIP786-12|BOLD:AAG4877
Boletina sp. CJB04|[16109]|JWDCJ1052-11|BOLD:AAM8956
Boletina sp. CJB04|[16110]|JWDCA215-10|BOLD:AAM8956
Boletina sp. CJB04|[16111]|JWDCA219-10|BOLD:AAM8956
Boletina sp. CJB04|[16112]|JWDCI753-11|BOLD:AAM8956
Boletina|[16113]|CHDIP469-12|BOLD:AAM8956
Boletina groenlandica|[16114]|JWDCA213-10|BOLD:AAL9142
Boletina groenlandica|[16115]|JWDCD006-10|BOLD:AAL9142
Boletina groenlandica|[16116]|JWDCL394-11|BOLD:AAL9142
Boletina groenlandica|[16117]|CHDIP867-12|BOLD:AAL9142
Coelosia sp. CJB1|[16118]|JWDCI062-10|BOLD:AAP9082
Coelosia gracilis|[16119]|JWDCG428-10|BOLD:AAP9089
Coelosia gracilis|[16120]|JWDCG427-10|BOLD:AAP9089
[illegible]

Coelosia sp. CJB1|[16118]|JWDCI062-10|BOLD:AAP9082
Coelosia gracilis|[16119]|JWDCG428-10|BOLD:AAP9089
Coelosia gracilis|[16120]|JWDCG427-10|BOLD:AAP9089
Coelosia gracilis|[16121]|JWDCG320-10|BOLD:AAP9089
Coelosia gracilis|[16122]|JWDCH886-10|BOLD:AAP9089
Coelosia gracilis|[16123]|JWDCG526-10|BOLD:AAP9089
Coelosia gracilis|[16124]|JWDCI123-10|BOLD:AAM8963
Coelosia gracilis|[16125]|JWDCG406-10|BOLD:AAM8963
Coelosia gracilis|[16126]|JWDCH580-10|BOLD:AAM8963
Coelosia gracilis|[16127]|JWDCI500-10|BOLD:AAM8963
Coelosia gracilis|[16128]|JWDCG408-10|BOLD:AAM8963
Coelosia gracilis|[16129]|CHDIP1216-12|BOLD:AAM8963
Boletina sp. CJB09|[16130]|JWDCL047-11|BOLD:AAP8898
Boletina sp. CJB09|[16131]|JWDCL104-11|BOLD:AAP8898
Boletina sp. CJB09|[16132]|JWDCH739-10|BOLD:AAP8898
Gnoriste sp. CJB1|[16133]|JWDCG648-10|BOLD:ACW2312
Gnoriste|[16134]|CHDIP813-12|BOLD:ACW2312
Boletina digitata|[16135]|JWDCB179-10|BOLD:AAG4874
Boletina digitata|[16136]|JWDCL396-11|BOLD:AAG4874
Boletina digitata|[16137]|JWDCA217-10|BOLD:AAG4874
Boletina digitata|[16138]|MTDIC023-09|BOLD:AAG4874
Boletina digitata|[16139]|JWDCB178-10|BOLD:AAG4874
Boletina digitata|[16140]|MTDIC203-09|BOLD:AAG4874
Boletina digitata|[16141]|TWDIP256-09|BOLD:AAG4874
Boletina digitata|[16142]|CHDIP822-12|BOLD:AAG4874
Boletina brevicornis|[16143]|JWDCD188-10|BOLD:AAL9149
Boletina palmata|[16144]|JWDCJ067-11|BOLD:AAG4924
Boletina palmata|[16145]|JWDCG044-10|BOLD:AAG4924
Boletina|[16146]|JWDCG051-10|BOLD:ABZ2061
Boletina minuta|[16147]|JWDCJ1584-11|BOLD:AAQ3546
Boletina|[16148]|CHDIP495-12|BOLD:AAN8582
Boletina|[16149]|CHDIP1203-12|BOLD:AAN8582
Boletina nitiduloides|[16150]|JWDCJ655-11|BOLD:AAU4924
Boletina populina|[16151]|JWDCL239-11|BOLD:AAU4901
Boletina|[16152]|MTDIC024-09|BOLD:AAG4875
Boletina|[16153]|TWDIP235-09|BOLD:AAG4875
Boletina|[16154]|JWDCA227-10|BOLD:AAM8958
Boletina sp. CJB08|[16155]|JWDCI867-11|BOLD:AAP9953
Boletina sp. CJB08|[16156]|JWDCI629-10|BOLD:AAP9953
Boletina sp. CJB08|[16157]|JWDCI628-10|BOLD:AAP9953
Boletina sp. CJB08|[16158]|JWDCJ1585-11|BOLD:AAP9953
Boletina|[16159]|CHDIP480-12|BOLD:AAP9953
Boletina pinusia|[16160]|JWDCJ379-11|BOLD:ABZ2146
Boletina pinusia|[16161]|CHDIP481-12|BOLD:ABZ2146
Boletina dissipata|[16162]|JWDCH051-10|BOLD:ACZ1770
Boletina dissipata|[16163]|JWDCJ1051-11|BOLD:ACZ1770
Boletina dissipata|[16164]|JWDCL238-11|BOLD:ACZ1770
Boletina dissipata|[16165]|JWDCG322-10|BOLD:ACZ1770
Boletina dissipata|[16166]|JWDCG807-10|BOLD:ACZ1770
Boletina dissipata|[16167]|JWDCH576-10|BOLD:ACZ1770
Boletina dissipata|[16168]|JWDCH684-10|BOLD:ACZ1770
Boletina dissipata|[16169]|JWDCH577-10|BOLD:ACZ1770
Boletina dissipata|[16170]|CHDIP447-12|BOLD:ACZ1770
Boletina dissipata|[16171]|CHDIP498-12|BOLD:ACZ1770
Boletina dissipata|[16172]|CHDIP1198-12|BOLD:ACZ1770
Boletina dissipata|[16173]|JWDCL305-11|BOLD:ACZ1770
Boletina dissipata|[16174]|JWDCH573-10|BOLD:ACZ1770
Boletina dissipata|[16175]|JWDCH054-10|BOLD:ACZ1770
Boletina dissipata|[16176]|JWDCH184-10|BOLD:ACZ1770
Boletina dissipata|[16177]|JWDCA222-10|BOLD:ACZ1770
Boletina dissipata|[16178]|JWDCA223-10|BOLD:ACZ1770
Boletina dissipata|[16179]|JWDCA226-10|BOLD:ACZ1770
Boletina dissipata|[16180]|JWDCH182-10|BOLD:ACZ1770
Boletina dissipata|[16181]|JWDCG246-10|BOLD:ACZ1770
Boletina dissipata|[16182]|JWDCI865-11|BOLD:ACZ1770
Boletina dissipata|[16183]|CHDIP1224-12|BOLD:ACZ1770
Katatopygia sahlbergi|[16184]|CHDIP870-12|BOLD:AAP6486
Syntemna sp. CJB2|[16185]|JWDCC647-10|BOLD:AAL9134
Palaeodocosia|[16186]|JWDCH489-10|BOLD:AAP9086
Palaeodocosia|[16187]|JWDCH956-10|BOLD:AAP9086
Palaeodocosia|[16188]|JWDCI526-10|BOLD:AAP9086
Palaeodocosia|[16189]|JWDCH288-10|BOLD:AAP9086
Palaeodocosia|[16190]|JWDCF902-10|BOLD:AAP9086
Palaeodocosia|[16191]|CHDIP1197-12|BOLD:AAP9086
Katatopygia|[16192]|JWDCE667-10|BOLD:AAP1823
Katatopygia|[16193]|JWDCE668-10|BOLD:AAP1823
Katatopygia|[16194]|JWDCJ473-11|BOLD:AAP1823
Katatopygia|[16195]|JWDCE052-10|BOLD:AAP1823
Katatopygia|[16196]|JWDCE384-10|BOLD:AAP1823
Katatopygia|[16197]|JWDCE138-10|BOLD:AAP1823
Katatopygia|[16198]|CHDIP847-12|BOLD:AAP1823
Katatopygia|[16199]|JWDCE318-10|BOLD:AAP1823
Katatopygia|[16200]|CHDIP848-12|BOLD:AAP1823
Mycetophilidae|[16201]|CHDIP1232-12|BOLD:ACA4481
Docosia gilvipes|[16202]|CHDIP485-12|BOLD:AAG4981
Ectrepesthoneura|[16203]|JWDCC546-10|BOLD:AAL9130
Ectrepesthoneura|[16204]|TWDIP237-09|BOLD:AAL9130
Ectrepesthoneura|[16205]|JWDCG045-10|BOLD:AAP9085
Mycomya maculata|[16206]|JWDCI064-10|BOLD:AAP9083
Mycomya maculata|[16207]|JWDCH955-10|BOLD:AAP9083
Mycomya maculata|[16208]|JWDCG461-10|BOLD:AAP9083
Mycomya maculata|[16209]|JWDCH681-10|BOLD:AAP9083
Mycomya maculata|[16210]|JWDCG557-10|BOLD:AAP9083
Mycomya maculata|[16211]|JWDCG583-10|BOLD:AAP9083
Mycomya maculata|[16212]|JWDCH488-10|BOLD:AAP9083
Mycomya sp. CJB2|[16213]|JWDCH487-10|BOLD:AAP9087
Mycomya sp. CJB4|[16214]|TWDIP310-09|BOLD:AAG4890
Mycomya|[16215]|CHDIP1221-12|BOLD:AAG4890
Mycomya|[16216]|CHDIP831-12|BOLD:ACB1125
Mycomya sp. CJB1|[16217]|JWDCB180-10|BOLD:AAG4986
Mycomya sp. CJB1|[16218]|JWDCH680-10|BOLD:AAG4986
Mycomya sp. CJB1|[16219]|JWDCI122-10|BOLD:AAG4986
Mycomya|[16220]|CHDIP1204-12|BOLD:AAG4986

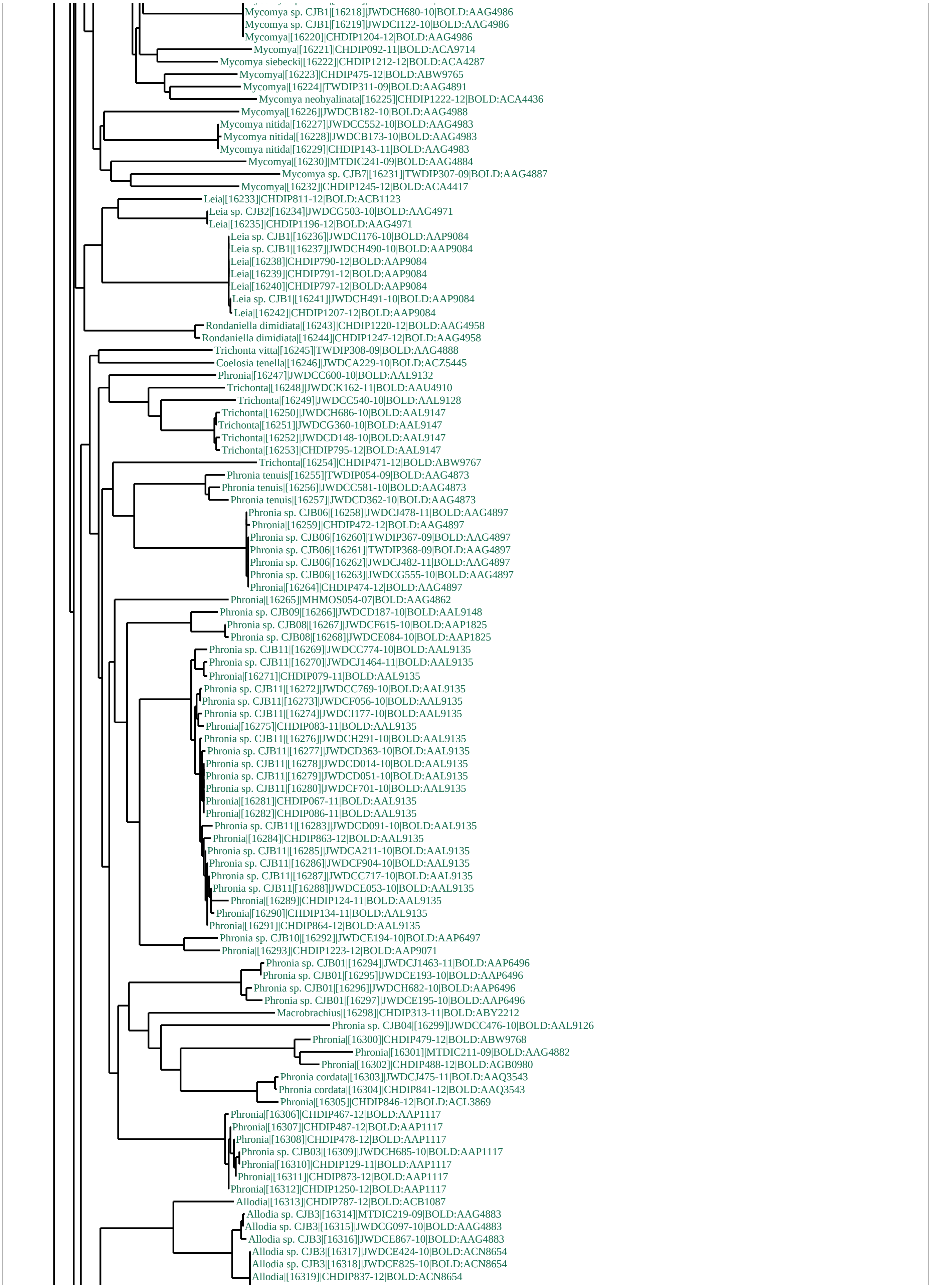
Mycomya sp. CJB1|[16218]|JWDCH680-10|BOLD:AAG4986
Mycomya sp. CJB1|[16219]|JWDCI122-10|BOLD:AAG4986
Mycomya|[16220]|CHDIP1204-12|BOLD:AAG4986
Mycomya|[16221]|CHDIP092-11|BOLD:ACA9714
Mycomya siebecki|[16222]|CHDIP1212-12|BOLD:ACA4287
Mycomya|[16223]|CHDIP475-12|BOLD:ABW9765
Mycomya|[16224]|TWDIP311-09|BOLD:AAG4891
Mycomya neohyalinata|[16225]|CHDIP1222-12|BOLD:ACA4436
Mycomya|[16226]|JWDCB182-10|BOLD:AAG4988
Mycomya nitida|[16227]|JWDCC552-10|BOLD:AAG4983
Mycomya nitida|[16228]|JWDCB173-10|BOLD:AAG4983
Mycomya nitida|[16229]|CHDIP143-11|BOLD:AAG4983
Mycomya|[16230]|MTDIC241-09|BOLD:AAG4884
Mycomya sp. CJB7|[16231]|TWDIP307-09|BOLD:AAG4887
Mycomya|[16232]|CHDIP1245-12|BOLD:ACA4417
Leia|[16233]|CHDIP811-12|BOLD:ACB1123
Leia sp. CJB2|[16234]|JWDCG503-10|BOLD:AAG4971
Leia|[16235]|CHDIP1196-12|BOLD:AAG4971
Leia sp. CJB1|[16236]|JWDCI176-10|BOLD:AAP9084
Leia sp. CJB1|[16237]|JWDCH490-10|BOLD:AAP9084
Leia|[16238]|CHDIP790-12|BOLD:AAP9084
Leia|[16239]|CHDIP791-12|BOLD:AAP9084
Leia|[16240]|CHDIP797-12|BOLD:AAP9084
Leia sp. CJB1|[16241]|JWDCH491-10|BOLD:AAP9084
Leia|[16242]|CHDIP1207-12|BOLD:AAP9084
Rondaniella dimidiata|[16243]|CHDIP1220-12|BOLD:AAG4958
Rondaniella dimidiata|[16244]|CHDIP1247-12|BOLD:AAG4958
Trichonta vitta|[16245]|TWDIP308-09|BOLD:AAG4888
Coelosia tenella|[16246]|JWDCA229-10|BOLD:ACZ5445
Phronia|[16247]|JWDCC600-10|BOLD:AAL9132
Trichonta|[16248]|JWDCK162-11|BOLD:AAU4910
Trichonta|[16249]|JWDCC540-10|BOLD:AAL9128
Trichonta|[16250]|JWDCH686-10|BOLD:AAL9147
Trichonta|[16251]|JWDCG360-10|BOLD:AAL9147
Trichonta|[16252]|JWDCD148-10|BOLD:AAL9147
Trichonta|[16253]|CHDIP795-12|BOLD:AAL9147
Trichonta|[16254]|CHDIP471-12|BOLD:ABW9767
Phronia tenuis|[16255]|TWDIP054-09|BOLD:AAG4873
Phronia tenuis|[16256]|JWDCC581-10|BOLD:AAG4873
Phronia tenuis|[16257]|JWDCD362-10|BOLD:AAG4873
Phronia sp. CJB06|[16258]|JWDCJ478-11|BOLD:AAG4897
Phronia|[16259]|CHDIP472-12|BOLD:AAG4897
Phronia sp. CJB06|[16260]|TWDIP367-09|BOLD:AAG4897
Phronia sp. CJB06|[16261]|TWDIP368-09|BOLD:AAG4897
Phronia sp. CJB06|[16262]|JWDCJ482-11|BOLD:AAG4897
Phronia sp. CJB06|[16263]|JWDCG555-10|BOLD:AAG4897
Phronia|[16264]|CHDIP474-12|BOLD:AAG4897
Phronia|[16265]|MHMOS054-07|BOLD:AAG4862
Phronia sp. CJB09|[16266]|JWDCD187-10|BOLD:AAL9148
Phronia sp. CJB08|[16267]|JWDCF615-10|BOLD:AAP1825
Phronia sp. CJB08|[16268]|JWDCE084-10|BOLD:AAP1825
Phronia sp. CJB11|[16269]|JWDCC774-10|BOLD:AAL9135
Phronia sp. CJB11|[16270]|JWDCJ1464-11|BOLD:AAL9135
Phronia|[16271]|CHDIP079-11|BOLD:AAL9135
Phronia sp. CJB11|[16272]|JWDCC769-10|BOLD:AAL9135
Phronia sp. CJB11|[16273]|JWDCF056-10|BOLD:AAL9135
Phronia sp. CJB11|[16274]|JWDCI177-10|BOLD:AAL9135
Phronia|[16275]|CHDIP083-11|BOLD:AAL9135
Phronia sp. CJB11|[16276]|JWDCH291-10|BOLD:AAL9135
Phronia sp. CJB11|[16277]|JWDCD363-10|BOLD:AAL9135
Phronia sp. CJB11|[16278]|JWDCD014-10|BOLD:AAL9135
Phronia sp. CJB11|[16279]|JWDCD051-10|BOLD:AAL9135
Phronia sp. CJB11|[16280]|JWDCF701-10|BOLD:AAL9135
Phronia|[16281]|CHDIP067-11|BOLD:AAL9135
Phronia|[16282]|CHDIP086-11|BOLD:AAL9135
Phronia sp. CJB11|[16283]|JWDCD091-10|BOLD:AAL9135
Phronia|[16284]|CHDIP863-12|BOLD:AAL9135
Phronia sp. CJB11|[16285]|JWDCA211-10|BOLD:AAL9135
Phronia sp. CJB11|[16286]|JWDCF904-10|BOLD:AAL9135
Phronia sp. CJB11|[16287]|JWDCC717-10|BOLD:AAL9135
Phronia sp. CJB11|[16288]|JWDCE053-10|BOLD:AAL9135
Phronia|[16289]|CHDIP124-11|BOLD:AAL9135
Phronia|[16290]|CHDIP134-11|BOLD:AAL9135
Phronia|[16291]|CHDIP864-12|BOLD:AAL9135
Phronia sp. CJB10|[16292]|JWDCE194-10|BOLD:AAP6497
Phronia|[16293]|CHDIP1223-12|BOLD:AAP9071
Phronia sp. CJB01|[16294]|JWDCJ1463-11|BOLD:AAP6496
Phronia sp. CJB01|[16295]|JWDCE193-10|BOLD:AAP6496
Phronia sp. CJB01|[16296]|JWDCH682-10|BOLD:AAP6496
Phronia sp. CJB01|[16297]|JWDCE195-10|BOLD:AAP6496
Macrobrachius|[16298]|CHDIP313-11|BOLD:ABY2212
Phronia sp. CJB04|[16299]|JWDCC476-10|BOLD:AAL9126
Phronia|[16300]|CHDIP479-12|BOLD:ABW9768
Phronia|[16301]|MTDIC211-09|BOLD:AAG4882
Phronia|[16302]|CHDIP488-12|BOLD:AGB0980
Phronia cordata|[16303]|JWDCJ475-11|BOLD:AAQ3543
Phronia cordata|[16304]|CHDIP841-12|BOLD:AAQ3543
Phronia|[16305]|CHDIP846-12|BOLD:ACL3869
Phronia|[16306]|CHDIP467-12|BOLD:AAP1117
Phronia|[16307]|CHDIP487-12|BOLD:AAP1117
Phronia|[16308]|CHDIP478-12|BOLD:AAP1117
Phronia sp. CJB03|[16309]|JWDCH685-10|BOLD:AAP1117
Phronia|[16310]|CHDIP129-11|BOLD:AAP1117
Phronia|[16311]|CHDIP873-12|BOLD:AAP1117
Phronia|[16312]|CHDIP1250-12|BOLD:AAP1117
Allodia|[16313]|CHDIP787-12|BOLD:ACB1087
Allodia sp. CJB3|[16314]|MTDIC219-09|BOLD:AAG4883
Allodia sp. CJB3|[16315]|JWDCG097-10|BOLD:AAG4883
Allodia sp. CJB3|[16316]|JWDCE867-10|BOLD:AAG4883
Allodia sp. CJB3|[16317]|JWDCE424-10|BOLD:ACN8654
Allodia sp. CJB3|[16318]|JWDCE825-10|BOLD:ACN8654
Allodia|[16319]|CHDIP837-12|BOLD:ACN8654

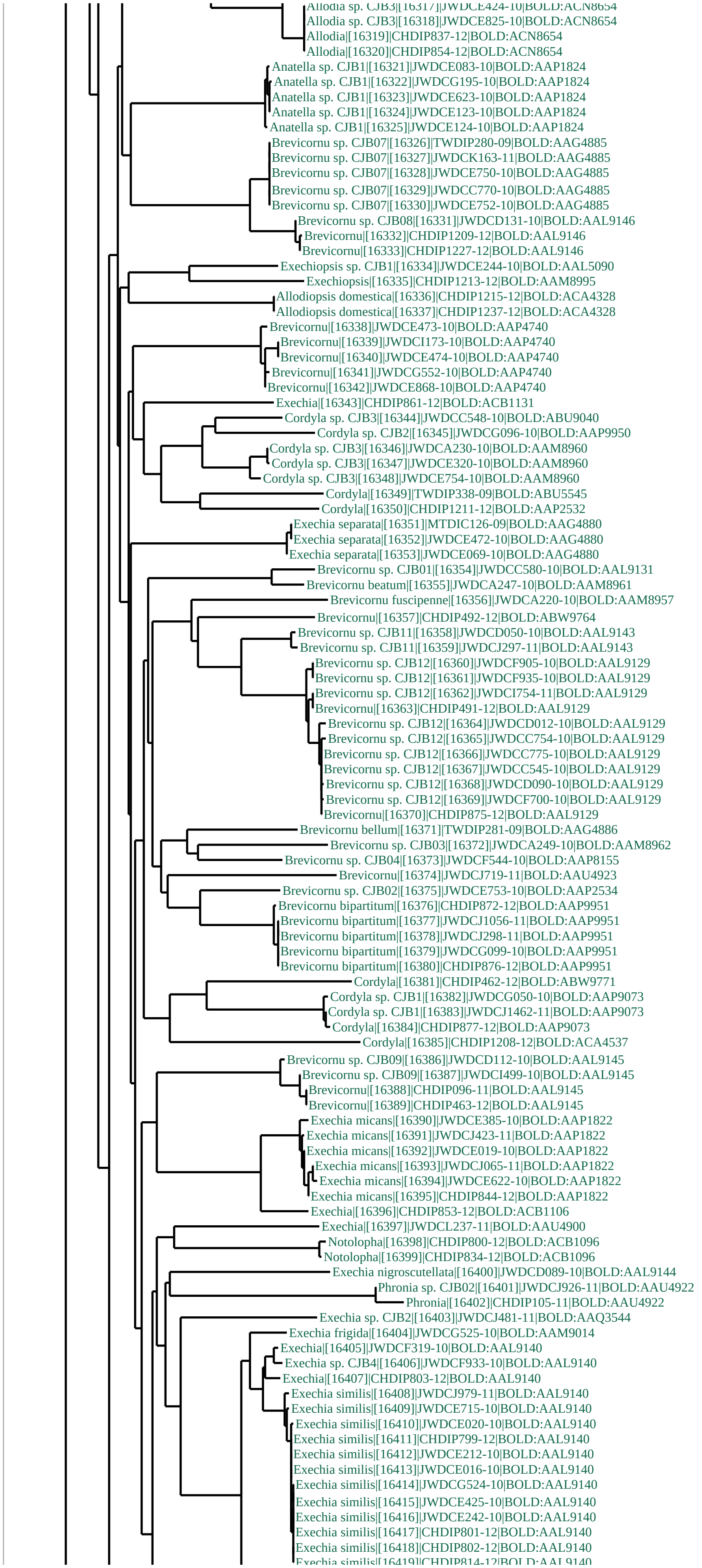
Allodia sp. CJB3|[16317]|JWDCE424-10|BOLD:ACN8654
Allodia sp. CJB3|[16318]|JWDCE825-10|BOLD:ACN8654
Allodia|[16319]|CHDIP837-12|BOLD:ACN8654
Allodia|[16320]|CHDIP854-12|BOLD:ACN8654
Anatella sp. CJB1|[16321]|JWDCE083-10|BOLD:AAP1824
Anatella sp. CJB1|[16322]|JWDCG195-10|BOLD:AAP1824
Anatella sp. CJB1|[16323]|JWDCE623-10|BOLD:AAP1824
Anatella sp. CJB1|[16324]|JWDCE123-10|BOLD:AAP1824
Anatella sp. CJB1|[16325]|JWDCE124-10|BOLD:AAP1824
Brevicornu sp. CJB07|[16326]|TWDIP280-09|BOLD:AAG4885
Brevicornu sp. CJB07|[16327]|JWDCK163-11|BOLD:AAG4885
Brevicornu sp. CJB07|[16328]|JWDCE750-10|BOLD:AAG4885
Brevicornu sp. CJB07|[16329]|JWDCC770-10|BOLD:AAG4885
Brevicornu sp. CJB07|[16330]|JWDCE752-10|BOLD:AAG4885
Brevicornu sp. CJB08|[16331]|JWDCD131-10|BOLD:AAL9146
Brevicornu|[16332]|CHDIP1209-12|BOLD:AAL9146
Brevicornu|[16333]|CHDIP1227-12|BOLD:AAL9146
Exechiopsis sp. CJB1|[16334]|JWDCE244-10|BOLD:AAL5090
Exechiopsis|[16335]|CHDIP1213-12|BOLD:AAM8995
Allodiopsis domestica|[16336]|CHDIP1215-12|BOLD:ACA4328
Allodiopsis domestica|[16337]|CHDIP1237-12|BOLD:ACA4328
Brevicornu|[16338]|JWDCE473-10|BOLD:AAP4740
Brevicornu|[16339]|JWDCI173-10|BOLD:AAP4740
Brevicornu|[16340]|JWDCE474-10|BOLD:AAP4740
Brevicornu|[16341]|JWDCG552-10|BOLD:AAP4740
Brevicornu|[16342]|JWDCE868-10|BOLD:AAP4740
Exechia|[16343]|CHDIP861-12|BOLD:ACB1131
Cordyla sp. CJB3|[16344]|JWDCC548-10|BOLD:ABU9040
Cordyla sp. CJB2|[16345]|JWDCG096-10|BOLD:AAP9950
Cordyla sp. CJB3|[16346]|JWDCA230-10|BOLD:AAM8960
Cordyla sp. CJB3|[16347]|JWDCE320-10|BOLD:AAM8960
Cordyla sp. CJB3|[16348]|JWDCE754-10|BOLD:AAM8960
Cordyla|[16349]|TWDIP338-09|BOLD:ABU5545
Cordyla|[16350]|CHDIP1211-12|BOLD:AAP2532
Exechia separata|[16351]|MTDIC126-09|BOLD:AAG4880
Exechia separata|[16352]|JWDCE472-10|BOLD:AAG4880
Exechia separata|[16353]|JWDCE069-10|BOLD:AAG4880
Brevicornu sp. CJB01|[16354]|JWDCC580-10|BOLD:AAL9131
Brevicornu beatum|[16355]|JWDCA247-10|BOLD:AAM8961
Brevicornu fuscipenne|[16356]|JWDCA220-10|BOLD:AAM8957
Brevicornu|[16357]|CHDIP492-12|BOLD:ABW9764
Brevicornu sp. CJB11|[16358]|JWDCD050-10|BOLD:AAL9143
Brevicornu sp. CJB11|[16359]|JWDCJ297-11|BOLD:AAL9143
Brevicornu sp. CJB12|[16360]|JWDCF905-10|BOLD:AAL9129
Brevicornu sp. CJB12|[16361]|JWDCF935-10|BOLD:AAL9129
Brevicornu sp. CJB12|[16362]|JWDCI754-11|BOLD:AAL9129
Brevicornu|[16363]|CHDIP491-12|BOLD:AAL9129
Brevicornu sp. CJB12|[16364]|JWDCD012-10|BOLD:AAL9129
Brevicornu sp. CJB12|[16365]|JWDCC754-10|BOLD:AAL9129
Brevicornu sp. CJB12|[16366]|JWDCC775-10|BOLD:AAL9129
Brevicornu sp. CJB12|[16367]|JWDCC545-10|BOLD:AAL9129
Brevicornu sp. CJB12|[16368]|JWDCD090-10|BOLD:AAL9129
Brevicornu sp. CJB12|[16369]|JWDCF700-10|BOLD:AAL9129
Brevicornu|[16370]|CHDIP875-12|BOLD:AAL9129
Brevicornu bellum|[16371]|TWDIP281-09|BOLD:AAG4886
Brevicornu sp. CJB03|[16372]|JWDCA249-10|BOLD:AAM8962
Brevicornu sp. CJB04|[16373]|JWDCF544-10|BOLD:AAP8155
Brevicornu|[16374]|JWDCJ719-11|BOLD:AAU4923
Brevicornu sp. CJB02|[16375]|JWDCE753-10|BOLD:AAP2534
Brevicornu bipartitum|[16376]|CHDIP872-12|BOLD:AAP9951
Brevicornu bipartitum|[16377]|JWDCJ1056-11|BOLD:AAP9951
Brevicornu bipartitum|[16378]|JWDCJ298-11|BOLD:AAP9951
Brevicornu bipartitum|[16379]|JWDCG099-10|BOLD:AAP9951
Brevicornu bipartitum|[16380]|CHDIP876-12|BOLD:AAP9951
Cordyla|[16381]|CHDIP462-12|BOLD:ABW9771
Cordyla sp. CJB1|[16382]|JWDCG050-10|BOLD:AAP9073
Cordyla sp. CJB1|[16383]|JWDCJ1462-11|BOLD:AAP9073
Cordyla|[16384]|CHDIP877-12|BOLD:AAP9073
Cordyla|[16385]|CHDIP1208-12|BOLD:ACA4537
Brevicornu sp. CJB09|[16386]|JWDCD112-10|BOLD:AAL9145
Brevicornu sp. CJB09|[16387]|JWDCI499-10|BOLD:AAL9145
Brevicornu|[16388]|CHDIP096-11|BOLD:AAL9145
Brevicornu|[16389]|CHDIP463-12|BOLD:AAL9145
Exechia micans|[16390]|JWDCE385-10|BOLD:AAP1822
Exechia micans|[16391]|JWDCJ423-11|BOLD:AAP1822
Exechia micans|[16392]|JWDCE019-10|BOLD:AAP1822
Exechia micans|[16393]|JWDCJ065-11|BOLD:AAP1822
Exechia micans|[16394]|JWDCE622-10|BOLD:AAP1822
Exechia micans|[16395]|CHDIP844-12|BOLD:AAP1822
Exechia|[16396]|CHDIP853-12|BOLD:ACB1106
Exechia|[16397]|JWDCL237-11|BOLD:AAU4900
Notolopha|[16398]|CHDIP800-12|BOLD:ACB1096
Notolopha|[16399]|CHDIP834-12|BOLD:ACB1096
Exechia nigroscutellata|[16400]|JWDCD089-10|BOLD:AAL9144
Phronia sp. CJB02|[16401]|JWDCJ926-11|BOLD:AAU4922
Phronia|[16402]|CHDIP105-11|BOLD:AAU4922
Exechia sp. CJB2|[16403]|JWDCJ481-11|BOLD:AAQ3544
Exechia frigida|[16404]|JWDCG525-10|BOLD:AAM9014
Exechia|[16405]|JWDCF319-10|BOLD:AAL9140
Exechia sp. CJB4|[16406]|JWDCF933-10|BOLD:AAL9140
Exechia|[16407]|CHDIP803-12|BOLD:AAL9140
Exechia similis|[16408]|JWDCJ979-11|BOLD:AAL9140
Exechia similis|[16409]|JWDCE715-10|BOLD:AAL9140
Exechia similis|[16410]|JWDCE020-10|BOLD:AAL9140
Exechia similis|[16411]|CHDIP799-12|BOLD:AAL9140
Exechia similis|[16412]|JWDCE212-10|BOLD:AAL9140
Exechia similis|[16413]|JWDCE016-10|BOLD:AAL9140
Exechia similis|[16414]|JWDCG524-10|BOLD:AAL9140
Exechia similis|[16415]|JWDCE425-10|BOLD:AAL9140
Exechia similis|[16416]|JWDCE242-10|BOLD:AAL9140
Exechia similis|[16417]|CHDIP801-12|BOLD:AAL9140
Exechia similis|[16418]|CHDIP802-12|BOLD:AAL9140
Exechia similis|[16419]|CHDIP814-12|BOLD:AAL9140

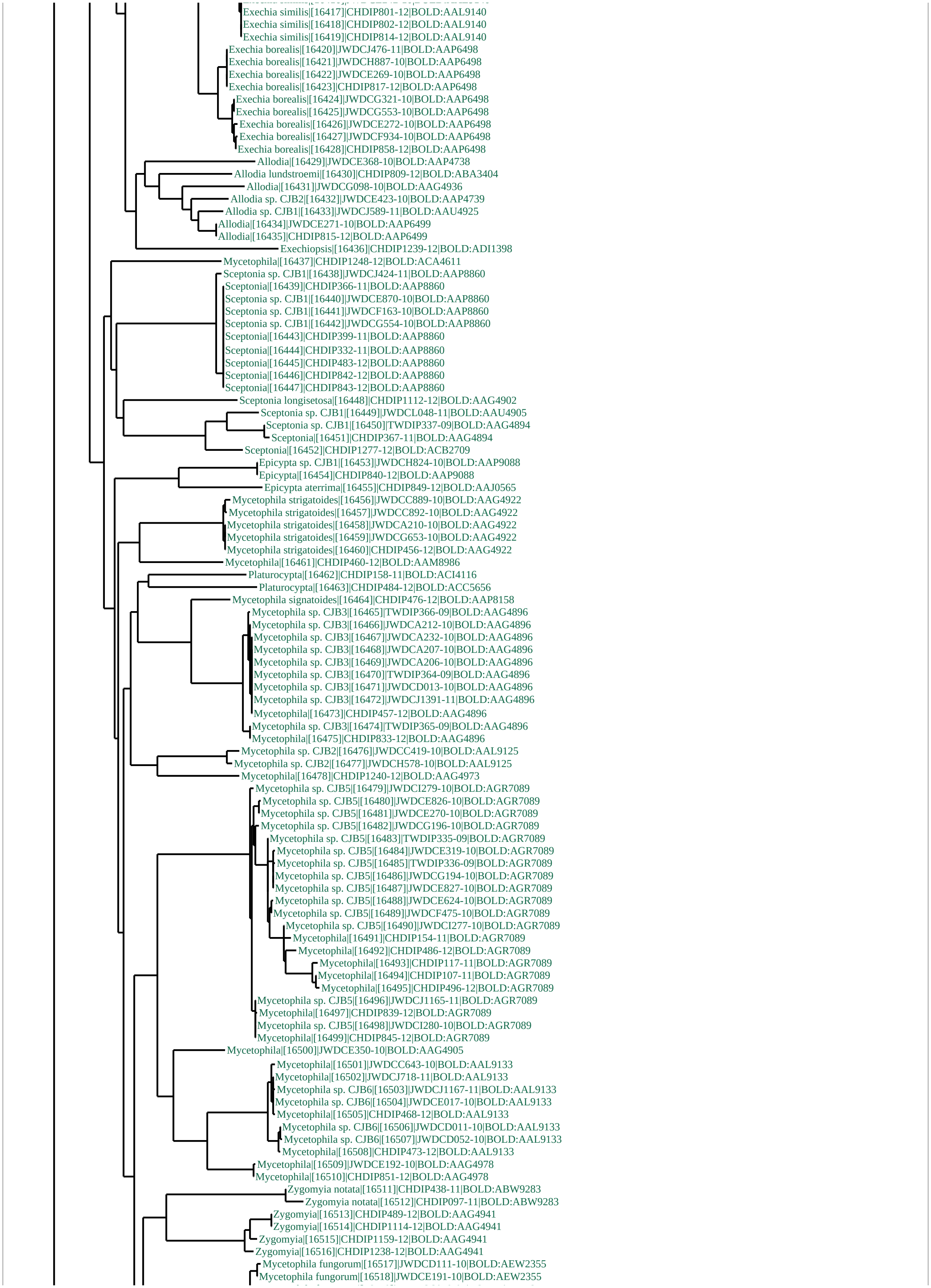
Exechia similis|[16417]|CHDIP801-12|BOLD:AAL9140
Exechia similis|[16418]|CHDIP802-12|BOLD:AAL9140
Exechia similis|[16419]|CHDIP814-12|BOLD:AAL9140
Exechia borealis|[16420]|JWDCJ476-11|BOLD:AAP6498
Exechia borealis|[16421]|JWDCH887-10|BOLD:AAP6498
Exechia borealis|[16422]|JWDCE269-10|BOLD:AAP6498
Exechia borealis|[16423]|CHDIP817-12|BOLD:AAP6498
Exechia borealis|[16424]|JWDCG321-10|BOLD:AAP6498
Exechia borealis|[16425]|JWDCG553-10|BOLD:AAP6498
Exechia borealis|[16426]|JWDCE272-10|BOLD:AAP6498
Exechia borealis|[16427]|JWDCF934-10|BOLD:AAP6498
Exechia borealis|[16428]|CHDIP858-12|BOLD:AAP6498
Allodia|[16429]|JWDCE368-10|BOLD:AAP4738
Allodia lundstroemi|[16430]|CHDIP809-12|BOLD:ABA3404
Allodia|[16431]|JWDCG098-10|BOLD:AAG4936
Allodia sp. CJB2|[16432]|JWDCE423-10|BOLD:AAP4739
Allodia sp. CJB1|[16433]|JWDCJ589-11|BOLD:AAU4925
Allodia|[16434]|JWDCE271-10|BOLD:AAP6499
Allodia|[16435]|CHDIP815-12|BOLD:AAP6499
Exechiopsis|[16436]|CHDIP1239-12|BOLD:ADI1398
Mycetophila|[16437]|CHDIP1248-12|BOLD:ACA4611
Sceptonia sp. CJB1|[16438]|JWDCJ424-11|BOLD:AAP8860
Sceptonia|[16439]|CHDIP366-11|BOLD:AAP8860
Sceptonia sp. CJB1|[16440]|JWDCE870-10|BOLD:AAP8860
Sceptonia sp. CJB1|[16441]|JWDCF163-10|BOLD:AAP8860
Sceptonia sp. CJB1|[16442]|JWDCG554-10|BOLD:AAP8860
Sceptonia|[16443]|CHDIP399-11|BOLD:AAP8860
Sceptonia|[16444]|CHDIP332-11|BOLD:AAP8860
Sceptonia|[16445]|CHDIP483-12|BOLD:AAP8860
Sceptonia|[16446]|CHDIP842-12|BOLD:AAP8860
Sceptonia|[16447]|CHDIP843-12|BOLD:AAP8860
Sceptonia longisetosa|[16448]|CHDIP1112-12|BOLD:AAG4902
Sceptonia sp. CJB1|[16449]|JWDCL048-11|BOLD:AAU4905
Sceptonia sp. CJB1|[16450]|TWDIP337-09|BOLD:AAG4894
Sceptonia|[16451]|CHDIP367-11|BOLD:AAG4894
Sceptonia|[16452]|CHDIP1277-12|BOLD:ACB2709
Epicypta sp. CJB1|[16453]|JWDCH824-10|BOLD:AAP9088
Epicypta|[16454]|CHDIP840-12|BOLD:AAP9088
Epicypta aterrima|[16455]|CHDIP849-12|BOLD:AAJ0565
Mycetophila strigatoides|[16456]|JWDCC889-10|BOLD:AAG4922
Mycetophila strigatoides|[16457]|JWDCC892-10|BOLD:AAG4922
Mycetophila strigatoides|[16458]|JWDCA210-10|BOLD:AAG4922
Mycetophila strigatoides|[16459]|JWDCG653-10|BOLD:AAG4922
Mycetophila strigatoides|[16460]|CHDIP456-12|BOLD:AAG4922
Mycetophila|[16461]|CHDIP460-12|BOLD:AAM8986
Platurocypta|[16462]|CHDIP158-11|BOLD:ACI4116
Platurocypta|[16463]|CHDIP484-12|BOLD:ACC5656
Mycetophila signatoides|[16464]|CHDIP476-12|BOLD:AAP8158
Mycetophila sp. CJB3|[16465]|TWDIP366-09|BOLD:AAG4896
Mycetophila sp. CJB3|[16466]|JWDCA212-10|BOLD:AAG4896
Mycetophila sp. CJB3|[16467]|JWDCA232-10|BOLD:AAG4896
Mycetophila sp. CJB3|[16468]|JWDCA207-10|BOLD:AAG4896
Mycetophila sp. CJB3|[16469]|JWDCA206-10|BOLD:AAG4896
Mycetophila sp. CJB3|[16470]|TWDIP364-09|BOLD:AAG4896
Mycetophila sp. CJB3|[16471]|JWDCD013-10|BOLD:AAG4896
Mycetophila sp. CJB3|[16472]|JWDCJ1391-11|BOLD:AAG4896
Mycetophila|[16473]|CHDIP457-12|BOLD:AAG4896
Mycetophila sp. CJB3|[16474]|TWDIP365-09|BOLD:AAG4896
Mycetophila|[16475]|CHDIP833-12|BOLD:AAG4896
Mycetophila sp. CJB2|[16476]|JWDCC419-10|BOLD:AAL9125
Mycetophila sp. CJB2|[16477]|JWDCH578-10|BOLD:AAL9125
Mycetophila|[16478]|CHDIP1240-12|BOLD:AAG4973
Mycetophila sp. CJB5|[16479]|JWDCI279-10|BOLD:AGR7089
Mycetophila sp. CJB5|[16480]|JWDCE826-10|BOLD:AGR7089
Mycetophila sp. CJB5|[16481]|JWDCE270-10|BOLD:AGR7089
Mycetophila sp. CJB5|[16482]|JWDCG196-10|BOLD:AGR7089
Mycetophila sp. CJB5|[16483]|TWDIP335-09|BOLD:AGR7089
Mycetophila sp. CJB5|[16484]|JWDCE319-10|BOLD:AGR7089
Mycetophila sp. CJB5|[16485]|TWDIP336-09|BOLD:AGR7089
Mycetophila sp. CJB5|[16486]|JWDCG194-10|BOLD:AGR7089
Mycetophila sp. CJB5|[16487]|JWDCE827-10|BOLD:AGR7089
Mycetophila sp. CJB5|[16488]|JWDCE624-10|BOLD:AGR7089
Mycetophila sp. CJB5|[16489]|JWDCF475-10|BOLD:AGR7089
Mycetophila sp. CJB5|[16490]|JWDCI277-10|BOLD:AGR7089
Mycetophila|[16491]|CHDIP154-11|BOLD:AGR7089
Mycetophila|[16492]|CHDIP486-12|BOLD:AGR7089
Mycetophila|[16493]|CHDIP117-11|BOLD:AGR7089
Mycetophila|[16494]|CHDIP107-11|BOLD:AGR7089
Mycetophila|[16495]|CHDIP496-12|BOLD:AGR7089
Mycetophila sp. CJB5|[16496]|JWDCJ1165-11|BOLD:AGR7089
Mycetophila|[16497]|CHDIP839-12|BOLD:AGR7089
Mycetophila sp. CJB5|[16498]|JWDCI280-10|BOLD:AGR7089
Mycetophila|[16499]|CHDIP845-12|BOLD:AGR7089
Mycetophila|[16500]|JWDCE350-10|BOLD:AAG4905
Mycetophila|[16501]|JWDCC643-10|BOLD:AAL9133
Mycetophila|[16502]|JWDCJ718-11|BOLD:AAL9133
Mycetophila sp. CJB6|[16503]|JWDCJ1167-11|BOLD:AAL9133
Mycetophila sp. CJB6|[16504]|JWDCE017-10|BOLD:AAL9133
Mycetophila|[16505]|CHDIP468-12|BOLD:AAL9133
Mycetophila sp. CJB6|[16506]|JWDCD011-10|BOLD:AAL9133
Mycetophila sp. CJB6|[16507]|JWDCD052-10|BOLD:AAL9133
Mycetophila|[16508]|CHDIP473-12|BOLD:AAL9133
Mycetophila|[16509]|JWDCE192-10|BOLD:AAG4978
Mycetophila|[16510]|CHDIP851-12|BOLD:AAG4978
Zygomyia notata|[16511]|CHDIP438-11|BOLD:ABW9283
Zygomyia notata|[16512]|CHDIP097-11|BOLD:ABW9283
Zygomyia|[16513]|CHDIP489-12|BOLD:AAG4941
Zygomyia|[16514]|CHDIP1114-12|BOLD:AAG4941
Zygomyia|[16515]|CHDIP1159-12|BOLD:AAG4941
Zygomyia|[16516]|CHDIP1238-12|BOLD:AAG4941
Mycetophila fungorum|[16517]|JWDCD111-10|BOLD:AEW2355
Mycetophila fungorum|[16518]|JWDCE191-10|BOLD:AEW2355

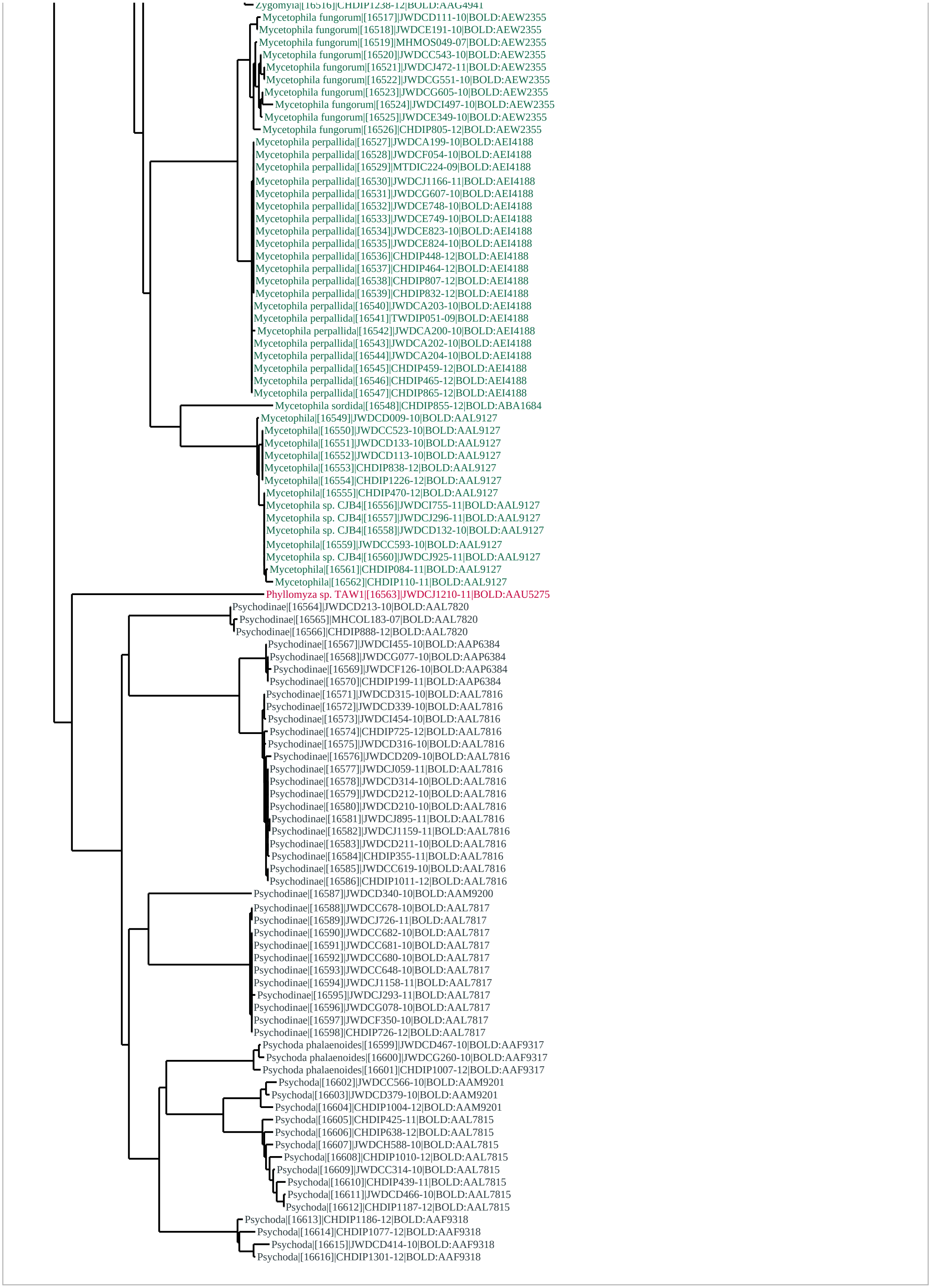
Zygomyia|[16516]|CHDIP1238-12|BOLD:AAG4941
Mycetophila fungorum|[16517]|JWDCD111-10|BOLD:AEW2355
Mycetophila fungorum|[16518]|JWDCE191-10|BOLD:AEW2355
Mycetophila fungorum|[16519]|MHMOS049-07|BOLD:AEW2355
Mycetophila fungorum|[16520]|JWDCC543-10|BOLD:AEW2355
Mycetophila fungorum|[16521]|JWDCJ472-11|BOLD:AEW2355
Mycetophila fungorum|[16522]|JWDCG551-10|BOLD:AEW2355
Mycetophila fungorum|[16523]|JWDCG605-10|BOLD:AEW2355
Mycetophila fungorum|[16524]|JWDCI497-10|BOLD:AEW2355
Mycetophila fungorum|[16525]|JWDCE349-10|BOLD:AEW2355
Mycetophila fungorum|[16526]|CHDIP805-12|BOLD:AEW2355
Mycetophila perpallida|[16527]|JWDCA199-10|BOLD:AEI4188
Mycetophila perpallida|[16528]|JWDCF054-10|BOLD:AEI4188
Mycetophila perpallida|[16529]|MTDIC224-09|BOLD:AEI4188
Mycetophila perpallida|[16530]|JWDCJ1166-11|BOLD:AEI4188
Mycetophila perpallida|[16531]|JWDCG607-10|BOLD:AEI4188
Mycetophila perpallida|[16532]|JWDCE748-10|BOLD:AEI4188
Mycetophila perpallida|[16533]|JWDCE749-10|BOLD:AEI4188
Mycetophila perpallida|[16534]|JWDCE823-10|BOLD:AEI4188
Mycetophila perpallida|[16535]|JWDCE824-10|BOLD:AEI4188
Mycetophila perpallida|[16536]|CHDIP448-12|BOLD:AEI4188
Mycetophila perpallida|[16537]|CHDIP464-12|BOLD:AEI4188
Mycetophila perpallida|[16538]|CHDIP807-12|BOLD:AEI4188
Mycetophila perpallida|[16539]|CHDIP832-12|BOLD:AEI4188
Mycetophila perpallida|[16540]|JWDCA203-10|BOLD:AEI4188
Mycetophila perpallida|[16541]|TWDIP051-09|BOLD:AEI4188
Mycetophila perpallida|[16542]|JWDCA200-10|BOLD:AEI4188
Mycetophila perpallida|[16543]|JWDCA202-10|BOLD:AEI4188
Mycetophila perpallida|[16544]|JWDCA204-10|BOLD:AEI4188
Mycetophila perpallida|[16545]|CHDIP459-12|BOLD:AEI4188
Mycetophila perpallida|[16546]|CHDIP465-12|BOLD:AEI4188
Mycetophila perpallida|[16547]|CHDIP865-12|BOLD:AEI4188
Mycetophila sordida|[16548]|CHDIP855-12|BOLD:ABA1684
Mycetophila|[16549]|JWDCD009-10|BOLD:AAL9127
Mycetophila|[16550]|JWDCC523-10|BOLD:AAL9127
Mycetophila|[16551]|JWDCD133-10|BOLD:AAL9127
Mycetophila|[16552]|JWDCD113-10|BOLD:AAL9127
Mycetophila|[16553]|CHDIP838-12|BOLD:AAL9127
Mycetophila|[16554]|CHDIP1226-12|BOLD:AAL9127
Mycetophila|[16555]|CHDIP470-12|BOLD:AAL9127
Mycetophila sp. CJB4|[16556]|JWDCI755-11|BOLD:AAL9127
Mycetophila sp. CJB4|[16557]|JWDCJ296-11|BOLD:AAL9127
Mycetophila sp. CJB4|[16558]|JWDCD132-10|BOLD:AAL9127
Mycetophila|[16559]|JWDCC593-10|BOLD:AAL9127
Mycetophila sp. CJB4|[16560]|JWDCJ925-11|BOLD:AAL9127
Mycetophila|[16561]|CHDIP084-11|BOLD:AAL9127
Mycetophila|[16562]|CHDIP110-11|BOLD:AAL9127
Phyllomyza sp. TAW1|[16563]|JWDCJ1210-11|BOLD:AAU5275
Psychodinae|[16564]|JWDCD213-10|BOLD:AAL7820
Psychodinae|[16565]|MHCOL183-07|BOLD:AAL7820
Psychodinae|[16566]|CHDIP888-12|BOLD:AAL7820
Psychodinae|[16567]|JWDCI455-10|BOLD:AAP6384
Psychodinae|[16568]|JWDCG077-10|BOLD:AAP6384
Psychodinae|[16569]|JWDCF126-10|BOLD:AAP6384
Psychodinae|[16570]|CHDIP199-11|BOLD:AAP6384
Psychodinae|[16571]|JWDCD315-10|BOLD:AAL7816
Psychodinae|[16572]|JWDCD339-10|BOLD:AAL7816
Psychodinae|[16573]|JWDCI454-10|BOLD:AAL7816
Psychodinae|[16574]|CHDIP725-12|BOLD:AAL7816
Psychodinae|[16575]|JWDCD316-10|BOLD:AAL7816
Psychodinae|[16576]|JWDCD209-10|BOLD:AAL7816
Psychodinae|[16577]|JWDCJ059-11|BOLD:AAL7816
Psychodinae|[16578]|JWDCD314-10|BOLD:AAL7816
Psychodinae|[16579]|JWDCD212-10|BOLD:AAL7816
Psychodinae|[16580]|JWDCD210-10|BOLD:AAL7816
Psychodinae|[16581]|JWDCJ895-11|BOLD:AAL7816
Psychodinae|[16582]|JWDCJ1159-11|BOLD:AAL7816
Psychodinae|[16583]|JWDCD211-10|BOLD:AAL7816
Psychodinae|[16584]|CHDIP355-11|BOLD:AAL7816
Psychodinae|[16585]|JWDCC619-10|BOLD:AAL7816
Psychodinae|[16586]|CHDIP1011-12|BOLD:AAL7816
Psychodinae|[16587]|JWDCD340-10|BOLD:AAM9200
Psychodinae|[16588]|JWDCC678-10|BOLD:AAL7817
Psychodinae|[16589]|JWDCJ726-11|BOLD:AAL7817
Psychodinae|[16590]|JWDCC682-10|BOLD:AAL7817
Psychodinae|[16591]|JWDCC681-10|BOLD:AAL7817
Psychodinae|[16592]|JWDCC680-10|BOLD:AAL7817
Psychodinae|[16593]|JWDCC648-10|BOLD:AAL7817
Psychodinae|[16594]|JWDCJ1158-11|BOLD:AAL7817
Psychodinae|[16595]|JWDCJ293-11|BOLD:AAL7817
Psychodinae|[16596]|JWDCG078-10|BOLD:AAL7817
Psychodinae|[16597]|JWDCF350-10|BOLD:AAL7817
Psychodinae|[16598]|CHDIP726-12|BOLD:AAL7817
Psychoda phalaenoides|[16599]|JWDCD467-10|BOLD:AAF9317
Psychoda phalaenoides|[16600]|JWDCG260-10|BOLD:AAF9317
Psychoda phalaenoides|[16601]|CHDIP1007-12|BOLD:AAF9317
Psychoda|[16602]|JWDCC566-10|BOLD:AAM9201
Psychoda|[16603]|JWDCD379-10|BOLD:AAM9201
Psychoda|[16604]|CHDIP1004-12|BOLD:AAM9201
Psychoda|[16605]|CHDIP425-11|BOLD:AAL7815
Psychoda|[16606]|CHDIP638-12|BOLD:AAL7815
Psychoda|[16607]|JWDCH588-10|BOLD:AAL7815
Psychoda|[16608]|CHDIP1010-12|BOLD:AAL7815
Psychoda|[16609]|JWDCC314-10|BOLD:AAL7815
Psychoda|[16610]|CHDIP439-11|BOLD:AAL7815
Psychoda|[16611]|JWDCD466-10|BOLD:AAL7815
Psychoda|[16612]|CHDIP1187-12|BOLD:AAL7815
Psychoda|[16613]|CHDIP1186-12|BOLD:AAF9318
Psychoda|[16614]|CHDIP1077-12|BOLD:AAF9318
Psychoda|[16615]|JWDCD414-10|BOLD:AAF9318
Psychoda|[16616]|CHDIP1301-12|BOLD:AAF9318
