## Supplementary material for "A Collaborative DNA Barcode Library for the Diptera of Churchill, Canada: Exploration of Biodiversity and Biogeography": Sampling intensity varied across years, with the most sampling occurring in 2010 and the least occurring in 2011 (Supplementary File 1

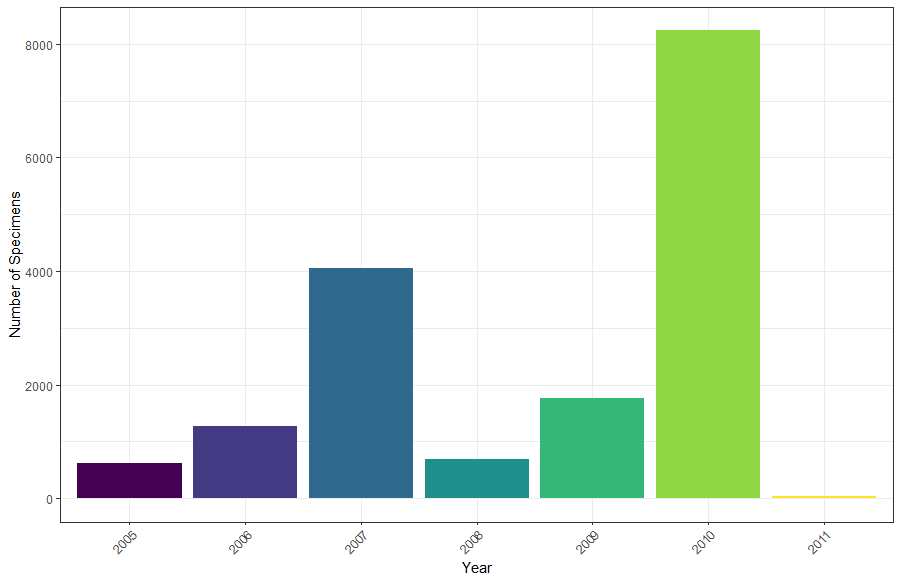

Supplementary Figure 1. Number of specimens collected in each year of this dataset.

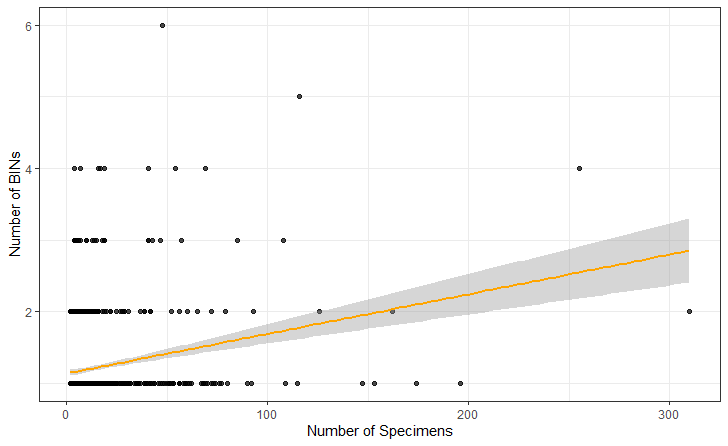

Supplementary Figure 2. The relationship between the number of specimens of each species and the number of BINs associated with each species. Each dot represents a species. Considering only those species with 2 or more specimens, there was a significant relationship between these factors (p-value = 2.2e x10^-16^, R^2^ = 0.07), and species with more specimens are more likely to be split across BINs, although the pattern accounts for a small amount of variability in the dataset. This relationship persists even when excluding the smaller sample of species with a greater than 200 specimens and greater than 3 BINs.

**Land Use Category Definitions**

Definitions for land use categories were taken from the EOSD Land Cover Classification Report (Wulder and Nelson, 2003) which includes recommendations for custom categories for Canada not found in the standard EOSD categories. For the simplicity of visualization purposes, certain categories were combined based on logical groupings. These were:

All Coniferous categories were merged.

All Shrub categories were merged (this entailed only a signal specimen and site).

All Wetland categories were merged.

Exposed/Barren, Non-Vegetated Land, Rock/Rubble and Developed categories were merged.

Bryoids

Land occupied by Bryophytes (mosses, liverworts, and hornworts) and lichen (foliose or fruticose, not crustose). Minimum 20% ground cover or 1/3rd of total vegetation must be bryophyte or lichen.

Coniferous-open

20-60% crown closure. Coniferous trees are 75% or more of total basal area.

Coniferous-sparse

10-25% crown closure. Coniferous trees are 75% or more of total basal area.

Exposed/Barren Land

<5% vegetation. This includes river sediments, exposed soils, pond or lake sediments, reservoir margins, beaches, landings, burned areas, road surfaces, mudflat sediments, cutbanks, moraine, gravel pits, tailings, railway surfaces, building and parking, or other non-vegetated surfaces.

Non-Vegetated Land

<1% vegetation. Analogous to Snow/Ice, Rock/Rubble, Exposed Land or Water.

Rock/Rubble

Land occupied by bedrock, rubble, talus blockfield, rubbly mine spoils or lava beds.

Developed

Land occupied by clearings for human settlement, economic activity, and major transportation routes. Analogous to Exposed/Barren Land

Shrub Low

At least 20% ground cover which is at least 1/3rd shrub (medium-sized perennial woody plant). Average shrub height of <2 m.

Shrubland

Area dominated by shrubs (<6 m tall). Shrub canopy accounts for 25-100% of the cover. Shrub cover is generally >25% when tree cover is <25%. Shrubs cover may be <25% in cases when the cover of other life forms (eg. Herbaceous or tree) is <25% and shrubs cover exceed the cover of the other life forms. Analogous to Shrub Tall and/or Shrub Low

Water

Lakes, reservoirs, rivers, streams, or salt water.

Wetland-herb

Land with a water table near, at, or above the soil surface for enough time to promote wetland or aquatic processes. The majority of vegetation is herb. Herbs are vascular plants without woody stems (grasses, crops, forbs, gramminoids). Minimum of 20% ground cover or 1/3rd of total vegetation must be herb.

Wetland-treed

Land with a water table near/at/above soil surface for enough time to promote wetland or aquatic processes. The majority of vegetation is coniferous, broadleaf, or mixed wood.

Wetland-shrubs

Land with a water table near/at/above soil surface for enough time to promote wetland or
